# Neuron-specific coding and regulatory sequences are the most highly conserved in amniote brains despite neuron-specific cell size diversity

**DOI:** 10.1101/2021.08.20.457147

**Authors:** Linhe Xu, Suzana Herculano-Houzel

## Abstract

Neurons are unique in that they are the only cell type in the body to display massive diversity in cell size, morphology, phenotype, and function both within individuals and across species. Here we use datasets encompassing up to 92 mammalian and 31 sauropsidian species to examine whether neuron-specific diversity occurs with higher evolutionary variation of neuron-specific coding and regulatory sequences compared to non-neuronal cell-specific sequences. We find that the opposite is true: Neuronal diversity in mammalian and sauropsidian evolution arose despite extreme levels of negative selection on neuron-specific protein-coding sequences on par with ATPase coding sequences, the benchmark of evolutionary conservation. We propose that such strong evolutionary conservation is imposed by excitability, which continually exposes cells to the risk of excitotoxic death, and speculate that neuronal cell size diversity is a self-organized consequence of variability in levels of activity, possibly constrained by energy supply to the developing brain.

## Introduction

Most single-nucleated cells in the mammalian body, including glial cells in the brain, maintain a constant size across species from mice to elephants^1,2^. The exception is neurons, whose size (the sum of the soma, axon, and dendrite) varies widely both within each brain and across species, often becoming larger in larger brains within each clade, as gauged by neuronal densities spanning at least 1,000-fold variation across mammalian brain structures and species^3^. Moreover, neuronal variation extends beyond simple cell size: Single-cell transcriptomic studies have confirmed that at least several dozen different types of neuronal cells exist within a single cerebral cortex^4,5^, onto which different physiological properties can be mapped^6^. In contrast, despite sharing a progenitor cell type with neurons^7^, astrocyte and oligodendrocyte cells are much more restricted in subtypes and cell size (all arbors included), varying in density by less than 10-fold within brains and across mammalian species^3,8,9^, with consistently only a few cell types defined transcriptomically^4,5^.

We have suggested that the high conservation of glial cell types and densities compared to neurons is evidence of strict evolutionary constraints against glial cell variation, whereas the immense phenotypic diversity of neurons suggests evolutionary variation^9^. Brain-specific protein-coding genes have been shown to be under stronger selective pressure than widely expressed genes^10^. However, brains are composed of not only neurons but also glial cells and endothelial cells. A later study that examined all mRNA expressed in neuronal-and non-neuronal cells in mouse reported that neuron-expressed gene sequences tend to show stronger negative correlation between a measure of evolutionary conservation and levels of gene expression than gene sequences expressed in other cell types^11^, which the authors attribute to stronger selection pressure against transcriptional and translational errors in more highly expressed sequences^12,13^. However, the explanatory power of gene expression level over coding sequence variation reported in the literature is very low, always less than 10% both in whole tissues^14^ (e.g., calculated from R values reported from Additional Data File 6 of *Tuller et al., 2008*) and in single cells^11^ (calculated as Spearman correlation coefficients in *Hu et al., 2020*). Such a low explanatory power casts doubts over the relevance of the authors’ conclusion that there is stronger selective pressure against variation in neuron-expressed genes compared to genes expressed in other cells. More recently, an analysis between human and mouse of the levels of conservation of gene co-expression in different cell types found less divergence in neurons compared to glial cells^15^. Most importantly, all of these studies have been restricted to mouse-human or human-chimpanzee comparisons, leaving unaddressed the wider question of whether the enormous diversity in neuronal cell size compared to all other cell types throughout brain evolution that we have documented across ∼100 species, including dinosaurs^16^, birds^17^, human^18^, and other mammals^19^, can be accounted for by higher variability of neuron-specific genes, or, alternatively, whether that diversity occurs despite strong conservation of these genes, in which case other mechanisms must be sought.

Here we compare protein-coding and upstream promoter sequences of neuron-, glial-and endothelial cell-specific genes across 92 mammalian and 31 sauropsidian species (**Supplementary Data S1**) to test our initial hypothesis that neuron-specific DNA sequences are more diverse than those specific to other brain cell types on the levels of protein-coding genes, upstream promoter sequences, and cis-regulatory elements. To ensure that our results were not biased or limited to any particular definition of cell type-specific genes, we performed all of our analyses on four datasets, each generated with a different technique, tissue specificity, animal age and sex, and data clustering method. The first is the Barres mouse brain transcriptomic database^20^ that lists the expression level of genes in collections of purified neurons, astrocytes, microglia, endothelial cells, and three developmental stages of oligodendrocytes. The second and third are the Linnarsson single cell transcriptomic databases obtained respectively with single cells isolated with microfluidic chip sequenced by Illumina HiSeq 2000^21^ or dissociated cell populations sequenced by the droplet-based 10X Genomics Chromium^22^, and which also differed in being restricted to regions of cerebral cortical tissue^21^ or extending to the entire CNS and PNS^22^ of male and female juvenile mice. Finally, the fourth dataset is another single cell transcriptomic database obtained with a different droplet-based method, Drop-seq, in male adult mice^23^.

As a proxy for evolutionary selective pressure acting on protein-coding sequences, we compared dN/dS ratios^24^ of neuron-, glia- and endothelium-specific genes calculated from dN and dS values in the Ensembl 98 database^25^, using either mouse, rat, human, or chicken as the reference genome. The dN/dS ratios are calculated pairwise for each gene between two species and represent the ratio between non-synonymous and synonymous substitution rates in protein-coding sequences for that gene. A ratio of 1 indicates neutral evolution of the coding sequence. Values of dN/dS greater than 1 indicate positive/directional selection, and values of dN/dS smaller than 1 indicate negative/purifying selection. We expected that if the coding sequences of both neuron-and glia-specific genes are under negative/purifying selection, they should consistently exhibit dN/dS<1, consistent with previous findings in the literature^10^. Additionally, we predicted that neuron-specific genes should have larger values of dN/dS than glial cell-specific genes and endothelium-specific genes given the extreme diversity in neuronal morphologies and cell sizes across mammalian species.

We used two approaches to compare evolutionary conservation of regulatory elements of glial and neuronal cell-specific sequences. First, we examined phastCons scores for the 2,000 bp upstream of coding sequences, as used in comparable studies^26,27^, employing the same definition of cell type-specific genes as in the dN/dS analysis. The phastCons scores indicate the probability that an individual base pair in a sequence is evolutionarily conserved, and those scores can be averaged to provide a mean phastCons score for an entire promoter region across all species in the dataset^28–30^. Second, we examined phastCons scores for cis-regulatory elements uncovered by single-cell chromatin accessibility essays in neuronal or non-neuronal cells in the mouse cerebrum^31^. Because cis-regulatory elements are not confined to any particular upstream sequence, this analysis complements any shortcomings of the comparison of regulatory elements in the 2,000 bp upstream region of coding sequences.

## Results

Contrary to our expectation that the large evolutionary diversity of neuronal cell size would correspond to less evolutionary conservation of neuron-specific sequences compared to glial cell-specific sequences, we found that in all four datasets, using the mouse genome as reference, the distribution of dN/dS ratios in mammals is sharply shifted towards lower values for neuron-specific genes compared to glial-or endothelial cell-specific genes. This is true whether the dN/dS ratio for *individual* neuronal-, glial-or endothelial-specific genes in each species is considered (**Figure 1A**; **Supplementary Table S1** and **Supplementary Table S2**); whether the *averaged* dN/dS ratio across *all* genes specific to neuronal, glial or endothelial cells of all subtypes pooled together is plotted in each species (**Figure 1C**; **Supplementary Table S3** and **Supplementary Table S4**), which circumvents the fact that some genes have missing orthologs in some species; and whether the *averaged* dN/dS across all genes specific to *specific* non-overlapping cell subtypes is plotted (**Figure 1E**; **Supplementary Table S3** and **Supplementary Table S4**). The same data are depicted as histograms in **Figures 1B and 1D**, which show that the distribution of average dN/dS ratios is significantly shifted towards lower values for neuron-specific genes compared to other cell subtype-specific genes. We find that genes specific and exclusive to each glial subtype all have a distribution of dN/dS ratios that is significantly shifted towards higher values than neuron-specific genes, with the highest median found for microglia-specific genes (**Figure 1D,E**). Similar results were obtained when using the human genome (**Supplementary Figure S1**) or the rat genome (**Supplementary Figure S2**) as reference for the calculation of dN/dS, and also across 31 sauropsidian species, using the chicken genome as reference (**Figure 2**), which indicates that our findings in mammalian species generalize to amniotes.

**Figure 1:**
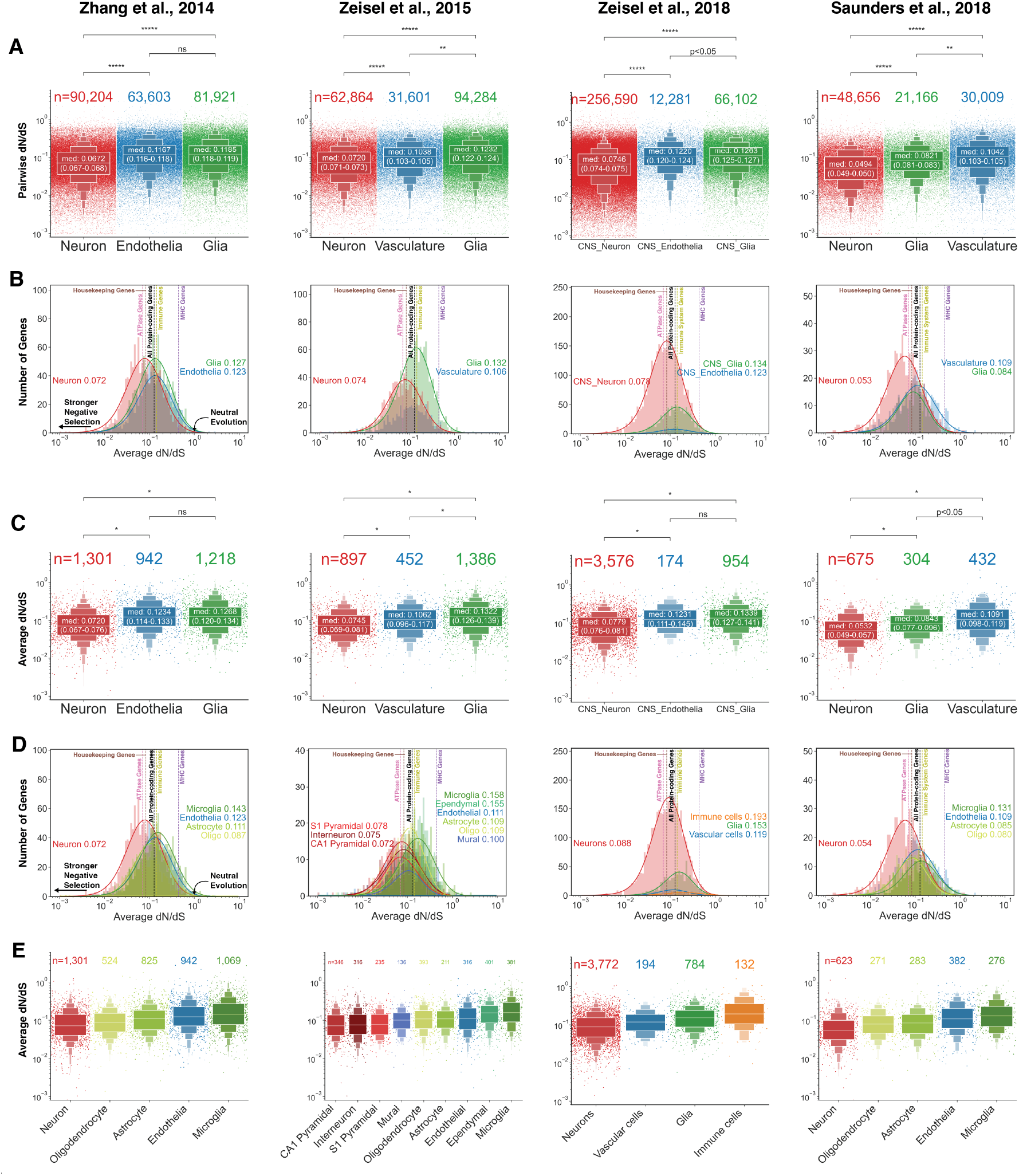
Neuron-specific genes have lower dN/dS ratios than genes specific to glial and endothelial cell types, calculated using the **mouse** genome as reference. **(A)** Enhanced box-strip plots (letter-value plots^32^ overlaying on top of strip plots) of **pairwise** dN/dS ratios of neuron-, glial cell-, and endothelial/vascular cell-specific protein-coding genes in 92 mammalian species against the mouse ortholog. Each dot represents one gene’s pairwise dN/dS in one species relative to the mouse ortholog. **(B, C)** Distribution plots and enhanced box-strip plots of dN/dS ratios **averaged** across 92 mammalian species against mouse reference genome for neuron-, glial cell-, and endothelial/vascular cell-specific protein-coding genes. Each dot represents averaged dN/dS ratios of one gene. **(D, E)** Distribution plots and enhanced box-strip plots of dN/dS ratios **averaged** across 92 mammalian species against mouse reference genome for cell-**subtype**-specific genes. Each dot represents averaged dN/dS ratios of one gene. Each panel shows cell-type-specific genes defined with expression data from four studies, labeled at the top of this figure. For enhanced box plots: the p-values indicate the significance of Mann-Whitney U-tests within each pair of cell types; “ns”: p≥0.05, “p<0.05”: p<0.05, “*”: p<10^-4^, “**”: p<10^-84^, “***”: p<10^-164^, “****”: p<10^-244^, “*****”: p<5×10^-324^. Numbers in the box for each cell type are the medians of pairwise dN/dS ratios, with a 95% confidence interval for the medians in parentheses. The sample size (number of genes) is labeled for each cell type at the top. For distribution plots: a median dN/dS ratio is labeled for each cell type. ATPases and housekeeping genes are considered benchmarks for very low dN/dS (and strong negative selective pressure), while immune system-related genes, and especially MHCs, represent a benchmark for high dN/dS (and strong positive selective pressure). The median of all protein-coding genes is also included as a benchmark.

**Figure 2:**
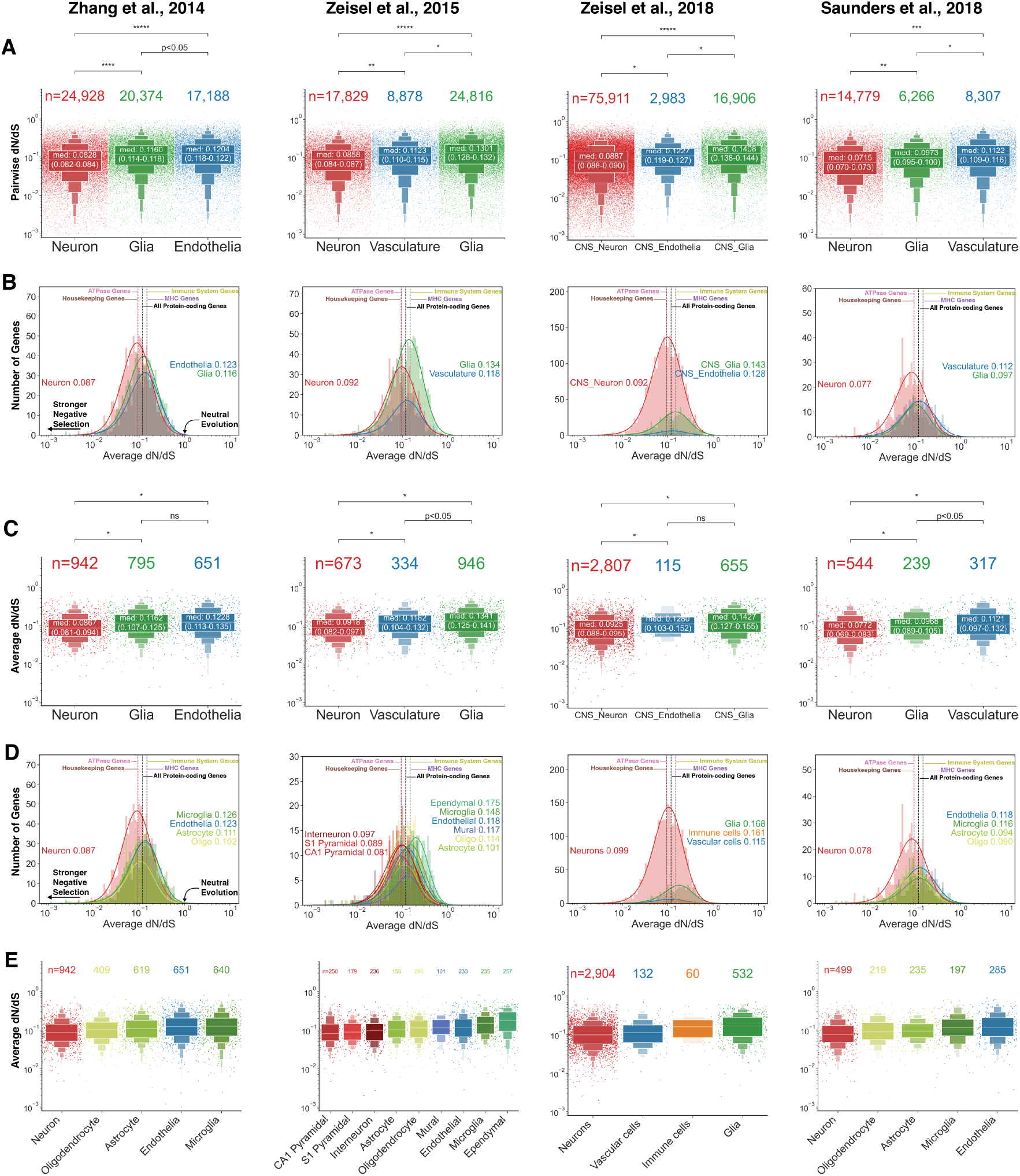
Neuron-specific genes have lower dN/dS ratios than genes specific to glial and endothelial cell types, calculated using the **chicken** genome as reference. **(A)** Enhanced box-strip plots of **pairwise** dN/dS ratios of neuron-, glial cell-, and endothelial/vascular cell-specific protein-coding genes in 31 **sauropsidian** species against the chicken ortholog. Each dot represents one gene’s pairwise dN/dS in one species relative to the chicken ortholog. **(B, C)** Distribution plots and enhanced box-strip plots of dN/dS ratios **averaged** across 31 sauropsidian species against chicken reference genome for neuron-, glial cell-, and endothelial/vascular cell-specific protein-coding genes. Each dot represents averaged dN/dS ratios of one gene. **(D, E)** Distribution plots and enhanced box-strip plots of dN/dS ratios **averaged** across 31 sauropsidian species against chicken reference genome for cell-**subtype**-specific genes. Each dot represents averaged dN/dS ratios of one gene. Each panel shows cell-type-specific genes defined with expression data from four studies, labeled at the top of this figure. For enhanced box plots: the p-values indicate the significance of Mann-Whitney U-tests within each pair of cell types; “ns”: p≥0.05, “p<0.05”: p<0.05, “*”: p<10^-4^, “**”: p<10^-84^, “***”: p<10^-164^, “****”: p<10^-244^, “*****”: p<5×10^-324^. Numbers in the box for each cell type are the medians of pairwise dN/dS ratios, with a 95% confidence interval for the medians in parentheses. The sample size (number of genes) is labeled for each cell type at the top. For distribution plots: a median dN/dS ratio is labeled for each cell type.

The 92 mammalian species in our analyses encompass all major mammalian clades (marsupials, afrotherians, xenarthrans, glires, primates, carnivorans, chiropterans, eulipotyphlans, perissodactyls, artiodactyls), with the exception of monotremes. To address whether our mammal-wide results apply to all clades, we compared pairwise dN/dS ratios for neuronal and non-neuronal cell type-specific genes defined with the Barres expression database^20^ in individual representative species from different clades: primates (human), glires (rat), marsupials (opossum and Tasmanian devil), chiroptera (large flying fox), carnivorans (cat) and artiodactyls (pig). In all of these species, we found significantly lower values of dN/dS for neuron-specific genes compared to genes specific to glial cells (**Table 1**), glial subtypes (**Supplementary Table S5**), and endothelial cells with the exception of oligodendrocyte-specific genes in rat and flying fox (**Supplementary Figure S3**, upper panel). These results hold across the four gene-specificity-defining methods, the four reference genomes, and the individual target species (92 mammalian species and 31 sauropsidian species; **Supplementary Table S6** and **Supplementary Table S7**).

A similar analysis using all genes expressed by each cell type (whether or not exclusively), defined as all protein-coding genes with FPKM higher than 1 for each cell type in the Barres dataset, also confirmed that neuron-expressed genes have a distribution of dN/dS ratios in these select species shifted towards lower values than other cell type-expressed genes (**Supplementary Figure S3**, lower panel; **Supplementary Table S8**).

**Table 1:** Neuron-specific genes have lower dN/dS ratios than glia-specific genes in representative mammalian species. Inferential statistics for cell type-specific genes dN/dS ratios’ calculated against the mouse reference genome for species representative of the different mammalian clades examined, with cell-type-specificity defined according to expression data from the Barres expression database^20^. To validate the result of lower neuron-specific genes’ dN/dS ratios, neuron-specific genes’ dN/dS are compared to glia-specific genes’ dN/dS between mouse and one of seven representative mammalian species. Median dN/dS and 95% confidence interval (CI_95%_), as well as the Mann-Whitney U test results (p-value and common language effect size), are reported here. Values excerpted from **Supplementary Table S6** and **Supplementary Table S7**.

| *Species* | *Neuron-specific*<br>*Genes median*<br>*dN/dS (CI<sub>95%</sub>)* | *Glia-specific*<br>*Genes median*<br>*dN/dS (CI<sub>95%</sub>)* | *Mann-*<br>*Whitney U*<br>*Test p-value* | *CLES* |
| --- | --- | --- | --- | --- |
| Human | 0.063 (0.059-0.068) | 0.117 (0.112-0.123) | $3.24 \times 10^{-40}$ | 66.27% |
| Rat | 0.075 (0.068-0.083) | 0.135 (0.126-0.143) | $4.69 \times 10^{-37}$ | 65.30% |
| Opossum | 0.061 (0.058-0.065) | 0.085(0.079-0.090) | $4.67 \times 10^{-12}$ | 59.23% |
| Megabat | 0.084 (0.074-0.089) | 0.130 (0.118-0.141) | $4.01 \times 10^{-16}$ | 64.18% |
| Tasmanian Devil | 0.057 (0.054-0.061) | 0.087 (0.080-0.091) | $7.03 \times 10^{-19}$ | 62.23% |
| Cat | 0.067 (0.062-0.071) | 0.114 (0.109-0.123) | $3.99 \times 10^{-38}$ | 66.22% |
| Pig | 0.064 (0.061-0.069) | 0.114 (0.107-0.124) | $1.96 \times 10^{-41}$ | 67.00% |

To provide benchmarks against which to compare the values of dN/dS for neuron-specific genes, we calculated median dN/dS ratios for genes known to be highly or poorly conserved, as well as for all protein-coding genes, across the same 92 mammalian species compared to the mouse (**Figure 1B,D**), human (**Supplementary Figure S1**), and rat reference genomes (**Supplementary Figure S2**), and across the same 31 sauropsidian species compared to the chicken reference genome (**Figure 2B,D**). We found that almost half of all neuron-specific genes have values of dN/dS smaller than the median dN/dS of ATPases and housekeeping genes (low dN/dS), which are the gold standards for highly conserved gene sequences^33,34^. Remarkably, neuron-specific genes are as evolutionarily conserved as the benchmarks of conservations, if not more, regardless of the reference genome used, and across four independent gene expression datasets (**Supplementary Table S3** and **Supplementary Table S4**). Conversely, with human, rat, and mouse reference genomes, microglia-specific genes have a similar distribution of dN/dS ratios to immune system-specific genes, which are known to be highly variable (high dN/dS)^35^, although still smaller than the high dN/dS ratio benchmark for major histocompatibility complex (MHC) genes^36^.

To put into context the low dN/dS ratios of neuron-specific genes (defined according to *Zhang et al., 2014*)^20^, we compared them to the dN/dS ratios of organ-specific genes in mouse, defined as genes with expression detected only in that organ but not anywhere else in the MGI GXD mouse gene expression database^37^. Interestingly, except for heart-and musculature-specific genes, we found that liver, lung, skin, pancreas, kidney, and brain-specific genes as a whole all have significantly higher average dN/dS ratios than neuron-specific genes (pancreas<0.05, all other p<0.0001; **Figure 3**; **Table 2**).

**Figure 3.**
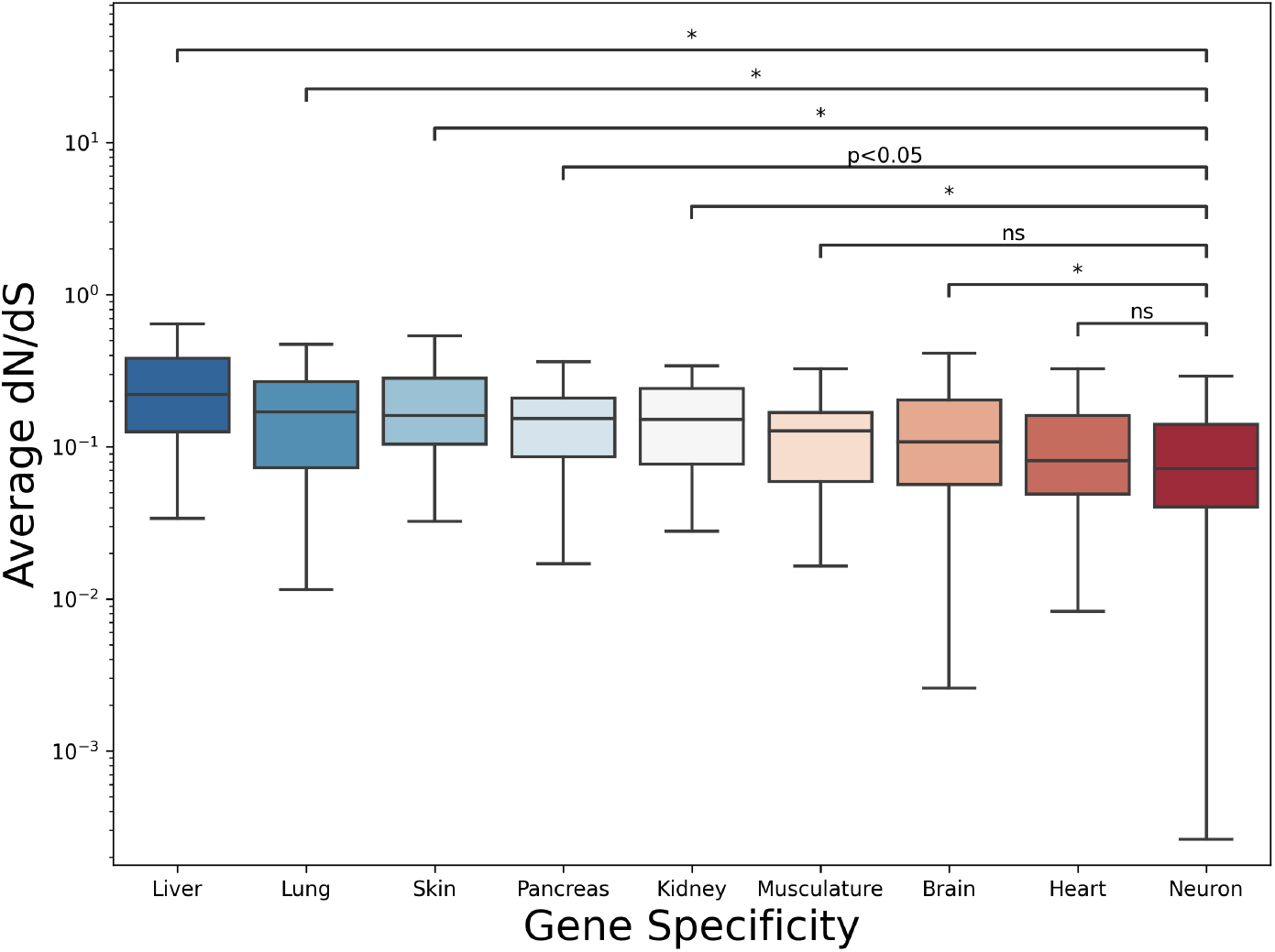
All organ-specific genes (including the brain) other than heart-specific and musculature-specific genes have significantly higher dN/dS than neuron-specific genes. The box plot shows a comparison of dN/dS between neuron-specific genes, as defined by the Barres dataset, and organ-specific genes in the body, using the mouse genome as reference. The p-values were generated by the Mann-Whitney U test between neuron-specific genes and each gene specificity. Note that brain-specific genes overlap with both glia-specific and neuron-specific genes previously analyzed. Data points that extend the 1.5 interquartile range past the low and high quartiles are identified as outliers and are not shown.

**Table 2.** Organ-specific dN/dS compared to neuron-specific genes defined using the Barres dataset as in **Figure 3**. The dN/dS ratios listed are the average across 92 mammalian target genomes against the mouse reference genome. Organ-specificity is based on the MGI expression database^37^. The 95% confidence intervals are provided alongside the median dN/dS. Mann-Whitney U-test p-values and common language effect sizes (CLES) are reported.

| Organ | # Genes | Median dN/dS<br>(CI<sub>95%</sub>) | M.W.U. p-value<br>Against<br>Neuron | CLES |
| --- | --- | --- | --- | --- |
| **Neuron** | 1,301 | 0.07 (0.07-0.08) | N/A | N/A |
| **Heart** | 48 | 0.08 (0.06-0.11) | 0.21 | 55.37% |
| **Brain** | 412 | 0.11 (0.09-0.12) | **$5.94 \times 10^{-11}$** | 60.68% |
| **Musculature** | 17 | 0.13 (0.06-0.16) | $7.85 \times 10^{-2}$ | 62.40% |
| **Kidney** | 43 | 0.15 (0.09-0.21) | **$9.83 \times 10^{-6}$** | 69.79% |
| **Pancreas** | 42 | 0.15 (0.11-0.18) | **$2.48 \times 10^{-4}$** | 66.59% |
| **Skin** | 34 | 0.16 (0.10-0.25) | **$1.08 \times 10^{-6}$** | 74.46% |
| **Lung** | 37 | 0.17 (0.12-0.23) | **$1.73 \times 10^{-6}$** | 73.02% |
| **Liver** | 77 | 0.22 (0.17-0.27) | **$1.68 \times 10^{-20}$** | 81.43% |

Next, we sought to determine, still using the Barres dataset to define cell type-specific genes^20^, whether the neuron-specific genes with low dN/dS and the microglial cell-specific genes with high dN/dS are disproportionately related to particular biological processes. We first identified the 71 slim gene ontology (GO) categories^38^ to which the cell type-specific genes belong (**Supplementary Data S2**). For each GO category, we then performed a contingency analysis between cell type-specificity and the bottom and top 25% values of the distribution of dN/dS using the χ^2^ test of independence (concatenated stats in **Supplementary Table S9**; visualization and detailed contingency tables in **Supplementary Data S3**). This test allows us to find whether any GO category is particularly overrepresented amongst neuron-specific genes with low dN/dS ratios, or amongst microglia-specific genes with high dN/dS ratios. As expected given the mesodermal origin of microglial cells^39^, we found highly significant associations between microglial cell-specific genes with the top 25% values of dN/dS and immune system-related GOs (χ^2^=36.048, p=1.92×10^-9^), in agreement with previous reports of enrichment of immune system-related genes in microglial cells^40^. In contrast, neuron-specific genes with low dN/dS ratios were significantly associated with transport-related and signal transduction-related GOs (transport: χ^2^=78.41, p=8.39×10^-19^; signal transduction: χ^2^=54.20, p<1.82×10^-13^). Nonetheless, virtually all GO categories with more than 225 cell type-specific genes represented showed a significant association between cell type being neuronal and dN/dS ratios in the lowest 25% of the distribution.

Finally, we tested whether the large evolutionary variability of neuronal cell size might be attributed to a particularly high evolutionary variability of neuronal cell type-specific **regulatory regions**, including the promoters and the cis-regulatory regions of neuron-specific genes, despite the extreme conservation of the coding regions. We first defined promoter regions as the 2,000 bp upstream of the coding sequences for each cell type-specific gene, according to common usage in the literature^26,27^, and defining cell-specificity of genes according to each of the four databases used in the dN/dS analysis^20–23^. We used phastCons scores^41^ to quantify the evolutionary conservation level of non-coding regulatory elements per gene. The phastCons score represents the probability of negative selection for each base pair given its flanking base pairs, based on probabilities of state transitions in phylogenetic hidden Markov models, with values ranging from 0 to 1 such that the higher the phastCons value for a nucleotide in the sequence, the more likely that that base pair went through negative selection.

Average phastCons scores for the 2,000 bp upstream **promoter** region for cell type-specific genes were sourced from the UCSC Genome Browser^29^ using 1) phastCons60way,which is a multiple alignment of 60 vertebrate species, including 44 mammals, 6 sauropsids (birds and non-avian reptiles), 9 fish, and 1 amphibian, centered around the mouse GRCm38 reference genome, and 2) phastCons77way, which is a multiple alignment of 77 vertebrate species, including 62 sauropsids, 10 fish, 3 amphibians, and 2 mammals. Promoters with less than 80% coverage were filtered out. At the same p<0.0001 level employed for coding sequences, we found no significant differences in promoter region phastCons scores between neuron- and endothelial cell-specific genes; between phastCons of endothelial cell- and glial cell-specific genes; or between neuron- and glia-specific genes. The only exception was found in the Saunders dataset using the chicken-centered phastCons77way score, where promoter regions of neuron-specific genes had significantly higher phastCons scores (therefore more conserved) than those of glial-specific promoter regions (**Figure 4A, C**). Adopting a less stringent p-value threshold of 0.05 would yield promoter phastCons scores that were actually higher for neuron-specific genes compared to glial-cell-specific genes (except in the *Zeisel et al., 2015* dataset with phastCons60way), which would suggest that neuronal promoter sequences are more evolutionarily conserved than glial promoter sequences, not less. Importantly, log-transformed dN/dS ratios of the coding region of cell type-specific genes have virtually no explanatory power over phastCons for the promoter region of the same genes (**Figure 4B, D**). The low correlation that fails to reach statistical significance indicates that the strong conservation of coding sequences unveiled by dN/dS ratios is not compensated by an increased diversity at the 2,000-bp promoter region of neuron-specific genes.

**Figure 4:**
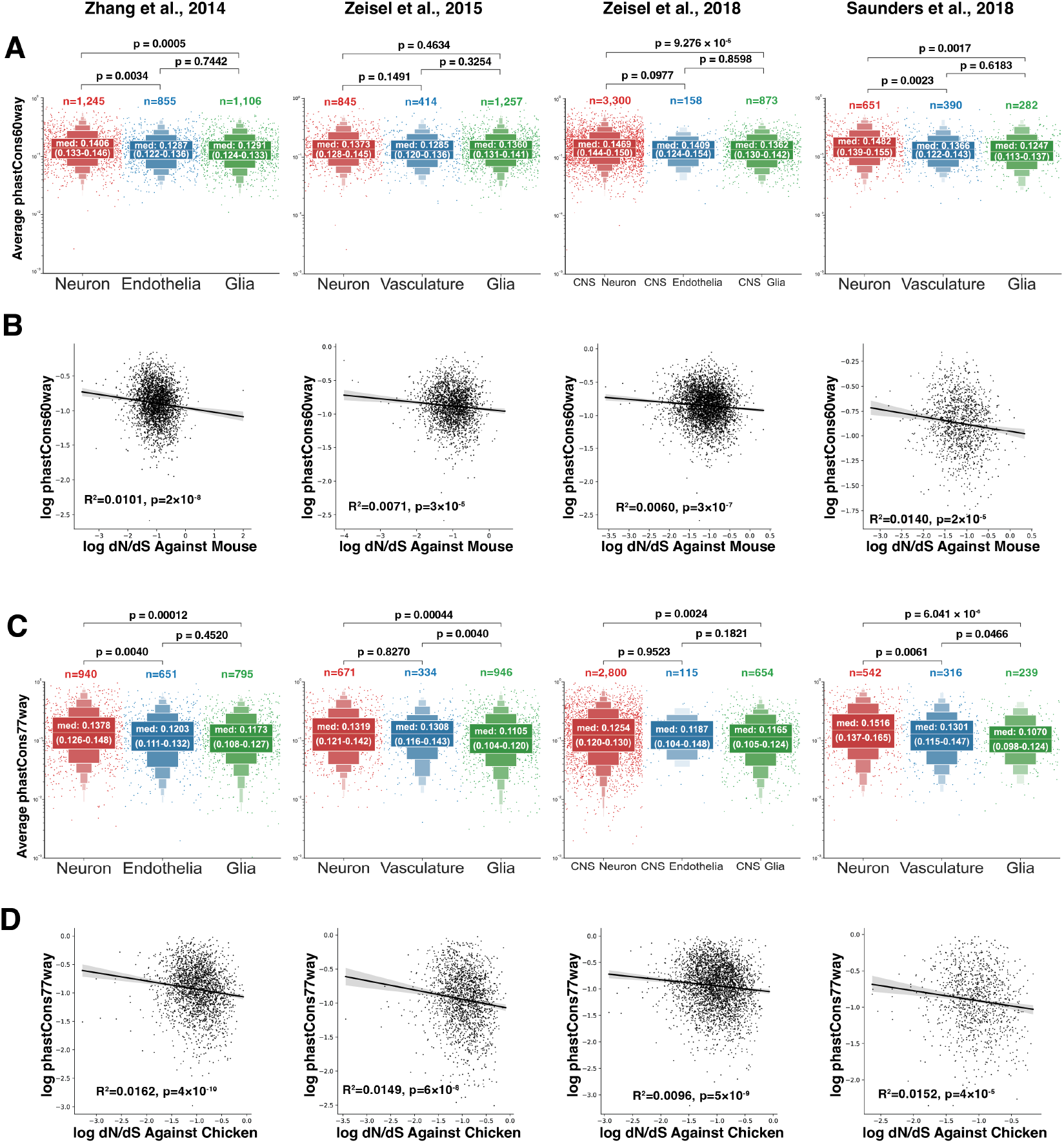
Upstream 2,000 bp regions of neuron-specific genes are not more conserved than those of glial-specific genes. With cell type-specific genes defined with four different datasets, we compared phastCons scores of the 2,000 bp upstream promoter region of cell type-specific calculated with (**A, B**) mouse-referenced multiple alignment of the genomes of 60 vertebrate species and (**C, D**) chicken-referenced multiple alignment of the genomes of 77 vertebrate species. (**A, C**) The phastCons scores show no difference at the p<0.0001 level across neuronal-, glial- and endothelial cell-specific genes, except with phastCons77way score in neuron and glial cell-specific genes defined with the Saunders dataset, where neuron-specific genes’ promoters has significantly higher dN/dS (aka, more conserved) than that of glial cell-specific genes. The p-values reported are from Mann-Whitney U tests; sample size n (i.e., number of cell type-specific genes with high-coverage phastCons) are also reported; in the middle boxes of enhanced box plots, median and 95% confidence interval (in parentheses) are reported. **(B, D)** Virtually no explanatory power was discovered between log-transformed average dN/dS ratios and phastCons scores of brain cell type-specific genes with linear regression (all R^2^ less than 0.02). R^2^ and p-value for the linear fits are reported.

The 2,000 bp sequence upstream of a coding region does however not necessarily function as its promoter. In contrast, chromatin accessibility assays reveal functionally defined cis-regulatory regions, which have been performed in single cell nuclei of neurons and non-neuronal cells in the mouse cerebrum^31^. We calculated the average phastCons scores for these cis-regulatory regions, again sourced from the UCSC Genome Browser^29^, using a multiple alignment of 60 vertebrate species centered around the mouse GRCm38 reference genome (phastCons60ways). In line with our dN/dS observations, we found that functionally defined neuronal cis-regulatory DNA elements have significantly higher phastCons than non-neuronal cis-regulatory DNA elements (**Figure 5**), which indicates that both neuronal-specific cis-regulatory DNA elements and coding sequences are more highly conserved than non-neuronal elements and sequences.

**Figure 5.**
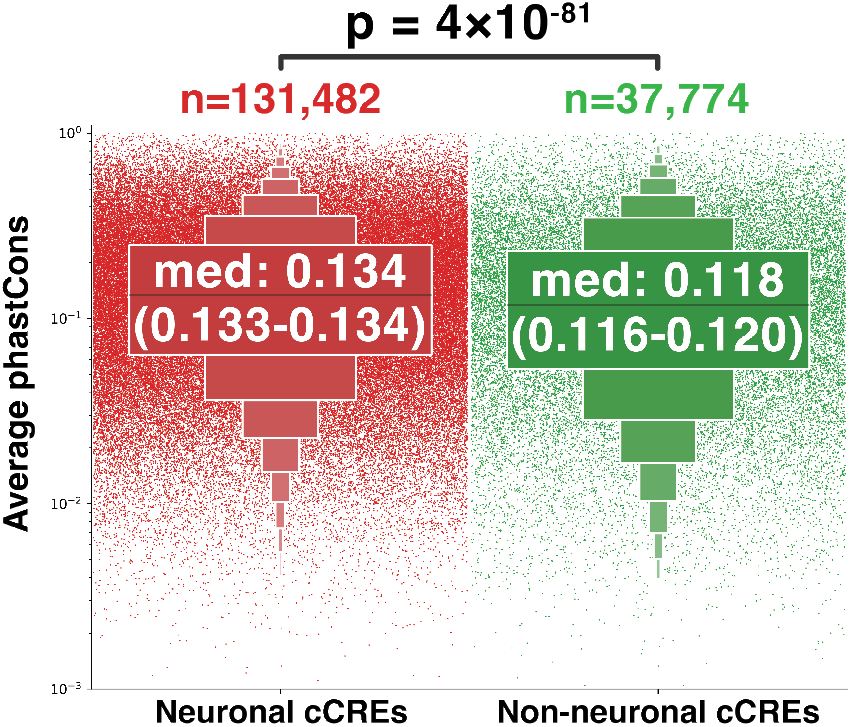
Neuronal cis-regulatory elements (cCREs) have higher phastCons (mouse-centered phastCons60way) than non-neuronal cCREs, indicating they are evolutionarily more conserved. The p-value is reported for Mann-Whitney U test; n: sample size, i.e., number of cCREs; median of phastCons as well as the 95% confidence interval in parentheses reported in the middle box of the enhanced boxplots.

We then tested whether the high conservation of neuron-specific sequences is associated with high levels of gene expression in these cells compared to other cell types. Crossing dN/dS scores for each gene with average expression levels per cell defined as fragments per kilobase of transcript sequence per million mapped fragments^20^ (FPKM) or as unique molecular identifiers (UMI) counts^22,23^, we found that gene expression level has virtually no explanatory power over dN/dS across genes (all three R^2^ values smaller than 0.04; **Figure 6**). Additionally, ANCOVA tests for all three datasets reveal statistically significant differences in dN/dS ratios between neuronal, endothelial/vascular cells, and glial cell types even after controlling for expression level (**Supplementary Table S10**), with lower dN/dS scores for neuron-specific genes compared to non-neuronal-specific genes with similar expression levels (**Figure 6**).

**Figure 6.**
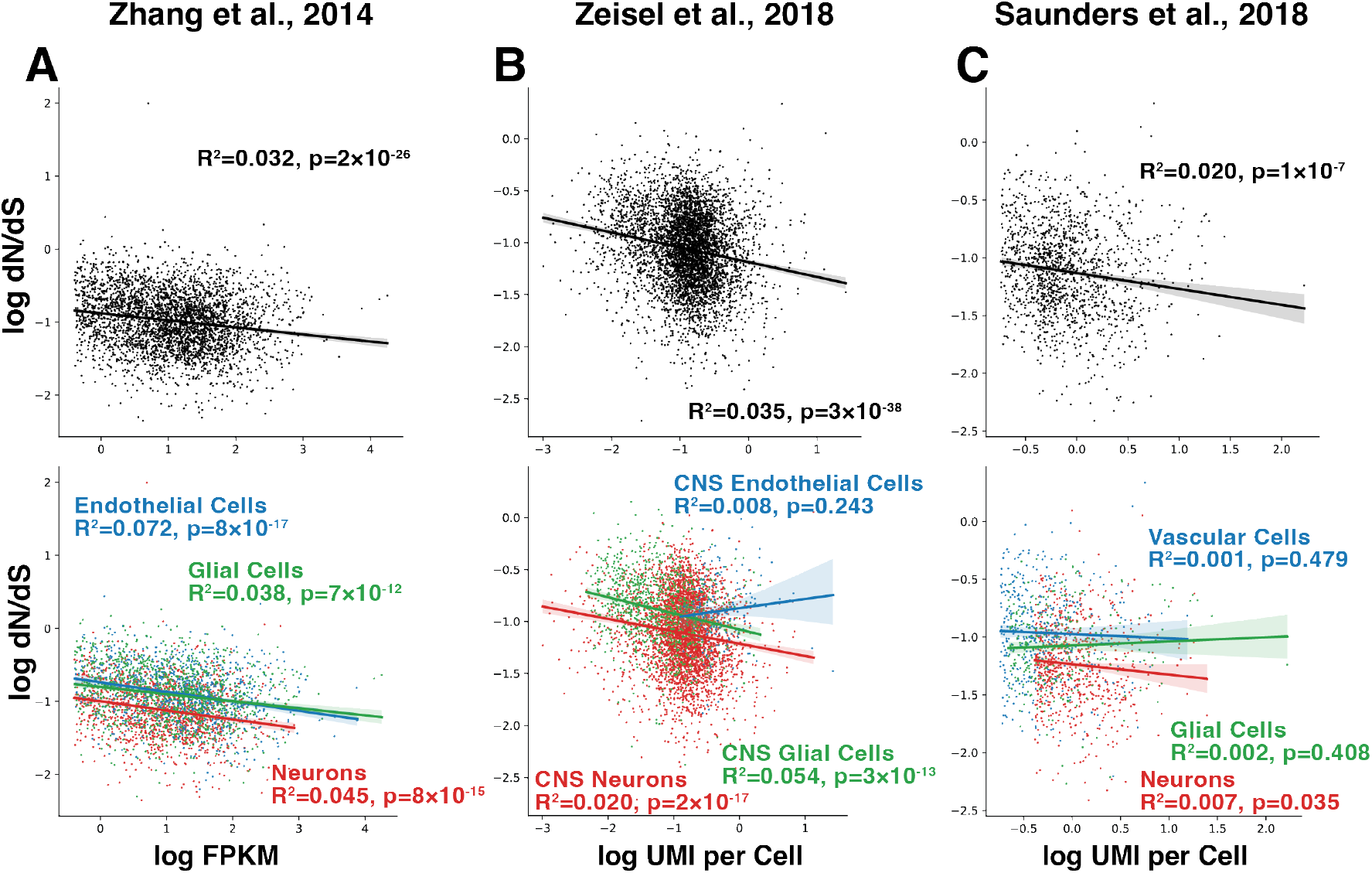
Expression level has virtually no explanatory power over dN/dS of cell type-specific genes, and neuron-specific coding sequences have lower dN/dS values than glial-specific sequences of similar expression level. The upper panel is all cell type-specific genes’ dN/dS and their expression level in corresponding cell types. The lower panel is the same data points color coded with cell type. R^2^ and p-value for each linear regression between log-transformed dN/dS ratios and expression levels are labeled.

It is paradoxical that neuron-specific coding sequences and cis-regulatory DNA elements are more evolutionarily conserved than their non-neuronal counterparts given the much more evolutionarily diverse morphology and cell sizes of neurons, a paradox that indicates that the source of neuronal cell size variability lies elsewhere than in the genome. How could diversity in mammalian neuronal cell size arise both in development and in evolution without building on genomic diversity? We hypothesize, building on our recent demonstration that energy supply to the adult brain is limiting to brain metabolism^42^, that in a scenario where neuronal excitation is energetically costly^43^ and the metabolic rate of the developing brain is also supply-limited, there should be a trade-off between neuronal firing rates and growing cell size, such that neurons with lower firing rates would be the ones to have more energy available for cell growth and thus attain larger adult cell sizes.

To test the hypothesis that variations in cell size should be correlated with (gene-based or not) neuron-specific differences in firing activity, such that larger neurons have lower firing rates, we examined the relationship between cell volume and firing rate across human cortical neurons in the Allen Brain Atlas Cell Type Database^44^, which covers a range of more than 30-fold variation in reconstructed cell volume, including all arbors in the tissue (but no spines). We restricted our analysis to human cortical neurons, 75% of which in the database are excitatory, because the vast majority of mouse cortical neurons in the database were inhibitory. We analyzed all 131 human cortical neurons with both morphological reconstruction and electrophysiological recordings available and calculated their cell volume in the tissue (including soma and all dendritic and axonal arbors) and representative firing rates (see “Materials and Methods”). We found that firing rate and cell size are indeed negatively correlated amongst excitatory neurons (Spearman ρ=-0.59, p<0.0001; **Figure 7**), such that larger neurons in the human cerebral cortex have significantly lower firing rates.

**Figure 7.**
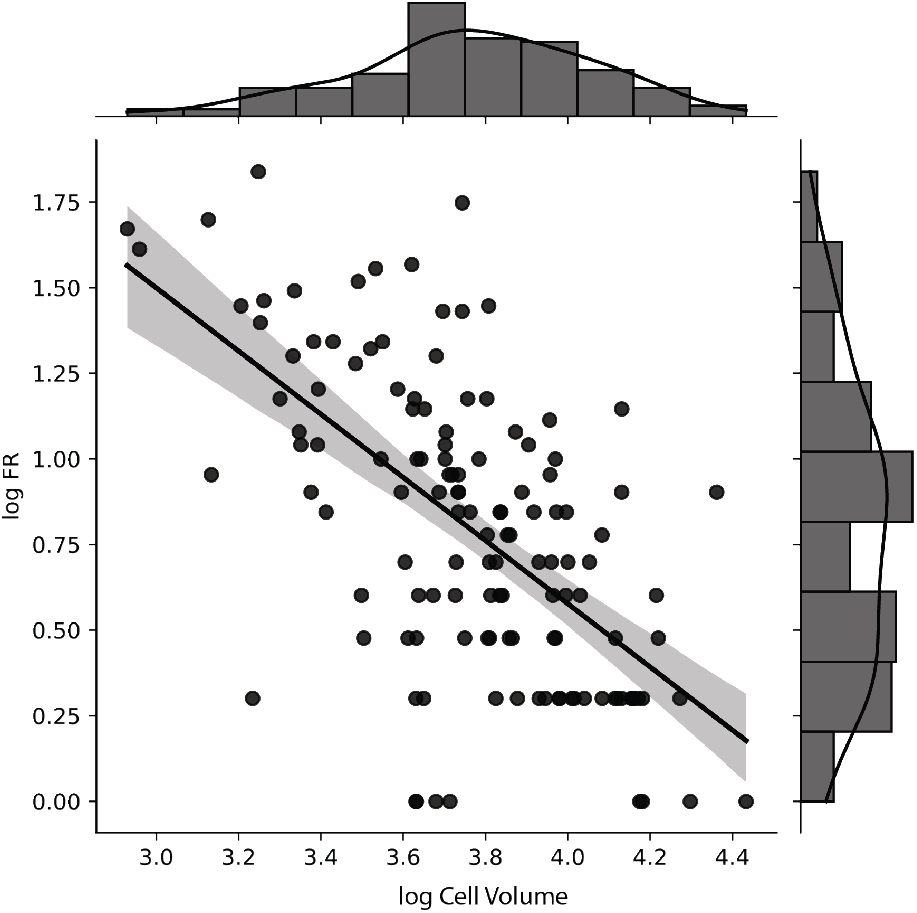
Firing rates and cell volume are negatively correlated across individual neurons in the human cerebral cortex. Linear regression on log-transformed firing rate and cell volume: *y* = −0.92*x* + 4.27; Pearson’s R=-0.61, p=6.814 × 10⁻¹⁵; Spearman’s ρ=-0.59, p=1.13 × 10⁻¹³.

## Discussion

Our initial motivation for this study was to explain the source of the enormous cell size and morphological diversity specific to neurons. Here we refute our initial hypothesis that the enormous evolutionary diversity in neuronal cell size (and therefore density) is due to higher variability in neuron-specific DNA sequences (protein-coding or regulatory) accumulated from mutations in molecular evolution, compared to other cell types. Although it has long been established that differences in protein-coding DNA sequences cannot explain all the morphological differences cross-species^45^, changes in gene expression and regulation are believed to be major players in establishing a species’ biological uniqueness^46^. However, we find evidence that, across vertebrate species, both neuron-specific coding sequences and cis-regulatory sequences are remarkably more conserved than their endothelial cell- and glial cell-specific counterparts, a higher conservation that cannot be explained simply by higher levels of expression, contrary to what has been suggested for protein-coding genes in yeast^12,13^. Our findings go beyond an earlier discovery that brain-specific genes as a whole are highly conserved^10^ by showing that it is specifically the neuron-specific sequences that admit very little evolutionary variation, showing as much conservation as ATPases, the benchmark for evolutionary conservation^33^. Importantly, our findings are reproducible across four methods of defining cell type-specific gene expression in brain cells that range from quantifying gene expression levels in cells isolated by immunopanning to unsupervised clustering of single cell RNA expression, despite the imperfect alignment across methods, as shown in the Venn diagram in **Supplementary Figure S4**. Additionally, we show that the distinctly higher conservation of neuron-specific coding sequences over glial-specific coding sequences occurred in parallel in both reptilian (including birds) and mammalian brain evolution, which implies that fundamental constraints to variation of neuron-specific sequences have been at work across more than 300 million years of evolution^47^. Outside the nervous system, we only find similarly low dN/dS values of genes expressed specifically in other tissues with excitable cells (heart and musculature; **Figure 3** and **Table 2**). Interestingly, we finding that the most conserved neuron-specific genes are enriched in transport-related and signal transduction-related GOs, categories that include genes involved in mediating neuronal excitability, such as ionic channels and metabotropic receptors (**Supplementary Table S9**). We thus hypothesize that molecular pathways associated with electrical excitability events, such as action potentials and membrane repolarization, are particularly sensitive to disruptions that can lead to excitotoxic cell death (for example, calcium overload from prolonged opening of voltage-gated calcium channels leading to mitochondrial release of cytochrome c, triggering the cell death pathway^48^). We postulate that, in the nervous system, this sensitivity strongly restricts the universe of possible functional configurations of those genes expressed specifically in neurons, but not in the other, non-excitable cells that compose brain tissue.

Our results have the important implication that the enormous morphological and functional diversity of neurons within and across mammalian brains arises in a context of strong regulatory and coding sequence conservation. Thus, unless post-transcriptional and/or post-translational mechanisms is responsible for neuronal cell volumes that are at least 500 times more variable than non-neuronal cell volume in adults^3^, or these are due to evolutionary variation in a very particular subset of genes, both cross-species and within-brain differences in neuronal size must arise through self-organizing mechanisms that build on cell-intrinsic variations in levels of gene expression.

However it happens, one expected requirement for the development and evolution of larger neuronal cell sizes is higher energy use, simply because of the larger volume of biosynthesis that builds larger dendritic arbors, axons, and neuronal cell bodies^49^. In neurons, the total energy costs of cell function are compounded by the inflating energetic cost of individual action potentials as neurons become larger, due to the larger membrane capacitance and ATP used to reestablish membrane potential, and further compounded by increased numbers of synapses – unless levels of activity can undergo compensatory regulation^43,50^. Importantly, we have recently demonstrated that brain energetics is constrained by a limited energy supply from blood flow that depends on capillary density in the brain^42,51^, such that oxygen delivery rates have very little room to increase even with an increase in blood flow to local tissue due to oxygen diffusivity from the capillary to brain tissues negatively correlating to blood flow^42^. As a result, neurons within a territory supplied by the same capillary and astrocyte share the same limiting energy and oxygen supply.

It is in the context of such a limiting energy supply extending to the developing brain that we propose that diversity in neuronal cell size arises from the imposed balance between excitability (firing activity) and cell growth. Specifically, we hypothesize that in a scenario where neuronal excitation is energetically costly^43^ and the metabolic rate of the developing brain is also supply-limited, there should be a trade-off between neuronal firing rates and growing cell size, such that neurons with lower firing rates (for instance due to varying firing rate set points, which are characteristic of each neuron through mechanisms yet to be specified^52,53^) would be the ones to have more energy available for cell growth and thus attain larger adult cell sizes. We further propose that differences in combinatorial gene expression across individual neurons in early development^6^ directly modulate neuronal excitability and indirectly affect the maximal cell size that a neuron can attain given a limiting blood supply. Such a central role of excitability in self-regulating neuronal cell size would solve the seeming paradox of how come neurons are the only single-nucleus cell type that occurs in a wide range of cell sizes within individuals, when all cells share the same genome (that is, other than variation from aneuploidies^54^).

Finally, we speculate that species- and clade-specific supply-limited energetics of the developing nervous system during those stages when neurons are in the process of establishing their adult cell sizes links together cell size variation in development and in evolution. For instance, any factors that modulate the blood supply to the uterus during gestation might impact the rate at which energy is supplied to the embryo, and thus allow for a larger range of adult neuronal cell sizes to be attained through the self-regulation of neuronal cell size proposed above. What we propose is thus a testable mechanism of self-regulation of neuronal cell size that solves the seeming paradox of high variability in neuronal cell size in the presence of extreme conservation of neuron-specific sequences.

To summarize, we propose that it is excitability that allows neurons to be uniquely variable in their cell size amongst single-nucleated cells. Neurons are continuously and rapidly excitable thanks to their ability to regenerate the resting membrane potential, without which cells would die^55^. The fact that only neurons can dynamically vary their activity and thus the rate of energy use is, we propose, what sets neurons apart from other cells in the body in their enormous elasticity in neuronal cell volume (soma and all neurites included). Regulation of neuronal cell size through electrical excitability would also explain the surprisingly extreme level of conservation of neuron-specific genes, which we show to be on par with the highly conserved ATPase genes. We submit that viability under conditions of fast and relentless excitability is restricted to cells that express only very particular protein sequences, and most evolutionary tampering with them is swiftly met with cell death.

## Materials and Methods

### Definition of cell-type-specific genes

Cell type-specific genes were selected based on gene expression levels in different cell types in the brain according to each of four different studies, configuring four datasets that we examined to test for congruity in our results. First, we used data from a mouse brain cell type enrichment study from the Ben Barres lab (*Zhang et al., 2014*)^20^ with expression levels in FPKM for 22,458 genes in neurons; astrocytes; oligodendrocytes in three states (precursor cell, newly formed, myelinating); microglia; and endothelial cells.^*^ For the purpose of this study, we averaged the expression level of cells in all three oligodendrocytic states and used the mean FPKM as the expression level for oligodendrocytes. We arbitrarily defined as cell type-specific genes those genes with expression levels in one brain cell type greater than the sum of expression levels in the other four cell types. For example, if a gene’s expression level in neurons is greater than the sum of its expression level in astrocytes, microglia, oligodendrocytes, and endothelial cells, we considered this gene to be a neuron-specific gene in the mouse brain. We repeated the same process to determine astrocyte-, oligodendrocyte-, microglia- and endothelial cell-specific genes. We then averaged the FPKM values in astrocytes, oligodendrocytes, and microglial cells and used that value as the expression level in glial cells as a whole. Effectively, that means that for the five-cell-types (NEAMO) analysis we use a threshold of four times the average expression level across four other cell types. We then identified glial cell-specific genes as genes with expression levels in glial cells greater than four times its average expression level in neurons and endothelial cells, arithmetically similar to our threshold for the five-cell-types analysis. Therefore, cell type-specific genes identified with the Barres dataset are exclusive, meaning that neuronal-, glial-, and endothelial cell-specific genes are non-overlapping. Alternatively, using the same database, we identified cell type-expressed genes, are protein-coding genes with their expression level (FPKM) in one cell type greater than 1, whatever their expression level in other cell types.

Three additional scRNA-seq studies were used to select cell type-specific genes to test the generalizability and robustness of our results. The first, a study from the Linnarsson group on the somatosensory cortex and hippocampal CA1 in P21-P31 mice of both sexes, *Zeisel et al.,* 2015, which has nine major cell types after clustering, provided lists of cell type-specific genes for each cell type in their supplement^21^. This study selected cell type-specific genes based on two criteria: 1) the gene was enriched with 99.9% posterior probability in that cell type and 2) the gene is not enriched in any other cell type. To make this analysis comparable to our analysis with the Barres data, we merged the nine cell types into three cell types: neurons (originally CA1 pyramidal neurons, S1 pyramidal neurons, and interneurons), glial cells (originally oligodendrocytes, astrocytes, microglial cells, and ependymal cells), and capillary cells (originally endothelial cells and mural cells) and did three-group analyses. Like the neurons, the mural cells are also excitable cells^56,57^, including vascular smooth muscle cells and pericytes^21^ that regulate the rate of blood flow^58^. As single-cell expression level values for *Zeisel et al., 2015* were not disclosed in the dataset, we could not include a regression analysis between dN/dS and FPKM or UMI count for it as we did for other studies in **Figure 6**.

The Linnarsson group later published a more comprehensive scRNA-seq dataset of the entire mouse nervous system, including not only the CNS but also the PNS and the enteric nervous system (*Zeisel et al., 2018*)^22^. After dimensionality reduction through manifold learning and clustering, they generated a curated classification of cells in the mouse nervous system, with 265 clusters in total. These clusters were automatically assigned to one of seven major classes, namely neurons, oligodendrocytes, astrocytes, ependymal cells, peripheral glia, immune cells, and vascular cells. We accessed expression level (UMI counts) and additional meta-data of all 265 cell clusters through a multidimensional matrix in loom format^†^. One data layer that *Zeisel et al., 2018* provided is a “trinary score” which indicates whether a gene can be confidently categorized as expressed in a specific cluster. We used a 0.999 trinary score cut-off to define cell type-specific genes. For CNS neuronal, CNS glial, and CNS endothelial cell-specific genes, if a gene has a “trinary score” greater or equal to 0.999 in at least one cluster under the umbrella cell type, but not in any cluster under the other two umbrella cell types, it is defined as cell type-specific. To make this three-cell-type analysis comparable to our analysis using the Barres database, we included microglial clusters as glial clusters. With the same 0.999 trinary score threshold, we also defined cell type-specific genes on umbrella clusters defined for the entire nervous system (i.e., not just CNS) by *Zeisel et al., 2018*, recorded as Rank 1 Taxonomy in their metadata labeling. In this case, we use the Linnarsson criterion and clustered microglial clusters into the immune cell group.

The third scRNA-seq dataset (and the fourth RNA-seq study) we used was a study from the McCarroll group (*Saunders et al., 2018*)^23^, which developed the Drop-seq sequencing method, that collected ∼700,000 single cells in nine regions of the adult mouse brain and clustered them into 565 clusters (referred to as “metacells”) that are transcriptionally distinct^‡^. The “metacells” can be categorized into three master classes: neurons, glial cells, or vascular cells. For each “metacell”, we calculated an “expression percentage” by dividing each gene’s aggregate UMI counts by the total UMI counts in that “metacell”. We then define cell type-specific genes as genes with more than 0.1% of the total expression of metacells annotated to be in that cell type that at the same time have the expression percentage value greater than four times the average expression percentage in other cell types.

### Benchmark Genes

ATPase genes, housekeeping genes, immune genes, and MHC genes were used as benchmarks in this study. Human “ATPase genes” used were retrieved from the HUGO Gene Nomenclature Committee database^59§^; mouse, rat, and chicken ATPase genes are orthologs of the human genes. “Housekeeping genes” used were human housekeeping genes^60**^ or mouse, rat, and chicken orthologs of these genes for the other three reference genome. “Immune genes” were mouse or human genes associated with GO:0045087 (innate immune response), selected from the InnateDB database^61††^; rat and chicken immune genes were retrieved from RGD^62^ and Ensembl 98^25^, respectively. For the mouse and human reference genomes, “MHC genes” were retrieved from the literature^63^; for the rat and chicken reference genomes, “MHC genes” were genes associated with the GO term MHC protein complex from the RGD database or from Ensembl 98, respectively.

We defined liver-, kidney-(metanephros-), lung-, skin-, brain-, heart-, pancreas-, and musculature-specific genes as genes with expression recorded only in that organ and no other organ in the MGI mouse gene expression database (GXD)^37^.

### The dN/dS Ratios

Pairwise dN/dS ratios of the genes of interest for each of 92 mammalian species (excluding the reference species) using as reference the mouse (GRCm38.p6) or human (GRCh38.p13) or rat (Rnor 6.0) genome and were calculated from dN and dS values retrieved from the Ensembl 98 database^25‡‡^. A similar procedure was used for the genes of interest expressed in each of 31 sauropsidian (including birds) species using the chicken reference genome (GRCg6a) as reference. Orthologues that are not one-to-one orthologues across pairs of species were filtered out for that target species.

### Gene Ontology

Gene Ontology (GO) slim terms associated with each gene of interest are biological processing slim ontology terms restricted to the mouse species. The genes mapped to each GO slim term were retrieved from the Princeton University Lewis-Sigler Institute for Integrative Genomics’ Generic GO Term Mapper^38§§^. The ontology aspects selected were “biological processing” and restricted to the species *Mus musculus* (MGI). Additional ontology terms “regulation of cell size”, “regulation of membrane potential”, “membrane depolarization”, “membrane hyperpolarization”, and “membrane repolarization”, which are not included in the slim terms, were also added on top of the slim terms because of our interest in cell size regulation and potential roles of unique properties of neurons. The list of genes belonging to each GO term can be found in **Supplementary Data S2**.

### Regulatory Elements’ PhastCons Scores

“Promoter sequences” of cell type-specific genes were defined as the 2,000 base pairs of DNA upstream of each coding sequence in the mouse or chicken genome. Coordinates of these promoters on the autosomes, sex chromosomes, or mitochondria genome were calculated based on the coordinates of the coding sequence on the mouse or chicken genome, and whether the gene is on the positive or negative strand. We downloaded bigWig files for both phastCons60way^***^ (from a mammal-rich vertebrate multiple alignment centered around the mouse genome) and phastCons77way^†††^ (from a sauropsid-rich multiple alignment centered around the chicken genome), which contain the phastCons scores for all available loci on the mouse and chicken genome. We then fed the start and end position of the promoters and the bigWig files to the Unix executable program bigWigSummary^64^ to estimate the average phastCons scores of each promoter, and proceeded to filter out entries with less than 80% phastCons coverage for quality control. These phastCons scores may also be retrieved with the UCSC Genome Browser’s Table Browser utility^29^. For the list of species included in the two multiple alignments, see **Supplementary Data S1**.

The coordinates for neuronal and non-neuronal candidate cis-regulatory sequences were defined based on a chromatin accessibility study on the mouse cerebrum using single nucleus ATAC-seq^31^. Neuronal cis-regulatory DNA elements are those associated with both GABAergic and glutamatergic neurons but not non-neuronal cells; non-neuronal cis-regulatory DNA elements are those associated with non-neuronal cell types but not associated with any of the neuronal cell types. The coordinates of these cis-regulatory elements and the mouse-centered phastCons60way bigwig file were similarly fed to bigWigSummary for calculating the average phastCons scores of each element. After filtering out cis-regulatory DNA elements with less than 80% coverage of phastCons scores for quality control, we analyzed the phastCons of a resulting 131,482 neuronal cis-regulatory DNA elements and 37,774 non-neuronal cis-regulatory DNA elements.

### Single Cell Electrophysiology and Morphology Data

Neuronal cell volume and representative firing rate are calculated based on the Allen Brain Atlas Cell Type Database^44^. Data from 156 cerebral cortical neurons from 35 human subjects that had both morphology reconstructions and electrophysiology recordings are publicly available, which we accessed through the Allen SDK^65^ with our custom Python scripts. Of the 156 human neurons, 131 has a valid “representative sweep” (see below). The human cortex database consists of both inhibitory (with dendrite type “aspiny”) and excitatory neurons (with dendrite type “sparsely spiny” or “spiny”), but predominantly the latter. The mouse counterpart of the dataset consists almost exclusively of inhibitory neurons and for this reason is not included in this study.

From the downloaded SWC files denoting the morphology of each cell, we calculated whole cell volumes (soma and all dendritic and axonal arbors in the volume) with the Python package “neuron_morphology”, which assumes that each compartment is a conic frustum^‡‡‡^. Dendritic spines are not included in original morphological models and therefore are not included in our volume estimates.

For those cells that had morphological reconstructions in the database, we retrieved and studied the quality-controlled current clamp recordings. For each neuron, we first identified the rheobase (the lowest long-square stimulus amplitude at which a cell starts to have spikes) from the cell electrophysiology metadata. We then used the Allen criterion of selecting the current clamp recording with stimulus amplitude 40 to 60 pA above rheobase to select a representative long-square sweep for each neuron. This gave us 131 human cortical neurons with a qualified representative sweep (32 inhibitory neurons and 99 excitatory neurons). We then calculated the firing rate as the number of spikes during the 1-second stimulation interval for that representative sweep for each cell^§§§^.

### Statistical Analysis and the Sharing of Reproducible Codes

We conducted all statistical analyses with custom Python scripts, which are publicly available at https://github.com/VeritatemAmo/neuron-glia-dNdS together with all codes and data needed for replicating this study.

Because dN/dS are log-normally distributed, we chose to use a median-based description of dN/dS values. Confidence intervals are based on ranked dN/dS values of each gene group. A binomial interval was first fitted to the number of genes with the Python SciPy package’s “stats.binom.interval” package, with alpha set to 0.95. After ranking the genes in each group based on dN/dS value, the genes at the low and high interval threshold of each group are set as the lower and higher bound of the confidence interval.

Mann Whitney U tests were performed with the “mwu” function of the pingouin package or “stats.mannwhitneyu” function of the SciPy package. The p-values that these packages return are in the float64 double-precision floating-point format. This means the smallest positive number that can be represented is 4.9406564584124654 × 10⁻³²⁴. Therefore, when we get a p-value of “0.0” from these packages, we report it as p<5 × 10⁻³²⁴ in this paper. We try to include a common language effect size (CLES) to assist our readers in understanding the result whenever possible, but with the huge amount of entries for all 92 species pairwise dN/dS ratios (i.e., when we use raw pairwise dN/dS ratios instead of dN/dS ratios average across the 92 mammalian species), that calculation of CLES exceed our computational capability, and the standard U statistics were reported instead in **Supplementary Table S2**.

For contingency analyses, we developed a custom statistical and visualization analysis pipeline. Essentially, we generated a crosstable with the Python crosstab function from the pandas library between a cell type and a gene ontology. Then, with the SciPy library, odds ratios were calculated with the “fisher_exact” function, and χ^2^ values and p values were generated using the “chi2_contingency” function. The contingency table between each GO term and the genes with the lowest or highest 25% dN/dS ratios, along with distribution plots of cell type-specific genes that belong to each GO term, are included in **Supplementary Data S3** and **Supplementary Table S9**.

Before running linear regression analyses, we first confirmed the normality of the distribution of log-transformed phastCons and dN/dS values with Q-Q plots. We then ran a least square linear regression of log-transformed phastCons and dN/dS scores of cell-type-specific genes. Since zero values cannot be log-transformed, they were excluded from the linear regression.

## Acknowledgments

We thank Profs. John A Capra (now at UCSF) and Jon H Kaas for in-person advice and the Vanderbilt University’s Advanced Computing Center for Research and Education (ACCRE) for technical support. We thank Drs. Nathan Gouwens and Jeremy Miller at the Allen Brain Institute for assistance in using their dataset. We thank Prof. Arpiar Saunders for assisting our understanding and providing further information about their DropViz data. We thank Prof. Amit Zeisel for helping us interpret the trinary scores reported in their 2018 paper. We thank Prof. Ye Zhang for directing us to the new data portal of the Barres dataset. This work was supported exclusively by Vanderbilt University start-up funds to SHH.

## Footnotes

* The original data portal https://web.stanford.edu/group/barres_lab/brain_rnaseq.html is no longer accessible, and the interactive portal was transferred to a new site http://www.brainrnaseq.org/.

† The aggregated loom file can be downloaded from https://storage.googleapis.com/linnarsson-lab-loom/l5_all.agg.loom. The data portal can be accessed at http://mousebrain.org/. This file type can be processed with the loompy python package developed the Linnarsson group: https://linnarssonlab.org/loompy/.

‡ Data retrieved from stable data portal http://dropviz.org/. Aggregate UMI counts in https://storage.googleapis.com/dropviz-downloads/static/metacells.BrainCellAtlas_Saunders_version_2018.04.01.csv. Annotations for the 565 clusters in https://storage.googleapis.com/dropviz-downloads/static/annotation.BrainCellAtlas_Saunders_version_2018.04.01.xlsx.

§ Human ATPase gene names from the HGNC portal at https://www.genenames.org/data/genegroup/#!/group/412. Mouse orthologs matched to human ATPase genes with the online BioMart GUI of Ensembl 98; same for mouse orthologs of other human genes below.

** Retrieved from the stable link: https://www.tau.ac.il/~elieis/HKG/.

†† Retrieved from the stable link: https://www.innatedb.com/annotatedGenes.do?type=innatedb.

‡‡ We access the Ensembl data with a custom bash script that loop through available species with XML BioMart RESTful access. This data can also be accessed through the online BioMart GUI (https://useast.ensembl.org/biomart/martview). Older versions of Ensembl are now archived. To access version 98 used in this study, go to http://sep2019.archive.ensembl.org/index.html.

§§ Access at https://go.princeton.edu/cgi-bin/GOTermMapper on October 13^th^, 2019.

*** Stable link at: http://hgdownload.cse.ucsc.edu/goldenpath/mm10/phastCons60way/.

††† Stable link at: https://hgdownload.soe.ucsc.edu/goldenPath/galGal6/phastCons77way/.

‡‡‡ The pre-calculated volume included in the cell metadata was estimated with L-Measure, which assumes each component to be a cylinder. Since the SWC files include the radius at each node, we reasoned that the conic frustum-based volume estimation would better capture each neuron’s volume in 3D space.

§§§ The pre-calculated firing rate in the Allen electrophysiology metadata is the inverse of inter-spike interval, which yields artificially high firing rates for bursting neurons. Because we are interested in estimating energetic cost of firing over longer time intervals, we calculated firing rates as indicated instead.

