## Supplementary Materials for "Neuron-specific coding and regulatory sequences are the most highly conserved in amniote brains despite neuron-specific cell size diversity"

### Supplementary Material

#### Legends to Supplementary Data Files

**Supplementary Data S1.** Species included in this study. Sheet 1 includes the mammalian species used for calculating pairwise dN/dS values against mouse, rat, and human reference genomes. Sheet 2 includes the reptile species used for calculating pairwise dN/dS values against the chicken reference genome. Sheet 3 includes the 60 vertebrate species of the 60-way multiple alignment used for phastCons analysis based on mouse genes. Sheet 4 includes the 77 vertebrate species of the 77-way multiple alignment used for phastCons analysis based on chicken genes.

**Supplementary Data S2.** Genes belonging to each GO slim term.

**Supplementary Data S3.** Visualization of the distribution of dN/dS ratios for genes belonging to each of the 71 GO slim terms and specific to each cell type, based on *Zhang et al., 2014* expression data. Below each figure are contingency tables analyzing the relationship between GO slim terms and cell type-specific genes with high and low dN/dS. In each cell of the contingency table is the actual count followed by the expected value in parentheses. Below each table are the $\chi^2$, odds ratios, and p values corresponding to the contingency table.

#### Legends to Supplementary Tables

**Supplementary Table S1.** Complete set of all tables with descriptive statistics of either cell-type-specific or cell-subtype-specific genes, defined with four RNA-seq studies, listing the *pairwise* dN/dS ratios between 92 mammalian target genomes against the human, rat, and mouse reference genomes or 31 sauropsidian target genomes against the chicken reference genome. *n: sample size; med: median dN/dS ratio. The 95% confidence intervals are reported in parentheses.

**Supplementary Table S2.** Mann-Whitney U test inferential statistics for the *pairwise* dN/dS ratios listed in Table S1. With mouse, rat, human, and chicken genomes as reference genome, cell-type-specific and cell-subtype-specific genes' dN/dS ratios were calculated between the reference species and one of many target genomes (92 mammalian species for mouse, rat, and human reference genome; 31 sauropsidian species for the chicken reference genome). That is, each gene is represented multiple times since there are multiple target genomes with orthologs to that gene. Cell type specificities are defined with four RNA-seq studies. Mann-Whitney U tests between pairs of cell types reports the U statistics and the p-value. Significant p-values (<0.0001) are bolded.

**Supplementary Table S3.** Complete set of all tables with descriptive statistics of either cell-type-specific or cell-subtype-specific genes, defined with four RNA-seq studies, listing the dN/dS ratios *averaged* across target genomes for each reference genome (92 mammalian target genomes against the human, rat, and mouse reference genomes or 31 sauropsidian target genomes against the chicken reference genome). Benchmarks are also included. *n: sample size; med: median dN/dS ratio. The 95% confidence intervals are reported in parentheses.

**Supplementary Table S4.** Mann-Whitney U test inferential statistics for dN/dS ratios *averaged* across orthologs. With mouse, rat, human, and chicken genomes as reference genome, cell-type-specific, cell-subtype-specific, and benchmark genes' dN/dS ratios were averaged across all orthologs from target genomes (92 mammalian species for mouse, rat, and human reference genome; 31 sauropsidian species for the chicken reference genome). Mann-Whitney U tests between pairs of cell types and/or benchmarks reports the U statistics, the p-value, and the common language effect size (CLES). Significant p-values (<0.0001) are bolded.

**Supplementary Table S5.** Mann-Whitney U test values between neuron-specific genes and glial cell subtype-specific genes (astrocyte-, microglia-, and oligodendrocyte-specific genes) for dN/dS calculated for each available target species using either chicken, human, rat, or mouse as reference genome and cell-type-specific genes defined using the expression data from Zhang et al., 2014 (Barres dataset). Tables include U statistics, p-value, and common language effect size (CLES) for each cell type comparison in each species.

**Supplementary Table S6.** Descriptive statistics of cell type and cell subtype-specific genes' dN/dS ratios between all available target genomes against mouse, rat, human, and chicken reference genome, with cell type specificity defined using expression data from four studies. *n: number of genes; med: median dN/dS ratio, with 95% confidence intervals in parentheses.

**Supplementary Table S7.** Mann-Whitney U test results on dN/dS ratios between pairs of major CNS cell types (neurons, glial cells, endothelial/vascular cells) for 92 mammalian target genomes against the mouse, rat, and human reference genome, and 31 sauropsidian target genomes against the chicken reference genome, with cell type specificity defined using expression data from four studies.

**Supplementary Table S8.** Inferential statistics for dN/dS ratios for genes expressed (whether or not specifically) in neuronal or glial cell types. Similar to cell type-specific genes' inferential statistics, median dN/dS of neuron- and glia-specific genes between mouse and seven representative species are reported along with the Mann-Whitney U test between glia- and neuron-specific genes. Averaged dN/dS ratios of 92 mammalian species are also reported here.

**Supplementary Table S9.** Concatenated statistics of contingency analysis between cell type-specificity (as determined with *Zhang et al., 2014* expression data) and the bottom and top 25% values of dN/dS ratios for genes specific to each GO. The GO terms are ranked by the odds ratio of contingency between neuron-specific genes and genes with the lowest 25% dN/dS ratios within the GO. The lowest two p values between neuron-specific genes and the lowest 25% dN/dS ratios are highlighted in yellow, and they are "Transport" and "Signal Transduction". num_genes: number of brain cell type-specific genes that belong to each GO term; n_med: median dN/dS ratio for neuron-specific genes that belong to that GO term; low_n_chi2: the $\chi^2$ calculated by a $\chi^2$ test of independence between neuron-specific genes and the genes with lowest 25% dN/dS ratios within a given GO term; low_n_p: the p-value calculated by a $\chi^2$ test of independence between neuron-specific genes and the genes with lowest 25% dN/dS ratios within a given GO term; low_n_OR: the odds ratio calculated by a Fisher's exact test between neuron-specific genes and the lowest 25% dN/dS ratios within a given GO term; low_n_fisher_p: the p-value calculated by a Fisher's exact test between neuron-specific genes and the lowest 25% dN/dS ratios within a given GO term.

**Supplementary Table S10.** ANCOVA shows the dN/dS of cell type-specific genes are significantly different across cell types (neuron, glial cells, and endothelial/vascular cells) even when controlling for expression level for all three studies that we have expression level data for.

### Supplementary Figures and Legends

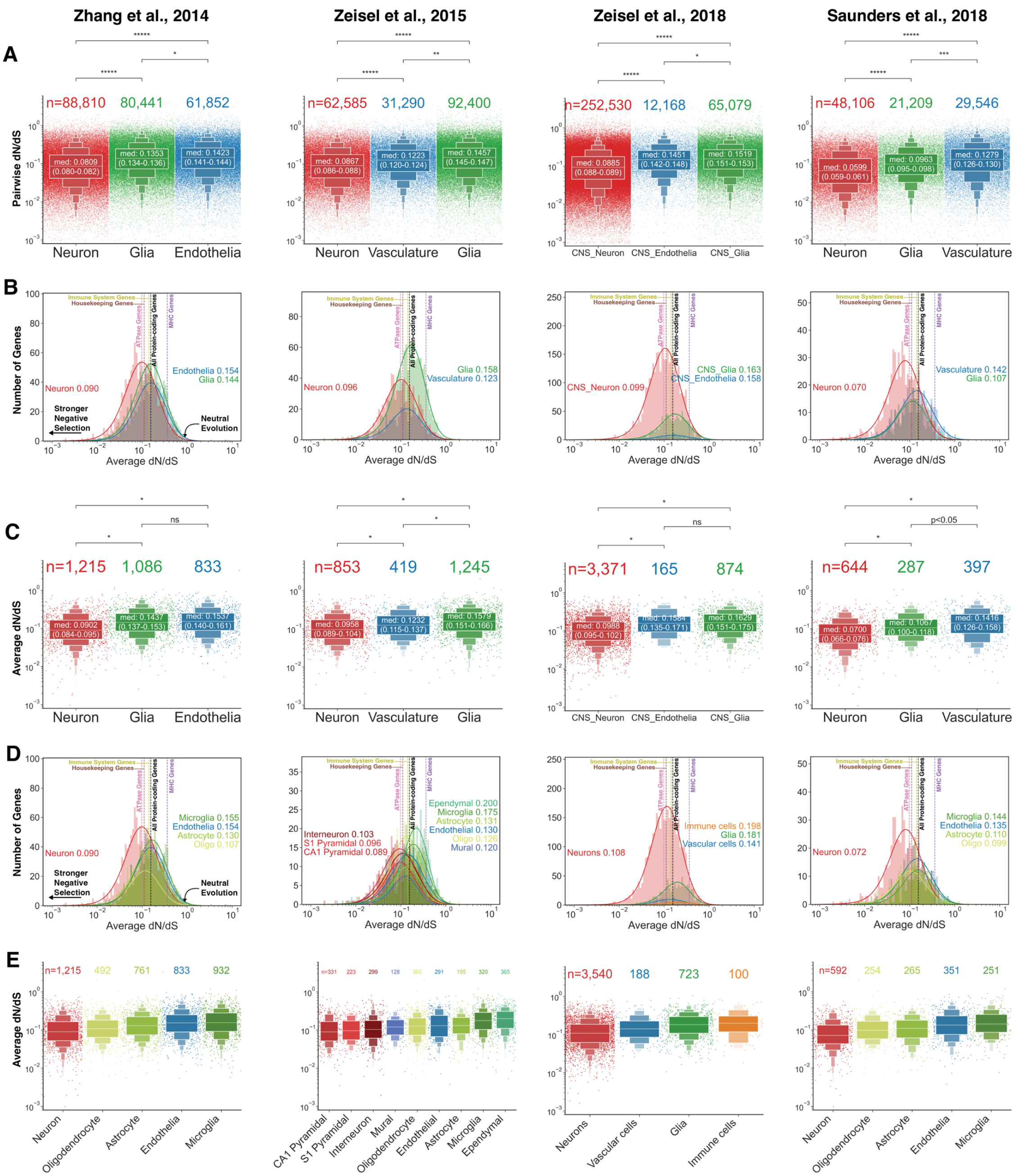


**Supplementary Figure S1:** Neuron-specific genes have lower dN/dS ratios than genes specific to glial and endothelial cell types, calculated using the human genome as reference. (A) Enhanced box-strip plots of pairwise dN/dS ratios of neuron-, glial cell-, and endothelial/vascular cell-specific protein-coding genes in 92 mammalian species against the human ortholog. Each dot represents one gene's pairwise dN/dS in one species relative to the human ortholog. (B, C) Distribution plots and enhanced box-strip plots of dN/dS ratios averaged across 92 mammalian species against human reference genome for neuron-, glial cell-, and endothelial/vascular cell-specific protein-coding genes. Each dot represents the averaged dN/dS ratios of one gene. (D, E) Distribution plots and enhanced box-strip plots of dN/dS ratios averaged across 92 mammalian species against human reference genome for cell-subtype-specific genes. Each dot represents the averaged dN/dS ratios of one gene. Each panel shows cell-type-specific genes defined with expression data from four studies, labeled at the top of this figure. For enhanced box plots: the p-values indicate the significance of Mann-Whitney U-tests within each pair of cell types; "ns": $p \geq 0.05$, "p<0.05": $p<0.05$, "*": $p<10^{-4}$, "**": $p<10^{-84}$, "***": $p<10^{-164}$, "****": $p<10^{-244}$, "*****": $p<5\times10^{-324}$. Numbers in the box for each cell type are the medians of pairwise dN/dS ratios, with a 95% confidence interval for the medians in parentheses. The sample size (number of genes) is labeled for each cell type at the top. For distribution plots: a median dN/dS ratio is labeled for each cell type.

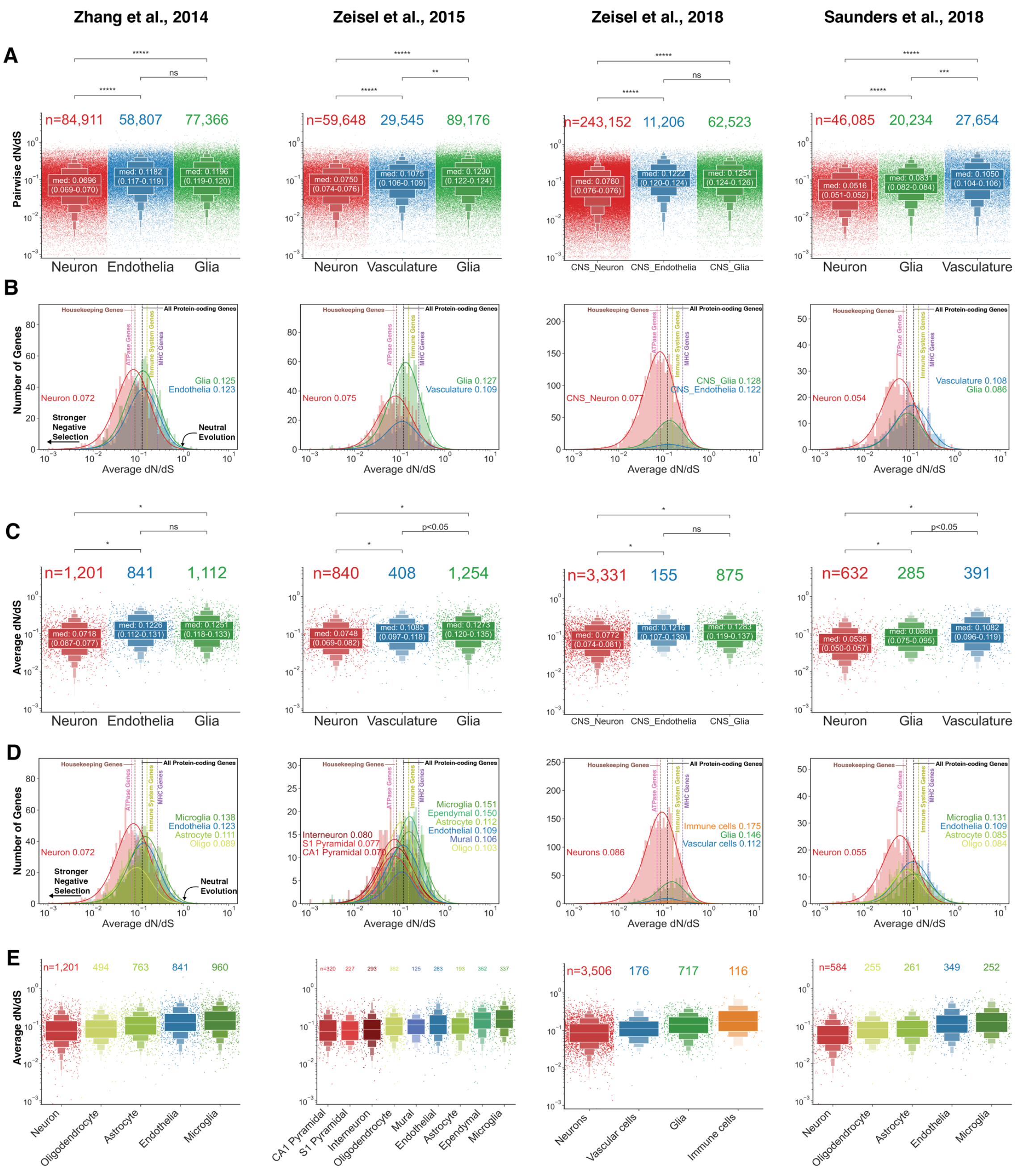


**Supplementary Figure S2:** Neuron-specific genes have lower dN/dS ratios than genes specific to glial and endothelial cell types, calculated using the rat genome as reference. (A) Enhanced box-strip plots of pairwise dN/dS ratios of neuron-, glial cell-, and endothelial/vascular cell-specific protein-coding genes in 92 mammalian species against the rat ortholog. Each dot represents one gene's pairwise dN/dS in one species relative to the rat ortholog. (B, C) Distribution plots and enhanced box-strip plots of dN/dS ratios averaged across 92 mammalian species against rat reference genome for neuron-, glial cell-, and endothelial/vascular cell-specific protein-coding genes. Each dot represents the averaged dN/dS ratios of one gene. (D, E) Distribution plots and enhanced box-strip plots of dN/dS ratios averaged across 92 mammalian species against rat reference genome for cell-subtype-specific genes. Each dot represents the averaged dN/dS ratios of one gene. Each panel shows cell-type-specific genes defined with expression data from four studies, labeled at the top of this figure. For enhanced box plots: the p-values indicate the significance of Mann-Whitney U-tests within each pair of cell types; "ns": $p \geq 0.05$, "p<0.05": $p<0.05$, "*": $p<10^{-4}$, "**": $p<10^{-84}$, "***": $p<10^{-164}$, "****": $p<10^{-244}$, "*****": $p<5\times10^{-324}$. Numbers in the box for each cell type are the medians of pairwise dN/dS ratios, with a 95% confidence interval for the medians in parentheses. The sample size (number of genes) is labeled for each cell type at the top. For distribution plots: a median dN/dS ratio is labeled for each cell type.

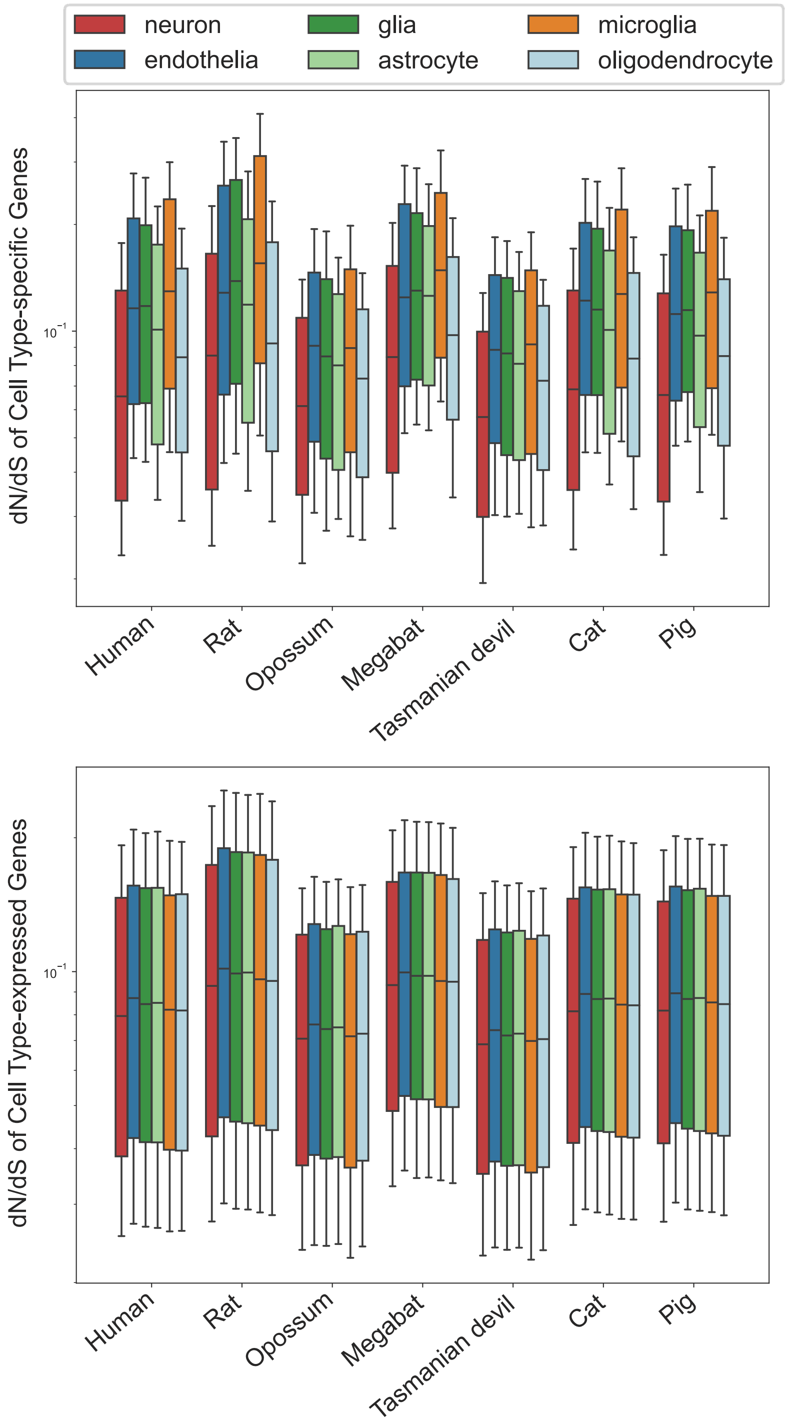


**Supplementary Figure S3**: Box plots of pairwise dN/dS of seven representative species calculated against mouse reference genome for brain cell type-specific (upper panel) or cell type-expressed (lower panel) genes, defined with expression data from *Zhang et al., 2014*. Whiskers show the 15th percentile to the 85th percentile of each distribution.

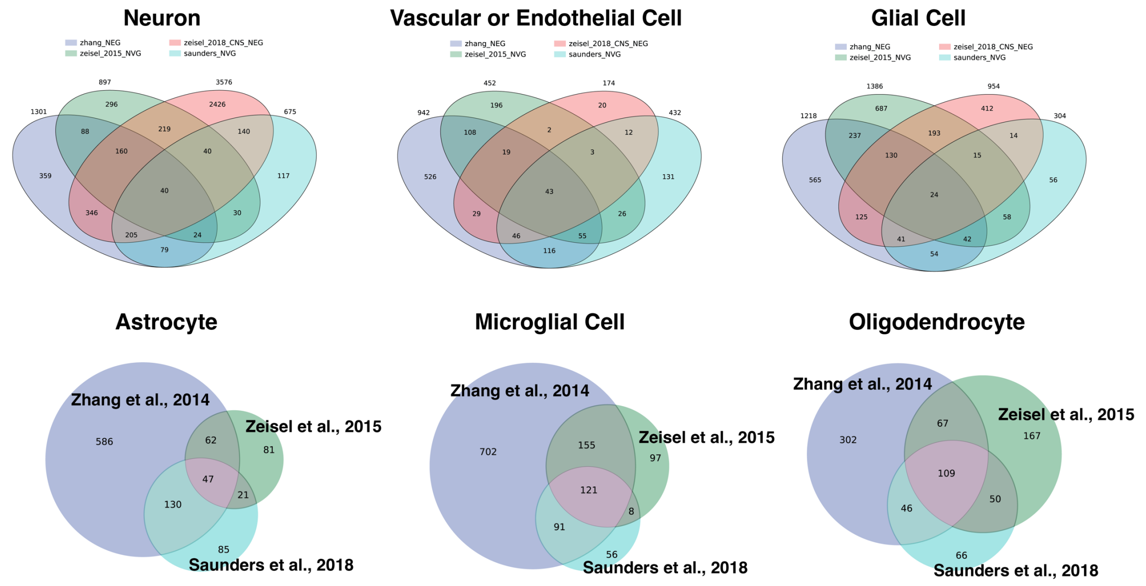


**Supplementary Figure S4:** Venn diagram showing the overlap between the cell-type-specific genes defined with different methods. Numbers indicate the number of genes in each set.
