## Supplementary Data S3 for "Neuron-specific coding and regulatory sequences are the most highly conserved in amniote brains despite neuron-specific cell size diversity"

### Five Celltypes Distribution of Average dN/dS Scores of Genes Related to aging

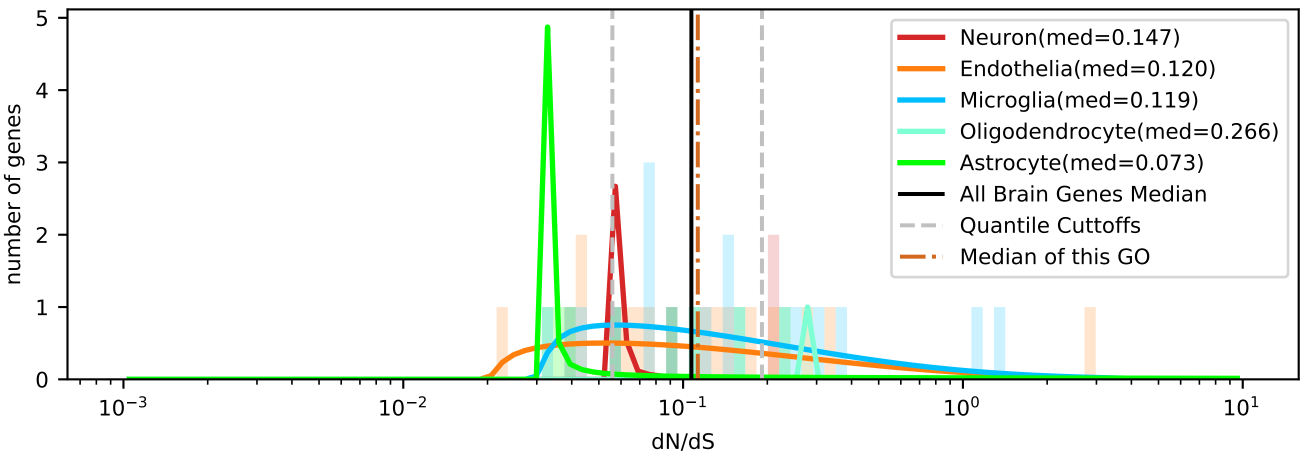


| | < cutoff | other genes | Total |
|---|---|---|---|
| In Astrocyte | 3 (1.70) | 5 (6.30) | 8 |
| Not IN Astrocyte | 7 (8.30) | 32 (30.70) | 39 |
| Total | 10 | 37 | 47 |

chi2: 0.573, p: 0.4492522677, OR: 2.74286

| | > cutoff | other genes | Total |
|---|---|---|---|
| In Astrocyte | 1 (2.38) | 7 (5.62) | 8 |
| Not IN Astrocyte | 13 (11.62) | 26 (27.38) | 39 |
| Total | 14 | 33 | 47 |

chi2: 0.562, p: 0.4536316926, OR: 0.28571

| | < cutoff | other genes | Total |
|---|---|---|---|
| In Microglia | 3 (4.04) | 16 (14.96) | 19 |
| Not IN Microglia | 7 (5.96) | 21 (22.04) | 28 |
| Total | 10 | 37 | 47 |

chi2: 0.155, p: 0.6935567855, OR: 0.56250

| | > cutoff | other genes | Total |
|---|---|---|---|
| In Microglia | 6 (5.66) | 13 (13.34) | 19 |
| Not IN Microglia | 8 (8.34) | 20 (19.66) | 28 |
| Total | 14 | 33 | 47 |

chi2: 0.011, p: 0.9173970735, OR: 1.15385

| | < cutoff | other genes | Total |
|---|---|---|---|
| In Oligodendrocyte | 0 (0.21) | 1 (0.79) | 1 |
| Not IN Oligodendrocyte | 10 (9.79) | 36 (36.21) | 46 |
| Total | 10 | 37 | 47 |

chi2: 0.503, p: 0.4780643953, OR: 0.00000

| | > cutoff | other genes | Total |
|---|---|---|---|
| In Oligodendrocyte | 1 (0.30) | 0 (0.70) | 1 |
| Not IN Oligodendrocyte | 13 (13.70) | 33 (32.30) | 46 |
| Total | 14 | 33 | 47 |

chi2: 0.200, p: 0.6550495820, OR: inf

| | < cutoff | other genes | Total |
|---|---|---|---|
| In Neuron | 0 (0.85) | 4 (3.15) | 4 |
| Not IN Neuron | 10 (9.15) | 33 (33.85) | 43 |
| Total | 10 | 37 | 47 |

chi2: 0.201, p: 0.6538629149, OR: 0.00000

| | > cutoff | other genes | Total |
|---|---|---|---|
| In Neuron | 2 (1.19) | 2 (2.81) | 4 |
| Not IN Neuron | 12 (12.81) | 31 (30.19) | 43 |
| Total | 14 | 33 | 47 |

chi2: 0.124, p: 0.7243582564, OR: 2.58333

| | < cutoff | other genes | Total |
|---|---|---|---|
| In Endothelia | 4 (3.19) | 11 (11.81) | 15 |
| Not IN Endothelia | 6 (6.81) | 26 (25.19) | 32 |
| Total | 10 | 37 | 47 |

chi2: 0.056, p: 0.8135239625, OR: 1.57576

| | > cutoff | other genes | Total |
|---|---|---|---|
| In Endothelia | 4 (4.47) | 11 (10.53) | 15 |
| Not IN Endothelia | 10 (9.53) | 22 (22.47) | 32 |
| Total | 14 | 33 | 47 |

chi2: 0.000, p: 0.9825777546, OR: 0.80000

### Five Celltypes Distribution of Average dN/dS Scores of Genes Related to anatomical structure development

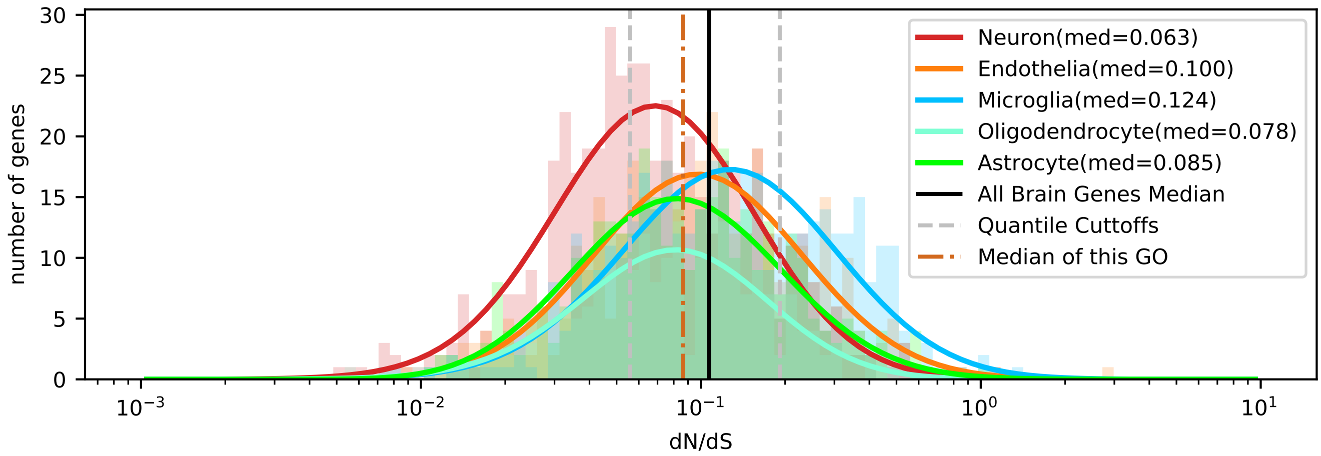


| | < cutoff | other genes | Total |
|---|---|---|---|
| In Astrocyte | 109 (104.11) | 230 (234.89) | 339 |
| Not IN Astrocyte | 465 (469.89) | 1065 (1060.11) | 1530 |
| Total | 574 | 1295 | 1869 |

chi2: 0.326, p: 0.5680241687, OR: 1.08541

| | > cutoff | other genes | Total |
|---|---|---|---|
| In Astrocyte | 57 (66.57) | 282 (272.43) | 339 |
| Not IN Astrocyte | 310 (300.43) | 1220 (1229.57) | 1530 |
| Total | 367 | 1502 | 1869 |

chi2: 1.877, p: 0.1706623745, OR: 0.79547

| | < cutoff | other genes | Total |
|---|---|---|---|
| In Microglia | 80 (127.15) | 334 (286.85) | 414 |
| Not IN Microglia | 494 (446.85) | 961 (1008.15) | 1455 |
| Total | 574 | 1295 | 1869 |

chi2: 31.726, p: 0.0000000178, OR: 0.46595

| | > cutoff | other genes | Total |
|---|---|---|---|
| In Microglia | 140 (81.29) | 274 (332.71) | 414 |
| Not IN Microglia | 227 (285.71) | 1228 (1169.29) | 1455 |
| Total | 367 | 1502 | 1869 |

chi2: 66.614, p: 0.0000000000, OR: 2.76408

| | < cutoff | other genes | Total |
|---|---|---|---|
| In Oligodendrocyte | 75 (68.79) | 149 (155.21) | 224 |
| Not IN Oligodendrocyte | 499 (505.21) | 1146 (1139.79) | 1645 |
| Total | 574 | 1295 | 1869 |

chi2: 0.776, p: 0.3783491667, OR: 1.15600

| | > cutoff | other genes | Total |
|---|---|---|---|
| In Oligodendrocyte | 28 (43.99) | 196 (180.01) | 224 |
| Not IN Oligodendrocyte | 339 (323.01) | 1306 (1321.99) | 1645 |
| Total | 367 | 1502 | 1869 |

chi2: 7.707, p: 0.0054998388, OR: 0.55036

| | < cutoff | other genes | Total |
|---|---|---|---|
| In Neuron | 216 (154.48) | 287 (348.52) | 503 |
| Not IN Neuron | 358 (419.52) | 1008 (946.48) | 1366 |
| Total | 574 | 1295 | 1869 |

chi2: 47.597, p: 0.0000000000, OR: 2.11909

| | > cutoff | other genes | Total |
|---|---|---|---|
| In Neuron | 56 (98.77) | 447 (404.23) | 503 |
| Not IN Neuron | 311 (268.23) | 1055 (1097.77) | 1366 |
| Total | 367 | 1502 | 1869 |

chi2: 30.799, p: 0.0000000286, OR: 0.42498

| | < cutoff | other genes | Total |
|---|---|---|---|
| In Endothelia | 94 (119.47) | 295 (269.53) | 389 |
| Not IN Endothelia | 480 (454.53) | 1000 (1025.47) | 1480 |
| Total | 574 | 1295 | 1869 |

chi2: 9.511, p: 0.0020428773, OR: 0.66384

| | > cutoff | other genes | Total |
|---|---|---|---|
| In Endothelia | 86 (76.38) | 303 (312.62) | 389 |
| Not IN Endothelia | 281 (290.62) | 1199 (1189.38) | 1480 |
| Total | 367 | 1502 | 1869 |

chi2: 1.709, p: 0.1910739433, OR: 1.21107

### Five Celltypes Distribution of Average dN/dS Scores of Genes Related to anatomical structure formation involved in morphogenesis

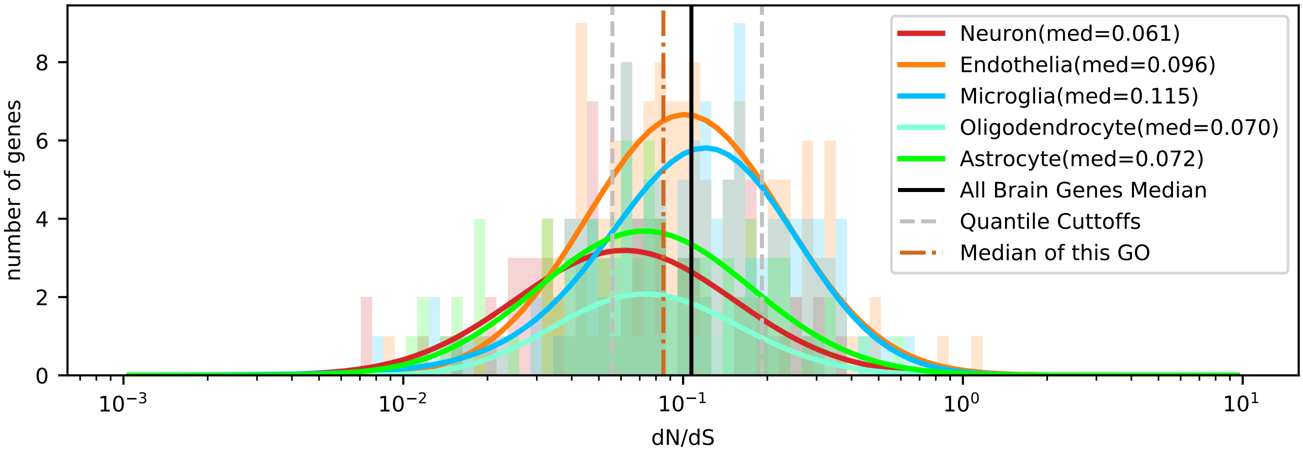


| | < cutoff | other genes | Total |
|---|---|---|---|
| In Astrocyte | 33 (25.83) | 54 (61.17) | 87 |
| Not IN Astrocyte | 108 (115.17) | 280 (272.83) | 388 |
| Total | 141 | 334 | 475 |

chi2: 3.004, p: 0.0830831009, OR: 1.58436

| | > cutoff | other genes | Total |
|---|---|---|---|
| In Astrocyte | 13 (17.22) | 74 (69.78) | 87 |
| Not IN Astrocyte | 81 (76.78) | 307 (311.22) | 388 |
| Total | 94 | 381 | 475 |

chi2: 1.225, p: 0.2684427077, OR: 0.66583

| | < cutoff | other genes | Total |
|---|---|---|---|
| In Microglia | 24 (36.81) | 100 (87.19) | 124 |
| Not IN Microglia | 117 (104.19) | 234 (246.81) | 351 |
| Total | 141 | 334 | 475 |

chi2: 7.921, p: 0.0048858777, OR: 0.48000

| | > cutoff | other genes | Total |
|---|---|---|---|
| In Microglia | 32 (24.54) | 92 (99.46) | 124 |
| Not IN Microglia | 62 (69.46) | 289 (281.54) | 351 |
| Total | 94 | 381 | 475 |

chi2: 3.332, p: 0.0679619428, OR: 1.62132

| | < cutoff | other genes | Total |
|---|---|---|---|
| In Oligodendrocyte | 15 (12.47) | 27 (29.53) | 42 |
| Not IN Oligodendrocyte | 126 (128.53) | 307 (304.47) | 433 |
| Total | 141 | 334 | 475 |

chi2: 0.517, p: 0.4721217326, OR: 1.35362

| | > cutoff | other genes | Total |
|---|---|---|---|
| In Oligodendrocyte | 3 (8.31) | 39 (33.69) | 42 |
| Not IN Oligodendrocyte | 91 (85.69) | 342 (347.31) | 433 |
| Total | 94 | 381 | 475 |

chi2: 3.809, p: 0.0509631773, OR: 0.28910

| | < cutoff | other genes | Total |
|---|---|---|---|
| In Neuron | 37 (22.86) | 40 (54.14) | 77 |
| Not IN Neuron | 104 (118.14) | 294 (279.86) | 398 |
| Total | 141 | 334 | 475 |

chi2: 13.822, p: 0.0002009679, OR: 2.61490

| | > cutoff | other genes | Total |
|---|---|---|---|
| In Neuron | 10 (15.24) | 67 (61.76) | 77 |
| Not IN Neuron | 84 (78.76) | 314 (319.24) | 398 |
| Total | 94 | 381 | 475 |

chi2: 2.192, p: 0.1387361361, OR: 0.55792

| | < cutoff | other genes | Total |
|---|---|---|---|
| In Endothelia | 32 (43.04) | 113 (101.96) | 145 |
| Not IN Endothelia | 109 (97.96) | 221 (232.04) | 330 |
| Total | 141 | 334 | 475 |

chi2: 5.286, p: 0.0215034271, OR: 0.57417

| | > cutoff | other genes | Total |
|---|---|---|---|
| In Endothelia | 36 (28.69) | 109 (116.31) | 145 |
| Not IN Endothelia | 58 (65.31) | 272 (264.69) | 330 |
| Total | 94 | 381 | 475 |

chi2: 2.896, p: 0.0887859349, OR: 1.54888

### Five Celltypes Distribution of Average dN/dS Scores of Genes Related to autophagy

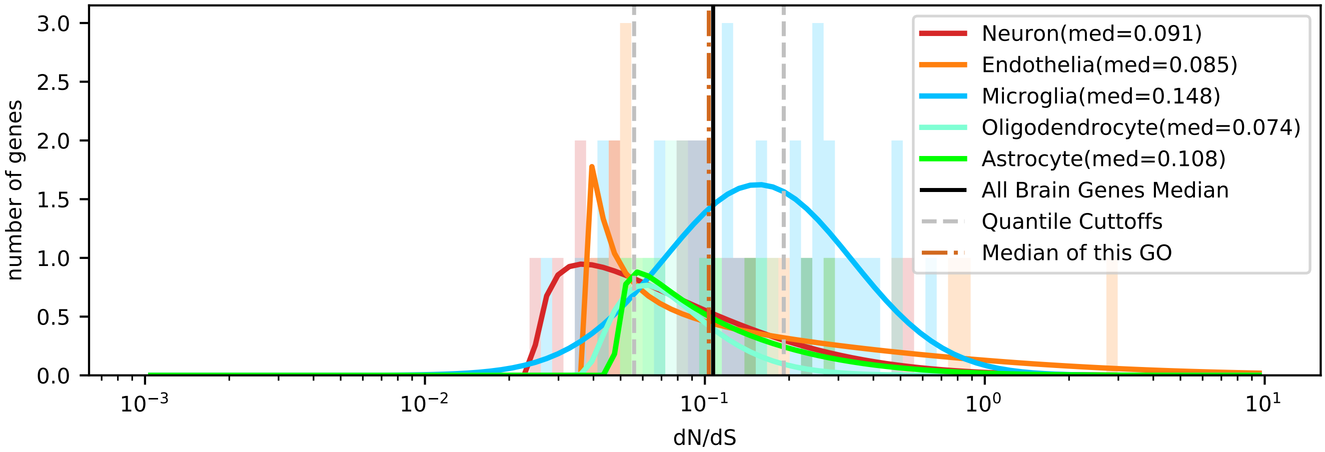


| | < cutoff | other genes | Total |
|---|---|---|---|
| In Astrocyte | 1 (2.56) | 9 (7.44) | 10 |
| Not IN Astrocyte | 21 (19.44) | 55 (56.56) | 76 |
| Total | 22 | 64 | 86 |

chi2: 0.666, p: 0.4146158298, OR: 0.29101

| | > cutoff | other genes | Total |
|---|---|---|---|
| In Astrocyte | 2 (2.79) | 8 (7.21) | 10 |
| Not IN Astrocyte | 22 (21.21) | 54 (54.79) | 76 |
| Total | 24 | 62 | 86 |

chi2: 0.048, p: 0.8274195373, OR: 0.61364

| | < cutoff | other genes | Total |
|---|---|---|---|
| In Microglia | 4 (8.70) | 30 (25.30) | 34 |
| Not IN Microglia | 18 (13.30) | 34 (38.70) | 52 |
| Total | 22 | 64 | 86 |

chi2: 4.502, p: 0.0338506274, OR: 0.25185

| | > cutoff | other genes | Total |
|---|---|---|---|
| In Microglia | 15 (9.49) | 19 (24.51) | 34 |
| Not IN Microglia | 9 (14.51) | 43 (37.49) | 52 |
| Total | 24 | 62 | 86 |

chi2: 6.073, p: 0.0137302189, OR: 3.77193

| | < cutoff | other genes | Total |
|---|---|---|---|
| In Oligodendrocyte | 2 (2.05) | 6 (5.95) | 8 |
| Not IN Oligodendrocyte | 20 (19.95) | 58 (58.05) | 78 |
| Total | 22 | 64 | 86 |

chi2: 0.149, p: 0.6996066401, OR: 0.96667

| | > cutoff | other genes | Total |
|---|---|---|---|
| In Oligodendrocyte | 0 (2.23) | 8 (5.77) | 8 |
| Not IN Oligodendrocyte | 24 (21.77) | 54 (56.23) | 78 |
| Total | 24 | 62 | 86 |

chi2: 2.056, p: 0.1515792524, OR: 0.00000

| | < cutoff | other genes | Total |
|---|---|---|---|
| In Neuron | 8 (4.35) | 9 (12.65) | 17 |
| Not IN Neuron | 14 (17.65) | 55 (51.35) | 69 |
| Total | 22 | 64 | 86 |

chi2: 3.824, p: 0.0505185815, OR: 3.49206

| | > cutoff | other genes | Total |
|---|---|---|---|
| In Neuron | 2 (4.74) | 15 (12.26) | 17 |
| Not IN Neuron | 22 (19.26) | 47 (49.74) | 69 |
| Total | 24 | 62 | 86 |

chi2: 1.835, p: 0.1755003005, OR: 0.28485

| | < cutoff | other genes | Total |
|---|---|---|---|
| In Endothelia | 7 (4.35) | 10 (12.65) | 17 |
| Not IN Endothelia | 15 (17.65) | 54 (51.35) | 69 |
| Total | 22 | 64 | 86 |

chi2: 1.782, p: 0.1818870564, OR: 2.52000

| | > cutoff | other genes | Total |
|---|---|---|---|
| In Endothelia | 5 (4.74) | 12 (12.26) | 17 |
| Not IN Endothelia | 19 (19.26) | 50 (49.74) | 69 |
| Total | 24 | 62 | 86 |

chi2: 0.022, p: 0.8828107965, OR: 1.09649

### Five Celltypes Distribution of Average dN/dS Scores of Genes Related to biosynthetic process

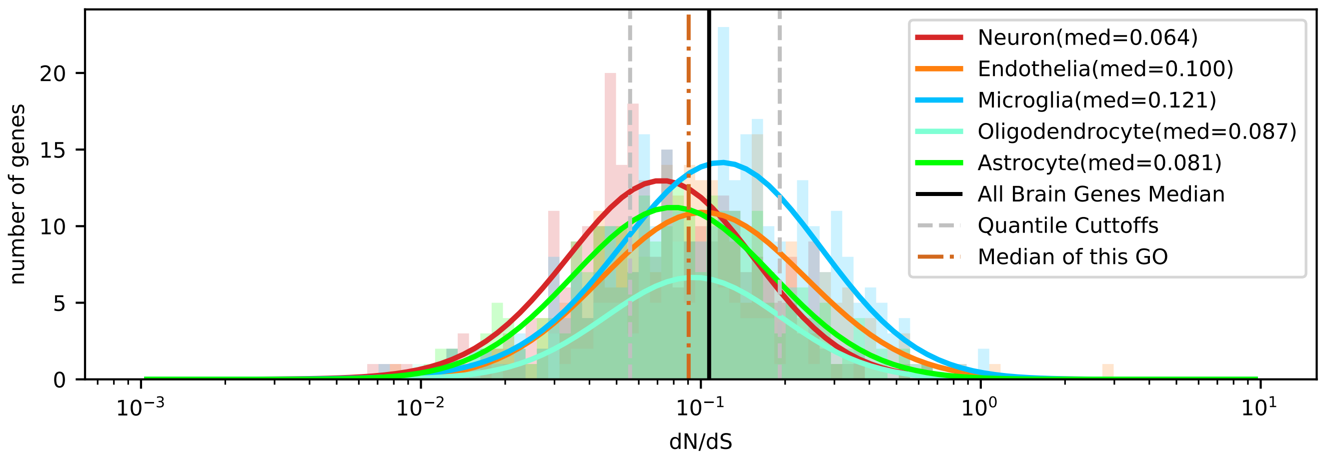


| | < cutoff | other genes | Total |
|---|---|---|---|
| In Astrocyte | 82 (69.32) | 165 (177.68) | 247 |
| Not IN Astrocyte | 259 (271.68) | 709 (696.32) | 968 |
| Total | 341 | 874 | 1215 |

chi2: 3.732, p: 0.0533641171, OR: 1.36043

| | > cutoff | other genes | Total |
|---|---|---|---|
| In Astrocyte | 35 (46.55) | 212 (200.45) | 247 |
| Not IN Astrocyte | 194 (182.45) | 774 (785.55) | 968 |
| Total | 229 | 986 | 1215 |

chi2: 4.060, p: 0.0439226797, OR: 0.65868

| | < cutoff | other genes | Total |
|---|---|---|---|
| In Microglia | 63 (89.53) | 256 (229.47) | 319 |
| Not IN Microglia | 278 (251.47) | 618 (644.53) | 896 |
| Total | 341 | 874 | 1215 |

chi2: 14.266, p: 0.0001586731, OR: 0.54707

| | > cutoff | other genes | Total |
|---|---|---|---|
| In Microglia | 85 (60.12) | 234 (258.88) | 319 |
| Not IN Microglia | 144 (168.88) | 752 (727.12) | 896 |
| Total | 229 | 986 | 1215 |

chi2: 16.513, p: 0.0000483112, OR: 1.89696

| | < cutoff | other genes | Total |
|---|---|---|---|
| In Oligodendrocyte | 29 (36.77) | 102 (94.23) | 131 |
| Not IN Oligodendrocyte | 312 (304.23) | 772 (779.77) | 1084 |
| Total | 341 | 874 | 1215 |

chi2: 2.238, p: 0.1346893191, OR: 0.70349

| | > cutoff | other genes | Total |
|---|---|---|---|
| In Oligodendrocyte | 22 (24.69) | 109 (106.31) | 131 |
| Not IN Oligodendrocyte | 207 (204.31) | 877 (879.69) | 1084 |
| Total | 229 | 986 | 1215 |

chi2: 0.268, p: 0.6043930532, OR: 0.85512

| | < cutoff | other genes | Total |
|---|---|---|---|
| In Neuron | 109 (76.34) | 163 (195.66) | 272 |
| Not IN Neuron | 232 (264.66) | 711 (678.34) | 943 |
| Total | 341 | 874 | 1215 |

chi2: 24.268, p: 0.0000008381, OR: 2.04937

| | > cutoff | other genes | Total |
|---|---|---|---|
| In Neuron | 32 (51.27) | 240 (220.73) | 272 |
| Not IN Neuron | 197 (177.73) | 746 (765.27) | 943 |
| Total | 229 | 986 | 1215 |

chi2: 10.906, p: 0.0009584479, OR: 0.50491

| | < cutoff | other genes | Total |
|---|---|---|---|
| In Endothelia | 58 (69.04) | 188 (176.96) | 246 |
| Not IN Endothelia | 283 (271.96) | 686 (697.04) | 969 |
| Total | 341 | 874 | 1215 |

chi2: 2.806, p: 0.0939272304, OR: 0.74784

| | > cutoff | other genes | Total |
|---|---|---|---|
| In Endothelia | 55 (46.37) | 191 (199.63) | 246 |
| Not IN Endothelia | 174 (182.63) | 795 (786.37) | 969 |
| Total | 229 | 986 | 1215 |

chi2: 2.205, p: 0.1375557705, OR: 1.31567

Five Celltypes Distribution of Average dN/dS Scores of Genes Related to carbohydrate metabolic process

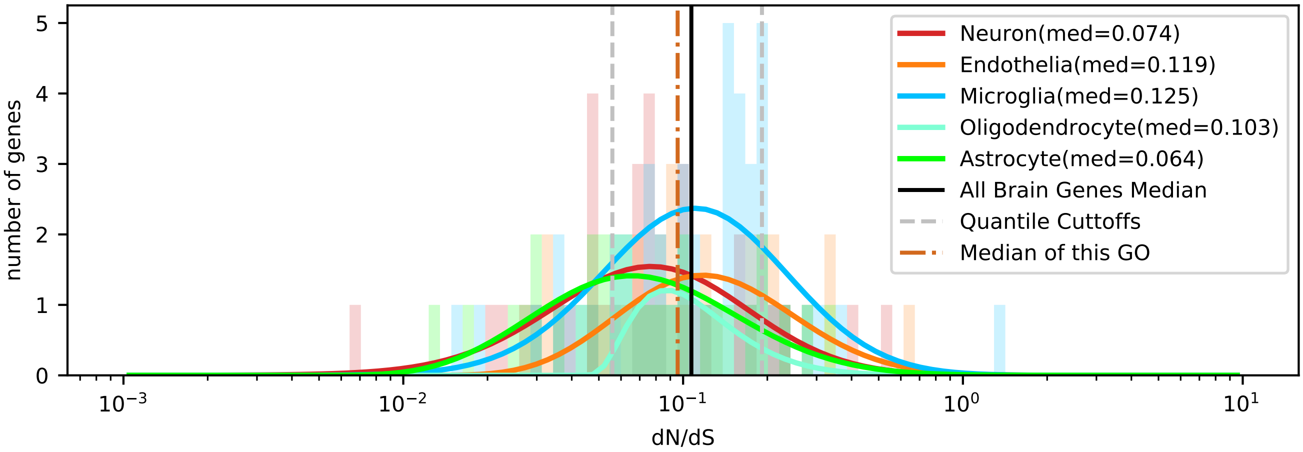


| | < cutoff | other genes | Total |
|---|---|---|---|
| In Astrocyte | 13 (7.85) | 18 (23.15) | 31 |
| Not IN Astrocyte | 26 (31.15) | 97 (91.85) | 123 |
| Total | 39 | 115 | 154 |

chi2: 4.617, p: 0.0316648716, OR: 2.69444

| | > cutoff | other genes | Total |
|---|---|---|---|
| In Astrocyte | 5 (5.23) | 26 (25.77) | 31 |
| Not IN Astrocyte | 21 (20.77) | 102 (102.23) | 123 |
| Total | 26 | 128 | 154 |

chi2: 0.020, p: 0.8864245163, OR: 0.93407

| | < cutoff | other genes | Total |
|---|---|---|---|
| In Microglia | 10 (12.41) | 39 (36.59) | 49 |
| Not IN Microglia | 29 (26.59) | 76 (78.41) | 105 |
| Total | 39 | 115 | 154 |

chi2: 0.577, p: 0.4475472402, OR: 0.67197

| | > cutoff | other genes | Total |
|---|---|---|---|
| In Microglia | 8 (8.27) | 41 (40.73) | 49 |
| Not IN Microglia | 18 (17.73) | 87 (87.27) | 105 |
| Total | 26 | 128 | 154 |

chi2: 0.011, p: 0.9164036682, OR: 0.94309

| | < cutoff | other genes | Total |
|---|---|---|---|
| In Oligodendrocyte | 0 (3.29) | 13 (9.71) | 13 |
| Not IN Oligodendrocyte | 39 (35.71) | 102 (105.29) | 141 |
| Total | 39 | 115 | 154 |

chi2: 3.464, p: 0.0627320828, OR: 0.00000

| | > cutoff | other genes | Total |
|---|---|---|---|
| In Oligodendrocyte | 1 (2.19) | 12 (10.81) | 13 |
| Not IN Oligodendrocyte | 25 (23.81) | 116 (117.19) | 141 |
| Total | 26 | 128 | 154 |

chi2: 0.289, p: 0.5908429840, OR: 0.38667

| | < cutoff | other genes | Total |
|---|---|---|---|
| In Neuron | 13 (8.61) | 21 (25.39) | 34 |
| Not IN Neuron | 26 (30.39) | 94 (89.61) | 120 |
| Total | 39 | 115 | 154 |

chi2: 3.020, p: 0.0822629294, OR: 2.23810

| | > cutoff | other genes | Total |
|---|---|---|---|
| In Neuron | 4 (5.74) | 30 (28.26) | 34 |
| Not IN Neuron | 22 (20.26) | 98 (99.74) | 120 |
| Total | 26 | 128 | 154 |

chi2: 0.414, p: 0.5200693967, OR: 0.59394

| | < cutoff | other genes | Total |
|---|---|---|---|
| In Endothelia | 3 (6.84) | 24 (20.16) | 27 |
| Not IN Endothelia | 36 (32.16) | 91 (94.84) | 127 |
| Total | 39 | 115 | 154 |

chi2: 2.646, p: 0.1038397945, OR: 0.31597

| | > cutoff | other genes | Total |
|---|---|---|---|
| In Endothelia | 8 (4.56) | 19 (22.44) | 27 |
| Not IN Endothelia | 18 (21.44) | 109 (105.56) | 127 |
| Total | 26 | 128 | 154 |

chi2: 2.769, p: 0.0960895834, OR: 2.54971

### Five Celltypes Distribution of Average dN/dS Scores of Genes Related to catabolic process

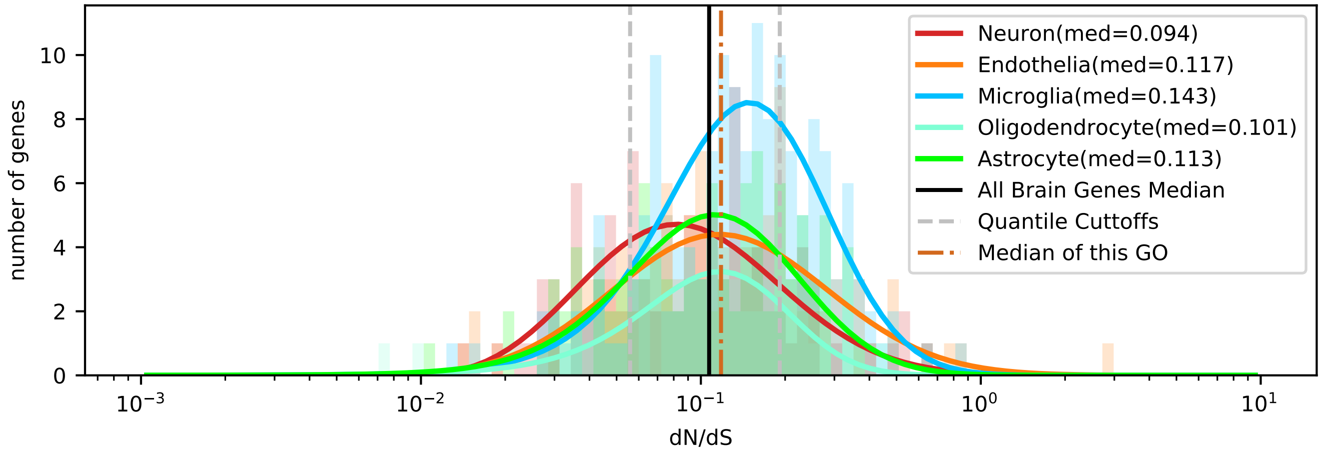


| | < cutoff | other genes | Total |
|---|---|---|---|
| In Astrocyte | 22 (19.03) | 76 (78.97) | 98 |
| Not IN Astrocyte | 78 (80.97) | 339 (336.03) | 417 |
| Total | 100 | 415 | 515 |

chi2: 0.492, p: 0.4831637179, OR: 1.25810

| | > cutoff | other genes | Total |
|---|---|---|---|
| In Astrocyte | 24 (24.36) | 74 (73.64) | 98 |
| Not IN Astrocyte | 104 (103.64) | 313 (313.36) | 417 |
| Total | 128 | 387 | 515 |

chi2: 0.001, p: 0.9704273657, OR: 0.97609

| | < cutoff | other genes | Total |
|---|---|---|---|
| In Microglia | 18 (30.49) | 139 (126.51) | 157 |
| Not IN Microglia | 82 (69.51) | 276 (288.49) | 358 |
| Total | 100 | 415 | 515 |

chi2: 8.412, p: 0.0037275481, OR: 0.43587

| | > cutoff | other genes | Total |
|---|---|---|---|
| In Microglia | 51 (39.02) | 106 (117.98) | 157 |
| Not IN Microglia | 77 (88.98) | 281 (269.02) | 358 |
| Total | 128 | 387 | 515 |

chi2: 6.464, p: 0.0110083733, OR: 1.75582

| | < cutoff | other genes | Total |
|---|---|---|---|
| In Oligodendrocyte | 11 (10.68) | 44 (44.32) | 55 |
| Not IN Oligodendrocyte | 89 (89.32) | 371 (370.68) | 460 |
| Total | 100 | 415 | 515 |

chi2: 0.004, p: 0.9483467791, OR: 1.04213

| | > cutoff | other genes | Total |
|---|---|---|---|
| In Oligodendrocyte | 10 (13.67) | 45 (41.33) | 55 |
| Not IN Oligodendrocyte | 118 (114.33) | 342 (345.67) | 460 |
| Total | 128 | 387 | 515 |

chi2: 1.095, p: 0.2953337544, OR: 0.64407

| | < cutoff | other genes | Total |
|---|---|---|---|
| In Neuron | 30 (19.61) | 71 (81.39) | 101 |
| Not IN Neuron | 70 (80.39) | 344 (333.61) | 414 |
| Total | 100 | 415 | 515 |

chi2: 7.697, p: 0.0055324631, OR: 2.07646

| | > cutoff | other genes | Total |
|---|---|---|---|
| In Neuron | 15 (25.10) | 86 (75.90) | 101 |
| Not IN Neuron | 113 (102.90) | 301 (311.10) | 414 |
| Total | 128 | 387 | 515 |

chi2: 6.081, p: 0.0136632583, OR: 0.46460

| | < cutoff | other genes | Total |
|---|---|---|---|
| In Endothelia | 19 (20.19) | 85 (83.81) | 104 |
| Not IN Endothelia | 81 (79.81) | 330 (331.19) | 411 |
| Total | 100 | 415 | 515 |

chi2: 0.037, p: 0.8472507955, OR: 0.91068

| | > cutoff | other genes | Total |
|---|---|---|---|
| In Endothelia | 28 (25.85) | 76 (78.15) | 104 |
| Not IN Endothelia | 100 (102.15) | 311 (308.85) | 411 |
| Total | 128 | 387 | 515 |

chi2: 0.176, p: 0.6748875108, OR: 1.14579

### Five Celltypes Distribution of Average dN/dS Scores of Genes Related to cell adhesion

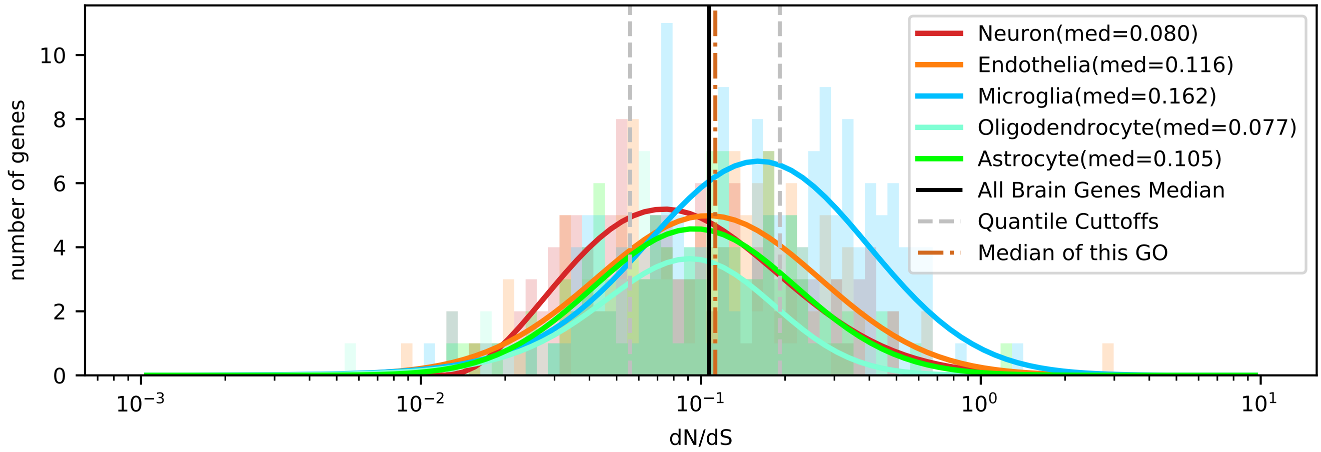


| | < cutoff | other genes | Total |
|---|---|---|---|
| In Astrocyte | 28 (26.33) | 74 (75.67) | 102 |
| Not IN Astrocyte | 123 (124.67) | 360 (358.33) | 483 |
| Total | 151 | 434 | 585 |

chi2: 0.085, p: 0.7704430824, OR: 1.10745

| | > cutoff | other genes | Total |
|---|---|---|---|
| In Astrocyte | 18 (28.25) | 84 (73.75) | 102 |
| Not IN Astrocyte | 144 (133.75) | 339 (349.25) | 483 |
| Total | 162 | 423 | 585 |

chi2: 5.633, p: 0.0176265261, OR: 0.50446

| | < cutoff | other genes | Total |
|---|---|---|---|
| In Microglia | 29 (43.36) | 139 (124.64) | 168 |
| Not IN Microglia | 122 (107.64) | 295 (309.36) | 417 |
| Total | 151 | 434 | 585 |

chi2: 8.382, p: 0.0037898958, OR: 0.50448

| | > cutoff | other genes | Total |
|---|---|---|---|
| In Microglia | 76 (46.52) | 92 (121.48) | 168 |
| Not IN Microglia | 86 (115.48) | 331 (301.52) | 417 |
| Total | 162 | 423 | 585 |

chi2: 35.016, p: 0.0000000033, OR: 3.17947

| | < cutoff | other genes | Total |
|---|---|---|---|
| In Oligodendrocyte | 21 (18.58) | 51 (53.42) | 72 |
| Not IN Oligodendrocyte | 130 (132.42) | 383 (380.58) | 513 |
| Total | 151 | 434 | 585 |

chi2: 0.303, p: 0.5817380958, OR: 1.21312

| | > cutoff | other genes | Total |
|---|---|---|---|
| In Oligodendrocyte | 10 (19.94) | 62 (52.06) | 72 |
| Not IN Oligodendrocyte | 152 (142.06) | 361 (370.94) | 513 |
| Total | 162 | 423 | 585 |

chi2: 7.046, p: 0.0079426532, OR: 0.38306

| | < cutoff | other genes | Total |
|---|---|---|---|
| In Neuron | 43 (30.72) | 76 (88.28) | 119 |
| Not IN Neuron | 108 (120.28) | 358 (345.72) | 466 |
| Total | 151 | 434 | 585 |

chi2: 7.650, p: 0.0056786494, OR: 1.87549

| | > cutoff | other genes | Total |
|---|---|---|---|
| In Neuron | 23 (32.95) | 96 (86.05) | 119 |
| Not IN Neuron | 139 (129.05) | 327 (336.95) | 466 |
| Total | 162 | 423 | 585 |

chi2: 4.709, p: 0.0300112121, OR: 0.56362

| | < cutoff | other genes | Total |
|---|---|---|---|
| In Endothelia | 30 (32.01) | 94 (91.99) | 124 |
| Not IN Endothelia | 121 (118.99) | 340 (342.01) | 461 |
| Total | 151 | 434 | 585 |

chi2: 0.121, p: 0.7275834967, OR: 0.89678

| | > cutoff | other genes | Total |
|---|---|---|---|
| In Endothelia | 35 (34.34) | 89 (89.66) | 124 |
| Not IN Endothelia | 127 (127.66) | 334 (333.34) | 461 |
| Total | 162 | 423 | 585 |

chi2: 0.001, p: 0.9708684037, OR: 1.03424

Five Celltypes Distribution of Average dN/dS Scores of Genes Related to cell cycle

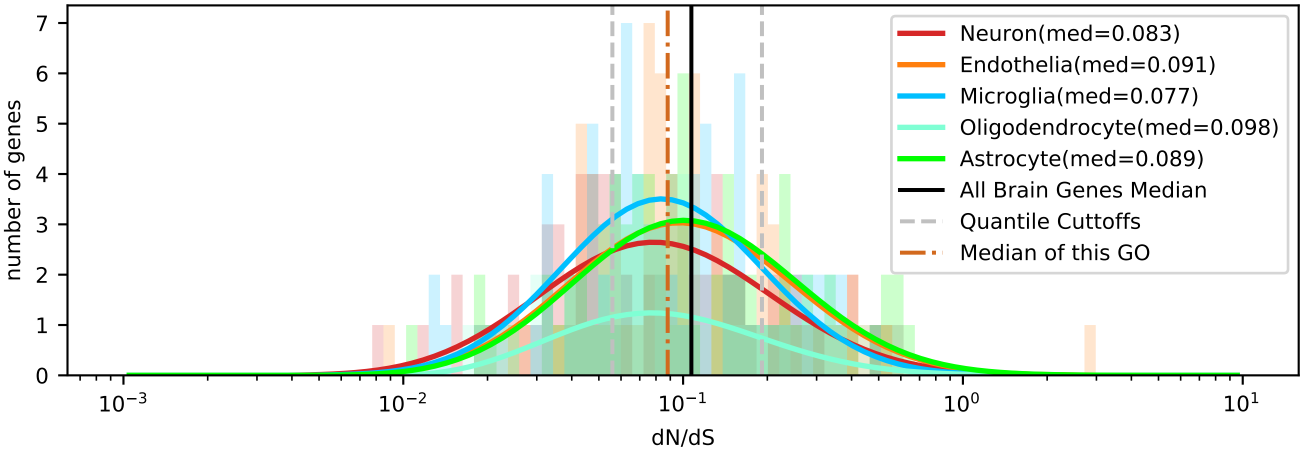


| | < cutoff | other genes | Total |
|---|---|---|---|
| In Astrocyte | 16 (21.66) | 59 (53.34) | 75 |
| Not IN Astrocyte | 77 (71.34) | 170 (175.66) | 247 |
| Total | 93 | 229 | 322 |

chi2: 2.254, p: 0.1332304909, OR: 0.59872

| | > cutoff | other genes | Total |
|---|---|---|---|
| In Astrocyte | 19 (15.61) | 56 (59.39) | 75 |
| Not IN Astrocyte | 48 (51.39) | 199 (195.61) | 247 |
| Total | 67 | 255 | 322 |

chi2: 0.884, p: 0.3471849257, OR: 1.40662

| | < cutoff | other genes | Total |
|---|---|---|---|
| In Microglia | 22 (22.53) | 56 (55.47) | 78 |
| Not IN Microglia | 71 (70.47) | 173 (173.53) | 244 |
| Total | 93 | 229 | 322 |

chi2: 0.000, p: 0.9935996398, OR: 0.95724

| | > cutoff | other genes | Total |
|---|---|---|---|
| In Microglia | 13 (16.23) | 65 (61.77) | 78 |
| Not IN Microglia | 54 (50.77) | 190 (193.23) | 244 |
| Total | 67 | 255 | 322 |

chi2: 0.765, p: 0.3817284146, OR: 0.70370

| | < cutoff | other genes | Total |
|---|---|---|---|
| In Oligodendrocyte | 9 (8.09) | 19 (19.91) | 28 |
| Not IN Oligodendrocyte | 84 (84.91) | 210 (209.09) | 294 |
| Total | 93 | 229 | 322 |

chi2: 0.032, p: 0.8569587808, OR: 1.18421

| | > cutoff | other genes | Total |
|---|---|---|---|
| In Oligodendrocyte | 4 (5.83) | 24 (22.17) | 28 |
| Not IN Oligodendrocyte | 63 (61.17) | 231 (232.83) | 294 |
| Total | 67 | 255 | 322 |

chi2: 0.417, p: 0.5182185478, OR: 0.61111

| | < cutoff | other genes | Total |
|---|---|---|---|
| In Neuron | 28 (19.35) | 39 (47.65) | 67 |
| Not IN Neuron | 65 (73.65) | 190 (181.35) | 255 |
| Total | 93 | 229 | 322 |

chi2: 6.093, p: 0.0135698046, OR: 2.09862

| | > cutoff | other genes | Total |
|---|---|---|---|
| In Neuron | 13 (13.94) | 54 (53.06) | 67 |
| Not IN Neuron | 54 (53.06) | 201 (201.94) | 255 |
| Total | 67 | 255 | 322 |

chi2: 0.022, p: 0.8814412237, OR: 0.89609

| | < cutoff | other genes | Total |
|---|---|---|---|
| In Endothelia | 18 (21.37) | 56 (52.63) | 74 |
| Not IN Endothelia | 75 (71.63) | 173 (176.37) | 248 |
| Total | 93 | 229 | 322 |

chi2: 0.705, p: 0.4011366863, OR: 0.74143

| | > cutoff | other genes | Total |
|---|---|---|---|
| In Endothelia | 18 (15.40) | 56 (58.60) | 74 |
| Not IN Endothelia | 49 (51.60) | 199 (196.40) | 248 |
| Total | 67 | 255 | 322 |

chi2: 0.471, p: 0.4926699803, OR: 1.30539

### Five Celltypes Distribution of Average dN/dS Scores of Genes Related to cell death

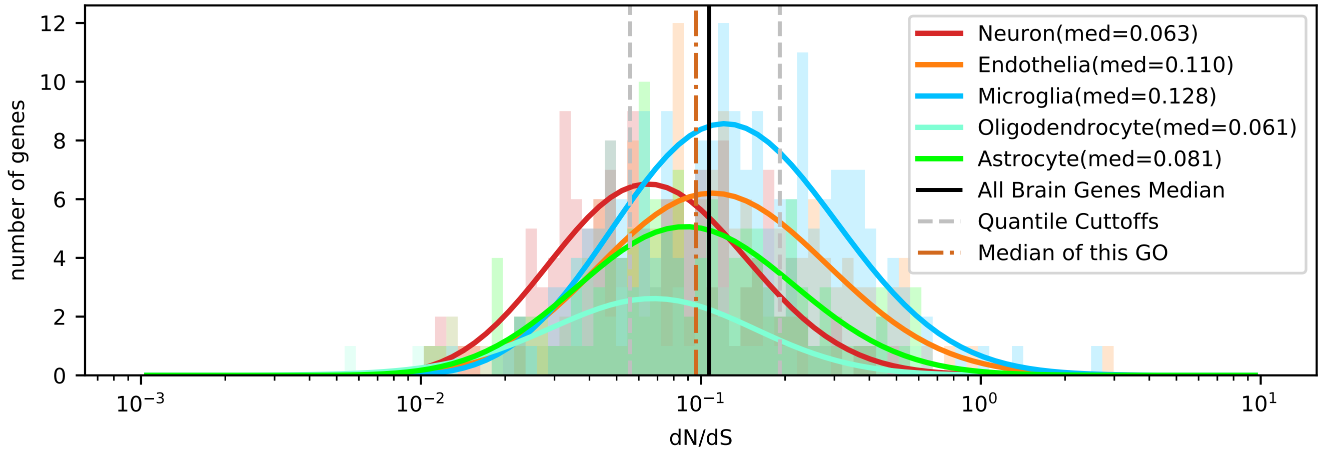


| | < cutoff | other genes | Total |
|---|---|---|---|
| In Astrocyte | 30 (32.95) | 90 (87.05) | 120 |
| Not IN Astrocyte | 157 (154.05) | 404 (406.95) | 561 |
| Total | 187 | 494 | 681 |

chi2: 0.305, p: 0.5806307850, OR: 0.85775

| | > cutoff | other genes | Total |
|---|---|---|---|
| In Astrocyte | 25 (28.02) | 95 (91.98) | 120 |
| Not IN Astrocyte | 134 (130.98) | 427 (430.02) | 561 |
| Total | 159 | 522 | 681 |

chi2: 0.358, p: 0.5494686467, OR: 0.83857

| | < cutoff | other genes | Total |
|---|---|---|---|
| In Microglia | 40 (56.84) | 167 (150.16) | 207 |
| Not IN Microglia | 147 (130.16) | 327 (343.84) | 474 |
| Total | 187 | 494 | 681 |

chi2: 9.305, p: 0.0022856471, OR: 0.53281

| | > cutoff | other genes | Total |
|---|---|---|---|
| In Microglia | 71 (48.33) | 136 (158.67) | 207 |
| Not IN Microglia | 88 (110.67) | 386 (363.33) | 474 |
| Total | 159 | 522 | 681 |

chi2: 19.061, p: 0.0000126620, OR: 2.28994

| | < cutoff | other genes | Total |
|---|---|---|---|
| In Oligodendrocyte | 26 (16.48) | 34 (43.52) | 60 |
| Not IN Oligodendrocyte | 161 (170.52) | 460 (450.48) | 621 |
| Total | 187 | 494 | 681 |

chi2: 7.472, p: 0.0062657940, OR: 2.18487

| | > cutoff | other genes | Total |
|---|---|---|---|
| In Oligodendrocyte | 5 (14.01) | 55 (45.99) | 60 |
| Not IN Oligodendrocyte | 154 (144.99) | 467 (476.01) | 621 |
| Total | 159 | 522 | 681 |

chi2: 7.394, p: 0.0065448534, OR: 0.27568

| | < cutoff | other genes | Total |
|---|---|---|---|
| In Neuron | 59 (38.17) | 80 (100.83) | 139 |
| Not IN Neuron | 128 (148.83) | 414 (393.17) | 542 |
| Total | 187 | 494 | 681 |

chi2: 18.758, p: 0.0000148416, OR: 2.38535

| | > cutoff | other genes | Total |
|---|---|---|---|
| In Neuron | 15 (32.45) | 124 (106.55) | 139 |
| Not IN Neuron | 144 (126.55) | 398 (415.45) | 542 |
| Total | 159 | 522 | 681 |

chi2: 14.517, p: 0.0001388659, OR: 0.33434

| | < cutoff | other genes | Total |
|---|---|---|---|
| In Endothelia | 32 (42.56) | 123 (112.44) | 155 |
| Not IN Endothelia | 155 (144.44) | 371 (381.56) | 526 |
| Total | 187 | 494 | 681 |

chi2: 4.246, p: 0.0393476022, OR: 0.62271

| | > cutoff | other genes | Total |
|---|---|---|---|
| In Endothelia | 43 (36.19) | 112 (118.81) | 155 |
| Not IN Endothelia | 116 (122.81) | 410 (403.19) | 526 |
| Total | 159 | 522 | 681 |

chi2: 1.859, p: 0.1727825709, OR: 1.35699

### Five Celltypes Distribution of Average dN/dS Scores of Genes Related to cell differentiation

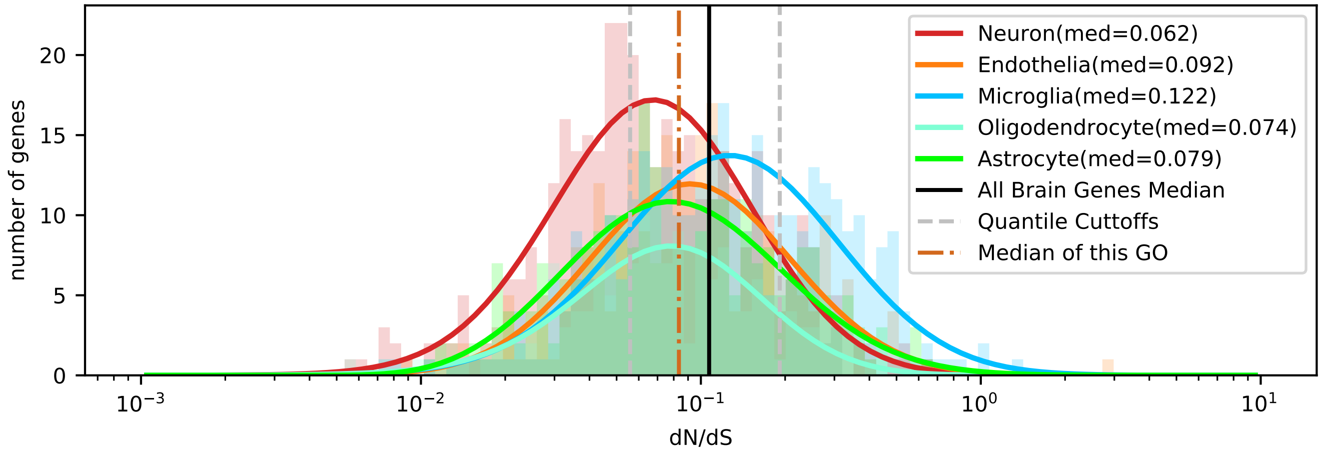


| | < cutoff | other genes | Total |
|---|---|---|---|
| In Astrocyte | 86 (82.15) | 172 (175.85) | 258 |
| Not IN Astrocyte | 362 (365.85) | 787 (783.15) | 1149 |
| Total | 448 | 959 | 1407 |

chi2: 0.246, p: 0.6202304323, OR: 1.08702

| | > cutoff | other genes | Total |
|---|---|---|---|
| In Astrocyte | 48 (49.69) | 210 (208.31) | 258 |
| Not IN Astrocyte | 223 (221.31) | 926 (927.69) | 1149 |
| Total | 271 | 1136 | 1407 |

chi2: 0.043, p: 0.8349065401, OR: 0.94914

| | < cutoff | other genes | Total |
|---|---|---|---|
| In Microglia | 65 (106.67) | 270 (228.33) | 335 |
| Not IN Microglia | 383 (341.33) | 689 (730.67) | 1072 |
| Total | 448 | 959 | 1407 |

chi2: 30.594, p: 0.0000000318, OR: 0.43308

| | > cutoff | other genes | Total |
|---|---|---|---|
| In Microglia | 115 (64.52) | 220 (270.48) | 335 |
| Not IN Microglia | 156 (206.48) | 916 (865.52) | 1072 |
| Total | 271 | 1136 | 1407 |

chi2: 62.925, p: 0.0000000000, OR: 3.06935

| | < cutoff | other genes | Total |
|---|---|---|---|
| In Oligodendrocyte | 58 (52.86) | 108 (113.14) | 166 |
| Not IN Oligodendrocyte | 390 (395.14) | 851 (845.86) | 1241 |
| Total | 448 | 959 | 1407 |

chi2: 0.679, p: 0.4099999780, OR: 1.17184

| | > cutoff | other genes | Total |
|---|---|---|---|
| In Oligodendrocyte | 18 (31.97) | 148 (134.03) | 166 |
| Not IN Oligodendrocyte | 253 (239.03) | 988 (1001.97) | 1241 |
| Total | 271 | 1136 | 1407 |

chi2: 7.972, p: 0.0047498894, OR: 0.47495

| | < cutoff | other genes | Total |
|---|---|---|---|
| In Neuron | 169 (121.95) | 214 (261.05) | 383 |
| Not IN Neuron | 279 (326.05) | 745 (697.95) | 1024 |
| Total | 448 | 959 | 1407 |

chi2: 35.820, p: 0.0000000022, OR: 2.10875

| | > cutoff | other genes | Total |
|---|---|---|---|
| In Neuron | 42 (73.77) | 341 (309.23) | 383 |
| Not IN Neuron | 229 (197.23) | 795 (826.77) | 1024 |
| Total | 271 | 1136 | 1407 |

chi2: 22.556, p: 0.0000020409, OR: 0.42759

| | < cutoff | other genes | Total |
|---|---|---|---|
| In Endothelia | 70 (84.38) | 195 (180.62) | 265 |
| Not IN Endothelia | 378 (363.62) | 764 (778.38) | 1142 |
| Total | 448 | 959 | 1407 |

chi2: 4.126, p: 0.0422278497, OR: 0.72555

| | > cutoff | other genes | Total |
|---|---|---|---|
| In Endothelia | 48 (51.04) | 217 (213.96) | 265 |
| Not IN Endothelia | 223 (219.96) | 919 (922.04) | 1142 |
| Total | 271 | 1136 | 1407 |

chi2: 0.193, p: 0.6603761401, OR: 0.91157

#### Five Celltypes Distribution of Average dN/dS Scores of Genes Related to cell division

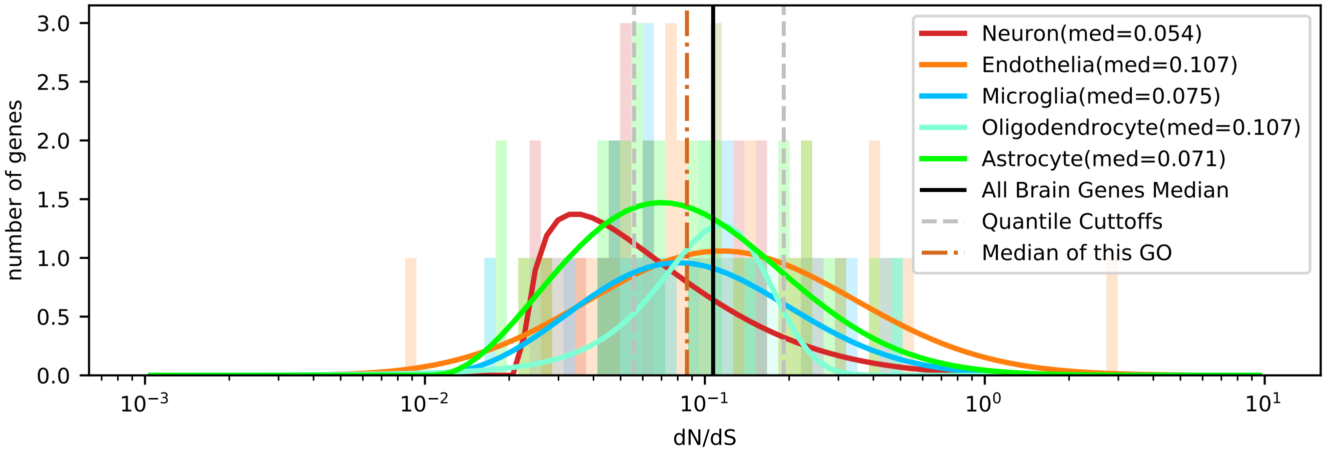


| | < cutoff | other genes | Total |
|---|---|---|---|
| In Astrocyte | 11 (10.94) | 24 (24.06) | 35 |
| Not IN Astrocyte | 29 (29.06) | 64 (63.94) | 93 |
| Total | 40 | 88 | 128 |

chi2: 0.035, p: 0.8515239858, OR: 1.01149

| | > cutoff | other genes | Total |
|---|---|---|---|
| In Astrocyte | 8 (6.84) | 27 (28.16) | 35 |
| Not IN Astrocyte | 17 (18.16) | 76 (74.84) | 93 |
| Total | 25 | 103 | 128 |

chi2: 0.110, p: 0.7397612472, OR: 1.32462

| | < cutoff | other genes | Total |
|---|---|---|---|
| In Microglia | 6 (6.88) | 16 (15.12) | 22 |
| Not IN Microglia | 34 (33.12) | 72 (72.88) | 106 |
| Total | 40 | 88 | 128 |

chi2: 0.036, p: 0.8496662760, OR: 0.79412

| | > cutoff | other genes | Total |
|---|---|---|---|
| In Microglia | 4 (4.30) | 18 (17.70) | 22 |
| Not IN Microglia | 21 (20.70) | 85 (85.30) | 106 |
| Total | 25 | 103 | 128 |

chi2: 0.014, p: 0.9044515807, OR: 0.89947

| | < cutoff | other genes | Total |
|---|---|---|---|
| In Oligodendrocyte | 3 (5.31) | 14 (11.69) | 17 |
| Not IN Oligodendrocyte | 37 (34.69) | 74 (76.31) | 111 |
| Total | 40 | 88 | 128 |

chi2: 1.037, p: 0.3084686341, OR: 0.42857

| | > cutoff | other genes | Total |
|---|---|---|---|
| In Oligodendrocyte | 1 (3.32) | 16 (13.68) | 17 |
| Not IN Oligodendrocyte | 24 (21.68) | 87 (89.32) | 111 |
| Total | 25 | 103 | 128 |

chi2: 1.430, p: 0.2317447491, OR: 0.22656

| | < cutoff | other genes | Total |
|---|---|---|---|
| In Neuron | 13 (7.19) | 10 (15.81) | 23 |
| Not IN Neuron | 27 (32.81) | 78 (72.19) | 105 |
| Total | 40 | 88 | 128 |

chi2: 6.963, p: 0.0083233570, OR: 3.75556

| | > cutoff | other genes | Total |
|---|---|---|---|
| In Neuron | 2 (4.49) | 21 (18.51) | 23 |
| Not IN Neuron | 23 (20.51) | 82 (84.49) | 105 |
| Total | 25 | 103 | 128 |

chi2: 1.338, p: 0.2473108677, OR: 0.33954

| | < cutoff | other genes | Total |
|---|---|---|---|
| In Endothelia | 7 (9.69) | 24 (21.31) | 31 |
| Not IN Endothelia | 33 (30.31) | 64 (66.69) | 97 |
| Total | 40 | 88 | 128 |

chi2: 0.948, p: 0.3302057396, OR: 0.56566

| | > cutoff | other genes | Total |
|---|---|---|---|
| In Endothelia | 10 (6.05) | 21 (24.95) | 31 |
| Not IN Endothelia | 15 (18.95) | 82 (78.05) | 97 |
| Total | 25 | 103 | 128 |

chi2: 3.215, p: 0.0729676734, OR: 2.60317

### Five Celltypes Distribution of Average dN/dS Scores of Genes Related to cell junction organization

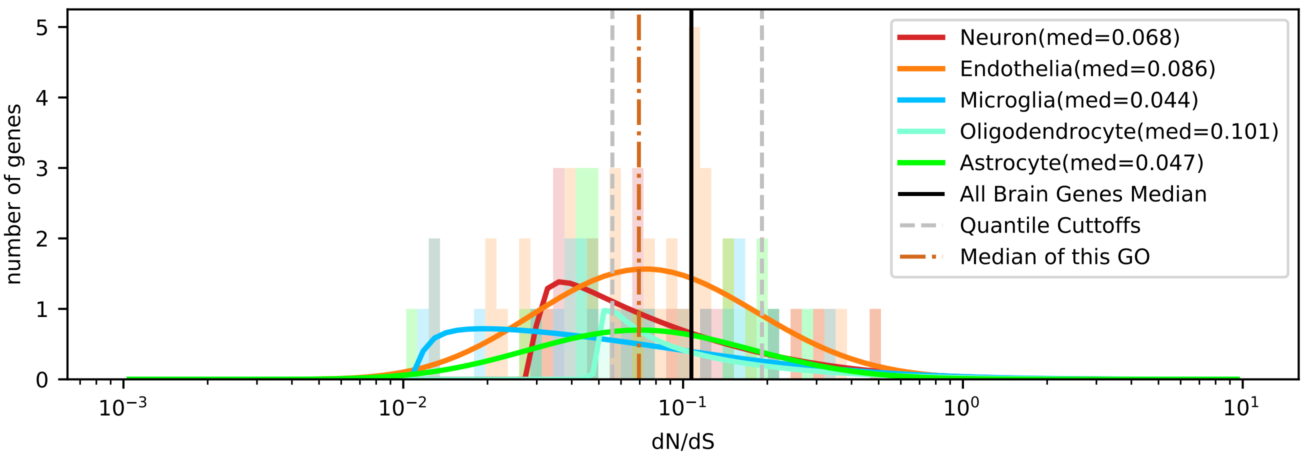


| | < cutoff | other genes | Total |
|---|---|---|---|
| In Astrocyte | 9 (6.54) | 8 (10.46) | 17 |
| Not IN Astrocyte | 31 (33.46) | 56 (53.54) | 87 |
| Total | 40 | 64 | 104 |

chi2: 1.143, p: 0.2849975071, OR: 2.03226

| | > cutoff | other genes | Total |
|---|---|---|---|
| In Astrocyte | 2 (2.45) | 15 (14.55) | 17 |
| Not IN Astrocyte | 13 (12.55) | 74 (74.45) | 87 |
| Total | 15 | 89 | 104 |

chi2: 0.001, p: 0.9710527901, OR: 0.75897

| | < cutoff | other genes | Total |
|---|---|---|---|
| In Microglia | 11 (7.31) | 8 (11.69) | 19 |
| Not IN Microglia | 29 (32.69) | 56 (52.31) | 85 |
| Total | 40 | 64 | 104 |

chi2: 2.773, p: 0.0958865458, OR: 2.65517

| | > cutoff | other genes | Total |
|---|---|---|---|
| In Microglia | 2 (2.74) | 17 (16.26) | 19 |
| Not IN Microglia | 13 (12.26) | 72 (72.74) | 85 |
| Total | 15 | 89 | 104 |

chi2: 0.030, p: 0.8621546771, OR: 0.65158

| | < cutoff | other genes | Total |
|---|---|---|---|
| In Oligodendrocyte | 1 (3.46) | 8 (5.54) | 9 |
| Not IN Oligodendrocyte | 39 (36.54) | 56 (58.46) | 95 |
| Total | 40 | 64 | 104 |

chi2: 1.977, p: 0.1596679128, OR: 0.17949

| | > cutoff | other genes | Total |
|---|---|---|---|
| In Oligodendrocyte | 2 (1.30) | 7 (7.70) | 9 |
| Not IN Oligodendrocyte | 13 (13.70) | 82 (81.30) | 95 |
| Total | 15 | 89 | 104 |

chi2: 0.040, p: 0.8411264158, OR: 1.80220

| | < cutoff | other genes | Total |
|---|---|---|---|
| In Neuron | 8 (8.08) | 13 (12.92) | 21 |
| Not IN Neuron | 32 (31.92) | 51 (51.08) | 83 |
| Total | 40 | 64 | 104 |

chi2: 0.045, p: 0.8317773750, OR: 0.98077

| | > cutoff | other genes | Total |
|---|---|---|---|
| In Neuron | 4 (3.03) | 17 (17.97) | 21 |
| Not IN Neuron | 11 (11.97) | 72 (71.03) | 83 |
| Total | 15 | 89 | 104 |

chi2: 0.107, p: 0.7432257981, OR: 1.54011

| | < cutoff | other genes | Total |
|---|---|---|---|
| In Endothelia | 11 (14.62) | 27 (23.38) | 38 |
| Not IN Endothelia | 29 (25.38) | 37 (40.62) | 66 |
| Total | 40 | 64 | 104 |

chi2: 1.700, p: 0.1922332346, OR: 0.51980

| | > cutoff | other genes | Total |
|---|---|---|---|
| In Endothelia | 5 (5.48) | 33 (32.52) | 38 |
| Not IN Endothelia | 10 (9.52) | 56 (56.48) | 66 |
| Total | 15 | 89 | 104 |

chi2: 0.000, p: 0.9911064872, OR: 0.84848

### Five Celltypes Distribution of Average dN/dS Scores of Genes Related to cell morphogenesis

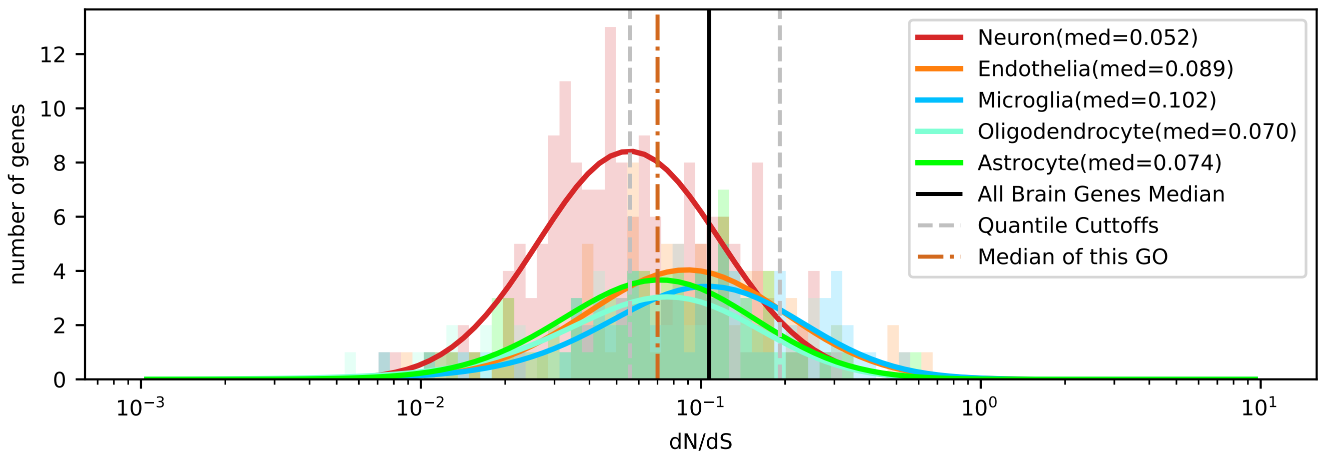


| | < cutoff | other genes | Total |
|---|---|---|---|
| In Astrocyte | 29 (31.27) | 50 (47.73) | 79 |
| Not IN Astrocyte | 161 (158.73) | 240 (242.27) | 401 |
| Total | 190 | 290 | 480 |

chi2: 0.199, p: 0.6557877332, OR: 0.86460

| | > cutoff | other genes | Total |
|---|---|---|---|
| In Astrocyte | 5 (8.56) | 74 (70.44) | 79 |
| Not IN Astrocyte | 47 (43.44) | 354 (357.56) | 401 |
| Total | 52 | 428 | 480 |

chi2: 1.467, p: 0.2257957626, OR: 0.50891

| | < cutoff | other genes | Total |
|---|---|---|---|
| In Microglia | 21 (30.08) | 55 (45.92) | 76 |
| Not IN Microglia | 169 (159.92) | 235 (244.08) | 404 |
| Total | 190 | 290 | 480 |

chi2: 4.816, p: 0.0281962286, OR: 0.53093

| | > cutoff | other genes | Total |
|---|---|---|---|
| In Microglia | 18 (8.23) | 58 (67.77) | 76 |
| Not IN Microglia | 34 (43.77) | 370 (360.23) | 404 |
| Total | 52 | 428 | 480 |

chi2: 13.897, p: 0.0001930812, OR: 3.37728

| | < cutoff | other genes | Total |
|---|---|---|---|
| In Oligodendrocyte | 25 (26.52) | 42 (40.48) | 67 |
| Not IN Oligodendrocyte | 165 (163.48) | 248 (249.52) | 413 |
| Total | 190 | 290 | 480 |

chi2: 0.076, p: 0.7833669347, OR: 0.89466

| | > cutoff | other genes | Total |
|---|---|---|---|
| In Oligodendrocyte | 7 (7.26) | 60 (59.74) | 67 |
| Not IN Oligodendrocyte | 45 (44.74) | 368 (368.26) | 413 |
| Total | 52 | 428 | 480 |

chi2: 0.010, p: 0.9184311982, OR: 0.95407

| | < cutoff | other genes | Total |
|---|---|---|---|
| In Neuron | 92 (66.90) | 77 (102.10) | 169 |
| Not IN Neuron | 98 (123.10) | 213 (187.90) | 311 |
| Total | 190 | 290 | 480 |

chi2: 23.118, p: 0.0000015239, OR: 2.59687

| | > cutoff | other genes | Total |
|---|---|---|---|
| In Neuron | 9 (18.31) | 160 (150.69) | 169 |
| Not IN Neuron | 43 (33.69) | 268 (277.31) | 311 |
| Total | 52 | 428 | 480 |

chi2: 7.335, p: 0.0067613961, OR: 0.35058

| | < cutoff | other genes | Total |
|---|---|---|---|
| In Endothelia | 23 (35.23) | 66 (53.77) | 89 |
| Not IN Endothelia | 167 (154.77) | 224 (236.23) | 391 |
| Total | 190 | 290 | 480 |

chi2: 7.935, p: 0.0048491185, OR: 0.46743

| | > cutoff | other genes | Total |
|---|---|---|---|
| In Endothelia | 13 (9.64) | 76 (79.36) | 89 |
| Not IN Endothelia | 39 (42.36) | 352 (348.64) | 391 |
| Total | 52 | 428 | 480 |

chi2: 1.167, p: 0.2800934155, OR: 1.54386

### Five Celltypes Distribution of Average dN/dS Scores of Genes Related to cell motility

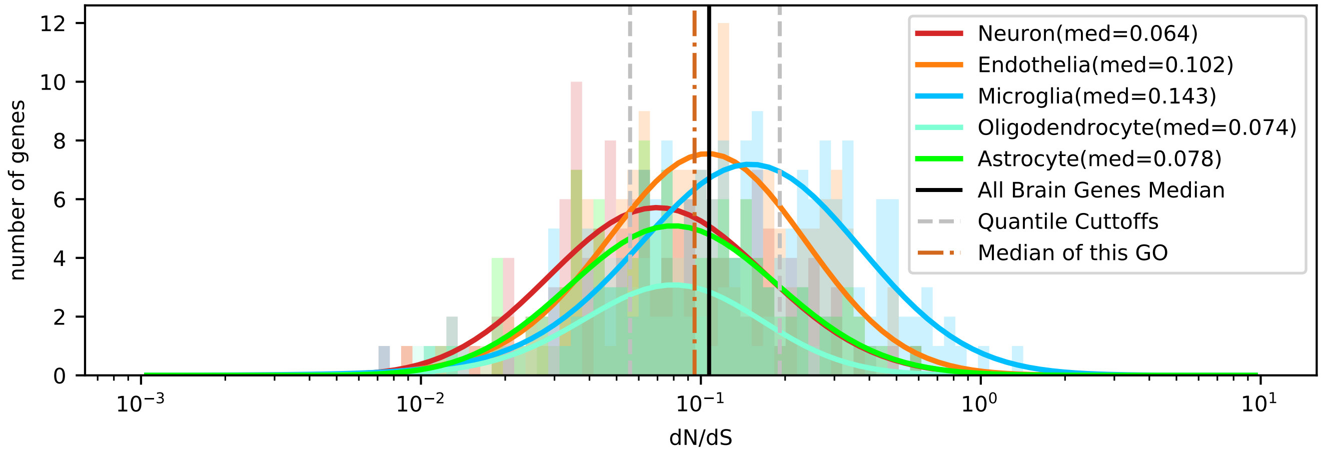


| | < cutoff | other genes | Total |
|---|---|---|---|
| In Astrocyte | 39 (32.69) | 77 (83.31) | 116 |
| Not IN Astrocyte | 147 (153.31) | 397 (390.69) | 544 |
| Total | 186 | 474 | 660 |

chi2: 1.744, p: 0.1866563849, OR: 1.36788

| | > cutoff | other genes | Total |
|---|---|---|---|
| In Astrocyte | 18 (28.47) | 98 (87.53) | 116 |
| Not IN Astrocyte | 144 (133.53) | 400 (410.47) | 544 |
| Total | 162 | 498 | 660 |

chi2: 5.616, p: 0.0177931066, OR: 0.51020

| | < cutoff | other genes | Total |
|---|---|---|---|
| In Microglia | 29 (50.73) | 151 (129.27) | 180 |
| Not IN Microglia | 157 (135.27) | 323 (344.73) | 480 |
| Total | 186 | 474 | 660 |

chi2: 17.007, p: 0.0000372519, OR: 0.39512

| | > cutoff | other genes | Total |
|---|---|---|---|
| In Microglia | 72 (44.18) | 108 (135.82) | 180 |
| Not IN Microglia | 90 (117.82) | 390 (362.18) | 480 |
| Total | 162 | 498 | 660 |

chi2: 30.781, p: 0.0000000289, OR: 2.88889

| | < cutoff | other genes | Total |
|---|---|---|---|
| In Oligodendrocyte | 21 (17.47) | 41 (44.53) | 62 |
| Not IN Oligodendrocyte | 165 (168.53) | 433 (429.47) | 598 |
| Total | 186 | 474 | 660 |

chi2: 0.806, p: 0.3692968465, OR: 1.34412

| | > cutoff | other genes | Total |
|---|---|---|---|
| In Oligodendrocyte | 7 (15.22) | 55 (46.78) | 62 |
| Not IN Oligodendrocyte | 155 (146.78) | 443 (451.22) | 598 |
| Total | 162 | 498 | 660 |

chi2: 5.726, p: 0.0167188731, OR: 0.36375

| | < cutoff | other genes | Total |
|---|---|---|---|
| In Neuron | 57 (37.76) | 77 (96.24) | 134 |
| Not IN Neuron | 129 (148.24) | 397 (377.76) | 526 |
| Total | 186 | 474 | 660 |

chi2: 16.241, p: 0.0000557653, OR: 2.27816

| | > cutoff | other genes | Total |
|---|---|---|---|
| In Neuron | 21 (32.89) | 113 (101.11) | 134 |
| Not IN Neuron | 141 (129.11) | 385 (396.89) | 526 |
| Total | 162 | 498 | 660 |

chi2: 6.560, p: 0.0104287656, OR: 0.50744

| | < cutoff | other genes | Total |
|---|---|---|---|
| In Endothelia | 40 (47.35) | 128 (120.65) | 168 |
| Not IN Endothelia | 146 (138.65) | 346 (353.35) | 492 |
| Total | 186 | 474 | 660 |

chi2: 1.849, p: 0.1739325481, OR: 0.74058

| | > cutoff | other genes | Total |
|---|---|---|---|
| In Endothelia | 44 (41.24) | 124 (126.76) | 168 |
| Not IN Endothelia | 118 (120.76) | 374 (371.24) | 492 |
| Total | 162 | 498 | 660 |

chi2: 0.221, p: 0.6383431595, OR: 1.12466

### Five Celltypes Distribution of Average dN/dS Scores of Genes Related to cell proliferation

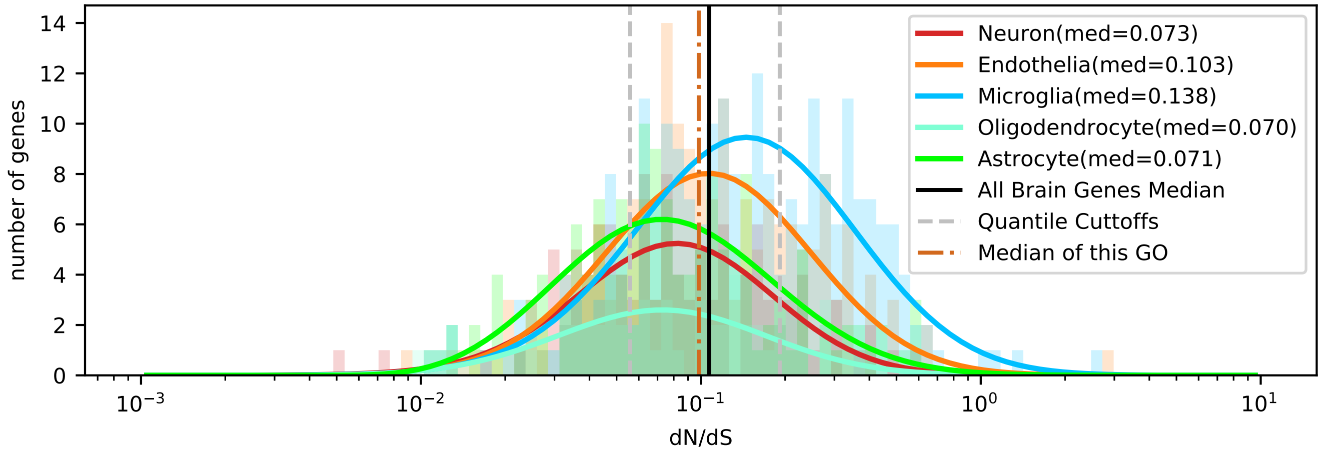


| | < cutoff | other genes | Total |
|---|---|---|---|
| In Astrocyte | 51 (37.34) | 95 (108.66) | 146 |
| Not IN Astrocyte | 137 (150.66) | 452 (438.34) | 589 |
| Total | 188 | 547 | 735 |

chi2: 7.771, p: 0.0053088947, OR: 1.77119

| | > cutoff | other genes | Total |
|---|---|---|---|
| In Astrocyte | 22 (36.75) | 124 (109.25) | 146 |
| Not IN Astrocyte | 163 (148.25) | 426 (440.75) | 589 |
| Total | 185 | 550 | 735 |

chi2: 9.213, p: 0.0024034592, OR: 0.46368

| | < cutoff | other genes | Total |
|---|---|---|---|
| In Microglia | 35 (59.34) | 197 (172.66) | 232 |
| Not IN Microglia | 153 (128.66) | 350 (374.34) | 503 |
| Total | 188 | 547 | 735 |

chi2: 18.807, p: 0.0000144608, OR: 0.40642

| | > cutoff | other genes | Total |
|---|---|---|---|
| In Microglia | 89 (58.39) | 143 (173.61) | 232 |
| Not IN Microglia | 96 (126.61) | 407 (376.39) | 503 |
| Total | 185 | 550 | 735 |

chi2: 30.308, p: 0.0000000369, OR: 2.63862

| | < cutoff | other genes | Total |
|---|---|---|---|
| In Oligodendrocyte | 24 (15.35) | 36 (44.65) | 60 |
| Not IN Oligodendrocyte | 164 (172.65) | 511 (502.35) | 675 |
| Total | 188 | 547 | 735 |

chi2: 6.337, p: 0.0118225721, OR: 2.07724

| | > cutoff | other genes | Total |
|---|---|---|---|
| In Oligodendrocyte | 9 (15.10) | 51 (44.90) | 60 |
| Not IN Oligodendrocyte | 176 (169.90) | 499 (505.10) | 675 |
| Total | 185 | 550 | 735 |

chi2: 3.024, p: 0.0820467547, OR: 0.50033

| | < cutoff | other genes | Total |
|---|---|---|---|
| In Neuron | 40 (29.16) | 74 (84.84) | 114 |
| Not IN Neuron | 148 (158.84) | 473 (462.16) | 621 |
| Total | 188 | 547 | 735 |

chi2: 5.832, p: 0.0157357180, OR: 1.72754

| | > cutoff | other genes | Total |
|---|---|---|---|
| In Neuron | 17 (28.69) | 97 (85.31) | 114 |
| Not IN Neuron | 168 (156.31) | 453 (464.69) | 621 |
| Total | 185 | 550 | 735 |

chi2: 6.907, p: 0.0085856795, OR: 0.47257

| | < cutoff | other genes | Total |
|---|---|---|---|
| In Endothelia | 38 (46.81) | 145 (136.19) | 183 |
| Not IN Endothelia | 150 (141.19) | 402 (410.81) | 552 |
| Total | 188 | 547 | 735 |

chi2: 2.638, p: 0.1043101097, OR: 0.70234

| | > cutoff | other genes | Total |
|---|---|---|---|
| In Endothelia | 48 (46.06) | 135 (136.94) | 183 |
| Not IN Endothelia | 137 (138.94) | 415 (413.06) | 552 |
| Total | 185 | 550 | 735 |

chi2: 0.080, p: 0.7773389660, OR: 1.07705

### Five Celltypes Distribution of Average dN/dS Scores of Genes Related to cell wall organization or biogenesis

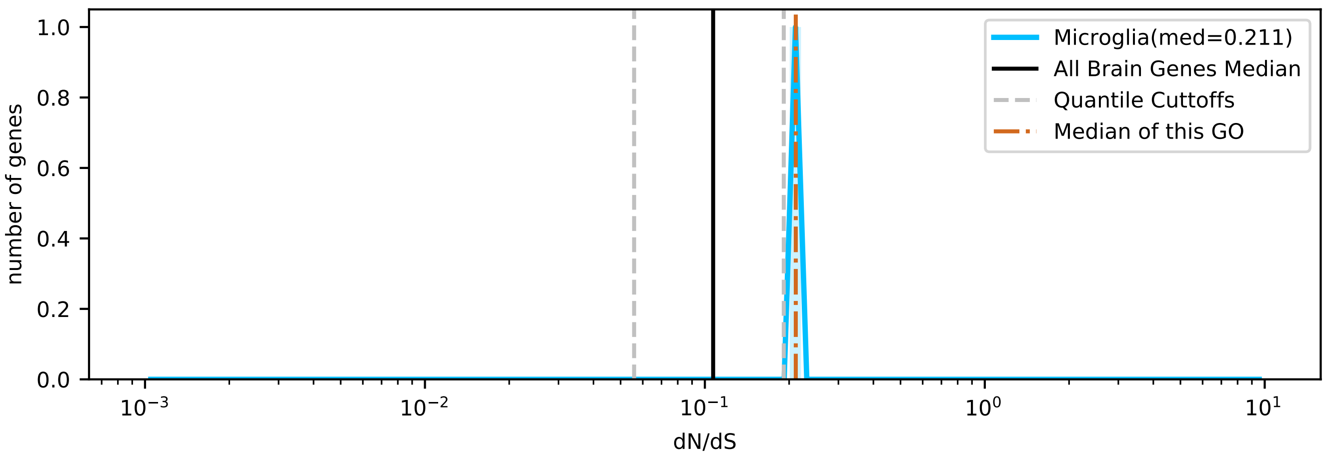

### Five Celltypes Distribution of Average dN/dS Scores of Genes Related to cell-cell signaling

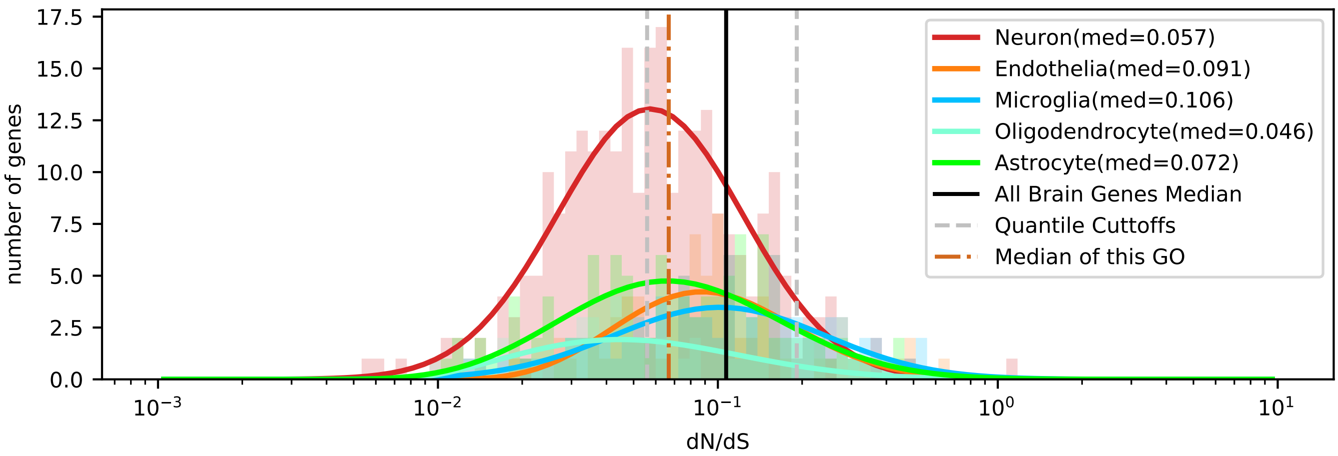


| | < cutoff | other genes | Total |
|---|---|---|---|
| In Astrocyte | 45 (47.07) | 68 (65.93) | 113 |
| Not IN Astrocyte | 202 (199.93) | 278 (280.07) | 480 |
| Total | 247 | 346 | 593 |

chi2: 0.111, p: 0.7395471556, OR: 0.91075

| | > cutoff | other genes | Total |
|---|---|---|---|
| In Astrocyte | 14 (12.39) | 99 (100.61) | 113 |
| Not IN Astrocyte | 51 (52.61) | 429 (427.39) | 480 |
| Total | 65 | 528 | 593 |

chi2: 0.139, p: 0.7093031906, OR: 1.18954

| | < cutoff | other genes | Total |
|---|---|---|---|
| In Microglia | 23 (34.16) | 59 (47.84) | 82 |
| Not IN Microglia | 224 (212.84) | 287 (298.16) | 511 |
| Total | 247 | 346 | 593 |

chi2: 6.611, p: 0.0101344051, OR: 0.49947

| | > cutoff | other genes | Total |
|---|---|---|---|
| In Microglia | 20 (8.99) | 62 (73.01) | 82 |
| Not IN Microglia | 45 (56.01) | 466 (454.99) | 511 |
| Total | 65 | 528 | 593 |

chi2: 16.023, p: 0.0000625865, OR: 3.34050

| | < cutoff | other genes | Total |
|---|---|---|---|
| In Oligodendrocyte | 27 (19.16) | 19 (26.84) | 46 |
| Not IN Oligodendrocyte | 220 (227.84) | 327 (319.16) | 547 |
| Total | 247 | 346 | 593 |

chi2: 5.224, p: 0.0222755765, OR: 2.11220

| | > cutoff | other genes | Total |
|---|---|---|---|
| In Oligodendrocyte | 5 (5.04) | 41 (40.96) | 46 |
| Not IN Oligodendrocyte | 60 (59.96) | 487 (487.04) | 547 |
| Total | 65 | 528 | 593 |

chi2: 0.051, p: 0.8219919921, OR: 0.98984

| | < cutoff | other genes | Total |
|---|---|---|---|
| In Neuron | 133 (112.46) | 137 (157.54) | 270 |
| Not IN Neuron | 114 (134.54) | 209 (188.46) | 323 |
| Total | 247 | 346 | 593 |

chi2: 11.234, p: 0.0008031619, OR: 1.77981

| | > cutoff | other genes | Total |
|---|---|---|---|
| In Neuron | 15 (29.60) | 255 (240.40) | 270 |
| Not IN Neuron | 50 (35.40) | 273 (287.60) | 323 |
| Total | 65 | 528 | 593 |

chi2: 13.842, p: 0.0001988449, OR: 0.32118

| | < cutoff | other genes | Total |
|---|---|---|---|
| In Endothelia | 19 (34.16) | 63 (47.84) | 82 |
| Not IN Endothelia | 228 (212.84) | 283 (298.16) | 511 |
| Total | 247 | 346 | 593 |

chi2: 12.507, p: 0.0004055347, OR: 0.37434

| | > cutoff | other genes | Total |
|---|---|---|---|
| In Endothelia | 11 (8.99) | 71 (73.01) | 82 |
| Not IN Endothelia | 54 (56.01) | 457 (454.99) | 511 |
| Total | 65 | 528 | 593 |

chi2: 0.331, p: 0.5648265826, OR: 1.31116

### Five Celltypes Distribution of Average dN/dS Scores of Genes Related to cellular amino acid metabolic process

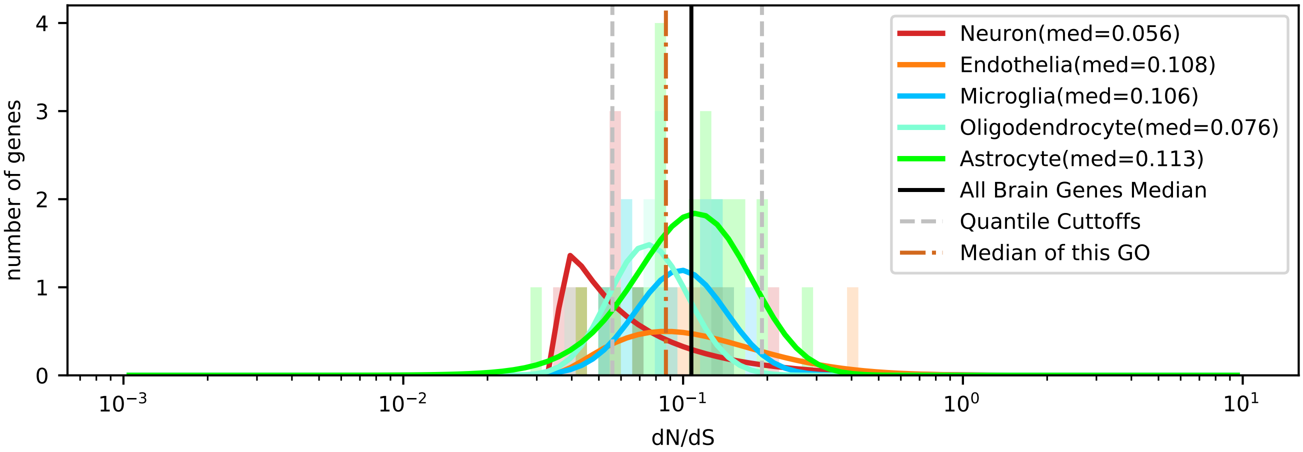


| | < cutoff | other genes | Total |
|---|---|---|---|
| In Astrocyte | 3 (4.24) | 21 (19.76) | 24 |
| Not IN Astrocyte | 9 (7.76) | 35 (36.24) | 44 |
| Total | 12 | 56 | 68 |

chi2: 0.240, p: 0.6245235180, OR: 0.55556

| | > cutoff | other genes | Total |
|---|---|---|---|
| In Astrocyte | 2 (1.41) | 22 (22.59) | 24 |
| Not IN Astrocyte | 2 (2.59) | 42 (41.41) | 44 |
| Total | 4 | 64 | 68 |

chi2: 0.009, p: 0.9241878575, OR: 1.90909

| | < cutoff | other genes | Total |
|---|---|---|---|
| In Microglia | 1 (2.12) | 11 (9.88) | 12 |
| Not IN Microglia | 11 (9.88) | 45 (46.12) | 56 |
| Total | 12 | 56 | 68 |

chi2: 0.266, p: 0.6062817743, OR: 0.37190

| | > cutoff | other genes | Total |
|---|---|---|---|
| In Microglia | 0 (0.71) | 12 (11.29) | 12 |
| Not IN Microglia | 4 (3.29) | 52 (52.71) | 56 |
| Total | 4 | 64 | 68 |

chi2: 0.077, p: 0.7807502664, OR: 0.00000

| | < cutoff | other genes | Total |
|---|---|---|---|
| In Oligodendrocyte | 2 (2.29) | 11 (10.71) | 13 |
| Not IN Oligodendrocyte | 10 (9.71) | 45 (45.29) | 55 |
| Total | 12 | 56 | 68 |

chi2: 0.028, p: 0.8677240786, OR: 0.81818

| | > cutoff | other genes | Total |
|---|---|---|---|
| In Oligodendrocyte | 0 (0.76) | 13 (12.24) | 13 |
| Not IN Oligodendrocyte | 4 (3.24) | 51 (51.76) | 55 |
| Total | 4 | 64 | 68 |

chi2: 0.120, p: 0.7286366450, OR: 0.00000

| | < cutoff | other genes | Total |
|---|---|---|---|
| In Neuron | 5 (1.94) | 6 (9.06) | 11 |
| Not IN Neuron | 7 (10.06) | 50 (46.94) | 57 |
| Total | 12 | 56 | 68 |

chi2: 4.886, p: 0.0270723230, OR: 5.95238

| | > cutoff | other genes | Total |
|---|---|---|---|
| In Neuron | 1 (0.65) | 10 (10.35) | 11 |
| Not IN Neuron | 3 (3.35) | 54 (53.65) | 57 |
| Total | 4 | 64 | 68 |

chi2: 0.042, p: 0.8369266572, OR: 1.80000

| | < cutoff | other genes | Total |
|---|---|---|---|
| In Endothelia | 1 (1.41) | 7 (6.59) | 8 |
| Not IN Endothelia | 11 (10.59) | 49 (49.41) | 60 |
| Total | 12 | 56 | 68 |

chi2: 0.008, p: 0.9305789049, OR: 0.63636

| | > cutoff | other genes | Total |
|---|---|---|---|
| In Endothelia | 1 (0.47) | 7 (7.53) | 8 |
| Not IN Endothelia | 3 (3.53) | 57 (56.47) | 60 |
| Total | 4 | 64 | 68 |

chi2: 0.002, p: 0.9624747396, OR: 2.71429

### Five Celltypes Distribution of Average dN/dS Scores of Genes Related to cellular component assembly

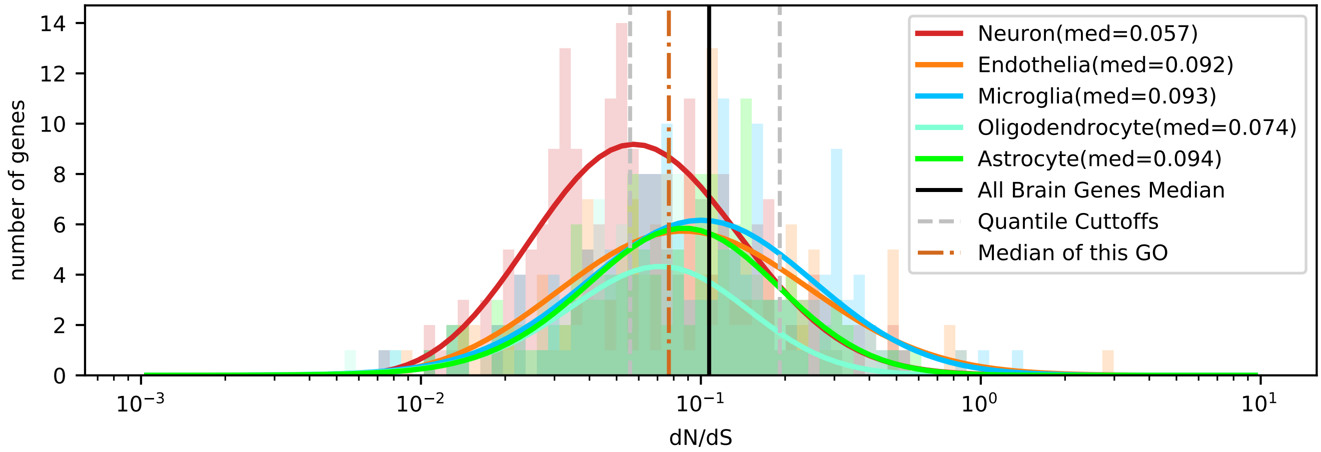


| | < cutoff | other genes | Total |
|---|---|---|---|
| In Astrocyte | 40 (45.46) | 86 (80.54) | 126 |
| Not IN Astrocyte | 223 (217.54) | 380 (385.46) | 603 |
| Total | 263 | 466 | 729 |

chi2: 1.022, p: 0.3119875654, OR: 0.79257

| | > cutoff | other genes | Total |
|---|---|---|---|
| In Astrocyte | 17 (19.36) | 109 (106.64) | 126 |
| Not IN Astrocyte | 95 (92.64) | 508 (510.36) | 603 |
| Total | 112 | 617 | 729 |

chi2: 0.255, p: 0.6137581754, OR: 0.83399

| | < cutoff | other genes | Total |
|---|---|---|---|
| In Microglia | 40 (55.92) | 115 (99.08) | 155 |
| Not IN Microglia | 223 (207.08) | 351 (366.92) | 574 |
| Total | 263 | 466 | 729 |

chi2: 8.447, p: 0.0036560294, OR: 0.54748

| | > cutoff | other genes | Total |
|---|---|---|---|
| In Microglia | 36 (23.81) | 119 (131.19) | 155 |
| Not IN Microglia | 76 (88.19) | 498 (485.81) | 574 |
| Total | 112 | 617 | 729 |

chi2: 8.606, p: 0.0033502956, OR: 1.98231

| | < cutoff | other genes | Total |
|---|---|---|---|
| In Oligodendrocyte | 37 (31.75) | 51 (56.25) | 88 |
| Not IN Oligodendrocyte | 226 (231.25) | 415 (409.75) | 641 |
| Total | 263 | 466 | 729 |

chi2: 1.266, p: 0.2605773725, OR: 1.33221

| | > cutoff | other genes | Total |
|---|---|---|---|
| In Oligodendrocyte | 6 (13.52) | 82 (74.48) | 88 |
| Not IN Oligodendrocyte | 106 (98.48) | 535 (542.52) | 641 |
| Total | 112 | 617 | 729 |

chi2: 4.898, p: 0.0268912322, OR: 0.36931

| | < cutoff | other genes | Total |
|---|---|---|---|
| In Neuron | 102 (74.32) | 104 (131.68) | 206 |
| Not IN Neuron | 161 (188.68) | 362 (334.32) | 523 |
| Total | 263 | 466 | 729 |

chi2: 21.678, p: 0.0000032240, OR: 2.20521

| | > cutoff | other genes | Total |
|---|---|---|---|
| In Neuron | 18 (31.65) | 188 (174.35) | 206 |
| Not IN Neuron | 94 (80.35) | 429 (442.65) | 523 |
| Total | 112 | 617 | 729 |

chi2: 8.997, p: 0.0027046223, OR: 0.43696

| | < cutoff | other genes | Total |
|---|---|---|---|
| In Endothelia | 44 (55.56) | 110 (98.44) | 154 |
| Not IN Endothelia | 219 (207.44) | 356 (367.56) | 575 |
| Total | 263 | 466 | 729 |

chi2: 4.365, p: 0.0366747963, OR: 0.65023

| | > cutoff | other genes | Total |
|---|---|---|---|
| In Endothelia | 35 (23.66) | 119 (130.34) | 154 |
| Not IN Endothelia | 77 (88.34) | 498 (486.66) | 575 |
| Total | 112 | 617 | 729 |

chi2: 7.440, p: 0.0063794839, OR: 1.90222

### Five Celltypes Distribution of Average dN/dS Scores of Genes Related to cellular nitrogen compound metabolic process

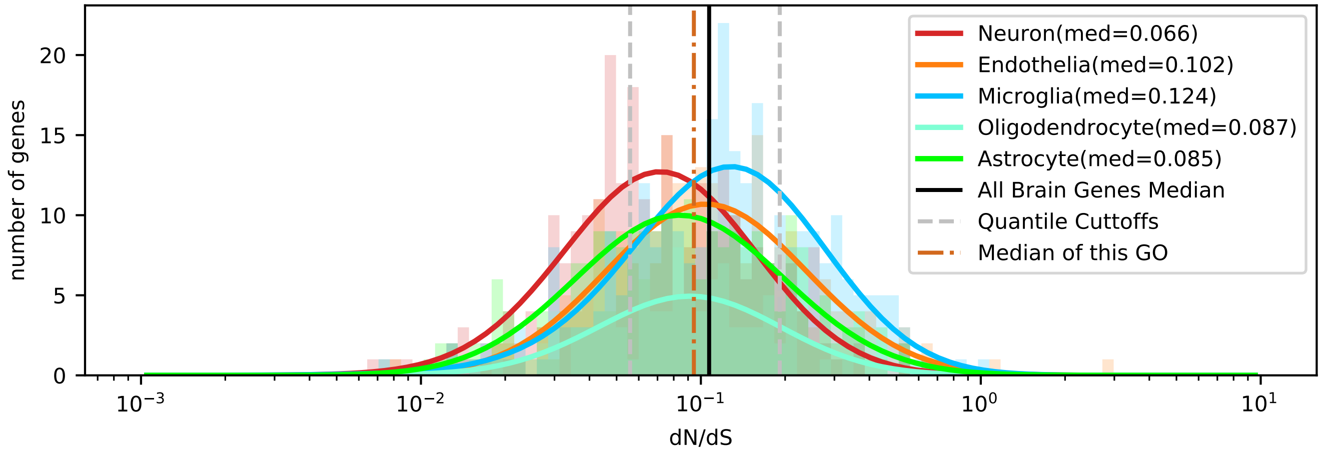


| | < cutoff | other genes | Total |
|---|---|---|---|
| In Astrocyte | 74 (64.20) | 156 (165.80) | 230 |
| Not IN Astrocyte | 240 (249.80) | 655 (645.20) | 895 |
| Total | 314 | 811 | 1125 |

chi2: 2.351, p: 0.1251659526, OR: 1.29460

| | > cutoff | other genes | Total |
|---|---|---|---|
| In Astrocyte | 42 (45.39) | 188 (184.61) | 230 |
| Not IN Astrocyte | 180 (176.61) | 715 (718.39) | 895 |
| Total | 222 | 903 | 1125 |

chi2: 0.288, p: 0.5918179332, OR: 0.88741

| | < cutoff | other genes | Total |
|---|---|---|---|
| In Microglia | 49 (79.27) | 235 (204.73) | 284 |
| Not IN Microglia | 265 (234.73) | 576 (606.27) | 841 |
| Total | 314 | 811 | 1125 |

chi2: 20.743, p: 0.0000052513, OR: 0.45322

| | > cutoff | other genes | Total |
|---|---|---|---|
| In Microglia | 79 (56.04) | 205 (227.96) | 284 |
| Not IN Microglia | 143 (165.96) | 698 (675.04) | 841 |
| Total | 222 | 903 | 1125 |

chi2: 14.998, p: 0.0001076537, OR: 1.88102

| | < cutoff | other genes | Total |
|---|---|---|---|
| In Oligodendrocyte | 27 (28.19) | 74 (72.81) | 101 |
| Not IN Oligodendrocyte | 287 (285.81) | 737 (738.19) | 1024 |
| Total | 314 | 811 | 1125 |

chi2: 0.026, p: 0.8724998056, OR: 0.93695

| | > cutoff | other genes | Total |
|---|---|---|---|
| In Oligodendrocyte | 15 (19.93) | 86 (81.07) | 101 |
| Not IN Oligodendrocyte | 207 (202.07) | 817 (821.93) | 1024 |
| Total | 222 | 903 | 1125 |

chi2: 1.348, p: 0.2456042690, OR: 0.68841

| | < cutoff | other genes | Total |
|---|---|---|---|
| In Neuron | 110 (75.92) | 162 (196.08) | 272 |
| Not IN Neuron | 204 (238.08) | 649 (614.92) | 853 |
| Total | 314 | 811 | 1125 |

chi2: 27.177, p: 0.0000001857, OR: 2.16019

| | > cutoff | other genes | Total |
|---|---|---|---|
| In Neuron | 32 (53.67) | 240 (218.33) | 272 |
| Not IN Neuron | 190 (168.33) | 663 (684.67) | 853 |
| Total | 222 | 903 | 1125 |

chi2: 13.726, p: 0.0002115486, OR: 0.46526

| | < cutoff | other genes | Total |
|---|---|---|---|
| In Endothelia | 54 (66.43) | 184 (171.57) | 238 |
| Not IN Endothelia | 260 (247.57) | 627 (639.43) | 887 |
| Total | 314 | 811 | 1125 |

chi2: 3.769, p: 0.0522249331, OR: 0.70773

| | > cutoff | other genes | Total |
|---|---|---|---|
| In Endothelia | 54 (46.97) | 184 (191.03) | 238 |
| Not IN Endothelia | 168 (175.03) | 719 (711.97) | 887 |
| Total | 222 | 903 | 1125 |

chi2: 1.437, p: 0.2306756820, OR: 1.25602

### Five Celltypes Distribution of Average dN/dS Scores of Genes Related to cellular protein modification process

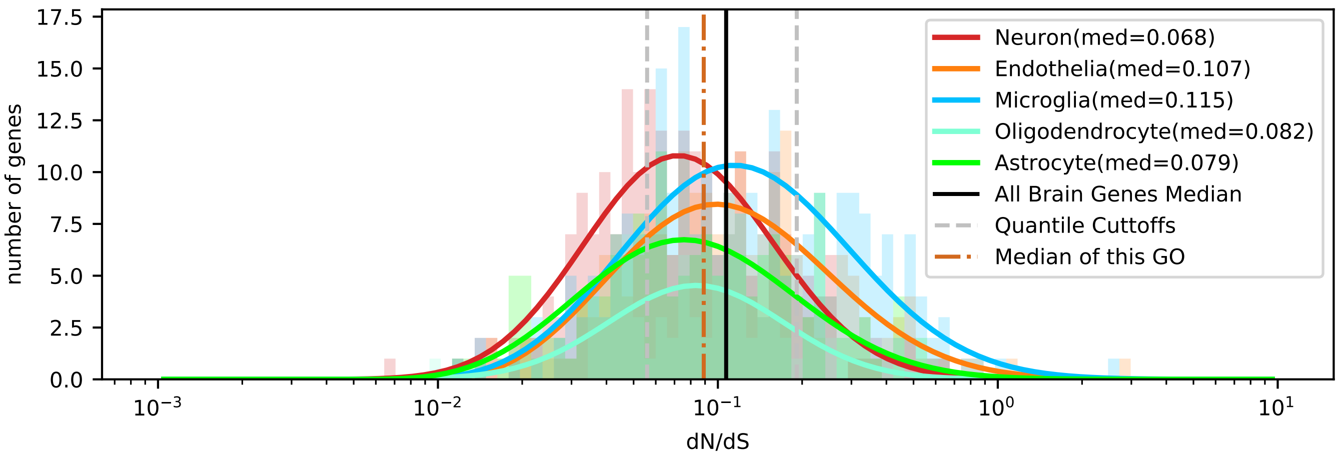


| | < cutoff | other genes | Total |
|---|---|---|---|
| In Astrocyte | 53 (44.53) | 105 (113.47) | 158 |
| Not IN Astrocyte | 208 (216.47) | 560 (551.53) | 768 |
| Total | 261 | 665 | 926 |

chi2: 2.393, p: 0.1219021303, OR: 1.35897

| | > cutoff | other genes | Total |
|---|---|---|---|
| In Astrocyte | 26 (31.57) | 132 (126.43) | 158 |
| Not IN Astrocyte | 159 (153.43) | 609 (614.57) | 768 |
| Total | 185 | 741 | 926 |

chi2: 1.225, p: 0.2683829936, OR: 0.75443

| | < cutoff | other genes | Total |
|---|---|---|---|
| In Microglia | 49 (71.87) | 206 (183.13) | 255 |
| Not IN Microglia | 212 (189.13) | 459 (481.87) | 671 |
| Total | 261 | 665 | 926 |

chi2: 13.384, p: 0.0002537972, OR: 0.51500

| | > cutoff | other genes | Total |
|---|---|---|---|
| In Microglia | 83 (50.94) | 172 (204.06) | 255 |
| Not IN Microglia | 102 (134.06) | 569 (536.94) | 671 |
| Total | 185 | 741 | 926 |

chi2: 33.707, p: 0.0000000064, OR: 2.69192

| | < cutoff | other genes | Total |
|---|---|---|---|
| In Oligodendrocyte | 30 (24.80) | 58 (63.20) | 88 |
| Not IN Oligodendrocyte | 231 (236.20) | 607 (601.80) | 838 |
| Total | 261 | 665 | 926 |

chi2: 1.368, p: 0.2420935394, OR: 1.35916

| | > cutoff | other genes | Total |
|---|---|---|---|
| In Oligodendrocyte | 10 (17.58) | 78 (70.42) | 88 |
| Not IN Oligodendrocyte | 175 (167.42) | 663 (670.58) | 838 |
| Total | 185 | 741 | 926 |

chi2: 3.938, p: 0.0471994868, OR: 0.48571

| | < cutoff | other genes | Total |
|---|---|---|---|
| In Neuron | 86 (63.70) | 140 (162.30) | 226 |
| Not IN Neuron | 175 (197.30) | 525 (502.70) | 700 |
| Total | 261 | 665 | 926 |

chi2: 13.743, p: 0.0002095850, OR: 1.84286

| | > cutoff | other genes | Total |
|---|---|---|---|
| In Neuron | 24 (45.15) | 202 (180.85) | 226 |
| Not IN Neuron | 161 (139.85) | 539 (560.15) | 700 |
| Total | 185 | 741 | 926 |

chi2: 15.614, p: 0.0000776590, OR: 0.39776

| | < cutoff | other genes | Total |
|---|---|---|---|
| In Endothelia | 43 (56.09) | 156 (142.91) | 199 |
| Not IN Endothelia | 218 (204.91) | 509 (522.09) | 727 |
| Total | 261 | 665 | 926 |

chi2: 5.012, p: 0.0251724807, OR: 0.64358

| | > cutoff | other genes | Total |
|---|---|---|---|
| In Endothelia | 42 (39.76) | 157 (159.24) | 199 |
| Not IN Endothelia | 143 (145.24) | 584 (581.76) | 727 |
| Total | 185 | 741 | 926 |

chi2: 0.122, p: 0.7272733283, OR: 1.09251

Five Celltypes Distribution of Average dN/dS Scores of Genes Related to chromosome organization

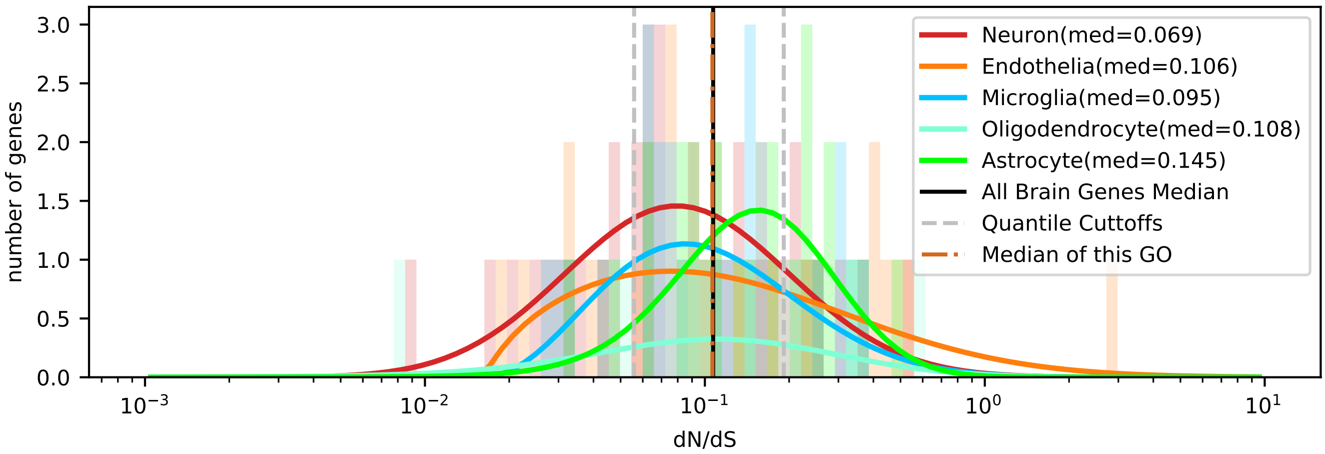


| | < cutoff | other genes | Total |
|---|---|---|---|
| In Astrocyte | 2 (5.53) | 23 (19.47) | 25 |
| Not IN Astrocyte | 25 (21.47) | 72 (75.53) | 97 |
| Total | 27 | 95 | 122 |

chi2: 2.685, p: 0.1012894115, OR: 0.25043

| | > cutoff | other genes | Total |
|---|---|---|---|
| In Astrocyte | 10 (6.76) | 15 (18.24) | 25 |
| Not IN Astrocyte | 23 (26.24) | 74 (70.76) | 97 |
| Total | 33 | 89 | 122 |

chi2: 1.911, p: 0.1668637520, OR: 2.14493

| | < cutoff | other genes | Total |
|---|---|---|---|
| In Microglia | 4 (5.31) | 20 (18.69) | 24 |
| Not IN Microglia | 23 (21.69) | 75 (76.31) | 98 |
| Total | 27 | 95 | 122 |

chi2: 0.198, p: 0.6561769965, OR: 0.65217

| | > cutoff | other genes | Total |
|---|---|---|---|
| In Microglia | 5 (6.49) | 19 (17.51) | 24 |
| Not IN Microglia | 28 (26.51) | 70 (71.49) | 98 |
| Total | 33 | 89 | 122 |

chi2: 0.259, p: 0.6110997046, OR: 0.65789

| | < cutoff | other genes | Total |
|---|---|---|---|
| In Oligodendrocyte | 3 (1.99) | 6 (7.01) | 9 |
| Not IN Oligodendrocyte | 24 (25.01) | 89 (87.99) | 113 |
| Total | 27 | 95 | 122 |

chi2: 0.180, p: 0.6715646833, OR: 1.85417

| | > cutoff | other genes | Total |
|---|---|---|---|
| In Oligodendrocyte | 3 (2.43) | 6 (6.57) | 9 |
| Not IN Oligodendrocyte | 30 (30.57) | 83 (82.43) | 113 |
| Total | 33 | 89 | 122 |

chi2: 0.003, p: 0.9592236512, OR: 1.38333

| | < cutoff | other genes | Total |
|---|---|---|---|
| In Neuron | 12 (7.97) | 24 (28.03) | 36 |
| Not IN Neuron | 15 (19.03) | 71 (66.97) | 86 |
| Total | 27 | 95 | 122 |

chi2: 2.854, p: 0.0911573024, OR: 2.36667

| | > cutoff | other genes | Total |
|---|---|---|---|
| In Neuron | 7 (9.74) | 29 (26.26) | 36 |
| Not IN Neuron | 26 (23.26) | 60 (62.74) | 86 |
| Total | 33 | 89 | 122 |

chi2: 1.000, p: 0.3173215250, OR: 0.55703

| | < cutoff | other genes | Total |
|---|---|---|---|
| In Endothelia | 6 (6.20) | 22 (21.80) | 28 |
| Not IN Endothelia | 21 (20.80) | 73 (73.20) | 94 |
| Total | 27 | 95 | 122 |

chi2: 0.025, p: 0.8750180421, OR: 0.94805

| | > cutoff | other genes | Total |
|---|---|---|---|
| In Endothelia | 8 (7.57) | 20 (20.43) | 28 |
| Not IN Endothelia | 25 (25.43) | 69 (68.57) | 94 |
| Total | 33 | 89 | 122 |

chi2: 0.001, p: 0.9714783237, OR: 1.10400

### Five Celltypes Distribution of Average dN/dS Scores of Genes Related to chromosome segregation

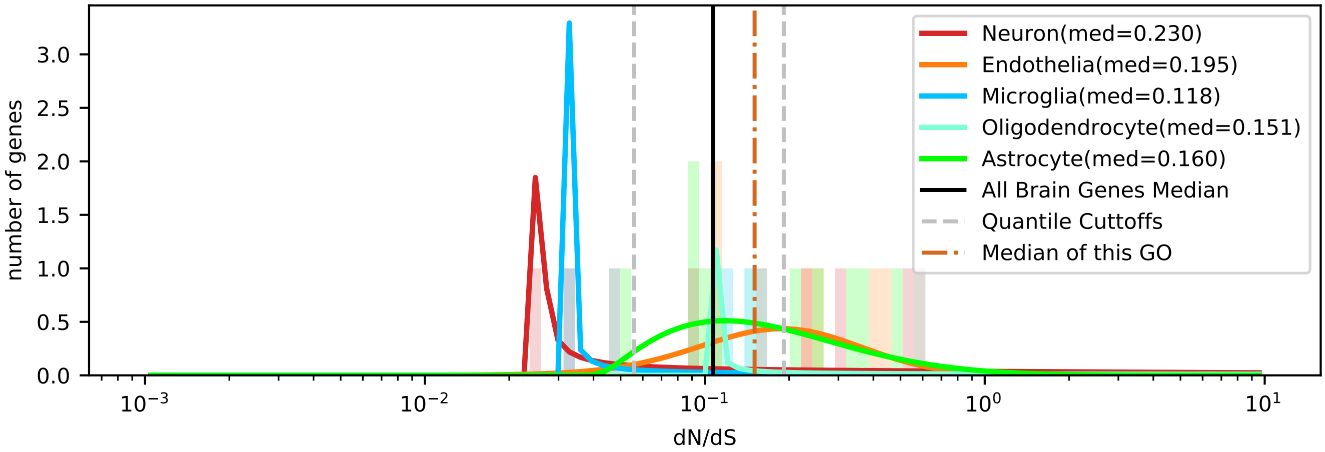


| | < cutoff | other genes | Total |
|---|---|---|---|
| In Astrocyte | 1 (1.82) | 9 (8.18) | 10 |
| Not IN Astrocyte | 5 (4.18) | 18 (18.82) | 23 |
| Total | 6 | 27 | 33 |

chi2: 0.098, p: 0.7546743291, OR: 0.40000

| | > cutoff | other genes | Total |
|---|---|---|---|
| In Astrocyte | 5 (4.24) | 5 (5.76) | 10 |
| Not IN Astrocyte | 9 (9.76) | 14 (13.24) | 23 |
| Total | 14 | 19 | 33 |

chi2: 0.039, p: 0.8435059895, OR: 1.55556

| | < cutoff | other genes | Total |
|---|---|---|---|
| In Microglia | 2 (0.91) | 3 (4.09) | 5 |
| Not IN Microglia | 4 (5.09) | 24 (22.91) | 28 |
| Total | 6 | 27 | 33 |

chi2: 0.553, p: 0.4569830942, OR: 4.00000

| | > cutoff | other genes | Total |
|---|---|---|---|
| In Microglia | 0 (2.12) | 5 (2.88) | 5 |
| Not IN Microglia | 14 (11.88) | 14 (16.12) | 28 |
| Total | 14 | 19 | 33 |

chi2: 2.536, p: 0.1112506085, OR: 0.00000

| | < cutoff | other genes | Total |
|---|---|---|---|
| In Oligodendrocyte | 0 (0.55) | 3 (2.45) | 3 |
| Not IN Oligodendrocyte | 6 (5.45) | 24 (24.55) | 30 |
| Total | 6 | 27 | 33 |

chi2: 0.005, p: 0.9431093312, OR: 0.00000

| | > cutoff | other genes | Total |
|---|---|---|---|
| In Oligodendrocyte | 1 (1.27) | 2 (1.73) | 3 |
| Not IN Oligodendrocyte | 13 (12.73) | 17 (17.27) | 30 |
| Total | 14 | 19 | 33 |

chi2: 0.078, p: 0.7806625434, OR: 0.65385

| | < cutoff | other genes | Total |
|---|---|---|---|
| In Neuron | 2 (1.27) | 5 (5.73) | 7 |
| Not IN Neuron | 4 (4.73) | 22 (21.27) | 26 |
| Total | 6 | 27 | 33 |

chi2: 0.063, p: 0.8018805384, OR: 2.20000

| | > cutoff | other genes | Total |
|---|---|---|---|
| In Neuron | 4 (2.97) | 3 (4.03) | 7 |
| Not IN Neuron | 10 (11.03) | 16 (14.97) | 26 |
| Total | 14 | 19 | 33 |

chi2: 0.209, p: 0.6477449293, OR: 2.13333

| | < cutoff | other genes | Total |
|---|---|---|---|
| In Endothelia | 1 (1.45) | 7 (6.55) | 8 |
| Not IN Endothelia | 5 (4.55) | 20 (20.45) | 25 |
| Total | 6 | 27 | 33 |

chi2: 0.002, p: 0.9618187683, OR: 0.57143

| | > cutoff | other genes | Total |
|---|---|---|---|
| In Endothelia | 4 (3.39) | 4 (4.61) | 8 |
| Not IN Endothelia | 10 (10.61) | 15 (14.39) | 25 |
| Total | 14 | 19 | 33 |

chi2: 0.008, p: 0.9305360417, OR: 1.50000

### Five Celltypes Distribution of Average dN/dS Scores of Genes Related to circulatory system process

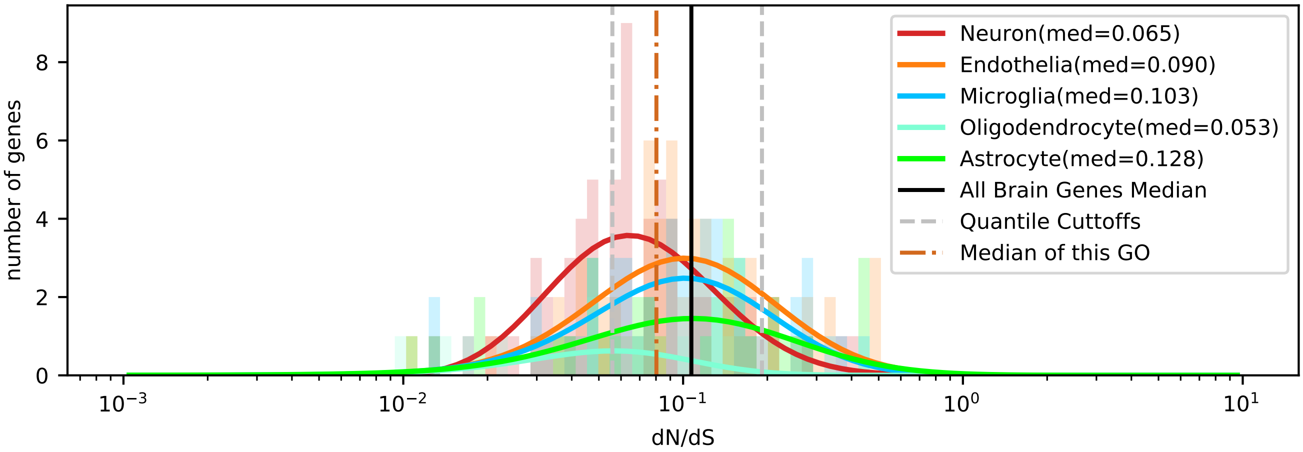


| | < cutoff | other genes | Total |
|---|---|---|---|
| In Astrocyte | 10 (10.11) | 25 (24.89) | 35 |
| Not IN Astrocyte | 55 (54.89) | 135 (135.11) | 190 |
| Total | 65 | 160 | 225 |

chi2: 0.025, p: 0.8745958376, OR: 0.98182

| | > cutoff | other genes | Total |
|---|---|---|---|
| In Astrocyte | 8 (5.13) | 27 (29.87) | 35 |
| Not IN Astrocyte | 25 (27.87) | 165 (162.13) | 190 |
| Total | 33 | 192 | 225 |

chi2: 1.514, p: 0.2184984264, OR: 1.95556

| | < cutoff | other genes | Total |
|---|---|---|---|
| In Microglia | 11 (14.73) | 40 (36.27) | 51 |
| Not IN Microglia | 54 (50.27) | 120 (123.73) | 174 |
| Total | 65 | 160 | 225 |

chi2: 1.290, p: 0.2559899062, OR: 0.61111

| | > cutoff | other genes | Total |
|---|---|---|---|
| In Microglia | 8 (7.48) | 43 (43.52) | 51 |
| Not IN Microglia | 25 (25.52) | 149 (148.48) | 174 |
| Total | 33 | 192 | 225 |

chi2: 0.000, p: 0.9928175821, OR: 1.10884

| | < cutoff | other genes | Total |
|---|---|---|---|
| In Oligodendrocyte | 6 (3.47) | 6 (8.53) | 12 |
| Not IN Oligodendrocyte | 59 (61.53) | 154 (151.47) | 213 |
| Total | 65 | 160 | 225 |

chi2: 1.772, p: 0.1831817427, OR: 2.61017

| | > cutoff | other genes | Total |
|---|---|---|---|
| In Oligodendrocyte | 0 (1.76) | 12 (10.24) | 12 |
| Not IN Oligodendrocyte | 33 (31.24) | 180 (181.76) | 213 |
| Total | 33 | 192 | 225 |

chi2: 1.117, p: 0.2906434088, OR: 0.00000

| | < cutoff | other genes | Total |
|---|---|---|---|
| In Neuron | 25 (18.78) | 40 (46.22) | 65 |
| Not IN Neuron | 40 (46.22) | 120 (113.78) | 160 |
| Total | 65 | 160 | 225 |

chi2: 3.448, p: 0.0633152582, OR: 1.87500

| | > cutoff | other genes | Total |
|---|---|---|---|
| In Neuron | 5 (9.53) | 60 (55.47) | 65 |
| Not IN Neuron | 28 (23.47) | 132 (136.53) | 160 |
| Total | 33 | 192 | 225 |

chi2: 2.812, p: 0.0935571492, OR: 0.39286

| | < cutoff | other genes | Total |
|---|---|---|---|
| In Endothelia | 13 (17.91) | 49 (44.09) | 62 |
| Not IN Endothelia | 52 (47.09) | 111 (115.91) | 163 |
| Total | 65 | 160 | 225 |

chi2: 2.109, p: 0.1464562035, OR: 0.56633

| | > cutoff | other genes | Total |
|---|---|---|---|
| In Endothelia | 12 (9.09) | 50 (52.91) | 62 |
| Not IN Endothelia | 21 (23.91) | 142 (139.09) | 163 |
| Total | 33 | 192 | 225 |

chi2: 1.030, p: 0.3100763643, OR: 1.62286

### Five Celltypes Distribution of Average dN/dS Scores of Genes Related to cofactor metabolic process

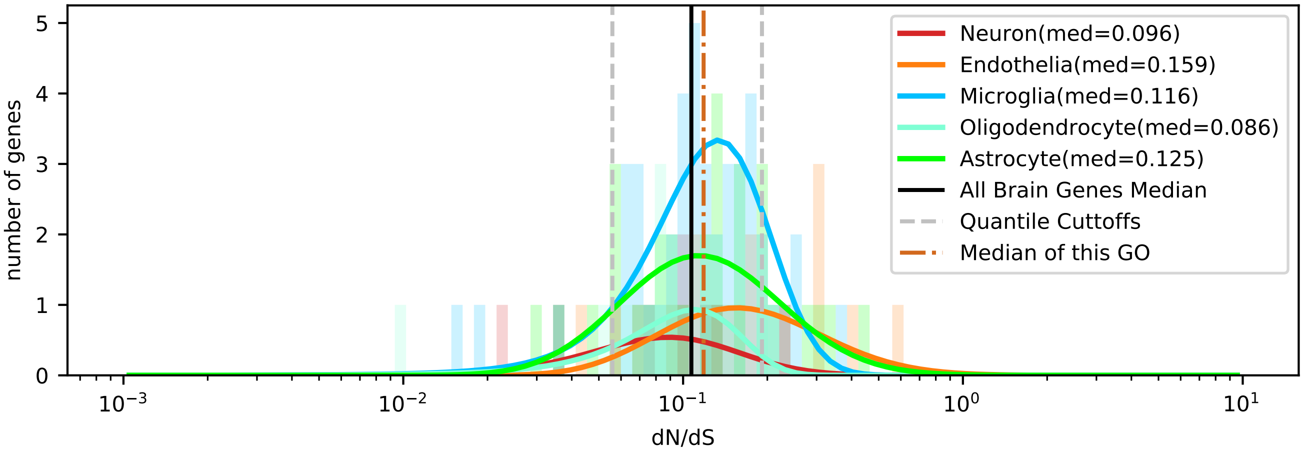


| | < cutoff | other genes | Total |
|---|---|---|---|
| In Astrocyte | 3 (3.21) | 27 (26.79) | 30 |
| Not IN Astrocyte | 9 (8.79) | 73 (73.21) | 82 |
| Total | 12 | 100 | 112 |

chi2: 0.039, p: 0.8437442443, OR: 0.90123

| | > cutoff | other genes | Total |
|---|---|---|---|
| In Astrocyte | 6 (4.82) | 24 (25.18) | 30 |
| Not IN Astrocyte | 12 (13.18) | 70 (68.82) | 82 |
| Total | 18 | 94 | 112 |

chi2: 0.155, p: 0.6934077535, OR: 1.45833

| | < cutoff | other genes | Total |
|---|---|---|---|
| In Microglia | 3 (4.71) | 41 (39.29) | 44 |
| Not IN Microglia | 9 (7.29) | 59 (60.71) | 68 |
| Total | 12 | 100 | 112 |

chi2: 0.577, p: 0.4475025603, OR: 0.47967

| | > cutoff | other genes | Total |
|---|---|---|---|
| In Microglia | 5 (7.07) | 39 (36.93) | 44 |
| Not IN Microglia | 13 (10.93) | 55 (57.07) | 68 |
| Total | 18 | 94 | 112 |

chi2: 0.685, p: 0.4077672743, OR: 0.54241

| | < cutoff | other genes | Total |
|---|---|---|---|
| In Oligodendrocyte | 2 (1.29) | 10 (10.71) | 12 |
| Not IN Oligodendrocyte | 10 (10.71) | 90 (89.29) | 100 |
| Total | 12 | 100 | 112 |

chi2: 0.045, p: 0.8323722146, OR: 1.80000

| | > cutoff | other genes | Total |
|---|---|---|---|
| In Oligodendrocyte | 0 (1.93) | 12 (10.07) | 12 |
| Not IN Oligodendrocyte | 18 (16.07) | 82 (83.93) | 100 |
| Total | 18 | 94 | 112 |

chi2: 1.412, p: 0.2347022048, OR: 0.00000

| | < cutoff | other genes | Total |
|---|---|---|---|
| In Neuron | 3 (0.96) | 6 (8.04) | 9 |
| Not IN Neuron | 9 (11.04) | 94 (91.96) | 103 |
| Total | 12 | 100 | 112 |

chi2: 2.979, p: 0.0843715978, OR: 5.22222

| | > cutoff | other genes | Total |
|---|---|---|---|
| In Neuron | 1 (1.45) | 8 (7.55) | 9 |
| Not IN Neuron | 17 (16.55) | 86 (86.45) | 103 |
| Total | 18 | 94 | 112 |

chi2: 0.003, p: 0.9595634365, OR: 0.63235

| | < cutoff | other genes | Total |
|---|---|---|---|
| In Endothelia | 1 (1.82) | 16 (15.18) | 17 |
| Not IN Endothelia | 11 (10.18) | 84 (84.82) | 95 |
| Total | 12 | 100 | 112 |

chi2: 0.075, p: 0.7843346343, OR: 0.47727

| | > cutoff | other genes | Total |
|---|---|---|---|
| In Endothelia | 6 (2.73) | 11 (14.27) | 17 |
| Not IN Endothelia | 12 (15.27) | 83 (79.73) | 95 |
| Total | 18 | 94 | 112 |

chi2: 3.939, p: 0.0471832554, OR: 3.77273

### Five Celltypes Distribution of Average dN/dS Scores of Genes Related to cytoskeleton organization

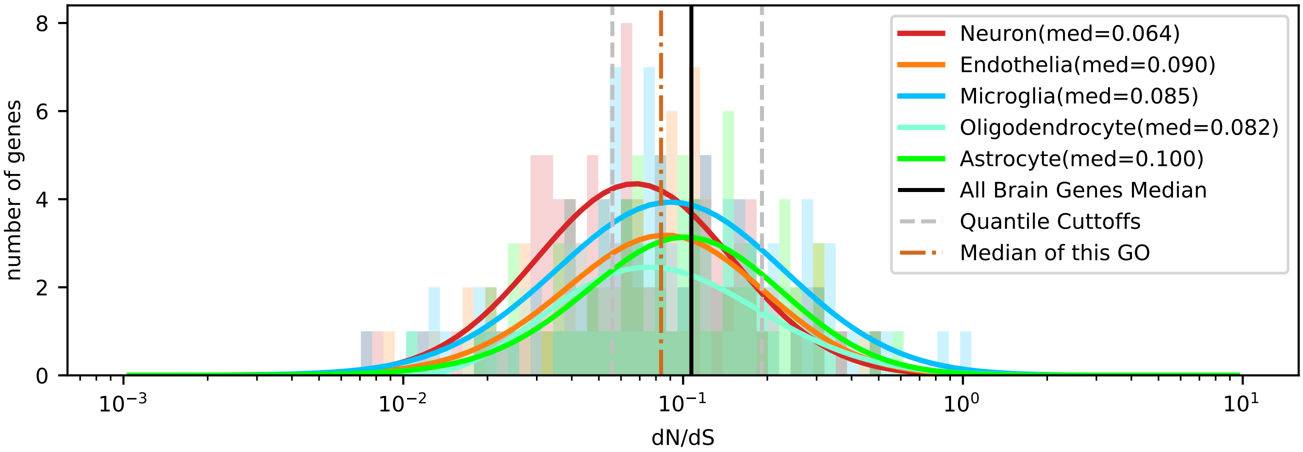


| | < cutoff | other genes | Total |
|---|---|---|---|
| In Astrocyte | 17 (22.12) | 52 (46.88) | 69 |
| Not IN Astrocyte | 108 (102.88) | 213 (218.12) | 321 |
| Total | 125 | 265 | 390 |

chi2: 1.722, p: 0.1894019385, OR: 0.64476

| | > cutoff | other genes | Total |
|---|---|---|---|
| In Astrocyte | 14 (11.15) | 55 (57.85) | 69 |
| Not IN Astrocyte | 49 (51.85) | 272 (269.15) | 321 |
| Total | 63 | 327 | 390 |

chi2: 0.720, p: 0.3960487140, OR: 1.41299

| | < cutoff | other genes | Total |
|---|---|---|---|
| In Microglia | 28 (32.05) | 72 (67.95) | 100 |
| Not IN Microglia | 97 (92.95) | 193 (197.05) | 290 |
| Total | 125 | 265 | 390 |

chi2: 0.779, p: 0.3775170018, OR: 0.77377

| | > cutoff | other genes | Total |
|---|---|---|---|
| In Microglia | 20 (16.15) | 80 (83.85) | 100 |
| Not IN Microglia | 43 (46.85) | 247 (243.15) | 290 |
| Total | 63 | 327 | 390 |

chi2: 1.112, p: 0.2917061911, OR: 1.43605

| | < cutoff | other genes | Total |
|---|---|---|---|
| In Oligodendrocyte | 18 (17.63) | 37 (37.37) | 55 |
| Not IN Oligodendrocyte | 107 (107.37) | 228 (227.63) | 335 |
| Total | 125 | 265 | 390 |

chi2: 0.002, p: 0.9681180366, OR: 1.03663

| | > cutoff | other genes | Total |
|---|---|---|---|
| In Oligodendrocyte | 10 (8.88) | 45 (46.12) | 55 |
| Not IN Oligodendrocyte | 53 (54.12) | 282 (280.88) | 335 |
| Total | 63 | 327 | 390 |

chi2: 0.059, p: 0.8077931662, OR: 1.18239

| | < cutoff | other genes | Total |
|---|---|---|---|
| In Neuron | 41 (30.45) | 54 (64.55) | 95 |
| Not IN Neuron | 84 (94.55) | 211 (200.45) | 295 |
| Total | 125 | 265 | 390 |

chi2: 6.456, p: 0.0110604466, OR: 1.90719

| | > cutoff | other genes | Total |
|---|---|---|---|
| In Neuron | 8 (15.35) | 87 (79.65) | 95 |
| Not IN Neuron | 55 (47.65) | 240 (247.35) | 295 |
| Total | 63 | 327 | 390 |

chi2: 4.816, p: 0.0282027433, OR: 0.40125

| | < cutoff | other genes | Total |
|---|---|---|---|
| In Endothelia | 21 (22.76) | 50 (48.24) | 71 |
| Not IN Endothelia | 104 (102.24) | 215 (216.76) | 319 |
| Total | 125 | 265 | 390 |

chi2: 0.125, p: 0.7238743471, OR: 0.86827

| | > cutoff | other genes | Total |
|---|---|---|---|
| In Endothelia | 11 (11.47) | 60 (59.53) | 71 |
| Not IN Endothelia | 52 (51.53) | 267 (267.47) | 319 |
| Total | 63 | 327 | 390 |

chi2: 0.000, p: 0.9912466109, OR: 0.94135

Five Celltypes Distribution of Average dN/dS Scores of Genes Related to cytoskeleton-dependent intracellular transport

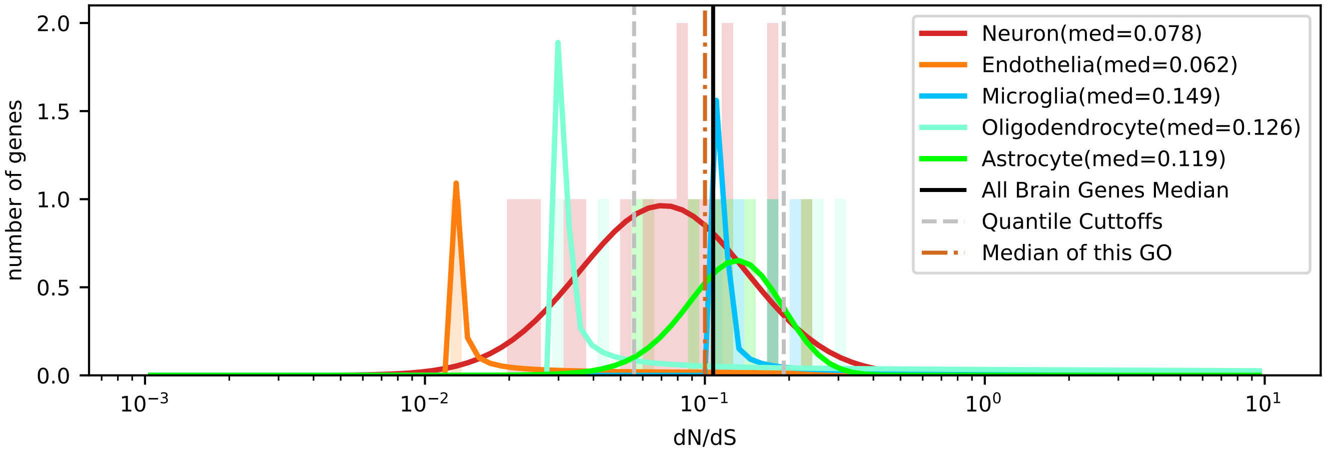


| | < cutoff | other genes | Total |
|---|---|---|---|
| In Astrocyte | 0 (1.62) | 7 (5.38) | 7 |
| Not IN Astrocyte | 9 (7.38) | 23 (24.62) | 32 |
| Total | 9 | 30 | 39 |

chi2: 1.220, p: 0.2693212303, OR: 0.00000

| | > cutoff | other genes | Total |
|---|---|---|---|
| In Astrocyte | 1 (0.90) | 6 (6.10) | 7 |
| Not IN Astrocyte | 4 (4.10) | 28 (27.90) | 32 |
| Total | 5 | 34 | 39 |

chi2: 0.246, p: 0.6198671302, OR: 1.16667

| | < cutoff | other genes | Total |
|---|---|---|---|
| In Microglia | 0 (0.92) | 4 (3.08) | 4 |
| Not IN Microglia | 9 (8.08) | 26 (26.92) | 35 |
| Total | 9 | 30 | 39 |

chi2: 0.281, p: 0.5961166029, OR: 0.00000

| | > cutoff | other genes | Total |
|---|---|---|---|
| In Microglia | 1 (0.51) | 3 (3.49) | 4 |
| Not IN Microglia | 4 (4.49) | 31 (30.51) | 35 |
| Total | 5 | 34 | 39 |

chi2: 0.000, p: 0.9838517956, OR: 2.58333

| | < cutoff | other genes | Total |
|---|---|---|---|
| In Oligodendrocyte | 2 (1.62) | 5 (5.38) | 7 |
| Not IN Oligodendrocyte | 7 (7.38) | 25 (24.62) | 32 |
| Total | 9 | 30 | 39 |

chi2: 0.013, p: 0.9090223661, OR: 1.42857

| | > cutoff | other genes | Total |
|---|---|---|---|
| In Oligodendrocyte | 2 (0.90) | 5 (6.10) | 7 |
| Not IN Oligodendrocyte | 3 (4.10) | 29 (27.90) | 32 |
| Total | 5 | 34 | 39 |

chi2: 0.566, p: 0.4520158046, OR: 3.86667

| | < cutoff | other genes | Total |
|---|---|---|---|
| In Neuron | 6 (4.15) | 12 (13.85) | 18 |
| Not IN Neuron | 3 (4.85) | 18 (16.15) | 21 |
| Total | 9 | 30 | 39 |

chi2: 1.053, p: 0.3047618940, OR: 3.00000

| | > cutoff | other genes | Total |
|---|---|---|---|
| In Neuron | 1 (2.31) | 17 (15.69) | 18 |
| Not IN Neuron | 4 (2.69) | 17 (18.31) | 21 |
| Total | 5 | 34 | 39 |

chi2: 0.602, p: 0.4377376176, OR: 0.25000

| | < cutoff | other genes | Total |
|---|---|---|---|
| In Endothelia | 1 (0.69) | 2 (2.31) | 3 |
| Not IN Endothelia | 8 (8.31) | 28 (27.69) | 36 |
| Total | 9 | 30 | 39 |

chi2: 0.075, p: 0.7838667032, OR: 1.75000

| | > cutoff | other genes | Total |
|---|---|---|---|
| In Endothelia | 0 (0.38) | 3 (2.62) | 3 |
| Not IN Endothelia | 5 (4.62) | 31 (31.38) | 36 |
| Total | 5 | 34 | 39 |

chi2: 0.043, p: 0.8356975845, OR: 0.00000

### Five Celltypes Distribution of Average dN/dS Scores of Genes Related to developmental maturation

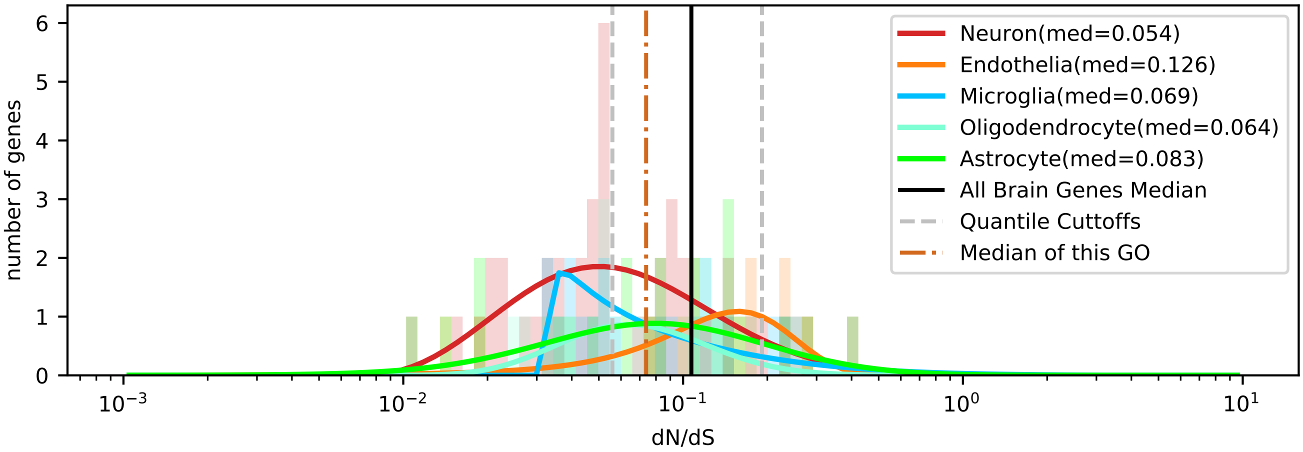


| | < cutoff | other genes | Total |
|---|---|---|---|
| In Astrocyte | 7 (9.65) | 15 (12.35) | 22 |
| Not IN Astrocyte | 43 (40.35) | 49 (51.65) | 92 |
| Total | 50 | 64 | 114 |

chi2: 1.057, p: 0.3040108983, OR: 0.53178

| | > cutoff | other genes | Total |
|---|---|---|---|
| In Astrocyte | 3 (2.70) | 19 (19.30) | 22 |
| Not IN Astrocyte | 11 (11.30) | 81 (80.70) | 92 |
| Total | 14 | 100 | 114 |

chi2: 0.021, p: 0.8840120803, OR: 1.16268

| | < cutoff | other genes | Total |
|---|---|---|---|
| In Microglia | 10 (8.77) | 10 (11.23) | 20 |
| Not IN Microglia | 40 (41.23) | 54 (52.77) | 94 |
| Total | 50 | 64 | 114 |

chi2: 0.131, p: 0.7178698193, OR: 1.35000

| | > cutoff | other genes | Total |
|---|---|---|---|
| In Microglia | 3 (2.46) | 17 (17.54) | 20 |
| Not IN Microglia | 11 (11.54) | 83 (82.46) | 94 |
| Total | 14 | 100 | 114 |

chi2: 0.001, p: 0.9737492826, OR: 1.33155

| | < cutoff | other genes | Total |
|---|---|---|---|
| In Oligodendrocyte | 6 (6.14) | 8 (7.86) | 14 |
| Not IN Oligodendrocyte | 44 (43.86) | 56 (56.14) | 100 |
| Total | 50 | 64 | 114 |

chi2: 0.043, p: 0.8361486725, OR: 0.95455

| | > cutoff | other genes | Total |
|---|---|---|---|
| In Oligodendrocyte | 0 (1.72) | 14 (12.28) | 14 |
| Not IN Oligodendrocyte | 14 (12.28) | 86 (87.72) | 100 |
| Total | 14 | 100 | 114 |

chi2: 1.124, p: 0.2891075958, OR: 0.00000

| | < cutoff | other genes | Total |
|---|---|---|---|
| In Neuron | 24 (17.98) | 17 (23.02) | 41 |
| Not IN Neuron | 26 (32.02) | 47 (40.98) | 73 |
| Total | 50 | 64 | 114 |

chi2: 4.709, p: 0.0300011724, OR: 2.55204

| | > cutoff | other genes | Total |
|---|---|---|---|
| In Neuron | 4 (5.04) | 37 (35.96) | 41 |
| Not IN Neuron | 10 (8.96) | 63 (64.04) | 73 |
| Total | 14 | 100 | 114 |

chi2: 0.101, p: 0.7503528445, OR: 0.68108

| | < cutoff | other genes | Total |
|---|---|---|---|
| In Endothelia | 3 (7.46) | 14 (9.54) | 17 |
| Not IN Endothelia | 47 (42.54) | 50 (54.46) | 97 |
| Total | 50 | 64 | 114 |

chi2: 4.394, p: 0.0360597736, OR: 0.22796

| | > cutoff | other genes | Total |
|---|---|---|---|
| In Endothelia | 4 (2.09) | 13 (14.91) | 17 |
| Not IN Endothelia | 10 (11.91) | 87 (85.09) | 97 |
| Total | 14 | 100 | 114 |

chi2: 1.280, p: 0.2579001628, OR: 2.67692

### Five Celltypes Distribution of Average dN/dS Scores of Genes Related to DNA metabolic process

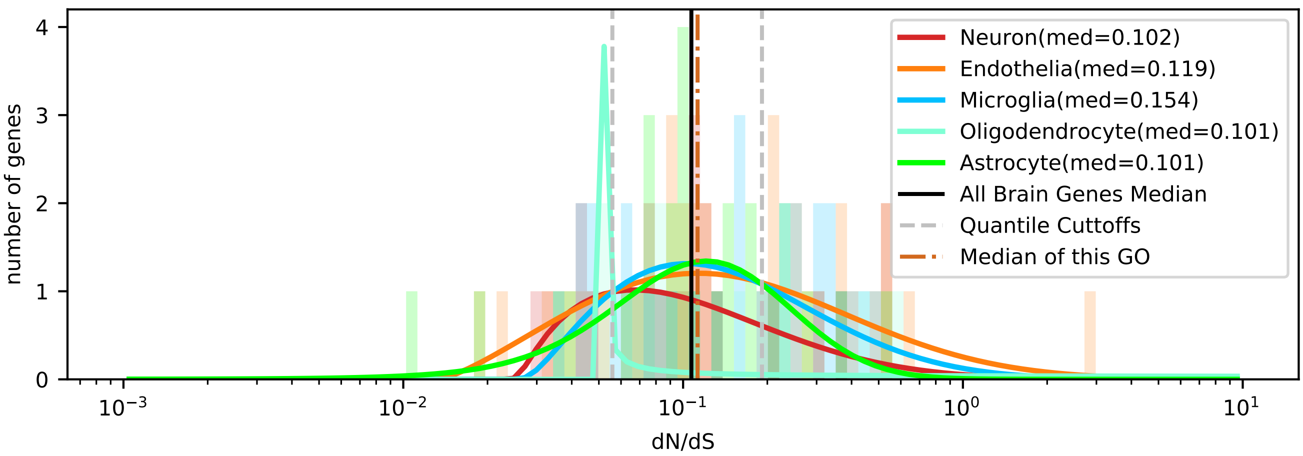


| | < cutoff | other genes | Total |
|---|---|---|---|
| In Astrocyte | 4 (5.18) | 23 (21.82) | 27 |
| Not IN Astrocyte | 20 (18.82) | 78 (79.18) | 98 |
| Total | 24 | 101 | 125 |

chi2: 0.142, p: 0.7058389157, OR: 0.67826

| | > cutoff | other genes | Total |
|---|---|---|---|
| In Astrocyte | 6 (9.29) | 21 (17.71) | 27 |
| Not IN Astrocyte | 37 (33.71) | 61 (64.29) | 98 |
| Total | 43 | 82 | 125 |

chi2: 1.627, p: 0.2020894701, OR: 0.47104

| | < cutoff | other genes | Total |
|---|---|---|---|
| In Microglia | 5 (5.76) | 25 (24.24) | 30 |
| Not IN Microglia | 19 (18.24) | 76 (76.76) | 95 |
| Total | 24 | 101 | 125 |

chi2: 0.019, p: 0.8900467421, OR: 0.80000

| | > cutoff | other genes | Total |
|---|---|---|---|
| In Microglia | 13 (10.32) | 17 (19.68) | 30 |
| Not IN Microglia | 30 (32.68) | 65 (62.32) | 95 |
| Total | 43 | 82 | 125 |

chi2: 0.924, p: 0.3365137369, OR: 1.65686

| | < cutoff | other genes | Total |
|---|---|---|---|
| In Oligodendrocyte | 2 (1.92) | 8 (8.08) | 10 |
| Not IN Oligodendrocyte | 22 (22.08) | 93 (92.92) | 115 |
| Total | 24 | 101 | 125 |

chi2: 0.124, p: 0.7251684948, OR: 1.05682

| | > cutoff | other genes | Total |
|---|---|---|---|
| In Oligodendrocyte | 3 (3.44) | 7 (6.56) | 10 |
| Not IN Oligodendrocyte | 40 (39.56) | 75 (75.44) | 115 |
| Total | 43 | 82 | 125 |

chi2: 0.002, p: 0.9667845027, OR: 0.80357

| | < cutoff | other genes | Total |
|---|---|---|---|
| In Neuron | 5 (4.22) | 17 (17.78) | 22 |
| Not IN Neuron | 19 (19.78) | 84 (83.22) | 103 |
| Total | 24 | 101 | 125 |

chi2: 0.027, p: 0.8692743406, OR: 1.30031

| | > cutoff | other genes | Total |
|---|---|---|---|
| In Neuron | 5 (7.57) | 17 (14.43) | 22 |
| Not IN Neuron | 38 (35.43) | 65 (67.57) | 103 |
| Total | 43 | 82 | 125 |

chi2: 1.045, p: 0.3065653287, OR: 0.50310

| | < cutoff | other genes | Total |
|---|---|---|---|
| In Endothelia | 8 (6.91) | 28 (29.09) | 36 |
| Not IN Endothelia | 16 (17.09) | 73 (71.91) | 89 |
| Total | 24 | 101 | 125 |

chi2: 0.087, p: 0.7680936079, OR: 1.30357

| | > cutoff | other genes | Total |
|---|---|---|---|
| In Endothelia | 16 (12.38) | 20 (23.62) | 36 |
| Not IN Endothelia | 27 (30.62) | 62 (58.38) | 89 |
| Total | 43 | 82 | 125 |

chi2: 1.679, p: 0.1951090909, OR: 1.83704

### Five Celltypes Distribution of Average dN/dS Scores of Genes Related to embryo development

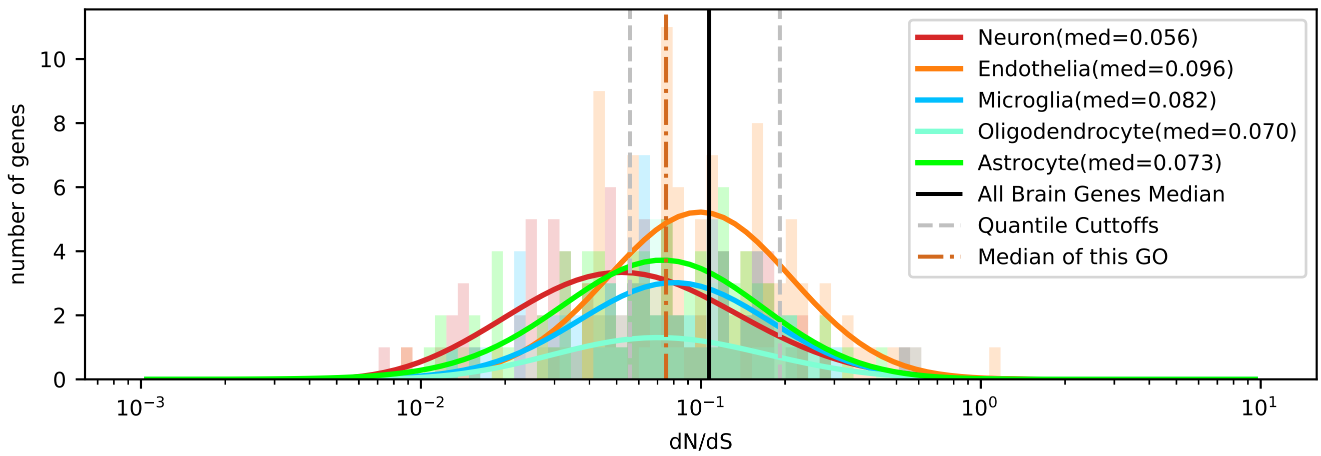


| | < cutoff | other genes | Total |
|---|---|---|---|
| In Astrocyte | 31 (29.45) | 53 (54.55) | 84 |
| Not IN Astrocyte | 98 (99.55) | 186 (184.45) | 284 |
| Total | 129 | 239 | 368 |

chi2: 0.075, p: 0.7837389293, OR: 1.11013

| | > cutoff | other genes | Total |
|---|---|---|---|
| In Astrocyte | 9 (10.73) | 75 (73.27) | 84 |
| Not IN Astrocyte | 38 (36.27) | 246 (247.73) | 284 |
| Total | 47 | 321 | 368 |

chi2: 0.209, p: 0.6476363384, OR: 0.77684

| | < cutoff | other genes | Total |
|---|---|---|---|
| In Microglia | 19 (22.43) | 45 (41.57) | 64 |
| Not IN Microglia | 110 (106.57) | 194 (197.43) | 304 |
| Total | 129 | 239 | 368 |

chi2: 0.716, p: 0.3975997789, OR: 0.74465

| | > cutoff | other genes | Total |
|---|---|---|---|
| In Microglia | 9 (8.17) | 55 (55.83) | 64 |
| Not IN Microglia | 38 (38.83) | 266 (265.17) | 304 |
| Total | 47 | 321 | 368 |

chi2: 0.018, p: 0.8931162133, OR: 1.14545

| | < cutoff | other genes | Total |
|---|---|---|---|
| In Oligodendrocyte | 11 (10.52) | 19 (19.48) | 30 |
| Not IN Oligodendrocyte | 118 (118.48) | 220 (219.52) | 338 |
| Total | 129 | 239 | 368 |

chi2: 0.000, p: 0.9948060308, OR: 1.07939

| | > cutoff | other genes | Total |
|---|---|---|---|
| In Oligodendrocyte | 4 (3.83) | 26 (26.17) | 30 |
| Not IN Oligodendrocyte | 43 (43.17) | 295 (294.83) | 338 |
| Total | 47 | 321 | 368 |

chi2: 0.036, p: 0.8499216636, OR: 1.05546

| | < cutoff | other genes | Total |
|---|---|---|---|
| In Neuron | 43 (29.10) | 40 (53.90) | 83 |
| Not IN Neuron | 86 (99.90) | 199 (185.10) | 285 |
| Total | 129 | 239 | 368 |

chi2: 12.279, p: 0.0004581081, OR: 2.48750

| | > cutoff | other genes | Total |
|---|---|---|---|
| In Neuron | 6 (10.60) | 77 (72.40) | 83 |
| Not IN Neuron | 41 (36.40) | 244 (248.60) | 285 |
| Total | 47 | 321 | 368 |

chi2: 2.348, p: 0.1254428287, OR: 0.46373

| | < cutoff | other genes | Total |
|---|---|---|---|
| In Endothelia | 25 (37.51) | 82 (69.49) | 107 |
| Not IN Endothelia | 104 (91.49) | 157 (169.51) | 261 |
| Total | 129 | 239 | 368 |

chi2: 8.346, p: 0.0038651469, OR: 0.46025

| | > cutoff | other genes | Total |
|---|---|---|---|
| In Endothelia | 19 (13.67) | 88 (93.33) | 107 |
| Not IN Endothelia | 28 (33.33) | 233 (227.67) | 261 |
| Total | 47 | 321 | 368 |

chi2: 2.764, p: 0.0963936863, OR: 1.79667

### Five Celltypes Distribution of Average dN/dS Scores of Genes Related to extracellular matrix organization

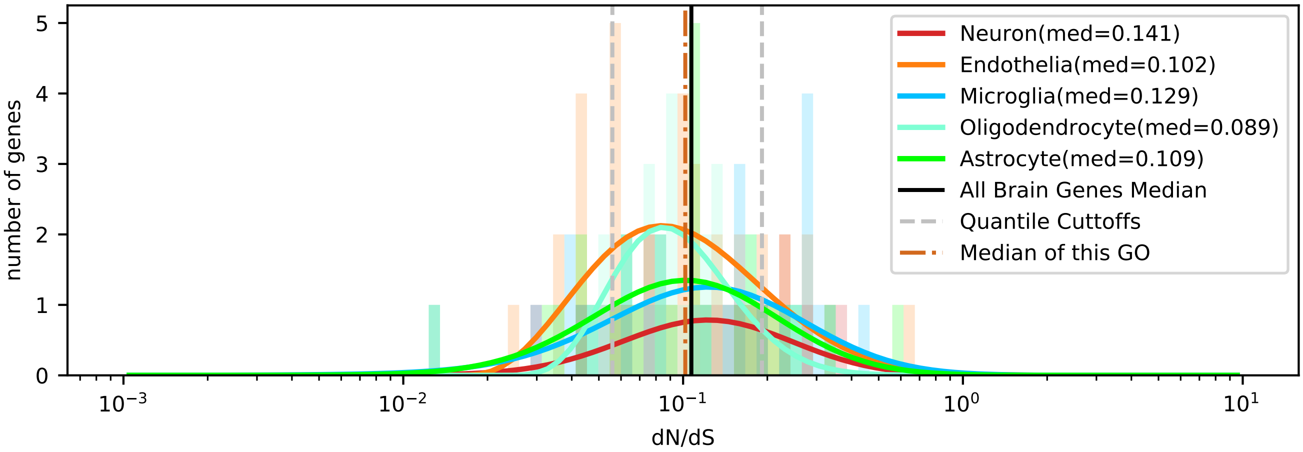


| | < cutoff | other genes | Total |
|---|---|---|---|
| In Astrocyte | 7 (5.64) | 21 (22.36) | 28 |
| Not IN Astrocyte | 21 (22.36) | 90 (88.64) | 111 |
| Total | 28 | 111 | 139 |

chi2: 0.205, p: 0.6503262878, OR: 1.42857

| | > cutoff | other genes | Total |
|---|---|---|---|
| In Astrocyte | 5 (5.84) | 23 (22.16) | 28 |
| Not IN Astrocyte | 24 (23.16) | 87 (87.84) | 111 |
| Total | 29 | 110 | 139 |

chi2: 0.032, p: 0.8588371466, OR: 0.78804

| | < cutoff | other genes | Total |
|---|---|---|---|
| In Microglia | 7 (5.64) | 21 (22.36) | 28 |
| Not IN Microglia | 21 (22.36) | 90 (88.64) | 111 |
| Total | 28 | 111 | 139 |

chi2: 0.205, p: 0.6503262878, OR: 1.42857

| | > cutoff | other genes | Total |
|---|---|---|---|
| In Microglia | 9 (5.84) | 19 (22.16) | 28 |
| Not IN Microglia | 20 (23.16) | 91 (87.84) | 111 |
| Total | 29 | 110 | 139 |

chi2: 1.914, p: 0.1665049378, OR: 2.15526

| | < cutoff | other genes | Total |
|---|---|---|---|
| In Oligodendrocyte | 4 (5.44) | 23 (21.56) | 27 |
| Not IN Oligodendrocyte | 24 (22.56) | 88 (89.44) | 112 |
| Total | 28 | 111 | 139 |

chi2: 0.252, p: 0.6157628084, OR: 0.63768

| | > cutoff | other genes | Total |
|---|---|---|---|
| In Oligodendrocyte | 2 (5.63) | 25 (21.37) | 27 |
| Not IN Oligodendrocyte | 27 (23.37) | 85 (88.63) | 112 |
| Total | 29 | 110 | 139 |

chi2: 2.733, p: 0.0983026142, OR: 0.25185

| | < cutoff | other genes | Total |
|---|---|---|---|
| In Neuron | 2 (3.02) | 13 (11.98) | 15 |
| Not IN Neuron | 26 (24.98) | 98 (99.02) | 124 |
| Total | 28 | 111 | 139 |

chi2: 0.126, p: 0.7222096343, OR: 0.57988

| | > cutoff | other genes | Total |
|---|---|---|---|
| In Neuron | 4 (3.13) | 11 (11.87) | 15 |
| Not IN Neuron | 25 (25.87) | 99 (98.13) | 124 |
| Total | 29 | 110 | 139 |

chi2: 0.062, p: 0.8031551311, OR: 1.44000

| | < cutoff | other genes | Total |
|---|---|---|---|
| In Endothelia | 8 (8.26) | 33 (32.74) | 41 |
| Not IN Endothelia | 20 (19.74) | 78 (78.26) | 98 |
| Total | 28 | 111 | 139 |

chi2: 0.012, p: 0.9110095532, OR: 0.94545

| | > cutoff | other genes | Total |
|---|---|---|---|
| In Endothelia | 9 (8.55) | 32 (32.45) | 41 |
| Not IN Endothelia | 20 (20.45) | 78 (77.55) | 98 |
| Total | 29 | 110 | 139 |

chi2: 0.001, p: 0.9802955524, OR: 1.09688

### Five Celltypes Distribution of Average dN/dS Scores of Genes Related to generation of precursor metabolites and energy

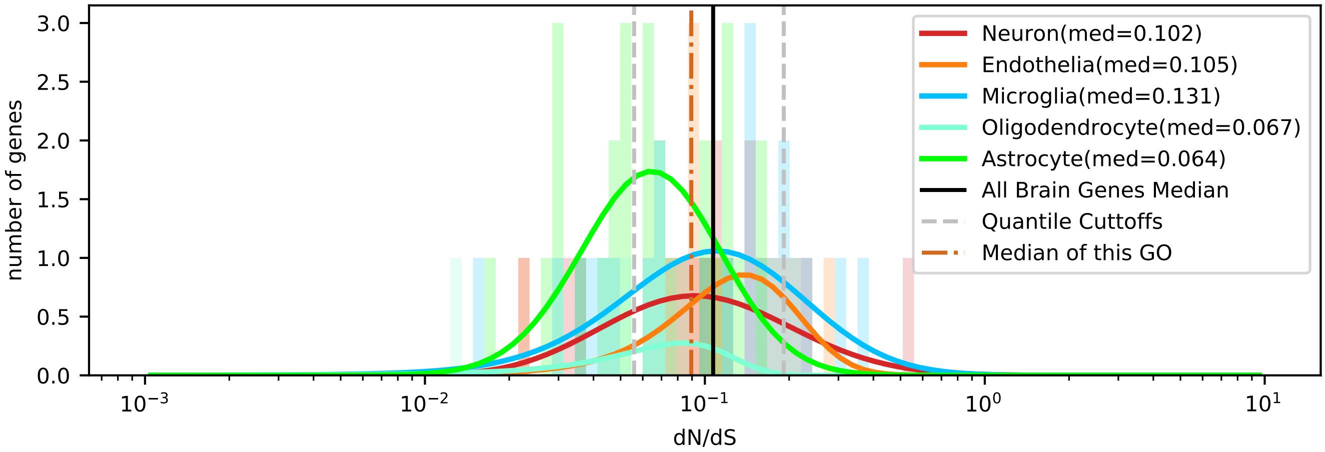


| | < cutoff | other genes | Total |
|---|---|---|---|
| In Astrocyte | 12 (8.20) | 15 (18.80) | 27 |
| Not IN Astrocyte | 12 (15.80) | 40 (36.20) | 52 |
| Total | 24 | 55 | 79 |

chi2: 2.893, p: 0.0889823835, OR: 2.66667

| | > cutoff | other genes | Total |
|---|---|---|---|
| In Astrocyte | 0 (3.42) | 27 (23.58) | 27 |
| Not IN Astrocyte | 10 (6.58) | 42 (45.42) | 52 |
| Total | 10 | 69 | 79 |

chi2: 4.333, p: 0.0373881812, OR: 0.00000

| | < cutoff | other genes | Total |
|---|---|---|---|
| In Microglia | 6 (6.68) | 16 (15.32) | 22 |
| Not IN Microglia | 18 (17.32) | 39 (39.68) | 57 |
| Total | 24 | 55 | 79 |

chi2: 0.010, p: 0.9202078721, OR: 0.81250

| | > cutoff | other genes | Total |
|---|---|---|---|
| In Microglia | 5 (2.78) | 17 (19.22) | 22 |
| Not IN Microglia | 5 (7.22) | 52 (49.78) | 57 |
| Total | 10 | 69 | 79 |

chi2: 1.676, p: 0.1954130619, OR: 3.05882

| | < cutoff | other genes | Total |
|---|---|---|---|
| In Oligodendrocyte | 2 (1.22) | 2 (2.78) | 4 |
| Not IN Oligodendrocyte | 22 (22.78) | 53 (52.22) | 75 |
| Total | 24 | 55 | 79 |

chi2: 0.101, p: 0.7506396605, OR: 2.40909

| | > cutoff | other genes | Total |
|---|---|---|---|
| In Oligodendrocyte | 0 (0.51) | 4 (3.49) | 4 |
| Not IN Oligodendrocyte | 10 (9.49) | 65 (65.51) | 75 |
| Total | 10 | 69 | 79 |

chi2: 0.000, p: 0.9922065181, OR: 0.00000

| | < cutoff | other genes | Total |
|---|---|---|---|
| In Neuron | 3 (4.25) | 11 (9.75) | 14 |
| Not IN Neuron | 21 (19.75) | 44 (45.25) | 65 |
| Total | 24 | 55 | 79 |

chi2: 0.233, p: 0.6294306522, OR: 0.57143

| | > cutoff | other genes | Total |
|---|---|---|---|
| In Neuron | 2 (1.77) | 12 (12.23) | 14 |
| Not IN Neuron | 8 (8.23) | 57 (56.77) | 65 |
| Total | 10 | 69 | 79 |

chi2: 0.058, p: 0.8094304707, OR: 1.18750

| | < cutoff | other genes | Total |
|---|---|---|---|
| In Endothelia | 1 (3.65) | 11 (8.35) | 12 |
| Not IN Endothelia | 23 (20.35) | 44 (46.65) | 67 |
| Total | 24 | 55 | 79 |

chi2: 2.139, p: 0.1436297114, OR: 0.17391

| | > cutoff | other genes | Total |
|---|---|---|---|
| In Endothelia | 3 (1.52) | 9 (10.48) | 12 |
| Not IN Endothelia | 7 (8.48) | 60 (58.52) | 67 |
| Total | 10 | 69 | 79 |

chi2: 0.855, p: 0.3550534907, OR: 2.85714

### Five Celltypes Distribution of Average dN/dS Scores of Genes Related to growth

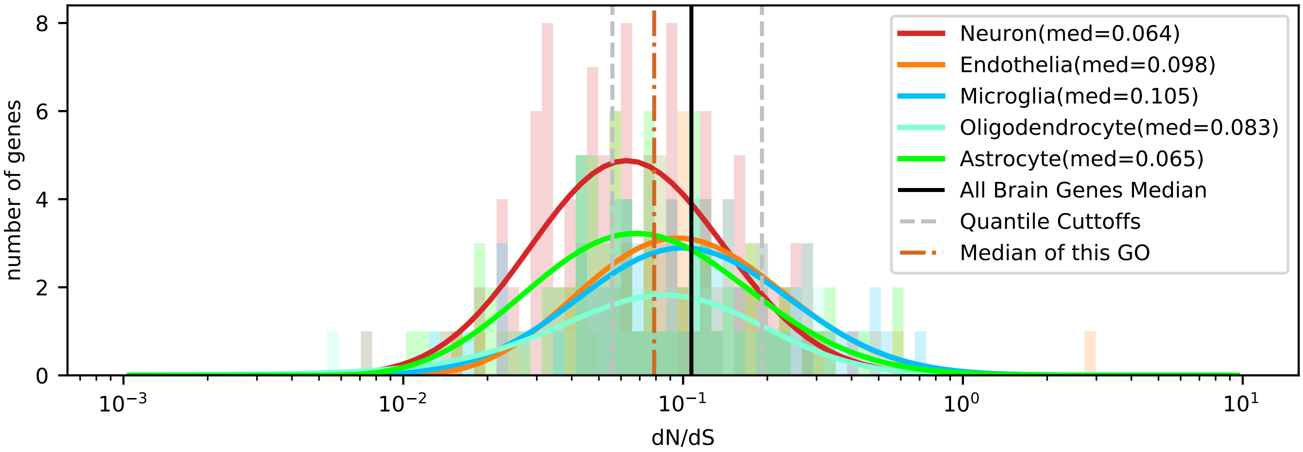


| | < cutoff | other genes | Total |
|---|---|---|---|
| In Astrocyte | 31 (25.74) | 46 (51.26) | 77 |
| Not IN Astrocyte | 89 (94.26) | 193 (187.74) | 282 |
| Total | 120 | 239 | 359 |

chi2: 1.685, p: 0.1943056710, OR: 1.46141

| | > cutoff | other genes | Total |
|---|---|---|---|
| In Astrocyte | 10 (11.80) | 67 (65.20) | 77 |
| Not IN Astrocyte | 45 (43.20) | 237 (238.80) | 282 |
| Total | 55 | 304 | 359 |

chi2: 0.214, p: 0.6434424077, OR: 0.78607

| | < cutoff | other genes | Total |
|---|---|---|---|
| In Microglia | 15 (22.40) | 52 (44.60) | 67 |
| Not IN Microglia | 105 (97.60) | 187 (194.40) | 292 |
| Total | 120 | 239 | 359 |

chi2: 3.921, p: 0.0476899213, OR: 0.51374

| | > cutoff | other genes | Total |
|---|---|---|---|
| In Microglia | 16 (10.26) | 51 (56.74) | 67 |
| Not IN Microglia | 39 (44.74) | 253 (247.26) | 292 |
| Total | 55 | 304 | 359 |

chi2: 3.877, p: 0.0489547183, OR: 2.03519

| | < cutoff | other genes | Total |
|---|---|---|---|
| In Oligodendrocyte | 14 (14.37) | 29 (28.63) | 43 |
| Not IN Oligodendrocyte | 106 (105.63) | 210 (210.37) | 316 |
| Total | 120 | 239 | 359 |

chi2: 0.002, p: 0.9651668048, OR: 0.95641

| | > cutoff | other genes | Total |
|---|---|---|---|
| In Oligodendrocyte | 7 (6.59) | 36 (36.41) | 43 |
| Not IN Oligodendrocyte | 48 (48.41) | 268 (267.59) | 316 |
| Total | 55 | 304 | 359 |

chi2: 0.002, p: 0.9684144491, OR: 1.08565

| | < cutoff | other genes | Total |
|---|---|---|---|
| In Neuron | 48 (34.43) | 55 (68.57) | 103 |
| Not IN Neuron | 72 (85.57) | 184 (170.43) | 256 |
| Total | 120 | 239 | 359 |

chi2: 10.453, p: 0.0012244129, OR: 2.23030

| | > cutoff | other genes | Total |
|---|---|---|---|
| In Neuron | 10 (15.78) | 93 (87.22) | 103 |
| Not IN Neuron | 45 (39.22) | 211 (216.78) | 256 |
| Total | 55 | 304 | 359 |

chi2: 2.926, p: 0.0871798551, OR: 0.50418

| | < cutoff | other genes | Total |
|---|---|---|---|
| In Endothelia | 12 (23.06) | 57 (45.94) | 69 |
| Not IN Endothelia | 108 (96.94) | 182 (193.06) | 290 |
| Total | 120 | 239 | 359 |

chi2: 8.997, p: 0.0027035673, OR: 0.35478

| | > cutoff | other genes | Total |
|---|---|---|---|
| In Endothelia | 12 (10.57) | 57 (58.43) | 69 |
| Not IN Endothelia | 43 (44.43) | 247 (245.57) | 290 |
| Total | 55 | 304 | 359 |

chi2: 0.119, p: 0.7297464165, OR: 1.20930

### Five Celltypes Distribution of Average dN/dS Scores of Genes Related to homeostatic process

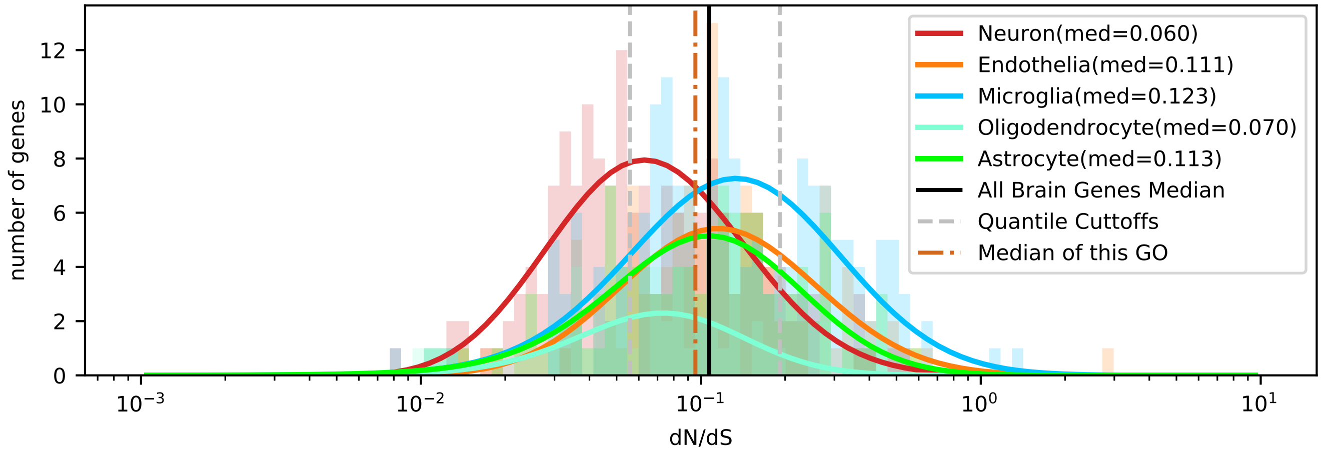


| | < cutoff | other genes | Total |
|---|---|---|---|
| In Astrocyte | 28 (30.60) | 83 (80.40) | 111 |
| Not IN Astrocyte | 141 (138.40) | 361 (363.60) | 502 |
| Total | 169 | 444 | 613 |

chi2: 0.243, p: 0.6217562024, OR: 0.86371

| | > cutoff | other genes | Total |
|---|---|---|---|
| In Astrocyte | 23 (23.36) | 88 (87.64) | 111 |
| Not IN Astrocyte | 106 (105.64) | 396 (396.36) | 502 |
| Total | 129 | 484 | 613 |

chi2: 0.001, p: 0.9710358955, OR: 0.97642

| | < cutoff | other genes | Total |
|---|---|---|---|
| In Microglia | 28 (47.42) | 144 (124.58) | 172 |
| Not IN Microglia | 141 (121.58) | 300 (319.42) | 441 |
| Total | 169 | 444 | 613 |

chi2: 14.486, p: 0.0001411950, OR: 0.41371

| | > cutoff | other genes | Total |
|---|---|---|---|
| In Microglia | 61 (36.20) | 111 (135.80) | 172 |
| Not IN Microglia | 68 (92.80) | 373 (348.20) | 441 |
| Total | 129 | 484 | 613 |

chi2: 28.731, p: 0.0000000832, OR: 3.01444

| | < cutoff | other genes | Total |
|---|---|---|---|
| In Oligodendrocyte | 17 (11.85) | 26 (31.15) | 43 |
| Not IN Oligodendrocyte | 152 (157.15) | 418 (412.85) | 570 |
| Total | 169 | 444 | 613 |

chi2: 2.703, p: 0.1001874125, OR: 1.79808

| | > cutoff | other genes | Total |
|---|---|---|---|
| In Oligodendrocyte | 3 (9.05) | 40 (33.95) | 43 |
| Not IN Oligodendrocyte | 126 (119.95) | 444 (450.05) | 570 |
| Total | 129 | 484 | 613 |

chi2: 4.635, p: 0.0313313461, OR: 0.26429

| | < cutoff | other genes | Total |
|---|---|---|---|
| In Neuron | 80 (46.87) | 90 (123.13) | 170 |
| Not IN Neuron | 89 (122.13) | 354 (320.87) | 443 |
| Total | 169 | 444 | 613 |

chi2: 43.406, p: 0.0000000000, OR: 3.53558

| | > cutoff | other genes | Total |
|---|---|---|---|
| In Neuron | 16 (35.77) | 154 (134.23) | 170 |
| Not IN Neuron | 113 (93.23) | 330 (349.77) | 443 |
| Total | 129 | 484 | 613 |

chi2: 18.200, p: 0.0000198854, OR: 0.30341

| | < cutoff | other genes | Total |
|---|---|---|---|
| In Endothelia | 16 (32.26) | 101 (84.74) | 117 |
| Not IN Endothelia | 153 (136.74) | 343 (359.26) | 496 |
| Total | 169 | 444 | 613 |

chi2: 13.132, p: 0.0002902397, OR: 0.35514

| | > cutoff | other genes | Total |
|---|---|---|---|
| In Endothelia | 26 (24.62) | 91 (92.38) | 117 |
| Not IN Endothelia | 103 (104.38) | 393 (391.62) | 496 |
| Total | 129 | 484 | 613 |

chi2: 0.049, p: 0.8247067463, OR: 1.09015

### Five Celltypes Distribution of Average dN/dS Scores of Genes Related to immune system process

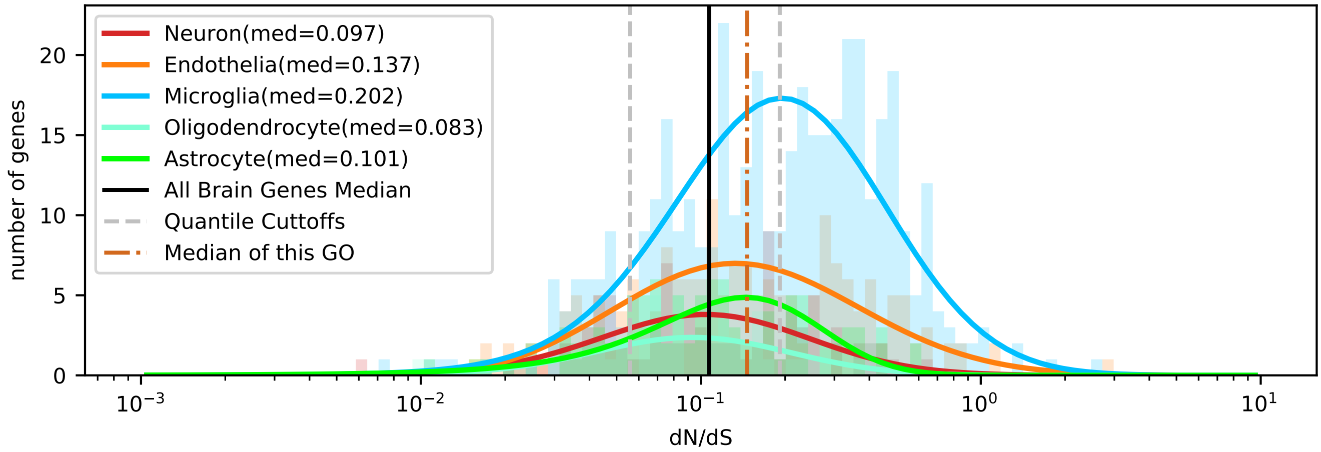


| | < cutoff | other genes | Total |
|---|---|---|---|
| In Astrocyte | 22 (16.83) | 77 (82.17) | 99 |
| Not IN Astrocyte | 121 (126.17) | 621 (615.83) | 742 |
| Total | 143 | 698 | 841 |

chi2: 1.767, p: 0.1838054975, OR: 1.46635

| | > cutoff | other genes | Total |
|---|---|---|---|
| In Astrocyte | 26 (40.85) | 73 (58.15) | 99 |
| Not IN Astrocyte | 321 (306.15) | 421 (435.85) | 742 |
| Total | 347 | 494 | 841 |

chi2: 9.724, p: 0.0018183703, OR: 0.46712

| | < cutoff | other genes | Total |
|---|---|---|---|
| In Microglia | 48 (70.73) | 368 (345.27) | 416 |
| Not IN Microglia | 95 (72.27) | 330 (352.73) | 425 |
| Total | 143 | 698 | 841 |

chi2: 16.664, p: 0.0000446171, OR: 0.45309

| | > cutoff | other genes | Total |
|---|---|---|---|
| In Microglia | 215 (171.64) | 201 (244.36) | 416 |
| Not IN Microglia | 132 (175.36) | 293 (249.64) | 425 |
| Total | 347 | 494 | 841 |

chi2: 36.048, p: 0.0000000019, OR: 2.37430

| | < cutoff | other genes | Total |
|---|---|---|---|
| In Oligodendrocyte | 14 (8.84) | 38 (43.16) | 52 |
| Not IN Oligodendrocyte | 129 (134.16) | 660 (654.84) | 789 |
| Total | 143 | 698 | 841 |

chi2: 3.152, p: 0.0758488063, OR: 1.88494

| | > cutoff | other genes | Total |
|---|---|---|---|
| In Oligodendrocyte | 9 (21.46) | 43 (30.54) | 52 |
| Not IN Oligodendrocyte | 338 (325.54) | 451 (463.46) | 789 |
| Total | 347 | 494 | 841 |

chi2: 12.089, p: 0.0005072789, OR: 0.27928

| | < cutoff | other genes | Total |
|---|---|---|---|
| In Neuron | 26 (15.13) | 63 (73.87) | 89 |
| Not IN Neuron | 117 (127.87) | 635 (624.13) | 752 |
| Total | 143 | 698 | 841 |

chi2: 9.569, p: 0.0019785806, OR: 2.23986

| | > cutoff | other genes | Total |
|---|---|---|---|
| In Neuron | 24 (36.72) | 65 (52.28) | 89 |
| Not IN Neuron | 323 (310.28) | 429 (441.72) | 752 |
| Total | 347 | 494 | 841 |

chi2: 7.744, p: 0.0053877535, OR: 0.49040

| | < cutoff | other genes | Total |
|---|---|---|---|
| In Endothelia | 33 (31.46) | 152 (153.54) | 185 |
| Not IN Endothelia | 110 (111.54) | 546 (544.46) | 656 |
| Total | 143 | 698 | 841 |

chi2: 0.053, p: 0.8171495956, OR: 1.07763

| | > cutoff | other genes | Total |
|---|---|---|---|
| In Endothelia | 73 (76.33) | 112 (108.67) | 185 |
| Not IN Endothelia | 274 (270.67) | 382 (385.33) | 656 |
| Total | 347 | 494 | 841 |

chi2: 0.229, p: 0.6320586033, OR: 0.90869

### Five Celltypes Distribution of Average dN/dS Scores of Genes Related to lipid metabolic process

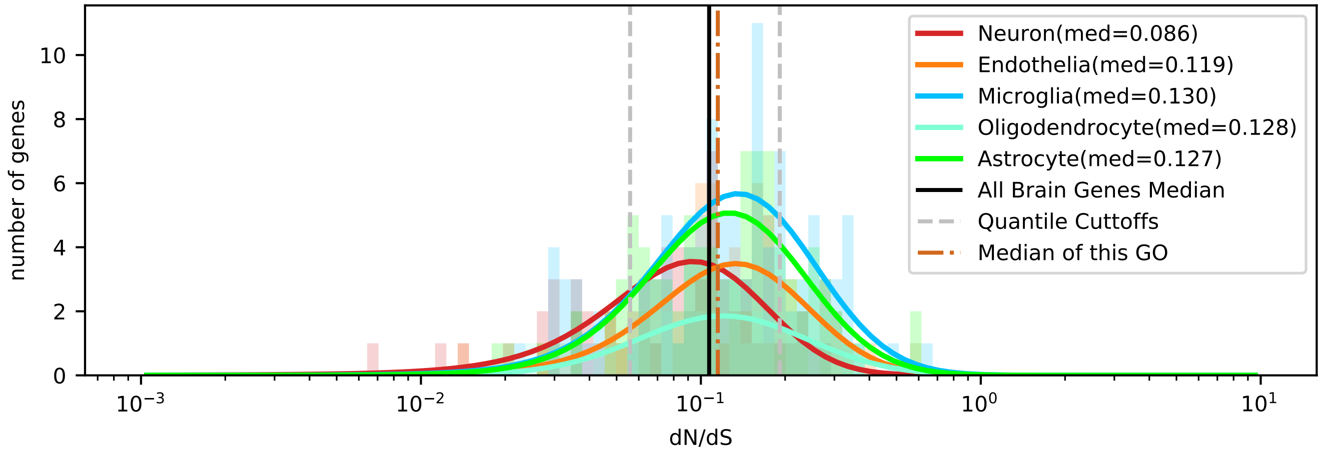


| | < cutoff | other genes | Total |
|---|---|---|---|
| In Astrocyte | 12 (15.09) | 78 (74.91) | 90 |
| Not IN Astrocyte | 47 (43.91) | 215 (218.09) | 262 |
| Total | 59 | 293 | 352 |

chi2: 0.715, p: 0.3977585043, OR: 0.70376

| | > cutoff | other genes | Total |
|---|---|---|---|
| In Astrocyte | 20 (20.45) | 70 (69.55) | 90 |
| Not IN Astrocyte | 60 (59.55) | 202 (202.45) | 262 |
| Total | 80 | 272 | 352 |

chi2: 0.000, p: 0.9894265353, OR: 0.96190

| | < cutoff | other genes | Total |
|---|---|---|---|
| In Microglia | 15 (17.26) | 88 (85.74) | 103 |
| Not IN Microglia | 44 (41.74) | 205 (207.26) | 249 |
| Total | 59 | 293 | 352 |

chi2: 0.306, p: 0.5800369274, OR: 0.79416

| | > cutoff | other genes | Total |
|---|---|---|---|
| In Microglia | 29 (23.41) | 74 (79.59) | 103 |
| Not IN Microglia | 51 (56.59) | 198 (192.41) | 249 |
| Total | 80 | 272 | 352 |

chi2: 2.025, p: 0.1546822973, OR: 1.52146

| | < cutoff | other genes | Total |
|---|---|---|---|
| In Oligodendrocyte | 5 (5.87) | 30 (29.13) | 35 |
| Not IN Oligodendrocyte | 54 (53.13) | 263 (263.87) | 317 |
| Total | 59 | 293 | 352 |

chi2: 0.031, p: 0.8612696312, OR: 0.81173

| | > cutoff | other genes | Total |
|---|---|---|---|
| In Oligodendrocyte | 10 (7.95) | 25 (27.05) | 35 |
| Not IN Oligodendrocyte | 70 (72.05) | 247 (244.95) | 317 |
| Total | 80 | 272 | 352 |

chi2: 0.431, p: 0.5112667417, OR: 1.41143

| | < cutoff | other genes | Total |
|---|---|---|---|
| In Neuron | 19 (10.73) | 45 (53.27) | 64 |
| Not IN Neuron | 40 (48.27) | 248 (239.73) | 288 |
| Total | 59 | 293 | 352 |

chi2: 8.270, p: 0.0040315126, OR: 2.61778

| | > cutoff | other genes | Total |
|---|---|---|---|
| In Neuron | 6 (14.55) | 58 (49.45) | 64 |
| Not IN Neuron | 74 (65.45) | 214 (222.55) | 288 |
| Total | 80 | 272 | 352 |

chi2: 7.039, p: 0.0079763045, OR: 0.29916

| | < cutoff | other genes | Total |
|---|---|---|---|
| In Endothelia | 8 (10.06) | 52 (49.94) | 60 |
| Not IN Endothelia | 51 (48.94) | 241 (243.06) | 292 |
| Total | 59 | 293 | 352 |

chi2: 0.349, p: 0.5546685311, OR: 0.72700

| | > cutoff | other genes | Total |
|---|---|---|---|
| In Endothelia | 15 (13.64) | 45 (46.36) | 60 |
| Not IN Endothelia | 65 (66.36) | 227 (225.64) | 292 |
| Total | 80 | 272 | 352 |

chi2: 0.085, p: 0.7702014956, OR: 1.16410

### Five Celltypes Distribution of Average dN/dS Scores of Genes Related to locomotion

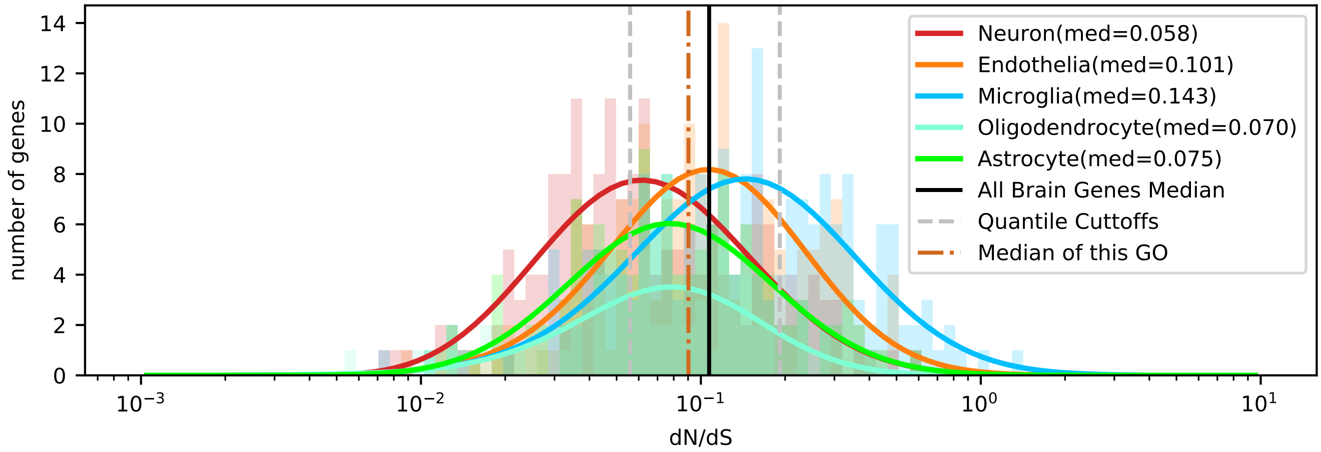


| | < cutoff | other genes | Total |
|---|---|---|---|
| In Astrocyte | 47 (41.51) | 88 (93.49) | 135 |
| Not IN Astrocyte | 187 (192.49) | 439 (433.51) | 626 |
| Total | 234 | 527 | 761 |

chi2: 1.052, p: 0.3049349191, OR: 1.25383

| | > cutoff | other genes | Total |
|---|---|---|---|
| In Astrocyte | 18 (29.63) | 117 (105.37) | 135 |
| Not IN Astrocyte | 149 (137.37) | 477 (488.63) | 626 |
| Total | 167 | 594 | 761 |

chi2: 6.507, p: 0.0107450042, OR: 0.49251

| | < cutoff | other genes | Total |
|---|---|---|---|
| In Microglia | 33 (59.65) | 161 (134.35) | 194 |
| Not IN Microglia | 201 (174.35) | 366 (392.65) | 567 |
| Total | 234 | 527 | 761 |

chi2: 22.222, p: 0.0000024284, OR: 0.37323

| | > cutoff | other genes | Total |
|---|---|---|---|
| In Microglia | 74 (42.57) | 120 (151.43) | 194 |
| Not IN Microglia | 93 (124.43) | 474 (442.57) | 567 |
| Total | 167 | 594 | 761 |

chi2: 38.632, p: 0.0000000005, OR: 3.14301

| | < cutoff | other genes | Total |
|---|---|---|---|
| In Oligodendrocyte | 27 (22.45) | 46 (50.55) | 73 |
| Not IN Oligodendrocyte | 207 (211.55) | 481 (476.45) | 688 |
| Total | 234 | 527 | 761 |

chi2: 1.169, p: 0.2796057439, OR: 1.36389

| | > cutoff | other genes | Total |
|---|---|---|---|
| In Oligodendrocyte | 7 (16.02) | 66 (56.98) | 73 |
| Not IN Oligodendrocyte | 160 (150.98) | 528 (537.02) | 688 |
| Total | 167 | 594 | 761 |

chi2: 6.421, p: 0.0112791030, OR: 0.35000

| | < cutoff | other genes | Total |
|---|---|---|---|
| In Neuron | 86 (55.35) | 94 (124.65) | 180 |
| Not IN Neuron | 148 (178.65) | 433 (402.35) | 581 |
| Total | 234 | 527 | 761 |

chi2: 31.067, p: 0.0000000249, OR: 2.67668

| | > cutoff | other genes | Total |
|---|---|---|---|
| In Neuron | 23 (39.50) | 157 (140.50) | 180 |
| Not IN Neuron | 144 (127.50) | 437 (453.50) | 581 |
| Total | 167 | 594 | 761 |

chi2: 10.876, p: 0.0009740676, OR: 0.44458

| | < cutoff | other genes | Total |
|---|---|---|---|
| In Endothelia | 41 (55.04) | 138 (123.96) | 179 |
| Not IN Endothelia | 193 (178.96) | 389 (403.04) | 582 |
| Total | 234 | 527 | 761 |

chi2: 6.290, p: 0.0121435568, OR: 0.59882

| | > cutoff | other genes | Total |
|---|---|---|---|
| In Endothelia | 45 (39.28) | 134 (139.72) | 179 |
| Not IN Endothelia | 122 (127.72) | 460 (454.28) | 582 |
| Total | 167 | 594 | 761 |

chi2: 1.161, p: 0.2811570831, OR: 1.26621

### Five Celltypes Distribution of Average dN/dS Scores of Genes Related to membrane organization

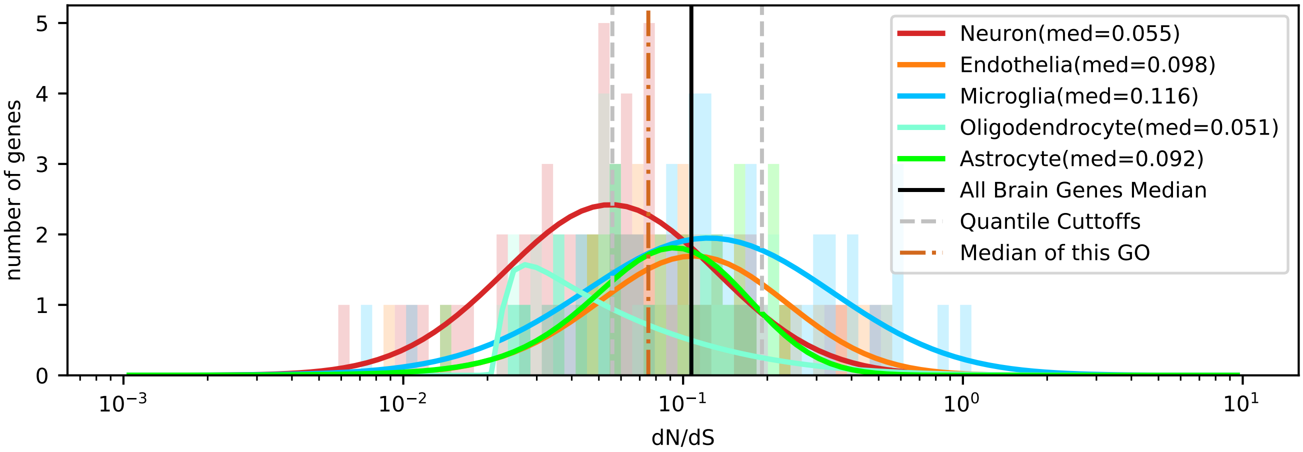


| | < cutoff | other genes | Total |
|---|---|---|---|
| In Astrocyte | 11 (11.30) | 21 (20.70) | 32 |
| Not IN Astrocyte | 60 (59.70) | 109 (109.30) | 169 |
| Total | 71 | 130 | 201 |

chi2: 0.006, p: 0.9368226476, OR: 0.95159

| | > cutoff | other genes | Total |
|---|---|---|---|
| In Astrocyte | 4 (5.09) | 28 (26.91) | 32 |
| Not IN Astrocyte | 28 (26.91) | 141 (142.09) | 169 |
| Total | 32 | 169 | 201 |

chi2: 0.098, p: 0.7540698172, OR: 0.71939

| | < cutoff | other genes | Total |
|---|---|---|---|
| In Microglia | 11 (19.07) | 43 (34.93) | 54 |
| Not IN Microglia | 60 (51.93) | 87 (95.07) | 147 |
| Total | 71 | 130 | 201 |

chi2: 6.359, p: 0.0116778635, OR: 0.37093

| | > cutoff | other genes | Total |
|---|---|---|---|
| In Microglia | 15 (8.60) | 39 (45.40) | 54 |
| Not IN Microglia | 17 (23.40) | 130 (123.60) | 147 |
| Total | 32 | 169 | 201 |

chi2: 6.591, p: 0.0102467229, OR: 2.94118

| | < cutoff | other genes | Total |
|---|---|---|---|
| In Oligodendrocyte | 14 (7.77) | 8 (14.23) | 22 |
| Not IN Oligodendrocyte | 57 (63.23) | 122 (115.77) | 179 |
| Total | 71 | 130 | 201 |

chi2: 7.332, p: 0.0067722325, OR: 3.74561

| | > cutoff | other genes | Total |
|---|---|---|---|
| In Oligodendrocyte | 2 (3.50) | 20 (18.50) | 22 |
| Not IN Oligodendrocyte | 30 (28.50) | 149 (150.50) | 179 |
| Total | 32 | 169 | 201 |

chi2: 0.383, p: 0.5358917271, OR: 0.49667

| | < cutoff | other genes | Total |
|---|---|---|---|
| In Neuron | 30 (20.13) | 27 (36.87) | 57 |
| Not IN Neuron | 41 (50.87) | 103 (93.13) | 144 |
| Total | 71 | 130 | 201 |

chi2: 9.402, p: 0.0021673165, OR: 2.79133

| | > cutoff | other genes | Total |
|---|---|---|---|
| In Neuron | 4 (9.07) | 53 (47.93) | 57 |
| Not IN Neuron | 28 (22.93) | 116 (121.07) | 144 |
| Total | 32 | 169 | 201 |

chi2: 3.828, p: 0.0503888444, OR: 0.31267

| | < cutoff | other genes | Total |
|---|---|---|---|
| In Endothelia | 5 (12.72) | 31 (23.28) | 36 |
| Not IN Endothelia | 66 (58.28) | 99 (106.72) | 165 |
| Total | 71 | 130 | 201 |

chi2: 7.713, p: 0.0054813754, OR: 0.24194

| | > cutoff | other genes | Total |
|---|---|---|---|
| In Endothelia | 7 (5.73) | 29 (30.27) | 36 |
| Not IN Endothelia | 25 (26.27) | 140 (138.73) | 165 |
| Total | 32 | 169 | 201 |

chi2: 0.149, p: 0.6991491692, OR: 1.35172

### Five Celltypes Distribution of Average dN/dS Scores of Genes Related to membrane_depolarization

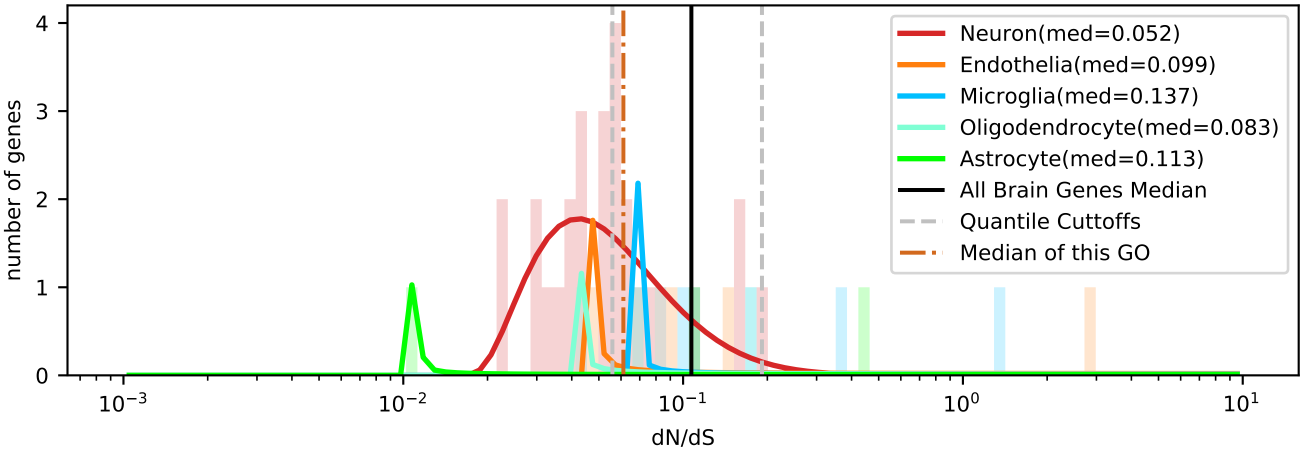


| | < cutoff | other genes | Total |
|---|---|---|---|
| In Astrocyte | 1 (1.26) | 2 (1.74) | 3 |
| Not IN Astrocyte | 17 (16.74) | 23 (23.26) | 40 |
| Total | 18 | 25 | 43 |

chi2: 0.088, p: 0.7670035932, OR: 0.67647

| | > cutoff | other genes | Total |
|---|---|---|---|
| In Astrocyte | 1 (0.35) | 2 (2.65) | 3 |
| Not IN Astrocyte | 4 (4.65) | 36 (35.35) | 40 |
| Total | 5 | 38 | 43 |

chi2: 0.080, p: 0.7777287940, OR: 4.50000

| | < cutoff | other genes | Total |
|---|---|---|---|
| In Microglia | 0 (2.51) | 6 (3.49) | 6 |
| Not IN Microglia | 18 (15.49) | 19 (21.51) | 37 |
| Total | 18 | 25 | 43 |

chi2: 3.221, p: 0.0727175008, OR: 0.00000

| | > cutoff | other genes | Total |
|---|---|---|---|
| In Microglia | 2 (0.70) | 4 (5.30) | 6 |
| Not IN Microglia | 3 (4.30) | 34 (32.70) | 37 |
| Total | 5 | 38 | 43 |

chi2: 1.213, p: 0.2706619670, OR: 5.66667

| | < cutoff | other genes | Total |
|---|---|---|---|
| In Oligodendrocyte | 1 (1.26) | 2 (1.74) | 3 |
| Not IN Oligodendrocyte | 17 (16.74) | 23 (23.26) | 40 |
| Total | 18 | 25 | 43 |

chi2: 0.088, p: 0.7670035932, OR: 0.67647

| | > cutoff | other genes | Total |
|---|---|---|---|
| In Oligodendrocyte | 0 (0.35) | 3 (2.65) | 3 |
| Not IN Oligodendrocyte | 5 (4.65) | 35 (35.35) | 40 |
| Total | 5 | 38 | 43 |

chi2: 0.080, p: 0.7777287940, OR: 0.00000

| | < cutoff | other genes | Total |
|---|---|---|---|
| In Neuron | 15 (10.47) | 10 (14.53) | 25 |
| Not IN Neuron | 3 (7.53) | 15 (10.47) | 18 |
| Total | 18 | 25 | 43 |

chi2: 6.392, p: 0.0114630456, OR: 7.50000

| | > cutoff | other genes | Total |
|---|---|---|---|
| In Neuron | 1 (2.91) | 24 (22.09) | 25 |
| Not IN Neuron | 4 (2.09) | 14 (15.91) | 18 |
| Total | 5 | 38 | 43 |

chi2: 1.841, p: 0.1748539919, OR: 0.14583

| | < cutoff | other genes | Total |
|---|---|---|---|
| In Endothelia | 1 (2.51) | 5 (3.49) | 6 |
| Not IN Endothelia | 17 (15.49) | 20 (21.51) | 37 |
| Total | 18 | 25 | 43 |

chi2: 0.814, p: 0.3667984604, OR: 0.23529

| | > cutoff | other genes | Total |
|---|---|---|---|
| In Endothelia | 1 (0.70) | 5 (5.30) | 6 |
| Not IN Endothelia | 4 (4.30) | 33 (32.70) | 37 |
| Total | 5 | 38 | 43 |

chi2: 0.074, p: 0.7860881970, OR: 1.65000

### Five Celltypes Distribution of Average dN/dS Scores of Genes Related to membrane_hyperpolarization

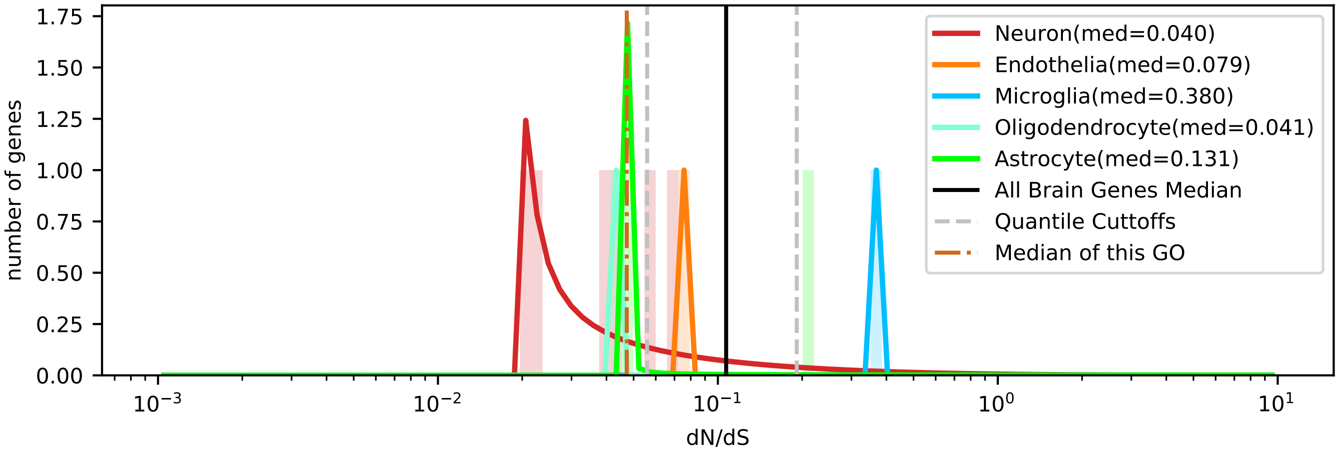


| | < cutoff | other genes | Total |
|---|---|---|---|
| In Astrocyte | 1 (1.09) | 1 (0.91) | 2 |
| Not IN Astrocyte | 5 (4.91) | 4 (4.09) | 9 |
| Total | 6 | 5 | 11 |

chi2: 0.413, p: 0.5207033238, OR: 0.80000

| | > cutoff | other genes | Total |
|---|---|---|---|
| In Astrocyte | 1 (0.36) | 1 (1.64) | 2 |
| Not IN Astrocyte | 1 (1.64) | 8 (7.36) | 9 |
| Total | 2 | 9 | 11 |

chi2: 0.076, p: 0.7822520699, OR: 8.00000

| | < cutoff | other genes | Total |
|---|---|---|---|
| In Microglia | 0 (0.55) | 1 (0.45) | 1 |
| Not IN Microglia | 6 (5.45) | 4 (4.55) | 10 |
| Total | 6 | 5 | 11 |

chi2: 0.009, p: 0.9237249184, OR: 0.00000

| | > cutoff | other genes | Total |
|---|---|---|---|
| In Microglia | 1 (0.18) | 0 (0.82) | 1 |
| Not IN Microglia | 1 (1.82) | 9 (8.18) | 10 |
| Total | 2 | 9 | 11 |

chi2: 0.749, p: 0.3869163178, OR: inf

| | < cutoff | other genes | Total |
|---|---|---|---|
| In Oligodendrocyte | 1 (0.55) | 0 (0.45) | 1 |
| Not IN Oligodendrocyte | 5 (5.45) | 5 (4.55) | 10 |
| Total | 6 | 5 | 11 |

chi2: 0.009, p: 0.9237249184, OR: inf

| | > cutoff | other genes | Total |
|---|---|---|---|
| In Oligodendrocyte | 0 (0.18) | 1 (0.82) | 1 |
| Not IN Oligodendrocyte | 2 (1.82) | 8 (8.18) | 10 |
| Total | 2 | 9 | 11 |

chi2: 0.749, p: 0.3869163178, OR: 0.00000

| | < cutoff | other genes | Total |
|---|---|---|---|
| In Neuron | 4 (3.27) | 2 (2.73) | 6 |
| Not IN Neuron | 2 (2.73) | 3 (2.27) | 5 |
| Total | 6 | 5 | 11 |

chi2: 0.076, p: 0.7822520699, OR: 3.00000

| | > cutoff | other genes | Total |
|---|---|---|---|
| In Neuron | 0 (1.09) | 6 (4.91) | 6 |
| Not IN Neuron | 2 (0.91) | 3 (4.09) | 5 |
| Total | 2 | 9 | 11 |

chi2: 0.861, p: 0.3535573755, OR: 0.00000

| | < cutoff | other genes | Total |
|---|---|---|---|
| In Endothelia | 0 (0.55) | 1 (0.45) | 1 |
| Not IN Endothelia | 6 (5.45) | 4 (4.55) | 10 |
| Total | 6 | 5 | 11 |

chi2: 0.009, p: 0.9237249184, OR: 0.00000

| | > cutoff | other genes | Total |
|---|---|---|---|
| In Endothelia | 0 (0.18) | 1 (0.82) | 1 |
| Not IN Endothelia | 2 (1.82) | 8 (8.18) | 10 |
| Total | 2 | 9 | 11 |

chi2: 0.749, p: 0.3869163178, OR: 0.00000

Five Celltypes Distribution of Average dN/dS Scores of Genes Related to membrane_repolarization

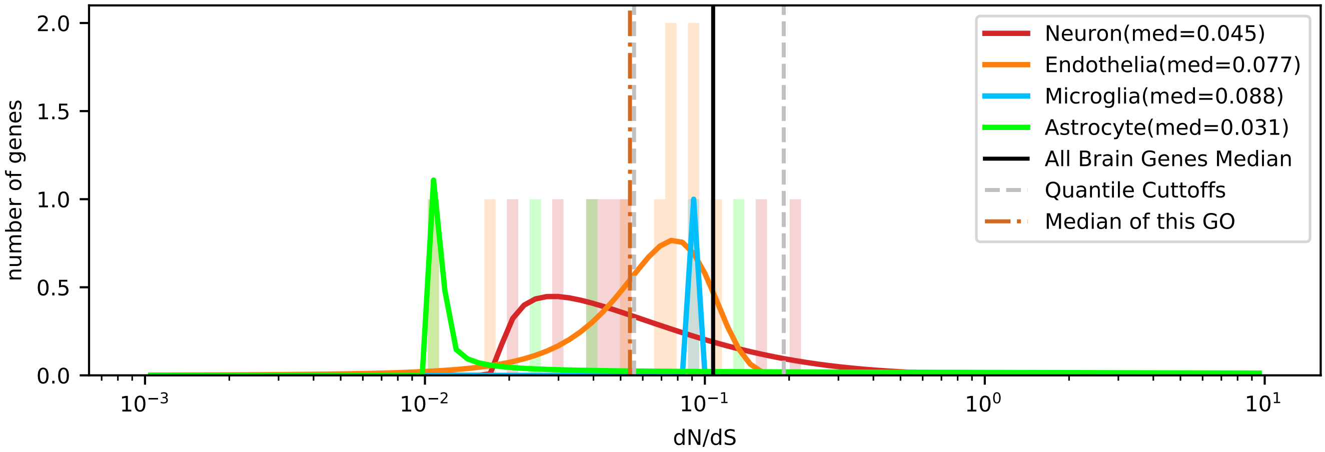

### Five Celltypes Distribution of Average dN/dS Scores of Genes Related to mitochondrion organization

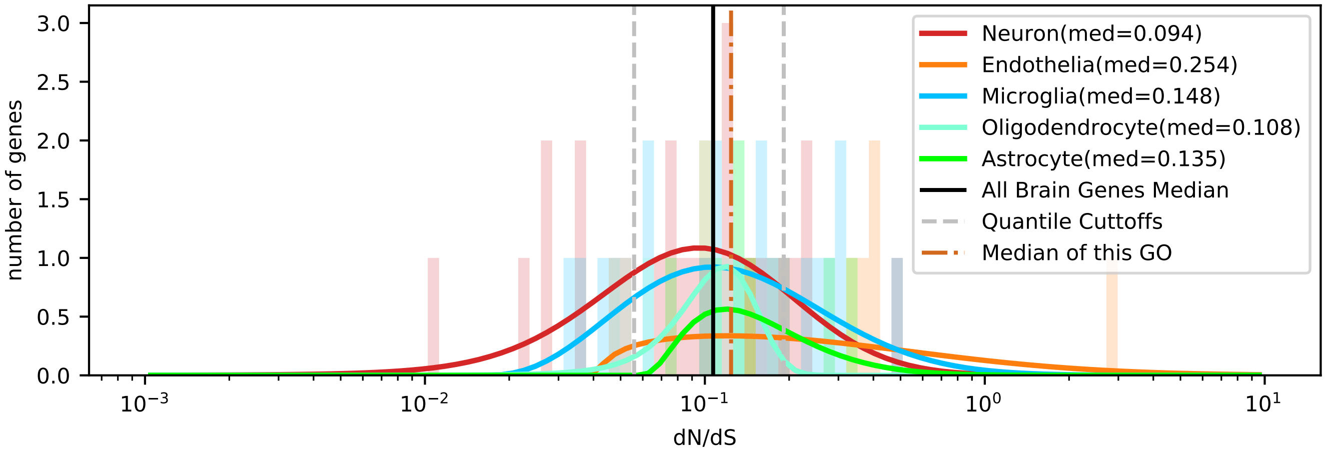


| | < cutoff | other genes | Total |
|---|---|---|---|
| In Astrocyte | 0 (1.32) | 7 (5.68) | 7 |
| Not IN Astrocyte | 13 (11.68) | 49 (50.32) | 62 |
| Total | 13 | 56 | 69 |

chi2: 0.697, p: 0.4037437287, OR: 0.00000

| | > cutoff | other genes | Total |
|---|---|---|---|
| In Astrocyte | 2 (1.83) | 5 (5.17) | 7 |
| Not IN Astrocyte | 16 (16.17) | 46 (45.83) | 62 |
| Total | 18 | 51 | 69 |

chi2: 0.088, p: 0.7671524380, OR: 1.15000

| | < cutoff | other genes | Total |
|---|---|---|---|
| In Microglia | 4 (3.77) | 16 (16.23) | 20 |
| Not IN Microglia | 9 (9.23) | 40 (39.77) | 49 |
| Total | 13 | 56 | 69 |

chi2: 0.033, p: 0.8556333123, OR: 1.11111

| | > cutoff | other genes | Total |
|---|---|---|---|
| In Microglia | 6 (5.22) | 14 (14.78) | 20 |
| Not IN Microglia | 12 (12.78) | 37 (36.22) | 49 |
| Total | 18 | 51 | 69 |

chi2: 0.029, p: 0.8644005376, OR: 1.32143

| | < cutoff | other genes | Total |
|---|---|---|---|
| In Oligodendrocyte | 1 (1.51) | 7 (6.49) | 8 |
| Not IN Oligodendrocyte | 12 (11.49) | 49 (49.51) | 61 |
| Total | 13 | 56 | 69 |

chi2: 0.000, p: 0.9944402470, OR: 0.58333

| | > cutoff | other genes | Total |
|---|---|---|---|
| In Oligodendrocyte | 0 (2.09) | 8 (5.91) | 8 |
| Not IN Oligodendrocyte | 18 (15.91) | 43 (45.09) | 61 |
| Total | 18 | 51 | 69 |

chi2: 1.847, p: 0.1741585313, OR: 0.00000

| | < cutoff | other genes | Total |
|---|---|---|---|
| In Neuron | 7 (4.52) | 17 (19.48) | 24 |
| Not IN Neuron | 6 (8.48) | 39 (36.52) | 45 |
| Total | 13 | 56 | 69 |

chi2: 1.635, p: 0.2009913475, OR: 2.67647

| | > cutoff | other genes | Total |
|---|---|---|---|
| In Neuron | 5 (6.26) | 19 (17.74) | 24 |
| Not IN Neuron | 13 (11.74) | 32 (33.26) | 45 |
| Total | 18 | 51 | 69 |

chi2: 0.192, p: 0.6614034491, OR: 0.64777

| | < cutoff | other genes | Total |
|---|---|---|---|
| In Endothelia | 1 (1.88) | 9 (8.12) | 10 |
| Not IN Endothelia | 12 (11.12) | 47 (47.88) | 59 |
| Total | 13 | 56 | 69 |

chi2: 0.113, p: 0.7369645210, OR: 0.43519

| | > cutoff | other genes | Total |
|---|---|---|---|
| In Endothelia | 5 (2.61) | 5 (7.39) | 10 |
| Not IN Endothelia | 13 (15.39) | 46 (43.61) | 59 |
| Total | 18 | 51 | 69 |

chi2: 2.170, p: 0.1407647874, OR: 3.53846

### Five Celltypes Distribution of Average dN/dS Scores of Genes Related to mitotic cell cycle

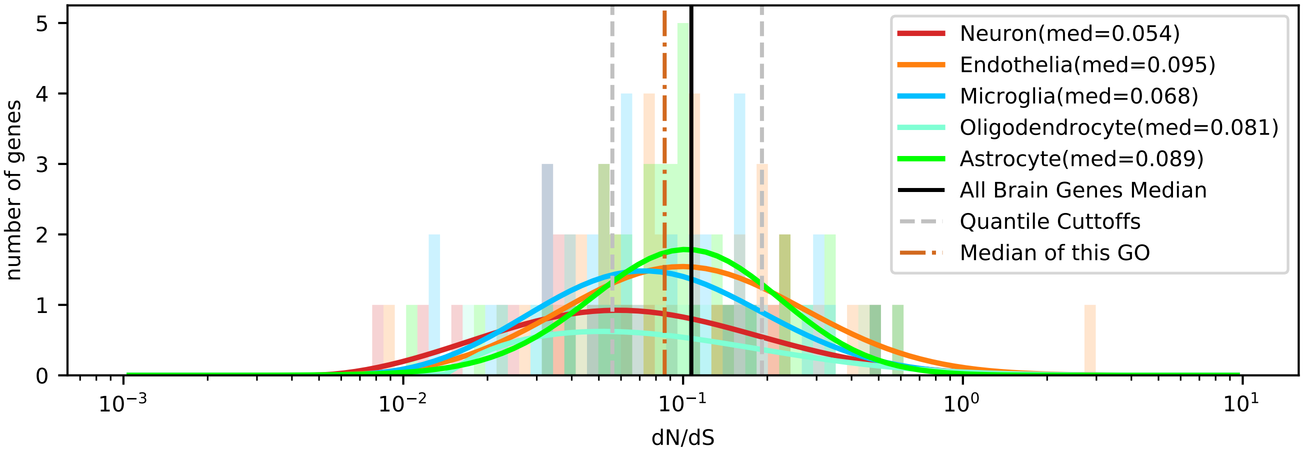


| | < cutoff | other genes | Total |
|---|---|---|---|
| In Astrocyte | 7 (12.35) | 31 (25.65) | 38 |
| Not IN Astrocyte | 45 (39.65) | 77 (82.35) | 122 |
| Total | 52 | 108 | 160 |

chi2: 3.701, p: 0.0543926956, OR: 0.38638

| | > cutoff | other genes | Total |
|---|---|---|---|
| In Astrocyte | 8 (7.36) | 30 (30.64) | 38 |
| Not IN Astrocyte | 23 (23.64) | 99 (98.36) | 122 |
| Total | 31 | 129 | 160 |

chi2: 0.004, p: 0.9484684677, OR: 1.14783

| | < cutoff | other genes | Total |
|---|---|---|---|
| In Microglia | 13 (12.03) | 24 (24.98) | 37 |
| Not IN Microglia | 39 (39.98) | 84 (83.03) | 123 |
| Total | 52 | 108 | 160 |

chi2: 0.036, p: 0.8491880612, OR: 1.16667

| | > cutoff | other genes | Total |
|---|---|---|---|
| In Microglia | 5 (7.17) | 32 (29.83) | 37 |
| Not IN Microglia | 26 (23.83) | 97 (99.17) | 123 |
| Total | 31 | 129 | 160 |

chi2: 0.627, p: 0.4285553040, OR: 0.58293

| | < cutoff | other genes | Total |
|---|---|---|---|
| In Oligodendrocyte | 7 (5.53) | 10 (11.47) | 17 |
| Not IN Oligodendrocyte | 45 (46.48) | 98 (96.53) | 143 |
| Total | 52 | 108 | 160 |

chi2: 0.285, p: 0.5933097989, OR: 1.52444

| | > cutoff | other genes | Total |
|---|---|---|---|
| In Oligodendrocyte | 3 (3.29) | 14 (13.71) | 17 |
| Not IN Oligodendrocyte | 28 (27.71) | 115 (115.29) | 143 |
| Total | 31 | 129 | 160 |

chi2: 0.018, p: 0.8934998920, OR: 0.88010

| | < cutoff | other genes | Total |
|---|---|---|---|
| In Neuron | 15 (8.78) | 12 (18.23) | 27 |
| Not IN Neuron | 37 (43.23) | 96 (89.78) | 133 |
| Total | 52 | 108 | 160 |

chi2: 6.657, p: 0.0098775661, OR: 3.24324

| | > cutoff | other genes | Total |
|---|---|---|---|
| In Neuron | 5 (5.23) | 22 (21.77) | 27 |
| Not IN Neuron | 26 (25.77) | 107 (107.23) | 133 |
| Total | 31 | 129 | 160 |

chi2: 0.021, p: 0.8858710047, OR: 0.93531

| | < cutoff | other genes | Total |
|---|---|---|---|
| In Endothelia | 10 (13.32) | 31 (27.68) | 41 |
| Not IN Endothelia | 42 (38.67) | 77 (80.33) | 119 |
| Total | 52 | 108 | 160 |

chi2: 1.193, p: 0.2747260886, OR: 0.59140

| | > cutoff | other genes | Total |
|---|---|---|---|
| In Endothelia | 10 (7.94) | 31 (33.06) | 41 |
| Not IN Endothelia | 21 (23.06) | 98 (95.94) | 119 |
| Total | 31 | 129 | 160 |

chi2: 0.508, p: 0.4758166027, OR: 1.50538

### Five Celltypes Distribution of Average dN/dS Scores of Genes Related to mitotic nuclear division

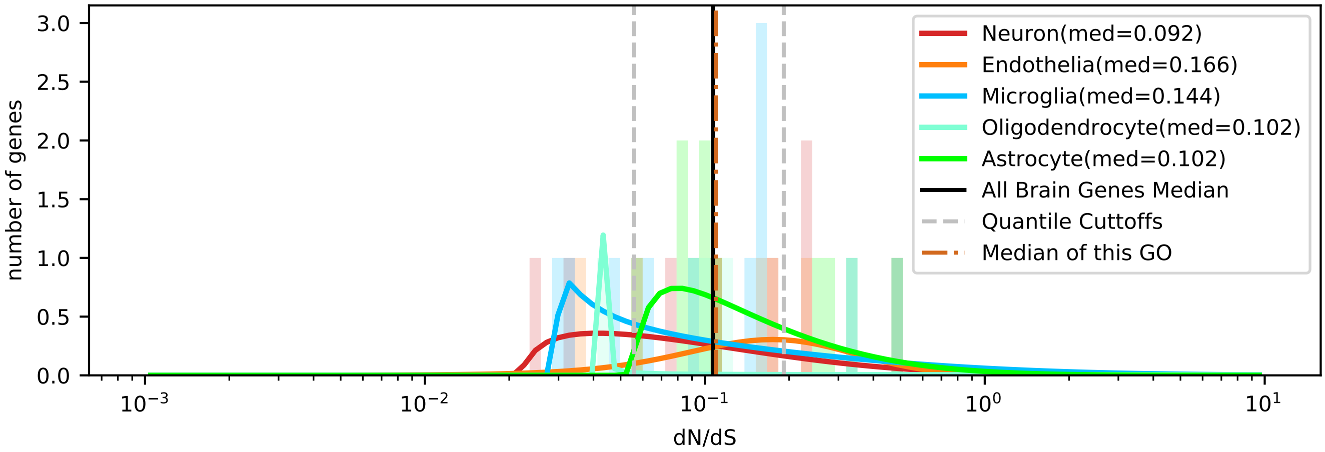


| | < cutoff | other genes | Total |
|---|---|---|---|
| In Astrocyte | 0 (2.26) | 11 (8.74) | 11 |
| Not IN Astrocyte | 8 (5.74) | 20 (22.26) | 28 |
| Total | 8 | 31 | 39 |

chi2: 2.396, p: 0.1216645907, OR: 0.00000

| | > cutoff | other genes | Total |
|---|---|---|---|
| In Astrocyte | 4 (2.82) | 7 (8.18) | 11 |
| Not IN Astrocyte | 6 (7.18) | 22 (20.82) | 28 |
| Total | 10 | 29 | 39 |

chi2: 0.307, p: 0.5797586079, OR: 2.09524

| | < cutoff | other genes | Total |
|---|---|---|---|
| In Microglia | 3 (2.26) | 8 (8.74) | 11 |
| Not IN Microglia | 5 (5.74) | 23 (22.26) | 28 |
| Total | 8 | 31 | 39 |

chi2: 0.046, p: 0.8300310292, OR: 1.72500

| | > cutoff | other genes | Total |
|---|---|---|---|
| In Microglia | 2 (2.82) | 9 (8.18) | 11 |
| Not IN Microglia | 8 (7.18) | 20 (20.82) | 28 |
| Total | 10 | 29 | 39 |

chi2: 0.068, p: 0.7939406959, OR: 0.55556

| | < cutoff | other genes | Total |
|---|---|---|---|
| In Oligodendrocyte | 1 (0.62) | 2 (2.38) | 3 |
| Not IN Oligodendrocyte | 7 (7.38) | 29 (28.62) | 36 |
| Total | 8 | 31 | 39 |

chi2: 0.029, p: 0.8636619455, OR: 2.07143

| | > cutoff | other genes | Total |
|---|---|---|---|
| In Oligodendrocyte | 0 (0.77) | 3 (2.23) | 3 |
| Not IN Oligodendrocyte | 10 (9.23) | 26 (26.77) | 36 |
| Total | 10 | 29 | 39 |

chi2: 0.137, p: 0.7109956682, OR: 0.00000

| | < cutoff | other genes | Total |
|---|---|---|---|
| In Neuron | 3 (1.64) | 5 (6.36) | 8 |
| Not IN Neuron | 5 (6.36) | 26 (24.64) | 31 |
| Total | 8 | 31 | 39 |

chi2: 0.712, p: 0.3989050617, OR: 3.12000

| | > cutoff | other genes | Total |
|---|---|---|---|
| In Neuron | 2 (2.05) | 6 (5.95) | 8 |
| Not IN Neuron | 8 (7.95) | 23 (23.05) | 31 |
| Total | 10 | 29 | 39 |

chi2: 0.166, p: 0.6836284608, OR: 0.95833

| | < cutoff | other genes | Total |
|---|---|---|---|
| In Endothelia | 1 (1.23) | 5 (4.77) | 6 |
| Not IN Endothelia | 7 (6.77) | 26 (26.23) | 33 |
| Total | 8 | 31 | 39 |

chi2: 0.088, p: 0.7672969797, OR: 0.74286

| | > cutoff | other genes | Total |
|---|---|---|---|
| In Endothelia | 2 (1.54) | 4 (4.46) | 6 |
| Not IN Endothelia | 8 (8.46) | 25 (24.54) | 33 |
| Total | 10 | 29 | 39 |

chi2: 0.002, p: 0.9688167465, OR: 1.56250

### Five Celltypes Distribution of Average dN/dS Scores of Genes Related to mRNA processing

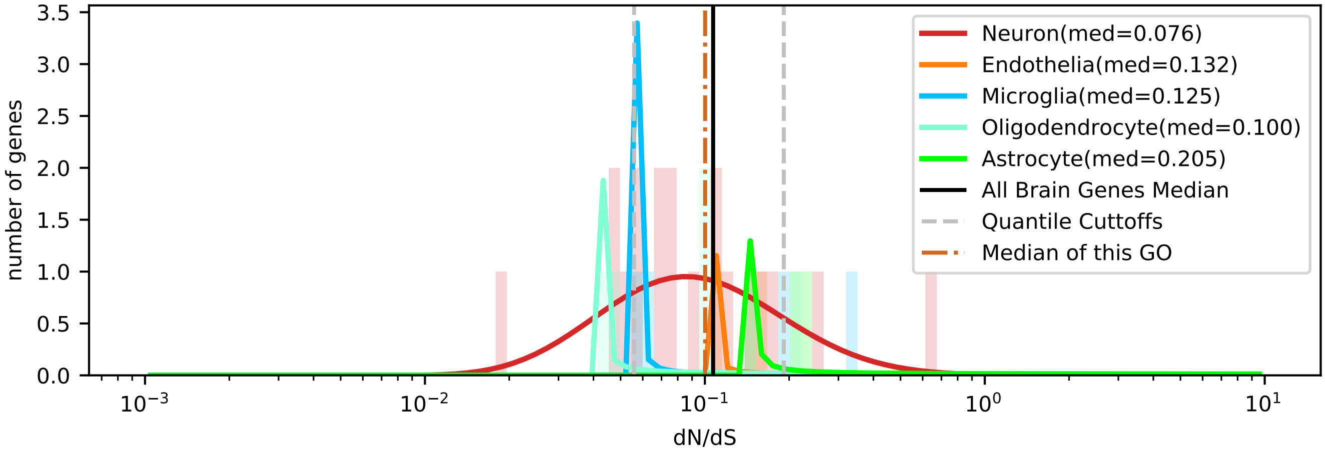


| | < cutoff | other genes | Total |
|---|---|---|---|
| In Astrocyte | 0 (0.55) | 3 (2.45) | 3 |
| Not IN Astrocyte | 6 (5.45) | 24 (24.55) | 30 |
| Total | 6 | 27 | 33 |

chi2: 0.005, p: 0.9431093312, OR: 0.00000

| | > cutoff | other genes | Total |
|---|---|---|---|
| In Astrocyte | 2 (0.55) | 1 (2.45) | 3 |
| Not IN Astrocyte | 4 (5.45) | 26 (24.55) | 30 |
| Total | 6 | 27 | 33 |

chi2: 2.246, p: 0.1339747156, OR: 13.00000

| | < cutoff | other genes | Total |
|---|---|---|---|
| In Microglia | 0 (0.73) | 4 (3.27) | 4 |
| Not IN Microglia | 6 (5.27) | 23 (23.73) | 29 |
| Total | 6 | 27 | 33 |

chi2: 0.099, p: 0.7533001504, OR: 0.00000

| | > cutoff | other genes | Total |
|---|---|---|---|
| In Microglia | 1 (0.73) | 3 (3.27) | 4 |
| Not IN Microglia | 5 (5.27) | 24 (23.73) | 29 |
| Total | 6 | 27 | 33 |

chi2: 0.099, p: 0.7533001504, OR: 1.60000

| | < cutoff | other genes | Total |
|---|---|---|---|
| In Oligodendrocyte | 2 (0.91) | 3 (4.09) | 5 |
| Not IN Oligodendrocyte | 4 (5.09) | 24 (22.91) | 28 |
| Total | 6 | 27 | 33 |

chi2: 0.553, p: 0.4569830942, OR: 4.00000

| | > cutoff | other genes | Total |
|---|---|---|---|
| In Oligodendrocyte | 1 (0.91) | 4 (4.09) | 5 |
| Not IN Oligodendrocyte | 5 (5.09) | 23 (22.91) | 28 |
| Total | 6 | 27 | 33 |

chi2: 0.265, p: 0.6065845206, OR: 1.15000

| | < cutoff | other genes | Total |
|---|---|---|---|
| In Neuron | 4 (3.45) | 15 (15.55) | 19 |
| Not IN Neuron | 2 (2.55) | 12 (11.45) | 14 |
| Total | 6 | 27 | 33 |

chi2: 0.002, p: 0.9668895365, OR: 1.60000

| | > cutoff | other genes | Total |
|---|---|---|---|
| In Neuron | 2 (3.45) | 17 (15.55) | 19 |
| Not IN Neuron | 4 (2.55) | 10 (11.45) | 14 |
| Total | 6 | 27 | 33 |

chi2: 0.760, p: 0.3833697031, OR: 0.29412

| | < cutoff | other genes | Total |
|---|---|---|---|
| In Endothelia | 0 (0.36) | 2 (1.64) | 2 |
| Not IN Endothelia | 6 (5.64) | 25 (25.36) | 31 |
| Total | 6 | 27 | 33 |

chi2: 0.067, p: 0.7964543888, OR: 0.00000

| | > cutoff | other genes | Total |
|---|---|---|---|
| In Endothelia | 0 (0.36) | 2 (1.64) | 2 |
| Not IN Endothelia | 6 (5.64) | 25 (25.36) | 31 |
| Total | 6 | 27 | 33 |

chi2: 0.067, p: 0.7964543888, OR: 0.00000

### Five Celltypes Distribution of Average dN/dS Scores of Genes Related to nervous system process

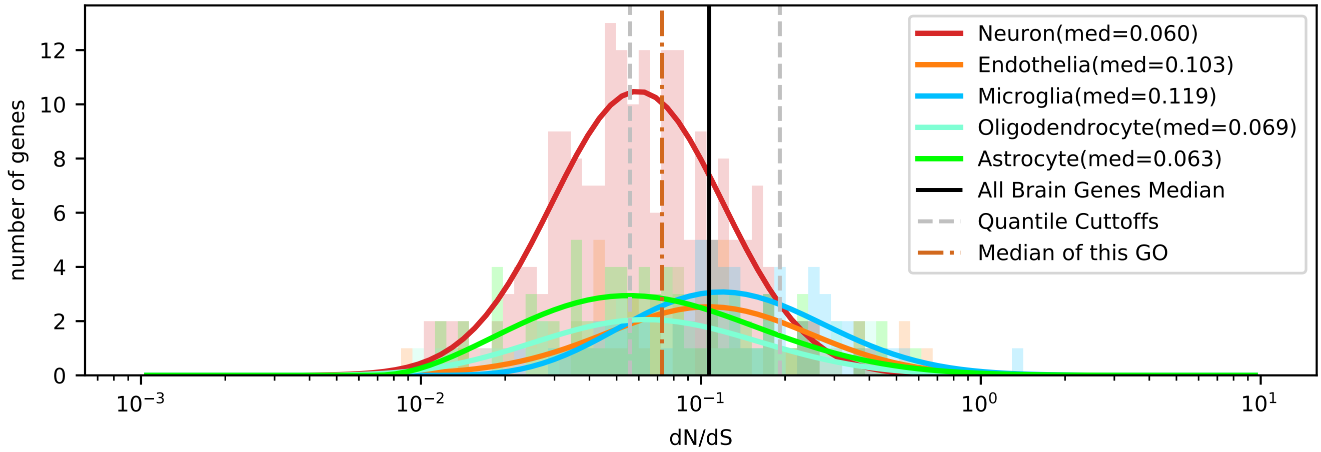


| | < cutoff | other genes | Total |
|---|---|---|---|
| In Astrocyte | 34 (29.05) | 42 (46.95) | 76 |
| Not IN Astrocyte | 138 (142.95) | 236 (231.05) | 374 |
| Total | 172 | 278 | 450 |

chi2: 1.328, p: 0.2490970840, OR: 1.38440

| | > cutoff | other genes | Total |
|---|---|---|---|
| In Astrocyte | 10 (9.80) | 66 (66.20) | 76 |
| Not IN Astrocyte | 48 (48.20) | 326 (325.80) | 374 |
| Total | 58 | 392 | 450 |

chi2: 0.012, p: 0.9116294606, OR: 1.02904

| | < cutoff | other genes | Total |
|---|---|---|---|
| In Microglia | 12 (25.61) | 55 (41.39) | 67 |
| Not IN Microglia | 160 (146.39) | 223 (236.61) | 383 |
| Total | 172 | 278 | 450 |

chi2: 12.762, p: 0.0003537117, OR: 0.30409

| | > cutoff | other genes | Total |
|---|---|---|---|
| In Microglia | 20 (8.64) | 47 (58.36) | 67 |
| Not IN Microglia | 38 (49.36) | 345 (333.64) | 383 |
| Total | 58 | 392 | 450 |

chi2: 18.436, p: 0.0000175714, OR: 3.86338

| | < cutoff | other genes | Total |
|---|---|---|---|
| In Oligodendrocyte | 20 (19.11) | 30 (30.89) | 50 |
| Not IN Oligodendrocyte | 152 (152.89) | 248 (247.11) | 400 |
| Total | 172 | 278 | 450 |

chi2: 0.014, p: 0.9044478051, OR: 1.08772

| | > cutoff | other genes | Total |
|---|---|---|---|
| In Oligodendrocyte | 6 (6.44) | 44 (43.56) | 50 |
| Not IN Oligodendrocyte | 52 (51.56) | 348 (348.44) | 400 |
| Total | 58 | 392 | 450 |

chi2: 0.001, p: 0.9801587271, OR: 0.91259

| | < cutoff | other genes | Total |
|---|---|---|---|
| In Neuron | 93 (76.06) | 106 (122.94) | 199 |
| Not IN Neuron | 79 (95.94) | 172 (155.06) | 251 |
| Total | 172 | 278 | 450 |

chi2: 10.309, p: 0.0013237079, OR: 1.91020

| | > cutoff | other genes | Total |
|---|---|---|---|
| In Neuron | 8 (25.65) | 191 (173.35) | 199 |
| Not IN Neuron | 50 (32.35) | 201 (218.65) | 251 |
| Total | 58 | 392 | 450 |

chi2: 23.598, p: 0.0000011873, OR: 0.16838

| | < cutoff | other genes | Total |
|---|---|---|---|
| In Endothelia | 13 (22.17) | 45 (35.83) | 58 |
| Not IN Endothelia | 159 (149.83) | 233 (242.17) | 392 |
| Total | 172 | 278 | 450 |

chi2: 6.299, p: 0.0120800663, OR: 0.42334

| | > cutoff | other genes | Total |
|---|---|---|---|
| In Endothelia | 14 (7.48) | 44 (50.52) | 58 |
| Not IN Endothelia | 44 (50.52) | 348 (341.48) | 392 |
| Total | 58 | 392 | 450 |

chi2: 6.398, p: 0.0114249765, OR: 2.51653

Five Celltypes Distribution of Average dN/dS Scores of Genes Related to nitrogen cycle metabolic process

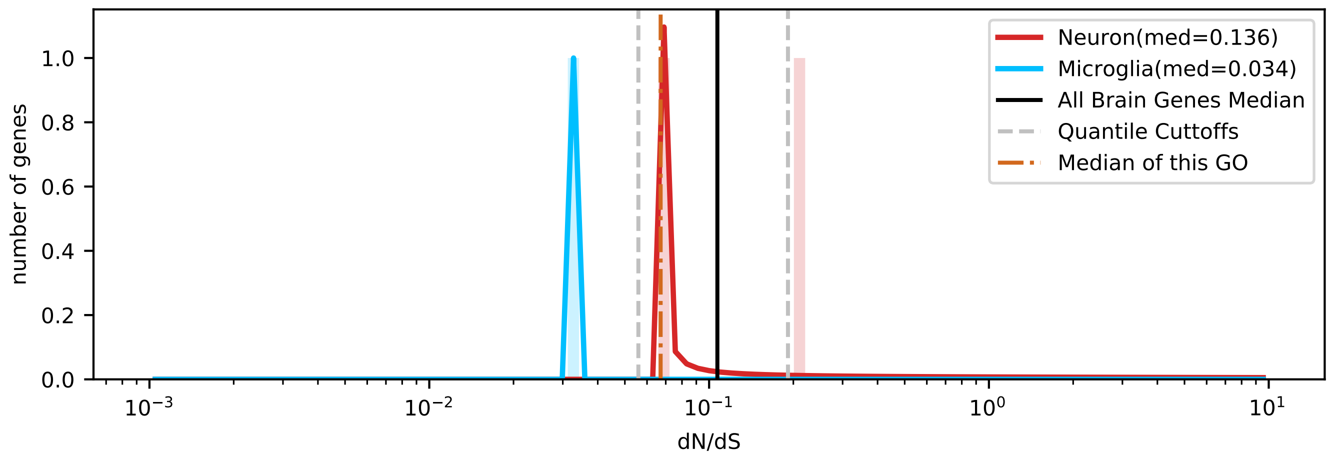

### Five Celltypes Distribution of Average dN/dS Scores of Genes Related to nucleobase-containing compound catabolic process

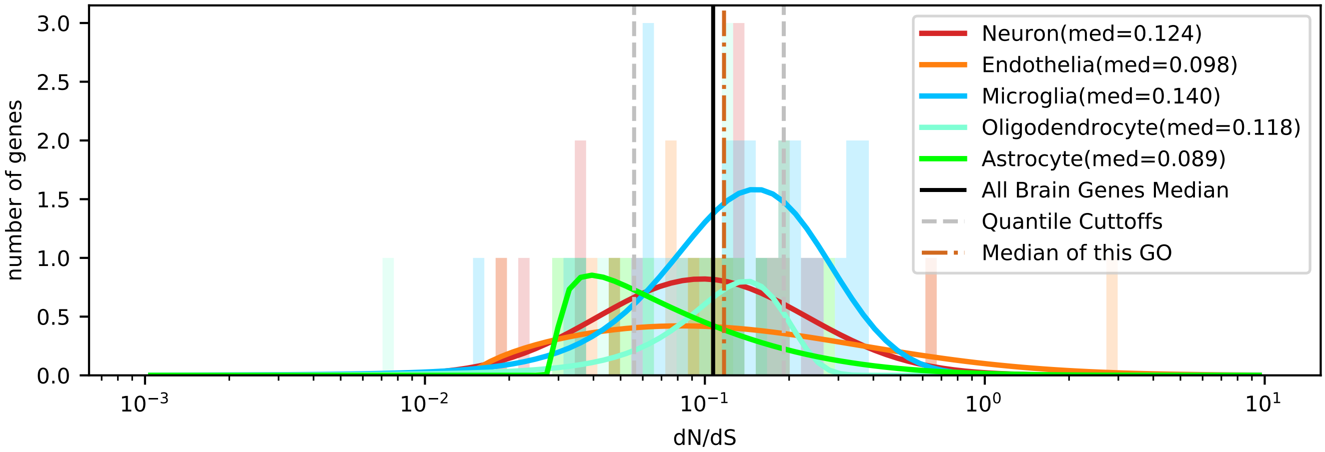


| | < cutoff | other genes | Total |
|---|---|---|---|
| In Astrocyte | 5 (2.75) | 8 (10.25) | 13 |
| Not IN Astrocyte | 13 (15.25) | 59 (56.75) | 72 |
| Total | 18 | 67 | 85 |

chi2: 1.661, p: 0.1975307731, OR: 2.83654

| | > cutoff | other genes | Total |
|---|---|---|---|
| In Astrocyte | 1 (3.06) | 12 (9.94) | 13 |
| Not IN Astrocyte | 19 (16.94) | 53 (55.06) | 72 |
| Total | 20 | 65 | 85 |

chi2: 1.226, p: 0.2681086604, OR: 0.23246

| | < cutoff | other genes | Total |
|---|---|---|---|
| In Microglia | 3 (6.14) | 26 (22.86) | 29 |
| Not IN Microglia | 15 (11.86) | 41 (44.14) | 56 |
| Total | 18 | 67 | 85 |

chi2: 2.187, p: 0.1391485358, OR: 0.31538

| | > cutoff | other genes | Total |
|---|---|---|---|
| In Microglia | 10 (6.82) | 19 (22.18) | 29 |
| Not IN Microglia | 10 (13.18) | 46 (42.82) | 56 |
| Total | 20 | 65 | 85 |

chi2: 2.084, p: 0.1488708898, OR: 2.42105

| | < cutoff | other genes | Total |
|---|---|---|---|
| In Oligodendrocyte | 2 (2.12) | 8 (7.88) | 10 |
| Not IN Oligodendrocyte | 16 (15.88) | 59 (59.12) | 75 |
| Total | 18 | 67 | 85 |

chi2: 0.099, p: 0.7527187039, OR: 0.92188

| | > cutoff | other genes | Total |
|---|---|---|---|
| In Oligodendrocyte | 2 (2.35) | 8 (7.65) | 10 |
| Not IN Oligodendrocyte | 18 (17.65) | 57 (57.35) | 75 |
| Total | 20 | 65 | 85 |

chi2: 0.014, p: 0.9070879278, OR: 0.79167

| | < cutoff | other genes | Total |
|---|---|---|---|
| In Neuron | 5 (4.02) | 14 (14.98) | 19 |
| Not IN Neuron | 13 (13.98) | 53 (52.02) | 66 |
| Total | 18 | 67 | 85 |

chi2: 0.092, p: 0.7614111866, OR: 1.45604

| | > cutoff | other genes | Total |
|---|---|---|---|
| In Neuron | 3 (4.47) | 16 (14.53) | 19 |
| Not IN Neuron | 17 (15.53) | 49 (50.47) | 66 |
| Total | 20 | 65 | 85 |

chi2: 0.355, p: 0.5513615745, OR: 0.54044

| | < cutoff | other genes | Total |
|---|---|---|---|
| In Endothelia | 3 (2.96) | 11 (11.04) | 14 |
| Not IN Endothelia | 15 (15.04) | 56 (55.96) | 71 |
| Total | 18 | 67 | 85 |

chi2: 0.111, p: 0.7394261440, OR: 1.01818

| | > cutoff | other genes | Total |
|---|---|---|---|
| In Endothelia | 4 (3.29) | 10 (10.71) | 14 |
| Not IN Endothelia | 16 (16.71) | 55 (54.29) | 71 |
| Total | 20 | 65 | 85 |

chi2: 0.020, p: 0.8871331188, OR: 1.37500

### Five Celltypes Distribution of Average dN/dS Scores of Genes Related to nucleocytoplasmic transport

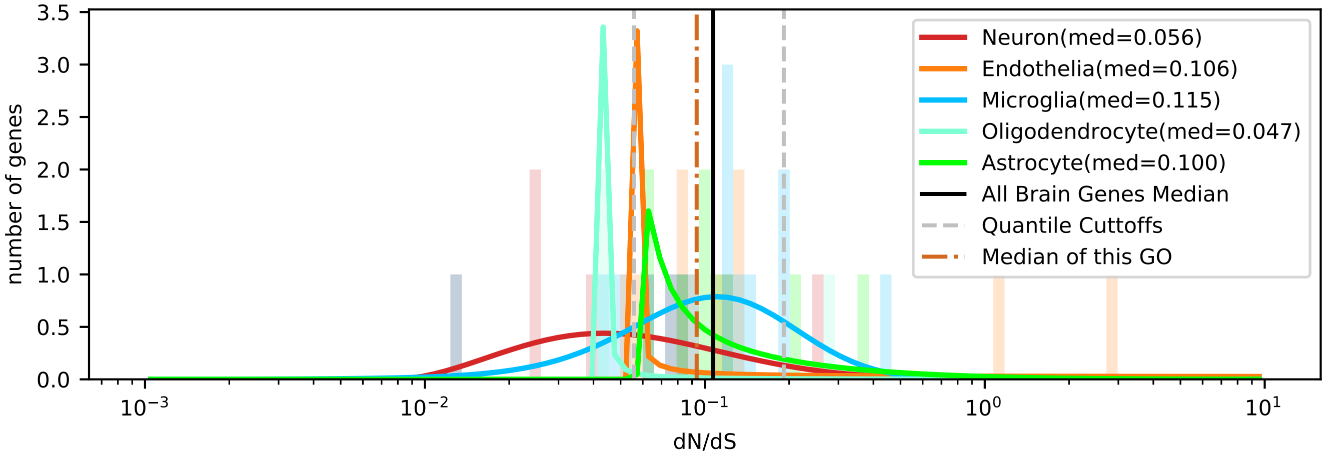


| | < cutoff | other genes | Total |
|---|---|---|---|
| In Astrocyte | 0 (2.30) | 9 (6.70) | 9 |
| Not IN Astrocyte | 12 (9.70) | 26 (28.30) | 38 |
| Total | 12 | 35 | 47 |

chi2: 2.336, p: 0.1263862636, OR: 0.00000

| | > cutoff | other genes | Total |
|---|---|---|---|
| In Astrocyte | 2 (1.34) | 7 (7.66) | 9 |
| Not IN Astrocyte | 5 (5.66) | 33 (32.34) | 38 |
| Total | 7 | 40 | 47 |

chi2: 0.028, p: 0.8680335542, OR: 1.88571

| | < cutoff | other genes | Total |
|---|---|---|---|
| In Microglia | 3 (3.83) | 12 (11.17) | 15 |
| Not IN Microglia | 9 (8.17) | 23 (23.83) | 32 |
| Total | 12 | 35 | 47 |

chi2: 0.056, p: 0.8129161283, OR: 0.63889

| | > cutoff | other genes | Total |
|---|---|---|---|
| In Microglia | 1 (2.23) | 14 (12.77) | 15 |
| Not IN Microglia | 6 (4.77) | 26 (27.23) | 32 |
| Total | 7 | 40 | 47 |

chi2: 0.416, p: 0.5188225841, OR: 0.30952

| | < cutoff | other genes | Total |
|---|---|---|---|
| In Oligodendrocyte | 3 (1.02) | 1 (2.98) | 4 |
| Not IN Oligodendrocyte | 9 (10.98) | 34 (32.02) | 43 |
| Total | 12 | 35 | 47 |

chi2: 3.143, p: 0.0762719210, OR: 11.33333

| | > cutoff | other genes | Total |
|---|---|---|---|
| In Oligodendrocyte | 1 (0.60) | 3 (3.40) | 4 |
| Not IN Oligodendrocyte | 6 (6.40) | 37 (36.60) | 43 |
| Total | 7 | 40 | 47 |

chi2: 0.020, p: 0.8882031134, OR: 2.05556

| | < cutoff | other genes | Total |
|---|---|---|---|
| In Neuron | 5 (2.55) | 5 (7.45) | 10 |
| Not IN Neuron | 7 (9.45) | 30 (27.55) | 37 |
| Total | 12 | 35 | 47 |

chi2: 2.532, p: 0.1115482972, OR: 4.28571

| | > cutoff | other genes | Total |
|---|---|---|---|
| In Neuron | 1 (1.49) | 9 (8.51) | 10 |
| Not IN Neuron | 6 (5.51) | 31 (31.49) | 37 |
| Total | 7 | 40 | 47 |

chi2: 0.000, p: 0.9915028965, OR: 0.57407

| | < cutoff | other genes | Total |
|---|---|---|---|
| In Endothelia | 1 (2.30) | 8 (6.70) | 9 |
| Not IN Endothelia | 11 (9.70) | 27 (28.30) | 38 |
| Total | 12 | 35 | 47 |

chi2: 0.460, p: 0.4975610222, OR: 0.30682

| | > cutoff | other genes | Total |
|---|---|---|---|
| In Endothelia | 2 (1.34) | 7 (7.66) | 9 |
| Not IN Endothelia | 5 (5.66) | 33 (32.34) | 38 |
| Total | 7 | 40 | 47 |

chi2: 0.028, p: 0.8680335542, OR: 1.88571

Five Celltypes Distribution of Average dN/dS Scores of Genes Related to photosynthesis

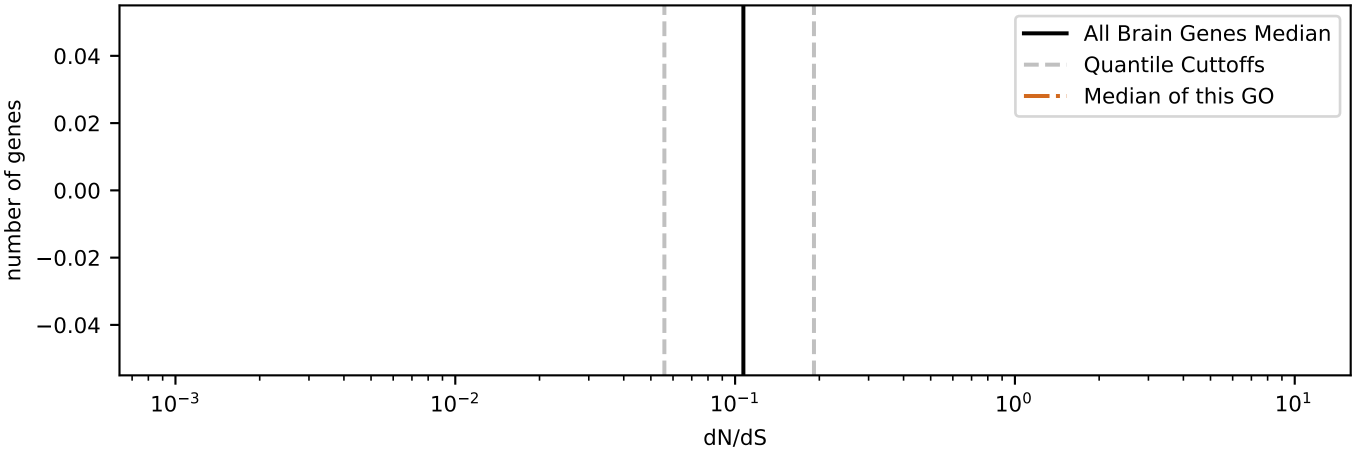

### Five Celltypes Distribution of Average dN/dS Scores of Genes Related to pigmentation

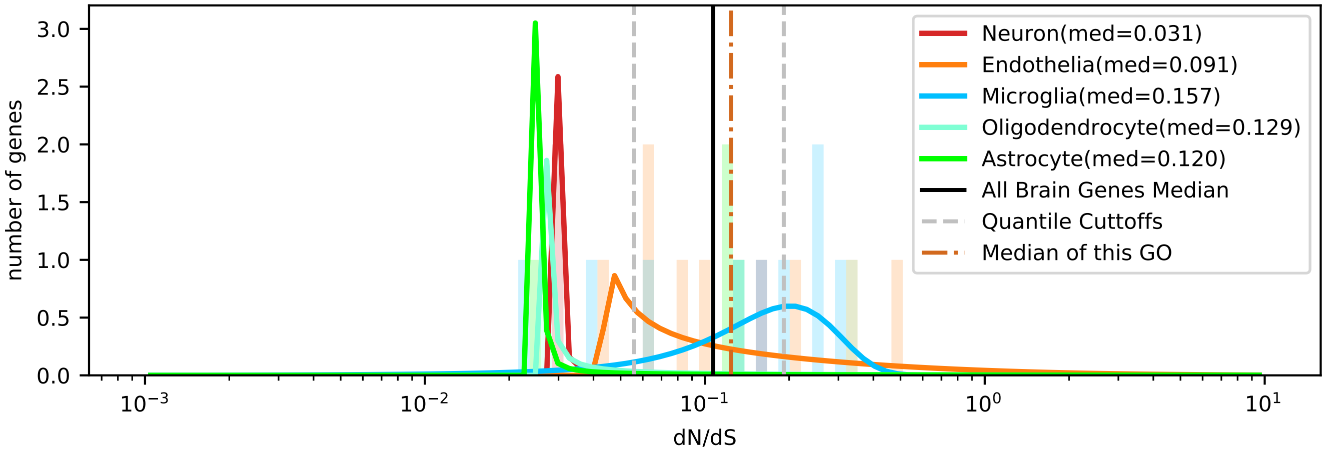


| | < cutoff | other genes | Total |
|---|---|---|---|
| In Astrocyte | 1 (1.04) | 3 (2.96) | 4 |
| Not IN Astrocyte | 6 (5.96) | 17 (17.04) | 23 |
| Total | 7 | 20 | 27 |

chi2: 0.328, p: 0.5671096294, OR: 0.94444

| | > cutoff | other genes | Total |
|---|---|---|---|
| In Astrocyte | 0 (1.19) | 4 (2.81) | 4 |
| Not IN Astrocyte | 8 (6.81) | 15 (16.19) | 23 |
| Total | 8 | 19 | 27 |

chi2: 0.661, p: 0.4162745582, OR: 0.00000

| | < cutoff | other genes | Total |
|---|---|---|---|
| In Microglia | 2 (2.33) | 7 (6.67) | 9 |
| Not IN Microglia | 5 (4.67) | 13 (13.33) | 18 |
| Total | 7 | 20 | 27 |

chi2: 0.024, p: 0.8766126033, OR: 0.74286

| | > cutoff | other genes | Total |
|---|---|---|---|
| In Microglia | 4 (2.67) | 5 (6.33) | 9 |
| Not IN Microglia | 4 (5.33) | 14 (12.67) | 18 |
| Total | 8 | 19 | 27 |

chi2: 0.555, p: 0.4562418259, OR: 2.80000

| | < cutoff | other genes | Total |
|---|---|---|---|
| In Oligodendrocyte | 1 (0.78) | 2 (2.22) | 3 |
| Not IN Oligodendrocyte | 6 (6.22) | 18 (17.78) | 24 |
| Total | 7 | 20 | 27 |

chi2: 0.151, p: 0.6978962433, OR: 1.50000

| | > cutoff | other genes | Total |
|---|---|---|---|
| In Oligodendrocyte | 1 (0.89) | 2 (2.11) | 3 |
| Not IN Oligodendrocyte | 7 (7.11) | 17 (16.89) | 24 |
| Total | 8 | 19 | 27 |

chi2: 0.272, p: 0.6019943981, OR: 1.21429

| | < cutoff | other genes | Total |
|---|---|---|---|
| In Neuron | 2 (0.78) | 1 (2.22) | 3 |
| Not IN Neuron | 5 (6.22) | 19 (17.78) | 24 |
| Total | 7 | 20 | 27 |

chi2: 1.019, p: 0.3128687148, OR: 7.60000

| | > cutoff | other genes | Total |
|---|---|---|---|
| In Neuron | 0 (0.89) | 3 (2.11) | 3 |
| Not IN Neuron | 8 (7.11) | 16 (16.89) | 24 |
| Total | 8 | 19 | 27 |

chi2: 0.272, p: 0.6019943981, OR: 0.00000

| | < cutoff | other genes | Total |
|---|---|---|---|
| In Endothelia | 1 (2.07) | 7 (5.93) | 8 |
| Not IN Endothelia | 6 (4.93) | 13 (14.07) | 19 |
| Total | 7 | 20 | 27 |

chi2: 0.305, p: 0.5808711140, OR: 0.30952

| | > cutoff | other genes | Total |
|---|---|---|---|
| In Endothelia | 3 (2.37) | 5 (5.63) | 8 |
| Not IN Endothelia | 5 (5.63) | 14 (13.37) | 19 |
| Total | 8 | 19 | 27 |

chi2: 0.014, p: 0.9047617990, OR: 1.68000

### Five Celltypes Distribution of Average dN/dS Scores of Genes Related to plasma membrane organization

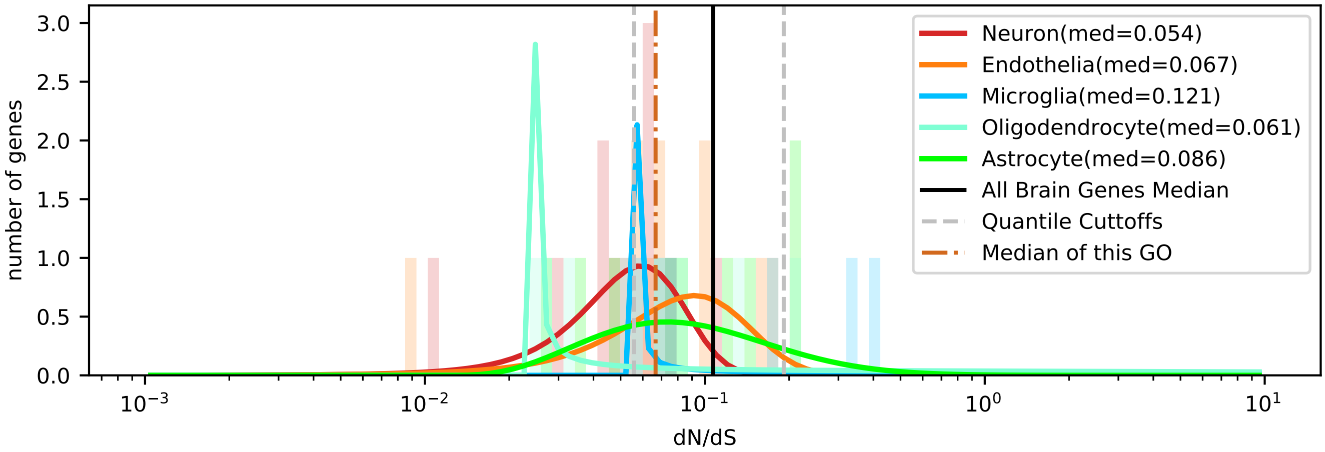


| | < cutoff | other genes | Total |
|---|---|---|---|
| In Astrocyte | 3 (2.86) | 6 (6.14) | 9 |
| Not IN Astrocyte | 11 (11.14) | 24 (23.86) | 35 |
| Total | 14 | 30 | 44 |

chi2: 0.085, p: 0.7704492491, OR: 1.09091

| | > cutoff | other genes | Total |
|---|---|---|---|
| In Astrocyte | 2 (1.02) | 7 (7.98) | 9 |
| Not IN Astrocyte | 3 (3.98) | 32 (31.02) | 35 |
| Total | 5 | 39 | 44 |

chi2: 0.316, p: 0.5740836301, OR: 3.04762

| | < cutoff | other genes | Total |
|---|---|---|---|
| In Microglia | 0 (1.91) | 6 (4.09) | 6 |
| Not IN Microglia | 14 (12.09) | 24 (25.91) | 38 |
| Total | 14 | 30 | 44 |

chi2: 1.766, p: 0.1838469824, OR: 0.00000

| | > cutoff | other genes | Total |
|---|---|---|---|
| In Microglia | 2 (0.68) | 4 (5.32) | 6 |
| Not IN Microglia | 3 (4.32) | 35 (33.68) | 38 |
| Total | 5 | 39 | 44 |

chi2: 1.283, p: 0.2574178207, OR: 5.83333

| | < cutoff | other genes | Total |
|---|---|---|---|
| In Oligodendrocyte | 4 (2.55) | 4 (5.45) | 8 |
| Not IN Oligodendrocyte | 10 (11.45) | 26 (24.55) | 36 |
| Total | 14 | 30 | 44 |

chi2: 0.642, p: 0.4231079167, OR: 2.60000

| | > cutoff | other genes | Total |
|---|---|---|---|
| In Oligodendrocyte | 1 (0.91) | 7 (7.09) | 8 |
| Not IN Oligodendrocyte | 4 (4.09) | 32 (31.91) | 36 |
| Total | 5 | 39 | 44 |

chi2: 0.254, p: 0.6143798085, OR: 1.14286

| | < cutoff | other genes | Total |
|---|---|---|---|
| In Neuron | 6 (3.50) | 5 (7.50) | 11 |
| Not IN Neuron | 8 (10.50) | 25 (22.50) | 33 |
| Total | 14 | 30 | 44 |

chi2: 2.235, p: 0.1349235520, OR: 3.75000

| | > cutoff | other genes | Total |
|---|---|---|---|
| In Neuron | 0 (1.25) | 11 (9.75) | 11 |
| Not IN Neuron | 5 (3.75) | 28 (29.25) | 33 |
| Total | 5 | 39 | 44 |

chi2: 0.677, p: 0.4106482693, OR: 0.00000

| | < cutoff | other genes | Total |
|---|---|---|---|
| In Endothelia | 1 (3.18) | 9 (6.82) | 10 |
| Not IN Endothelia | 13 (10.82) | 21 (23.18) | 34 |
| Total | 14 | 30 | 44 |

chi2: 1.687, p: 0.1939595286, OR: 0.17949

| | > cutoff | other genes | Total |
|---|---|---|---|
| In Endothelia | 0 (1.14) | 10 (8.86) | 10 |
| Not IN Endothelia | 5 (3.86) | 29 (30.14) | 34 |
| Total | 5 | 39 | 44 |

chi2: 0.520, p: 0.4707130312, OR: 0.00000

### Five Celltypes Distribution of Average dN/dS Scores of Genes Related to protein folding

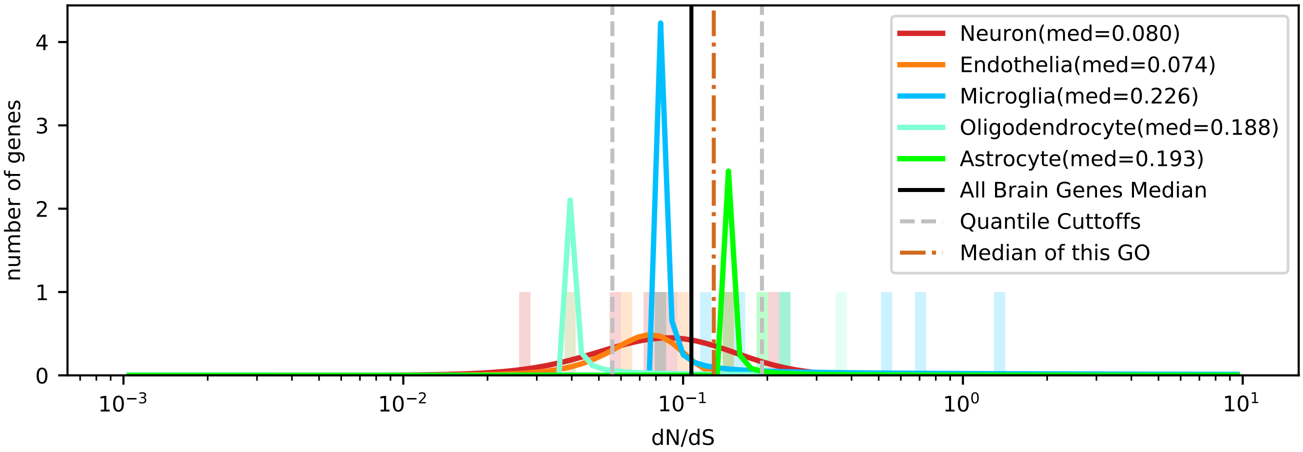


| | < cutoff | other genes | Total |
|---|---|---|---|
| In Astrocyte | 0 (0.38) | 3 (2.62) | 3 |
| Not IN Astrocyte | 3 (2.62) | 18 (18.38) | 21 |
| Total | 3 | 21 | 24 |

chi2: 0.054, p: 0.8155403095, OR: 0.00000

| | > cutoff | other genes | Total |
|---|---|---|---|
| In Astrocyte | 2 (1.00) | 1 (2.00) | 3 |
| Not IN Astrocyte | 6 (7.00) | 15 (14.00) | 21 |
| Total | 8 | 16 | 24 |

chi2: 0.429, p: 0.5126907603, OR: 5.00000

| | < cutoff | other genes | Total |
|---|---|---|---|
| In Microglia | 0 (0.88) | 7 (6.12) | 7 |
| Not IN Microglia | 3 (2.12) | 14 (14.88) | 17 |
| Total | 3 | 21 | 24 |

chi2: 0.259, p: 0.6105989114, OR: 0.00000

| | > cutoff | other genes | Total |
|---|---|---|---|
| In Microglia | 4 (2.33) | 3 (4.67) | 7 |
| Not IN Microglia | 4 (5.67) | 13 (11.33) | 17 |
| Total | 8 | 16 | 24 |

chi2: 1.235, p: 0.2663799233, OR: 4.33333

| | < cutoff | other genes | Total |
|---|---|---|---|
| In Oligodendrocyte | 1 (0.38) | 2 (2.62) | 3 |
| Not IN Oligodendrocyte | 2 (2.62) | 19 (18.38) | 21 |
| Total | 3 | 21 | 24 |

chi2: 0.054, p: 0.8155403095, OR: 4.75000

| | > cutoff | other genes | Total |
|---|---|---|---|
| In Oligodendrocyte | 1 (1.00) | 2 (2.00) | 3 |
| Not IN Oligodendrocyte | 7 (7.00) | 14 (14.00) | 21 |
| Total | 8 | 16 | 24 |

chi2: 0.000, p: 1.0000000000, OR: 1.00000

| | < cutoff | other genes | Total |
|---|---|---|---|
| In Neuron | 1 (0.88) | 6 (6.12) | 7 |
| Not IN Neuron | 2 (2.12) | 15 (14.88) | 17 |
| Total | 3 | 21 | 24 |

chi2: 0.259, p: 0.6105989114, OR: 1.25000

| | > cutoff | other genes | Total |
|---|---|---|---|
| In Neuron | 1 (2.33) | 6 (4.67) | 7 |
| Not IN Neuron | 7 (5.67) | 10 (11.33) | 17 |
| Total | 8 | 16 | 24 |

chi2: 0.630, p: 0.4272628567, OR: 0.23810

| | < cutoff | other genes | Total |
|---|---|---|---|
| In Endothelia | 1 (0.50) | 3 (3.50) | 4 |
| Not IN Endothelia | 2 (2.50) | 18 (17.50) | 20 |
| Total | 3 | 21 | 24 |

chi2: 0.000, p: 1.0000000000, OR: 3.00000

| | > cutoff | other genes | Total |
|---|---|---|---|
| In Endothelia | 0 (1.33) | 4 (2.67) | 4 |
| Not IN Endothelia | 8 (6.67) | 12 (13.33) | 20 |
| Total | 8 | 16 | 24 |

chi2: 0.938, p: 0.3329216081, OR: 0.00000

### Five Celltypes Distribution of Average dN/dS Scores of Genes Related to protein maturation

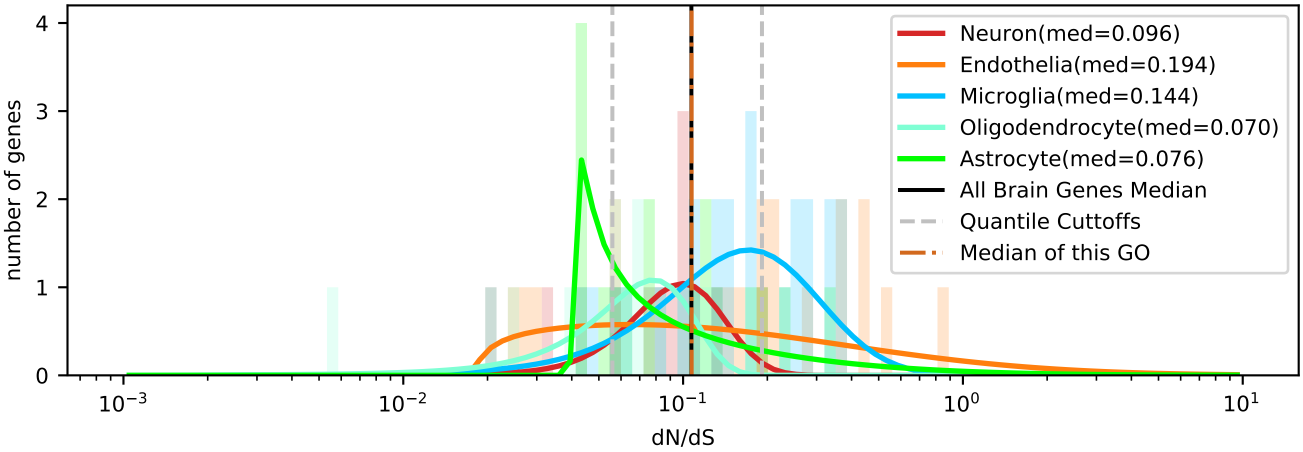


| | < cutoff | other genes | Total |
|---|---|---|---|
| In Astrocyte | 5 (3.95) | 12 (13.05) | 17 |
| Not IN Astrocyte | 15 (16.05) | 54 (52.95) | 69 |
| Total | 20 | 66 | 86 |

chi2: 0.123, p: 0.7261311330, OR: 1.50000

| | > cutoff | other genes | Total |
|---|---|---|---|
| In Astrocyte | 2 (4.35) | 15 (12.65) | 17 |
| Not IN Astrocyte | 20 (17.65) | 49 (51.35) | 69 |
| Total | 22 | 64 | 86 |

chi2: 1.316, p: 0.2512368917, OR: 0.32667

| | < cutoff | other genes | Total |
|---|---|---|---|
| In Microglia | 3 (5.81) | 22 (19.19) | 25 |
| Not IN Microglia | 17 (14.19) | 44 (46.81) | 61 |
| Total | 20 | 66 | 86 |

chi2: 1.692, p: 0.1933578043, OR: 0.35294

| | > cutoff | other genes | Total |
|---|---|---|---|
| In Microglia | 9 (6.40) | 16 (18.60) | 25 |
| Not IN Microglia | 13 (15.60) | 48 (45.40) | 61 |
| Total | 22 | 64 | 86 |

chi2: 1.312, p: 0.2520059636, OR: 2.07692

| | < cutoff | other genes | Total |
|---|---|---|---|
| In Oligodendrocyte | 4 (3.02) | 9 (9.98) | 13 |
| Not IN Oligodendrocyte | 16 (16.98) | 57 (56.02) | 73 |
| Total | 20 | 66 | 86 |

chi2: 0.115, p: 0.7340721736, OR: 1.58333

| | > cutoff | other genes | Total |
|---|---|---|---|
| In Oligodendrocyte | 0 (3.33) | 13 (9.67) | 13 |
| Not IN Oligodendrocyte | 22 (18.67) | 51 (54.33) | 73 |
| Total | 22 | 64 | 86 |

chi2: 3.801, p: 0.0512369071, OR: 0.00000

| | < cutoff | other genes | Total |
|---|---|---|---|
| In Neuron | 1 (2.56) | 10 (8.44) | 11 |
| Not IN Neuron | 19 (17.44) | 56 (57.56) | 75 |
| Total | 20 | 66 | 86 |

chi2: 0.654, p: 0.4186992612, OR: 0.29474

| | > cutoff | other genes | Total |
|---|---|---|---|
| In Neuron | 0 (2.81) | 11 (8.19) | 11 |
| Not IN Neuron | 22 (19.19) | 53 (55.81) | 75 |
| Total | 22 | 64 | 86 |

chi2: 2.932, p: 0.0868456497, OR: 0.00000

| | < cutoff | other genes | Total |
|---|---|---|---|
| In Endothelia | 7 (4.65) | 13 (15.35) | 20 |
| Not IN Endothelia | 13 (15.35) | 53 (50.65) | 66 |
| Total | 20 | 66 | 86 |

chi2: 1.248, p: 0.2639731608, OR: 2.19527

| | > cutoff | other genes | Total |
|---|---|---|---|
| In Endothelia | 11 (5.12) | 9 (14.88) | 20 |
| Not IN Endothelia | 11 (16.88) | 55 (49.12) | 66 |
| Total | 22 | 64 | 86 |

chi2: 9.919, p: 0.0016354885, OR: 6.11111

### Five Celltypes Distribution of Average dN/dS Scores of Genes Related to protein targeting

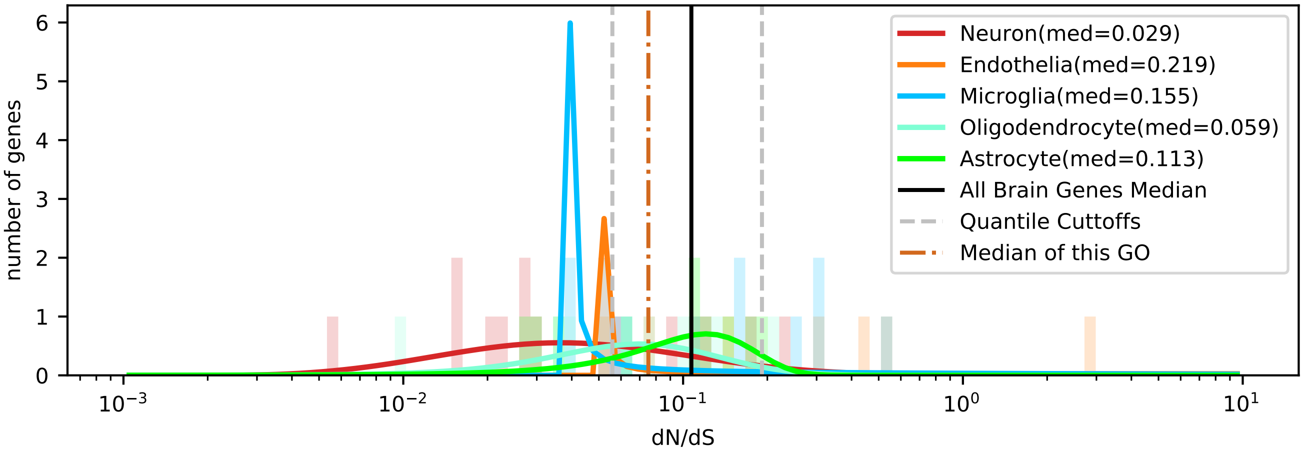


| | < cutoff | other genes | Total |
|---|---|---|---|
| In Astrocyte | 3 (3.89) | 7 (6.11) | 10 |
| Not IN Astrocyte | 18 (17.11) | 26 (26.89) | 44 |
| Total | 21 | 33 | 54 |

chi2: 0.078, p: 0.7798900710, OR: 0.61905

| | > cutoff | other genes | Total |
|---|---|---|---|
| In Astrocyte | 0 (1.85) | 10 (8.15) | 10 |
| Not IN Astrocyte | 10 (8.15) | 34 (35.85) | 44 |
| Total | 10 | 44 | 54 |

chi2: 1.486, p: 0.2227766973, OR: 0.00000

| | < cutoff | other genes | Total |
|---|---|---|---|
| In Microglia | 2 (4.28) | 9 (6.72) | 11 |
| Not IN Microglia | 19 (16.72) | 24 (26.28) | 43 |
| Total | 21 | 33 | 54 |

chi2: 1.518, p: 0.2178852751, OR: 0.28070

| | > cutoff | other genes | Total |
|---|---|---|---|
| In Microglia | 4 (2.04) | 7 (8.96) | 11 |
| Not IN Microglia | 6 (7.96) | 37 (35.04) | 43 |
| Total | 10 | 44 | 54 |

chi2: 1.619, p: 0.2031858781, OR: 3.52381

| | < cutoff | other genes | Total |
|---|---|---|---|
| In Oligodendrocyte | 5 (3.89) | 5 (6.11) | 10 |
| Not IN Oligodendrocyte | 16 (17.11) | 28 (26.89) | 44 |
| Total | 21 | 33 | 54 |

chi2: 0.193, p: 0.6605492052, OR: 1.75000

| | > cutoff | other genes | Total |
|---|---|---|---|
| In Oligodendrocyte | 1 (1.85) | 9 (8.15) | 10 |
| Not IN Oligodendrocyte | 9 (8.15) | 35 (35.85) | 44 |
| Total | 10 | 44 | 54 |

chi2: 0.101, p: 0.7510006043, OR: 0.43210

| | < cutoff | other genes | Total |
|---|---|---|---|
| In Neuron | 10 (5.83) | 5 (9.17) | 15 |
| Not IN Neuron | 11 (15.17) | 28 (23.83) | 39 |
| Total | 21 | 33 | 54 |

chi2: 5.222, p: 0.0223031682, OR: 5.09091

| | > cutoff | other genes | Total |
|---|---|---|---|
| In Neuron | 1 (2.78) | 14 (12.22) | 15 |
| Not IN Neuron | 9 (7.22) | 30 (31.78) | 39 |
| Total | 10 | 44 | 54 |

chi2: 0.999, p: 0.3175983365, OR: 0.23810

| | < cutoff | other genes | Total |
|---|---|---|---|
| In Endothelia | 1 (3.11) | 7 (4.89) | 8 |
| Not IN Endothelia | 20 (17.89) | 26 (28.11) | 46 |
| Total | 21 | 33 | 54 |

chi2: 1.603, p: 0.2055215357, OR: 0.18571

| | > cutoff | other genes | Total |
|---|---|---|---|
| In Endothelia | 4 (1.48) | 4 (6.52) | 8 |
| Not IN Endothelia | 6 (8.52) | 40 (37.48) | 46 |
| Total | 10 | 44 | 54 |

chi2: 3.962, p: 0.0465303620, OR: 6.66667

### Five Celltypes Distribution of Average dN/dS Scores of Genes Related to protein-containing complex assembly

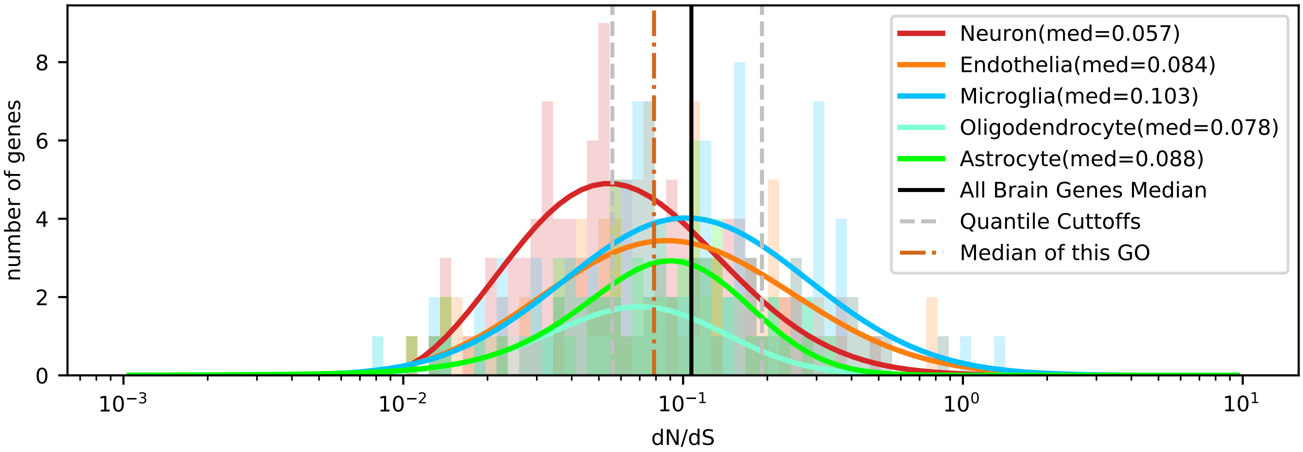


| | < cutoff | other genes | Total |
|---|---|---|---|
| In Astrocyte | 16 (19.45) | 40 (36.55) | 56 |
| Not IN Astrocyte | 124 (120.55) | 223 (226.45) | 347 |
| Total | 140 | 263 | 403 |

chi2: 0.798, p: 0.3716040936, OR: 0.71935

| | > cutoff | other genes | Total |
|---|---|---|---|
| In Astrocyte | 6 (10.00) | 50 (46.00) | 56 |
| Not IN Astrocyte | 66 (62.00) | 281 (285.00) | 347 |
| Total | 72 | 331 | 403 |

chi2: 1.736, p: 0.1876188914, OR: 0.51091

| | < cutoff | other genes | Total |
|---|---|---|---|
| In Microglia | 26 (37.52) | 82 (70.48) | 108 |
| Not IN Microglia | 114 (102.48) | 181 (192.52) | 295 |
| Total | 140 | 263 | 403 |

chi2: 6.774, p: 0.0092500112, OR: 0.50342

| | > cutoff | other genes | Total |
|---|---|---|---|
| In Microglia | 29 (19.30) | 79 (88.70) | 108 |
| Not IN Microglia | 43 (52.70) | 252 (242.30) | 295 |
| Total | 72 | 331 | 403 |

chi2: 7.303, p: 0.0068821389, OR: 2.15131

| | < cutoff | other genes | Total |
|---|---|---|---|
| In Oligodendrocyte | 15 (12.16) | 20 (22.84) | 35 |
| Not IN Oligodendrocyte | 125 (127.84) | 243 (240.16) | 368 |
| Total | 140 | 263 | 403 |

chi2: 0.756, p: 0.3844368590, OR: 1.45800

| | > cutoff | other genes | Total |
|---|---|---|---|
| In Oligodendrocyte | 2 (6.25) | 33 (28.75) | 35 |
| Not IN Oligodendrocyte | 70 (65.75) | 298 (302.25) | 368 |
| Total | 72 | 331 | 403 |

chi2: 3.003, p: 0.0830877488, OR: 0.25801

| | < cutoff | other genes | Total |
|---|---|---|---|
| In Neuron | 55 (38.56) | 56 (72.44) | 111 |
| Not IN Neuron | 85 (101.44) | 207 (190.56) | 292 |
| Total | 140 | 263 | 403 |

chi2: 13.933, p: 0.0001893971, OR: 2.39181

| | > cutoff | other genes | Total |
|---|---|---|---|
| In Neuron | 10 (19.83) | 101 (91.17) | 111 |
| Not IN Neuron | 62 (52.17) | 230 (239.83) | 292 |
| Total | 72 | 331 | 403 |

chi2: 7.378, p: 0.0066032007, OR: 0.36729

| | < cutoff | other genes | Total |
|---|---|---|---|
| In Endothelia | 28 (32.31) | 65 (60.69) | 93 |
| Not IN Endothelia | 112 (107.69) | 198 (202.31) | 310 |
| Total | 140 | 263 | 403 |

chi2: 0.894, p: 0.3444107038, OR: 0.76154

| | > cutoff | other genes | Total |
|---|---|---|---|
| In Endothelia | 25 (16.62) | 68 (76.38) | 93 |
| Not IN Endothelia | 47 (55.38) | 263 (254.62) | 310 |
| Total | 72 | 331 | 403 |

chi2: 5.922, p: 0.0149526714, OR: 2.05726

### Five Celltypes Distribution of Average dN/dS Scores of Genes Related to regulation_of_cell_size

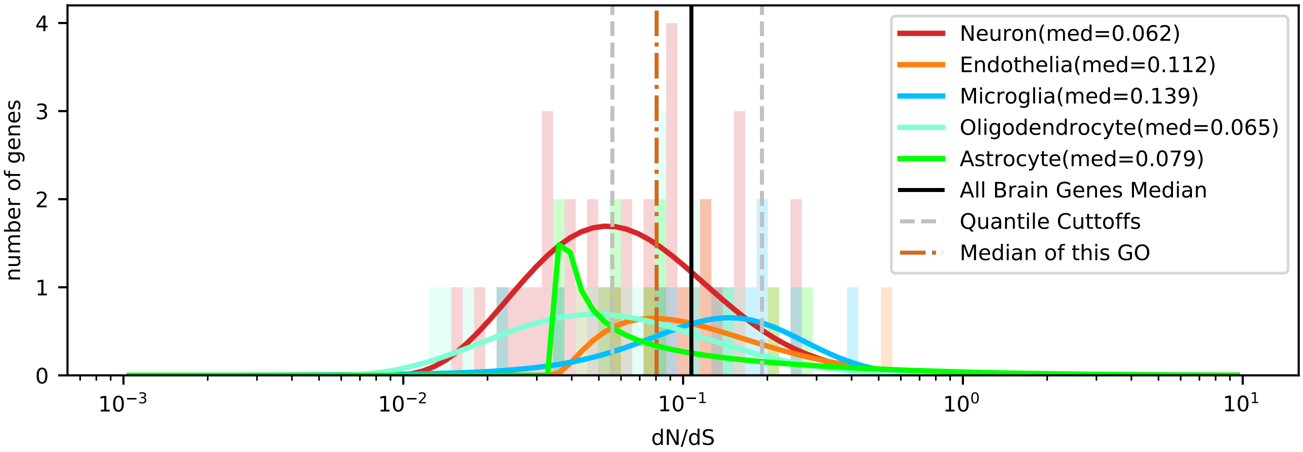


| | < cutoff | other genes | Total |
|---|---|---|---|
| In Astrocyte | 5 (4.06) | 6 (6.94) | 11 |
| Not IN Astrocyte | 26 (26.94) | 47 (46.06) | 73 |
| Total | 31 | 53 | 84 |

chi2: 0.087, p: 0.7678159397, OR: 1.50641

| | > cutoff | other genes | Total |
|---|---|---|---|
| In Astrocyte | 2 (1.31) | 9 (9.69) | 11 |
| Not IN Astrocyte | 8 (8.69) | 65 (64.31) | 73 |
| Total | 10 | 74 | 84 |

chi2: 0.036, p: 0.8491265062, OR: 1.80556

| | < cutoff | other genes | Total |
|---|---|---|---|
| In Microglia | 2 (4.43) | 10 (7.57) | 12 |
| Not IN Microglia | 29 (26.57) | 43 (45.43) | 72 |
| Total | 31 | 53 | 84 |

chi2: 1.553, p: 0.2127001865, OR: 0.29655

| | > cutoff | other genes | Total |
|---|---|---|---|
| In Microglia | 3 (1.43) | 9 (10.57) | 12 |
| Not IN Microglia | 7 (8.57) | 65 (63.43) | 72 |
| Total | 10 | 74 | 84 |

chi2: 1.064, p: 0.3022616690, OR: 3.09524

| | < cutoff | other genes | Total |
|---|---|---|---|
| In Oligodendrocyte | 7 (5.90) | 9 (10.10) | 16 |
| Not IN Oligodendrocyte | 24 (25.10) | 44 (42.90) | 68 |
| Total | 31 | 53 | 84 |

chi2: 0.117, p: 0.7317871648, OR: 1.42593

| | > cutoff | other genes | Total |
|---|---|---|---|
| In Oligodendrocyte | 1 (1.90) | 15 (14.10) | 16 |
| Not IN Oligodendrocyte | 9 (8.10) | 59 (59.90) | 68 |
| Total | 10 | 74 | 84 |

chi2: 0.121, p: 0.7283758809, OR: 0.43704

| | < cutoff | other genes | Total |
|---|---|---|---|
| In Neuron | 16 (12.55) | 18 (21.45) | 34 |
| Not IN Neuron | 15 (18.45) | 35 (31.55) | 50 |
| Total | 31 | 53 | 84 |

chi2: 1.850, p: 0.1738208338, OR: 2.07407

| | > cutoff | other genes | Total |
|---|---|---|---|
| In Neuron | 2 (4.05) | 32 (29.95) | 34 |
| Not IN Neuron | 8 (5.95) | 42 (44.05) | 50 |
| Total | 10 | 74 | 84 |

chi2: 1.128, p: 0.2881045172, OR: 0.32812

| | < cutoff | other genes | Total |
|---|---|---|---|
| In Endothelia | 1 (4.06) | 10 (6.94) | 11 |
| Not IN Endothelia | 30 (26.94) | 43 (46.06) | 73 |
| Total | 31 | 53 | 84 |

chi2: 2.943, p: 0.0862457618, OR: 0.14333

| | > cutoff | other genes | Total |
|---|---|---|---|
| In Endothelia | 2 (1.31) | 9 (9.69) | 11 |
| Not IN Endothelia | 8 (8.69) | 65 (64.31) | 73 |
| Total | 10 | 74 | 84 |

chi2: 0.036, p: 0.8491265062, OR: 1.80556

### Five Celltypes Distribution of Average dN/dS Scores of Genes Related to regulation_of_membrane_potential

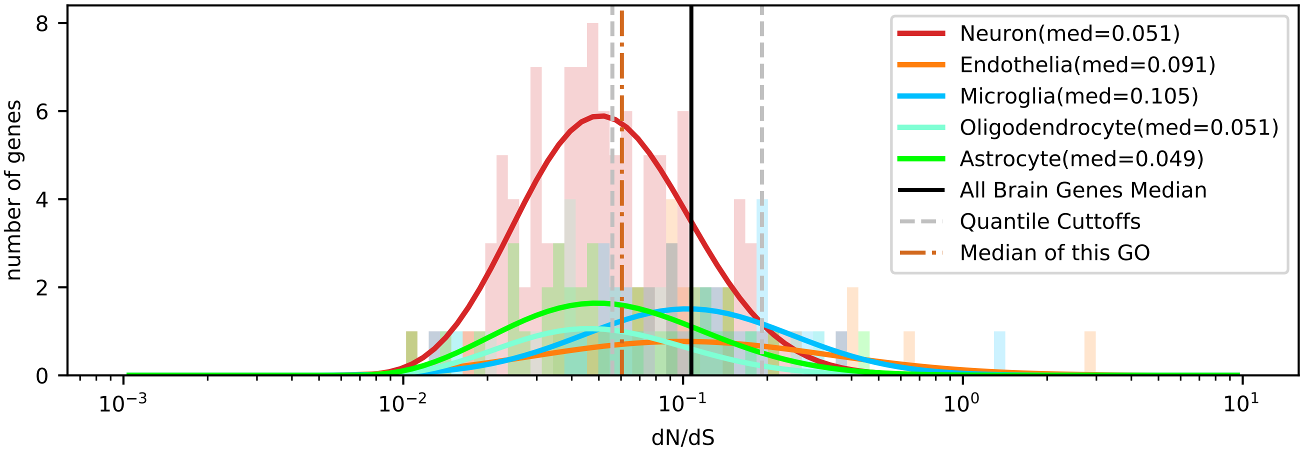


| | < cutoff | other genes | Total |
|---|---|---|---|
| In Astrocyte | 21 (16.81) | 15 (19.19) | 36 |
| Not IN Astrocyte | 85 (89.19) | 106 (101.81) | 191 |
| Total | 106 | 121 | 227 |

chi2: 1.805, p: 0.1790635587, OR: 1.74588

| | > cutoff | other genes | Total |
|---|---|---|---|
| In Astrocyte | 2 (3.17) | 34 (32.83) | 36 |
| Not IN Astrocyte | 18 (16.83) | 173 (174.17) | 191 |
| Total | 20 | 207 | 227 |

chi2: 0.185, p: 0.6667299349, OR: 0.56536

| | < cutoff | other genes | Total |
|---|---|---|---|
| In Microglia | 8 (16.34) | 27 (18.66) | 35 |
| Not IN Microglia | 98 (89.66) | 94 (102.34) | 192 |
| Total | 106 | 121 | 227 |

chi2: 8.349, p: 0.0038583906, OR: 0.28420

| | > cutoff | other genes | Total |
|---|---|---|---|
| In Microglia | 7 (3.08) | 28 (31.92) | 35 |
| Not IN Microglia | 13 (16.92) | 179 (175.08) | 192 |
| Total | 20 | 207 | 227 |

chi2: 4.907, p: 0.0267473964, OR: 3.44231

| | < cutoff | other genes | Total |
|---|---|---|---|
| In Oligodendrocyte | 11 (9.81) | 10 (11.19) | 21 |
| Not IN Oligodendrocyte | 95 (96.19) | 111 (109.81) | 206 |
| Total | 106 | 121 | 227 |

chi2: 0.101, p: 0.7500527525, OR: 1.28526

| | > cutoff | other genes | Total |
|---|---|---|---|
| In Oligodendrocyte | 1 (1.85) | 20 (19.15) | 21 |
| Not IN Oligodendrocyte | 19 (18.15) | 187 (187.85) | 206 |
| Total | 20 | 207 | 227 |

chi2: 0.080, p: 0.7771517631, OR: 0.49211

| | < cutoff | other genes | Total |
|---|---|---|---|
| In Neuron | 61 (51.83) | 50 (59.17) | 111 |
| Not IN Neuron | 45 (54.17) | 71 (61.83) | 116 |
| Total | 106 | 121 | 227 |

chi2: 5.321, p: 0.0210714163, OR: 1.92489

| | > cutoff | other genes | Total |
|---|---|---|---|
| In Neuron | 4 (9.78) | 107 (101.22) | 111 |
| Not IN Neuron | 16 (10.22) | 100 (105.78) | 116 |
| Total | 20 | 207 | 227 |

chi2: 6.117, p: 0.0133906887, OR: 0.23364

| | < cutoff | other genes | Total |
|---|---|---|---|
| In Endothelia | 5 (11.21) | 19 (12.79) | 24 |
| Not IN Endothelia | 101 (94.79) | 102 (108.21) | 203 |
| Total | 106 | 121 | 227 |

chi2: 6.097, p: 0.0135426656, OR: 0.26576

| | > cutoff | other genes | Total |
|---|---|---|---|
| In Endothelia | 6 (2.11) | 18 (21.89) | 24 |
| Not IN Endothelia | 14 (17.89) | 189 (185.11) | 203 |
| Total | 20 | 207 | 227 |

chi2: 6.647, p: 0.0099339729, OR: 4.50000

### Five Celltypes Distribution of Average dN/dS Scores of Genes Related to reproduction

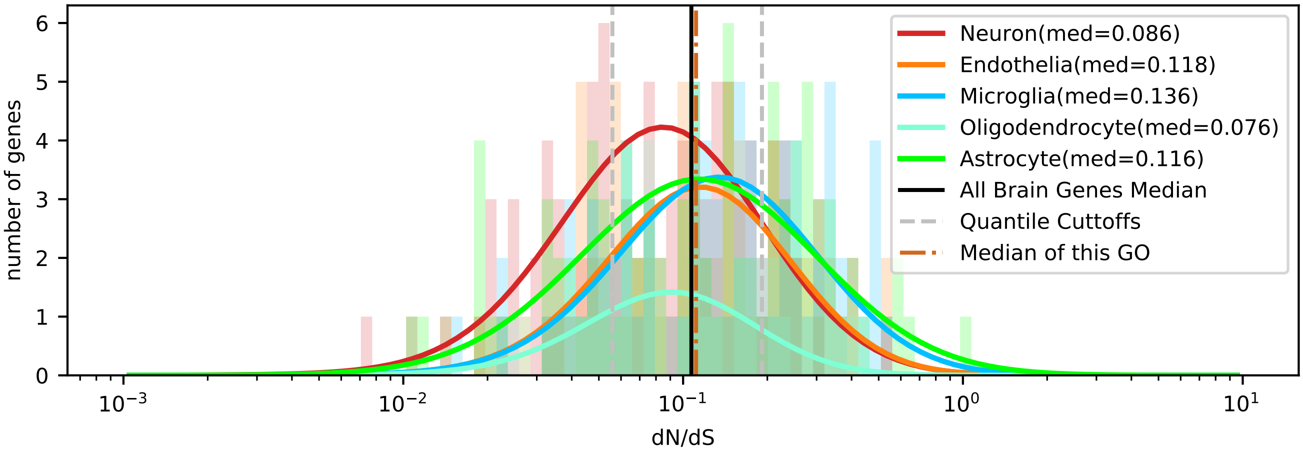


| | < cutoff | other genes | Total |
|---|---|---|---|
| In Astrocyte | 23 (23.00) | 65 (65.00) | 88 |
| Not IN Astrocyte | 69 (69.00) | 195 (195.00) | 264 |
| Total | 92 | 260 | 352 |

chi2: 0.000, p: 1.0000000000, OR: 1.00000

| | > cutoff | other genes | Total |
|---|---|---|---|
| In Astrocyte | 32 (24.00) | 56 (64.00) | 88 |
| Not IN Astrocyte | 64 (72.00) | 200 (192.00) | 264 |
| Total | 96 | 256 | 352 |

chi2: 4.297, p: 0.0381824719, OR: 1.78571

| | < cutoff | other genes | Total |
|---|---|---|---|
| In Microglia | 14 (19.60) | 61 (55.40) | 75 |
| Not IN Microglia | 78 (72.40) | 199 (204.60) | 277 |
| Total | 92 | 260 | 352 |

chi2: 2.285, p: 0.1306447979, OR: 0.58554

| | > cutoff | other genes | Total |
|---|---|---|---|
| In Microglia | 26 (20.45) | 49 (54.55) | 75 |
| Not IN Microglia | 70 (75.55) | 207 (201.45) | 277 |
| Total | 96 | 256 | 352 |

chi2: 2.175, p: 0.1403073750, OR: 1.56910

| | < cutoff | other genes | Total |
|---|---|---|---|
| In Oligodendrocyte | 6 (7.06) | 21 (19.94) | 27 |
| Not IN Oligodendrocyte | 86 (84.94) | 239 (240.06) | 325 |
| Total | 92 | 260 | 352 |

chi2: 0.064, p: 0.7996357896, OR: 0.79402

| | > cutoff | other genes | Total |
|---|---|---|---|
| In Oligodendrocyte | 4 (7.36) | 23 (19.64) | 27 |
| Not IN Oligodendrocyte | 92 (88.64) | 233 (236.36) | 325 |
| Total | 96 | 256 | 352 |

chi2: 1.658, p: 0.1978111719, OR: 0.44045

| | < cutoff | other genes | Total |
|---|---|---|---|
| In Neuron | 35 (25.09) | 61 (70.91) | 96 |
| Not IN Neuron | 57 (66.91) | 199 (189.09) | 256 |
| Total | 92 | 260 | 352 |

chi2: 6.568, p: 0.0103812859, OR: 2.00316

| | > cutoff | other genes | Total |
|---|---|---|---|
| In Neuron | 16 (26.18) | 80 (69.82) | 96 |
| Not IN Neuron | 80 (69.82) | 176 (186.18) | 256 |
| Total | 96 | 256 | 352 |

chi2: 6.769, p: 0.0092758904, OR: 0.44000

| | < cutoff | other genes | Total |
|---|---|---|---|
| In Endothelia | 14 (17.25) | 52 (48.75) | 66 |
| Not IN Endothelia | 78 (74.75) | 208 (211.25) | 286 |
| Total | 92 | 260 | 352 |

chi2: 0.731, p: 0.3927200697, OR: 0.71795

| | > cutoff | other genes | Total |
|---|---|---|---|
| In Endothelia | 18 (18.00) | 48 (48.00) | 66 |
| Not IN Endothelia | 78 (78.00) | 208 (208.00) | 286 |
| Total | 96 | 256 | 352 |

chi2: 0.000, p: 1.0000000000, OR: 1.00000

### Five Celltypes Distribution of Average dN/dS Scores of Genes Related to response to stress

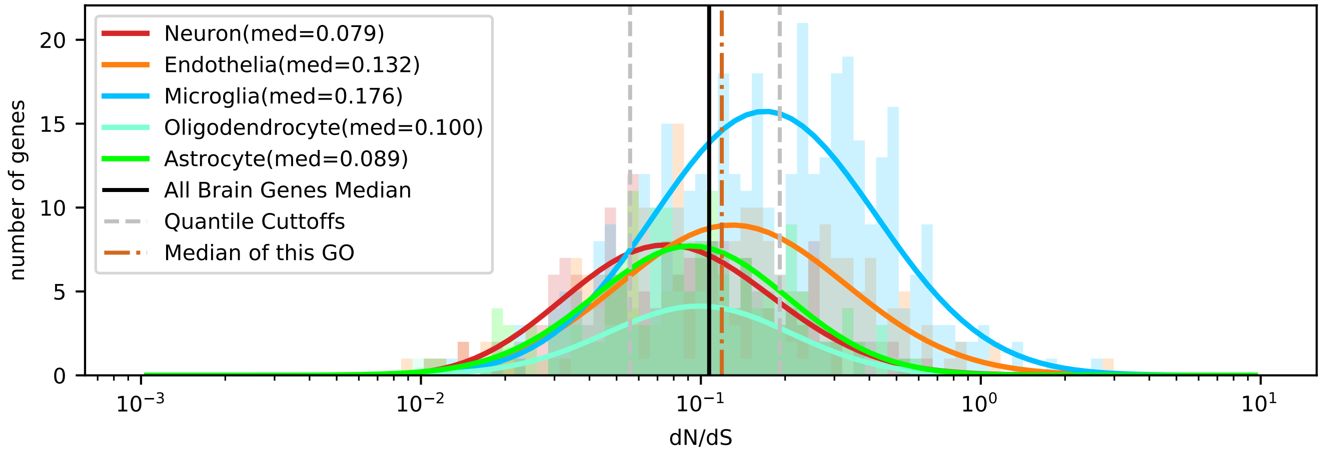


| | < cutoff | other genes | Total |
|---|---|---|---|
| In Astrocyte | 42 (34.16) | 124 (131.84) | 166 |
| Not IN Astrocyte | 171 (178.84) | 698 (690.16) | 869 |
| Total | 213 | 822 | 1035 |

chi2: 2.364, p: 0.1242015879, OR: 1.38257

| | > cutoff | other genes | Total |
|---|---|---|---|
| In Astrocyte | 32 (54.21) | 134 (111.79) | 166 |
| Not IN Astrocyte | 306 (283.79) | 563 (585.21) | 869 |
| Total | 338 | 697 | 1035 |

chi2: 15.378, p: 0.0000880258, OR: 0.43937

| | < cutoff | other genes | Total |
|---|---|---|---|
| In Microglia | 47 (79.44) | 339 (306.56) | 386 |
| Not IN Microglia | 166 (133.56) | 483 (515.44) | 649 |
| Total | 213 | 822 | 1035 |

chi2: 25.784, p: 0.0000003819, OR: 0.40340

| | > cutoff | other genes | Total |
|---|---|---|---|
| In Microglia | 186 (126.06) | 200 (259.94) | 386 |
| Not IN Microglia | 152 (211.94) | 497 (437.06) | 649 |
| Total | 338 | 697 | 1035 |

chi2: 66.383, p: 0.0000000000, OR: 3.04086

| | < cutoff | other genes | Total |
|---|---|---|---|
| In Oligodendrocyte | 21 (17.49) | 64 (67.51) | 85 |
| Not IN Oligodendrocyte | 192 (195.51) | 758 (754.49) | 950 |
| Total | 213 | 822 | 1035 |

chi2: 0.709, p: 0.3997114788, OR: 1.29541

| | > cutoff | other genes | Total |
|---|---|---|---|
| In Oligodendrocyte | 15 (27.76) | 70 (57.24) | 85 |
| Not IN Oligodendrocyte | 323 (310.24) | 627 (639.76) | 950 |
| Total | 338 | 697 | 1035 |

chi2: 8.758, p: 0.0030826220, OR: 0.41597

| | < cutoff | other genes | Total |
|---|---|---|---|
| In Neuron | 62 (35.81) | 112 (138.19) | 174 |
| Not IN Neuron | 151 (177.19) | 710 (683.81) | 861 |
| Total | 213 | 822 | 1035 |

chi2: 27.899, p: 0.0000001278, OR: 2.60289

| | > cutoff | other genes | Total |
|---|---|---|---|
| In Neuron | 29 (56.82) | 145 (117.18) | 174 |
| Not IN Neuron | 309 (281.18) | 552 (579.82) | 861 |
| Total | 338 | 697 | 1035 |

chi2: 23.452, p: 0.0000012806, OR: 0.35728

| | < cutoff | other genes | Total |
|---|---|---|---|
| In Endothelia | 41 (46.10) | 183 (177.90) | 224 |
| Not IN Endothelia | 172 (166.90) | 639 (644.10) | 811 |
| Total | 213 | 822 | 1035 |

chi2: 0.737, p: 0.3905825417, OR: 0.83235

| | > cutoff | other genes | Total |
|---|---|---|---|
| In Endothelia | 76 (73.15) | 148 (150.85) | 224 |
| Not IN Endothelia | 262 (264.85) | 549 (546.15) | 811 |
| Total | 338 | 697 | 1035 |

chi2: 0.143, p: 0.7054533365, OR: 1.07603

### Five Celltypes Distribution of Average dN/dS Scores of Genes Related to ribonucleoprotein complex assembly

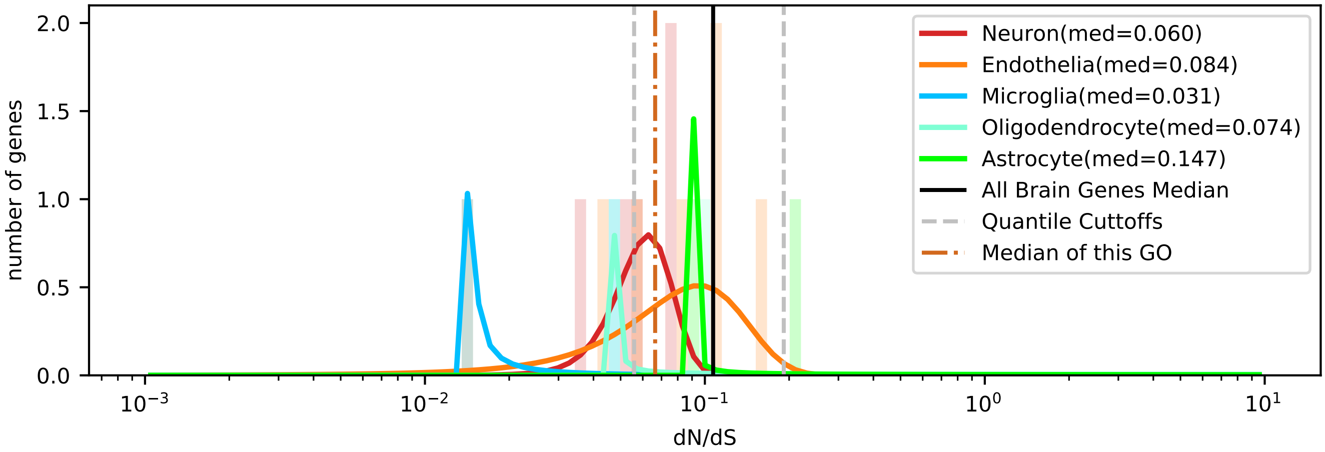


| | < cutoff | other genes | Total |
|---|---|---|---|
| In Astrocyte | 0 (0.78) | 2 (1.22) | 2 |
| Not IN Astrocyte | 7 (6.22) | 9 (9.78) | 16 |
| Total | 7 | 11 | 18 |

chi2: 0.183, p: 0.6691228454, OR: 0.00000

| | > cutoff | other genes | Total |
|---|---|---|---|
| In Astrocyte | 1 (0.11) | 1 (1.89) | 2 |
| Not IN Astrocyte | 0 (0.89) | 16 (15.11) | 16 |
| Total | 1 | 17 | 18 |

chi2: 1.621, p: 0.2029073390, OR: inf

| | < cutoff | other genes | Total |
|---|---|---|---|
| In Microglia | 2 (0.78) | 0 (1.22) | 2 |
| Not IN Microglia | 5 (6.22) | 11 (9.78) | 16 |
| Total | 7 | 11 | 18 |

chi2: 1.235, p: 0.2665185850, OR: inf

| | > cutoff | other genes | Total |
|---|---|---|---|
| In Microglia | 0 (0.11) | 2 (1.89) | 2 |
| Not IN Microglia | 1 (0.89) | 15 (15.11) | 16 |
| Total | 1 | 17 | 18 |

chi2: 1.621, p: 0.2029073390, OR: 0.00000

| | < cutoff | other genes | Total |
|---|---|---|---|
| In Oligodendrocyte | 1 (0.78) | 1 (1.22) | 2 |
| Not IN Oligodendrocyte | 6 (6.22) | 10 (9.78) | 16 |
| Total | 7 | 11 | 18 |

chi2: 0.183, p: 0.6691228454, OR: 1.66667

| | > cutoff | other genes | Total |
|---|---|---|---|
| In Oligodendrocyte | 0 (0.11) | 2 (1.89) | 2 |
| Not IN Oligodendrocyte | 1 (0.89) | 15 (15.11) | 16 |
| Total | 1 | 17 | 18 |

chi2: 1.621, p: 0.2029073390, OR: 0.00000

| | < cutoff | other genes | Total |
|---|---|---|---|
| In Neuron | 2 (1.94) | 3 (3.06) | 5 |
| Not IN Neuron | 5 (5.06) | 8 (7.94) | 13 |
| Total | 7 | 11 | 18 |

chi2: 0.230, p: 0.6313979306, OR: 1.06667

| | > cutoff | other genes | Total |
|---|---|---|---|
| In Neuron | 0 (0.28) | 5 (4.72) | 5 |
| Not IN Neuron | 1 (0.72) | 12 (12.28) | 13 |
| Total | 1 | 17 | 18 |

chi2: 0.261, p: 0.6096852764, OR: 0.00000

| | < cutoff | other genes | Total |
|---|---|---|---|
| In Endothelia | 2 (2.72) | 5 (4.28) | 7 |
| Not IN Endothelia | 5 (4.28) | 6 (6.72) | 11 |
| Total | 7 | 11 | 18 |

chi2: 0.049, p: 0.8255620449, OR: 0.48000

| | > cutoff | other genes | Total |
|---|---|---|---|
| In Endothelia | 0 (0.39) | 7 (6.61) | 7 |
| Not IN Endothelia | 1 (0.61) | 10 (10.39) | 11 |
| Total | 1 | 17 | 18 |

chi2: 0.055, p: 0.8145743753, OR: 0.00000

Five Celltypes Distribution of Average dN/dS Scores of Genes Related to ribosome biogenesis

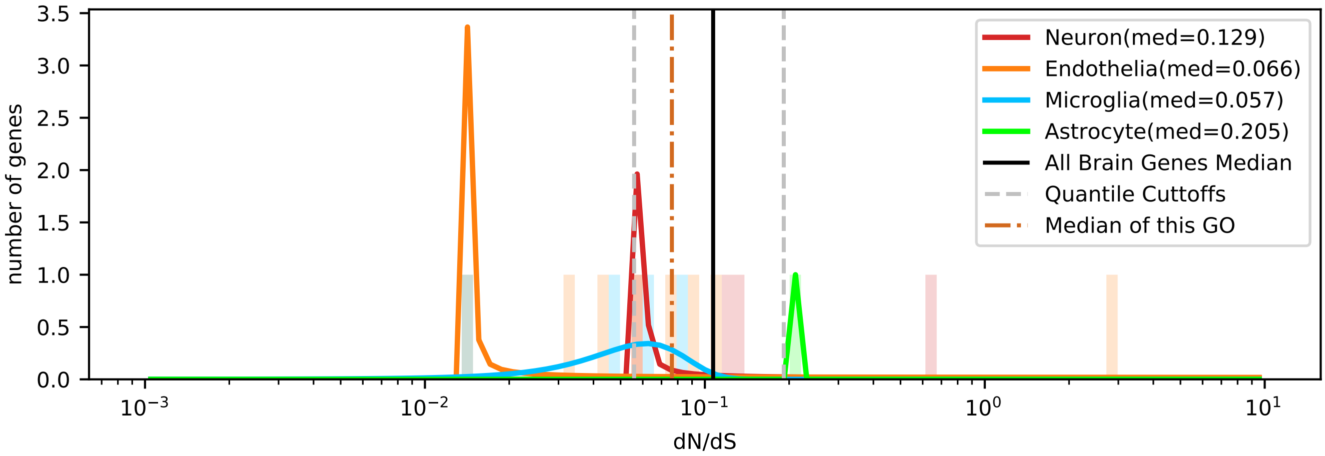

Five Celltypes Distribution of Average dN/dS Scores of Genes Related to secondary metabolic process

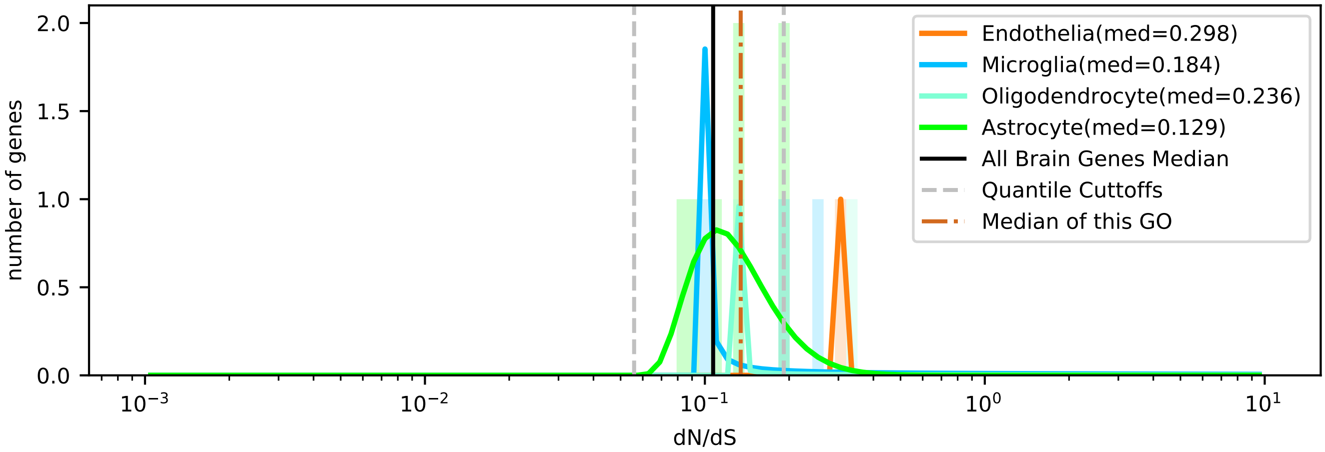

### Five Celltypes Distribution of Average dN/dS Scores of Genes Related to signal transduction

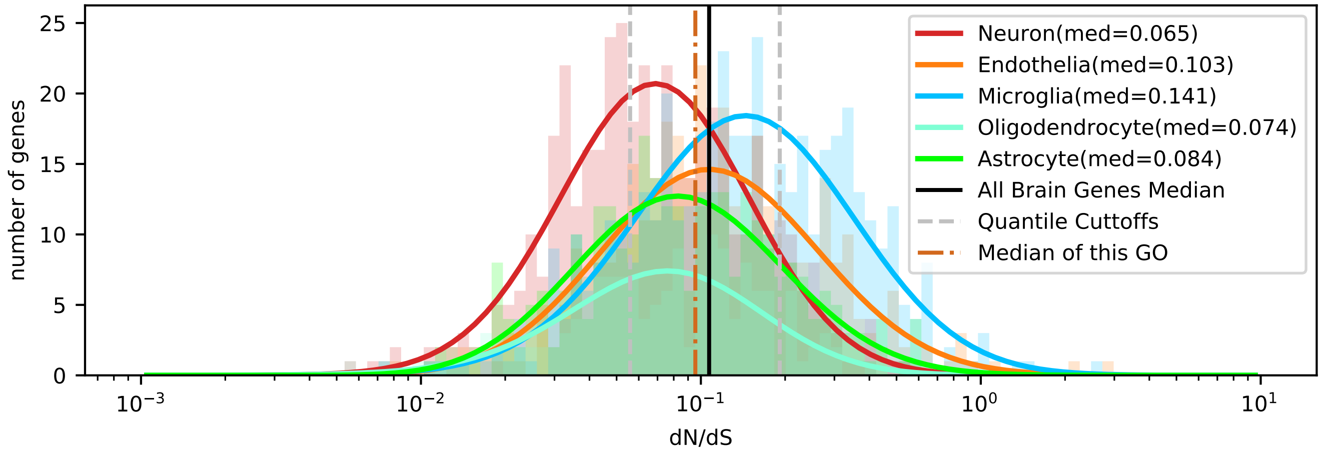


| | < cutoff | other genes | Total |
|---|---|---|---|
| In Astrocyte | 89 (82.47) | 200 (206.53) | 289 |
| Not IN Astrocyte | 391 (397.53) | 1002 (995.47) | 1393 |
| Total | 480 | 1202 | 1682 |

chi2: 0.744, p: 0.3883403270, OR: 1.14038

| | > cutoff | other genes | Total |
|---|---|---|---|
| In Astrocyte | 49 (63.06) | 240 (225.94) | 289 |
| Not IN Astrocyte | 318 (303.94) | 1075 (1089.06) | 1393 |
| Total | 367 | 1315 | 1682 |

chi2: 4.502, p: 0.0338548671, OR: 0.69019

| | < cutoff | other genes | Total |
|---|---|---|---|
| In Microglia | 73 (127.85) | 375 (320.15) | 448 |
| Not IN Microglia | 407 (352.15) | 827 (881.85) | 1234 |
| Total | 480 | 1202 | 1682 |

chi2: 44.066, p: 0.0000000000, OR: 0.39555

| | > cutoff | other genes | Total |
|---|---|---|---|
| In Microglia | 170 (97.75) | 278 (350.25) | 448 |
| Not IN Microglia | 197 (269.25) | 1037 (964.75) | 1234 |
| Total | 367 | 1315 | 1682 |

chi2: 91.819, p: 0.0000000000, OR: 3.21897

| | < cutoff | other genes | Total |
|---|---|---|---|
| In Oligodendrocyte | 61 (45.66) | 99 (114.34) | 160 |
| Not IN Oligodendrocyte | 419 (434.34) | 1103 (1087.66) | 1522 |
| Total | 480 | 1202 | 1682 |

chi2: 7.459, p: 0.0063126683, OR: 1.62202

| | > cutoff | other genes | Total |
|---|---|---|---|
| In Oligodendrocyte | 16 (34.91) | 144 (125.09) | 160 |
| Not IN Oligodendrocyte | 351 (332.09) | 1171 (1189.91) | 1522 |
| Total | 367 | 1315 | 1682 |

chi2: 13.725, p: 0.0002116706, OR: 0.37069

| | < cutoff | other genes | Total |
|---|---|---|---|
| In Neuron | 185 (124.71) | 252 (312.29) | 437 |
| Not IN Neuron | 295 (355.29) | 950 (889.71) | 1245 |
| Total | 480 | 1202 | 1682 |

chi2: 54.195, p: 0.0000000000, OR: 2.36414

| | > cutoff | other genes | Total |
|---|---|---|---|
| In Neuron | 45 (95.35) | 392 (341.65) | 437 |
| Not IN Neuron | 322 (271.65) | 923 (973.35) | 1245 |
| Total | 367 | 1315 | 1682 |

chi2: 45.037, p: 0.0000000000, OR: 0.32906

| | < cutoff | other genes | Total |
|---|---|---|---|
| In Endothelia | 72 (99.31) | 276 (248.69) | 348 |
| Not IN Endothelia | 408 (380.69) | 926 (953.31) | 1334 |
| Total | 480 | 1202 | 1682 |

chi2: 12.770, p: 0.0003521614, OR: 0.59207

| | > cutoff | other genes | Total |
|---|---|---|---|
| In Endothelia | 87 (75.93) | 261 (272.07) | 348 |
| Not IN Endothelia | 280 (291.07) | 1054 (1042.93) | 1334 |
| Total | 367 | 1315 | 1682 |

chi2: 2.373, p: 0.1234849043, OR: 1.25476

### Five Celltypes Distribution of Average dN/dS Scores of Genes Related to small molecule metabolic process

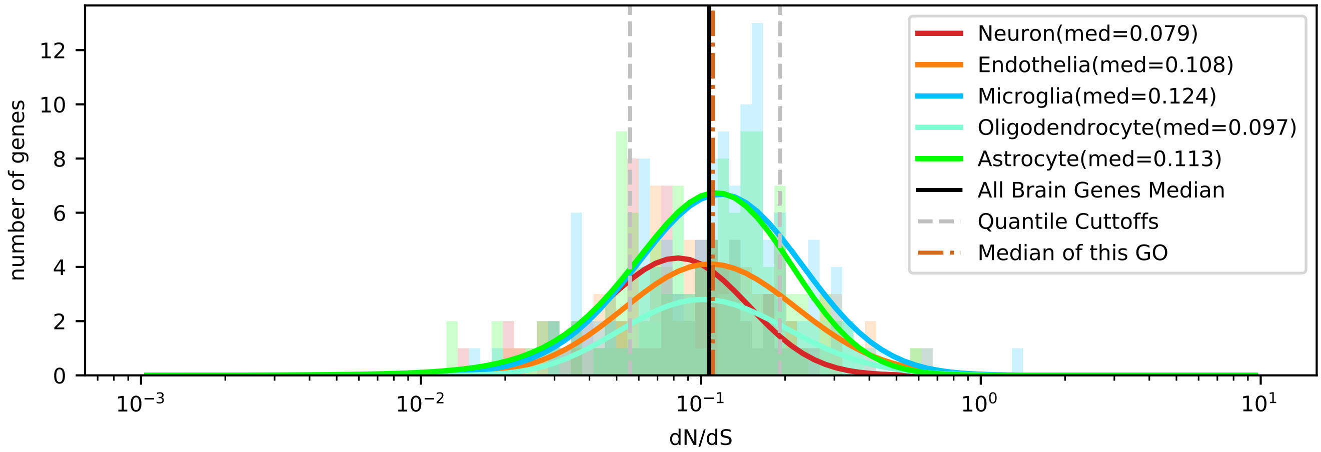


| | < cutoff | other genes | Total |
|---|---|---|---|
| In Astrocyte | 25 (23.11) | 96 (97.89) | 121 |
| Not IN Astrocyte | 60 (61.89) | 264 (262.11) | 324 |
| Total | 85 | 360 | 445 |

chi2: 0.141, p: 0.7068504532, OR: 1.14583

| | > cutoff | other genes | Total |
|---|---|---|---|
| In Astrocyte | 20 (19.58) | 101 (101.42) | 121 |
| Not IN Astrocyte | 52 (52.42) | 272 (271.58) | 324 |
| Total | 72 | 373 | 445 |

chi2: 0.001, p: 0.9821056101, OR: 1.03580

| | < cutoff | other genes | Total |
|---|---|---|---|
| In Microglia | 19 (23.69) | 105 (100.31) | 124 |
| Not IN Microglia | 66 (61.31) | 255 (259.69) | 321 |
| Total | 85 | 360 | 445 |

chi2: 1.267, p: 0.2602600939, OR: 0.69913

| | > cutoff | other genes | Total |
|---|---|---|---|
| In Microglia | 21 (20.06) | 103 (103.94) | 124 |
| Not IN Microglia | 51 (51.94) | 270 (269.06) | 321 |
| Total | 72 | 373 | 445 |

chi2: 0.016, p: 0.9001341616, OR: 1.07938

| | < cutoff | other genes | Total |
|---|---|---|---|
| In Oligodendrocyte | 7 (9.74) | 44 (41.26) | 51 |
| Not IN Oligodendrocyte | 78 (75.26) | 316 (318.74) | 394 |
| Total | 85 | 360 | 445 |

chi2: 0.720, p: 0.3961082899, OR: 0.64452

| | > cutoff | other genes | Total |
|---|---|---|---|
| In Oligodendrocyte | 10 (8.25) | 41 (42.75) | 51 |
| Not IN Oligodendrocyte | 62 (63.75) | 332 (330.25) | 394 |
| Total | 72 | 373 | 445 |

chi2: 0.254, p: 0.6139516533, OR: 1.30606

| | < cutoff | other genes | Total |
|---|---|---|---|
| In Neuron | 22 (13.56) | 49 (57.44) | 71 |
| Not IN Neuron | 63 (71.44) | 311 (302.56) | 374 |
| Total | 85 | 360 | 445 |

chi2: 6.834, p: 0.0089440326, OR: 2.21639

| | > cutoff | other genes | Total |
|---|---|---|---|
| In Neuron | 5 (11.49) | 66 (59.51) | 71 |
| Not IN Neuron | 67 (60.51) | 307 (313.49) | 374 |
| Total | 72 | 373 | 445 |

chi2: 4.430, p: 0.0353090354, OR: 0.34713

| | < cutoff | other genes | Total |
|---|---|---|---|
| In Endothelia | 12 (14.90) | 66 (63.10) | 78 |
| Not IN Endothelia | 73 (70.10) | 294 (296.90) | 367 |
| Total | 85 | 360 | 445 |

chi2: 0.579, p: 0.4467385056, OR: 0.73225

| | > cutoff | other genes | Total |
|---|---|---|---|
| In Endothelia | 16 (12.62) | 62 (65.38) | 78 |
| Not IN Endothelia | 56 (59.38) | 311 (307.62) | 367 |
| Total | 72 | 373 | 445 |

chi2: 0.951, p: 0.3295680522, OR: 1.43318

### Five Celltypes Distribution of Average dN/dS Scores of Genes Related to sulfur compound metabolic process

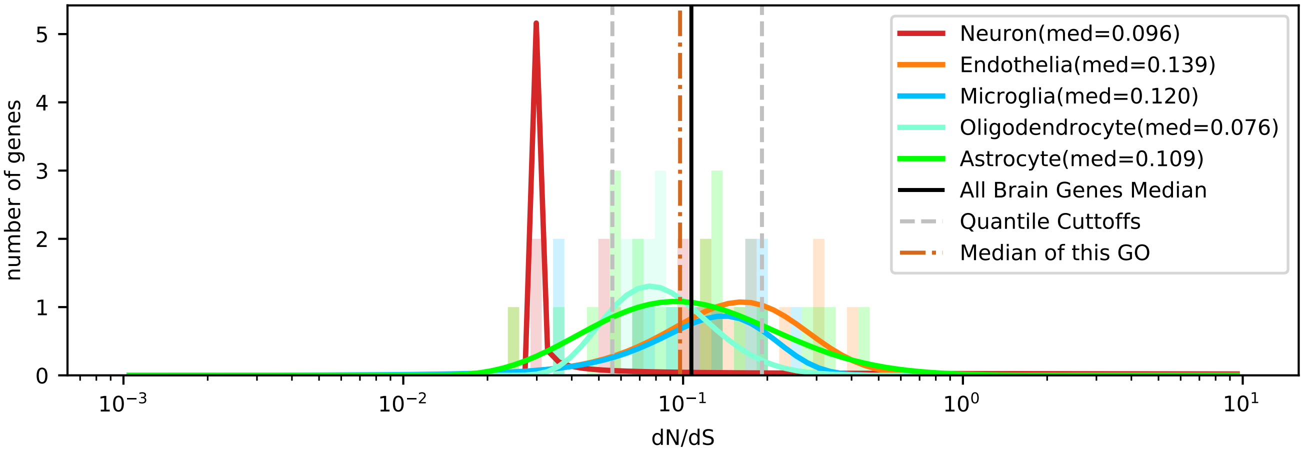


| | < cutoff | other genes | Total |
|---|---|---|---|
| In Astrocyte | 4 (3.52) | 18 (18.48) | 22 |
| Not IN Astrocyte | 8 (8.48) | 45 (44.52) | 53 |
| Total | 12 | 63 | 75 |

chi2: 0.000, p: 0.9889607918, OR: 1.25000

| | > cutoff | other genes | Total |
|---|---|---|---|
| In Astrocyte | 4 (3.23) | 18 (18.77) | 22 |
| Not IN Astrocyte | 7 (7.77) | 46 (45.23) | 53 |
| Total | 11 | 64 | 75 |

chi2: 0.038, p: 0.8446480230, OR: 1.46032

| | < cutoff | other genes | Total |
|---|---|---|---|
| In Microglia | 2 (1.92) | 10 (10.08) | 12 |
| Not IN Microglia | 10 (10.08) | 53 (52.92) | 63 |
| Total | 12 | 63 | 75 |

chi2: 0.130, p: 0.7182161295, OR: 1.06000

| | > cutoff | other genes | Total |
|---|---|---|---|
| In Microglia | 2 (1.76) | 10 (10.24) | 12 |
| Not IN Microglia | 9 (9.24) | 54 (53.76) | 63 |
| Total | 11 | 64 | 75 |

chi2: 0.054, p: 0.8169400407, OR: 1.20000

| | < cutoff | other genes | Total |
|---|---|---|---|
| In Oligodendrocyte | 1 (2.40) | 14 (12.60) | 15 |
| Not IN Oligodendrocyte | 11 (9.60) | 49 (50.40) | 60 |
| Total | 12 | 63 | 75 |

chi2: 0.502, p: 0.4785209765, OR: 0.31818

| | > cutoff | other genes | Total |
|---|---|---|---|
| In Oligodendrocyte | 0 (2.20) | 15 (12.80) | 15 |
| Not IN Oligodendrocyte | 11 (8.80) | 49 (51.20) | 60 |
| Total | 11 | 64 | 75 |

chi2: 1.924, p: 0.1653864798, OR: 0.00000

| | < cutoff | other genes | Total |
|---|---|---|---|
| In Neuron | 4 (1.44) | 5 (7.56) | 9 |
| Not IN Neuron | 8 (10.56) | 58 (55.44) | 66 |
| Total | 12 | 63 | 75 |

chi2: 3.987, p: 0.0458616916, OR: 5.80000

| | > cutoff | other genes | Total |
|---|---|---|---|
| In Neuron | 0 (1.32) | 9 (7.68) | 9 |
| Not IN Neuron | 11 (9.68) | 55 (56.32) | 66 |
| Total | 11 | 64 | 75 |

chi2: 0.678, p: 0.4101562506, OR: 0.00000

| | < cutoff | other genes | Total |
|---|---|---|---|
| In Endothelia | 1 (2.72) | 16 (14.28) | 17 |
| Not IN Endothelia | 11 (9.28) | 47 (48.72) | 58 |
| Total | 12 | 63 | 75 |

chi2: 0.842, p: 0.3587188049, OR: 0.26705

| | > cutoff | other genes | Total |
|---|---|---|---|
| In Endothelia | 5 (2.49) | 12 (14.51) | 17 |
| Not IN Endothelia | 6 (8.51) | 52 (49.49) | 58 |
| Total | 11 | 64 | 75 |

chi2: 2.447, p: 0.1177282790, OR: 3.61111

### Five Celltypes Distribution of Average dN/dS Scores of Genes Related to symbiont process

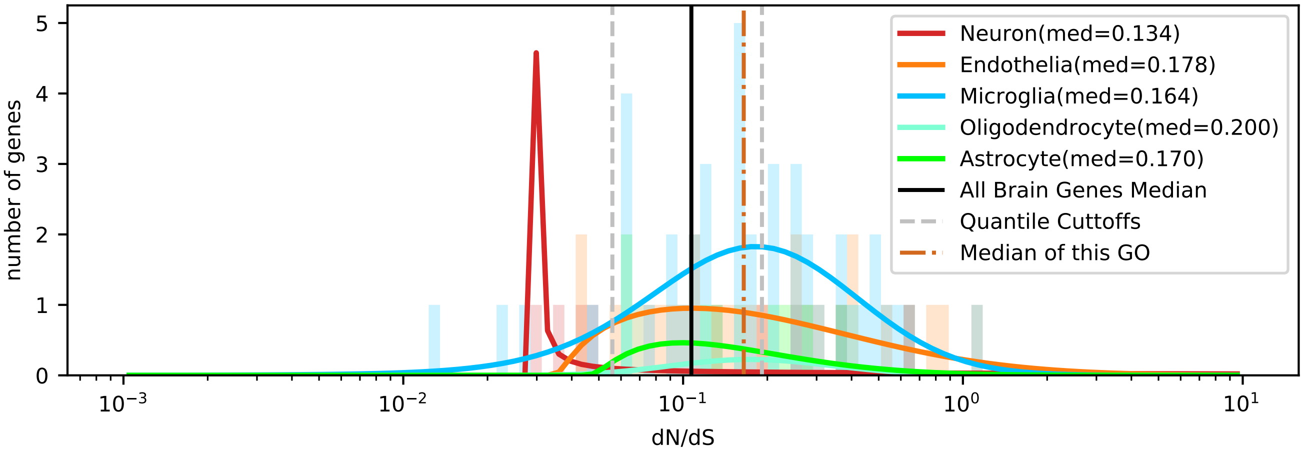


| | < cutoff | other genes | Total |
|---|---|---|---|
| In Astrocyte | 0 (0.90) | 8 (7.10) | 8 |
| Not IN Astrocyte | 10 (9.10) | 71 (71.90) | 81 |
| Total | 10 | 79 | 89 |

chi2: 0.219, p: 0.6397256680, OR: 0.00000

| | > cutoff | other genes | Total |
|---|---|---|---|
| In Astrocyte | 4 (3.60) | 4 (4.40) | 8 |
| Not IN Astrocyte | 36 (36.40) | 45 (44.60) | 81 |
| Total | 40 | 49 | 89 |

chi2: 0.005, p: 0.9432753124, OR: 1.25000

| | < cutoff | other genes | Total |
|---|---|---|---|
| In Microglia | 4 (4.83) | 39 (38.17) | 43 |
| Not IN Microglia | 6 (5.17) | 40 (40.83) | 46 |
| Total | 10 | 79 | 89 |

chi2: 0.050, p: 0.8238208349, OR: 0.68376

| | > cutoff | other genes | Total |
|---|---|---|---|
| In Microglia | 19 (19.33) | 24 (23.67) | 43 |
| Not IN Microglia | 21 (20.67) | 25 (25.33) | 46 |
| Total | 40 | 49 | 89 |

chi2: 0.006, p: 0.9407993442, OR: 0.94246

| | < cutoff | other genes | Total |
|---|---|---|---|
| In Oligodendrocyte | 0 (0.45) | 4 (3.55) | 4 |
| Not IN Oligodendrocyte | 10 (9.55) | 75 (75.45) | 85 |
| Total | 10 | 79 | 89 |

chi2: 0.007, p: 0.9347156445, OR: 0.00000

| | > cutoff | other genes | Total |
|---|---|---|---|
| In Oligodendrocyte | 2 (1.80) | 2 (2.20) | 4 |
| Not IN Oligodendrocyte | 38 (38.20) | 47 (46.80) | 85 |
| Total | 40 | 49 | 89 |

chi2: 0.094, p: 0.7594157627, OR: 1.23684

| | < cutoff | other genes | Total |
|---|---|---|---|
| In Neuron | 4 (1.01) | 5 (7.99) | 9 |
| Not IN Neuron | 6 (8.99) | 74 (71.01) | 80 |
| Total | 10 | 79 | 89 |

chi2: 7.677, p: 0.0055936555, OR: 9.86667

| | > cutoff | other genes | Total |
|---|---|---|---|
| In Neuron | 4 (4.04) | 5 (4.96) | 9 |
| Not IN Neuron | 36 (35.96) | 44 (44.04) | 80 |
| Total | 40 | 49 | 89 |

chi2: 0.103, p: 0.7477334672, OR: 0.97778

| | < cutoff | other genes | Total |
|---|---|---|---|
| In Endothelia | 2 (2.81) | 23 (22.19) | 25 |
| Not IN Endothelia | 8 (7.19) | 56 (56.81) | 64 |
| Total | 10 | 79 | 89 |

chi2: 0.053, p: 0.8175038876, OR: 0.60870

| | > cutoff | other genes | Total |
|---|---|---|---|
| In Endothelia | 11 (11.24) | 14 (13.76) | 25 |
| Not IN Endothelia | 29 (28.76) | 35 (35.24) | 64 |
| Total | 40 | 49 | 89 |

chi2: 0.016, p: 0.9003719473, OR: 0.94828

### Five Celltypes Distribution of Average dN/dS Scores of Genes Related to translation

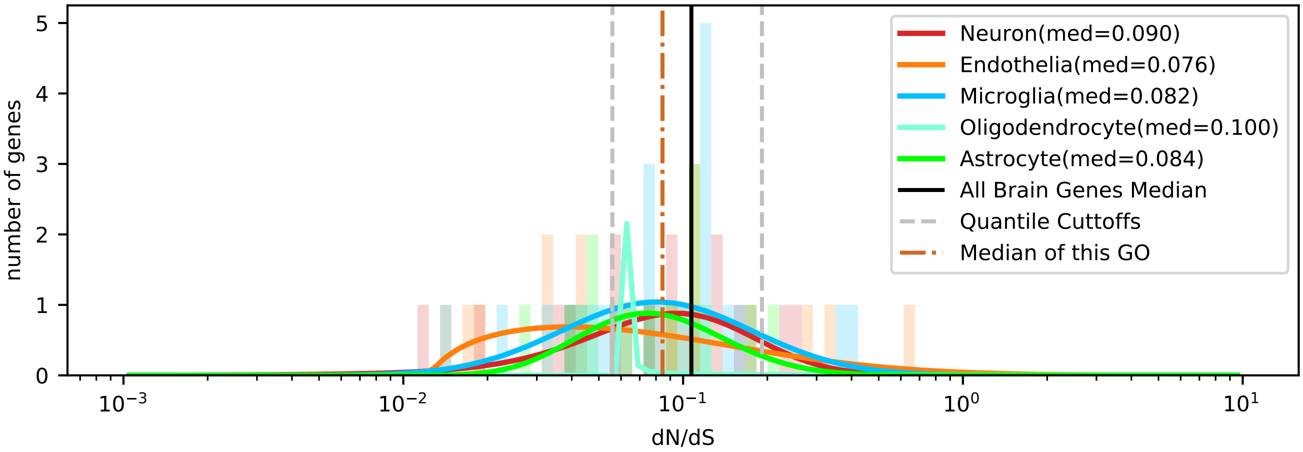


| | < cutoff | other genes | Total |
|---|---|---|---|
| In Astrocyte | 5 (4.67) | 9 (9.33) | 14 |
| Not IN Astrocyte | 20 (20.33) | 41 (40.67) | 61 |
| Total | 25 | 50 | 75 |

chi2: 0.011, p: 0.9165545330, OR: 1.13889

| | > cutoff | other genes | Total |
|---|---|---|---|
| In Astrocyte | 1 (1.49) | 13 (12.51) | 14 |
| Not IN Astrocyte | 7 (6.51) | 54 (54.49) | 61 |
| Total | 8 | 67 | 75 |

chi2: 0.000, p: 0.9948934639, OR: 0.59341

| | < cutoff | other genes | Total |
|---|---|---|---|
| In Microglia | 7 (7.33) | 15 (14.67) | 22 |
| Not IN Microglia | 18 (17.67) | 35 (35.33) | 53 |
| Total | 25 | 50 | 75 |

chi2: 0.008, p: 0.9285512296, OR: 0.90741

| | > cutoff | other genes | Total |
|---|---|---|---|
| In Microglia | 2 (2.35) | 20 (19.65) | 22 |
| Not IN Microglia | 6 (5.65) | 47 (47.35) | 53 |
| Total | 8 | 67 | 75 |

chi2: 0.016, p: 0.8997488588, OR: 0.78333

| | < cutoff | other genes | Total |
|---|---|---|---|
| In Oligodendrocyte | 0 (1.00) | 3 (2.00) | 3 |
| Not IN Oligodendrocyte | 25 (24.00) | 47 (48.00) | 72 |
| Total | 25 | 50 | 75 |

chi2: 0.391, p: 0.5319710581, OR: 0.00000

| | > cutoff | other genes | Total |
|---|---|---|---|
| In Oligodendrocyte | 0 (0.32) | 3 (2.68) | 3 |
| Not IN Oligodendrocyte | 8 (7.68) | 64 (64.32) | 72 |
| Total | 8 | 67 | 75 |

chi2: 0.118, p: 0.7311459497, OR: 0.00000

| | < cutoff | other genes | Total |
|---|---|---|---|
| In Neuron | 5 (5.67) | 12 (11.33) | 17 |
| Not IN Neuron | 20 (19.33) | 38 (38.67) | 58 |
| Total | 25 | 50 | 75 |

chi2: 0.010, p: 0.9223217455, OR: 0.79167

| | > cutoff | other genes | Total |
|---|---|---|---|
| In Neuron | 2 (1.81) | 15 (15.19) | 17 |
| Not IN Neuron | 6 (6.19) | 52 (51.81) | 58 |
| Total | 8 | 67 | 75 |

chi2: 0.078, p: 0.7795172655, OR: 1.15556

| | < cutoff | other genes | Total |
|---|---|---|---|
| In Endothelia | 8 (6.33) | 11 (12.67) | 19 |
| Not IN Endothelia | 17 (18.67) | 39 (37.33) | 56 |
| Total | 25 | 50 | 75 |

chi2: 0.432, p: 0.5111347158, OR: 1.66845

| | > cutoff | other genes | Total |
|---|---|---|---|
| In Endothelia | 3 (2.03) | 16 (16.97) | 19 |
| Not IN Endothelia | 5 (5.97) | 51 (50.03) | 56 |
| Total | 8 | 67 | 75 |

chi2: 0.166, p: 0.6839313259, OR: 1.91250

### Five Celltypes Distribution of Average dN/dS Scores of Genes Related to transmembrane transport

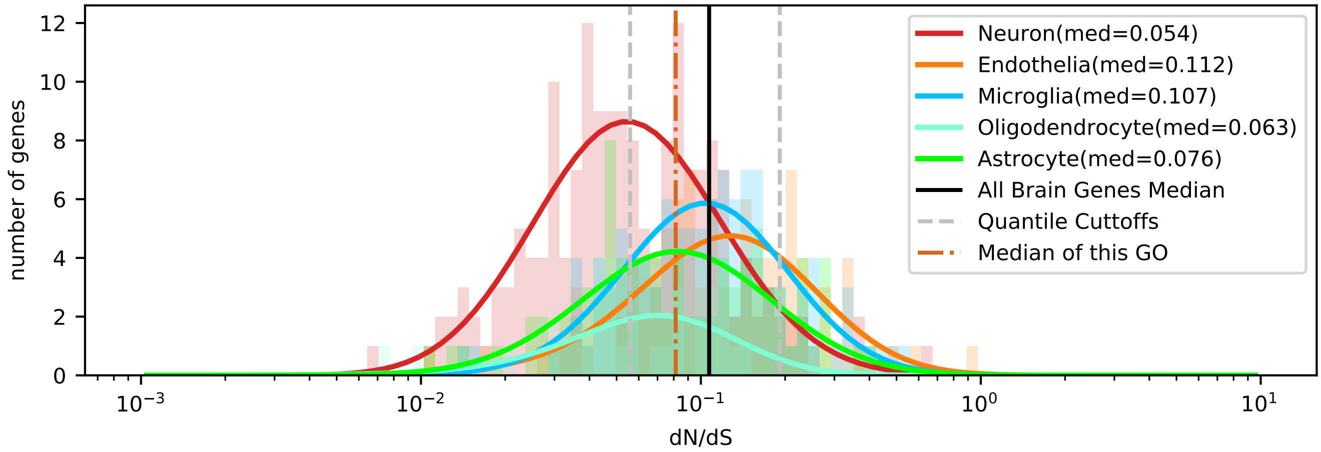


| | < cutoff | other genes | Total |
|---|---|---|---|
| In Astrocyte | 32 (31.19) | 57 (57.81) | 89 |
| Not IN Astrocyte | 146 (146.81) | 273 (272.19) | 419 |
| Total | 178 | 330 | 508 |

chi2: 0.006, p: 0.9385823279, OR: 1.04975

| | > cutoff | other genes | Total |
|---|---|---|---|
| In Astrocyte | 12 (11.04) | 77 (77.96) | 89 |
| Not IN Astrocyte | 51 (51.96) | 368 (367.04) | 419 |
| Total | 63 | 445 | 508 |

chi2: 0.027, p: 0.8698786367, OR: 1.12452

| | < cutoff | other genes | Total |
|---|---|---|---|
| In Microglia | 23 (37.84) | 85 (70.16) | 108 |
| Not IN Microglia | 155 (140.16) | 245 (259.84) | 400 |
| Total | 178 | 330 | 508 |

chi2: 10.627, p: 0.0011142820, OR: 0.42770

| | > cutoff | other genes | Total |
|---|---|---|---|
| In Microglia | 17 (13.39) | 91 (94.61) | 108 |
| Not IN Microglia | 46 (49.61) | 354 (350.39) | 400 |
| Total | 63 | 445 | 508 |

chi2: 1.044, p: 0.3067855721, OR: 1.43765

| | < cutoff | other genes | Total |
|---|---|---|---|
| In Oligodendrocyte | 17 (13.31) | 21 (24.69) | 38 |
| Not IN Oligodendrocyte | 161 (164.69) | 309 (305.31) | 470 |
| Total | 178 | 330 | 508 |

chi2: 1.268, p: 0.2602047151, OR: 1.55368

| | > cutoff | other genes | Total |
|---|---|---|---|
| In Oligodendrocyte | 1 (4.71) | 37 (33.29) | 38 |
| Not IN Oligodendrocyte | 62 (58.29) | 408 (411.71) | 470 |
| Total | 63 | 445 | 508 |

chi2: 2.702, p: 0.1002081155, OR: 0.17786

| | < cutoff | other genes | Total |
|---|---|---|---|
| In Neuron | 93 (62.37) | 85 (115.63) | 178 |
| Not IN Neuron | 85 (115.63) | 245 (214.37) | 330 |
| Total | 178 | 330 | 508 |

chi2: 34.492, p: 0.0000000043, OR: 3.15363

| | > cutoff | other genes | Total |
|---|---|---|---|
| In Neuron | 8 (22.07) | 170 (155.93) | 178 |
| Not IN Neuron | 55 (40.93) | 275 (289.07) | 330 |
| Total | 63 | 445 | 508 |

chi2: 14.670, p: 0.0001280832, OR: 0.23529

| | < cutoff | other genes | Total |
|---|---|---|---|
| In Endothelia | 13 (33.29) | 82 (61.71) | 95 |
| Not IN Endothelia | 165 (144.71) | 248 (268.29) | 413 |
| Total | 178 | 330 | 508 |

chi2: 22.272, p: 0.0000023662, OR: 0.23829

| | > cutoff | other genes | Total |
|---|---|---|---|
| In Endothelia | 25 (11.78) | 70 (83.22) | 95 |
| Not IN Endothelia | 38 (51.22) | 375 (361.78) | 413 |
| Total | 63 | 445 | 508 |

chi2: 19.279, p: 0.0000112930, OR: 3.52444

Five Celltypes Distribution of Average dN/dS Scores of Genes Related to transport

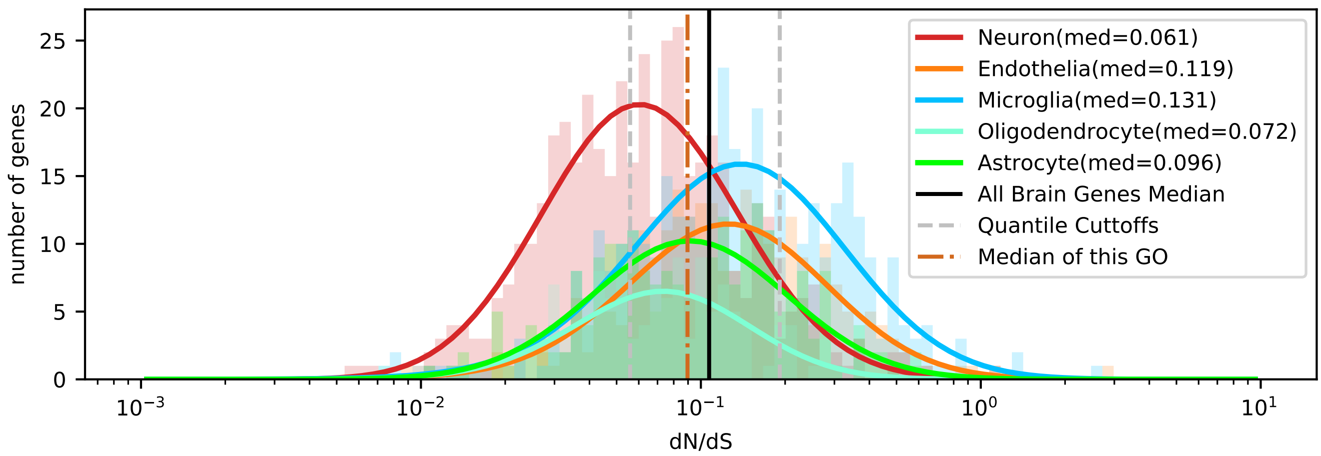


| | < cutoff | other genes | Total |
|---|---|---|---|
| In Astrocyte | 66 (66.38) | 156 (155.62) | 222 |
| Not IN Astrocyte | 358 (357.62) | 838 (838.38) | 1196 |
| Total | 424 | 994 | 1418 |

chi2: 0.000, p: 0.9848217924, OR: 0.99033

| | > cutoff | other genes | Total |
|---|---|---|---|
| In Astrocyte | 38 (43.84) | 184 (178.16) | 222 |
| Not IN Astrocyte | 242 (236.16) | 954 (959.84) | 1196 |
| Total | 280 | 1138 | 1418 |

chi2: 0.960, p: 0.3272603821, OR: 0.81414

| | < cutoff | other genes | Total |
|---|---|---|---|
| In Microglia | 66 (112.43) | 310 (263.57) | 376 |
| Not IN Microglia | 358 (311.57) | 684 (730.43) | 1042 |
| Total | 424 | 994 | 1418 |

chi2: 36.424, p: 0.0000000016, OR: 0.40678

| | > cutoff | other genes | Total |
|---|---|---|---|
| In Microglia | 122 (74.25) | 254 (301.75) | 376 |
| Not IN Microglia | 158 (205.75) | 884 (836.25) | 1042 |
| Total | 280 | 1138 | 1418 |

chi2: 50.999, p: 0.0000000000, OR: 2.68733

| | < cutoff | other genes | Total |
|---|---|---|---|
| In Oligodendrocyte | 51 (37.97) | 76 (89.03) | 127 |
| Not IN Oligodendrocyte | 373 (386.03) | 918 (904.97) | 1291 |
| Total | 424 | 994 | 1418 |

chi2: 6.473, p: 0.0109504142, OR: 1.65155

| | > cutoff | other genes | Total |
|---|---|---|---|
| In Oligodendrocyte | 14 (25.08) | 113 (101.92) | 127 |
| Not IN Oligodendrocyte | 266 (254.92) | 1025 (1036.08) | 1291 |
| Total | 280 | 1138 | 1418 |

chi2: 6.106, p: 0.0134708924, OR: 0.47741

| | < cutoff | other genes | Total |
|---|---|---|---|
| In Neuron | 202 (130.97) | 236 (307.03) | 438 |
| Not IN Neuron | 222 (293.03) | 758 (686.97) | 980 |
| Total | 424 | 994 | 1418 |

chi2: 78.407, p: 0.0000000000, OR: 2.92251

| | > cutoff | other genes | Total |
|---|---|---|---|
| In Neuron | 31 (86.49) | 407 (351.51) | 438 |
| Not IN Neuron | 249 (193.51) | 731 (786.49) | 980 |
| Total | 280 | 1138 | 1418 |

chi2: 63.032, p: 0.0000000000, OR: 0.22361

| | < cutoff | other genes | Total |
|---|---|---|---|
| In Endothelia | 39 (76.25) | 216 (178.75) | 255 |
| Not IN Endothelia | 385 (347.75) | 778 (815.25) | 1163 |
| Total | 424 | 994 | 1418 |

chi2: 30.806, p: 0.0000000285, OR: 0.36486

| | > cutoff | other genes | Total |
|---|---|---|---|
| In Endothelia | 75 (50.35) | 180 (204.65) | 255 |
| Not IN Endothelia | 205 (229.65) | 958 (933.35) | 1163 |
| Total | 280 | 1138 | 1418 |

chi2: 17.593, p: 0.0000273541, OR: 1.94715

Five Celltypes Distribution of Average dN/dS Scores of Genes Related to transposition

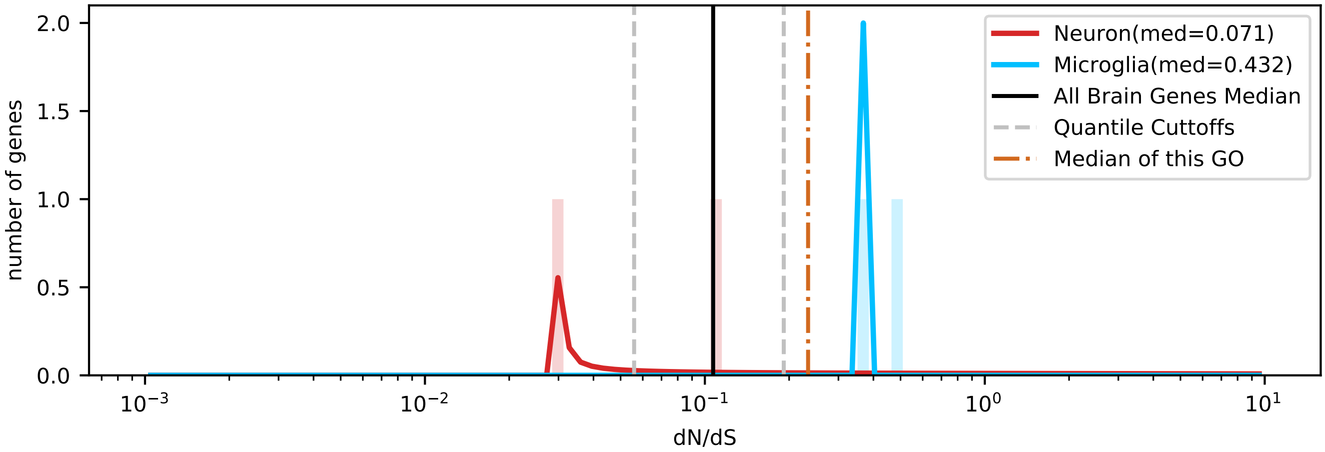

Five Celltypes Distribution of Average dN/dS Scores of Genes Related to tRNA metabolic process

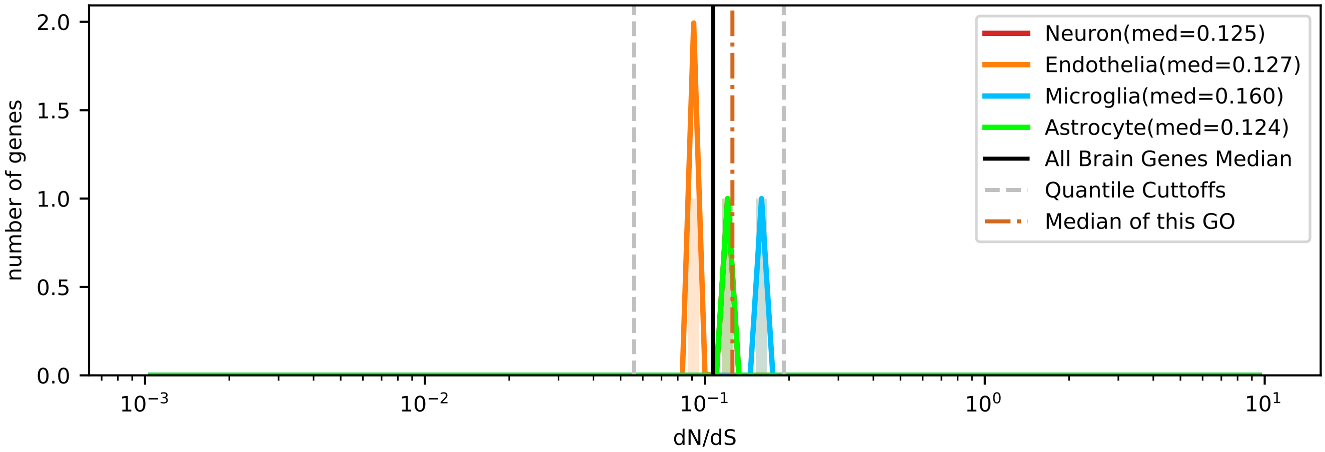

### Five Celltypes Distribution of Average dN/dS Scores of Genes Related to vacuolar transport

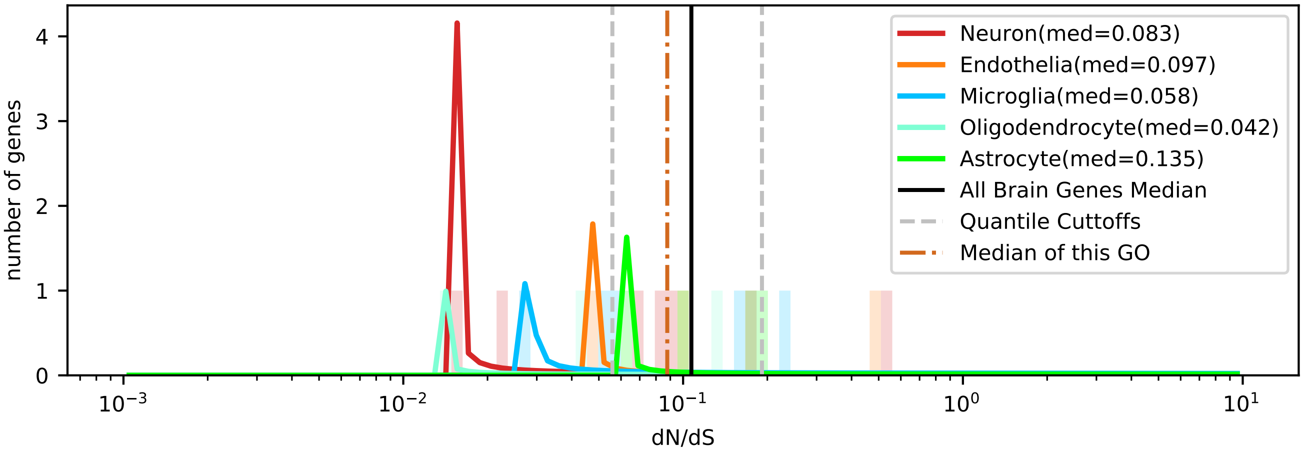


| | < cutoff | other genes | Total |
|---|---|---|---|
| In Astrocyte | 0 (1.27) | 4 (2.73) | 4 |
| Not IN Astrocyte | 7 (5.73) | 11 (12.27) | 18 |
| Total | 7 | 15 | 22 |

chi2: 0.841, p: 0.3591094153, OR: 0.00000

| | > cutoff | other genes | Total |
|---|---|---|---|
| In Astrocyte | 0 (0.55) | 4 (3.45) | 4 |
| Not IN Astrocyte | 3 (2.45) | 15 (15.55) | 18 |
| Total | 3 | 19 | 22 |

chi2: 0.005, p: 0.9416340109, OR: 0.00000

| | < cutoff | other genes | Total |
|---|---|---|---|
| In Microglia | 2 (1.59) | 3 (3.41) | 5 |
| Not IN Microglia | 5 (5.41) | 12 (11.59) | 17 |
| Total | 7 | 15 | 22 |

chi2: 0.010, p: 0.9209022613, OR: 1.60000

| | > cutoff | other genes | Total |
|---|---|---|---|
| In Microglia | 1 (0.68) | 4 (4.32) | 5 |
| Not IN Microglia | 2 (2.32) | 15 (14.68) | 17 |
| Total | 3 | 19 | 22 |

chi2: 0.073, p: 0.7875135841, OR: 1.87500

| | < cutoff | other genes | Total |
|---|---|---|---|
| In Oligodendrocyte | 2 (0.95) | 1 (2.05) | 3 |
| Not IN Oligodendrocyte | 5 (6.05) | 14 (12.95) | 19 |
| Total | 7 | 15 | 22 |

chi2: 0.529, p: 0.4668914937, OR: 5.60000

| | > cutoff | other genes | Total |
|---|---|---|---|
| In Oligodendrocyte | 0 (0.41) | 3 (2.59) | 3 |
| Not IN Oligodendrocyte | 3 (2.59) | 16 (16.41) | 19 |
| Total | 3 | 19 | 22 |

chi2: 0.027, p: 0.8692777295, OR: 0.00000

| | < cutoff | other genes | Total |
|---|---|---|---|
| In Neuron | 2 (2.23) | 5 (4.77) | 7 |
| Not IN Neuron | 5 (4.77) | 10 (10.23) | 15 |
| Total | 7 | 15 | 22 |

chi2: 0.072, p: 0.7886810316, OR: 0.80000

| | > cutoff | other genes | Total |
|---|---|---|---|
| In Neuron | 1 (0.95) | 6 (6.05) | 7 |
| Not IN Neuron | 2 (2.05) | 13 (12.95) | 15 |
| Total | 3 | 19 | 22 |

chi2: 0.368, p: 0.5443232564, OR: 1.08333

| | < cutoff | other genes | Total |
|---|---|---|---|
| In Endothelia | 1 (0.95) | 2 (2.05) | 3 |
| Not IN Endothelia | 6 (6.05) | 13 (12.95) | 19 |
| Total | 7 | 15 | 22 |

chi2: 0.368, p: 0.5443232564, OR: 1.08333

| | > cutoff | other genes | Total |
|---|---|---|---|
| In Endothelia | 1 (0.41) | 2 (2.59) | 3 |
| Not IN Endothelia | 2 (2.59) | 17 (16.41) | 19 |
| Total | 3 | 19 | 22 |

chi2: 0.027, p: 0.8692777295, OR: 4.25000

### Five Celltypes Distribution of Average dN/dS Scores of Genes Related to vesicle-mediated transport

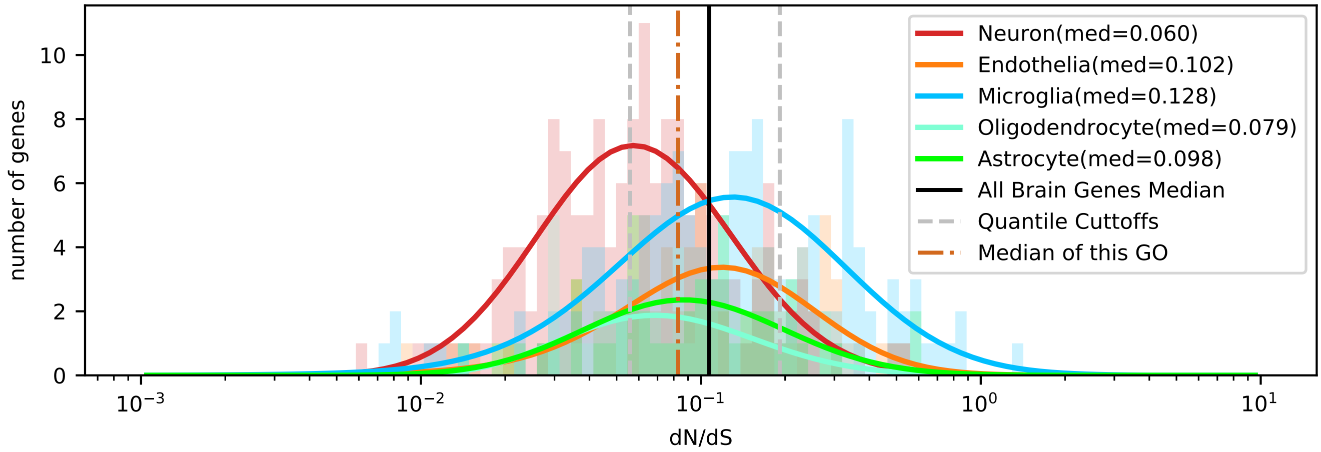


| | < cutoff | other genes | Total |
|---|---|---|---|
| In Astrocyte | 14 (16.10) | 38 (35.90) | 52 |
| Not IN Astrocyte | 129 (126.90) | 281 (283.10) | 410 |
| Total | 143 | 319 | 462 |

chi2: 0.258, p: 0.6114797543, OR: 0.80253

| | > cutoff | other genes | Total |
|---|---|---|---|
| In Astrocyte | 8 (10.35) | 44 (41.65) | 52 |
| Not IN Astrocyte | 84 (81.65) | 326 (328.35) | 410 |
| Total | 92 | 370 | 462 |

chi2: 0.468, p: 0.4941170410, OR: 0.70563

| | < cutoff | other genes | Total |
|---|---|---|---|
| In Microglia | 28 (44.57) | 116 (99.43) | 144 |
| Not IN Microglia | 115 (98.43) | 203 (219.57) | 318 |
| Total | 143 | 319 | 462 |

chi2: 12.193, p: 0.0004796341, OR: 0.42609

| | > cutoff | other genes | Total |
|---|---|---|---|
| In Microglia | 45 (28.68) | 99 (115.32) | 144 |
| Not IN Microglia | 47 (63.32) | 271 (254.68) | 318 |
| Total | 92 | 370 | 462 |

chi2: 15.842, p: 0.0000688486, OR: 2.62089

| | < cutoff | other genes | Total |
|---|---|---|---|
| In Oligodendrocyte | 15 (12.07) | 24 (26.93) | 39 |
| Not IN Oligodendrocyte | 128 (130.93) | 295 (292.07) | 423 |
| Total | 143 | 319 | 462 |

chi2: 0.773, p: 0.3793363938, OR: 1.44043

| | > cutoff | other genes | Total |
|---|---|---|---|
| In Oligodendrocyte | 7 (7.77) | 32 (31.23) | 39 |
| Not IN Oligodendrocyte | 85 (84.23) | 338 (338.77) | 423 |
| Total | 92 | 370 | 462 |

chi2: 0.012, p: 0.9111681482, OR: 0.86985

| | < cutoff | other genes | Total |
|---|---|---|---|
| In Neuron | 72 (47.67) | 82 (106.33) | 154 |
| Not IN Neuron | 71 (95.33) | 237 (212.67) | 308 |
| Total | 143 | 319 | 462 |

chi2: 25.888, p: 0.0000003618, OR: 2.93095

| | > cutoff | other genes | Total |
|---|---|---|---|
| In Neuron | 11 (30.67) | 143 (123.33) | 154 |
| Not IN Neuron | 81 (61.33) | 227 (246.67) | 308 |
| Total | 92 | 370 | 462 |

chi2: 22.437, p: 0.0000021719, OR: 0.21557

| | < cutoff | other genes | Total |
|---|---|---|---|
| In Endothelia | 14 (22.60) | 59 (50.40) | 73 |
| Not IN Endothelia | 129 (120.40) | 260 (268.60) | 389 |
| Total | 143 | 319 | 462 |

chi2: 4.989, p: 0.0255136048, OR: 0.47826

| | > cutoff | other genes | Total |
|---|---|---|---|
| In Endothelia | 21 (14.54) | 52 (58.46) | 73 |
| Not IN Endothelia | 71 (77.46) | 318 (311.54) | 389 |
| Total | 92 | 370 | 462 |

chi2: 3.628, p: 0.0568276342, OR: 1.80878
