## Supplementary Table S1 for "Neuron-specific coding and regulatory sequences are the most highly conserved in amniote brains despite neuron-specific cell size diversity"

### Zhang et al., 2014, three cell types

| Sample Size and Median dN/dS (CI95%) | Chicken Reference Genome | Human Reference Genome | Rat Reference Genome | Mouse Reference Genome |
|---|---|---|---|---|
| Endothelia | n=17,188<br>med:0.1204<br>(0.1184-0.1220) | n=61,852<br>med:0.1423<br>(0.1411-0.1436) | n=58,807<br>med:0.1182<br>(0.1171-0.1191) | n=63,603<br>med:0.1167<br>(0.1157-0.1178) |
| Glia | n=20,374<br>med:0.1160<br>(0.1142-0.1177) | n=80,441<br>med:0.1353<br>(0.1344-0.1364) | n=77,366<br>med:0.1196<br>(0.1189-0.1204) | n=81,921<br>med:0.1185<br>(0.1177-0.1194) |
| Neuron | n=24,928<br>med:0.0828<br>(0.0816-0.0841) | n=88,810<br>med:0.0809<br>(0.0802-0.0816) | n=84,911<br>med:0.0696<br>(0.0690-0.0702) | n=90,204<br>med:0.0672<br>(0.0667-0.0678) |

### Zhang et al., 2014, cell subtypes

| Sample Size and Median dN/dS (CI95%) | Chicken Reference Genome | Human Reference Genome | Rat Reference Genome | Mouse Reference Genome |
|---|---|---|---|---|
| Microglia | n=16,467<br>med:0.1276<br>(0.1254-0.1297) | n=68,785<br>med:0.1472<br>(0.1459-0.1485) | n=66,058<br>med:0.1312<br>(0.1302-0.1321) | n=69,722<br>med:0.1303<br>(0.1293-0.1313) |
| Endothelia | n=17,188<br>med:0.1204<br>(0.1184-0.1220) | n=61,852<br>med:0.1423<br>(0.1411-0.1436) | n=58,807<br>med:0.1182<br>(0.1171-0.1191) | n=63,603<br>med:0.1167<br>(0.1157-0.1178) |
| Astrocyte | n=16,384<br>med:0.1080<br>(0.1060-0.1099) | n=56,982<br>med:0.1227<br>(0.1215-0.1238) | n=54,685<br>med:0.1047<br>(0.1038-0.1057) | n=57,614<br>med:0.1024<br>(0.1015-0.1035) |
| Oligodendrocyte | n=10,476<br>med:0.0977<br>(0.0960-0.0994) | n=36,410<br>med:0.1009<br>(0.0996-0.1020) | n=35,179<br>med:0.0869<br>(0.0860-0.0878) | n=36,842<br>med:0.0860<br>(0.0850-0.0870) |
| Neuron | n=24,928<br>med:0.0828<br>(0.0816-0.0841) | n=88,810<br>med:0.0809<br>(0.0802-0.0816) | n=84,911<br>med:0.0696<br>(0.0690-0.0702) | n=90,204<br>med:0.0672<br>(0.0667-0.0678) |

### Zeisel et al., 2015, three cell types

| Sample Size and Median dN/dS (CI95%) | Chicken Reference Genome | Human Reference Genome | Rat Reference Genome | Mouse Reference Genome |
|---|---|---|---|---|
| Glia | n=24,816 med:0.1301 (0.1283-0.1318) | n=92,400 med:0.1457 (0.1447-0.1467) | n=89,176 med:0.1230 (0.1220-0.1238) | n=94,284 med:0.1232 (0.1223-0.1240) |
| Vasculature | n=8,878 med:0.1123 (0.1100-0.1150) | n=31,290 med:0.1223 (0.1204-0.1238) | n=29,545 med:0.1075 (0.1061-0.1086) | n=31,601 med:0.1038 (0.1026-0.1051) |
| Neuron | n=17,829 med:0.0858 (0.0842-0.0871) | n=62,585 med:0.0867 (0.0858-0.0876) | n=59,648 med:0.0750 (0.0742-0.0759) | n=62,864 med:0.0720 (0.0712-0.0727) |

### Zeisel et al., 2015, cell subtypes

| Sample Size and Median dN/dS (CI95%) | Chicken Reference Genome | Human Reference Genome | Rat Reference Genome | Mouse Reference Genome |
|---|---|---|---|---|
| Ependymal | n=6,539<br>med:0.1735<br>(0.1688-0.1780) | n=26,981<br>med:0.1863<br>(0.1838-0.1886) | n=25,840<br>med:0.1462<br>(0.1443-0.1480) | n=27,458<br>med:0.1486<br>(0.1468-0.1503) |
| Microglia | n=6,227<br>med:0.1431<br>(0.1397-0.1467) | n=23,791<br>med:0.1622<br>(0.1602-0.1641) | n=23,327<br>med:0.1427<br>(0.1408-0.1446) | n=24,642<br>med:0.1402<br>(0.1385-0.1420) |
| Endothelial | n=6,174<br>med:0.1113<br>(0.1085-0.1145) | n=21,835<br>med:0.1287<br>(0.1263-0.1306) | n=20,494<br>med:0.1110<br>(0.1093-0.1128) | n=21,937<br>med:0.1093<br>(0.1077-0.1111) |
| Mural | n=2,704<br>med:0.1144<br>(0.1109-0.1180) | n=9,455<br>med:0.1112<br>(0.1090-0.1139) | n=9,051<br>med:0.1005<br>(0.0982-0.1025) | n=9,664<br>med:0.0948<br>(0.0925-0.0964) |
| Oligodendrocyte | n=7,919<br>med:0.1088<br>(0.1062-0.1119) | n=26,957<br>med:0.1148<br>(0.1131-0.1163) | n=25,907<br>med:0.0998<br>(0.0986-0.1011) | n=27,528<br>med:0.1020<br>(0.1008-0.1033) |
| Astrocyte | n=4,131<br>med:0.1018<br>(0.0978-0.1047) | n=14,671<br>med:0.1217<br>(0.1196-0.1234) | n=14,102<br>med:0.1045<br>(0.1029-0.1059) | n=14,656<br>med:0.0998<br>(0.0983-0.1013) |
| Interneuron | n=6,213<br>med:0.0901<br>(0.0874-0.0926) | n=21,961<br>med:0.0923<br>(0.0907-0.0942) | n=20,630<br>med:0.0793<br>(0.0779-0.0806) | n=22,103<br>med:0.0750<br>(0.0737-0.0763) |
| S1 Pyramidal | n=4,665 | n=16,450 | n=16,256 | n=16,679 |

| | | | | |
|---|---|---|---|---|
| CA1 Pyramidal | med:0.0859<br>(0.0832-0.0887) | med:0.0867<br>(0.0850-0.0887) | med:0.0791<br>(0.0772-0.0807) | med:0.0740<br>(0.0727-0.0756) |
| | n=6,951<br>med:0.0815<br>(0.0790-0.0839) | n=24,174<br>med:0.0811<br>(0.0797-0.0825) | n=22,762<br>med:0.0685<br>(0.0672-0.0697) | n=24,082<br>med:0.0680<br>(0.0667-0.0692) |

### Zeisel et al., 2018, three cell types

| Sample Size and Median dN/dS (CI95%) | Chicken Reference Genome | Human Reference Genome | Rat Reference Genome | Mouse Reference Genome |
|---|---|---|---|---|
| CNS_Glia | n=16,906<br>med:0.1408<br>(0.1380-0.1435) | n=65,079<br>med:0.1519<br>(0.1506-0.1532) | n=62,523<br>med:0.1254<br>(0.1243-0.1264) | n=66,102<br>med:0.1263<br>(0.1253-0.1274) |
| CNS_Endothelia | n=2,983<br>med:0.1227<br>(0.1191-0.1267) | n=12,168<br>med:0.1451<br>(0.1423-0.1483) | n=11,206<br>med:0.1222<br>(0.1198-0.1239) | n=12,281<br>med:0.1220<br>(0.1201-0.1242) |
| CNS_Neuron | n=75,911<br>med:0.0887<br>(0.0880-0.0895) | n=252,530<br>med:0.0885<br>(0.0881-0.0890) | n=243,152<br>med:0.0760<br>(0.0756-0.0763) | n=256,590<br>med:0.0746<br>(0.0743-0.0750) |

### Zeisel et al., 2018, cell subtypes

| Sample Size and Median dN/dS (CI95%) | Chicken Reference Genome | Human Reference Genome | Rat Reference Genome | Mouse Reference Genome |
|---|---|---|---|---|
| Glia | n=13,707 med:0.1636 (0.1605-0.1668) | n=53,560 med:0.1721 (0.1703-0.1739) | n=51,344 med:0.1429 (0.1418-0.1441) | n=54,589 med:0.1464 (0.1453-0.1476) |
| Immune cells | n=1,560 med:0.1618 (0.1545-0.1685) | n=7,257 med:0.1761 (0.1708-0.1806) | n=7,184 med:0.1506 (0.1465-0.1544) | n=7,461 med:0.1524 (0.1493-0.1554) |
| Vascular cells | n=3,305 med:0.1108 (0.1074-0.1148) | n=13,924 med:0.1325 (0.1302-0.1346) | n=12,792 med:0.1131 (0.1113-0.1150) | n=13,849 med:0.1149 (0.1127-0.1168) |
| Neurons | n=77,966 med:0.0964 (0.0955-0.0972) | n=265,605 med:0.0990 (0.0985-0.0995) | n=255,785 med:0.0848 (0.0844-0.0852) | n=270,578 med:0.0835 (0.0832-0.0839) |

### Saunders et al., 2018, three cell types

| Sample Size and Median dN/dS (CI95%) | Chicken Reference Genome | Human Reference Genome | Rat Reference Genome | Mouse Reference Genome |
|---|---|---|---|---|
| Vasculature | n=8,307 med:0.1122 (0.1090-0.1157) | n=29,546 med:0.1279 (0.1258-0.1299) | n=27,654 med:0.1050 (0.1038-0.1063) | n=30,009 med:0.1042 (0.1029-0.1054) |
| Glia | n=6,266 med:0.0973 (0.0951-0.1001) | n=21,209 med:0.0963 (0.0948-0.0979) | n=20,234 med:0.0831 (0.0820-0.0841) | n=21,166 med:0.0821 (0.0809-0.0831) |
| Neuron | n=14,779 med:0.0715 (0.0698-0.0732) | n=48,106 med:0.0599 (0.0591-0.0608) | n=46,085 med:0.0516 (0.0510-0.0522) | n=48,656 med:0.0494 (0.0488-0.0500) |

### Saunders et al., 2018, cell subtypes

| Sample Size and Median dN/dS (CI95%) | Chicken Reference Genome | Human Reference Genome | Rat Reference Genome | Mouse Reference Genome |
|---|---|---|---|---|
| Endothelia | n=7,607<br>med:0.1157<br>(0.1122-0.1182) | n=26,101<br>med:0.1244<br>(0.1224-0.1263) | n=24,613<br>med:0.1053<br>(0.1039-0.1067) | n=26,274<br>med:0.1042<br>(0.1029-0.1055) |
| Microglia | n=5,125<br>med:0.1169<br>(0.1125-0.1205) | n=18,798<br>med:0.1388<br>(0.1369-0.1410) | n=17,973<br>med:0.1249<br>(0.1227-0.1271) | n=18,986<br>med:0.1212<br>(0.1197-0.1230) |
| Astrocyte | n=6,136<br>med:0.0971<br>(0.0949-0.0998) | n=19,694<br>med:0.0988<br>(0.0971-0.1002) | n=18,746<br>med:0.0835<br>(0.0825-0.0849) | n=19,780<br>med:0.0822<br>(0.0810-0.0834) |
| Oligodendrocyte | n=5,835<br>med:0.0897<br>(0.0871-0.0922) | n=18,767<br>med:0.0880<br>(0.0863-0.0896) | n=18,184<br>med:0.0801<br>(0.0788-0.0815) | n=19,122<br>med:0.0767<br>(0.0753-0.0778) |
| Neuron | n=13,525<br>med:0.0718<br>(0.0700-0.0736) | n=44,247<br>med:0.0620<br>(0.0611-0.0630) | n=42,654<br>med:0.0533<br>(0.0527-0.0539) | n=44,885<br>med:0.0508<br>(0.0501-0.0514) |
