## Supplementary Table S2 for "Neuron-specific coding and regulatory sequences are the most highly conserved in amniote brains despite neuron-specific cell size diversity"

### Chicken, Zhang et al., 2014, three cell types

| MWU Test | Glia | Neuron |
|---|---|---|
| Endothelia | U=178820666.5<br>$p=3.719 \times 10^{-4}$ | U=260663884.5<br>**$p<5 \times 10^{-324}$** |
| Glia | - | U=304898539.5<br>**$p=1.836 \times 10^{-296}$** |

### Chicken, Zhang et al., 2014, cell subtypes

| MWU Test | Endothelia | Astrocyte | Oligodendrocyte | Neuron |
|---|---|---|---|---|
| Microglia | U=144731413.0<br>$p=3.093 \times 10^{-4}$ | U=147393240.0<br>**$p=6.81 \times 10^{-48}$** | U=99114995.5<br>**$p=7.232 \times 10^{-95}$** | U=254149455.0<br>**$p<5 \times 10^{-324}$** |
| Endothelia | - | U=150632096.0<br>**$p=1.709 \times 10^{-28}$** | U=101350603.5<br>**$p=4.227 \times 10^{-69}$** | U=260663884.5<br>**$p<5 \times 10^{-324}$** |
| Astrocyte | - | - | U=90618746.5<br>**$p=9.715 \times 10^{-15}$** | U=235490078.0<br>**$p=2.381 \times 10^{-153}$** |
| Oligodendrocyte | - | - | - | U=143903349.5<br>**$p=4.331 \times 10^{-52}$** |

### Chicken, Zeisel et al., 2015, three cell types

| MWU Test | Vasculature | Neuron |
| --- | --- | --- |
| Glia | U=119182969.0<br>**p=1.78 × $10^{-30}$** | U=272972438.0<br>**p<5 × $10^{-324}$** |
| Vasculature | - | U=91695966.5<br>**p=2.79 × $10^{-99}$** |

### Chicken, Zeisel et al., 2015, cell subtypes

| MWU Test | Microglia | Mural | Endothelial | Oligodendrocyte | Astrocyte | Interneuron | S1 Pyramidal | CA1 Pyramidal |
|---|---|---|---|---|---|---|---|---|
| Ependymal | U=22751561.0<br>**$p=1.409 \times 10^{-30}$** | U=11123611.0<br>**$p=3.34 \times 10^{-85}$** | U=25086392.0<br>**$p=4.089 \times 10^{-124}$** | U=33160476.5<br>**$p=3.379 \times 10^{-186}$** | U=17521974.5<br>**$p=5.173 \times 10^{-148}$** | U=27943036.0<br>**$p=3.719 \times 10^{-295}$** | U=21152721.0<br>**$p=8.601 \times 10^{-268}$** | U=31368608.5<br>**$p<5 \times 10^{-324}$** |
| Microglia | - | U=9681274.5<br>**$p=1.721 \times 10^{-29}$** | U=21932364.5<br>**$p=4.379 \times 10^{-42}$** | U=29043110.0<br>**$p=5.527 \times 10^{-74}$** | U=15425891.0<br>**$p=2.383 \times 10^{-66}$** | U=24893020.0<br>**$p=5.882 \times 10^{-169}$** | U=18870794.5<br>**$p=8.182 \times 10^{-158}$** | U=28120554.5<br>**$p=4.975 \times 10^{-194}$** |
| Mural | - | - | U=8334086.0<br>p=0.906 | U=11041995.5<br>$p=1.482 \times 10^{-2}$ | U=5899607.0<br>**$p=8.064 \times 10^{-5}$** | U=9665038.5<br>**$p=1.022 \times 10^{-29}$** | U=7327075.0<br>**$p=4.715 \times 10^{-31}$** | U=11002523.0<br>**$p=6.427 \times 10^{-39}$** |
| Endothelial | - | - | - | U=25225205.5<br>$p=1.146 \times 10^{-3}$ | U=13453765.0<br>**$p=2.148 \times 10^{-6}$** | U=21998975.0<br>**$p=1.437 \times 10^{-45}$** | U=16659301.5<br>**$p=1.525 \times 10^{-44}$** | U=25043053.5<br>**$p=1.654 \times 10^{-61}$** |
| Oligodendrocyte | - | - | - | - | U=16739409.5<br>$p=3.473 \times 10^{-2}$ | U=27544987.5<br>**$p=2.082 \times 10^{-34}$** | U=20852619.5<br>**$p=1.064 \times 10^{-33}$** | U=31395165.5<br>**$p=9.693 \times 10^{-50}$** |
| Astrocyte | - | - | - | - | - | U=14078281.0<br>**$p=5.661 \times 10^{-17}$** | U=10661458.0<br>**$p=6.067 \times 10^{-18}$** | U=16091192.0<br>**$p=1.796 \times 10^{-26}$** |
| Interneuron | - | - | - | - | - | - | U=14575546.5<br>p=0.606 | U=22128714.0<br>$p=1.39 \times 10^{-2}$ |
| S1 Pyramidal | - | - | - | - | - | - | - | U=16556757.0<br>$p=5.25 \times 10^{-2}$ |

### Chicken, Saunders et al., 2018, three cell types

| MWU Test | Glia | Neuron |
| --- | --- | --- |
| Vasculature | U=28033471.0 **$p=1.406 \times 10^{-15}$** | U=75723891.0 **$p=2.495 \times 10^{-191}$** |
| Glia | - | U=54350043.0 **$p=1.033 \times 10^{-88}$** |

### Chicken, Saunders et al., 2018, cell subtypes

| MWU Test | Endothelia | Astrocyte | Oligodendrocyte | Neuron |
|---|---|---|---|---|
| Microglia | U=19477593.0 p=0.94 | U=17723557.0 **$p=2.514 \times 10^{-31}$** | U=17052470.5 **$p=5.339 \times 10^{-37}$** | U=43853825.0 **$p=1.002 \times 10^{-172}$** |
| Endothelia | - | U=26201478.0 **$p=3.218 \times 10^{-35}$** | U=25227126.5 **$p=3.754 \times 10^{-42}$** | U=64698275.0 **$p=6.501 \times 10^{-213}$** |
| Astrocyte | - | - | U=18289221.0 $p=4.037 \times 10^{-2}$ | U=48022936.5 **$p=3.928 \times 10^{-70}$** |
| Oligodendrocyte | - | - | - | U=44651241.0 **$p=5.792 \times 10^{-48}$** |

### Chicken, Zeisel et al., 2018, three cell types

| MWU Test | CNS_Endothelia | CNS_Neuron |
|---|---|---|
| CNS_Glia | U=27465398.5 **$p=7.102 \times 10^{-15}$** | U=809587112.0 **$p<5 \times 10^{-324}$** |
| CNS_Endothelia | - | U=134494874.0 **$p=4.452 \times 10^{-68}$** |

### Chicken, Zeisel et al., 2018, cell subtypes

| MWU Test | Immune cells | Vascular cells | Neurons |
|---|---|---|---|
| Glia | U=11247599.0<br>p=7.471 × $10^{-4}$ | U=28119105.0<br>**p=2.943 × $10^{-103}$** | U=699512394.5<br>**p<5 × $10^{-324}$** |
| Immune cells | - | U=3106638.0<br>**p=6.294 × $10^{-31}$** | U=77489834.5<br>**p=5.156 × $10^{-77}$** |
| Vascular cells | - | - | U=139107771.0<br>**p=7.643 × $10^{-15}$** |

### Mouse, Zhang et al., 2014, three cell types

| MWU Test | Endothelia | Neuron |
|---|---|---|
| Glia | U=2611899751.0<br>p=0.4 | U=4846129544.0<br>**p<5 × 10$^{-324}$** |
| Endothelia | - | U=3746814246.5<br>**p<5 × 10$^{-324}$** |

### Mouse, Zhang et al., 2014, cell subtypes

| MWU Test | Endothelia | Astrocyte | Oligodendrocyte | Neuron |
|---|---|---|---|---|
| Microglia | U=2364905819.0<br>**p=3.214 × $10^{-98}$** | U=2348590757.0<br>**p<5 × $10^{-324}$** | U=1622114081.5<br>**p<5 × $10^{-324}$** | U=4286291963.0<br>**p<5 × $10^{-324}$** |
| Endothelia | - | U=2022080139.0<br>**p=8.448 × $10^{-214}$** | U=1405036410.0<br>**p<5 × $10^{-324}$** | U=3746814246.5<br>**p<5 × $10^{-324}$** |
| Astrocyte | - | - | U=1165202849.5<br>**p=1.61 × $10^{-142}$** | U=3157612190.5<br>**p<5 × $10^{-324}$** |
| Oligodendrocyte | - | - | - | U=1869563755.5<br>**p=3.584 × $10^{-269}$** |

### Mouse, Zeisel et al., 2015, three cell types

| MWU Test | Vasculature | Neuron |
|---|---|---|
| Glia | U=1633174372.0 **$p=3.541 \times 10^{-145}$** | U=3816277403.5 **$p<5 \times 10^{-324}$** |
| Vasculature | - | U=1188412862.5 **$p<5 \times 10^{-324}$** |

### Mouse, Zeisel et al., 2015, cell subtypes

| MWU Test | Microglia | Endothelial | Oligodendrocyte | Astrocyte | Mural | Interneuron | S1 Pyramidal | CA1 Pyramidal |
|---|---|---|---|---|---|---|---|---|
| Ependymal | U=342275232.5<br>$p=2.07 \times 10^{-2}$ | U=350695988.5<br>**$p=4.118 \times 10^{-217}$** | U=463113072.0<br>**$p<5 \times 10^{-324}$** | U=249035621.0<br>**$p<5 \times 10^{-324}$** | U=169741863.5<br>**$p<5 \times 10^{-324}$** | U=415270757.5<br>**$p<5 \times 10^{-324}$** | U=316408632.5<br>**$p<5 \times 10^{-324}$** | U=460449177.5<br>**$p<5 \times 10^{-324}$** |
| Microglia | - | U=311400596.5<br>**$p=3.247 \times 10^{-177}$** | U=410486312.5<br>**$p<5 \times 10^{-324}$** | U=220328905.0<br>**$p=1.627 \times 10^{-292}$** | U=150282122.5<br>**$p<5 \times 10^{-324}$** | U=368954509.0<br>**$p<5 \times 10^{-324}$** | U=280884916.0<br>**$p<5 \times 10^{-324}$** | U=409657333.0<br>**$p<5 \times 10^{-324}$** |
| Endothelial | - | - | U=317813133.0<br>**$p=8.291 \times 10^{-24}$** | U=169844860.5<br>**$p=4.28 \times 10^{-20}$** | U=116477527.5<br>**$p=1.126 \times 10^{-44}$** | U=292929247.0<br>**$p<5 \times 10^{-324}$** | U=222289596.0<br>**$p=6.876 \times 10^{-288}$** | U=326456348.0<br>**$p<5 \times 10^{-324}$** |
| Oligodendrocyte | - | - | - | U=202361344.5<br>$p=0.593$ | U=139277757.5<br>**$p=5.322 \times 10^{-12}$** | U=354459357.0<br>**$p=4.538 \times 10^{-220}$** | U=268802328.5<br>**$p=6.629 \times 10^{-200}$** | U=396010483.0<br>**$p<5 \times 10^{-324}$** |
| Astrocyte | - | - | - | - | U=74043629.0<br>**$p=1.735 \times 10^{-9}$** | U=189519300.0<br>**$p=2.442 \times 10^{-168}$** | U=143758527.5<br>**$p=5.164 \times 10^{-160}$** | U=212102082.0<br>**$p=2.784 \times 10^{-244}$** |
| Mural | - | - | - | - | - | U=120459665.0<br>**$p=1.017 \times 10^{-73}$** | U=91284939.0<br>**$p=3.108 \times 10^{-72}$** | U=134993067.5<br>**$p=2.498 \times 10^{-117}$** |
| Interneuron | - | - | - | - | - | - | U=184152051.0<br>$p=0.872$ | U=273792061.0<br>**$p=9.058 \times 10^{-8}$** |
| S1 Pyramidal | - | - | - | - | - | - | - | U=206924406.5<br>**$p=1.829 \times 10^{-7}$** |

### Mouse, Saunders et al., 2018, three cell types

| MWU Test | Glia | Neuron |
|---|---|---|
| Vasculature | U=362093691.5<br>**$p=4.591 \times 10^{-161}$** | U=1000486018.5<br>**$p<5 \times 10^{-324}$** |
| Glia | - | U=651117111.0<br>**$p<5 \times 10^{-324}$** |

### Mouse, Saunders et al., 2018, cell subtypes

| MWU Test | Endothelia | Astrocyte | Oligodendrocyte | Neuron |
|---|---|---|---|---|
| Microglia | U=269376399.5 **$p=5.873 \times 10^{-48}$** | U=233858004.0 **$p<5 \times 10^{-324}$** | U=230876538.0 **$p<5 \times 10^{-324}$** | U=619864175.5 **$p<5 \times 10^{-324}$** |
| Endothelia | - | U=302468455.5 **$p=4.739 \times 10^{-200}$** | U=299612455.0 **$p=4.412 \times 10^{-270}$** | U=816532886.5 **$p<5 \times 10^{-324}$** |
| Astrocyte | - | - | U=195227440.5 **$p=3.418 \times 10^{-8}$** | U=553231754.5 **$p<5 \times 10^{-324}$** |
| Oligodendrocyte | - | - | - | U=521112665.5 **$p<5 \times 10^{-324}$** |

### Mouse, Zeisel et al., 2018, three cell types

| MWU Test | CNS_Endothelia | CNS_Neuron |
|---|---|---|
| CNS_Glia | U=411451329.0<br>$p=1.591 \times 10^{-2}$ | U=11188504876.0<br>$\mathbf{p<5 \times 10^{-324}}$ |
| CNS_Endothelia | - | U=2054703896.0<br>$\mathbf{p<5 \times 10^{-324}}$ |

### Mouse, Zeisel et al., 2018, cell subtypes

| MWU Test | Glia | Vascular cells | Neurons |
|---|---|---|---|
| Immune cells | U=215704545.0 **$p=9.537 \times 10^{-17}$** | U=62390412.5 **$p=2.2 \times 10^{-138}$** | U=1389089079.5 **$p<5 \times 10^{-324}$** |
| Glia | - | U=439596861.5 **$p=2.26 \times 10^{-193}$** | U=9935523351.0 **$p<5 \times 10^{-324}$** |
| Vascular cells | - | - | U=2250080985.5 **$p<5 \times 10^{-324}$** |

### Rat, Zhang et al., 2014, three cell types

| MWU Test | Endothelia | Neuron |
|---|---|---|
| Glia | U=2271288736.0 p=0.622 | U=4302746845.0 **$p<5 \times 10^{-324}$** |
| Endothelia | - | U=3272130865.0 **$p<5 \times 10^{-324}$** |

### Rat, Zhang et al., 2014, cell subtypes

| MWU Test | Endothelia | Astrocyte | Oligodendrocyte | Neuron |
|---|---|---|---|---|
| Microglia | U=2076506098.5 **$p=7.453 \times 10^{-99}$** | U=2118607361.0 **$p<5 \times 10^{-324}$** | U=1490535384.0 **$p<5 \times 10^{-324}$** | U=3833165109.0 **$p<5 \times 10^{-324}$** |
| Endothelia | - | U=1778264220.0 **$p=1.855 \times 10^{-209}$** | U=1260771512.0 **$p<5 \times 10^{-324}$** | U=3272130865.0 **$p<5 \times 10^{-324}$** |
| Astrocyte | - | - | U=1071494749.0 **$p=2.195 \times 10^{-183}$** | U=2819987309.5 **$p<5 \times 10^{-324}$** |
| Oligodendrocyte | - | - | - | U=1660996773.5 **$p=5.279 \times 10^{-206}$** |

### Rat, Zeisel et al., 2015, three cell types

| MWU Test | Vasculature | Neuron |
| --- | --- | --- |
| Glia | U=1430669832.0<br>**p=3.836 × $10^{-109}$** | U=3397310246.0<br>**p<5 × $10^{-324}$** |
| Vasculature | - | U=1057580299.5<br>**p<5 × $10^{-324}$** |

### Rat, Zeisel et al., 2015, cell subtypes

| MWU Test | Microglia | Endothelial | Astrocyte | Mural | Oligodendrocyte | Interneuron | S1 Pyramidal | CA1 Pyramidal |
|---|---|---|---|---|---|---|---|---|
| Ependymal | U=295949241.5<br>$p=5.427 \times 10^{-4}$ | U=304316952.0<br>**$p=3.008 \times 10^{-168}$** | U=221694690.5<br>**$p=1.154 \times 10^{-281}$** | U=147021286.5<br>**$p=2.368 \times 10^{-291}$** | U=412308002.0<br>**$p<5 \times 10^{-324}$** | U=358262066.0<br>**$p<5 \times 10^{-324}$** | U=286335450.5<br>**$p<5 \times 10^{-324}$** | U=404446089.0<br>**$p<5 \times 10^{-324}$** |
| Microglia | - | U=278869803.0<br>**$p=1.045 \times 10^{-199}$** | U=202616273.5<br>**$p=3.28 \times 10^{-310}$** | U=134248867.0<br>**$p<5 \times 10^{-324}$** | U=377137698.5<br>**$p<5 \times 10^{-324}$** | U=327575970.5<br>**$p<5 \times 10^{-324}$** | U=261872094.5<br>**$p<5 \times 10^{-324}$** | U=370334129.5<br>**$p<5 \times 10^{-324}$** |
| Endothelial | - | - | U=152821118.5<br>**$p=8.058 \times 10^{-20}$** | U=101864038.5<br>**$p=1.726 \times 10^{-41}$** | U=286597676.5<br>**$p=3.268 \times 10^{-49}$** | U=255189213.5<br>**$p=8.216 \times 10^{-290}$** | U=203713975.0<br>**$p=6.127 \times 10^{-296}$** | U=290538821.0<br>**$p<5 \times 10^{-324}$** |
| Astrocyte | - | - | - | U=66479414.5<br>**$p=8.244 \times 10^{-8}$** | U=187171677.0<br>**$p=4.532 \times 10^{-5}$** | U=169012225.0<br>**$p=2.957 \times 10^{-145}$** | U=134874808.0<br>**$p=7.689 \times 10^{-156}$** | U=193210480.5<br>**$p=4.94 \times 10^{-238}$** |
| Mural | - | - | - | - | U=115276399.0<br>$p=1.739 \times 10^{-2}$ | U=104761270.0<br>**$p=3.724 \times 10^{-63}$** | U=83562977.0<br>**$p=5.216 \times 10^{-72}$** | U=119814043.0<br>**$p=1.875 \times 10^{-114}$** |
| Oligodendrocyte | - | - | - | - | - | U=303144374.5<br>**$p=2.399 \times 10^{-137}$** | U=241875261.0<br>**$p=4.978 \times 10^{-146}$** | U=346367297.0<br>**$p=2.461 \times 10^{-243}$** |
| Interneuron | - | - | - | - | - | - | U=169139806.5<br>$p=0.151$ | U=244000470.0<br>**$p=1.57 \times 10^{-12}$** |
| S1 Pyramidal | - | - | - | - | - | - | - | U=190750232.0<br>**$p=1.662 \times 10^{-7}$** |

### Rat, Saunders et al., 2018, three cell types

| MWU Test | Glia | Neuron |
|---|---|---|
| Vasculature | U=321745571.0<br>**$p=1.445 \times 10^{-173}$** | U=873418011.5<br>**$p<5 \times 10^{-324}$** |
| Glia | - | U=584389992.0<br>**$p<5 \times 10^{-324}$** |

### Rat, Saunders et al., 2018, cell subtypes

| MWU Test | Endothelia | Astrocyte | Oligodendrocyte | Neuron |
|---|---|---|---|---|
| Microglia | U=240447130.0 **$p=2.468 \times 10^{-53}$** | U=212646066.5 **$p<5 \times 10^{-324}$** | U=209899113.5 **$p<5 \times 10^{-324}$** | U=559711972.5 **$p<5 \times 10^{-324}$** |
| Endothelia | - | U=270483821.0 **$p=1.725 \times 10^{-208}$** | U=267787574.5 **$p=8.262 \times 10^{-266}$** | U=726589157.0 **$p<5 \times 10^{-324}$** |
| Astrocyte | - | - | U=175287626.5 **$p=2.199 \times 10^{-6}$** | U=496063591.5 **$p<5 \times 10^{-324}$** |
| Oligodendrocyte | - | - | - | U=469914559.0 **$p<5 \times 10^{-324}$** |

### Rat, Zeisel et al., 2018, three cell types

| MWU Test | CNS_Endothelia | CNS_Neuron |
|---|---|---|
| CNS_Glia | U=352618273.0<br>p=0.267 | U=9975829672.0<br>**$p<5 \times 10^{-324}$** |
| CNS_Endothelia | - | U=1780557678.0<br>**$p<5 \times 10^{-324}$** |

### Rat, Zeisel et al., 2018, cell subtypes

| MWU Test | Glia | Vascular cells | Neurons |
|---|---|---|---|
| Immune cells | U=198553984.0 **$p=6.158 \times 10^{-26}$** | U=56118634.5 **$p=4.869 \times 10^{-149}$** | U=1258775709.5 **$p<5 \times 10^{-324}$** |
| Glia | - | U=382649347.5 **$p=2.32 \times 10^{-184}$** | U=8705097254.0 **$p<5 \times 10^{-324}$** |
| Vascular cells | - | - | U=1932908227.0 **$p=9.294 \times 10^{-264}$** |

### Human, Zhang et al., 2014, three cell types

| MWU Test | Glia | Neuron |
|---|---|---|
| Endothelia | U=2536870798.0<br>**p=1.562 × $10^{-10}$** | U=3527136160.5<br>**p<5 × $10^{-324}$** |
| Glia | - | U=4549211852.0<br>**p<5 × $10^{-324}$** |

### Human, Zhang et al., 2014, cell subtypes

| MWU Test | Endothelia | Astrocyte | Oligodendrocyte | Neuron |
|---|---|---|---|---|
| Microglia | U=2195430707.0 **$p=1.257 \times 10^{-23}$** | U=2192605527.5 **$p=5.348 \times 10^{-289}$** | U=1535429100.0 **$p<5 \times 10^{-324}$** | U=4019382133.0 **$p<5 \times 10^{-324}$** |
| Endothelia | - | U=1913559227.0 **$p=1.013 \times 10^{-144}$** | U=1342974083.5 **$p<5 \times 10^{-324}$** | U=3527136160.5 **$p<5 \times 10^{-324}$** |
| Astrocyte | - | - | U=1150213151.5 **$p=1.477 \times 10^{-173}$** | U=3060452601.0 **$p<5 \times 10^{-324}$** |
| Oligodendrocyte | - | - | - | U=1796317933.0 **$p=9.425 \times 10^{-210}$** |

### Human, Zeisel et al., 2015, three cell types

| MWU Test | Vasculature | Neuron |
|---|---|---|
| Glia | U=1587577513.0<br>**p=3.99 × $10^{-149}$** | U=3667528817.5<br>**p<5 × $10^{-324}$** |
| Vasculature | - | U=1149444130.0<br>**p<5 × $10^{-324}$** |

### Human, Zeisel et al., 2015, cell subtypes

| MWU Test | Microglia | Endothelial | Astrocyte | Oligodendrocyte | Mural | Interneuron | S1 Pyramidal | CA1 Pyramidal |
|---|---|---|---|---|---|---|---|---|
| Ependymal | U=339936855.0 **$p=1.051 \times 10^{-30}$** | U=351338103.5 **$p=1.976 \times 10^{-294}$** | U=246096848.0 **$p<5 \times 10^{-324}$** | U=462809032.5 **$p<5 \times 10^{-324}$** | U=166558264.5 **$p<5 \times 10^{-324}$** | U=404368675.0 **$p<5 \times 10^{-324}$** | U=308914154.0 **$p<5 \times 10^{-324}$** | U=456181295.0 **$p<5 \times 10^{-324}$** |
| Microglia | - | U=296486826.5 **$p=1.035 \times 10^{-150}$** | U=207277606.5 **$p=1.308 \times 10^{-210}$** | U=391112058.0 **$p<5 \times 10^{-324}$** | U=140596510.0 **$p=5.582 \times 10^{-278}$** | U=344727530.0 **$p<5 \times 10^{-324}$** | U=263250893.5 **$p<5 \times 10^{-324}$** | U=390038154.5 **$p<5 \times 10^{-324}$** |
| Endothelial | - | - | U=165376441.5 **$p=1.339 \times 10^{-7}$** | U=314428113.5 **$p=1.087 \times 10^{-38}$** | U=112456197.5 **$p=2.662 \times 10^{-36}$** | U=281064821.0 **$p=5.099 \times 10^{-214}$** | U=215067328.0 **$p=8.275 \times 10^{-241}$** | U=320133566.0 **$p<5 \times 10^{-324}$** |
| Astrocyte | - | - | - | U=205349135.0 **$p=8.376 \times 10^{-11}$** | U=73528046.0 **$p=2.837 \times 10^{-15}$** | U=185995868.5 **$p=4.027 \times 10^{-139}$** | U=142312339.0 **$p=8.859 \times 10^{-165}$** | U=212554654.5 **$p=4.723 \times 10^{-237}$** |
| Oligodendrocyte | - | - | - | - | U=129926937.5 $p=4.672 \times 10^{-3}$ | U=329699810.5 **$p=2.417 \times 10^{-104}$** | U=252389253.0 **$p=1.558 \times 10^{-129}$** | U=377096190.0 **$p=7.256 \times 10^{-208}$** |
| Mural | - | - | - | - | - | U=114376106.5 **$p=1.727 \times 10^{-46}$** | U=87580974.5 **$p=2.443 \times 10^{-64}$** | U=131221334.0 **$p=2.011 \times 10^{-99}$** |
| Interneuron | - | - | - | - | - | - | U=184645794.5 $p=1.876 \times 10^{-4}$ | U=277383821.0 **$p=6.348 \times 10^{-17}$** |
| S1 Pyramidal | - | - | - | - | - | - | - | U=203287912.0 $p=1.224 \times 10^{-4}$ |

### Human, Saunders et al., 2018, three cell types

| MWU Test | Glia | Neuron |
|---|---|---|
| Vasculature | U=358450826.5 **$p=3.876 \times 10^{-169}$** | U=953147105.0 **$p<5 \times 10^{-324}$** |
| Glia | - | U=627904477.5 **$p<5 \times 10^{-324}$** |

### Human, Saunders et al., 2018, cell subtypes

| MWU Test | Endothelia | Astrocyte | Oligodendrocyte | Neuron |
|---|---|---|---|---|
| Microglia | U=258662327.5 **$p=7.215 \times 10^{-23}$** | U=223723942.0 **$p=4.139 \times 10^{-275}$** | U=221284426.0 **$p<5 \times 10^{-324}$** | U=581654183.0 **$p<5 \times 10^{-324}$** |
| Endothelia | - | U=294665581.0 **$p=3.693 \times 10^{-159}$** | U=291798687.5 **$p=6.191 \times 10^{-263}$** | U=772597797.0 **$p<5 \times 10^{-324}$** |
| Astrocyte | - | - | U=193749937.0 **$p=1.96 \times 10^{-16}$** | U=528295358.5 **$p<5 \times 10^{-324}$** |
| Oligodendrocyte | - | - | - | U=485760462.5 **$p=2.107 \times 10^{-250}$** |

### Human, Zeisel et al., 2018, three cell types

| MWU Test | CNS_Endothelia | CNS_Neuron |
| --- | --- | --- |
| CNS_Glia | U=405686516.5<br>**p=1.585 × $10^{-5}$** | U=10725783659.5<br>**p<5 × $10^{-324}$** |
| CNS_Endothelia | - | U=1970617384.5<br>**p<5 × $10^{-324}$** |

### Human, Zeisel et al., 2018, cell subtypes

| MWU Test | Glia | Vascular cells | Neurons |
|---|---|---|---|
| Immune cells | U=200372463.5 **$p=1.737 \times 10^{-5}$** | U=59732419.0 **$p=2.053 \times 10^{-105}$** | U=1292996053.0 **$p<5 \times 10^{-324}$** |
| Glia | - | U=430906096.0 **$p=1.406 \times 10^{-176}$** | U=9401647826.5 **$p<5 \times 10^{-324}$** |
| Vascular cells | - | - | U=2190749751.5 **$p=1.519 \times 10^{-296}$** |
