## Supplementary Table S3 for "Neuron-specific coding and regulatory sequences are the most highly conserved in amniote brains despite neuron-specific cell size diversity"

### Benchmarks

| Sample Size and Median dN/dS (CI95%) | Chicken Reference Genome | Human Reference Genome | Rat Reference Genome | Mouse Reference Genome |
|---|---|---|---|---|
| MHC | n=7<br>med:0.1478<br>(0.0481-0.2474) | n=20<br>med:0.3627<br>(0.2952-0.4451) | n=13<br>med:0.2648<br>(0.2026-0.3398) | n=26<br>med:0.4454<br>(0.3369-0.5110) |
| Immune | n=430<br>med:0.1502<br>(0.1328-0.1778) | n=1,271<br>med:0.1426<br>(0.1336-0.1570) | n=725<br>med:0.1575<br>(0.1454-0.1803) | n=713<br>med:0.1440<br>(0.1298-0.1622) |
| protein-coding | n=14,032<br>med:0.1182<br>(0.1159-0.1211) | n=18,412<br>med:0.1562<br>(0.1536-0.1589) | n=17,812<br>med:0.1227<br>(0.1204-0.1247) | n=19,595<br>med:0.1280<br>(0.1257-0.1301) |
| Housekeeping | n=2,716<br>med:0.0941<br>(0.0899-0.0989) | n=3,493<br>med:0.1109<br>(0.1072-0.1159) | n=3,058<br>med:0.0852<br>(0.0820-0.0885) | n=3,248<br>med:0.0841<br>(0.0809-0.0877) |
| ATPase | n=107<br>med:0.0948<br>(0.0698-0.1063) | n=136<br>med:0.0984<br>(0.0839-0.1189) | n=114<br>med:0.0717<br>(0.0599-0.0855) | n=128<br>med:0.0706<br>(0.0578-0.0831) |

### Zhang et al., 2014, three cell types

| Sample Size and Median dN/dS (CI95%) | Chicken Reference Genome | Human Reference Genome | Rat Reference Genome | Mouse Reference Genome |
|---|---|---|---|---|
| Endothelia | n=651<br>med:0.1228<br>(0.1128-0.1348) | n=833<br>med:0.1537<br>(0.1405-0.1612) | n=841<br>med:0.1226<br>(0.1117-0.1309) | n=942<br>med:0.1234<br>(0.1144-0.1329) |
| Glia | n=795<br>med:0.1162<br>(0.1072-0.1249) | n=1,086<br>med:0.1437<br>(0.1372-0.1527) | n=1,112<br>med:0.1251<br>(0.1185-0.1329) | n=1,218<br>med:0.1268<br>(0.1204-0.1339) |
| Neuron | n=942<br>med:0.0867<br>(0.0807-0.0939) | n=1,215<br>med:0.0902<br>(0.0836-0.0946) | n=1,201<br>med:0.0718<br>(0.0675-0.0772) | n=1,301<br>med:0.0720<br>(0.0671-0.0759) |

### Zhang et al., 2014, cell subtypes

| Sample Size and Median dN/dS (CI95%) | Chicken Reference Genome | Human Reference Genome | Rat Reference Genome | Mouse Reference Genome |
|---|---|---|---|---|
| Microglia | n=640<br>med:0.1262<br>(0.1162-0.1367) | n=932<br>med:0.1555<br>(0.1462-0.1666) | n=960<br>med:0.1378<br>(0.1311-0.1453) | n=1,069<br>med:0.1428<br>(0.1350-0.1529) |
| Endothelia | n=651<br>med:0.1228<br>(0.1128-0.1348) | n=833<br>med:0.1537<br>(0.1405-0.1612) | n=841<br>med:0.1226<br>(0.1117-0.1309) | n=942<br>med:0.1234<br>(0.1144-0.1329) |
| Astrocyte | n=619<br>med:0.1111<br>(0.1008-0.1207) | n=761<br>med:0.1299<br>(0.1230-0.1398) | n=763<br>med:0.1106<br>(0.0993-0.1181) | n=825<br>med:0.1106<br>(0.1019-0.1173) |
| Oligodendrocyte | n=409<br>med:0.1018<br>(0.0881-0.1071) | n=492<br>med:0.1075<br>(0.1017-0.1147) | n=494<br>med:0.0890<br>(0.0804-0.0962) | n=524<br>med:0.0867<br>(0.0808-0.0958) |
| Neuron | n=942<br>med:0.0867<br>(0.0807-0.0939) | n=1,215<br>med:0.0902<br>(0.0836-0.0946) | n=1,201<br>med:0.0718<br>(0.0675-0.0772) | n=1,301<br>med:0.0720<br>(0.0671-0.0759) |

### Zeisel et al., 2015, three cell types

| Sample Size and Median dN/dS (CI95%) | Chicken Reference Genome | Human Reference Genome | Rat Reference Genome | Mouse Reference Genome |
|---|---|---|---|---|
| Glia | n=946<br>med:0.1341<br>(0.1250-0.1409) | n=1,245<br>med:0.1579<br>(0.1507-0.1657) | n=1,254<br>med:0.1273<br>(0.1195-0.1348) | n=1,386<br>med:0.1322<br>(0.1259-0.1388) |
| Vasculature | n=334<br>med:0.1182<br>(0.1045-0.1317) | n=419<br>med:0.1232<br>(0.1152-0.1371) | n=408<br>med:0.1085<br>(0.0972-0.1185) | n=452<br>med:0.1062<br>(0.0957-0.1171) |
| Neuron | n=673<br>med:0.0918<br>(0.0816-0.0966) | n=853<br>med:0.0958<br>(0.0888-0.1036) | n=840<br>med:0.0748<br>(0.0693-0.0820) | n=897<br>med:0.0745<br>(0.0690-0.0809) |

### Zeisel et al., 2015, cell subtypes

| Sample Size and Median dN/dS (CI95%) | Chicken Reference Genome | Human Reference Genome | Rat Reference Genome | Mouse Reference Genome |
|---|---|---|---|---|
| Ependymal | n=257<br>med:0.1753<br>(0.1633-0.1923) | n=365<br>med:0.1996<br>(0.1833-0.2206) | n=362<br>med:0.1505<br>(0.1370-0.1647) | n=401<br>med:0.1550<br>(0.1427-0.1761) |
| Microglia | n=235<br>med:0.1482<br>(0.1296-0.1587) | n=320<br>med:0.1746<br>(0.1536-0.1951) | n=337<br>med:0.1508<br>(0.1353-0.1692) | n=381<br>med:0.1584<br>(0.1389-0.1752) |
| Endothelial | n=233<br>med:0.1184<br>(0.1037-0.1347) | n=291<br>med:0.1296<br>(0.1158-0.1566) | n=283<br>med:0.1093<br>(0.0972-0.1307) | n=316<br>med:0.1107<br>(0.0976-0.1262) |
| Mural | n=101<br>med:0.1172<br>(0.0994-0.1343) | n=128<br>med:0.1200<br>(0.1034-0.1320) | n=125<br>med:0.1063<br>(0.0887-0.1157) | n=136<br>med:0.1005<br>(0.0834-0.1138) |
| Oligodendrocyte | n=298<br>med:0.1143<br>(0.0991-0.1256) | n=365<br>med:0.1256<br>(0.1160-0.1387) | n=362<br>med:0.1032<br>(0.0939-0.1137) | n=393<br>med:0.1086<br>(0.0959-0.1181) |
| Astrocyte | n=156<br>med:0.1009<br>(0.0889-0.1166) | n=195<br>med:0.1306<br>(0.1128-0.1511) | n=193<br>med:0.1116<br>(0.0947-0.1209) | n=211<br>med:0.1092<br>(0.0953-0.1242) |
| Interneuron | n=236<br>med:0.0967<br>(0.0816-0.1028) | n=299<br>med:0.1031<br>(0.0913-0.1156) | n=293<br>med:0.0800<br>(0.0687-0.0901) | n=316<br>med:0.0752<br>(0.0671-0.0872) |
| S1 Pyramidal | n=179 | n=223 | n=227 | n=235 |

| | | | | |
|---|---|---|---|---|
| CA1 Pyramidal | med:0.0889 (0.0804-0.1002) | med:0.0959 (0.0765-0.1120) | med:0.0768 (0.0665-0.0958) | med:0.0779 (0.0628-0.0905) |
| | n=258 med:0.0810 (0.0756-0.0967) | n=331 med:0.0893 (0.0800-0.1009) | n=320 med:0.0704 (0.0631-0.0774) | n=346 med:0.0721 (0.0638-0.0809) |

### Zeisel et al., 2018, three cell types

| Sample Size and Median dN/dS (CI95%) | Chicken Reference Genome | Human Reference Genome | Rat Reference Genome | Mouse Reference Genome |
|---|---|---|---|---|
| CNS_Glia | n=655 med:0.1427 (0.1268-0.1552) | n=874 med:0.1629 (0.1510-0.1751) | n=875 med:0.1283 (0.1192-0.1367) | n=954 med:0.1339 (0.1268-0.1415) |
| CNS_Endothelia | n=115 med:0.1280 (0.1033-0.1517) | n=165 med:0.1584 (0.1354-0.1711) | n=155 med:0.1216 (0.1073-0.1392) | n=174 med:0.1231 (0.1107-0.1449) |
| CNS_Neuron | n=2,807 med:0.0925 (0.0883-0.0954) | n=3,371 med:0.0988 (0.0951-0.1022) | n=3,331 med:0.0772 (0.0744-0.0806) | n=3,576 med:0.0779 (0.0756-0.0810) |

### Zeisel et al., 2018, cell subtypes

| Sample Size and Median dN/dS (CI95%) | Chicken Reference Genome | Human Reference Genome | Rat Reference Genome | Mouse Reference Genome |
|---|---|---|---|---|
| Glia | n=532<br>med:0.1684<br>(0.1541-0.1834) | n=723<br>med:0.1807<br>(0.1694-0.1992) | n=717<br>med:0.1457<br>(0.1366-0.1544) | n=784<br>med:0.1528<br>(0.1426-0.1626) |
| Immune cells | n=60<br>med:0.1613<br>(0.1176-0.1936) | n=100<br>med:0.1979<br>(0.1643-0.2248) | n=116<br>med:0.1746<br>(0.1335-0.2155) | n=132<br>med:0.1930<br>(0.1529-0.2232) |
| Vascular cells | n=132<br>med:0.1146<br>(0.0927-0.1317) | n=188<br>med:0.1406<br>(0.1199-0.1607) | n=176<br>med:0.1116<br>(0.0963-0.1307) | n=194<br>med:0.1185<br>(0.1013-0.1373) |
| Neurons | n=2,904<br>med:0.0994<br>(0.0960-0.1024) | n=3,540<br>med:0.1081<br>(0.1048-0.1120) | n=3,506<br>med:0.0861<br>(0.0834-0.0892) | n=3,772<br>med:0.0881<br>(0.0841-0.0915) |

### Saunders et al., 2018, three cell types

| Sample Size and Median dN/dS (CI95%) | Chicken Reference Genome | Human Reference Genome | Rat Reference Genome | Mouse Reference Genome |
|---|---|---|---|---|
| Vasculature | n=317<br>med:0.1121<br>(0.0967-0.1322) | n=397<br>med:0.1416<br>(0.1259-0.1580) | n=391<br>med:0.1082<br>(0.0959-0.1189) | n=432<br>med:0.1091<br>(0.0981-0.1193) |
| Glia | n=239<br>med:0.0968<br>(0.0889-0.1053) | n=287<br>med:0.1067<br>(0.0999-0.1180) | n=285<br>med:0.0860<br>(0.0755-0.0955) | n=304<br>med:0.0843<br>(0.0774-0.0962) |
| Neuron | n=544<br>med:0.0772<br>(0.0693-0.0835) | n=644<br>med:0.0700<br>(0.0663-0.0756) | n=632<br>med:0.0536<br>(0.0500-0.0574) | n=675<br>med:0.0532<br>(0.0494-0.0568) |

### Saunders et al., 2018, cell subtypes

| Sample Size and Median dN/dS (CI95%) | Chicken Reference Genome | Human Reference Genome | Rat Reference Genome | Mouse Reference Genome |
|---|---|---|---|---|
| Endothelia | n=285<br>med:0.1184<br>(0.1010-0.1367) | n=351<br>med:0.1351<br>(0.1217-0.1545) | n=349<br>med:0.1085<br>(0.0959-0.1216) | n=382<br>med:0.1095<br>(0.1003-0.1193) |
| Microglia | n=197<br>med:0.1162<br>(0.1003-0.1407) | n=251<br>med:0.1436<br>(0.1302-0.1612) | n=252<br>med:0.1308<br>(0.1135-0.1457) | n=276<br>med:0.1311<br>(0.1117-0.1493) |
| Astrocyte | n=235<br>med:0.0941<br>(0.0867-0.1076) | n=265<br>med:0.1097<br>(0.0939-0.1197) | n=261<br>med:0.0848<br>(0.0729-0.0948) | n=283<br>med:0.0851<br>(0.0741-0.0983) |
| Oligodendrocyte | n=219<br>med:0.0904<br>(0.0800-0.0997) | n=254<br>med:0.0987<br>(0.0852-0.1095) | n=255<br>med:0.0839<br>(0.0745-0.0945) | n=271<br>med:0.0799<br>(0.0679-0.0901) |
| Neuron | n=499<br>med:0.0775<br>(0.0721-0.0837) | n=592<br>med:0.0720<br>(0.0672-0.0786) | n=584<br>med:0.0548<br>(0.0512-0.0608) | n=623<br>med:0.0537<br>(0.0502-0.0580) |
