## Supplementary Table S4 for "Neuron-specific coding and regulatory sequences are the most highly conserved in amniote brains despite neuron-specific cell size diversity"

### Chicken, Zhang et al., 2014, three cell types

| MWU Test | MHC | Endothelia | protein-coding | Glia | ATPase | Housekeeping | Neuron |
|---|---|---|---|---|---|---|---|
| Immune | U=1520.0<br>p=0.965<br>CLES:50.50% | U=158054.0<br>$p=3.177\times10^{-4}$<br>CLES:56.46% | U=3476414.0<br>**$p=7.097\times10^{-8}$**<br>CLES:57.62% | U=199845.5<br>**$p=9.901\times10^{-7}$**<br>CLES:58.46% | U=30600.0<br>**$p=1.237\times10^{-7}$**<br>CLES:66.51% | U=756534.0<br>**$p=6.079\times10^{-23}$**<br>CLES:64.78% | U=271456.5<br>**$p=4.295\times10^{-24}$**<br>CLES:67.02% |
| MHC | - | U=2654.0<br>p=0.453<br>CLES:58.24% | U=57958.5<br>p=0.409<br>CLES:59.01% | U=3342.0<br>p=0.36<br>CLES:60.05% | U=515.0<br>$p=9.844\times10^{-2}$<br>CLES:68.76% | U=12710.0<br>p=0.123<br>CLES:66.85% | U=4583.0<br>$p=7.521\times10^{-2}$<br>CLES:69.50% |
| Endothelia | - | - | U=4700267.5<br>p=0.209<br>CLES:51.45% | U=269424.0<br>p=0.178<br>CLES:52.06% | U=42457.0<br>$p=2.789\times10^{-4}$<br>CLES:60.95% | U=1046593.5<br>**$p=2.96\times10^{-13}$**<br>CLES:59.19% | U=380132.0<br>**$p=3.799\times10^{-16}$**<br>CLES:61.99% |
| protein-coding | - | - | - | U=5626691.5<br>p=0.677<br>CLES:50.44% | U=888022.5<br>$p=1.097\times10^{-3}$<br>CLES:59.15% | U=21896988.0<br>**$p=7.036\times10^{-35}$**<br>CLES:57.46% | U=7921184.0<br>**$p=1.677\times10^{-24}$**<br>CLES:59.93% |
| Glia | - | - | - | - | U=50377.0<br>$p=1.933\times10^{-3}$<br>CLES:59.22% | U=1239698.5<br>**$p=1.911\times10^{-10}$**<br>CLES:57.41% | U=451223.0<br>**$p=1.68\times10^{-13}$**<br>CLES:60.25% |
| ATPase | - | - | - | - | - | U=140837.5<br>p=0.589<br>CLES:48.46% | U=51232.0<br>p=0.779<br>CLES:50.83% |
| Housekeeping | - | - | - | - | - | - | U=1338778.5<br>$p=3.302\times10^{-2}$<br>CLES:52.33% |

### Chicken, Zhang et al., 2014, cell subtypes

| MWU Test | MHC | Microglia | Endothelia | protein-coding | Astrocyte | Oligodendrocyte | ATPase | Housekeeping | Neuron |
|---|---|---|---|---|---|---|---|---|---|
| Immune | U=1520.0<br>p=0.965<br>CLES:50.50% | U=155701.0<br>p=$2.599 \times 10^{-4}$<br>CLES:56.58% | U=158054.0<br>p=$3.177 \times 10^{-4}$<br>CLES:56.46% | U=3476414.0<br>**p=$7.097 \times 10^{-8}$**<br>CLES:57.62% | U=160154.5<br>**p=$2.034 \times 10^{-8}$**<br>CLES:60.17% | U=111660.0<br>**p=$1.364 \times 10^{-11}$**<br>CLES:63.49% | U=30600.0<br>**p=$1.237 \times 10^{-7}$**<br>CLES:66.51% | U=756534.0<br>**p=$6.079 \times 10^{-23}$**<br>CLES:64.78% | U=271456.5<br>**p=$4.295 \times 10^{-24}$**<br>CLES:67.02% |
| MHC | - | U=2615.5<br>p=0.446<br>CLES:58.38% | U=2654.0<br>p=0.453<br>CLES:58.24% | U=57958.5<br>p=0.409<br>CLES:59.01% | U=2674.0<br>p=0.287<br>CLES:61.71% | U=1871.0<br>p=0.164<br>CLES:65.35% | U=515.0<br>p=$9.844 \times 10^{-2}$<br>CLES:68.76% | U=12710.0<br>p=0.123<br>CLES:66.85% | U=4583.0<br>p=$7.521 \times 10^{-2}$<br>CLES:69.50% |
| Microglia | - | - | U=208145.0<br>p=0.979<br>CLES:49.96% | U=4613129.0<br>p=0.241<br>CLES:51.37% | U=213384.0<br>p=$1.765 \times 10^{-2}$<br>CLES:53.86% | U=150889.0<br>**p=$2.905 \times 10^{-5}$**<br>CLES:57.64% | U=41722.5<br>p=$2.93 \times 10^{-4}$<br>CLES:60.93% | U=1026503.0<br>**p=$9.538 \times 10^{-13}$**<br>CLES:59.05% | U=373227.0<br>**p=$8.302 \times 10^{-16}$**<br>CLES:61.91% |
| Endothelia | - | - | - | U=4700267.5<br>p=0.209<br>CLES:51.45% | U=217469.0<br>p=$1.442 \times 10^{-2}$<br>CLES:53.97% | U=153683.0<br>**p=$2.275 \times 10^{-5}$**<br>CLES:57.72% | U=42457.0<br>p=$2.789 \times 10^{-4}$<br>CLES:60.95% | U=1046593.5<br>**p=$2.96 \times 10^{-13}$**<br>CLES:59.19% | U=380132.0<br>**p=$3.799 \times 10^{-16}$**<br>CLES:61.99% |
| protein-coding | - | - | - | - | U=4535817.5<br>p=$6.103 \times 10^{-2}$<br>CLES:52.22% | U=3202004.5<br>**p=$6.326 \times 10^{-5}$**<br>CLES:55.79% | U=888022.5<br>p=$1.097 \times 10^{-3}$<br>CLES:59.15% | U=21896988.0<br>**p=$7.036 \times 10^{-35}$**<br>CLES:57.46% | U=7921184.0<br>**p=$1.677 \times 10^{-24}$**<br>CLES:59.93% |
| Astrocyte | - | - | - | - | - | U=136160.0<br>p=$3.99 \times 10^{-2}$<br>CLES:53.78% | U=38064.0<br>p=$1.353 \times 10^{-2}$<br>CLES:57.47% | U=936057.0<br>**p=$1.009 \times 10^{-5}$**<br>CLES:55.68% | U=341115.0<br>**p=$1.276 \times 10^{-8}$**<br>CLES:58.50% |
| Oligodendrocyte | - | - | - | - | - | - | U=23597.0<br>p=0.212<br>CLES:53.92% | U=578355.0<br>p=0.178<br>CLES:52.06% | U=211549.0<br>p=$4.104 \times 10^{-3}$<br>CLES:54.91% |
| ATPase | - | - | - | - | - | - | - | U=140837.5<br>p=0.589<br>CLES:48.46% | U=51232.0<br>p=0.779<br>CLES:50.83% |
| Housekeeping | - | - | - | - | - | - | - | - | U=1338778.5<br>p=$3.302 \times 10^{-2}$<br>CLES:52.33% |

### Chicken, Zeisel et al., 2015, three cell types

| MWU Test | MHC | Glia | protein-coding | Vasculature | ATPase | Housekeeping | Neuron |
|---|---|---|---|---|---|---|---|
| Immune | U=1520.0<br>p=0.965<br>CLES:50.50% | U=223549.0<br>$p=3.172 \times 10^{-3}$<br>CLES:54.96% | U=3476414.0<br>**$p=7.097 \times 10^{-8}$**<br>CLES:57.62% | U=83996.5<br>**$p=5.641 \times 10^{-5}$**<br>CLES:58.49% | U=30600.0<br>**$p=1.237 \times 10^{-7}$**<br>CLES:66.51% | U=756534.0<br>**$p=6.079 \times 10^{-23}$**<br>CLES:64.78% | U=191525.0<br>**$p=1.128 \times 10^{-19}$**<br>CLES:66.18% |
| MHC | - | U=3717.0<br>p=0.576<br>CLES:56.13% | U=57958.5<br>p=0.409<br>CLES:59.01% | U=1416.0<br>p=0.34<br>CLES:60.56% | U=515.0<br>$p=9.844 \times 10^{-2}$<br>CLES:68.76% | U=12710.0<br>p=0.123<br>CLES:66.85% | U=3222.5<br>$p=9.377 \times 10^{-2}$<br>CLES:68.40% |
| Glia | - | - | U=7063233.0<br>$p=9.324 \times 10^{-4}$<br>CLES:53.21% | U=170287.0<br>$p=3.412 \times 10^{-2}$<br>CLES:53.89% | U=63656.5<br>**$p=1.214 \times 10^{-5}$**<br>CLES:62.89% | U=1567139.5<br>**$p=6.346 \times 10^{-24}$**<br>CLES:60.99% | U=399400.0<br>**$p=2.237 \times 10^{-18}$**<br>CLES:62.73% |
| protein-coding | - | - | - | U=2364208.0<br>p=0.781<br>CLES:50.45% | U=888022.5<br>$p=1.097 \times 10^{-3}$<br>CLES:59.15% | U=21896988.0<br>**$p=7.036 \times 10^{-35}$**<br>CLES:57.46% | U=5561376.5<br>**$p=5.967 \times 10^{-15}$**<br>CLES:58.89% |
| Vasculature | - | - | - | - | U=21150.5<br>$p=4.24 \times 10^{-3}$<br>CLES:59.18% | U=521203.5<br>**$p=8.458 \times 10^{-6}$**<br>CLES:57.46% | U=133246.0<br>**$p=1.592 \times 10^{-6}$**<br>CLES:59.28% |
| ATPase | - | - | - | - | - | U=140837.5<br>p=0.589<br>CLES:48.46% | U=35757.0<br>p=0.909<br>CLES:49.65% |
| Housekeeping | - | - | - | - | - | - | U=936021.0<br>p=0.331<br>CLES:51.21% |

### Chicken, Zeisel et al., 2015, cell subtypes

| MWU Test | Immune | Microglia | MHC | Endothelial | protein-coding | Mural | Oligodendrocyte | Astrocyte | Interneuron | ATPase | Housekeeping | S1 Pyramidal | CA1 Pyramidal |
|---|---|---|---|---|---|---|---|---|---|---|---|---|---|
| Ependymal | U=59393.0<br>p=0.1<br>CLES:53.74% | U=34413.0<br>p=$7.454 \times 10^{-3}$<br>CLES:56.98% | U=969.0<br>p=0.729<br>CLES:53.86% | U=37665.0<br>**p=$8.034 \times 10^{-7}$**<br>CLES:62.90% | U=2227264.5<br>**p=$9.642 \times 10^{-11}$**<br>CLES:61.76% | U=16596.0<br>**p=$4.051 \times 10^{-5}$**<br>CLES:63.94% | U=50205.0<br>**p=$2.56 \times 10^{-10}$**<br>CLES:65.55% | U=26549.0<br>**p=$3.222 \times 10^{-8}$**<br>CLES:66.22% | U=42884.0<br>**p=$1.91 \times 10^{-15}$**<br>CLES:70.71% | U=19499.5<br>**p=$3.245 \times 10^{-10}$**<br>CLES:70.91% | U=482245.0<br>**p=$4.058 \times 10^{-24}$**<br>CLES:69.09% | U=32747.0<br>**p=$5.112 \times 10^{-14}$**<br>CLES:71.18% | U=46717.0<br>**p=$9.536 \times 10^{-16}$**<br>CLES:70.46% |
| Immune | - | U=53230.5<br>p=0.253<br>CLES:52.68% | U=1520.0<br>p=0.965<br>CLES:50.50% | U=58374.5<br>p=$4.377 \times 10^{-4}$<br>CLES:58.26% | U=3476414.0<br>**p=$7.097 \times 10^{-8}$**<br>CLES:57.62% | U=25622.0<br>p=$4.871 \times 10^{-3}$<br>CLES:59.00% | U=77916.0<br>**p=$6.959 \times 10^{-7}$**<br>CLES:60.81% | U=41285.5<br>**p=$1.905 \times 10^{-5}$**<br>CLES:61.55% | U=66973.5<br>**p=$8.195 \times 10^{-12}$**<br>CLES:66.00% | U=30600.0<br>**p=$1.237 \times 10^{-7}$**<br>CLES:66.51% | U=756534.0<br>**p=$6.079 \times 10^{-23}$**<br>CLES:64.78% | U=51165.0<br>**p=$1.452 \times 10^{-10}$**<br>CLES:66.47% | U=73386.5<br>**p=$1.258 \times 10^{-12}$**<br>CLES:66.15% |
| Microglia | - | - | U=768.0<br>p=0.767<br>CLES:46.69% | U=30967.0<br>p=$1.415 \times 10^{-2}$<br>CLES:56.56% | U=1840233.5<br>p=$2.229 \times 10^{-3}$<br>CLES:55.81% | U=13498.0<br>p=$4.588 \times 10^{-2}$<br>CLES:56.87% | U=41531.0<br>p=$2.235 \times 10^{-4}$<br>CLES:59.30% | U=22071.0<br>p=$6.307 \times 10^{-4}$<br>CLES:60.20% | U=36176.0<br>**p=$1.077 \times 10^{-8}$**<br>CLES:65.23% | U=16505.0<br>**p=$3.518 \times 10^{-6}$**<br>CLES:65.64% | U=406464.5<br>**p=$3.175 \times 10^{-12}$**<br>CLES:63.68% | U=27656.0<br>**p=$3.994 \times 10^{-8}$**<br>CLES:65.75% | U=39837.0<br>**p=$1.672 \times 10^{-9}$**<br>CLES:65.71% |
| MHC | - | - | - | U=979.0<br>p=0.368<br>CLES:60.02% | U=57958.5<br>p=0.409<br>CLES:59.01% | U=437.0<br>p=0.3<br>CLES:61.81% | U=1315.0<br>p=0.239<br>CLES:63.04% | U=695.0<br>p=0.224<br>CLES:63.64% | U=1128.0<br>p=$9.996 \times 10^{-2}$<br>CLES:68.28% | U=515.0<br>p=$9.844 \times 10^{-2}$<br>CLES:68.76% | U=12710.0<br>p=0.123<br>CLES:66.85% | U=861.0<br>p=$9.401 \times 10^{-2}$<br>CLES:68.72% | U=1233.5<br>p=$9.908 \times 10^{-2}$<br>CLES:68.30% |
| Endothelial | - | - | - | - | U=1621238.5<br>p=0.829<br>CLES:49.59% | U=11744.0<br>p=0.978<br>CLES:49.90% | U=36319.0<br>p=0.361<br>CLES:52.31% | U=19469.0<br>p=0.234<br>CLES:53.56% | U=32253.0<br>p=$1.185 \times 10^{-3}$<br>CLES:58.65% | U=14709.5<br>p=$7.689 \times 10^{-3}$<br>CLES:59.00% | U=363088.5<br>p=$1.825 \times 10^{-4}$<br>CLES:57.38% | U=24652.0<br>p=$1.524 \times 10^{-3}$<br>CLES:59.11% | U=35732.0<br>p=$3.009 \times 10^{-4}$<br>CLES:59.44% |
| protein-coding | - | - | - | - | - | U=715990.5<br>p=0.857<br>CLES:50.52% | U=2199763.0<br>p=0.123<br>CLES:52.61% | U=1175030.0<br>p=0.113<br>CLES:53.68% | U=1941153.0<br>**p=$5.422 \times 10^{-6}$**<br>CLES:58.62% | U=888022.5<br>p=$1.097 \times 10^{-3}$<br>CLES:59.15% | U=21896988.0<br>**p=$7.036 \times 10^{-35}$**<br>CLES:57.46% | U=1478658.5<br>**p=$4.41 \times 10^{-5}$**<br>CLES:58.87% | U=2141565.0<br>**p=$4.473 \times 10^{-7}$**<br>CLES:59.16% |
| Mural | - | - | - | - | - | - | U=15757.0<br>p=0.48<br>CLES:52.35% | U=8530.0<br>p=0.263<br>CLES:54.14% | U=14124.0<br>p=$7.109 \times 10^{-3}$<br>CLES:59.25% | U=6441.0<br>p=$1.684 \times 10^{-2}$<br>CLES:59.60% | U=158115.0<br>p=$9.027 \times 10^{-3}$<br>CLES:57.64% | U=10821.0<br>p=$6.196 \times 10^{-3}$<br>CLES:59.85% | U=15664.0<br>p=$2.886 \times 10^{-3}$<br>CLES:60.11% |
| Oligodendrocyte | - | - | - | - | - | - | - | U=23827.0<br>p=0.661<br>CLES:51.25% | U=40020.0<br>p=$6.105 \times 10^{-3}$<br>CLES:56.90% | U=18300.0<br>p=$2.328 \times 10^{-2}$<br>CLES:57.39% | U=448490.5<br>p=$2.127 \times 10^{-3}$<br>CLES:55.41% | U=30402.0<br>p=$1.049 \times 10^{-2}$<br>CLES:56.99% | U=44332.0<br>p=$1.823 \times 10^{-3}$<br>CLES:57.66% |
| Astrocyte | - | - | - | - | - | - | - | - | U=20340.0<br>p=$7.857 \times 10^{-2}$<br>CLES:55.25% | U=9352.0<br>p=$9.706 \times 10^{-2}$<br>CLES:56.03% | U=229939.5<br>p=$7.246 \times 10^{-2}$<br>CLES:54.27% | U=15605.0<br>p=$6.323 \times 10^{-2}$<br>CLES:55.88% | U=22684.0<br>p=$3.005 \times 10^{-2}$<br>CLES:56.36% |
| Interneuron | - | - | - | - | - | - | - | - | - | U=12777.0<br>p=0.86<br>CLES:50.60% | U=314815.5<br>p=0.652<br>CLES:49.12% | U=21193.0<br>p=0.954<br>CLES:50.17% | U=31070.0<br>p=0.693<br>CLES:51.03% |
| ATPase | - | - | - | - | - | - | - | - | - | - | U=140837.5<br>p=0.589<br>CLES:48.46% | U=9459.0<br>p=0.863<br>CLES:49.39% | U=13823.0<br>p=0.983<br>CLES:50.07% |
| Housekeeping | - | - | - | - | - | - | - | - | - | - | - | U=247584.0<br>p=0.678<br>CLES:50.93% | U=362276.5<br>p=0.366<br>CLES:51.70% |
| S1 Pyramidal | - | - | - | - | - | - | - | - | - | - | - | - | U=23598.0<br>p=0.696<br>CLES:51.10% |

### Chicken, Saunders et al., 2018, three cell types

| MWU Test | MHC | protein-coding | Vasculature | Glia | ATPase | Housekeeping | Neuron |
|---|---|---|---|---|---|---|---|
| Immune | U=1520.0<br>p=0.965<br>CLES:50.50% | U=3476414.0<br>**p=7.097 × $10^{-8}$**<br>CLES:57.62% | U=79501.0<br>**p=9.93 × $10^{-5}$**<br>CLES:58.32% | U=65437.0<br>**p=4.461 × $10^{-9}$**<br>CLES:63.67% | U=30600.0<br>**p=1.237 × $10^{-7}$**<br>CLES:66.51% | U=756534.0<br>**p=6.079 × $10^{-23}$**<br>CLES:64.78% | U=164091.0<br>**p=3.059 × $10^{-27}$**<br>CLES:70.15% |
| MHC | - | U=57958.5<br>p=0.409<br>CLES:59.01% | U=1320.5<br>p=0.391<br>CLES:59.51% | U=1093.0<br>p=0.168<br>CLES:65.33% | U=515.0<br>p=9.844 × $10^{-2}$<br>CLES:68.76% | U=12710.0<br>p=0.123<br>CLES:66.85% | U=2772.0<br>p=3.82 × $10^{-2}$<br>CLES:72.79% |
| protein-coding | - | - | U=2262104.5<br>p=0.602<br>CLES:50.86% | U=1876430.5<br>p=1.575 × $10^{-3}$<br>CLES:55.95% | U=888022.5<br>p=1.097 × $10^{-3}$<br>CLES:59.15% | U=21896988.0<br>**p=7.036 × $10^{-35}$**<br>CLES:57.46% | U=4842041.0<br>**p=1.784 × $10^{-26}$**<br>CLES:63.43% |
| Vasculature | - | - | - | U=41685.0<br>p=4.256 × $10^{-2}$<br>CLES:55.02% | U=19768.5<br>p=1.039 × $10^{-2}$<br>CLES:58.28% | U=487777.0<br>p=1.032 × $10^{-4}$<br>CLES:56.65% | U=108150.0<br>**p=4.677 × $10^{-10}$**<br>CLES:62.71% |
| Glia | - | - | - | - | U=13754.0<br>p=0.261<br>CLES:53.78% | U=337715.5<br>p=0.298<br>CLES:52.03% | U=76684.0<br>**p=6.176 × $10^{-5}$**<br>CLES:58.98% |
| ATPase | - | - | - | - | - | U=140837.5<br>p=0.589<br>CLES:48.46% | U=31970.5<br>p=0.107<br>CLES:54.92% |
| Housekeeping | - | - | - | - | - | - | U=829263.5<br>**p=6.272 × $10^{-6}$**<br>CLES:56.13% |

### Chicken, Saunders et al., 2018, cell subtypes

| MWU Test | MHC | Endothelia | protein-coding | Microglia | ATPase | Housekeeping | Astrocyte | Oligodendrocyte | Neuron |
|---|---|---|---|---|---|---|---|---|---|
| Immune | U=1520.0<br>p=0.965<br>CLES:50.50% | U=69836.5<br>p=$1.546 \times 10^{-3}$<br>CLES:56.99% | U=3476414.0<br>**p=$7.097 \times 10^{-8}$**<br>CLES:57.62% | U=49307.5<br>p=$9.603 \times 10^{-4}$<br>CLES:58.21% | U=30600.0<br>**p=$1.237 \times 10^{-7}$**<br>CLES:66.51% | U=756534.0<br>**p=$6.079 \times 10^{-23}$**<br>CLES:64.78% | U=65381.5<br>**p=$3.535 \times 10^{-10}$**<br>CLES:64.70% | U=61175.5<br>**p=$4.411 \times 10^{-10}$**<br>CLES:64.96% | U=150006.5<br>**p=$1.11 \times 10^{-25}$**<br>CLES:69.91% |
| MHC | - | U=1149.0<br>p=0.494<br>CLES:57.59% | U=57958.5<br>p=0.409<br>CLES:59.01% | U=818.5<br>p=0.402<br>CLES:59.35% | U=515.0<br>p=$9.844 \times 10^{-2}$<br>CLES:68.76% | U=12710.0<br>p=0.123<br>CLES:66.85% | U=1107.0<br>p=0.12<br>CLES:67.29% | U=1012.0<br>p=0.15<br>CLES:66.01% | U=2531.0<br>p=$4.127 \times 10^{-2}$<br>CLES:72.46% |
| Endothelia | - | - | U=2026163.5<br>p=0.7<br>CLES:50.67% | U=28670.0<br>p=0.691<br>CLES:51.06% | U=18285.5<br>p=$2.37 \times 10^{-3}$<br>CLES:59.96% | U=451147.0<br>**p=$4.075 \times 10^{-6}$**<br>CLES:58.28% | U=38573.0<br>p=$2.864 \times 10^{-3}$<br>CLES:57.59% | U=36392.0<br>p=$1.381 \times 10^{-3}$<br>CLES:58.31% | U=91306.0<br>**p=$3.543 \times 10^{-11}$**<br>CLES:64.20% |
| protein-coding | - | - | - | U=1393775.5<br>p=0.839<br>CLES:50.42% | U=888022.5<br>p=$1.097 \times 10^{-3}$<br>CLES:59.15% | U=21896988.0<br>**p=$7.036 \times 10^{-35}$**<br>CLES:57.46% | U=1874862.5<br>p=$3.051 \times 10^{-4}$<br>CLES:56.86% | U=1766551.5<br>p=$1.401 \times 10^{-4}$<br>CLES:57.49% | U=4424337.5<br>**p=$1.156 \times 10^{-23}$**<br>CLES:63.19% |
| Microglia | - | - | - | - | U=12441.5<br>p=$9.381 \times 10^{-3}$<br>CLES:59.02% | U=306519.5<br>p=$6.242 \times 10^{-4}$<br>CLES:57.29% | U=26333.0<br>p=$1.373 \times 10^{-2}$<br>CLES:56.88% | U=24787.0<br>p=$8.647 \times 10^{-3}$<br>CLES:57.45% | U=62399.0<br>**p=$2.96 \times 10^{-8}$**<br>CLES:63.48% |
| ATPase | - | - | - | - | - | U=140837.5<br>p=0.589<br>CLES:48.46% | U=11747.0<br>p=0.33<br>CLES:46.72% | U=11265.5<br>p=0.573<br>CLES:48.08% | U=29141.5<br>p=0.137<br>CLES:54.58% |
| Housekeeping | - | - | - | - | - | - | U=310744.5<br>p=0.503<br>CLES:48.69% | U=296105.5<br>p=0.914<br>CLES:49.78% | U=756935.0<br>**p=$3.175 \times 10^{-5}$**<br>CLES:55.85% |
| Astrocyte | - | - | - | - | - | - | - | U=26606.0<br>p=0.532<br>CLES:51.70% | U=68665.0<br>p=$1.816 \times 10^{-4}$<br>CLES:58.56% |
| Oligodendrocyte | - | - | - | - | - | - | - | - | U=61685.0<br>p=$5.909 \times 10^{-3}$<br>CLES:56.45% |

### Chicken, Zeisel et al., 2018, three cell types

| MWU Test | MHC | CNS_Glia | CNS_Endothelia | protein-coding | ATPase | Housekeeping | CNS_Neuron |
|---|---|---|---|---|---|---|---|
| Immune | U=1520.0<br>p=0.965<br>CLES:50.50% | U=148640.5<br>p=0.122<br>CLES:52.77% | U=27868.5<br>$p=3.613 \times 10^{-2}$<br>CLES:56.36% | U=3476414.0<br>**$p=7.097 \times 10^{-8}$**<br>CLES:57.62% | U=30600.0<br>**$p=1.237 \times 10^{-7}$**<br>CLES:66.51% | U=756534.0<br>**$p=6.079 \times 10^{-23}$**<br>CLES:64.78% | U=788690.5<br>**$p=1.053 \times 10^{-24}$**<br>CLES:65.34% |
| MHC | - | U=2454.0<br>p=0.749<br>CLES:53.52% | U=470.0<br>p=0.461<br>CLES:58.39% | U=57958.5<br>p=0.409<br>CLES:59.01% | U=515.0<br>$p=9.844 \times 10^{-2}$<br>CLES:68.76% | U=12710.0<br>p=0.123<br>CLES:66.85% | U=13265.5<br>p=0.109<br>CLES:67.51% |
| CNS_Glia | - | - | U=40814.0<br>p=0.152<br>CLES:54.18% | U=5093076.5<br>**$p=2.713 \times 10^{-6}$**<br>CLES:55.41% | U=45591.5<br>**$p=5.824 \times 10^{-7}$**<br>CLES:65.05% | U=1123152.5<br>**$p=1.453 \times 10^{-25}$**<br>CLES:63.13% | U=1174502.0<br>**$p=1.578 \times 10^{-28}$**<br>CLES:63.88% |
| CNS_Endothelia | - | - | - | U=836291.5<br>p=0.5<br>CLES:51.83% | U=7622.5<br>$p=2.119 \times 10^{-3}$<br>CLES:61.95% | U=186987.0<br>$p=3.315 \times 10^{-4}$<br>CLES:59.87% | U=196191.0<br>**$p=8.74 \times 10^{-5}$**<br>CLES:60.78% |
| protein-coding | - | - | - | - | U=888022.5<br>$p=1.097 \times 10^{-3}$<br>CLES:59.15% | U=21896988.0<br>**$p=7.036 \times 10^{-35}$**<br>CLES:57.46% | U=22851219.5<br>**$p=4.069 \times 10^{-41}$**<br>CLES:58.02% |
| ATPase | - | - | - | - | - | U=140837.5<br>p=0.589<br>CLES:48.46% | U=146746.5<br>p=0.688<br>CLES:48.86% |
| Housekeeping | - | - | - | - | - | - | U=3841366.5<br>p=0.619<br>CLES:50.39% |

### Chicken, Zeisel et al., 2018, cell subtypes

| MWU Test | Nervous Sys Immune cells | Immune (benchmark) | MHC | protein-coding | Vascular cells | Neurons | ATPase | Housekeeping |
|---|---|---|---|---|---|---|---|---|
| Glia | U=16966.0<br>p=0.423<br>CLES:53.15% | U=119723.5<br>p=0.212<br>CLES:52.34% | U=1945.0<br>p=0.84<br>CLES:52.23% | U=4521814.0<br>**p=1.113 × $10^{-16}$**<br>CLES:60.57% | U=44574.0<br>**p=1.617 × $10^{-6}$**<br>CLES:63.47% | U=1031336.0<br>**p=8.367 × $10^{-35}$**<br>CLES:66.76% | U=39842.0<br>**p=6.536 × $10^{-11}$**<br>CLES:69.99% | U=983727.5<br>**p=7.734 × $10^{-40}$**<br>CLES:68.08% |
| Nervous Sys Immune cells | - | U=12809.5<br>p=0.93<br>CLES:49.65% | U=212.0<br>p=0.975<br>CLES:50.48% | U=491560.0<br>p=2.476 × $10^{-2}$<br>CLES:58.39% | U=4889.0<br>p=9.28 × $10^{-3}$<br>CLES:61.73% | U=113068.0<br>**p=7.667 × $10^{-5}$**<br>CLES:64.89% | U=4397.0<br>**p=7.569 × $10^{-5}$**<br>CLES:68.49% | U=108043.0<br>**p=1.522 × $10^{-5}$**<br>CLES:66.30% |
| Immune (benchmark) | - | - | U=1520.0<br>p=0.965<br>CLES:50.50% | U=3476414.0<br>**p=7.097 × $10^{-8}$**<br>CLES:57.62% | U=34059.5<br>p=5.011 × $10^{-4}$<br>CLES:60.01% | U=790551.0<br>**p=4.618 × $10^{-19}$**<br>CLES:63.31% | U=30600.0<br>**p=1.237 × $10^{-7}$**<br>CLES:66.51% | U=756534.0<br>**p=6.079 × $10^{-23}$**<br>CLES:64.78% |
| MHC | - | - | - | U=57958.5<br>p=0.409<br>CLES:59.01% | U=577.0<br>p=0.27<br>CLES:62.45% | U=13292.0<br>p=0.159<br>CLES:65.39% | U=515.0<br>p=9.844 × $10^{-2}$<br>CLES:68.76% | U=12710.0<br>p=0.123<br>CLES:66.85% |
| protein-coding | - | - | - | - | U=969687.0<br>p=0.351<br>CLES:52.35% | U=22709819.0<br>**p=2.077 × $10^{-22}$**<br>CLES:55.73% | U=888022.5<br>p=1.097 × $10^{-3}$<br>CLES:59.15% | U=21896988.0<br>**p=7.036 × $10^{-35}$**<br>CLES:57.46% |
| Vascular cells | - | - | - | - | - | U=205252.0<br>p=0.168<br>CLES:53.54% | U=8069.5<br>p=5.813 × $10^{-2}$<br>CLES:57.13% | U=198661.0<br>p=3.544 × $10^{-2}$<br>CLES:55.41% |
| Neurons | - | - | - | - | - | - | U=166731.5<br>p=0.198<br>CLES:53.66% | U=4102385.0<br>p=9.006 × $10^{-3}$<br>CLES:52.01% |
| ATPase | - | - | - | - | - | - | - | U=140837.5<br>p=0.589<br>CLES:48.46% |

### Mouse, Zhang et al., 2014, three cell types

| MWU Test | Immune | protein-coding | Glia | Endothelia | Housekeeping | Neuron | ATPase |
|---|---|---|---|---|---|---|---|
| MHC | U=15488.0<br>**p=6.025 × $10^{-9}$**<br>CLES:83.55% | U=444853.0<br>**p=4.491 × $10^{-11}$**<br>CLES:87.32% | U=28113.5<br>**p=1.251 × $10^{-11}$**<br>CLES:88.78% | U=21783.5<br>**p=1.189 × $10^{-11}$**<br>CLES:88.94% | U=81185.0<br>**p=4.838 × $10^{-16}$**<br>CLES:96.14% | U=32231.5<br>**p=2.43 × $10^{-15}$**<br>CLES:95.29% | U=3221.0<br>**p=6.037 × $10^{-14}$**<br>CLES:96.78% |
| Immune | - | U=7509472.0<br>p=6.574 × $10^{-4}$<br>CLES:53.75% | U=466825.0<br>p=5.822 × $10^{-3}$<br>CLES:53.75% | U=362451.0<br>p=5.678 × $10^{-3}$<br>CLES:53.96% | U=1528126.0<br>**p=7.056 × $10^{-41}$**<br>CLES:65.99% | U=625478.5<br>**p=2.231 × $10^{-38}$**<br>CLES:67.43% | U=63162.0<br>**p=4.294 × $10^{-12}$**<br>CLES:69.21% |
| protein-coding | - | - | U=11896177.5<br>p=0.855<br>CLES:49.84% | U=9228600.5<br>p=0.997<br>CLES:50.00% | U=40096032.0<br>**p=6.851 × $10^{-125}$**<br>CLES:63.00% | U=16560869.5<br>**p=3.008 × $10^{-73}$**<br>CLES:64.96% | U=1673449.0<br>**p=6.512 × $10^{-11}$**<br>CLES:66.72% |
| Glia | - | - | - | U=575311.0<br>p=0.91<br>CLES:50.14% | U=2525556.5<br>**p=3.478 × $10^{-46}$**<br>CLES:63.84% | U=1046595.5<br>**p=3.639 × $10^{-44}$**<br>CLES:66.05% | U=105559.5<br>**p=4.131 × $10^{-11}$**<br>CLES:67.71% |
| Endothelia | - | - | - | - | U=1949715.5<br>**p=9.115 × $10^{-38}$**<br>CLES:63.72% | U=808108.0<br>**p=4.316 × $10^{-38}$**<br>CLES:65.94% | U=81615.5<br>**p=7.967 × $10^{-11}$**<br>CLES:67.69% |
| Housekeeping | - | - | - | - | - | U=2226800.0<br>p=4.408 × $10^{-3}$<br>CLES:52.70% | U=226816.0<br>p=7.989 × $10^{-2}$<br>CLES:54.56% |
| Neuron | - | - | - | - | - | - | U=86207.0<br>p=0.509<br>CLES:51.77% |

### Mouse, Zhang et al., 2014, cell subtypes

| MWU Test | Immune | Microglia | protein-coding | Endothelia | Astrocyte | Oligodendrocyte | Housekeeping | Neuron | ATPase |
|---|---|---|---|---|---|---|---|---|---|
| MHC | U=15488.0<br>**$p=6.025 \times 10^{-9}$**<br>CLES:83.55% | U=23698.5<br>**$p=7.676 \times 10^{-10}$**<br>CLES:85.26% | U=444853.0<br>**$p=4.491 \times 10^{-11}$**<br>CLES:87.32% | U=21783.5<br>**$p=1.189 \times 10^{-11}$**<br>CLES:88.94% | U=19874.0<br>**$p=1.233 \times 10^{-13}$**<br>CLES:92.65% | U=12970.0<br>**$p=6.963 \times 10^{-15}$**<br>CLES:95.20% | U=81185.0<br>**$p=4.838 \times 10^{-16}$**<br>CLES:96.14% | U=32231.5<br>**$p=2.43 \times 10^{-15}$**<br>CLES:95.29% | U=3221.0<br>**$p=6.037 \times 10^{-14}$**<br>CLES:96.78% |
| Immune | - | U=379623.0<br>$p=0.89$<br>CLES:49.81% | U=7509472.0<br>$p=6.574 \times 10^{-4}$<br>CLES:53.75% | U=362451.0<br>$p=5.678 \times 10^{-3}$<br>CLES:53.96% | U=347228.0<br>**$p=9.638 \times 10^{-10}$**<br>CLES:59.03% | U=240267.5<br>**$p=7.238 \times 10^{-18}$**<br>CLES:64.31% | U=1528126.0<br>**$p=7.056 \times 10^{-41}$**<br>CLES:65.99% | U=625478.5<br>**$p=2.231 \times 10^{-38}$**<br>CLES:67.43% | U=63162.0<br>**$p=4.294 \times 10^{-12}$**<br>CLES:69.21% |
| Microglia | - | - | U=11368035.0<br>**$p=2.481 \times 10^{-6}$**<br>CLES:54.27% | U=549070.0<br>$p=4.53 \times 10^{-4}$<br>CLES:54.53% | U=529749.0<br>**$p=5.337 \times 10^{-14}$**<br>CLES:60.07% | U=370192.0<br>**$p=1.516 \times 10^{-25}$**<br>CLES:66.09% | U=2349037.5<br>**$p=2.267 \times 10^{-67}$**<br>CLES:67.65% | U=966460.0<br>**$p=4.168 \times 10^{-60}$**<br>CLES:69.49% | U=97422.5<br>**$p=4.229 \times 10^{-15}$**<br>CLES:71.20% |
| protein-coding | - | - | - | U=9228600.5<br>$p=0.997$<br>CLES:50.00% | U=8956964.0<br>**$p=1.367 \times 10^{-7}$**<br>CLES:55.41% | U=6294244.5<br>**$p=9.269 \times 10^{-19}$**<br>CLES:61.30% | U=40096032.0<br>**$p=6.851 \times 10^{-125}$**<br>CLES:63.00% | U=16560869.5<br>**$p=3.008 \times 10^{-73}$**<br>CLES:64.96% | U=1673449.0<br>**$p=6.512 \times 10^{-11}$**<br>CLES:66.72% |
| Endothelia | - | - | - | - | U=433364.0<br>**$p=2.843 \times 10^{-5}$**<br>CLES:55.76% | U=306272.0<br>**$p=1.928 \times 10^{-14}$**<br>CLES:62.05% | U=1949715.5<br>**$p=9.115 \times 10^{-38}$**<br>CLES:63.72% | U=808108.0<br>**$p=4.316 \times 10^{-38}$**<br>CLES:65.94% | U=81615.5<br>**$p=7.967 \times 10^{-11}$**<br>CLES:67.69% |
| Astrocyte | - | - | - | - | - | U=244037.0<br>**$p=6.368 \times 10^{-5}$**<br>CLES:56.45% | U=1562499.0<br>**$p=1.542 \times 10^{-13}$**<br>CLES:58.31% | U=652851.0<br>**$p=3.648 \times 10^{-17}$**<br>CLES:60.83% | U=66032.5<br>**$p=4.953 \times 10^{-6}$**<br>CLES:62.53% |
| Oligodendrocyte | - | - | - | - | - | - | U=885563.5<br>$p=0.135$<br>CLES:52.03% | U=374299.5<br>$p=1.027 \times 10^{-3}$<br>CLES:54.90% | U=37991.0<br>$p=1.972 \times 10^{-2}$<br>CLES:56.64% |
| Housekeeping | - | - | - | - | - | - | - | U=2226800.0<br>$p=4.408 \times 10^{-3}$<br>CLES:52.70% | U=226816.0<br>$p=7.989 \times 10^{-2}$<br>CLES:54.56% |
| Neuron | - | - | - | - | - | - | - | - | U=86207.0<br>$p=0.509$<br>CLES:51.77% |

### Mouse, Zeisel et al., 2015, three cell types

| MWU Test | Immune | Glia | protein-coding | Vasculature | Housekeeping | Neuron | ATPase |
|---|---|---|---|---|---|---|---|
| MHC | U=15488.0<br>**$p=6.025 \times 10^{-9}$**<br>CLES:83.55% | U=32459.0<br>**$p=2.379 \times 10^{-12}$**<br>CLES:90.07% | U=444853.0<br>**$p=4.491 \times 10^{-11}$**<br>CLES:87.32% | U=10863.5<br>**$p=3.307 \times 10^{-13}$**<br>CLES:92.44% | U=81185.0<br>**$p=4.838 \times 10^{-16}$**<br>CLES:96.14% | U=22276.5<br>**$p=2.352 \times 10^{-15}$**<br>CLES:95.52% | U=3221.0<br>**$p=6.037 \times 10^{-14}$**<br>CLES:96.78% |
| Immune | - | U=532783.5<br>$p=3.273 \times 10^{-3}$<br>CLES:53.91% | U=7509472.0<br>$p=6.574 \times 10^{-4}$<br>CLES:53.75% | U=190992.5<br>**$p=9.557 \times 10^{-8}$**<br>CLES:59.26% | U=1528126.0<br>**$p=7.056 \times 10^{-41}$**<br>CLES:65.99% | U=429534.5<br>**$p=2.295 \times 10^{-32}$**<br>CLES:67.16% | U=63162.0<br>**$p=4.294 \times 10^{-12}$**<br>CLES:69.21% |
| Glia | - | - | U=13676802.0<br>p=0.655<br>CLES:50.36% | U=353535.0<br>**$p=3.909 \times 10^{-5}$**<br>CLES:56.43% | U=2893645.5<br>**$p=1.301 \times 10^{-53}$**<br>CLES:64.28% | U=820156.5<br>**$p=4.147 \times 10^{-38}$**<br>CLES:65.97% | U=120871.5<br>**$p=1.069 \times 10^{-11}$**<br>CLES:68.13% |
| protein-coding | - | - | - | U=4927969.5<br>**$p=4.021 \times 10^{-5}$**<br>CLES:55.64% | U=40096032.0<br>**$p=6.851 \times 10^{-125}$**<br>CLES:63.00% | U=11353329.5<br>**$p=1.365 \times 10^{-49}$**<br>CLES:64.59% | U=1673449.0<br>**$p=6.512 \times 10^{-11}$**<br>CLES:66.72% |
| Vasculature | - | - | - | - | U=848956.0<br>**$p=6.66 \times 10^{-8}$**<br>CLES:57.83% | U=242414.0<br>**$p=4.178 \times 10^{-9}$**<br>CLES:59.79% | U=35873.5<br>**$p=3.331 \times 10^{-5}$**<br>CLES:62.00% |
| Housekeeping | - | - | - | - | - | U=1517920.0<br>$p=5.377 \times 10^{-2}$<br>CLES:52.10% | U=226816.0<br>$p=7.989 \times 10^{-2}$<br>CLES:54.56% |
| Neuron | - | - | - | - | - | - | U=60222.0<br>p=0.369<br>CLES:52.45% |

### Mouse, Zeisel et al., 2015, cell subtypes

| MWU Test | Microglia | Ependymal | Immune | protein-coding | Endothelial | Astrocyte | Oligodendrocyte | Mural | Housekeeping | S1 Pyramidal | Interneuron | CA1 Pyramidal | ATPase |
|---|---|---|---|---|---|---|---|---|---|---|---|---|---|
| MHC | U=8349.0<br>**$p=4.892 \times 10^{-9}$**<br>CLES:84.28% | U=9331.0<br>**$p=1.456 \times 10^{-11}$**<br>CLES:89.50% | U=15488.0<br>**$p=6.025 \times 10^{-9}$**<br>CLES:83.55% | U=444853.0<br>**$p=4.491 \times 10^{-11}$**<br>CLES:87.32% | U=7487.5<br>**$p=3.109 \times 10^{-12}$**<br>CLES:91.13% | U=5167.0<br>**$p=2.025 \times 10^{-13}$**<br>CLES:94.19% | U=9612.0<br>**$p=5.113 \times 10^{-14}$**<br>CLES:94.07% | U=3376.0<br>**$p=2.22 \times 10^{-13}$**<br>CLES:95.48% | U=81185.0<br>**$p=4.838 \times 10^{-16}$**<br>CLES:96.14% | U=5892.0<br>**$p=8.095 \times 10^{-15}$**<br>CLES:96.43% | U=7767.0<br>**$p=4.37 \times 10^{-14}$**<br>CLES:94.54% | U=8617.5<br>**$p=6.733 \times 10^{-15}$**<br>CLES:95.79% | U=3221.0<br>**$p=6.037 \times 10^{-14}$**<br>CLES:96.78% |
| Microglia | - | U=80811.5<br>p=0.161<br>CLES:52.89% | U=142173.0<br>p=0.202<br>CLES:52.34% | U=4234494.0<br>**$p=6.804 \times 10^{-6}$**<br>CLES:56.72% | U=73846.0<br>**$p=2.507 \times 10^{-7}$**<br>CLES:61.34% | U=51255.0<br>**$p=2.881 \times 10^{-8}$**<br>CLES:63.76% | U=95438.5<br>**$p=3.709 \times 10^{-11}$**<br>CLES:63.74% | U=34344.0<br>**$p=1.697 \times 10^{-8}$**<br>CLES:66.28% | U=868005.5<br>**$p=5.602 \times 10^{-38}$**<br>CLES:70.14% | U=64369.0<br>**$p=6.514 \times 10^{-20}$**<br>CLES:71.89% | U=84981.5<br>**$p=7.6 \times 10^{-21}$**<br>CLES:70.58% | U=95031.0<br>**$p=7.327 \times 10^{-25}$**<br>CLES:72.09% | U=35852.0<br>**$p=1.647 \times 10^{-15}$**<br>CLES:73.52% |
| Ependymal | - | - | U=141004.5<br>p=0.705<br>CLES:49.32% | U=4266448.0<br>$p=3.17 \times 10^{-3}$<br>CLES:54.30% | U=74821.0<br>**$p=3.143 \times 10^{-5}$**<br>CLES:59.05% | U=52519.0<br>**$p=8.993 \times 10^{-7}$**<br>CLES:62.07% | U=97379.5<br>**$p=8.872 \times 10^{-9}$**<br>CLES:61.79% | U=35201.0<br>**$p=3.915 \times 10^{-7}$**<br>CLES:64.55% | U=892649.5<br>**$p=7.361 \times 10^{-34}$**<br>CLES:68.54% | U=66244.0<br>**$p=1.216 \times 10^{-17}$**<br>CLES:70.30% | U=87420.5<br>**$p=2.359 \times 10^{-18}$**<br>CLES:68.99% | U=97651.0<br>**$p=6.879 \times 10^{-22}$**<br>CLES:70.38% | U=36993.5<br>**$p=5.3 \times 10^{-14}$**<br>CLES:72.07% |
| Immune | - | - | - | U=7509472.0<br>$p=6.574 \times 10^{-4}$<br>CLES:53.75% | U=130815.5<br>**$p=3.632 \times 10^{-5}$**<br>CLES:58.06% | U=90189.0<br>**$p=1.107 \times 10^{-5}$**<br>CLES:59.95% | U=168206.0<br>**$p=3.256 \times 10^{-8}$**<br>CLES:60.03% | U=60177.0<br>**$p=8.141 \times 10^{-6}$**<br>CLES:62.06% | U=1528126.0<br>**$p=7.056 \times 10^{-41}$**<br>CLES:65.99% | U=112855.5<br>**$p=1.372 \times 10^{-15}$**<br>CLES:67.35% | U=149603.0<br>**$p=4.391 \times 10^{-17}$**<br>CLES:66.40% | U=167076.0<br>**$p=7.464 \times 10^{-21}$**<br>CLES:67.72% | U=63162.0<br>**$p=4.294 \times 10^{-12}$**<br>CLES:69.21% |
| protein-coding | - | - | - | - | U=3368384.5<br>$p=7.208 \times 10^{-3}$<br>CLES:54.40% | U=2323738.5<br>$p=1.906 \times 10^{-3}$<br>CLES:56.20% | U=4335781.5<br>**$p=1.824 \times 10^{-5}$**<br>CLES:56.30% | U=1559585.0<br>$p=6.013 \times 10^{-4}$<br>CLES:58.52% | U=40096032.0<br>**$p=6.851 \times 10^{-125}$**<br>CLES:63.00% | U=2980982.5<br>**$p=7.328 \times 10^{-15}$**<br>CLES:64.74% | U=3951508.0<br>**$p=3.177 \times 10^{-17}$**<br>CLES:63.82% | U=4420839.0<br>**$p=2.677 \times 10^{-22}$**<br>CLES:65.21% | U=1673449.0<br>**$p=6.512 \times 10^{-11}$**<br>CLES:66.72% |
| Endothelial | - | - | - | - | - | U=34403.0<br>p=0.534<br>CLES:51.60% | U=64244.0<br>p=0.428<br>CLES:51.73% | U=23201.0<br>p=0.179<br>CLES:53.99% | U=603075.0<br>**$p=2.635 \times 10^{-7}$**<br>CLES:58.76% | U=44979.0<br>**$p=2.171 \times 10^{-5}$**<br>CLES:60.57% | U=59701.0<br>**$p=2.062 \times 10^{-5}$**<br>CLES:59.79% | U=66896.0<br>**$p=6.526 \times 10^{-7}$**<br>CLES:61.18% | U=25365.5<br>**$p=2.697 \times 10^{-5}$**<br>CLES:62.71% |
| Astrocyte | - | - | - | - | - | - | U=41599.0<br>p=0.947<br>CLES:50.17% | U=15115.0<br>p=0.401<br>CLES:52.67% | U=397077.0<br>$p=1.085 \times 10^{-4}$<br>CLES:57.94% | U=29829.0<br>$p=2.11 \times 10^{-4}$<br>CLES:60.16% | U=39570.0<br>$p=2.746 \times 10^{-4}$<br>CLES:59.35% | U=44459.0<br>**$p=1.576 \times 10^{-5}$**<br>CLES:60.90% | U=16855.0<br>$p=1.281 \times 10^{-4}$<br>CLES:62.41% |
| Oligodendrocyte | - | - | - | - | - | - | - | U=27991.0<br>p=0.41<br>CLES:52.37% | U=735913.5<br>**$p=6.949 \times 10^{-7}$**<br>CLES:57.65% | U=55181.0<br>**$p=4.279 \times 10^{-5}$**<br>CLES:59.75% | U=73145.5<br>**$p=4.565 \times 10^{-5}$**<br>CLES:58.90% | U=82275.0<br>**$p=8.086 \times 10^{-7}$**<br>CLES:60.51% | U=31171.0<br>**$p=4.73 \times 10^{-5}$**<br>CLES:61.97% |
| Mural | - | - | - | - | - | - | - | - | U=245881.0<br>$p=2.502 \times 10^{-2}$<br>CLES:55.66% | U=18524.0<br>$p=1.061 \times 10^{-2}$<br>CLES:57.96% | U=24612.0<br>$p=1.42 \times 10^{-2}$<br>CLES:57.27% | U=27702.0<br>$p=2.425 \times 10^{-3}$<br>CLES:58.87% | U=10508.0<br>$p=3.629 \times 10^{-3}$<br>CLES:60.36% |
| Housekeeping | - | - | - | - | - | - | - | - | - | U=396037.5<br>p=0.333<br>CLES:51.89% | U=528191.5<br>p=0.39<br>CLES:51.46% | U=593691.0<br>$p=8.321 \times 10^{-2}$<br>CLES:52.83% | U=226816.0<br>$p=7.989 \times 10^{-2}$<br>CLES:54.56% |
| S1 Pyramidal | - | - | - | - | - | - | - | - | - | - | U=36828.0<br>p=0.87<br>CLES:49.59% | U=41511.0<br>p=0.667<br>CLES:51.05% | U=15877.0<br>p=0.381<br>CLES:52.78% |
| Interneuron | - | - | - | - | - | - | - | - | - | - | - | U=56099.0<br>p=0.561<br>CLES:51.31% | U=21432.0<br>p=0.324<br>CLES:52.99% |
| CA1 Pyramidal | - | - | - | - | - | - | - | - | - | - | - | - | U=22913.0<br>p=0.562<br>CLES:51.74% |

### Mouse, Saunders et al., 2018, three cell types

| MWU Test | Immune | protein-coding | Vasculature | Glia | Housekeeping | ATPase | Neuron |
|---|---|---|---|---|---|---|---|
| MHC | U=15488.0<br>**$p=6.025 \times 10^{-9}$**<br>CLES:83.55% | U=444853.0<br>**$p=4.491 \times 10^{-11}$**<br>CLES:87.32% | U=10170.5<br>**$p=3.71 \times 10^{-12}$**<br>CLES:90.55% | U=7602.0<br>**$p=5.451 \times 10^{-15}$**<br>CLES:96.18% | U=81185.0<br>**$p=4.838 \times 10^{-16}$**<br>CLES:96.14% | U=3221.0<br>**$p=6.037 \times 10^{-14}$**<br>CLES:96.78% | U=17115.0<br>**$p=1.865 \times 10^{-16}$**<br>CLES:97.52% |
| Immune | - | U=7509472.0<br>$p=6.574 \times 10^{-4}$<br>CLES:53.75% | U=178318.0<br>**$p=7.389 \times 10^{-6}$**<br>CLES:57.89% | U=142009.5<br>**$p=4.387 \times 10^{-15}$**<br>CLES:65.52% | U=1528126.0<br>**$p=7.056 \times 10^{-41}$**<br>CLES:65.99% | U=63162.0<br>**$p=4.294 \times 10^{-12}$**<br>CLES:69.21% | U=358771.5<br>**$p=2.006 \times 10^{-56}$**<br>CLES:74.55% |
| protein-coding | - | - | U=4602005.5<br>$p=1.88 \times 10^{-3}$<br>CLES:54.36% | U=3728701.5<br>**$p=4.399 \times 10^{-14}$**<br>CLES:62.59% | U=40096032.0<br>**$p=6.851 \times 10^{-125}$**<br>CLES:63.00% | U=1673449.0<br>**$p=6.512 \times 10^{-11}$**<br>CLES:66.72% | U=9644544.5<br>**$p=1.969 \times 10^{-91}$**<br>CLES:72.92% |
| Vasculature | - | - | - | U=76651.0<br>$p=1.095 \times 10^{-4}$<br>CLES:58.37% | U=825800.5<br>**$p=2.123 \times 10^{-9}$**<br>CLES:58.85% | U=34831.5<br>**$p=7.913 \times 10^{-6}$**<br>CLES:62.99% | U=203302.0<br>**$p=1.539 \times 10^{-28}$**<br>CLES:69.72% |
| Glia | - | - | - | - | U=502745.5<br>p=0.597<br>CLES:50.92% | U=21660.0<br>$p=6.294 \times 10^{-2}$<br>CLES:55.66% | U=131156.0<br>**$p=3.047 \times 10^{-12}$**<br>CLES:63.92% |
| Housekeeping | - | - | - | - | - | U=226816.0<br>$p=7.989 \times 10^{-2}$<br>CLES:54.56% | U=1360605.0<br>**$p=5.346 \times 10^{-23}$**<br>CLES:62.06% |
| ATPase | - | - | - | - | - | - | U=49662.0<br>$p=7.24 \times 10^{-3}$<br>CLES:57.48% |

### Mouse, Saunders et al., 2018, cell subtypes

| MWU Test | Immune | Microglia | protein-coding | Endothelia | Astrocyte | Housekeeping | Oligodendrocyte | ATPase | Neuron |
|---|---|---|---|---|---|---|---|---|---|
| MHC | U=15488.0<br>**$p=6.025 \times 10^{-9}$**<br>CLES:83.55% | U=6336.5<br>**$p=1.077 \times 10^{-10}$**<br>CLES:88.30% | U=444853.0<br>**$p=4.491 \times 10^{-11}$**<br>CLES:87.32% | U=8976.5<br>**$p=5.495 \times 10^{-12}$**<br>CLES:90.38% | U=7041.0<br>**$p=1.257 \times 10^{-14}$**<br>CLES:95.69% | U=81185.0<br>**$p=4.838 \times 10^{-16}$**<br>CLES:96.14% | U=6794.0<br>**$p=5.344 \times 10^{-15}$**<br>CLES:96.42% | U=3221.0<br>**$p=6.037 \times 10^{-14}$**<br>CLES:96.78% | U=15796.0<br>**$p=2.092 \times 10^{-16}$**<br>CLES:97.52% |
| Immune | - | U=104548.0<br>p=0.127<br>CLES:53.13% | U=7509472.0<br>$p=6.574 \times 10^{-4}$<br>CLES:53.75% | U=155617.0<br>**$p=9.765 \times 10^{-5}$**<br>CLES:57.14% | U=132212.5<br>**$p=2.01 \times 10^{-14}$**<br>CLES:65.52% | U=1528126.0<br>**$p=7.056 \times 10^{-41}$**<br>CLES:65.99% | U=128850.0<br>**$p=5.731 \times 10^{-16}$**<br>CLES:66.68% | U=63162.0<br>**$p=4.294 \times 10^{-12}$**<br>CLES:69.21% | U=329825.5<br>**$p=6.28 \times 10^{-53}$**<br>CLES:74.25% |
| Microglia | - | - | U=2750945.5<br>p=0.621<br>CLES:50.87% | U=57532.0<br>$p=4.536 \times 10^{-2}$<br>CLES:54.57% | U=50200.0<br>**$p=5.29 \times 10^{-9}$**<br>CLES:64.27% | U=578965.5<br>**$p=7.834 \times 10^{-16}$**<br>CLES:64.58% | U=49165.0<br>**$p=1.932 \times 10^{-10}$**<br>CLES:65.73% | U=24200.5<br>**$p=2.154 \times 10^{-9}$**<br>CLES:68.50% | U=128432.0<br>**$p=2.971 \times 10^{-32}$**<br>CLES:74.69% |
| protein-coding | - | - | - | U=4006025.5<br>$p=1.831 \times 10^{-2}$<br>CLES:53.52% | U=3474563.5<br>**$p=2.426 \times 10^{-13}$**<br>CLES:62.66% | U=40096032.0<br>**$p=6.851 \times 10^{-125}$**<br>CLES:63.00% | U=3395414.0<br>**$p=2.896 \times 10^{-15}$**<br>CLES:63.94% | U=1673449.0<br>**$p=6.512 \times 10^{-11}$**<br>CLES:66.72% | U=8858843.5<br>**$p=3.083 \times 10^{-82}$**<br>CLES:72.57% |
| Endothelia | - | - | - | - | U=64381.0<br>**$p=2.484 \times 10^{-5}$**<br>CLES:59.55% | U=743694.0<br>**$p=1.954 \times 10^{-10}$**<br>CLES:59.94% | U=63115.0<br>**$p=1.754 \times 10^{-6}$**<br>CLES:60.97% | U=31345.5<br>**$p=1.755 \times 10^{-6}$**<br>CLES:64.11% | U=168139.0<br>**$p=3.708 \times 10^{-28}$**<br>CLES:70.65% |
| Astrocyte | - | - | - | - | - | U=464781.5<br>p=0.752<br>CLES:50.56% | U=39532.0<br>p=0.529<br>CLES:51.55% | U=19965.0<br>$p=9.669 \times 10^{-2}$<br>CLES:55.12% | U=110604.0<br>**$p=7.768 \times 10^{-10}$**<br>CLES:62.73% |
| Housekeeping | - | - | - | - | - | - | U=447024.5<br>p=0.667<br>CLES:50.79% | U=226816.0<br>$p=7.989 \times 10^{-2}$<br>CLES:54.56% | U=1246393.0<br>**$p=4.201 \times 10^{-20}$**<br>CLES:61.60% |
| Oligodendrocyte | - | - | - | - | - | - | - | U=18712.5<br>p=0.203<br>CLES:53.95% | U=103930.0<br>**$p=3.822 \times 10^{-8}$**<br>CLES:61.56% |
| ATPase | - | - | - | - | - | - | - | - | U=45454.0<br>$p=1.253 \times 10^{-2}$<br>CLES:57.00% |

### Mouse, Zeisel et al., 2018, three cell types

| MWU Test | Immune | CNS_Glia | protein-coding | CNS_Endothelia | Housekeeping | CNS_Neuron | ATPase |
|---|---|---|---|---|---|---|---|
| MHC | U=15488.0<br>**$p=6.025 \times 10^{-9}$**<br>CLES:83.55% | U=22374.0<br>**$p=2.514 \times 10^{-12}$**<br>CLES:90.20% | U=444853.0<br>**$p=4.491 \times 10^{-11}$**<br>CLES:87.32% | U=4048.0<br>**$p=8.802 \times 10^{-11}$**<br>CLES:89.48% | U=81185.0<br>**$p=4.838 \times 10^{-16}$**<br>CLES:96.14% | U=89218.0<br>**$p=6.103 \times 10^{-16}$**<br>CLES:95.96% | U=3221.0<br>**$p=6.037 \times 10^{-14}$**<br>CLES:96.78% |
| Immune | - | U=359699.5<br>$p=4.385 \times 10^{-2}$<br>CLES:52.88% | U=7509472.0<br>$p=6.574 \times 10^{-4}$<br>CLES:53.75% | U=66449.0<br>p=0.145<br>CLES:53.56% | U=1528126.0<br>**$p=7.056 \times 10^{-41}$**<br>CLES:65.99% | U=1708674.0<br>**$p=8.057 \times 10^{-47}$**<br>CLES:67.02% | U=63162.0<br>**$p=4.294 \times 10^{-12}$**<br>CLES:69.21% |
| CNS_Glia | - | - | U=9642767.5<br>$p=9.811 \times 10^{-2}$<br>CLES:51.58% | U=85338.0<br>p=0.554<br>CLES:51.41% | U=2045678.5<br>**$p=2.645 \times 10^{-51}$**<br>CLES:66.02% | U=2307337.5<br>**$p=4.653 \times 10^{-63}$**<br>CLES:67.63% | U=85493.5<br>**$p=1.823 \times 10^{-13}$**<br>CLES:70.01% |
| protein-coding | - | - | - | U=1688790.5<br>p=0.831<br>CLES:49.53% | U=40096032.0<br>**$p=6.851 \times 10^{-125}$**<br>CLES:63.00% | U=45085316.0<br>**$p=2.458 \times 10^{-164}$**<br>CLES:64.34% | U=1673449.0<br>**$p=6.512 \times 10^{-11}$**<br>CLES:66.72% |
| CNS_Endothelia | - | - | - | - | U=365736.5<br>**$p=5.767 \times 10^{-11}$**<br>CLES:64.71% | U=413471.0<br>**$p=2.144 \times 10^{-13}$**<br>CLES:66.45% | U=15338.5<br>**$p=2.103 \times 10^{-8}$**<br>CLES:68.87% |
| Housekeeping | - | - | - | - | - | U=5989824.0<br>$p=2.482 \times 10^{-2}$<br>CLES:51.57% | U=226816.0<br>$p=7.989 \times 10^{-2}$<br>CLES:54.56% |
| CNS_Neuron | - | - | - | - | - | - | U=242763.0<br>p=0.242<br>CLES:53.04% |

### Mouse, Zeisel et al., 2018, cell subtypes

| MWU Test | Nervous Sys Immune cells | Glia | Immune (benchmark) | protein-coding | Vascular cells | Neurons | Housekeeping | ATPase |
|---|---|---|---|---|---|---|---|---|
| MHC | U=2668.0<br>**p=$8.121 \times 10^{-6}$**<br>CLES:77.74% | U=18202.5<br>**p=$8.826 \times 10^{-12}$**<br>CLES:89.30% | U=15488.0<br>**p=$6.025 \times 10^{-9}$**<br>CLES:83.55% | U=444853.0<br>**p=$4.491 \times 10^{-11}$**<br>CLES:87.32% | U=4689.0<br>**p=$1.175 \times 10^{-12}$**<br>CLES:92.96% | U=93292.5<br>**p=$1.982 \times 10^{-15}$**<br>CLES:95.13% | U=81185.0<br>**p=$4.838 \times 10^{-16}$**<br>CLES:96.14% | U=3221.0<br>**p=$6.037 \times 10^{-14}$**<br>CLES:96.78% |
| Nervous Sys Immune cells | - | U=61404.0<br>p=$5.927 \times 10^{-4}$<br>CLES:59.33% | U=55092.0<br>p=$1.816 \times 10^{-3}$<br>CLES:58.54% | U=1625361.0<br>**p=$3.53 \times 10^{-7}$**<br>CLES:62.84% | U=17183.0<br>**p=$1.593 \times 10^{-7}$**<br>CLES:67.10% | U=370324.0<br>**p=$1.498 \times 10^{-21}$**<br>CLES:74.38% | U=324496.0<br>**p=$1.245 \times 10^{-23}$**<br>CLES:75.69% | U=13308.0<br>**p=$1.09 \times 10^{-15}$**<br>CLES:78.76% |
| Glia | - | - | U=280410.5<br>p=0.913<br>CLES:50.16% | U=8467249.0<br>**p=$1.138 \times 10^{-6}$**<br>CLES:55.12% | U=90333.0<br>**p=$5.011 \times 10^{-5}$**<br>CLES:59.39% | U=2020056.0<br>**p=$1.02 \times 10^{-58}$**<br>CLES:68.31% | U=1777562.0<br>**p=$1.329 \times 10^{-66}$**<br>CLES:69.81% | U=73775.5<br>**p=$1.336 \times 10^{-17}$**<br>CLES:73.52% |
| Immune (benchmark) | - | - | - | U=7509472.0<br>p=$6.574 \times 10^{-4}$<br>CLES:53.75% | U=78824.0<br>p=$2.82 \times 10^{-3}$<br>CLES:56.99% | U=1733721.0<br>**p=$1.341 \times 10^{-34}$**<br>CLES:64.46% | U=1528126.0<br>**p=$7.056 \times 10^{-41}$**<br>CLES:65.99% | U=63162.0<br>**p=$4.294 \times 10^{-12}$**<br>CLES:69.21% |
| protein-coding | - | - | - | - | U=2013034.0<br>p=0.156<br>CLES:52.95% | U=45421033.5<br>**p=$2.817 \times 10^{-110}$**<br>CLES:61.45% | U=40096032.0<br>**p=$6.851 \times 10^{-125}$**<br>CLES:63.00% | U=1673449.0<br>**p=$6.512 \times 10^{-11}$**<br>CLES:66.72% |
| Vascular cells | - | - | - | - | - | U=439054.0<br>**p=$2.546 \times 10^{-6}$**<br>CLES:60.00% | U=388292.0<br>**p=$5.131 \times 10^{-8}$**<br>CLES:61.62% | U=16372.0<br>**p=$1.31 \times 10^{-6}$**<br>CLES:65.93% |
| Neurons | - | - | - | - | - | - | U=6342114.5<br>p=$1.059 \times 10^{-2}$<br>CLES:51.77% | U=271790.5<br>p=$1.53 \times 10^{-2}$<br>CLES:56.29% |
| Housekeeping | - | - | - | - | - | - | - | U=226816.0<br>p=$7.989 \times 10^{-2}$<br>CLES:54.56% |

### Rat, Zhang et al., 2014, three cell types

| MWU Test | Immune | Glia | protein-coding | Endothelia | Housekeeping | Neuron | ATPase |
|---|---|---|---|---|---|---|---|
| MHC | U=6451.0<br>$p=2.253 \times 10^{-2}$<br>CLES:68.45% | U=11884.0<br>**$p=6.408 \times 10^{-5}$**<br>CLES:82.21% | U=187211.5<br>$p=1.174 \times 10^{-4}$<br>CLES:80.85% | U=8955.5<br>**$p=7.732 \times 10^{-5}$**<br>CLES:81.91% | U=36850.0<br>**$p=1.036 \times 10^{-7}$**<br>CLES:92.70% | U=14471.0<br>**$p=1.157 \times 10^{-7}$**<br>CLES:92.69% | U=1389.0<br>**$p=2.606 \times 10^{-7}$**<br>CLES:93.72% |
| Immune | - | U=460968.5<br>**$p=1.913 \times 10^{-7}$**<br>CLES:57.18% | U=7449466.5<br>**$p=2.099 \times 10^{-12}$**<br>CLES:57.69% | U=347972.0<br>**$p=1.357 \times 10^{-6}$**<br>CLES:57.07% | U=1511573.5<br>**$p=1.819 \times 10^{-52}$**<br>CLES:68.18% | U=613026.5<br>**$p=5.055 \times 10^{-51}$**<br>CLES:70.40% | U=58828.0<br>**$p=3.423 \times 10^{-13}$**<br>CLES:71.18% |
| Glia | - | - | U=10075222.0<br>p=0.331<br>CLES:50.87% | U=466223.0<br>p=0.911<br>CLES:49.85% | U=2139077.0<br>**$p=2.597 \times 10^{-37}$**<br>CLES:62.90% | U=883352.0<br>**$p=3.791 \times 10^{-41}$**<br>CLES:66.14% | U=84439.5<br>**$p=4.971 \times 10^{-9}$**<br>CLES:66.61% |
| protein-coding | - | - | - | U=7340076.0<br>p=0.326<br>CLES:49.00% | U=33447597.0<br>**$p=1.302 \times 10^{-90}$**<br>CLES:61.41% | U=13762536.0<br>**$p=2.758 \times 10^{-62}$**<br>CLES:64.33% | U=1319678.0<br>**$p=3.264 \times 10^{-8}$**<br>CLES:64.99% |
| Endothelia | - | - | - | - | U=1622416.5<br>**$p=2.573 \times 10^{-31}$**<br>CLES:63.09% | U=669969.0<br>**$p=2.767 \times 10^{-36}$**<br>CLES:66.33% | U=64148.5<br>**$p=4.472 \times 10^{-9}$**<br>CLES:66.91% |
| Housekeeping | - | - | - | - | - | U=1963065.5<br>$p=4.483 \times 10^{-4}$<br>CLES:53.45% | U=188824.0<br>p=0.131<br>CLES:54.16% |
| Neuron | - | - | - | - | - | - | U=69525.0<br>p=0.783<br>CLES:50.78% |

### Rat, Zhang et al., 2014, cell subtypes

| MWU Test | Immune | Microglia | protein-coding | Endothelia | Astrocyte | Oligodendrocyte | Housekeeping | Neuron | ATPase |
|---|---|---|---|---|---|---|---|---|---|
| MHC | U=6451.0<br>p=$2.253 \times 10^{-2}$<br>CLES:68.45% | U=9672.0<br>p=$6.509 \times 10^{-4}$<br>CLES:77.50% | U=187211.5<br>p=$1.174 \times 10^{-4}$<br>CLES:80.85% | U=8955.5<br>**p=$7.732 \times 10^{-5}$**<br>CLES:81.91% | U=8720.0<br>**p=$2.709 \times 10^{-6}$**<br>CLES:87.91% | U=5931.0<br>**p=$1.831 \times 10^{-7}$**<br>CLES:92.35% | U=36850.0<br>**p=$1.036 \times 10^{-7}$**<br>CLES:92.70% | U=14471.0<br>**p=$1.157 \times 10^{-7}$**<br>CLES:92.69% | U=1389.0<br>**p=$2.606 \times 10^{-7}$**<br>CLES:93.72% |
| Immune | - | U=374303.5<br>p=$7.817 \times 10^{-3}$<br>CLES:53.78% | U=7449466.5<br>**p=$2.099 \times 10^{-12}$**<br>CLES:57.69% | U=347972.0<br>**p=$1.357 \times 10^{-6}$**<br>CLES:57.07% | U=343320.0<br>**p=$7.971 \times 10^{-16}$**<br>CLES:62.06% | U=240952.5<br>**p=$1.132 \times 10^{-24}$**<br>CLES:67.28% | U=1511573.5<br>**p=$1.819 \times 10^{-52}$**<br>CLES:68.18% | U=613026.5<br>**p=$5.055 \times 10^{-51}$**<br>CLES:70.40% | U=58828.0<br>**p=$3.423 \times 10^{-13}$**<br>CLES:71.18% |
| Microglia | - | - | U=9326463.0<br>**p=$2.046 \times 10^{-6}$**<br>CLES:54.54% | U=434154.0<br>p=$5.647 \times 10^{-3}$<br>CLES:53.77% | U=434881.0<br>**p=$2.213 \times 10^{-11}$**<br>CLES:59.37% | U=311351.0<br>**p=$1.254 \times 10^{-22}$**<br>CLES:65.65% | U=1948670.5<br>**p=$4.499 \times 10^{-53}$**<br>CLES:66.38% | U=798831.0<br>**p=$1.07 \times 10^{-53}$**<br>CLES:69.29% | U=76417.5<br>**p=$4.225 \times 10^{-12}$**<br>CLES:69.83% |
| protein-coding | - | - | - | U=7340076.0<br>p=0.326<br>CLES:49.00% | U=7393672.5<br>**p=$3.698 \times 10^{-5}$**<br>CLES:54.40% | U=5320988.5<br>**p=$1.823 \times 10^{-15}$**<br>CLES:60.47% | U=33447597.0<br>**p=$1.302 \times 10^{-90}$**<br>CLES:61.41% | U=13762536.0<br>**p=$2.758 \times 10^{-62}$**<br>CLES:64.33% | U=1319678.0<br>**p=$3.264 \times 10^{-8}$**<br>CLES:64.99% |
| Endothelia | - | - | - | - | U=357647.0<br>**p=$7.102 \times 10^{-5}$**<br>CLES:55.74% | U=258464.0<br>**p=$8.644 \times 10^{-14}$**<br>CLES:62.21% | U=1622416.5<br>**p=$2.573 \times 10^{-31}$**<br>CLES:63.09% | U=669969.0<br>**p=$2.767 \times 10^{-36}$**<br>CLES:66.33% | U=64148.5<br>**p=$4.472 \times 10^{-9}$**<br>CLES:66.91% |
| Astrocyte | - | - | - | - | - | U=213533.0<br>**p=$6.65 \times 10^{-5}$**<br>CLES:56.65% | U=1345497.5<br>**p=$5.328 \times 10^{-11}$**<br>CLES:57.67% | U=560209.0<br>**p=$8.156 \times 10^{-17}$**<br>CLES:61.13% | U=53606.5<br>**p=$6.084 \times 10^{-5}$**<br>CLES:61.63% |
| Oligodendrocyte | - | - | - | - | - | - | U=775114.5<br>p=0.349<br>CLES:51.31% | U=326496.0<br>p=$1.116 \times 10^{-3}$<br>CLES:55.03% | U=31317.0<br>p=$6.172 \times 10^{-2}$<br>CLES:55.61% |
| Housekeeping | - | - | - | - | - | - | - | U=1963065.5<br>p=$4.483 \times 10^{-4}$<br>CLES:53.45% | U=188824.0<br>p=0.131<br>CLES:54.16% |
| Neuron | - | - | - | - | - | - | - | - | U=69525.0<br>p=0.783<br>CLES:50.78% |

### Rat, Zeisel et al., 2015, three cell types

| MWU Test | Immune | Glia | protein-coding | Vasculature | Housekeeping | Neuron | ATPase |
|---|---|---|---|---|---|---|---|
| MHC | U=6451.0<br>$p=2.253 \times 10^{-2}$<br>CLES:68.45% | U=13497.5<br>**$p=4.636 \times 10^{-5}$**<br>CLES:82.80% | U=187211.5<br>$p=1.174 \times 10^{-4}$<br>CLES:80.85% | U=4625.0<br>**$p=4.943 \times 10^{-6}$**<br>CLES:87.20% | U=36850.0<br>**$p=1.036 \times 10^{-7}$**<br>CLES:92.70% | U=10078.0<br>**$p=1.624 \times 10^{-7}$**<br>CLES:92.29% | U=1389.0<br>**$p=2.606 \times 10^{-7}$**<br>CLES:93.72% |
| Immune | - | U=521389.0<br>**$p=4.894 \times 10^{-8}$**<br>CLES:57.35% | U=7449466.5<br>**$p=2.099 \times 10^{-12}$**<br>CLES:57.69% | U=182015.0<br>**$p=1.101 \times 10^{-10}$**<br>CLES:61.53% | U=1511573.5<br>**$p=1.819 \times 10^{-52}$**<br>CLES:68.18% | U=422865.5<br>**$p=3.135 \times 10^{-40}$**<br>CLES:69.44% | U=58828.0<br>**$p=3.423 \times 10^{-13}$**<br>CLES:71.18% |
| Glia | - | - | U=11384385.5<br>p=0.251<br>CLES:50.97% | U=282544.0<br>$p=1.503 \times 10^{-3}$<br>CLES:55.22% | U=2419771.0<br>**$p=1.002 \times 10^{-41}$**<br>CLES:63.10% | U=683498.0<br>**$p=6.278 \times 10^{-31}$**<br>CLES:64.89% | U=95406.5<br>**$p=3.121 \times 10^{-9}$**<br>CLES:66.74% |
| protein-coding | - | - | - | U=3912209.0<br>$p=8.007 \times 10^{-3}$<br>CLES:53.83% | U=33447597.0<br>**$p=1.302 \times 10^{-90}$**<br>CLES:61.41% | U=9443780.0<br>**$p=6.629 \times 10^{-38}$**<br>CLES:63.12% | U=1319678.0<br>**$p=3.264 \times 10^{-8}$**<br>CLES:64.99% |
| Vasculature | - | - | - | - | U=726493.5<br>**$p=6.403 \times 10^{-8}$**<br>CLES:58.23% | U=206598.0<br>**$p=3.637 \times 10^{-9}$**<br>CLES:60.28% | U=28972.5<br>**$p=5.953 \times 10^{-5}$**<br>CLES:62.29% |
| Housekeeping | - | - | - | - | - | U=1336864.0<br>$p=6.916 \times 10^{-2}$<br>CLES:52.04% | U=188824.0<br>p=0.131<br>CLES:54.16% |
| Neuron | - | - | - | - | - | - | U=49878.0<br>p=0.469<br>CLES:52.09% |

### Rat, Zeisel et al., 2015, cell subtypes

| MWU Test | Immune | Microglia | Ependymal | protein-coding | Astrocyte | Endothelial | Mural | Oligodendrocyte | Housekeeping | Interneuron | S1 Pyramidal | ATPase | CA1 Pyramidal |
|---|---|---|---|---|---|---|---|---|---|---|---|---|---|
| MHC | U=6451.0<br>p=$2.253 \times 10^{-2}$<br>CLES:68.45% | U=3267.5<br>p=$2.637 \times 10^{-3}$<br>CLES:74.58% | U=3750.0<br>p=$2.761 \times 10^{-4}$<br>CLES:79.69% | U=187211.5<br>p=$1.174 \times 10^{-4}$<br>CLES:80.85% | U=2256.0<br>**p=$1.497 \times 10^{-6}$**<br>CLES:89.92% | U=3111.0<br>**p=$2.531 \times 10^{-5}$**<br>CLES:84.56% | U=1514.0<br>**p=$3.231 \times 10^{-7}$**<br>CLES:93.17% | U=4224.0<br>**p=$1.11 \times 10^{-6}$**<br>CLES:89.76% | U=36850.0<br>**p=$1.036 \times 10^{-7}$**<br>CLES:92.70% | U=3486.0<br>**p=$4.092 \times 10^{-7}$**<br>CLES:91.52% | U=2753.0<br>**p=$1.558 \times 10^{-7}$**<br>CLES:93.29% | U=1389.0<br>**p=$2.606 \times 10^{-7}$**<br>CLES:93.72% | U=3839.0<br>**p=$2.368 \times 10^{-7}$**<br>CLES:92.28% |
| Immune | - | U=124613.5<br>p=0.598<br>CLES:51.00% | U=141908.5<br>p=$2.852 \times 10^{-2}$<br>CLES:54.07% | U=7449466.5<br>**p=$2.099 \times 10^{-12}$**<br>CLES:57.69% | U=87669.0<br>**p=$6.343 \times 10^{-8}$**<br>CLES:62.65% | U=123103.5<br>**p=$7.841 \times 10^{-7}$**<br>CLES:60.00% | U=58911.5<br>**p=$8.138 \times 10^{-8}$**<br>CLES:65.01% | U=167198.0<br>**p=$1.652 \times 10^{-13}$**<br>CLES:63.71% | U=1511573.5<br>**p=$1.819 \times 10^{-52}$**<br>CLES:68.18% | U=145692.0<br>**p=$1.467 \times 10^{-20}$**<br>CLES:68.59% | U=114648.0<br>**p=$3.523 \times 10^{-19}$**<br>CLES:69.66% | U=58828.0<br>**p=$3.423 \times 10^{-13}$**<br>CLES:71.18% | U=162525.5<br>**p=$4.369 \times 10^{-25}$**<br>CLES:70.05% |
| Microglia | - | - | U=64622.0<br>p=0.174<br>CLES:52.97% | U=3461141.0<br>**p=$1.395 \times 10^{-6}$**<br>CLES:57.66% | U=41323.0<br>**p=$2.121 \times 10^{-7}$**<br>CLES:63.53% | U=57584.0<br>**p=$8.375 \times 10^{-6}$**<br>CLES:60.38% | U=27901.0<br>**p=$8.156 \times 10^{-8}$**<br>CLES:66.23% | U=78831.0<br>**p=$2.307 \times 10^{-11}$**<br>CLES:64.62% | U=716662.0<br>**p=$4.264 \times 10^{-32}$**<br>CLES:69.54% | U=69237.0<br>**p=$2.821 \times 10^{-18}$**<br>CLES:70.12% | U=54611.0<br>**p=$6.648 \times 10^{-18}$**<br>CLES:71.39% | U=27969.0<br>**p=$3.296 \times 10^{-13}$**<br>CLES:72.80% | U=77310.0<br>**p=$6.678 \times 10^{-22}$**<br>CLES:71.69% |
| Ependymal | - | - | - | U=3544184.0<br>p=$1.194 \times 10^{-3}$<br>CLES:54.97% | U=42709.0<br>**p=$1.549 \times 10^{-5}$**<br>CLES:61.13% | U=59140.0<br>p=$7.489 \times 10^{-4}$<br>CLES:57.73% | U=28912.0<br>**p=$3.582 \times 10^{-6}$**<br>CLES:63.89% | U=81377.0<br>**p=$1.754 \times 10^{-8}$**<br>CLES:62.10% | U=743729.0<br>**p=$9.304 \times 10^{-27}$**<br>CLES:67.18% | U=71892.0<br>**p=$4.812 \times 10^{-15}$**<br>CLES:67.78% | U=56779.0<br>**p=$5.875 \times 10^{-15}$**<br>CLES:69.10% | U=29081.5<br>**p=$4.247 \times 10^{-11}$**<br>CLES:70.47% | U=80209.0<br>**p=$3.949 \times 10^{-18}$**<br>CLES:69.24% |
| protein-coding | - | - | - | - | U=1887667.5<br>p=$1.875 \times 10^{-2}$<br>CLES:54.91% | U=2633935.5<br>p=0.193<br>CLES:52.25% | U=1278273.5<br>p=$4.23 \times 10^{-3}$<br>CLES:57.41% | U=3618932.0<br>**p=$6.425 \times 10^{-5}$**<br>CLES:56.13% | U=33447597.0<br>**p=$1.302 \times 10^{-90}$**<br>CLES:61.41% | U=3249764.5<br>**p=$5.367 \times 10^{-13}$**<br>CLES:62.27% | U=2557139.5<br>**p=$6.501 \times 10^{-12}$**<br>CLES:63.24% | U=1319678.0<br>**p=$3.264 \times 10^{-8}$**<br>CLES:64.99% | U=3636876.0<br>**p=$2.26 \times 10^{-17}$**<br>CLES:63.81% |
| Astrocyte | - | - | - | - | - | U=25942.0<br>p=0.354<br>CLES:47.50% | U=12711.0<br>p=0.418<br>CLES:52.69% | U=36011.0<br>p=0.549<br>CLES:51.54% | U=339320.5<br>p=$4.709 \times 10^{-4}$<br>CLES:57.49% | U=33241.0<br>p=$1.045 \times 10^{-3}$<br>CLES:58.78% | U=26167.0<br>p=$5.884 \times 10^{-4}$<br>CLES:59.73% | U=13562.0<br>p=$6.561 \times 10^{-4}$<br>CLES:61.64% | U=37399.0<br>**p=$6.129 \times 10^{-5}$**<br>CLES:60.56% |
| Endothelial | - | - | - | - | - | - | U=19483.0<br>p=0.102<br>CLES:55.08% | U=55248.0<br>p=$8.658 \times 10^{-2}$<br>CLES:53.93% | U=515825.5<br>**p=$8.608 \times 10^{-8}$**<br>CLES:59.60% | U=50272.0<br>**p=$1.019 \times 10^{-5}$**<br>CLES:60.63% | U=39651.0<br>**p=$5.296 \times 10^{-6}$**<br>CLES:61.72% | U=20484.5<br>**p=$2.575 \times 10^{-5}$**<br>CLES:63.49% | U=56436.0<br>**p=$1.741 \times 10^{-7}$**<br>CLES:62.32% |
| Mural | - | - | - | - | - | - | - | U=22094.0<br>p=0.696<br>CLES:48.83% | U=210668.0<br>p=$5.232 \times 10^{-2}$<br>CLES:55.11% | U=20663.0<br>p=$3.77 \times 10^{-2}$<br>CLES:56.42% | U=16263.0<br>p=$2.314 \times 10^{-2}$<br>CLES:57.31% | U=8488.0<br>p=$1.07 \times 10^{-2}$<br>CLES:59.56% | U=23313.0<br>p=$6.593 \times 10^{-3}$<br>CLES:58.28% |
| Oligodendrocyte | - | - | - | - | - | - | - | - | U=620059.5<br>p=$1.791 \times 10^{-4}$<br>CLES:56.01% | U=60634.0<br>p=$1.597 \times 10^{-3}$<br>CLES:57.17% | U=47761.0<br>p=$8.999 \times 10^{-4}$<br>CLES:58.12% | U=24794.0<br>p=$1.164 \times 10^{-3}$<br>CLES:60.08% | U=68258.0<br>**p=$5.675 \times 10^{-5}$**<br>CLES:58.92% |
| Housekeeping | - | - | - | - | - | - | - | - | - | U=459338.0<br>p=0.473<br>CLES:51.27% | U=360295.5<br>p=0.338<br>CLES:51.90% | U=188824.0<br>p=0.131<br>CLES:54.16% | U=517230.5<br>p=$9.222 \times 10^{-2}$<br>CLES:52.86% |
| Interneuron | - | - | - | - | - | - | - | - | - | - | U=33697.0<br>p=0.795<br>CLES:50.66% | U=17632.0<br>p=0.383<br>CLES:52.79% | U=48358.0<br>p=0.5<br>CLES:51.58% |
| S1 Pyramidal | - | - | - | - | - | - | - | - | - | - | - | U=13516.0<br>p=0.502<br>CLES:52.23% | U=37067.0<br>p=0.682<br>CLES:51.03% |
| ATPase | - | - | - | - | - | - | - | - | - | - | - | - | U=17750.0<br>p=0.67<br>CLES:48.66% |

### Rat, Saunders et al., 2018, three cell types

| MWU Test | Immune | protein-coding | Vasculature | Glia | Housekeeping | ATPase | Neuron |
|---|---|---|---|---|---|---|---|
| MHC | U=6451.0<br>$p=2.253 \times 10^{-2}$<br>CLES:68.45% | U=187211.5<br>$p=1.174 \times 10^{-4}$<br>CLES:80.85% | U=4279.5<br>**$p=2.729 \times 10^{-5}$**<br>CLES:84.19% | U=3429.0<br>**$p=2.137 \times 10^{-7}$**<br>CLES:92.55% | U=36850.0<br>**$p=1.036 \times 10^{-7}$**<br>CLES:92.70% | U=1389.0<br>**$p=2.606 \times 10^{-7}$**<br>CLES:93.72% | U=7922.0<br>**$p=9.801 \times 10^{-9}$**<br>CLES:96.42% |
| Immune | - | U=7449466.5<br>**$p=2.099 \times 10^{-12}$**<br>CLES:57.69% | U=172144.5<br>**$p=3.232 \times 10^{-9}$**<br>CLES:60.73% | U=140376.0<br>**$p=6.503 \times 10^{-19}$**<br>CLES:67.94% | U=1511573.5<br>**$p=1.819 \times 10^{-52}$**<br>CLES:68.18% | U=58828.0<br>**$p=3.423 \times 10^{-13}$**<br>CLES:71.18% | U=352281.5<br>**$p=1.331 \times 10^{-65}$**<br>CLES:76.88% |
| protein-coding | - | - | U=3706959.5<br>$p=2.88 \times 10^{-2}$<br>CLES:53.23% | U=3111703.5<br>**$p=5.591 \times 10^{-11}$**<br>CLES:61.30% | U=33447597.0<br>**$p=1.302 \times 10^{-90}$**<br>CLES:61.41% | U=1319678.0<br>**$p=3.264 \times 10^{-8}$**<br>CLES:64.99% | U=8105674.0<br>**$p=4.196 \times 10^{-79}$**<br>CLES:72.00% |
| Vasculature | - | - | - | U=64882.0<br>$p=2.573 \times 10^{-4}$<br>CLES:58.22% | U=698474.0<br>**$p=5.706 \times 10^{-8}$**<br>CLES:58.42% | U=27767.5<br>**$p=6.41 \times 10^{-5}$**<br>CLES:62.30% | U=172392.0<br>**$p=2.053 \times 10^{-26}$**<br>CLES:69.76% |
| Glia | - | - | - | - | U=440070.5<br>p=0.782<br>CLES:50.49% | U=17851.0<br>p=0.123<br>CLES:54.94% | U=114582.0<br>**$p=3.95 \times 10^{-11}$**<br>CLES:63.61% |
| Housekeeping | - | - | - | - | - | U=188824.0<br>p=0.131<br>CLES:54.16% | U=1201320.0<br>**$p=5.518 \times 10^{-22}$**<br>CLES:62.16% |
| ATPase | - | - | - | - | - | - | U=41696.0<br>$p=7.405 \times 10^{-3}$<br>CLES:57.87% |

### Rat, Saunders et al., 2018, cell subtypes

| MWU Test | Immune | Microglia | protein-coding | Endothelia | Housekeeping | Astrocyte | Oligodendrocyte | ATPase | Neuron |
|---|---|---|---|---|---|---|---|---|---|
| MHC | U=6451.0<br>p=$2.253 \times 10^{-2}$<br>CLES:68.45% | U=2633.0<br>p=$2.238 \times 10^{-4}$<br>CLES:80.37% | U=187211.5<br>p=$1.174 \times 10^{-4}$<br>CLES:80.85% | U=3793.0<br>**p=$3.893 \times 10^{-5}$**<br>CLES:83.60% | U=36850.0<br>**p=$1.036 \times 10^{-7}$**<br>CLES:92.70% | U=3183.0<br>**p=$9.872 \times 10^{-8}$**<br>CLES:93.81% | U=3108.0<br>**p=$1.043 \times 10^{-7}$**<br>CLES:93.76% | U=1389.0<br>**p=$2.606 \times 10^{-7}$**<br>CLES:93.72% | U=7311.0<br>**p=$1.105 \times 10^{-8}$**<br>CLES:96.30% |
| Immune | - | U=102410.5<br>p=$4.154 \times 10^{-3}$<br>CLES:56.05% | U=7449466.5<br>**p=$2.099 \times 10^{-12}$**<br>CLES:57.69% | U=151099.5<br>**p=$2.415 \times 10^{-7}$**<br>CLES:59.72% | U=1511573.5<br>**p=$1.819 \times 10^{-52}$**<br>CLES:68.18% | U=129851.5<br>**p=$4.175 \times 10^{-19}$**<br>CLES:68.62% | U=127752.0<br>**p=$1.049 \times 10^{-19}$**<br>CLES:69.10% | U=58828.0<br>**p=$3.423 \times 10^{-13}$**<br>CLES:71.18% | U=323813.5<br>**p=$4.289 \times 10^{-61}$**<br>CLES:76.48% |
| Microglia | - | - | U=2334246.5<br>p=0.274<br>CLES:52.00% | U=47589.0<br>p=$8.529 \times 10^{-2}$<br>CLES:54.11% | U=492640.0<br>**p=$1.829 \times 10^{-13}$**<br>CLES:63.93% | U=42464.0<br>**p=$1.156 \times 10^{-8}$**<br>CLES:64.56% | U=41987.0<br>**p=$2.287 \times 10^{-9}$**<br>CLES:65.34% | U=19445.5<br>**p=$5.937 \times 10^{-8}$**<br>CLES:67.69% | U=109485.0<br>**p=$3.843 \times 10^{-29}$**<br>CLES:74.39% |
| protein-coding | - | - | - | U=3235662.5<br>p=0.189<br>CLES:52.05% | U=33447597.0<br>**p=$1.302 \times 10^{-90}$**<br>CLES:61.41% | U=2880084.5<br>**p=$3.138 \times 10^{-11}$**<br>CLES:61.95% | U=2844850.5<br>**p=$3.954 \times 10^{-12}$**<br>CLES:62.63% | U=1319678.0<br>**p=$3.264 \times 10^{-8}$**<br>CLES:64.99% | U=7437596.0<br>**p=$3.503 \times 10^{-70}$**<br>CLES:71.50% |
| Endothelia | - | - | - | - | U=638166.0<br>**p=$1.914 \times 10^{-9}$**<br>CLES:59.80% | U=54912.0<br>**p=$1.365 \times 10^{-5}$**<br>CLES:60.28% | U=54285.0<br>**p=$3.83 \times 10^{-6}$**<br>CLES:61.00% | U=25367.5<br>**p=$1.018 \times 10^{-5}$**<br>CLES:63.76% | U=144279.0<br>**p=$1.983 \times 10^{-26}$**<br>CLES:70.79% |
| Housekeeping | - | - | - | - | - | U=400201.0<br>p=0.939<br>CLES:50.14% | U=397158.5<br>p=0.621<br>CLES:50.93% | U=188824.0<br>p=0.131<br>CLES:54.16% | U=1098694.0<br>**p=$9.856 \times 10^{-19}$**<br>CLES:61.52% |
| Astrocyte | - | - | - | - | - | - | U=33920.0<br>p=0.705<br>CLES:50.97% | U=16180.0<br>p=0.177<br>CLES:54.38% | U=95147.0<br>**p=$7.646 \times 10^{-9}$**<br>CLES:62.42% |
| Oligodendrocyte | - | - | - | - | - | - | - | U=15548.5<br>p=0.285<br>CLES:53.49% | U=91599.0<br>**p=$1.107 \times 10^{-7}$**<br>CLES:61.51% |
| ATPase | - | - | - | - | - | - | - | - | U=38069.0<br>p=$1.52 \times 10^{-2}$<br>CLES:57.18% |

### Rat, Zeisel et al., 2018, three cell types

| MWU Test | Immune | CNS_Glia | protein-coding | CNS_Endothelia | Housekeeping | CNS_Neuron | ATPase |
|---|---|---|---|---|---|---|---|
| MHC | U=6451.0<br>$p=2.253 \times 10^{-2}$<br>CLES:68.45% | U=9532.0<br>**$p=2.821 \times 10^{-5}$**<br>CLES:83.80% | U=187211.5<br>$p=1.174 \times 10^{-4}$<br>CLES:80.85% | U=1696.0<br>**$p=4.424 \times 10^{-5}$**<br>CLES:84.17% | U=36850.0<br>**$p=1.036 \times 10^{-7}$**<br>CLES:92.70% | U=40468.0<br>**$p=6.102 \times 10^{-8}$**<br>CLES:93.45% | U=1389.0<br>**$p=2.606 \times 10^{-7}$**<br>CLES:93.72% |
| Immune | - | U=361513.0<br>**$p=1.45 \times 10^{-6}$**<br>CLES:56.99% | U=7449466.5<br>**$p=2.099 \times 10^{-12}$**<br>CLES:57.69% | U=63980.5<br>$p=6.668 \times 10^{-3}$<br>CLES:56.93% | U=1511573.5<br>**$p=1.819 \times 10^{-52}$**<br>CLES:68.18% | U=1684304.0<br>**$p=1.625 \times 10^{-62}$**<br>CLES:69.74% | U=58828.0<br>**$p=3.423 \times 10^{-13}$**<br>CLES:71.18% |
| CNS_Glia | - | - | U=8039192.0<br>p=0.114<br>CLES:51.58% | U=68411.0<br>p=0.861<br>CLES:50.44% | U=1717335.5<br>**$p=1.405 \times 10^{-37}$**<br>CLES:64.18% | U=1939586.0<br>**$p=1.969 \times 10^{-51}$**<br>CLES:66.55% | U=67915.5<br>**$p=3.204 \times 10^{-10}$**<br>CLES:68.09% |
| protein-coding | - | - | - | U=1342371.5<br>p=0.554<br>CLES:48.62% | U=33447597.0<br>**$p=1.302 \times 10^{-90}$**<br>CLES:61.41% | U=37595320.5<br>**$p=8.146 \times 10^{-133}$**<br>CLES:63.36% | U=1319678.0<br>**$p=3.264 \times 10^{-8}$**<br>CLES:64.99% |
| CNS_Endothelia | - | - | - | - | U=303396.0<br>**$p=3.787 \times 10^{-9}$**<br>CLES:64.01% | U=343400.0<br>**$p=3.41 \times 10^{-12}$**<br>CLES:66.51% | U=12017.5<br>**$p=4.5 \times 10^{-7}$**<br>CLES:68.01% |
| Housekeeping | - | - | - | - | - | U=5301006.0<br>$p=4.758 \times 10^{-3}$<br>CLES:52.04% | U=188824.0<br>p=0.131<br>CLES:54.16% |
| CNS_Neuron | - | - | - | - | - | - | U=198510.0<br>p=0.408<br>CLES:52.28% |

### Rat, Zeisel et al., 2018, cell subtypes

| MWU Test | Nervous Sys Immune cells | Immune (benchmark) | Glia | protein-coding | Vascular cells | Neurons | Housekeeping | ATPase |
|---|---|---|---|---|---|---|---|---|
| MHC | U=1000.5<br>p=$5.427 \times 10^{-2}$<br>CLES:66.35% | U=6451.0<br>p=$2.253 \times 10^{-2}$<br>CLES:68.45% | U=7499.5<br>p=$1.653 \times 10^{-4}$<br>CLES:80.46% | U=187211.5<br>p=$1.174 \times 10^{-4}$<br>CLES:80.85% | U=2029.0<br>**p=$3.366 \times 10^{-6}$**<br>CLES:88.68% | U=41847.0<br>**p=$1.868 \times 10^{-7}$**<br>CLES:91.81% | U=36850.0<br>**p=$1.036 \times 10^{-7}$**<br>CLES:92.70% | U=1389.0<br>**p=$2.606 \times 10^{-7}$**<br>CLES:93.72% |
| Nervous Sys Immune cells | - | U=45276.5<br>p=0.184<br>CLES:53.84% | U=48304.0<br>p=$5.206 \times 10^{-3}$<br>CLES:58.08% | U=1279591.5<br>**p=$9.146 \times 10^{-6}$**<br>CLES:61.93% | U=13376.0<br>**p=$7.247 \times 10^{-6}$**<br>CLES:65.52% | U=293211.0<br>**p=$5.078 \times 10^{-16}$**<br>CLES:72.10% | U=258492.0<br>**p=$5.564 \times 10^{-17}$**<br>CLES:72.87% | U=10042.0<br>**p=$1.066 \times 10^{-11}$**<br>CLES:75.94% |
| Immune (benchmark) | - | - | U=279962.5<br>p=$1.122 \times 10^{-2}$<br>CLES:53.86% | U=7449466.5<br>**p=$2.099 \times 10^{-12}$**<br>CLES:57.69% | U=77402.5<br>**p=$1.123 \times 10^{-5}$**<br>CLES:60.66% | U=1707893.5<br>**p=$3.039 \times 10^{-48}$**<br>CLES:67.19% | U=1511573.5<br>**p=$1.819 \times 10^{-52}$**<br>CLES:68.18% | U=58828.0<br>**p=$3.423 \times 10^{-13}$**<br>CLES:71.18% |
| Glia | - | - | - | U=7054799.5<br>**p=$1.886 \times 10^{-6}$**<br>CLES:55.24% | U=74738.0<br>p=$1.466 \times 10^{-4}$<br>CLES:59.23% | U=1685417.0<br>**p=$4.775 \times 10^{-47}$**<br>CLES:67.05% | U=1489234.5<br>**p=$1.346 \times 10^{-50}$**<br>CLES:67.92% | U=58560.5<br>**p=$1.075 \times 10^{-13}$**<br>CLES:71.64% |
| protein-coding | - | - | - | - | U=1651050.0<br>p=0.223<br>CLES:52.67% | U=37694318.5<br>**p=$4.826 \times 10^{-84}$**<br>CLES:60.36% | U=33447597.0<br>**p=$1.302 \times 10^{-90}$**<br>CLES:61.41% | U=1319678.0<br>**p=$3.264 \times 10^{-8}$**<br>CLES:64.99% |
| Vascular cells | - | - | - | - | - | U=364427.0<br>**p=$4.868 \times 10^{-5}$**<br>CLES:59.06% | U=323120.0<br>**p=$7.315 \times 10^{-6}$**<br>CLES:60.04% | U=12901.0<br>**p=$3.916 \times 10^{-5}$**<br>CLES:64.30% |
| Neurons | - | - | - | - | - | - | U=5498604.5<br>p=$7.171 \times 10^{-2}$<br>CLES:51.29% | U=221982.0<br>p=$4.38 \times 10^{-2}$<br>CLES:55.54% |
| Housekeeping | - | - | - | - | - | - | - | U=188824.0<br>p=0.131<br>CLES:54.16% |

### Human, Zhang et al., 2014, three cell types

| MWU Test | protein-coding | Endothelia | Glia | Immune | Housekeeping | ATPase | Neuron |
|---|---|---|---|---|---|---|---|
| MHC | U=307170.0 **$p=2.294\times10^{-7}$** CLES:83.42% | U=14589.0 **$p=9.046\times10^{-9}$** CLES:87.57% | U=19217.0 **$p=3.558\times10^{-9}$** CLES:88.48% | U=20628.0 **$p=1.703\times10^{-6}$** CLES:81.15% | U=64943.0 **$p=3.233\times10^{-11}$** CLES:92.96% | U=2571.0 **$p=1.391\times10^{-10}$** CLES:94.52% | U=22985.0 **$p=7.47\times10^{-12}$** CLES:94.59% |
| protein-coding | - | U=7837525.0 p=0.281 CLES:51.10% | U=10466118.5 $p=9.361\times10^{-3}$ CLES:52.34% | U=12067086.0 $p=6.157\times10^{-2}$ CLES:51.57% | U=38906080.0 **$p=2.223\times10^{-86}$** CLES:60.49% | U=1562822.0 **$p=5.861\times10^{-7}$** CLES:62.41% | U=14630631.5 **$p=1.587\times10^{-72}$** CLES:65.40% |
| Endothelia | - | - | U=464836.0 p=0.298 CLES:51.38% | U=536981.0 p=0.577 CLES:50.72% | U=1752945.5 **$p=3.465\times10^{-20}$** CLES:60.25% | U=70667.5 **$p=3.587\times10^{-6}$** CLES:62.38% | U=665153.0 **$p=1.018\times10^{-33}$** CLES:65.72% |
| Glia | - | - | - | U=684687.5 p=0.74 CLES:49.60% | U=2236530.0 **$p=4.217\times10^{-19}$** CLES:58.96% | U=90177.5 **$p=2.568\times10^{-5}$** CLES:61.06% | U=851536.0 **$p=1.829\times10^{-33}$** CLES:64.54% |
| Immune | - | - | - | - | U=2578216.0 **$p=1.385\times10^{-17}$** CLES:58.07% | U=103391.0 $p=1.656\times10^{-4}$ CLES:59.81% | U=965899.5 **$p=2.453\times10^{-27}$** CLES:62.55% |
| Housekeeping | - | - | - | - | - | U=246623.5 p=0.448 CLES:51.92% | U=2353940.0 **$p=1.319\times10^{-8}$** CLES:55.47% |
| ATPase | - | - | - | - | - | - | U=88750.0 p=0.155 CLES:53.71% |

### Human, Zhang et al., 2014, cell subtypes

| MWU Test | protein-coding | Microglia | Endothelia | Immune | Astrocyte | Housekeeping | Oligodendrocyte | ATPase | Neuron |
|---|---|---|---|---|---|---|---|---|---|
| MHC | U=307170.0<br>**p=2.294 × $10^{-7}$**<br>CLES:83.42% | U=15990.5<br>**p=4.203 × $10^{-8}$**<br>CLES:85.79% | U=14589.0<br>**p=9.046 × $10^{-9}$**<br>CLES:87.57% | U=20628.0<br>**p=1.703 × $10^{-6}$**<br>CLES:81.15% | U=13876.0<br>**p=3.15 × $10^{-10}$**<br>CLES:91.17% | U=64943.0<br>**p=3.233 × $10^{-11}$**<br>CLES:92.96% | U=9269.0<br>**p=2.019 × $10^{-11}$**<br>CLES:94.20% | U=2571.0<br>**p=1.391 × $10^{-10}$**<br>CLES:94.52% | U=22985.0<br>**p=7.47 × $10^{-12}$**<br>CLES:94.59% |
| protein-coding | - | U=8509934.0<br>p=0.674<br>CLES:49.59% | U=7837525.0<br>p=0.281<br>CLES:51.10% | U=12067086.0<br>p=6.157 × $10^{-2}$<br>CLES:51.57% | U=7801208.5<br>**p=1.06 × $10^{-7}$**<br>CLES:55.68% | U=38906080.0<br>**p=2.223 × $10^{-86}$**<br>CLES:60.49% | U=5545010.0<br>**p=1.865 × $10^{-17}$**<br>CLES:61.21% | U=1562822.0<br>**p=5.861 × $10^{-7}$**<br>CLES:62.41% | U=14630631.5<br>**p=1.587 × $10^{-72}$**<br>CLES:65.40% |
| Microglia | - | - | U=400665.0<br>p=0.243<br>CLES:51.61% | U=616934.0<br>p=9.472 × $10^{-2}$<br>CLES:52.08% | U=401705.0<br>**p=2.539 × $10^{-6}$**<br>CLES:56.64% | U=2008567.5<br>**p=4.257 × $10^{-28}$**<br>CLES:61.70% | U=287524.0<br>**p=2.925 × $10^{-15}$**<br>CLES:62.70% | U=80780.5<br>**p=2.227 × $10^{-7}$**<br>CLES:63.73% | U=759640.0<br>**p=4.739 × $10^{-42}$**<br>CLES:67.08% |
| Endothelia | - | - | - | U=536981.0<br>p=0.577<br>CLES:50.72% | U=348749.0<br>p=5.332 × $10^{-4}$<br>CLES:55.02% | U=1752945.5<br>**p=3.465 × $10^{-20}$**<br>CLES:60.25% | U=250888.0<br>**p=8.433 × $10^{-12}$**<br>CLES:61.22% | U=70667.5<br>**p=3.587 × $10^{-6}$**<br>CLES:62.38% | U=665153.0<br>**p=1.018 × $10^{-33}$**<br>CLES:65.72% |
| Immune | - | - | - | - | U=517012.5<br>p=9.083 × $10^{-3}$<br>CLES:53.45% | U=2578216.0<br>**p=1.385 × $10^{-17}$**<br>CLES:58.07% | U=365373.0<br>**p=3.856 × $10^{-8}$**<br>CLES:58.43% | U=103391.0<br>p=1.656 × $10^{-4}$<br>CLES:59.81% | U=965899.5<br>**p=2.453 × $10^{-27}$**<br>CLES:62.55% |
| Astrocyte | - | - | - | - | - | U=1476614.0<br>**p=1.545 × $10^{-6}$**<br>CLES:55.55% | U=210498.0<br>p=1.964 × $10^{-4}$<br>CLES:56.22% | U=59697.5<br>p=4.286 × $10^{-3}$<br>CLES:57.68% | U=566217.0<br>**p=3.797 × $10^{-17}$**<br>CLES:61.24% |
| Housekeeping | - | - | - | - | - | - | U=865323.5<br>p=0.8<br>CLES:50.35% | U=246623.5<br>p=0.448<br>CLES:51.92% | U=2353940.0<br>**p=1.319 × $10^{-8}$**<br>CLES:55.47% |
| Oligodendrocyte | - | - | - | - | - | - | - | U=34652.0<br>p=0.523<br>CLES:51.79% | U=332040.0<br>p=3.259 × $10^{-4}$<br>CLES:55.55% |
| ATPase | - | - | - | - | - | - | - | - | U=88750.0<br>p=0.155<br>CLES:53.71% |

### Human, Zeisel et al., 2015, three cell types

| MWU Test | Glia | protein-coding | Immune | Vasculature | Housekeeping | ATPase | Neuron |
|---|---|---|---|---|---|---|---|
| MHC | U=21805.0 **$p=7.853 \times 10^{-9}$** CLES:87.57% | U=307170.0 **$p=2.294 \times 10^{-7}$** CLES:83.42% | U=20628.0 **$p=1.703 \times 10^{-6}$** CLES:81.15% | U=7616.5 **$p=6.386 \times 10^{-10}$** CLES:90.89% | U=64943.0 **$p=3.233 \times 10^{-11}$** CLES:92.96% | U=2571.0 **$p=1.391 \times 10^{-10}$** CLES:94.52% | U=16166.5 **$p=7.368 \times 10^{-12}$** CLES:94.76% |
| Glia | - | U=11354376.0 p=0.581 CLES:49.53% | U=809750.5 p=0.309 CLES:51.17% | U=294177.5 **$p=8.857 \times 10^{-5}$** CLES:56.39% | U=2651040.5 **$p=1.294 \times 10^{-30}$** CLES:60.96% | U=106727.5 **$p=5.817 \times 10^{-7}$** CLES:63.03% | U=692310.0 **$p=2.543 \times 10^{-32}$** CLES:65.19% |
| protein-coding | - | - | U=12067086.0 $p=6.157 \times 10^{-2}$ CLES:51.57% | U=4337629.0 **$p=1.27 \times 10^{-5}$** CLES:56.23% | U=38906080.0 **$p=2.223 \times 10^{-86}$** CLES:60.49% | U=1562822.0 **$p=5.861 \times 10^{-7}$** CLES:62.41% | U=10096561.5 **$p=2.458 \times 10^{-45}$** CLES:64.29% |
| Immune | - | - | - | U=287440.5 $p=1.456 \times 10^{-2}$ CLES:53.97% | U=2578216.0 **$p=1.385 \times 10^{-17}$** CLES:58.07% | U=103391.0 $p=1.656 \times 10^{-4}$ CLES:59.81% | U=666491.0 **$p=2.739 \times 10^{-19}$** CLES:61.48% |
| Vasculature | - | - | - | - | U=803173.5 $p=1.084 \times 10^{-3}$ CLES:54.88% | U=32443.5 $p=1.503 \times 10^{-2}$ CLES:56.93% | U=211492.0 **$p=1.01 \times 10^{-7}$** CLES:59.17% |
| Housekeeping | - | - | - | - | - | U=246623.5 p=0.448 CLES:51.92% | U=1611743.0 $p=2.05 \times 10^{-4}$ CLES:54.09% |
| ATPase | - | - | - | - | - | - | U=60614.0 p=0.399 CLES:52.25% |

### Human, Zeisel et al., 2015, cell subtypes

| MWU Test | Ependymal | Microglia | protein-coding | Immune | Astrocyte | Endothelial | Oligodendrocyte | Mural | Housekeeping | Interneuron | ATPase | S1 Pyramidal | CA1 Pyramidal |
|---|---|---|---|---|---|---|---|---|---|---|---|---|---|
| MHC | U=6013.0<br>**$p=1.086 \times 10^{-6}$**<br>CLES:82.37% | U=5410.0<br>**$p=2.206 \times 10^{-7}$**<br>CLES:84.53% | U=307170.0<br>**$p=2.294 \times 10^{-7}$**<br>CLES:83.42% | U=20628.0<br>**$p=1.703 \times 10^{-6}$**<br>CLES:81.15% | U=3618.0<br>**$p=3.102 \times 10^{-10}$**<br>CLES:92.77% | U=5157.5<br>**$p=7.633 \times 10^{-9}$**<br>CLES:88.62% | U=6764.0<br>**$p=1.317 \times 10^{-10}$**<br>CLES:92.66% | U=2459.0<br>**$p=3.841 \times 10^{-11}$**<br>CLES:96.05% | U=64943.0<br>**$p=3.233 \times 10^{-11}$**<br>CLES:92.96% | U=5607.0<br>**$p=5.67 \times 10^{-11}$**<br>CLES:93.76% | U=2571.0<br>**$p=1.391 \times 10^{-10}$**<br>CLES:94.52% | U=4307.0<br>**$p=5.371 \times 10^{-12}$**<br>CLES:96.57% | U=6252.5<br>**$p=2.451 \times 10^{-11}$**<br>CLES:94.45% |
| Ependymal | - | U=62697.0<br>$p=9.637 \times 10^{-2}$<br>CLES:53.68% | U=3784176.5<br>**$p=3.558 \times 10^{-5}$**<br>CLES:56.31% | U=264858.0<br>**$p=3.54 \times 10^{-5}$**<br>CLES:57.09% | U=45732.0<br>**$p=2.682 \times 10^{-8}$**<br>CLES:64.25% | U=65499.0<br>**$p=2.772 \times 10^{-7}$**<br>CLES:61.67% | U=86955.0<br>**$p=9.295 \times 10^{-13}$**<br>CLES:65.27% | U=31934.0<br>**$p=6.327 \times 10^{-10}$**<br>CLES:68.35% | U=865553.5<br>**$p=1.974 \times 10^{-29}$**<br>CLES:67.89% | U=76745.0<br>**$p=1.919 \times 10^{-19}$**<br>CLES:70.32% | U=34698.5<br>**$p=7.143 \times 10^{-12}$**<br>CLES:69.90% | U=59199.0<br>**$p=2.12 \times 10^{-20}$**<br>CLES:72.73% | U=87574.0<br>**$p=1.123 \times 10^{-24}$**<br>CLES:72.49% |
| Microglia | - | - | U=3134307.0<br>$p=4.949 \times 10^{-2}$<br>CLES:53.20% | U=221796.5<br>$p=1.208 \times 10^{-2}$<br>CLES:54.53% | U=37783.0<br>**$p=5.857 \times 10^{-5}$**<br>CLES:60.55% | U=54462.5<br>$p=2.878 \times 10^{-4}$<br>CLES:58.49% | U=72100.0<br>**$p=1.148 \times 10^{-7}$**<br>CLES:61.73% | U=26470.0<br>**$p=1.311 \times 10^{-6}$**<br>CLES:64.62% | U=722494.0<br>**$p=3.94 \times 10^{-18}$**<br>CLES:64.64% | U=64374.0<br>**$p=1.037 \times 10^{-13}$**<br>CLES:67.28% | U=29034.0<br>**$p=1.607 \times 10^{-8}$**<br>CLES:66.71% | U=49571.0<br>**$p=1.137 \times 10^{-14}$**<br>CLES:69.47% | U=73773.0<br>**$p=4.112 \times 10^{-18}$**<br>CLES:69.65% |
| protein-coding | - | - | - | U=12067086.0<br>$p=6.157 \times 10^{-2}$<br>CLES:51.57% | U=2018165.5<br>$p=2.803 \times 10^{-3}$<br>CLES:56.21% | U=2932601.0<br>$p=5.509 \times 10^{-3}$<br>CLES:54.73% | U=3856662.0<br>**$p=1.29 \times 10^{-6}$**<br>CLES:57.39% | U=1405028.0<br>$p=1.725 \times 10^{-4}$<br>CLES:59.62% | U=38906080.0<br>**$p=2.223 \times 10^{-86}$**<br>CLES:60.49% | U=3454205.5<br>**$p=3.66 \times 10^{-14}$**<br>CLES:62.74% | U=1562822.0<br>**$p=5.861 \times 10^{-7}$**<br>CLES:62.41% | U=2663897.5<br>**$p=1.993 \times 10^{-14}$**<br>CLES:64.88% | U=3978458.5<br>**$p=1.362 \times 10^{-21}$**<br>CLES:65.28% |
| Immune | - | - | - | - | U=133447.0<br>$p=8.359 \times 10^{-2}$<br>CLES:53.84% | U=195138.5<br>p=0.141<br>CLES:52.76% | U=255217.0<br>$p=3.459 \times 10^{-3}$<br>CLES:55.01% | U=92302.0<br>$p=1.19 \times 10^{-2}$<br>CLES:56.74% | U=2578216.0<br>**$p=1.385 \times 10^{-17}$**<br>CLES:58.07% | U=227900.5<br>**$p=7.824 \times 10^{-8}$**<br>CLES:59.97% | U=103391.0<br>$p=1.656 \times 10^{-4}$<br>CLES:59.81% | U=175608.5<br>**$p=1.174 \times 10^{-8}$**<br>CLES:61.96% | U=262982.0<br>**$p=2.208 \times 10^{-12}$**<br>CLES:62.51% |
| Astrocyte | - | - | - | - | - | U=27657.0<br>p=0.638<br>CLES:48.74% | U=36641.0<br>p=0.564<br>CLES:51.48% | U=13431.0<br>p=0.247<br>CLES:53.81% | U=376366.0<br>$p=1.337 \times 10^{-2}$<br>CLES:55.26% | U=33861.0<br>$p=2.399 \times 10^{-3}$<br>CLES:58.08% | U=15264.0<br>$p=1.934 \times 10^{-2}$<br>CLES:57.56% | U=26182.0<br>$p=3.152 \times 10^{-4}$<br>CLES:60.21% | U=39426.0<br>**$p=2.151 \times 10^{-5}$**<br>CLES:61.08% |
| Endothelial | - | - | - | - | - | - | U=55833.0<br>p=0.258<br>CLES:52.57% | U=20341.0<br>p=0.133<br>CLES:54.61% | U=570454.5<br>$p=5.108 \times 10^{-4}$<br>CLES:56.12% | U=50779.0<br>$p=4.416 \times 10^{-4}$<br>CLES:58.36% | U=23017.5<br>$p=6.571 \times 10^{-3}$<br>CLES:58.16% | U=39355.0<br>**$p=3.482 \times 10^{-5}$**<br>CLES:60.65% | U=59091.0<br>**$p=1.02 \times 10^{-6}$**<br>CLES:61.35% |
| Oligodendrocyte | - | - | - | - | - | - | - | U=24342.0<br>p=0.479<br>CLES:52.10% | U=686627.0<br>$p=1.52 \times 10^{-2}$<br>CLES:53.86% | U=61791.0<br>$p=3.313 \times 10^{-3}$<br>CLES:56.62% | U=27731.0<br>$p=4.341 \times 10^{-2}$<br>CLES:55.86% | U=47779.0<br>$p=3.961 \times 10^{-4}$<br>CLES:58.70% | U=72035.0<br>**$p=1.138 \times 10^{-5}$**<br>CLES:59.62% |
| Mural | - | - | - | - | - | - | - | - | U=232719.0<br>p=0.43<br>CLES:52.05% | U=21063.0<br>$p=9.917 \times 10^{-2}$<br>CLES:55.04% | U=9426.0<br>p=0.245<br>CLES:54.15% | U=16411.0<br>$p=1.944 \times 10^{-2}$<br>CLES:57.49% | U=24793.0<br>$p=4.633 \times 10^{-3}$<br>CLES:58.52% |
| Housekeeping | - | - | - | - | - | - | - | - | - | U=547680.5<br>p=0.161<br>CLES:52.44% | U=246623.5<br>p=0.448<br>CLES:51.92% | U=423831.5<br>$p=2.696 \times 10^{-2}$<br>CLES:54.41% | U=640231.0<br>$p=1.208 \times 10^{-3}$<br>CLES:55.37% |
| Interneuron | - | - | - | - | - | - | - | - | - | - | U=20126.0<br>p=0.866<br>CLES:49.49% | U=34703.0<br>p=0.424<br>CLES:52.05% | U=52672.0<br>p=0.162<br>CLES:53.22% |
| ATPase | - | - | - | - | - | - | - | - | - | - | - | U=15991.0<br>p=0.386<br>CLES:52.73% | U=24085.0<br>p=0.234<br>CLES:53.50% |
| S1 Pyramidal | - | - | - | - | - | - | - | - | - | - | - | - | U=37800.0<br>p=0.629<br>CLES:51.21% |

### Human, Saunders et al., 2018, three cell types

| MWU Test | protein-coding | Immune | Vasculature | Housekeeping | Glia | ATPase | Neuron |
|---|---|---|---|---|---|---|---|
| MHC | U=307170.0 **$p=2.294 \times 10^{-7}$** CLES:83.42% | U=20628.0 **$p=1.703 \times 10^{-6}$** CLES:81.15% | U=7033.0 **$p=5.77 \times 10^{-9}$** CLES:88.58% | U=64943.0 **$p=3.233 \times 10^{-11}$** CLES:92.96% | U=5393.0 **$p=4.968 \times 10^{-11}$** CLES:93.95% | U=2571.0 **$p=1.391 \times 10^{-10}$** CLES:94.52% | U=12524.0 **$p=5.99 \times 10^{-13}$** CLES:97.24% |
| protein-coding | - | U=12067086.0 $p=6.157 \times 10^{-2}$ CLES:51.57% | U=3954390.5 $p=5.126 \times 10^{-3}$ CLES:54.10% | U=38906080.0 **$p=2.223 \times 10^{-86}$** CLES:60.49% | U=3236110.0 **$p=5.922 \times 10^{-11}$** CLES:61.24% | U=1562822.0 **$p=5.861 \times 10^{-7}$** CLES:62.41% | U=8532796.0 **$p=2.631 \times 10^{-80}$** CLES:71.96% |
| Immune | - | - | U=263439.5 p=0.183 CLES:52.21% | U=2578216.0 **$p=1.385 \times 10^{-17}$** CLES:58.07% | U=212671.5 **$p=1.088 \times 10^{-5}$** CLES:58.30% | U=103391.0 $p=1.656 \times 10^{-4}$ CLES:59.81% | U=563912.5 **$p=1.072 \times 10^{-41}$** CLES:68.89% |
| Vasculature | - | - | - | U=787735.5 **$p=8.562 \times 10^{-6}$** CLES:56.81% | U=65608.0 $p=7.065 \times 10^{-4}$ CLES:57.58% | U=31771.5 $p=2.066 \times 10^{-3}$ CLES:58.84% | U=176762.0 **$p=2.927 \times 10^{-25}$** CLES:69.14% |
| Housekeeping | - | - | - | - | U=504630.5 p=0.849 CLES:50.34% | U=246623.5 p=0.448 CLES:51.92% | U=1416058.0 **$p=1.326 \times 10^{-25}$** CLES:62.95% |
| Glia | - | - | - | - | - | U=20256.0 p=0.529 CLES:51.90% | U=118004.0 **$p=1.438 \times 10^{-11}$** CLES:63.85% |
| ATPase | - | - | - | - | - | - | U=53758.0 **$p=2.993 \times 10^{-5}$** CLES:61.38% |

### Human, Saunders et al., 2018, cell subtypes

| MWU Test | protein-coding | Microglia | Immune | Endothelia | Housekeeping | Astrocyte | Oligodendrocyte | ATPase | Neuron |
|---|---|---|---|---|---|---|---|---|---|
| MHC | U=307170.0<br>**p=$2.294 \times 10^{-7}$**<br>CLES:83.42% | U=4430.5<br>**p=$1.256 \times 10^{-8}$**<br>CLES:88.26% | U=20628.0<br>**p=$1.703 \times 10^{-6}$**<br>CLES:81.15% | U=6210.0<br>**p=$7.177 \times 10^{-9}$**<br>CLES:88.46% | U=64943.0<br>**p=$3.233 \times 10^{-11}$**<br>CLES:92.96% | U=5017.0<br>**p=$2.766 \times 10^{-11}$**<br>CLES:94.66% | U=4815.0<br>**p=$2.625 \times 10^{-11}$**<br>CLES:94.78% | U=2571.0<br>**p=$1.391 \times 10^{-10}$**<br>CLES:94.52% | U=11485.0<br>**p=$8.366 \times 10^{-13}$**<br>CLES:97.00% |
| protein-coding | - | U=2381060.0<br>p=0.407<br>CLES:51.52% | U=12067086.0<br>p=$6.157 \times 10^{-2}$<br>CLES:51.57% | U=3496502.5<br>p=$8.337 \times 10^{-3}$<br>CLES:54.10% | U=38906080.0<br>**p=$2.223 \times 10^{-86}$**<br>CLES:60.49% | U=3027110.5<br>**p=$1.565 \times 10^{-11}$**<br>CLES:62.04% | U=2977352.5<br>**p=$6.776 \times 10^{-14}$**<br>CLES:63.66% | U=1562822.0<br>**p=$5.861 \times 10^{-7}$**<br>CLES:62.41% | U=7769112.0<br>**p=$9.949 \times 10^{-70}$**<br>CLES:71.28% |
| Microglia | - | - | U=160810.5<br>p=0.838<br>CLES:50.41% | U=46592.0<br>p=0.227<br>CLES:52.88% | U=525236.5<br>**p=$1.51 \times 10^{-7}$**<br>CLES:59.91% | U=41107.0<br>**p=$3.542 \times 10^{-6}$**<br>CLES:61.80% | U=40634.0<br>**p=$9.263 \times 10^{-8}$**<br>CLES:63.74% | U=21156.5<br>**p=$9.977 \times 10^{-5}$**<br>CLES:61.98% | U=107062.0<br>**p=$3.85 \times 10^{-24}$**<br>CLES:72.05% |
| Immune | - | - | - | U=232259.0<br>p=0.236<br>CLES:52.06% | U=2578216.0<br>**p=$1.385 \times 10^{-17}$**<br>CLES:58.07% | U=199644.0<br>**p=$1.978 \times 10^{-6}$**<br>CLES:59.27% | U=195840.5<br>**p=$7.767 \times 10^{-8}$**<br>CLES:60.66% | U=103391.0<br>p=$1.656 \times 10^{-4}$<br>CLES:59.81% | U=513226.0<br>**p=$8.33 \times 10^{-37}$**<br>CLES:68.21% |
| Endothelia | - | - | - | - | U=697638.0<br>**p=$1.962 \times 10^{-5}$**<br>CLES:56.90% | U=54479.0<br>p=$2.675 \times 10^{-4}$<br>CLES:58.57% | U=53900.0<br>**p=$1.115 \times 10^{-5}$**<br>CLES:60.46% | U=28142.5<br>p=$2.158 \times 10^{-3}$<br>CLES:58.95% | U=143111.0<br>**p=$3.038 \times 10^{-22}$**<br>CLES:68.87% |
| Housekeeping | - | - | - | - | - | U=474969.5<br>p=0.476<br>CLES:51.31% | U=471841.0<br>p=$8.992 \times 10^{-2}$<br>CLES:53.18% | U=246623.5<br>p=0.448<br>CLES:51.92% | U=1284969.0<br>**p=$3.058 \times 10^{-21}$**<br>CLES:62.14% |
| Astrocyte | - | - | - | - | - | - | U=35147.0<br>p=0.382<br>CLES:52.22% | U=18347.0<br>p=0.766<br>CLES:50.91% | U=96670.0<br>**p=$5.239 \times 10^{-8}$**<br>CLES:61.62% |
| Oligodendrocyte | - | - | - | - | - | - | - | U=16833.5<br>p=0.68<br>CLES:48.73% | U=90093.0<br>**p=$4.735 \times 10^{-6}$**<br>CLES:59.92% |
| ATPase | - | - | - | - | - | - | - | - | U=48764.0<br>p=$1.197 \times 10^{-4}$<br>CLES:60.57% |

### Human, Zeisel et al., 2018, three cell types

| MWU Test | CNS_Glia | CNS_Endothelia | protein-coding | Immune | Housekeeping | CNS_Neuron | ATPase |
|---|---|---|---|---|---|---|---|
| MHC | U=15160.0<br>**$p=1.885\times10^{-8}$**<br>CLES:86.73% | U=2869.0<br>**$p=7.137\times10^{-8}$**<br>CLES:86.94% | U=307170.0<br>**$p=2.294\times10^{-7}$**<br>CLES:83.42% | U=20628.0<br>**$p=1.703\times10^{-6}$**<br>CLES:81.15% | U=64943.0<br>**$p=3.233\times10^{-11}$**<br>CLES:92.96% | U=64215.5<br>**$p=2.793\times10^{-12}$**<br>CLES:95.25% | U=2571.0<br>**$p=1.391\times10^{-10}$**<br>CLES:94.52% |
| CNS_Glia | - | U=75056.0<br>p=0.404<br>CLES:52.05% | U=8268303.5<br>p=0.167<br>CLES:51.38% | U=587803.0<br>$p=2.162\times10^{-2}$<br>CLES:52.91% | U=1925349.0<br>**$p=5.324\times10^{-33}$**<br>CLES:63.07% | U=1976732.0<br>**$p=7.569\times10^{-55}$**<br>CLES:67.09% | U=77656.5<br>**$p=8.47\times10^{-9}$**<br>CLES:65.33% |
| CNS_Endothelia | - | - | U=1507205.5<br>p=0.864<br>CLES:49.61% | U=108068.5<br>p=0.522<br>CLES:51.53% | U=352006.5<br>**$p=1.47\times10^{-6}$**<br>CLES:61.08% | U=362302.0<br>**$p=4.847\times10^{-11}$**<br>CLES:65.14% | U=14187.5<br>**$p=7.876\times10^{-5}$**<br>CLES:63.22% |
| protein-coding | - | - | - | U=12067086.0<br>$p=6.157\times10^{-2}$<br>CLES:51.57% | U=38906080.0<br>**$p=2.223\times10^{-86}$**<br>CLES:60.49% | U=39723834.5<br>**$p=8.63\times10^{-148}$**<br>CLES:64.00% | U=1562822.0<br>**$p=5.861\times10^{-7}$**<br>CLES:62.41% |
| Immune | - | - | - | - | U=2578216.0<br>**$p=1.385\times10^{-17}$**<br>CLES:58.07% | U=2619708.5<br>**$p=9.361\times10^{-32}$**<br>CLES:61.14% | U=103391.0<br>$p=1.656\times10^{-4}$<br>CLES:59.81% |
| Housekeeping | - | - | - | - | - | U=6308982.0<br>**$p=2.807\times10^{-7}$**<br>CLES:53.58% | U=246623.5<br>p=0.448<br>CLES:51.92% |
| CNS_Neuron | - | - | - | - | - | - | U=221488.0<br>p=0.504<br>CLES:48.31% |

### Human, Zeisel et al., 2018, cell subtypes

| MWU Test | Nervous Sys Immune cells | Glia | protein-coding | Immune (benchmark) | Vascular cells | Housekeeping | Neurons | ATPase |
|---|---|---|---|---|---|---|---|---|
| MHC | U=1624.0<br>**p=1.131 × $10^{-5}$**<br>CLES:81.20% | U=12177.0<br>**p=1.749 × $10^{-7}$**<br>CLES:84.21% | U=307170.0<br>**p=2.294 × $10^{-7}$**<br>CLES:83.42% | U=20628.0<br>**p=1.703 × $10^{-6}$**<br>CLES:81.15% | U=3387.0<br>**p=3.933 × $10^{-9}$**<br>CLES:90.08% | U=64943.0<br>**p=3.233 × $10^{-11}$**<br>CLES:92.96% | U=66623.0<br>**p=9.645 × $10^{-12}$**<br>CLES:94.10% | U=2571.0<br>**p=1.391 × $10^{-10}$**<br>CLES:94.52% |
| Nervous Sys Immune cells | - | U=38211.0<br>p=0.355<br>CLES:52.85% | U=1049808.0<br>p=1.534 × $10^{-2}$<br>CLES:57.02% | U=73667.5<br>p=7.956 × $10^{-3}$<br>CLES:57.96% | U=11613.0<br>p=1.009 × $10^{-3}$<br>CLES:61.77% | U=238564.0<br>**p=4.137 × $10^{-10}$**<br>CLES:68.30% | U=245881.0<br>**p=3.007 × $10^{-11}$**<br>CLES:69.46% | U=9555.0<br>**p=1.068 × $10^{-7}$**<br>CLES:70.26% |
| Glia | - | - | U=7285125.0<br>**p=1.572 × $10^{-5}$**<br>CLES:54.73% | U=513435.0<br>**p=1.264 × $10^{-5}$**<br>CLES:55.87% | U=80738.0<br>**p=7.042 × $10^{-5}$**<br>CLES:59.40% | U=1679751.5<br>**p=1.586 × $10^{-44}$**<br>CLES:66.51% | U=1733044.0<br>**p=4.502 × $10^{-51}$**<br>CLES:67.71% | U=67581.5<br>**p=3.983 × $10^{-12}$**<br>CLES:68.73% |
| protein-coding | - | - | - | U=12067086.0<br>p=6.157 × $10^{-2}$<br>CLES:51.57% | U=1848777.0<br>p=0.107<br>CLES:53.41% | U=38906080.0<br>**p=2.223 × $10^{-86}$**<br>CLES:60.49% | U=39966751.0<br>**p=2.833 × $10^{-101}$**<br>CLES:61.32% | U=1562822.0<br>**p=5.861 × $10^{-7}$**<br>CLES:62.41% |
| Immune (benchmark) | - | - | - | - | U=122284.5<br>p=0.602<br>CLES:51.18% | U=2578216.0<br>**p=1.385 × $10^{-17}$**<br>CLES:58.07% | U=2637877.5<br>**p=6.281 × $10^{-20}$**<br>CLES:58.63% | U=103391.0<br>p=1.656 × $10^{-4}$<br>CLES:59.81% |
| Vascular cells | - | - | - | - | - | U=382072.0<br>p=1.536 × $10^{-4}$<br>CLES:58.18% | U=393457.0<br>**p=2.436 × $10^{-5}$**<br>CLES:59.12% | U=15419.0<br>p=1.546 × $10^{-3}$<br>CLES:60.31% |
| Housekeeping | - | - | - | - | - | - | U=6255564.5<br>p=0.391<br>CLES:50.59% | U=246623.5<br>p=0.448<br>CLES:51.92% |
| Neurons | - | - | - | - | - | - | - | U=247190.5<br>p=0.594<br>CLES:51.34% |
