## Supplementary Table S5 for "Neuron-specific coding and regulatory sequences are the most highly conserved in amniote brains despite neuron-specific cell size diversity"

### Chicken Reference Genome, Zhang et al., 2014, cell subtypes

| MWU Test | Microglia vs Neuron | Astrocyte vs Neuron | Oligodendrocyte vs Neuron |
|---|---|---|---|
| Spoon-billed sandpiper | U=301517.0<br>**$p=8.961 \times 10^{-18}$**<br>CLES:63.54% | U=284655.0<br>**$p=6.204 \times 10^{-7}$**<br>CLES:57.77% | U=172843.0<br>$p=1.842 \times 10^{-2}$<br>CLES:54.24% |
| White-throated sparrow | U=234524.0<br>**$p=1.965 \times 10^{-13}$**<br>CLES:62.25% | U=213026.0<br>**$p=1.117 \times 10^{-5}$**<br>CLES:57.35% | U=133470.0<br>$p=1.477 \times 10^{-2}$<br>CLES:54.68% |
| Abingdon island giant tortoise | U=238666.5<br>**$p=2.387 \times 10^{-10}$**<br>CLES:60.42% | U=221739.0<br>**$p=2.466 \times 10^{-6}$**<br>CLES:57.82% | U=137103.5<br>$p=1.447 \times 10^{-2}$<br>CLES:54.66% |
| Japanese quail | U=315011.0<br>**$p=3.558 \times 10^{-11}$**<br>CLES:60.16% | U=291597.0<br>$p=1.01 \times 10^{-4}$<br>CLES:55.98% | U=178046.5<br>p=0.153<br>CLES:52.53% |
| Budgerigar | U=267857.5<br>**$p=1.562 \times 10^{-17}$**<br>CLES:63.81% | U=244416.5<br>**$p=7.528 \times 10^{-7}$**<br>CLES:58.01% | U=147395.5<br>$p=3.796 \times 10^{-4}$<br>CLES:56.74% |
| Turkey | U=226252.5<br>**$p=6.239 \times 10^{-7}$**<br>CLES:58.20% | U=199697.5<br>$p=2.367 \times 10^{-4}$<br>CLES:56.21% | U=128287.5<br>$p=3.512 \times 10^{-2}$<br>CLES:54.05% |
| Tuatara | U=234204.5<br>**$p=3.905 \times 10^{-11}$**<br>CLES:60.94% | U=213880.0<br>**$p=2.231 \times 10^{-6}$**<br>CLES:57.93% | U=131589.0<br>$p=2.56 \times 10^{-2}$<br>CLES:54.30% |
| Bengalese finch | U=297283.0<br>**$p=4.477 \times 10^{-17}$**<br>CLES:63.27% | U=275183.5<br>**$p=2.822 \times 10^{-8}$**<br>CLES:58.78% | U=168129.0<br>$p=1.223 \times 10^{-3}$<br>CLES:55.92% |
| Agassiz's desert tortoise | U=253730.0<br>**$p=8.497 \times 10^{-12}$**<br>CLES:61.10% | U=234342.0<br>**$p=9.884 \times 10^{-5}$**<br>CLES:56.32% | U=137537.0<br>$p=3.438 \times 10^{-2}$<br>CLES:54.04% |
| Common canary | U=266554.0<br>**$p=1.382 \times 10^{-15}$**<br>CLES:62.91% | U=240105.5<br>**$p=4.004 \times 10^{-8}$**<br>CLES:58.98% | U=144193.5<br>$p=1.558 \times 10^{-3}$<br>CLES:56.02% |
| Flycatcher | U=297374.0<br>**$p=1.786 \times 10^{-19}$**<br>CLES:64.31% | U=266707.0<br>**$p=4.751 \times 10^{-7}$**<br>CLES:58.00% | U=156091.0<br>$p=5.848 \times 10^{-3}$<br>CLES:55.15% |
| Painted turtle | U=281274.0<br>**$p=3.003 \times 10^{-13}$**<br>CLES:61.60% | U=273744.0<br>**$p=1.785 \times 10^{-7}$**<br>CLES:58.23% | U=168222.0<br>$p=1.036 \times 10^{-3}$<br>CLES:55.99% |
| Great Tit | U=304204.5<br>**$p=5.901 \times 10^{-19}$**<br>CLES:64.00% | U=274371.5<br>**$p=8.568 \times 10^{-8}$**<br>CLES:58.46% | U=169007.0<br>$p=1.979 \times 10^{-3}$<br>CLES:55.64% |
| Duck | U=260479.5<br>**$p=1.046 \times 10^{-11}$**<br>CLES:60.97% | U=239005.5<br>**$p=4.797 \times 10^{-5}$**<br>CLES:56.58% | U=150497.0<br>$p=1.493 \times 10^{-2}$<br>CLES:54.53% |
| Blue tit | U=241698.0<br>**$p=1.647 \times 10^{-12}$**<br>CLES:61.63% | U=227853.5<br>**$p=6.229 \times 10^{-7}$**<br>CLES:58.21% | U=131219.5<br>$p=3.987 \times 10^{-2}$<br>CLES:53.96% |

| | | | |
|---|---|---|---|
| Pink-footed goose | U=310619.0<br>**p=4.798 × $10^{-14}$**<br>CLES:61.67% | U=288347.0<br>**p=8.397 × $10^{-7}$**<br>CLES:57.64% | U=171825.5<br>p=3.011 × $10^{-2}$<br>CLES:53.89% |
| Okarito brown kiwi | U=281412.5<br>**p=3.92 × $10^{-13}$**<br>CLES:61.50% | U=246696.5<br>**p=1.104 × $10^{-5}$**<br>CLES:57.09% | U=157176.5<br>p=1.536 × $10^{-3}$<br>CLES:55.88% |
| Anole lizard | U=208195.0<br>**p=7.928 × $10^{-6}$**<br>CLES:57.54% | U=213230.0<br>p=7.983 × $10^{-4}$<br>CLES:55.56% | U=126190.0<br>p=9.324 × $10^{-2}$<br>CLES:53.26% |
| Chilean tinamou | U=250588.5<br>**p=2.581 × $10^{-12}$**<br>CLES:61.43% | U=227161.0<br>p=3.077 × $10^{-4}$<br>CLES:55.90% | U=146172.0<br>p=8.906 × $10^{-3}$<br>CLES:54.91% |
| Emu | U=289256.0<br>**p=8.57 × $10^{-15}$**<br>CLES:62.27% | U=274747.5<br>**p=4.964 × $10^{-9}$**<br>CLES:59.25% | U=162510.5<br>p=6.815 × $10^{-3}$<br>CLES:54.96% |
| Australian saltwater crocodile | U=230625.0<br>**p=7.872 × $10^{-18}$**<br>CLES:64.55% | U=223440.5<br>**p=8.423 × $10^{-7}$**<br>CLES:58.15% | U=139953.5<br>p=5.197 × $10^{-4}$<br>CLES:56.62% |
| Ruff | U=293258.0<br>**p=1.14 × $10^{-15}$**<br>CLES:62.64% | U=276084.0<br>**p=1.596 × $10^{-7}$**<br>CLES:58.24% | U=169692.5<br>p=5.514 × $10^{-4}$<br>CLES:56.29% |
| Blue-crowned manakin | U=296655.0<br>**p=2.037 × $10^{-17}$**<br>CLES:63.41% | U=274308.0<br>**p=1.513 × $10^{-8}$**<br>CLES:58.95% | U=166527.5<br>p=1.639 × $10^{-3}$<br>CLES:55.76% |
| Argentine black and white tegu | U=266090.5<br>**p=5 × $10^{-7}$**<br>CLES:58.00% | U=278555.0<br>**p=2.859 × $10^{-8}$**<br>CLES:58.74% | U=173934.0<br>p=1.273 × $10^{-3}$<br>CLES:55.82% |
| Dark-eyed junco | U=262962.0<br>**p=3.95 × $10^{-17}$**<br>CLES:63.69% | U=221647.0<br>**p=6.447 × $10^{-6}$**<br>CLES:57.49% | U=144287.0<br>**p=9.345 × $10^{-5}$**<br>CLES:57.48% |
| Golden-collared manakin | U=236114.0<br>**p=8.52 × $10^{-17}$**<br>CLES:63.95% | U=229023.0<br>**p=1.519 × $10^{-8}$**<br>CLES:59.36% | U=138194.0<br>**p=2.851 × $10^{-5}$**<br>CLES:58.11% |
| Great spotted kiwi | U=265560.0<br>**p=3.706 × $10^{-14}$**<br>CLES:62.21% | U=243101.5<br>**p=5.356 × $10^{-7}$**<br>CLES:58.13% | U=146157.0<br>p=1.301 × $10^{-3}$<br>CLES:56.09% |
| Zebra Finch | U=199805.0<br>**p=1.428 × $10^{-17}$**<br>CLES:64.96% | U=174314.0<br>**p=2.39 × $10^{-7}$**<br>CLES:59.18% | U=108172.5<br>p=4.482 × $10^{-3}$<br>CLES:55.80% |
| Chinese softshell turtle | U=229205.5<br>**p=2.14 × $10^{-14}$**<br>CLES:62.87% | U=235006.5<br>**p=9.454 × $10^{-10}$**<br>CLES:60.08% | U=134074.0<br>p=7.931 × $10^{-4}$<br>CLES:56.51% |
| Little spotted kiwi | U=276821.0<br>**p=2.614 × $10^{-14}$**<br>CLES:62.13% | U=245526.5<br>**p=3.68 × $10^{-7}$**<br>CLES:58.24% | U=151548.0<br>p=1.144 × $10^{-3}$<br>CLES:56.09% |
| Helmeted guineafowl | U=314248.5<br>**p=2.039 × $10^{-10}$**<br>CLES:59.75% | U=284961.5<br>p=3.706 × $10^{-3}$<br>CLES:54.45% | U=175432.0<br>p=0.205<br>CLES:52.25% |

### Human Reference Genome, Zhang et al., 2014, cell subtypes

| MWU Test | Microglia vs Neuron | Astrocyte vs Neuron | Oligodendrocyte vs Neuron |
|---|---|---|---|
| Microbat | U=425182.5<br>**p=9.582 × $10^{-25}$**<br>CLES:64.80% | U=329914.0<br>**p=9.172 × $10^{-9}$**<br>CLES:58.67% | U=192014.5<br>p=3.397 × $10^{-3}$<br>CLES:55.18% |
| Hedgehog | U=30284.0<br>**p=3.859 × $10^{-7}$**<br>CLES:64.07% | U=16438.0<br>p=3.117 × $10^{-2}$<br>CLES:56.89% | U=11049.5<br>p=0.486<br>CLES:52.48% |
| Naked mole-rat male | U=511009.0<br>**p=7.775 × $10^{-36}$**<br>CLES:67.37% | U=395045.5<br>**p=3.894 × $10^{-17}$**<br>CLES:62.34% | U=236206.0<br>**p=7.389 × $10^{-6}$**<br>CLES:57.59% |
| Sooty mangabey | U=649276.0<br>**p=1.653 × $10^{-27}$**<br>CLES:64.05% | U=513680.0<br>**p=2.995 × $10^{-14}$**<br>CLES:60.34% | U=298110.0<br>p=2.224 × $10^{-3}$<br>CLES:54.83% |
| Olive baboon | U=645887.5<br>**p=5.503 × $10^{-25}$**<br>CLES:63.32% | U=515902.0<br>**p=2.219 × $10^{-16}$**<br>CLES:61.20% | U=300453.0<br>p=3.085 × $10^{-4}$<br>CLES:55.71% |
| American bison | U=428571.5<br>**p=2.255 × $10^{-36}$**<br>CLES:68.31% | U=310167.5<br>**p=3.242 × $10^{-14}$**<br>CLES:61.75% | U=187455.0<br>**p=1.338 × $10^{-5}$**<br>CLES:57.77% |
| Leopard | U=633556.0<br>**p=1.883 × $10^{-44}$**<br>CLES:68.52% | U=493698.0<br>**p=2.992 × $10^{-17}$**<br>CLES:61.67% | U=274906.5<br>p=8.943 × $10^{-4}$<br>CLES:55.38% |
| Bolivian squirrel monkey | U=623809.0<br>**p=5.261 × $10^{-38}$**<br>CLES:67.01% | U=474635.5<br>**p=5.424 × $10^{-17}$**<br>CLES:61.70% | U=268822.5<br>p=3.558 × $10^{-3}$<br>CLES:54.72% |
| Crab-eating macaque | U=642136.0<br>**p=9.07 × $10^{-25}$**<br>CLES:63.28% | U=509624.5<br>**p=1.007 × $10^{-14}$**<br>CLES:60.56% | U=291060.5<br>p=6.77 × $10^{-3}$<br>CLES:54.29% |
| Red fox | U=556986.0<br>**p=1.016 × $10^{-29}$**<br>CLES:65.26% | U=398180.0<br>**p=9.131 × $10^{-9}$**<br>CLES:58.27% | U=237793.0<br>p=2.577 × $10^{-2}$<br>CLES:53.70% |
| Shrew mouse | U=667504.0<br>**p=1.694 × $10^{-48}$**<br>CLES:69.17% | U=503646.5<br>**p=8.341 × $10^{-16}$**<br>CLES:61.04% | U=293076.0<br>**p=1.388 × $10^{-5}$**<br>CLES:56.98% |
| Arctic ground squirrel | U=647865.5<br>**p=3.23 × $10^{-48}$**<br>CLES:69.25% | U=503512.5<br>**p=1.617 × $10^{-19}$**<br>CLES:62.46% | U=287892.0<br>**p=6.951 × $10^{-5}$**<br>CLES:56.38% |
| Chinese hamster CriGri | U=552188.0<br>**p=1.622 × $10^{-40}$**<br>CLES:68.21% | U=413026.0<br>**p=6.941 × $10^{-16}$**<br>CLES:61.65% | U=233196.0<br>p=5.896 × $10^{-4}$<br>CLES:55.79% |
| Opossum | U=416229.0<br>**p=1.961 × $10^{-12}$**<br>CLES:60.02% | U=362265.0<br>**p=1.967 × $10^{-9}$**<br>CLES:58.85% | U=212764.0<br>p=1.397 × $10^{-2}$<br>CLES:54.20% |
| Dolphin | U=161765.0<br>**p=1.042 × $10^{-17}$** | U=105034.5<br>**p=3.681 × $10^{-5}$** | U=65058.0<br>p=2.653 × $10^{-2}$ |

| | | | |
|---|---|---|---|
| | CLES:65.73% | CLES:58.28% | CLES:55.11% |
| Capuchin | U=661717.0<br>**p=$1.354 \times 10^{-42}$**<br>CLES:67.86% | U=508418.5<br>**p=$2.336 \times 10^{-21}$**<br>CLES:63.10% | U=295837.5<br>**p=$3.991 \times 10^{-5}$**<br>CLES:56.54% |
| Mouse Lemur | U=626018.0<br>**p=$1.755 \times 10^{-43}$**<br>CLES:68.34% | U=477414.5<br>**p=$5.202 \times 10^{-21}$**<br>CLES:63.21% | U=275882.0<br>p=$7.509 \times 10^{-4}$<br>CLES:55.44% |
| Damara mole rat | U=512538.0<br>**p=$3.867 \times 10^{-37}$**<br>CLES:67.74% | U=416046.5<br>**p=$1.407 \times 10^{-18}$**<br>CLES:62.74% | U=250639.0<br>**p=$2.461 \times 10^{-6}$**<br>CLES:57.85% |
| Marmoset | U=629053.5<br>**p=$3.555 \times 10^{-37}$**<br>CLES:66.78% | U=486929.5<br>**p=$1.907 \times 10^{-16}$**<br>CLES:61.41% | U=284185.5<br>p=$3.28 \times 10^{-4}$<br>CLES:55.77% |
| Goat | U=629316.0<br>**p=$1.067 \times 10^{-37}$**<br>CLES:66.91% | U=480243.0<br>**p=$2.605 \times 10^{-15}$**<br>CLES:60.98% | U=286357.0<br>p=$1.434 \times 10^{-4}$<br>CLES:56.10% |
| Common wombat | U=409566.0<br>**p=$7.558 \times 10^{-18}$**<br>CLES:62.40% | U=343615.0<br>**p=$1.218 \times 10^{-11}$**<br>CLES:60.17% | U=204361.0<br>**p=$6.748 \times 10^{-5}$**<br>CLES:56.96% |
| American black bear | U=451585.0<br>**p=$1.2 \times 10^{-21}$**<br>CLES:63.45% | U=318082.5<br>**p=$5.6 \times 10^{-10}$**<br>CLES:59.47% | U=184778.5<br>p=$3.275 \times 10^{-2}$<br>CLES:53.77% |
| Sloth | U=8182.0<br>**p=$5.753 \times 10^{-6}$**<br>CLES:67.71% | U=6125.0<br>p=0.137<br>CLES:55.97% | U=3901.0<br>p=0.256<br>CLES:55.19% |
| Polar bear | U=440306.5<br>**p=$5.279 \times 10^{-29}$**<br>CLES:66.00% | U=312391.5<br>**p=$7.648 \times 10^{-8}$**<br>CLES:58.18% | U=179877.0<br>p=$7.503 \times 10^{-2}$<br>CLES:53.15% |
| Alpine marmot | U=576363.0<br>**p=$9.648 \times 10^{-41}$**<br>CLES:68.03% | U=413474.5<br>**p=$9.54 \times 10^{-13}$**<br>CLES:60.23% | U=234713.5<br>p=$2.437 \times 10^{-3}$<br>CLES:55.08% |
| Prairie vole | U=635955.0<br>**p=$7.141 \times 10^{-47}$**<br>CLES:69.05% | U=467118.5<br>**p=$1.402 \times 10^{-16}$**<br>CLES:61.60% | U=280202.0<br>**p=$3.745 \times 10^{-6}$**<br>CLES:57.53% |
| Donkey | U=643817.0<br>**p=$2.411 \times 10^{-34}$**<br>CLES:65.94% | U=471978.0<br>**p=$1.644 \times 10^{-12}$**<br>CLES:59.81% | U=279004.5<br>p=$9.869 \times 10^{-4}$<br>CLES:55.31% |
| Golden Hamster | U=569052.0<br>**p=$8.191 \times 10^{-45}$**<br>CLES:69.15% | U=431139.0<br>**p=$6.949 \times 10^{-16}$**<br>CLES:61.55% | U=254746.5<br>**p=$6.661 \times 10^{-7}$**<br>CLES:58.33% |
| Armadillo | U=512552.0<br>**p=$1.521 \times 10^{-31}$**<br>CLES:66.14% | U=398959.5<br>**p=$1.362 \times 10^{-13}$**<br>CLES:60.72% | U=234494.0<br>p=$6.993 \times 10^{-4}$<br>CLES:55.68% |
| Ugandan red Colobus | U=648910.0<br>**p=$7.511 \times 10^{-37}$**<br>CLES:66.56% | U=490866.0<br>**p=$4.459 \times 10^{-15}$**<br>CLES:60.83% | U=295961.0<br>p=$1.766 \times 10^{-4}$<br>CLES:55.96% |
| Pig-tailed macaque | U=648175.5<br>**p=$8.366 \times 10^{-26}$** | U=508668.0<br>**p=$3.821 \times 10^{-14}$** | U=297864.5<br>p=$7.871 \times 10^{-3}$ |

| | | | |
|---|---|---|---|
| | CLES:63.56% | CLES:60.33% | CLES:54.18% |
| Panda | U=618350.5<br>**$p=1.06 \times 10^{-38}$**<br>CLES:67.22% | U=448520.0<br>**$p=4.164 \times 10^{-14}$**<br>CLES:60.66% | U=272493.5<br>$p=1.124 \times 10^{-4}$<br>CLES:56.27% |
| Ferret | U=615581.5<br>**$p=3.843 \times 10^{-36}$**<br>CLES:66.59% | U=447822.0<br>**$p=3.28 \times 10^{-14}$**<br>CLES:60.71% | U=262216.0<br>$p=1.951 \times 10^{-3}$<br>CLES:55.06% |
| Koala | U=476321.0<br>**$p=2.225 \times 10^{-18}$**<br>CLES:62.14% | U=396262.0<br>**$p=3.937 \times 10^{-12}$**<br>CLES:60.07% | U=239474.5<br>$p=3.328 \times 10^{-4}$<br>CLES:56.01% |
| Wallaby | U=43323.0<br>$p=1.738 \times 10^{-3}$<br>CLES:57.73% | U=28603.0<br>$p=3.822 \times 10^{-3}$<br>CLES:58.04% | U=21340.5<br>$p=0.107$<br>CLES:54.83% |
| Tarsier | U=403572.0<br>**$p=1.555 \times 10^{-26}$**<br>CLES:65.59% | U=309816.0<br>**$p=2.943 \times 10^{-10}$**<br>CLES:59.68% | U=171903.0<br>$p=1.638 \times 10^{-2}$<br>CLES:54.33% |
| Degu | U=510418.5<br>**$p=8.927 \times 10^{-37}$**<br>CLES:67.69% | U=403785.5<br>**$p=5.338 \times 10^{-17}$**<br>CLES:62.24% | U=246444.5<br>**$p=2.378 \times 10^{-5}$**<br>CLES:57.07% |
| Chinese hamster CHOK1GS | U=677528.5<br>**$p=1.155 \times 10^{-48}$**<br>CLES:69.13% | U=517588.0<br>**$p=1.298 \times 10^{-17}$**<br>CLES:61.67% | U=297947.0<br>**$p=1.963 \times 10^{-5}$**<br>CLES:56.82% |
| Drill | U=594062.5<br>**$p=3.08 \times 10^{-22}$**<br>CLES:62.73% | U=459226.5<br>**$p=1.348 \times 10^{-12}$**<br>CLES:59.90% | U=257438.5<br>$p=4.418 \times 10^{-2}$<br>CLES:53.27% |
| Gibbon | U=579647.5<br>**$p=3.495 \times 10^{-20}$**<br>CLES:62.14% | U=456039.5<br>**$p=1.774 \times 10^{-10}$**<br>CLES:58.89% | U=265368.5<br>$p=1.422 \times 10^{-2}$<br>CLES:53.97% |
| Rat | U=629522.5<br>**$p=6.37 \times 10^{-45}$**<br>CLES:68.64% | U=471394.0<br>**$p=9.44 \times 10^{-16}$**<br>CLES:61.21% | U=272148.0<br>$p=1.223 \times 10^{-4}$<br>CLES:56.25% |
| Macaque | U=649810.5<br>**$p=8.779 \times 10^{-31}$**<br>CLES:64.97% | U=511676.5<br>**$p=9.354 \times 10^{-16}$**<br>CLES:60.98% | U=297739.5<br>$p=1.224 \times 10^{-4}$<br>CLES:56.11% |
| American mink | U=591865.0<br>**$p=3.536 \times 10^{-38}$**<br>CLES:67.28% | U=439137.0<br>**$p=1.404 \times 10^{-14}$**<br>CLES:60.91% | U=254395.0<br>$p=1.354 \times 10^{-3}$<br>CLES:55.26% |
| Gelada | U=666914.0<br>**$p=4.732 \times 10^{-29}$**<br>CLES:64.39% | U=529838.0<br>**$p=1.218 \times 10^{-17}$**<br>CLES:61.62% | U=306023.0<br>$p=4.733 \times 10^{-4}$<br>CLES:55.51% |
| Sheep | U=575695.5<br>**$p=1.104 \times 10^{-37}$**<br>CLES:67.30% | U=425415.0<br>**$p=2.913 \times 10^{-12}$**<br>CLES:59.95% | U=246974.0<br>$p=5.007 \times 10^{-3}$<br>CLES:54.64% |
| Tasmanian devil | U=389129.0<br>**$p=5.854 \times 10^{-18}$**<br>CLES:62.61% | U=315606.5<br>**$p=1.976 \times 10^{-12}$**<br>CLES:60.83% | U=193764.5<br>**$p=8.386 \times 10^{-5}$**<br>CLES:56.95% |
| Elephant | U=588182.5<br>**$p=1.024 \times 10^{-35}$** | U=441714.5<br>**$p=2.387 \times 10^{-12}$** | U=257473.0<br>$p=5.948 \times 10^{-3}$ |

| | | | |
|---|---|---|---|
| | CLES:66.69% | CLES:59.88% | CLES:54.49% |
| Algerian mouse | U=681549.0 | U=513922.0 | U=300365.5 |
| | **$p=3.387 \times 10^{-47}$** | **$p=1.259 \times 10^{-15}$** | **$p=6.077 \times 10^{-6}$** |
| | CLES:68.77% | CLES:60.91% | CLES:57.23% |
| Dingo | U=593367.5 | U=434397.5 | U=250507.5 |
| | **$p=1.185 \times 10^{-33}$** | **$p=1.813 \times 10^{-11}$** | $p=1.094 \times 10^{-2}$ |
| | CLES:66.10% | CLES:59.51% | CLES:54.19% |
| Golden snub-nosed monkey | U=584604.0 | U=457389.0 | U=268022.0 |
| | **$p=3.11 \times 10^{-25}$** | **$p=1.716 \times 10^{-14}$** | $p=5.832 \times 10^{-4}$ |
| | CLES:63.74% | CLES:60.75% | CLES:55.58% |
| Horse | U=657677.5 | U=491592.5 | U=283856.5 |
| | **$p=2.703 \times 10^{-41}$** | **$p=3.321 \times 10^{-15}$** | $p=2.844 \times 10^{-4}$ |
| | CLES:67.59% | CLES:60.89% | CLES:55.86% |
| Black snub-nosed monkey | U=576628.5 | U=445075.5 | U=273971.5 |
| | **$p=6.163 \times 10^{-19}$** | **$p=1.175 \times 10^{-10}$** | $p=1.064 \times 10^{-2}$ |
| | CLES:61.72% | CLES:59.05% | CLES:54.09% |
| Brazilian guinea pig | U=313142.5 | U=244713.0 | U=155732.0 |
| | **$p=6.511 \times 10^{-22}$** | **$p=1.26 \times 10^{-11}$** | $p=4.384 \times 10^{-4}$ |
| | CLES:64.97% | CLES:61.12% | CLES:56.52% |
| Orangutan | U=561174.5 | U=447876.0 | U=270268.0 |
| | **$p=2.947 \times 10^{-14}$** | **$p=9.56 \times 10^{-9}$** | $p=2.309 \times 10^{-2}$ |
| | CLES:60.02% | CLES:58.00% | CLES:53.65% |
| Northern American deer | U=613269.0 | U=468362.0 | U=273737.0 |
| mouse | **$p=2.546 \times 10^{-45}$** | **$p=1.059 \times 10^{-16}$** | **$p=3.119 \times 10^{-6}$** |
| | CLES:68.85% | CLES:61.59% | CLES:57.60% |
| Naked mole-rat female | U=593527.0 | U=456671.5 | U=275461.5 |
| | **$p=1.735 \times 10^{-40}$** | **$p=4.889 \times 10^{-18}$** | **$p=7.372 \times 10^{-6}$** |
| | CLES:67.90% | CLES:62.25% | CLES:57.31% |
| Mouse | U=723163.5 | U=539483.0 | U=312250.5 |
| | **$p=5.191 \times 10^{-48}$** | **$p=1.442 \times 10^{-16}$** | **$p=7.219 \times 10^{-6}$** |
| | CLES:68.65% | CLES:61.15% | CLES:57.10% |
| Daurian ground squirrel | U=414552.0 | U=301015.5 | U=174113.0 |
| | **$p=1.075 \times 10^{-33}$** | **$p=3.958 \times 10^{-11}$** | $p=3.183 \times 10^{-3}$ |
| | CLES:67.70% | CLES:60.24% | CLES:55.32% |
| Tiger | U=378872.5 | U=295015.5 | U=161696.0 |
| | **$p=2.731 \times 10^{-34}$** | **$p=2.448 \times 10^{-11}$** | $p=9.121 \times 10^{-4}$ |
| | CLES:68.33% | CLES:60.40% | CLES:56.13% |
| Lesser hedgehog tenrec | U=28016.0 | U=18199.0 | U=10086.0 |
| | **$p=5.479 \times 10^{-10}$** | $p=2.768 \times 10^{-3}$ | $p=2.507 \times 10^{-2}$ |
| | CLES:67.80% | CLES:59.33% | CLES:58.46% |
| Mongolian gerbil | U=500036.5 | U=356412.5 | U=212723.5 |
| | **$p=4.784 \times 10^{-38}$** | **$p=1.936 \times 10^{-14}$** | $p=1.202 \times 10^{-4}$ |
| | CLES:68.03% | CLES:61.46% | CLES:56.64% |
| Ma's night monkey | U=626241.5 | U=482392.0 | U=281643.0 |
| | **$p=8.648 \times 10^{-35}$** | **$p=7.414 \times 10^{-15}$** | $p=6.799 \times 10^{-4}$ |
| | CLES:66.19% | CLES:60.78% | CLES:55.46% |
| Upper Galilee mountains | U=609907.5 | U=469921.0 | U=286718.0 |
| blind mole rat | **$p=3.664 \times 10^{-44}$** | **$p=6.331 \times 10^{-17}$** | **$p=1.109 \times 10^{-6}$** |

| | | | |
|---|---|---|---|
| | CLES:68.67% | CLES:61.73% | CLES:57.89% |
| Kangaroo rat | U=491470.0 | U=393684.0 | U=235500.5 |
| | **p=$2.524 \times 10^{-33}$** | **p=$6.472 \times 10^{-16}$** | p=$1.257 \times 10^{-3}$ |
| | CLES:66.89% | CLES:61.85% | CLES:55.41% |
| Chimpanzee | U=536984.0 | U=448968.5 | U=261096.5 |
| | **p=$7.344 \times 10^{-11}$** | **p=$2.482 \times 10^{-8}$** | p=0.16 |
| | CLES:58.60% | CLES:57.72% | CLES:52.25% |
| Guinea Pig | U=596336.5 | U=450317.0 | U=262755.0 |
| | **p=$1.272 \times 10^{-40}$** | **p=$8.407 \times 10^{-18}$** | p=$2.069 \times 10^{-4}$ |
| | CLES:67.87% | CLES:62.18% | CLES:56.07% |
| American beaver | U=539023.5 | U=371927.5 | U=219637.0 |
| | **p=$7.116 \times 10^{-34}$** | **p=$6.242 \times 10^{-13}$** | p=$3.457 \times 10^{-4}$ |
| | CLES:66.55% | CLES:60.63% | CLES:56.14% |
| Wild yak | U=460787.5 | U=344658.0 | U=198598.5 |
| | **p=$2.535 \times 10^{-34}$** | **p=$5.042 \times 10^{-15}$** | p=$3.636 \times 10^{-4}$ |
| | CLES:67.38% | CLES:61.80% | CLES:56.23% |
| Coquerel's sifaka | U=600871.0 | U=472434.0 | U=261509.5 |
| | **p=$4.18 \times 10^{-38}$** | **p=$1.279 \times 10^{-19}$** | p=$2.126 \times 10^{-3}$ |
| | CLES:67.23% | CLES:62.72% | CLES:55.03% |
| Alpaca | U=12924.0 | U=14167.0 | U=5414.0 |
| | **p=$1.024 \times 10^{-8}$** | p=$6.325 \times 10^{-3}$ | p=$1.512 \times 10^{-2}$ |
| | CLES:70.21% | CLES:58.97% | CLES:60.89% |
| Lesser Egyptian jerboa | U=506029.0 | U=405920.0 | U=247320.0 |
| | **p=$2.218 \times 10^{-39}$** | **p=$3.792 \times 10^{-19}$** | **p=$6.932 \times 10^{-7}$** |
| | CLES:68.40% | CLES:63.07% | CLES:58.33% |
| Long-tailed chinchilla | U=604070.0 | U=463806.0 | U=273784.0 |
| | **p=$1.48 \times 10^{-38}$** | **p=$7 \times 10^{-16}$** | p=$4.619 \times 10^{-4}$ |
| | CLES:67.32% | CLES:61.32% | CLES:55.66% |
| Angola colobus | U=605510.0 | U=464151.5 | U=277631.5 |
| | **p=$3.479 \times 10^{-26}$** | **p=$2.253 \times 10^{-12}$** | p=$9.424 \times 10^{-4}$ |
| | CLES:63.91% | CLES:59.77% | CLES:55.32% |
| Ryukyu mouse | U=678183.0 | U=507050.0 | U=292115.0 |
| | **p=$2.966 \times 10^{-49}$** | **p=$1.547 \times 10^{-15}$** | **p=$1.202 \times 10^{-5}$** |
| | CLES:69.26% | CLES:60.92% | CLES:57.05% |
| Chinese hamster PICR | U=664247.0 | U=497233.5 | U=284100.5 |
| | **p=$1.018 \times 10^{-50}$** | **p=$1.668 \times 10^{-17}$** | **p=$4.422 \times 10^{-5}$** |
| | CLES:69.67% | CLES:61.75% | CLES:56.60% |
| Vervet-AGM | U=644058.0 | U=499369.0 | U=296763.5 |
| | **p=$9.665 \times 10^{-26}$** | **p=$1.726 \times 10^{-13}$** | p=$1.821 \times 10^{-3}$ |
| | CLES:63.55% | CLES:60.09% | CLES:54.92% |
| Cow | U=636625.5 | U=467379.0 | U=283502.0 |
| | **p=$2.121 \times 10^{-42}$** | **p=$1.954 \times 10^{-14}$** | p=$1.055 \times 10^{-4}$ |
| | CLES:68.00% | CLES:60.69% | CLES:56.23% |
| Bushbaby | U=673312.5 | U=493688.0 | U=296495.5 |
| | **p=$3.171 \times 10^{-48}$** | **p=$1.067 \times 10^{-16}$** | **p=$1.877 \times 10^{-5}$** |
| | CLES:69.06% | CLES:61.48% | CLES:56.84% |
| Bonobo | U=511353.0 | U=427081.5 | U=253010.0 |
| | **p=$5.962 \times 10^{-5}$** | **p=$1.223 \times 10^{-5}$** | p=0.529 |

| | | | |
|---|---|---|---|
| | CLES:55.29% | CLES:56.10% | CLES:51.01% |
| Hyrax | U=51137.0<br>**p=7.225 × $10^{-7}$**<br>CLES:61.95% | U=25107.0<br>p=2.792 × $10^{-2}$<br>CLES:56.30% | U=17310.0<br>p=0.352<br>CLES:52.96% |
| Rabbit | U=451450.0<br>**p=7.397 × $10^{-24}$**<br>CLES:64.27% | U=353177.0<br>**p=3.971 × $10^{-8}$**<br>CLES:58.12% | U=200005.0<br>p=0.167<br>CLES:52.38% |
| Megabat | U=156110.5<br>**p=1.889 × $10^{-17}$**<br>CLES:65.73% | U=100544.0<br>**p=5.85 × $10^{-8}$**<br>CLES:61.12% | U=58903.0<br>p=0.188<br>CLES:53.07% |
| Dog | U=557173.5<br>**p=7.817 × $10^{-28}$**<br>CLES:64.73% | U=434081.5<br>**p=3.114 × $10^{-10}$**<br>CLES:58.87% | U=244601.5<br>p=2.471 × $10^{-2}$<br>CLES:53.71% |
| Steppe mouse | U=635609.5<br>**p=3.225 × $10^{-41}$**<br>CLES:67.72% | U=474689.0<br>**p=2.176 × $10^{-14}$**<br>CLES:60.61% | U=275972.0<br>**p=2.936 × $10^{-6}$**<br>CLES:57.65% |
| Greater bamboo lemur | U=634860.5<br>**p=5.777 × $10^{-36}$**<br>CLES:66.42% | U=491711.5<br>**p=2.887 × $10^{-18}$**<br>CLES:62.08% | U=283868.5<br>p=2.487 × $10^{-4}$<br>CLES:55.88% |
| Pig | U=654037.0<br>**p=1.175 × $10^{-45}$**<br>CLES:68.61% | U=482498.0<br>**p=1.181 × $10^{-16}$**<br>CLES:61.51% | U=281537.0<br>p=1.24 × $10^{-4}$<br>CLES:56.19% |
| Tree Shrew | U=18406.0<br>**p=1.946 × $10^{-5}$**<br>CLES:63.38% | U=11098.0<br>p=8.13 × $10^{-2}$<br>CLES:56.05% | U=7305.0<br>p=6.053 × $10^{-2}$<br>CLES:57.50% |
| Pika | U=70857.0<br>**p=1.721 × $10^{-13}$**<br>CLES:66.68% | U=45042.0<br>**p=1.03 × $10^{-5}$**<br>CLES:61.03% | U=25385.0<br>p=0.121<br>CLES:54.50% |
| Shrew | U=15987.0<br>**p=5.019 × $10^{-6}$**<br>CLES:64.90% | U=9517.0<br>p=2.025 × $10^{-2}$<br>CLES:58.53% | U=6300.0<br>p=2.813 × $10^{-2}$<br>CLES:59.29% |
| Squirrel | U=624497.0<br>**p=4.506 × $10^{-42}$**<br>CLES:68.04% | U=490012.5<br>**p=1.231 × $10^{-17}$**<br>CLES:61.86% | U=280007.0<br>**p=9.262 × $10^{-5}$**<br>CLES:56.33% |
| Gorilla | U=555270.0<br>**p=4.291 × $10^{-10}$**<br>CLES:58.17% | U=462743.0<br>**p=8.83 × $10^{-8}$**<br>CLES:57.36% | U=277182.5<br>p=6.876 × $10^{-3}$<br>CLES:54.32% |
| Cat | U=639878.0<br>**p=5.955 × $10^{-40}$**<br>CLES:67.38% | U=485432.0<br>**p=8.901 × $10^{-15}$**<br>CLES:60.71% | U=273848.5<br>p=2.1 × $10^{-3}$<br>CLES:54.97% |

### Rat Reference Genome, Zhang et al., 2014, cell subtypes

| MWU Test | Microglia vs Neuron | Astrocyte vs Neuron | Oligodendrocyte vs Neuron |
|---|---|---|---|
| Microbat | U=392519.0<br>**p=$7.665 \times 10^{-30}$**<br>CLES:66.81% | U=303011.0<br>**p=$3.642 \times 10^{-11}$**<br>CLES:60.26% | U=169321.0<br>p=$3.039 \times 10^{-3}$<br>CLES:55.42% |
| Hedgehog | U=27910.0<br>**p=$1.071 \times 10^{-9}$**<br>CLES:67.49% | U=14311.0<br>p=$4.861 \times 10^{-2}$<br>CLES:56.52% | U=9501.5<br>p=0.993<br>CLES:50.03% |
| Naked mole-rat male | U=468518.0<br>**p=$1.017 \times 10^{-39}$**<br>CLES:68.84% | U=351908.5<br>**p=$4.92 \times 10^{-14}$**<br>CLES:61.31% | U=213551.0<br>**p=$1.856 \times 10^{-5}$**<br>CLES:57.42% |
| Sooty mangabey | U=604720.0<br>**p=$5.369 \times 10^{-45}$**<br>CLES:68.87% | U=454942.5<br>**p=$2.549 \times 10^{-15}$**<br>CLES:61.14% | U=261257.5<br>p=$2.646 \times 10^{-4}$<br>CLES:56.00% |
| Olive baboon | U=602692.5<br>**p=$6.483 \times 10^{-45}$**<br>CLES:68.86% | U=451696.5<br>**p=$1.143 \times 10^{-16}$**<br>CLES:61.72% | U=266199.5<br>**p=$2.366 \times 10^{-5}$**<br>CLES:56.93% |
| American bison | U=367250.5<br>**p=$9.799 \times 10^{-35}$**<br>CLES:68.60% | U=267270.5<br>**p=$5.608 \times 10^{-11}$**<br>CLES:60.48% | U=159001.0<br>p=$9.254 \times 10^{-4}$<br>CLES:56.11% |
| Leopard | U=544015.0<br>**p=$4.432 \times 10^{-41}$**<br>CLES:68.46% | U=425718.5<br>**p=$6.555 \times 10^{-14}$**<br>CLES:60.70% | U=238034.0<br>p=$2.33 \times 10^{-3}$<br>CLES:55.10% |
| Bolivian squirrel monkey | U=557023.0<br>**p=$7.733 \times 10^{-46}$**<br>CLES:69.49% | U=419021.0<br>**p=$7.457 \times 10^{-16}$**<br>CLES:61.61% | U=239460.0<br>p=$5.333 \times 10^{-4}$<br>CLES:55.81% |
| Crab-eating macaque | U=604557.0<br>**p=$1.42 \times 10^{-43}$**<br>CLES:68.52% | U=446992.5<br>**p=$3.383 \times 10^{-15}$**<br>CLES:61.14% | U=253912.5<br>p=$1.218 \times 10^{-3}$<br>CLES:55.34% |
| Human | U=632872.5<br>**p=$4.314 \times 10^{-45}$**<br>CLES:68.66% | U=473061.0<br>**p=$6.843 \times 10^{-16}$**<br>CLES:61.26% | U=273115.0<br>p=$1.083 \times 10^{-4}$<br>CLES:56.30% |
| Red fox | U=502309.5<br>**p=$2.414 \times 10^{-36}$**<br>CLES:67.57% | U=354363.5<br>**p=$1.589 \times 10^{-10}$**<br>CLES:59.52% | U=215535.0<br>p=$1.729 \times 10^{-2}$<br>CLES:54.04% |
| Shrew mouse | U=725592.0<br>**p=$1.375 \times 10^{-47}$**<br>CLES:68.51% | U=509602.5<br>**p=$7.096 \times 10^{-13}$**<br>CLES:59.77% | U=303412.5<br>p=$6.404 \times 10^{-3}$<br>CLES:54.27% |
| Arctic ground squirrel | U=603841.0<br>**p=$6.743 \times 10^{-52}$**<br>CLES:70.44% | U=449047.0<br>**p=$1.373 \times 10^{-16}$**<br>CLES:61.70% | U=259699.0<br>p=$2.502 \times 10^{-4}$<br>CLES:56.01% |
| Chinese hamster CriGri | U=623020.5<br>**p=$7.582 \times 10^{-52}$**<br>CLES:70.20% | U=421360.0<br>**p=$2.244 \times 10^{-14}$**<br>CLES:60.94% | U=248487.5<br>p=$1.535 \times 10^{-3}$<br>CLES:55.23% |
| Opossum | U=403884.0<br>**p=$2.011 \times 10^{-14}$** | U=334796.0<br>**p=$9.763 \times 10^{-8}$** | U=203524.0<br>p=$1.561 \times 10^{-2}$ |

| | | | |
|---|---|---|---|
| | CLES:61.02% | CLES:58.00% | CLES:54.18% |
| Dolphin | U=160338.0 | U=109103.0 | U=61585.0 |
| | **p=$2.452 \times 10^{-21}$** | **p=$9.344 \times 10^{-7}$** | p=$1.587 \times 10^{-2}$ |
| | CLES:67.57% | CLES:59.78% | CLES:55.66% |
| Capuchin | U=588156.0 | U=437150.0 | U=255984.5 |
| | **p=$6.101 \times 10^{-50}$** | **p=$2.185 \times 10^{-17}$** | p=$1.711 \times 10^{-4}$ |
| | CLES:70.15% | CLES:62.11% | CLES:56.20% |
| Mouse Lemur | U=548419.0 | U=408353.0 | U=241099.5 |
| | **p=$5.268 \times 10^{-42}$** | **p=$2.706 \times 10^{-15}$** | p=$1.662 \times 10^{-3}$ |
| | CLES:68.64% | CLES:61.46% | CLES:55.24% |
| Damara mole rat | U=467088.0 | U=368300.5 | U=220892.5 |
| | **p=$3.11 \times 10^{-41}$** | **p=$2.107 \times 10^{-16}$** | **p=$1.474 \times 10^{-5}$** |
| | CLES:69.28% | CLES:62.24% | CLES:57.44% |
| Marmoset | U=589523.5 | U=439243.0 | U=255938.0 |
| | **p=$2.201 \times 10^{-47}$** | **p=$3.68 \times 10^{-15}$** | p=$4.751 \times 10^{-4}$ |
| | CLES:69.55% | CLES:61.16% | CLES:55.75% |
| Goat | U=549238.5 | U=423001.0 | U=242504.5 |
| | **p=$4.257 \times 10^{-41}$** | **p=$3.215 \times 10^{-16}$** | p=$2.77 \times 10^{-4}$ |
| | CLES:68.41% | CLES:61.73% | CLES:56.08% |
| Common wombat | U=401460.5 | U=318784.5 | U=200648.0 |
| | **p=$8.517 \times 10^{-22}$** | **p=$1.415 \times 10^{-10}$** | **p=$3.618 \times 10^{-5}$** |
| | CLES:63.99% | CLES:59.81% | CLES:57.25% |
| American black bear | U=399412.0 | U=281902.0 | U=163191.0 |
| | **p=$8.474 \times 10^{-29}$** | **p=$3.544 \times 10^{-13}$** | p=$3.929 \times 10^{-3}$ |
| | CLES:66.34% | CLES:61.53% | CLES:55.27% |
| Sloth | U=8668.0 | U=5798.5 | U=4215.5 |
| | **p=$3.319 \times 10^{-6}$** | p=$9.95 \times 10^{-3}$ | p=$7.149 \times 10^{-2}$ |
| | CLES:67.90% | CLES:60.75% | CLES:58.18% |
| Polar bear | U=400542.0 | U=286166.0 | U=163723.0 |
| | **p=$5.735 \times 10^{-32}$** | **p=$7.829 \times 10^{-9}$** | p=$6.673 \times 10^{-2}$ |
| | CLES:67.33% | CLES:59.01% | CLES:53.32% |
| Alpine marmot | U=548908.5 | U=388285.5 | U=225680.5 |
| | **p=$8.028 \times 10^{-48}$** | **p=$8.364 \times 10^{-13}$** | p=$2.583 \times 10^{-4}$ |
| | CLES:69.99% | CLES:60.43% | CLES:56.21% |
| Prairie vole | U=676340.5 | U=481311.0 | U=294156.5 |
| | **p=$1.255 \times 10^{-49}$** | **p=$4.67 \times 10^{-17}$** | **p=$7.99 \times 10^{-5}$** |
| | CLES:69.32% | CLES:61.70% | CLES:56.28% |
| Donkey | U=582782.0 | U=426717.0 | U=245683.0 |
| | **p=$4.11 \times 10^{-41}$** | **p=$5.048 \times 10^{-15}$** | p=$1.806 \times 10^{-3}$ |
| | CLES:68.10% | CLES:61.20% | CLES:55.18% |
| Golden Hamster | U=613388.5 | U=434011.0 | U=266877.5 |
| | **p=$5.856 \times 10^{-46}$** | **p=$1.698 \times 10^{-13}$** | **p=$4.981 \times 10^{-5}$** |
| | CLES:69.00% | CLES:60.48% | CLES:56.63% |
| Armadillo | U=443594.0 | U=347406.5 | U=200954.0 |
| | **p=$1.267 \times 10^{-32}$** | **p=$1.069 \times 10^{-13}$** | p=$2.216 \times 10^{-3}$ |
| | CLES:67.10% | CLES:61.17% | CLES:55.32% |
| Ugandan red Colobus | U=587470.5 | U=434814.0 | U=261243.5 |
| | **p=$3.253 \times 10^{-47}$** | **p=$1.419 \times 10^{-14}$** | **p=$5.16 \times 10^{-5}$** |

| | | | |
|---|---|---|---|
| | CLES:69.54% | CLES:60.95% | CLES:56.66% |
| Pig-tailed macaque | U=603298.0<br>**$p=1.845 \times 10^{-41}$**<br>CLES:68.04% | U=447027.0<br>**$p=3.062 \times 10^{-14}$**<br>CLES:60.73% | U=262020.5<br>$p=6.824 \times 10^{-4}$<br>CLES:55.57% |
| Panda | U=548180.5<br>**$p=8.945 \times 10^{-44}$**<br>CLES:69.06% | U=399546.5<br>**$p=2.914 \times 10^{-15}$**<br>CLES:61.51% | U=238493.0<br>$p=6.847 \times 10^{-4}$<br>CLES:55.68% |
| Ferret | U=537937.0<br>**$p=1.33 \times 10^{-42}$**<br>CLES:68.87% | U=393423.5<br>**$p=2.44 \times 10^{-14}$**<br>CLES:61.13% | U=224031.5<br>$p=2.72 \times 10^{-3}$<br>CLES:55.10% |
| Koala | U=461780.0<br>**$p=8.119 \times 10^{-21}$**<br>CLES:63.14% | U=371872.0<br>**$p=4.498 \times 10^{-11}$**<br>CLES:59.70% | U=235623.0<br>$p=1.066 \times 10^{-4}$<br>CLES:56.52% |
| Wallaby | U=40823.0<br>$p=5.957 \times 10^{-4}$<br>CLES:58.64% | U=25343.0<br>$p=2.074 \times 10^{-2}$<br>CLES:56.60% | U=20719.5<br>$p=7.452 \times 10^{-2}$<br>CLES:55.40% |
| Tarsier | U=384972.5<br>**$p=5.582 \times 10^{-37}$**<br>CLES:69.04% | U=276912.5<br>**$p=3.432 \times 10^{-10}$**<br>CLES:59.93% | U=155239.0<br>$p=3.772 \times 10^{-2}$<br>CLES:53.84% |
| Degu | U=454313.0<br>**$p=1.453 \times 10^{-38}$**<br>CLES:68.76% | U=354662.0<br>**$p=2.714 \times 10^{-13}$**<br>CLES:60.97% | U=223475.0<br>**$p=3.401 \times 10^{-5}$**<br>CLES:57.10% |
| Chinese hamster CHOK1GS | U=740194.0<br>**$p=3.871 \times 10^{-54}$**<br>CLES:69.79% | U=525956.5<br>**$p=6.385 \times 10^{-17}$**<br>CLES:61.36% | U=313451.0<br>$p=1.162 \times 10^{-4}$<br>CLES:56.03% |
| Drill | U=560639.0<br>**$p=1.298 \times 10^{-42}$**<br>CLES:68.65% | U=408019.5<br>**$p=1.922 \times 10^{-14}$**<br>CLES:61.07% | U=232012.0<br>$p=5.05 \times 10^{-3}$<br>CLES:54.71% |
| Gibbon | U=545468.0<br>**$p=2.584 \times 10^{-42}$**<br>CLES:68.74% | U=409576.0<br>**$p=9.033 \times 10^{-14}$**<br>CLES:60.75% | U=234753.0<br>$p=1.17 \times 10^{-3}$<br>CLES:55.46% |
| Macaque | U=642960.0<br>**$p=2.441 \times 10^{-46}$**<br>CLES:68.86% | U=475225.5<br>**$p=1.035 \times 10^{-15}$**<br>CLES:61.18% | U=272826.0<br>$p=1.22 \times 10^{-3}$<br>CLES:55.24% |
| American mink | U=501295.5<br>**$p=4.778 \times 10^{-37}$**<br>CLES:67.78% | U=380444.0<br>**$p=5.965 \times 10^{-13}$**<br>CLES:60.56% | U=218307.5<br>$p=8.105 \times 10^{-3}$<br>CLES:54.50% |
| Gelada | U=618106.0<br>**$p=1.323 \times 10^{-44}$**<br>CLES:68.67% | U=461816.5<br>**$p=1.407 \times 10^{-15}$**<br>CLES:61.21% | U=268511.5<br>$p=1.35 \times 10^{-4}$<br>CLES:56.24% |
| Sheep | U=505488.5<br>**$p=5.311 \times 10^{-38}$**<br>CLES:68.01% | U=376317.5<br>**$p=2.439 \times 10^{-11}$**<br>CLES:59.80% | U=215112.5<br>$p=1.927 \times 10^{-2}$<br>CLES:54.00% |
| Tasmanian devil | U=370501.5<br>**$p=2.783 \times 10^{-19}$**<br>CLES:63.30% | U=291247.0<br>**$p=1.122 \times 10^{-11}$**<br>CLES:60.66% | U=184799.0<br>**$p=4.229 \times 10^{-5}$**<br>CLES:57.32% |
| Elephant | U=523509.0<br>**$p=2.69 \times 10^{-38}$** | U=390364.0<br>**$p=2.294 \times 10^{-12}$** | U=230204.0<br>$p=6.843 \times 10^{-3}$ |

| | | | |
|---|---|---|---|
| | CLES:67.91% | CLES:60.22% | CLES:54.54% |
| Algerian mouse | U=731043.0<br>**$p=3.635 \times 10^{-46}$**<br>CLES:68.16% | U=511829.0<br>**$p=1.246 \times 10^{-11}$**<br>CLES:59.19% | U=305653.0<br>$p=6.823 \times 10^{-3}$<br>CLES:54.23% |
| Dingo | U=539576.5<br>**$p=6.573 \times 10^{-40}$**<br>CLES:68.17% | U=396517.0<br>**$p=3.289 \times 10^{-14}$**<br>CLES:61.05% | U=225449.0<br>$p=1.89 \times 10^{-3}$<br>CLES:55.28% |
| Golden snub-nosed monkey | U=547894.0<br>**$p=4.983 \times 10^{-44}$**<br>CLES:69.12% | U=412071.0<br>**$p=7.765 \times 10^{-15}$**<br>CLES:61.20% | U=240435.0<br>$p=1.071 \times 10^{-4}$<br>CLES:56.49% |
| Horse | U=596646.0<br>**$p=1.75 \times 10^{-41}$**<br>CLES:68.10% | U=436733.5<br>**$p=3.122 \times 10^{-12}$**<br>CLES:59.88% | U=256898.5<br>$p=1.479 \times 10^{-4}$<br>CLES:56.29% |
| Black snub-nosed monkey | U=545413.0<br>**$p=3.168 \times 10^{-39}$**<br>CLES:67.96% | U=399888.5<br>**$p=1.044 \times 10^{-11}$**<br>CLES:59.84% | U=244888.5<br>$p=2.89 \times 10^{-3}$<br>CLES:54.92% |
| Brazilian guinea pig | U=279984.0<br>**$p=3.556 \times 10^{-22}$**<br>CLES:65.52% | U=215818.5<br>**$p=1.509 \times 10^{-10}$**<br>CLES:60.83% | U=142473.0<br>$p=2.178 \times 10^{-4}$<br>CLES:57.00% |
| Orangutan | U=560589.5<br>**$p=1.364 \times 10^{-45}$**<br>CLES:69.39% | U=416999.0<br>**$p=2.108 \times 10^{-15}$**<br>CLES:61.42% | U=236923.0<br>$p=8.201 \times 10^{-4}$<br>CLES:55.62% |
| Northern American deer mouse | U=657856.0<br>**$p=1.684 \times 10^{-50}$**<br>CLES:69.63% | U=472107.5<br>**$p=1.063 \times 10^{-15}$**<br>CLES:61.17% | U=281016.0<br>**$p=9.412 \times 10^{-5}$**<br>CLES:56.27% |
| Naked mole-rat female | U=532519.5<br>**$p=4.486 \times 10^{-41}$**<br>CLES:68.56% | U=398788.0<br>**$p=3.703 \times 10^{-14}$**<br>CLES:61.02% | U=241836.0<br>**$p=8.923 \times 10^{-5}$**<br>CLES:56.56% |
| Mouse | U=785083.5<br>**$p=8.813 \times 10^{-50}$**<br>CLES:68.58% | U=548245.0<br>**$p=6.347 \times 10^{-14}$**<br>CLES:60.04% | U=319423.0<br>$p=7.945 \times 10^{-3}$<br>CLES:54.10% |
| Daurian ground squirrel | U=399611.0<br>**$p=6.082 \times 10^{-39}$**<br>CLES:69.39% | U=279293.0<br>**$p=5.114 \times 10^{-10}$**<br>CLES:59.80% | U=168742.5<br>$p=1.436 \times 10^{-3}$<br>CLES:55.79% |
| Tiger | U=345640.0<br>**$p=1.18 \times 10^{-36}$**<br>CLES:69.52% | U=269274.0<br>**$p=7.165 \times 10^{-10}$**<br>CLES:59.78% | U=148810.0<br>$p=1.029 \times 10^{-3}$<br>CLES:56.18% |
| Lesser hedgehog tenrec | U=24441.0<br>**$p=1.641 \times 10^{-7}$**<br>CLES:65.40% | U=16896.0<br>$p=1.407 \times 10^{-3}$<br>CLES:60.18% | U=9285.0<br>p=0.154<br>CLES:55.38% |
| Mongolian gerbil | U=550813.5<br>**$p=5.441 \times 10^{-42}$**<br>CLES:68.55% | U=365096.0<br>**$p=3.366 \times 10^{-12}$**<br>CLES:60.31% | U=225591.5<br>$p=1.718 \times 10^{-3}$<br>CLES:55.28% |
| Ma's night monkey | U=565184.5<br>**$p=1.908 \times 10^{-46}$**<br>CLES:69.56% | U=429959.0<br>**$p=1.456 \times 10^{-15}$**<br>CLES:61.40% | U=243963.0<br>$p=3.017 \times 10^{-4}$<br>CLES:56.04% |
| Upper Galilee mountains blind mole rat | U=623629.0<br>**$p=1.928 \times 10^{-44}$** | U=447988.0<br>**$p=1.401 \times 10^{-13}$** | U=284646.0<br>**$p=4.481 \times 10^{-5}$** |

| | | | |
|---|---|---|---|
| | CLES:68.60% | CLES:60.46% | CLES:56.56% |
| Kangaroo rat | U=445347.0 | U=354355.5 | U=215760.0 |
| | **$p=1.391 \times 10^{-35}$** | **$p=1.702 \times 10^{-14}$** | $p=8.268 \times 10^{-4}$ |
| | CLES:68.01% | CLES:61.53% | CLES:55.73% |
| Chimpanzee | U=567028.0 | U=436900.0 | U=258592.0 |
| | **$p=5.142 \times 10^{-46}$** | **$p=1.04 \times 10^{-16}$** | **$p=2.046 \times 10^{-6}$** |
| | CLES:69.46% | CLES:61.84% | CLES:57.89% |
| Guinea Pig | U=542452.0 | U=399081.0 | U=233801.5 |
| | **$p=6.548 \times 10^{-43}$** | **$p=2.878 \times 10^{-15}$** | $p=4.545 \times 10^{-4}$ |
| | CLES:68.90% | CLES:61.50% | CLES:55.90% |
| American beaver | U=508527.5 | U=341436.0 | U=208361.0 |
| | **$p=7.284 \times 10^{-35}$** | **$p=2.541 \times 10^{-10}$** | $p=3.057 \times 10^{-3}$ |
| | CLES:67.08% | CLES:59.50% | CLES:55.11% |
| Wild yak | U=400229.5 | U=295318.5 | U=166837.0 |
| | **$p=2.191 \times 10^{-36}$** | **$p=9.186 \times 10^{-13}$** | $p=2.925 \times 10^{-3}$ |
| | CLES:68.65% | CLES:61.16% | CLES:55.41% |
| Coquerel's sifaka | U=524393.0 | U=398455.0 | U=225813.0 |
| | **$p=5.992 \times 10^{-42}$** | **$p=1.464 \times 10^{-15}$** | $p=2.821 \times 10^{-3}$ |
| | CLES:68.86% | CLES:61.64% | CLES:55.07% |
| Alpaca | U=14410.0 | U=17120.5 | U=6645.0 |
| | **$p=6.775 \times 10^{-10}$** | $p=1.491 \times 10^{-3}$ | $p=6.746 \times 10^{-3}$ |
| | CLES:71.39% | CLES:60.00% | CLES:61.60% |
| Lesser Egyptian jerboa | U=496301.0 | U=389945.5 | U=237364.5 |
| | **$p=4.233 \times 10^{-42}$** | **$p=2.836 \times 10^{-18}$** | **$p=6.346 \times 10^{-6}$** |
| | CLES:69.19% | CLES:62.85% | CLES:57.61% |
| Long-tailed chinchilla | U=543426.5 | U=408907.5 | U=245155.0 |
| | **$p=5.025 \times 10^{-43}$** | **$p=6.463 \times 10^{-15}$** | $p=1.65 \times 10^{-4}$ |
| | CLES:68.94% | CLES:61.28% | CLES:56.28% |
| Angola colobus | U=565022.0 | U=416336.5 | U=246754.5 |
| | **$p=1.648 \times 10^{-45}$** | **$p=9.484 \times 10^{-15}$** | **$p=8.023 \times 10^{-5}$** |
| | CLES:69.32% | CLES:61.14% | CLES:56.57% |
| Ryukyu mouse | U=721739.5 | U=508989.0 | U=299120.5 |
| | **$p=3.275 \times 10^{-46}$** | **$p=3.055 \times 10^{-12}$** | $p=1.061 \times 10^{-2}$ |
| | CLES:68.24% | CLES:59.49% | CLES:54.01% |
| Chinese hamster PICR | U=730224.5 | U=502592.5 | U=297433.0 |
| | **$p=2.8 \times 10^{-60}$** | **$p=3.822 \times 10^{-15}$** | $p=3.376 \times 10^{-4}$ |
| | CLES:71.08% | CLES:60.78% | CLES:55.68% |
| Vervet-AGM | U=607926.0 | U=458923.0 | U=260681.0 |
| | **$p=1.912 \times 10^{-49}$** | **$p=2.268 \times 10^{-18}$** | $p=1.623 \times 10^{-4}$ |
| | CLES:69.85% | CLES:62.33% | CLES:56.19% |
| Cow | U=561720.5 | U=405904.5 | U=239949.0 |
| | **$p=7.222 \times 10^{-49}$** | **$p=2.079 \times 10^{-14}$** | $p=1.639 \times 10^{-4}$ |
| | CLES:70.15% | CLES:61.08% | CLES:56.33% |
| Bushbaby | U=610578.0 | U=437265.5 | U=263097.0 |
| | **$p=7.883 \times 10^{-51}$** | **$p=2.941 \times 10^{-15}$** | **$p=4.85 \times 10^{-5}$** |
| | CLES:70.14% | CLES:61.25% | CLES:56.68% |
| Bonobo | U=568825.0 | U=423081.5 | U=245333.5 |
| | **$p=6.044 \times 10^{-44}$** | **$p=1.347 \times 10^{-15}$** | $p=6.682 \times 10^{-4}$ |

| | | | |
|---|---|---|---|
| | CLES:68.90% | CLES:61.45% | CLES:55.64% |
| Hyrax | U=51197.0<br>**$p=9.61 \times 10^{-9}$**<br>CLES:63.94% | U=26234.0<br>$p=1.384 \times 10^{-2}$<br>CLES:56.99% | U=17051.5<br>p=0.668<br>CLES:51.36% |
| Rabbit | U=427676.0<br>**$p=1.97 \times 10^{-25}$**<br>CLES:65.01% | U=319906.0<br>**$p=1.712 \times 10^{-6}$**<br>CLES:57.23% | U=186966.0<br>p=0.435<br>CLES:51.36% |
| Megabat | U=167223.0<br>**$p=1.799 \times 10^{-23}$**<br>CLES:68.35% | U=110840.0<br>**$p=1.141 \times 10^{-10}$**<br>CLES:63.02% | U=64546.0<br>$p=6.983 \times 10^{-2}$<br>CLES:54.16% |
| Dog | U=525356.5<br>**$p=1.63 \times 10^{-33}$**<br>CLES:66.61% | U=407069.0<br>**$p=1.384 \times 10^{-11}$**<br>CLES:59.71% | U=226812.5<br>$p=3.694 \times 10^{-2}$<br>CLES:53.51% |
| Steppe mouse | U=694077.5<br>**$p=1.166 \times 10^{-42}$**<br>CLES:67.61% | U=478327.0<br>**$p=2.278 \times 10^{-10}$**<br>CLES:58.72% | U=288762.5<br>$p=1.678 \times 10^{-3}$<br>CLES:54.99% |
| Greater bamboo lemur | U=563006.5<br>**$p=7.264 \times 10^{-40}$**<br>CLES:67.96% | U=428410.5<br>**$p=7.303 \times 10^{-16}$**<br>CLES:61.53% | U=248371.5<br>$p=1.073 \times 10^{-4}$<br>CLES:56.44% |
| Pig | U=576536.5<br>**$p=5.366 \times 10^{-43}$**<br>CLES:68.62% | U=427467.0<br>**$p=3.991 \times 10^{-15}$**<br>CLES:61.24% | U=248778.0<br>$p=1.54 \times 10^{-3}$<br>CLES:55.24% |
| Tree Shrew | U=20505.5<br>**$p=5.302 \times 10^{-9}$**<br>CLES:68.12% | U=12457.5<br>$p=2.507 \times 10^{-2}$<br>CLES:57.61% | U=7613.0<br>$p=4.723 \times 10^{-2}$<br>CLES:57.90% |
| Pika | U=66477.5<br>**$p=1.543 \times 10^{-13}$**<br>CLES:67.00% | U=41068.5<br>**$p=2.734 \times 10^{-5}$**<br>CLES:60.73% | U=24468.5<br>$p=9.308 \times 10^{-2}$<br>CLES:54.92% |
| Shrew | U=16474.5<br>**$p=4.729 \times 10^{-7}$**<br>CLES:66.41% | U=9672.0<br>$p=2.888 \times 10^{-3}$<br>CLES:61.00% | U=5525.5<br>p=0.245<br>CLES:54.99% |
| Squirrel | U=578993.0<br>**$p=3.59 \times 10^{-46}$**<br>CLES:69.39% | U=436177.5<br>**$p=1.014 \times 10^{-14}$**<br>CLES:61.01% | U=257082.5<br>$p=1.392 \times 10^{-4}$<br>CLES:56.29% |
| Gorilla | U=577627.5<br>**$p=8.142 \times 10^{-45}$**<br>CLES:69.07% | U=439645.5<br>**$p=6.147 \times 10^{-16}$**<br>CLES:61.50% | U=253833.0<br>$p=1.963 \times 10^{-4}$<br>CLES:56.16% |
| Cat | U=585170.0<br>**$p=3.832 \times 10^{-41}$**<br>CLES:68.10% | U=439703.0<br>**$p=5.141 \times 10^{-13}$**<br>CLES:60.20% | U=248800.0<br>$p=8.127 \times 10^{-3}$<br>CLES:54.36% |

### Mouse Reference Genome, Zhang et al., 2014, cell subtypes

| MWU Test | Microglia vs Neuron | Astrocyte vs Neuron | Oligodendrocyte vs Neuron |
|---|---|---|---|
| Microbat | U=430043.0<br>**$p=8.301 \times 10^{-30}$**<br>CLES:66.40% | U=329337.0<br>**$p=6.019 \times 10^{-10}$**<br>CLES:59.37% | U=191738.0<br>$p=6.858 \times 10^{-4}$<br>CLES:56.02% |
| Hedgehog | U=30823.0<br>**$p=1.103 \times 10^{-9}$**<br>CLES:67.01% | U=15068.0<br>$p=4.181 \times 10^{-2}$<br>CLES:56.66% | U=9786.5<br>$p=0.63$<br>CLES:48.27% |
| Naked mole-rat male | U=526352.0<br>**$p=4.475 \times 10^{-43}$**<br>CLES:69.12% | U=393951.5<br>**$p=1.544 \times 10^{-15}$**<br>CLES:61.67% | U=237765.0<br>**$p=2.905 \times 10^{-6}$**<br>CLES:57.93% |
| Sooty mangabey | U=682906.5<br>**$p=2.055 \times 10^{-48}$**<br>CLES:69.03% | U=512533.0<br>**$p=2.754 \times 10^{-16}$**<br>CLES:61.19% | U=298818.0<br>**$p=1.082 \times 10^{-5}$**<br>CLES:57.03% |
| Olive baboon | U=681648.0<br>**$p=1.421 \times 10^{-47}$**<br>CLES:68.85% | U=507956.0<br>**$p=2.829 \times 10^{-17}$**<br>CLES:61.61% | U=302070.0<br>**$p=1.759 \times 10^{-6}$**<br>CLES:57.63% |
| American bison | U=414826.0<br>**$p=4.671 \times 10^{-36}$**<br>CLES:68.38% | U=301166.0<br>**$p=4.713 \times 10^{-13}$**<br>CLES:61.26% | U=181116.5<br>**$p=2.359 \times 10^{-5}$**<br>CLES:57.61% |
| Leopard | U=611818.0<br>**$p=2.157 \times 10^{-44}$**<br>CLES:68.69% | U=480357.0<br>**$p=4.635 \times 10^{-16}$**<br>CLES:61.28% | U=271479.0<br>**$p=8.322 \times 10^{-5}$**<br>CLES:56.42% |
| Bolivian squirrel monkey | U=630822.0<br>**$p=1.896 \times 10^{-50}$**<br>CLES:69.89% | U=472555.0<br>**$p=1.599 \times 10^{-17}$**<br>CLES:61.92% | U=272950.0<br>**$p=1.772 \times 10^{-5}$**<br>CLES:57.02% |
| Crab-eating macaque | U=679590.5<br>**$p=9.191 \times 10^{-47}$**<br>CLES:68.68% | U=505208.0<br>**$p=1.906 \times 10^{-16}$**<br>CLES:61.30% | U=292860.0<br>**$p=3.976 \times 10^{-5}$**<br>CLES:56.58% |
| Human | U=724827.5<br>**$p=2.984 \times 10^{-48}$**<br>CLES:68.69% | U=540828.0<br>**$p=1.04 \times 10^{-16}$**<br>CLES:61.20% | U=313065.5<br>**$p=6.149 \times 10^{-6}$**<br>CLES:57.16% |
| Red fox | U=557989.0<br>**$p=2.954 \times 10^{-36}$**<br>CLES:67.06% | U=391354.0<br>**$p=4.458 \times 10^{-10}$**<br>CLES:59.05% | U=239276.0<br>$p=3.438 \times 10^{-3}$<br>CLES:54.87% |
| Shrew mouse | U=826654.0<br>**$p=9.309 \times 10^{-48}$**<br>CLES:67.91% | U=591854.0<br>**$p=8.485 \times 10^{-16}$**<br>CLES:60.59% | U=344439.5<br>$p=5.579 \times 10^{-3}$<br>CLES:54.20% |
| Arctic ground squirrel | U=680172.0<br>**$p=3.043 \times 10^{-57}$**<br>CLES:70.90% | U=503320.0<br>**$p=4.505 \times 10^{-19}$**<br>CLES:62.30% | U=293073.5<br>**$p=1.579 \times 10^{-5}$**<br>CLES:56.91% |
| Chinese hamster CriGri | U=694009.0<br>**$p=2.277 \times 10^{-53}$**<br>CLES:69.95% | U=472600.5<br>**$p=6.177 \times 10^{-14}$**<br>CLES:60.43% | U=273804.5<br>$p=4.351 \times 10^{-3}$<br>CLES:54.59% |
| Opossum | U=443282.5<br>**$p=7.515 \times 10^{-13}$** | U=372957.5<br>**$p=1.531 \times 10^{-7}$** | U=225767.0<br>$p=1.47 \times 10^{-2}$ |

| | | | |
|---|---|---|---|
| | CLES:60.05% | CLES:57.65% | CLES:54.11% |
| Dolphin | U=179396.0<br>**$p=1.661 \times 10^{-23}$**<br>CLES:68.03% | U=121319.5<br>**$p=2.56 \times 10^{-8}$**<br>CLES:60.88% | U=71306.0<br>$p=3.946 \times 10^{-3}$<br>CLES:56.54% |
| Capuchin | U=663474.5<br>**$p=2.263 \times 10^{-53}$**<br>CLES:70.25% | U=498376.0<br>**$p=2.886 \times 10^{-19}$**<br>CLES:62.42% | U=293788.5<br>**$p=2.989 \times 10^{-6}$**<br>CLES:57.50% |
| Mouse Lemur | U=617676.5<br>**$p=4.035 \times 10^{-47}$**<br>CLES:69.25% | U=459372.0<br>**$p=1.509 \times 10^{-17}$**<br>CLES:62.03% | U=270692.5<br>$p=2.33 \times 10^{-4}$<br>CLES:55.98% |
| Damara mole rat | U=525356.5<br>**$p=4.131 \times 10^{-43}$**<br>CLES:69.15% | U=409952.5<br>**$p=5.457 \times 10^{-16}$**<br>CLES:61.73% | U=249902.0<br>**$p=3.81 \times 10^{-7}$**<br>CLES:58.51% |
| Marmoset | U=660537.0<br>**$p=1.638 \times 10^{-49}$**<br>CLES:69.44% | U=497057.0<br>**$p=1.888 \times 10^{-16}$**<br>CLES:61.34% | U=289892.5<br>**$p=8.675 \times 10^{-5}$**<br>CLES:56.29% |
| Goat | U=616759.0<br>**$p=1.635 \times 10^{-42}$**<br>CLES:68.19% | U=470768.0<br>**$p=9.27 \times 10^{-17}$**<br>CLES:61.63% | U=277637.0<br>**$p=4.18 \times 10^{-5}$**<br>CLES:56.64% |
| Common wombat | U=441263.0<br>**$p=1.731 \times 10^{-21}$**<br>CLES:63.53% | U=355166.0<br>**$p=1.579 \times 10^{-10}$**<br>CLES:59.51% | U=223486.0<br>**$p=8.965 \times 10^{-6}$**<br>CLES:57.60% |
| American black bear | U=449047.0<br>**$p=1.148 \times 10^{-28}$**<br>CLES:65.81% | U=318077.0<br>**$p=6.247 \times 10^{-13}$**<br>CLES:61.06% | U=185932.0<br>$p=3.705 \times 10^{-3}$<br>CLES:55.14% |
| Sloth | U=10241.0<br>**$p=3.756 \times 10^{-7}$**<br>CLES:68.82% | U=6843.0<br>$p=4.752 \times 10^{-3}$<br>CLES:61.32% | U=4748.0<br>$p=4.459 \times 10^{-2}$<br>CLES:58.91% |
| Polar bear | U=449285.0<br>**$p=1.556 \times 10^{-32}$**<br>CLES:66.98% | U=321285.5<br>**$p=7.94 \times 10^{-10}$**<br>CLES:59.34% | U=184267.5<br>$p=2.093 \times 10^{-2}$<br>CLES:54.08% |
| Alpine marmot | U=611725.5<br>**$p=1.862 \times 10^{-50}$**<br>CLES:70.01% | U=426965.0<br>**$p=2.404 \times 10^{-13}$**<br>CLES:60.43% | U=249137.0<br>**$p=8.223 \times 10^{-5}$**<br>CLES:56.55% |
| Prairie vole | U=765498.5<br>**$p=8.246 \times 10^{-54}$**<br>CLES:69.55% | U=540340.0<br>**$p=7.658 \times 10^{-17}$**<br>CLES:61.27% | U=329798.0<br>**$p=1.735 \times 10^{-5}$**<br>CLES:56.67% |
| Donkey | U=643882.0<br>**$p=8.664 \times 10^{-42}$**<br>CLES:67.80% | U=471428.5<br>**$p=3.977 \times 10^{-13}$**<br>CLES:60.09% | U=279105.5<br>$p=2.325 \times 10^{-4}$<br>CLES:55.95% |
| Golden Hamster | U=693927.0<br>**$p=5.636 \times 10^{-54}$**<br>CLES:70.11% | U=493203.0<br>**$p=1.34 \times 10^{-15}$**<br>CLES:61.03% | U=302591.0<br>**$p=1.328 \times 10^{-6}$**<br>CLES:57.71% |
| Armadillo | U=501814.0<br>**$p=2.232 \times 10^{-34}$**<br>CLES:67.04% | U=394341.0<br>**$p=8.428 \times 10^{-16}$**<br>CLES:61.75% | U=225076.5<br>**$p=9.649 \times 10^{-5}$**<br>CLES:56.65% |
| Ugandan red Colobus | U=663370.5<br>**$p=6.025 \times 10^{-50}$** | U=488073.0<br>**$p=9.941 \times 10^{-16}$** | U=291858.5<br>**$p=1.859 \times 10^{-5}$** |

| | | | |
|---|---|---|---|
| | CLES:69.51% | CLES:61.11% | CLES:56.87% |
| Pig-tailed macaque | U=681214.5 | U=502659.0 | U=302224.0 |
| | **$p=2.514 \times 10^{-46}$** | **$p=8.111 \times 10^{-15}$** | **$p=1.729 \times 10^{-5}$** |
| | CLES:68.58% | CLES:60.65% | CLES:56.83% |
| Panda | U=603121.0 | U=439190.0 | U=265113.0 |
| | **$p=1.401 \times 10^{-43}$** | **$p=2.124 \times 10^{-15}$** | **$p=3.337 \times 10^{-5}$** |
| | CLES:68.53% | CLES:61.29% | CLES:56.80% |
| Ferret | U=595748.0 | U=439360.5 | U=258695.0 |
| | **$p=5.966 \times 10^{-45}$** | **$p=8.262 \times 10^{-16}$** | **$p=7.927 \times 10^{-5}$** |
| | CLES:68.93% | CLES:61.46% | CLES:56.52% |
| Koala | U=510019.0 | U=409517.0 | U=267400.0 |
| | **$p=6.016 \times 10^{-20}$** | **$p=4.577 \times 10^{-11}$** | **$p=2.53 \times 10^{-6}$** |
| | CLES:62.50% | CLES:59.47% | CLES:57.71% |
| Wallaby | U=46311.0 | U=30144.0 | U=24378.5 |
| | $p=1.579 \times 10^{-4}$ | $p=3.417 \times 10^{-3}$ | $p=2.4 \times 10^{-2}$ |
| | CLES:59.24% | CLES:58.06% | CLES:56.60% |
| Tarsier | U=423720.0 | U=305079.5 | U=175837.0 |
| | **$p=1.435 \times 10^{-40}$** | **$p=2.973 \times 10^{-11}$** | $p=1.517 \times 10^{-3}$ |
| | CLES:69.56% | CLES:60.29% | CLES:55.73% |
| Degu | U=517161.0 | U=406313.0 | U=252788.5 |
| | **$p=1.151 \times 10^{-43}$** | **$p=1.192 \times 10^{-15}$** | **$p=6.171 \times 10^{-6}$** |
| | CLES:69.44% | CLES:61.66% | CLES:57.54% |
| Chinese hamster CHOK1GS | U=828486.5 | U=587524.5 | U=346307.5 |
| | **$p=3.667 \times 10^{-58}$** | **$p=6.301 \times 10^{-16}$** | $p=4.717 \times 10^{-4}$ |
| | CLES:69.98% | CLES:60.66% | CLES:55.33% |
| Drill | U=627429.5 | U=454892.5 | U=263119.5 |
| | **$p=2.902 \times 10^{-43}$** | **$p=3.196 \times 10^{-14}$** | $p=5.235 \times 10^{-4}$ |
| | CLES:68.25% | CLES:60.66% | CLES:55.67% |
| Gibbon | U=620418.0 | U=465275.5 | U=267497.0 |
| | **$p=6.64 \times 10^{-47}$** | **$p=8.401 \times 10^{-16}$** | $p=1.016 \times 10^{-4}$ |
| | CLES:69.18% | CLES:61.27% | CLES:56.36% |
| Rat | U=786285.0 | U=549410.0 | U=320622.0 |
| | **$p=1.228 \times 10^{-49}$** | **$p=4.697 \times 10^{-14}$** | $p=6.598 \times 10^{-3}$ |
| | CLES:68.54% | CLES:60.08% | CLES:54.20% |
| Macaque | U=714570.0 | U=528868.5 | U=309641.5 |
| | **$p=7.77 \times 10^{-47}$** | **$p=2.916 \times 10^{-16}$** | $p=1.241 \times 10^{-4}$ |
| | CLES:68.45% | CLES:61.09% | CLES:56.05% |
| American mink | U=566380.5 | U=428035.0 | U=246364.5 |
| | **$p=1.434 \times 10^{-40}$** | **$p=1.729 \times 10^{-14}$** | $p=9.248 \times 10^{-4}$ |
| | CLES:68.11% | CLES:60.95% | CLES:55.50% |
| Gelada | U=703318.5 | U=520379.5 | U=301761.5 |
| | **$p=4.19 \times 10^{-49}$** | **$p=3.331 \times 10^{-16}$** | **$p=3.18 \times 10^{-5}$** |
| | CLES:69.04% | CLES:61.13% | CLES:56.64% |
| Sheep | U=562413.0 | U=414931.0 | U=241864.5 |
| | **$p=9.06 \times 10^{-40}$** | **$p=1.219 \times 10^{-11}$** | $p=2.835 \times 10^{-3}$ |
| | CLES:67.95% | CLES:59.71% | CLES:54.98% |
| Tasmanian devil | U=412521.5 | U=324118.0 | U=207423.0 |
| | **$p=7.967 \times 10^{-20}$** | **$p=1.811 \times 10^{-12}$** | **$p=1.371 \times 10^{-5}$** |

| | | | |
|---|---|---|---|
| | CLES:63.14% | CLES:60.78% | CLES:57.56% |
| Elephant | U=585206.5<br>**$p=2.124 \times 10^{-41}$**<br>CLES:68.17% | U=439295.0<br>**$p=2.367 \times 10^{-12}$**<br>CLES:59.90% | U=260223.5<br>$p=6.638 \times 10^{-4}$<br>CLES:55.58% |
| Algerian mouse | U=783624.5<br>**$p=4.116 \times 10^{-30}$**<br>CLES:63.90% | U=591663.0<br>**$p=1.098 \times 10^{-13}$**<br>CLES:59.63% | U=341238.0<br>$p=6.725 \times 10^{-3}$<br>CLES:54.06% |
| Dingo | U=602391.0<br>**$p=3.406 \times 10^{-41}$**<br>CLES:67.97% | U=439444.0<br>**$p=2.059 \times 10^{-13}$**<br>CLES:60.41% | U=256487.0<br>$p=1.192 \times 10^{-3}$<br>CLES:55.33% |
| Golden snub-nosed monkey | U=613275.5<br>**$p=4.171 \times 10^{-46}$**<br>CLES:69.04% | U=457574.0<br>**$p=1.081 \times 10^{-16}$**<br>CLES:61.68% | U=271386.0<br>**$p=1.178 \times 10^{-6}$**<br>CLES:57.96% |
| Horse | U=669442.5<br>**$p=1.924 \times 10^{-44}$**<br>CLES:68.24% | U=491696.0<br>**$p=2.298 \times 10^{-13}$**<br>CLES:60.09% | U=287981.5<br>**$p=2.247 \times 10^{-5}$**<br>CLES:56.85% |
| Black snub-nosed monkey | U=613290.5<br>**$p=1.998 \times 10^{-43}$**<br>CLES:68.42% | U=445261.0<br>**$p=1.36 \times 10^{-13}$**<br>CLES:60.45% | U=276749.0<br>**$p=9.045 \times 10^{-5}$**<br>CLES:56.32% |
| Brazilian guinea pig | U=315813.0<br>**$p=3.563 \times 10^{-26}$**<br>CLES:66.53% | U=243528.0<br>**$p=2.715 \times 10^{-11}$**<br>CLES:60.94% | U=159201.0<br>**$p=3.432 \times 10^{-5}$**<br>CLES:57.67% |
| Orangutan | U=628156.0<br>**$p=9.22 \times 10^{-48}$**<br>CLES:69.31% | U=469426.5<br>**$p=5.376 \times 10^{-16}$**<br>CLES:61.32% | U=270876.0<br>**$p=2.512 \times 10^{-5}$**<br>CLES:56.89% |
| Northern American deer mouse | U=757440.0<br>**$p=1.493 \times 10^{-55}$**<br>CLES:69.93% | U=533804.0<br>**$p=1.446 \times 10^{-16}$**<br>CLES:61.16% | U=322299.5<br>**$p=6.675 \times 10^{-6}$**<br>CLES:57.02% |
| Naked mole-rat female | U=602033.0<br>**$p=2.855 \times 10^{-45}$**<br>CLES:68.97% | U=450995.5<br>**$p=8.787 \times 10^{-16}$**<br>CLES:61.38% | U=273228.0<br>**$p=1.254 \times 10^{-5}$**<br>CLES:57.13% |
| Daurian ground squirrel | U=450717.0<br>**$p=7.724 \times 10^{-45}$**<br>CLES:70.32% | U=309188.5<br>**$p=8.602 \times 10^{-11}$**<br>CLES:59.99% | U=188778.0<br>$p=1.075 \times 10^{-4}$<br>CLES:56.88% |
| Tiger | U=384637.0<br>**$p=2.327 \times 10^{-37}$**<br>CLES:69.17% | U=295147.0<br>**$p=3.32 \times 10^{-10}$**<br>CLES:59.76% | U=167844.0<br>**$p=8.025 \times 10^{-5}$**<br>CLES:57.25% |
| Lesser hedgehog tenrec | U=29021.0<br>**$p=1.077 \times 10^{-8}$**<br>CLES:66.15% | U=18123.0<br>$p=9.904 \times 10^{-3}$<br>CLES:58.00% | U=10789.0<br>$p=6.848 \times 10^{-2}$<br>CLES:56.65% |
| Mongolian gerbil | U=622448.0<br>**$p=1.319 \times 10^{-45}$**<br>CLES:68.79% | U=413667.0<br>**$p=7.743 \times 10^{-14}$**<br>CLES:60.75% | U=251514.5<br>$p=5.823 \times 10^{-4}$<br>CLES:55.66% |
| Ma's night monkey | U=638758.0<br>**$p=4.254 \times 10^{-48}$**<br>CLES:69.32% | U=486609.0<br>**$p=2.508 \times 10^{-16}$**<br>CLES:61.36% | U=281704.0<br>**$p=1.364 \times 10^{-5}$**<br>CLES:57.06% |
| Upper Galilee mountains blind mole rat | U=696648.5<br>**$p=2.617 \times 10^{-47}$** | U=501477.0<br>**$p=1.127 \times 10^{-13}$** | U=318734.5<br>**$p=3.13 \times 10^{-5}$** |

| | | | |
|---|---|---|---|
| | CLES:68.70% | CLES:60.20% | CLES:56.51% |
| Kangaroo rat | U=504027.0 | U=397079.0 | U=240233.5 |
| | **$p=3.424 \times 10^{-39}$** | **$p=3.552 \times 10^{-14}$** | $p=2.205 \times 10^{-4}$ |
| | CLES:68.39% | CLES:61.04% | CLES:56.19% |
| Chimpanzee | U=638618.5 | U=486794.0 | U=293331.0 |
| | **$p=2.76 \times 10^{-50}$** | **$p=1.778 \times 10^{-17}$** | **$p=9.49 \times 10^{-8}$** |
| | CLES:69.81% | CLES:61.82% | CLES:58.62% |
| Guinea Pig | U=601671.0 | U=455164.0 | U=264145.0 |
| | **$p=7.342 \times 10^{-47}$** | **$p=1.583 \times 10^{-17}$** | **$p=7.279 \times 10^{-5}$** |
| | CLES:69.32% | CLES:62.03% | CLES:56.50% |
| American beaver | U=579765.0 | U=387793.0 | U=233185.5 |
| | **$p=1.138 \times 10^{-37}$** | **$p=3.225 \times 10^{-11}$** | $p=6.06 \times 10^{-4}$ |
| | CLES:67.23% | CLES:59.67% | CLES:55.79% |
| Wild yak | U=452344.0 | U=335634.0 | U=190073.0 |
| | **$p=5.602 \times 10^{-37}$** | **$p=3.223 \times 10^{-14}$** | $p=9.85 \times 10^{-4}$ |
| | CLES:68.22% | CLES:61.51% | CLES:55.83% |
| Coquerel's sifaka | U=606617.5 | U=467088.5 | U=264013.5 |
| | **$p=3.723 \times 10^{-44}$** | **$p=9.946 \times 10^{-18}$** | $p=1.713 \times 10^{-4}$ |
| | CLES:68.66% | CLES:62.04% | CLES:56.17% |
| Alpaca | U=16327.5 | U=18806.5 | U=7300.5 |
| | **$p=1.88 \times 10^{-10}$** | $p=2.162 \times 10^{-3}$ | $p=3.335 \times 10^{-3}$ |
| | CLES:71.38% | CLES:59.40% | CLES:62.33% |
| Lesser Egyptian jerboa | U=553805.5 | U=428266.0 | U=263264.0 |
| | **$p=3.989 \times 10^{-46}$** | **$p=7.053 \times 10^{-18}$** | **$p=2.545 \times 10^{-6}$** |
| | CLES:69.62% | CLES:62.39% | CLES:57.75% |
| Long-tailed chinchilla | U=609275.0 | U=462806.5 | U=273620.0 |
| | **$p=4.534 \times 10^{-46}$** | **$p=1.548 \times 10^{-15}$** | $p=1.082 \times 10^{-4}$ |
| | CLES:69.09% | CLES:61.18% | CLES:56.28% |
| Angola colobus | U=637871.0 | U=465619.5 | U=283448.5 |
| | **$p=9.669 \times 10^{-48}$** | **$p=8.845 \times 10^{-16}$** | **$p=2.962 \times 10^{-6}$** |
| | CLES:69.21% | CLES:61.26% | CLES:57.55% |
| Ryukyu mouse | U=818712.5 | U=591961.0 | U=341693.5 |
| | **$p=1.202 \times 10^{-43}$** | **$p=4.175 \times 10^{-15}$** | $p=2.939 \times 10^{-3}$ |
| | CLES:67.05% | CLES:60.29% | CLES:54.52% |
| Chinese hamster PICR | U=820443.5 | U=568202.0 | U=330783.0 |
| | **$p=1.056 \times 10^{-66}$** | **$p=5.049 \times 10^{-17}$** | $p=2.026 \times 10^{-4}$ |
| | CLES:71.62% | CLES:61.17% | CLES:55.74% |
| Vervet-AGM | U=679538.0 | U=502482.0 | U=293706.0 |
| | **$p=5.306 \times 10^{-51}$** | **$p=2.206 \times 10^{-17}$** | **$p=3.383 \times 10^{-5}$** |
| | CLES:69.60% | CLES:61.67% | CLES:56.63% |
| Cow | U=628470.0 | U=457881.5 | U=276976.5 |
| | **$p=1.386 \times 10^{-49}$** | **$p=2.594 \times 10^{-16}$** | **$p=2.176 \times 10^{-5}$** |
| | CLES:69.72% | CLES:61.55% | CLES:56.89% |
| Bushbaby | U=685274.5 | U=494790.5 | U=298221.0 |
| | **$p=9.155 \times 10^{-56}$** | **$p=1.438 \times 10^{-16}$** | **$p=1.085 \times 10^{-6}$** |
| | CLES:70.57% | CLES:61.42% | CLES:57.82% |
| Bonobo | U=636947.0 | U=467117.5 | U=277217.0 |
| | **$p=1.228 \times 10^{-45}$** | **$p=3.577 \times 10^{-14}$** | **$p=9.534 \times 10^{-5}$** |

| | | | |
|---|---|---|---|
| | CLES:68.74% | CLES:60.56% | CLES:56.31% |
| Hyrax | U=55034.5 | U=27069.0 | U=17951.5 |
| | **$p=5.396 \times 10^{-9}$** | $p=2.873 \times 10^{-2}$ | p=0.685 |
| | CLES:63.92% | CLES:56.14% | CLES:51.27% |
| Rabbit | U=473913.0 | U=357529.0 | U=207348.0 |
| | **$p=1.898 \times 10^{-26}$** | **$p=2.992 \times 10^{-7}$** | p=0.159 |
| | CLES:64.94% | CLES:57.54% | CLES:52.40% |
| Megabat | U=183554.0 | U=120891.0 | U=66779.0 |
| | **$p=8.315 \times 10^{-24}$** | **$p=7.739 \times 10^{-11}$** | $p=5.149 \times 10^{-2}$ |
| | CLES:68.04% | CLES:62.84% | CLES:54.46% |
| Dog | U=591532.0 | U=450058.0 | U=255301.0 |
| | **$p=8.644 \times 10^{-37}$** | **$p=1.314 \times 10^{-12}$** | $p=8.096 \times 10^{-3}$ |
| | CLES:66.96% | CLES:59.96% | CLES:54.34% |
| Steppe mouse | U=703960.0 | U=524240.5 | U=305426.0 |
| | **$p=5.437 \times 10^{-14}$** | **$p=4.554 \times 10^{-7}$** | $p=9.264 \times 10^{-2}$ |
| | CLES:59.26% | CLES:56.68% | CLES:52.59% |
| Greater bamboo lemur | U=638724.5 | U=485636.0 | U=281282.5 |
| | **$p=9.989 \times 10^{-44}$** | **$p=2.477 \times 10^{-17}$** | **$p=4.614 \times 10^{-6}$** |
| | CLES:68.29% | CLES:61.75% | CLES:57.44% |
| Pig | U=656468.5 | U=484332.5 | U=287541.0 |
| | **$p=2.965 \times 10^{-48}$** | **$p=1.044 \times 10^{-16}$** | **$p=1.662 \times 10^{-5}$** |
| | CLES:69.18% | CLES:61.53% | CLES:56.93% |
| Tree Shrew | U=22717.5 | U=13352.5 | U=8575.5 |
| | **$p=1.379 \times 10^{-8}$** | $p=3.897 \times 10^{-2}$ | $p=4.433 \times 10^{-3}$ |
| | CLES:67.11% | CLES:56.87% | CLES:61.19% |
| Pika | U=73413.0 | U=44897.0 | U=25669.0 |
| | **$p=6.929 \times 10^{-14}$** | **$p=1.768 \times 10^{-5}$** | p=0.129 |
| | CLES:66.81% | CLES:60.74% | CLES:54.39% |
| Shrew | U=17960.5 | U=10113.0 | U=6532.5 |
| | **$p=2.756 \times 10^{-7}$** | $p=3.321 \times 10^{-2}$ | $p=7.075 \times 10^{-2}$ |
| | CLES:66.39% | CLES:57.67% | CLES:57.52% |
| Squirrel | U=649874.0 | U=490017.0 | U=291064.0 |
| | **$p=1.364 \times 10^{-49}$** | **$p=2.376 \times 10^{-17}$** | **$p=2.326 \times 10^{-6}$** |
| | CLES:69.57% | CLES:61.76% | CLES:57.63% |
| Gorilla | U=642966.5 | U=491089.5 | U=284349.0 |
| | **$p=3.819 \times 10^{-46}$** | **$p=1.723 \times 10^{-15}$** | **$p=2.585 \times 10^{-5}$** |
| | CLES:68.83% | CLES:60.98% | CLES:56.80% |
| Cat | U=653263.0 | U=491180.0 | U=279281.0 |
| | **$p=3.469 \times 10^{-45}$** | **$p=4.796 \times 10^{-15}$** | $p=3.05 \times 10^{-4}$ |
| | CLES:68.52% | CLES:60.80% | CLES:55.83% |
