## Supplementary Table S6 for "Neuron-specific coding and regulatory sequences are the most highly conserved in amniote brains despite neuron-specific cell size diversity"

#### Chicken Reference Genome, Zhang et al., 2014, three cell types

| Sample Size and Median dN/dS (CI95%) | Endothelia-specific Genes | Glia-specific Genes | Neuron-specific Genes |
|---|---|---|---|
| Spoon-billed sandpiper | n=581 med:0.120 (0.111-0.128) | n=707 med:0.115 (0.105-0.124) | n=866 med:0.081 (0.074-0.088) |
| White-throated sparrow | n=519 med:0.120 (0.107-0.135) | n=614 med:0.119 (0.108-0.126) | n=758 med:0.087 (0.080-0.093) |
| Abingdon island giant tortoise | n=537 med:0.115 (0.106-0.123) | n=616 med:0.109 (0.102-0.118) | n=767 med:0.081 (0.074-0.087) |
| Japanese quail | n=608 med:0.152 (0.137-0.167) | n=738 med:0.152 (0.136-0.167) | n=892 med:0.112 (0.101-0.121) |
| Budgerigar | n=565 med:0.140 (0.125-0.148) | n=645 med:0.130 (0.124-0.139) | n=792 med:0.093 (0.087-0.100) |
| Turkey | n=551 med:0.162 (0.143-0.182) | n=643 med:0.175 (0.158-0.190) | n=728 med:0.124 (0.115-0.138) |
| Tuatara | n=523 med:0.099 (0.091-0.109) | n=625 med:0.095 (0.090-0.103) | n=755 med:0.073 (0.068-0.079) |
| Bengalese finch | n=573 med:0.117 (0.106-0.125) | n=680 med:0.110 (0.103-0.119) | n=859 med:0.074 (0.068-0.082) |
| Agassiz's desert tortoise | n=551 med:0.116 (0.106-0.124) | n=643 med:0.112 (0.104-0.120) | n=788 med:0.086 (0.079-0.091) |
| Common canary | n=556 med:0.119 (0.110-0.128) | n=656 med:0.121 (0.108-0.131) | n=792 med:0.083 (0.079-0.090) |
| Flycatcher | n=573 med:0.111 (0.099-0.121) | n=660 med:0.109 (0.097-0.116) | n=850 med:0.076 (0.069-0.083) |
| Painted turtle | n=577 med:0.114 (0.103-0.124) | n=676 med:0.112 (0.103-0.119) | n=844 med:0.079 (0.072-0.084) |
| Great Tit | n=581 med:0.119 (0.109-0.130) | n=682 med:0.120 (0.109-0.129) | n=858 med:0.079 (0.071-0.084) |
| Duck | n=542 med:0.138 | n=662 med:0.138 | n=800 med:0.095 |

| | | | |
|---|---|---|---|
| | (0.124-0.152) | (0.126-0.146) | (0.089-0.105) |
| Blue tit | n=524<br>med:0.130<br>(0.119-0.146) | n=635<br>med:0.123<br>(0.116-0.134) | n=760<br>med:0.093<br>(0.084-0.101) |
| Pink-footed goose | n=602<br>med:0.134<br>(0.121-0.150) | n=734<br>med:0.125<br>(0.114-0.136) | n=864<br>med:0.089<br>(0.083-0.097) |
| Okarito brown kiwi | n=569<br>med:0.131<br>(0.117-0.141) | n=676<br>med:0.128<br>(0.114-0.139) | n=820<br>med:0.094<br>(0.088-0.099) |
| Anole lizard | n=501<br>med:0.083<br>(0.076-0.091) | n=600<br>med:0.077<br>(0.071-0.084) | n=757<br>med:0.062<br>(0.057-0.066) |
| Chilean tinamou | n=532<br>med:0.110<br>(0.099-0.121) | n=645<br>med:0.106<br>(0.096-0.115) | n=783<br>med:0.077<br>(0.071-0.082) |
| Emu | n=583<br>med:0.131<br>(0.120-0.141) | n=683<br>med:0.124<br>(0.113-0.133) | n=840<br>med:0.085<br>(0.076-0.091) |
| Australian saltwater crocodile | n=524<br>med:0.113<br>(0.105-0.123) | n=599<br>med:0.107<br>(0.099-0.115) | n=749<br>med:0.074<br>(0.069-0.080) |
| Ruff | n=581<br>med:0.130<br>(0.114-0.139) | n=685<br>med:0.124<br>(0.113-0.133) | n=842<br>med:0.087<br>(0.078-0.092) |
| Blue-crowned manakin | n=580<br>med:0.123<br>(0.111-0.132) | n=695<br>med:0.118<br>(0.107-0.128) | n=846<br>med:0.079<br>(0.073-0.086) |
| Argentine black and white tegu | n=562<br>med:0.081<br>(0.075-0.087) | n=670<br>med:0.077<br>(0.072-0.082) | n=856<br>med:0.056<br>(0.051-0.060) |
| Dark-eyed junco | n=534<br>med:0.116<br>(0.103-0.127) | n=650<br>med:0.111<br>(0.098-0.119) | n=782<br>med:0.077<br>(0.070-0.083) |
| Golden-collared manakin | n=528<br>med:0.129<br>(0.119-0.142) | n=626<br>med:0.127<br>(0.117-0.135) | n=755<br>med:0.088<br>(0.080-0.095) |
| Great spotted kiwi | n=553<br>med:0.136<br>(0.123-0.151) | n=665<br>med:0.132<br>(0.119-0.143) | n=792<br>med:0.093<br>(0.087-0.099) |
| Zebra Finch | n=469<br>med:0.109<br>(0.099-0.120) | n=536<br>med:0.106<br>(0.097-0.122) | n=685<br>med:0.071<br>(0.065-0.078) |
| Chinese softshell turtle | n=531<br>med:0.108<br>(0.099-0.123) | n=613<br>med:0.103<br>(0.097-0.110) | n=758<br>med:0.075<br>(0.069-0.081) |
| Little spotted kiwi | n=571<br>med:0.130 | n=677<br>med:0.133 | n=797<br>med:0.093 |

| | | | |
|---|---|---|---|
| | (0.119-0.141) | (0.118-0.143) | (0.085-0.097) |
| Helmeted guineafowl | n=607<br>med:0.153<br>(0.140-0.177) | n=738<br>med:0.147<br>(0.134-0.165) | n=893<br>med:0.108<br>(0.095-0.118) |

#### Chicken Reference Genome, Zhang et al., 2014, cell subtypes

| Sample Size and Median dN/dS (CI95%) | Microglia-specific Genes | Endothelia-specific Genes | Astrocyte-specific Genes | Oligodendrocyte-specific Genes | Neuron-specific Genes |
|---|---|---|---|---|---|
| Spoon-billed sandpiper | n=548 med:0.131 (0.119-0.143) | n=581 med:0.120 (0.111-0.128) | n=569 med:0.106 (0.097-0.116) | n=368 med:0.095 (0.081-0.103) | n=866 med:0.081 (0.074-0.088) |
| White-throated sparrow | n=497 med:0.134 (0.120-0.146) | n=519 med:0.120 (0.107-0.135) | n=490 med:0.114 (0.103-0.128) | n=322 med:0.101 (0.091-0.109) | n=758 med:0.087 (0.080-0.093) |
| Abingdon island giant tortoise | n=515 med:0.117 (0.107-0.125) | n=537 med:0.115 (0.106-0.123) | n=500 med:0.105 (0.093-0.115) | n=327 med:0.093 (0.084-0.101) | n=767 med:0.081 (0.074-0.087) |
| Japanese quail | n=587 med:0.170 (0.150-0.188) | n=608 med:0.152 (0.137-0.167) | n=584 med:0.141 (0.128-0.157) | n=380 med:0.125 (0.109-0.143) | n=892 med:0.112 (0.101-0.121) |
| Budgerigar | n=530 med:0.143 (0.135-0.158) | n=565 med:0.140 (0.125-0.148) | n=532 med:0.121 (0.110-0.132) | n=328 med:0.113 (0.101-0.125) | n=792 med:0.093 (0.087-0.100) |
| Turkey | n=534 med:0.178 (0.161-0.196) | n=551 med:0.162 (0.143-0.182) | n=488 med:0.155 (0.136-0.178) | n=326 med:0.138 (0.119-0.173) | n=728 med:0.124 (0.115-0.138) |
| Tuatara | n=509 med:0.101 (0.094-0.109) | n=523 med:0.099 (0.091-0.109) | n=489 med:0.093 (0.087-0.100) | n=321 med:0.086 (0.078-0.093) | n=755 med:0.073 (0.068-0.079) |
| Bengalese finch | n=547 med:0.124 (0.115-0.134) | n=573 med:0.117 (0.106-0.125) | n=545 med:0.102 (0.092-0.111) | n=350 med:0.093 (0.081-0.103) | n=859 med:0.074 (0.068-0.082) |
| Agassiz's desert tortoise | n=527 med:0.122 (0.112-0.130) | n=551 med:0.116 (0.106-0.124) | n=528 med:0.104 (0.093-0.116) | n=323 med:0.099 (0.087-0.107) | n=788 med:0.086 (0.079-0.091) |
| Common canary | n=535 med:0.131 (0.116-0.142) | n=556 med:0.119 (0.110-0.128) | n=514 med:0.115 (0.103-0.123) | n=325 med:0.097 (0.088-0.112) | n=792 med:0.083 (0.079-0.090) |
| Flycatcher | n=544 med:0.123 (0.109-0.133) | n=573 med:0.111 (0.099-0.121) | n=541 med:0.098 (0.091-0.108) | n=333 med:0.090 (0.080-0.098) | n=850 med:0.076 (0.069-0.083) |
| Painted turtle | n=541 med:0.119 (0.110-0.125) | n=577 med:0.114 (0.103-0.124) | n=557 med:0.103 (0.095-0.113) | n=356 med:0.096 (0.086-0.104) | n=844 med:0.079 (0.072-0.084) |

| | | | | | |
|---|---|---|---|---|---|
| Great Tit | n=554 med:0.133 (0.122-0.145) | n=581 med:0.119 (0.109-0.130) | n=547 med:0.107 (0.093-0.119) | n=354 med:0.097 (0.085-0.105) | n=858 med:0.079 (0.071-0.084) |
| Duck | n=534 med:0.142 (0.131-0.155) | n=542 med:0.138 (0.124-0.152) | n=528 med:0.123 (0.110-0.138) | n=345 med:0.114 (0.101-0.130) | n=800 med:0.095 (0.089-0.105) |
| Blue tit | n=516 med:0.136 (0.123-0.145) | n=524 med:0.130 (0.119-0.146) | n=515 med:0.125 (0.114-0.139) | n=320 med:0.105 (0.090-0.117) | n=760 med:0.093 (0.084-0.101) |
| Pink-footed goose | n=583 med:0.143 (0.126-0.153) | n=602 med:0.134 (0.121-0.150) | n=579 med:0.117 (0.106-0.130) | n=369 med:0.105 (0.092-0.115) | n=864 med:0.089 (0.083-0.097) |
| Okarito brown kiwi | n=558 med:0.142 (0.127-0.154) | n=569 med:0.131 (0.117-0.141) | n=527 med:0.116 (0.106-0.129) | n=343 med:0.116 (0.100-0.128) | n=820 med:0.094 (0.088-0.099) |
| Anole lizard | n=478 med:0.078 (0.073-0.087) | n=501 med:0.083 (0.076-0.091) | n=507 med:0.075 (0.067-0.084) | n=313 med:0.068 (0.063-0.075) | n=757 med:0.062 (0.057-0.066) |
| Chilean tinamou | n=521 med:0.120 (0.106-0.130) | n=532 med:0.110 (0.099-0.121) | n=519 med:0.090 (0.083-0.101) | n=340 med:0.089 (0.082-0.098) | n=783 med:0.077 (0.071-0.082) |
| Emu | n=553 med:0.133 (0.123-0.145) | n=583 med:0.131 (0.120-0.141) | n=552 med:0.115 (0.103-0.129) | n=352 med:0.103 (0.089-0.112) | n=840 med:0.085 (0.076-0.091) |
| Australian saltwater crocodile | n=477 med:0.118 (0.107-0.128) | n=524 med:0.113 (0.105-0.123) | n=513 med:0.096 (0.086-0.105) | n=330 med:0.092 (0.081-0.100) | n=749 med:0.074 (0.069-0.080) |
| Ruff | n=556 med:0.133 (0.123-0.145) | n=581 med:0.130 (0.114-0.139) | n=563 med:0.115 (0.105-0.127) | n=358 med:0.103 (0.093-0.115) | n=842 med:0.087 (0.078-0.092) |
| Blue-crowned manakin | n=553 med:0.128 (0.118-0.138) | n=580 med:0.123 (0.111-0.132) | n=550 med:0.108 (0.098-0.120) | n=353 med:0.094 (0.084-0.107) | n=846 med:0.079 (0.073-0.086) |
| Argentine black and white tegu | n=536 med:0.076 (0.070-0.085) | n=562 med:0.081 (0.075-0.087) | n=554 med:0.075 (0.069-0.082) | n=364 med:0.070 (0.063-0.076) | n=856 med:0.056 (0.051-0.060) |
| Dark-eyed junco | n=528 med:0.127 (0.114-0.137) | n=534 med:0.116 (0.103-0.127) | n=493 med:0.097 (0.087-0.105) | n=321 med:0.095 (0.083-0.106) | n=782 med:0.077 (0.070-0.083) |

| | | | | | |
|---|---|---|---|---|---|
| Golden-collared manakin | n=489 med:0.142 (0.131-0.152) | n=528 med:0.129 (0.119-0.142) | n=511 med:0.119 (0.110-0.131) | n=315 med:0.121 (0.106-0.133) | n=755 med:0.088 (0.080-0.095) |
| Great spotted kiwi | n=539 med:0.144 (0.134-0.160) | n=553 med:0.136 (0.123-0.151) | n=528 med:0.120 (0.109-0.131) | n=329 med:0.116 (0.099-0.128) | n=792 med:0.093 (0.087-0.099) |
| Zebra Finch | n=449 med:0.116 (0.106-0.133) | n=469 med:0.109 (0.099-0.120) | n=430 med:0.097 (0.087-0.113) | n=283 med:0.084 (0.077-0.096) | n=685 med:0.071 (0.065-0.078) |
| Chinese softshell turtle | n=481 med:0.109 (0.098-0.117) | n=531 med:0.108 (0.099-0.123) | n=516 med:0.097 (0.092-0.107) | n=313 med:0.091 (0.082-0.101) | n=758 med:0.075 (0.069-0.081) |
| Little spotted kiwi | n=559 med:0.137 (0.130-0.158) | n=571 med:0.130 (0.119-0.141) | n=529 med:0.122 (0.106-0.134) | n=339 med:0.111 (0.099-0.124) | n=797 med:0.093 (0.085-0.097) |
| Helmeted guineafowl | n=589 med:0.176 (0.152-0.188) | n=607 med:0.153 (0.140-0.177) | n=586 med:0.136 (0.120-0.149) | n=376 med:0.123 (0.109-0.140) | n=893 med:0.108 (0.095-0.118) |

#### Chicken Reference Genome, Zeisel et al., 2015, three cell types

| Sample Size and Median dN/dS (CI95%) | Glia-specific Genes | Vasculature-specific Genes | Neuron-specific Genes |
|---|---|---|---|
| Spoon-billed sandpiper | n=856 med:0.138 (0.127-0.146) | n=305 med:0.117 (0.105-0.128) | n=618 med:0.084 (0.078-0.092) |
| White-throated sparrow | n=755 med:0.137 (0.126-0.147) | n=277 med:0.117 (0.103-0.131) | n=538 med:0.088 (0.080-0.098) |
| Abingdon island giant tortoise | n=766 med:0.122 (0.111-0.130) | n=275 med:0.106 (0.093-0.117) | n=559 med:0.083 (0.072-0.091) |
| Japanese quail | n=880 med:0.170 (0.159-0.185) | n=316 med:0.152 (0.134-0.165) | n=632 med:0.122 (0.107-0.134) |
| Budgerigar | n=789 med:0.147 (0.136-0.159) | n=282 med:0.127 (0.111-0.147) | n=554 med:0.099 (0.089-0.108) |
| Turkey | n=806 med:0.189 (0.170-0.199) | n=276 med:0.182 (0.156-0.208) | n=543 med:0.139 (0.125-0.155) |
| Tuatara | n=769 med:0.103 (0.096-0.112) | n=264 med:0.088 (0.080-0.100) | n=544 med:0.077 (0.068-0.083) |
| Bengalese finch | n=804 med:0.120 (0.109-0.131) | n=297 med:0.107 (0.097-0.123) | n=597 med:0.078 (0.069-0.089) |
| Agassiz's desert tortoise | n=806 med:0.123 (0.116-0.135) | n=282 med:0.107 (0.100-0.116) | n=565 med:0.092 (0.084-0.099) |
| Common canary | n=783 med:0.131 (0.121-0.143) | n=282 med:0.110 (0.097-0.121) | n=577 med:0.087 (0.079-0.097) |
| Flycatcher | n=829 med:0.121 (0.112-0.129) | n=298 med:0.106 (0.095-0.119) | n=589 med:0.082 (0.075-0.089) |
| Painted turtle | n=837 med:0.123 (0.115-0.131) | n=295 med:0.109 (0.097-0.119) | n=597 med:0.083 (0.076-0.091) |
| Great Tit | n=836 med:0.133 (0.124-0.144) | n=295 med:0.115 (0.100-0.127) | n=613 med:0.082 (0.075-0.088) |
| Duck | n=787 med:0.152 (0.138-0.167) | n=289 med:0.117 (0.105-0.133) | n=570 med:0.101 (0.089-0.110) |
| Blue tit | n=778 | n=275 | n=550 |

| | | | |
|---|---|---|---|
| | med:0.138 (0.129-0.148) | med:0.122 (0.108-0.138) | med:0.093 (0.084-0.101) |
| Pink-footed goose | n=870 med:0.148 (0.137-0.160) | n=312 med:0.118 (0.109-0.132) | n=623 med:0.096 (0.086-0.104) |
| Okarito brown kiwi | n=827 med:0.149 (0.138-0.160) | n=299 med:0.121 (0.110-0.136) | n=583 med:0.093 (0.085-0.100) |
| Anole lizard | n=742 med:0.082 (0.075-0.089) | n=259 med:0.081 (0.075-0.090) | n=526 med:0.059 (0.054-0.064) |
| Chilean tinamou | n=771 med:0.119 (0.110-0.128) | n=281 med:0.097 (0.085-0.116) | n=557 med:0.074 (0.066-0.081) |
| Emu | n=835 med:0.140 (0.131-0.150) | n=301 med:0.125 (0.113-0.137) | n=594 med:0.084 (0.079-0.090) |
| Australian saltwater crocodile | n=758 med:0.116 (0.108-0.126) | n=272 med:0.102 (0.092-0.113) | n=547 med:0.076 (0.069-0.084) |
| Ruff | n=850 med:0.139 (0.126-0.149) | n=304 med:0.114 (0.102-0.128) | n=608 med:0.085 (0.078-0.093) |
| Blue-crowned manakin | n=821 med:0.132 (0.124-0.144) | n=296 med:0.108 (0.095-0.126) | n=606 med:0.086 (0.079-0.095) |
| Argentine black and white tegu | n=823 med:0.086 (0.078-0.091) | n=291 med:0.078 (0.070-0.086) | n=614 med:0.059 (0.052-0.064) |
| Dark-eyed junco | n=762 med:0.125 (0.112-0.137) | n=289 med:0.105 (0.096-0.121) | n=565 med:0.079 (0.071-0.086) |
| Golden-collared manakin | n=750 med:0.137 (0.129-0.148) | n=276 med:0.130 (0.113-0.143) | n=549 med:0.092 (0.083-0.099) |
| Great spotted kiwi | n=806 med:0.148 (0.139-0.164) | n=283 med:0.125 (0.110-0.138) | n=570 med:0.096 (0.087-0.102) |
| Zebra Finch | n=644 med:0.115 (0.108-0.125) | n=228 med:0.108 (0.084-0.125) | n=497 med:0.075 (0.068-0.084) |
| Chinese softshell turtle | n=756 med:0.114 (0.106-0.120) | n=274 med:0.098 (0.089-0.108) | n=545 med:0.075 (0.070-0.081) |
| Little spotted kiwi | n=833 med:0.148 (0.136-0.160) | n=293 med:0.120 (0.112-0.137) | n=565 med:0.096 (0.087-0.103) |
| Helmeted guineafowl | n=887 | n=312 | n=634 |

| | | |
|---|---|---|
| med:0.170<br>(0.156-0.186) | med:0.148<br>(0.131-0.166) | med:0.117<br>(0.103-0.131) |

### Chicken Reference Genome, Zeisel et al., 2015, cell subtypes

| Sample Size and Median dN/dS (CI95%) | Ependymal-specific Genes | Microglia-specific Genes | Endothelial-specific Genes | Mural-specific Genes | Oligodendrocyte-specific Genes | Astrocyte-specific Genes | Interneuron-specific Genes | S1 Pyramidal-specific Genes | CA1 Pyramidal-specific Genes |
|---|---|---|---|---|---|---|---|---|---|
| Spoon-billed sandpiper | n=228 med:0.195 (0.160-0.216) | n=211 med:0.149 (0.127-0.171) | n=213 med:0.118 (0.105-0.131) | n=92 med:0.115 (0.086-0.139) | n=277 med:0.118 (0.097-0.133) | n=140 med:0.103 (0.082-0.126) | n=212 med:0.089 (0.078-0.098) | n=160 med:0.080 (0.062-0.097) | n=246 med:0.083 (0.069-0.094) |
| White-throated sparrow | n=201 med:0.164 (0.145-0.202) | n=189 med:0.159 (0.133-0.181) | n=189 med:0.117 (0.100-0.134) | n=88 med:0.118 (0.098-0.142) | n=243 med:0.114 (0.103-0.126) | n=122 med:0.113 (0.093-0.146) | n=188 med:0.096 (0.080-0.106) | n=133 med:0.087 (0.068-0.116) | n=217 med:0.086 (0.074-0.100) |
| Abingdon island giant tortoise | n=205 med:0.156 (0.134-0.171) | n=197 med:0.128 (0.107-0.140) | n=195 med:0.103 (0.093-0.114) | n=80 med:0.113 (0.082-0.126) | n=245 med:0.101 (0.091-0.117) | n=119 med:0.106 (0.093-0.126) | n=204 med:0.087 (0.069-0.099) | n=141 med:0.086 (0.067-0.099) | n=214 med:0.074 (0.063-0.086) |
| Japanese quail | n=236 med:0.235 (0.212-0.264) | n=213 med:0.180 (0.167-0.212) | n=220 med:0.154 (0.135-0.172) | n=96 med:0.132 (0.088-0.178) | n=284 med:0.143 (0.123-0.164) | n=147 med:0.122 (0.093-0.154) | n=221 med:0.113 (0.095-0.134) | n=171 med:0.130 (0.097-0.147) | n=240 med:0.121 (0.102-0.143) |
| Budgerigar | n=206 med:0.213 (0.176-0.246) | n=204 med:0.161 (0.141-0.181) | n=197 med:0.126 (0.108-0.148) | n=85 med:0.131 (0.104-0.172) | n=253 med:0.120 (0.103-0.137) | n=126 med:0.115 (0.089-0.135) | n=198 med:0.099 (0.084-0.118) | n=144 med:0.099 (0.083-0.116) | n=212 med:0.098 (0.081-0.109) |
| Turkey | n=217 med:0.225 (0.199-0.276) | n=205 med:0.203 (0.177-0.234) | n=192 med:0.174 (0.141-0.211) | n=84 med:0.191 (0.160-0.234) | n=251 med:0.154 (0.134-0.185) | n=133 med:0.110 (0.092-0.136) | n=199 med:0.134 (0.109-0.165) | n=141 med:0.128 (0.114-0.155) | n=203 med:0.146 (0.120-0.167) |
| Tuatara | n=207 med:0.130 (0.114-0.143) | n=191 med:0.112 (0.092-0.132) | n=184 med:0.086 (0.079-0.099) | n=80 med:0.099 (0.076-0.118) | n=245 med:0.092 (0.082-0.100) | n=126 med:0.085 (0.073-0.105) | n=193 med:0.076 (0.065-0.087) | n=143 med:0.074 (0.058-0.091) | n=208 med:0.078 (0.062-0.091) |
| Bengalese finch | n=208 med:0.162 (0.130-0.187) | n=200 med:0.132 (0.112-0.152) | n=204 med:0.103 (0.089-0.124) | n=93 med:0.113 (0.096-0.137) | n=259 med:0.106 (0.095-0.120) | n=137 med:0.090 (0.071-0.108) | n=198 med:0.080 (0.066-0.102) | n=165 med:0.091 (0.071-0.103) | n=234 med:0.066 (0.058-0.081) |
| Agassiz's desert tortoise | n=211 med:0.160 (0.144-0.176) | n=206 med:0.136 (0.114-0.147) | n=196 med:0.103 (0.096-0.113) | n=86 med:0.120 (0.093-0.130) | n=253 med:0.105 (0.089-0.117) | n=136 med:0.107 (0.095-0.120) | n=198 med:0.099 (0.087-0.107) | n=150 med:0.085 (0.072-0.102) | n=217 med:0.085 (0.073-0.097) |
| Common canary | n=195 med:0.177 (0.153-0.197) | n=203 med:0.138 (0.121-0.161) | n=191 med:0.108 (0.089-0.117) | n=91 med:0.120 (0.097-0.150) | n=255 med:0.110 (0.093-0.125) | n=130 med:0.108 (0.090-0.123) | n=198 med:0.096 (0.080-0.114) | n=149 med:0.092 (0.079-0.113) | n=230 med:0.075 (0.062-0.088) |
| Flycatcher | n=221 med:0.160 (0.145-0.188) | n=211 med:0.124 (0.109-0.146) | n=206 med:0.102 (0.089-0.119) | n=92 med:0.116 (0.090-0.131) | n=258 med:0.104 (0.087-0.116) | n=139 med:0.091 (0.070-0.104) | n=205 med:0.090 (0.073-0.101) | n=156 med:0.085 (0.073-0.099) | n=228 med:0.076 (0.061-0.087) |
| Painted turtle | n=221 med:0.160 (0.145-0.178) | n=211 med:0.128 (0.113-0.139) | n=206 med:0.105 (0.095-0.121) | n=89 med:0.113 (0.086-0.123) | n=266 med:0.099 (0.088-0.117) | n=139 med:0.105 (0.091-0.117) | n=205 med:0.086 (0.071-0.096) | n=157 med:0.082 (0.068-0.101) | n=235 med:0.079 (0.069-0.094) |
| Great Tit | n=218 med:0.182 (0.153-0.202) | n=212 med:0.151 (0.129-0.172) | n=205 med:0.114 (0.094-0.127) | n=90 med:0.122 (0.090-0.148) | n=267 med:0.108 (0.099-0.124) | n=139 med:0.092 (0.077-0.121) | n=212 med:0.087 (0.079-0.102) | n=161 med:0.074 (0.061-0.088) | n=240 med:0.079 (0.069-0.092) |
| Duck | n=201 med:0.204 (0.180-0.232) | n=196 med:0.177 (0.150-0.194) | n=201 med:0.122 (0.107-0.145) | n=88 med:0.110 (0.095-0.131) | n=252 med:0.127 (0.111-0.138) | n=138 med:0.117 (0.100-0.139) | n=199 med:0.105 (0.088-0.116) | n=151 med:0.097 (0.072-0.116) | n=220 med:0.096 (0.076-0.117) |
| Blue tit | n=205 med:0.187 (0.161-0.210) | n=197 med:0.151 (0.137-0.174) | n=187 med:0.125 (0.104-0.145) | n=88 med:0.118 (0.103-0.130) | n=245 med:0.114 (0.102-0.130) | n=131 med:0.104 (0.090-0.133) | n=191 med:0.101 (0.086-0.118) | n=140 med:0.088 (0.073-0.102) | n=219 med:0.091 (0.077-0.106) |
| Pink-footed goose | n=232 med:0.202 (0.178-0.228) | n=222 med:0.155 (0.136-0.184) | n=219 med:0.120 (0.109-0.139) | n=93 med:0.117 (0.096-0.142) | n=274 med:0.126 (0.112-0.147) | n=142 med:0.117 (0.094-0.134) | n=215 med:0.099 (0.080-0.111) | n=171 med:0.100 (0.087-0.121) | n=237 med:0.089 (0.077-0.104) |
| Okarito brown kiwi | n=223 med:0.209 (0.190-0.229) | n=207 med:0.155 (0.137-0.181) | n=210 med:0.123 (0.106-0.139) | n=89 med:0.115 (0.092-0.144) | n=264 med:0.119 (0.100-0.138) | n=133 med:0.112 (0.091-0.141) | n=204 med:0.095 (0.081-0.101) | n=153 med:0.100 (0.083-0.119) | n=226 med:0.089 (0.076-0.103) |
| Anole lizard | n=204 med:0.105 (0.085-0.120) | n=180 med:0.091 (0.083-0.110) | n=179 med:0.086 (0.069-0.094) | n=80 med:0.079 (0.067-0.099) | n=229 med:0.072 (0.063-0.079) | n=129 med:0.067 (0.060-0.084) | n=183 med:0.063 (0.056-0.073) | n=134 med:0.059 (0.050-0.067) | n=209 med:0.057 (0.048-0.067) |
| Chilean tinamou | n=205 med:0.171 (0.140-0.193) | n=185 med:0.138 (0.113-0.158) | n=202 med:0.093 (0.078-0.116) | n=79 med:0.104 (0.085-0.136) | n=253 med:0.098 (0.086-0.114) | n=128 med:0.090 (0.076-0.113) | n=193 med:0.075 (0.063-0.086) | n=144 med:0.076 (0.059-0.091) | n=220 med:0.073 (0.063-0.084) |
| Emu | n=223 med:0.197 (0.176-0.222) | n=211 med:0.141 (0.126-0.172) | n=208 med:0.125 (0.112-0.143) | n=93 med:0.126 (0.101-0.158) | n=260 med:0.118 (0.099-0.131) | n=141 med:0.102 (0.083-0.132) | n=208 med:0.085 (0.073-0.100) | n=158 med:0.087 (0.074-0.102) | n=228 med:0.083 (0.068-0.090) |
| Australian saltwater crocodile | n=200 med:0.148 (0.131-0.173) | n=189 med:0.128 (0.115-0.148) | n=196 med:0.102 (0.092-0.114) | n=76 med:0.103 (0.080-0.126) | n=247 med:0.102 (0.089-0.115) | n=122 med:0.089 (0.078-0.106) | n=182 med:0.086 (0.072-0.092) | n=144 med:0.074 (0.065-0.088) | n=221 med:0.070 (0.060-0.081) |
| Ruff | n=231 med:0.184 (0.161-0.206) | n=207 med:0.157 (0.135-0.184) | n=211 med:0.112 (0.093-0.131) | n=93 med:0.116 (0.100-0.147) | n=268 med:0.113 (0.098-0.128) | n=144 med:0.097 (0.075-0.119) | n=215 med:0.094 (0.082-0.108) | n=160 med:0.081 (0.072-0.098) | n=233 med:0.076 (0.066-0.088) |
| Blue-crowned manakin | n=210 med:0.175 (0.156-0.193) | n=207 med:0.137 (0.124-0.160) | n=211 med:0.115 (0.094-0.130) | n=85 med:0.104 (0.081-0.132) | n=266 med:0.118 (0.098-0.131) | n=138 med:0.095 (0.081-0.118) | n=208 med:0.094 (0.080-0.110) | n=164 med:0.086 (0.068-0.100) | n=234 med:0.082 (0.068-0.094) |
| Argentine black and white tegu | n=224 med:0.109 (0.089-0.123) | n=201 med:0.097 (0.085-0.110) | n=201 med:0.079 (0.068-0.092) | n=90 med:0.075 (0.065-0.088) | n=258 med:0.069 (0.063-0.078) | n=140 med:0.075 (0.065-0.089) | n=214 med:0.059 (0.051-0.072) | n=160 med:0.049 (0.041-0.059) | n=240 med:0.065 (0.054-0.070) |
| Dark-eyed junco | n=191 med:0.178 (0.144-0.198) | n=203 med:0.139 (0.125-0.160) | n=200 med:0.101 (0.085-0.115) | n=89 med:0.121 | n=242 med:0.102 (0.089-0.123) | n=126 med:0.083 (0.066-0.097) | n=197 med:0.081 (0.066-0.096) | n=147 med:0.080 (0.070-0.093) | n=221 med:0.073 (0.061-0.090) |

| | | | | | | | | | |
|---|---|---|---|---|---|---|---|---|---|
| | | | | (0.096-0.140) | | | | | |
| Golden-collared manakin | n=184 med:0.189 (0.161-0.208) | n=184 med:0.153 (0.135-0.168) | n=186 med:0.131 (0.110-0.148) | n=90 med:0.120 (0.103-0.147) | n=249 med:0.121 (0.107-0.135) | n=133 med:0.102 (0.084-0.123) | n=192 med:0.101 (0.081-0.124) | n=142 med:0.088 (0.070-0.096) | n=215 med:0.087 (0.073-0.109) |
| Great spotted kiwi | n=213 med:0.198 (0.173-0.220) | n=204 med:0.169 (0.140-0.187) | n=198 med:0.128 (0.104-0.147) | n=85 med:0.124 (0.106-0.153) | n=256 med:0.120 (0.106-0.142) | n=133 med:0.123 (0.103-0.139) | n=193 med:0.099 (0.083-0.108) | n=153 med:0.096 (0.083-0.119) | n=224 med:0.093 (0.077-0.103) |
| Zebra Finch | n=160 med:0.151 (0.126-0.180) | n=153 med:0.138 (0.115-0.162) | n=153 med:0.097 (0.078-0.122) | n=75 med:0.122 (0.086-0.140) | n=222 med:0.096 (0.085-0.113) | n=109 med:0.084 (0.067-0.113) | n=177 med:0.082 (0.061-0.099) | n=122 med:0.078 (0.065-0.095) | n=198 med:0.074 (0.061-0.081) |
| Chinese softshell turtle | n=203 med:0.144 (0.128-0.157) | n=194 med:0.123 (0.101-0.145) | n=194 med:0.096 (0.086-0.106) | n=80 med:0.108 (0.089-0.126) | n=232 med:0.102 (0.088-0.111) | n=127 med:0.097 (0.091-0.111) | n=188 med:0.077 (0.068-0.089) | n=137 med:0.073 (0.065-0.085) | n=220 med:0.075 (0.062-0.087) |
| Little spotted kiwi | n=220 med:0.210 (0.183-0.225) | n=213 med:0.154 (0.134-0.180) | n=205 med:0.123 (0.109-0.140) | n=88 med:0.118 (0.107-0.154) | n=264 med:0.118 (0.102-0.145) | n=136 med:0.112 (0.095-0.136) | n=202 med:0.100 (0.085-0.111) | n=144 med:0.095 (0.083-0.116) | n=219 med:0.093 (0.077-0.103) |
| Helmeted guineafowl | n=236 med:0.231 (0.197-0.257) | n=221 med:0.183 (0.163-0.210) | n=215 med:0.152 (0.135-0.179) | n=97 med:0.135 (0.099-0.174) | n=282 med:0.139 (0.122-0.152) | n=148 med:0.130 (0.107-0.168) | n=221 med:0.120 (0.105-0.142) | n=170 med:0.129 (0.089-0.159) | n=243 med:0.107 (0.087-0.130) |

#### Chicken Reference Genome, Saunders et al., 2018, three cell types

| Sample Size and Median dN/dS (CI95%) | Vasculature-specific Genes | Glia-specific Genes | Neuron-specific Genes |
|---|---|---|---|
| Spoon-billed sandpiper | n=274 med:0.103 (0.088-0.121) | n=220 med:0.098 (0.084-0.113) | n=504 med:0.066 (0.056-0.073) |
| White-throated sparrow | n=255 med:0.104 (0.095-0.126) | n=184 med:0.100 (0.084-0.111) | n=457 med:0.078 (0.067-0.083) |
| Abingdon island giant tortoise | n=263 med:0.108 (0.091-0.130) | n=193 med:0.093 (0.078-0.108) | n=461 med:0.074 (0.065-0.084) |
| Japanese quail | n=297 med:0.143 (0.119-0.164) | n=229 med:0.142 (0.123-0.158) | n=508 med:0.107 (0.093-0.121) |
| Budgerigar | n=267 med:0.125 (0.115-0.139) | n=196 med:0.108 (0.097-0.124) | n=473 med:0.085 (0.073-0.091) |
| Turkey | n=257 med:0.149 (0.128-0.182) | n=191 med:0.156 (0.121-0.185) | n=438 med:0.122 (0.108-0.140) |
| Tuatara | n=267 med:0.091 (0.073-0.101) | n=191 med:0.086 (0.076-0.095) | n=446 med:0.066 (0.059-0.073) |
| Bengalese finch | n=286 med:0.116 (0.097-0.128) | n=215 med:0.090 (0.079-0.103) | n=502 med:0.062 (0.058-0.073) |
| Agassiz's desert tortoise | n=264 med:0.111 (0.096-0.128) | n=198 med:0.107 (0.083-0.116) | n=468 med:0.074 (0.065-0.084) |
| Common canary | n=265 med:0.108 (0.096-0.121) | n=196 med:0.096 (0.086-0.113) | n=484 med:0.070 (0.059-0.078) |
| Flycatcher | n=279 med:0.109 (0.090-0.119) | n=203 med:0.084 (0.071-0.098) | n=497 med:0.060 (0.054-0.068) |
| Painted turtle | n=269 med:0.109 (0.093-0.128) | n=212 med:0.099 (0.086-0.108) | n=501 med:0.071 (0.062-0.079) |
| Great Tit | n=287 med:0.109 (0.093-0.126) | n=211 med:0.099 (0.087-0.109) | n=496 med:0.066 (0.059-0.075) |
| Duck | n=257 med:0.125 (0.107-0.141) | n=202 med:0.121 (0.100-0.137) | n=461 med:0.088 (0.075-0.100) |
| Blue tit | n=257 | n=187 | n=463 |

| | | | |
|---|---|---|---|
| | med:0.123 (0.113-0.142) | med:0.102 (0.088-0.117) | med:0.083 (0.075-0.093) |
| Pink-footed goose | n=291 med:0.114 (0.093-0.133) | n=224 med:0.109 (0.090-0.126) | n=501 med:0.078 (0.073-0.090) |
| Okarito brown kiwi | n=272 med:0.126 (0.104-0.141) | n=214 med:0.109 (0.094-0.122) | n=485 med:0.072 (0.065-0.077) |
| Anole lizard | n=248 med:0.083 (0.075-0.093) | n=190 med:0.068 (0.057-0.077) | n=446 med:0.056 (0.050-0.062) |
| Chilean tinamou | n=252 med:0.097 (0.079-0.116) | n=201 med:0.080 (0.069-0.093) | n=459 med:0.064 (0.058-0.073) |
| Emu | n=282 med:0.126 (0.108-0.149) | n=205 med:0.104 (0.083-0.125) | n=495 med:0.070 (0.060-0.076) |
| Australian saltwater crocodile | n=249 med:0.111 (0.101-0.121) | n=187 med:0.089 (0.075-0.103) | n=463 med:0.066 (0.057-0.074) |
| Ruff | n=277 med:0.113 (0.098-0.132) | n=209 med:0.107 (0.091-0.129) | n=496 med:0.073 (0.064-0.082) |
| Blue-crowned manakin | n=274 med:0.113 (0.099-0.127) | n=216 med:0.090 (0.082-0.106) | n=499 med:0.070 (0.061-0.076) |
| Argentine black and white tegu | n=276 med:0.072 (0.061-0.081) | n=206 med:0.064 (0.058-0.069) | n=504 med:0.051 (0.048-0.056) |
| Dark-eyed junco | n=270 med:0.104 (0.089-0.123) | n=196 med:0.095 (0.083-0.107) | n=461 med:0.068 (0.062-0.075) |
| Golden-collared manakin | n=258 med:0.129 (0.116-0.143) | n=189 med:0.118 (0.102-0.131) | n=456 med:0.073 (0.065-0.082) |
| Great spotted kiwi | n=272 med:0.119 (0.105-0.143) | n=205 med:0.113 (0.094-0.131) | n=475 med:0.078 (0.069-0.089) |
| Zebra Finch | n=230 med:0.099 (0.082-0.118) | n=168 med:0.081 (0.073-0.089) | n=414 med:0.060 (0.055-0.070) |
| Chinese softshell turtle | n=250 med:0.115 (0.089-0.129) | n=193 med:0.092 (0.082-0.101) | n=470 med:0.061 (0.054-0.070) |
| Little spotted kiwi | n=275 med:0.120 (0.102-0.145) | n=208 med:0.100 (0.092-0.124) | n=480 med:0.075 (0.067-0.088) |
| Helmeted guineafowl | n=287 | n=227 | n=516 |

| | | |
|---|---|---|
| med:0.149 (0.123-0.174) | med:0.123 (0.104-0.143) | med:0.099 (0.084-0.114) |

#### Chicken Reference Genome, Saunders et al., 2018, cell subtypes

| Sample Size and Median dN/dS (CI95%) | Endothelia-specific Genes | Microglia-specific Genes | Astrocyte-specific Genes | Oligodendrocyte-specific Genes | Neuron-specific Genes |
|---|---|---|---|---|---|
| Spoon-billed sandpiper | n=254 med:0.111 (0.097-0.129) | n=171 med:0.118 (0.097-0.143) | n=216 med:0.100 (0.077-0.116) | n=201 med:0.092 (0.078-0.106) | n=462 med:0.066 (0.057-0.076) |
| White-throated sparrow | n=234 med:0.109 (0.099-0.136) | n=153 med:0.128 (0.109-0.158) | n=181 med:0.107 (0.091-0.121) | n=179 med:0.088 (0.077-0.102) | n=416 med:0.078 (0.066-0.084) |
| Abingdon island giant tortoise | n=239 med:0.105 (0.091-0.130) | n=161 med:0.109 (0.087-0.125) | n=183 med:0.096 (0.082-0.110) | n=183 med:0.088 (0.075-0.097) | n=421 med:0.078 (0.066-0.085) |
| Japanese quail | n=270 med:0.141 (0.119-0.164) | n=185 med:0.158 (0.120-0.179) | n=225 med:0.132 (0.099-0.152) | n=208 med:0.130 (0.114-0.160) | n=472 med:0.106 (0.091-0.121) |
| Budgerigar | n=246 med:0.126 (0.110-0.139) | n=164 med:0.144 (0.118-0.163) | n=196 med:0.106 (0.090-0.123) | n=184 med:0.101 (0.089-0.120) | n=434 med:0.085 (0.073-0.090) |
| Turkey | n=234 med:0.152 (0.123-0.188) | n=166 med:0.176 (0.149-0.193) | n=189 med:0.129 (0.101-0.156) | n=179 med:0.158 (0.127-0.206) | n=403 med:0.124 (0.110-0.142) |
| Tuatara | n=240 med:0.091 (0.076-0.102) | n=162 med:0.095 (0.078-0.103) | n=198 med:0.087 (0.073-0.098) | n=174 med:0.082 (0.074-0.093) | n=408 med:0.065 (0.058-0.071) |
| Bengalese finch | n=259 med:0.117 (0.098-0.128) | n=171 med:0.109 (0.097-0.128) | n=212 med:0.093 (0.077-0.105) | n=197 med:0.083 (0.070-0.098) | n=461 med:0.062 (0.058-0.074) |
| Agassiz's desert tortoise | n=244 med:0.110 (0.096-0.125) | n=163 med:0.106 (0.094-0.119) | n=194 med:0.101 (0.083-0.110) | n=186 med:0.087 (0.076-0.104) | n=434 med:0.075 (0.066-0.085) |
| Common canary | n=240 med:0.117 (0.099-0.130) | n=163 med:0.112 (0.093-0.132) | n=189 med:0.102 (0.084-0.116) | n=183 med:0.088 (0.073-0.097) | n=442 med:0.072 (0.060-0.079) |
| Flycatcher | n=256 med:0.108 (0.086-0.118) | n=167 med:0.116 (0.097-0.138) | n=210 med:0.088 (0.070-0.106) | n=185 med:0.075 (0.064-0.093) | n=454 med:0.060 (0.054-0.068) |
| Painted turtle | n=251 med:0.107 (0.093-0.127) | n=173 med:0.104 (0.089-0.119) | n=204 med:0.102 (0.095-0.112) | n=203 med:0.082 (0.071-0.098) | n=457 med:0.072 (0.063-0.080) |
| Great Tit | n=263 med:0.112 (0.094-0.123) | n=170 med:0.116 (0.097-0.133) | n=208 med:0.097 (0.082-0.110) | n=197 med:0.089 (0.073-0.104) | n=456 med:0.066 (0.059-0.075) |
| Duck | n=235 med:0.126 (0.111-0.145) | n=165 med:0.138 (0.112-0.156) | n=195 med:0.113 (0.103-0.133) | n=185 med:0.113 (0.092-0.133) | n=418 med:0.088 (0.075-0.100) |
| Blue tit | n=235 med:0.128 | n=165 med:0.122 | n=178 med:0.112 | n=184 med:0.089 | n=423 med:0.087 |

| | | | | | |
|---|---|---|---|---|---|
| | (0.113-0.152) | (0.104-0.137) | (0.100-0.124) | (0.077-0.103) | (0.077-0.096) |
| Pink-footed goose | n=265<br>med:0.118<br>(0.101-0.136) | n=180<br>med:0.129<br>(0.106-0.153) | n=215<br>med:0.102<br>(0.088-0.117) | n=205<br>med:0.099<br>(0.083-0.120) | n=459<br>med:0.078<br>(0.069-0.091) |
| Okarito brown kiwi | n=248<br>med:0.129<br>(0.109-0.148) | n=177<br>med:0.133<br>(0.103-0.158) | n=202<br>med:0.097<br>(0.086-0.112) | n=197<br>med:0.098<br>(0.084-0.114) | n=440<br>med:0.074<br>(0.067-0.080) |
| Anole lizard | n=229<br>med:0.085<br>(0.075-0.099) | n=144<br>med:0.079<br>(0.069-0.089) | n=185<br>med:0.061<br>(0.054-0.073) | n=174<br>med:0.064<br>(0.056-0.072) | n=408<br>med:0.057<br>(0.050-0.061) |
| Chilean tinamou | n=231<br>med:0.099<br>(0.085-0.119) | n=155<br>med:0.112<br>(0.095-0.138) | n=192<br>med:0.081<br>(0.071-0.094) | n=188<br>med:0.081<br>(0.073-0.093) | n=418<br>med:0.063<br>(0.057-0.074) |
| Emu | n=258<br>med:0.129<br>(0.111-0.153) | n=172<br>med:0.126<br>(0.106-0.141) | n=203<br>med:0.099<br>(0.083-0.111) | n=196<br>med:0.094<br>(0.078-0.116) | n=452<br>med:0.070<br>(0.060-0.076) |
| Australian saltwater crocodile | n=228<br>med:0.119<br>(0.104-0.126) | n=151<br>med:0.104<br>(0.089-0.117) | n=185<br>med:0.088<br>(0.079-0.099) | n=182<br>med:0.084<br>(0.073-0.095) | n=427<br>med:0.067<br>(0.058-0.075) |
| Ruff | n=253<br>med:0.121<br>(0.100-0.133) | n=173<br>med:0.126<br>(0.097-0.154) | n=209<br>med:0.105<br>(0.086-0.115) | n=192<br>med:0.094<br>(0.081-0.110) | n=452<br>med:0.073<br>(0.064-0.085) |
| Blue-crowned manakin | n=258<br>med:0.118<br>(0.107-0.132) | n=169<br>med:0.115<br>(0.102-0.129) | n=209<br>med:0.099<br>(0.089-0.114) | n=200<br>med:0.084<br>(0.073-0.110) | n=454<br>med:0.070<br>(0.061-0.076) |
| Argentine black and white tegu | n=252<br>med:0.079<br>(0.069-0.087) | n=166<br>med:0.073<br>(0.062-0.085) | n=201<br>med:0.067<br>(0.061-0.078) | n=190<br>med:0.062<br>(0.055-0.065) | n=461<br>med:0.050<br>(0.045-0.055) |
| Dark-eyed junco | n=244<br>med:0.106<br>(0.091-0.124) | n=160<br>med:0.124<br>(0.095-0.135) | n=194<br>med:0.096<br>(0.078-0.107) | n=173<br>med:0.088<br>(0.075-0.098) | n=421<br>med:0.067<br>(0.062-0.074) |
| Golden-collared manakin | n=234<br>med:0.131<br>(0.118-0.151) | n=154<br>med:0.129<br>(0.117-0.152) | n=197<br>med:0.108<br>(0.092-0.127) | n=179<br>med:0.100<br>(0.082-0.125) | n=416<br>med:0.073<br>(0.066-0.082) |
| Great spotted kiwi | n=248<br>med:0.129<br>(0.105-0.152) | n=169<br>med:0.132<br>(0.106-0.154) | n=202<br>med:0.107<br>(0.094-0.122) | n=188<br>med:0.099<br>(0.088-0.115) | n=435<br>med:0.081<br>(0.070-0.092) |
| Zebra Finch | n=212<br>med:0.100<br>(0.082-0.122) | n=136<br>med:0.106<br>(0.089-0.129) | n=161<br>med:0.081<br>(0.070-0.094) | n=164<br>med:0.077<br>(0.068-0.085) | n=377<br>med:0.060<br>(0.055-0.070) |
| Chinese softshell turtle | n=233<br>med:0.104<br>(0.084-0.129) | n=166<br>med:0.104<br>(0.082-0.122) | n=183<br>med:0.094<br>(0.088-0.102) | n=177<br>med:0.085<br>(0.070-0.097) | n=431<br>med:0.062<br>(0.055-0.071) |
| Little spotted kiwi | n=252<br>med:0.125<br>(0.105-0.154) | n=170<br>med:0.134<br>(0.103-0.157) | n=201<br>med:0.096<br>(0.086-0.111) | n=194<br>med:0.097<br>(0.085-0.114) | n=440<br>med:0.075<br>(0.067-0.088) |
| Helmeted guineafowl | n=262<br>med:0.149<br>(0.131-0.178) | n=184<br>med:0.147<br>(0.114-0.181) | n=219<br>med:0.129<br>(0.105-0.146) | n=208<br>med:0.112<br>(0.096-0.135) | n=473<br>med:0.099<br>(0.084-0.115) |

#### Chicken Reference Genome, Zeisel et al., 2018, three cell types

| Sample Size and Median dN/dS (CI95%) | CNS_Glia-specific Genes | CNS_Endothelia-specific Genes | CNS_Neuron-specific Genes |
|---|---|---|---|
| Spoon-billed sandpiper | n=582 med:0.146 (0.135-0.159) | n=103 med:0.127 (0.107-0.163) | n=2,582 med:0.089 (0.085-0.095) |
| White-throated sparrow | n=506 med:0.151 (0.139-0.162) | n=88 med:0.146 (0.107-0.167) | n=2,324 med:0.095 (0.091-0.099) |
| Abingdon island giant tortoise | n=527 med:0.126 (0.116-0.139) | n=96 med:0.121 (0.092-0.141) | n=2,390 med:0.088 (0.084-0.091) |
| Japanese quail | n=621 med:0.185 (0.164-0.199) | n=106 med:0.154 (0.117-0.176) | n=2,657 med:0.118 (0.111-0.124) |
| Budgerigar | n=533 med:0.161 (0.148-0.173) | n=98 med:0.131 (0.105-0.158) | n=2,454 med:0.102 (0.097-0.107) |
| Turkey | n=523 med:0.212 (0.193-0.231) | n=96 med:0.185 (0.141-0.210) | n=2,334 med:0.131 (0.124-0.139) |
| Tuatara | n=532 med:0.109 (0.098-0.119) | n=87 med:0.107 (0.084-0.123) | n=2,298 med:0.076 (0.073-0.078) |
| Bengalese finch | n=549 med:0.136 (0.123-0.156) | n=99 med:0.118 (0.095-0.131) | n=2,553 med:0.082 (0.078-0.087) |
| Agassiz's desert tortoise | n=551 med:0.132 (0.121-0.141) | n=101 med:0.123 (0.090-0.137) | n=2,417 med:0.091 (0.087-0.095) |
| Common canary | n=513 med:0.148 (0.134-0.160) | n=94 med:0.117 (0.091-0.141) | n=2,443 med:0.088 (0.085-0.093) |
| Flycatcher | n=547 med:0.141 (0.129-0.155) | n=101 med:0.116 (0.099-0.132) | n=2,537 med:0.081 (0.075-0.084) |
| Painted turtle | n=580 med:0.130 (0.118-0.140) | n=104 med:0.105 (0.088-0.128) | n=2,551 med:0.087 (0.083-0.090) |
| Great Tit | n=566 med:0.145 (0.128-0.156) | n=99 med:0.125 (0.099-0.153) | n=2,533 med:0.087 (0.082-0.090) |
| Duck | n=540 med:0.164 (0.148-0.180) | n=93 med:0.123 (0.107-0.167) | n=2,396 med:0.102 (0.097-0.108) |
| Blue tit | n=523 | n=86 | n=2,383 |

| | | | |
|---|---|---|---|
| | med:0.156 (0.142-0.169) | med:0.141 (0.118-0.171) | med:0.097 (0.093-0.102) |
| Pink-footed goose | n=608 med:0.160 (0.147-0.175) | n=104 med:0.123 (0.099-0.175) | n=2,597 med:0.099 (0.094-0.103) |
| Okarito brown kiwi | n=572 med:0.153 (0.139-0.163) | n=100 med:0.135 (0.112-0.164) | n=2,510 med:0.099 (0.095-0.103) |
| Anole lizard | n=500 med:0.088 (0.077-0.093) | n=87 med:0.082 (0.060-0.101) | n=2,298 med:0.062 (0.060-0.065) |
| Chilean tinamou | n=521 med:0.124 (0.111-0.135) | n=90 med:0.118 (0.075-0.146) | n=2,323 med:0.081 (0.077-0.084) |
| Emu | n=577 med:0.151 (0.143-0.166) | n=105 med:0.141 (0.106-0.160) | n=2,550 med:0.093 (0.088-0.099) |
| Australian saltwater crocodile | n=523 med:0.129 (0.115-0.141) | n=94 med:0.124 (0.103-0.138) | n=2,327 med:0.081 (0.078-0.085) |
| Ruff | n=580 med:0.151 (0.135-0.164) | n=99 med:0.131 (0.112-0.152) | n=2,556 med:0.092 (0.087-0.096) |
| Blue-crowned manakin | n=549 med:0.140 (0.130-0.150) | n=102 med:0.117 (0.083-0.141) | n=2,542 med:0.083 (0.080-0.088) |
| Argentine black and white tegu | n=581 med:0.088 (0.081-0.095) | n=99 med:0.089 (0.075-0.108) | n=2,608 med:0.060 (0.058-0.063) |
| Dark-eyed junco | n=507 med:0.135 (0.121-0.152) | n=96 med:0.121 (0.085-0.146) | n=2,361 med:0.082 (0.078-0.085) |
| Golden-collared manakin | n=517 med:0.155 (0.146-0.164) | n=86 med:0.138 (0.101-0.165) | n=2,329 med:0.094 (0.090-0.099) |
| Great spotted kiwi | n=560 med:0.154 (0.143-0.170) | n=95 med:0.144 (0.108-0.177) | n=2,449 med:0.101 (0.094-0.105) |
| Zebra Finch | n=414 med:0.127 (0.115-0.146) | n=81 med:0.111 (0.089-0.137) | n=2,101 med:0.080 (0.074-0.084) |
| Chinese softshell turtle | n=523 med:0.119 (0.109-0.131) | n=88 med:0.104 (0.079-0.133) | n=2,359 med:0.081 (0.078-0.085) |
| Little spotted kiwi | n=562 med:0.155 (0.141-0.168) | n=100 med:0.132 (0.104-0.161) | n=2,485 med:0.101 (0.096-0.106) |
| Helmeted guineafowl | n=619 | n=106 | n=2,664 |

| | | |
|---|---|---|
| med:0.200 (0.176-0.216) | med:0.148 (0.102-0.180) | med:0.115 (0.108-0.122) |

#### Chicken Reference Genome, Zeisel et al., 2018, cell subtypes

| Sample Size and Median dN/dS (CI95%) | Glia-specific Genes | Immune cells-specific Genes | Vascular cells-specific Genes | Neurons-specific Genes |
|---|---|---|---|---|
| Spoon-billed sandpiper | n=463<br>med:0.167<br>(0.152-0.179) | n=54<br>med:0.172<br>(0.124-0.196) | n=119<br>med:0.117<br>(0.087-0.128) | n=2,672<br>med:0.098<br>(0.093-0.103) |
| White-throated sparrow | n=410<br>med:0.178<br>(0.159-0.197) | n=51<br>med:0.188<br>(0.130-0.221) | n=91<br>med:0.108<br>(0.094-0.154) | n=2,369<br>med:0.102<br>(0.098-0.107) |
| Abingdon island giant tortoise | n=428<br>med:0.145<br>(0.129-0.157) | n=43<br>med:0.140<br>(0.085-0.171) | n=104<br>med:0.106<br>(0.080-0.125) | n=2,465<br>med:0.094<br>(0.090-0.098) |
| Japanese quail | n=500<br>med:0.213<br>(0.193-0.231) | n=56<br>med:0.204<br>(0.167-0.252) | n=120<br>med:0.119<br>(0.094-0.142) | n=2,744<br>med:0.128<br>(0.124-0.133) |
| Budgerigar | n=438<br>med:0.187<br>(0.169-0.208) | n=49<br>med:0.189<br>(0.134-0.225) | n=104<br>med:0.121<br>(0.102-0.145) | n=2,518<br>med:0.111<br>(0.106-0.116) |
| Turkey | n=435<br>med:0.241<br>(0.203-0.261) | n=52<br>med:0.217<br>(0.165-0.260) | n=99<br>med:0.162<br>(0.129-0.190) | n=2,394<br>med:0.141<br>(0.132-0.149) |
| Tuatara | n=414<br>med:0.128<br>(0.115-0.138) | n=46<br>med:0.102<br>(0.080-0.154) | n=96<br>med:0.090<br>(0.070-0.109) | n=2,363<br>med:0.081<br>(0.078-0.085) |
| Bengalese finch | n=427<br>med:0.169<br>(0.147-0.185) | n=52<br>med:0.153<br>(0.109-0.180) | n=115<br>med:0.107<br>(0.087-0.126) | n=2,613<br>med:0.091<br>(0.087-0.095) |
| Agassiz's desert tortoise | n=456<br>med:0.149<br>(0.135-0.162) | n=44<br>med:0.127<br>(0.084-0.176) | n=117<br>med:0.096<br>(0.081-0.121) | n=2,485<br>med:0.096<br>(0.091-0.099) |
| Common canary | n=415<br>med:0.173<br>(0.154-0.191) | n=54<br>med:0.160<br>(0.104-0.208) | n=106<br>med:0.126<br>(0.097-0.137) | n=2,487<br>med:0.099<br>(0.093-0.104) |
| Flycatcher | n=446<br>med:0.166<br>(0.149-0.187) | n=55<br>med:0.149<br>(0.103-0.182) | n=110<br>med:0.107<br>(0.083-0.125) | n=2,598<br>med:0.089<br>(0.085-0.092) |
| Painted turtle | n=474<br>med:0.152<br>(0.139-0.167) | n=46<br>med:0.130<br>(0.086-0.183) | n=117<br>med:0.096<br>(0.086-0.125) | n=2,627<br>med:0.092<br>(0.088-0.096) |
| Great Tit | n=460<br>med:0.166<br>(0.147-0.184) | n=54<br>med:0.175<br>(0.112-0.210) | n=111<br>med:0.112<br>(0.083-0.129) | n=2,601<br>med:0.094<br>(0.090-0.099) |
| Duck | n=433<br>med:0.185<br>(0.171-0.205) | n=53<br>med:0.196<br>(0.150-0.232) | n=109<br>med:0.124<br>(0.106-0.165) | n=2,467<br>med:0.110<br>(0.106-0.116) |
| Blue tit | n=419 | n=53 | n=99 | n=2,422 |

| | | | | |
|---|---|---|---|---|
| | med:0.179 (0.161-0.197) | med:0.165 (0.104-0.185) | med:0.122 (0.113-0.149) | med:0.105 (0.099-0.110) |
| Pink-footed goose | n=497 med:0.185 (0.169-0.202) | n=55 med:0.183 (0.143-0.218) | n=119 med:0.122 (0.099-0.151) | n=2,678 med:0.108 (0.103-0.113) |
| Okarito brown kiwi | n=464 med:0.179 (0.157-0.194) | n=55 med:0.161 (0.127-0.192) | n=105 med:0.112 (0.092-0.134) | n=2,602 med:0.105 (0.101-0.109) |
| Anole lizard | n=415 med:0.107 (0.093-0.114) | n=38 med:0.093 (0.069-0.145) | n=100 med:0.082 (0.065-0.097) | n=2,356 med:0.065 (0.062-0.068) |
| Chilean tinamou | n=427 med:0.142 (0.123-0.168) | n=47 med:0.162 (0.106-0.186) | n=98 med:0.112 (0.071-0.138) | n=2,385 med:0.087 (0.084-0.090) |
| Emu | n=480 med:0.176 (0.153-0.195) | n=53 med:0.140 (0.097-0.186) | n=111 med:0.119 (0.088-0.146) | n=2,629 med:0.101 (0.096-0.105) |
| Australian saltwater crocodile | n=427 med:0.150 (0.135-0.161) | n=45 med:0.124 (0.096-0.159) | n=93 med:0.099 (0.085-0.111) | n=2,377 med:0.088 (0.084-0.092) |
| Ruff | n=470 med:0.176 (0.161-0.200) | n=49 med:0.174 (0.122-0.220) | n=113 med:0.116 (0.092-0.133) | n=2,644 med:0.101 (0.095-0.105) |
| Blue-crowned manakin | n=434 med:0.159 (0.144-0.173) | n=53 med:0.168 (0.124-0.213) | n=111 med:0.106 (0.084-0.137) | n=2,605 med:0.093 (0.088-0.098) |
| Argentine black and white tegu | n=478 med:0.105 (0.093-0.115) | n=45 med:0.091 (0.069-0.132) | n=111 med:0.080 (0.066-0.091) | n=2,689 med:0.064 (0.062-0.067) |
| Dark-eyed junco | n=412 med:0.160 (0.138-0.181) | n=54 med:0.153 (0.104-0.206) | n=103 med:0.123 (0.101-0.133) | n=2,412 med:0.089 (0.085-0.093) |
| Golden-collared manakin | n=412 med:0.182 (0.163-0.205) | n=51 med:0.175 (0.130-0.206) | n=102 med:0.129 (0.096-0.150) | n=2,383 med:0.103 (0.098-0.109) |
| Great spotted kiwi | n=455 med:0.181 (0.160-0.197) | n=52 med:0.169 (0.137-0.208) | n=104 med:0.123 (0.102-0.144) | n=2,535 med:0.108 (0.103-0.114) |
| Zebra Finch | n=322 med:0.153 (0.128-0.174) | n=40 med:0.159 (0.098-0.186) | n=87 med:0.099 (0.083-0.116) | n=2,139 med:0.086 (0.082-0.090) |
| Chinese softshell turtle | n=432 med:0.138 (0.128-0.156) | n=47 med:0.146 (0.091-0.182) | n=102 med:0.098 (0.081-0.124) | n=2,407 med:0.087 (0.084-0.090) |
| Little spotted kiwi | n=463 med:0.186 (0.167-0.202) | n=56 med:0.161 (0.127-0.180) | n=109 med:0.112 (0.084-0.140) | n=2,570 med:0.109 (0.105-0.114) |

| | | | | |
|---|---|---|---|---|
| Helmeted guineafowl | n=501<br>med:0.223<br>(0.203-0.240) | n=58<br>med:0.240<br>(0.192-0.264) | n=120<br>med:0.124<br>(0.100-0.149) | n=2,730<br>med:0.127<br>(0.121-0.134) |

#### Mouse Reference Genome, Zhang et al., 2014, three cell types

| Sample Size and Median dN/dS (CI95%) | Endothelia-specific Genes | Glia-specific Genes | Neuron-specific Genes |
|---|---|---|---|
| Microbat | n=662 med:0.113 (0.105-0.126) | n=824 med:0.120 (0.112-0.130) | n=920 med:0.078 (0.071-0.084) |
| Hedgehog | n=186 med:0.130 (0.110-0.150) | n=224 med:0.131 (0.114-0.148) | n=218 med:0.086 (0.074-0.103) |
| Naked mole-rat male | n=689 med:0.119 (0.110-0.127) | n=918 med:0.120 (0.115-0.127) | n=1,006 med:0.066 (0.061-0.071) |
| Sooty mangabey | n=783 med:0.119 (0.111-0.127) | n=1,028 med:0.119 (0.113-0.126) | n=1,149 med:0.066 (0.060-0.071) |
| Olive baboon | n=791 med:0.119 (0.112-0.126) | n=1,034 med:0.119 (0.114-0.128) | n=1,142 med:0.066 (0.062-0.071) |
| American bison | n=648 med:0.116 (0.104-0.126) | n=808 med:0.118 (0.109-0.129) | n=852 med:0.069 (0.064-0.075) |
| Leopard | n=760 med:0.106 (0.097-0.119) | n=967 med:0.108 (0.101-0.114) | n=1,101 med:0.062 (0.058-0.067) |
| Bolivian squirrel monkey | n=755 med:0.121 (0.113-0.129) | n=964 med:0.123 (0.115-0.129) | n=1,098 med:0.066 (0.061-0.070) |
| Crab-eating macaque | n=784 med:0.119 (0.112-0.124) | n=1,029 med:0.118 (0.113-0.127) | n=1,140 med:0.066 (0.062-0.071) |
| Human | n=815 med:0.114 (0.108-0.123) | n=1,046 med:0.117 (0.112-0.123) | n=1,183 med:0.063 (0.059-0.068) |
| Red fox | n=739 med:0.117 (0.110-0.125) | n=917 med:0.121 (0.110-0.127) | n=1,026 med:0.076 (0.070-0.081) |
| Shrew mouse | n=875 med:0.119 (0.108-0.131) | n=1,143 med:0.127 (0.118-0.135) | n=1,246 med:0.065 (0.058-0.071) |
| Arctic ground squirrel | n=786 med:0.113 (0.104-0.126) | n=1,019 med:0.113 (0.105-0.122) | n=1,122 med:0.058 (0.053-0.064) |
| Chinese hamster CriGri | n=780 med:0.131 (0.120-0.139) | n=1,028 med:0.140 (0.132-0.148) | n=1,100 med:0.075 (0.070-0.083) |
| Opossum | n=694 | n=878 | n=1,003 |

| | | | |
|---|---|---|---|
| | med:0.090<br>(0.083-0.096) | med:0.085<br>(0.079-0.090) | med:0.061<br>(0.058-0.065) |
| Dolphin | n=459<br>med:0.115<br>(0.106-0.124) | n=534<br>med:0.132<br>(0.121-0.139) | n=546<br>med:0.083<br>(0.071-0.090) |
| Capuchin | n=764<br>med:0.121<br>(0.113-0.129) | n=1,009<br>med:0.121<br>(0.114-0.127) | n=1,123<br>med:0.065<br>(0.060-0.070) |
| Mouse Lemur | n=761<br>med:0.116<br>(0.106-0.122) | n=976<br>med:0.113<br>(0.106-0.120) | n=1,089<br>med:0.062<br>(0.057-0.067) |
| Damara mole rat | n=703<br>med:0.114<br>(0.109-0.121) | n=906<br>med:0.115<br>(0.108-0.122) | n=1,017<br>med:0.062<br>(0.057-0.067) |
| Marmoset | n=764<br>med:0.124<br>(0.114-0.135) | n=1,005<br>med:0.122<br>(0.116-0.130) | n=1,127<br>med:0.069<br>(0.064-0.075) |
| Goat | n=752<br>med:0.113<br>(0.105-0.122) | n=954<br>med:0.113<br>(0.107-0.120) | n=1,099<br>med:0.065<br>(0.060-0.070) |
| Common wombat | n=677<br>med:0.090<br>(0.084-0.096) | n=847<br>med:0.088<br>(0.082-0.094) | n=958<br>med:0.059<br>(0.056-0.062) |
| American black bear | n=684<br>med:0.121<br>(0.113-0.129) | n=874<br>med:0.118<br>(0.111-0.127) | n=892<br>med:0.077<br>(0.071-0.083) |
| Sloth | n=104<br>med:0.166<br>(0.141-0.192) | n=142<br>med:0.175<br>(0.140-0.211) | n=124<br>med:0.092<br>(0.078-0.118) |
| Polar bear | n=688<br>med:0.127<br>(0.118-0.137) | n=848<br>med:0.128<br>(0.120-0.138) | n=892<br>med:0.085<br>(0.078-0.092) |
| Alpine marmot | n=754<br>med:0.114<br>(0.108-0.126) | n=966<br>med:0.121<br>(0.111-0.128) | n=1,039<br>med:0.064<br>(0.059-0.072) |
| Prairie vole | n=816<br>med:0.115<br>(0.104-0.125) | n=1,084<br>med:0.123<br>(0.117-0.133) | n=1,195<br>med:0.061<br>(0.057-0.065) |
| Donkey | n=761<br>med:0.115<br>(0.107-0.123) | n=988<br>med:0.121<br>(0.117-0.127) | n=1,116<br>med:0.073<br>(0.067-0.078) |
| Golden Hamster | n=779<br>med:0.127<br>(0.117-0.136) | n=1,028<br>med:0.138<br>(0.129-0.149) | n=1,135<br>med:0.066<br>(0.062-0.070) |
| Armadillo | n=692<br>med:0.119<br>(0.109-0.130) | n=908<br>med:0.116<br>(0.109-0.125) | n=981<br>med:0.071<br>(0.066-0.076) |
| Ugandan red Colobus | n=771 | n=1,007 | n=1,128 |

| | | | |
|---|---|---|---|
| | med:0.119<br>(0.112-0.129) | med:0.121<br>(0.113-0.127) | med:0.066<br>(0.061-0.071) |
| Pig-tailed macaque | n=795<br>med:0.120<br>(0.114-0.128) | n=1,021<br>med:0.120<br>(0.115-0.129) | n=1,151<br>med:0.067<br>(0.063-0.073) |
| Panda | n=771<br>med:0.117<br>(0.105-0.125) | n=968<br>med:0.116<br>(0.108-0.123) | n=1,068<br>med:0.064<br>(0.061-0.069) |
| Ferret | n=749<br>med:0.107<br>(0.099-0.116) | n=953<br>med:0.116<br>(0.108-0.123) | n=1,067<br>med:0.065<br>(0.060-0.068) |
| Koala | n=721<br>med:0.092<br>(0.087-0.097) | n=914<br>med:0.088<br>(0.083-0.094) | n=1,053<br>med:0.062<br>(0.058-0.065) |
| Wallaby | n=208<br>med:0.085<br>(0.074-0.091) | n=293<br>med:0.083<br>(0.077-0.091) | n=295<br>med:0.062<br>(0.054-0.071) |
| Tarsier | n=640<br>med:0.123<br>(0.114-0.130) | n=813<br>med:0.125<br>(0.118-0.132) | n=874<br>med:0.075<br>(0.066-0.082) |
| Degu | n=660<br>med:0.112<br>(0.103-0.119) | n=874<br>med:0.112<br>(0.105-0.119) | n=1,046<br>med:0.063<br>(0.057-0.069) |
| Chinese hamster CHOK1GS | n=862<br>med:0.123<br>(0.112-0.132) | n=1,129<br>med:0.130<br>(0.122-0.136) | n=1,237<br>med:0.065<br>(0.061-0.071) |
| Drill | n=753<br>med:0.120<br>(0.113-0.126) | n=989<br>med:0.122<br>(0.116-0.131) | n=1,079<br>med:0.068<br>(0.064-0.074) |
| Gibbon | n=753<br>med:0.124<br>(0.117-0.134) | n=986<br>med:0.127<br>(0.121-0.134) | n=1,091<br>med:0.073<br>(0.069-0.080) |
| Rat | n=842<br>med:0.126<br>(0.114-0.137) | n=1,108<br>med:0.135<br>(0.126-0.143) | n=1,200<br>med:0.075<br>(0.068-0.083) |
| Macaque | n=812<br>med:0.120<br>(0.112-0.129) | n=1,058<br>med:0.121<br>(0.115-0.130) | n=1,173<br>med:0.066<br>(0.061-0.071) |
| American mink | n=739<br>med:0.108<br>(0.100-0.115) | n=938<br>med:0.111<br>(0.104-0.121) | n=1,042<br>med:0.066<br>(0.061-0.069) |
| Gelada | n=794<br>med:0.116<br>(0.109-0.123) | n=1,034<br>med:0.117<br>(0.111-0.127) | n=1,171<br>med:0.063<br>(0.058-0.067) |
| Sheep | n=730<br>med:0.122<br>(0.113-0.128) | n=926<br>med:0.120<br>(0.112-0.130) | n=1,045<br>med:0.075<br>(0.070-0.081) |
| Tasmanian devil | n=660 | n=847 | n=910 |

| | | | |
|---|---|---|---|
| | med:0.088 (0.080-0.092) | med:0.087 (0.080-0.091) | med:0.057 (0.054-0.061) |
| Elephant | n=759 med:0.115 (0.107-0.125) | n=945 med:0.118 (0.112-0.126) | n=1,069 med:0.070 (0.065-0.075) |
| Algerian mouse | n=863 med:0.132 (0.120-0.148) | n=1,151 med:0.130 (0.119-0.139) | n=1,245 med:0.063 (0.055-0.069) |
| Dingo | n=758 med:0.109 (0.102-0.119) | n=964 med:0.115 (0.109-0.124) | n=1,073 med:0.067 (0.062-0.072) |
| Golden snub-nosed monkey | n=784 med:0.119 (0.113-0.127) | n=988 med:0.123 (0.118-0.129) | n=1,069 med:0.067 (0.063-0.072) |
| Horse | n=773 med:0.114 (0.107-0.124) | n=993 med:0.118 (0.113-0.124) | n=1,146 med:0.071 (0.064-0.076) |
| Black snub-nosed monkey | n=765 med:0.123 (0.114-0.132) | n=985 med:0.125 (0.118-0.133) | n=1,080 med:0.073 (0.068-0.078) |
| Brazilian guinea pig | n=532 med:0.127 (0.118-0.134) | n=727 med:0.125 (0.116-0.133) | n=782 med:0.083 (0.077-0.089) |
| Orangutan | n=775 med:0.118 (0.109-0.127) | n=977 med:0.119 (0.112-0.127) | n=1,092 med:0.066 (0.060-0.072) |
| Northern American deer mouse | n=839 med:0.115 (0.105-0.122) | n=1,106 med:0.123 (0.113-0.132) | n=1,156 med:0.061 (0.056-0.065) |
| Naked mole-rat female | n=742 med:0.117 (0.111-0.128) | n=973 med:0.116 (0.109-0.125) | n=1,087 med:0.064 (0.059-0.069) |
| Daurian ground squirrel | n=678 med:0.127 (0.117-0.139) | n=836 med:0.126 (0.118-0.133) | n=878 med:0.074 (0.067-0.080) |
| Tiger | n=633 med:0.121 (0.112-0.130) | n=799 med:0.125 (0.118-0.133) | n=840 med:0.077 (0.068-0.083) |
| Lesser hedgehog tenrec | n=204 med:0.134 (0.118-0.156) | n=244 med:0.124 (0.108-0.133) | n=214 med:0.083 (0.073-0.096) |
| Mongolian gerbil | n=774 med:0.122 (0.114-0.134) | n=1,007 med:0.129 (0.119-0.140) | n=1,027 med:0.069 (0.063-0.076) |
| Ma's night monkey | n=758 med:0.126 (0.116-0.134) | n=996 med:0.122 (0.115-0.131) | n=1,117 med:0.069 (0.066-0.074) |

| | | | |
|---|---|---|---|
| Upper Galilee mountains blind mole rat | n=778 med:0.122 (0.113-0.132) | n=1,032 med:0.123 (0.115-0.129) | n=1,175 med:0.063 (0.057-0.069) |
| Kangaroo rat | n=680 med:0.106 (0.100-0.113) | n=879 med:0.108 (0.100-0.113) | n=1,018 med:0.061 (0.056-0.068) |
| Chimpanzee | n=761 med:0.118 (0.111-0.129) | n=992 med:0.121 (0.115-0.125) | n=1,117 med:0.065 (0.061-0.070) |
| Guinea Pig | n=766 med:0.110 (0.104-0.121) | n=963 med:0.116 (0.110-0.122) | n=1,065 med:0.068 (0.062-0.072) |
| American beaver | n=744 med:0.121 (0.110-0.131) | n=955 med:0.121 (0.114-0.129) | n=1,017 med:0.072 (0.066-0.078) |
| Wild yak | n=690 med:0.115 (0.108-0.126) | n=847 med:0.118 (0.111-0.128) | n=896 med:0.072 (0.067-0.077) |
| Coquerel's sifaka | n=745 med:0.118 (0.108-0.126) | n=951 med:0.115 (0.109-0.122) | n=1,088 med:0.064 (0.059-0.068) |
| Alpaca | n=165 med:0.166 (0.138-0.193) | n=166 med:0.150 (0.126-0.179) | n=183 med:0.089 (0.077-0.103) |
| Lesser Egyptian jerboa | n=698 med:0.114 (0.108-0.121) | n=916 med:0.117 (0.108-0.122) | n=1,048 med:0.059 (0.053-0.064) |
| Long-tailed chinchilla | n=748 med:0.106 (0.099-0.116) | n=967 med:0.113 (0.105-0.120) | n=1,090 med:0.060 (0.056-0.065) |
| Angola colobus | n=768 med:0.122 (0.113-0.132) | n=995 med:0.124 (0.116-0.130) | n=1,092 med:0.066 (0.060-0.072) |
| Ryukyu mouse | n=882 med:0.129 (0.113-0.138) | n=1,128 med:0.124 (0.113-0.137) | n=1,246 med:0.059 (0.053-0.064) |
| Chinese hamster PICR | n=854 med:0.132 (0.122-0.144) | n=1,110 med:0.136 (0.128-0.142) | n=1,211 med:0.065 (0.060-0.071) |
| Vervet-AGM | n=800 med:0.117 (0.110-0.125) | n=1,029 med:0.119 (0.113-0.126) | n=1,130 med:0.064 (0.059-0.068) |
| Cow | n=767 med:0.110 (0.104-0.119) | n=966 med:0.113 (0.105-0.122) | n=1,094 med:0.065 (0.060-0.069) |
| Bushbaby | n=769 med:0.119 (0.109-0.129) | n=1,014 med:0.124 (0.116-0.132) | n=1,141 med:0.066 (0.060-0.071) |

| | | | |
|---|---|---|---|
| Bonobo | n=762<br>med:0.125<br>(0.114-0.132) | n=1,006<br>med:0.121<br>(0.115-0.127) | n=1,094<br>med:0.070<br>(0.066-0.074) |
| Hyrax | n=225<br>med:0.110<br>(0.097-0.125) | n=294<br>med:0.125<br>(0.116-0.147) | n=287<br>med:0.088<br>(0.078-0.100) |
| Rabbit | n=690<br>med:0.106<br>(0.101-0.116) | n=890<br>med:0.108<br>(0.101-0.114) | n=977<br>med:0.070<br>(0.065-0.080) |
| Megabat | n=419<br>med:0.125<br>(0.108-0.136) | n=553<br>med:0.130<br>(0.118-0.141) | n=545<br>med:0.084<br>(0.074-0.089) |
| Dog | n=751<br>med:0.115<br>(0.108-0.126) | n=978<br>med:0.117<br>(0.107-0.127) | n=1,080<br>med:0.073<br>(0.068-0.077) |
| Steppe mouse | n=850<br>med:0.165<br>(0.145-0.180) | n=1,130<br>med:0.146<br>(0.129-0.159) | n=1,195<br>med:0.088<br>(0.080-0.098) |
| Greater bamboo lemur | n=793<br>med:0.115<br>(0.109-0.127) | n=997<br>med:0.116<br>(0.110-0.122) | n=1,103<br>med:0.066<br>(0.059-0.070) |
| Pig | n=776<br>med:0.111<br>(0.105-0.118) | n=990<br>med:0.114<br>(0.107-0.124) | n=1,115<br>med:0.064<br>(0.061-0.069) |
| Tree Shrew | n=184<br>med:0.129<br>(0.111-0.148) | n=207<br>med:0.135<br>(0.116-0.153) | n=182<br>med:0.086<br>(0.076-0.096) |
| Pika | n=276<br>med:0.108<br>(0.097-0.119) | n=366<br>med:0.121<br>(0.110-0.133) | n=330<br>med:0.075<br>(0.067-0.085) |
| Shrew | n=147<br>med:0.122<br>(0.109-0.137) | n=165<br>med:0.108<br>(0.091-0.118) | n=167<br>med:0.075<br>(0.062-0.091) |
| Squirrel | n=766<br>med:0.112<br>(0.104-0.120) | n=985<br>med:0.108<br>(0.102-0.114) | n=1,135<br>med:0.060<br>(0.055-0.066) |
| Gorilla | n=762<br>med:0.120<br>(0.113-0.129) | n=1,005<br>med:0.121<br>(0.114-0.128) | n=1,120<br>med:0.069<br>(0.063-0.073) |
| Cat | n=785<br>med:0.120<br>(0.111-0.126) | n=1,000<br>med:0.114<br>(0.109-0.123) | n=1,119<br>med:0.067<br>(0.062-0.071) |

#### Mouse Reference Genome, Zhang et al., 2014, cell subtypes

| Sample Size and Median dN/dS (CI95%) | Microglia-specific Genes | Endothelia-specific Genes | Astrocyte-specific Genes | Oligodendrocyte-specific Genes | Neuron-specific Genes |
|---|---|---|---|---|---|
| Microbat | n=704 med:0.128 (0.120-0.135) | n=662 med:0.113 (0.105-0.126) | n=603 med:0.106 (0.099-0.112) | n=372 med:0.092 (0.085-0.102) | n=920 med:0.078 (0.071-0.084) |
| Hedgehog | n=211 med:0.157 (0.137-0.168) | n=186 med:0.130 (0.110-0.150) | n=122 med:0.125 (0.092-0.139) | n=93 med:0.087 (0.072-0.103) | n=218 med:0.086 (0.074-0.103) |
| Naked mole-rat male | n=757 med:0.128 (0.120-0.137) | n=689 med:0.119 (0.110-0.127) | n=635 med:0.105 (0.097-0.111) | n=408 med:0.090 (0.081-0.099) | n=1,006 med:0.066 (0.061-0.071) |
| Sooty mangabey | n=861 med:0.132 (0.124-0.140) | n=783 med:0.119 (0.111-0.127) | n=729 med:0.104 (0.093-0.110) | n=456 med:0.087 (0.081-0.094) | n=1,149 med:0.066 (0.060-0.071) |
| Olive baboon | n=867 med:0.133 (0.126-0.141) | n=791 med:0.119 (0.112-0.126) | n=722 med:0.104 (0.097-0.114) | n=459 med:0.092 (0.083-0.099) | n=1,142 med:0.066 (0.062-0.071) |
| American bison | n=712 med:0.125 (0.115-0.135) | n=648 med:0.116 (0.104-0.126) | n=577 med:0.100 (0.092-0.110) | n=369 med:0.089 (0.080-0.097) | n=852 med:0.069 (0.064-0.075) |
| Leopard | n=809 med:0.121 (0.113-0.130) | n=760 med:0.106 (0.097-0.119) | n=712 med:0.095 (0.086-0.102) | n=437 med:0.078 (0.069-0.084) | n=1,101 med:0.062 (0.058-0.067) |
| Bolivian squirrel monkey | n=822 med:0.135 (0.127-0.145) | n=755 med:0.121 (0.113-0.129) | n=695 med:0.106 (0.095-0.114) | n=436 med:0.088 (0.079-0.093) | n=1,098 med:0.066 (0.061-0.070) |
| Crab-eating macaque | n=868 med:0.131 (0.123-0.139) | n=784 med:0.119 (0.112-0.124) | n=723 med:0.105 (0.094-0.113) | n=454 med:0.087 (0.079-0.096) | n=1,140 med:0.066 (0.062-0.071) |
| Human | n=892 med:0.127 (0.120-0.138) | n=815 med:0.114 (0.108-0.123) | n=747 med:0.101 (0.093-0.111) | n=463 med:0.084 (0.076-0.095) | n=1,183 med:0.063 (0.059-0.068) |
| Red fox | n=811 med:0.131 (0.122-0.140) | n=739 med:0.117 (0.110-0.125) | n=646 med:0.103 (0.097-0.110) | n=425 med:0.087 (0.081-0.098) | n=1,026 med:0.076 (0.070-0.081) |
| Shrew mouse | n=977 med:0.145 (0.133-0.160) | n=875 med:0.119 (0.108-0.131) | n=784 med:0.105 (0.096-0.115) | n=510 med:0.076 (0.066-0.088) | n=1,246 med:0.065 (0.058-0.071) |
| Arctic ground squirrel | n=855 med:0.130 (0.121-0.136) | n=786 med:0.113 (0.104-0.126) | n=720 med:0.095 (0.087-0.103) | n=459 med:0.077 (0.071-0.087) | n=1,122 med:0.058 (0.053-0.064) |
| Chinese hamster CriGri | n=902 med:0.163 (0.153-0.174) | n=780 med:0.131 (0.120-0.139) | n=711 med:0.114 (0.107-0.126) | n=456 med:0.092 (0.081-0.102) | n=1,100 med:0.075 (0.070-0.083) |
| Opossum | n=736 med:0.089 (0.082-0.095) | n=694 med:0.090 (0.083-0.096) | n=645 med:0.080 (0.073-0.086) | n=416 med:0.073 (0.066-0.081) | n=1,003 med:0.061 (0.058-0.065) |
| Dolphin | n=483 med:0.138 (0.131-0.152) | n=459 med:0.115 (0.106-0.124) | n=365 med:0.121 (0.108-0.132) | n=231 med:0.102 (0.092-0.114) | n=546 med:0.083 (0.071-0.090) |
| Capuchin | n=841 med:0.133 (0.124-0.141) | n=764 med:0.121 (0.113-0.129) | n=711 med:0.107 (0.095-0.114) | n=455 med:0.085 (0.077-0.093) | n=1,123 med:0.065 (0.060-0.070) |
| Mouse Lemur | n=819 med:0.126 | n=761 med:0.116 | n=680 med:0.100 | n=444 med:0.077 | n=1,089 med:0.062 |

| | | | | | |
|---|---|---|---|---|---|
| | (0.117-0.133) | (0.106-0.122) | (0.091-0.106) | (0.071-0.089) | (0.057-0.067) |
| Damara mole rat | n=747<br>med:0.122<br>(0.112-0.129) | n=703<br>med:0.114<br>(0.109-0.121) | n=653<br>med:0.097<br>(0.086-0.107) | n=420<br>med:0.084<br>(0.077-0.095) | n=1,017<br>med:0.062<br>(0.057-0.067) |
| Marmoset | n=844<br>med:0.137<br>(0.129-0.147) | n=764<br>med:0.124<br>(0.114-0.135) | n=719<br>med:0.112<br>(0.101-0.118) | n=457<br>med:0.089<br>(0.080-0.098) | n=1,127<br>med:0.069<br>(0.064-0.075) |
| Goat | n=823<br>med:0.120<br>(0.114-0.130) | n=752<br>med:0.113<br>(0.105-0.122) | n=695<br>med:0.097<br>(0.090-0.105) | n=446<br>med:0.084<br>(0.076-0.091) | n=1,099<br>med:0.065<br>(0.060-0.070) |
| Common wombat | n=725<br>med:0.092<br>(0.084-0.099) | n=677<br>med:0.090<br>(0.084-0.096) | n=623<br>med:0.083<br>(0.075-0.088) | n=405<br>med:0.077<br>(0.072-0.083) | n=958<br>med:0.059<br>(0.056-0.062) |
| American black bear | n=765<br>med:0.123<br>(0.115-0.132) | n=684<br>med:0.121<br>(0.113-0.129) | n=584<br>med:0.112<br>(0.100-0.120) | n=378<br>med:0.088<br>(0.079-0.096) | n=892<br>med:0.077<br>(0.071-0.083) |
| Sloth | n=120<br>med:0.190<br>(0.144-0.231) | n=104<br>med:0.166<br>(0.141-0.192) | n=90<br>med:0.139<br>(0.109-0.160) | n=65<br>med:0.133<br>(0.073-0.172) | n=124<br>med:0.092<br>(0.078-0.118) |
| Polar bear | n=752<br>med:0.135<br>(0.127-0.146) | n=688<br>med:0.127<br>(0.118-0.137) | n=607<br>med:0.114<br>(0.103-0.124) | n=382<br>med:0.090<br>(0.085-0.100) | n=892<br>med:0.085<br>(0.078-0.092) |
| Alpine marmot | n=841<br>med:0.133<br>(0.125-0.143) | n=754<br>med:0.114<br>(0.108-0.126) | n=680<br>med:0.095<br>(0.088-0.105) | n=424<br>med:0.088<br>(0.076-0.096) | n=1,039<br>med:0.064<br>(0.059-0.072) |
| Prairie vole | n=921<br>med:0.141<br>(0.127-0.151) | n=816<br>med:0.115<br>(0.104-0.125) | n=738<br>med:0.102<br>(0.092-0.114) | n=487<br>med:0.079<br>(0.074-0.089) | n=1,195<br>med:0.061<br>(0.057-0.065) |
| Donkey | n=851<br>med:0.128<br>(0.124-0.137) | n=761<br>med:0.115<br>(0.107-0.123) | n=703<br>med:0.106<br>(0.098-0.113) | n=447<br>med:0.090<br>(0.081-0.100) | n=1,116<br>med:0.073<br>(0.067-0.078) |
| Golden Hamster | n=872<br>med:0.157<br>(0.144-0.167) | n=779<br>med:0.127<br>(0.117-0.136) | n=712<br>med:0.115<br>(0.104-0.122) | n=462<br>med:0.096<br>(0.085-0.107) | n=1,135<br>med:0.066<br>(0.062-0.070) |
| Armadillo | n=763<br>med:0.126<br>(0.118-0.134) | n=692<br>med:0.119<br>(0.109-0.130) | n=651<br>med:0.108<br>(0.096-0.113) | n=405<br>med:0.087<br>(0.079-0.094) | n=981<br>med:0.071<br>(0.066-0.076) |
| Ugandan red Colobus | n=846<br>med:0.135<br>(0.127-0.144) | n=771<br>med:0.119<br>(0.112-0.129) | n=708<br>med:0.104<br>(0.094-0.110) | n=455<br>med:0.085<br>(0.078-0.095) | n=1,128<br>med:0.066<br>(0.061-0.071) |
| Pig-tailed macaque | n=863<br>med:0.133<br>(0.124-0.142) | n=795<br>med:0.120<br>(0.114-0.128) | n=720<br>med:0.102<br>(0.094-0.112) | n=462<br>med:0.086<br>(0.078-0.096) | n=1,151<br>med:0.067<br>(0.063-0.073) |
| Panda | n=824<br>med:0.123<br>(0.114-0.131) | n=771<br>med:0.117<br>(0.105-0.125) | n=671<br>med:0.098<br>(0.090-0.104) | n=437<br>med:0.082<br>(0.073-0.090) | n=1,068<br>med:0.064<br>(0.061-0.069) |
| Ferret | n=810<br>med:0.122<br>(0.116-0.133) | n=749<br>med:0.107<br>(0.099-0.116) | n=670<br>med:0.097<br>(0.088-0.104) | n=429<br>med:0.079<br>(0.071-0.088) | n=1,067<br>med:0.065<br>(0.060-0.068) |
| Koala | n=775<br>med:0.093<br>(0.086-0.097) | n=721<br>med:0.092<br>(0.087-0.097) | n=654<br>med:0.084<br>(0.078-0.090) | n=440<br>med:0.077<br>(0.072-0.082) | n=1,053<br>med:0.062<br>(0.058-0.065) |
| Wallaby | n=265<br>med:0.085<br>(0.076-0.092) | n=208<br>med:0.085<br>(0.074-0.091) | n=176<br>med:0.085<br>(0.072-0.101) | n=146<br>med:0.077<br>(0.067-0.091) | n=295<br>med:0.062<br>(0.054-0.071) |
| Tarsier | n=697<br>med:0.137<br>(0.129-0.146) | n=640<br>med:0.123<br>(0.114-0.130) | n=579<br>med:0.106<br>(0.095-0.114) | n=361<br>med:0.088<br>(0.080-0.097) | n=874<br>med:0.075<br>(0.066-0.082) |

| | | | | | |
|---|---|---|---|---|---|
| Degu | n=712 med:0.121 (0.112-0.125) | n=660 med:0.112 (0.103-0.119) | n=630 med:0.096 (0.089-0.105) | n=420 med:0.083 (0.073-0.089) | n=1,046 med:0.063 (0.057-0.069) |
| Chinese hamster CHOK1GS | n=957 med:0.150 (0.141-0.161) | n=862 med:0.123 (0.112-0.132) | n=783 med:0.107 (0.098-0.116) | n=506 med:0.083 (0.075-0.094) | n=1,237 med:0.065 (0.061-0.071) |
| Drill | n=852 med:0.135 (0.124-0.144) | n=753 med:0.120 (0.113-0.126) | n=695 med:0.106 (0.094-0.114) | n=438 med:0.086 (0.078-0.096) | n=1,079 med:0.068 (0.064-0.074) |
| Gibbon | n=822 med:0.143 (0.134-0.154) | n=753 med:0.124 (0.117-0.134) | n=696 med:0.111 (0.099-0.119) | n=435 med:0.093 (0.085-0.105) | n=1,091 med:0.073 (0.069-0.080) |
| Rat | n=956 med:0.149 (0.138-0.162) | n=842 med:0.126 (0.114-0.137) | n=762 med:0.116 (0.104-0.124) | n=493 med:0.085 (0.076-0.096) | n=1,200 med:0.075 (0.068-0.083) |
| Macaque | n=890 med:0.134 (0.123-0.143) | n=812 med:0.120 (0.112-0.129) | n=738 med:0.107 (0.096-0.114) | n=471 med:0.086 (0.079-0.094) | n=1,173 med:0.066 (0.061-0.071) |
| American mink | n=798 med:0.123 (0.113-0.129) | n=739 med:0.108 (0.100-0.115) | n=674 med:0.092 (0.086-0.101) | n=426 med:0.079 (0.073-0.086) | n=1,042 med:0.066 (0.061-0.069) |
| Gelada | n=870 med:0.134 (0.125-0.141) | n=794 med:0.116 (0.109-0.123) | n=727 med:0.101 (0.092-0.110) | n=455 med:0.084 (0.075-0.093) | n=1,171 med:0.063 (0.058-0.067) |
| Sheep | n=792 med:0.130 (0.120-0.138) | n=730 med:0.122 (0.113-0.128) | n=665 med:0.102 (0.093-0.110) | n=421 med:0.088 (0.080-0.098) | n=1,045 med:0.075 (0.070-0.081) |
| Tasmanian devil | n=718 med:0.092 (0.086-0.096) | n=660 med:0.088 (0.080-0.092) | n=586 med:0.081 (0.075-0.088) | n=396 med:0.072 (0.066-0.078) | n=910 med:0.057 (0.054-0.061) |
| Elephant | n=803 med:0.131 (0.124-0.139) | n=759 med:0.115 (0.107-0.125) | n=686 med:0.102 (0.093-0.108) | n=438 med:0.089 (0.081-0.095) | n=1,069 med:0.070 (0.065-0.075) |
| Algerian mouse | n=985 med:0.141 (0.130-0.158) | n=863 med:0.132 (0.120-0.148) | n=797 med:0.105 (0.094-0.119) | n=507 med:0.083 (0.070-0.097) | n=1,245 med:0.063 (0.055-0.069) |
| Dingo | n=826 med:0.124 (0.117-0.133) | n=758 med:0.109 (0.102-0.119) | n=678 med:0.098 (0.090-0.108) | n=432 med:0.083 (0.076-0.090) | n=1,073 med:0.067 (0.062-0.072) |
| Golden snub-nosed monkey | n=831 med:0.132 (0.126-0.142) | n=784 med:0.119 (0.113-0.127) | n=694 med:0.108 (0.097-0.118) | n=438 med:0.089 (0.080-0.100) | n=1,069 med:0.067 (0.063-0.072) |
| Horse | n=856 med:0.129 (0.121-0.137) | n=773 med:0.114 (0.107-0.124) | n=714 med:0.101 (0.092-0.110) | n=442 med:0.091 (0.080-0.102) | n=1,146 med:0.071 (0.064-0.076) |
| Black snub-nosed monkey | n=830 med:0.136 (0.128-0.145) | n=765 med:0.123 (0.114-0.132) | n=682 med:0.112 (0.103-0.120) | n=455 med:0.089 (0.083-0.099) | n=1,080 med:0.073 (0.068-0.078) |
| Brazilian guinea pig | n=607 med:0.126 (0.118-0.138) | n=532 med:0.127 (0.118-0.134) | n=511 med:0.117 (0.107-0.127) | n=353 med:0.100 (0.091-0.112) | n=782 med:0.083 (0.077-0.089) |
| Orangutan | n=830 med:0.131 (0.124-0.139) | n=775 med:0.118 (0.109-0.127) | n=701 med:0.103 (0.095-0.112) | n=436 med:0.082 (0.077-0.091) | n=1,092 med:0.066 (0.060-0.072) |
| Northern American deer mouse | n=937 med:0.145 (0.132-0.154) | n=839 med:0.115 (0.105-0.122) | n=755 med:0.103 (0.094-0.113) | n=489 med:0.084 (0.072-0.093) | n=1,156 med:0.061 (0.056-0.065) |

| | | | | | |
|---|---|---|---|---|---|
| Naked mole-rat female | n=803 med:0.127 (0.118-0.135) | n=742 med:0.117 (0.111-0.128) | n=676 med:0.101 (0.094-0.108) | n=440 med:0.085 (0.078-0.096) | n=1,087 med:0.064 (0.059-0.069) |
| Daurian ground squirrel | n=730 med:0.138 (0.130-0.149) | n=678 med:0.127 (0.117-0.139) | n=587 med:0.103 (0.092-0.114) | n=378 med:0.092 (0.083-0.099) | n=878 med:0.074 (0.067-0.080) |
| Tiger | n=662 med:0.140 (0.128-0.152) | n=633 med:0.121 (0.112-0.130) | n=588 med:0.106 (0.097-0.116) | n=349 med:0.095 (0.086-0.107) | n=840 med:0.077 (0.068-0.083) |
| Lesser hedgehog tenrec | n=205 med:0.128 (0.116-0.150) | n=204 med:0.134 (0.118-0.156) | n=146 med:0.116 (0.095-0.125) | n=89 med:0.104 (0.080-0.122) | n=214 med:0.083 (0.073-0.096) |
| Mongolian gerbil | n=881 med:0.150 (0.137-0.161) | n=774 med:0.122 (0.114-0.134) | n=663 med:0.111 (0.102-0.123) | n=440 med:0.085 (0.076-0.097) | n=1,027 med:0.069 (0.063-0.076) |
| Ma's night monkey | n=825 med:0.136 (0.128-0.144) | n=758 med:0.126 (0.116-0.134) | n=710 med:0.107 (0.098-0.115) | n=442 med:0.090 (0.083-0.096) | n=1,117 med:0.069 (0.066-0.074) |
| Upper Galilee mountains blind mole rat | n=863 med:0.138 (0.129-0.147) | n=778 med:0.122 (0.113-0.132) | n=709 med:0.104 (0.093-0.112) | n=480 med:0.085 (0.076-0.096) | n=1,175 med:0.063 (0.057-0.069) |
| Kangaroo rat | n=724 med:0.115 (0.110-0.125) | n=680 med:0.106 (0.100-0.113) | n=639 med:0.094 (0.084-0.102) | n=420 med:0.080 (0.072-0.088) | n=1,018 med:0.061 (0.056-0.068) |
| Chimpanzee | n=819 med:0.133 (0.123-0.142) | n=761 med:0.118 (0.111-0.129) | n=705 med:0.106 (0.097-0.115) | n=448 med:0.091 (0.082-0.100) | n=1,117 med:0.065 (0.061-0.070) |
| Guinea Pig | n=815 med:0.125 (0.117-0.133) | n=766 med:0.110 (0.104-0.121) | n=689 med:0.101 (0.094-0.109) | n=439 med:0.083 (0.075-0.090) | n=1,065 med:0.068 (0.062-0.072) |
| American beaver | n=848 med:0.131 (0.124-0.144) | n=744 med:0.121 (0.110-0.131) | n=639 med:0.106 (0.096-0.116) | n=411 med:0.096 (0.084-0.104) | n=1,017 med:0.072 (0.066-0.078) |
| Wild yak | n=740 med:0.128 (0.119-0.138) | n=690 med:0.115 (0.108-0.126) | n=609 med:0.102 (0.094-0.110) | n=380 med:0.086 (0.080-0.095) | n=896 med:0.072 (0.067-0.077) |
| Coquerel's sifaka | n=812 med:0.130 (0.121-0.135) | n=745 med:0.118 (0.108-0.126) | n=692 med:0.105 (0.093-0.111) | n=432 med:0.083 (0.073-0.093) | n=1,088 med:0.064 (0.059-0.068) |
| Alpaca | n=125 med:0.184 (0.152-0.231) | n=165 med:0.166 (0.138-0.193) | n=173 med:0.119 (0.107-0.140) | n=64 med:0.145 (0.114-0.165) | n=183 med:0.089 (0.077-0.103) |
| Lesser Egyptian jerboa | n=759 med:0.124 (0.119-0.133) | n=698 med:0.114 (0.108-0.121) | n=655 med:0.098 (0.091-0.106) | n=435 med:0.086 (0.077-0.097) | n=1,048 med:0.059 (0.053-0.064) |
| Long-tailed chinchilla | n=809 med:0.122 (0.113-0.132) | n=748 med:0.106 (0.099-0.116) | n=694 med:0.093 (0.086-0.102) | n=446 med:0.078 (0.070-0.086) | n=1,090 med:0.060 (0.056-0.065) |
| Angola colobus | n=844 med:0.134 (0.126-0.144) | n=768 med:0.122 (0.113-0.132) | n=696 med:0.106 (0.096-0.115) | n=451 med:0.087 (0.082-0.097) | n=1,092 med:0.066 (0.060-0.072) |
| Ryukyu mouse | n=980 med:0.146 (0.131-0.158) | n=882 med:0.129 (0.113-0.138) | n=788 med:0.106 (0.094-0.117) | n=503 med:0.082 (0.070-0.093) | n=1,246 med:0.059 (0.053-0.064) |
| Chinese hamster PICR | n=946 med:0.159 (0.151-0.170) | n=854 med:0.132 (0.122-0.144) | n=767 med:0.110 (0.101-0.118) | n=490 med:0.088 (0.077-0.096) | n=1,211 med:0.065 (0.060-0.071) |
| Vervet-AGM | n=864 | n=800 | n=721 | n=459 | n=1,130 |

| | | | | | |
|---|---|---|---|---|---|
| | med:0.135<br>(0.123-0.141) | med:0.117<br>(0.110-0.125) | med:0.102<br>(0.093-0.110) | med:0.084<br>(0.075-0.092) | med:0.064<br>(0.059-0.068) |
| Cow | n=824<br>med:0.127<br>(0.117-0.134) | n=767<br>med:0.110<br>(0.104-0.119) | n=680<br>med:0.095<br>(0.088-0.104) | n=445<br>med:0.085<br>(0.076-0.092) | n=1,094<br>med:0.065<br>(0.060-0.069) |
| Bushbaby | n=851<br>med:0.135<br>(0.127-0.145) | n=769<br>med:0.119<br>(0.109-0.129) | n=706<br>med:0.102<br>(0.091-0.108) | n=452<br>med:0.085<br>(0.079-0.096) | n=1,141<br>med:0.066<br>(0.060-0.071) |
| Bonobo | n=847<br>med:0.138<br>(0.128-0.144) | n=762<br>med:0.125<br>(0.114-0.132) | n=705<br>med:0.107<br>(0.097-0.116) | n=450<br>med:0.088<br>(0.080-0.098) | n=1,094<br>med:0.070<br>(0.066-0.074) |
| Hyrax | n=300<br>med:0.146<br>(0.126-0.162) | n=225<br>med:0.110<br>(0.097-0.125) | n=168<br>med:0.112<br>(0.093-0.133) | n=122<br>med:0.102<br>(0.087-0.118) | n=287<br>med:0.088<br>(0.078-0.100) |
| Rabbit | n=747<br>med:0.119<br>(0.111-0.127) | n=690<br>med:0.106<br>(0.101-0.116) | n=636<br>med:0.096<br>(0.088-0.104) | n=405<br>med:0.081<br>(0.069-0.089) | n=977<br>med:0.070<br>(0.065-0.080) |
| Megabat | n=495<br>med:0.148<br>(0.137-0.158) | n=419<br>med:0.125<br>(0.108-0.136) | n=353<br>med:0.126<br>(0.116-0.146) | n=225<br>med:0.097<br>(0.083-0.108) | n=545<br>med:0.084<br>(0.074-0.089) |
| Dog | n=818<br>med:0.127<br>(0.117-0.137) | n=751<br>med:0.115<br>(0.108-0.126) | n=695<br>med:0.102<br>(0.095-0.109) | n=435<br>med:0.084<br>(0.078-0.092) | n=1,080<br>med:0.073<br>(0.068-0.077) |
| Steppe mouse | n=994<br>med:0.162<br>(0.147-0.181) | n=850<br>med:0.165<br>(0.145-0.180) | n=774<br>med:0.136<br>(0.120-0.151) | n=486<br>med:0.111<br>(0.093-0.131) | n=1,195<br>med:0.088<br>(0.080-0.098) |
| Greater bamboo lemur | n=848<br>med:0.128<br>(0.119-0.135) | n=793<br>med:0.115<br>(0.109-0.127) | n=713<br>med:0.102<br>(0.092-0.110) | n=444<br>med:0.088<br>(0.076-0.098) | n=1,103<br>med:0.066<br>(0.059-0.070) |
| Pig | n=851<br>med:0.127<br>(0.120-0.134) | n=776<br>med:0.111<br>(0.105-0.118) | n=706<br>med:0.097<br>(0.092-0.107) | n=453<br>med:0.084<br>(0.075-0.093) | n=1,115<br>med:0.064<br>(0.061-0.069) |
| Tree Shrew | n=186<br>med:0.152<br>(0.128-0.179) | n=184<br>med:0.129<br>(0.111-0.148) | n=129<br>med:0.113<br>(0.086-0.135) | n=77<br>med:0.112<br>(0.094-0.142) | n=182<br>med:0.086<br>(0.076-0.096) |
| Pika | n=333<br>med:0.136<br>(0.117-0.147) | n=276<br>med:0.108<br>(0.097-0.119) | n=224<br>med:0.105<br>(0.094-0.122) | n=143<br>med:0.086<br>(0.074-0.105) | n=330<br>med:0.075<br>(0.067-0.085) |
| Shrew | n=162<br>med:0.135<br>(0.108-0.172) | n=147<br>med:0.122<br>(0.109-0.137) | n=105<br>med:0.098<br>(0.073-0.116) | n=68<br>med:0.097<br>(0.073-0.109) | n=167<br>med:0.075<br>(0.062-0.091) |
| Squirrel | n=823<br>med:0.119<br>(0.111-0.130) | n=766<br>med:0.112<br>(0.104-0.120) | n=699<br>med:0.093<br>(0.086-0.104) | n=445<br>med:0.082<br>(0.075-0.090) | n=1,135<br>med:0.060<br>(0.055-0.066) |
| Gorilla | n=834<br>med:0.134<br>(0.123-0.144) | n=762<br>med:0.120<br>(0.113-0.129) | n=719<br>med:0.105<br>(0.097-0.116) | n=447<br>med:0.089<br>(0.080-0.098) | n=1,120<br>med:0.069<br>(0.063-0.073) |
| Cat | n=852<br>med:0.127<br>(0.119-0.135) | n=785<br>med:0.120<br>(0.111-0.126) | n=722<br>med:0.101<br>(0.091-0.106) | n=447<br>med:0.081<br>(0.074-0.091) | n=1,119<br>med:0.067<br>(0.062-0.071) |

#### Mouse Reference Genome, Zeisel et al., 2015, three cell types

| Sample Size and Median dN/dS (CI95%) | Glia-specific Genes | Vasculature-specific Genes | Neuron-specific Genes |
|---|---|---|---|
| Microbat | n=1,015 med:0.122 (0.113-0.131) | n=345 med:0.108 (0.096-0.117) | n=663 med:0.079 (0.073-0.089) |
| Hedgehog | n=246 med:0.121 (0.104-0.148) | n=78 med:0.115 (0.078-0.140) | n=142 med:0.110 (0.084-0.133) |
| Naked mole-rat male | n=1,000 med:0.124 (0.116-0.133) | n=345 med:0.103 (0.093-0.118) | n=678 med:0.072 (0.064-0.079) |
| Sooty mangabey | n=1,176 med:0.124 (0.118-0.131) | n=398 med:0.103 (0.092-0.115) | n=797 med:0.072 (0.066-0.077) |
| Olive baboon | n=1,173 med:0.125 (0.117-0.134) | n=396 med:0.103 (0.093-0.119) | n=786 med:0.073 (0.066-0.078) |
| American bison | n=978 med:0.117 (0.109-0.127) | n=327 med:0.098 (0.088-0.111) | n=608 med:0.073 (0.066-0.079) |
| Leopard | n=1,120 med:0.112 (0.106-0.122) | n=374 med:0.094 (0.084-0.105) | n=778 med:0.066 (0.061-0.071) |
| Bolivian squirrel monkey | n=1,120 med:0.129 (0.120-0.136) | n=380 med:0.104 (0.096-0.120) | n=765 med:0.069 (0.063-0.077) |
| Crab-eating macaque | n=1,185 med:0.124 (0.117-0.131) | n=397 med:0.105 (0.094-0.116) | n=779 med:0.072 (0.062-0.077) |
| Human | n=1,204 med:0.122 (0.115-0.128) | n=412 med:0.099 (0.089-0.108) | n=823 med:0.066 (0.060-0.072) |
| Red fox | n=1,097 med:0.124 (0.114-0.131) | n=368 med:0.104 (0.098-0.117) | n=723 med:0.079 (0.072-0.087) |
| Shrew mouse | n=1,293 med:0.135 (0.126-0.146) | n=423 med:0.105 (0.092-0.119) | n=856 med:0.069 (0.061-0.075) |
| Arctic ground squirrel | n=1,146 med:0.120 (0.108-0.127) | n=395 med:0.100 (0.091-0.112) | n=807 med:0.060 (0.054-0.067) |
| Chinese hamster CriGri | n=1,182 med:0.141 (0.132-0.149) | n=380 med:0.113 (0.105-0.130) | n=768 med:0.083 (0.074-0.095) |
| Opossum | n=1,034 | n=353 | n=708 |

| | | | |
|---|---|---|---|
| | med:0.097<br>(0.088-0.102) | med:0.082<br>(0.070-0.094) | med:0.065<br>(0.059-0.071) |
| Dolphin | n=640<br>med:0.130<br>(0.119-0.140) | n=209<br>med:0.117<br>(0.098-0.133) | n=399<br>med:0.076<br>(0.071-0.087) |
| Capuchin | n=1,142<br>med:0.125<br>(0.119-0.133) | n=391<br>med:0.103<br>(0.094-0.116) | n=773<br>med:0.070<br>(0.063-0.077) |
| Mouse Lemur | n=1,134<br>med:0.120<br>(0.113-0.128) | n=380<br>med:0.106<br>(0.095-0.118) | n=763<br>med:0.067<br>(0.061-0.074) |
| Damara mole rat | n=1,049<br>med:0.117<br>(0.109-0.124) | n=360<br>med:0.103<br>(0.091-0.112) | n=727<br>med:0.066<br>(0.059-0.074) |
| Marmoset | n=1,141<br>med:0.130<br>(0.121-0.137) | n=394<br>med:0.104<br>(0.097-0.115) | n=799<br>med:0.074<br>(0.066-0.082) |
| Goat | n=1,151<br>med:0.112<br>(0.105-0.118) | n=373<br>med:0.100<br>(0.090-0.112) | n=750<br>med:0.071<br>(0.066-0.079) |
| Common wombat | n=972<br>med:0.095<br>(0.090-0.100) | n=337<br>med:0.082<br>(0.075-0.089) | n=683<br>med:0.061<br>(0.056-0.067) |
| American black bear | n=1,017<br>med:0.125<br>(0.115-0.134) | n=340<br>med:0.109<br>(0.097-0.124) | n=635<br>med:0.079<br>(0.071-0.086) |
| Sloth | n=185<br>med:0.167<br>(0.145-0.194) | n=49<br>med:0.138<br>(0.104-0.170) | n=96<br>med:0.087<br>(0.058-0.108) |
| Polar bear | n=1,025<br>med:0.130<br>(0.122-0.137) | n=338<br>med:0.111<br>(0.100-0.125) | n=644<br>med:0.084<br>(0.076-0.090) |
| Alpine marmot | n=1,099<br>med:0.122<br>(0.112-0.129) | n=379<br>med:0.100<br>(0.091-0.110) | n=736<br>med:0.065<br>(0.059-0.074) |
| Prairie vole | n=1,204<br>med:0.131<br>(0.123-0.138) | n=394<br>med:0.106<br>(0.095-0.118) | n=824<br>med:0.065<br>(0.061-0.070) |
| Donkey | n=1,160<br>med:0.128<br>(0.120-0.132) | n=390<br>med:0.100<br>(0.092-0.109) | n=783<br>med:0.077<br>(0.070-0.082) |
| Golden Hamster | n=1,154<br>med:0.144<br>(0.134-0.150) | n=382<br>med:0.123<br>(0.102-0.134) | n=797<br>med:0.072<br>(0.066-0.082) |
| Armadillo | n=1,060<br>med:0.120<br>(0.115-0.127) | n=357<br>med:0.101<br>(0.093-0.112) | n=703<br>med:0.071<br>(0.067-0.078) |
| Ugandan red Colobus | n=1,135 | n=376 | n=776 |

| | | | |
|---|---|---|---|
| | med:0.124<br>(0.117-0.133) | med:0.102<br>(0.093-0.116) | med:0.072<br>(0.065-0.079) |
| Pig-tailed macaque | n=1,173<br>med:0.126<br>(0.119-0.135) | n=403<br>med:0.109<br>(0.094-0.121) | n=795<br>med:0.070<br>(0.063-0.077) |
| Panda | n=1,129<br>med:0.112<br>(0.107-0.120) | n=376<br>med:0.095<br>(0.086-0.108) | n=746<br>med:0.067<br>(0.061-0.074) |
| Ferret | n=1,121<br>med:0.115<br>(0.109-0.124) | n=367<br>med:0.099<br>(0.089-0.110) | n=748<br>med:0.070<br>(0.063-0.075) |
| Koala | n=1,053<br>med:0.097<br>(0.093-0.104) | n=364<br>med:0.080<br>(0.070-0.089) | n=747<br>med:0.064<br>(0.059-0.069) |
| Wallaby | n=291<br>med:0.086<br>(0.078-0.098) | n=112<br>med:0.083<br>(0.072-0.093) | n=211<br>med:0.066<br>(0.054-0.074) |
| Tarsier | n=962<br>med:0.123<br>(0.115-0.129) | n=306<br>med:0.111<br>(0.098-0.124) | n=607<br>med:0.084<br>(0.075-0.090) |
| Degu | n=974<br>med:0.117<br>(0.109-0.124) | n=340<br>med:0.107<br>(0.097-0.119) | n=685<br>med:0.069<br>(0.061-0.075) |
| Chinese hamster<br>CHOK1GS | n=1,273<br>med:0.135<br>(0.128-0.143) | n=417<br>med:0.110<br>(0.098-0.126) | n=854<br>med:0.073<br>(0.065-0.079) |
| Drill | n=1,149<br>med:0.126<br>(0.120-0.133) | n=380<br>med:0.104<br>(0.091-0.119) | n=749<br>med:0.073<br>(0.066-0.080) |
| Gibbon | n=1,124<br>med:0.134<br>(0.127-0.143) | n=384<br>med:0.107<br>(0.097-0.120) | n=756<br>med:0.080<br>(0.073-0.086) |
| Rat | n=1,250<br>med:0.134<br>(0.125-0.142) | n=409<br>med:0.117<br>(0.105-0.129) | n=837<br>med:0.077<br>(0.066-0.084) |
| Macaque | n=1,189<br>med:0.127<br>(0.119-0.136) | n=406<br>med:0.102<br>(0.092-0.117) | n=813<br>med:0.073<br>(0.065-0.079) |
| American mink | n=1,072<br>med:0.112<br>(0.106-0.121) | n=357<br>med:0.095<br>(0.082-0.110) | n=723<br>med:0.069<br>(0.063-0.075) |
| Gelada | n=1,183<br>med:0.121<br>(0.115-0.130) | n=398<br>med:0.101<br>(0.091-0.117) | n=798<br>med:0.068<br>(0.062-0.075) |
| Sheep | n=1,119<br>med:0.122<br>(0.113-0.128) | n=359<br>med:0.111<br>(0.102-0.119) | n=720<br>med:0.080<br>(0.074-0.085) |
| Tasmanian devil | n=971 | n=336 | n=639 |

| | | | |
|---|---|---|---|
| | med:0.092 (0.087-0.098) | med:0.079 (0.073-0.089) | med:0.062 (0.057-0.068) |
| Elephant | n=1,138 med:0.120 (0.112-0.127) | n=377 med:0.105 (0.097-0.114) | n=742 med:0.077 (0.069-0.083) |
| Algerian mouse | n=1,311 med:0.137 (0.128-0.150) | n=422 med:0.106 (0.092-0.119) | n=862 med:0.064 (0.057-0.074) |
| Dingo | n=1,100 med:0.118 (0.112-0.124) | n=364 med:0.100 (0.090-0.108) | n=746 med:0.070 (0.065-0.077) |
| Golden snub-nosed monkey | n=1,126 med:0.127 (0.120-0.134) | n=380 med:0.106 (0.096-0.118) | n=753 med:0.071 (0.064-0.077) |
| Horse | n=1,161 med:0.128 (0.119-0.133) | n=396 med:0.105 (0.097-0.113) | n=784 med:0.076 (0.067-0.081) |
| Black snub-nosed monkey | n=1,129 med:0.132 (0.123-0.139) | n=381 med:0.111 (0.100-0.126) | n=754 med:0.077 (0.070-0.083) |
| Brazilian guinea pig | n=831 med:0.125 (0.119-0.131) | n=283 med:0.118 (0.104-0.131) | n=548 med:0.091 (0.084-0.099) |
| Orangutan | n=1,122 med:0.123 (0.113-0.129) | n=391 med:0.099 (0.090-0.113) | n=761 med:0.068 (0.061-0.074) |
| Northern American deer mouse | n=1,235 med:0.132 (0.122-0.142) | n=403 med:0.101 (0.091-0.118) | n=803 med:0.066 (0.061-0.073) |
| Naked mole-rat female | n=1,054 med:0.124 (0.116-0.133) | n=360 med:0.103 (0.095-0.118) | n=734 med:0.071 (0.062-0.079) |
| Daurian ground squirrel | n=1,002 med:0.128 (0.120-0.137) | n=338 med:0.105 (0.093-0.120) | n=616 med:0.078 (0.069-0.086) |
| Tiger | n=953 med:0.126 (0.120-0.135) | n=305 med:0.114 (0.097-0.125) | n=603 med:0.079 (0.070-0.085) |
| Lesser hedgehog tenrec | n=277 med:0.122 (0.108-0.145) | n=92 med:0.118 (0.100-0.142) | n=154 med:0.085 (0.070-0.100) |
| Mongolian gerbil | n=1,147 med:0.139 (0.130-0.150) | n=372 med:0.113 (0.090-0.127) | n=722 med:0.074 (0.065-0.084) |
| Ma's night monkey | n=1,116 med:0.130 (0.122-0.138) | n=380 med:0.110 (0.100-0.120) | n=769 med:0.074 (0.067-0.083) |

| | | | |
|---|---|---|---|
| Upper Galilee mountains blind mole rat | n=1,163<br>med:0.126<br>(0.115-0.131) | n=376<br>med:0.110<br>(0.098-0.124) | n=788<br>med:0.064<br>(0.058-0.071) |
| Kangaroo rat | n=1,007<br>med:0.111<br>(0.105-0.117) | n=343<br>med:0.101<br>(0.083-0.111) | n=694<br>med:0.067<br>(0.061-0.075) |
| Chimpanzee | n=1,114<br>med:0.128<br>(0.121-0.135) | n=379<br>med:0.102<br>(0.093-0.119) | n=763<br>med:0.069<br>(0.063-0.077) |
| Guinea Pig | n=1,091<br>med:0.116<br>(0.107-0.123) | n=379<br>med:0.103<br>(0.090-0.111) | n=763<br>med:0.070<br>(0.065-0.078) |
| American beaver | n=1,059<br>med:0.125<br>(0.118-0.133) | n=366<br>med:0.112<br>(0.095-0.130) | n=686<br>med:0.074<br>(0.067-0.081) |
| Wild yak | n=1,042<br>med:0.118<br>(0.110-0.127) | n=338<br>med:0.100<br>(0.086-0.111) | n=631<br>med:0.075<br>(0.071-0.083) |
| Coquerel's sifaka | n=1,113<br>med:0.121<br>(0.115-0.130) | n=372<br>med:0.103<br>(0.094-0.114) | n=760<br>med:0.073<br>(0.066-0.080) |
| Alpaca | n=254<br>med:0.174<br>(0.150-0.193) | n=74<br>med:0.121<br>(0.109-0.147) | n=148<br>med:0.081<br>(0.068-0.097) |
| Lesser Egyptian jerboa | n=1,064<br>med:0.121<br>(0.113-0.129) | n=342<br>med:0.101<br>(0.087-0.115) | n=731<br>med:0.068<br>(0.062-0.076) |
| Long-tailed chinchilla | n=1,102<br>med:0.115<br>(0.107-0.122) | n=376<br>med:0.093<br>(0.087-0.104) | n=766<br>med:0.064<br>(0.058-0.071) |
| Angola colobus | n=1,157<br>med:0.125<br>(0.117-0.131) | n=385<br>med:0.109<br>(0.094-0.124) | n=763<br>med:0.071<br>(0.065-0.079) |
| Ryukyu mouse | n=1,281<br>med:0.131<br>(0.120-0.141) | n=425<br>med:0.110<br>(0.093-0.124) | n=858<br>med:0.064<br>(0.058-0.073) |
| Chinese hamster PICR | n=1,248<br>med:0.141<br>(0.134-0.150) | n=408<br>med:0.120<br>(0.107-0.135) | n=849<br>med:0.074<br>(0.066-0.081) |
| Vervet-AGM | n=1,184<br>med:0.121<br>(0.113-0.128) | n=396<br>med:0.100<br>(0.089-0.114) | n=787<br>med:0.066<br>(0.060-0.074) |
| Cow | n=1,159<br>med:0.115<br>(0.107-0.122) | n=381<br>med:0.102<br>(0.088-0.110) | n=748<br>med:0.069<br>(0.063-0.075) |
| Bushbaby | n=1,155<br>med:0.127<br>(0.120-0.135) | n=391<br>med:0.108<br>(0.095-0.122) | n=777<br>med:0.072<br>(0.064-0.077) |

| | | | |
|---|---|---|---|
| Bonobo | n=1,150<br>med:0.129<br>(0.124-0.135) | n=387<br>med:0.108<br>(0.096-0.124) | n=751<br>med:0.075<br>(0.068-0.082) |
| Hyrax | n=332<br>med:0.131<br>(0.118-0.151) | n=113<br>med:0.110<br>(0.089-0.129) | n=198<br>med:0.093<br>(0.080-0.107) |
| Rabbit | n=1,031<br>med:0.117<br>(0.110-0.125) | n=348<br>med:0.099<br>(0.088-0.107) | n=691<br>med:0.078<br>(0.069-0.085) |
| Megabat | n=637<br>med:0.132<br>(0.122-0.144) | n=232<br>med:0.125<br>(0.108-0.136) | n=376<br>med:0.091<br>(0.081-0.103) |
| Dog | n=1,117<br>med:0.119<br>(0.112-0.129) | n=372<br>med:0.108<br>(0.097-0.115) | n=749<br>med:0.077<br>(0.067-0.082) |
| Steppe mouse | n=1,257<br>med:0.176<br>(0.160-0.194) | n=415<br>med:0.144<br>(0.116-0.162) | n=846<br>med:0.091<br>(0.083-0.105) |
| Greater bamboo lemur | n=1,158<br>med:0.119<br>(0.112-0.126) | n=386<br>med:0.104<br>(0.091-0.114) | n=776<br>med:0.073<br>(0.063-0.079) |
| Pig | n=1,155<br>med:0.120<br>(0.113-0.128) | n=391<br>med:0.100<br>(0.087-0.108) | n=779<br>med:0.067<br>(0.062-0.073) |
| Tree Shrew | n=222<br>med:0.137<br>(0.117-0.164) | n=79<br>med:0.115<br>(0.088-0.145) | n=145<br>med:0.089<br>(0.080-0.112) |
| Pika | n=409<br>med:0.122<br>(0.105-0.136) | n=149<br>med:0.096<br>(0.081-0.117) | n=216<br>med:0.075<br>(0.067-0.095) |
| Shrew | n=201<br>med:0.117<br>(0.101-0.135) | n=57<br>med:0.122<br>(0.070-0.132) | n=126<br>med:0.091<br>(0.064-0.112) |
| Squirrel | n=1,135<br>med:0.115<br>(0.109-0.124) | n=390<br>med:0.096<br>(0.086-0.109) | n=789<br>med:0.063<br>(0.057-0.067) |
| Gorilla | n=1,142<br>med:0.127<br>(0.120-0.134) | n=387<br>med:0.107<br>(0.095-0.121) | n=772<br>med:0.072<br>(0.066-0.080) |
| Cat | n=1,173<br>med:0.117<br>(0.111-0.126) | n=386<br>med:0.102<br>(0.092-0.116) | n=790<br>med:0.069<br>(0.063-0.077) |

### Mouse Reference Genome, Zeisel et al., 2015, cell subtypes

| Sample Size and Median dN/dS (CI95%) | Ependymal-specific Genes | Microglia-specific Genes | Endothelial-specific Genes | Mural-specific Genes | Oligodendrocyte-specific Genes | Astrocyte-specific Genes | Interneuron-specific Genes | S1 Pyramidal-specific Genes | CA1 Pyramidal-specific Genes |
|---|---|---|---|---|---|---|---|---|---|
| Microbat | n=298 med:0.153 (0.134-0.164) | n=263 med:0.139 (0.125-0.157) | n=242 med:0.112 (0.100-0.127) | n=103 med:0.090 (0.073-0.114) | n=295 med:0.099 (0.091-0.110) | n=159 med:0.104 (0.087-0.118) | n=241 med:0.090 (0.077-0.104) | n=177 med:0.078 (0.064-0.101) | n=245 med:0.073 (0.063-0.086) |
| Hedgehog | n=57 med:0.150 (0.104-0.179) | n=72 med:0.159 (0.120-0.185) | n=61 med:0.115 (0.071-0.140) | n=17 med:0.115 (0.056-0.149) | n=77 med:0.104 (0.080-0.122) | n=40 med:0.080 (0.044-0.116) | n=46 med:0.124 (0.084-0.159) | n=36 med:0.079 (0.063-0.131) | n=60 med:0.123 (0.069-0.154) |
| Naked mole-rat male | n=292 med:0.149 (0.130-0.167) | n=263 med:0.140 (0.122-0.155) | n=234 med:0.106 (0.093-0.121) | n=111 med:0.096 (0.080-0.119) | n=290 med:0.102 (0.095-0.118) | n=155 med:0.101 (0.086-0.116) | n=241 med:0.077 (0.061-0.086) | n=187 med:0.073 (0.059-0.088) | n=250 med:0.069 (0.060-0.079) |
| Sooty mangabey | n=338 med:0.155 (0.143-0.175) | n=311 med:0.140 (0.124-0.150) | n=278 med:0.109 (0.093-0.126) | n=120 med:0.094 (0.072-0.111) | n=339 med:0.106 (0.094-0.118) | n=188 med:0.100 (0.087-0.110) | n=279 med:0.076 (0.064-0.088) | n=217 med:0.071 (0.059-0.085) | n=301 med:0.069 (0.057-0.077) |
| Olive baboon | n=338 med:0.153 (0.137-0.172) | n=309 med:0.142 (0.123-0.151) | n=277 med:0.114 (0.094-0.128) | n=119 med:0.092 (0.079-0.115) | n=340 med:0.106 (0.095-0.117) | n=186 med:0.100 (0.089-0.110) | n=273 med:0.078 (0.066-0.093) | n=212 med:0.074 (0.060-0.081) | n=301 med:0.068 (0.059-0.078) |
| American bison | n=304 med:0.142 (0.124-0.159) | n=257 med:0.141 (0.118-0.155) | n=230 med:0.110 (0.089-0.127) | n=97 med:0.086 (0.073-0.103) | n=273 med:0.100 (0.091-0.108) | n=144 med:0.094 (0.080-0.107) | n=210 med:0.080 (0.067-0.093) | n=162 med:0.071 (0.060-0.082) | n=236 med:0.067 (0.056-0.078) |
| Leopard | n=340 med:0.134 (0.124-0.154) | n=280 med:0.131 (0.112-0.142) | n=256 med:0.097 (0.084-0.120) | n=118 med:0.088 (0.072-0.102) | n=324 med:0.093 (0.083-0.106) | n=176 med:0.093 (0.081-0.105) | n=279 med:0.069 (0.057-0.078) | n=201 med:0.065 (0.055-0.077) | n=298 med:0.065 (0.056-0.072) |
| Bolivian squirrel monkey | n=339 med:0.153 (0.138-0.170) | n=285 med:0.147 (0.128-0.159) | n=265 med:0.117 (0.097-0.128) | n=115 med:0.097 (0.081-0.112) | n=325 med:0.099 (0.089-0.118) | n=171 med:0.104 (0.090-0.122) | n=269 med:0.070 (0.062-0.091) | n=202 med:0.071 (0.059-0.086) | n=294 med:0.065 (0.057-0.078) |
| Crab-eating macaque | n=345 med:0.153 (0.139-0.171) | n=312 med:0.140 (0.123-0.147) | n=277 med:0.109 (0.094-0.123) | n=120 med:0.097 (0.073-0.116) | n=342 med:0.105 (0.094-0.117) | n=186 med:0.098 (0.089-0.110) | n=271 med:0.073 (0.061-0.083) | n=207 med:0.075 (0.056-0.090) | n=301 med:0.065 (0.058-0.076) |
| Human | n=352 med:0.153 (0.134-0.168) | n=309 med:0.136 (0.121-0.151) | n=288 med:0.102 (0.091-0.125) | n=124 med:0.089 (0.077-0.107) | n=351 med:0.099 (0.089-0.109) | n=192 med:0.100 (0.086-0.117) | n=289 med:0.072 (0.061-0.083) | n=214 med:0.067 (0.056-0.081) | n=320 med:0.062 (0.053-0.071) |
| Red fox | n=319 med:0.150 (0.133-0.162) | n=288 med:0.142 (0.123-0.155) | n=256 med:0.109 (0.100-0.124) | n=112 med:0.100 (0.090-0.108) | n=322 med:0.105 (0.098-0.113) | n=168 med:0.101 (0.087-0.116) | n=252 med:0.082 (0.065-0.094) | n=193 med:0.081 (0.067-0.099) | n=278 med:0.074 (0.064-0.085) |
| Shrew mouse | n=369 med:0.174 (0.148-0.191) | n=346 med:0.165 (0.146-0.189) | n=290 med:0.113 (0.093-0.135) | n=133 med:0.094 (0.078-0.113) | n=375 med:0.102 (0.090-0.119) | n=203 med:0.107 (0.091-0.122) | n=298 med:0.067 (0.054-0.078) | n=223 med:0.074 (0.057-0.093) | n=335 med:0.067 (0.059-0.076) |
| Arctic ground squirrel | n=324 med:0.141 (0.128-0.153) | n=294 med:0.137 (0.118-0.155) | n=273 med:0.107 (0.093-0.115) | n=122 med:0.087 (0.072-0.106) | n=343 med:0.093 (0.083-0.108) | n=185 med:0.096 (0.078-0.109) | n=283 med:0.066 (0.056-0.076) | n=211 med:0.063 (0.050-0.078) | n=313 med:0.053 (0.047-0.065) |
| Chinese hamster CriGri | n=342 med:0.167 (0.148-0.185) | n=317 med:0.165 (0.148-0.195) | n=261 med:0.121 (0.106-0.140) | n=119 med:0.106 (0.088-0.114) | n=346 med:0.115 (0.104-0.127) | n=177 med:0.113 (0.100-0.130) | n=278 med:0.083 (0.069-0.098) | n=198 med:0.083 (0.070-0.108) | n=292 med:0.084 (0.070-0.102) |
| Opossum | n=299 med:0.112 (0.103-0.129) | n=258 med:0.102 (0.086-0.113) | n=240 med:0.088 (0.072-0.101) | n=113 med:0.069 (0.055-0.093) | n=313 med:0.084 (0.070-0.096) | n=164 med:0.080 (0.067-0.096) | n=250 med:0.064 (0.055-0.073) | n=191 med:0.068 (0.056-0.074) | n=267 med:0.063 (0.054-0.073) |
| Dolphin | n=196 med:0.150 (0.130-0.175) | n=163 med:0.144 (0.122-0.160) | n=143 med:0.123 (0.103-0.152) | n=66 med:0.099 (0.077-0.123) | n=182 med:0.112 (0.101-0.135) | n=99 med:0.101 (0.079-0.123) | n=141 med:0.094 (0.074-0.111) | n=108 med:0.071 (0.054-0.085) | n=150 med:0.074 (0.062-0.086) |
| Capuchin | n=335 med:0.153 (0.139-0.166) | n=296 med:0.140 (0.125-0.151) | n=270 med:0.107 (0.095-0.125) | n=121 med:0.095 (0.083-0.108) | n=332 med:0.102 (0.090-0.116) | n=179 med:0.103 (0.086-0.121) | n=272 med:0.072 (0.063-0.085) | n=207 med:0.067 (0.055-0.084) | n=294 med:0.068 (0.059-0.080) |
| Mouse Lemur | n=336 med:0.148 (0.136-0.165) | n=298 med:0.138 (0.124-0.151) | n=267 med:0.113 (0.099-0.127) | n=113 med:0.096 (0.072-0.113) | n=323 med:0.097 (0.084-0.108) | n=177 med:0.101 (0.084-0.113) | n=271 med:0.072 (0.059-0.085) | n=206 med:0.072 (0.060-0.085) | n=286 med:0.061 (0.053-0.072) |
| Damara mole rat | n=305 med:0.144 (0.126-0.161) | n=269 med:0.137 (0.120-0.156) | n=248 med:0.106 (0.093-0.120) | n=112 med:0.095 (0.076-0.111) | n=312 med:0.095 (0.081-0.108) | n=163 med:0.092 (0.082-0.110) | n=264 med:0.067 (0.057-0.079) | n=189 med:0.073 (0.059-0.092) | n=274 med:0.060 (0.052-0.076) |
| Marmoset | n=341 med:0.164 (0.142-0.175) | n=298 med:0.143 (0.128-0.156) | n=275 med:0.112 (0.098-0.134) | n=119 med:0.095 (0.079-0.110) | n=326 med:0.104 (0.090-0.117) | n=176 med:0.114 (0.094-0.134) | n=283 med:0.082 (0.066-0.096) | n=211 med:0.080 (0.063-0.098) | n=305 med:0.064 (0.055-0.078) |
| Goat | n=337 med:0.128 (0.119-0.146) | n=294 med:0.134 (0.117-0.154) | n=258 med:0.105 (0.091-0.124) | n=115 med:0.088 (0.079-0.108) | n=344 med:0.095 (0.083-0.105) | n=176 med:0.097 (0.088-0.109) | n=262 med:0.077 (0.066-0.085) | n=200 med:0.070 (0.060-0.085) | n=288 med:0.067 (0.061-0.079) |
| Common wombat | n=267 med:0.110 (0.100-0.123) | n=261 med:0.104 (0.096-0.117) | n=232 med:0.084 (0.078-0.097) | n=105 med:0.072 (0.059-0.089) | n=292 med:0.082 (0.072-0.090) | n=152 med:0.083 (0.068-0.095) | n=226 med:0.060 (0.055-0.069) | n=186 med:0.067 (0.056-0.075) | n=271 med:0.058 (0.051-0.071) |
| American black bear | n=289 med:0.148 (0.130-0.166) | n=261 med:0.146 (0.126-0.158) | n=233 med:0.119 (0.102-0.135) | n=107 med:0.092 (0.080-0.108) | n=303 med:0.104 (0.088-0.115) | n=164 med:0.106 (0.091-0.123) | n=224 med:0.086 (0.075-0.099) | n=161 med:0.072 (0.063-0.095) | n=250 med:0.073 (0.064-0.082) |
| Sloth | n=64 med:0.195 (0.155-0.236) | n=49 med:0.160 (0.111-0.226) | n=39 med:0.139 (0.099-0.178) | n=10 med:0.137 (0.055-0.276) | n=54 med:0.158 (0.097-0.186) | n=18 med:0.130 (0.064-0.211) | n=38 med:0.089 (0.051-0.125) | n=26 med:0.100 (0.043-0.152) | n=32 med:0.084 (0.049-0.115) |
| Polar bear | n=301 med:0.163 (0.147-0.173) | n=270 med:0.143 (0.125-0.161) | n=239 med:0.120 (0.109-0.141) | n=99 med:0.092 | n=304 med:0.105 (0.097-0.119) | n=150 med:0.100 (0.089-0.120) | n=230 med:0.086 (0.071-0.100) | n=160 med:0.088 (0.072-0.101) | n=254 med:0.082 (0.072-0.089) |

| | | | | | | | | | |
|---|---|---|---|---|---|---|---|---|---|
| | | | | (0.077-0.107) | | | | | |
| Alpine marmot | n=315 med:0.149 (0.133-0.162) | n=299 med:0.133 (0.115-0.153) | n=262 med:0.105 (0.096-0.130) | n=117 med:0.091 (0.076-0.102) | n=315 med:0.102 (0.088-0.115) | n=170 med:0.096 (0.081-0.119) | n=251 med:0.071 (0.058-0.079) | n=192 med:0.074 (0.058-0.088) | n=293 med:0.058 (0.050-0.070) |
| Prairie vole | n=347 med:0.154 (0.136-0.172) | n=317 med:0.165 (0.141-0.181) | n=272 med:0.108 (0.091-0.122) | n=122 med:0.100 (0.082-0.125) | n=356 med:0.098 (0.079-0.118) | n=184 med:0.103 (0.082-0.117) | n=289 med:0.065 (0.057-0.078) | n=220 med:0.065 (0.056-0.077) | n=315 med:0.065 (0.056-0.074) |
| Donkey | n=339 med:0.155 (0.139-0.169) | n=299 med:0.141 (0.127-0.157) | n=265 med:0.106 (0.096-0.119) | n=125 med:0.091 (0.081-0.103) | n=342 med:0.105 (0.093-0.117) | n=180 med:0.103 (0.087-0.117) | n=269 med:0.082 (0.066-0.093) | n=208 med:0.081 (0.070-0.092) | n=306 med:0.070 (0.061-0.080) |
| Golden Hamster | n=332 med:0.161 (0.144-0.180) | n=302 med:0.161 (0.150-0.185) | n=263 med:0.130 (0.109-0.143) | n=119 med:0.097 (0.077-0.128) | n=343 med:0.115 (0.100-0.131) | n=177 med:0.118 (0.100-0.146) | n=277 med:0.069 (0.059-0.081) | n=209 med:0.079 (0.060-0.111) | n=311 med:0.076 (0.061-0.088) |
| Armadillo | n=304 med:0.148 (0.133-0.162) | n=286 med:0.135 (0.121-0.147) | n=249 med:0.106 (0.086-0.123) | n=108 med:0.099 (0.077-0.109) | n=304 med:0.101 (0.091-0.114) | n=166 med:0.103 (0.086-0.118) | n=241 med:0.068 (0.063-0.086) | n=186 med:0.080 (0.073-0.089) | n=276 med:0.067 (0.060-0.073) |
| Ugandan red Colobus | n=327 med:0.157 (0.137-0.172) | n=296 med:0.136 (0.124-0.150) | n=266 med:0.110 (0.094-0.129) | n=110 med:0.094 (0.077-0.107) | n=328 med:0.105 (0.092-0.115) | n=184 med:0.103 (0.091-0.115) | n=278 med:0.079 (0.067-0.095) | n=204 med:0.070 (0.056-0.083) | n=294 med:0.067 (0.058-0.077) |
| Pig-tailed macaque | n=339 med:0.156 (0.144-0.170) | n=310 med:0.145 (0.129-0.153) | n=282 med:0.115 (0.096-0.126) | n=121 med:0.094 (0.075-0.115) | n=342 med:0.106 (0.095-0.118) | n=182 med:0.098 (0.086-0.109) | n=278 med:0.073 (0.062-0.088) | n=211 med:0.073 (0.056-0.085) | n=306 med:0.066 (0.057-0.077) |
| Panda | n=327 med:0.143 (0.120-0.159) | n=293 med:0.136 (0.122-0.150) | n=267 med:0.106 (0.090-0.121) | n=109 med:0.086 (0.077-0.099) | n=334 med:0.094 (0.084-0.101) | n=175 med:0.093 (0.077-0.103) | n=262 med:0.077 (0.061-0.087) | n=197 med:0.067 (0.060-0.084) | n=287 med:0.061 (0.054-0.072) |
| Ferret | n=334 med:0.146 (0.125-0.160) | n=290 med:0.135 (0.117-0.153) | n=258 med:0.104 (0.091-0.123) | n=109 med:0.090 (0.072-0.106) | n=323 med:0.092 (0.079-0.102) | n=174 med:0.095 (0.084-0.105) | n=260 med:0.076 (0.065-0.087) | n=200 med:0.070 (0.059-0.088) | n=288 med:0.065 (0.052-0.073) |
| Koala | n=296 med:0.122 (0.104-0.133) | n=269 med:0.106 (0.094-0.119) | n=249 med:0.081 (0.069-0.090) | n=115 med:0.076 (0.066-0.095) | n=325 med:0.089 (0.076-0.095) | n=163 med:0.084 (0.072-0.097) | n=264 med:0.066 (0.058-0.075) | n=201 med:0.066 (0.057-0.072) | n=282 med:0.059 (0.053-0.072) |
| Wallaby | n=80 med:0.095 (0.075-0.124) | n=76 med:0.085 (0.069-0.119) | n=75 med:0.083 (0.071-0.100) | n=37 med:0.081 (0.063-0.103) | n=98 med:0.083 (0.061-0.103) | n=37 med:0.086 (0.063-0.110) | n=78 med:0.068 (0.048-0.080) | n=62 med:0.063 (0.048-0.075) | n=71 med:0.066 (0.047-0.089) |
| Tarsier | n=282 med:0.160 (0.145-0.172) | n=247 med:0.137 (0.121-0.156) | n=219 med:0.115 (0.098-0.128) | n=87 med:0.109 (0.074-0.129) | n=288 med:0.099 (0.090-0.112) | n=145 med:0.105 (0.091-0.119) | n=220 med:0.086 (0.071-0.093) | n=159 med:0.086 (0.070-0.096) | n=228 med:0.080 (0.064-0.092) |
| Degu | n=287 med:0.136 (0.123-0.151) | n=239 med:0.128 (0.116-0.145) | n=228 med:0.111 (0.094-0.128) | n=112 med:0.101 (0.087-0.119) | n=295 med:0.100 (0.086-0.112) | n=153 med:0.092 (0.079-0.112) | n=240 med:0.069 (0.056-0.079) | n=183 med:0.076 (0.060-0.094) | n=262 med:0.064 (0.053-0.073) |
| Chinese hamster CHOK1GS | n=365 med:0.160 (0.144-0.183) | n=342 med:0.159 (0.141-0.181) | n=287 med:0.120 (0.102-0.137) | n=130 med:0.098 (0.083-0.112) | n=367 med:0.108 (0.093-0.120) | n=199 med:0.106 (0.089-0.119) | n=303 med:0.071 (0.058-0.087) | n=227 med:0.070 (0.055-0.083) | n=324 med:0.075 (0.062-0.091) |
| Drill | n=336 med:0.158 (0.144-0.174) | n=303 med:0.137 (0.124-0.152) | n=263 med:0.113 (0.092-0.125) | n=117 med:0.091 (0.069-0.107) | n=328 med:0.109 (0.096-0.118) | n=182 med:0.099 (0.086-0.115) | n=260 med:0.081 (0.066-0.095) | n=203 med:0.074 (0.060-0.088) | n=286 med:0.069 (0.058-0.079) |
| Gibbon | n=334 med:0.167 (0.152-0.179) | n=294 med:0.151 (0.135-0.170) | n=268 med:0.110 (0.098-0.133) | n=116 med:0.098 (0.081-0.115) | n=317 med:0.111 (0.098-0.123) | n=179 med:0.109 (0.091-0.125) | n=265 med:0.088 (0.077-0.098) | n=200 med:0.082 (0.069-0.099) | n=291 med:0.071 (0.062-0.080) |
| Rat | n=362 med:0.162 (0.143-0.191) | n=337 med:0.170 (0.147-0.195) | n=284 med:0.120 (0.102-0.134) | n=125 med:0.115 (0.090-0.131) | n=359 med:0.103 (0.090-0.119) | n=192 med:0.107 (0.090-0.121) | n=292 med:0.079 (0.064-0.091) | n=226 med:0.080 (0.059-0.089) | n=319 med:0.070 (0.060-0.083) |
| Macaque | n=340 med:0.152 (0.136-0.167) | n=318 med:0.144 (0.131-0.152) | n=282 med:0.104 (0.092-0.123) | n=124 med:0.098 (0.080-0.121) | n=342 med:0.104 (0.090-0.116) | n=189 med:0.103 (0.091-0.127) | n=287 med:0.079 (0.062-0.096) | n=214 med:0.075 (0.064-0.091) | n=312 med:0.066 (0.056-0.075) |
| American mink | n=325 med:0.134 (0.118-0.146) | n=265 med:0.133 (0.119-0.147) | n=244 med:0.102 (0.088-0.119) | n=113 med:0.079 (0.067-0.099) | n=318 med:0.094 (0.082-0.107) | n=164 med:0.095 (0.088-0.102) | n=253 med:0.075 (0.064-0.082) | n=197 med:0.068 (0.058-0.079) | n=273 med:0.068 (0.059-0.075) |
| Gelada | n=340 med:0.154 (0.136-0.169) | n=311 med:0.138 (0.121-0.146) | n=278 med:0.108 (0.092-0.125) | n=120 med:0.090 (0.072-0.116) | n=346 med:0.101 (0.090-0.114) | n=186 med:0.096 (0.083-0.106) | n=277 med:0.072 (0.057-0.084) | n=213 med:0.070 (0.055-0.080) | n=308 med:0.066 (0.058-0.078) |
| Sheep | n=339 med:0.144 (0.128-0.159) | n=280 med:0.143 (0.123-0.160) | n=249 med:0.115 (0.102-0.127) | n=110 med:0.102 (0.085-0.112) | n=325 med:0.101 (0.094-0.111) | n=175 med:0.103 (0.091-0.113) | n=251 med:0.079 (0.071-0.089) | n=189 med:0.079 (0.067-0.091) | n=280 med:0.082 (0.072-0.089) |
| Tasmanian devil | n=280 med:0.105 (0.094-0.118) | n=250 med:0.099 (0.087-0.111) | n=234 med:0.079 (0.072-0.089) | n=102 med:0.079 (0.053-0.094) | n=295 med:0.081 (0.071-0.093) | n=146 med:0.084 (0.072-0.094) | n=223 med:0.065 (0.055-0.075) | n=170 med:0.059 (0.047-0.071) | n=246 med:0.061 (0.054-0.073) |
| Elephant | n=331 med:0.146 (0.129-0.157) | n=288 med:0.135 (0.118-0.152) | n=260 med:0.108 (0.100-0.132) | n=117 med:0.096 (0.082-0.112) | n=345 med:0.099 (0.087-0.112) | n=174 med:0.097 (0.083-0.111) | n=269 med:0.082 (0.067-0.091) | n=194 med:0.083 (0.069-0.096) | n=279 med:0.070 (0.060-0.078) |
| Algerian mouse | n=375 med:0.167 (0.145-0.195) | n=356 med:0.173 (0.142-0.213) | n=292 med:0.112 (0.097-0.135) | n=130 med:0.083 (0.058-0.116) | n=372 med:0.114 (0.095-0.133) | n=208 med:0.096 (0.079-0.119) | n=302 med:0.064 (0.043-0.076) | n=228 med:0.059 (0.044-0.080) | n=332 med:0.072 (0.057-0.094) |
| Dingo | n=320 med:0.141 (0.124-0.156) | n=291 med:0.133 (0.120-0.146) | n=255 med:0.101 (0.090-0.114) | n=109 med:0.092 (0.079-0.112) | n=318 med:0.099 (0.085-0.110) | n=171 med:0.098 (0.080-0.114) | n=260 med:0.067 (0.059-0.081) | n=194 med:0.076 (0.062-0.086) | n=292 med:0.070 (0.058-0.077) |
| Golden snub-nosed monkey | n=332 med:0.156 (0.143-0.172) | n=300 med:0.140 (0.126-0.152) | n=269 med:0.111 (0.095-0.127) | n=111 med:0.099 (0.083-0.125) | n=322 med:0.107 (0.096-0.119) | n=172 med:0.101 (0.088-0.114) | n=262 med:0.075 (0.063-0.090) | n=199 med:0.071 (0.058-0.083) | n=292 med:0.066 (0.058-0.079) |

| | | | | | | | | | |
|---|---|---|---|---|---|---|---|---|---|
| Horse | n=345 med:0.144 (0.131-0.166) | n=299 med:0.140 (0.128-0.155) | n=273 med:0.108 (0.097-0.125) | n=123 med:0.097 (0.084-0.107) | n=333 med:0.114 (0.096-0.124) | n=184 med:0.106 (0.088-0.121) | n=273 med:0.081 (0.071-0.097) | n=210 med:0.077 (0.060-0.086) | n=301 med:0.068 (0.058-0.079) |
| Black snub-nosed monkey | n=333 med:0.166 (0.147-0.175) | n=290 med:0.147 (0.133-0.160) | n=267 med:0.116 (0.105-0.135) | n=114 med:0.103 (0.084-0.118) | n=334 med:0.107 (0.098-0.119) | n=172 med:0.104 (0.088-0.120) | n=265 med:0.077 (0.067-0.094) | n=199 med:0.079 (0.064-0.092) | n=290 med:0.074 (0.065-0.082) |
| Brazilian guinea pig | n=235 med:0.136 (0.122-0.148) | n=210 med:0.142 (0.127-0.154) | n=188 med:0.125 (0.108-0.137) | n=95 med:0.107 (0.093-0.131) | n=258 med:0.115 (0.107-0.122) | n=128 med:0.105 (0.092-0.122) | n=193 med:0.087 (0.077-0.099) | n=144 med:0.097 (0.082-0.105) | n=211 med:0.090 (0.081-0.105) |
| Orangutan | n=313 med:0.144 (0.133-0.164) | n=299 med:0.136 (0.122-0.152) | n=272 med:0.105 (0.090-0.122) | n=119 med:0.092 (0.082-0.119) | n=328 med:0.097 (0.084-0.112) | n=182 med:0.109 (0.095-0.121) | n=269 med:0.073 (0.060-0.084) | n=199 med:0.072 (0.060-0.084) | n=293 med:0.061 (0.052-0.070) |
| Northern American deer mouse | n=358 med:0.156 (0.144-0.174) | n=326 med:0.156 (0.134-0.175) | n=277 med:0.109 (0.093-0.128) | n=126 med:0.091 (0.072-0.111) | n=358 med:0.105 (0.092-0.113) | n=193 med:0.098 (0.086-0.122) | n=277 med:0.066 (0.056-0.073) | n=219 med:0.064 (0.058-0.081) | n=307 med:0.067 (0.056-0.081) |
| Naked mole-rat female | n=311 med:0.151 (0.134-0.171) | n=272 med:0.137 (0.124-0.154) | n=244 med:0.108 (0.097-0.131) | n=116 med:0.096 (0.083-0.116) | n=307 med:0.104 (0.089-0.117) | n=164 med:0.098 (0.086-0.109) | n=258 med:0.075 (0.061-0.085) | n=205 med:0.071 (0.058-0.091) | n=271 med:0.065 (0.057-0.077) |
| Daurian ground squirrel | n=294 med:0.153 (0.137-0.166) | n=263 med:0.141 (0.125-0.161) | n=234 med:0.114 (0.094-0.132) | n=104 med:0.096 (0.080-0.108) | n=295 med:0.113 (0.098-0.123) | n=150 med:0.098 (0.084-0.118) | n=218 med:0.078 (0.069-0.090) | n=169 med:0.085 (0.063-0.097) | n=229 med:0.071 (0.056-0.089) |
| Tiger | n=295 med:0.149 (0.130-0.167) | n=250 med:0.144 (0.123-0.161) | n=209 med:0.118 (0.097-0.133) | n=96 med:0.103 (0.084-0.121) | n=265 med:0.101 (0.086-0.117) | n=143 med:0.107 (0.090-0.124) | n=213 med:0.089 (0.072-0.104) | n=159 med:0.079 (0.063-0.092) | n=231 med:0.071 (0.061-0.082) |
| Lesser hedgehog tenrec | n=82 med:0.158 (0.119-0.186) | n=73 med:0.146 (0.107-0.177) | n=67 med:0.131 (0.105-0.159) | n=25 med:0.104 (0.056-0.112) | n=83 med:0.088 (0.069-0.119) | n=39 med:0.114 (0.078-0.153) | n=61 med:0.092 (0.068-0.127) | n=31 med:0.086 (0.058-0.121) | n=62 med:0.079 (0.056-0.100) |
| Mongolian gerbil | n=321 med:0.188 (0.158-0.208) | n=318 med:0.164 (0.145-0.182) | n=258 med:0.116 (0.090-0.134) | n=114 med:0.095 (0.076-0.133) | n=332 med:0.106 (0.091-0.129) | n=176 med:0.116 (0.096-0.130) | n=254 med:0.070 (0.055-0.086) | n=190 med:0.088 (0.067-0.104) | n=278 med:0.069 (0.061-0.086) |
| Ma's night monkey | n=324 med:0.157 (0.141-0.177) | n=291 med:0.148 (0.129-0.158) | n=267 med:0.116 (0.101-0.133) | n=113 med:0.099 (0.081-0.112) | n=322 med:0.104 (0.095-0.120) | n=179 med:0.103 (0.089-0.124) | n=270 med:0.080 (0.066-0.094) | n=207 med:0.074 (0.061-0.087) | n=292 med:0.070 (0.056-0.084) |
| Upper Galilee mountains blind mole rat | n=335 med:0.144 (0.127-0.156) | n=304 med:0.146 (0.133-0.164) | n=256 med:0.112 (0.096-0.131) | n=120 med:0.104 (0.087-0.129) | n=345 med:0.099 (0.085-0.111) | n=179 med:0.093 (0.078-0.112) | n=282 med:0.064 (0.055-0.076) | n=210 med:0.065 (0.055-0.082) | n=296 med:0.060 (0.056-0.072) |
| Kangaroo rat | n=294 med:0.138 (0.126-0.145) | n=255 med:0.122 (0.105-0.140) | n=239 med:0.101 (0.080-0.116) | n=104 med:0.100 (0.076-0.116) | n=291 med:0.092 (0.079-0.104) | n=167 med:0.093 (0.072-0.114) | n=250 med:0.068 (0.056-0.079) | n=184 med:0.071 (0.055-0.085) | n=260 med:0.064 (0.054-0.074) |
| Chimpanzee | n=324 med:0.155 (0.144-0.175) | n=287 med:0.138 (0.122-0.152) | n=263 med:0.111 (0.096-0.131) | n=116 med:0.091 (0.078-0.107) | n=323 med:0.109 (0.098-0.123) | n=180 med:0.097 (0.089-0.119) | n=263 med:0.075 (0.065-0.085) | n=203 med:0.071 (0.058-0.081) | n=297 med:0.062 (0.054-0.076) |
| Guinea Pig | n=313 med:0.136 (0.121-0.150) | n=277 med:0.135 (0.120-0.148) | n=261 med:0.104 (0.092-0.116) | n=118 med:0.094 (0.076-0.117) | n=335 med:0.098 (0.084-0.107) | n=166 med:0.098 (0.080-0.108) | n=271 med:0.070 (0.059-0.081) | n=201 med:0.083 (0.068-0.089) | n=291 med:0.066 (0.057-0.074) |
| American beaver | n=298 med:0.144 (0.133-0.157) | n=283 med:0.146 (0.124-0.167) | n=260 med:0.120 (0.098-0.138) | n=106 med:0.102 (0.081-0.124) | n=314 med:0.108 (0.097-0.120) | n=164 med:0.099 (0.082-0.118) | n=238 med:0.072 (0.058-0.081) | n=180 med:0.085 (0.065-0.102) | n=268 med:0.071 (0.062-0.083) |
| Wild yak | n=315 med:0.142 (0.126-0.155) | n=275 med:0.144 (0.116-0.159) | n=234 med:0.108 (0.092-0.126) | n=104 med:0.081 (0.068-0.103) | n=294 med:0.098 (0.090-0.108) | n=158 med:0.096 (0.082-0.111) | n=221 med:0.082 (0.072-0.098) | n=178 med:0.071 (0.063-0.078) | n=232 med:0.075 (0.060-0.091) |
| Coquerel's sifaka | n=336 med:0.148 (0.133-0.164) | n=283 med:0.132 (0.121-0.158) | n=261 med:0.110 (0.095-0.119) | n=111 med:0.097 (0.079-0.108) | n=321 med:0.099 (0.087-0.112) | n=173 med:0.104 (0.082-0.115) | n=265 med:0.077 (0.063-0.089) | n=203 med:0.077 (0.065-0.088) | n=292 med:0.070 (0.055-0.080) |
| Alpaca | n=81 med:0.201 (0.167-0.235) | n=62 med:0.194 (0.156-0.236) | n=55 med:0.146 (0.112-0.221) | n=19 med:0.101 (0.074-0.125) | n=70 med:0.144 (0.121-0.179) | n=41 med:0.103 (0.079-0.163) | n=59 med:0.080 (0.060-0.102) | n=30 med:0.088 (0.063-0.143) | n=59 med:0.070 (0.052-0.099) |
| Lesser Egyptian jerboa | n=320 med:0.138 (0.122-0.150) | n=264 med:0.138 (0.124-0.155) | n=226 med:0.102 (0.087-0.122) | n=116 med:0.099 (0.076-0.121) | n=318 med:0.100 (0.089-0.113) | n=162 med:0.108 (0.093-0.127) | n=260 med:0.066 (0.060-0.083) | n=196 med:0.072 (0.058-0.078) | n=275 med:0.064 (0.057-0.077) |
| Long-tailed chinchilla | n=323 med:0.139 (0.121-0.156) | n=285 med:0.137 (0.120-0.147) | n=256 med:0.098 (0.089-0.114) | n=120 med:0.086 (0.072-0.102) | n=330 med:0.098 (0.082-0.109) | n=164 med:0.094 (0.079-0.106) | n=274 med:0.065 (0.058-0.081) | n=200 med:0.067 (0.053-0.084) | n=292 med:0.059 (0.049-0.071) |
| Angola colobus | n=332 med:0.151 (0.134-0.168) | n=309 med:0.139 (0.127-0.150) | n=273 med:0.112 (0.097-0.130) | n=112 med:0.095 (0.078-0.125) | n=335 med:0.104 (0.091-0.117) | n=181 med:0.101 (0.086-0.113) | n=265 med:0.080 (0.067-0.091) | n=204 med:0.071 (0.062-0.083) | n=294 med:0.066 (0.056-0.075) |
| Ryukyu mouse | n=371 med:0.145 (0.129-0.177) | n=347 med:0.177 (0.156-0.197) | n=297 med:0.114 (0.093-0.132) | n=128 med:0.104 (0.079-0.120) | n=364 med:0.103 (0.089-0.118) | n=199 med:0.102 (0.084-0.119) | n=300 med:0.064 (0.053-0.074) | n=226 med:0.070 (0.047-0.086) | n=332 med:0.063 (0.056-0.077) |
| Chinese hamster PICR | n=363 med:0.170 (0.154-0.200) | n=326 med:0.163 (0.144-0.190) | n=282 med:0.132 (0.115-0.147) | n=126 med:0.105 (0.086-0.118) | n=359 med:0.114 (0.097-0.132) | n=200 med:0.107 (0.092-0.128) | n=300 med:0.075 (0.061-0.090) | n=219 med:0.074 (0.061-0.090) | n=330 med:0.074 (0.063-0.082) |
| Vervet-AGM | n=348 med:0.147 (0.131-0.161) | n=301 med:0.138 (0.123-0.149) | n=279 med:0.101 (0.087-0.119) | n=117 med:0.093 (0.076-0.106) | n=347 med:0.100 (0.090-0.114) | n=188 med:0.097 (0.085-0.110) | n=280 med:0.071 (0.061-0.080) | n=207 med:0.071 (0.056-0.086) | n=300 med:0.060 (0.052-0.068) |
| Cow | n=352 med:0.135 (0.118-0.152) | n=300 med:0.141 (0.123-0.154) | n=265 med:0.109 (0.089-0.120) | n=116 med:0.095 (0.074-0.105) | n=332 med:0.096 (0.088-0.106) | n=175 med:0.095 (0.085-0.104) | n=257 med:0.075 (0.066-0.086) | n=195 med:0.066 (0.061-0.084) | n=296 med:0.066 (0.054-0.073) |
| Bushbaby | n=336 | n=293 | n=272 | n=119 | n=341 | n=185 | n=274 | n=211 | n=292 |

| | | | | | | | | | |
|---|---|---|---|---|---|---|---|---|---|
| | med:0.152<br>(0.130-0.172) | med:0.147<br>(0.131-0.160) | med:0.111<br>(0.094-0.127) | med:0.102<br>(0.083-0.120) | med:0.104<br>(0.092-0.119) | med:0.103<br>(0.088-0.125) | med:0.076<br>(0.066-0.085) | med:0.073<br>(0.061-0.091) | med:0.066<br>(0.058-0.074) |
| Bonobo | n=333<br>med:0.156<br>(0.141-0.173) | n=298<br>med:0.141<br>(0.127-0.153) | n=269<br>med:0.114<br>(0.099-0.135) | n=118<br>med:0.097<br>(0.081-0.117) | n=338<br>med:0.103<br>(0.093-0.118) | n=181<br>med:0.115<br>(0.090-0.128) | n=262<br>med:0.078<br>(0.068-0.093) | n=202<br>med:0.073<br>(0.062-0.085) | n=287<br>med:0.072<br>(0.061-0.083) |
| Hyrax | n=83<br>med:0.175<br>(0.137-0.217) | n=98<br>med:0.163<br>(0.122-0.195) | n=81<br>med:0.111<br>(0.085-0.134) | n=32<br>med:0.110<br>(0.086-0.178) | n=103<br>med:0.116<br>(0.106-0.128) | n=48<br>med:0.096<br>(0.072-0.124) | n=72<br>med:0.083<br>(0.068-0.111) | n=58<br>med:0.112<br>(0.088-0.137) | n=68<br>med:0.086<br>(0.068-0.103) |
| Rabbit | n=306<br>med:0.142<br>(0.118-0.155) | n=263<br>med:0.132<br>(0.116-0.148) | n=238<br>med:0.104<br>(0.092-0.121) | n=110<br>med:0.088<br>(0.074-0.101) | n=298<br>med:0.106<br>(0.091-0.118) | n=164<br>med:0.095<br>(0.080-0.107) | n=244<br>med:0.081<br>(0.067-0.095) | n=181<br>med:0.082<br>(0.064-0.094) | n=266<br>med:0.070<br>(0.061-0.084) |
| Megabat | n=179<br>med:0.156<br>(0.142-0.173) | n=176<br>med:0.156<br>(0.131-0.185) | n=157<br>med:0.129<br>(0.106-0.144) | n=75<br>med:0.117<br>(0.091-0.132) | n=180<br>med:0.104<br>(0.087-0.117) | n=102<br>med:0.120<br>(0.089-0.151) | n=123<br>med:0.102<br>(0.087-0.113) | n=97<br>med:0.106<br>(0.082-0.122) | n=156<br>med:0.079<br>(0.063-0.091) |
| Dog | n=320<br>med:0.146<br>(0.132-0.166) | n=292<br>med:0.135<br>(0.118-0.146) | n=261<br>med:0.110<br>(0.099-0.129) | n=111<br>med:0.100<br>(0.085-0.111) | n=334<br>med:0.099<br>(0.091-0.106) | n=171<br>med:0.098<br>(0.088-0.113) | n=264<br>med:0.079<br>(0.067-0.093) | n=200<br>med:0.084<br>(0.061-0.101) | n=285<br>med:0.070<br>(0.059-0.079) |
| Steppe mouse | n=354<br>med:0.232<br>(0.198-0.266) | n=356<br>med:0.197<br>(0.167-0.235) | n=289<br>med:0.155<br>(0.118-0.177) | n=126<br>med:0.112<br>(0.071-0.152) | n=356<br>med:0.137<br>(0.119-0.156) | n=191<br>med:0.138<br>(0.102-0.167) | n=291<br>med:0.085<br>(0.066-0.119) | n=225<br>med:0.097<br>(0.074-0.113) | n=330<br>med:0.099<br>(0.077-0.124) |
| Greater bamboo lemur | n=339<br>med:0.143<br>(0.128-0.164) | n=302<br>med:0.133<br>(0.118-0.148) | n=266<br>med:0.112<br>(0.098-0.124) | n=120<br>med:0.084<br>(0.067-0.110) | n=336<br>med:0.102<br>(0.090-0.110) | n=181<br>med:0.104<br>(0.092-0.114) | n=275<br>med:0.078<br>(0.063-0.089) | n=207<br>med:0.073<br>(0.057-0.089) | n=294<br>med:0.068<br>(0.054-0.078) |
| Pig | n=333<br>med:0.144<br>(0.128-0.161) | n=297<br>med:0.141<br>(0.129-0.154) | n=275<br>med:0.106<br>(0.088-0.112) | n=116<br>med:0.087<br>(0.076-0.103) | n=340<br>med:0.097<br>(0.087-0.108) | n=185<br>med:0.106<br>(0.086-0.119) | n=278<br>med:0.073<br>(0.061-0.083) | n=205<br>med:0.071<br>(0.059-0.081) | n=296<br>med:0.066<br>(0.056-0.071) |
| Tree Shrew | n=69<br>med:0.175<br>(0.126-0.199) | n=61<br>med:0.150<br>(0.116-0.219) | n=56<br>med:0.126<br>(0.094-0.172) | n=23<br>med:0.097<br>(0.055-0.117) | n=54<br>med:0.122<br>(0.094-0.165) | n=38<br>med:0.100<br>(0.062-0.133) | n=59<br>med:0.089<br>(0.072-0.142) | n=37<br>med:0.089<br>(0.069-0.122) | n=49<br>med:0.096<br>(0.078-0.130) |
| Pika | n=116<br>med:0.139<br>(0.115-0.174) | n=120<br>med:0.131<br>(0.104-0.161) | n=107<br>med:0.103<br>(0.091-0.135) | n=42<br>med:0.065<br>(0.046-0.109) | n=110<br>med:0.087<br>(0.074-0.111) | n=63<br>med:0.120<br>(0.082-0.140) | n=80<br>med:0.076<br>(0.060-0.105) | n=56<br>med:0.091<br>(0.069-0.120) | n=80<br>med:0.069<br>(0.053-0.092) |
| Shrew | n=53<br>med:0.135<br>(0.101-0.173) | n=67<br>med:0.110<br>(0.080-0.156) | n=41<br>med:0.122<br>(0.060-0.163) | n=16<br>med:0.121<br>(0.028-0.153) | n=60<br>med:0.121<br>(0.090-0.165) | n=21<br>med:0.098<br>(0.080-0.135) | n=43<br>med:0.083<br>(0.055-0.121) | n=34<br>med:0.101<br>(0.046-0.148) | n=49<br>med:0.094<br>(0.058-0.136) |
| Squirrel | n=327<br>med:0.147<br>(0.134-0.160) | n=288<br>med:0.125<br>(0.109-0.143) | n=271<br>med:0.105<br>(0.090-0.119) | n=119<br>med:0.085<br>(0.072-0.100) | n=338<br>med:0.099<br>(0.088-0.111) | n=182<br>med:0.094<br>(0.077-0.111) | n=278<br>med:0.065<br>(0.055-0.073) | n=211<br>med:0.064<br>(0.053-0.080) | n=300<br>med:0.059<br>(0.050-0.069) |
| Gorilla | n=333<br>med:0.153<br>(0.141-0.174) | n=296<br>med:0.142<br>(0.124-0.157) | n=271<br>med:0.109<br>(0.095-0.134) | n=116<br>med:0.099<br>(0.078-0.120) | n=334<br>med:0.103<br>(0.093-0.117) | n=179<br>med:0.100<br>(0.086-0.123) | n=267<br>med:0.080<br>(0.068-0.094) | n=206<br>med:0.069<br>(0.056-0.082) | n=299<br>med:0.070<br>(0.058-0.080) |
| Cat | n=346<br>med:0.139<br>(0.129-0.164) | n=305<br>med:0.133<br>(0.117-0.150) | n=268<br>med:0.116<br>(0.096-0.132) | n=118<br>med:0.090<br>(0.079-0.106) | n=339<br>med:0.094<br>(0.088-0.110) | n=183<br>med:0.099<br>(0.080-0.111) | n=276<br>med:0.071<br>(0.063-0.081) | n=210<br>med:0.075<br>(0.061-0.089) | n=304<br>med:0.065<br>(0.058-0.072) |

#### Mouse Reference Genome, Saunders et al., 2018, three cell types

| Sample Size and Median dN/dS (CI95%) | Vasculature-specific Genes | Glia-specific Genes | Neuron-specific Genes |
|---|---|---|---|
| Microbat | n=314 med:0.103 (0.096-0.116) | n=210 med:0.088 (0.072-0.099) | n=503 med:0.059 (0.052-0.062) |
| Hedgehog | n=85 med:0.137 (0.106-0.160) | n=57 med:0.095 (0.066-0.136) | n=117 med:0.064 (0.050-0.080) |
| Naked mole-rat male | n=330 med:0.101 (0.090-0.111) | n=225 med:0.089 (0.076-0.101) | n=548 med:0.048 (0.043-0.054) |
| Sooty mangabey | n=369 med:0.111 (0.097-0.123) | n=263 med:0.084 (0.075-0.092) | n=612 med:0.051 (0.047-0.055) |
| Olive baboon | n=369 med:0.110 (0.093-0.123) | n=260 med:0.085 (0.075-0.097) | n=616 med:0.051 (0.046-0.055) |
| American bison | n=319 med:0.102 (0.089-0.123) | n=210 med:0.085 (0.078-0.094) | n=482 med:0.049 (0.044-0.054) |
| Leopard | n=369 med:0.093 (0.081-0.104) | n=255 med:0.075 (0.064-0.083) | n=591 med:0.046 (0.040-0.051) |
| Bolivian squirrel monkey | n=356 med:0.107 (0.098-0.122) | n=252 med:0.083 (0.074-0.091) | n=594 med:0.050 (0.046-0.055) |
| Crab-eating macaque | n=371 med:0.105 (0.096-0.114) | n=254 med:0.084 (0.075-0.096) | n=608 med:0.050 (0.045-0.055) |
| Human | n=386 med:0.100 (0.087-0.114) | n=275 med:0.077 (0.065-0.089) | n=634 med:0.046 (0.041-0.049) |
| Red fox | n=348 med:0.105 (0.099-0.117) | n=233 med:0.088 (0.080-0.101) | n=576 med:0.059 (0.049-0.066) |
| Shrew mouse | n=412 med:0.099 (0.087-0.117) | n=296 med:0.079 (0.065-0.092) | n=651 med:0.042 (0.037-0.047) |
| Arctic ground squirrel | n=372 med:0.097 (0.089-0.112) | n=265 med:0.070 (0.064-0.082) | n=612 med:0.042 (0.037-0.046) |
| Chinese hamster CriGri | n=368 med:0.116 (0.105-0.130) | n=262 med:0.089 (0.074-0.102) | n=579 med:0.057 (0.051-0.064) |
| Opossum | n=339 | n=236 | n=534 |

| | | | |
|---|---|---|---|
| | med:0.080<br>(0.068-0.088) | med:0.072<br>(0.062-0.081) | med:0.050<br>(0.047-0.056) |
| Dolphin | n=216<br>med:0.111<br>(0.101-0.126) | n=128<br>med:0.098<br>(0.084-0.113) | n=305<br>med:0.055<br>(0.046-0.064) |
| Capuchin | n=354<br>med:0.110<br>(0.096-0.123) | n=261<br>med:0.082<br>(0.071-0.089) | n=602<br>med:0.049<br>(0.043-0.054) |
| Mouse Lemur | n=360<br>med:0.100<br>(0.086-0.111) | n=259<br>med:0.075<br>(0.069-0.085) | n=595<br>med:0.047<br>(0.043-0.053) |
| Damara mole rat | n=339<br>med:0.100<br>(0.084-0.111) | n=234<br>med:0.075<br>(0.069-0.085) | n=585<br>med:0.044<br>(0.040-0.050) |
| Marmoset | n=356<br>med:0.111<br>(0.100-0.123) | n=253<br>med:0.083<br>(0.074-0.093) | n=607<br>med:0.050<br>(0.045-0.057) |
| Goat | n=364<br>med:0.103<br>(0.093-0.117) | n=251<br>med:0.080<br>(0.070-0.090) | n=591<br>med:0.048<br>(0.043-0.053) |
| Common wombat | n=323<br>med:0.083<br>(0.074-0.091) | n=218<br>med:0.069<br>(0.063-0.075) | n=520<br>med:0.046<br>(0.041-0.053) |
| American black bear | n=328<br>med:0.105<br>(0.094-0.121) | n=232<br>med:0.080<br>(0.070-0.089) | n=494<br>med:0.065<br>(0.053-0.071) |
| Sloth | n=48<br>med:0.170<br>(0.099-0.213) | n=34<br>med:0.160<br>(0.080-0.247) | n=70<br>med:0.080<br>(0.050-0.096) |
| Polar bear | n=330<br>med:0.111<br>(0.100-0.130) | n=226<br>med:0.095<br>(0.085-0.107) | n=528<br>med:0.065<br>(0.061-0.073) |
| Alpine marmot | n=357<br>med:0.098<br>(0.087-0.110) | n=242<br>med:0.081<br>(0.069-0.091) | n=564<br>med:0.044<br>(0.040-0.052) |
| Prairie vole | n=380<br>med:0.098<br>(0.081-0.108) | n=286<br>med:0.079<br>(0.071-0.091) | n=632<br>med:0.041<br>(0.037-0.048) |
| Donkey | n=372<br>med:0.104<br>(0.094-0.113) | n=265<br>med:0.089<br>(0.077-0.095) | n=591<br>med:0.055<br>(0.051-0.063) |
| Golden Hamster | n=370<br>med:0.114<br>(0.096-0.131) | n=269<br>med:0.093<br>(0.084-0.113) | n=593<br>med:0.052<br>(0.045-0.058) |
| Armadillo | n=339<br>med:0.105<br>(0.095-0.120) | n=228<br>med:0.080<br>(0.071-0.094) | n=535<br>med:0.058<br>(0.053-0.064) |
| Ugandan red Colobus | n=361 | n=258 | n=595 |

| | | | |
|---|---|---|---|
| | med:0.107 (0.095-0.117) | med:0.080 (0.071-0.094) | med:0.048 (0.044-0.055) |
| Pig-tailed macaque | n=371 med:0.110 (0.097-0.121) | n=266 med:0.084 (0.075-0.095) | n=616 med:0.050 (0.045-0.055) |
| Panda | n=362 med:0.103 (0.091-0.113) | n=250 med:0.078 (0.069-0.088) | n=568 med:0.046 (0.042-0.049) |
| Ferret | n=354 med:0.099 (0.090-0.108) | n=240 med:0.075 (0.068-0.085) | n=580 med:0.045 (0.040-0.051) |
| Koala | n=344 med:0.084 (0.074-0.094) | n=247 med:0.076 (0.067-0.083) | n=568 med:0.048 (0.043-0.054) |
| Wallaby | n=101 med:0.067 (0.057-0.083) | n=90 med:0.075 (0.060-0.086) | n=139 med:0.048 (0.041-0.060) |
| Tarsier | n=306 med:0.113 (0.100-0.121) | n=224 med:0.088 (0.079-0.099) | n=484 med:0.049 (0.044-0.057) |
| Degu | n=309 med:0.099 (0.086-0.109) | n=220 med:0.077 (0.067-0.088) | n=569 med:0.043 (0.039-0.048) |
| Chinese hamster CHOK1GS | n=408 med:0.110 (0.097-0.122) | n=287 med:0.075 (0.063-0.088) | n=648 med:0.045 (0.039-0.051) |
| Drill | n=353 med:0.112 (0.100-0.122) | n=246 med:0.085 (0.075-0.098) | n=574 med:0.052 (0.047-0.058) |
| Gibbon | n=348 med:0.114 (0.107-0.126) | n=247 med:0.088 (0.080-0.099) | n=596 med:0.056 (0.051-0.062) |
| Rat | n=391 med:0.111 (0.092-0.130) | n=285 med:0.090 (0.076-0.103) | n=632 med:0.050 (0.043-0.058) |
| Macaque | n=374 med:0.112 (0.097-0.126) | n=270 med:0.084 (0.073-0.092) | n=622 med:0.049 (0.043-0.053) |
| American mink | n=363 med:0.095 (0.087-0.107) | n=248 med:0.079 (0.068-0.088) | n=566 med:0.045 (0.040-0.052) |
| Gelada | n=367 med:0.108 (0.093-0.118) | n=267 med:0.080 (0.070-0.089) | n=622 med:0.048 (0.041-0.052) |
| Sheep | n=350 med:0.108 (0.101-0.122) | n=240 med:0.079 (0.073-0.089) | n=572 med:0.056 (0.049-0.061) |
| Tasmanian devil | n=316 | n=227 | n=514 |

| | | | |
|---|---|---|---|
| | med:0.074 (0.068-0.088) | med:0.071 (0.057-0.078) | med:0.044 (0.040-0.052) |
| Elephant | n=360 med:0.105 (0.095-0.117) | n=259 med:0.086 (0.073-0.089) | n=579 med:0.055 (0.048-0.060) |
| Algerian mouse | n=408 med:0.099 (0.084-0.124) | n=293 med:0.073 (0.052-0.091) | n=645 med:0.037 (0.027-0.047) |
| Dingo | n=350 med:0.100 (0.088-0.109) | n=251 med:0.078 (0.068-0.087) | n=586 med:0.052 (0.047-0.060) |
| Golden snub-nosed monkey | n=354 med:0.109 (0.097-0.119) | n=243 med:0.090 (0.078-0.106) | n=568 med:0.050 (0.046-0.055) |
| Horse | n=369 med:0.106 (0.094-0.113) | n=253 med:0.084 (0.076-0.096) | n=614 med:0.053 (0.048-0.059) |
| Black snub-nosed monkey | n=358 med:0.114 (0.102-0.125) | n=253 med:0.088 (0.077-0.098) | n=577 med:0.057 (0.052-0.061) |
| Brazilian guinea pig | n=257 med:0.117 (0.096-0.131) | n=192 med:0.090 (0.084-0.103) | n=447 med:0.071 (0.064-0.079) |
| Orangutan | n=364 med:0.104 (0.089-0.119) | n=255 med:0.087 (0.078-0.097) | n=596 med:0.048 (0.044-0.052) |
| Northern American deer mouse | n=399 med:0.107 (0.092-0.118) | n=286 med:0.074 (0.065-0.088) | n=628 med:0.040 (0.034-0.046) |
| Naked mole-rat female | n=356 med:0.102 (0.093-0.111) | n=243 med:0.078 (0.068-0.091) | n=589 med:0.046 (0.042-0.050) |
| Daurian ground squirrel | n=314 med:0.108 (0.095-0.123) | n=207 med:0.093 (0.077-0.104) | n=496 med:0.049 (0.041-0.056) |
| Tiger | n=305 med:0.103 (0.093-0.125) | n=198 med:0.086 (0.075-0.106) | n=478 med:0.053 (0.045-0.061) |
| Lesser hedgehog tenrec | n=84 med:0.127 (0.100-0.149) | n=75 med:0.088 (0.071-0.108) | n=104 med:0.065 (0.047-0.084) |
| Mongolian gerbil | n=369 med:0.111 (0.097-0.127) | n=253 med:0.068 (0.062-0.084) | n=541 med:0.050 (0.043-0.056) |
| Ma's night monkey | n=348 med:0.116 (0.101-0.132) | n=256 med:0.084 (0.074-0.095) | n=606 med:0.053 (0.047-0.059) |

| | | | |
|---|---|---|---|
| Upper Galilee mountains blind mole rat | n=366 med:0.105 (0.088-0.119) | n=270 med:0.075 (0.066-0.082) | n=622 med:0.044 (0.039-0.049) |
| Kangaroo rat | n=323 med:0.089 (0.080-0.106) | n=240 med:0.077 (0.067-0.090) | n=562 med:0.047 (0.041-0.052) |
| Chimpanzee | n=351 med:0.108 (0.094-0.121) | n=256 med:0.083 (0.076-0.094) | n=595 med:0.050 (0.046-0.054) |
| Guinea Pig | n=359 med:0.098 (0.092-0.108) | n=253 med:0.085 (0.076-0.095) | n=598 med:0.047 (0.043-0.052) |
| American beaver | n=363 med:0.107 (0.093-0.118) | n=250 med:0.084 (0.076-0.097) | n=502 med:0.054 (0.048-0.060) |
| Wild yak | n=327 med:0.103 (0.093-0.119) | n=214 med:0.083 (0.074-0.091) | n=515 med:0.048 (0.042-0.055) |
| Coquerel's sifaka | n=359 med:0.106 (0.093-0.115) | n=250 med:0.079 (0.068-0.091) | n=595 med:0.049 (0.045-0.055) |
| Alpaca | n=64 med:0.172 (0.137-0.219) | n=44 med:0.110 (0.069-0.161) | n=107 med:0.066 (0.051-0.079) |
| Lesser Egyptian jerboa | n=331 med:0.095 (0.080-0.105) | n=248 med:0.086 (0.077-0.093) | n=576 med:0.042 (0.039-0.047) |
| Long-tailed chinchilla | n=354 med:0.097 (0.089-0.105) | n=245 med:0.078 (0.064-0.086) | n=596 med:0.044 (0.040-0.049) |
| Angola colobus | n=358 med:0.108 (0.098-0.121) | n=257 med:0.084 (0.074-0.096) | n=590 med:0.049 (0.044-0.053) |
| Ryukyu mouse | n=413 med:0.104 (0.091-0.126) | n=293 med:0.083 (0.060-0.097) | n=647 med:0.035 (0.029-0.040) |
| Chinese hamster PICR | n=401 med:0.122 (0.110-0.135) | n=281 med:0.080 (0.070-0.094) | n=640 med:0.046 (0.040-0.052) |
| Vervet-AGM | n=377 med:0.101 (0.090-0.118) | n=272 med:0.077 (0.066-0.086) | n=616 med:0.045 (0.041-0.051) |
| Cow | n=359 med:0.103 (0.089-0.111) | n=254 med:0.078 (0.066-0.084) | n=591 med:0.047 (0.042-0.050) |
| Bushbaby | n=363 med:0.105 (0.093-0.115) | n=269 med:0.079 (0.071-0.091) | n=612 med:0.049 (0.045-0.053) |

| | | | |
|---|---|---|---|
| Bonobo | n=358<br>med:0.107<br>(0.095-0.125) | n=253<br>med:0.088<br>(0.078-0.098) | n=587<br>med:0.051<br>(0.047-0.057) |
| Hyrax | n=107<br>med:0.104<br>(0.087-0.132) | n=77<br>med:0.086<br>(0.062-0.114) | n=136<br>med:0.058<br>(0.046-0.071) |
| Rabbit | n=322<br>med:0.098<br>(0.085-0.107) | n=223<br>med:0.082<br>(0.068-0.093) | n=546<br>med:0.056<br>(0.050-0.063) |
| Megabat | n=191<br>med:0.123<br>(0.099-0.133) | n=136<br>med:0.089<br>(0.072-0.102) | n=281<br>med:0.057<br>(0.051-0.064) |
| Dog | n=358<br>med:0.107<br>(0.098-0.119) | n=251<br>med:0.087<br>(0.074-0.100) | n=591<br>med:0.053<br>(0.047-0.060) |
| Steppe mouse | n=386<br>med:0.137<br>(0.109-0.158) | n=286<br>med:0.108<br>(0.083-0.131) | n=629<br>med:0.057<br>(0.043-0.074) |
| Greater bamboo lemur | n=373<br>med:0.105<br>(0.092-0.113) | n=263<br>med:0.088<br>(0.075-0.100) | n=595<br>med:0.047<br>(0.044-0.054) |
| Pig | n=369<br>med:0.105<br>(0.093-0.112) | n=266<br>med:0.084<br>(0.072-0.096) | n=598<br>med:0.050<br>(0.045-0.056) |
| Tree Shrew | n=81<br>med:0.135<br>(0.104-0.155) | n=41<br>med:0.090<br>(0.071-0.128) | n=81<br>med:0.080<br>(0.058-0.093) |
| Pika | n=134<br>med:0.101<br>(0.086-0.115) | n=94<br>med:0.081<br>(0.063-0.097) | n=153<br>med:0.053<br>(0.044-0.067) |
| Shrew | n=54<br>med:0.125<br>(0.109-0.178) | n=38<br>med:0.081<br>(0.046-0.105) | n=89<br>med:0.061<br>(0.037-0.080) |
| Squirrel | n=360<br>med:0.096<br>(0.081-0.109) | n=259<br>med:0.073<br>(0.065-0.087) | n=610<br>med:0.043<br>(0.039-0.047) |
| Gorilla | n=357<br>med:0.110<br>(0.095-0.124) | n=259<br>med:0.085<br>(0.076-0.097) | n=601<br>med:0.049<br>(0.044-0.054) |
| Cat | n=375<br>med:0.100<br>(0.092-0.116) | n=256<br>med:0.079<br>(0.071-0.092) | n=607<br>med:0.051<br>(0.046-0.059) |

#### Mouse Reference Genome, Saunders et al., 2018, cell subtypes

| Sample Size and Median dN/dS (CI95%) | Endothelia-specific Genes | Microglia-specific Genes | Astrocyte-specific Genes | Oligodendrocyte-specific Genes | Neuron-specific Genes |
|---|---|---|---|---|---|
| Microbat | n=277 med:0.104 (0.097-0.116) | n=203 med:0.123 (0.112-0.134) | n=197 med:0.090 (0.070-0.104) | n=189 med:0.080 (0.067-0.094) | n=459 med:0.059 (0.052-0.064) |
| Hedgehog | n=77 med:0.134 (0.106-0.163) | n=47 med:0.121 (0.089-0.134) | n=46 med:0.083 (0.050-0.115) | n=54 med:0.078 (0.063-0.107) | n=109 med:0.064 (0.050-0.081) |
| Naked mole-rat male | n=292 med:0.101 (0.091-0.110) | n=212 med:0.121 (0.106-0.138) | n=218 med:0.082 (0.073-0.094) | n=201 med:0.080 (0.069-0.099) | n=507 med:0.050 (0.046-0.055) |
| Sooty mangabey | n=322 med:0.108 (0.091-0.118) | n=237 med:0.121 (0.104-0.140) | n=257 med:0.083 (0.069-0.092) | n=235 med:0.078 (0.065-0.089) | n=564 med:0.051 (0.047-0.056) |
| Olive baboon | n=322 med:0.109 (0.090-0.120) | n=239 med:0.126 (0.107-0.141) | n=252 med:0.082 (0.073-0.093) | n=232 med:0.077 (0.066-0.093) | n=568 med:0.052 (0.047-0.057) |
| American bison | n=272 med:0.101 (0.088-0.122) | n=203 med:0.107 (0.096-0.130) | n=192 med:0.083 (0.072-0.098) | n=193 med:0.085 (0.078-0.094) | n=441 med:0.050 (0.044-0.057) |
| Leopard | n=318 med:0.091 (0.080-0.100) | n=221 med:0.114 (0.101-0.132) | n=248 med:0.072 (0.062-0.082) | n=226 med:0.069 (0.058-0.080) | n=546 med:0.047 (0.042-0.053) |
| Bolivian squirrel monkey | n=312 med:0.106 (0.096-0.121) | n=214 med:0.125 (0.107-0.135) | n=233 med:0.081 (0.068-0.092) | n=235 med:0.075 (0.066-0.084) | n=547 med:0.051 (0.047-0.057) |
| Crab-eating macaque | n=326 med:0.101 (0.091-0.112) | n=238 med:0.120 (0.103-0.133) | n=248 med:0.082 (0.070-0.093) | n=230 med:0.081 (0.067-0.094) | n=561 med:0.052 (0.047-0.058) |
| Human | n=341 med:0.102 (0.090-0.113) | n=244 med:0.121 (0.110-0.132) | n=259 med:0.075 (0.065-0.086) | n=244 med:0.074 (0.063-0.086) | n=584 med:0.047 (0.043-0.052) |
| Red fox | n=301 med:0.104 (0.097-0.115) | n=223 med:0.127 (0.107-0.142) | n=214 med:0.098 (0.082-0.107) | n=222 med:0.084 (0.070-0.099) | n=530 med:0.061 (0.053-0.069) |
| Shrew mouse | n=363 med:0.106 (0.091-0.120) | n=265 med:0.137 (0.117-0.157) | n=273 med:0.084 (0.066-0.095) | n=261 med:0.073 (0.060-0.087) | n=602 med:0.043 (0.038-0.050) |
| Arctic ground squirrel | n=328 med:0.099 (0.092-0.112) | n=232 med:0.110 (0.092-0.129) | n=244 med:0.075 (0.063-0.087) | n=236 med:0.070 (0.060-0.079) | n=564 med:0.043 (0.038-0.047) |
| Chinese hamster CriGri | n=327 med:0.118 (0.106-0.130) | n=233 med:0.149 (0.133-0.171) | n=241 med:0.096 (0.077-0.110) | n=238 med:0.088 (0.069-0.102) | n=534 med:0.062 (0.053-0.070) |
| Opossum | n=302 med:0.080 (0.072-0.091) | n=210 med:0.090 (0.076-0.103) | n=228 med:0.073 (0.061-0.082) | n=217 med:0.067 (0.053-0.075) | n=492 med:0.050 (0.046-0.056) |
| Dolphin | n=192 med:0.114 (0.103-0.135) | n=127 med:0.123 (0.099-0.139) | n=115 med:0.102 (0.082-0.116) | n=119 med:0.096 (0.074-0.114) | n=283 med:0.055 (0.046-0.064) |
| Capuchin | n=319 med:0.108 (0.095-0.121) | n=224 med:0.121 (0.102-0.137) | n=244 med:0.079 (0.069-0.090) | n=237 med:0.076 (0.066-0.086) | n=553 med:0.051 (0.044-0.055) |
| Mouse Lemur | n=318 med:0.102 | n=226 med:0.113 | n=235 med:0.078 | n=231 med:0.073 | n=549 med:0.049 |

| | | | | | |
|---|---|---|---|---|---|
| | (0.087-0.111) | (0.095-0.127) | (0.068-0.090) | (0.060-0.081) | (0.044-0.055) |
| Damara mole rat | n=294 med:0.100 (0.089-0.112) | n=209 med:0.115 (0.093-0.135) | n=226 med:0.081 (0.072-0.093) | n=218 med:0.068 (0.058-0.077) | n=539 med:0.045 (0.040-0.052) |
| Marmoset | n=313 med:0.109 (0.100-0.122) | n=223 med:0.123 (0.110-0.136) | n=239 med:0.085 (0.068-0.099) | n=224 med:0.077 (0.065-0.083) | n=561 med:0.052 (0.046-0.058) |
| Goat | n=318 med:0.104 (0.093-0.117) | n=233 med:0.116 (0.099-0.131) | n=234 med:0.078 (0.071-0.089) | n=231 med:0.078 (0.069-0.088) | n=544 med:0.049 (0.044-0.054) |
| Common wombat | n=284 med:0.087 (0.078-0.096) | n=213 med:0.089 (0.079-0.105) | n=211 med:0.072 (0.064-0.083) | n=200 med:0.069 (0.058-0.076) | n=476 med:0.048 (0.042-0.053) |
| American black bear | n=290 med:0.105 (0.098-0.119) | n=200 med:0.130 (0.111-0.144) | n=201 med:0.086 (0.078-0.098) | n=213 med:0.076 (0.067-0.088) | n=454 med:0.066 (0.055-0.071) |
| Sloth | n=41 med:0.162 (0.112-0.213) | n=34 med:0.129 (0.075-0.166) | n=23 med:0.150 (0.087-0.241) | n=29 med:0.141 (0.072-0.248) | n=63 med:0.072 (0.051-0.093) |
| Polar bear | n=282 med:0.114 (0.101-0.131) | n=202 med:0.137 (0.114-0.154) | n=210 med:0.099 (0.089-0.109) | n=213 med:0.087 (0.075-0.098) | n=480 med:0.069 (0.063-0.075) |
| Alpine marmot | n=306 med:0.099 (0.090-0.111) | n=225 med:0.106 (0.093-0.128) | n=222 med:0.079 (0.067-0.088) | n=219 med:0.073 (0.064-0.089) | n=521 med:0.046 (0.040-0.052) |
| Prairie vole | n=339 med:0.097 (0.079-0.108) | n=247 med:0.128 (0.113-0.158) | n=266 med:0.076 (0.065-0.088) | n=253 med:0.075 (0.062-0.081) | n=584 med:0.044 (0.037-0.050) |
| Donkey | n=324 med:0.106 (0.095-0.118) | n=233 med:0.123 (0.113-0.134) | n=244 med:0.087 (0.077-0.100) | n=237 med:0.081 (0.069-0.093) | n=547 med:0.055 (0.052-0.064) |
| Golden Hamster | n=325 med:0.115 (0.102-0.130) | n=236 med:0.142 (0.123-0.166) | n=252 med:0.100 (0.078-0.118) | n=237 med:0.083 (0.071-0.092) | n=555 med:0.053 (0.045-0.059) |
| Armadillo | n=288 med:0.103 (0.095-0.117) | n=218 med:0.119 (0.107-0.134) | n=220 med:0.084 (0.071-0.096) | n=208 med:0.080 (0.068-0.091) | n=493 med:0.060 (0.054-0.066) |
| Ugandan red Colobus | n=322 med:0.107 (0.094-0.121) | n=233 med:0.121 (0.102-0.136) | n=238 med:0.079 (0.067-0.091) | n=232 med:0.077 (0.068-0.094) | n=547 med:0.049 (0.045-0.056) |
| Pig-tailed macaque | n=322 med:0.109 (0.094-0.119) | n=235 med:0.126 (0.106-0.141) | n=252 med:0.085 (0.069-0.095) | n=238 med:0.082 (0.067-0.094) | n=569 med:0.051 (0.046-0.057) |
| Panda | n=315 med:0.103 (0.092-0.113) | n=228 med:0.119 (0.100-0.141) | n=234 med:0.077 (0.068-0.093) | n=236 med:0.072 (0.060-0.084) | n=523 med:0.046 (0.042-0.050) |
| Ferret | n=309 med:0.099 (0.090-0.109) | n=227 med:0.119 (0.099-0.134) | n=231 med:0.080 (0.067-0.090) | n=223 med:0.072 (0.063-0.079) | n=531 med:0.046 (0.040-0.054) |
| Koala | n=308 med:0.083 (0.073-0.093) | n=224 med:0.091 (0.079-0.101) | n=228 med:0.072 (0.065-0.084) | n=225 med:0.074 (0.066-0.081) | n=526 med:0.049 (0.044-0.055) |
| Wallaby | n=79 med:0.076 (0.057-0.085) | n=68 med:0.087 (0.071-0.102) | n=63 med:0.067 (0.055-0.086) | n=77 med:0.075 (0.056-0.091) | n=125 med:0.048 (0.041-0.060) |
| Tarsier | n=266 med:0.113 (0.100-0.122) | n=191 med:0.126 (0.112-0.142) | n=198 med:0.088 (0.075-0.103) | n=202 med:0.084 (0.071-0.091) | n=448 med:0.052 (0.046-0.059) |

| | | | | | |
|---|---|---|---|---|---|
| Degu | n=272<br>med:0.095<br>(0.084-0.107) | n=194<br>med:0.119<br>(0.103-0.131) | n=220<br>med:0.077<br>(0.067-0.084) | n=203<br>med:0.067<br>(0.055-0.080) | n=527<br>med:0.044<br>(0.040-0.048) |
| Chinese hamster CHOK1GS | n=361<br>med:0.110<br>(0.096-0.122) | n=255<br>med:0.139<br>(0.116-0.155) | n=269<br>med:0.077<br>(0.068-0.095) | n=257<br>med:0.071<br>(0.063-0.083) | n=600<br>med:0.046<br>(0.039-0.052) |
| Drill | n=307<br>med:0.112<br>(0.099-0.121) | n=230<br>med:0.125<br>(0.113-0.138) | n=247<br>med:0.083<br>(0.072-0.093) | n=228<br>med:0.078<br>(0.064-0.088) | n=529<br>med:0.053<br>(0.047-0.060) |
| Gibbon | n=302<br>med:0.113<br>(0.105-0.126) | n=225<br>med:0.134<br>(0.118-0.155) | n=237<br>med:0.091<br>(0.082-0.103) | n=223<br>med:0.082<br>(0.071-0.098) | n=548<br>med:0.059<br>(0.053-0.065) |
| Rat | n=348<br>med:0.110<br>(0.092-0.132) | n=252<br>med:0.140<br>(0.117-0.168) | n=261<br>med:0.084<br>(0.073-0.098) | n=253<br>med:0.078<br>(0.067-0.096) | n=583<br>med:0.051<br>(0.044-0.059) |
| Macaque | n=333<br>med:0.106<br>(0.093-0.119) | n=246<br>med:0.125<br>(0.105-0.141) | n=258<br>med:0.087<br>(0.073-0.100) | n=242<br>med:0.076<br>(0.068-0.087) | n=574<br>med:0.050<br>(0.045-0.055) |
| American mink | n=313<br>med:0.100<br>(0.087-0.109) | n=214<br>med:0.121<br>(0.101-0.133) | n=215<br>med:0.079<br>(0.062-0.090) | n=225<br>med:0.070<br>(0.062-0.081) | n=522<br>med:0.046<br>(0.041-0.054) |
| Gelada | n=325<br>med:0.108<br>(0.093-0.118) | n=238<br>med:0.119<br>(0.102-0.136) | n=256<br>med:0.081<br>(0.069-0.093) | n=240<br>med:0.076<br>(0.063-0.087) | n=572<br>med:0.049<br>(0.043-0.054) |
| Sheep | n=300<br>med:0.112<br>(0.101-0.124) | n=226<br>med:0.120<br>(0.105-0.140) | n=228<br>med:0.083<br>(0.073-0.096) | n=221<br>med:0.081<br>(0.071-0.095) | n=522<br>med:0.057<br>(0.050-0.061) |
| Tasmanian devil | n=276<br>med:0.077<br>(0.069-0.089) | n=202<br>med:0.093<br>(0.082-0.103) | n=211<br>med:0.071<br>(0.059-0.081) | n=205<br>med:0.064<br>(0.052-0.073) | n=478<br>med:0.045<br>(0.041-0.053) |
| Elephant | n=316<br>med:0.105<br>(0.093-0.121) | n=225<br>med:0.121<br>(0.110-0.133) | n=237<br>med:0.086<br>(0.077-0.097) | n=238<br>med:0.078<br>(0.065-0.089) | n=533<br>med:0.056<br>(0.049-0.061) |
| Algerian mouse | n=359<br>med:0.108<br>(0.087-0.129) | n=265<br>med:0.134<br>(0.096-0.167) | n=276<br>med:0.071<br>(0.057-0.084) | n=255<br>med:0.073<br>(0.050-0.091) | n=594<br>med:0.035<br>(0.026-0.045) |
| Dingo | n=309<br>med:0.099<br>(0.085-0.106) | n=226<br>med:0.122<br>(0.101-0.132) | n=233<br>med:0.080<br>(0.071-0.090) | n=224<br>med:0.068<br>(0.060-0.078) | n=542<br>med:0.054<br>(0.048-0.061) |
| Golden snub-nosed monkey | n=312<br>med:0.104<br>(0.091-0.113) | n=221<br>med:0.125<br>(0.106-0.140) | n=238<br>med:0.088<br>(0.077-0.104) | n=223<br>med:0.078<br>(0.069-0.098) | n=528<br>med:0.052<br>(0.048-0.058) |
| Horse | n=324<br>med:0.105<br>(0.093-0.116) | n=227<br>med:0.118<br>(0.107-0.132) | n=241<br>med:0.087<br>(0.074-0.100) | n=224<br>med:0.082<br>(0.073-0.099) | n=566<br>med:0.056<br>(0.051-0.062) |
| Black snub-nosed monkey | n=318<br>med:0.110<br>(0.100-0.124) | n=214<br>med:0.121<br>(0.104-0.141) | n=232<br>med:0.086<br>(0.080-0.107) | n=230<br>med:0.077<br>(0.064-0.088) | n=536<br>med:0.057<br>(0.052-0.063) |
| Brazilian guinea pig | n=214<br>med:0.118<br>(0.097-0.131) | n=164<br>med:0.123<br>(0.112-0.143) | n=179<br>med:0.104<br>(0.089-0.113) | n=188<br>med:0.090<br>(0.084-0.104) | n=420<br>med:0.072<br>(0.064-0.081) |
| Orangutan | n=325<br>med:0.104<br>(0.090-0.117) | n=237<br>med:0.127<br>(0.111-0.139) | n=242<br>med:0.085<br>(0.076-0.098) | n=233<br>med:0.077<br>(0.061-0.084) | n=548<br>med:0.049<br>(0.045-0.054) |
| Northern American deer mouse | n=346<br>med:0.105<br>(0.091-0.118) | n=249<br>med:0.127<br>(0.112-0.147) | n=262<br>med:0.080<br>(0.062-0.090) | n=250<br>med:0.068<br>(0.057-0.077) | n=577<br>med:0.041<br>(0.035-0.048) |

| | | | | | |
|---|---|---|---|---|---|
| Naked mole-rat female | n=315 med:0.102 (0.092-0.108) | n=217 med:0.115 (0.096-0.132) | n=232 med:0.079 (0.072-0.091) | n=217 med:0.075 (0.063-0.089) | n=546 med:0.047 (0.043-0.051) |
| Daurian ground squirrel | n=265 med:0.113 (0.101-0.128) | n=199 med:0.123 (0.108-0.131) | n=184 med:0.083 (0.069-0.095) | n=186 med:0.085 (0.073-0.095) | n=455 med:0.050 (0.042-0.056) |
| Tiger | n=253 med:0.107 (0.095-0.126) | n=181 med:0.125 (0.111-0.154) | n=201 med:0.091 (0.075-0.106) | n=175 med:0.081 (0.070-0.090) | n=440 med:0.055 (0.047-0.063) |
| Lesser hedgehog tenrec | n=80 med:0.127 (0.100-0.149) | n=48 med:0.110 (0.081-0.161) | n=60 med:0.107 (0.080-0.122) | n=57 med:0.085 (0.065-0.104) | n=96 med:0.065 (0.047-0.083) |
| Mongolian gerbil | n=327 med:0.109 (0.096-0.125) | n=239 med:0.130 (0.112-0.155) | n=236 med:0.082 (0.067-0.102) | n=227 med:0.069 (0.063-0.085) | n=497 med:0.049 (0.042-0.057) |
| Ma's night monkey | n=312 med:0.111 (0.100-0.130) | n=221 med:0.126 (0.100-0.139) | n=239 med:0.081 (0.071-0.089) | n=226 med:0.079 (0.064-0.093) | n=559 med:0.055 (0.048-0.060) |
| Upper Galilee mountains blind mole rat | n=322 med:0.105 (0.088-0.121) | n=237 med:0.128 (0.113-0.141) | n=263 med:0.077 (0.065-0.087) | n=245 med:0.074 (0.067-0.084) | n=575 med:0.046 (0.040-0.052) |
| Kangaroo rat | n=280 med:0.091 (0.083-0.106) | n=202 med:0.113 (0.093-0.122) | n=226 med:0.084 (0.070-0.100) | n=214 med:0.073 (0.059-0.085) | n=520 med:0.049 (0.042-0.054) |
| Chimpanzee | n=301 med:0.108 (0.094-0.118) | n=224 med:0.121 (0.109-0.139) | n=233 med:0.082 (0.074-0.095) | n=229 med:0.082 (0.068-0.093) | n=550 med:0.050 (0.046-0.055) |
| Guinea Pig | n=309 med:0.098 (0.091-0.108) | n=223 med:0.116 (0.100-0.125) | n=238 med:0.084 (0.071-0.096) | n=233 med:0.075 (0.063-0.085) | n=553 med:0.048 (0.043-0.055) |
| American beaver | n=317 med:0.108 (0.095-0.120) | n=223 med:0.122 (0.109-0.143) | n=234 med:0.087 (0.076-0.100) | n=222 med:0.082 (0.069-0.094) | n=464 med:0.055 (0.050-0.062) |
| Wild yak | n=276 med:0.106 (0.093-0.119) | n=208 med:0.114 (0.100-0.140) | n=195 med:0.084 (0.076-0.094) | n=201 med:0.082 (0.068-0.092) | n=475 med:0.049 (0.043-0.057) |
| Coquerel's sifaka | n=316 med:0.106 (0.093-0.115) | n=222 med:0.120 (0.106-0.138) | n=235 med:0.081 (0.072-0.094) | n=235 med:0.073 (0.066-0.083) | n=546 med:0.049 (0.045-0.057) |
| Alpaca | n=61 med:0.166 (0.114-0.219) | n=37 med:0.195 (0.130-0.260) | n=52 med:0.101 (0.085-0.142) | n=44 med:0.107 (0.069-0.122) | n=98 med:0.067 (0.052-0.080) |
| Lesser Egyptian jerboa | n=294 med:0.100 (0.084-0.107) | n=198 med:0.122 (0.101-0.131) | n=237 med:0.080 (0.072-0.091) | n=226 med:0.075 (0.062-0.085) | n=534 med:0.043 (0.039-0.048) |
| Long-tailed chinchilla | n=304 med:0.098 (0.090-0.106) | n=227 med:0.115 (0.097-0.133) | n=241 med:0.076 (0.062-0.085) | n=225 med:0.069 (0.058-0.081) | n=551 med:0.047 (0.042-0.051) |
| Angola colobus | n=312 med:0.109 (0.098-0.120) | n=231 med:0.121 (0.106-0.143) | n=240 med:0.082 (0.070-0.094) | n=234 med:0.077 (0.064-0.091) | n=545 med:0.050 (0.045-0.055) |
| Ryukyu mouse | n=360 med:0.105 (0.091-0.130) | n=259 med:0.144 (0.115-0.162) | n=270 med:0.069 (0.057-0.079) | n=257 med:0.072 (0.057-0.088) | n=598 med:0.034 (0.029-0.041) |
| Chinese hamster PICR | n=356 med:0.120 (0.107-0.132) | n=255 med:0.149 (0.132-0.171) | n=263 med:0.083 (0.072-0.096) | n=253 med:0.080 (0.069-0.095) | n=589 med:0.046 (0.041-0.053) |
| Vervet-AGM | n=335 | n=235 | n=252 | n=241 | n=567 |

| | | | | | |
|---|---|---|---|---|---|
| | med:0.101<br>(0.089-0.117) | med:0.125<br>(0.107-0.139) | med:0.077<br>(0.068-0.089) | med:0.076<br>(0.064-0.084) | med:0.047<br>(0.042-0.052) |
| Cow | n=311<br>med:0.105<br>(0.091-0.114) | n=238<br>med:0.123<br>(0.100-0.141) | n=223<br>med:0.078<br>(0.067-0.086) | n=233<br>med:0.077<br>(0.063-0.087) | n=549<br>med:0.047<br>(0.043-0.052) |
| Bushbaby | n=326<br>med:0.106<br>(0.094-0.120) | n=234<br>med:0.123<br>(0.109-0.135) | n=244<br>med:0.084<br>(0.071-0.096) | n=243<br>med:0.078<br>(0.068-0.087) | n=566<br>med:0.050<br>(0.046-0.055) |
| Bonobo | n=311<br>med:0.112<br>(0.099-0.130) | n=232<br>med:0.124<br>(0.112-0.140) | n=236<br>med:0.085<br>(0.073-0.097) | n=229<br>med:0.078<br>(0.066-0.091) | n=543<br>med:0.052<br>(0.047-0.057) |
| Hyrax | n=94<br>med:0.107<br>(0.087-0.132) | n=63<br>med:0.150<br>(0.088-0.197) | n=60<br>med:0.089<br>(0.065-0.110) | n=75<br>med:0.076<br>(0.062-0.114) | n=127<br>med:0.058<br>(0.044-0.071) |
| Rabbit | n=274<br>med:0.098<br>(0.085-0.108) | n=201<br>med:0.116<br>(0.104-0.133) | n=209<br>med:0.077<br>(0.066-0.086) | n=206<br>med:0.071<br>(0.060-0.087) | n=505<br>med:0.058<br>(0.053-0.064) |
| Megabat | n=181<br>med:0.113<br>(0.096-0.130) | n=134<br>med:0.138<br>(0.115-0.156) | n=125<br>med:0.102<br>(0.081-0.116) | n=114<br>med:0.081<br>(0.065-0.096) | n=264<br>med:0.058<br>(0.052-0.065) |
| Dog | n=319<br>med:0.105<br>(0.096-0.114) | n=227<br>med:0.113<br>(0.097-0.127) | n=231<br>med:0.087<br>(0.073-0.100) | n=225<br>med:0.075<br>(0.066-0.094) | n=546<br>med:0.054<br>(0.048-0.062) |
| Steppe mouse | n=344<br>med:0.138<br>(0.111-0.169) | n=263<br>med:0.161<br>(0.130-0.188) | n=263<br>med:0.084<br>(0.063-0.101) | n=254<br>med:0.108<br>(0.081-0.132) | n=576<br>med:0.060<br>(0.042-0.078) |
| Greater bamboo lemur | n=323<br>med:0.105<br>(0.093-0.115) | n=230<br>med:0.114<br>(0.100-0.131) | n=246<br>med:0.083<br>(0.071-0.099) | n=234<br>med:0.079<br>(0.069-0.089) | n=548<br>med:0.051<br>(0.045-0.057) |
| Pig | n=324<br>med:0.102<br>(0.091-0.109) | n=227<br>med:0.123<br>(0.105-0.143) | n=239<br>med:0.082<br>(0.073-0.098) | n=239<br>med:0.074<br>(0.067-0.087) | n=552<br>med:0.052<br>(0.046-0.057) |
| Tree Shrew | n=65<br>med:0.137<br>(0.104-0.165) | n=40<br>med:0.116<br>(0.098-0.217) | n=53<br>med:0.081<br>(0.055-0.119) | n=29<br>med:0.093<br>(0.062-0.169) | n=76<br>med:0.077<br>(0.058-0.088) |
| Pika | n=112<br>med:0.102<br>(0.080-0.119) | n=90<br>med:0.106<br>(0.087-0.122) | n=81<br>med:0.083<br>(0.067-0.104) | n=78<br>med:0.072<br>(0.057-0.097) | n=139<br>med:0.056<br>(0.041-0.070) |
| Shrew | n=48<br>med:0.134<br>(0.109-0.216) | n=40<br>med:0.092<br>(0.067-0.108) | n=29<br>med:0.088<br>(0.047-0.110) | n=37<br>med:0.084<br>(0.065-0.098) | n=82<br>med:0.062<br>(0.038-0.080) |
| Squirrel | n=312<br>med:0.098<br>(0.085-0.110) | n=225<br>med:0.102<br>(0.086-0.120) | n=246<br>med:0.076<br>(0.068-0.087) | n=233<br>med:0.069<br>(0.062-0.078) | n=560<br>med:0.045<br>(0.039-0.049) |
| Gorilla | n=312<br>med:0.109<br>(0.095-0.125) | n=229<br>med:0.124<br>(0.110-0.143) | n=241<br>med:0.080<br>(0.070-0.093) | n=235<br>med:0.082<br>(0.067-0.095) | n=557<br>med:0.050<br>(0.046-0.056) |
| Cat | n=329<br>med:0.101<br>(0.093-0.116) | n=240<br>med:0.119<br>(0.105-0.134) | n=245<br>med:0.080<br>(0.071-0.099) | n=234<br>med:0.072<br>(0.062-0.081) | n=559<br>med:0.052<br>(0.047-0.060) |

#### Mouse Reference Genome, Zeisel et al., 2018, three cell types

| Sample Size and Median dN/dS (CI95%) | CNS_Glia-specific Genes | CNS_Endothelia-specific Genes | CNS_Neuron-specific Genes |
|---|---|---|---|
| Microbat | n=681<br>med:0.128<br>(0.120-0.138) | n=119<br>med:0.133<br>(0.110-0.147) | n=2,785<br>med:0.080<br>(0.076-0.084) |
| Hedgehog | n=155<br>med:0.128<br>(0.111-0.141) | n=35<br>med:0.130<br>(0.094-0.157) | n=616<br>med:0.091<br>(0.084-0.100) |
| Naked mole-rat male | n=725<br>med:0.129<br>(0.121-0.137) | n=136<br>med:0.122<br>(0.108-0.143) | n=2,757<br>med:0.075<br>(0.073-0.079) |
| Sooty mangabey | n=830<br>med:0.131<br>(0.120-0.140) | n=155<br>med:0.120<br>(0.098-0.142) | n=3,192<br>med:0.076<br>(0.073-0.079) |
| Olive baboon | n=823<br>med:0.130<br>(0.120-0.139) | n=156<br>med:0.122<br>(0.102-0.142) | n=3,186<br>med:0.074<br>(0.071-0.077) |
| American bison | n=680<br>med:0.117<br>(0.108-0.135) | n=124<br>med:0.122<br>(0.107-0.138) | n=2,583<br>med:0.073<br>(0.069-0.076) |
| Leopard | n=805<br>med:0.115<br>(0.107-0.125) | n=149<br>med:0.125<br>(0.103-0.130) | n=3,140<br>med:0.067<br>(0.065-0.070) |
| Bolivian squirrel monkey | n=791<br>med:0.133<br>(0.121-0.140) | n=142<br>med:0.138<br>(0.115-0.159) | n=3,064<br>med:0.074<br>(0.072-0.077) |
| Crab-eating macaque | n=833<br>med:0.128<br>(0.119-0.135) | n=154<br>med:0.122<br>(0.098-0.144) | n=3,153<br>med:0.073<br>(0.071-0.076) |
| Human | n=836<br>med:0.123<br>(0.117-0.134) | n=160<br>med:0.120<br>(0.102-0.143) | n=3,287<br>med:0.069<br>(0.066-0.072) |
| Red fox | n=775<br>med:0.125<br>(0.115-0.133) | n=139<br>med:0.122<br>(0.112-0.145) | n=3,010<br>med:0.079<br>(0.077-0.082) |
| Shrew mouse | n=902<br>med:0.139<br>(0.129-0.155) | n=163<br>med:0.122<br>(0.101-0.152) | n=3,396<br>med:0.074<br>(0.070-0.078) |
| Arctic ground squirrel | n=814<br>med:0.120<br>(0.111-0.129) | n=156<br>med:0.116<br>(0.100-0.136) | n=3,187<br>med:0.067<br>(0.064-0.071) |
| Chinese hamster CriGri | n=823<br>med:0.143<br>(0.134-0.156) | n=147<br>med:0.141<br>(0.122-0.162) | n=3,105<br>med:0.084<br>(0.080-0.088) |
| Opossum | n=734 | n=138 | n=2,877 |

| | | | |
|---|---|---|---|
| | med:0.094 (0.086-0.102) | med:0.082 (0.068-0.103) | med:0.067 (0.065-0.069) |
| Dolphin | n=446 med:0.128 (0.116-0.144) | n=92 med:0.124 (0.100-0.151) | n=1,721 med:0.082 (0.077-0.086) |
| Capuchin | n=815 med:0.130 (0.121-0.139) | n=150 med:0.131 (0.108-0.152) | n=3,116 med:0.075 (0.072-0.078) |
| Mouse Lemur | n=784 med:0.126 (0.115-0.135) | n=149 med:0.121 (0.099-0.133) | n=3,079 med:0.072 (0.069-0.074) |
| Damara mole rat | n=736 med:0.120 (0.113-0.130) | n=130 med:0.119 (0.105-0.134) | n=2,969 med:0.071 (0.068-0.074) |
| Marmoset | n=810 med:0.131 (0.122-0.138) | n=151 med:0.120 (0.101-0.151) | n=3,142 med:0.078 (0.075-0.080) |
| Goat | n=788 med:0.115 (0.107-0.123) | n=137 med:0.120 (0.103-0.133) | n=3,101 med:0.070 (0.068-0.073) |
| Common wombat | n=699 med:0.097 (0.090-0.101) | n=130 med:0.091 (0.081-0.105) | n=2,799 med:0.064 (0.062-0.067) |
| American black bear | n=710 med:0.125 (0.115-0.132) | n=134 med:0.124 (0.105-0.144) | n=2,750 med:0.082 (0.079-0.085) |
| Sloth | n=113 med:0.167 (0.139-0.194) | n=20 med:0.191 (0.120-0.264) | n=435 med:0.105 (0.093-0.118) |
| Polar bear | n=702 med:0.135 (0.128-0.143) | n=128 med:0.151 (0.121-0.174) | n=2,775 med:0.086 (0.083-0.090) |
| Alpine marmot | n=765 med:0.125 (0.115-0.136) | n=146 med:0.131 (0.103-0.155) | n=3,049 med:0.071 (0.068-0.074) |
| Prairie vole | n=849 med:0.130 (0.120-0.139) | n=160 med:0.120 (0.101-0.137) | n=3,263 med:0.070 (0.067-0.073) |
| Donkey | n=801 med:0.125 (0.118-0.131) | n=142 med:0.121 (0.104-0.139) | n=3,189 med:0.078 (0.075-0.081) |
| Golden Hamster | n=819 med:0.144 (0.134-0.154) | n=154 med:0.143 (0.119-0.158) | n=3,096 med:0.079 (0.075-0.084) |
| Armadillo | n=729 med:0.127 (0.120-0.134) | n=129 med:0.123 (0.106-0.137) | n=2,863 med:0.078 (0.075-0.082) |
| Ugandan red Colobus | n=808 | n=150 | n=3,099 |

| | | | |
|---|---|---|---|
| | med:0.130<br>(0.121-0.140) | med:0.127<br>(0.107-0.149) | med:0.074<br>(0.072-0.077) |
| Pig-tailed macaque | n=828<br>med:0.130<br>(0.122-0.140) | n=158<br>med:0.123<br>(0.110-0.142) | n=3,184<br>med:0.075<br>(0.072-0.078) |
| Panda | n=781<br>med:0.120<br>(0.110-0.128) | n=144<br>med:0.123<br>(0.097-0.145) | n=3,122<br>med:0.071<br>(0.069-0.075) |
| Ferret | n=779<br>med:0.122<br>(0.112-0.131) | n=138<br>med:0.110<br>(0.098-0.129) | n=3,093<br>med:0.068<br>(0.066-0.071) |
| Koala | n=761<br>med:0.096<br>(0.090-0.104) | n=146<br>med:0.088<br>(0.075-0.104) | n=3,060<br>med:0.068<br>(0.065-0.070) |
| Wallaby | n=219<br>med:0.081<br>(0.075-0.094) | n=56<br>med:0.086<br>(0.069-0.116) | n=870<br>med:0.073<br>(0.067-0.077) |
| Tarsier | n=687<br>med:0.133<br>(0.121-0.145) | n=123<br>med:0.132<br>(0.115-0.153) | n=2,595<br>med:0.078<br>(0.074-0.081) |
| Degu | n=693<br>med:0.120<br>(0.112-0.126) | n=133<br>med:0.103<br>(0.089-0.120) | n=2,776<br>med:0.071<br>(0.068-0.073) |
| Chinese hamster CHOK1GS | n=885<br>med:0.139<br>(0.130-0.149) | n=166<br>med:0.136<br>(0.111-0.154) | n=3,376<br>med:0.076<br>(0.073-0.080) |
| Drill | n=804<br>med:0.130<br>(0.121-0.141) | n=145<br>med:0.128<br>(0.112-0.144) | n=3,079<br>med:0.077<br>(0.073-0.080) |
| Gibbon | n=796<br>med:0.138<br>(0.130-0.151) | n=146<br>med:0.130<br>(0.110-0.150) | n=3,055<br>med:0.083<br>(0.079-0.086) |
| Rat | n=872<br>med:0.140<br>(0.131-0.151) | n=155<br>med:0.132<br>(0.113-0.164) | n=3,324<br>med:0.076<br>(0.073-0.080) |
| Macaque | n=847<br>med:0.128<br>(0.118-0.137) | n=158<br>med:0.121<br>(0.104-0.145) | n=3,241<br>med:0.076<br>(0.072-0.078) |
| American mink | n=767<br>med:0.118<br>(0.110-0.128) | n=140<br>med:0.111<br>(0.098-0.124) | n=3,016<br>med:0.069<br>(0.066-0.072) |
| Gelada | n=824<br>med:0.129<br>(0.118-0.135) | n=155<br>med:0.121<br>(0.100-0.141) | n=3,194<br>med:0.070<br>(0.068-0.073) |
| Sheep | n=775<br>med:0.123<br>(0.115-0.133) | n=134<br>med:0.122<br>(0.111-0.139) | n=3,038<br>med:0.080<br>(0.077-0.082) |
| Tasmanian devil | n=698 | n=131 | n=2,739 |

| | | | |
|---|---|---|---|
| | med:0.094 (0.087-0.101) | med:0.086 (0.068-0.097) | med:0.063 (0.061-0.065) |
| Elephant | n=765 med:0.121 (0.112-0.130) | n=139 med:0.126 (0.105-0.139) | n=3,076 med:0.077 (0.074-0.081) |
| Algerian mouse | n=905 med:0.151 (0.137-0.167) | n=164 med:0.113 (0.088-0.153) | n=3,423 med:0.074 (0.069-0.081) |
| Dingo | n=764 med:0.118 (0.110-0.128) | n=147 med:0.121 (0.100-0.136) | n=3,089 med:0.073 (0.069-0.076) |
| Golden snub-nosed monkey | n=803 med:0.132 (0.124-0.142) | n=157 med:0.119 (0.104-0.143) | n=3,022 med:0.075 (0.071-0.078) |
| Horse | n=805 med:0.129 (0.120-0.138) | n=145 med:0.124 (0.107-0.143) | n=3,204 med:0.076 (0.073-0.080) |
| Black snub-nosed monkey | n=796 med:0.133 (0.124-0.144) | n=149 med:0.133 (0.114-0.151) | n=3,064 med:0.080 (0.077-0.083) |
| Brazilian guinea pig | n=581 med:0.127 (0.120-0.135) | n=101 med:0.132 (0.114-0.148) | n=2,287 med:0.096 (0.092-0.099) |
| Orangutan | n=783 med:0.125 (0.116-0.132) | n=147 med:0.130 (0.105-0.153) | n=3,118 med:0.073 (0.070-0.076) |
| Northern American deer mouse | n=876 med:0.130 (0.121-0.140) | n=159 med:0.126 (0.103-0.159) | n=3,228 med:0.067 (0.064-0.071) |
| Naked mole-rat female | n=769 med:0.127 (0.118-0.137) | n=152 med:0.117 (0.102-0.136) | n=2,941 med:0.073 (0.070-0.077) |
| Daurian ground squirrel | n=687 med:0.131 (0.119-0.143) | n=126 med:0.140 (0.118-0.163) | n=2,681 med:0.076 (0.073-0.079) |
| Tiger | n=694 med:0.130 (0.123-0.138) | n=117 med:0.135 (0.125-0.169) | n=2,555 med:0.079 (0.075-0.082) |
| Lesser hedgehog tenrec | n=180 med:0.124 (0.114-0.144) | n=39 med:0.126 (0.104-0.189) | n=688 med:0.084 (0.078-0.092) |
| Mongolian gerbil | n=798 med:0.140 (0.127-0.147) | n=155 med:0.122 (0.104-0.154) | n=2,988 med:0.080 (0.076-0.084) |
| Ma's night monkey | n=804 med:0.134 (0.124-0.143) | n=144 med:0.124 (0.104-0.150) | n=3,078 med:0.078 (0.075-0.082) |

| | | | |
|---|---|---|---|
| Upper Galilee mountains blind mole rat | n=799<br>med:0.126<br>(0.117-0.135) | n=153<br>med:0.125<br>(0.104-0.138) | n=3,164<br>med:0.070<br>(0.066-0.073) |
| Kangaroo rat | n=696<br>med:0.115<br>(0.106-0.123) | n=130<br>med:0.111<br>(0.089-0.132) | n=2,884<br>med:0.070<br>(0.067-0.073) |
| Chimpanzee | n=801<br>med:0.130<br>(0.122-0.138) | n=147<br>med:0.113<br>(0.096-0.139) | n=3,041<br>med:0.074<br>(0.071-0.077) |
| Guinea Pig | n=761<br>med:0.118<br>(0.109-0.126) | n=146<br>med:0.108<br>(0.093-0.133) | n=3,125<br>med:0.073<br>(0.070-0.076) |
| American beaver | n=750<br>med:0.127<br>(0.119-0.138) | n=154<br>med:0.127<br>(0.108-0.143) | n=2,900<br>med:0.078<br>(0.075-0.082) |
| Wild yak | n=727<br>med:0.119<br>(0.112-0.131) | n=129<br>med:0.123<br>(0.108-0.145) | n=2,722<br>med:0.074<br>(0.071-0.077) |
| Coquerel's sifaka | n=788<br>med:0.126<br>(0.116-0.136) | n=143<br>med:0.123<br>(0.104-0.136) | n=3,079<br>med:0.076<br>(0.072-0.079) |
| Alpaca | n=173<br>med:0.179<br>(0.151-0.201) | n=28<br>med:0.148<br>(0.096-0.232) | n=686<br>med:0.089<br>(0.082-0.098) |
| Lesser Egyptian jerboa | n=739<br>med:0.121<br>(0.113-0.129) | n=141<br>med:0.115<br>(0.099-0.136) | n=2,909<br>med:0.072<br>(0.069-0.076) |
| Long-tailed chinchilla | n=788<br>med:0.117<br>(0.110-0.127) | n=143<br>med:0.105<br>(0.093-0.118) | n=3,130<br>med:0.068<br>(0.065-0.070) |
| Angola colobus | n=812<br>med:0.136<br>(0.125-0.144) | n=147<br>med:0.140<br>(0.114-0.150) | n=3,102<br>med:0.075<br>(0.072-0.079) |
| Ryukyu mouse | n=890<br>med:0.142<br>(0.129-0.154) | n=168<br>med:0.128<br>(0.099-0.146) | n=3,399<br>med:0.072<br>(0.068-0.076) |
| Chinese hamster PICR | n=871<br>med:0.144<br>(0.135-0.155) | n=165<br>med:0.141<br>(0.122-0.160) | n=3,339<br>med:0.080<br>(0.076-0.084) |
| Vervet-AGM | n=824<br>med:0.124<br>(0.117-0.134) | n=159<br>med:0.123<br>(0.107-0.136) | n=3,216<br>med:0.070<br>(0.066-0.073) |
| Cow | n=799<br>med:0.113<br>(0.106-0.122) | n=143<br>med:0.114<br>(0.107-0.132) | n=3,118<br>med:0.072<br>(0.068-0.074) |
| Bushbaby | n=812<br>med:0.129<br>(0.119-0.140) | n=145<br>med:0.133<br>(0.115-0.145) | n=3,157<br>med:0.076<br>(0.072-0.079) |

| | | | |
|---|---|---|---|
| Bonobo | n=811<br>med:0.132<br>(0.124-0.141) | n=150<br>med:0.132<br>(0.110-0.147) | n=3,068<br>med:0.077<br>(0.074-0.080) |
| Hyrax | n=219<br>med:0.130<br>(0.115-0.155) | n=57<br>med:0.125<br>(0.090-0.150) | n=827<br>med:0.084<br>(0.078-0.093) |
| Rabbit | n=712<br>med:0.116<br>(0.106-0.123) | n=126<br>med:0.121<br>(0.106-0.134) | n=2,827<br>med:0.076<br>(0.072-0.080) |
| Megabat | n=442<br>med:0.136<br>(0.122-0.147) | n=85<br>med:0.142<br>(0.110-0.162) | n=1,575<br>med:0.085<br>(0.080-0.088) |
| Dog | n=798<br>med:0.124<br>(0.115-0.134) | n=149<br>med:0.130<br>(0.110-0.143) | n=3,072<br>med:0.077<br>(0.074-0.080) |
| Steppe mouse | n=879<br>med:0.176<br>(0.165-0.196) | n=157<br>med:0.108<br>(0.079-0.170) | n=3,325<br>med:0.102<br>(0.095-0.111) |
| Greater bamboo lemur | n=820<br>med:0.124<br>(0.114-0.132) | n=149<br>med:0.125<br>(0.107-0.142) | n=3,149<br>med:0.075<br>(0.072-0.078) |
| Pig | n=796<br>med:0.120<br>(0.110-0.129) | n=149<br>med:0.115<br>(0.106-0.131) | n=3,169<br>med:0.074<br>(0.071-0.076) |
| Tree Shrew | n=159<br>med:0.148<br>(0.119-0.160) | n=31<br>med:0.122<br>(0.087-0.172) | n=564<br>med:0.089<br>(0.085-0.099) |
| Pika | n=285<br>med:0.113<br>(0.096-0.120) | n=65<br>med:0.132<br>(0.098-0.148) | n=990<br>med:0.076<br>(0.072-0.081) |
| Shrew | n=124<br>med:0.117<br>(0.097-0.136) | n=25<br>med:0.126<br>(0.086-0.221) | n=570<br>med:0.074<br>(0.068-0.082) |
| Squirrel | n=786<br>med:0.119<br>(0.112-0.131) | n=151<br>med:0.116<br>(0.097-0.139) | n=3,173<br>med:0.067<br>(0.064-0.070) |
| Gorilla | n=805<br>med:0.132<br>(0.122-0.140) | n=147<br>med:0.132<br>(0.108-0.155) | n=3,104<br>med:0.075<br>(0.072-0.078) |
| Cat | n=829<br>med:0.124<br>(0.116-0.132) | n=155<br>med:0.124<br>(0.109-0.144) | n=3,218<br>med:0.073<br>(0.071-0.076) |

#### Mouse Reference Genome, Zeisel et al., 2018, cell subtypes

| Sample Size and Median dN/dS (CI95%) | Glia-specific Genes | Immune cells-specific Genes | Vascular cells-specific Genes | Neurons-specific Genes |
|---|---|---|---|---|
| Microbat | n=572 med:0.147 (0.132-0.157) | n=74 med:0.138 (0.105-0.169) | n=144 med:0.111 (0.095-0.137) | n=2,907 med:0.090 (0.086-0.093) |
| Hedgehog | n=129 med:0.134 (0.116-0.163) | n=26 med:0.132 (0.089-0.168) | n=29 med:0.147 (0.117-0.179) | n=677 med:0.103 (0.094-0.112) |
| Naked mole-rat male | n=607 med:0.149 (0.136-0.160) | n=78 med:0.150 (0.106-0.178) | n=156 med:0.111 (0.105-0.129) | n=2,905 med:0.085 (0.082-0.089) |
| Sooty mangabey | n=674 med:0.153 (0.143-0.165) | n=90 med:0.156 (0.126-0.190) | n=175 med:0.119 (0.103-0.136) | n=3,365 med:0.084 (0.081-0.087) |
| Olive baboon | n=675 med:0.151 (0.140-0.166) | n=86 med:0.149 (0.119-0.181) | n=174 med:0.119 (0.100-0.141) | n=3,360 med:0.082 (0.078-0.085) |
| American bison | n=564 med:0.143 (0.134-0.153) | n=75 med:0.150 (0.125-0.181) | n=135 med:0.115 (0.096-0.131) | n=2,731 med:0.080 (0.077-0.084) |
| Leopard | n=663 med:0.135 (0.124-0.146) | n=88 med:0.133 (0.105-0.166) | n=166 med:0.106 (0.093-0.125) | n=3,301 med:0.075 (0.072-0.078) |
| Bolivian squirrel monkey | n=664 med:0.150 (0.140-0.164) | n=83 med:0.165 (0.120-0.186) | n=157 med:0.125 (0.108-0.147) | n=3,225 med:0.084 (0.080-0.087) |
| Crab-eating macaque | n=683 med:0.147 (0.135-0.158) | n=91 med:0.149 (0.120-0.189) | n=172 med:0.116 (0.099-0.139) | n=3,326 med:0.081 (0.078-0.084) |
| Human | n=695 med:0.148 (0.137-0.156) | n=93 med:0.148 (0.115-0.182) | n=184 med:0.116 (0.104-0.126) | n=3,444 med:0.079 (0.075-0.082) |
| Red fox | n=643 med:0.144 (0.133-0.156) | n=85 med:0.155 (0.120-0.189) | n=161 med:0.115 (0.100-0.136) | n=3,152 med:0.086 (0.084-0.089) |
| Shrew mouse | n=733 med:0.172 (0.156-0.183) | n=118 med:0.199 (0.155-0.257) | n=185 med:0.108 (0.095-0.138) | n=3,602 med:0.088 (0.082-0.092) |
| Arctic ground squirrel | n=666 med:0.140 (0.131-0.150) | n=88 med:0.152 (0.113-0.192) | n=177 med:0.112 (0.092-0.132) | n=3,364 med:0.077 (0.074-0.080) |
| Chinese hamster CriGri | n=671 med:0.173 (0.161-0.185) | n=106 med:0.184 (0.142-0.243) | n=165 med:0.139 (0.116-0.155) | n=3,298 med:0.096 (0.091-0.099) |
| Opossum | n=607 | n=74 | n=162 | n=3,031 |

| | | | | |
|---|---|---|---|---|
| | med:0.107<br>(0.098-0.113) | med:0.089<br>(0.067-0.113) | med:0.088<br>(0.070-0.097) | med:0.073<br>(0.070-0.075) |
| Dolphin | n=380<br>med:0.149<br>(0.136-0.162) | n=58<br>med:0.162<br>(0.135-0.229) | n=98<br>med:0.119<br>(0.100-0.144) | n=1,810<br>med:0.092<br>(0.087-0.096) |
| Capuchin | n=676<br>med:0.154<br>(0.142-0.161) | n=90<br>med:0.153<br>(0.118-0.197) | n=167<br>med:0.116<br>(0.104-0.134) | n=3,288<br>med:0.082<br>(0.079-0.086) |
| Mouse Lemur | n=651<br>med:0.144<br>(0.135-0.154) | n=91<br>med:0.148<br>(0.109-0.178) | n=165<br>med:0.115<br>(0.093-0.133) | n=3,260<br>med:0.081<br>(0.077-0.085) |
| Damara mole rat | n=608<br>med:0.139<br>(0.130-0.150) | n=76<br>med:0.141<br>(0.093-0.169) | n=151<br>med:0.117<br>(0.105-0.132) | n=3,104<br>med:0.080<br>(0.076-0.084) |
| Marmoset | n=677<br>med:0.155<br>(0.141-0.166) | n=92<br>med:0.153<br>(0.118-0.174) | n=175<br>med:0.123<br>(0.108-0.142) | n=3,316<br>med:0.087<br>(0.083-0.090) |
| Goat | n=652<br>med:0.136<br>(0.124-0.144) | n=87<br>med:0.154<br>(0.128-0.179) | n=158<br>med:0.100<br>(0.090-0.119) | n=3,260<br>med:0.079<br>(0.075-0.082) |
| Common wombat | n=578<br>med:0.111<br>(0.103-0.120) | n=70<br>med:0.103<br>(0.081-0.124) | n=149<br>med:0.090<br>(0.083-0.104) | n=2,947<br>med:0.070<br>(0.067-0.073) |
| American black bear | n=586<br>med:0.141<br>(0.130-0.151) | n=80<br>med:0.157<br>(0.122-0.181) | n=152<br>med:0.113<br>(0.092-0.135) | n=2,908<br>med:0.090<br>(0.086-0.093) |
| Sloth | n=82<br>med:0.203<br>(0.172-0.241) | n=14<br>med:0.328<br>(0.126-0.477) | n=19<br>med:0.156<br>(0.114-0.245) | n=467<br>med:0.128<br>(0.116-0.140) |
| Polar bear | n=584<br>med:0.154<br>(0.143-0.165) | n=76<br>med:0.155<br>(0.115-0.210) | n=139<br>med:0.133<br>(0.111-0.154) | n=2,908<br>med:0.095<br>(0.092-0.099) |
| Alpine marmot | n=628<br>med:0.143<br>(0.135-0.151) | n=90<br>med:0.155<br>(0.111-0.191) | n=162<br>med:0.121<br>(0.098-0.141) | n=3,225<br>med:0.081<br>(0.078-0.085) |
| Prairie vole | n=707<br>med:0.151<br>(0.141-0.166) | n=104<br>med:0.179<br>(0.133-0.234) | n=178<br>med:0.127<br>(0.109-0.144) | n=3,438<br>med:0.080<br>(0.076-0.084) |
| Donkey | n=667<br>med:0.145<br>(0.132-0.155) | n=87<br>med:0.135<br>(0.122-0.160) | n=163<br>med:0.115<br>(0.098-0.127) | n=3,365<br>med:0.086<br>(0.083-0.090) |
| Golden Hamster | n=680<br>med:0.167<br>(0.153-0.177) | n=98<br>med:0.187<br>(0.145-0.233) | n=166<br>med:0.126<br>(0.108-0.149) | n=3,273<br>med:0.091<br>(0.086-0.096) |
| Armadillo | n=609<br>med:0.139<br>(0.130-0.155) | n=87<br>med:0.131<br>(0.107-0.169) | n=149<br>med:0.111<br>(0.095-0.135) | n=3,007<br>med:0.087<br>(0.084-0.091) |

| | | | | |
|---|---|---|---|---|
| Ugandan red Colobus | n=664 med:0.154 (0.140-0.163) | n=88 med:0.145 (0.118-0.162) | n=165 med:0.115 (0.104-0.140) | n=3,268 med:0.082 (0.078-0.085) |
| Pig-tailed macaque | n=684 med:0.151 (0.137-0.162) | n=87 med:0.153 (0.124-0.199) | n=176 med:0.119 (0.102-0.138) | n=3,360 med:0.083 (0.079-0.086) |
| Panda | n=644 med:0.137 (0.128-0.148) | n=92 med:0.157 (0.124-0.180) | n=159 med:0.113 (0.091-0.127) | n=3,269 med:0.080 (0.076-0.083) |
| Ferret | n=654 med:0.139 (0.129-0.149) | n=89 med:0.156 (0.125-0.176) | n=159 med:0.103 (0.088-0.118) | n=3,243 med:0.078 (0.074-0.081) |
| Koala | n=632 med:0.114 (0.107-0.125) | n=78 med:0.100 (0.077-0.124) | n=166 med:0.086 (0.071-0.096) | n=3,204 med:0.073 (0.071-0.075) |
| Wallaby | n=155 med:0.095 (0.076-0.109) | n=19 med:0.080 (0.063-0.114) | n=65 med:0.081 (0.070-0.090) | n=948 med:0.076 (0.072-0.080) |
| Tarsier | n=581 med:0.159 (0.150-0.165) | n=78 med:0.158 (0.121-0.187) | n=131 med:0.126 (0.110-0.152) | n=2,736 med:0.090 (0.086-0.093) |
| Degu | n=570 med:0.139 (0.128-0.147) | n=70 med:0.126 (0.095-0.148) | n=158 med:0.107 (0.093-0.124) | n=2,959 med:0.080 (0.076-0.083) |
| Chinese hamster CHOK1GS | n=730 med:0.167 (0.153-0.179) | n=111 med:0.190 (0.143-0.251) | n=187 med:0.122 (0.101-0.143) | n=3,576 med:0.087 (0.083-0.091) |
| Drill | n=668 med:0.152 (0.139-0.164) | n=88 med:0.155 (0.127-0.202) | n=170 med:0.119 (0.107-0.139) | n=3,246 med:0.084 (0.081-0.088) |
| Gibbon | n=662 med:0.162 (0.151-0.170) | n=82 med:0.166 (0.132-0.209) | n=165 med:0.123 (0.110-0.140) | n=3,219 med:0.091 (0.087-0.094) |
| Rat | n=717 med:0.163 (0.151-0.177) | n=116 med:0.216 (0.147-0.273) | n=176 med:0.114 (0.093-0.137) | n=3,500 med:0.088 (0.083-0.091) |
| Macaque | n=704 med:0.148 (0.139-0.159) | n=94 med:0.147 (0.120-0.171) | n=180 med:0.120 (0.103-0.143) | n=3,412 med:0.083 (0.080-0.088) |
| American mink | n=639 med:0.136 (0.125-0.149) | n=79 med:0.153 (0.122-0.177) | n=161 med:0.104 (0.089-0.118) | n=3,166 med:0.078 (0.075-0.082) |
| Gelada | n=680 med:0.149 (0.136-0.160) | n=91 med:0.150 (0.119-0.185) | n=174 med:0.115 (0.103-0.139) | n=3,383 med:0.079 (0.075-0.081) |
| Sheep | n=646 med:0.142 (0.133-0.149) | n=82 med:0.168 (0.124-0.189) | n=155 med:0.114 (0.103-0.136) | n=3,175 med:0.088 (0.085-0.092) |

| | | | | |
|---|---|---|---|---|
| Tasmanian devil | n=581 med:0.109 (0.101-0.116) | n=74 med:0.109 (0.078-0.128) | n=148 med:0.072 (0.063-0.085) | n=2,863 med:0.069 (0.066-0.072) |
| Elephant | n=637 med:0.140 (0.128-0.148) | n=82 med:0.151 (0.113-0.194) | n=161 med:0.111 (0.101-0.145) | n=3,235 med:0.087 (0.083-0.090) |
| Algerian mouse | n=751 med:0.178 (0.164-0.197) | n=123 med:0.207 (0.153-0.260) | n=177 med:0.095 (0.078-0.136) | n=3,625 med:0.086 (0.081-0.091) |
| Dingo | n=621 med:0.136 (0.125-0.147) | n=88 med:0.145 (0.117-0.168) | n=167 med:0.109 (0.088-0.125) | n=3,255 med:0.082 (0.079-0.085) |
| Golden snub-nosed monkey | n=663 med:0.155 (0.144-0.164) | n=88 med:0.157 (0.126-0.186) | n=168 med:0.119 (0.104-0.143) | n=3,184 med:0.083 (0.079-0.086) |
| Horse | n=679 med:0.147 (0.138-0.159) | n=84 med:0.150 (0.121-0.178) | n=167 med:0.113 (0.098-0.129) | n=3,363 med:0.085 (0.082-0.088) |
| Black snub-nosed monkey | n=659 med:0.156 (0.146-0.166) | n=85 med:0.153 (0.116-0.179) | n=170 med:0.127 (0.108-0.144) | n=3,240 med:0.087 (0.084-0.092) |
| Brazilian guinea pig | n=471 med:0.147 (0.139-0.153) | n=60 med:0.128 (0.107-0.170) | n=126 med:0.134 (0.114-0.152) | n=2,383 med:0.102 (0.099-0.106) |
| Orangutan | n=642 med:0.145 (0.131-0.155) | n=91 med:0.155 (0.122-0.181) | n=161 med:0.117 (0.100-0.130) | n=3,273 med:0.080 (0.077-0.084) |
| Northern American deer mouse | n=711 med:0.153 (0.139-0.166) | n=109 med:0.188 (0.142-0.250) | n=178 med:0.120 (0.100-0.137) | n=3,426 med:0.080 (0.076-0.084) |
| Naked mole-rat female | n=644 med:0.147 (0.137-0.159) | n=81 med:0.154 (0.115-0.193) | n=178 med:0.115 (0.106-0.130) | n=3,094 med:0.082 (0.078-0.085) |
| Daurian ground squirrel | n=570 med:0.146 (0.139-0.159) | n=80 med:0.158 (0.118-0.198) | n=143 med:0.129 (0.103-0.147) | n=2,834 med:0.086 (0.083-0.090) |
| Tiger | n=590 med:0.146 (0.137-0.157) | n=77 med:0.150 (0.109-0.187) | n=129 med:0.127 (0.100-0.141) | n=2,697 med:0.087 (0.083-0.091) |
| Lesser hedgehog tenrec | n=140 med:0.134 (0.118-0.156) | n=19 med:0.172 (0.109-0.351) | n=40 med:0.114 (0.093-0.180) | n=759 med:0.094 (0.084-0.101) |
| Mongolian gerbil | n=656 med:0.165 (0.149-0.179) | n=100 med:0.184 (0.145-0.232) | n=171 med:0.122 (0.104-0.136) | n=3,173 med:0.091 (0.087-0.095) |
| Ma's night monkey | n=661 med:0.153 (0.144-0.164) | n=89 med:0.153 (0.121-0.184) | n=169 med:0.118 (0.103-0.137) | n=3,235 med:0.086 (0.083-0.091) |

| | | | | |
|---|---|---|---|---|
| Upper Galilee mountains blind mole rat | n=663 med:0.150 (0.140-0.162) | n=95 med:0.158 (0.128-0.203) | n=169 med:0.122 (0.104-0.144) | n=3,359 med:0.083 (0.078-0.086) |
| Kangaroo rat | n=567 med:0.133 (0.123-0.141) | n=74 med:0.120 (0.094-0.147) | n=152 med:0.106 (0.089-0.125) | n=3,017 med:0.079 (0.075-0.083) |
| Chimpanzee | n=654 med:0.150 (0.137-0.161) | n=85 med:0.152 (0.120-0.197) | n=167 med:0.114 (0.098-0.131) | n=3,222 med:0.082 (0.079-0.086) |
| Guinea Pig | n=635 med:0.143 (0.133-0.154) | n=77 med:0.142 (0.104-0.176) | n=168 med:0.111 (0.094-0.130) | n=3,266 med:0.082 (0.079-0.085) |
| American beaver | n=594 med:0.149 (0.137-0.157) | n=92 med:0.167 (0.123-0.200) | n=171 med:0.118 (0.104-0.133) | n=3,083 med:0.088 (0.084-0.091) |
| Wild yak | n=600 med:0.142 (0.131-0.152) | n=83 med:0.146 (0.113-0.167) | n=144 med:0.115 (0.100-0.136) | n=2,871 med:0.082 (0.079-0.086) |
| Coquerel's sifaka | n=655 med:0.148 (0.138-0.160) | n=84 med:0.153 (0.115-0.181) | n=162 med:0.112 (0.096-0.129) | n=3,233 med:0.084 (0.081-0.088) |
| Alpaca | n=148 med:0.201 (0.176-0.229) | n=13 med:0.203 (0.111-0.334) | n=28 med:0.144 (0.095-0.192) | n=718 med:0.104 (0.098-0.113) |
| Lesser Egyptian jerboa | n=607 med:0.143 (0.133-0.151) | n=74 med:0.142 (0.122-0.177) | n=157 med:0.112 (0.097-0.131) | n=3,040 med:0.082 (0.078-0.085) |
| Long-tailed chinchilla | n=653 med:0.140 (0.130-0.150) | n=83 med:0.139 (0.097-0.156) | n=160 med:0.104 (0.096-0.122) | n=3,280 med:0.076 (0.073-0.080) |
| Angola colobus | n=666 med:0.153 (0.142-0.165) | n=89 med:0.150 (0.116-0.200) | n=162 med:0.130 (0.109-0.143) | n=3,249 med:0.084 (0.081-0.088) |
| Ryukyu mouse | n=729 med:0.161 (0.148-0.182) | n=118 med:0.208 (0.158-0.256) | n=188 med:0.121 (0.092-0.139) | n=3,595 med:0.083 (0.080-0.088) |
| Chinese hamster PICR | n=724 med:0.170 (0.158-0.183) | n=110 med:0.206 (0.154-0.269) | n=182 med:0.127 (0.098-0.145) | n=3,535 med:0.092 (0.088-0.096) |
| Vervet-AGM | n=679 med:0.149 (0.135-0.160) | n=88 med:0.147 (0.119-0.163) | n=179 med:0.119 (0.103-0.132) | n=3,386 med:0.078 (0.075-0.082) |
| Cow | n=662 med:0.135 (0.123-0.142) | n=88 med:0.151 (0.140-0.197) | n=167 med:0.114 (0.091-0.127) | n=3,288 med:0.079 (0.076-0.082) |
| Bushbaby | n=661 med:0.150 (0.141-0.158) | n=90 med:0.142 (0.113-0.194) | n=160 med:0.128 (0.111-0.141) | n=3,342 med:0.085 (0.082-0.089) |

| | | | | |
|---|---|---|---|---|
| Bonobo | n=664 med:0.154 (0.143-0.163) | n=88 med:0.150 (0.118-0.182) | n=169 med:0.124 (0.106-0.140) | n=3,236 med:0.085 (0.081-0.088) |
| Hyrax | n=178 med:0.165 (0.137-0.178) | n=35 med:0.235 (0.147-0.248) | n=58 med:0.115 (0.080-0.156) | n=885 med:0.095 (0.087-0.101) |
| Rabbit | n=595 med:0.133 (0.124-0.142) | n=79 med:0.138 (0.109-0.168) | n=148 med:0.106 (0.084-0.126) | n=2,958 med:0.083 (0.081-0.087) |
| Megabat | n=349 med:0.146 (0.131-0.161) | n=53 med:0.190 (0.146-0.227) | n=92 med:0.137 (0.111-0.154) | n=1,704 med:0.094 (0.088-0.098) |
| Dog | n=660 med:0.146 (0.135-0.157) | n=88 med:0.131 (0.107-0.163) | n=167 med:0.112 (0.097-0.130) | n=3,220 med:0.085 (0.082-0.088) |
| Steppe mouse | n=729 med:0.214 (0.188-0.242) | n=124 med:0.198 (0.143-0.249) | n=176 med:0.116 (0.089-0.156) | n=3,505 med:0.113 (0.105-0.120) |
| Greater bamboo lemur | n=676 med:0.144 (0.135-0.154) | n=90 med:0.146 (0.109-0.170) | n=171 med:0.113 (0.101-0.132) | n=3,328 med:0.083 (0.080-0.087) |
| Pig | n=671 med:0.139 (0.130-0.150) | n=85 med:0.138 (0.118-0.168) | n=171 med:0.107 (0.089-0.123) | n=3,330 med:0.080 (0.077-0.083) |
| Tree Shrew | n=135 med:0.158 (0.136-0.188) | n=20 med:0.280 (0.160-0.402) | n=30 med:0.155 (0.087-0.189) | n=610 med:0.101 (0.089-0.110) |
| Pika | n=218 med:0.127 (0.113-0.136) | n=41 med:0.152 (0.104-0.191) | n=72 med:0.112 (0.086-0.142) | n=1,095 med:0.086 (0.078-0.093) |
| Shrew | n=91 med:0.142 (0.112-0.170) | n=18 med:0.200 (0.136-0.325) | n=34 med:0.128 (0.101-0.165) | n=648 med:0.087 (0.079-0.093) |
| Squirrel | n=656 med:0.141 (0.131-0.151) | n=84 med:0.135 (0.099-0.168) | n=169 med:0.111 (0.091-0.132) | n=3,353 med:0.076 (0.073-0.079) |
| Gorilla | n=665 med:0.153 (0.141-0.165) | n=85 med:0.161 (0.118-0.197) | n=168 med:0.122 (0.107-0.143) | n=3,277 med:0.084 (0.080-0.088) |
| Cat | n=688 med:0.137 (0.130-0.151) | n=99 med:0.138 (0.107-0.165) | n=175 med:0.112 (0.098-0.128) | n=3,378 med:0.081 (0.078-0.085) |

#### Rat Reference Genome, Zhang et al., 2014, three cell types

| Sample Size and Median dN/dS (CI95%) | Endothelia-specific Genes | Glia-specific Genes | Neuron-specific Genes |
|---|---|---|---|
| Microbat | n=619<br>med:0.117<br>(0.106-0.128) | n=782<br>med:0.121<br>(0.112-0.128) | n=873<br>med:0.076<br>(0.071-0.081) |
| Hedgehog | n=170<br>med:0.132<br>(0.107-0.149) | n=214<br>med:0.120<br>(0.107-0.134) | n=211<br>med:0.079<br>(0.069-0.099) |
| Naked mole-rat male | n=639<br>med:0.119<br>(0.110-0.131) | n=877<br>med:0.118<br>(0.110-0.125) | n=944<br>med:0.068<br>(0.063-0.072) |
| Sooty mangabey | n=720<br>med:0.122<br>(0.111-0.131) | n=960<br>med:0.123<br>(0.116-0.128) | n=1,080<br>med:0.067<br>(0.063-0.074) |
| Olive baboon | n=727<br>med:0.121<br>(0.111-0.128) | n=971<br>med:0.125<br>(0.119-0.130) | n=1,070<br>med:0.068<br>(0.063-0.076) |
| American bison | n=599<br>med:0.122<br>(0.105-0.131) | n=761<br>med:0.119<br>(0.111-0.127) | n=805<br>med:0.070<br>(0.064-0.076) |
| Leopard | n=704<br>med:0.109<br>(0.099-0.118) | n=918<br>med:0.107<br>(0.099-0.114) | n=1,036<br>med:0.062<br>(0.058-0.068) |
| Bolivian squirrel monkey | n=700<br>med:0.122<br>(0.112-0.131) | n=903<br>med:0.122<br>(0.116-0.131) | n=1,029<br>med:0.068<br>(0.062-0.074) |
| Crab-eating macaque | n=721<br>med:0.120<br>(0.111-0.128) | n=966<br>med:0.122<br>(0.115-0.127) | n=1,072<br>med:0.069<br>(0.064-0.076) |
| Human | n=743<br>med:0.116<br>(0.108-0.124) | n=982<br>med:0.119<br>(0.112-0.125) | n=1,100<br>med:0.065<br>(0.060-0.071) |
| Red fox | n=689<br>med:0.121<br>(0.115-0.127) | n=880<br>med:0.117<br>(0.110-0.125) | n=968<br>med:0.075<br>(0.069-0.081) |
| Shrew mouse | n=797<br>med:0.122<br>(0.112-0.132) | n=1,067<br>med:0.133<br>(0.125-0.143) | n=1,160<br>med:0.076<br>(0.071-0.083) |
| Arctic ground squirrel | n=721<br>med:0.114<br>(0.105-0.125) | n=959<br>med:0.114<br>(0.106-0.125) | n=1,061<br>med:0.062<br>(0.057-0.066) |
| Chinese hamster CriGri | n=720<br>med:0.134<br>(0.124-0.148) | n=971<br>med:0.140<br>(0.132-0.147) | n=1,032<br>med:0.076<br>(0.070-0.086) |
| Opossum | n=640 | n=827 | n=951 |

| | | | |
|---|---|---|---|
| | med:0.091<br>(0.084-0.100) | med:0.087<br>(0.079-0.092) | med:0.060<br>(0.057-0.063) |
| Dolphin | n=426<br>med:0.120<br>(0.108-0.130) | n=503<br>med:0.127<br>(0.121-0.138) | n=517<br>med:0.083<br>(0.073-0.092) |
| Capuchin | n=713<br>med:0.122<br>(0.112-0.130) | n=947<br>med:0.122<br>(0.115-0.128) | n=1,052<br>med:0.067<br>(0.061-0.072) |
| Mouse Lemur | n=704<br>med:0.116<br>(0.109-0.122) | n=926<br>med:0.115<br>(0.105-0.122) | n=1,027<br>med:0.066<br>(0.060-0.071) |
| Damara mole rat | n=650<br>med:0.116<br>(0.106-0.125) | n=850<br>med:0.112<br>(0.105-0.122) | n=959<br>med:0.063<br>(0.058-0.069) |
| Marmoset | n=714<br>med:0.124<br>(0.116-0.136) | n=945<br>med:0.122<br>(0.116-0.129) | n=1,053<br>med:0.071<br>(0.066-0.078) |
| Goat | n=707<br>med:0.114<br>(0.105-0.124) | n=904<br>med:0.114<br>(0.106-0.120) | n=1,032<br>med:0.065<br>(0.061-0.070) |
| Common wombat | n=630<br>med:0.090<br>(0.083-0.097) | n=800<br>med:0.090<br>(0.085-0.096) | n=908<br>med:0.059<br>(0.056-0.063) |
| American black bear | n=632<br>med:0.123<br>(0.112-0.131) | n=832<br>med:0.116<br>(0.109-0.123) | n=827<br>med:0.076<br>(0.070-0.081) |
| Sloth | n=97<br>med:0.159<br>(0.140-0.186) | n=127<br>med:0.184<br>(0.148-0.223) | n=115<br>med:0.090<br>(0.075-0.113) |
| Polar bear | n=636<br>med:0.128<br>(0.117-0.137) | n=805<br>med:0.125<br>(0.117-0.134) | n=839<br>med:0.083<br>(0.076-0.091) |
| Alpine marmot | n=698<br>med:0.117<br>(0.106-0.127) | n=915<br>med:0.119<br>(0.108-0.127) | n=984<br>med:0.067<br>(0.061-0.073) |
| Prairie vole | n=751<br>med:0.121<br>(0.110-0.131) | n=1,019<br>med:0.125<br>(0.118-0.133) | n=1,124<br>med:0.067<br>(0.060-0.072) |
| Donkey | n=704<br>med:0.116<br>(0.111-0.125) | n=940<br>med:0.120<br>(0.113-0.126) | n=1,050<br>med:0.072<br>(0.065-0.078) |
| Golden Hamster | n=716<br>med:0.131<br>(0.122-0.141) | n=971<br>med:0.138<br>(0.128-0.144) | n=1,071<br>med:0.072<br>(0.066-0.078) |
| Armadillo | n=646<br>med:0.119<br>(0.110-0.126) | n=861<br>med:0.114<br>(0.105-0.120) | n=922<br>med:0.072<br>(0.066-0.076) |
| Ugandan red Colobus | n=710 | n=943 | n=1,060 |

| | | | |
|---|---|---|---|
| | med:0.121 (0.111-0.130) | med:0.123 (0.118-0.129) | med:0.068 (0.063-0.074) |
| Pig-tailed macaque | n=732 med:0.123 (0.115-0.132) | n=959 med:0.123 (0.117-0.129) | n=1,084 med:0.070 (0.065-0.076) |
| Panda | n=709 med:0.119 (0.108-0.127) | n=920 med:0.115 (0.106-0.123) | n=1,015 med:0.065 (0.060-0.069) |
| Ferret | n=697 med:0.107 (0.101-0.115) | n=905 med:0.114 (0.107-0.121) | n=1,004 med:0.066 (0.060-0.070) |
| Koala | n=675 med:0.093 (0.085-0.099) | n=867 med:0.088 (0.084-0.093) | n=995 med:0.063 (0.059-0.066) |
| Wallaby | n=191 med:0.080 (0.070-0.094) | n=275 med:0.082 (0.075-0.092) | n=273 med:0.063 (0.056-0.071) |
| Tarsier | n=595 med:0.124 (0.111-0.133) | n=772 med:0.124 (0.114-0.132) | n=831 med:0.073 (0.068-0.081) |
| Degu | n=614 med:0.112 (0.106-0.123) | n=824 med:0.112 (0.103-0.120) | n=976 med:0.065 (0.059-0.071) |
| Chinese hamster CHOK1GS | n=794 med:0.130 (0.121-0.142) | n=1,063 med:0.133 (0.125-0.142) | n=1,163 med:0.070 (0.064-0.077) |
| Drill | n=697 med:0.121 (0.112-0.129) | n=927 med:0.125 (0.118-0.132) | n=1,017 med:0.070 (0.066-0.076) |
| Gibbon | n=686 med:0.122 (0.114-0.131) | n=927 med:0.129 (0.123-0.137) | n=1,020 med:0.077 (0.072-0.083) |
| Macaque | n=755 med:0.122 (0.115-0.130) | n=999 med:0.123 (0.118-0.128) | n=1,105 med:0.070 (0.064-0.075) |
| American mink | n=679 med:0.107 (0.100-0.114) | n=891 med:0.111 (0.104-0.118) | n=977 med:0.067 (0.062-0.072) |
| Gelada | n=729 med:0.116 (0.108-0.127) | n=969 med:0.122 (0.115-0.128) | n=1,095 med:0.065 (0.059-0.072) |
| Sheep | n=674 med:0.122 (0.114-0.129) | n=872 med:0.121 (0.114-0.128) | n=991 med:0.078 (0.071-0.083) |
| Tasmanian devil | n=618 med:0.088 (0.084-0.094) | n=797 med:0.086 (0.079-0.091) | n=862 med:0.056 (0.053-0.060) |
| Elephant | n=694 | n=899 | n=1,005 |

| | | | |
|---|---|---|---|
| | med:0.118 (0.110-0.126) | med:0.121 (0.113-0.129) | med:0.071 (0.066-0.078) |
| Algerian mouse | n=799 med:0.127 (0.117-0.138) | n=1,069 med:0.133 (0.125-0.142) | n=1,167 med:0.079 (0.068-0.086) |
| Dingo | n=701 med:0.112 (0.105-0.120) | n=908 med:0.112 (0.105-0.121) | n=1,007 med:0.066 (0.061-0.071) |
| Golden snub-nosed monkey | n=717 med:0.118 (0.110-0.127) | n=929 med:0.126 (0.119-0.134) | n=1,011 med:0.069 (0.063-0.075) |
| Horse | n=718 med:0.118 (0.112-0.126) | n=938 med:0.119 (0.112-0.127) | n=1,079 med:0.073 (0.067-0.079) |
| Black snub-nosed monkey | n=707 med:0.125 (0.114-0.136) | n=932 med:0.127 (0.120-0.135) | n=1,025 med:0.075 (0.069-0.082) |
| Brazilian guinea pig | n=491 med:0.124 (0.117-0.133) | n=698 med:0.121 (0.115-0.129) | n=733 med:0.085 (0.078-0.092) |
| Orangutan | n=716 med:0.117 (0.108-0.124) | n=922 med:0.121 (0.113-0.129) | n=1,024 med:0.069 (0.063-0.074) |
| Northern American deer mouse | n=765 med:0.119 (0.110-0.130) | n=1,036 med:0.124 (0.116-0.132) | n=1,081 med:0.066 (0.060-0.071) |
| Naked mole-rat female | n=689 med:0.119 (0.109-0.129) | n=925 med:0.117 (0.109-0.123) | n=1,018 med:0.067 (0.062-0.073) |
| Mouse | n=839 med:0.126 (0.114-0.137) | n=1,105 med:0.135 (0.126-0.142) | n=1,200 med:0.075 (0.068-0.083) |
| Daurian ground squirrel | n=630 med:0.129 (0.116-0.138) | n=786 med:0.126 (0.117-0.134) | n=831 med:0.075 (0.069-0.081) |
| Tiger | n=589 med:0.121 (0.114-0.129) | n=760 med:0.123 (0.115-0.133) | n=793 med:0.076 (0.069-0.085) |
| Lesser hedgehog tenrec | n=184 med:0.136 (0.117-0.154) | n=222 med:0.116 (0.106-0.129) | n=202 med:0.087 (0.068-0.105) |
| Mongolian gerbil | n=711 med:0.128 (0.113-0.143) | n=949 med:0.134 (0.126-0.145) | n=967 med:0.075 (0.069-0.084) |
| Ma's night monkey | n=702 med:0.124 (0.115-0.137) | n=936 med:0.123 (0.115-0.133) | n=1,039 med:0.072 (0.066-0.077) |

| | | | |
|---|---|---|---|
| Upper Galilee mountains blind mole rat | n=720<br>med:0.129<br>(0.115-0.136) | n=981<br>med:0.124<br>(0.115-0.134) | n=1,106<br>med:0.068<br>(0.062-0.073) |
| Kangaroo rat | n=624<br>med:0.105<br>(0.099-0.111) | n=823<br>med:0.107<br>(0.101-0.115) | n=963<br>med:0.064<br>(0.059-0.069) |
| Chimpanzee | n=705<br>med:0.121<br>(0.112-0.127) | n=934<br>med:0.123<br>(0.116-0.128) | n=1,056<br>med:0.067<br>(0.063-0.073) |
| Guinea Pig | n=707<br>med:0.115<br>(0.106-0.122) | n=913<br>med:0.118<br>(0.110-0.122) | n=1,003<br>med:0.071<br>(0.065-0.075) |
| American beaver | n=683<br>med:0.120<br>(0.113-0.132) | n=903<br>med:0.122<br>(0.114-0.131) | n=950<br>med:0.072<br>(0.068-0.079) |
| Wild yak | n=650<br>med:0.119<br>(0.107-0.129) | n=807<br>med:0.117<br>(0.111-0.125) | n=834<br>med:0.072<br>(0.066-0.077) |
| Coquerel's sifaka | n=681<br>med:0.120<br>(0.112-0.129) | n=880<br>med:0.120<br>(0.111-0.128) | n=1,010<br>med:0.066<br>(0.060-0.072) |
| Alpaca | n=152<br>med:0.149<br>(0.135-0.183) | n=156<br>med:0.148<br>(0.125-0.182) | n=174<br>med:0.086<br>(0.072-0.101) |
| Lesser Egyptian jerboa | n=648<br>med:0.115<br>(0.106-0.124) | n=877<br>med:0.118<br>(0.108-0.125) | n=988<br>med:0.062<br>(0.056-0.068) |
| Long-tailed chinchilla | n=690<br>med:0.108<br>(0.099-0.116) | n=902<br>med:0.110<br>(0.103-0.117) | n=1,025<br>med:0.062<br>(0.058-0.067) |
| Angola colobus | n=705<br>med:0.121<br>(0.113-0.131) | n=937<br>med:0.125<br>(0.118-0.133) | n=1,024<br>med:0.068<br>(0.063-0.076) |
| Ryukyu mouse | n=798<br>med:0.125<br>(0.116-0.137) | n=1,050<br>med:0.133<br>(0.121-0.142) | n=1,161<br>med:0.076<br>(0.069-0.084) |
| Chinese hamster PICR | n=788<br>med:0.137<br>(0.127-0.150) | n=1,053<br>med:0.140<br>(0.132-0.147) | n=1,139<br>med:0.072<br>(0.065-0.080) |
| Vervet-AGM | n=735<br>med:0.120<br>(0.109-0.127) | n=972<br>med:0.122<br>(0.117-0.130) | n=1,064<br>med:0.063<br>(0.058-0.069) |
| Cow | n=715<br>med:0.112<br>(0.101-0.123) | n=915<br>med:0.116<br>(0.110-0.123) | n=1,024<br>med:0.066<br>(0.061-0.071) |
| Bushbaby | n=710<br>med:0.120<br>(0.110-0.129) | n=961<br>med:0.124<br>(0.116-0.131) | n=1,072<br>med:0.067<br>(0.062-0.074) |

| | | | |
|---|---|---|---|
| Bonobo | n=708 med:0.123 (0.115-0.129) | n=951 med:0.124 (0.116-0.129) | n=1,023 med:0.071 (0.066-0.075) |
| Hyrax | n=202 med:0.106 (0.096-0.128) | n=281 med:0.129 (0.114-0.145) | n=279 med:0.087 (0.076-0.101) |
| Rabbit | n=636 med:0.110 (0.101-0.119) | n=853 med:0.111 (0.103-0.118) | n=924 med:0.073 (0.069-0.079) |
| Megabat | n=388 med:0.127 (0.115-0.135) | n=524 med:0.133 (0.123-0.143) | n=525 med:0.084 (0.075-0.093) |
| Dog | n=702 med:0.119 (0.109-0.126) | n=922 med:0.119 (0.109-0.126) | n=1,019 med:0.073 (0.069-0.079) |
| Steppe mouse | n=784 med:0.136 (0.124-0.150) | n=1,057 med:0.145 (0.134-0.153) | n=1,122 med:0.085 (0.079-0.092) |
| Greater bamboo lemur | n=721 med:0.118 (0.109-0.126) | n=943 med:0.117 (0.110-0.125) | n=1,033 med:0.066 (0.061-0.073) |
| Pig | n=722 med:0.112 (0.107-0.120) | n=932 med:0.117 (0.110-0.126) | n=1,045 med:0.066 (0.062-0.073) |
| Tree Shrew | n=165 med:0.133 (0.103-0.146) | n=200 med:0.133 (0.117-0.153) | n=173 med:0.090 (0.076-0.107) |
| Pika | n=259 med:0.100 (0.093-0.113) | n=345 med:0.122 (0.106-0.130) | n=316 med:0.076 (0.069-0.083) |
| Shrew | n=131 med:0.117 (0.106-0.136) | n=157 med:0.102 (0.082-0.118) | n=157 med:0.071 (0.055-0.086) |
| Squirrel | n=705 med:0.113 (0.104-0.126) | n=928 med:0.110 (0.101-0.118) | n=1,067 med:0.064 (0.059-0.069) |
| Gorilla | n=703 med:0.122 (0.114-0.131) | n=949 med:0.122 (0.116-0.130) | n=1,056 med:0.070 (0.064-0.075) |
| Cat | n=730 med:0.122 (0.112-0.129) | n=953 med:0.113 (0.108-0.122) | n=1,057 med:0.069 (0.063-0.076) |

#### Rat Reference Genome, Zhang et al., 2014, cell subtypes

| Sample Size and Median dN/dS (CI95%) | Microglia-specific Genes | Endothelia-specific Genes | Astrocyte-specific Genes | Oligodendrocyte-specific Genes | Neuron-specific Genes |
|---|---|---|---|---|---|
| Microbat | n=673 med:0.127 (0.120-0.133) | n=619 med:0.117 (0.106-0.128) | n=576 med:0.105 (0.096-0.118) | n=350 med:0.090 (0.082-0.097) | n=873 med:0.076 (0.071-0.081) |
| Hedgehog | n=196 med:0.144 (0.134-0.156) | n=170 med:0.132 (0.107-0.149) | n=120 med:0.113 (0.085-0.131) | n=90 med:0.086 (0.075-0.099) | n=211 med:0.079 (0.069-0.099) |
| Naked mole-rat male | n=721 med:0.128 (0.118-0.137) | n=639 med:0.119 (0.110-0.131) | n=608 med:0.107 (0.096-0.112) | n=394 med:0.090 (0.081-0.099) | n=944 med:0.068 (0.063-0.072) |
| Sooty mangabey | n=813 med:0.133 (0.126-0.145) | n=720 med:0.122 (0.111-0.131) | n=689 med:0.104 (0.095-0.114) | n=432 med:0.089 (0.080-0.097) | n=1,080 med:0.067 (0.063-0.074) |
| Olive baboon | n=818 med:0.135 (0.127-0.147) | n=727 med:0.121 (0.111-0.128) | n=684 med:0.107 (0.098-0.116) | n=437 med:0.090 (0.082-0.099) | n=1,070 med:0.068 (0.063-0.076) |
| American bison | n=665 med:0.128 (0.120-0.135) | n=599 med:0.122 (0.105-0.131) | n=549 med:0.099 (0.089-0.109) | n=352 med:0.086 (0.080-0.097) | n=805 med:0.070 (0.064-0.076) |
| Leopard | n=767 med:0.121 (0.114-0.128) | n=704 med:0.109 (0.099-0.118) | n=677 med:0.095 (0.084-0.101) | n=417 med:0.079 (0.072-0.084) | n=1,036 med:0.062 (0.058-0.068) |
| Bolivian squirrel monkey | n=779 med:0.135 (0.125-0.146) | n=700 med:0.122 (0.112-0.131) | n=661 med:0.109 (0.100-0.119) | n=417 med:0.088 (0.079-0.095) | n=1,029 med:0.068 (0.062-0.074) |
| Crab-eating macaque | n=823 med:0.131 (0.125-0.143) | n=721 med:0.120 (0.111-0.128) | n=682 med:0.106 (0.099-0.114) | n=428 med:0.086 (0.079-0.096) | n=1,072 med:0.069 (0.064-0.076) |
| Human | n=838 med:0.128 (0.121-0.138) | n=743 med:0.116 (0.108-0.124) | n=702 med:0.105 (0.096-0.113) | n=441 med:0.088 (0.080-0.095) | n=1,100 med:0.065 (0.060-0.071) |
| Red fox | n=768 med:0.132 (0.124-0.141) | n=689 med:0.121 (0.115-0.127) | n=615 med:0.104 (0.096-0.111) | n=412 med:0.086 (0.078-0.095) | n=968 med:0.075 (0.069-0.081) |
| Shrew mouse | n=913 med:0.152 (0.141-0.164) | n=797 med:0.122 (0.112-0.132) | n=735 med:0.112 (0.101-0.120) | n=482 med:0.090 (0.080-0.101) | n=1,160 med:0.076 (0.071-0.083) |
| Arctic ground squirrel | n=808 med:0.129 (0.119-0.135) | n=721 med:0.114 (0.105-0.125) | n=686 med:0.097 (0.087-0.105) | n=437 med:0.080 (0.072-0.087) | n=1,061 med:0.062 (0.057-0.066) |
| Chinese hamster CriGri | n=860 med:0.161 (0.150-0.174) | n=720 med:0.134 (0.124-0.148) | n=670 med:0.116 (0.105-0.126) | n=436 med:0.097 (0.085-0.107) | n=1,032 med:0.076 (0.070-0.086) |
| Opossum | n=696 med:0.089 (0.081-0.098) | n=640 med:0.091 (0.084-0.100) | n=607 med:0.079 (0.072-0.086) | n=395 med:0.070 (0.064-0.078) | n=951 med:0.060 (0.057-0.063) |
| Dolphin | n=459 med:0.137 (0.126-0.149) | n=426 med:0.120 (0.108-0.130) | n=353 med:0.118 (0.106-0.129) | n=214 med:0.103 (0.090-0.113) | n=517 med:0.083 (0.073-0.092) |
| Capuchin | n=797 med:0.134 (0.125-0.144) | n=713 med:0.122 (0.112-0.130) | n=669 med:0.107 (0.098-0.116) | n=433 med:0.086 (0.078-0.094) | n=1,052 med:0.067 (0.061-0.072) |
| Mouse Lemur | n=778 med:0.129 | n=704 med:0.116 | n=647 med:0.100 | n=425 med:0.081 | n=1,027 med:0.066 |

| | | | | | |
|---|---|---|---|---|---|
| | (0.120-0.135) | (0.109-0.122) | (0.092-0.109) | (0.072-0.090) | (0.060-0.071) |
| Damara mole rat | n=703 med:0.122 (0.111-0.133) | n=650 med:0.116 (0.106-0.125) | n=617 med:0.102 (0.092-0.110) | n=401 med:0.085 (0.078-0.092) | n=959 med:0.063 (0.058-0.069) |
| Marmoset | n=805 med:0.140 (0.127-0.151) | n=714 med:0.124 (0.116-0.136) | n=682 med:0.110 (0.099-0.118) | n=436 med:0.091 (0.083-0.098) | n=1,053 med:0.071 (0.066-0.078) |
| Goat | n=778 med:0.122 (0.114-0.128) | n=707 med:0.114 (0.105-0.124) | n=664 med:0.097 (0.089-0.105) | n=419 med:0.084 (0.074-0.091) | n=1,032 med:0.065 (0.061-0.070) |
| Common wombat | n=691 med:0.094 (0.085-0.099) | n=630 med:0.090 (0.083-0.097) | n=587 med:0.083 (0.075-0.089) | n=386 med:0.077 (0.069-0.083) | n=908 med:0.059 (0.056-0.063) |
| American black bear | n=728 med:0.121 (0.115-0.128) | n=632 med:0.123 (0.112-0.131) | n=554 med:0.113 (0.103-0.121) | n=357 med:0.089 (0.079-0.095) | n=827 med:0.076 (0.070-0.081) |
| Sloth | n=111 med:0.188 (0.143-0.234) | n=97 med:0.159 (0.140-0.186) | n=83 med:0.146 (0.111-0.172) | n=63 med:0.127 (0.084-0.171) | n=115 med:0.090 (0.075-0.113) |
| Polar bear | n=709 med:0.134 (0.127-0.142) | n=636 med:0.128 (0.117-0.137) | n=578 med:0.109 (0.099-0.120) | n=366 med:0.090 (0.083-0.100) | n=839 med:0.083 (0.076-0.091) |
| Alpine marmot | n=797 med:0.132 (0.122-0.143) | n=698 med:0.117 (0.106-0.127) | n=653 med:0.097 (0.086-0.107) | n=408 med:0.086 (0.079-0.094) | n=984 med:0.067 (0.061-0.073) |
| Prairie vole | n=868 med:0.140 (0.128-0.152) | n=751 med:0.121 (0.110-0.131) | n=694 med:0.109 (0.097-0.120) | n=465 med:0.086 (0.076-0.097) | n=1,124 med:0.067 (0.060-0.072) |
| Donkey | n=815 med:0.132 (0.123-0.139) | n=704 med:0.116 (0.111-0.125) | n=664 med:0.109 (0.100-0.115) | n=424 med:0.089 (0.079-0.096) | n=1,050 med:0.072 (0.065-0.078) |
| Golden Hamster | n=830 med:0.155 (0.144-0.165) | n=716 med:0.131 (0.122-0.141) | n=670 med:0.118 (0.108-0.126) | n=440 med:0.096 (0.084-0.106) | n=1,071 med:0.072 (0.066-0.078) |
| Armadillo | n=717 med:0.128 (0.118-0.136) | n=646 med:0.119 (0.110-0.126) | n=616 med:0.104 (0.094-0.113) | n=394 med:0.084 (0.078-0.092) | n=922 med:0.072 (0.066-0.076) |
| Ugandan red Colobus | n=797 med:0.134 (0.127-0.147) | n=710 med:0.121 (0.111-0.130) | n=673 med:0.107 (0.096-0.114) | n=435 med:0.089 (0.080-0.098) | n=1,060 med:0.068 (0.063-0.074) |
| Pig-tailed macaque | n=818 med:0.132 (0.123-0.143) | n=732 med:0.123 (0.115-0.132) | n=679 med:0.106 (0.097-0.116) | n=435 med:0.085 (0.079-0.097) | n=1,084 med:0.070 (0.065-0.076) |
| Panda | n=782 med:0.121 (0.115-0.133) | n=709 med:0.119 (0.108-0.127) | n=640 med:0.101 (0.091-0.107) | n=422 med:0.082 (0.073-0.089) | n=1,015 med:0.065 (0.060-0.069) |
| Ferret | n=778 med:0.123 (0.115-0.131) | n=697 med:0.107 (0.101-0.115) | n=641 med:0.098 (0.088-0.107) | n=405 med:0.077 (0.071-0.085) | n=1,004 med:0.066 (0.060-0.070) |
| Koala | n=735 med:0.092 (0.086-0.099) | n=675 med:0.093 (0.085-0.099) | n=626 med:0.086 (0.078-0.092) | n=419 med:0.076 (0.069-0.082) | n=995 med:0.063 (0.059-0.066) |
| Wallaby | n=255 med:0.084 (0.075-0.093) | n=191 med:0.080 (0.070-0.094) | n=164 med:0.084 (0.069-0.097) | n=137 med:0.077 (0.067-0.088) | n=273 med:0.063 (0.056-0.071) |
| Tarsier | n=671 med:0.138 (0.129-0.146) | n=595 med:0.124 (0.111-0.133) | n=556 med:0.107 (0.100-0.115) | n=347 med:0.087 (0.079-0.093) | n=831 med:0.073 (0.068-0.081) |

| | | | | | |
|---|---|---|---|---|---|
| Degu | n=677 med:0.120 (0.113-0.128) | n=614 med:0.112 (0.106-0.123) | n=596 med:0.095 (0.085-0.103) | n=401 med:0.083 (0.076-0.090) | n=976 med:0.065 (0.059-0.071) |
| Chinese hamster CHOK1GS | n=912 med:0.150 (0.142-0.162) | n=794 med:0.130 (0.121-0.142) | n=737 med:0.113 (0.102-0.124) | n=481 med:0.092 (0.082-0.103) | n=1,163 med:0.070 (0.064-0.077) |
| Drill | n=803 med:0.137 (0.128-0.149) | n=697 med:0.121 (0.112-0.129) | n=657 med:0.108 (0.098-0.117) | n=417 med:0.089 (0.079-0.097) | n=1,017 med:0.070 (0.066-0.076) |
| Gibbon | n=778 med:0.145 (0.135-0.152) | n=686 med:0.122 (0.114-0.131) | n=661 med:0.111 (0.100-0.123) | n=415 med:0.096 (0.086-0.107) | n=1,020 med:0.077 (0.072-0.083) |
| Macaque | n=845 med:0.134 (0.124-0.145) | n=755 med:0.122 (0.115-0.130) | n=703 med:0.107 (0.099-0.116) | n=447 med:0.088 (0.080-0.096) | n=1,105 med:0.070 (0.064-0.075) |
| American mink | n=757 med:0.120 (0.114-0.130) | n=679 med:0.107 (0.100-0.114) | n=643 med:0.097 (0.087-0.107) | n=410 med:0.078 (0.073-0.084) | n=977 med:0.067 (0.062-0.072) |
| Gelada | n=822 med:0.133 (0.124-0.144) | n=729 med:0.116 (0.108-0.127) | n=689 med:0.104 (0.096-0.111) | n=436 med:0.088 (0.078-0.096) | n=1,095 med:0.065 (0.059-0.072) |
| Sheep | n=750 med:0.129 (0.122-0.138) | n=674 med:0.122 (0.114-0.129) | n=635 med:0.105 (0.095-0.113) | n=402 med:0.086 (0.080-0.095) | n=991 med:0.078 (0.071-0.083) |
| Tasmanian devil | n=679 med:0.090 (0.085-0.097) | n=618 med:0.088 (0.084-0.094) | n=557 med:0.078 (0.070-0.086) | n=374 med:0.071 (0.064-0.079) | n=862 med:0.056 (0.053-0.060) |
| Elephant | n=767 med:0.132 (0.123-0.140) | n=694 med:0.118 (0.110-0.126) | n=645 med:0.103 (0.095-0.112) | n=420 med:0.087 (0.081-0.094) | n=1,005 med:0.071 (0.066-0.078) |
| Algerian mouse | n=919 med:0.153 (0.141-0.166) | n=799 med:0.127 (0.117-0.138) | n=741 med:0.112 (0.102-0.122) | n=483 med:0.090 (0.078-0.101) | n=1,167 med:0.079 (0.068-0.086) |
| Dingo | n=786 med:0.124 (0.115-0.131) | n=701 med:0.112 (0.105-0.120) | n=645 med:0.100 (0.090-0.109) | n=405 med:0.083 (0.077-0.089) | n=1,007 med:0.066 (0.061-0.071) |
| Golden snub-nosed monkey | n=784 med:0.136 (0.127-0.146) | n=717 med:0.118 (0.110-0.127) | n=666 med:0.108 (0.100-0.119) | n=421 med:0.091 (0.082-0.099) | n=1,011 med:0.069 (0.063-0.075) |
| Horse | n=812 med:0.132 (0.123-0.142) | n=718 med:0.118 (0.112-0.126) | n=676 med:0.106 (0.094-0.112) | n=423 med:0.092 (0.084-0.098) | n=1,079 med:0.073 (0.067-0.079) |
| Black snub-nosed monkey | n=783 med:0.137 (0.129-0.149) | n=707 med:0.125 (0.114-0.136) | n=652 med:0.109 (0.102-0.118) | n=435 med:0.092 (0.084-0.099) | n=1,025 med:0.075 (0.069-0.082) |
| Brazilian guinea pig | n=583 med:0.126 (0.119-0.135) | n=491 med:0.124 (0.117-0.133) | n=484 med:0.118 (0.110-0.125) | n=341 med:0.101 (0.094-0.109) | n=733 med:0.085 (0.078-0.092) |
| Orangutan | n=789 med:0.134 (0.125-0.143) | n=716 med:0.117 (0.108-0.124) | n=663 med:0.105 (0.097-0.114) | n=416 med:0.082 (0.076-0.089) | n=1,024 med:0.069 (0.063-0.074) |
| Northern American deer mouse | n=874 med:0.144 (0.134-0.153) | n=765 med:0.119 (0.110-0.130) | n=714 med:0.110 (0.100-0.117) | n=462 med:0.086 (0.077-0.096) | n=1,081 med:0.066 (0.060-0.071) |
| Naked mole-rat female | n=763 med:0.129 (0.118-0.137) | n=689 med:0.119 (0.109-0.129) | n=642 med:0.105 (0.092-0.112) | n=420 med:0.088 (0.080-0.098) | n=1,018 med:0.067 (0.062-0.073) |
| Mouse | n=954 | n=839 | n=761 | n=492 | n=1,200 |

| | | | | | |
|---|---|---|---|---|---|
| | med:0.149 (0.138-0.162) | med:0.126 (0.114-0.137) | med:0.116 (0.104-0.124) | med:0.085 (0.075-0.096) | med:0.075 (0.068-0.083) |
| Daurian ground squirrel | n=693 med:0.137 (0.128-0.144) | n=630 med:0.129 (0.116-0.138) | n=562 med:0.105 (0.095-0.116) | n=364 med:0.091 (0.083-0.098) | n=831 med:0.075 (0.069-0.081) |
| Tiger | n=627 med:0.140 (0.127-0.150) | n=589 med:0.121 (0.114-0.129) | n=568 med:0.109 (0.099-0.119) | n=334 med:0.094 (0.083-0.102) | n=793 med:0.076 (0.069-0.085) |
| Lesser hedgehog tenrec | n=185 med:0.127 (0.112-0.145) | n=184 med:0.136 (0.117-0.154) | n=139 med:0.111 (0.098-0.135) | n=83 med:0.091 (0.078-0.116) | n=202 med:0.087 (0.068-0.105) |
| Mongolian gerbil | n=831 med:0.152 (0.141-0.164) | n=711 med:0.128 (0.113-0.143) | n=626 med:0.118 (0.104-0.124) | n=422 med:0.093 (0.084-0.103) | n=967 med:0.075 (0.069-0.084) |
| Ma's night monkey | n=782 med:0.140 (0.131-0.148) | n=702 med:0.124 (0.115-0.137) | n=674 med:0.109 (0.100-0.118) | n=419 med:0.089 (0.083-0.099) | n=1,039 med:0.072 (0.066-0.077) |
| Upper Galilee mountains blind mole rat | n=822 med:0.140 (0.130-0.151) | n=720 med:0.129 (0.115-0.136) | n=670 med:0.106 (0.098-0.114) | n=455 med:0.089 (0.080-0.096) | n=1,106 med:0.068 (0.062-0.073) |
| Kangaroo rat | n=680 med:0.117 (0.109-0.125) | n=624 med:0.105 (0.099-0.111) | n=598 med:0.095 (0.087-0.105) | n=402 med:0.081 (0.073-0.089) | n=963 med:0.064 (0.059-0.069) |
| Chimpanzee | n=773 med:0.135 (0.126-0.145) | n=705 med:0.121 (0.112-0.127) | n=669 med:0.109 (0.100-0.116) | n=423 med:0.091 (0.084-0.101) | n=1,056 med:0.067 (0.063-0.073) |
| Guinea Pig | n=785 med:0.126 (0.115-0.133) | n=707 med:0.115 (0.106-0.122) | n=647 med:0.107 (0.097-0.115) | n=417 med:0.084 (0.076-0.091) | n=1,003 med:0.071 (0.065-0.075) |
| American beaver | n=798 med:0.135 (0.125-0.146) | n=683 med:0.120 (0.113-0.132) | n=604 med:0.106 (0.096-0.119) | n=398 med:0.093 (0.084-0.105) | n=950 med:0.072 (0.068-0.079) |
| Wild yak | n=699 med:0.128 (0.120-0.135) | n=650 med:0.119 (0.107-0.129) | n=579 med:0.103 (0.097-0.111) | n=361 med:0.086 (0.077-0.091) | n=834 med:0.072 (0.066-0.077) |
| Coquerel's sifaka | n=754 med:0.132 (0.126-0.141) | n=681 med:0.120 (0.112-0.129) | n=640 med:0.103 (0.096-0.115) | n=406 med:0.083 (0.075-0.092) | n=1,010 med:0.066 (0.060-0.072) |
| Alpaca | n=116 med:0.185 (0.148-0.229) | n=152 med:0.149 (0.135-0.183) | n=164 med:0.125 (0.107-0.147) | n=62 med:0.138 (0.100-0.168) | n=174 med:0.086 (0.072-0.101) |
| Lesser Egyptian jerboa | n=726 med:0.127 (0.119-0.137) | n=648 med:0.115 (0.106-0.124) | n=628 med:0.102 (0.094-0.111) | n=417 med:0.092 (0.081-0.101) | n=988 med:0.062 (0.056-0.068) |
| Long-tailed chinchilla | n=769 med:0.117 (0.109-0.128) | n=690 med:0.108 (0.099-0.116) | n=651 med:0.096 (0.088-0.104) | n=425 med:0.077 (0.071-0.085) | n=1,025 med:0.062 (0.058-0.067) |
| Angola colobus | n=796 med:0.139 (0.130-0.151) | n=705 med:0.121 (0.113-0.131) | n=665 med:0.107 (0.100-0.116) | n=426 med:0.090 (0.083-0.098) | n=1,024 med:0.068 (0.063-0.076) |
| Ryukyu mouse | n=911 med:0.153 (0.139-0.164) | n=798 med:0.125 (0.116-0.137) | n=737 med:0.114 (0.106-0.123) | n=477 med:0.088 (0.079-0.097) | n=1,161 med:0.076 (0.069-0.084) |
| Chinese hamster PICR | n=902 med:0.161 (0.151-0.174) | n=788 med:0.137 (0.127-0.150) | n=726 med:0.113 (0.102-0.126) | n=469 med:0.095 (0.083-0.105) | n=1,139 med:0.072 (0.065-0.080) |
| Vervet-AGM | n=818 med:0.134 | n=735 med:0.120 | n=692 med:0.105 | n=436 med:0.085 | n=1,064 med:0.063 |

| | | | | | |
|---|---|---|---|---|---|
| | (0.124-0.144) | (0.109-0.127) | (0.095-0.114) | (0.076-0.094) | (0.058-0.069) |
| Cow | n=782<br>med:0.127<br>(0.121-0.133) | n=715<br>med:0.112<br>(0.101-0.123) | n=649<br>med:0.097<br>(0.088-0.106) | n=416<br>med:0.084<br>(0.077-0.092) | n=1,024<br>med:0.066<br>(0.061-0.071) |
| Bushbaby | n=812<br>med:0.133<br>(0.123-0.145) | n=710<br>med:0.120<br>(0.110-0.129) | n=666<br>med:0.105<br>(0.094-0.113) | n=433<br>med:0.085<br>(0.079-0.095) | n=1,072<br>med:0.067<br>(0.062-0.074) |
| Bonobo | n=807<br>med:0.137<br>(0.129-0.146) | n=708<br>med:0.123<br>(0.115-0.129) | n=673<br>med:0.110<br>(0.100-0.117) | n=431<br>med:0.087<br>(0.081-0.096) | n=1,023<br>med:0.071<br>(0.066-0.075) |
| Hyrax | n=287<br>med:0.138<br>(0.125-0.167) | n=202<br>med:0.106<br>(0.096-0.128) | n=165<br>med:0.114<br>(0.090-0.135) | n=119<br>med:0.098<br>(0.086-0.121) | n=279<br>med:0.087<br>(0.076-0.101) |
| Rabbit | n=712<br>med:0.119<br>(0.113-0.128) | n=636<br>med:0.110<br>(0.101-0.119) | n=605<br>med:0.096<br>(0.085-0.105) | n=394<br>med:0.078<br>(0.070-0.087) | n=924<br>med:0.073<br>(0.069-0.079) |
| Megabat | n=466<br>med:0.148<br>(0.136-0.159) | n=388<br>med:0.127<br>(0.115-0.135) | n=335<br>med:0.132<br>(0.116-0.143) | n=227<br>med:0.097<br>(0.084-0.109) | n=525<br>med:0.084<br>(0.075-0.093) |
| Dog | n=774<br>med:0.127<br>(0.120-0.135) | n=702<br>med:0.119<br>(0.109-0.126) | n=669<br>med:0.106<br>(0.097-0.112) | n=416<br>med:0.086<br>(0.080-0.090) | n=1,019<br>med:0.073<br>(0.069-0.079) |
| Steppe mouse | n=915<br>med:0.165<br>(0.151-0.182) | n=784<br>med:0.136<br>(0.124-0.150) | n=726<br>med:0.123<br>(0.113-0.132) | n=468<br>med:0.105<br>(0.092-0.118) | n=1,122<br>med:0.085<br>(0.079-0.092) |
| Greater bamboo lemur | n=802<br>med:0.126<br>(0.118-0.136) | n=721<br>med:0.118<br>(0.109-0.126) | n=674<br>med:0.104<br>(0.096-0.113) | n=426<br>med:0.091<br>(0.081-0.095) | n=1,033<br>med:0.066<br>(0.061-0.073) |
| Pig | n=804<br>med:0.127<br>(0.118-0.135) | n=722<br>med:0.112<br>(0.107-0.120) | n=668<br>med:0.102<br>(0.092-0.109) | n=431<br>med:0.083<br>(0.074-0.093) | n=1,045<br>med:0.066<br>(0.062-0.073) |
| Tree Shrew | n=174<br>med:0.154<br>(0.135-0.175) | n=165<br>med:0.133<br>(0.103-0.146) | n=125<br>med:0.118<br>(0.101-0.143) | n=76<br>med:0.106<br>(0.088-0.134) | n=173<br>med:0.090<br>(0.076-0.107) |
| Pika | n=314<br>med:0.133<br>(0.123-0.148) | n=259<br>med:0.100<br>(0.093-0.113) | n=214<br>med:0.107<br>(0.091-0.119) | n=141<br>med:0.091<br>(0.078-0.103) | n=316<br>med:0.076<br>(0.069-0.083) |
| Shrew | n=158<br>med:0.121<br>(0.097-0.164) | n=131<br>med:0.117<br>(0.106-0.136) | n=101<br>med:0.102<br>(0.078-0.119) | n=64<br>med:0.079<br>(0.059-0.105) | n=157<br>med:0.071<br>(0.055-0.086) |
| Squirrel | n=782<br>med:0.120<br>(0.112-0.128) | n=705<br>med:0.113<br>(0.104-0.126) | n=670<br>med:0.094<br>(0.086-0.104) | n=428<br>med:0.082<br>(0.073-0.089) | n=1,067<br>med:0.064<br>(0.059-0.069) |
| Gorilla | n=792<br>med:0.136<br>(0.126-0.144) | n=703<br>med:0.122<br>(0.114-0.131) | n=677<br>med:0.106<br>(0.101-0.116) | n=428<br>med:0.088<br>(0.082-0.097) | n=1,056<br>med:0.070<br>(0.064-0.075) |
| Cat | n=813<br>med:0.125<br>(0.118-0.134) | n=730<br>med:0.122<br>(0.112-0.129) | n=691<br>med:0.101<br>(0.090-0.108) | n=433<br>med:0.082<br>(0.077-0.089) | n=1,057<br>med:0.069<br>(0.063-0.076) |

#### Rat Reference Genome, Zeisel et al., 2015, three cell types

| Sample Size and Median dN/dS (CI95%) | Glia-specific Genes | Vasculature-specific Genes | Neuron-specific Genes |
|---|---|---|---|
| Microbat | n=955 med:0.122 (0.113-0.129) | n=328 med:0.110 (0.098-0.120) | n=629 med:0.084 (0.074-0.092) |
| Hedgehog | n=230 med:0.132 (0.106-0.146) | n=71 med:0.124 (0.086-0.143) | n=142 med:0.109 (0.084-0.134) |
| Naked mole-rat male | n=945 med:0.120 (0.113-0.131) | n=324 med:0.109 (0.097-0.117) | n=642 med:0.070 (0.066-0.078) |
| Sooty mangabey | n=1,108 med:0.127 (0.119-0.133) | n=369 med:0.104 (0.096-0.120) | n=749 med:0.076 (0.067-0.083) |
| Olive baboon | n=1,106 med:0.125 (0.119-0.133) | n=368 med:0.104 (0.094-0.123) | n=743 med:0.075 (0.068-0.082) |
| American bison | n=924 med:0.117 (0.110-0.127) | n=306 med:0.102 (0.091-0.117) | n=589 med:0.073 (0.065-0.082) |
| Leopard | n=1,065 med:0.111 (0.103-0.117) | n=352 med:0.098 (0.091-0.110) | n=740 med:0.069 (0.064-0.077) |
| Bolivian squirrel monkey | n=1,065 med:0.127 (0.119-0.135) | n=360 med:0.109 (0.098-0.126) | n=724 med:0.074 (0.066-0.081) |
| Crab-eating macaque | n=1,118 med:0.125 (0.117-0.132) | n=371 med:0.104 (0.093-0.115) | n=735 med:0.074 (0.066-0.081) |
| Human | n=1,134 med:0.122 (0.115-0.128) | n=378 med:0.105 (0.093-0.117) | n=776 med:0.073 (0.063-0.079) |
| Red fox | n=1,042 med:0.123 (0.116-0.132) | n=341 med:0.110 (0.099-0.122) | n=693 med:0.079 (0.071-0.088) |
| Shrew mouse | n=1,200 med:0.136 (0.128-0.147) | n=391 med:0.115 (0.104-0.133) | n=808 med:0.076 (0.070-0.085) |
| Arctic ground squirrel | n=1,090 med:0.119 (0.110-0.125) | n=371 med:0.104 (0.094-0.116) | n=764 med:0.065 (0.058-0.074) |
| Chinese hamster CriGri | n=1,115 med:0.141 (0.132-0.150) | n=360 med:0.122 (0.110-0.136) | n=726 med:0.086 (0.076-0.100) |
| Opossum | n=975 | n=328 | n=671 |

| | | | |
|---|---|---|---|
| | med:0.098 (0.091-0.103) | med:0.085 (0.072-0.100) | med:0.066 (0.060-0.070) |
| Dolphin | n=600 med:0.129 (0.120-0.139) | n=192 med:0.110 (0.096-0.132) | n=381 med:0.083 (0.072-0.094) |
| Capuchin | n=1,081 med:0.126 (0.117-0.133) | n=368 med:0.105 (0.095-0.120) | n=734 med:0.072 (0.066-0.086) |
| Mouse Lemur | n=1,071 med:0.121 (0.113-0.130) | n=356 med:0.107 (0.098-0.117) | n=717 med:0.073 (0.065-0.078) |
| Damara mole rat | n=986 med:0.118 (0.109-0.126) | n=338 med:0.104 (0.093-0.116) | n=693 med:0.071 (0.062-0.078) |
| Marmoset | n=1,086 med:0.128 (0.119-0.136) | n=366 med:0.110 (0.098-0.125) | n=753 med:0.078 (0.069-0.086) |
| Goat | n=1,090 med:0.113 (0.106-0.119) | n=348 med:0.101 (0.093-0.113) | n=713 med:0.073 (0.068-0.081) |
| Common wombat | n=927 med:0.093 (0.089-0.101) | n=318 med:0.084 (0.075-0.095) | n=647 med:0.062 (0.058-0.069) |
| American black bear | n=968 med:0.124 (0.116-0.134) | n=322 med:0.112 (0.096-0.126) | n=601 med:0.077 (0.071-0.083) |
| Sloth | n=175 med:0.164 (0.134-0.204) | n=49 med:0.139 (0.094-0.159) | n=92 med:0.103 (0.058-0.127) |
| Polar bear | n=976 med:0.127 (0.118-0.136) | n=314 med:0.116 (0.107-0.131) | n=610 med:0.084 (0.077-0.092) |
| Alpine marmot | n=1,042 med:0.117 (0.109-0.128) | n=357 med:0.106 (0.095-0.118) | n=701 med:0.069 (0.060-0.076) |
| Prairie vole | n=1,131 med:0.133 (0.123-0.141) | n=367 med:0.110 (0.100-0.119) | n=778 med:0.070 (0.065-0.075) |
| Donkey | n=1,098 med:0.126 (0.118-0.133) | n=360 med:0.106 (0.092-0.115) | n=742 med:0.082 (0.073-0.088) |
| Golden Hamster | n=1,093 med:0.142 (0.133-0.151) | n=354 med:0.122 (0.103-0.136) | n=753 med:0.080 (0.072-0.089) |
| Armadillo | n=1,008 med:0.120 (0.114-0.128) | n=334 med:0.103 (0.093-0.117) | n=664 med:0.075 (0.070-0.083) |
| Ugandan red Colobus | n=1,075 | n=352 | n=739 |

| | | | |
|---|---|---|---|
| | med:0.126<br>(0.118-0.132) | med:0.109<br>(0.095-0.120) | med:0.078<br>(0.069-0.087) |
| Pig-tailed macaque | n=1,104<br>med:0.127<br>(0.120-0.135) | n=372<br>med:0.109<br>(0.098-0.120) | n=747<br>med:0.073<br>(0.066-0.081) |
| Panda | n=1,070<br>med:0.114<br>(0.106-0.121) | n=349<br>med:0.101<br>(0.092-0.112) | n=712<br>med:0.071<br>(0.063-0.079) |
| Ferret | n=1,067<br>med:0.115<br>(0.107-0.122) | n=349<br>med:0.100<br>(0.091-0.113) | n=713<br>med:0.072<br>(0.066-0.078) |
| Koala | n=1,004<br>med:0.097<br>(0.094-0.105) | n=342<br>med:0.085<br>(0.076-0.092) | n=711<br>med:0.066<br>(0.060-0.072) |
| Wallaby | n=276<br>med:0.087<br>(0.082-0.097) | n=103<br>med:0.092<br>(0.076-0.098) | n=196<br>med:0.064<br>(0.051-0.073) |
| Tarsier | n=918<br>med:0.124<br>(0.113-0.132) | n=290<br>med:0.111<br>(0.095-0.126) | n=583<br>med:0.084<br>(0.078-0.092) |
| Degu | n=924<br>med:0.118<br>(0.109-0.122) | n=321<br>med:0.108<br>(0.096-0.119) | n=646<br>med:0.073<br>(0.067-0.080) |
| Chinese hamster CHOK1GS | n=1,199<br>med:0.138<br>(0.129-0.146) | n=391<br>med:0.121<br>(0.111-0.134) | n=809<br>med:0.079<br>(0.070-0.087) |
| Drill | n=1,083<br>med:0.127<br>(0.122-0.133) | n=356<br>med:0.106<br>(0.095-0.115) | n=708<br>med:0.076<br>(0.066-0.083) |
| Gibbon | n=1,059<br>med:0.135<br>(0.127-0.143) | n=357<br>med:0.110<br>(0.099-0.121) | n=716<br>med:0.083<br>(0.075-0.092) |
| Macaque | n=1,122<br>med:0.128<br>(0.119-0.136) | n=378<br>med:0.106<br>(0.094-0.116) | n=775<br>med:0.077<br>(0.069-0.088) |
| American mink | n=1,018<br>med:0.113<br>(0.107-0.121) | n=334<br>med:0.096<br>(0.084-0.110) | n=691<br>med:0.069<br>(0.063-0.076) |
| Gelada | n=1,119<br>med:0.123<br>(0.116-0.130) | n=371<br>med:0.104<br>(0.094-0.116) | n=753<br>med:0.073<br>(0.065-0.080) |
| Sheep | n=1,060<br>med:0.122<br>(0.114-0.128) | n=333<br>med:0.111<br>(0.100-0.120) | n=691<br>med:0.081<br>(0.075-0.088) |
| Tasmanian devil | n=916<br>med:0.094<br>(0.088-0.099) | n=308<br>med:0.083<br>(0.073-0.089) | n=613<br>med:0.062<br>(0.057-0.070) |
| Elephant | n=1,071 | n=345 | n=702 |

| | | | |
|---|---|---|---|
| | med:0.121 (0.111-0.128) | med:0.108 (0.098-0.124) | med:0.079 (0.073-0.087) |
| Algerian mouse | n=1,214 med:0.136 (0.127-0.144) | n=394 med:0.119 (0.097-0.133) | n=817 med:0.079 (0.067-0.087) |
| Dingo | n=1,046 med:0.115 (0.109-0.122) | n=342 med:0.101 (0.093-0.109) | n=710 med:0.071 (0.062-0.078) |
| Golden snub-nosed monkey | n=1,058 med:0.127 (0.120-0.137) | n=357 med:0.106 (0.098-0.115) | n=710 med:0.073 (0.066-0.082) |
| Horse | n=1,099 med:0.128 (0.121-0.134) | n=366 med:0.105 (0.094-0.119) | n=742 med:0.079 (0.070-0.088) |
| Black snub-nosed monkey | n=1,068 med:0.131 (0.123-0.139) | n=353 med:0.113 (0.102-0.128) | n=715 med:0.081 (0.073-0.087) |
| Brazilian guinea pig | n=788 med:0.122 (0.117-0.130) | n=270 med:0.118 (0.103-0.128) | n=514 med:0.094 (0.086-0.099) |
| Orangutan | n=1,070 med:0.123 (0.114-0.130) | n=366 med:0.111 (0.097-0.122) | n=719 med:0.072 (0.064-0.078) |
| Northern American deer mouse | n=1,147 med:0.132 (0.120-0.140) | n=376 med:0.107 (0.096-0.124) | n=753 med:0.068 (0.062-0.075) |
| Naked mole-rat female | n=1,004 med:0.123 (0.115-0.132) | n=337 med:0.109 (0.099-0.120) | n=699 med:0.073 (0.066-0.080) |
| Mouse | n=1,250 med:0.134 (0.125-0.142) | n=408 med:0.117 (0.102-0.129) | n=837 med:0.077 (0.066-0.084) |
| Daurian ground squirrel | n=947 med:0.127 (0.119-0.133) | n=323 med:0.112 (0.099-0.128) | n=582 med:0.077 (0.067-0.089) |
| Tiger | n=910 med:0.121 (0.116-0.130) | n=291 med:0.117 (0.097-0.131) | n=573 med:0.077 (0.072-0.085) |
| Lesser hedgehog tenrec | n=254 med:0.121 (0.104-0.140) | n=87 med:0.119 (0.090-0.143) | n=148 med:0.085 (0.075-0.102) |
| Mongolian gerbil | n=1,084 med:0.142 (0.128-0.154) | n=344 med:0.112 (0.102-0.131) | n=682 med:0.078 (0.070-0.089) |
| Ma's night monkey | n=1,061 med:0.129 (0.120-0.138) | n=359 med:0.112 (0.099-0.125) | n=728 med:0.079 (0.069-0.088) |

| | | | |
|---|---|---|---|
| Upper Galilee mountains blind mole rat | n=1,097 med:0.126 (0.117-0.135) | n=355 med:0.111 (0.099-0.124) | n=743 med:0.069 (0.063-0.078) |
| Kangaroo rat | n=954 med:0.109 (0.102-0.118) | n=317 med:0.100 (0.089-0.114) | n=659 med:0.069 (0.062-0.077) |
| Chimpanzee | n=1,054 med:0.128 (0.121-0.136) | n=353 med:0.108 (0.094-0.120) | n=718 med:0.075 (0.066-0.081) |
| Guinea Pig | n=1,035 med:0.117 (0.109-0.124) | n=351 med:0.109 (0.097-0.118) | n=721 med:0.073 (0.066-0.079) |
| American beaver | n=996 med:0.126 (0.117-0.135) | n=340 med:0.116 (0.103-0.133) | n=644 med:0.077 (0.070-0.087) |
| Wild yak | n=988 med:0.116 (0.109-0.125) | n=317 med:0.102 (0.090-0.113) | n=602 med:0.077 (0.070-0.086) |
| Coquerel's sifaka | n=1,033 med:0.121 (0.115-0.130) | n=348 med:0.106 (0.096-0.123) | n=708 med:0.077 (0.070-0.087) |
| Alpaca | n=239 med:0.172 (0.146-0.194) | n=69 med:0.120 (0.107-0.147) | n=147 med:0.081 (0.070-0.098) |
| Lesser Egyptian jerboa | n=1,008 med:0.123 (0.114-0.129) | n=324 med:0.103 (0.091-0.119) | n=695 med:0.071 (0.063-0.076) |
| Long-tailed chinchilla | n=1,039 med:0.112 (0.105-0.118) | n=352 med:0.097 (0.090-0.106) | n=725 med:0.069 (0.061-0.076) |
| Angola colobus | n=1,080 med:0.126 (0.119-0.133) | n=359 med:0.111 (0.096-0.118) | n=718 med:0.075 (0.067-0.083) |
| Ryukyu mouse | n=1,192 med:0.134 (0.125-0.144) | n=391 med:0.120 (0.100-0.132) | n=810 med:0.073 (0.065-0.083) |
| Chinese hamster PICR | n=1,177 med:0.144 (0.135-0.151) | n=382 med:0.123 (0.112-0.135) | n=807 med:0.080 (0.071-0.089) |
| Vervet-AGM | n=1,117 med:0.121 (0.114-0.130) | n=365 med:0.107 (0.091-0.119) | n=748 med:0.068 (0.062-0.076) |
| Cow | n=1,093 med:0.115 (0.109-0.123) | n=353 med:0.105 (0.090-0.116) | n=715 med:0.073 (0.066-0.080) |
| Bushbaby | n=1,089 med:0.125 (0.116-0.134) | n=362 med:0.112 (0.102-0.129) | n=734 med:0.075 (0.068-0.082) |

| | | | |
|---|---|---|---|
| Bonobo | n=1,093 med:0.127 (0.121-0.135) | n=359 med:0.112 (0.102-0.126) | n=712 med:0.078 (0.070-0.086) |
| Hyrax | n=326 med:0.133 (0.117-0.152) | n=102 med:0.111 (0.090-0.145) | n=195 med:0.093 (0.082-0.106) |
| Rabbit | n=987 med:0.116 (0.109-0.123) | n=328 med:0.098 (0.091-0.112) | n=660 med:0.079 (0.071-0.087) |
| Megabat | n=608 med:0.133 (0.122-0.143) | n=217 med:0.122 (0.108-0.138) | n=362 med:0.094 (0.083-0.103) |
| Dog | n=1,058 med:0.121 (0.112-0.131) | n=355 med:0.108 (0.097-0.125) | n=711 med:0.077 (0.071-0.086) |
| Steppe mouse | n=1,177 med:0.152 (0.144-0.164) | n=390 med:0.126 (0.111-0.141) | n=794 med:0.088 (0.079-0.094) |
| Greater bamboo lemur | n=1,096 med:0.120 (0.114-0.128) | n=358 med:0.106 (0.095-0.118) | n=734 med:0.077 (0.067-0.083) |
| Pig | n=1,097 med:0.121 (0.112-0.128) | n=363 med:0.099 (0.091-0.110) | n=740 med:0.074 (0.066-0.080) |
| Tree Shrew | n=211 med:0.135 (0.119-0.159) | n=69 med:0.110 (0.091-0.147) | n=139 med:0.109 (0.084-0.127) |
| Pika | n=387 med:0.126 (0.107-0.136) | n=139 med:0.098 (0.088-0.124) | n=209 med:0.082 (0.069-0.097) |
| Shrew | n=186 med:0.123 (0.104-0.143) | n=52 med:0.113 (0.073-0.129) | n=119 med:0.080 (0.054-0.106) |
| Squirrel | n=1,071 med:0.118 (0.110-0.124) | n=367 med:0.101 (0.089-0.112) | n=750 med:0.065 (0.058-0.072) |
| Gorilla | n=1,081 med:0.126 (0.118-0.135) | n=357 med:0.109 (0.095-0.122) | n=729 med:0.077 (0.069-0.084) |
| Cat | n=1,118 med:0.118 (0.111-0.124) | n=366 med:0.108 (0.095-0.122) | n=756 med:0.073 (0.068-0.079) |

### Rat Reference Genome, Zeisel et al., 2015, cell subtypes

| Sample Size and Median dN/dS (CI95%) | Ependymal-specific Genes | Microglia-specific Genes | Endothelial-specific Genes | Mural-specific Genes | Oligodendrocyte-specific Genes | Astrocyte-specific Genes | Interneuron-specific Genes | S1 Pyramidal-specific Genes | CA1 Pyramidal-specific Genes |
|---|---|---|---|---|---|---|---|---|---|
| Microbat | n=275 med:0.153 (0.135-0.164) | n=248 med:0.136 (0.126-0.153) | n=230 med:0.116 (0.100-0.132) | n=98 med:0.099 (0.076-0.116) | n=277 med:0.098 (0.081-0.105) | n=155 med:0.105 (0.089-0.120) | n=225 med:0.093 (0.080-0.102) | n=172 med:0.083 (0.068-0.100) | n=232 med:0.071 (0.063-0.086) |
| Hedgehog | n=52 med:0.148 (0.132-0.199) | n=69 med:0.152 (0.123-0.194) | n=55 med:0.124 (0.083-0.144) | n=16 med:0.113 (0.057-0.157) | n=73 med:0.101 (0.082-0.137) | n=36 med:0.089 (0.046-0.116) | n=48 med:0.127 (0.092-0.151) | n=39 med:0.078 (0.056-0.115) | n=55 med:0.132 (0.065-0.162) |
| Naked mole-rat male | n=276 med:0.146 (0.130-0.163) | n=245 med:0.139 (0.125-0.148) | n=222 med:0.110 (0.097-0.128) | n=102 med:0.106 (0.075-0.120) | n=270 med:0.104 (0.091-0.114) | n=154 med:0.097 (0.086-0.110) | n=224 med:0.077 (0.069-0.095) | n=178 med:0.071 (0.061-0.089) | n=240 med:0.064 (0.056-0.072) |
| Sooty mangabey | n=321 med:0.152 (0.134-0.171) | n=287 med:0.149 (0.132-0.162) | n=260 med:0.107 (0.094-0.130) | n=109 med:0.103 (0.070-0.125) | n=320 med:0.102 (0.091-0.112) | n=180 med:0.105 (0.093-0.122) | n=259 med:0.083 (0.066-0.097) | n=208 med:0.080 (0.067-0.093) | n=282 med:0.068 (0.059-0.077) |
| Olive baboon | n=321 med:0.148 (0.124-0.169) | n=289 med:0.147 (0.133-0.158) | n=260 med:0.110 (0.094-0.131) | n=108 med:0.098 (0.083-0.120) | n=320 med:0.103 (0.091-0.116) | n=176 med:0.109 (0.094-0.122) | n=254 med:0.082 (0.065-0.098) | n=206 med:0.076 (0.067-0.091) | n=283 med:0.069 (0.058-0.081) |
| American bison | n=283 med:0.137 (0.118-0.158) | n=246 med:0.140 (0.119-0.159) | n=214 med:0.115 (0.094-0.131) | n=92 med:0.086 (0.078-0.102) | n=255 med:0.104 (0.090-0.113) | n=140 med:0.097 (0.085-0.108) | n=199 med:0.083 (0.069-0.089) | n=160 med:0.073 (0.060-0.088) | n=230 med:0.063 (0.052-0.078) |
| Leopard | n=321 med:0.129 (0.117-0.146) | n=265 med:0.129 (0.115-0.146) | n=241 med:0.104 (0.091-0.121) | n=111 med:0.092 (0.074-0.111) | n=305 med:0.090 (0.079-0.103) | n=174 med:0.094 (0.080-0.112) | n=258 med:0.073 (0.060-0.085) | n=198 med:0.074 (0.054-0.085) | n=284 med:0.066 (0.057-0.074) |
| Bolivian squirrel monkey | n=323 med:0.145 (0.130-0.164) | n=269 med:0.147 (0.128-0.168) | n=250 med:0.114 (0.098-0.134) | n=110 med:0.103 (0.085-0.120) | n=306 med:0.100 (0.089-0.114) | n=167 med:0.111 (0.094-0.129) | n=246 med:0.079 (0.066-0.097) | n=197 med:0.074 (0.063-0.087) | n=281 med:0.066 (0.057-0.078) |
| Crab-eating macaque | n=325 med:0.150 (0.129-0.168) | n=294 med:0.144 (0.128-0.157) | n=258 med:0.105 (0.092-0.128) | n=113 med:0.101 (0.071-0.115) | n=322 med:0.100 (0.090-0.112) | n=177 med:0.104 (0.089-0.126) | n=251 med:0.079 (0.064-0.094) | n=201 med:0.079 (0.068-0.095) | n=283 med:0.065 (0.057-0.077) |
| Human | n=333 med:0.145 (0.133-0.169) | n=292 med:0.140 (0.126-0.153) | n=264 med:0.104 (0.092-0.126) | n=114 med:0.105 (0.086-0.116) | n=326 med:0.097 (0.087-0.110) | n=183 med:0.108 (0.096-0.124) | n=267 med:0.082 (0.066-0.096) | n=206 med:0.074 (0.061-0.086) | n=303 med:0.063 (0.054-0.076) |
| Red fox | n=300 med:0.145 (0.125-0.162) | n=273 med:0.144 (0.125-0.156) | n=237 med:0.110 (0.098-0.127) | n=104 med:0.107 (0.090-0.126) | n=302 med:0.104 (0.093-0.114) | n=167 med:0.109 (0.095-0.122) | n=238 med:0.080 (0.068-0.093) | n=189 med:0.086 (0.068-0.099) | n=266 med:0.073 (0.062-0.089) |
| Shrew mouse | n=344 med:0.159 (0.140-0.182) | n=322 med:0.172 (0.149-0.196) | n=267 med:0.114 (0.101-0.135) | n=124 med:0.127 (0.104-0.136) | n=348 med:0.115 (0.098-0.128) | n=186 med:0.104 (0.089-0.121) | n=278 med:0.077 (0.066-0.090) | n=218 med:0.080 (0.063-0.090) | n=312 med:0.072 (0.061-0.088) |
| Arctic ground squirrel | n=307 med:0.137 (0.122-0.154) | n=281 med:0.141 (0.119-0.159) | n=258 med:0.107 (0.095-0.124) | n=113 med:0.098 (0.073-0.110) | n=321 med:0.092 (0.083-0.105) | n=181 med:0.104 (0.083-0.114) | n=263 med:0.070 (0.060-0.082) | n=206 med:0.075 (0.053-0.088) | n=295 med:0.058 (0.049-0.069) |
| Chinese hamster CriGri | n=320 med:0.156 (0.140-0.184) | n=301 med:0.173 (0.153-0.201) | n=249 med:0.123 (0.110-0.143) | n=111 med:0.117 (0.096-0.136) | n=327 med:0.115 (0.102-0.132) | n=167 med:0.113 (0.095-0.132) | n=257 med:0.086 (0.073-0.101) | n=194 med:0.087 (0.068-0.112) | n=275 med:0.086 (0.070-0.106) |
| Opossum | n=283 med:0.117 (0.108-0.129) | n=243 med:0.104 (0.088-0.112) | n=222 med:0.096 (0.074-0.106) | n=106 med:0.078 (0.058-0.096) | n=291 med:0.083 (0.070-0.093) | n=158 med:0.080 (0.070-0.096) | n=233 med:0.066 (0.060-0.073) | n=190 med:0.065 (0.055-0.077) | n=248 med:0.065 (0.054-0.073) |
| Dolphin | n=176 med:0.157 (0.127-0.176) | n=157 med:0.139 (0.125-0.162) | n=131 med:0.128 (0.100-0.152) | n=61 med:0.095 (0.084-0.119) | n=168 med:0.111 (0.091-0.132) | n=99 med:0.111 (0.081-0.124) | n=134 med:0.101 (0.077-0.123) | n=106 med:0.080 (0.058-0.092) | n=141 med:0.076 (0.065-0.091) |
| Capuchin | n=317 med:0.153 (0.132-0.170) | n=277 med:0.141 (0.130-0.159) | n=256 med:0.108 (0.095-0.133) | n=112 med:0.100 (0.085-0.117) | n=314 med:0.101 (0.089-0.107) | n=173 med:0.109 (0.091-0.126) | n=254 med:0.085 (0.064-0.100) | n=200 med:0.073 (0.063-0.093) | n=280 med:0.067 (0.060-0.078) |
| Mouse Lemur | n=315 med:0.149 (0.130-0.164) | n=282 med:0.140 (0.125-0.160) | n=248 med:0.111 (0.099-0.124) | n=108 med:0.099 (0.073-0.118) | n=305 med:0.093 (0.080-0.106) | n=169 med:0.102 (0.084-0.117) | n=250 med:0.077 (0.071-0.084) | n=198 med:0.079 (0.064-0.091) | n=269 med:0.062 (0.054-0.073) |
| Damara mole rat | n=286 med:0.145 (0.124-0.157) | n=251 med:0.137 (0.127-0.156) | n=233 med:0.108 (0.093-0.121) | n=105 med:0.101 (0.085-0.116) | n=294 med:0.092 (0.082-0.106) | n=155 med:0.100 (0.086-0.110) | n=244 med:0.072 (0.063-0.084) | n=187 med:0.077 (0.058-0.096) | n=262 med:0.063 (0.051-0.079) |
| Marmoset | n=323 med:0.157 (0.136-0.173) | n=285 med:0.140 (0.124-0.155) | n=256 med:0.120 (0.097-0.139) | n=110 med:0.101 (0.087-0.116) | n=310 med:0.098 (0.088-0.108) | n=168 med:0.123 (0.106-0.141) | n=265 med:0.087 (0.071-0.105) | n=203 med:0.083 (0.067-0.099) | n=285 med:0.068 (0.057-0.079) |
| Goat | n=319 med:0.129 (0.114-0.146) | n=280 med:0.138 (0.122-0.159) | n=240 med:0.108 (0.097-0.121) | n=108 med:0.085 (0.077-0.107) | n=320 med:0.091 (0.081-0.102) | n=171 med:0.097 (0.087-0.107) | n=245 med:0.077 (0.069-0.087) | n=195 med:0.074 (0.063-0.085) | n=273 med:0.067 (0.059-0.080) |
| Common wombat | n=253 med:0.115 (0.099-0.122) | n=250 med:0.108 (0.091-0.118) | n=220 med:0.090 (0.079-0.103) | n=98 med:0.071 (0.057-0.099) | n=279 med:0.082 (0.072-0.089) | n=145 med:0.087 (0.074-0.092) | n=212 med:0.064 (0.058-0.072) | n=181 med:0.064 (0.050-0.077) | n=254 med:0.060 (0.051-0.069) |
| American black bear | n=275 med:0.150 (0.130-0.164) | n=249 med:0.136 (0.121-0.153) | n=220 med:0.119 (0.101-0.142) | n=102 med:0.096 (0.088-0.114) | n=284 med:0.105 (0.090-0.121) | n=160 med:0.111 (0.088-0.121) | n=208 med:0.088 (0.074-0.103) | n=157 med:0.075 (0.061-0.091) | n=236 med:0.073 (0.059-0.080) |
| Sloth | n=60 med:0.199 (0.134-0.241) | n=46 med:0.182 (0.118-0.274) | n=37 med:0.149 (0.094-0.174) | n=12 med:0.123 (0.052-0.159) | n=52 med:0.138 (0.088-0.191) | n=17 med:0.128 (0.068-0.228) | n=32 med:0.112 (0.041-0.167) | n=28 med:0.111 (0.040-0.164) | n=32 med:0.078 (0.049-0.157) |
| Polar bear | n=285 med:0.159 (0.144-0.169) | n=256 med:0.141 (0.127-0.170) | n=220 med:0.125 (0.110-0.144) | n=94 med:0.107 (0.085-0.117) | n=288 med:0.106 (0.094-0.116) | n=147 med:0.107 (0.086-0.119) | n=214 med:0.086 (0.076-0.103) | n=155 med:0.089 (0.070-0.110) | n=241 med:0.080 (0.072-0.090) |
| Alpine marmot | n=296 med:0.142 (0.129-0.152) | n=281 med:0.136 (0.120-0.158) | n=248 med:0.113 (0.098-0.133) | n=109 med:0.097 (0.072-0.115) | n=296 med:0.096 (0.084-0.108) | n=169 med:0.101 (0.079-0.112) | n=235 med:0.071 (0.060-0.085) | n=191 med:0.087 (0.062-0.094) | n=275 med:0.058 (0.052-0.070) |
| Prairie vole | n=325 med:0.152 (0.139-0.176) | n=299 med:0.166 (0.147-0.186) | n=252 med:0.109 (0.092-0.119) | n=115 med:0.115 (0.098-0.131) | n=334 med:0.096 (0.084-0.114) | n=173 med:0.105 (0.090-0.127) | n=270 med:0.073 (0.062-0.085) | n=213 med:0.074 (0.057-0.091) | n=295 med:0.068 (0.056-0.074) |
| Donkey | n=319 med:0.154 (0.141-0.165) | n=287 med:0.139 (0.129-0.156) | n=245 med:0.111 (0.099-0.124) | n=115 med:0.092 (0.085-0.111) | n=318 med:0.101 (0.090-0.113) | n=174 med:0.109 (0.091-0.123) | n=251 med:0.086 (0.076-0.095) | n=201 med:0.085 (0.071-0.092) | n=290 med:0.072 (0.059-0.087) |
| Golden Hamster | n=313 med:0.158 (0.143-0.178) | n=286 med:0.175 (0.162-0.198) | n=243 med:0.133 (0.109-0.147) | n=111 med:0.102 (0.081-0.127) | n=323 med:0.110 (0.096-0.126) | n=171 med:0.125 (0.104-0.142) | n=258 med:0.076 (0.068-0.090) | n=204 med:0.089 (0.066-0.115) | n=291 med:0.079 (0.065-0.094) |
| Armadillo | n=287 med:0.146 (0.131-0.163) | n=273 med:0.138 (0.124-0.151) | n=233 med:0.112 (0.094-0.126) | n=101 med:0.095 (0.078-0.113) | n=290 med:0.100 (0.084-0.107) | n=158 med:0.106 (0.095-0.117) | n=223 med:0.076 (0.069-0.090) | n=179 med:0.088 (0.074-0.103) | n=262 med:0.068 (0.058-0.077) |
| Ugandan red Colobus | n=313 med:0.152 (0.128-0.169) | n=277 med:0.142 (0.129-0.159) | n=249 med:0.114 (0.095-0.134) | n=103 med:0.106 (0.084-0.118) | n=308 med:0.101 (0.093-0.114) | n=177 med:0.111 (0.096-0.124) | n=261 med:0.089 (0.074-0.099) | n=199 med:0.078 (0.063-0.094) | n=279 med:0.068 (0.059-0.082) |
| Pig-tailed macaque | n=321 med:0.156 (0.134-0.171) | n=291 med:0.150 (0.133-0.159) | n=261 med:0.114 (0.099-0.131) | n=111 med:0.103 (0.071-0.118) | n=319 med:0.100 (0.090-0.115) | n=173 med:0.108 (0.089-0.124) | n=257 med:0.080 (0.065-0.097) | n=204 med:0.078 (0.064-0.093) | n=286 med:0.066 (0.056-0.079) |
| Panda | n=305 med:0.135 (0.121-0.157) | n=278 med:0.136 (0.118-0.154) | n=245 med:0.107 (0.093-0.121) | n=104 med:0.094 (0.079-0.108) | n=315 med:0.091 (0.081-0.100) | n=172 med:0.096 (0.081-0.112) | n=249 med:0.078 (0.062-0.089) | n=193 med:0.072 (0.062-0.090) | n=270 med:0.064 (0.054-0.072) |

| | | | | | | | | | |
|---|---|---|---|---|---|---|---|---|---|
| Ferret | n=315<br>med:0.139<br>(0.121-0.157) | n=279<br>med:0.134<br>(0.121-0.156) | n=245<br>med:0.108<br>(0.093-0.124) | n=104<br>med:0.091<br>(0.069-0.105) | n=302<br>med:0.088<br>(0.076-0.099) | n=171<br>med:0.096<br>(0.079-0.112) | n=244<br>med:0.078<br>(0.069-0.091) | n=197<br>med:0.076<br>(0.060-0.090) | n=272<br>med:0.063<br>(0.054-0.072) |
| Koala | n=281<br>med:0.119<br>(0.105-0.131) | n=254<br>med:0.110<br>(0.097-0.124) | n=233<br>med:0.086<br>(0.077-0.095) | n=109<br>med:0.077<br>(0.065-0.097) | n=310<br>med:0.089<br>(0.075-0.095) | n=159<br>med:0.086<br>(0.077-0.095) | n=246<br>med:0.068<br>(0.060-0.075) | n=195<br>med:0.066<br>(0.054-0.078) | n=270<br>med:0.064<br>(0.053-0.075) |
| Wallaby | n=74<br>med:0.096<br>(0.080-0.124) | n=70<br>med:0.089<br>(0.074-0.118) | n=68<br>med:0.092<br>(0.074-0.107) | n=35<br>med:0.083<br>(0.052-0.099) | n=92<br>med:0.078<br>(0.069-0.095) | n=40<br>med:0.093<br>(0.077-0.106) | n=70<br>med:0.052<br>(0.043-0.073) | n=63<br>med:0.063<br>(0.049-0.074) | n=63<br>med:0.074<br>(0.050-0.094) |
| Tarsier | n=266<br>med:0.156<br>(0.141-0.171) | n=236<br>med:0.139<br>(0.127-0.157) | n=208<br>med:0.114<br>(0.095-0.132) | n=82<br>med:0.106<br>(0.076-0.127) | n=274<br>med:0.098<br>(0.088-0.105) | n=142<br>med:0.106<br>(0.091-0.120) | n=207<br>med:0.090<br>(0.079-0.102) | n=159<br>med:0.087<br>(0.066-0.095) | n=217<br>med:0.079<br>(0.063-0.094) |
| Degu | n=273<br>med:0.136<br>(0.121-0.147) | n=226<br>med:0.134<br>(0.115-0.147) | n=220<br>med:0.110<br>(0.095-0.127) | n=101<br>med:0.104<br>(0.086-0.119) | n=276<br>med:0.100<br>(0.087-0.111) | n=149<br>med:0.089<br>(0.075-0.111) | n=225<br>med:0.072<br>(0.064-0.081) | n=174<br>med:0.085<br>(0.068-0.105) | n=247<br>med:0.067<br>(0.057-0.077) |
| Chinese hamster CHOK1GS | n=343<br>med:0.159<br>(0.144-0.184) | n=324<br>med:0.172<br>(0.150-0.200) | n=270<br>med:0.128<br>(0.112-0.142) | n=121<br>med:0.116<br>(0.097-0.126) | n=345<br>med:0.106<br>(0.090-0.120) | n=187<br>med:0.104<br>(0.091-0.127) | n=284<br>med:0.079<br>(0.068-0.093) | n=221<br>med:0.086<br>(0.062-0.102) | n=304<br>med:0.074<br>(0.062-0.092) |
| Drill | n=319<br>med:0.156<br>(0.135-0.172) | n=283<br>med:0.148<br>(0.131-0.161) | n=247<br>med:0.112<br>(0.096-0.130) | n=109<br>med:0.099<br>(0.067-0.112) | n=307<br>med:0.103<br>(0.092-0.115) | n=174<br>med:0.102<br>(0.090-0.126) | n=241<br>med:0.082<br>(0.066-0.097) | n=199<br>med:0.077<br>(0.065-0.096) | n=268<br>med:0.068<br>(0.059-0.080) |
| Gibbon | n=318<br>med:0.160<br>(0.141-0.177) | n=278<br>med:0.159<br>(0.137-0.168) | n=247<br>med:0.114<br>(0.101-0.136) | n=110<br>med:0.099<br>(0.082-0.120) | n=293<br>med:0.111<br>(0.099-0.118) | n=170<br>med:0.114<br>(0.097-0.135) | n=248<br>med:0.094<br>(0.080-0.105) | n=193<br>med:0.089<br>(0.075-0.101) | n=275<br>med:0.069<br>(0.060-0.079) |
| Macaque | n=319<br>med:0.156<br>(0.136-0.170) | n=301<br>med:0.147<br>(0.131-0.162) | n=262<br>med:0.104<br>(0.092-0.122) | n=116<br>med:0.108<br>(0.085-0.128) | n=322<br>med:0.101<br>(0.089-0.113) | n=180<br>med:0.112<br>(0.097-0.131) | n=270<br>med:0.086<br>(0.072-0.102) | n=208<br>med:0.081<br>(0.068-0.097) | n=297<br>med:0.066<br>(0.059-0.078) |
| American mink | n=306<br>med:0.129<br>(0.115-0.142) | n=252<br>med:0.130<br>(0.113-0.155) | n=228<br>med:0.103<br>(0.089-0.120) | n=106<br>med:0.084<br>(0.074-0.108) | n=300<br>med:0.094<br>(0.082-0.107) | n=160<br>med:0.099<br>(0.090-0.111) | n=235<br>med:0.076<br>(0.063-0.090) | n=191<br>med:0.068<br>(0.057-0.084) | n=265<br>med:0.065<br>(0.058-0.072) |
| Gelada | n=325<br>med:0.148<br>(0.130-0.166) | n=289<br>med:0.147<br>(0.130-0.160) | n=260<br>med:0.107<br>(0.093-0.124) | n=111<br>med:0.101<br>(0.071-0.119) | n=325<br>med:0.101<br>(0.089-0.109) | n=180<br>med:0.105<br>(0.089-0.120) | n=260<br>med:0.080<br>(0.065-0.094) | n=203<br>med:0.075<br>(0.057-0.085) | n=290<br>med:0.068<br>(0.058-0.081) |
| Sheep | n=317<br>med:0.139<br>(0.124-0.158) | n=272<br>med:0.144<br>(0.128-0.166) | n=228<br>med:0.117<br>(0.104-0.127) | n=105<br>med:0.092<br>(0.082-0.118) | n=302<br>med:0.096<br>(0.087-0.108) | n=169<br>med:0.101<br>(0.089-0.114) | n=239<br>med:0.081<br>(0.070-0.095) | n=185<br>med:0.078<br>(0.069-0.094) | n=267<br>med:0.081<br>(0.071-0.093) |
| Tasmanian devil | n=262<br>med:0.111<br>(0.097-0.122) | n=237<br>med:0.100<br>(0.083-0.115) | n=213<br>med:0.084<br>(0.073-0.094) | n=95<br>med:0.080<br>(0.059-0.093) | n=280<br>med:0.080<br>(0.069-0.093) | n=137<br>med:0.088<br>(0.078-0.097) | n=210<br>med:0.064<br>(0.056-0.073) | n=169<br>med:0.058<br>(0.049-0.072) | n=234<br>med:0.064<br>(0.055-0.075) |
| Elephant | n=310<br>med:0.141<br>(0.127-0.161) | n=272<br>med:0.137<br>(0.124-0.154) | n=238<br>med:0.107<br>(0.097-0.128) | n=107<br>med:0.111<br>(0.088-0.134) | n=322<br>med:0.102<br>(0.090-0.112) | n=167<br>med:0.105<br>(0.088-0.112) | n=252<br>med:0.088<br>(0.074-0.100) | n=186<br>med:0.086<br>(0.070-0.102) | n=264<br>med:0.070<br>(0.059-0.079) |
| Algerian mouse | n=348<br>med:0.165<br>(0.143-0.191) | n=327<br>med:0.173<br>(0.153-0.192) | n=272<br>med:0.121<br>(0.096-0.135) | n=122<br>med:0.112<br>(0.091-0.139) | n=347<br>med:0.114<br>(0.095-0.127) | n=192<br>med:0.105<br>(0.091-0.125) | n=283<br>med:0.082<br>(0.059-0.091) | n=223<br>med:0.080<br>(0.062-0.090) | n=311<br>med:0.073<br>(0.062-0.088) |
| Dingo | n=300<br>med:0.136<br>(0.122-0.155) | n=278<br>med:0.131<br>(0.115-0.148) | n=240<br>med:0.101<br>(0.092-0.112) | n=102<br>med:0.101<br>(0.088-0.113) | n=303<br>med:0.092<br>(0.083-0.104) | n=165<br>med:0.105<br>(0.089-0.120) | n=244<br>med:0.071<br>(0.059-0.083) | n=190<br>med:0.075<br>(0.060-0.090) | n=276<br>med:0.068<br>(0.055-0.080) |
| Golden snub-nosed monkey | n=315<br>med:0.152<br>(0.135-0.169) | n=278<br>med:0.149<br>(0.132-0.164) | n=253<br>med:0.106<br>(0.095-0.126) | n=104<br>med:0.106<br>(0.081-0.116) | n=303<br>med:0.104<br>(0.092-0.117) | n=162<br>med:0.110<br>(0.093-0.126) | n=241<br>med:0.082<br>(0.068-0.096) | n=193<br>med:0.078<br>(0.064-0.094) | n=276<br>med:0.066<br>(0.056-0.076) |
| Horse | n=324<br>med:0.149<br>(0.128-0.166) | n=286<br>med:0.139<br>(0.127-0.159) | n=255<br>med:0.111<br>(0.096-0.125) | n=111<br>med:0.094<br>(0.085-0.116) | n=314<br>med:0.107<br>(0.093-0.122) | n=175<br>med:0.113<br>(0.093-0.128) | n=256<br>med:0.089<br>(0.074-0.100) | n=204<br>med:0.079<br>(0.065-0.094) | n=282<br>med:0.069<br>(0.058-0.085) |
| Black snub-nosed monkey | n=316<br>med:0.158<br>(0.145-0.178) | n=272<br>med:0.152<br>(0.133-0.163) | n=246<br>med:0.122<br>(0.103-0.143) | n=107<br>med:0.106<br>(0.092-0.123) | n=317<br>med:0.101<br>(0.090-0.113) | n=163<br>med:0.115<br>(0.100-0.128) | n=247<br>med:0.086<br>(0.071-0.099) | n=190<br>med:0.087<br>(0.067-0.098) | n=278<br>med:0.075<br>(0.065-0.084) |
| Brazilian guinea pig | n=214<br>med:0.136<br>(0.122-0.142) | n=204<br>med:0.137<br>(0.130-0.152) | n=181<br>med:0.122<br>(0.106-0.134) | n=89<br>med:0.107<br>(0.088-0.126) | n=247<br>med:0.112<br>(0.102-0.120) | n=123<br>med:0.103<br>(0.093-0.115) | n=181<br>med:0.091<br>(0.076-0.103) | n=136<br>med:0.098<br>(0.084-0.110) | n=197<br>med:0.094<br>(0.083-0.104) |
| Orangutan | n=297<br>med:0.143<br>(0.128-0.155) | n=288<br>med:0.139<br>(0.123-0.156) | n=253<br>med:0.110<br>(0.091-0.132) | n=113<br>med:0.111<br>(0.090-0.123) | n=309<br>med:0.095<br>(0.084-0.109) | n=176<br>med:0.113<br>(0.095-0.132) | n=251<br>med:0.076<br>(0.065-0.093) | n=194<br>med:0.077<br>(0.062-0.097) | n=274<br>med:0.062<br>(0.051-0.073) |
| Northern American deer mouse | n=333<br>med:0.153<br>(0.137-0.168) | n=305<br>med:0.161<br>(0.141-0.181) | n=259<br>med:0.115<br>(0.098-0.127) | n=117<br>med:0.097<br>(0.085-0.123) | n=331<br>med:0.099<br>(0.085-0.116) | n=178<br>med:0.108<br>(0.093-0.126) | n=258<br>med:0.068<br>(0.057-0.079) | n=212<br>med:0.069<br>(0.058-0.092) | n=283<br>med:0.065<br>(0.058-0.087) |
| Naked mole-rat female | n=299<br>med:0.150<br>(0.133-0.168) | n=255<br>med:0.139<br>(0.120-0.150) | n=230<br>med:0.110<br>(0.096-0.127) | n=107<br>med:0.108<br>(0.088-0.127) | n=289<br>med:0.100<br>(0.086-0.113) | n=161<br>med:0.097<br>(0.088-0.113) | n=243<br>med:0.080<br>(0.068-0.088) | n=198<br>med:0.076<br>(0.057-0.094) | n=258<br>med:0.065<br>(0.054-0.078) |
| Mouse | n=362<br>med:0.162<br>(0.143-0.191) | n=337<br>med:0.170<br>(0.147-0.195) | n=283<br>med:0.119<br>(0.102-0.134) | n=125<br>med:0.115<br>(0.090-0.131) | n=359<br>med:0.103<br>(0.090-0.119) | n=192<br>med:0.107<br>(0.090-0.121) | n=292<br>med:0.079<br>(0.064-0.091) | n=226<br>med:0.080<br>(0.059-0.089) | n=319<br>med:0.070<br>(0.060-0.083) |
| Daurian ground squirrel | n=273<br>med:0.147<br>(0.129-0.158) | n=248<br>med:0.141<br>(0.126-0.162) | n=223<br>med:0.120<br>(0.103-0.134) | n=100<br>med:0.103<br>(0.078-0.120) | n=279<br>med:0.110<br>(0.095-0.122) | n=147<br>med:0.104<br>(0.088-0.120) | n=206<br>med:0.081<br>(0.070-0.095) | n=164<br>med:0.083<br>(0.060-0.101) | n=212<br>med:0.072<br>(0.056-0.084) |
| Tiger | n=281<br>med:0.138<br>(0.121-0.162) | n=240<br>med:0.141<br>(0.121-0.167) | n=198<br>med:0.119<br>(0.097-0.136) | n=93<br>med:0.111<br>(0.090-0.135) | n=250<br>med:0.100<br>(0.087-0.113) | n=139<br>med:0.116<br>(0.090-0.126) | n=198<br>med:0.087<br>(0.073-0.103) | n=157<br>med:0.083<br>(0.067-0.099) | n=218<br>med:0.071<br>(0.057-0.077) |
| Lesser hedgehog tenrec | n=70<br>med:0.140<br>(0.107-0.194) | n=68<br>med:0.144<br>(0.111-0.196) | n=64<br>med:0.133<br>(0.098-0.169) | n=23<br>med:0.080<br>(0.056-0.118) | n=80<br>med:0.080<br>(0.066-0.116) | n=36<br>med:0.124<br>(0.089-0.149) | n=57<br>med:0.097<br>(0.068-0.131) | n=33<br>med:0.085<br>(0.060-0.107) | n=58<br>med:0.077<br>(0.057-0.111) |
| Mongolian gerbil | n=301<br>med:0.182<br>(0.153-0.202) | n=302<br>med:0.174<br>(0.155-0.202) | n=238<br>med:0.118<br>(0.103-0.133) | n=106<br>med:0.106<br>(0.081-0.139) | n=315<br>med:0.107<br>(0.095-0.119) | n=166<br>med:0.109<br>(0.090-0.125) | n=237<br>med:0.082<br>(0.065-0.106) | n=185<br>med:0.084<br>(0.066-0.115) | n=260<br>med:0.073<br>(0.059-0.089) |
| Ma's night monkey | n=308<br>med:0.155<br>(0.137-0.175) | n=278<br>med:0.145<br>(0.134-0.158) | n=252<br>med:0.117<br>(0.099-0.138) | n=107<br>med:0.102<br>(0.086-0.122) | n=304<br>med:0.100<br>(0.090-0.111) | n=171<br>med:0.112<br>(0.095-0.129) | n=250<br>med:0.091<br>(0.072-0.102) | n=200<br>med:0.080<br>(0.065-0.094) | n=278<br>med:0.066<br>(0.059-0.082) |
| Upper Galilee mountains blind mole rat | n=314<br>med:0.146<br>(0.132-0.161) | n=289<br>med:0.148<br>(0.131-0.166) | n=242<br>med:0.110<br>(0.092-0.128) | n=113<br>med:0.114<br>(0.081-0.135) | n=324<br>med:0.101<br>(0.088-0.115) | n=170<br>med:0.100<br>(0.079-0.119) | n=260<br>med:0.072<br>(0.057-0.083) | n=203<br>med:0.075<br>(0.057-0.097) | n=280<br>med:0.066<br>(0.054-0.077) |
| Kangaroo rat | n=272<br>med:0.138<br>(0.126-0.154) | n=241<br>med:0.126<br>(0.105-0.142) | n=220<br>med:0.103<br>(0.088-0.120) | n=97<br>med:0.096<br>(0.082-0.117) | n=279<br>med:0.092<br>(0.080-0.100) | n=162<br>med:0.098<br>(0.079-0.113) | n=233<br>med:0.069<br>(0.060-0.085) | n=179<br>med:0.078<br>(0.061-0.093) | n=247<br>med:0.064<br>(0.054-0.074) |
| Chimpanzee | n=309<br>med:0.153<br>(0.137-0.175) | n=272<br>med:0.143<br>(0.129-0.157) | n=244<br>med:0.113<br>(0.097-0.134) | n=109<br>med:0.105<br>(0.071-0.116) | n=301<br>med:0.109<br>(0.097-0.118) | n=172<br>med:0.109<br>(0.098-0.124) | n=243<br>med:0.083<br>(0.066-0.099) | n=196<br>med:0.076<br>(0.066-0.093) | n=279<br>med:0.066<br>(0.056-0.078) |
| Guinea Pig | n=296<br>med:0.136<br>(0.126-0.154) | n=267<br>med:0.135<br>(0.122-0.149) | n=244<br>med:0.109<br>(0.093-0.119) | n=107<br>med:0.108<br>(0.087-0.124) | n=310<br>med:0.097<br>(0.084-0.106) | n=162<br>med:0.095<br>(0.084-0.112) | n=250<br>med:0.076<br>(0.066-0.089) | n=195<br>med:0.083<br>(0.067-0.092) | n=276<br>med:0.064<br>(0.055-0.073) |
| American beaver | n=275<br>med:0.149<br>(0.134-0.161) | n=267<br>med:0.145<br>(0.127-0.171) | n=243<br>med:0.125<br>(0.103-0.144) | n=97<br>med:0.107<br>(0.082-0.122) | n=295<br>med:0.110<br>(0.097-0.125) | n=159<br>med:0.101<br>(0.079-0.120) | n=220<br>med:0.077<br>(0.065-0.098) | n=174<br>med:0.088<br>(0.062-0.101) | n=250<br>med:0.073<br>(0.060-0.086) |
| Wild yak | n=296<br>med:0.138<br>(0.119-0.155) | n=265<br>med:0.140<br>(0.117-0.156) | n=219<br>med:0.109<br>(0.098-0.131) | n=98<br>med:0.082<br>(0.070-0.106) | n=275<br>med:0.103<br>(0.087-0.112) | n=152<br>med:0.099<br>(0.085-0.111) | n=206<br>med:0.086<br>(0.074-0.098) | n=175<br>med:0.071<br>(0.061-0.085) | n=221<br>med:0.076<br>(0.058-0.088) |
| Coquerel's sifaka | n=306 | n=263 | n=242 | n=106 | n=299 | n=165 | n=244 | n=195 | n=269 |

| | | | | | | | | | |
|---|---|---|---|---|---|---|---|---|---|
| | med:0.149 (0.132-0.160) | med:0.139 (0.125-0.157) | med:0.116 (0.100-0.134) | med:0.096 (0.081-0.107) | med:0.096 (0.087-0.108) | med:0.105 (0.083-0.121) | med:0.082 (0.064-0.094) | med:0.084 (0.067-0.095) | med:0.074 (0.055-0.082) |
| Alpaca | n=75 med:0.192 (0.145-0.229) | n=61 med:0.216 (0.170-0.244) | n=49 med:0.146 (0.107-0.214) | n=20 med:0.112 (0.077-0.120) | n=65 med:0.147 (0.123-0.176) | n=38 med:0.105 (0.090-0.154) | n=56 med:0.095 (0.071-0.106) | n=34 med:0.081 (0.057-0.124) | n=57 med:0.071 (0.045-0.106) |
| Lesser Egyptian jerboa | n=299 med:0.145 (0.126-0.157) | n=251 med:0.142 (0.127-0.154) | n=214 med:0.106 (0.087-0.128) | n=110 med:0.101 (0.081-0.119) | n=298 med:0.098 (0.088-0.108) | n=160 med:0.110 (0.083-0.125) | n=243 med:0.072 (0.060-0.083) | n=193 med:0.072 (0.057-0.085) | n=259 med:0.067 (0.059-0.082) |
| Long-tailed chinchilla | n=308 med:0.129 (0.114-0.143) | n=265 med:0.130 (0.117-0.148) | n=241 med:0.097 (0.089-0.111) | n=111 med:0.095 (0.073-0.109) | n=304 med:0.094 (0.085-0.106) | n=162 med:0.095 (0.077-0.106) | n=255 med:0.073 (0.061-0.080) | n=192 med:0.076 (0.060-0.088) | n=278 med:0.059 (0.051-0.074) |
| Angola colobus | n=312 med:0.148 (0.130-0.165) | n=286 med:0.147 (0.131-0.163) | n=252 med:0.112 (0.096-0.129) | n=107 med:0.108 (0.083-0.118) | n=310 med:0.100 (0.093-0.111) | n=172 med:0.105 (0.089-0.117) | n=245 med:0.085 (0.069-0.097) | n=198 med:0.077 (0.063-0.091) | n=275 med:0.068 (0.059-0.076) |
| Ryukyu mouse | n=342 med:0.160 (0.135-0.190) | n=322 med:0.174 (0.149-0.195) | n=272 med:0.121 (0.100-0.135) | n=119 med:0.118 (0.086-0.136) | n=341 med:0.105 (0.092-0.120) | n=187 med:0.116 (0.096-0.131) | n=279 med:0.078 (0.062-0.093) | n=220 med:0.075 (0.064-0.088) | n=311 med:0.070 (0.059-0.090) |
| Chinese hamster PICR | n=339 med:0.175 (0.150-0.199) | n=312 med:0.171 (0.150-0.198) | n=264 med:0.130 (0.113-0.142) | n=118 med:0.116 (0.085-0.128) | n=340 med:0.109 (0.097-0.125) | n=186 med:0.108 (0.094-0.133) | n=283 med:0.080 (0.071-0.096) | n=214 med:0.085 (0.064-0.103) | n=310 med:0.077 (0.065-0.093) |
| Vervet-AGM | n=331 med:0.143 (0.123-0.162) | n=283 med:0.144 (0.130-0.153) | n=257 med:0.105 (0.088-0.122) | n=108 med:0.108 (0.084-0.124) | n=323 med:0.099 (0.086-0.107) | n=180 med:0.102 (0.088-0.117) | n=262 med:0.075 (0.062-0.087) | n=201 med:0.076 (0.062-0.091) | n=285 med:0.062 (0.048-0.069) |
| Cow | n=327 med:0.138 (0.112-0.154) | n=288 med:0.142 (0.127-0.161) | n=247 med:0.109 (0.095-0.127) | n=106 med:0.090 (0.075-0.112) | n=308 med:0.096 (0.084-0.110) | n=170 med:0.101 (0.086-0.111) | n=241 med:0.077 (0.066-0.087) | n=192 med:0.075 (0.060-0.088) | n=282 med:0.069 (0.057-0.080) |
| Bushbaby | n=317 med:0.153 (0.135-0.172) | n=280 med:0.145 (0.130-0.166) | n=251 med:0.114 (0.097-0.135) | n=111 med:0.110 (0.090-0.131) | n=317 med:0.099 (0.091-0.111) | n=175 med:0.105 (0.087-0.121) | n=255 med:0.079 (0.066-0.092) | n=206 med:0.084 (0.070-0.092) | n=273 med:0.067 (0.058-0.076) |
| Bonobo | n=318 med:0.150 (0.134-0.170) | n=283 med:0.145 (0.132-0.158) | n=249 med:0.121 (0.102-0.136) | n=110 med:0.108 (0.088-0.117) | n=316 med:0.101 (0.090-0.113) | n=176 med:0.117 (0.098-0.130) | n=243 med:0.082 (0.070-0.101) | n=195 med:0.079 (0.066-0.091) | n=274 med:0.072 (0.061-0.085) |
| Hyrax | n=80 med:0.168 (0.104-0.198) | n=96 med:0.169 (0.129-0.206) | n=72 med:0.110 (0.082-0.146) | n=30 med:0.120 (0.084-0.159) | n=103 med:0.114 (0.092-0.131) | n=47 med:0.094 (0.069-0.143) | n=68 med:0.088 (0.066-0.113) | n=61 med:0.111 (0.090-0.133) | n=66 med:0.083 (0.062-0.104) |
| Rabbit | n=287 med:0.136 (0.120-0.158) | n=256 med:0.133 (0.116-0.146) | n=225 med:0.107 (0.092-0.121) | n=103 med:0.091 (0.075-0.102) | n=284 med:0.102 (0.087-0.111) | n=160 med:0.095 (0.076-0.105) | n=227 med:0.083 (0.067-0.092) | n=178 med:0.083 (0.070-0.094) | n=255 med:0.072 (0.059-0.085) |
| Megabat | n=171 med:0.156 (0.135-0.174) | n=165 med:0.154 (0.138-0.183) | n=146 med:0.129 (0.108-0.145) | n=71 med:0.114 (0.101-0.138) | n=172 med:0.092 (0.078-0.107) | n=100 med:0.122 (0.100-0.144) | n=119 med:0.100 (0.086-0.120) | n=98 med:0.100 (0.081-0.119) | n=145 med:0.080 (0.065-0.097) |
| Dog | n=300 med:0.148 (0.129-0.165) | n=277 med:0.136 (0.122-0.148) | n=248 med:0.111 (0.099-0.133) | n=107 med:0.104 (0.086-0.120) | n=315 med:0.099 (0.086-0.107) | n=166 med:0.104 (0.087-0.122) | n=249 med:0.080 (0.067-0.094) | n=193 med:0.089 (0.073-0.103) | n=269 med:0.071 (0.059-0.078) |
| Steppe mouse | n=332 med:0.181 (0.150-0.209) | n=328 med:0.182 (0.162-0.212) | n=272 med:0.127 (0.111-0.153) | n=118 med:0.121 (0.094-0.148) | n=337 med:0.128 (0.111-0.142) | n=180 med:0.125 (0.104-0.155) | n=269 med:0.089 (0.070-0.102) | n=219 med:0.088 (0.076-0.100) | n=306 med:0.087 (0.068-0.098) |
| Greater bamboo lemur | n=319 med:0.147 (0.127-0.161) | n=283 med:0.137 (0.120-0.157) | n=248 med:0.108 (0.101-0.122) | n=110 med:0.096 (0.080-0.114) | n=318 med:0.099 (0.087-0.107) | n=176 med:0.107 (0.092-0.120) | n=256 med:0.084 (0.068-0.095) | n=198 med:0.078 (0.062-0.092) | n=280 med:0.067 (0.056-0.079) |
| Pig | n=311 med:0.143 (0.124-0.158) | n=285 med:0.139 (0.129-0.153) | n=254 med:0.105 (0.092-0.116) | n=109 med:0.090 (0.073-0.106) | n=322 med:0.097 (0.087-0.106) | n=179 med:0.109 (0.096-0.119) | n=260 med:0.078 (0.068-0.086) | n=200 med:0.077 (0.063-0.084) | n=280 med:0.066 (0.057-0.074) |
| Tree Shrew | n=62 med:0.173 (0.123-0.219) | n=58 med:0.160 (0.125-0.228) | n=49 med:0.147 (0.099-0.160) | n=20 med:0.080 (0.057-0.109) | n=53 med:0.109 (0.082-0.141) | n=38 med:0.099 (0.062-0.134) | n=52 med:0.123 (0.081-0.158) | n=40 med:0.104 (0.067-0.134) | n=47 med:0.095 (0.069-0.143) |
| Pika | n=105 med:0.146 (0.111-0.184) | n=112 med:0.134 (0.117-0.167) | n=100 med:0.114 (0.092-0.133) | n=39 med:0.086 (0.055-0.120) | n=109 med:0.092 (0.073-0.129) | n=61 med:0.115 (0.085-0.136) | n=73 med:0.087 (0.067-0.110) | n=57 med:0.092 (0.067-0.126) | n=79 med:0.070 (0.058-0.094) |
| Shrew | n=48 med:0.140 (0.107-0.216) | n=59 med:0.127 (0.094-0.178) | n=36 med:0.120 (0.070-0.161) | n=16 med:0.087 (0.029-0.129) | n=58 med:0.109 (0.075-0.138) | n=21 med:0.097 (0.073-0.149) | n=37 med:0.067 (0.050-0.122) | n=35 med:0.082 (0.038-0.133) | n=47 med:0.085 (0.043-0.111) |
| Squirrel | n=305 med:0.138 (0.124-0.158) | n=273 med:0.128 (0.113-0.147) | n=254 med:0.107 (0.090-0.123) | n=113 med:0.090 (0.074-0.107) | n=316 med:0.097 (0.086-0.111) | n=177 med:0.094 (0.076-0.111) | n=261 med:0.067 (0.057-0.074) | n=205 med:0.074 (0.056-0.089) | n=284 med:0.058 (0.051-0.068) |
| Gorilla | n=317 med:0.152 (0.138-0.171) | n=280 med:0.142 (0.130-0.158) | n=248 med:0.110 (0.095-0.136) | n=109 med:0.105 (0.085-0.122) | n=315 med:0.102 (0.088-0.113) | n=169 med:0.114 (0.094-0.122) | n=246 med:0.085 (0.075-0.104) | n=199 med:0.074 (0.062-0.089) | n=284 med:0.069 (0.058-0.081) |
| Cat | n=326 med:0.141 (0.126-0.159) | n=292 med:0.131 (0.117-0.147) | n=254 med:0.119 (0.098-0.133) | n=112 med:0.093 (0.083-0.113) | n=323 med:0.095 (0.083-0.106) | n=177 med:0.106 (0.093-0.121) | n=258 med:0.078 (0.067-0.089) | n=207 med:0.080 (0.065-0.092) | n=291 med:0.068 (0.059-0.073) |

#### Rat Reference Genome, Saunders et al., 2018, three cell types

| Sample Size and Median dN/dS (CI95%) | Vasculature-specific Genes | Glia-specific Genes | Neuron-specific Genes |
|---|---|---|---|
| Microbat | n=290 med:0.102 (0.093-0.118) | n=198 med:0.088 (0.072-0.098) | n=481 med:0.057 (0.051-0.061) |
| Hedgehog | n=78 med:0.139 (0.107-0.154) | n=54 med:0.092 (0.069-0.117) | n=109 med:0.059 (0.046-0.077) |
| Naked mole-rat male | n=302 med:0.101 (0.090-0.109) | n=217 med:0.090 (0.074-0.099) | n=516 med:0.050 (0.045-0.054) |
| Sooty mangabey | n=340 med:0.112 (0.096-0.124) | n=249 med:0.081 (0.070-0.091) | n=575 med:0.052 (0.048-0.058) |
| Olive baboon | n=341 med:0.109 (0.094-0.123) | n=247 med:0.084 (0.073-0.097) | n=581 med:0.052 (0.047-0.056) |
| American bison | n=293 med:0.099 (0.089-0.118) | n=195 med:0.084 (0.079-0.096) | n=462 med:0.049 (0.044-0.054) |
| Leopard | n=341 med:0.096 (0.085-0.105) | n=243 med:0.074 (0.064-0.083) | n=557 med:0.047 (0.042-0.051) |
| Bolivian squirrel monkey | n=330 med:0.109 (0.101-0.124) | n=243 med:0.085 (0.072-0.095) | n=560 med:0.052 (0.048-0.056) |
| Crab-eating macaque | n=339 med:0.104 (0.094-0.113) | n=243 med:0.081 (0.071-0.090) | n=571 med:0.053 (0.046-0.058) |
| Human | n=353 med:0.103 (0.091-0.112) | n=260 med:0.079 (0.064-0.087) | n=594 med:0.047 (0.043-0.051) |
| Red fox | n=321 med:0.107 (0.098-0.118) | n=229 med:0.092 (0.081-0.103) | n=544 med:0.058 (0.049-0.064) |
| Shrew mouse | n=378 med:0.115 (0.101-0.133) | n=279 med:0.092 (0.080-0.102) | n=613 med:0.051 (0.046-0.059) |
| Arctic ground squirrel | n=339 med:0.101 (0.088-0.111) | n=251 med:0.077 (0.067-0.085) | n=578 med:0.045 (0.041-0.049) |
| Chinese hamster CriGri | n=340 med:0.120 (0.108-0.136) | n=252 med:0.086 (0.077-0.108) | n=550 med:0.060 (0.053-0.065) |
| Opossum | n=312 | n=225 | n=508 |

| | | | |
|---|---|---|---|
| | med:0.076<br>(0.068-0.087) | med:0.070<br>(0.059-0.078) | med:0.051<br>(0.047-0.058) |
| Dolphin | n=195<br>med:0.110<br>(0.098-0.125) | n=122<br>med:0.103<br>(0.084-0.113) | n=298<br>med:0.054<br>(0.046-0.062) |
| Capuchin | n=330<br>med:0.109<br>(0.098-0.124) | n=249<br>med:0.080<br>(0.070-0.089) | n=568<br>med:0.051<br>(0.045-0.055) |
| Mouse Lemur | n=339<br>med:0.100<br>(0.089-0.112) | n=247<br>med:0.075<br>(0.066-0.087) | n=567<br>med:0.050<br>(0.045-0.056) |
| Damara mole rat | n=308<br>med:0.099<br>(0.092-0.109) | n=220<br>med:0.077<br>(0.064-0.089) | n=551<br>med:0.046<br>(0.041-0.055) |
| Marmoset | n=333<br>med:0.111<br>(0.101-0.124) | n=239<br>med:0.082<br>(0.071-0.093) | n=574<br>med:0.051<br>(0.046-0.055) |
| Goat | n=337<br>med:0.101<br>(0.088-0.114) | n=241<br>med:0.080<br>(0.067-0.086) | n=563<br>med:0.050<br>(0.045-0.055) |
| Common wombat | n=304<br>med:0.082<br>(0.072-0.089) | n=210<br>med:0.069<br>(0.060-0.079) | n=494<br>med:0.048<br>(0.043-0.054) |
| American black bear | n=304<br>med:0.104<br>(0.090-0.116) | n=224<br>med:0.083<br>(0.067-0.094) | n=471<br>med:0.058<br>(0.051-0.067) |
| Sloth | n=47<br>med:0.157<br>(0.099-0.209) | n=34<br>med:0.148<br>(0.088-0.248) | n=65<br>med:0.074<br>(0.046-0.097) |
| Polar bear | n=303<br>med:0.113<br>(0.102-0.124) | n=219<br>med:0.094<br>(0.081-0.109) | n=501<br>med:0.063<br>(0.058-0.071) |
| Alpine marmot | n=329<br>med:0.103<br>(0.093-0.114) | n=233<br>med:0.081<br>(0.071-0.094) | n=530<br>med:0.047<br>(0.042-0.054) |
| Prairie vole | n=351<br>med:0.100<br>(0.086-0.113) | n=270<br>med:0.085<br>(0.074-0.098) | n=597<br>med:0.044<br>(0.039-0.049) |
| Donkey | n=339<br>med:0.101<br>(0.092-0.114) | n=257<br>med:0.086<br>(0.075-0.095) | n=565<br>med:0.055<br>(0.051-0.061) |
| Golden Hamster | n=339<br>med:0.122<br>(0.099-0.134) | n=256<br>med:0.095<br>(0.084-0.107) | n=561<br>med:0.056<br>(0.050-0.061) |
| Armadillo | n=310<br>med:0.108<br>(0.095-0.120) | n=221<br>med:0.081<br>(0.070-0.099) | n=516<br>med:0.060<br>(0.054-0.066) |
| Ugandan red Colobus | n=335 | n=248 | n=559 |

| | | | |
|---|---|---|---|
| | med:0.106<br>(0.093-0.118) | med:0.079<br>(0.069-0.091) | med:0.050<br>(0.045-0.055) |
| Pig-tailed macaque | n=341<br>med:0.112<br>(0.093-0.123) | n=249<br>med:0.082<br>(0.070-0.093) | n=581<br>med:0.052<br>(0.047-0.058) |
| Panda | n=332<br>med:0.101<br>(0.092-0.116) | n=246<br>med:0.081<br>(0.070-0.091) | n=542<br>med:0.049<br>(0.044-0.053) |
| Ferret | n=323<br>med:0.095<br>(0.085-0.105) | n=233<br>med:0.071<br>(0.063-0.081) | n=554<br>med:0.046<br>(0.041-0.052) |
| Koala | n=317<br>med:0.080<br>(0.071-0.093) | n=240<br>med:0.077<br>(0.068-0.084) | n=537<br>med:0.050<br>(0.044-0.055) |
| Wallaby | n=92<br>med:0.071<br>(0.052-0.077) | n=89<br>med:0.070<br>(0.049-0.085) | n=131<br>med:0.052<br>(0.042-0.060) |
| Tarsier | n=288<br>med:0.110<br>(0.101-0.128) | n=215<br>med:0.089<br>(0.076-0.098) | n=465<br>med:0.052<br>(0.045-0.059) |
| Degu | n=282<br>med:0.100<br>(0.086-0.108) | n=204<br>med:0.081<br>(0.066-0.088) | n=535<br>med:0.047<br>(0.042-0.050) |
| Chinese hamster CHOK1GS | n=377<br>med:0.120<br>(0.102-0.131) | n=274<br>med:0.086<br>(0.076-0.095) | n=612<br>med:0.051<br>(0.044-0.056) |
| Drill | n=326<br>med:0.110<br>(0.099-0.123) | n=232<br>med:0.088<br>(0.077-0.099) | n=548<br>med:0.053<br>(0.047-0.060) |
| Gibbon | n=320<br>med:0.114<br>(0.103-0.125) | n=236<br>med:0.089<br>(0.079-0.101) | n=560<br>med:0.058<br>(0.053-0.065) |
| Macaque | n=346<br>med:0.113<br>(0.097-0.129) | n=257<br>med:0.083<br>(0.072-0.095) | n=587<br>med:0.050<br>(0.046-0.055) |
| American mink | n=334<br>med:0.096<br>(0.085-0.107) | n=240<br>med:0.075<br>(0.065-0.085) | n=537<br>med:0.047<br>(0.042-0.053) |
| Gelada | n=338<br>med:0.105<br>(0.093-0.115) | n=253<br>med:0.081<br>(0.069-0.092) | n=580<br>med:0.049<br>(0.044-0.054) |
| Sheep | n=318<br>med:0.108<br>(0.097-0.124) | n=226<br>med:0.084<br>(0.074-0.092) | n=546<br>med:0.055<br>(0.050-0.059) |
| Tasmanian devil | n=291<br>med:0.076<br>(0.067-0.088) | n=217<br>med:0.067<br>(0.056-0.076) | n=492<br>med:0.045<br>(0.042-0.051) |
| Elephant | n=333 | n=247 | n=550 |

| | | | |
|---|---|---|---|
| | med:0.106 (0.094-0.116) | med:0.083 (0.070-0.095) | med:0.054 (0.048-0.059) |
| Algerian mouse | n=377 med:0.122 (0.097-0.138) | n=277 med:0.089 (0.075-0.107) | n=613 med:0.052 (0.044-0.058) |
| Dingo | n=324 med:0.102 (0.090-0.110) | n=240 med:0.082 (0.070-0.090) | n=558 med:0.054 (0.045-0.058) |
| Golden snub-nosed monkey | n=327 med:0.105 (0.095-0.116) | n=232 med:0.081 (0.072-0.099) | n=546 med:0.052 (0.048-0.056) |
| Horse | n=341 med:0.107 (0.095-0.117) | n=244 med:0.086 (0.078-0.095) | n=583 med:0.055 (0.050-0.060) |
| Black snub-nosed monkey | n=331 med:0.112 (0.102-0.124) | n=242 med:0.083 (0.073-0.092) | n=548 med:0.056 (0.053-0.062) |
| Brazilian guinea pig | n=234 med:0.113 (0.096-0.125) | n=187 med:0.091 (0.081-0.101) | n=420 med:0.073 (0.068-0.080) |
| Orangutan | n=338 med:0.106 (0.095-0.120) | n=247 med:0.086 (0.074-0.103) | n=557 med:0.050 (0.045-0.054) |
| Northern American deer mouse | n=360 med:0.110 (0.095-0.123) | n=269 med:0.082 (0.072-0.099) | n=590 med:0.048 (0.043-0.055) |
| Naked mole-rat female | n=323 med:0.101 (0.092-0.115) | n=234 med:0.084 (0.071-0.091) | n=552 med:0.049 (0.044-0.053) |
| Mouse | n=391 med:0.111 (0.092-0.130) | n=285 med:0.090 (0.076-0.103) | n=631 med:0.050 (0.043-0.058) |
| Daurian ground squirrel | n=295 med:0.112 (0.097-0.123) | n=198 med:0.095 (0.082-0.106) | n=470 med:0.047 (0.043-0.055) |
| Tiger | n=280 med:0.105 (0.096-0.119) | n=190 med:0.093 (0.076-0.101) | n=452 med:0.052 (0.047-0.058) |
| Lesser hedgehog tenrec | n=77 med:0.116 (0.089-0.156) | n=68 med:0.086 (0.076-0.110) | n=103 med:0.066 (0.055-0.091) |
| Mongolian gerbil | n=339 med:0.118 (0.102-0.136) | n=246 med:0.077 (0.068-0.095) | n=513 med:0.054 (0.046-0.063) |
| Ma's night monkey | n=324 med:0.115 (0.105-0.130) | n=244 med:0.087 (0.074-0.093) | n=570 med:0.055 (0.049-0.059) |

| | | | |
|---|---|---|---|
| Upper Galilee mountains blind mole rat | n=338 med:0.110 (0.095-0.128) | n=260 med:0.079 (0.070-0.090) | n=589 med:0.049 (0.043-0.054) |
| Kangaroo rat | n=291 med:0.095 (0.081-0.106) | n=231 med:0.081 (0.072-0.089) | n=535 med:0.049 (0.044-0.055) |
| Chimpanzee | n=319 med:0.107 (0.096-0.121) | n=241 med:0.084 (0.074-0.099) | n=563 med:0.050 (0.045-0.057) |
| Guinea Pig | n=328 med:0.101 (0.091-0.110) | n=238 med:0.084 (0.071-0.095) | n=558 med:0.052 (0.046-0.056) |
| American beaver | n=330 med:0.107 (0.098-0.122) | n=239 med:0.084 (0.075-0.094) | n=473 med:0.056 (0.048-0.065) |
| Wild yak | n=304 med:0.103 (0.090-0.119) | n=205 med:0.083 (0.073-0.094) | n=491 med:0.048 (0.044-0.056) |
| Coquerel's sifaka | n=335 med:0.107 (0.096-0.121) | n=236 med:0.079 (0.069-0.091) | n=562 med:0.051 (0.046-0.056) |
| Alpaca | n=59 med:0.158 (0.120-0.200) | n=47 med:0.123 (0.079-0.156) | n=107 med:0.062 (0.046-0.069) |
| Lesser Egyptian jerboa | n=302 med:0.091 (0.080-0.106) | n=239 med:0.083 (0.075-0.094) | n=547 med:0.047 (0.042-0.051) |
| Long-tailed chinchilla | n=322 med:0.095 (0.086-0.105) | n=232 med:0.078 (0.068-0.091) | n=564 med:0.046 (0.040-0.051) |
| Angola colobus | n=328 med:0.110 (0.098-0.119) | n=246 med:0.084 (0.072-0.095) | n=552 med:0.050 (0.045-0.055) |
| Ryukyu mouse | n=376 med:0.111 (0.102-0.134) | n=279 med:0.090 (0.080-0.103) | n=610 med:0.052 (0.046-0.060) |
| Chinese hamster PICR | n=369 med:0.127 (0.113-0.139) | n=267 med:0.088 (0.079-0.099) | n=601 med:0.052 (0.046-0.057) |
| Vervet-AGM | n=345 med:0.100 (0.091-0.119) | n=260 med:0.073 (0.064-0.085) | n=582 med:0.046 (0.041-0.051) |
| Cow | n=330 med:0.098 (0.085-0.110) | n=241 med:0.075 (0.065-0.086) | n=561 med:0.048 (0.044-0.052) |
| Bushbaby | n=337 med:0.103 (0.095-0.115) | n=254 med:0.083 (0.075-0.093) | n=576 med:0.052 (0.047-0.058) |

| | | | |
|---|---|---|---|
| Bonobo | n=328 med:0.109 (0.101-0.123) | n=238 med:0.089 (0.078-0.100) | n=553 med:0.051 (0.044-0.057) |
| Hyrax | n=99 med:0.107 (0.084-0.126) | n=76 med:0.088 (0.064-0.109) | n=136 med:0.057 (0.048-0.069) |
| Rabbit | n=300 med:0.096 (0.088-0.110) | n=213 med:0.082 (0.070-0.093) | n=512 med:0.057 (0.051-0.065) |
| Megabat | n=173 med:0.121 (0.102-0.132) | n=134 med:0.086 (0.070-0.104) | n=267 med:0.061 (0.052-0.069) |
| Dog | n=331 med:0.108 (0.099-0.118) | n=242 med:0.086 (0.075-0.099) | n=560 med:0.053 (0.046-0.059) |
| Steppe mouse | n=357 med:0.122 (0.107-0.142) | n=272 med:0.105 (0.088-0.122) | n=593 med:0.060 (0.052-0.070) |
| Greater bamboo lemur | n=342 med:0.107 (0.096-0.118) | n=252 med:0.089 (0.076-0.099) | n=564 med:0.049 (0.045-0.055) |
| Pig | n=344 med:0.102 (0.092-0.112) | n=255 med:0.084 (0.072-0.096) | n=562 med:0.050 (0.046-0.053) |
| Tree Shrew | n=73 med:0.115 (0.096-0.147) | n=41 med:0.092 (0.065-0.112) | n=79 med:0.088 (0.060-0.107) |
| Pika | n=124 med:0.097 (0.090-0.116) | n=90 med:0.082 (0.065-0.099) | n=146 med:0.058 (0.041-0.070) |
| Shrew | n=48 med:0.127 (0.090-0.188) | n=35 med:0.059 (0.041-0.102) | n=87 med:0.048 (0.034-0.063) |
| Squirrel | n=333 med:0.101 (0.084-0.111) | n=244 med:0.076 (0.066-0.086) | n=572 med:0.047 (0.042-0.052) |
| Gorilla | n=325 med:0.111 (0.101-0.126) | n=243 med:0.084 (0.076-0.101) | n=566 med:0.051 (0.045-0.057) |
| Cat | n=345 med:0.103 (0.094-0.114) | n=248 med:0.080 (0.071-0.091) | n=575 med:0.051 (0.047-0.055) |

#### Rat Reference Genome, Saunders et al., 2018, cell subtypes

| Sample Size and Median dN/dS (CI95%) | Endothelia-specific Genes | Microglia-specific Genes | Astrocyte-specific Genes | Oligodendrocyte-specific Genes | Neuron-specific Genes |
|---|---|---|---|---|---|
| Microbat | n=260 med:0.103 (0.095-0.119) | n=193 med:0.120 (0.107-0.139) | n=185 med:0.088 (0.070-0.102) | n=182 med:0.079 (0.068-0.094) | n=441 med:0.058 (0.054-0.062) |
| Hedgehog | n=71 med:0.139 (0.111-0.154) | n=43 med:0.128 (0.096-0.141) | n=43 med:0.081 (0.054-0.100) | n=50 med:0.092 (0.060-0.110) | n=102 med:0.062 (0.047-0.077) |
| Naked mole-rat male | n=272 med:0.101 (0.090-0.111) | n=199 med:0.117 (0.102-0.145) | n=212 med:0.080 (0.070-0.095) | n=191 med:0.086 (0.071-0.099) | n=479 med:0.051 (0.047-0.058) |
| Sooty mangabey | n=303 med:0.106 (0.094-0.122) | n=225 med:0.128 (0.111-0.148) | n=243 med:0.083 (0.071-0.095) | n=224 med:0.082 (0.070-0.093) | n=530 med:0.054 (0.049-0.059) |
| Olive baboon | n=303 med:0.104 (0.092-0.123) | n=226 med:0.135 (0.113-0.151) | n=236 med:0.084 (0.072-0.095) | n=221 med:0.082 (0.069-0.096) | n=536 med:0.055 (0.049-0.059) |
| American bison | n=257 med:0.102 (0.088-0.121) | n=190 med:0.122 (0.099-0.135) | n=182 med:0.084 (0.075-0.089) | n=180 med:0.084 (0.077-0.097) | n=425 med:0.051 (0.045-0.058) |
| Leopard | n=297 med:0.096 (0.085-0.104) | n=210 med:0.115 (0.100-0.136) | n=234 med:0.074 (0.064-0.078) | n=213 med:0.073 (0.063-0.081) | n=516 med:0.049 (0.044-0.052) |
| Bolivian squirrel monkey | n=294 med:0.109 (0.098-0.126) | n=205 med:0.131 (0.110-0.151) | n=223 med:0.083 (0.070-0.105) | n=227 med:0.076 (0.069-0.089) | n=518 med:0.053 (0.049-0.058) |
| Crab-eating macaque | n=303 med:0.102 (0.092-0.114) | n=226 med:0.129 (0.114-0.148) | n=236 med:0.084 (0.072-0.096) | n=220 med:0.083 (0.070-0.094) | n=528 med:0.055 (0.049-0.060) |
| Human | n=318 med:0.100 (0.091-0.112) | n=230 med:0.125 (0.111-0.140) | n=243 med:0.074 (0.066-0.086) | n=231 med:0.078 (0.063-0.087) | n=549 med:0.049 (0.045-0.054) |
| Red fox | n=287 med:0.105 (0.098-0.117) | n=215 med:0.137 (0.108-0.149) | n=207 med:0.094 (0.078-0.102) | n=213 med:0.086 (0.074-0.101) | n=503 med:0.060 (0.054-0.067) |
| Shrew mouse | n=335 med:0.112 (0.099-0.123) | n=247 med:0.151 (0.128-0.170) | n=255 med:0.090 (0.075-0.101) | n=245 med:0.092 (0.076-0.101) | n=568 med:0.054 (0.047-0.061) |
| Arctic ground squirrel | n=304 med:0.101 (0.088-0.111) | n=222 med:0.109 (0.093-0.135) | n=233 med:0.077 (0.068-0.089) | n=222 med:0.074 (0.065-0.084) | n=535 med:0.046 (0.042-0.051) |
| Chinese hamster CriGri | n=307 med:0.123 (0.108-0.140) | n=219 med:0.154 (0.121-0.177) | n=228 med:0.093 (0.085-0.102) | n=228 med:0.087 (0.074-0.107) | n=508 med:0.061 (0.056-0.067) |
| Opossum | n=278 med:0.078 (0.070-0.091) | n=200 med:0.090 (0.074-0.107) | n=214 med:0.069 (0.060-0.081) | n=204 med:0.064 (0.052-0.072) | n=469 med:0.052 (0.048-0.059) |
| Dolphin | n=180 med:0.113 (0.104-0.129) | n=118 med:0.126 (0.101-0.146) | n=117 med:0.098 (0.079-0.111) | n=109 med:0.103 (0.082-0.113) | n=276 med:0.055 (0.047-0.063) |
| Capuchin | n=302 med:0.107 (0.096-0.126) | n=210 med:0.128 (0.109-0.149) | n=229 med:0.082 (0.072-0.096) | n=227 med:0.078 (0.067-0.091) | n=523 med:0.052 (0.046-0.057) |
| Mouse Lemur | n=300 med:0.102 | n=213 med:0.115 | n=224 med:0.080 | n=223 med:0.072 | n=525 med:0.052 |

| | | | | | |
|---|---|---|---|---|---|
| | (0.089-0.114) | (0.100-0.141) | (0.071-0.085) | (0.064-0.082) | (0.047-0.058) |
| Damara mole rat | n=274<br>med:0.101<br>(0.093-0.112) | n=195<br>med:0.112<br>(0.097-0.136) | n=212<br>med:0.086<br>(0.072-0.097) | n=204<br>med:0.073<br>(0.060-0.082) | n=509<br>med:0.053<br>(0.042-0.057) |
| Marmoset | n=297<br>med:0.114<br>(0.102-0.131) | n=214<br>med:0.123<br>(0.113-0.145) | n=231<br>med:0.086<br>(0.071-0.100) | n=212<br>med:0.080<br>(0.067-0.093) | n=530<br>med:0.052<br>(0.047-0.057) |
| Goat | n=301<br>med:0.105<br>(0.093-0.116) | n=220<br>med:0.120<br>(0.096-0.135) | n=224<br>med:0.076<br>(0.068-0.086) | n=217<br>med:0.077<br>(0.067-0.086) | n=520<br>med:0.052<br>(0.047-0.057) |
| Common wombat | n=268<br>med:0.085<br>(0.077-0.093) | n=198<br>med:0.089<br>(0.075-0.103) | n=199<br>med:0.077<br>(0.067-0.087) | n=192<br>med:0.068<br>(0.058-0.075) | n=453<br>med:0.050<br>(0.043-0.054) |
| American black bear | n=269<br>med:0.110<br>(0.095-0.122) | n=193<br>med:0.117<br>(0.101-0.136) | n=195<br>med:0.087<br>(0.072-0.100) | n=202<br>med:0.081<br>(0.067-0.093) | n=435<br>med:0.061<br>(0.053-0.070) |
| Sloth | n=39<br>med:0.150<br>(0.099-0.195) | n=31<br>med:0.126<br>(0.084-0.157) | n=20<br>med:0.140<br>(0.081-0.228) | n=29<br>med:0.136<br>(0.077-0.208) | n=59<br>med:0.062<br>(0.046-0.086) |
| Polar bear | n=262<br>med:0.113<br>(0.102-0.128) | n=193<br>med:0.137<br>(0.115-0.163) | n=203<br>med:0.093<br>(0.084-0.108) | n=201<br>med:0.086<br>(0.075-0.102) | n=458<br>med:0.067<br>(0.059-0.073) |
| Alpine marmot | n=289<br>med:0.103<br>(0.091-0.114) | n=212<br>med:0.114<br>(0.098-0.137) | n=216<br>med:0.080<br>(0.069-0.093) | n=205<br>med:0.080<br>(0.066-0.093) | n=493<br>med:0.048<br>(0.043-0.055) |
| Prairie vole | n=314<br>med:0.102<br>(0.086-0.114) | n=230<br>med:0.133<br>(0.112-0.159) | n=250<br>med:0.090<br>(0.072-0.097) | n=238<br>med:0.077<br>(0.067-0.088) | n=554<br>med:0.045<br>(0.041-0.052) |
| Donkey | n=301<br>med:0.108<br>(0.096-0.117) | n=225<br>med:0.128<br>(0.108-0.139) | n=232<br>med:0.087<br>(0.078-0.100) | n=224<br>med:0.078<br>(0.068-0.095) | n=526<br>med:0.056<br>(0.052-0.062) |
| Golden Hamster | n=302<br>med:0.124<br>(0.106-0.139) | n=221<br>med:0.151<br>(0.123-0.172) | n=242<br>med:0.094<br>(0.083-0.109) | n=224<br>med:0.089<br>(0.077-0.099) | n=527<br>med:0.058<br>(0.051-0.063) |
| Armadillo | n=272<br>med:0.107<br>(0.097-0.117) | n=206<br>med:0.124<br>(0.105-0.136) | n=206<br>med:0.081<br>(0.069-0.096) | n=204<br>med:0.079<br>(0.073-0.087) | n=477<br>med:0.061<br>(0.055-0.067) |
| Ugandan red Colobus | n=302<br>med:0.108<br>(0.093-0.126) | n=221<br>med:0.129<br>(0.116-0.149) | n=226<br>med:0.079<br>(0.070-0.093) | n=222<br>med:0.083<br>(0.069-0.093) | n=515<br>med:0.051<br>(0.046-0.057) |
| Pig-tailed macaque | n=300<br>med:0.111<br>(0.092-0.123) | n=224<br>med:0.131<br>(0.115-0.150) | n=235<br>med:0.083<br>(0.072-0.097) | n=223<br>med:0.083<br>(0.069-0.095) | n=537<br>med:0.054<br>(0.049-0.060) |
| Panda | n=293<br>med:0.103<br>(0.093-0.116) | n=215<br>med:0.117<br>(0.100-0.141) | n=222<br>med:0.077<br>(0.065-0.088) | n=225<br>med:0.073<br>(0.066-0.084) | n=501<br>med:0.050<br>(0.044-0.055) |
| Ferret | n=287<br>med:0.098<br>(0.091-0.110) | n=216<br>med:0.118<br>(0.100-0.132) | n=216<br>med:0.077<br>(0.062-0.088) | n=215<br>med:0.071<br>(0.063-0.080) | n=509<br>med:0.049<br>(0.043-0.055) |
| Koala | n=282<br>med:0.084<br>(0.075-0.094) | n=211<br>med:0.087<br>(0.075-0.108) | n=217<br>med:0.075<br>(0.065-0.086) | n=220<br>med:0.076<br>(0.067-0.089) | n=499<br>med:0.051<br>(0.045-0.057) |
| Wallaby | n=75<br>med:0.075<br>(0.059-0.084) | n=58<br>med:0.079<br>(0.068-0.105) | n=62<br>med:0.061<br>(0.037-0.093) | n=73<br>med:0.074<br>(0.059-0.086) | n=116<br>med:0.052<br>(0.041-0.060) |
| Tarsier | n=252<br>med:0.111<br>(0.102-0.129) | n=183<br>med:0.127<br>(0.108-0.145) | n=192<br>med:0.092<br>(0.072-0.101) | n=192<br>med:0.082<br>(0.072-0.097) | n=432<br>med:0.054<br>(0.046-0.063) |

| | | | | | |
|---|---|---|---|---|---|
| Degu | n=249 med:0.098 (0.084-0.109) | n=182 med:0.119 (0.096-0.137) | n=207 med:0.075 (0.060-0.084) | n=187 med:0.076 (0.063-0.087) | n=497 med:0.047 (0.043-0.052) |
| Chinese hamster CHOK1GS | n=337 med:0.119 (0.100-0.130) | n=239 med:0.135 (0.117-0.171) | n=255 med:0.088 (0.075-0.096) | n=244 med:0.082 (0.070-0.099) | n=569 med:0.051 (0.044-0.057) |
| Drill | n=288 med:0.110 (0.098-0.126) | n=219 med:0.132 (0.120-0.149) | n=230 med:0.086 (0.074-0.098) | n=215 med:0.085 (0.069-0.098) | n=505 med:0.056 (0.048-0.062) |
| Gibbon | n=283 med:0.113 (0.102-0.127) | n=216 med:0.144 (0.127-0.160) | n=225 med:0.092 (0.080-0.102) | n=211 med:0.088 (0.075-0.101) | n=517 med:0.061 (0.054-0.067) |
| Macaque | n=314 med:0.103 (0.093-0.121) | n=237 med:0.131 (0.116-0.151) | n=245 med:0.091 (0.077-0.104) | n=230 med:0.078 (0.069-0.091) | n=544 med:0.052 (0.047-0.057) |
| American mink | n=292 med:0.097 (0.088-0.107) | n=204 med:0.119 (0.102-0.138) | n=204 med:0.080 (0.062-0.089) | n=221 med:0.072 (0.064-0.083) | n=499 med:0.048 (0.043-0.054) |
| Gelada | n=304 med:0.104 (0.093-0.116) | n=224 med:0.129 (0.107-0.144) | n=243 med:0.083 (0.071-0.095) | n=228 med:0.079 (0.065-0.091) | n=535 med:0.050 (0.046-0.056) |
| Sheep | n=282 med:0.115 (0.103-0.127) | n=212 med:0.126 (0.105-0.141) | n=213 med:0.080 (0.072-0.092) | n=208 med:0.083 (0.072-0.089) | n=501 med:0.056 (0.051-0.060) |
| Tasmanian devil | n=257 med:0.080 (0.070-0.090) | n=192 med:0.089 (0.078-0.103) | n=198 med:0.070 (0.062-0.082) | n=196 med:0.062 (0.055-0.070) | n=461 med:0.046 (0.042-0.052) |
| Elephant | n=295 med:0.110 (0.096-0.119) | n=212 med:0.129 (0.111-0.143) | n=221 med:0.089 (0.078-0.096) | n=224 med:0.079 (0.069-0.095) | n=508 med:0.056 (0.049-0.060) |
| Algerian mouse | n=335 med:0.122 (0.097-0.134) | n=243 med:0.143 (0.117-0.169) | n=255 med:0.089 (0.073-0.099) | n=242 med:0.087 (0.073-0.101) | n=566 med:0.053 (0.046-0.060) |
| Dingo | n=291 med:0.101 (0.089-0.108) | n=215 med:0.121 (0.105-0.139) | n=220 med:0.082 (0.069-0.090) | n=212 med:0.072 (0.064-0.082) | n=518 med:0.055 (0.046-0.059) |
| Golden snub-nosed monkey | n=291 med:0.102 (0.091-0.114) | n=209 med:0.133 (0.112-0.149) | n=228 med:0.087 (0.073-0.104) | n=213 med:0.078 (0.068-0.092) | n=509 med:0.054 (0.050-0.059) |
| Horse | n=305 med:0.110 (0.097-0.119) | n=218 med:0.125 (0.103-0.145) | n=228 med:0.090 (0.075-0.104) | n=216 med:0.086 (0.076-0.096) | n=539 med:0.058 (0.053-0.064) |
| Black snub-nosed monkey | n=297 med:0.112 (0.098-0.125) | n=204 med:0.128 (0.112-0.154) | n=221 med:0.090 (0.079-0.106) | n=219 med:0.081 (0.068-0.089) | n=512 med:0.057 (0.053-0.063) |
| Brazilian guinea pig | n=201 med:0.120 (0.104-0.133) | n=161 med:0.121 (0.109-0.133) | n=173 med:0.095 (0.086-0.106) | n=181 med:0.096 (0.085-0.111) | n=397 med:0.073 (0.068-0.081) |
| Orangutan | n=308 med:0.104 (0.092-0.119) | n=228 med:0.135 (0.115-0.143) | n=228 med:0.087 (0.074-0.098) | n=220 med:0.079 (0.067-0.088) | n=514 med:0.052 (0.047-0.056) |
| Northern American deer mouse | n=314 med:0.106 (0.095-0.125) | n=231 med:0.139 (0.112-0.150) | n=244 med:0.082 (0.070-0.097) | n=232 med:0.079 (0.066-0.090) | n=544 med:0.048 (0.042-0.056) |
| Naked mole-rat female | n=292 med:0.102 (0.092-0.116) | n=204 med:0.114 (0.095-0.139) | n=225 med:0.081 (0.070-0.091) | n=205 med:0.080 (0.068-0.090) | n=512 med:0.051 (0.045-0.056) |
| Mouse | n=348 | n=252 | n=261 | n=253 | n=582 |

| | | | | | |
|---|---|---|---|---|---|
| | med:0.110 (0.092-0.132) | med:0.140 (0.117-0.168) | med:0.084 (0.073-0.098) | med:0.078 (0.067-0.096) | med:0.051 (0.044-0.060) |
| Daurian ground squirrel | n=256 med:0.115 (0.102-0.130) | n=190 med:0.118 (0.108-0.144) | n=174 med:0.080 (0.070-0.100) | n=177 med:0.088 (0.075-0.100) | n=432 med:0.048 (0.044-0.057) |
| Tiger | n=235 med:0.109 (0.097-0.124) | n=175 med:0.130 (0.109-0.149) | n=191 med:0.089 (0.074-0.103) | n=165 med:0.081 (0.069-0.096) | n=418 med:0.054 (0.049-0.065) |
| Lesser hedgehog tenrec | n=74 med:0.116 (0.090-0.158) | n=46 med:0.113 (0.091-0.149) | n=55 med:0.102 (0.087-0.120) | n=50 med:0.078 (0.057-0.086) | n=95 med:0.066 (0.055-0.097) |
| Mongolian gerbil | n=306 med:0.120 (0.102-0.143) | n=224 med:0.147 (0.118-0.162) | n=224 med:0.082 (0.066-0.102) | n=219 med:0.085 (0.070-0.105) | n=473 med:0.055 (0.046-0.063) |
| Ma's night monkey | n=295 med:0.111 (0.102-0.124) | n=212 med:0.133 (0.105-0.151) | n=228 med:0.086 (0.073-0.098) | n=216 med:0.081 (0.070-0.092) | n=528 med:0.056 (0.050-0.061) |
| Upper Galilee mountains blind mole rat | n=301 med:0.107 (0.092-0.124) | n=224 med:0.129 (0.110-0.148) | n=247 med:0.077 (0.067-0.090) | n=233 med:0.081 (0.073-0.093) | n=546 med:0.052 (0.044-0.058) |
| Kangaroo rat | n=256 med:0.095 (0.081-0.106) | n=185 med:0.109 (0.092-0.128) | n=213 med:0.086 (0.072-0.097) | n=210 med:0.080 (0.067-0.087) | n=499 med:0.054 (0.046-0.058) |
| Chimpanzee | n=282 med:0.105 (0.094-0.122) | n=213 med:0.135 (0.111-0.146) | n=225 med:0.083 (0.073-0.097) | n=215 med:0.084 (0.076-0.100) | n=522 med:0.052 (0.047-0.057) |
| Guinea Pig | n=293 med:0.104 (0.094-0.112) | n=215 med:0.118 (0.101-0.135) | n=220 med:0.085 (0.071-0.096) | n=219 med:0.078 (0.067-0.090) | n=518 med:0.054 (0.048-0.057) |
| American beaver | n=293 med:0.113 (0.100-0.126) | n=210 med:0.129 (0.106-0.150) | n=217 med:0.085 (0.079-0.102) | n=213 med:0.083 (0.071-0.093) | n=438 med:0.056 (0.050-0.066) |
| Wild yak | n=268 med:0.104 (0.092-0.122) | n=197 med:0.117 (0.103-0.137) | n=184 med:0.084 (0.077-0.091) | n=191 med:0.078 (0.067-0.089) | n=456 med:0.052 (0.044-0.060) |
| Coquerel's sifaka | n=295 med:0.107 (0.094-0.120) | n=206 med:0.128 (0.112-0.144) | n=213 med:0.079 (0.068-0.090) | n=227 med:0.075 (0.068-0.089) | n=515 med:0.052 (0.046-0.058) |
| Alpaca | n=58 med:0.149 (0.105-0.200) | n=36 med:0.203 (0.141-0.260) | n=49 med:0.098 (0.090-0.156) | n=44 med:0.108 (0.079-0.127) | n=98 med:0.065 (0.048-0.075) |
| Lesser Egyptian jerboa | n=271 med:0.095 (0.081-0.107) | n=183 med:0.123 (0.100-0.138) | n=225 med:0.081 (0.073-0.095) | n=217 med:0.080 (0.066-0.089) | n=510 med:0.048 (0.043-0.053) |
| Long-tailed chinchilla | n=283 med:0.098 (0.089-0.108) | n=214 med:0.113 (0.094-0.131) | n=217 med:0.075 (0.066-0.086) | n=212 med:0.075 (0.066-0.085) | n=522 med:0.048 (0.041-0.054) |
| Angola colobus | n=291 med:0.107 (0.096-0.122) | n=218 med:0.131 (0.114-0.152) | n=229 med:0.083 (0.070-0.095) | n=221 med:0.083 (0.069-0.092) | n=510 med:0.052 (0.047-0.058) |
| Ryukyu mouse | n=333 med:0.107 (0.096-0.130) | n=239 med:0.153 (0.120-0.172) | n=253 med:0.082 (0.070-0.097) | n=244 med:0.082 (0.071-0.094) | n=565 med:0.053 (0.046-0.063) |
| Chinese hamster PICR | n=332 med:0.127 (0.110-0.140) | n=239 med:0.154 (0.131-0.179) | n=248 med:0.087 (0.076-0.095) | n=241 med:0.095 (0.079-0.105) | n=555 med:0.053 (0.045-0.059) |
| Vervet-AGM | n=313 med:0.103 | n=222 med:0.130 | n=240 med:0.080 | n=227 med:0.076 | n=538 med:0.048 |

| | | | | | |
|---|---|---|---|---|---|
| | (0.092-0.120) | (0.113-0.149) | (0.069-0.091) | (0.064-0.090) | (0.044-0.053) |
| Cow | n=295<br>med:0.102<br>(0.087-0.113) | n=226<br>med:0.127<br>(0.107-0.138) | n=214<br>med:0.077<br>(0.069-0.084) | n=217<br>med:0.077<br>(0.064-0.085) | n=523<br>med:0.049<br>(0.045-0.054) |
| Bushbaby | n=307<br>med:0.106<br>(0.096-0.119) | n=224<br>med:0.122<br>(0.107-0.137) | n=226<br>med:0.083<br>(0.074-0.095) | n=227<br>med:0.083<br>(0.072-0.093) | n=534<br>med:0.053<br>(0.048-0.058) |
| Bonobo | n=293<br>med:0.111<br>(0.102-0.127) | n=221<br>med:0.135<br>(0.113-0.145) | n=226<br>med:0.084<br>(0.072-0.098) | n=215<br>med:0.082<br>(0.072-0.100) | n=513<br>med:0.052<br>(0.045-0.060) |
| Hyrax | n=87<br>med:0.098<br>(0.076-0.126) | n=62<br>med:0.140<br>(0.088-0.202) | n=56<br>med:0.087<br>(0.065-0.104) | n=72<br>med:0.084<br>(0.066-0.109) | n=127<br>med:0.057<br>(0.048-0.069) |
| Rabbit | n=259<br>med:0.100<br>(0.088-0.112) | n=192<br>med:0.119<br>(0.096-0.132) | n=200<br>med:0.074<br>(0.066-0.087) | n=198<br>med:0.080<br>(0.067-0.092) | n=477<br>med:0.058<br>(0.053-0.068) |
| Megabat | n=170<br>med:0.111<br>(0.091-0.127) | n=125<br>med:0.146<br>(0.128-0.163) | n=123<br>med:0.096<br>(0.084-0.114) | n=111<br>med:0.083<br>(0.069-0.104) | n=253<br>med:0.062<br>(0.053-0.069) |
| Dog | n=299<br>med:0.108<br>(0.097-0.119) | n=216<br>med:0.112<br>(0.098-0.136) | n=224<br>med:0.088<br>(0.077-0.100) | n=217<br>med:0.082<br>(0.068-0.098) | n=520<br>med:0.055<br>(0.048-0.062) |
| Steppe mouse | n=320<br>med:0.119<br>(0.103-0.140) | n=242<br>med:0.165<br>(0.135-0.182) | n=248<br>med:0.090<br>(0.077-0.113) | n=243<br>med:0.105<br>(0.086-0.116) | n=545<br>med:0.063<br>(0.052-0.072) |
| Greater bamboo lemur | n=298<br>med:0.107<br>(0.096-0.118) | n=218<br>med:0.119<br>(0.098-0.134) | n=232<br>med:0.087<br>(0.076-0.100) | n=224<br>med:0.082<br>(0.070-0.096) | n=521<br>med:0.052<br>(0.047-0.058) |
| Pig | n=306<br>med:0.100<br>(0.093-0.111) | n=216<br>med:0.131<br>(0.101-0.150) | n=229<br>med:0.085<br>(0.075-0.101) | n=226<br>med:0.074<br>(0.065-0.088) | n=522<br>med:0.051<br>(0.046-0.055) |
| Tree Shrew | n=61<br>med:0.115<br>(0.095-0.147) | n=39<br>med:0.132<br>(0.097-0.228) | n=49<br>med:0.088<br>(0.056-0.110) | n=30<br>med:0.095<br>(0.063-0.117) | n=75<br>med:0.088<br>(0.057-0.108) |
| Pika | n=109<br>med:0.095<br>(0.082-0.116) | n=85<br>med:0.115<br>(0.090-0.130) | n=77<br>med:0.087<br>(0.058-0.099) | n=76<br>med:0.085<br>(0.066-0.103) | n=132<br>med:0.059<br>(0.043-0.072) |
| Shrew | n=43<br>med:0.120<br>(0.087-0.191) | n=37<br>med:0.106<br>(0.062-0.126) | n=29<br>med:0.085<br>(0.055-0.107) | n=34<br>med:0.077<br>(0.043-0.119) | n=80<br>med:0.051<br>(0.037-0.063) |
| Squirrel | n=297<br>med:0.099<br>(0.084-0.111) | n=214<br>med:0.107<br>(0.093-0.124) | n=233<br>med:0.077<br>(0.066-0.088) | n=217<br>med:0.071<br>(0.065-0.086) | n=528<br>med:0.048<br>(0.044-0.054) |
| Gorilla | n=293<br>med:0.110<br>(0.096-0.126) | n=217<br>med:0.132<br>(0.110-0.145) | n=227<br>med:0.081<br>(0.070-0.093) | n=223<br>med:0.083<br>(0.074-0.099) | n=526<br>med:0.054<br>(0.047-0.059) |
| Cat | n=308<br>med:0.104<br>(0.095-0.122) | n=232<br>med:0.117<br>(0.099-0.140) | n=233<br>med:0.082<br>(0.072-0.098) | n=224<br>med:0.076<br>(0.068-0.085) | n=530<br>med:0.052<br>(0.048-0.057) |

#### Rat Reference Genome, Zeisel et al., 2018, three cell types

| Sample Size and Median dN/dS (CI95%) | CNS_Glia-specific Genes | CNS_Endothelia-specific Genes | CNS_Neuron-specific Genes |
|---|---|---|---|
| Microbat | n=648 med:0.127 (0.119-0.137) | n=109 med:0.128 (0.112-0.155) | n=2,635 med:0.081 (0.078-0.084) |
| Hedgehog | n=151 med:0.127 (0.102-0.144) | n=33 med:0.134 (0.088-0.158) | n=577 med:0.089 (0.082-0.096) |
| Naked mole-rat male | n=691 med:0.129 (0.121-0.137) | n=126 med:0.118 (0.104-0.136) | n=2,616 med:0.076 (0.073-0.079) |
| Sooty mangabey | n=779 med:0.133 (0.122-0.144) | n=139 med:0.120 (0.097-0.137) | n=3,022 med:0.078 (0.075-0.080) |
| Olive baboon | n=773 med:0.133 (0.122-0.142) | n=141 med:0.126 (0.103-0.136) | n=3,018 med:0.076 (0.073-0.079) |
| American bison | n=645 med:0.118 (0.108-0.129) | n=112 med:0.127 (0.099-0.144) | n=2,460 med:0.072 (0.070-0.076) |
| Leopard | n=769 med:0.111 (0.102-0.120) | n=133 med:0.122 (0.096-0.132) | n=2,978 med:0.068 (0.065-0.071) |
| Bolivian squirrel monkey | n=750 med:0.131 (0.122-0.142) | n=134 med:0.137 (0.117-0.156) | n=2,907 med:0.075 (0.072-0.078) |
| Crab-eating macaque | n=779 med:0.130 (0.119-0.141) | n=140 med:0.120 (0.096-0.140) | n=2,985 med:0.075 (0.072-0.078) |
| Human | n=794 med:0.126 (0.116-0.135) | n=145 med:0.116 (0.100-0.137) | n=3,105 med:0.072 (0.069-0.075) |
| Red fox | n=744 med:0.124 (0.114-0.135) | n=125 med:0.130 (0.104-0.143) | n=2,865 med:0.079 (0.076-0.082) |
| Shrew mouse | n=846 med:0.142 (0.133-0.154) | n=147 med:0.132 (0.111-0.150) | n=3,194 med:0.079 (0.075-0.083) |
| Arctic ground squirrel | n=774 med:0.121 (0.111-0.129) | n=139 med:0.123 (0.097-0.139) | n=3,017 med:0.069 (0.066-0.072) |
| Chinese hamster CriGri | n=775 med:0.141 (0.134-0.152) | n=138 med:0.143 (0.120-0.161) | n=2,944 med:0.089 (0.084-0.092) |
| Opossum | n=697 | n=126 | n=2,731 |

| | | | |
|---|---|---|---|
| | med:0.095 (0.089-0.102) | med:0.094 (0.072-0.109) | med:0.067 (0.064-0.069) |
| Dolphin | n=417 med:0.128 (0.116-0.141) | n=85 med:0.123 (0.100-0.146) | n=1,614 med:0.084 (0.080-0.089) |
| Capuchin | n=766 med:0.130 (0.122-0.137) | n=138 med:0.132 (0.113-0.153) | n=2,963 med:0.076 (0.073-0.079) |
| Mouse Lemur | n=738 med:0.124 (0.115-0.132) | n=136 med:0.117 (0.100-0.128) | n=2,927 med:0.073 (0.069-0.076) |
| Damara mole rat | n=689 med:0.121 (0.109-0.131) | n=121 med:0.111 (0.100-0.136) | n=2,817 med:0.073 (0.070-0.076) |
| Marmoset | n=774 med:0.129 (0.118-0.137) | n=137 med:0.127 (0.105-0.147) | n=2,976 med:0.078 (0.075-0.082) |
| Goat | n=749 med:0.115 (0.104-0.122) | n=129 med:0.114 (0.100-0.131) | n=2,935 med:0.071 (0.068-0.074) |
| Common wombat | n=663 med:0.097 (0.090-0.102) | n=116 med:0.095 (0.079-0.110) | n=2,656 med:0.065 (0.062-0.067) |
| American black bear | n=668 med:0.119 (0.113-0.127) | n=121 med:0.125 (0.103-0.139) | n=2,601 med:0.081 (0.078-0.084) |
| Sloth | n=104 med:0.172 (0.139-0.205) | n=18 med:0.174 (0.102-0.255) | n=404 med:0.099 (0.087-0.114) |
| Polar bear | n=668 med:0.126 (0.120-0.139) | n=116 med:0.150 (0.120-0.168) | n=2,631 med:0.086 (0.082-0.089) |
| Alpine marmot | n=721 med:0.124 (0.116-0.133) | n=135 med:0.131 (0.104-0.151) | n=2,881 med:0.072 (0.069-0.076) |
| Prairie vole | n=798 med:0.129 (0.120-0.139) | n=145 med:0.123 (0.109-0.141) | n=3,092 med:0.074 (0.071-0.078) |
| Donkey | n=760 med:0.124 (0.115-0.132) | n=125 med:0.124 (0.101-0.135) | n=3,022 med:0.078 (0.076-0.081) |
| Golden Hamster | n=772 med:0.147 (0.137-0.160) | n=140 med:0.139 (0.120-0.163) | n=2,931 med:0.085 (0.080-0.089) |
| Armadillo | n=684 med:0.121 (0.114-0.130) | n=116 med:0.121 (0.109-0.132) | n=2,728 med:0.079 (0.076-0.082) |
| Ugandan red Colobus | n=760 | n=136 | n=2,935 |

| | | | |
|---|---|---|---|
| | med:0.131 (0.121-0.141) | med:0.129 (0.107-0.148) | med:0.076 (0.073-0.080) |
| Pig-tailed macaque | n=775 med:0.132 (0.122-0.143) | n=143 med:0.127 (0.106-0.143) | n=3,017 med:0.076 (0.073-0.079) |
| Panda | n=740 med:0.121 (0.111-0.128) | n=133 med:0.123 (0.105-0.142) | n=2,956 med:0.073 (0.070-0.076) |
| Ferret | n=737 med:0.114 (0.109-0.123) | n=128 med:0.109 (0.097-0.128) | n=2,942 med:0.070 (0.067-0.073) |
| Koala | n=722 med:0.097 (0.092-0.104) | n=132 med:0.090 (0.076-0.104) | n=2,901 med:0.067 (0.065-0.069) |
| Wallaby | n=210 med:0.086 (0.076-0.093) | n=52 med:0.091 (0.061-0.108) | n=810 med:0.069 (0.065-0.075) |
| Tarsier | n=653 med:0.133 (0.118-0.140) | n=114 med:0.132 (0.109-0.148) | n=2,472 med:0.079 (0.075-0.082) |
| Degu | n=657 med:0.117 (0.109-0.124) | n=125 med:0.106 (0.092-0.115) | n=2,627 med:0.072 (0.069-0.075) |
| Chinese hamster CHOK1GS | n=834 med:0.138 (0.130-0.149) | n=151 med:0.136 (0.113-0.155) | n=3,190 med:0.081 (0.077-0.085) |
| Drill | n=757 med:0.133 (0.123-0.143) | n=132 med:0.124 (0.107-0.137) | n=2,917 med:0.077 (0.075-0.080) |
| Gibbon | n=750 med:0.141 (0.134-0.151) | n=134 med:0.129 (0.109-0.148) | n=2,900 med:0.083 (0.081-0.087) |
| Macaque | n=798 med:0.129 (0.117-0.139) | n=144 med:0.124 (0.098-0.140) | n=3,074 med:0.077 (0.075-0.080) |
| American mink | n=727 med:0.115 (0.108-0.122) | n=128 med:0.107 (0.100-0.127) | n=2,853 med:0.069 (0.066-0.072) |
| Gelada | n=776 med:0.130 (0.119-0.140) | n=138 med:0.120 (0.099-0.135) | n=3,014 med:0.072 (0.069-0.076) |
| Sheep | n=741 med:0.123 (0.115-0.129) | n=125 med:0.124 (0.109-0.143) | n=2,885 med:0.081 (0.077-0.084) |
| Tasmanian devil | n=663 med:0.091 (0.084-0.098) | n=117 med:0.086 (0.071-0.097) | n=2,606 med:0.063 (0.061-0.066) |
| Elephant | n=719 | n=128 | n=2,917 |

| | | | |
|---|---|---|---|
| | med:0.124 (0.112-0.131) | med:0.125 (0.110-0.136) | med:0.078 (0.075-0.082) |
| Algerian mouse | n=843 med:0.141 (0.132-0.154) | n=150 med:0.137 (0.112-0.162) | n=3,234 med:0.079 (0.075-0.083) |
| Dingo | n=728 med:0.115 (0.107-0.123) | n=133 med:0.113 (0.098-0.131) | n=2,938 med:0.073 (0.069-0.076) |
| Golden snub-nosed monkey | n=761 med:0.136 (0.125-0.142) | n=144 med:0.118 (0.102-0.136) | n=2,862 med:0.076 (0.072-0.079) |
| Horse | n=766 med:0.128 (0.119-0.139) | n=133 med:0.124 (0.102-0.138) | n=3,026 med:0.078 (0.075-0.081) |
| Black snub-nosed monkey | n=760 med:0.136 (0.124-0.145) | n=136 med:0.132 (0.113-0.148) | n=2,900 med:0.081 (0.078-0.084) |
| Brazilian guinea pig | n=551 med:0.127 (0.119-0.137) | n=91 med:0.130 (0.117-0.142) | n=2,170 med:0.096 (0.091-0.099) |
| Orangutan | n=742 med:0.126 (0.117-0.137) | n=137 med:0.119 (0.104-0.152) | n=2,949 med:0.074 (0.072-0.077) |
| Northern American deer mouse | n=813 med:0.128 (0.119-0.137) | n=142 med:0.127 (0.111-0.148) | n=3,034 med:0.072 (0.069-0.076) |
| Naked mole-rat female | n=735 med:0.127 (0.119-0.134) | n=138 med:0.118 (0.101-0.134) | n=2,782 med:0.075 (0.072-0.078) |
| Mouse | n=871 med:0.140 (0.132-0.151) | n=155 med:0.132 (0.113-0.164) | n=3,322 med:0.076 (0.073-0.080) |
| Daurian ground squirrel | n=647 med:0.128 (0.119-0.139) | n=119 med:0.135 (0.113-0.159) | n=2,537 med:0.077 (0.072-0.080) |
| Tiger | n=664 med:0.129 (0.119-0.139) | n=104 med:0.132 (0.119-0.146) | n=2,424 med:0.080 (0.076-0.083) |
| Lesser hedgehog tenrec | n=166 med:0.130 (0.111-0.149) | n=37 med:0.124 (0.097-0.183) | n=648 med:0.084 (0.077-0.092) |
| Mongolian gerbil | n=753 med:0.137 (0.126-0.148) | n=140 med:0.129 (0.102-0.152) | n=2,824 med:0.083 (0.080-0.088) |
| Ma's night monkey | n=763 med:0.133 (0.122-0.142) | n=132 med:0.129 (0.108-0.156) | n=2,915 med:0.079 (0.077-0.082) |

| | | | |
|---|---|---|---|
| Upper Galilee mountains blind mole rat | n=757 med:0.129 (0.120-0.139) | n=140 med:0.125 (0.109-0.137) | n=3,001 med:0.072 (0.069-0.076) |
| Kangaroo rat | n=659 med:0.113 (0.104-0.124) | n=119 med:0.116 (0.095-0.126) | n=2,740 med:0.071 (0.067-0.074) |
| Chimpanzee | n=759 med:0.130 (0.120-0.139) | n=134 med:0.113 (0.097-0.132) | n=2,899 med:0.076 (0.073-0.079) |
| Guinea Pig | n=724 med:0.119 (0.112-0.129) | n=131 med:0.107 (0.094-0.132) | n=2,952 med:0.075 (0.072-0.078) |
| American beaver | n=714 med:0.126 (0.118-0.136) | n=140 med:0.125 (0.104-0.145) | n=2,741 med:0.081 (0.077-0.084) |
| Wild yak | n=688 med:0.116 (0.110-0.128) | n=121 med:0.123 (0.103-0.140) | n=2,576 med:0.074 (0.071-0.078) |
| Coquerel's sifaka | n=721 med:0.126 (0.118-0.137) | n=129 med:0.119 (0.103-0.138) | n=2,858 med:0.076 (0.073-0.079) |
| Alpaca | n=160 med:0.172 (0.146-0.197) | n=23 med:0.146 (0.092-0.244) | n=664 med:0.091 (0.081-0.099) |
| Lesser Egyptian jerboa | n=700 med:0.121 (0.114-0.129) | n=129 med:0.118 (0.103-0.136) | n=2,766 med:0.074 (0.070-0.077) |
| Long-tailed chinchilla | n=738 med:0.116 (0.109-0.125) | n=132 med:0.104 (0.096-0.117) | n=2,958 med:0.069 (0.066-0.072) |
| Angola colobus | n=767 med:0.135 (0.125-0.142) | n=134 med:0.135 (0.114-0.155) | n=2,920 med:0.077 (0.074-0.080) |
| Ryukyu mouse | n=832 med:0.141 (0.130-0.153) | n=149 med:0.127 (0.104-0.167) | n=3,198 med:0.079 (0.075-0.082) |
| Chinese hamster PICR | n=822 med:0.144 (0.136-0.157) | n=150 med:0.138 (0.124-0.164) | n=3,156 med:0.085 (0.081-0.089) |
| Vervet-AGM | n=780 med:0.126 (0.116-0.136) | n=143 med:0.121 (0.105-0.134) | n=3,044 med:0.070 (0.067-0.073) |
| Cow | n=758 med:0.111 (0.107-0.122) | n=130 med:0.114 (0.100-0.131) | n=2,943 med:0.072 (0.069-0.074) |
| Bushbaby | n=764 med:0.129 (0.116-0.136) | n=133 med:0.127 (0.108-0.142) | n=2,982 med:0.076 (0.074-0.080) |

| | | | |
|---|---|---|---|
| Bonobo | n=770<br>med:0.134<br>(0.125-0.140) | n=137<br>med:0.126<br>(0.106-0.141) | n=2,908<br>med:0.078<br>(0.075-0.081) |
| Hyrax | n=209<br>med:0.131<br>(0.113-0.149) | n=50<br>med:0.106<br>(0.090-0.151) | n=803<br>med:0.084<br>(0.078-0.090) |
| Rabbit | n=675<br>med:0.113<br>(0.105-0.121) | n=115<br>med:0.115<br>(0.100-0.130) | n=2,680<br>med:0.077<br>(0.073-0.080) |
| Megabat | n=421<br>med:0.133<br>(0.120-0.146) | n=76<br>med:0.133<br>(0.109-0.169) | n=1,492<br>med:0.085<br>(0.080-0.090) |
| Dog | n=761<br>med:0.122<br>(0.114-0.133) | n=134<br>med:0.131<br>(0.106-0.150) | n=2,920<br>med:0.078<br>(0.075-0.082) |
| Steppe mouse | n=824<br>med:0.157<br>(0.144-0.168) | n=144<br>med:0.140<br>(0.119-0.174) | n=3,116<br>med:0.088<br>(0.083-0.092) |
| Greater bamboo lemur | n=764<br>med:0.125<br>(0.116-0.133) | n=135<br>med:0.119<br>(0.107-0.134) | n=2,984<br>med:0.076<br>(0.072-0.079) |
| Pig | n=753<br>med:0.120<br>(0.111-0.129) | n=138<br>med:0.108<br>(0.098-0.123) | n=3,002<br>med:0.075<br>(0.072-0.078) |
| Tree Shrew | n=155<br>med:0.149<br>(0.117-0.168) | n=29<br>med:0.103<br>(0.076-0.160) | n=535<br>med:0.088<br>(0.081-0.097) |
| Pika | n=273<br>med:0.106<br>(0.092-0.121) | n=58<br>med:0.118<br>(0.095-0.131) | n=956<br>med:0.075<br>(0.071-0.081) |
| Shrew | n=114<br>med:0.116<br>(0.092-0.135) | n=22<br>med:0.116<br>(0.073-0.243) | n=543<br>med:0.074<br>(0.064-0.081) |
| Squirrel | n=749<br>med:0.120<br>(0.110-0.127) | n=140<br>med:0.124<br>(0.097-0.133) | n=3,008<br>med:0.069<br>(0.065-0.071) |
| Gorilla | n=756<br>med:0.132<br>(0.122-0.140) | n=135<br>med:0.126<br>(0.107-0.145) | n=2,942<br>med:0.077<br>(0.074-0.080) |
| Cat | n=788<br>med:0.121<br>(0.111-0.128) | n=140<br>med:0.126<br>(0.106-0.138) | n=3,052<br>med:0.075<br>(0.072-0.078) |

#### Rat Reference Genome, Zeisel et al., 2018, cell subtypes

| Sample Size and Median dN/dS (CI95%) | Glia-specific Genes | Immune cells-specific Genes | Vascular cells-specific Genes | Neurons-specific Genes |
|---|---|---|---|---|
| Microbat | n=543 med:0.140 (0.132-0.155) | n=73 med:0.130 (0.100-0.178) | n=134 med:0.112 (0.093-0.130) | n=2,754 med:0.090 (0.086-0.093) |
| Hedgehog | n=121 med:0.131 (0.108-0.160) | n=24 med:0.123 (0.087-0.156) | n=28 med:0.139 (0.083-0.158) | n=634 med:0.100 (0.091-0.111) |
| Naked mole-rat male | n=572 med:0.141 (0.132-0.155) | n=73 med:0.137 (0.098-0.167) | n=142 med:0.113 (0.100-0.131) | n=2,751 med:0.087 (0.083-0.091) |
| Sooty mangabey | n=629 med:0.151 (0.140-0.161) | n=87 med:0.167 (0.121-0.187) | n=160 med:0.115 (0.099-0.128) | n=3,174 med:0.085 (0.081-0.089) |
| Olive baboon | n=627 med:0.151 (0.139-0.161) | n=83 med:0.154 (0.126-0.186) | n=162 med:0.110 (0.092-0.128) | n=3,174 med:0.083 (0.079-0.086) |
| American bison | n=538 med:0.136 (0.124-0.147) | n=75 med:0.144 (0.127-0.179) | n=128 med:0.119 (0.094-0.135) | n=2,595 med:0.082 (0.078-0.086) |
| Leopard | n=629 med:0.130 (0.117-0.143) | n=87 med:0.124 (0.097-0.166) | n=155 med:0.104 (0.090-0.119) | n=3,126 med:0.076 (0.073-0.079) |
| Bolivian squirrel monkey | n=624 med:0.147 (0.137-0.158) | n=80 med:0.150 (0.115-0.189) | n=148 med:0.121 (0.107-0.144) | n=3,049 med:0.085 (0.080-0.087) |
| Crab-eating macaque | n=634 med:0.147 (0.138-0.159) | n=88 med:0.158 (0.119-0.194) | n=159 med:0.110 (0.095-0.123) | n=3,142 med:0.083 (0.079-0.086) |
| Human | n=653 med:0.144 (0.135-0.154) | n=89 med:0.141 (0.120-0.185) | n=167 med:0.113 (0.099-0.123) | n=3,249 med:0.081 (0.077-0.084) |
| Red fox | n=617 med:0.137 (0.127-0.152) | n=81 med:0.162 (0.123-0.189) | n=147 med:0.117 (0.098-0.137) | n=2,986 med:0.087 (0.083-0.091) |
| Shrew mouse | n=684 med:0.161 (0.150-0.175) | n=110 med:0.202 (0.163-0.240) | n=170 med:0.116 (0.099-0.134) | n=3,374 med:0.090 (0.086-0.094) |
| Arctic ground squirrel | n=628 med:0.137 (0.124-0.146) | n=83 med:0.160 (0.110-0.189) | n=161 med:0.105 (0.086-0.130) | n=3,177 med:0.079 (0.075-0.083) |
| Chinese hamster CriGri | n=632 med:0.165 (0.152-0.177) | n=102 med:0.217 (0.147-0.264) | n=156 med:0.141 (0.123-0.157) | n=3,115 med:0.099 (0.095-0.102) |
| Opossum | n=577 | n=70 | n=150 | n=2,877 |

| | | | | |
|---|---|---|---|---|
| | med:0.106 (0.098-0.114) | med:0.079 (0.065-0.110) | med:0.089 (0.069-0.104) | med:0.072 (0.069-0.075) |
| Dolphin | n=353 med:0.151 (0.136-0.162) | n=55 med:0.162 (0.125-0.232) | n=91 med:0.117 (0.098-0.143) | n=1,706 med:0.094 (0.089-0.099) |
| Capuchin | n=627 med:0.147 (0.136-0.157) | n=85 med:0.162 (0.111-0.203) | n=156 med:0.114 (0.101-0.133) | n=3,112 med:0.083 (0.080-0.087) |
| Mouse Lemur | n=613 med:0.143 (0.126-0.151) | n=87 med:0.140 (0.108-0.163) | n=153 med:0.109 (0.095-0.125) | n=3,087 med:0.082 (0.078-0.085) |
| Damara mole rat | n=569 med:0.138 (0.124-0.147) | n=73 med:0.135 (0.098-0.166) | n=140 med:0.115 (0.102-0.132) | n=2,938 med:0.082 (0.078-0.086) |
| Marmoset | n=637 med:0.150 (0.139-0.158) | n=92 med:0.136 (0.109-0.177) | n=160 med:0.127 (0.111-0.142) | n=3,136 med:0.087 (0.084-0.090) |
| Goat | n=619 med:0.133 (0.121-0.142) | n=87 med:0.143 (0.124-0.182) | n=152 med:0.100 (0.085-0.117) | n=3,080 med:0.079 (0.076-0.084) |
| Common wombat | n=547 med:0.108 (0.100-0.119) | n=67 med:0.103 (0.078-0.129) | n=136 med:0.091 (0.079-0.103) | n=2,794 med:0.071 (0.067-0.074) |
| American black bear | n=553 med:0.136 (0.123-0.146) | n=75 med:0.150 (0.109-0.183) | n=136 med:0.111 (0.092-0.131) | n=2,744 med:0.090 (0.087-0.094) |
| Sloth | n=74 med:0.228 (0.183-0.252) | n=14 med:0.300 (0.105-0.613) | n=18 med:0.152 (0.090-0.209) | n=431 med:0.130 (0.114-0.140) |
| Polar bear | n=551 med:0.148 (0.138-0.156) | n=75 med:0.164 (0.115-0.210) | n=127 med:0.130 (0.110-0.153) | n=2,760 med:0.096 (0.092-0.100) |
| Alpine marmot | n=590 med:0.139 (0.130-0.148) | n=86 med:0.150 (0.101-0.176) | n=150 med:0.117 (0.101-0.140) | n=3,040 med:0.082 (0.079-0.085) |
| Prairie vole | n=664 med:0.148 (0.138-0.161) | n=99 med:0.176 (0.128-0.236) | n=165 med:0.124 (0.110-0.141) | n=3,248 med:0.084 (0.080-0.089) |
| Donkey | n=625 med:0.139 (0.129-0.152) | n=85 med:0.134 (0.112-0.161) | n=148 med:0.112 (0.091-0.126) | n=3,186 med:0.088 (0.085-0.091) |
| Golden Hamster | n=637 med:0.165 (0.153-0.179) | n=94 med:0.195 (0.141-0.250) | n=153 med:0.131 (0.106-0.144) | n=3,090 med:0.096 (0.091-0.101) |
| Armadillo | n=568 med:0.138 (0.126-0.145) | n=86 med:0.127 (0.101-0.155) | n=136 med:0.108 (0.093-0.128) | n=2,863 med:0.087 (0.083-0.091) |

| | | | | |
|---|---|---|---|---|
| Ugandan red Colobus | n=617 med:0.150 (0.139-0.160) | n=83 med:0.148 (0.116-0.167) | n=153 med:0.114 (0.096-0.136) | n=3,090 med:0.084 (0.080-0.087) |
| Pig-tailed macaque | n=634 med:0.148 (0.139-0.161) | n=84 med:0.162 (0.125-0.209) | n=162 med:0.116 (0.099-0.127) | n=3,173 med:0.084 (0.081-0.088) |
| Panda | n=610 med:0.134 (0.126-0.142) | n=88 med:0.158 (0.106-0.172) | n=150 med:0.111 (0.092-0.123) | n=3,085 med:0.081 (0.078-0.084) |
| Ferret | n=618 med:0.134 (0.122-0.144) | n=88 med:0.153 (0.116-0.170) | n=149 med:0.102 (0.086-0.109) | n=3,074 med:0.078 (0.075-0.082) |
| Koala | n=599 med:0.108 (0.101-0.121) | n=73 med:0.103 (0.075-0.130) | n=151 med:0.086 (0.067-0.095) | n=3,036 med:0.072 (0.069-0.075) |
| Wallaby | n=148 med:0.097 (0.078-0.106) | n=18 med:0.080 (0.065-0.108) | n=59 med:0.080 (0.068-0.090) | n=874 med:0.073 (0.068-0.079) |
| Tarsier | n=549 med:0.152 (0.139-0.162) | n=75 med:0.146 (0.114-0.173) | n=122 med:0.123 (0.101-0.147) | n=2,601 med:0.089 (0.085-0.093) |
| Degu | n=542 med:0.134 (0.124-0.146) | n=66 med:0.119 (0.090-0.148) | n=144 med:0.109 (0.099-0.122) | n=2,799 med:0.081 (0.078-0.084) |
| Chinese hamster CHOK1GS | n=682 med:0.162 (0.150-0.177) | n=107 med:0.227 (0.147-0.262) | n=173 med:0.127 (0.105-0.142) | n=3,370 med:0.092 (0.088-0.095) |
| Drill | n=626 med:0.148 (0.138-0.157) | n=86 med:0.165 (0.127-0.190) | n=156 med:0.117 (0.103-0.130) | n=3,062 med:0.087 (0.083-0.090) |
| Gibbon | n=622 med:0.157 (0.145-0.164) | n=77 med:0.170 (0.139-0.224) | n=151 med:0.117 (0.108-0.139) | n=3,051 med:0.092 (0.088-0.095) |
| Macaque | n=658 med:0.145 (0.138-0.156) | n=92 med:0.149 (0.114-0.175) | n=166 med:0.118 (0.098-0.131) | n=3,220 med:0.086 (0.082-0.090) |
| American mink | n=607 med:0.133 (0.119-0.142) | n=79 med:0.145 (0.109-0.168) | n=147 med:0.100 (0.081-0.114) | n=2,989 med:0.078 (0.075-0.081) |
| Gelada | n=635 med:0.146 (0.136-0.157) | n=87 med:0.159 (0.115-0.187) | n=159 med:0.114 (0.099-0.127) | n=3,184 med:0.079 (0.076-0.083) |
| Sheep | n=611 med:0.133 (0.127-0.143) | n=84 med:0.153 (0.123-0.185) | n=146 med:0.116 (0.103-0.131) | n=3,004 med:0.089 (0.086-0.093) |
| Tasmanian devil | n=552 med:0.105 (0.097-0.114) | n=70 med:0.098 (0.071-0.127) | n=135 med:0.077 (0.066-0.090) | n=2,723 med:0.069 (0.067-0.072) |

| | | | | |
|---|---|---|---|---|
| Elephant | n=598 med:0.135 (0.125-0.145) | n=81 med:0.137 (0.120-0.190) | n=146 med:0.112 (0.094-0.142) | n=3,056 med:0.088 (0.084-0.091) |
| Algerian mouse | n=696 med:0.167 (0.154-0.184) | n=110 med:0.215 (0.154-0.268) | n=167 med:0.123 (0.096-0.139) | n=3,415 med:0.091 (0.086-0.094) |
| Dingo | n=592 med:0.129 (0.117-0.142) | n=85 med:0.158 (0.112-0.178) | n=154 med:0.103 (0.087-0.124) | n=3,081 med:0.081 (0.078-0.085) |
| Golden snub-nosed monkey | n=626 med:0.150 (0.139-0.159) | n=83 med:0.160 (0.120-0.189) | n=155 med:0.112 (0.095-0.127) | n=3,004 med:0.084 (0.080-0.087) |
| Horse | n=638 med:0.144 (0.132-0.159) | n=82 med:0.146 (0.114-0.176) | n=155 med:0.109 (0.092-0.131) | n=3,170 med:0.087 (0.083-0.090) |
| Black snub-nosed monkey | n=622 med:0.152 (0.143-0.163) | n=81 med:0.154 (0.113-0.182) | n=157 med:0.118 (0.101-0.142) | n=3,055 med:0.089 (0.085-0.093) |
| Brazilian guinea pig | n=440 med:0.143 (0.138-0.154) | n=61 med:0.131 (0.099-0.157) | n=115 med:0.125 (0.115-0.144) | n=2,266 med:0.104 (0.099-0.108) |
| Orangutan | n=601 med:0.145 (0.133-0.153) | n=89 med:0.144 (0.123-0.191) | n=153 med:0.116 (0.095-0.124) | n=3,086 med:0.082 (0.078-0.086) |
| Northern American deer mouse | n=660 med:0.151 (0.138-0.162) | n=102 med:0.197 (0.146-0.264) | n=158 med:0.115 (0.098-0.137) | n=3,210 med:0.083 (0.079-0.088) |
| Naked mole-rat female | n=607 med:0.141 (0.131-0.155) | n=77 med:0.140 (0.105-0.189) | n=160 med:0.114 (0.102-0.126) | n=2,915 med:0.083 (0.080-0.087) |
| Mouse | n=715 med:0.163 (0.151-0.177) | n=116 med:0.216 (0.147-0.273) | n=176 med:0.114 (0.093-0.137) | n=3,496 med:0.088 (0.083-0.091) |
| Daurian ground squirrel | n=533 med:0.143 (0.133-0.154) | n=76 med:0.152 (0.112-0.208) | n=135 med:0.127 (0.104-0.144) | n=2,672 med:0.088 (0.084-0.092) |
| Tiger | n=561 med:0.145 (0.132-0.156) | n=77 med:0.167 (0.114-0.214) | n=120 med:0.123 (0.099-0.137) | n=2,558 med:0.088 (0.084-0.093) |
| Lesser hedgehog tenrec | n=129 med:0.138 (0.119-0.160) | n=18 med:0.168 (0.107-0.324) | n=36 med:0.094 (0.076-0.124) | n=707 med:0.094 (0.085-0.105) |
| Mongolian gerbil | n=611 med:0.157 (0.139-0.172) | n=96 med:0.190 (0.151-0.230) | n=157 med:0.122 (0.104-0.139) | n=2,995 med:0.097 (0.092-0.101) |
| Ma's night monkey | n=621 med:0.150 (0.138-0.159) | n=86 med:0.144 (0.110-0.180) | n=154 med:0.116 (0.104-0.131) | n=3,060 med:0.087 (0.083-0.091) |

| | | | | |
|---|---|---|---|---|
| Upper Galilee mountains blind mole rat | n=625 med:0.153 (0.139-0.163) | n=90 med:0.156 (0.123-0.201) | n=158 med:0.129 (0.100-0.140) | n=3,172 med:0.085 (0.080-0.089) |
| Kangaroo rat | n=533 med:0.132 (0.121-0.143) | n=69 med:0.119 (0.089-0.150) | n=138 med:0.113 (0.088-0.126) | n=2,855 med:0.080 (0.077-0.083) |
| Chimpanzee | n=612 med:0.145 (0.134-0.156) | n=82 med:0.147 (0.126-0.200) | n=153 med:0.113 (0.091-0.127) | n=3,063 med:0.085 (0.081-0.089) |
| Guinea Pig | n=597 med:0.140 (0.131-0.152) | n=75 med:0.141 (0.101-0.175) | n=152 med:0.108 (0.095-0.122) | n=3,078 med:0.083 (0.080-0.086) |
| American beaver | n=560 med:0.142 (0.131-0.154) | n=87 med:0.151 (0.114-0.209) | n=154 med:0.114 (0.098-0.135) | n=2,898 med:0.090 (0.086-0.094) |
| Wild yak | n=564 med:0.136 (0.124-0.148) | n=82 med:0.130 (0.112-0.169) | n=137 med:0.120 (0.098-0.136) | n=2,707 med:0.084 (0.080-0.087) |
| Coquerel's sifaka | n=600 med:0.147 (0.137-0.157) | n=81 med:0.146 (0.118-0.172) | n=153 med:0.113 (0.098-0.129) | n=2,995 med:0.084 (0.080-0.087) |
| Alpaca | n=140 med:0.194 (0.169-0.214) | n=14 med:0.221 (0.122-0.352) | n=25 med:0.146 (0.092-0.169) | n=697 med:0.107 (0.099-0.114) |
| Lesser Egyptian jerboa | n=573 med:0.141 (0.131-0.150) | n=72 med:0.136 (0.121-0.170) | n=145 med:0.112 (0.103-0.128) | n=2,880 med:0.082 (0.079-0.086) |
| Long-tailed chinchilla | n=615 med:0.136 (0.127-0.146) | n=78 med:0.130 (0.096-0.162) | n=152 med:0.107 (0.096-0.117) | n=3,097 med:0.077 (0.074-0.080) |
| Angola colobus | n=621 med:0.148 (0.139-0.159) | n=85 med:0.156 (0.123-0.208) | n=151 med:0.117 (0.105-0.137) | n=3,054 med:0.086 (0.082-0.090) |
| Ryukyu mouse | n=680 med:0.164 (0.152-0.174) | n=108 med:0.229 (0.171-0.268) | n=171 med:0.116 (0.103-0.136) | n=3,370 med:0.089 (0.085-0.093) |
| Chinese hamster PICR | n=681 med:0.168 (0.153-0.179) | n=105 med:0.233 (0.151-0.286) | n=168 med:0.126 (0.106-0.142) | n=3,336 med:0.096 (0.092-0.100) |
| Vervet-AGM | n=636 med:0.145 (0.135-0.157) | n=85 med:0.148 (0.117-0.174) | n=165 med:0.118 (0.095-0.124) | n=3,196 med:0.079 (0.075-0.083) |
| Cow | n=623 med:0.132 (0.120-0.140) | n=88 med:0.148 (0.124-0.185) | n=156 med:0.113 (0.091-0.125) | n=3,100 med:0.080 (0.076-0.083) |
| Bushbaby | n=616 med:0.147 (0.136-0.155) | n=89 med:0.150 (0.116-0.191) | n=150 med:0.119 (0.101-0.136) | n=3,154 med:0.087 (0.083-0.091) |

| | | | | |
|---|---|---|---|---|
| Bonobo | n=621<br>med:0.147<br>(0.138-0.158) | n=86<br>med:0.146<br>(0.125-0.185) | n=154<br>med:0.114<br>(0.100-0.129) | n=3,060<br>med:0.086<br>(0.083-0.090) |
| Hyrax | n=173<br>med:0.155<br>(0.133-0.177) | n=34<br>med:0.216<br>(0.142-0.272) | n=52<br>med:0.106<br>(0.068-0.143) | n=854<br>med:0.092<br>(0.086-0.100) |
| Rabbit | n=564<br>med:0.127<br>(0.121-0.136) | n=76<br>med:0.139<br>(0.101-0.173) | n=135<br>med:0.101<br>(0.083-0.118) | n=2,797<br>med:0.084<br>(0.081-0.088) |
| Megabat | n=334<br>med:0.140<br>(0.126-0.159) | n=52<br>med:0.180<br>(0.146-0.242) | n=83<br>med:0.128<br>(0.115-0.145) | n=1,608<br>med:0.095<br>(0.091-0.100) |
| Dog | n=627<br>med:0.142<br>(0.129-0.156) | n=85<br>med:0.122<br>(0.102-0.169) | n=156<br>med:0.110<br>(0.098-0.130) | n=3,046<br>med:0.087<br>(0.084-0.090) |
| Steppe mouse | n=682<br>med:0.172<br>(0.161-0.189) | n=114<br>med:0.226<br>(0.182-0.276) | n=163<br>med:0.126<br>(0.100-0.154) | n=3,282<br>med:0.098<br>(0.094-0.103) |
| Greater bamboo lemur | n=628<br>med:0.144<br>(0.131-0.156) | n=85<br>med:0.128<br>(0.099-0.178) | n=159<br>med:0.116<br>(0.099-0.130) | n=3,153<br>med:0.085<br>(0.081-0.088) |
| Pig | n=633<br>med:0.137<br>(0.127-0.146) | n=83<br>med:0.138<br>(0.105-0.161) | n=159<br>med:0.107<br>(0.092-0.124) | n=3,147<br>med:0.082<br>(0.078-0.085) |
| Tree Shrew | n=130<br>med:0.171<br>(0.145-0.192) | n=20<br>med:0.276<br>(0.156-0.346) | n=25<br>med:0.120<br>(0.088-0.177) | n=577<br>med:0.097<br>(0.089-0.108) |
| Pika | n=206<br>med:0.121<br>(0.106-0.138) | n=39<br>med:0.130<br>(0.090-0.178) | n=63<br>med:0.105<br>(0.083-0.129) | n=1,053<br>med:0.084<br>(0.079-0.089) |
| Shrew | n=85<br>med:0.140<br>(0.107-0.168) | n=19<br>med:0.212<br>(0.118-0.313) | n=29<br>med:0.123<br>(0.095-0.160) | n=612<br>med:0.085<br>(0.079-0.094) |
| Squirrel | n=618<br>med:0.138<br>(0.126-0.148) | n=82<br>med:0.127<br>(0.097-0.176) | n=159<br>med:0.116<br>(0.093-0.131) | n=3,174<br>med:0.078<br>(0.074-0.081) |
| Gorilla | n=622<br>med:0.150<br>(0.139-0.158) | n=82<br>med:0.154<br>(0.123-0.219) | n=154<br>med:0.118<br>(0.103-0.135) | n=3,097<br>med:0.085<br>(0.081-0.089) |
| Cat | n=650<br>med:0.134<br>(0.126-0.148) | n=95<br>med:0.123<br>(0.097-0.167) | n=164<br>med:0.112<br>(0.098-0.128) | n=3,200<br>med:0.083<br>(0.080-0.086) |

#### Human Reference Genome, Zhang et al., 2014, three cell types

| Sample Size and Median dN/dS (CI95%) | Endothelia-specific Genes | Glia-specific Genes | Neuron-specific Genes |
|---|---|---|---|
| Microbat | n=665 med:0.145 (0.136-0.163) | n=817 med:0.145 (0.133-0.156) | n=928 med:0.095 (0.086-0.104) |
| Hedgehog | n=172 med:0.157 (0.129-0.179) | n=222 med:0.140 (0.122-0.161) | n=224 med:0.097 (0.075-0.109) |
| Naked mole-rat male | n=668 med:0.143 (0.131-0.154) | n=916 med:0.134 (0.126-0.145) | n=998 med:0.078 (0.070-0.084) |
| Sooty mangabey | n=782 med:0.173 (0.157-0.187) | n=1,039 med:0.162 (0.148-0.179) | n=1,152 med:0.098 (0.090-0.107) |
| Olive baboon | n=787 med:0.183 (0.169-0.197) | n=1,040 med:0.172 (0.157-0.186) | n=1,150 med:0.103 (0.093-0.112) |
| American bison | n=638 med:0.143 (0.131-0.157) | n=818 med:0.137 (0.129-0.146) | n=863 med:0.080 (0.073-0.087) |
| Leopard | n=749 med:0.129 (0.121-0.139) | n=980 med:0.121 (0.111-0.129) | n=1,118 med:0.068 (0.062-0.072) |
| Bolivian squirrel monkey | n=752 med:0.188 (0.169-0.205) | n=980 med:0.162 (0.147-0.175) | n=1,099 med:0.093 (0.086-0.101) |
| Crab-eating macaque | n=786 med:0.171 (0.160-0.186) | n=1,038 med:0.149 (0.138-0.163) | n=1,148 med:0.097 (0.090-0.106) |
| Red fox | n=739 med:0.143 (0.134-0.150) | n=929 med:0.134 (0.126-0.142) | n=1,037 med:0.089 (0.084-0.096) |
| Shrew mouse | n=777 med:0.114 (0.106-0.122) | n=1,006 med:0.119 (0.113-0.126) | n=1,138 med:0.064 (0.058-0.069) |
| Arctic ground squirrel | n=769 med:0.134 (0.123-0.144) | n=999 med:0.122 (0.114-0.130) | n=1,115 med:0.065 (0.059-0.070) |
| Chinese hamster CriGri | n=714 med:0.125 (0.118-0.133) | n=920 med:0.127 (0.119-0.133) | n=1,012 med:0.072 (0.067-0.079) |
| Opossum | n=663 med:0.096 (0.091-0.101) | n=848 med:0.085 (0.079-0.092) | n=974 med:0.065 (0.061-0.069) |
| Dolphin | n=437 | n=511 | n=527 |

| | | | |
|---|---|---|---|
| | med:0.161 (0.147-0.173) | med:0.152 (0.140-0.166) | med:0.103 (0.090-0.112) |
| Capuchin | n=765 med:0.181 (0.166-0.197) | n=1,020 med:0.155 (0.145-0.172) | n=1,130 med:0.085 (0.077-0.091) |
| Mouse Lemur | n=755 med:0.147 (0.132-0.160) | n=984 med:0.136 (0.126-0.145) | n=1,101 med:0.075 (0.069-0.081) |
| Damara mole rat | n=695 med:0.137 (0.127-0.147) | n=899 med:0.129 (0.118-0.136) | n=1,017 med:0.071 (0.067-0.075) |
| Marmoset | n=754 med:0.185 (0.170-0.200) | n=995 med:0.164 (0.152-0.175) | n=1,120 med:0.095 (0.086-0.104) |
| Goat | n=759 med:0.137 (0.127-0.143) | n=976 med:0.131 (0.122-0.138) | n=1,117 med:0.075 (0.069-0.081) |
| Common wombat | n=649 med:0.095 (0.088-0.103) | n=830 med:0.090 (0.086-0.094) | n=927 med:0.061 (0.057-0.066) |
| American black bear | n=678 med:0.147 (0.136-0.157) | n=890 med:0.133 (0.124-0.141) | n=902 med:0.097 (0.087-0.106) |
| Sloth | n=94 med:0.224 (0.185-0.256) | n=134 med:0.209 (0.157-0.271) | n=114 med:0.125 (0.102-0.150) |
| Polar bear | n=670 med:0.155 (0.143-0.166) | n=854 med:0.150 (0.143-0.159) | n=886 med:0.106 (0.100-0.115) |
| Alpine marmot | n=732 med:0.135 (0.122-0.149) | n=944 med:0.129 (0.118-0.137) | n=1,017 med:0.075 (0.069-0.081) |
| Prairie vole | n=744 med:0.114 (0.104-0.123) | n=977 med:0.120 (0.110-0.125) | n=1,107 med:0.063 (0.058-0.068) |
| Donkey | n=755 med:0.149 (0.139-0.162) | n=1,002 med:0.141 (0.130-0.149) | n=1,121 med:0.087 (0.080-0.093) |
| Golden Hamster | n=710 med:0.122 (0.113-0.132) | n=921 med:0.125 (0.117-0.131) | n=1,055 med:0.068 (0.062-0.072) |
| Armadillo | n=705 med:0.153 (0.140-0.163) | n=931 med:0.142 (0.131-0.151) | n=991 med:0.089 (0.082-0.098) |
| Ugandan red Colobus | n=770 med:0.177 (0.164-0.196) | n=1,012 med:0.159 (0.148-0.173) | n=1,135 med:0.091 (0.082-0.097) |
| Pig-tailed macaque | n=794 | n=1,033 | n=1,155 |

| | | | |
|---|---|---|---|
| | med:0.179 (0.166-0.196) | med:0.159 (0.147-0.173) | med:0.098 (0.091-0.107) |
| Panda | n=765 med:0.136 (0.128-0.145) | n=997 med:0.131 (0.123-0.140) | n=1,081 med:0.076 (0.068-0.082) |
| Ferret | n=746 med:0.123 (0.114-0.136) | n=990 med:0.121 (0.114-0.132) | n=1,080 med:0.072 (0.067-0.078) |
| Koala | n=686 med:0.100 (0.092-0.107) | n=887 med:0.092 (0.086-0.097) | n=1,018 med:0.065 (0.061-0.069) |
| Wallaby | n=199 med:0.088 (0.075-0.102) | n=290 med:0.084 (0.079-0.098) | n=280 med:0.063 (0.053-0.072) |
| Tarsier | n=639 med:0.147 (0.135-0.163) | n=818 med:0.147 (0.135-0.157) | n=874 med:0.093 (0.085-0.100) |
| Degu | n=652 med:0.127 (0.121-0.136) | n=890 med:0.120 (0.112-0.127) | n=1,033 med:0.067 (0.060-0.072) |
| Chinese hamster CHOK1GS | n=781 med:0.124 (0.115-0.132) | n=1,019 med:0.122 (0.113-0.128) | n=1,145 med:0.067 (0.062-0.072) |
| Drill | n=750 med:0.182 (0.170-0.197) | n=1,002 med:0.176 (0.163-0.188) | n=1,086 med:0.113 (0.102-0.121) |
| Gibbon | n=754 med:0.213 (0.199-0.225) | n=998 med:0.220 (0.201-0.236) | n=1,100 med:0.156 (0.139-0.166) |
| Rat | n=740 med:0.116 (0.107-0.124) | n=979 med:0.119 (0.112-0.126) | n=1,097 med:0.065 (0.060-0.071) |
| Macaque | n=786 med:0.177 (0.161-0.193) | n=1,036 med:0.178 (0.162-0.196) | n=1,151 med:0.100 (0.089-0.108) |
| American mink | n=743 med:0.125 (0.114-0.136) | n=975 med:0.119 (0.112-0.126) | n=1,051 med:0.071 (0.067-0.076) |
| Gelada | n=787 med:0.155 (0.135-0.169) | n=1,043 med:0.147 (0.136-0.162) | n=1,173 med:0.084 (0.075-0.092) |
| Sheep | n=731 med:0.146 (0.135-0.156) | n=940 med:0.141 (0.136-0.148) | n=1,056 med:0.092 (0.084-0.098) |
| Tasmanian devil | n=639 med:0.094 (0.085-0.103) | n=831 med:0.089 (0.084-0.095) | n=893 med:0.060 (0.057-0.064) |
| Elephant | n=745 | n=958 | n=1,069 |

| | | | |
|---|---|---|---|
| | med:0.146 (0.133-0.157) | med:0.138 (0.128-0.148) | med:0.087 (0.078-0.094) |
| Algerian mouse | n=777 med:0.116 (0.109-0.122) | n=1,018 med:0.117 (0.111-0.124) | n=1,151 med:0.064 (0.059-0.069) |
| Dingo | n=753 med:0.135 (0.124-0.144) | n=960 med:0.129 (0.121-0.138) | n=1,075 med:0.079 (0.072-0.085) |
| Golden snub-nosed monkey | n=781 med:0.184 (0.170-0.200) | n=997 med:0.165 (0.154-0.182) | n=1,074 med:0.106 (0.097-0.114) |
| Horse | n=765 med:0.151 (0.138-0.159) | n=993 med:0.142 (0.133-0.152) | n=1,142 med:0.081 (0.074-0.088) |
| Black snub-nosed monkey | n=765 med:0.195 (0.179-0.210) | n=991 med:0.175 (0.163-0.190) | n=1,094 med:0.125 (0.116-0.136) |
| Brazilian guinea pig | n=529 med:0.143 (0.132-0.151) | n=730 med:0.137 (0.128-0.144) | n=785 med:0.094 (0.086-0.104) |
| Orangutan | n=763 med:0.182 (0.164-0.199) | n=997 med:0.171 (0.157-0.190) | n=1,100 med:0.124 (0.113-0.136) |
| Northern American deer mouse | n=762 med:0.117 (0.107-0.125) | n=993 med:0.116 (0.110-0.123) | n=1,068 med:0.063 (0.058-0.067) |
| Naked mole-rat female | n=726 med:0.140 (0.130-0.152) | n=973 med:0.132 (0.124-0.142) | n=1,090 med:0.073 (0.067-0.079) |
| Mouse | n=811 med:0.114 (0.108-0.123) | n=1,046 med:0.117 (0.112-0.123) | n=1,181 med:0.063 (0.059-0.068) |
| Daurian ground squirrel | n=663 med:0.150 (0.139-0.163) | n=808 med:0.140 (0.129-0.148) | n=860 med:0.088 (0.077-0.096) |
| Tiger | n=622 med:0.152 (0.142-0.165) | n=793 med:0.147 (0.137-0.158) | n=835 med:0.091 (0.081-0.099) |
| Lesser hedgehog tenrec | n=194 med:0.171 (0.156-0.206) | n=229 med:0.155 (0.134-0.173) | n=213 med:0.097 (0.079-0.111) |
| Mongolian gerbil | n=696 med:0.119 (0.112-0.129) | n=897 med:0.119 (0.112-0.131) | n=946 med:0.070 (0.063-0.077) |
| Ma's night monkey | n=758 med:0.195 (0.182-0.212) | n=1,003 med:0.174 (0.156-0.189) | n=1,121 med:0.103 (0.096-0.113) |

| | | | |
|---|---|---|---|
| Upper Galilee mountains blind mole rat | n=729 med:0.134 (0.121-0.142) | n=967 med:0.126 (0.118-0.134) | n=1,113 med:0.066 (0.060-0.071) |
| Kangaroo rat | n=666 med:0.119 (0.110-0.126) | n=875 med:0.115 (0.108-0.125) | n=1,012 med:0.070 (0.065-0.076) |
| Chimpanzee | n=766 med:0.215 (0.188-0.239) | n=981 med:0.199 (0.180-0.214) | n=1,108 med:0.134 (0.111-0.152) |
| Guinea Pig | n=751 med:0.129 (0.118-0.139) | n=967 med:0.125 (0.116-0.131) | n=1,065 med:0.074 (0.067-0.081) |
| American beaver | n=726 med:0.142 (0.129-0.155) | n=926 med:0.134 (0.124-0.144) | n=983 med:0.082 (0.075-0.087) |
| Wild yak | n=687 med:0.142 (0.133-0.155) | n=866 med:0.137 (0.130-0.147) | n=901 med:0.081 (0.074-0.088) |
| Coquerel's sifaka | n=739 med:0.151 (0.139-0.164) | n=961 med:0.136 (0.129-0.146) | n=1,090 med:0.080 (0.071-0.086) |
| Alpaca | n=152 med:0.195 (0.175-0.217) | n=151 med:0.200 (0.165-0.271) | n=156 med:0.118 (0.099-0.140) |
| Lesser Egyptian jerboa | n=665 med:0.118 (0.109-0.129) | n=884 med:0.122 (0.115-0.129) | n=1,012 med:0.065 (0.059-0.071) |
| Long-tailed chinchilla | n=734 med:0.123 (0.110-0.134) | n=975 med:0.123 (0.116-0.132) | n=1,093 med:0.069 (0.064-0.073) |
| Angola colobus | n=774 med:0.193 (0.175-0.207) | n=999 med:0.169 (0.159-0.184) | n=1,103 med:0.105 (0.096-0.114) |
| Ryukyu mouse | n=777 med:0.117 (0.109-0.127) | n=995 med:0.119 (0.113-0.124) | n=1,148 med:0.064 (0.059-0.068) |
| Chinese hamster PICR | n=784 med:0.130 (0.120-0.139) | n=1,003 med:0.124 (0.116-0.131) | n=1,123 med:0.067 (0.061-0.072) |
| Vervet-AGM | n=798 med:0.165 (0.155-0.180) | n=1,037 med:0.156 (0.143-0.172) | n=1,140 med:0.091 (0.084-0.098) |
| Cow | n=759 med:0.131 (0.122-0.142) | n=983 med:0.130 (0.120-0.139) | n=1,108 med:0.073 (0.067-0.082) |
| Bushbaby | n=764 med:0.141 (0.130-0.150) | n=1,009 med:0.135 (0.125-0.144) | n=1,139 med:0.074 (0.067-0.081) |

| | | | |
|---|---|---|---|
| Bonobo | n=763<br>med:0.234<br>(0.207-0.266) | n=993<br>med:0.217<br>(0.199-0.234) | n=1,083<br>med:0.182<br>(0.165-0.206) |
| Hyrax | n=216<br>med:0.138<br>(0.119-0.162) | n=290<br>med:0.146<br>(0.130-0.165) | n=277<br>med:0.097<br>(0.084-0.118) |
| Rabbit | n=667<br>med:0.133<br>(0.121-0.144) | n=871<br>med:0.126<br>(0.119-0.135) | n=957<br>med:0.084<br>(0.078-0.091) |
| Megabat | n=399<br>med:0.156<br>(0.146-0.172) | n=526<br>med:0.158<br>(0.147-0.167) | n=500<br>med:0.101<br>(0.088-0.116) |
| Dog | n=731<br>med:0.147<br>(0.134-0.154) | n=961<br>med:0.136<br>(0.127-0.143) | n=1,064<br>med:0.088<br>(0.083-0.094) |
| Steppe mouse | n=749<br>med:0.120<br>(0.112-0.131) | n=993<br>med:0.121<br>(0.115-0.127) | n=1,103<br>med:0.070<br>(0.065-0.075) |
| Greater bamboo lemur | n=779<br>med:0.149<br>(0.139-0.161) | n=1,005<br>med:0.141<br>(0.131-0.149) | n=1,114<br>med:0.080<br>(0.074-0.087) |
| Pig | n=764<br>med:0.134<br>(0.126-0.143) | n=984<br>med:0.132<br>(0.122-0.141) | n=1,111<br>med:0.073<br>(0.067-0.079) |
| Tree Shrew | n=171<br>med:0.163<br>(0.135-0.176) | n=193<br>med:0.157<br>(0.133-0.177) | n=165<br>med:0.106<br>(0.089-0.120) |
| Pika | n=268<br>med:0.113<br>(0.101-0.132) | n=358<br>med:0.131<br>(0.119-0.147) | n=328<br>med:0.082<br>(0.072-0.088) |
| Shrew | n=141<br>med:0.138<br>(0.117-0.159) | n=164<br>med:0.116<br>(0.097-0.154) | n=161<br>med:0.090<br>(0.072-0.107) |
| Squirrel | n=755<br>med:0.133<br>(0.120-0.146) | n=969<br>med:0.120<br>(0.109-0.126) | n=1,122<br>med:0.066<br>(0.063-0.071) |
| Gorilla | n=769<br>med:0.212<br>(0.191-0.233) | n=1,010<br>med:0.202<br>(0.184-0.222) | n=1,119<br>med:0.130<br>(0.122-0.146) |
| Cat | n=774<br>med:0.140<br>(0.131-0.150) | n=999<br>med:0.132<br>(0.120-0.142) | n=1,112<br>med:0.076<br>(0.071-0.084) |

#### Human Reference Genome, Zhang et al., 2014, cell subtypes

| Sample Size and Median dN/dS (CI95%) | Microglia-specific Genes | Endothelia-specific Genes | Astrocyte-specific Genes | Oligodendrocyte-specific Genes | Neuron-specific Genes |
|---|---|---|---|---|---|
| Microbat | n=707 med:0.151 (0.140-0.165) | n=665 med:0.145 (0.136-0.163) | n=606 med:0.125 (0.116-0.134) | n=375 med:0.113 (0.101-0.126) | n=928 med:0.095 (0.086-0.104) |
| Hedgehog | n=211 med:0.161 (0.143-0.177) | n=172 med:0.157 (0.129-0.179) | n=129 med:0.124 (0.090-0.153) | n=94 med:0.104 (0.080-0.120) | n=224 med:0.097 (0.075-0.109) |
| Naked mole-rat male | n=760 med:0.143 (0.129-0.151) | n=668 med:0.143 (0.131-0.154) | n=635 med:0.123 (0.114-0.132) | n=411 med:0.096 (0.089-0.107) | n=998 med:0.078 (0.070-0.084) |
| Sooty mangabey | n=880 med:0.178 (0.162-0.194) | n=782 med:0.173 (0.157-0.187) | n=739 med:0.155 (0.142-0.169) | n=472 med:0.118 (0.102-0.134) | n=1,152 med:0.098 (0.090-0.107) |
| Olive baboon | n=887 med:0.174 (0.160-0.192) | n=787 med:0.183 (0.169-0.197) | n=733 med:0.164 (0.148-0.177) | n=469 med:0.131 (0.119-0.143) | n=1,150 med:0.103 (0.093-0.112) |
| American bison | n=727 med:0.149 (0.136-0.157) | n=638 med:0.143 (0.131-0.157) | n=582 med:0.118 (0.107-0.131) | n=376 med:0.103 (0.094-0.113) | n=863 med:0.080 (0.073-0.087) |
| Leopard | n=827 med:0.134 (0.125-0.147) | n=749 med:0.129 (0.121-0.139) | n=716 med:0.106 (0.098-0.115) | n=444 med:0.084 (0.075-0.090) | n=1,118 med:0.068 (0.062-0.072) |
| Bolivian squirrel monkey | n=847 med:0.182 (0.171-0.197) | n=752 med:0.188 (0.169-0.205) | n=700 med:0.147 (0.135-0.159) | n=447 med:0.111 (0.099-0.125) | n=1,099 med:0.093 (0.086-0.101) |
| Crab-eating macaque | n=884 med:0.165 (0.152-0.179) | n=786 med:0.171 (0.160-0.186) | n=733 med:0.151 (0.137-0.163) | n=467 med:0.118 (0.101-0.133) | n=1,148 med:0.097 (0.090-0.106) |
| Red fox | n=823 med:0.149 (0.139-0.160) | n=739 med:0.143 (0.134-0.150) | n=659 med:0.120 (0.110-0.130) | n=427 med:0.102 (0.093-0.109) | n=1,037 med:0.089 (0.084-0.096) |
| Shrew mouse | n=848 med:0.132 (0.124-0.140) | n=777 med:0.114 (0.106-0.122) | n=725 med:0.100 (0.092-0.108) | n=452 med:0.085 (0.076-0.094) | n=1,138 med:0.064 (0.058-0.069) |
| Arctic ground squirrel | n=839 med:0.137 (0.127-0.147) | n=769 med:0.134 (0.123-0.144) | n=723 med:0.105 (0.098-0.116) | n=458 med:0.088 (0.081-0.097) | n=1,115 med:0.065 (0.059-0.070) |
| Chinese hamster CriGri | n=800 med:0.135 (0.128-0.146) | n=714 med:0.125 (0.118-0.133) | n=662 med:0.111 (0.100-0.122) | n=413 med:0.093 (0.082-0.101) | n=1,012 med:0.072 (0.067-0.079) |
| Opossum | n=712 med:0.091 (0.083-0.099) | n=663 med:0.096 (0.091-0.101) | n=632 med:0.091 (0.082-0.097) | n=403 med:0.074 (0.069-0.081) | n=974 med:0.065 (0.061-0.069) |
| Dolphin | n=467 med:0.165 | n=437 med:0.161 | n=342 med:0.138 | n=224 med:0.124 | n=527 med:0.103 |

| | | | | | |
|---|---|---|---|---|---|
| | (0.149-0.179) | (0.147-0.173) | (0.123-0.153) | (0.103-0.135) | (0.090-0.112) |
| Capuchin | n=863<br>med:0.180<br>(0.163-0.193) | n=765<br>med:0.181<br>(0.166-0.197) | n=713<br>med:0.146<br>(0.134-0.162) | n=463<br>med:0.116<br>(0.099-0.126) | n=1,130<br>med:0.085<br>(0.077-0.091) |
| Mouse Lemur | n=832<br>med:0.148<br>(0.139-0.158) | n=755<br>med:0.147<br>(0.132-0.160) | n=686<br>med:0.123<br>(0.114-0.140) | n=452<br>med:0.094<br>(0.084-0.106) | n=1,101<br>med:0.075<br>(0.069-0.081) |
| Damara mole rat | n=744<br>med:0.131<br>(0.119-0.141) | n=695<br>med:0.137<br>(0.127-0.147) | n=652<br>med:0.115<br>(0.106-0.124) | n=426<br>med:0.094<br>(0.085-0.102) | n=1,017<br>med:0.071<br>(0.067-0.075) |
| Marmoset | n=841<br>med:0.186<br>(0.169-0.201) | n=754<br>med:0.185<br>(0.170-0.200) | n=708<br>med:0.157<br>(0.143-0.167) | n=455<br>med:0.126<br>(0.114-0.137) | n=1,120<br>med:0.095<br>(0.086-0.104) |
| Goat | n=842<br>med:0.136<br>(0.127-0.146) | n=759<br>med:0.137<br>(0.127-0.143) | n=705<br>med:0.111<br>(0.103-0.123) | n=457<br>med:0.096<br>(0.086-0.105) | n=1,117<br>med:0.075<br>(0.069-0.081) |
| Common wombat | n=708<br>med:0.098<br>(0.089-0.104) | n=649<br>med:0.095<br>(0.088-0.103) | n=616<br>med:0.089<br>(0.082-0.094) | n=387<br>med:0.080<br>(0.073-0.085) | n=927<br>med:0.061<br>(0.057-0.066) |
| American black bear | n=789<br>med:0.143<br>(0.133-0.151) | n=678<br>med:0.147<br>(0.136-0.157) | n=593<br>med:0.129<br>(0.118-0.142) | n=381<br>med:0.107<br>(0.096-0.119) | n=902<br>med:0.097<br>(0.087-0.106) |
| Sloth | n=106<br>med:0.256<br>(0.195-0.314) | n=94<br>med:0.224<br>(0.185-0.256) | n=96<br>med:0.142<br>(0.112-0.174) | n=62<br>med:0.156<br>(0.105-0.220) | n=114<br>med:0.125<br>(0.102-0.150) |
| Polar bear | n=753<br>med:0.161<br>(0.152-0.173) | n=670<br>med:0.155<br>(0.143-0.166) | n=606<br>med:0.133<br>(0.124-0.146) | n=382<br>med:0.114<br>(0.105-0.123) | n=886<br>med:0.106<br>(0.100-0.115) |
| Alpine marmot | n=833<br>med:0.140<br>(0.133-0.150) | n=732<br>med:0.135<br>(0.122-0.149) | n=675<br>med:0.110<br>(0.100-0.119) | n=419<br>med:0.096<br>(0.085-0.105) | n=1,017<br>med:0.075<br>(0.069-0.081) |
| Prairie vole | n=832<br>med:0.127<br>(0.120-0.138) | n=744<br>med:0.114<br>(0.104-0.123) | n=685<br>med:0.097<br>(0.089-0.107) | n=440<br>med:0.080<br>(0.073-0.093) | n=1,107<br>med:0.063<br>(0.058-0.068) |
| Donkey | n=871<br>med:0.154<br>(0.144-0.163) | n=755<br>med:0.149<br>(0.139-0.162) | n=704<br>med:0.128<br>(0.115-0.140) | n=450<br>med:0.109<br>(0.098-0.118) | n=1,121<br>med:0.087<br>(0.080-0.093) |
| Golden Hamster | n=780<br>med:0.136<br>(0.128-0.148) | n=710<br>med:0.122<br>(0.113-0.132) | n=664<br>med:0.104<br>(0.095-0.113) | n=414<br>med:0.097<br>(0.084-0.105) | n=1,055<br>med:0.068<br>(0.062-0.072) |
| Armadillo | n=782<br>med:0.153<br>(0.142-0.161) | n=705<br>med:0.153<br>(0.140-0.163) | n=663<br>med:0.132<br>(0.118-0.143) | n=425<br>med:0.106<br>(0.097-0.117) | n=991<br>med:0.089<br>(0.082-0.098) |
| Ugandan red Colobus | n=859<br>med:0.186<br>(0.172-0.202) | n=770<br>med:0.177<br>(0.164-0.196) | n=711<br>med:0.146<br>(0.135-0.159) | n=466<br>med:0.119<br>(0.106-0.131) | n=1,135<br>med:0.091<br>(0.082-0.097) |
| Pig-tailed macaque | n=883<br>med:0.176<br>(0.159-0.189) | n=794<br>med:0.179<br>(0.166-0.196) | n=730<br>med:0.155<br>(0.136-0.165) | n=476<br>med:0.118<br>(0.105-0.135) | n=1,155<br>med:0.098<br>(0.091-0.107) |
| Panda | n=851 | n=765 | n=684 | n=448 | n=1,081 |

| | | | | | |
|---|---|---|---|---|---|
| | med:0.141<br>(0.131-0.151) | med:0.136<br>(0.128-0.145) | med:0.112<br>(0.104-0.121) | med:0.097<br>(0.087-0.109) | med:0.076<br>(0.068-0.082) |
| Ferret | n=856<br>med:0.131<br>(0.120-0.143) | n=746<br>med:0.123<br>(0.114-0.136) | n=683<br>med:0.108<br>(0.096-0.118) | n=441<br>med:0.087<br>(0.078-0.096) | n=1,080<br>med:0.072<br>(0.067-0.078) |
| Koala | n=753<br>med:0.098<br>(0.090-0.108) | n=686<br>med:0.100<br>(0.092-0.107) | n=648<br>med:0.091<br>(0.084-0.099) | n=420<br>med:0.078<br>(0.071-0.083) | n=1,018<br>med:0.065<br>(0.061-0.069) |
| Wallaby | n=268<br>med:0.085<br>(0.074-0.096) | n=199<br>med:0.088<br>(0.075-0.102) | n=176<br>med:0.083<br>(0.072-0.097) | n=139<br>med:0.080<br>(0.067-0.084) | n=280<br>med:0.063<br>(0.053-0.072) |
| Tarsier | n=704<br>med:0.156<br>(0.147-0.165) | n=639<br>med:0.147<br>(0.135-0.163) | n=594<br>med:0.129<br>(0.116-0.144) | n=362<br>med:0.110<br>(0.092-0.122) | n=874<br>med:0.093<br>(0.085-0.100) |
| Degu | n=730<br>med:0.125<br>(0.116-0.135) | n=652<br>med:0.127<br>(0.121-0.136) | n=628<br>med:0.107<br>(0.098-0.119) | n=418<br>med:0.088<br>(0.078-0.094) | n=1,033<br>med:0.067<br>(0.060-0.072) |
| Chinese hamster CHOK1GS | n=856<br>med:0.134<br>(0.125-0.141) | n=781<br>med:0.124<br>(0.115-0.132) | n=733<br>med:0.107<br>(0.097-0.116) | n=458<br>med:0.089<br>(0.081-0.096) | n=1,145<br>med:0.067<br>(0.062-0.072) |
| Drill | n=872<br>med:0.184<br>(0.173-0.197) | n=750<br>med:0.182<br>(0.170-0.197) | n=706<br>med:0.167<br>(0.156-0.181) | n=445<br>med:0.122<br>(0.115-0.142) | n=1,086<br>med:0.113<br>(0.102-0.121) |
| Gibbon | n=848<br>med:0.239<br>(0.224-0.252) | n=754<br>med:0.213<br>(0.199-0.225) | n=704<br>med:0.208<br>(0.192-0.228) | n=447<br>med:0.171<br>(0.151-0.189) | n=1,100<br>med:0.156<br>(0.139-0.166) |
| Rat | n=836<br>med:0.128<br>(0.121-0.139) | n=740<br>med:0.116<br>(0.107-0.124) | n=702<br>med:0.105<br>(0.096-0.113) | n=441<br>med:0.088<br>(0.080-0.095) | n=1,097<br>med:0.065<br>(0.060-0.071) |
| Macaque | n=869<br>med:0.193<br>(0.174-0.210) | n=786<br>med:0.177<br>(0.161-0.193) | n=729<br>med:0.163<br>(0.151-0.175) | n=461<br>med:0.131<br>(0.117-0.147) | n=1,151<br>med:0.100<br>(0.089-0.108) |
| American mink | n=837<br>med:0.131<br>(0.121-0.144) | n=743<br>med:0.125<br>(0.114-0.136) | n=686<br>med:0.108<br>(0.096-0.118) | n=438<br>med:0.086<br>(0.079-0.094) | n=1,051<br>med:0.071<br>(0.067-0.076) |
| Gelada | n=883<br>med:0.159<br>(0.144-0.176) | n=787<br>med:0.155<br>(0.135-0.169) | n=733<br>med:0.143<br>(0.127-0.159) | n=470<br>med:0.107<br>(0.093-0.121) | n=1,173<br>med:0.084<br>(0.075-0.092) |
| Sheep | n=810<br>med:0.152<br>(0.144-0.163) | n=731<br>med:0.146<br>(0.135-0.156) | n=672<br>med:0.125<br>(0.114-0.136) | n=428<br>med:0.103<br>(0.097-0.113) | n=1,056<br>med:0.092<br>(0.084-0.098) |
| Tasmanian devil | n=696<br>med:0.094<br>(0.086-0.105) | n=639<br>med:0.094<br>(0.085-0.103) | n=581<br>med:0.089<br>(0.080-0.097) | n=381<br>med:0.074<br>(0.068-0.084) | n=893<br>med:0.060<br>(0.057-0.064) |
| Elephant | n=825<br>med:0.154<br>(0.145-0.163) | n=745<br>med:0.146<br>(0.133-0.157) | n=690<br>med:0.125<br>(0.114-0.133) | n=442<br>med:0.102<br>(0.090-0.111) | n=1,069<br>med:0.087<br>(0.078-0.094) |
| Algerian mouse | n=861<br>med:0.128<br>(0.120-0.140) | n=777<br>med:0.116<br>(0.109-0.122) | n=733<br>med:0.101<br>(0.092-0.110) | n=456<br>med:0.085<br>(0.076-0.095) | n=1,151<br>med:0.064<br>(0.059-0.069) |

| | | | | | |
|---|---|---|---|---|---|
| Dingo | n=835 med:0.138 (0.128-0.145) | n=753 med:0.135 (0.124-0.144) | n=679 med:0.109 (0.099-0.118) | n=430 med:0.091 (0.084-0.102) | n=1,075 med:0.079 (0.072-0.085) |
| Golden snub-nosed monkey | n=854 med:0.181 (0.167-0.193) | n=781 med:0.184 (0.170-0.200) | n=701 med:0.163 (0.146-0.172) | n=449 med:0.131 (0.119-0.141) | n=1,074 med:0.106 (0.097-0.114) |
| Horse | n=852 med:0.155 (0.146-0.166) | n=765 med:0.151 (0.138-0.159) | n=707 med:0.127 (0.111-0.140) | n=445 med:0.103 (0.095-0.115) | n=1,142 med:0.081 (0.074-0.088) |
| Black snub-nosed monkey | n=854 med:0.195 (0.183-0.211) | n=765 med:0.195 (0.179-0.210) | n=689 med:0.173 (0.160-0.189) | n=463 med:0.143 (0.129-0.160) | n=1,094 med:0.125 (0.116-0.136) |
| Brazilian guinea pig | n=614 med:0.140 (0.128-0.152) | n=529 med:0.143 (0.132-0.151) | n=510 med:0.137 (0.123-0.145) | n=351 med:0.110 (0.101-0.121) | n=785 med:0.094 (0.086-0.104) |
| Orangutan | n=850 med:0.190 (0.173-0.209) | n=763 med:0.182 (0.164-0.199) | n=702 med:0.180 (0.161-0.195) | n=458 med:0.143 (0.126-0.159) | n=1,100 med:0.124 (0.113-0.136) |
| Northern American deer mouse | n=834 med:0.132 (0.122-0.138) | n=762 med:0.117 (0.107-0.125) | n=712 med:0.101 (0.092-0.110) | n=445 med:0.085 (0.079-0.094) | n=1,068 med:0.063 (0.058-0.067) |
| Naked mole-rat female | n=802 med:0.143 (0.131-0.150) | n=726 med:0.140 (0.130-0.152) | n=673 med:0.119 (0.111-0.130) | n=441 med:0.095 (0.087-0.106) | n=1,090 med:0.073 (0.067-0.079) |
| Mouse | n=892 med:0.127 (0.120-0.138) | n=811 med:0.114 (0.108-0.123) | n=747 med:0.101 (0.093-0.111) | n=463 med:0.084 (0.076-0.095) | n=1,181 med:0.063 (0.059-0.068) |
| Daurian ground squirrel | n=712 med:0.151 (0.141-0.162) | n=663 med:0.150 (0.139-0.163) | n=581 med:0.123 (0.110-0.134) | n=366 med:0.103 (0.093-0.114) | n=860 med:0.088 (0.077-0.096) |
| Tiger | n=664 med:0.164 (0.154-0.176) | n=622 med:0.152 (0.142-0.165) | n=585 med:0.130 (0.116-0.137) | n=345 med:0.114 (0.098-0.128) | n=835 med:0.091 (0.081-0.099) |
| Lesser hedgehog tenrec | n=194 med:0.165 (0.137-0.187) | n=194 med:0.171 (0.156-0.206) | n=144 med:0.122 (0.107-0.154) | n=81 med:0.119 (0.089-0.142) | n=213 med:0.097 (0.079-0.111) |
| Mongolian gerbil | n=777 med:0.135 (0.125-0.144) | n=696 med:0.119 (0.112-0.129) | n=613 med:0.106 (0.101-0.116) | n=397 med:0.090 (0.083-0.100) | n=946 med:0.070 (0.063-0.077) |
| Ma's night monkey | n=844 med:0.194 (0.179-0.209) | n=758 med:0.195 (0.182-0.212) | n=708 med:0.155 (0.144-0.171) | n=453 med:0.123 (0.113-0.136) | n=1,121 med:0.103 (0.096-0.113) |
| Upper Galilee mountains blind mole rat | n=798 med:0.138 (0.126-0.145) | n=729 med:0.134 (0.121-0.142) | n=684 med:0.106 (0.096-0.115) | n=445 med:0.091 (0.081-0.099) | n=1,113 med:0.066 (0.060-0.071) |
| Kangaroo rat | n=726 med:0.128 (0.121-0.137) | n=666 med:0.119 (0.110-0.126) | n=629 med:0.106 (0.099-0.114) | n=420 med:0.084 (0.076-0.093) | n=1,012 med:0.070 (0.065-0.076) |
| Chimpanzee | n=827 | n=766 | n=702 | n=451 | n=1,108 |

| | | | | | |
|---|---|---|---|---|---|
| | med:0.204 (0.188-0.222) | med:0.215 (0.188-0.239) | med:0.198 (0.177-0.223) | med:0.144 (0.125-0.168) | med:0.134 (0.111-0.152) |
| Guinea Pig | n=825 med:0.133 (0.124-0.142) | n=751 med:0.129 (0.118-0.139) | n=680 med:0.117 (0.108-0.124) | n=440 med:0.088 (0.078-0.098) | n=1,065 med:0.074 (0.067-0.081) |
| American beaver | n=824 med:0.148 (0.138-0.158) | n=726 med:0.142 (0.129-0.155) | n=624 med:0.122 (0.111-0.132) | n=398 med:0.105 (0.095-0.115) | n=983 med:0.082 (0.075-0.087) |
| Wild yak | n=759 med:0.146 (0.136-0.155) | n=687 med:0.142 (0.133-0.155) | n=619 med:0.123 (0.111-0.131) | n=392 med:0.101 (0.090-0.109) | n=901 med:0.081 (0.074-0.088) |
| Coquerel's sifaka | n=820 med:0.150 (0.141-0.159) | n=739 med:0.151 (0.139-0.164) | n=691 med:0.128 (0.118-0.139) | n=436 med:0.098 (0.088-0.106) | n=1,090 med:0.080 (0.071-0.086) |
| Alpaca | n=118 med:0.227 (0.178-0.302) | n=152 med:0.195 (0.175-0.217) | n=154 med:0.148 (0.130-0.177) | n=57 med:0.169 (0.101-0.238) | n=156 med:0.118 (0.099-0.140) |
| Lesser Egyptian jerboa | n=731 med:0.129 (0.118-0.137) | n=665 med:0.118 (0.109-0.129) | n=636 med:0.106 (0.099-0.116) | n=419 med:0.094 (0.083-0.105) | n=1,012 med:0.065 (0.059-0.071) |
| Long-tailed chinchilla | n=821 med:0.131 (0.121-0.141) | n=734 med:0.123 (0.110-0.134) | n=692 med:0.108 (0.097-0.115) | n=450 med:0.084 (0.075-0.097) | n=1,093 med:0.069 (0.064-0.073) |
| Angola colobus | n=859 med:0.189 (0.173-0.201) | n=774 med:0.193 (0.175-0.207) | n=704 med:0.162 (0.140-0.171) | n=455 med:0.130 (0.118-0.143) | n=1,103 med:0.105 (0.096-0.114) |
| Ryukyu mouse | n=853 med:0.132 (0.124-0.140) | n=777 med:0.117 (0.109-0.127) | n=725 med:0.103 (0.092-0.109) | n=446 med:0.083 (0.077-0.094) | n=1,148 med:0.064 (0.059-0.068) |
| Chinese hamster PICR | n=849 med:0.137 (0.129-0.145) | n=784 med:0.130 (0.120-0.139) | n=717 med:0.107 (0.097-0.117) | n=447 med:0.089 (0.080-0.095) | n=1,123 med:0.067 (0.061-0.072) |
| Vervet-AGM | n=889 med:0.171 (0.155-0.183) | n=798 med:0.165 (0.155-0.180) | n=729 med:0.143 (0.131-0.161) | n=474 med:0.113 (0.100-0.129) | n=1,140 med:0.091 (0.084-0.098) |
| Cow | n=845 med:0.144 (0.132-0.152) | n=759 med:0.131 (0.122-0.142) | n=695 med:0.109 (0.101-0.120) | n=455 med:0.096 (0.086-0.104) | n=1,108 med:0.073 (0.067-0.082) |
| Bushbaby | n=856 med:0.153 (0.141-0.166) | n=764 med:0.141 (0.130-0.150) | n=705 med:0.116 (0.106-0.126) | n=458 med:0.094 (0.084-0.107) | n=1,139 med:0.074 (0.067-0.081) |
| Bonobo | n=854 med:0.226 (0.206-0.250) | n=763 med:0.234 (0.207-0.266) | n=703 med:0.239 (0.219-0.259) | n=458 med:0.186 (0.165-0.219) | n=1,083 med:0.182 (0.165-0.206) |
| Hyrax | n=298 med:0.159 (0.141-0.174) | n=216 med:0.138 (0.119-0.162) | n=161 med:0.127 (0.101-0.154) | n=118 med:0.121 (0.093-0.142) | n=277 med:0.097 (0.084-0.118) |
| Rabbit | n=734 med:0.139 (0.131-0.147) | n=667 med:0.133 (0.121-0.144) | n=635 med:0.114 (0.105-0.124) | n=399 med:0.088 (0.081-0.101) | n=957 med:0.084 (0.078-0.091) |

| | | | | | |
|---|---|---|---|---|---|
| Megabat | n=475 med:0.171 (0.158-0.185) | n=399 med:0.156 (0.146-0.172) | n=329 med:0.150 (0.137-0.172) | n=222 med:0.109 (0.095-0.132) | n=500 med:0.101 (0.088-0.116) |
| Dog | n=809 med:0.142 (0.132-0.152) | n=731 med:0.147 (0.134-0.154) | n=693 med:0.117 (0.108-0.129) | n=428 med:0.103 (0.092-0.111) | n=1,064 med:0.088 (0.083-0.094) |
| Steppe mouse | n=851 med:0.133 (0.125-0.142) | n=749 med:0.120 (0.112-0.131) | n=710 med:0.107 (0.099-0.117) | n=434 med:0.092 (0.084-0.102) | n=1,103 med:0.070 (0.065-0.075) |
| Greater bamboo lemur | n=858 med:0.151 (0.139-0.162) | n=779 med:0.149 (0.139-0.161) | n=711 med:0.132 (0.119-0.140) | n=456 med:0.103 (0.094-0.111) | n=1,114 med:0.080 (0.074-0.087) |
| Pig | n=858 med:0.149 (0.138-0.160) | n=764 med:0.134 (0.126-0.143) | n=706 med:0.117 (0.105-0.127) | n=451 med:0.092 (0.083-0.103) | n=1,111 med:0.073 (0.067-0.079) |
| Tree Shrew | n=176 med:0.162 (0.138-0.189) | n=171 med:0.163 (0.135-0.176) | n=120 med:0.136 (0.105-0.161) | n=77 med:0.154 (0.109-0.176) | n=165 med:0.106 (0.089-0.120) |
| Pika | n=324 med:0.147 (0.129-0.169) | n=268 med:0.113 (0.101-0.132) | n=225 med:0.115 (0.097-0.132) | n=142 med:0.093 (0.081-0.111) | n=328 med:0.082 (0.072-0.088) |
| Shrew | n=153 med:0.161 (0.117-0.196) | n=141 med:0.138 (0.117-0.159) | n=101 med:0.115 (0.088-0.141) | n=66 med:0.116 (0.089-0.160) | n=161 med:0.090 (0.072-0.107) |
| Squirrel | n=818 med:0.128 (0.121-0.137) | n=755 med:0.133 (0.120-0.146) | n=706 med:0.106 (0.098-0.116) | n=443 med:0.091 (0.080-0.097) | n=1,122 med:0.066 (0.063-0.071) |
| Gorilla | n=853 med:0.197 (0.177-0.218) | n=769 med:0.212 (0.191-0.233) | n=721 med:0.194 (0.167-0.213) | n=456 med:0.160 (0.141-0.189) | n=1,119 med:0.130 (0.122-0.146) |
| Cat | n=854 med:0.143 (0.130-0.156) | n=774 med:0.140 (0.131-0.150) | n=719 med:0.117 (0.104-0.126) | n=448 med:0.093 (0.086-0.101) | n=1,112 med:0.076 (0.071-0.084) |

#### Human Reference Genome, Zeisel et al., 2015, three cell types

| Sample Size and Median dN/dS (CI95%) | Glia-specific Genes | Vasculature-specific Genes | Neuron-specific Genes |
|---|---|---|---|
| Microbat | n=1,016 med:0.150 (0.141-0.158) | n=359 med:0.121 (0.112-0.139) | n=669 med:0.101 (0.091-0.111) |
| Hedgehog | n=238 med:0.147 (0.125-0.162) | n=79 med:0.135 (0.089-0.179) | n=146 med:0.109 (0.088-0.125) |
| Naked mole-rat male | n=992 med:0.147 (0.134-0.155) | n=341 med:0.112 (0.096-0.131) | n=665 med:0.085 (0.078-0.091) |
| Sooty mangabey | n=1,187 med:0.186 (0.168-0.200) | n=401 med:0.144 (0.124-0.162) | n=814 med:0.108 (0.092-0.118) |
| Olive baboon | n=1,192 med:0.181 (0.168-0.198) | n=398 med:0.165 (0.138-0.189) | n=806 med:0.120 (0.108-0.130) |
| American bison | n=982 med:0.143 (0.132-0.154) | n=332 med:0.115 (0.100-0.132) | n=623 med:0.081 (0.074-0.088) |
| Leopard | n=1,125 med:0.128 (0.119-0.135) | n=373 med:0.105 (0.095-0.118) | n=789 med:0.073 (0.065-0.079) |
| Bolivian squirrel monkey | n=1,130 med:0.178 (0.169-0.195) | n=383 med:0.163 (0.139-0.185) | n=777 med:0.097 (0.090-0.106) |
| Crab-eating macaque | n=1,192 med:0.178 (0.166-0.191) | n=399 med:0.143 (0.123-0.161) | n=800 med:0.105 (0.094-0.116) |
| Red fox | n=1,093 med:0.149 (0.140-0.157) | n=372 med:0.122 (0.108-0.139) | n=735 med:0.090 (0.085-0.100) |
| Shrew mouse | n=1,158 med:0.122 (0.114-0.131) | n=393 med:0.103 (0.094-0.118) | n=794 med:0.067 (0.060-0.075) |
| Arctic ground squirrel | n=1,123 med:0.132 (0.124-0.138) | n=398 med:0.110 (0.098-0.122) | n=804 med:0.071 (0.064-0.079) |
| Chinese hamster CriGri | n=1,066 med:0.131 (0.124-0.139) | n=365 med:0.112 (0.098-0.126) | n=722 med:0.078 (0.070-0.086) |
| Opossum | n=994 med:0.104 (0.097-0.111) | n=344 med:0.087 (0.077-0.097) | n=693 med:0.069 (0.064-0.073) |
| Dolphin | n=601 | n=195 | n=397 |

| | | | |
|---|---|---|---|
| | med:0.158<br>(0.146-0.177) | med:0.135<br>(0.118-0.165) | med:0.098<br>(0.088-0.109) |
| Capuchin | n=1,147<br>med:0.176<br>(0.160-0.189) | n=392<br>med:0.158<br>(0.136-0.172) | n=789<br>med:0.092<br>(0.087-0.103) |
| Mouse Lemur | n=1,137<br>med:0.148<br>(0.139-0.156) | n=381<br>med:0.125<br>(0.105-0.141) | n=773<br>med:0.082<br>(0.076-0.088) |
| Damara mole rat | n=1,047<br>med:0.132<br>(0.124-0.143) | n=362<br>med:0.112<br>(0.099-0.130) | n=724<br>med:0.076<br>(0.069-0.085) |
| Marmoset | n=1,136<br>med:0.188<br>(0.173-0.201) | n=395<br>med:0.153<br>(0.133-0.174) | n=805<br>med:0.101<br>(0.091-0.110) |
| Goat | n=1,152<br>med:0.136<br>(0.128-0.141) | n=381<br>med:0.116<br>(0.103-0.131) | n=772<br>med:0.080<br>(0.074-0.086) |
| Common wombat | n=947<br>med:0.104<br>(0.097-0.111) | n=333<br>med:0.087<br>(0.078-0.101) | n=670<br>med:0.066<br>(0.060-0.071) |
| American black bear | n=1,032<br>med:0.146<br>(0.135-0.158) | n=347<br>med:0.130<br>(0.111-0.147) | n=650<br>med:0.095<br>(0.085-0.104) |
| Sloth | n=172<br>med:0.219<br>(0.175-0.261) | n=49<br>med:0.187<br>(0.129-0.268) | n=87<br>med:0.129<br>(0.093-0.152) |
| Polar bear | n=1,025<br>med:0.154<br>(0.147-0.164) | n=335<br>med:0.139<br>(0.120-0.155) | n=641<br>med:0.103<br>(0.093-0.111) |
| Alpine marmot | n=1,071<br>med:0.137<br>(0.128-0.143) | n=376<br>med:0.111<br>(0.098-0.128) | n=730<br>med:0.075<br>(0.069-0.085) |
| Prairie vole | n=1,107<br>med:0.121<br>(0.115-0.129) | n=374<br>med:0.094<br>(0.081-0.105) | n=779<br>med:0.065<br>(0.060-0.073) |
| Donkey | n=1,154<br>med:0.154<br>(0.145-0.165) | n=390<br>med:0.122<br>(0.109-0.138) | n=792<br>med:0.095<br>(0.088-0.104) |
| Golden Hamster | n=1,057<br>med:0.126<br>(0.118-0.133) | n=361<br>med:0.110<br>(0.094-0.126) | n=751<br>med:0.071<br>(0.064-0.078) |
| Armadillo | n=1,062<br>med:0.152<br>(0.143-0.160) | n=359<br>med:0.133<br>(0.114-0.148) | n=716<br>med:0.094<br>(0.086-0.100) |
| Ugandan red Colobus | n=1,150<br>med:0.179<br>(0.171-0.192) | n=376<br>med:0.149<br>(0.128-0.169) | n=796<br>med:0.098<br>(0.090-0.110) |
| Pig-tailed macaque | n=1,181 | n=402 | n=812 |

| | | | |
|---|---|---|---|
| | med:0.184<br>(0.171-0.194) | med:0.153<br>(0.134-0.172) | med:0.108<br>(0.096-0.120) |
| Panda | n=1,134<br>med:0.132<br>(0.125-0.142) | n=380<br>med:0.114<br>(0.106-0.128) | n=761<br>med:0.079<br>(0.071-0.087) |
| Ferret | n=1,131<br>med:0.133<br>(0.124-0.143) | n=381<br>med:0.109<br>(0.093-0.122) | n=766<br>med:0.078<br>(0.070-0.084) |
| Koala | n=1,031<br>med:0.108<br>(0.102-0.115) | n=354<br>med:0.087<br>(0.080-0.096) | n=734<br>med:0.070<br>(0.063-0.076) |
| Wallaby | n=284<br>med:0.102<br>(0.088-0.110) | n=111<br>med:0.089<br>(0.076-0.103) | n=204<br>med:0.066<br>(0.056-0.076) |
| Tarsier | n=961<br>med:0.147<br>(0.141-0.162) | n=306<br>med:0.127<br>(0.106-0.148) | n=630<br>med:0.100<br>(0.089-0.109) |
| Degu | n=983<br>med:0.128<br>(0.121-0.138) | n=343<br>med:0.113<br>(0.097-0.126) | n=684<br>med:0.074<br>(0.067-0.083) |
| Chinese hamster CHOK1GS | n=1,152<br>med:0.129<br>(0.121-0.136) | n=399<br>med:0.108<br>(0.094-0.121) | n=800<br>med:0.071<br>(0.066-0.078) |
| Drill | n=1,153<br>med:0.194<br>(0.182-0.205) | n=380<br>med:0.154<br>(0.142-0.179) | n=767<br>med:0.121<br>(0.111-0.130) |
| Gibbon | n=1,149<br>med:0.245<br>(0.235-0.265) | n=387<br>med:0.212<br>(0.186-0.223) | n=775<br>med:0.169<br>(0.153-0.189) |
| Rat | n=1,131<br>med:0.122<br>(0.115-0.128) | n=378<br>med:0.105<br>(0.093-0.117) | n=774<br>med:0.073<br>(0.063-0.079) |
| Macaque | n=1,163<br>med:0.192<br>(0.179-0.207) | n=397<br>med:0.147<br>(0.132-0.168) | n=804<br>med:0.108<br>(0.099-0.120) |
| American mink | n=1,092<br>med:0.131<br>(0.122-0.139) | n=370<br>med:0.104<br>(0.091-0.120) | n=744<br>med:0.078<br>(0.071-0.087) |
| Gelada | n=1,190<br>med:0.167<br>(0.155-0.178) | n=398<br>med:0.121<br>(0.099-0.139) | n=813<br>med:0.093<br>(0.084-0.102) |
| Sheep | n=1,115<br>med:0.145<br>(0.140-0.155) | n=363<br>med:0.127<br>(0.115-0.139) | n=736<br>med:0.092<br>(0.084-0.100) |
| Tasmanian devil | n=950<br>med:0.102<br>(0.097-0.109) | n=333<br>med:0.085<br>(0.075-0.093) | n=629<br>med:0.068<br>(0.061-0.074) |
| Elephant | n=1,131 | n=376 | n=751 |

| | | | |
|---|---|---|---|
| | med:0.147 (0.138-0.156) | med:0.123 (0.113-0.135) | med:0.094 (0.084-0.101) |
| Algerian mouse | n=1,162 med:0.123 (0.115-0.130) | n=394 med:0.100 (0.090-0.113) | n=807 med:0.068 (0.060-0.072) |
| Dingo | n=1,097 med:0.136 (0.128-0.144) | n=368 med:0.115 (0.103-0.124) | n=750 med:0.080 (0.072-0.088) |
| Golden snub-nosed monkey | n=1,139 med:0.184 (0.171-0.198) | n=381 med:0.163 (0.138-0.181) | n=772 med:0.116 (0.105-0.128) |
| Horse | n=1,147 med:0.157 (0.145-0.167) | n=394 med:0.122 (0.111-0.135) | n=787 med:0.088 (0.082-0.096) |
| Black snub-nosed monkey | n=1,149 med:0.207 (0.185-0.221) | n=375 med:0.182 (0.165-0.200) | n=773 med:0.139 (0.125-0.154) |
| Brazilian guinea pig | n=829 med:0.142 (0.135-0.150) | n=292 med:0.126 (0.112-0.137) | n=555 med:0.106 (0.100-0.114) |
| Orangutan | n=1,144 med:0.207 (0.193-0.223) | n=390 med:0.160 (0.135-0.184) | n=771 med:0.129 (0.116-0.145) |
| Northern American deer mouse | n=1,118 med:0.126 (0.118-0.134) | n=385 med:0.097 (0.086-0.116) | n=749 med:0.066 (0.060-0.073) |
| Naked mole-rat female | n=1,052 med:0.147 (0.136-0.155) | n=358 med:0.122 (0.100-0.140) | n=723 med:0.084 (0.073-0.090) |
| Mouse | n=1,204 med:0.122 (0.115-0.128) | n=410 med:0.097 (0.088-0.108) | n=823 med:0.066 (0.060-0.072) |
| Daurian ground squirrel | n=963 med:0.144 (0.137-0.154) | n=335 med:0.117 (0.104-0.131) | n=610 med:0.089 (0.076-0.100) |
| Tiger | n=941 med:0.154 (0.142-0.163) | n=305 med:0.127 (0.107-0.147) | n=613 med:0.090 (0.081-0.098) |
| Lesser hedgehog tenrec | n=269 med:0.147 (0.123-0.162) | n=89 med:0.158 (0.110-0.190) | n=153 med:0.108 (0.086-0.132) |
| Mongolian gerbil | n=1,036 med:0.127 (0.119-0.134) | n=352 med:0.105 (0.094-0.121) | n=668 med:0.073 (0.066-0.082) |
| Ma's night monkey | n=1,124 med:0.193 (0.179-0.206) | n=382 med:0.164 (0.148-0.183) | n=784 med:0.114 (0.101-0.123) |

| | | | |
|---|---|---|---|
| Upper Galilee mountains blind mole rat | n=1,092<br>med:0.132<br>(0.124-0.139) | n=364<br>med:0.113<br>(0.098-0.124) | n=760<br>med:0.067<br>(0.062-0.073) |
| Kangaroo rat | n=1,003<br>med:0.121<br>(0.111-0.129) | n=341<br>med:0.105<br>(0.096-0.119) | n=689<br>med:0.073<br>(0.066-0.081) |
| Chimpanzee | n=1,122<br>med:0.226<br>(0.201-0.244) | n=380<br>med:0.184<br>(0.149-0.217) | n=776<br>med:0.136<br>(0.118-0.157) |
| Guinea Pig | n=1,097<br>med:0.129<br>(0.120-0.139) | n=381<br>med:0.105<br>(0.091-0.118) | n=765<br>med:0.077<br>(0.070-0.086) |
| American beaver | n=1,032<br>med:0.138<br>(0.131-0.148) | n=359<br>med:0.117<br>(0.105-0.133) | n=672<br>med:0.083<br>(0.077-0.092) |
| Wild yak | n=1,038<br>med:0.144<br>(0.133-0.152) | n=341<br>med:0.114<br>(0.103-0.130) | n=639<br>med:0.086<br>(0.079-0.093) |
| Coquerel's sifaka | n=1,114<br>med:0.154<br>(0.142-0.163) | n=373<br>med:0.121<br>(0.107-0.144) | n=766<br>med:0.090<br>(0.079-0.096) |
| Alpaca | n=220<br>med:0.228<br>(0.181-0.259) | n=65<br>med:0.167<br>(0.140-0.190) | n=135<br>med:0.099<br>(0.084-0.122) |
| Lesser Egyptian jerboa | n=1,022<br>med:0.129<br>(0.122-0.136) | n=334<br>med:0.097<br>(0.084-0.116) | n=716<br>med:0.071<br>(0.065-0.078) |
| Long-tailed chinchilla | n=1,110<br>med:0.131<br>(0.123-0.137) | n=373<br>med:0.098<br>(0.090-0.109) | n=763<br>med:0.070<br>(0.065-0.079) |
| Angola colobus | n=1,168<br>med:0.188<br>(0.177-0.198) | n=386<br>med:0.156<br>(0.135-0.175) | n=784<br>med:0.123<br>(0.107-0.136) |
| Ryukyu mouse | n=1,141<br>med:0.121<br>(0.114-0.128) | n=391<br>med:0.097<br>(0.089-0.112) | n=797<br>med:0.067<br>(0.061-0.074) |
| Chinese hamster PICR | n=1,132<br>med:0.133<br>(0.125-0.141) | n=393<br>med:0.112<br>(0.099-0.126) | n=799<br>med:0.073<br>(0.067-0.080) |
| Vervet-AGM | n=1,189<br>med:0.171<br>(0.158-0.185) | n=395<br>med:0.142<br>(0.125-0.166) | n=803<br>med:0.096<br>(0.087-0.106) |
| Cow | n=1,154<br>med:0.135<br>(0.128-0.145) | n=388<br>med:0.110<br>(0.100-0.128) | n=767<br>med:0.077<br>(0.072-0.083) |
| Bushbaby | n=1,150<br>med:0.151<br>(0.138-0.162) | n=391<br>med:0.120<br>(0.106-0.131) | n=787<br>med:0.081<br>(0.075-0.088) |

| | | | |
|---|---|---|---|
| Bonobo | n=1,146<br>med:0.239<br>(0.224-0.262) | n=381<br>med:0.247<br>(0.195-0.271) | n=770<br>med:0.200<br>(0.167-0.234) |
| Hyrax | n=322<br>med:0.158<br>(0.139-0.171) | n=116<br>med:0.124<br>(0.097-0.150) | n=199<br>med:0.100<br>(0.078-0.111) |
| Rabbit | n=1,008<br>med:0.141<br>(0.133-0.150) | n=342<br>med:0.116<br>(0.097-0.127) | n=689<br>med:0.086<br>(0.078-0.095) |
| Megabat | n=596<br>med:0.165<br>(0.154-0.179) | n=222<br>med:0.148<br>(0.131-0.166) | n=361<br>med:0.111<br>(0.096-0.127) |
| Dog | n=1,083<br>med:0.143<br>(0.137-0.152) | n=366<br>med:0.126<br>(0.109-0.137) | n=748<br>med:0.088<br>(0.081-0.096) |
| Steppe mouse | n=1,126<br>med:0.129<br>(0.122-0.136) | n=386<br>med:0.105<br>(0.095-0.117) | n=779<br>med:0.074<br>(0.068-0.081) |
| Greater bamboo lemur | n=1,162<br>med:0.155<br>(0.146-0.164) | n=385<br>med:0.117<br>(0.105-0.134) | n=786<br>med:0.089<br>(0.082-0.094) |
| Pig | n=1,145<br>med:0.145<br>(0.136-0.153) | n=393<br>med:0.107<br>(0.101-0.122) | n=775<br>med:0.080<br>(0.073-0.086) |
| Tree Shrew | n=205<br>med:0.170<br>(0.150-0.192) | n=64<br>med:0.164<br>(0.118-0.211) | n=125<br>med:0.129<br>(0.112-0.145) |
| Pika | n=409<br>med:0.134<br>(0.122-0.151) | n=148<br>med:0.096<br>(0.077-0.130) | n=210<br>med:0.085<br>(0.078-0.108) |
| Shrew | n=196<br>med:0.151<br>(0.118-0.174) | n=59<br>med:0.136<br>(0.095-0.160) | n=120<br>med:0.103<br>(0.081-0.130) |
| Squirrel | n=1,116<br>med:0.130<br>(0.124-0.137) | n=388<br>med:0.105<br>(0.095-0.116) | n=790<br>med:0.070<br>(0.063-0.076) |
| Gorilla | n=1,156<br>med:0.229<br>(0.205-0.248) | n=392<br>med:0.196<br>(0.168-0.227) | n=794<br>med:0.143<br>(0.125-0.158) |
| Cat | n=1,156<br>med:0.142<br>(0.130-0.152) | n=382<br>med:0.118<br>(0.105-0.129) | n=790<br>med:0.080<br>(0.074-0.087) |

### Human Reference Genome, Zeisel et al., 2015, cell subtypes

| Sample Size and Median dN/dS (CI95%) | Ependymal-specific Genes | Microglia-specific Genes | Endothelial-specific Genes | Mural-specific Genes | Oligodendrocyte-specific Genes | Astrocyte-specific Genes | Interneuron-specific Genes | S1 Pyramidal-specific Genes | CA1 Pyramidal-specific Genes |
|---|---|---|---|---|---|---|---|---|---|
| Microbat | n=301 med:0.199 (0.176-0.220) | n=259 med:0.174 (0.149-0.194) | n=252 med:0.128 (0.114-0.151) | n=107 med:0.115 (0.087-0.132) | n=292 med:0.113 (0.100-0.130) | n=164 med:0.121 (0.100-0.134) | n=239 med:0.113 (0.100-0.123) | n=182 med:0.096 (0.081-0.114) | n=248 med:0.089 (0.077-0.106) |
| Hedgehog | n=60 med:0.197 (0.120-0.222) | n=67 med:0.164 (0.134-0.210) | n=59 med:0.152 (0.089-0.197) | n=20 med:0.103 (0.055-0.148) | n=77 med:0.128 (0.092-0.149) | n=34 med:0.117 (0.071-0.185) | n=51 med:0.103 (0.078-0.134) | n=35 med:0.111 (0.075-0.145) | n=60 med:0.107 (0.071-0.169) |
| Naked mole-rat male | n=292 med:0.172 (0.158-0.193) | n=254 med:0.159 (0.142-0.182) | n=234 med:0.120 (0.100-0.147) | n=107 med:0.107 (0.087-0.131) | n=288 med:0.113 (0.103-0.128) | n=158 med:0.119 (0.102-0.147) | n=237 med:0.089 (0.076-0.099) | n=179 med:0.084 (0.073-0.101) | n=249 med:0.083 (0.067-0.091) |
| Sooty mangabey | n=345 med:0.250 (0.217-0.275) | n=307 med:0.201 (0.169-0.225) | n=282 med:0.153 (0.130-0.197) | n=119 med:0.120 (0.097-0.149) | n=345 med:0.139 (0.117-0.156) | n=190 med:0.151 (0.127-0.174) | n=288 med:0.116 (0.092-0.141) | n=218 med:0.103 (0.081-0.125) | n=308 med:0.098 (0.082-0.121) |
| Olive baboon | n=345 med:0.241 (0.218-0.265) | n=309 med:0.192 (0.169-0.226) | n=279 med:0.184 (0.142-0.208) | n=119 med:0.132 (0.112-0.166) | n=348 med:0.148 (0.127-0.169) | n=190 med:0.150 (0.125-0.166) | n=285 med:0.132 (0.112-0.156) | n=213 med:0.117 (0.081-0.138) | n=308 med:0.106 (0.088-0.126) |
| American bison | n=300 med:0.188 (0.163-0.207) | n=258 med:0.160 (0.139-0.182) | n=235 med:0.131 (0.101-0.154) | n=97 med:0.106 (0.084-0.116) | n=274 med:0.115 (0.102-0.127) | n=150 med:0.111 (0.094-0.128) | n=215 med:0.086 (0.076-0.100) | n=165 med:0.079 (0.063-0.094) | n=243 med:0.078 (0.061-0.088) |
| Leopard | n=336 med:0.170 (0.153-0.187) | n=284 med:0.148 (0.127-0.169) | n=255 med:0.107 (0.100-0.131) | n=118 med:0.095 (0.078-0.114) | n=324 med:0.092 (0.083-0.103) | n=181 med:0.111 (0.095-0.127) | n=279 med:0.076 (0.060-0.083) | n=203 med:0.073 (0.061-0.083) | n=307 med:0.070 (0.059-0.081) |
| Bolivian squirrel monkey | n=338 med:0.244 (0.218-0.266) | n=290 med:0.202 (0.179-0.231) | n=267 med:0.166 (0.139-0.196) | n=116 med:0.158 (0.115-0.190) | n=330 med:0.132 (0.113-0.147) | n=172 med:0.142 (0.124-0.163) | n=271 med:0.100 (0.087-0.115) | n=205 med:0.096 (0.085-0.129) | n=301 med:0.095 (0.085-0.106) |
| Crab-eating macaque | n=350 med:0.232 (0.208-0.259) | n=310 med:0.191 (0.166-0.219) | n=279 med:0.152 (0.127-0.181) | n=120 med:0.124 (0.100-0.155) | n=345 med:0.137 (0.117-0.157) | n=187 med:0.141 (0.112-0.169) | n=282 med:0.110 (0.096-0.127) | n=211 med:0.102 (0.080-0.126) | n=307 med:0.099 (0.086-0.119) |
| Red fox | n=320 med:0.184 (0.165-0.208) | n=281 med:0.162 (0.144-0.183) | n=257 med:0.124 (0.108-0.151) | n=115 med:0.118 (0.099-0.136) | n=320 med:0.115 (0.103-0.127) | n=172 med:0.129 (0.114-0.147) | n=258 med:0.089 (0.078-0.101) | n=194 med:0.098 (0.081-0.120) | n=283 med:0.087 (0.076-0.108) |
| Shrew mouse | n=338 med:0.145 (0.129-0.163) | n=295 med:0.138 (0.128-0.146) | n=274 med:0.104 (0.093-0.119) | n=119 med:0.102 (0.083-0.124) | n=337 med:0.103 (0.090-0.114) | n=188 med:0.103 (0.085-0.115) | n=274 med:0.072 (0.060-0.087) | n=208 med:0.072 (0.059-0.082) | n=312 med:0.061 (0.052-0.072) |
| Arctic ground squirrel | n=322 med:0.178 (0.152-0.191) | n=281 med:0.149 (0.133-0.161) | n=278 med:0.115 (0.098-0.137) | n=120 med:0.104 (0.080-0.113) | n=333 med:0.109 (0.095-0.120) | n=187 med:0.112 (0.096-0.126) | n=280 med:0.077 (0.066-0.093) | n=209 med:0.076 (0.062-0.089) | n=315 med:0.062 (0.051-0.075) |
| Chinese hamster CriGri | n=312 med:0.154 (0.139-0.172) | n=278 med:0.148 (0.134-0.162) | n=255 med:0.118 (0.093-0.130) | n=110 med:0.105 (0.085-0.124) | n=313 med:0.105 (0.097-0.124) | n=163 med:0.113 (0.095-0.132) | n=260 med:0.083 (0.070-0.099) | n=185 med:0.078 (0.065-0.098) | n=277 med:0.072 (0.061-0.085) |
| Opossum | n=287 med:0.135 (0.119-0.149) | n=247 med:0.112 (0.095-0.126) | n=235 med:0.093 (0.082-0.110) | n=109 med:0.075 (0.063-0.088) | n=297 med:0.085 (0.074-0.094) | n=163 med:0.093 (0.082-0.104) | n=246 med:0.072 (0.063-0.083) | n=185 med:0.070 (0.058-0.080) | n=262 med:0.066 (0.059-0.072) |
| Dolphin | n=180 med:0.215 (0.177-0.245) | n=152 med:0.174 (0.150-0.201) | n=131 med:0.154 (0.110-0.186) | n=64 med:0.129 (0.115-0.158) | n=171 med:0.129 (0.106-0.144) | n=98 med:0.128 (0.101-0.146) | n=137 med:0.108 (0.091-0.126) | n=106 med:0.081 (0.061-0.108) | n=154 med:0.095 (0.080-0.113) |
| Capuchin | n=339 med:0.235 (0.203-0.258) | n=295 med:0.202 (0.182-0.239) | n=272 med:0.165 (0.129-0.190) | n=120 med:0.150 (0.107-0.168) | n=332 med:0.121 (0.104-0.140) | n=181 med:0.154 (0.124-0.176) | n=280 med:0.102 (0.087-0.113) | n=207 med:0.089 (0.076-0.113) | n=302 med:0.089 (0.076-0.100) |
| Mouse Lemur | n=334 med:0.203 (0.177-0.229) | n=294 med:0.163 (0.149-0.184) | n=268 med:0.131 (0.112-0.163) | n=113 med:0.104 (0.088-0.126) | n=329 med:0.110 (0.087-0.128) | n=180 med:0.120 (0.105-0.136) | n=268 med:0.087 (0.074-0.107) | n=208 med:0.084 (0.074-0.098) | n=297 med:0.078 (0.070-0.085) |
| Damara mole rat | n=298 med:0.166 (0.149-0.184) | n=261 med:0.156 (0.137-0.176) | n=252 med:0.117 (0.102-0.139) | n=110 med:0.100 (0.083-0.129) | n=318 med:0.106 (0.096-0.116) | n=170 med:0.111 (0.096-0.127) | n=261 med:0.077 (0.066-0.100) | n=187 med:0.079 (0.066-0.104) | n=276 med:0.071 (0.058-0.084) |
| Marmoset | n=340 med:0.248 (0.221-0.271) | n=297 med:0.205 (0.180-0.232) | n=276 med:0.171 (0.145-0.200) | n=119 med:0.133 (0.116-0.155) | n=324 med:0.137 (0.114-0.160) | n=175 med:0.150 (0.134-0.173) | n=288 med:0.117 (0.101-0.136) | n=212 med:0.102 (0.083-0.121) | n=305 med:0.090 (0.076-0.101) |
| Goat | n=335 med:0.166 (0.146-0.181) | n=294 med:0.148 (0.135-0.174) | n=265 med:0.118 (0.103-0.138) | n=116 med:0.111 (0.092-0.128) | n=339 med:0.105 (0.094-0.118) | n=184 med:0.120 (0.107-0.134) | n=268 med:0.085 (0.074-0.092) | n=202 med:0.083 (0.069-0.097) | n=302 med:0.076 (0.066-0.084) |
| Common wombat | n=265 med:0.130 (0.118-0.140) | n=248 med:0.117 (0.101-0.127) | n=232 med:0.090 (0.078-0.104) | n=101 med:0.079 (0.062-0.109) | n=281 med:0.081 (0.073-0.092) | n=153 med:0.089 (0.078-0.100) | n=222 med:0.067 (0.057-0.079) | n=179 med:0.069 (0.053-0.080) | n=269 med:0.062 (0.052-0.071) |
| American black bear | n=293 med:0.186 (0.156-0.201) | n=265 med:0.165 (0.144-0.179) | n=240 med:0.143 (0.121-0.170) | n=107 med:0.110 (0.086-0.132) | n=305 med:0.114 (0.103-0.129) | n=169 med:0.132 (0.117-0.154) | n=231 med:0.106 (0.087-0.124) | n=164 med:0.095 (0.071-0.108) | n=255 med:0.085 (0.078-0.103) |
| Sloth | n=60 med:0.245 (0.180-0.284) | n=41 med:0.215 (0.154-0.337) | n=40 med:0.191 (0.095-0.278) | n=9 med:0.180 (0.105-0.357) | n=50 med:0.169 (0.112-0.262) | n=21 med:0.122 (0.069-0.280) | n=33 med:0.112 (0.085-0.152) | n=22 med:0.139 (0.066-0.223) | n=32 med:0.125 (0.081-0.173) |
| Polar bear | n=304 med:0.197 (0.180-0.213) | n=267 med:0.168 (0.149-0.186) | n=239 med:0.150 (0.121-0.169) | n=96 med:0.120 (0.095-0.148) | n=302 med:0.127 (0.114-0.139) | n=152 med:0.133 (0.112-0.150) | n=227 med:0.104 (0.089-0.120) | n=158 med:0.107 (0.090-0.129) | n=256 med:0.097 (0.081-0.111) |
| Alpine marmot | n=310 med:0.185 (0.160-0.200) | n=285 med:0.149 (0.134-0.164) | n=264 med:0.117 (0.098-0.141) | n=112 med:0.103 (0.084-0.120) | n=307 med:0.109 (0.095-0.125) | n=169 med:0.112 (0.086-0.124) | n=247 med:0.081 (0.073-0.099) | n=191 med:0.079 (0.066-0.099) | n=292 med:0.066 (0.057-0.082) |
| Prairie vole | n=321 med:0.141 (0.128-0.159) | n=282 med:0.142 (0.129-0.153) | n=260 med:0.098 (0.082-0.116) | n=114 med:0.088 (0.074-0.106) | n=329 med:0.099 (0.084-0.110) | n=175 med:0.097 (0.083-0.113) | n=272 med:0.072 (0.061-0.084) | n=203 med:0.068 (0.056-0.082) | n=304 med:0.061 (0.052-0.069) |
| Donkey | n=333 med:0.207 (0.182-0.226) | n=295 med:0.173 (0.161-0.193) | n=269 med:0.131 (0.110-0.152) | n=121 med:0.110 (0.102-0.125) | n=341 med:0.119 (0.110-0.130) | n=185 med:0.129 (0.106-0.148) | n=273 med:0.101 (0.084-0.114) | n=205 med:0.100 (0.089-0.118) | n=314 med:0.088 (0.077-0.101) |
| Golden Hamster | n=306 med:0.138 (0.124-0.157) | n=271 med:0.150 (0.131-0.167) | n=252 med:0.117 (0.098-0.144) | n=109 med:0.093 (0.072-0.120) | n=310 med:0.107 (0.097-0.118) | n=170 med:0.110 (0.092-0.130) | n=261 med:0.075 (0.066-0.087) | n=196 med:0.072 (0.060-0.093) | n=294 med:0.064 (0.055-0.080) |
| Armadillo | n=304 med:0.196 (0.180-0.206) | n=283 med:0.167 (0.152-0.189) | n=250 med:0.132 (0.110-0.153) | n=109 med:0.136 (0.101-0.157) | n=301 med:0.122 (0.107-0.129) | n=174 med:0.125 (0.112-0.145) | n=249 med:0.093 (0.081-0.103) | n=185 med:0.099 (0.085-0.115) | n=282 med:0.090 (0.077-0.101) |

| | | | | | | | | | |
|---|---|---|---|---|---|---|---|---|---|
| Ugandan red Colobus | n=333<br>med:0.233<br>(0.210-0.255) | n=296<br>med:0.193<br>(0.172-0.229) | n=266<br>med:0.165<br>(0.134-0.187) | n=110<br>med:0.125<br>(0.104-0.157) | n=338<br>med:0.147<br>(0.121-0.173) | n=183<br>med:0.136<br>(0.108-0.161) | n=284<br>med:0.119<br>(0.096-0.143) | n=210<br>med:0.091<br>(0.072-0.111) | n=302<br>med:0.089<br>(0.075-0.105) |
| Pig-tailed macaque | n=343<br>med:0.240<br>(0.210-0.272) | n=307<br>med:0.203<br>(0.174-0.229) | n=283<br>med:0.162<br>(0.136-0.200) | n=119<br>med:0.140<br>(0.109-0.162) | n=346<br>med:0.141<br>(0.115-0.162) | n=185<br>med:0.151<br>(0.121-0.176) | n=286<br>med:0.121<br>(0.098-0.142) | n=213<br>med:0.106<br>(0.090-0.126) | n=313<br>med:0.100<br>(0.088-0.119) |
| Panda | n=332<br>med:0.166<br>(0.149-0.193) | n=291<br>med:0.158<br>(0.140-0.177) | n=270<br>med:0.120<br>(0.106-0.137) | n=110<br>med:0.109<br>(0.091-0.122) | n=331<br>med:0.108<br>(0.092-0.116) | n=180<br>med:0.114<br>(0.093-0.131) | n=269<br>med:0.089<br>(0.071-0.097) | n=201<br>med:0.074<br>(0.067-0.094) | n=291<br>med:0.073<br>(0.064-0.083) |
| Ferret | n=334<br>med:0.175<br>(0.152-0.197) | n=295<br>med:0.149<br>(0.130-0.160) | n=270<br>med:0.110<br>(0.093-0.132) | n=111<br>med:0.106<br>(0.083-0.124) | n=321<br>med:0.096<br>(0.087-0.111) | n=181<br>med:0.106<br>(0.089-0.129) | n=269<br>med:0.082<br>(0.070-0.097) | n=201<br>med:0.082<br>(0.068-0.095) | n=296<br>med:0.069<br>(0.060-0.082) |
| Koala | n=294<br>med:0.136<br>(0.119-0.150) | n=259<br>med:0.123<br>(0.108-0.136) | n=244<br>med:0.089<br>(0.078-0.099) | n=110<br>med:0.084<br>(0.070-0.103) | n=312<br>med:0.087<br>(0.078-0.098) | n=166<br>med:0.097<br>(0.083-0.106) | n=259<br>med:0.075<br>(0.066-0.083) | n=194<br>med:0.069<br>(0.058-0.080) | n=281<br>med:0.066<br>(0.056-0.075) |
| Wallaby | n=78<br>med:0.117<br>(0.098-0.148) | n=71<br>med:0.102<br>(0.079-0.128) | n=73<br>med:0.088<br>(0.075-0.108) | n=38<br>med:0.097<br>(0.050-0.120) | n=92<br>med:0.089<br>(0.067-0.107) | n=43<br>med:0.086<br>(0.072-0.109) | n=75<br>med:0.067<br>(0.049-0.090) | n=60<br>med:0.068<br>(0.050-0.079) | n=69<br>med:0.061<br>(0.047-0.088) |
| Tarsier | n=283<br>med:0.214<br>(0.198-0.227) | n=246<br>med:0.164<br>(0.147-0.190) | n=220<br>med:0.132<br>(0.106-0.160) | n=86<br>med:0.114<br>(0.087-0.152) | n=283<br>med:0.118<br>(0.096-0.128) | n=149<br>med:0.124<br>(0.103-0.141) | n=227<br>med:0.105<br>(0.091-0.119) | n=170<br>med:0.093<br>(0.076-0.116) | n=233<br>med:0.097<br>(0.083-0.112) |
| Degu | n=286<br>med:0.160<br>(0.141-0.175) | n=241<br>med:0.143<br>(0.125-0.155) | n=235<br>med:0.121<br>(0.100-0.136) | n=108<br>med:0.098<br>(0.079-0.125) | n=298<br>med:0.106<br>(0.090-0.118) | n=158<br>med:0.103<br>(0.088-0.123) | n=236<br>med:0.073<br>(0.062-0.089) | n=183<br>med:0.085<br>(0.069-0.105) | n=265<br>med:0.067<br>(0.058-0.078) |
| Chinese hamster CHOK1GS | n=333<br>med:0.154<br>(0.136-0.175) | n=302<br>med:0.146<br>(0.134-0.159) | n=279<br>med:0.113<br>(0.093-0.130) | n=120<br>med:0.099<br>(0.077-0.121) | n=330<br>med:0.102<br>(0.092-0.116) | n=187<br>med:0.108<br>(0.089-0.126) | n=285<br>med:0.077<br>(0.064-0.090) | n=211<br>med:0.073<br>(0.060-0.089) | n=304<br>med:0.067<br>(0.056-0.074) |
| Drill | n=336<br>med:0.261<br>(0.229-0.281) | n=300<br>med:0.199<br>(0.182-0.223) | n=265<br>med:0.171<br>(0.136-0.197) | n=115<br>med:0.147<br>(0.126-0.166) | n=333<br>med:0.150<br>(0.129-0.181) | n=184<br>med:0.163<br>(0.138-0.186) | n=267<br>med:0.131<br>(0.108-0.151) | n=205<br>med:0.116<br>(0.087-0.131) | n=295<br>med:0.118<br>(0.100-0.133) |
| Gibbon | n=339<br>med:0.326<br>(0.293-0.359) | n=297<br>med:0.268<br>(0.241-0.291) | n=271<br>med:0.214<br>(0.186-0.229) | n=116<br>med:0.203<br>(0.157-0.236) | n=330<br>med:0.193<br>(0.173-0.228) | n=183<br>med:0.201<br>(0.154-0.236) | n=272<br>med:0.190<br>(0.148-0.217) | n=207<br>med:0.181<br>(0.159-0.220) | n=296<br>med:0.151<br>(0.129-0.172) |
| Rat | n=333<br>med:0.145<br>(0.134-0.169) | n=291<br>med:0.140<br>(0.127-0.153) | n=264<br>med:0.104<br>(0.092-0.126) | n=114<br>med:0.105<br>(0.086-0.116) | n=326<br>med:0.097<br>(0.087-0.110) | n=181<br>med:0.108<br>(0.090-0.123) | n=266<br>med:0.080<br>(0.065-0.096) | n=205<br>med:0.074<br>(0.061-0.088) | n=303<br>med:0.063<br>(0.054-0.076) |
| Macaque | n=335<br>med:0.242<br>(0.213-0.261) | n=307<br>med:0.217<br>(0.186-0.258) | n=277<br>med:0.161<br>(0.134-0.196) | n=120<br>med:0.137<br>(0.093-0.163) | n=333<br>med:0.147<br>(0.128-0.169) | n=188<br>med:0.164<br>(0.131-0.189) | n=282<br>med:0.116<br>(0.097-0.154) | n=210<br>med:0.117<br>(0.102-0.147) | n=312<br>med:0.097<br>(0.082-0.116) |
| American mink | n=323<br>med:0.165<br>(0.147-0.184) | n=274<br>med:0.143<br>(0.128-0.163) | n=254<br>med:0.110<br>(0.092-0.138) | n=116<br>med:0.096<br>(0.079-0.112) | n=322<br>med:0.104<br>(0.088-0.117) | n=173<br>med:0.110<br>(0.099-0.124) | n=264<br>med:0.087<br>(0.073-0.097) | n=196<br>med:0.072<br>(0.060-0.090) | n=284<br>med:0.072<br>(0.063-0.084) |
| Gelada | n=347<br>med:0.233<br>(0.211-0.254) | n=307<br>med:0.180<br>(0.155-0.206) | n=277<br>med:0.130<br>(0.102-0.161) | n=121<br>med:0.102<br>(0.085-0.133) | n=348<br>med:0.126<br>(0.107-0.141) | n=188<br>med:0.130<br>(0.107-0.159) | n=282<br>med:0.103<br>(0.091-0.128) | n=217<br>med:0.085<br>(0.063-0.107) | n=314<br>med:0.087<br>(0.072-0.103) |
| Sheep | n=328<br>med:0.183<br>(0.167-0.201) | n=287<br>med:0.167<br>(0.148-0.180) | n=253<br>med:0.135<br>(0.114-0.159) | n=110<br>med:0.122<br>(0.104-0.132) | n=323<br>med:0.116<br>(0.106-0.132) | n=177<br>med:0.131<br>(0.115-0.145) | n=254<br>med:0.096<br>(0.083-0.107) | n=191<br>med:0.091<br>(0.080-0.104) | n=291<br>med:0.090<br>(0.075-0.101) |
| Tasmanian devil | n=275<br>med:0.129<br>(0.116-0.145) | n=239<br>med:0.109<br>(0.090-0.121) | n=232<br>med:0.084<br>(0.074-0.097) | n=101<br>med:0.085<br>(0.063-0.095) | n=290<br>med:0.084<br>(0.071-0.092) | n=146<br>med:0.096<br>(0.087-0.106) | n=221<br>med:0.072<br>(0.061-0.084) | n=164<br>med:0.063<br>(0.050-0.074) | n=244<br>med:0.068<br>(0.055-0.078) |
| Elephant | n=328<br>med:0.191<br>(0.161-0.209) | n=292<br>med:0.165<br>(0.152-0.185) | n=260<br>med:0.128<br>(0.114-0.147) | n=116<br>med:0.114<br>(0.089-0.129) | n=335<br>med:0.109<br>(0.097-0.126) | n=176<br>med:0.120<br>(0.104-0.140) | n=272<br>med:0.099<br>(0.084-0.112) | n=193<br>med:0.087<br>(0.074-0.110) | n=286<br>med:0.092<br>(0.075-0.103) |
| Algerian mouse | n=340<br>med:0.154<br>(0.132-0.165) | n=294<br>med:0.141<br>(0.127-0.158) | n=275<br>med:0.103<br>(0.089-0.125) | n=119<br>med:0.092<br>(0.078-0.116) | n=336<br>med:0.103<br>(0.090-0.111) | n=192<br>med:0.103<br>(0.086-0.122) | n=280<br>med:0.071<br>(0.059-0.080) | n=213<br>med:0.070<br>(0.058-0.084) | n=314<br>med:0.061<br>(0.055-0.071) |
| Dingo | n=319<br>med:0.175<br>(0.153-0.197) | n=288<br>med:0.151<br>(0.131-0.166) | n=256<br>med:0.116<br>(0.104-0.129) | n=112<br>med:0.106<br>(0.091-0.134) | n=316<br>med:0.105<br>(0.093-0.118) | n=174<br>med:0.113<br>(0.100-0.126) | n=260<br>med:0.081<br>(0.067-0.096) | n=193<br>med:0.083<br>(0.071-0.099) | n=297<br>med:0.077<br>(0.065-0.089) |
| Golden snub-nosed monkey | n=333<br>med:0.247<br>(0.219-0.273) | n=301<br>med:0.205<br>(0.182-0.223) | n=269<br>med:0.169<br>(0.137-0.216) | n=112<br>med:0.158<br>(0.126-0.172) | n=329<br>med:0.150<br>(0.137-0.168) | n=176<br>med:0.146<br>(0.123-0.163) | n=269<br>med:0.119<br>(0.099-0.144) | n=202<br>med:0.113<br>(0.095-0.146) | n=301<br>med:0.113<br>(0.091-0.129) |
| Horse | n=339<br>med:0.199<br>(0.172-0.223) | n=295<br>med:0.177<br>(0.157-0.193) | n=274<br>med:0.128<br>(0.113-0.152) | n=120<br>med:0.113<br>(0.099-0.125) | n=329<br>med:0.123<br>(0.107-0.135) | n=184<br>med:0.123<br>(0.104-0.146) | n=273<br>med:0.095<br>(0.081-0.110) | n=206<br>med:0.088<br>(0.079-0.102) | n=308<br>med:0.086<br>(0.069-0.096) |
| Black snub-nosed monkey | n=335<br>med:0.264<br>(0.241-0.290) | n=290<br>med:0.230<br>(0.191-0.247) | n=261<br>med:0.197<br>(0.171-0.223) | n=114<br>med:0.158<br>(0.129-0.189) | n=346<br>med:0.154<br>(0.140-0.173) | n=178<br>med:0.161<br>(0.141-0.192) | n=270<br>med:0.138<br>(0.113-0.166) | n=203<br>med:0.151<br>(0.115-0.179) | n=300<br>med:0.136<br>(0.118-0.153) |
| Brazilian guinea pig | n=235<br>med:0.156<br>(0.144-0.168) | n=212<br>med:0.161<br>(0.142-0.177) | n=196<br>med:0.130<br>(0.116-0.147) | n=96<br>med:0.115<br>(0.097-0.136) | n=252<br>med:0.126<br>(0.115-0.139) | n=130<br>med:0.122<br>(0.109-0.142) | n=192<br>med:0.105<br>(0.087-0.115) | n=146<br>med:0.106<br>(0.092-0.126) | n=217<br>med:0.108<br>(0.091-0.115) |
| Orangutan | n=323<br>med:0.280<br>(0.238-0.321) | n=300<br>med:0.200<br>(0.172-0.227) | n=272<br>med:0.172<br>(0.148-0.198) | n=118<br>med:0.135<br>(0.108-0.180) | n=338<br>med:0.171<br>(0.146-0.194) | n=183<br>med:0.175<br>(0.139-0.200) | n=266<br>med:0.145<br>(0.116-0.169) | n=204<br>med:0.137<br>(0.104-0.168) | n=301<br>med:0.119<br>(0.097-0.133) |
| Northern American deer mouse | n=326<br>med:0.150<br>(0.137-0.165) | n=288<br>med:0.143<br>(0.129-0.160) | n=269<br>med:0.103<br>(0.082-0.125) | n=116<br>med:0.096<br>(0.079-0.115) | n=322<br>med:0.099<br>(0.090-0.116) | n=182<br>med:0.104<br>(0.087-0.123) | n=258<br>med:0.070<br>(0.059-0.086) | n=202<br>med:0.064<br>(0.054-0.080) | n=289<br>med:0.064<br>(0.055-0.074) |
| Naked mole-rat female | n=309<br>med:0.177<br>(0.162-0.197) | n=265<br>med:0.158<br>(0.143-0.179) | n=247<br>med:0.128<br>(0.100-0.147) | n=111<br>med:0.111<br>(0.091-0.139) | n=307<br>med:0.114<br>(0.100-0.129) | n=171<br>med:0.120<br>(0.105-0.140) | n=256<br>med:0.088<br>(0.073-0.096) | n=195<br>med:0.084<br>(0.069-0.095) | n=272<br>med:0.076<br>(0.066-0.091) |
| Mouse | n=352<br>med:0.153<br>(0.134-0.168) | n=309<br>med:0.136<br>(0.121-0.151) | n=286<br>med:0.102<br>(0.090-0.125) | n=124<br>med:0.089<br>(0.077-0.107) | n=351<br>med:0.099<br>(0.089-0.109) | n=192<br>med:0.100<br>(0.086-0.117) | n=289<br>med:0.072<br>(0.061-0.083) | n=214<br>med:0.067<br>(0.056-0.081) | n=320<br>med:0.062<br>(0.053-0.071) |
| Daurian ground squirrel | n=288<br>med:0.195<br>(0.168-0.211) | n=246<br>med:0.158<br>(0.141-0.184) | n=235<br>med:0.123<br>(0.099-0.140) | n=100<br>med:0.111<br>(0.094-0.127) | n=282<br>med:0.118<br>(0.102-0.129) | n=147<br>med:0.125<br>(0.093-0.147) | n=217<br>med:0.099<br>(0.084-0.118) | n=165<br>med:0.078<br>(0.070-0.107) | n=228<br>med:0.078<br>(0.062-0.107) |
| Tiger | n=291<br>med:0.193<br>(0.168-0.211) | n=249<br>med:0.177<br>(0.155-0.194) | n=208<br>med:0.133<br>(0.104-0.153) | n=97<br>med:0.118<br>(0.104-0.148) | n=257<br>med:0.120<br>(0.100-0.132) | n=144<br>med:0.127<br>(0.109-0.154) | n=218<br>med:0.092<br>(0.080-0.104) | n=158<br>med:0.094<br>(0.078-0.110) | n=237<br>med:0.083<br>(0.066-0.096) |
| Lesser hedgehog tenrec | n=81<br>med:0.186<br>(0.148-0.217) | n=66<br>med:0.159<br>(0.121-0.187) | n=65<br>med:0.176<br>(0.115-0.236) | n=24<br>med:0.108<br>(0.051-0.162) | n=81<br>med:0.096<br>(0.073-0.132) | n=41<br>med:0.118<br>(0.077-0.174) | n=62<br>med:0.114<br>(0.089-0.150) | n=31<br>med:0.120<br>(0.062-0.152) | n=60<br>med:0.088<br>(0.066-0.132) |
| Mongolian gerbil | n=294<br>med:0.150<br>(0.134-0.176) | n=275<br>med:0.146<br>(0.133-0.162) | n=248<br>med:0.107<br>(0.090-0.124) | n=104<br>med:0.101<br>(0.084-0.126) | n=300<br>med:0.103<br>(0.090-0.114) | n=167<br>med:0.104<br>(0.088-0.123) | n=233<br>med:0.076<br>(0.062-0.091) | n=174<br>med:0.083<br>(0.060-0.101) | n=261<br>med:0.070<br>(0.059-0.077) |

| | | | | | | | | | |
|---|---|---|---|---|---|---|---|---|---|
| Ma's night monkey | n=327<br>med:0.246<br>(0.222-0.279) | n=294<br>med:0.222<br>(0.193-0.251) | n=269<br>med:0.176<br>(0.150-0.208) | n=113<br>med:0.143<br>(0.127-0.177) | n=323<br>med:0.139<br>(0.121-0.157) | n=180<br>med:0.150<br>(0.129-0.183) | n=275<br>med:0.117<br>(0.097-0.138) | n=209<br>med:0.111<br>(0.091-0.136) | n=300<br>med:0.112<br>(0.097-0.124) |
| Upper Galilee mountains blind mole rat | n=319<br>med:0.158<br>(0.140-0.170) | n=276<br>med:0.149<br>(0.138-0.162) | n=253<br>med:0.113<br>(0.089-0.128) | n=111<br>med:0.112<br>(0.089-0.135) | n=323<br>med:0.102<br>(0.088-0.114) | n=174<br>med:0.103<br>(0.082-0.125) | n=273<br>med:0.073<br>(0.061-0.085) | n=199<br>med:0.067<br>(0.057-0.075) | n=288<br>med:0.064<br>(0.053-0.072) |
| Kangaroo rat | n=289<br>med:0.160<br>(0.140-0.182) | n=253<br>med:0.136<br>(0.118-0.149) | n=238<br>med:0.104<br>(0.093-0.123) | n=103<br>med:0.106<br>(0.091-0.122) | n=294<br>med:0.096<br>(0.082-0.106) | n=167<br>med:0.102<br>(0.090-0.123) | n=242<br>med:0.078<br>(0.062-0.091) | n=185<br>med:0.071<br>(0.059-0.087) | n=262<br>med:0.068<br>(0.060-0.079) |
| Chimpanzee | n=324<br>med:0.290<br>(0.261-0.331) | n=288<br>med:0.237<br>(0.198-0.279) | n=264<br>med:0.211<br>(0.152-0.247) | n=116<br>med:0.156<br>(0.099-0.190) | n=329<br>med:0.152<br>(0.126-0.196) | n=181<br>med:0.182<br>(0.138-0.226) | n=269<br>med:0.152<br>(0.119-0.185) | n=205<br>med:0.117<br>(0.088-0.142) | n=302<br>med:0.137<br>(0.101-0.170) |
| Guinea Pig | n=317<br>med:0.157<br>(0.145-0.170) | n=281<br>med:0.152<br>(0.133-0.170) | n=266<br>med:0.112<br>(0.091-0.125) | n=115<br>med:0.094<br>(0.077-0.116) | n=329<br>med:0.100<br>(0.087-0.114) | n=170<br>med:0.112<br>(0.094-0.129) | n=270<br>med:0.081<br>(0.069-0.092) | n=202<br>med:0.082<br>(0.069-0.100) | n=293<br>med:0.070<br>(0.065-0.084) |
| American beaver | n=291<br>med:0.181<br>(0.156-0.192) | n=269<br>med:0.157<br>(0.137-0.171) | n=256<br>med:0.128<br>(0.106-0.153) | n=103<br>med:0.103<br>(0.092-0.122) | n=309<br>med:0.120<br>(0.105-0.129) | n=163<br>med:0.110<br>(0.097-0.123) | n=230<br>med:0.084<br>(0.068-0.100) | n=176<br>med:0.091<br>(0.068-0.112) | n=266<br>med:0.078<br>(0.066-0.088) |
| Wild yak | n=308<br>med:0.179<br>(0.159-0.196) | n=273<br>med:0.154<br>(0.140-0.180) | n=238<br>med:0.120<br>(0.104-0.141) | n=103<br>med:0.107<br>(0.091-0.126) | n=294<br>med:0.117<br>(0.104-0.133) | n=163<br>med:0.114<br>(0.097-0.131) | n=224<br>med:0.096<br>(0.083-0.109) | n=177<br>med:0.082<br>(0.069-0.093) | n=238<br>med:0.081<br>(0.065-0.098) |
| Coquerel's sifaka | n=331<br>med:0.205<br>(0.183-0.234) | n=279<br>med:0.169<br>(0.150-0.190) | n=262<br>med:0.133<br>(0.112-0.154) | n=111<br>med:0.107<br>(0.092-0.123) | n=326<br>med:0.117<br>(0.102-0.126) | n=178<br>med:0.127<br>(0.107-0.151) | n=266<br>med:0.091<br>(0.076-0.102) | n=204<br>med:0.094<br>(0.076-0.108) | n=296<br>med:0.087<br>(0.070-0.097) |
| Alpaca | n=71<br>med:0.267<br>(0.199-0.343) | n=55<br>med:0.271<br>(0.177-0.312) | n=49<br>med:0.182<br>(0.142-0.233) | n=16<br>med:0.143<br>(0.100-0.169) | n=57<br>med:0.202<br>(0.117-0.243) | n=37<br>med:0.149<br>(0.120-0.227) | n=50<br>med:0.105<br>(0.076-0.133) | n=26<br>med:0.107<br>(0.049-0.207) | n=59<br>med:0.094<br>(0.062-0.133) |
| Lesser Egyptian jerboa | n=307<br>med:0.155<br>(0.140-0.171) | n=252<br>med:0.142<br>(0.128-0.152) | n=225<br>med:0.098<br>(0.081-0.123) | n=109<br>med:0.097<br>(0.076-0.127) | n=300<br>med:0.107<br>(0.093-0.118) | n=163<br>med:0.125<br>(0.106-0.138) | n=252<br>med:0.079<br>(0.065-0.088) | n=193<br>med:0.067<br>(0.057-0.078) | n=271<br>med:0.068<br>(0.057-0.080) |
| Long-tailed chinchilla | n=331<br>med:0.156<br>(0.138-0.170) | n=281<br>med:0.152<br>(0.138-0.163) | n=258<br>med:0.101<br>(0.090-0.122) | n=115<br>med:0.093<br>(0.071-0.111) | n=328<br>med:0.103<br>(0.088-0.114) | n=170<br>med:0.106<br>(0.093-0.123) | n=271<br>med:0.072<br>(0.063-0.088) | n=200<br>med:0.073<br>(0.063-0.087) | n=292<br>med:0.065<br>(0.057-0.081) |
| Angola colobus | n=336<br>med:0.252<br>(0.218-0.278) | n=310<br>med:0.214<br>(0.189-0.239) | n=273<br>med:0.164<br>(0.137-0.211) | n=113<br>med:0.137<br>(0.109-0.163) | n=339<br>med:0.145<br>(0.129-0.159) | n=183<br>med:0.154<br>(0.117-0.178) | n=275<br>med:0.129<br>(0.112-0.146) | n=208<br>med:0.128<br>(0.092-0.158) | n=301<br>med:0.112<br>(0.094-0.132) |
| Ryukyu mouse | n=337<br>med:0.145<br>(0.129-0.162) | n=293<br>med:0.139<br>(0.124-0.151) | n=274<br>med:0.104<br>(0.087-0.126) | n=117<br>med:0.094<br>(0.076-0.108) | n=328<br>med:0.101<br>(0.089-0.110) | n=183<br>med:0.099<br>(0.085-0.112) | n=280<br>med:0.069<br>(0.059-0.085) | n=208<br>med:0.071<br>(0.057-0.083) | n=309<br>med:0.063<br>(0.054-0.072) |
| Chinese hamster PICR | n=333<br>med:0.162<br>(0.144-0.183) | n=287<br>med:0.146<br>(0.135-0.164) | n=275<br>med:0.118<br>(0.100-0.135) | n=118<br>med:0.101<br>(0.075-0.121) | n=325<br>med:0.105<br>(0.097-0.121) | n=187<br>med:0.109<br>(0.092-0.128) | n=282<br>med:0.080<br>(0.070-0.093) | n=205<br>med:0.072<br>(0.060-0.085) | n=312<br>med:0.069<br>(0.057-0.078) |
| Vervet-AGM | n=351<br>med:0.226<br>(0.206-0.257) | n=303<br>med:0.187<br>(0.161-0.208) | n=278<br>med:0.156<br>(0.126-0.184) | n=117<br>med:0.135<br>(0.096-0.155) | n=349<br>med:0.128<br>(0.112-0.145) | n=186<br>med:0.129<br>(0.113-0.161) | n=284<br>med:0.109<br>(0.092-0.130) | n=211<br>med:0.092<br>(0.075-0.111) | n=308<br>med:0.084<br>(0.066-0.101) |
| Cow | n=344<br>med:0.171<br>(0.152-0.194) | n=295<br>med:0.158<br>(0.139-0.176) | n=270<br>med:0.116<br>(0.100-0.133) | n=118<br>med:0.106<br>(0.086-0.133) | n=333<br>med:0.110<br>(0.095-0.117) | n=182<br>med:0.115<br>(0.095-0.128) | n=261<br>med:0.081<br>(0.072-0.096) | n=203<br>med:0.079<br>(0.070-0.091) | n=303<br>med:0.073<br>(0.060-0.081) |
| Bushbaby | n=335<br>med:0.195<br>(0.174-0.215) | n=292<br>med:0.170<br>(0.150-0.188) | n=273<br>med:0.124<br>(0.107-0.145) | n=118<br>med:0.106<br>(0.093-0.131) | n=337<br>med:0.117<br>(0.102-0.127) | n=186<br>med:0.120<br>(0.105-0.143) | n=277<br>med:0.090<br>(0.080-0.099) | n=212<br>med:0.075<br>(0.066-0.092) | n=298<br>med:0.075<br>(0.064-0.085) |
| Bonobo | n=323<br>med:0.286<br>(0.260-0.333) | n=299<br>med:0.263<br>(0.221-0.316) | n=265<br>med:0.252<br>(0.206-0.279) | n=116<br>med:0.216<br>(0.159-0.269) | n=340<br>med:0.197<br>(0.159-0.228) | n=184<br>med:0.205<br>(0.175-0.254) | n=269<br>med:0.205<br>(0.165-0.246) | n=206<br>med:0.191<br>(0.123-0.254) | n=295<br>med:0.199<br>(0.156-0.241) |
| Hyrax | n=84<br>med:0.179<br>(0.154-0.236) | n=94<br>med:0.179<br>(0.163-0.204) | n=83<br>med:0.131<br>(0.096-0.182) | n=33<br>med:0.117<br>(0.083-0.150) | n=97<br>med:0.123<br>(0.093-0.142) | n=47<br>med:0.126<br>(0.094-0.144) | n=71<br>med:0.079<br>(0.059-0.113) | n=55<br>med:0.118<br>(0.083-0.156) | n=73<br>med:0.092<br>(0.063-0.119) |
| Rabbit | n=298<br>med:0.168<br>(0.152-0.195) | n=254<br>med:0.149<br>(0.134-0.168) | n=233<br>med:0.125<br>(0.103-0.137) | n=109<br>med:0.098<br>(0.082-0.119) | n=294<br>med:0.117<br>(0.099-0.135) | n=162<br>med:0.118<br>(0.095-0.139) | n=242<br>med:0.089<br>(0.080-0.103) | n=179<br>med:0.092<br>(0.071-0.107) | n=268<br>med:0.077<br>(0.067-0.095) |
| Megabat | n=178<br>med:0.206<br>(0.174-0.245) | n=158<br>med:0.182<br>(0.159-0.220) | n=151<br>med:0.156<br>(0.131-0.179) | n=71<br>med:0.135<br>(0.115-0.164) | n=162<br>med:0.117<br>(0.097-0.139) | n=98<br>med:0.159<br>(0.137-0.190) | n=126<br>med:0.117<br>(0.091-0.134) | n=89<br>med:0.135<br>(0.101-0.157) | n=146<br>med:0.093<br>(0.079-0.129) |
| Dog | n=311<br>med:0.190<br>(0.170-0.206) | n=284<br>med:0.157<br>(0.138-0.175) | n=256<br>med:0.132<br>(0.112-0.154) | n=110<br>med:0.119<br>(0.093-0.138) | n=316<br>med:0.111<br>(0.100-0.128) | n=172<br>med:0.122<br>(0.109-0.137) | n=267<br>med:0.091<br>(0.080-0.099) | n=197<br>med:0.098<br>(0.081-0.119) | n=284<br>med:0.080<br>(0.069-0.091) |
| Steppe mouse | n=331<br>med:0.154<br>(0.138-0.169) | n=295<br>med:0.141<br>(0.128-0.154) | n=270<br>med:0.109<br>(0.096-0.131) | n=116<br>med:0.094<br>(0.079-0.118) | n=321<br>med:0.110<br>(0.101-0.120) | n=179<br>med:0.110<br>(0.095-0.127) | n=269<br>med:0.080<br>(0.069-0.097) | n=207<br>med:0.072<br>(0.065-0.084) | n=303<br>med:0.070<br>(0.058-0.083) |
| Greater bamboo lemur | n=337<br>med:0.197<br>(0.179-0.218) | n=301<br>med:0.167<br>(0.151-0.187) | n=267<br>med:0.126<br>(0.107-0.149) | n=118<br>med:0.107<br>(0.091-0.126) | n=341<br>med:0.115<br>(0.094-0.133) | n=183<br>med:0.136<br>(0.118-0.152) | n=273<br>med:0.099<br>(0.083-0.113) | n=209<br>med:0.086<br>(0.076-0.098) | n=304<br>med:0.085<br>(0.073-0.091) |
| Pig | n=329<br>med:0.171<br>(0.157-0.187) | n=299<br>med:0.160<br>(0.143-0.175) | n=275<br>med:0.116<br>(0.102-0.131) | n=118<br>med:0.101<br>(0.088-0.118) | n=331<br>med:0.111<br>(0.097-0.120) | n=186<br>med:0.122<br>(0.106-0.153) | n=274<br>med:0.083<br>(0.072-0.092) | n=203<br>med:0.084<br>(0.070-0.097) | n=298<br>med:0.076<br>(0.062-0.084) |
| Tree Shrew | n=67<br>med:0.188<br>(0.159-0.240) | n=51<br>med:0.178<br>(0.130-0.232) | n=45<br>med:0.173<br>(0.142-0.247) | n=19<br>med:0.118<br>(0.083-0.173) | n=56<br>med:0.149<br>(0.120-0.175) | n=31<br>med:0.142<br>(0.059-0.200) | n=49<br>med:0.123<br>(0.087-0.143) | n=28<br>med:0.092<br>(0.058-0.146) | n=48<br>med:0.148<br>(0.115-0.189) |
| Pika | n=121<br>med:0.167<br>(0.130-0.197) | n=111<br>med:0.154<br>(0.127-0.174) | n=108<br>med:0.127<br>(0.089-0.148) | n=40<br>med:0.070<br>(0.060-0.095) | n=117<br>med:0.101<br>(0.081-0.128) | n=60<br>med:0.114<br>(0.087-0.155) | n=73<br>med:0.085<br>(0.065-0.109) | n=52<br>med:0.113<br>(0.078-0.152) | n=85<br>med:0.079<br>(0.065-0.109) |
| Shrew | n=55<br>med:0.210<br>(0.136-0.248) | n=59<br>med:0.137<br>(0.096-0.175) | n=45<br>med:0.131<br>(0.079-0.179) | n=14<br>med:0.136<br>(0.036-0.182) | n=62<br>med:0.121<br>(0.073-0.184) | n=20<br>med:0.171<br>(0.079-0.228) | n=44<br>med:0.089<br>(0.072-0.103) | n=28<br>med:0.116<br>(0.061-0.132) | n=48<br>med:0.143<br>(0.068-0.165) |
| Squirrel | n=325<br>med:0.179<br>(0.152-0.200) | n=277<br>med:0.139<br>(0.127-0.160) | n=272<br>med:0.109<br>(0.092-0.126) | n=116<br>med:0.101<br>(0.084-0.110) | n=329<br>med:0.110<br>(0.096-0.120) | n=185<br>med:0.110<br>(0.092-0.127) | n=276<br>med:0.073<br>(0.062-0.082) | n=206<br>med:0.074<br>(0.062-0.085) | n=308<br>med:0.064<br>(0.055-0.076) |
| Gorilla | n=338<br>med:0.301<br>(0.257-0.337) | n=299<br>med:0.222<br>(0.190-0.264) | n=272<br>med:0.184<br>(0.145-0.239) | n=120<br>med:0.212<br>(0.172-0.274) | n=337<br>med:0.182<br>(0.156-0.233) | n=182<br>med:0.165<br>(0.123-0.202) | n=277<br>med:0.162<br>(0.126-0.207) | n=209<br>med:0.110<br>(0.078-0.154) | n=308<br>med:0.144<br>(0.115-0.171) |
| Cat | n=340<br>med:0.184<br>(0.164-0.205) | n=299<br>med:0.160<br>(0.134-0.175) | n=267<br>med:0.127<br>(0.108-0.139) | n=115<br>med:0.104<br>(0.088-0.117) | n=332<br>med:0.104<br>(0.093-0.120) | n=185<br>med:0.118<br>(0.103-0.138) | n=275<br>med:0.079<br>(0.068-0.092) | n=205<br>med:0.087<br>(0.075-0.108) | n=310<br>med:0.075<br>(0.065-0.086) |

#### Human Reference Genome, Saunders et al., 2018, three cell types

| Sample Size and Median dN/dS (CI95%) | Vasculature-specific Genes | Glia-specific Genes | Neuron-specific Genes |
|---|---|---|---|
| Microbat | n=318 med:0.129 (0.119-0.144) | n=219 med:0.110 (0.091-0.121) | n=502 med:0.072 (0.065-0.078) |
| Hedgehog | n=76 med:0.167 (0.127-0.202) | n=63 med:0.088 (0.076-0.135) | n=110 med:0.074 (0.056-0.092) |
| Naked mole-rat male | n=323 med:0.122 (0.108-0.138) | n=227 med:0.097 (0.085-0.110) | n=543 med:0.056 (0.051-0.062) |
| Sooty mangabey | n=374 med:0.157 (0.130-0.184) | n=277 med:0.119 (0.103-0.141) | n=617 med:0.076 (0.066-0.090) |
| Olive baboon | n=377 med:0.168 (0.140-0.193) | n=271 med:0.125 (0.109-0.148) | n=620 med:0.077 (0.066-0.085) |
| American bison | n=315 med:0.130 (0.111-0.146) | n=214 med:0.103 (0.090-0.117) | n=493 med:0.052 (0.045-0.060) |
| Leopard | n=369 med:0.117 (0.106-0.130) | n=260 med:0.078 (0.069-0.089) | n=599 med:0.050 (0.046-0.055) |
| Bolivian squirrel monkey | n=364 med:0.174 (0.148-0.210) | n=259 med:0.123 (0.108-0.134) | n=593 med:0.070 (0.061-0.079) |
| Crab-eating macaque | n=378 med:0.151 (0.125-0.171) | n=271 med:0.122 (0.095-0.134) | n=615 med:0.073 (0.062-0.082) |
| Red fox | n=351 med:0.130 (0.115-0.143) | n=241 med:0.103 (0.088-0.114) | n=580 med:0.070 (0.062-0.078) |
| Shrew mouse | n=369 med:0.105 (0.090-0.117) | n=269 med:0.077 (0.065-0.091) | n=616 med:0.048 (0.043-0.051) |
| Arctic ground squirrel | n=363 med:0.122 (0.107-0.137) | n=267 med:0.082 (0.069-0.095) | n=607 med:0.046 (0.041-0.051) |
| Chinese hamster CriGri | n=342 med:0.112 (0.097-0.123) | n=242 med:0.087 (0.075-0.101) | n=547 med:0.054 (0.048-0.061) |
| Opossum | n=325 med:0.092 (0.083-0.100) | n=232 med:0.070 (0.060-0.083) | n=526 med:0.056 (0.050-0.062) |
| Dolphin | n=200 | n=125 | n=289 |

| | | | |
|---|---|---|---|
| | med:0.155<br>(0.132-0.171) | med:0.119<br>(0.098-0.134) | med:0.066<br>(0.059-0.074) |
| Capuchin | n=358<br>med:0.170<br>(0.151-0.197) | n=269<br>med:0.109<br>(0.088-0.127) | n=603<br>med:0.062<br>(0.055-0.073) |
| Mouse Lemur | n=362<br>med:0.135<br>(0.117-0.156) | n=268<br>med:0.089<br>(0.079-0.096) | n=599<br>med:0.058<br>(0.051-0.065) |
| Damara mole rat | n=338<br>med:0.119<br>(0.105-0.133) | n=246<br>med:0.085<br>(0.075-0.097) | n=580<br>med:0.052<br>(0.046-0.056) |
| Marmoset | n=363<br>med:0.170<br>(0.152-0.194) | n=252<br>med:0.115<br>(0.097-0.134) | n=602<br>med:0.070<br>(0.059-0.080) |
| Goat | n=364<br>med:0.124<br>(0.105-0.137) | n=259<br>med:0.086<br>(0.075-0.101) | n=600<br>med:0.054<br>(0.048-0.060) |
| Common wombat | n=314<br>med:0.090<br>(0.079-0.097) | n=210<br>med:0.075<br>(0.062-0.082) | n=508<br>med:0.050<br>(0.046-0.056) |
| American black bear | n=335<br>med:0.136<br>(0.111-0.152) | n=239<br>med:0.093<br>(0.078-0.109) | n=496<br>med:0.076<br>(0.066-0.087) |
| Sloth | n=48<br>med:0.211<br>(0.174-0.335) | n=35<br>med:0.126<br>(0.081-0.262) | n=65<br>med:0.105<br>(0.073-0.137) |
| Polar bear | n=325<br>med:0.144<br>(0.129-0.161) | n=233<br>med:0.114<br>(0.097-0.125) | n=526<br>med:0.088<br>(0.079-0.095) |
| Alpine marmot | n=351<br>med:0.124<br>(0.107-0.138) | n=242<br>med:0.085<br>(0.072-0.098) | n=558<br>med:0.051<br>(0.047-0.057) |
| Prairie vole | n=348<br>med:0.096<br>(0.083-0.106) | n=264<br>med:0.079<br>(0.069-0.091) | n=597<br>med:0.044<br>(0.040-0.048) |
| Donkey | n=368<br>med:0.141<br>(0.124-0.157) | n=273<br>med:0.100<br>(0.084-0.111) | n=597<br>med:0.069<br>(0.062-0.078) |
| Golden Hamster | n=339<br>med:0.104<br>(0.093-0.120) | n=243<br>med:0.092<br>(0.083-0.105) | n=563<br>med:0.053<br>(0.046-0.057) |
| Armadillo | n=342<br>med:0.144<br>(0.121-0.160) | n=240<br>med:0.105<br>(0.088-0.120) | n=541<br>med:0.076<br>(0.069-0.083) |
| Ugandan red Colobus | n=372<br>med:0.175<br>(0.145-0.201) | n=266<br>med:0.109<br>(0.089-0.126) | n=601<br>med:0.065<br>(0.052-0.071) |
| Pig-tailed macaque | n=374 | n=275 | n=619 |

| | | | |
|---|---|---|---|
| | med:0.156 (0.125-0.177) | med:0.118 (0.101-0.137) | med:0.072 (0.063-0.085) |
| Panda | n=369 med:0.124 (0.109-0.139) | n=259 med:0.088 (0.079-0.102) | n=578 med:0.054 (0.048-0.059) |
| Ferret | n=361 med:0.115 (0.104-0.124) | n=254 med:0.085 (0.077-0.097) | n=589 med:0.053 (0.047-0.060) |
| Koala | n=337 med:0.092 (0.083-0.103) | n=243 med:0.078 (0.068-0.085) | n=555 med:0.053 (0.049-0.060) |
| Wallaby | n=97 med:0.072 (0.052-0.087) | n=90 med:0.081 (0.059-0.088) | n=135 med:0.051 (0.040-0.064) |
| Tarsier | n=309 med:0.143 (0.121-0.160) | n=228 med:0.107 (0.088-0.122) | n=474 med:0.066 (0.057-0.078) |
| Degu | n=305 med:0.112 (0.100-0.125) | n=229 med:0.082 (0.067-0.094) | n=562 med:0.047 (0.043-0.052) |
| Chinese hamster CHOK1GS | n=372 med:0.112 (0.099-0.127) | n=265 med:0.081 (0.066-0.096) | n=612 med:0.048 (0.044-0.055) |
| Drill | n=363 med:0.171 (0.148-0.190) | n=257 med:0.138 (0.117-0.167) | n=584 med:0.089 (0.074-0.099) |
| Gibbon | n=360 med:0.202 (0.183-0.222) | n=258 med:0.170 (0.150-0.196) | n=597 med:0.120 (0.106-0.138) |
| Rat | n=349 med:0.103 (0.091-0.112) | n=258 med:0.079 (0.064-0.087) | n=593 med:0.047 (0.043-0.051) |
| Macaque | n=370 med:0.169 (0.145-0.191) | n=267 med:0.128 (0.110-0.147) | n=614 med:0.069 (0.059-0.082) |
| American mink | n=365 med:0.112 (0.101-0.124) | n=259 med:0.084 (0.073-0.093) | n=572 med:0.050 (0.044-0.058) |
| Gelada | n=373 med:0.129 (0.111-0.154) | n=275 med:0.108 (0.093-0.123) | n=624 med:0.052 (0.046-0.064) |
| Sheep | n=351 med:0.133 (0.117-0.144) | n=244 med:0.100 (0.090-0.109) | n=575 med:0.067 (0.060-0.078) |
| Tasmanian devil | n=305 med:0.088 (0.077-0.097) | n=224 med:0.068 (0.057-0.083) | n=502 med:0.050 (0.045-0.056) |
| Elephant | n=354 | n=259 | n=581 |

| | | | |
|---|---|---|---|
| | med:0.128 (0.119-0.150) | med:0.099 (0.088-0.111) | med:0.069 (0.061-0.078) |
| Algerian mouse | n=374 med:0.104 (0.091-0.117) | n=267 med:0.077 (0.069-0.088) | n=616 med:0.046 (0.042-0.050) |
| Dingo | n=352 med:0.122 (0.107-0.137) | n=253 med:0.082 (0.072-0.094) | n=581 med:0.063 (0.056-0.069) |
| Golden snub-nosed monkey | n=365 med:0.169 (0.144-0.188) | n=253 med:0.132 (0.114-0.145) | n=581 med:0.078 (0.066-0.089) |
| Horse | n=368 med:0.141 (0.122-0.155) | n=259 med:0.101 (0.090-0.118) | n=611 med:0.063 (0.055-0.072) |
| Black snub-nosed monkey | n=373 med:0.188 (0.159-0.210) | n=263 med:0.133 (0.110-0.152) | n=587 med:0.101 (0.086-0.114) |
| Brazilian guinea pig | n=266 med:0.128 (0.109-0.148) | n=197 med:0.103 (0.092-0.114) | n=441 med:0.082 (0.074-0.092) |
| Orangutan | n=374 med:0.156 (0.135-0.183) | n=269 med:0.144 (0.120-0.165) | n=596 med:0.101 (0.086-0.117) |
| Northern American deer mouse | n=371 med:0.101 (0.087-0.112) | n=261 med:0.083 (0.070-0.091) | n=590 med:0.045 (0.040-0.050) |
| Naked mole-rat female | n=347 med:0.126 (0.110-0.140) | n=244 med:0.089 (0.077-0.105) | n=585 med:0.055 (0.050-0.061) |
| Mouse | n=384 med:0.098 (0.087-0.114) | n=275 med:0.077 (0.065-0.089) | n=631 med:0.046 (0.041-0.050) |
| Daurian ground squirrel | n=308 med:0.133 (0.119-0.146) | n=207 med:0.101 (0.088-0.117) | n=488 med:0.056 (0.046-0.063) |
| Tiger | n=300 med:0.133 (0.119-0.149) | n=200 med:0.103 (0.087-0.115) | n=473 med:0.060 (0.053-0.071) |
| Lesser hedgehog tenrec | n=83 med:0.170 (0.133-0.222) | n=70 med:0.110 (0.079-0.166) | n=103 med:0.071 (0.057-0.098) |
| Mongolian gerbil | n=337 med:0.105 (0.096-0.117) | n=231 med:0.077 (0.067-0.087) | n=509 med:0.052 (0.045-0.064) |
| Ma's night monkey | n=354 med:0.188 (0.173-0.217) | n=266 med:0.121 (0.105-0.139) | n=605 med:0.081 (0.069-0.092) |

| | | | |
|---|---|---|---|
| Upper Galilee mountains blind mole rat | n=346 med:0.114 (0.105-0.126) | n=258 med:0.082 (0.066-0.096) | n=598 med:0.047 (0.043-0.053) |
| Kangaroo rat | n=315 med:0.106 (0.091-0.117) | n=244 med:0.086 (0.075-0.097) | n=564 med:0.052 (0.046-0.056) |
| Chimpanzee | n=353 med:0.173 (0.127-0.211) | n=264 med:0.134 (0.101-0.158) | n=599 med:0.095 (0.074-0.114) |
| Guinea Pig | n=357 med:0.114 (0.103-0.126) | n=257 med:0.097 (0.082-0.104) | n=589 med:0.056 (0.048-0.063) |
| American beaver | n=356 med:0.125 (0.113-0.143) | n=244 med:0.098 (0.085-0.107) | n=487 med:0.060 (0.051-0.068) |
| Wild yak | n=329 med:0.123 (0.116-0.139) | n=221 med:0.102 (0.087-0.112) | n=518 med:0.053 (0.047-0.060) |
| Coquerel's sifaka | n=361 med:0.139 (0.123-0.161) | n=261 med:0.093 (0.082-0.111) | n=592 med:0.058 (0.052-0.067) |
| Alpaca | n=61 med:0.199 (0.167-0.249) | n=37 med:0.118 (0.069-0.236) | n=99 med:0.088 (0.066-0.106) |
| Lesser Egyptian jerboa | n=311 med:0.097 (0.087-0.109) | n=243 med:0.094 (0.084-0.107) | n=563 med:0.046 (0.042-0.052) |
| Long-tailed chinchilla | n=350 med:0.106 (0.096-0.118) | n=254 med:0.083 (0.068-0.094) | n=590 med:0.050 (0.044-0.055) |
| Angola colobus | n=371 med:0.178 (0.147-0.209) | n=260 med:0.123 (0.106-0.139) | n=600 med:0.082 (0.068-0.092) |
| Ryukyu mouse | n=369 med:0.103 (0.092-0.117) | n=266 med:0.078 (0.070-0.092) | n=614 med:0.048 (0.043-0.052) |
| Chinese hamster PICR | n=372 med:0.115 (0.104-0.131) | n=260 med:0.081 (0.067-0.097) | n=601 med:0.048 (0.043-0.054) |
| Vervet-AGM | n=384 med:0.152 (0.125-0.170) | n=282 med:0.099 (0.086-0.113) | n=615 med:0.068 (0.059-0.077) |
| Cow | n=360 med:0.117 (0.105-0.136) | n=260 med:0.085 (0.074-0.102) | n=598 med:0.050 (0.045-0.058) |
| Bushbaby | n=362 med:0.131 (0.116-0.146) | n=272 med:0.096 (0.084-0.105) | n=607 med:0.055 (0.050-0.062) |

| | | | |
|---|---|---|---|
| Bonobo | n=361 med:0.241 (0.190-0.266) | n=259 med:0.180 (0.145-0.232) | n=591 med:0.136 (0.115-0.166) |
| Hyrax | n=101 med:0.136 (0.105-0.168) | n=74 med:0.096 (0.080-0.114) | n=138 med:0.058 (0.053-0.068) |
| Rabbit | n=316 med:0.126 (0.107-0.141) | n=224 med:0.091 (0.077-0.106) | n=534 med:0.066 (0.059-0.075) |
| Megabat | n=188 med:0.156 (0.137-0.181) | n=131 med:0.096 (0.087-0.125) | n=252 med:0.074 (0.060-0.089) |
| Dog | n=348 med:0.134 (0.120-0.148) | n=247 med:0.100 (0.085-0.115) | n=584 med:0.064 (0.056-0.075) |
| Steppe mouse | n=350 med:0.110 (0.096-0.119) | n=260 med:0.091 (0.078-0.099) | n=596 med:0.050 (0.044-0.057) |
| Greater bamboo lemur | n=373 med:0.142 (0.128-0.154) | n=272 med:0.101 (0.094-0.118) | n=598 med:0.060 (0.054-0.069) |
| Pig | n=363 med:0.130 (0.115-0.142) | n=262 med:0.096 (0.085-0.111) | n=597 med:0.057 (0.052-0.064) |
| Tree Shrew | n=72 med:0.173 (0.142-0.233) | n=47 med:0.113 (0.071-0.149) | n=79 med:0.099 (0.085-0.124) |
| Pika | n=133 med:0.116 (0.098-0.146) | n=94 med:0.081 (0.066-0.104) | n=153 med:0.061 (0.041-0.078) |
| Shrew | n=55 med:0.149 (0.119-0.185) | n=39 med:0.073 (0.051-0.122) | n=82 med:0.082 (0.050-0.105) |
| Squirrel | n=354 med:0.113 (0.100-0.128) | n=259 med:0.086 (0.072-0.096) | n=605 med:0.048 (0.042-0.055) |
| Gorilla | n=368 med:0.165 (0.132-0.189) | n=263 med:0.171 (0.131-0.202) | n=604 med:0.106 (0.082-0.127) |
| Cat | n=366 med:0.126 (0.117-0.142) | n=260 med:0.093 (0.077-0.104) | n=603 med:0.059 (0.053-0.066) |

#### Human Reference Genome, Saunders et al., 2018, cell subtypes

| Sample Size and Median dN/dS (CI95%) | Endothelia-specific Genes | Microglia-specific Genes | Astrocyte-specific Genes | Oligodendrocyte-specific Genes | Neuron-specific Genes |
|---|---|---|---|---|---|
| Microbat | n=278 med:0.128 (0.115-0.144) | n=203 med:0.150 (0.132-0.172) | n=202 med:0.106 (0.090-0.120) | n=193 med:0.096 (0.078-0.114) | n=459 med:0.072 (0.065-0.079) |
| Hedgehog | n=70 med:0.173 (0.139-0.204) | n=46 med:0.137 (0.101-0.160) | n=47 med:0.088 (0.067-0.123) | n=52 med:0.098 (0.075-0.143) | n=102 med:0.072 (0.056-0.092) |
| Naked mole-rat male | n=288 med:0.115 (0.101-0.132) | n=210 med:0.142 (0.119-0.156) | n=217 med:0.097 (0.086-0.112) | n=198 med:0.089 (0.078-0.104) | n=502 med:0.057 (0.052-0.064) |
| Sooty mangabey | n=328 med:0.152 (0.126-0.172) | n=242 med:0.158 (0.136-0.190) | n=260 med:0.118 (0.094-0.139) | n=243 med:0.104 (0.084-0.122) | n=566 med:0.077 (0.067-0.090) |
| Olive baboon | n=330 med:0.168 (0.139-0.188) | n=244 med:0.158 (0.136-0.189) | n=257 med:0.125 (0.105-0.141) | n=241 med:0.104 (0.078-0.125) | n=571 med:0.079 (0.072-0.089) |
| American bison | n=276 med:0.129 (0.108-0.146) | n=202 med:0.134 (0.111-0.152) | n=195 med:0.102 (0.086-0.112) | n=193 med:0.101 (0.085-0.114) | n=451 med:0.055 (0.046-0.062) |
| Leopard | n=322 med:0.107 (0.095-0.128) | n=227 med:0.125 (0.111-0.146) | n=251 med:0.085 (0.074-0.099) | n=224 med:0.076 (0.064-0.089) | n=552 med:0.052 (0.047-0.058) |
| Bolivian squirrel monkey | n=322 med:0.160 (0.138-0.199) | n=223 med:0.171 (0.147-0.193) | n=235 med:0.123 (0.110-0.145) | n=236 med:0.100 (0.084-0.117) | n=546 med:0.074 (0.063-0.081) |
| Crab-eating macaque | n=333 med:0.140 (0.121-0.167) | n=242 med:0.149 (0.130-0.172) | n=254 med:0.119 (0.101-0.133) | n=239 med:0.087 (0.075-0.122) | n=565 med:0.078 (0.066-0.088) |
| Red fox | n=309 med:0.124 (0.112-0.141) | n=229 med:0.139 (0.123-0.159) | n=222 med:0.102 (0.087-0.118) | n=223 med:0.095 (0.082-0.109) | n=533 med:0.073 (0.063-0.082) |
| Shrew mouse | n=329 med:0.104 (0.089-0.115) | n=238 med:0.123 (0.108-0.134) | n=254 med:0.074 (0.064-0.091) | n=238 med:0.076 (0.064-0.085) | n=570 med:0.049 (0.045-0.053) |
| Arctic ground squirrel | n=323 med:0.115 (0.103-0.128) | n=227 med:0.131 (0.112-0.146) | n=247 med:0.083 (0.072-0.096) | n=232 med:0.078 (0.063-0.093) | n=556 med:0.047 (0.041-0.052) |
| Chinese hamster CriGri | n=307 med:0.112 (0.096-0.125) | n=209 med:0.133 (0.118-0.149) | n=224 med:0.082 (0.074-0.097) | n=216 med:0.084 (0.067-0.101) | n=502 med:0.057 (0.052-0.063) |
| Opossum | n=290 med:0.092 (0.084-0.099) | n=208 med:0.090 (0.080-0.111) | n=222 med:0.078 (0.065-0.091) | n=210 med:0.068 (0.058-0.078) | n=482 med:0.056 (0.049-0.063) |
| Dolphin | n=186 med:0.150 | n=121 med:0.149 | n=112 med:0.119 | n=114 med:0.116 | n=267 med:0.067 |

| | | | | | |
|---|---|---|---|---|---|
| | (0.125-0.170) | (0.128-0.163) | (0.095-0.139) | (0.090-0.140) | (0.061-0.075) |
| Capuchin | n=322<br>med:0.155<br>(0.134-0.187) | n=228<br>med:0.170<br>(0.145-0.195) | n=246<br>med:0.124<br>(0.102-0.142) | n=237<br>med:0.087<br>(0.076-0.112) | n=553<br>med:0.067<br>(0.057-0.077) |
| Mouse Lemur | n=324<br>med:0.130<br>(0.113-0.155) | n=224<br>med:0.148<br>(0.131-0.167) | n=241<br>med:0.098<br>(0.089-0.119) | n=235<br>med:0.083<br>(0.071-0.094) | n=552<br>med:0.060<br>(0.053-0.067) |
| Damara mole rat | n=300<br>med:0.114<br>(0.101-0.132) | n=205<br>med:0.127<br>(0.101-0.147) | n=230<br>med:0.097<br>(0.086-0.112) | n=217<br>med:0.077<br>(0.067-0.092) | n=534<br>med:0.053<br>(0.047-0.058) |
| Marmoset | n=320<br>med:0.161<br>(0.145-0.186) | n=225<br>med:0.168<br>(0.144-0.192) | n=238<br>med:0.130<br>(0.107-0.150) | n=220<br>med:0.097<br>(0.080-0.117) | n=555<br>med:0.072<br>(0.062-0.083) |
| Goat | n=325<br>med:0.120<br>(0.103-0.136) | n=234<br>med:0.130<br>(0.112-0.143) | n=244<br>med:0.086<br>(0.076-0.098) | n=236<br>med:0.084<br>(0.072-0.102) | n=550<br>med:0.058<br>(0.050-0.064) |
| Common wombat | n=276<br>med:0.090<br>(0.079-0.099) | n=209<br>med:0.101<br>(0.084-0.117) | n=209<br>med:0.079<br>(0.068-0.089) | n=188<br>med:0.072<br>(0.061-0.081) | n=463<br>med:0.050<br>(0.046-0.056) |
| American black bear | n=297<br>med:0.137<br>(0.112-0.152) | n=208<br>med:0.144<br>(0.128-0.165) | n=205<br>med:0.104<br>(0.090-0.125) | n=211<br>med:0.096<br>(0.078-0.114) | n=456<br>med:0.076<br>(0.066-0.089) |
| Sloth | n=40<br>med:0.194<br>(0.147-0.341) | n=29<br>med:0.154<br>(0.128-0.239) | n=27<br>med:0.157<br>(0.093-0.241) | n=30<br>med:0.177<br>(0.087-0.262) | n=58<br>med:0.100<br>(0.073-0.135) |
| Polar bear | n=282<br>med:0.139<br>(0.124-0.156) | n=203<br>med:0.152<br>(0.143-0.175) | n=214<br>med:0.126<br>(0.105-0.135) | n=210<br>med:0.111<br>(0.096-0.123) | n=477<br>med:0.091<br>(0.080-0.097) |
| Alpine marmot | n=302<br>med:0.115<br>(0.107-0.135) | n=222<br>med:0.133<br>(0.117-0.150) | n=223<br>med:0.084<br>(0.069-0.100) | n=213<br>med:0.084<br>(0.066-0.097) | n=513<br>med:0.053<br>(0.046-0.058) |
| Prairie vole | n=312<br>med:0.094<br>(0.085-0.103) | n=224<br>med:0.120<br>(0.102-0.136) | n=246<br>med:0.074<br>(0.067-0.090) | n=232<br>med:0.072<br>(0.060-0.085) | n=550<br>med:0.045<br>(0.040-0.050) |
| Donkey | n=326<br>med:0.139<br>(0.118-0.155) | n=235<br>med:0.151<br>(0.134-0.172) | n=252<br>med:0.101<br>(0.088-0.111) | n=241<br>med:0.095<br>(0.075-0.112) | n=553<br>med:0.074<br>(0.063-0.081) |
| Golden Hamster | n=303<br>med:0.106<br>(0.094-0.121) | n=216<br>med:0.129<br>(0.120-0.150) | n=236<br>med:0.088<br>(0.075-0.100) | n=215<br>med:0.084<br>(0.069-0.096) | n=525<br>med:0.054<br>(0.048-0.060) |
| Armadillo | n=295<br>med:0.144<br>(0.115-0.156) | n=223<br>med:0.151<br>(0.123-0.163) | n=228<br>med:0.106<br>(0.090-0.117) | n=212<br>med:0.101<br>(0.088-0.113) | n=495<br>med:0.077<br>(0.070-0.085) |
| Ugandan red Colobus | n=335<br>med:0.157<br>(0.130-0.181) | n=241<br>med:0.171<br>(0.140-0.192) | n=240<br>med:0.112<br>(0.095-0.130) | n=234<br>med:0.094<br>(0.078-0.126) | n=550<br>med:0.068<br>(0.053-0.075) |
| Pig-tailed macaque | n=330<br>med:0.142<br>(0.119-0.169) | n=241<br>med:0.156<br>(0.140-0.189) | n=254<br>med:0.115<br>(0.099-0.133) | n=242<br>med:0.100<br>(0.080-0.116) | n=570<br>med:0.076<br>(0.064-0.087) |
| Panda | n=325 | n=230 | n=243 | n=234 | n=531 |

| | | | | | |
|---|---|---|---|---|---|
| | med:0.120 (0.106-0.137) | med:0.134 (0.119-0.158) | med:0.097 (0.086-0.109) | med:0.085 (0.073-0.097) | med:0.055 (0.048-0.061) |
| Ferret | n=319 med:0.113 (0.101-0.124) | n=235 med:0.119 (0.104-0.135) | n=236 med:0.091 (0.077-0.101) | n=228 med:0.080 (0.069-0.093) | n=537 med:0.055 (0.048-0.063) |
| Koala | n=300 med:0.091 (0.080-0.102) | n=223 med:0.105 (0.087-0.119) | n=228 med:0.079 (0.068-0.090) | n=217 med:0.074 (0.068-0.082) | n=513 med:0.055 (0.050-0.063) |
| Wallaby | n=74 med:0.079 (0.058-0.096) | n=65 med:0.087 (0.065-0.120) | n=66 med:0.073 (0.057-0.093) | n=71 med:0.081 (0.059-0.098) | n=121 med:0.050 (0.038-0.064) |
| Tarsier | n=271 med:0.135 (0.115-0.160) | n=195 med:0.144 (0.121-0.163) | n=204 med:0.105 (0.089-0.118) | n=197 med:0.093 (0.074-0.122) | n=437 med:0.068 (0.060-0.080) |
| Degu | n=271 med:0.102 (0.095-0.121) | n=199 med:0.125 (0.109-0.141) | n=219 med:0.088 (0.071-0.099) | n=206 med:0.069 (0.060-0.087) | n=519 med:0.049 (0.044-0.056) |
| Chinese hamster CHOK1GS | n=334 med:0.108 (0.094-0.118) | n=232 med:0.124 (0.109-0.139) | n=252 med:0.080 (0.071-0.088) | n=234 med:0.082 (0.066-0.097) | n=564 med:0.050 (0.045-0.057) |
| Drill | n=317 med:0.160 (0.141-0.187) | n=237 med:0.182 (0.156-0.200) | n=246 med:0.134 (0.118-0.156) | n=231 med:0.108 (0.089-0.143) | n=535 med:0.090 (0.081-0.101) |
| Gibbon | n=316 med:0.191 (0.177-0.215) | n=233 med:0.219 (0.187-0.253) | n=242 med:0.185 (0.160-0.198) | n=225 med:0.142 (0.119-0.175) | n=549 med:0.129 (0.110-0.150) |
| Rat | n=314 med:0.100 (0.091-0.112) | n=229 med:0.125 (0.112-0.140) | n=241 med:0.074 (0.066-0.086) | n=230 med:0.078 (0.062-0.087) | n=548 med:0.049 (0.045-0.054) |
| Macaque | n=330 med:0.161 (0.131-0.181) | n=242 med:0.168 (0.153-0.215) | n=251 med:0.139 (0.120-0.167) | n=236 med:0.105 (0.077-0.134) | n=566 med:0.072 (0.060-0.084) |
| American mink | n=320 med:0.111 (0.097-0.124) | n=226 med:0.120 (0.108-0.136) | n=226 med:0.084 (0.070-0.101) | n=230 med:0.081 (0.066-0.093) | n=527 med:0.052 (0.045-0.063) |
| Gelada | n=331 med:0.128 (0.104-0.154) | n=241 med:0.135 (0.118-0.174) | n=257 med:0.107 (0.093-0.123) | n=240 med:0.083 (0.075-0.106) | n=572 med:0.058 (0.048-0.068) |
| Sheep | n=307 med:0.135 (0.123-0.147) | n=227 med:0.141 (0.127-0.163) | n=228 med:0.100 (0.091-0.115) | n=222 med:0.092 (0.080-0.106) | n=525 med:0.067 (0.060-0.078) |
| Tasmanian devil | n=270 med:0.088 (0.078-0.097) | n=201 med:0.097 (0.083-0.108) | n=209 med:0.078 (0.066-0.087) | n=199 med:0.069 (0.060-0.077) | n=464 med:0.051 (0.045-0.058) |
| Elephant | n=312 med:0.128 (0.117-0.152) | n=230 med:0.154 (0.132-0.173) | n=239 med:0.103 (0.092-0.118) | n=233 med:0.094 (0.077-0.108) | n=533 med:0.071 (0.063-0.080) |
| Algerian mouse | n=328 med:0.106 (0.093-0.118) | n=235 med:0.121 (0.106-0.134) | n=254 med:0.076 (0.068-0.091) | n=232 med:0.074 (0.062-0.088) | n=568 med:0.048 (0.043-0.052) |

| | | | | | |
|---|---|---|---|---|---|
| Dingo | n=313 med:0.116 (0.102-0.130) | n=226 med:0.130 (0.112-0.146) | n=235 med:0.089 (0.077-0.102) | n=223 med:0.078 (0.067-0.093) | n=535 med:0.065 (0.057-0.071) |
| Golden snub-nosed monkey | n=321 med:0.150 (0.124-0.171) | n=229 med:0.171 (0.149-0.197) | n=242 med:0.128 (0.110-0.148) | n=227 med:0.115 (0.095-0.143) | n=537 med:0.084 (0.070-0.095) |
| Horse | n=325 med:0.134 (0.117-0.152) | n=228 med:0.155 (0.135-0.177) | n=242 med:0.101 (0.087-0.122) | n=226 med:0.095 (0.076-0.113) | n=561 med:0.068 (0.058-0.074) |
| Black snub-nosed monkey | n=330 med:0.178 (0.155-0.202) | n=224 med:0.178 (0.155-0.209) | n=238 med:0.143 (0.116-0.161) | n=235 med:0.111 (0.096-0.147) | n=543 med:0.104 (0.088-0.122) |
| Brazilian guinea pig | n=222 med:0.129 (0.108-0.150) | n=167 med:0.140 (0.121-0.159) | n=177 med:0.110 (0.093-0.130) | n=183 med:0.103 (0.091-0.122) | n=412 med:0.082 (0.074-0.092) |
| Orangutan | n=330 med:0.169 (0.139-0.197) | n=241 med:0.172 (0.145-0.218) | n=248 med:0.126 (0.107-0.153) | n=237 med:0.135 (0.112-0.162) | n=547 med:0.100 (0.084-0.117) |
| Northern American deer mouse | n=324 med:0.100 (0.086-0.112) | n=223 med:0.118 (0.101-0.133) | n=245 med:0.079 (0.064-0.088) | n=227 med:0.073 (0.062-0.087) | n=542 med:0.046 (0.041-0.052) |
| Naked mole-rat female | n=311 med:0.115 (0.103-0.130) | n=216 med:0.133 (0.113-0.147) | n=229 med:0.093 (0.080-0.105) | n=211 med:0.081 (0.070-0.104) | n=542 med:0.056 (0.051-0.062) |
| Mouse | n=339 med:0.102 (0.087-0.113) | n=244 med:0.121 (0.110-0.132) | n=259 med:0.075 (0.065-0.086) | n=244 med:0.074 (0.063-0.086) | n=581 med:0.047 (0.043-0.052) |
| Daurian ground squirrel | n=263 med:0.133 (0.118-0.143) | n=192 med:0.139 (0.125-0.156) | n=182 med:0.093 (0.079-0.105) | n=183 med:0.094 (0.076-0.109) | n=446 med:0.058 (0.046-0.065) |
| Tiger | n=252 med:0.132 (0.111-0.149) | n=183 med:0.148 (0.130-0.174) | n=201 med:0.105 (0.089-0.115) | n=171 med:0.098 (0.082-0.110) | n=435 med:0.064 (0.055-0.074) |
| Lesser hedgehog tenrec | n=77 med:0.158 (0.128-0.222) | n=45 med:0.132 (0.091-0.187) | n=66 med:0.119 (0.099-0.156) | n=50 med:0.091 (0.061-0.132) | n=95 med:0.071 (0.055-0.092) |
| Mongolian gerbil | n=301 med:0.104 (0.096-0.117) | n=212 med:0.117 (0.100-0.130) | n=217 med:0.080 (0.073-0.091) | n=209 med:0.080 (0.064-0.092) | n=468 med:0.053 (0.046-0.065) |
| Ma's night monkey | n=318 med:0.176 (0.159-0.192) | n=229 med:0.173 (0.152-0.200) | n=240 med:0.126 (0.111-0.147) | n=228 med:0.099 (0.079-0.125) | n=557 med:0.084 (0.072-0.094) |
| Upper Galilee mountains blind mole rat | n=305 med:0.114 (0.100-0.126) | n=222 med:0.121 (0.112-0.140) | n=252 med:0.085 (0.072-0.099) | n=228 med:0.079 (0.065-0.091) | n=550 med:0.048 (0.044-0.055) |
| Kangaroo rat | n=278 med:0.101 (0.090-0.114) | n=201 med:0.115 (0.100-0.127) | n=231 med:0.090 (0.079-0.104) | n=215 med:0.075 (0.061-0.090) | n=522 med:0.054 (0.047-0.059) |
| Chimpanzee | n=313 | n=230 | n=237 | n=235 | n=550 |

| | | | | | |
|---|---|---|---|---|---|
| | med:0.184 (0.140-0.224) | med:0.211 (0.156-0.261) | med:0.160 (0.117-0.204) | med:0.105 (0.081-0.139) | med:0.099 (0.074-0.120) |
| Guinea Pig | n=315 med:0.113 (0.102-0.125) | n=227 med:0.127 (0.105-0.144) | n=239 med:0.098 (0.076-0.107) | n=226 med:0.078 (0.064-0.096) | n=542 med:0.057 (0.049-0.064) |
| American beaver | n=310 med:0.125 (0.113-0.140) | n=220 med:0.144 (0.123-0.162) | n=227 med:0.102 (0.090-0.114) | n=215 med:0.093 (0.075-0.103) | n=449 med:0.062 (0.052-0.068) |
| Wild yak | n=288 med:0.124 (0.115-0.138) | n=209 med:0.140 (0.119-0.152) | n=203 med:0.099 (0.090-0.113) | n=202 med:0.096 (0.074-0.107) | n=477 med:0.056 (0.048-0.064) |
| Coquerel's sifaka | n=317 med:0.134 (0.116-0.157) | n=226 med:0.145 (0.124-0.166) | n=241 med:0.107 (0.093-0.120) | n=240 med:0.087 (0.078-0.102) | n=541 med:0.059 (0.052-0.071) |
| Alpaca | n=56 med:0.197 (0.141-0.249) | n=39 med:0.178 (0.129-0.271) | n=44 med:0.133 (0.108-0.166) | n=36 med:0.117 (0.076-0.178) | n=92 med:0.093 (0.077-0.108) |
| Lesser Egyptian jerboa | n=279 med:0.096 (0.086-0.106) | n=191 med:0.120 (0.106-0.135) | n=231 med:0.094 (0.081-0.106) | n=214 med:0.078 (0.066-0.090) | n=520 med:0.047 (0.042-0.055) |
| Long-tailed chinchilla | n=307 med:0.103 (0.095-0.116) | n=231 med:0.135 (0.115-0.154) | n=240 med:0.084 (0.073-0.098) | n=224 med:0.072 (0.061-0.086) | n=544 med:0.051 (0.046-0.060) |
| Angola colobus | n=325 med:0.164 (0.140-0.190) | n=235 med:0.185 (0.147-0.202) | n=242 med:0.115 (0.104-0.129) | n=232 med:0.101 (0.084-0.121) | n=550 med:0.086 (0.076-0.098) |
| Ryukyu mouse | n=320 med:0.101 (0.090-0.116) | n=229 med:0.125 (0.113-0.137) | n=250 med:0.075 (0.067-0.083) | n=233 med:0.074 (0.062-0.084) | n=568 med:0.049 (0.044-0.053) |
| Chinese hamster PICR | n=332 med:0.112 (0.102-0.129) | n=229 med:0.127 (0.115-0.145) | n=246 med:0.082 (0.075-0.095) | n=231 med:0.082 (0.067-0.096) | n=551 med:0.050 (0.044-0.057) |
| Vervet-AGM | n=342 med:0.152 (0.128-0.166) | n=242 med:0.156 (0.126-0.178) | n=255 med:0.107 (0.093-0.119) | n=244 med:0.090 (0.080-0.102) | n=565 med:0.072 (0.064-0.088) |
| Cow | n=318 med:0.114 (0.103-0.133) | n=237 med:0.136 (0.118-0.153) | n=234 med:0.089 (0.074-0.101) | n=236 med:0.078 (0.067-0.096) | n=552 med:0.052 (0.046-0.060) |
| Bushbaby | n=328 med:0.128 (0.113-0.146) | n=232 med:0.137 (0.125-0.161) | n=246 med:0.091 (0.079-0.106) | n=240 med:0.089 (0.076-0.103) | n=559 med:0.056 (0.050-0.064) |
| Bonobo | n=322 med:0.248 (0.203-0.281) | n=234 med:0.220 (0.177-0.265) | n=235 med:0.207 (0.153-0.233) | n=233 med:0.174 (0.122-0.222) | n=544 med:0.142 (0.117-0.171) |
| Hyrax | n=88 med:0.141 (0.103-0.182) | n=69 med:0.144 (0.089-0.187) | n=61 med:0.112 (0.086-0.132) | n=69 med:0.090 (0.068-0.119) | n=128 med:0.057 (0.051-0.066) |
| Rabbit | n=275 med:0.119 (0.100-0.141) | n=197 med:0.145 (0.130-0.157) | n=204 med:0.085 (0.073-0.096) | n=206 med:0.081 (0.074-0.099) | n=494 med:0.071 (0.061-0.081) |

| | | | | | |
|---|---|---|---|---|---|
| Megabat | n=179 med:0.149 (0.123-0.171) | n=131 med:0.150 (0.129-0.171) | n=118 med:0.134 (0.110-0.150) | n=107 med:0.088 (0.077-0.117) | n=234 med:0.074 (0.063-0.092) |
| Dog | n=314 med:0.131 (0.115-0.140) | n=226 med:0.130 (0.114-0.146) | n=228 med:0.100 (0.089-0.114) | n=216 med:0.094 (0.071-0.106) | n=538 med:0.067 (0.058-0.077) |
| Steppe mouse | n=314 med:0.109 (0.097-0.119) | n=234 med:0.124 (0.111-0.133) | n=246 med:0.088 (0.071-0.099) | n=228 med:0.088 (0.074-0.096) | n=546 med:0.053 (0.047-0.059) |
| Greater bamboo lemur | n=327 med:0.137 (0.117-0.150) | n=233 med:0.147 (0.123-0.161) | n=248 med:0.107 (0.096-0.120) | n=236 med:0.094 (0.084-0.103) | n=549 med:0.063 (0.055-0.072) |
| Pig | n=322 med:0.128 (0.110-0.139) | n=228 med:0.144 (0.123-0.162) | n=238 med:0.097 (0.086-0.108) | n=232 med:0.086 (0.074-0.096) | n=549 med:0.058 (0.053-0.066) |
| Tree Shrew | n=59 med:0.173 (0.122-0.225) | n=43 med:0.140 (0.100-0.189) | n=52 med:0.111 (0.067-0.192) | n=31 med:0.118 (0.060-0.188) | n=71 med:0.096 (0.071-0.121) |
| Pika | n=108 med:0.110 (0.081-0.142) | n=86 med:0.133 (0.097-0.155) | n=83 med:0.088 (0.065-0.101) | n=82 med:0.078 (0.064-0.097) | n=135 med:0.064 (0.042-0.079) |
| Shrew | n=47 med:0.149 (0.095-0.203) | n=40 med:0.102 (0.069-0.137) | n=32 med:0.082 (0.059-0.110) | n=38 med:0.097 (0.048-0.122) | n=76 med:0.082 (0.050-0.105) |
| Squirrel | n=314 med:0.108 (0.096-0.121) | n=220 med:0.126 (0.106-0.142) | n=244 med:0.087 (0.073-0.096) | n=229 med:0.079 (0.067-0.094) | n=554 med:0.049 (0.043-0.056) |
| Gorilla | n=326 med:0.180 (0.135-0.198) | n=239 med:0.185 (0.149-0.221) | n=246 med:0.134 (0.108-0.168) | n=236 med:0.136 (0.112-0.174) | n=557 med:0.114 (0.087-0.135) |
| Cat | n=328 med:0.124 (0.110-0.137) | n=241 med:0.133 (0.117-0.155) | n=247 med:0.100 (0.085-0.113) | n=229 med:0.082 (0.070-0.098) | n=552 med:0.061 (0.054-0.069) |

#### Human Reference Genome, Zeisel et al., 2018, three cell types

| Sample Size and Median dN/dS (CI95%) | CNS_Glia-specific Genes | CNS_Endothelia-specific Genes | CNS_Neuron-specific Genes |
|---|---|---|---|
| Microbat | n=685 med:0.157 (0.149-0.173) | n=122 med:0.174 (0.133-0.204) | n=2,770 med:0.099 (0.094-0.104) |
| Hedgehog | n=158 med:0.141 (0.128-0.170) | n=33 med:0.161 (0.139-0.211) | n=592 med:0.103 (0.093-0.112) |
| Naked mole-rat male | n=726 med:0.154 (0.141-0.161) | n=139 med:0.142 (0.122-0.166) | n=2,725 med:0.085 (0.081-0.088) |
| Sooty mangabey | n=841 med:0.198 (0.175-0.213) | n=156 med:0.160 (0.130-0.183) | n=3,233 med:0.113 (0.107-0.118) |
| Olive baboon | n=835 med:0.197 (0.183-0.215) | n=155 med:0.177 (0.139-0.197) | n=3,233 med:0.111 (0.107-0.117) |
| American bison | n=684 med:0.152 (0.138-0.164) | n=124 med:0.143 (0.129-0.184) | n=2,588 med:0.084 (0.080-0.089) |
| Leopard | n=809 med:0.139 (0.127-0.151) | n=147 med:0.134 (0.121-0.150) | n=3,142 med:0.073 (0.070-0.076) |
| Bolivian squirrel monkey | n=811 med:0.193 (0.178-0.210) | n=149 med:0.211 (0.156-0.237) | n=3,081 med:0.107 (0.100-0.113) |
| Crab-eating macaque | n=843 med:0.182 (0.168-0.199) | n=154 med:0.160 (0.125-0.178) | n=3,202 med:0.105 (0.101-0.111) |
| Red fox | n=784 med:0.146 (0.134-0.158) | n=142 med:0.149 (0.122-0.176) | n=2,998 med:0.092 (0.089-0.096) |
| Shrew mouse | n=813 med:0.126 (0.116-0.134) | n=152 med:0.121 (0.099-0.138) | n=3,164 med:0.071 (0.068-0.074) |
| Arctic ground squirrel | n=809 med:0.142 (0.130-0.150) | n=154 med:0.133 (0.112-0.154) | n=3,134 med:0.073 (0.070-0.077) |
| Chinese hamster CriGri | n=754 med:0.137 (0.124-0.146) | n=140 med:0.129 (0.112-0.155) | n=2,911 med:0.078 (0.074-0.081) |
| Opossum | n=707 med:0.103 (0.096-0.111) | n=132 med:0.093 (0.082-0.114) | n=2,789 med:0.073 (0.070-0.075) |
| Dolphin | n=422 | n=86 | n=1,620 |

| | | | |
|---|---|---|---|
| | med:0.162<br>(0.149-0.184) | med:0.161<br>(0.139-0.195) | med:0.099<br>(0.094-0.105) |
| Capuchin | n=830<br>med:0.194<br>(0.176-0.206) | n=153<br>med:0.186<br>(0.148-0.209) | n=3,133<br>med:0.101<br>(0.096-0.106) |
| Mouse Lemur | n=781<br>med:0.160<br>(0.148-0.172) | n=153<br>med:0.149<br>(0.120-0.173) | n=3,072<br>med:0.086<br>(0.082-0.091) |
| Damara mole rat | n=741<br>med:0.139<br>(0.126-0.149) | n=131<br>med:0.143<br>(0.117-0.172) | n=2,941<br>med:0.078<br>(0.074-0.082) |
| Marmoset | n=813<br>med:0.193<br>(0.176-0.210) | n=152<br>med:0.171<br>(0.151-0.194) | n=3,124<br>med:0.107<br>(0.102-0.113) |
| Goat | n=796<br>med:0.141<br>(0.130-0.149) | n=146<br>med:0.134<br>(0.115-0.152) | n=3,098<br>med:0.080<br>(0.077-0.084) |
| Common wombat | n=674<br>med:0.104<br>(0.098-0.113) | n=124<br>med:0.100<br>(0.081-0.123) | n=2,711<br>med:0.070<br>(0.066-0.072) |
| American black bear | n=706<br>med:0.142<br>(0.131-0.156) | n=135<br>med:0.153<br>(0.124-0.172) | n=2,739<br>med:0.098<br>(0.093-0.102) |
| Sloth | n=105<br>med:0.218<br>(0.179-0.273) | n=18<br>med:0.234<br>(0.156-0.412) | n=397<br>med:0.131<br>(0.119-0.149) |
| Polar bear | n=693<br>med:0.167<br>(0.155-0.180) | n=125<br>med:0.179<br>(0.154-0.203) | n=2,734<br>med:0.104<br>(0.101-0.108) |
| Alpine marmot | n=756<br>med:0.144<br>(0.135-0.156) | n=143<br>med:0.153<br>(0.137-0.174) | n=2,978<br>med:0.078<br>(0.074-0.081) |
| Prairie vole | n=772<br>med:0.123<br>(0.116-0.133) | n=151<br>med:0.115<br>(0.096-0.133) | n=3,070<br>med:0.069<br>(0.065-0.072) |
| Donkey | n=799<br>med:0.158<br>(0.146-0.169) | n=148<br>med:0.151<br>(0.125-0.175) | n=3,164<br>med:0.094<br>(0.090-0.098) |
| Golden Hamster | n=749<br>med:0.131<br>(0.125-0.144) | n=147<br>med:0.130<br>(0.110-0.153) | n=2,910<br>med:0.076<br>(0.073-0.079) |
| Armadillo | n=743<br>med:0.160<br>(0.152-0.169) | n=131<br>med:0.164<br>(0.146-0.191) | n=2,865<br>med:0.098<br>(0.094-0.104) |
| Ugandan red Colobus | n=824<br>med:0.195<br>(0.181-0.211) | n=149<br>med:0.171<br>(0.144-0.211) | n=3,120<br>med:0.106<br>(0.098-0.112) |
| Pig-tailed macaque | n=836 | n=157 | n=3,234 |

| | | | |
|---|---|---|---|
| | med:0.191<br>(0.174-0.205) | med:0.172<br>(0.142-0.197) | med:0.113<br>(0.107-0.119) |
| Panda | n=790<br>med:0.144<br>(0.135-0.152) | n=148<br>med:0.139<br>(0.116-0.168) | n=3,097<br>med:0.083<br>(0.078-0.086) |
| Ferret | n=790<br>med:0.136<br>(0.123-0.148) | n=146<br>med:0.120<br>(0.108-0.141) | n=3,105<br>med:0.077<br>(0.074-0.080) |
| Koala | n=739<br>med:0.105<br>(0.099-0.113) | n=138<br>med:0.104<br>(0.087-0.117) | n=2,970<br>med:0.072<br>(0.069-0.075) |
| Wallaby | n=211<br>med:0.088<br>(0.077-0.099) | n=52<br>med:0.088<br>(0.069-0.122) | n=841<br>med:0.075<br>(0.068-0.080) |
| Tarsier | n=693<br>med:0.165<br>(0.151-0.179) | n=125<br>med:0.163<br>(0.135-0.190) | n=2,601<br>med:0.094<br>(0.090-0.099) |
| Degu | n=701<br>med:0.131<br>(0.123-0.141) | n=136<br>med:0.115<br>(0.100-0.139) | n=2,736<br>med:0.075<br>(0.071-0.079) |
| Chinese hamster CHOK1GS | n=808<br>med:0.130<br>(0.119-0.141) | n=156<br>med:0.129<br>(0.113-0.144) | n=3,172<br>med:0.074<br>(0.071-0.077) |
| Drill | n=814<br>med:0.198<br>(0.184-0.213) | n=145<br>med:0.179<br>(0.151-0.209) | n=3,115<br>med:0.123<br>(0.116-0.128) |
| Gibbon | n=810<br>med:0.263<br>(0.246-0.281) | n=147<br>med:0.214<br>(0.177-0.253) | n=3,138<br>med:0.162<br>(0.155-0.172) |
| Rat | n=790<br>med:0.127<br>(0.116-0.135) | n=144<br>med:0.116<br>(0.099-0.137) | n=3,105<br>med:0.072<br>(0.069-0.075) |
| Macaque | n=830<br>med:0.197<br>(0.182-0.214) | n=154<br>med:0.169<br>(0.131-0.207) | n=3,179<br>med:0.114<br>(0.108-0.120) |
| American mink | n=781<br>med:0.137<br>(0.126-0.148) | n=145<br>med:0.130<br>(0.114-0.162) | n=3,020<br>med:0.077<br>(0.073-0.080) |
| Gelada | n=840<br>med:0.180<br>(0.167-0.193) | n=156<br>med:0.154<br>(0.124-0.180) | n=3,226<br>med:0.092<br>(0.087-0.096) |
| Sheep | n=776<br>med:0.156<br>(0.142-0.165) | n=140<br>med:0.149<br>(0.132-0.184) | n=3,037<br>med:0.095<br>(0.092-0.098) |
| Tasmanian devil | n=683<br>med:0.099<br>(0.091-0.106) | n=125<br>med:0.106<br>(0.089-0.119) | n=2,665<br>med:0.068<br>(0.065-0.071) |
| Elephant | n=767 | n=138 | n=3,049 |

| | | | |
|---|---|---|---|
| | med:0.150 (0.140-0.161) | med:0.154 (0.124-0.182) | med:0.094 (0.089-0.097) |
| Algerian mouse | n=811 med:0.127 (0.119-0.135) | n=153 med:0.118 (0.100-0.142) | n=3,197 med:0.070 (0.068-0.074) |
| Dingo | n=766 med:0.137 (0.124-0.147) | n=148 med:0.139 (0.121-0.163) | n=3,048 med:0.083 (0.079-0.086) |
| Golden snub-nosed monkey | n=817 med:0.200 (0.182-0.218) | n=157 med:0.188 (0.158-0.219) | n=3,085 med:0.116 (0.108-0.123) |
| Horse | n=801 med:0.168 (0.156-0.178) | n=148 med:0.161 (0.134-0.180) | n=3,167 med:0.093 (0.088-0.096) |
| Black snub-nosed monkey | n=813 med:0.213 (0.188-0.236) | n=150 med:0.202 (0.170-0.237) | n=3,102 med:0.139 (0.132-0.146) |
| Brazilian guinea pig | n=591 med:0.146 (0.136-0.158) | n=103 med:0.145 (0.125-0.168) | n=2,274 med:0.111 (0.105-0.115) |
| Orangutan | n=799 med:0.217 (0.195-0.230) | n=149 med:0.197 (0.154-0.227) | n=3,134 med:0.133 (0.127-0.139) |
| Northern American deer mouse | n=795 med:0.126 (0.117-0.136) | n=151 med:0.120 (0.104-0.133) | n=3,033 med:0.069 (0.066-0.072) |
| Naked mole-rat female | n=766 med:0.153 (0.139-0.161) | n=152 med:0.138 (0.115-0.159) | n=2,905 med:0.081 (0.078-0.085) |
| Mouse | n=836 med:0.123 (0.117-0.134) | n=159 med:0.118 (0.098-0.143) | n=3,287 med:0.069 (0.066-0.072) |
| Daurian ground squirrel | n=671 med:0.150 (0.140-0.163) | n=125 med:0.161 (0.134-0.181) | n=2,602 med:0.086 (0.080-0.091) |
| Tiger | n=693 med:0.162 (0.150-0.174) | n=110 med:0.160 (0.134-0.190) | n=2,534 med:0.092 (0.087-0.095) |
| Lesser hedgehog tenrec | n=175 med:0.166 (0.141-0.204) | n=37 med:0.179 (0.138-0.236) | n=673 med:0.102 (0.091-0.114) |
| Mongolian gerbil | n=728 med:0.129 (0.119-0.134) | n=142 med:0.118 (0.096-0.134) | n=2,784 med:0.075 (0.072-0.079) |
| Ma's night monkey | n=807 med:0.207 (0.192-0.226) | n=148 med:0.181 (0.165-0.225) | n=3,098 med:0.116 (0.111-0.122) |

| | | | |
|---|---|---|---|
| Upper Galilee mountains blind mole rat | n=752 med:0.137 (0.127-0.146) | n=146 med:0.125 (0.109-0.148) | n=3,010 med:0.072 (0.069-0.075) |
| Kangaroo rat | n=690 med:0.127 (0.119-0.140) | n=129 med:0.118 (0.096-0.144) | n=2,842 med:0.076 (0.072-0.080) |
| Chimpanzee | n=801 med:0.241 (0.218-0.263) | n=142 med:0.177 (0.129-0.240) | n=3,096 med:0.148 (0.136-0.156) |
| Guinea Pig | n=771 med:0.129 (0.118-0.143) | n=146 med:0.122 (0.100-0.140) | n=3,064 med:0.080 (0.077-0.083) |
| American beaver | n=727 med:0.146 (0.134-0.156) | n=155 med:0.141 (0.118-0.164) | n=2,810 med:0.087 (0.083-0.091) |
| Wild yak | n=731 med:0.150 (0.139-0.161) | n=131 med:0.157 (0.126-0.177) | n=2,717 med:0.086 (0.081-0.090) |
| Coquerel's sifaka | n=790 med:0.158 (0.148-0.169) | n=145 med:0.152 (0.121-0.183) | n=3,077 med:0.091 (0.088-0.098) |
| Alpaca | n=156 med:0.225 (0.164-0.254) | n=27 med:0.208 (0.130-0.290) | n=619 med:0.118 (0.108-0.126) |
| Lesser Egyptian jerboa | n=715 med:0.132 (0.124-0.140) | n=137 med:0.113 (0.092-0.140) | n=2,815 med:0.077 (0.073-0.079) |
| Long-tailed chinchilla | n=796 med:0.133 (0.123-0.146) | n=142 med:0.116 (0.101-0.139) | n=3,086 med:0.072 (0.069-0.075) |
| Angola colobus | n=821 med:0.204 (0.190-0.224) | n=147 med:0.199 (0.163-0.236) | n=3,140 med:0.121 (0.115-0.127) |
| Ryukyu mouse | n=798 med:0.125 (0.116-0.135) | n=153 med:0.127 (0.099-0.151) | n=3,163 med:0.071 (0.067-0.074) |
| Chinese hamster PICR | n=796 med:0.140 (0.129-0.148) | n=154 med:0.130 (0.118-0.155) | n=3,146 med:0.077 (0.074-0.081) |
| Vervet-AGM | n=830 med:0.180 (0.166-0.194) | n=163 med:0.165 (0.133-0.203) | n=3,220 med:0.096 (0.092-0.101) |
| Cow | n=803 med:0.140 (0.131-0.148) | n=148 med:0.126 (0.109-0.151) | n=3,124 med:0.081 (0.077-0.085) |
| Bushbaby | n=798 med:0.156 (0.142-0.166) | n=147 med:0.154 (0.129-0.173) | n=3,131 med:0.088 (0.084-0.092) |

| | | | |
|---|---|---|---|
| Bonobo | n=803 med:0.266 (0.239-0.283) | n=150 med:0.231 (0.174-0.273) | n=3,106 med:0.189 (0.176-0.201) |
| Hyrax | n=222 med:0.154 (0.128-0.177) | n=55 med:0.152 (0.103-0.212) | n=791 med:0.095 (0.086-0.104) |
| Rabbit | n=695 med:0.141 (0.132-0.151) | n=125 med:0.144 (0.125-0.161) | n=2,772 med:0.088 (0.084-0.092) |
| Megabat | n=419 med:0.164 (0.153-0.181) | n=81 med:0.181 (0.152-0.214) | n=1,473 med:0.107 (0.099-0.112) |
| Dog | n=784 med:0.153 (0.138-0.170) | n=143 med:0.160 (0.137-0.178) | n=3,020 med:0.090 (0.086-0.094) |
| Steppe mouse | n=799 med:0.132 (0.124-0.142) | n=148 med:0.121 (0.104-0.150) | n=3,067 med:0.076 (0.073-0.079) |
| Greater bamboo lemur | n=819 med:0.159 (0.149-0.171) | n=155 med:0.154 (0.128-0.180) | n=3,139 med:0.091 (0.086-0.095) |
| Pig | n=790 med:0.146 (0.139-0.153) | n=154 med:0.134 (0.113-0.170) | n=3,126 med:0.085 (0.081-0.089) |
| Tree Shrew | n=150 med:0.163 (0.140-0.188) | n=30 med:0.164 (0.105-0.211) | n=520 med:0.112 (0.100-0.121) |
| Pika | n=276 med:0.108 (0.097-0.131) | n=61 med:0.159 (0.102-0.178) | n=981 med:0.085 (0.079-0.092) |
| Shrew | n=127 med:0.118 (0.096-0.168) | n=27 med:0.128 (0.079-0.242) | n=553 med:0.086 (0.079-0.099) |
| Squirrel | n=785 med:0.138 (0.126-0.146) | n=150 med:0.133 (0.107-0.156) | n=3,128 med:0.073 (0.070-0.076) |
| Gorilla | n=828 med:0.248 (0.221-0.267) | n=150 med:0.185 (0.153-0.227) | n=3,178 med:0.150 (0.142-0.161) |
| Cat | n=816 med:0.147 (0.137-0.160) | n=152 med:0.138 (0.124-0.166) | n=3,165 med:0.084 (0.080-0.088) |

#### Human Reference Genome, Zeisel et al., 2018, cell subtypes

| Sample Size and Median dN/dS (CI95%) | Glia-specific Genes | Immune cells-specific Genes | Vascular cells-specific Genes | Neurons-specific Genes |
|---|---|---|---|---|
| Microbat | n=569 med:0.179 (0.164-0.193) | n=75 med:0.186 (0.144-0.207) | n=149 med:0.144 (0.114-0.171) | n=2,890 med:0.109 (0.104-0.114) |
| Hedgehog | n=133 med:0.158 (0.134-0.198) | n=25 med:0.148 (0.117-0.180) | n=31 med:0.164 (0.145-0.215) | n=659 med:0.112 (0.103-0.123) |
| Naked mole-rat male | n=602 med:0.168 (0.155-0.183) | n=77 med:0.165 (0.134-0.192) | n=157 med:0.138 (0.113-0.161) | n=2,865 med:0.098 (0.093-0.103) |
| Sooty mangabey | n=687 med:0.216 (0.199-0.233) | n=95 med:0.241 (0.198-0.276) | n=178 med:0.151 (0.129-0.177) | n=3,394 med:0.123 (0.118-0.129) |
| Olive baboon | n=684 med:0.222 (0.201-0.238) | n=94 med:0.221 (0.181-0.291) | n=176 med:0.169 (0.136-0.189) | n=3,398 med:0.124 (0.118-0.130) |
| American bison | n=562 med:0.169 (0.153-0.184) | n=80 med:0.190 (0.156-0.223) | n=138 med:0.138 (0.121-0.163) | n=2,735 med:0.094 (0.090-0.099) |
| Leopard | n=662 med:0.159 (0.144-0.173) | n=89 med:0.162 (0.121-0.195) | n=168 med:0.119 (0.103-0.132) | n=3,302 med:0.084 (0.081-0.087) |
| Bolivian squirrel monkey | n=667 med:0.218 (0.202-0.234) | n=90 med:0.228 (0.171-0.296) | n=165 med:0.192 (0.161-0.222) | n=3,236 med:0.120 (0.115-0.124) |
| Crab-eating macaque | n=686 med:0.211 (0.197-0.227) | n=96 med:0.228 (0.184-0.274) | n=176 med:0.142 (0.118-0.174) | n=3,370 med:0.118 (0.113-0.122) |
| Red fox | n=642 med:0.167 (0.149-0.181) | n=88 med:0.187 (0.138-0.209) | n=164 med:0.128 (0.112-0.154) | n=3,135 med:0.101 (0.097-0.106) |
| Shrew mouse | n=665 med:0.145 (0.135-0.157) | n=89 med:0.137 (0.111-0.169) | n=173 med:0.112 (0.102-0.128) | n=3,334 med:0.080 (0.077-0.084) |
| Arctic ground squirrel | n=657 med:0.154 (0.143-0.166) | n=86 med:0.158 (0.136-0.187) | n=179 med:0.123 (0.104-0.141) | n=3,284 med:0.082 (0.079-0.087) |
| Chinese hamster CriGri | n=619 med:0.156 (0.146-0.166) | n=83 med:0.152 (0.125-0.184) | n=159 med:0.128 (0.113-0.145) | n=3,083 med:0.088 (0.084-0.092) |
| Opossum | n=581 med:0.121 (0.110-0.132) | n=71 med:0.086 (0.066-0.131) | n=158 med:0.085 (0.073-0.097) | n=2,931 med:0.078 (0.074-0.081) |
| Dolphin | n=358 | n=48 | n=95 | n=1,706 |

| | | | | |
|---|---|---|---|---|
| | med:0.189 (0.161-0.212) | med:0.193 (0.158-0.268) | med:0.149 (0.117-0.180) | med:0.111 (0.104-0.117) |
| Capuchin | n=678 med:0.219 (0.203-0.236) | n=94 med:0.262 (0.191-0.308) | n=172 med:0.157 (0.141-0.189) | n=3,306 med:0.114 (0.108-0.119) |
| Mouse Lemur | n=649 med:0.178 (0.167-0.199) | n=90 med:0.173 (0.148-0.225) | n=172 med:0.130 (0.108-0.160) | n=3,251 med:0.097 (0.093-0.100) |
| Damara mole rat | n=610 med:0.156 (0.144-0.167) | n=74 med:0.165 (0.107-0.188) | n=156 med:0.139 (0.118-0.154) | n=3,069 med:0.088 (0.085-0.093) |
| Marmoset | n=672 med:0.222 (0.205-0.241) | n=93 med:0.230 (0.169-0.275) | n=177 med:0.174 (0.154-0.200) | n=3,294 med:0.119 (0.115-0.125) |
| Goat | n=649 med:0.160 (0.147-0.171) | n=92 med:0.181 (0.140-0.220) | n=167 med:0.117 (0.096-0.140) | n=3,268 med:0.090 (0.086-0.093) |
| Common wombat | n=552 med:0.121 (0.111-0.136) | n=66 med:0.116 (0.078-0.144) | n=145 med:0.095 (0.081-0.115) | n=2,842 med:0.074 (0.072-0.078) |
| American black bear | n=574 med:0.157 (0.142-0.174) | n=83 med:0.176 (0.132-0.191) | n=153 med:0.128 (0.106-0.152) | n=2,890 med:0.106 (0.102-0.111) |
| Sloth | n=81 med:0.273 (0.200-0.308) | n=9 med:0.480 (0.185-0.519) | n=17 med:0.223 (0.156-0.387) | n=419 med:0.162 (0.147-0.178) |
| Polar bear | n=580 med:0.190 (0.175-0.205) | n=78 med:0.180 (0.144-0.215) | n=140 med:0.154 (0.122-0.181) | n=2,873 med:0.117 (0.111-0.122) |
| Alpine marmot | n=617 med:0.153 (0.141-0.168) | n=87 med:0.154 (0.137-0.184) | n=162 med:0.135 (0.113-0.156) | n=3,132 med:0.091 (0.086-0.095) |
| Prairie vole | n=651 med:0.142 (0.132-0.152) | n=84 med:0.150 (0.120-0.167) | n=174 med:0.117 (0.100-0.134) | n=3,222 med:0.078 (0.075-0.081) |
| Donkey | n=658 med:0.179 (0.167-0.196) | n=91 med:0.173 (0.150-0.216) | n=167 med:0.130 (0.113-0.151) | n=3,333 med:0.104 (0.099-0.110) |
| Golden Hamster | n=628 med:0.152 (0.142-0.163) | n=84 med:0.163 (0.125-0.208) | n=162 med:0.114 (0.097-0.125) | n=3,064 med:0.085 (0.081-0.090) |
| Armadillo | n=609 med:0.180 (0.164-0.192) | n=89 med:0.168 (0.141-0.206) | n=151 med:0.145 (0.118-0.174) | n=3,018 med:0.110 (0.105-0.114) |
| Ugandan red Colobus | n=671 med:0.214 (0.196-0.229) | n=95 med:0.224 (0.172-0.279) | n=167 med:0.146 (0.112-0.177) | n=3,277 med:0.119 (0.112-0.123) |

| | | | | |
|---|---|---|---|---|
| Pig-tailed macaque | n=686 med:0.211 (0.189-0.230) | n=94 med:0.237 (0.207-0.275) | n=179 med:0.154 (0.129-0.182) | n=3,401 med:0.124 (0.119-0.130) |
| Panda | n=647 med:0.155 (0.145-0.170) | n=93 med:0.170 (0.148-0.192) | n=166 med:0.121 (0.105-0.139) | n=3,246 med:0.092 (0.088-0.096) |
| Ferret | n=657 med:0.158 (0.145-0.175) | n=91 med:0.162 (0.131-0.179) | n=168 med:0.106 (0.090-0.129) | n=3,260 med:0.086 (0.083-0.090) |
| Koala | n=608 med:0.124 (0.117-0.134) | n=76 med:0.108 (0.076-0.143) | n=162 med:0.087 (0.080-0.103) | n=3,099 med:0.078 (0.075-0.081) |
| Wallaby | n=154 med:0.086 (0.074-0.099) | n=18 med:0.094 (0.033-0.120) | n=63 med:0.082 (0.073-0.103) | n=927 med:0.078 (0.074-0.083) |
| Tarsier | n=578 med:0.192 (0.178-0.214) | n=78 med:0.175 (0.147-0.228) | n=137 med:0.142 (0.123-0.164) | n=2,750 med:0.106 (0.102-0.110) |
| Degu | n=572 med:0.148 (0.138-0.161) | n=73 med:0.141 (0.112-0.158) | n=160 med:0.122 (0.107-0.140) | n=2,911 med:0.084 (0.081-0.088) |
| Chinese hamster CHOK1GS | n=666 med:0.151 (0.143-0.161) | n=90 med:0.152 (0.125-0.184) | n=181 med:0.120 (0.107-0.135) | n=3,343 med:0.083 (0.080-0.087) |
| Drill | n=673 med:0.219 (0.202-0.245) | n=94 med:0.214 (0.185-0.269) | n=171 med:0.176 (0.144-0.196) | n=3,270 med:0.135 (0.128-0.142) |
| Gibbon | n=672 med:0.284 (0.264-0.305) | n=90 med:0.297 (0.243-0.348) | n=169 med:0.183 (0.164-0.211) | n=3,290 med:0.179 (0.171-0.188) |
| Rat | n=651 med:0.145 (0.136-0.154) | n=88 med:0.142 (0.121-0.185) | n=167 med:0.113 (0.099-0.123) | n=3,251 med:0.081 (0.077-0.084) |
| Macaque | n=687 med:0.221 (0.205-0.236) | n=95 med:0.263 (0.208-0.325) | n=177 med:0.154 (0.126-0.203) | n=3,345 med:0.126 (0.119-0.132) |
| American mink | n=645 med:0.155 (0.145-0.170) | n=88 med:0.153 (0.129-0.176) | n=167 med:0.115 (0.096-0.131) | n=3,182 med:0.087 (0.083-0.090) |
| Gelada | n=690 med:0.207 (0.183-0.223) | n=96 med:0.204 (0.152-0.246) | n=176 med:0.130 (0.105-0.161) | n=3,390 med:0.101 (0.095-0.105) |
| Sheep | n=641 med:0.170 (0.158-0.183) | n=88 med:0.200 (0.147-0.233) | n=162 med:0.146 (0.124-0.163) | n=3,171 med:0.103 (0.099-0.108) |
| Tasmanian devil | n=563 med:0.120 (0.103-0.131) | n=73 med:0.106 (0.083-0.140) | n=146 med:0.081 (0.070-0.099) | n=2,769 med:0.074 (0.072-0.078) |

| | | | | |
|---|---|---|---|---|
| Elephant | n=635 med:0.163 (0.149-0.177) | n=87 med:0.167 (0.148-0.197) | n=164 med:0.129 (0.106-0.169) | n=3,197 med:0.105 (0.100-0.109) |
| Algerian mouse | n=681 med:0.152 (0.140-0.159) | n=89 med:0.150 (0.121-0.185) | n=174 med:0.117 (0.102-0.126) | n=3,363 med:0.080 (0.076-0.083) |
| Dingo | n=618 med:0.155 (0.142-0.172) | n=91 med:0.164 (0.130-0.195) | n=170 med:0.116 (0.097-0.144) | n=3,213 med:0.091 (0.087-0.096) |
| Golden snub-nosed monkey | n=670 med:0.219 (0.201-0.240) | n=94 med:0.231 (0.184-0.278) | n=170 med:0.176 (0.141-0.217) | n=3,235 med:0.129 (0.123-0.135) |
| Horse | n=669 med:0.178 (0.168-0.193) | n=86 med:0.184 (0.162-0.224) | n=168 med:0.139 (0.113-0.165) | n=3,315 med:0.102 (0.097-0.106) |
| Black snub-nosed monkey | n=667 med:0.241 (0.219-0.264) | n=93 med:0.237 (0.189-0.269) | n=173 med:0.176 (0.150-0.225) | n=3,265 med:0.153 (0.145-0.160) |
| Brazilian guinea pig | n=479 med:0.165 (0.152-0.176) | n=62 med:0.154 (0.120-0.195) | n=135 med:0.145 (0.117-0.167) | n=2,357 med:0.117 (0.113-0.122) |
| Orangutan | n=657 med:0.236 (0.222-0.262) | n=93 med:0.235 (0.201-0.309) | n=166 med:0.160 (0.134-0.201) | n=3,285 med:0.144 (0.138-0.151) |
| Northern American deer mouse | n=648 med:0.145 (0.135-0.156) | n=87 med:0.151 (0.130-0.187) | n=172 med:0.118 (0.098-0.132) | n=3,194 med:0.079 (0.075-0.082) |
| Naked mole-rat female | n=636 med:0.168 (0.156-0.181) | n=79 med:0.165 (0.142-0.192) | n=178 med:0.137 (0.118-0.157) | n=3,051 med:0.092 (0.087-0.095) |
| Mouse | n=695 med:0.148 (0.137-0.156) | n=93 med:0.148 (0.115-0.182) | n=184 med:0.116 (0.104-0.126) | n=3,444 med:0.079 (0.075-0.082) |
| Daurian ground squirrel | n=559 med:0.163 (0.148-0.178) | n=76 med:0.163 (0.139-0.224) | n=143 med:0.135 (0.117-0.173) | n=2,740 med:0.098 (0.094-0.103) |
| Tiger | n=585 med:0.177 (0.165-0.194) | n=78 med:0.183 (0.162-0.238) | n=129 med:0.135 (0.120-0.172) | n=2,674 med:0.103 (0.098-0.107) |
| Lesser hedgehog tenrec | n=132 med:0.184 (0.145-0.208) | n=17 med:0.274 (0.166-0.402) | n=41 med:0.138 (0.093-0.190) | n=735 med:0.109 (0.102-0.121) |
| Mongolian gerbil | n=601 med:0.149 (0.137-0.159) | n=77 med:0.152 (0.115-0.191) | n=160 med:0.116 (0.101-0.133) | n=2,935 med:0.084 (0.081-0.089) |
| Ma's night monkey | n=660 med:0.231 (0.210-0.253) | n=92 med:0.221 (0.158-0.275) | n=172 med:0.180 (0.157-0.205) | n=3,259 med:0.128 (0.122-0.134) |

| | | | | |
|---|---|---|---|---|
| Upper Galilee mountains blind mole rat | n=628 med:0.159 (0.144-0.168) | n=82 med:0.157 (0.122-0.190) | n=162 med:0.126 (0.113-0.150) | n=3,189 med:0.083 (0.078-0.088) |
| Kangaroo rat | n=562 med:0.145 (0.133-0.162) | n=70 med:0.134 (0.106-0.164) | n=152 med:0.118 (0.091-0.129) | n=2,981 med:0.086 (0.081-0.090) |
| Chimpanzee | n=659 med:0.261 (0.236-0.281) | n=92 med:0.287 (0.195-0.361) | n=165 med:0.180 (0.153-0.230) | n=3,258 med:0.163 (0.153-0.172) |
| Guinea Pig | n=631 med:0.157 (0.145-0.168) | n=80 med:0.160 (0.124-0.193) | n=173 med:0.118 (0.101-0.135) | n=3,200 med:0.088 (0.085-0.091) |
| American beaver | n=572 med:0.162 (0.149-0.175) | n=90 med:0.160 (0.147-0.185) | n=170 med:0.130 (0.107-0.153) | n=2,980 med:0.098 (0.094-0.102) |
| Wild yak | n=598 med:0.167 (0.151-0.185) | n=87 med:0.173 (0.145-0.212) | n=147 med:0.141 (0.119-0.160) | n=2,872 med:0.096 (0.092-0.102) |
| Coquerel's sifaka | n=653 med:0.180 (0.165-0.202) | n=85 med:0.191 (0.157-0.231) | n=167 med:0.129 (0.110-0.152) | n=3,225 med:0.102 (0.098-0.107) |
| Alpaca | n=131 med:0.231 (0.195-0.278) | n=12 med:0.253 (0.150-0.365) | n=29 med:0.195 (0.124-0.290) | n=644 med:0.132 (0.121-0.143) |
| Lesser Egyptian jerboa | n=583 med:0.149 (0.138-0.161) | n=73 med:0.149 (0.120-0.171) | n=155 med:0.109 (0.089-0.135) | n=2,936 med:0.086 (0.082-0.090) |
| Long-tailed chinchilla | n=650 med:0.158 (0.149-0.168) | n=85 med:0.155 (0.132-0.180) | n=164 med:0.122 (0.104-0.143) | n=3,236 med:0.083 (0.078-0.087) |
| Angola colobus | n=673 med:0.227 (0.207-0.246) | n=93 med:0.233 (0.197-0.301) | n=168 med:0.174 (0.133-0.204) | n=3,284 med:0.137 (0.130-0.142) |
| Ryukyu mouse | n=664 med:0.150 (0.133-0.157) | n=87 med:0.158 (0.123-0.198) | n=175 med:0.120 (0.104-0.136) | n=3,320 med:0.078 (0.075-0.082) |
| Chinese hamster PICR | n=667 med:0.157 (0.146-0.167) | n=90 med:0.154 (0.137-0.187) | n=175 med:0.121 (0.108-0.135) | n=3,312 med:0.086 (0.083-0.091) |
| Vervet-AGM | n=681 med:0.210 (0.193-0.226) | n=94 med:0.198 (0.166-0.241) | n=185 med:0.140 (0.122-0.179) | n=3,390 med:0.110 (0.104-0.116) |
| Cow | n=665 med:0.151 (0.142-0.166) | n=90 med:0.177 (0.158-0.206) | n=172 med:0.124 (0.100-0.141) | n=3,289 med:0.091 (0.087-0.095) |
| Bushbaby | n=650 med:0.172 (0.161-0.192) | n=90 med:0.184 (0.141-0.217) | n=164 med:0.132 (0.119-0.154) | n=3,306 med:0.098 (0.093-0.103) |

| | | | | |
|---|---|---|---|---|
| Bonobo | n=664 med:0.286 (0.267-0.309) | n=95 med:0.267 (0.206-0.340) | n=168 med:0.209 (0.168-0.265) | n=3,243 med:0.200 (0.186-0.212) |
| Hyrax | n=177 med:0.179 (0.149-0.209) | n=32 med:0.221 (0.099-0.346) | n=55 med:0.138 (0.094-0.195) | n=863 med:0.106 (0.098-0.116) |
| Rabbit | n=578 med:0.154 (0.143-0.163) | n=75 med:0.177 (0.147-0.223) | n=148 med:0.121 (0.108-0.146) | n=2,901 med:0.095 (0.091-0.099) |
| Megabat | n=332 med:0.178 (0.155-0.200) | n=44 med:0.236 (0.170-0.315) | n=94 med:0.171 (0.140-0.192) | n=1,569 med:0.116 (0.110-0.122) |
| Dog | n=648 med:0.178 (0.160-0.191) | n=86 med:0.161 (0.128-0.196) | n=164 med:0.132 (0.106-0.156) | n=3,159 med:0.099 (0.095-0.104) |
| Steppe mouse | n=663 med:0.152 (0.143-0.162) | n=92 med:0.153 (0.125-0.177) | n=172 med:0.119 (0.107-0.138) | n=3,219 med:0.085 (0.081-0.088) |
| Greater bamboo lemur | n=674 med:0.178 (0.164-0.194) | n=92 med:0.182 (0.155-0.203) | n=175 med:0.136 (0.112-0.157) | n=3,308 med:0.102 (0.096-0.106) |
| Pig | n=662 med:0.162 (0.150-0.173) | n=88 med:0.157 (0.132-0.190) | n=175 med:0.128 (0.105-0.141) | n=3,278 med:0.093 (0.090-0.097) |
| Tree Shrew | n=127 med:0.200 (0.154-0.231) | n=16 med:0.288 (0.137-0.487) | n=27 med:0.165 (0.105-0.296) | n=560 med:0.121 (0.114-0.132) |
| Pika | n=224 med:0.128 (0.108-0.139) | n=37 med:0.175 (0.108-0.260) | n=71 med:0.139 (0.100-0.164) | n=1,075 med:0.093 (0.086-0.101) |
| Shrew | n=98 med:0.158 (0.104-0.204) | n=16 med:0.252 (0.136-0.292) | n=35 med:0.149 (0.113-0.185) | n=624 med:0.103 (0.094-0.112) |
| Squirrel | n=650 med:0.152 (0.140-0.164) | n=83 med:0.148 (0.129-0.170) | n=171 med:0.117 (0.106-0.143) | n=3,286 med:0.081 (0.078-0.085) |
| Gorilla | n=687 med:0.281 (0.254-0.303) | n=92 med:0.258 (0.199-0.318) | n=171 med:0.195 (0.174-0.247) | n=3,333 med:0.169 (0.156-0.181) |
| Cat | n=674 med:0.168 (0.155-0.180) | n=95 med:0.165 (0.126-0.207) | n=177 med:0.128 (0.105-0.151) | n=3,318 med:0.095 (0.091-0.098) |
