## Supplementary Table S7 for "Neuron-specific coding and regulatory sequences are the most highly conserved in amniote brains despite neuron-specific cell size diversity"

### Chicken Reference Genome, Zhang et al., 2014

| MWU Test | Endothelia vs Glia | Endothelia vs Neuron | Glia vs Neuron |
|---|---|---|---|
| Spoon-billed sandpiper | U=213548.0<br>p=0.219<br>CLES:51.99% | U=311970.0<br>**$p=9.098 \times 10^{-15}$**<br>CLES:62.00% | U=370071.0<br>**$p=9.684 \times 10^{-13}$**<br>CLES:60.44% |
| White-throated sparrow | U=161613.0<br>p=0.678<br>CLES:50.72% | U=236511.0<br>**$p=7.735 \times 10^{-10}$**<br>CLES:60.12% | U=277465.0<br>**$p=8.594 \times 10^{-10}$**<br>CLES:59.62% |
| Abingdon island giant tortoise | U=170031.5<br>p=0.411<br>CLES:51.40% | U=251325.5<br>**$p=1.19 \times 10^{-11}$**<br>CLES:61.02% | U=281664.0<br>**$p=7.558 \times 10^{-10}$**<br>CLES:59.61% |
| Japanese quail | U=227910.0<br>p=0.616<br>CLES:50.79% | U=313350.0<br>**$p=3.028 \times 10^{-7}$**<br>CLES:57.78% | U=376944.0<br>**$p=4.349 \times 10^{-7}$**<br>CLES:57.26% |
| Budgerigar | U=183495.0<br>p=0.833<br>CLES:50.35% | U=274265.5<br>**$p=1.247 \times 10^{-12}$**<br>CLES:61.29% | U=313157.5<br>**$p=1.59 \times 10^{-13}$**<br>CLES:61.30% |
| Turkey | U=173566.5<br>p=0.547<br>CLES:48.99% | U=228278.5<br>**$p=2.263 \times 10^{-5}$**<br>CLES:56.91% | U=271100.5<br>**$p=4.095 \times 10^{-7}$**<br>CLES:57.91% |
| Tuatara | U=167242.5<br>p=0.496<br>CLES:51.16% | U=237875.0<br>**$p=4.548 \times 10^{-10}$**<br>CLES:60.24% | U=280657.0<br>**$p=1.292 \times 10^{-9}$**<br>CLES:59.48% |
| Bengalese finch | U=199690.5<br>p=0.445<br>CLES:51.25% | U=303935.0<br>**$p=4.586 \times 10^{-14}$**<br>CLES:61.75% | U=355275.0<br>**$p=2.851 \times 10^{-13}$**<br>CLES:60.82% |
| Agassiz's desert tortoise | U=183158.0<br>p=0.312<br>CLES:51.70% | U=262241.5<br>**$p=8.945 \times 10^{-11}$**<br>CLES:60.40% | U=297833.0<br>**$p=1.055 \times 10^{-8}$**<br>CLES:58.78% |
| Common canary | U=184285.0<br>p=0.752<br>CLES:50.53% | U=270764.0<br>**$p=6.48 \times 10^{-13}$**<br>CLES:61.49% | U=317823.0<br>**$p=2.327 \times 10^{-13}$**<br>CLES:61.17% |
| Flycatcher | U=191753.5<br>p=0.669<br>CLES:50.70% | U=300480.0<br>**$p=6.801 \times 10^{-14}$**<br>CLES:61.69% | U=344054.0<br>**$p=3.979 \times 10^{-14}$**<br>CLES:61.33% |
| Painted turtle | U=200569.0<br>p=0.385<br>CLES:51.42% | U=301196.0<br>**$p=3.061 \times 10^{-14}$**<br>CLES:61.85% | U=346697.0<br>**$p=5.082 \times 10^{-13}$**<br>CLES:60.77% |
| Great Tit | U=201506.0<br>p=0.6<br>CLES:50.85% | U=311693.0<br>**$p=6.828 \times 10^{-16}$**<br>CLES:62.53% | U=364376.0<br>**$p=1.206 \times 10^{-16}$**<br>CLES:62.27% |
| Duck | U=184969.0<br>p=0.354<br>CLES:51.55% | U=263285.5<br>**$p=2.506 \times 10^{-11}$**<br>CLES:60.72% | U=314460.0<br>**$p=6.405 \times 10^{-10}$**<br>CLES:59.38% |
| Blue tit | U=171551.0<br>p=0.361<br>CLES:51.56% | U=241694.0<br>**$p=7.06 \times 10^{-11}$**<br>CLES:60.69% | U=287394.0<br>**$p=7.666 \times 10^{-10}$**<br>CLES:59.55% |

| | | | |
|---|---|---|---|
| Pink-footed goose | U=228775.0<br>p=0.264<br>CLES:51.77% | U=318690.0<br>**p=1.952 × 10$^{-13}$**<br>CLES:61.27% | U=379583.0<br>**p=1.058 × 10$^{-11}$**<br>CLES:59.85% |
| Okarito brown kiwi | U=196096.0<br>p=0.55<br>CLES:50.98% | U=281515.0<br>**p=5.389 × 10$^{-11}$**<br>CLES:60.34% | U=331248.5<br>**p=7.806 × 10$^{-11}$**<br>CLES:59.76% |
| Anole lizard | U=155578.0<br>p=0.315<br>CLES:51.76% | U=223746.0<br>**p=6.353 × 10$^{-8}$**<br>CLES:59.00% | U=257062.0<br>**p=2.927 × 10$^{-5}$**<br>CLES:56.60% |
| Chilean tinamou | U=174026.0<br>p=0.672<br>CLES:50.72% | U=248017.0<br>**p=4.116 × 10$^{-9}$**<br>CLES:59.54% | U=298530.5<br>**p=2.971 × 10$^{-9}$**<br>CLES:59.11% |
| Emu | U=206274.5<br>p=0.268<br>CLES:51.80% | U=302691.0<br>**p=3.296 × 10$^{-14}$**<br>CLES:61.81% | U=347031.0<br>**p=1.801 × 10$^{-12}$**<br>CLES:60.49% |
| Australian saltwater crocodile | U=164778.0<br>p=0.148<br>CLES:52.50% | U=249056.5<br>**p=2.782 × 10$^{-16}$**<br>CLES:63.46% | U=274212.5<br>**p=2.149 × 10$^{-12}$**<br>CLES:61.12% |
| Ruff | U=206261.0<br>p=0.262<br>CLES:51.83% | U=303758.5<br>**p=8.213 × 10$^{-15}$**<br>CLES:62.09% | U=349212.5<br>**p=1.267 × 10$^{-12}$**<br>CLES:60.55% |
| Blue-crowned manakin | U=207859.0<br>p=0.335<br>CLES:51.57% | U=311002.0<br>**p=8.261 × 10$^{-18}$**<br>CLES:63.38% | U=365130.0<br>**p=2.726 × 10$^{-16}$**<br>CLES:62.10% |
| Argentine black and white tegu | U=193385.0<br>p=0.411<br>CLES:51.36% | U=289911.0<br>**p=5.897 × 10$^{-11}$**<br>CLES:60.26% | U=337292.0<br>**p=3.318 × 10$^{-9}$**<br>CLES:58.81% |
| Dark-eyed junco | U=179744.0<br>p=0.29<br>CLES:51.78% | U=261481.0<br>**p=7.106 × 10$^{-15}$**<br>CLES:62.62% | U=311163.0<br>**p=2.521 × 10$^{-13}$**<br>CLES:61.22% |
| Golden-collared manakin | U=169366.0<br>p=0.467<br>CLES:51.24% | U=246178.0<br>**p=7.252 × 10$^{-13}$**<br>CLES:61.75% | U=289521.0<br>**p=5.529 × 10$^{-13}$**<br>CLES:61.26% |
| Great spotted kiwi | U=187489.5<br>p=0.554<br>CLES:50.98% | U=267163.5<br>**p=6.273 × 10$^{-12}$**<br>CLES:61.00% | U=318475.0<br>**p=5.491 × 10$^{-12}$**<br>CLES:60.47% |
| Zebra Finch | U=127482.5<br>p=0.697<br>CLES:50.71% | U=201155.5<br>**p=3.162 × 10$^{-13}$**<br>CLES:62.61% | U=229217.0<br>**p=8.426 × 10$^{-14}$**<br>CLES:62.43% |
| Chinese softshell turtle | U=165780.0<br>p=0.587<br>CLES:50.93% | U=252084.5<br>**p=1.091 × 10$^{-14}$**<br>CLES:62.63% | U=288316.0<br>**p=1.572 × 10$^{-14}$**<br>CLES:62.05% |
| Little spotted kiwi | U=195091.5<br>p=0.776<br>CLES:50.47% | U=276076.0<br>**p=1.634 × 10$^{-11}$**<br>CLES:60.66% | U=326908.5<br>**p=2.311 × 10$^{-12}$**<br>CLES:60.59% |
| Helmeted guineafowl | U=227229.5<br>p=0.647<br>CLES:50.72% | U=312992.5<br>**p=3.449 × 10$^{-7}$**<br>CLES:57.74% | U=376500.5<br>**p=6.941 × 10$^{-7}$**<br>CLES:57.13% |

### Chicken Reference Genome, Zeisel et al., 2015

| MWU Test | Glia vs Vasculature | Glia vs Neuron | Vasculature vs Neuron |
|---|---|---|---|
| Spoon-billed sandpiper | U=140326.5<br>$p=5.162 \times 10^{-2}$<br>CLES:53.75% | U=332824.0<br>**$p=2.403 \times 10^{-17}$**<br>CLES:62.91% | U=112779.0<br>**$p=1.145 \times 10^{-6}$**<br>CLES:59.83% |
| White-throated sparrow | U=114887.0<br>$p=1.502 \times 10^{-2}$<br>CLES:54.93% | U=257291.0<br>**$p=2.637 \times 10^{-16}$**<br>CLES:63.34% | U=87628.0<br>**$p=3.793 \times 10^{-5}$**<br>CLES:58.80% |
| Abingdon island giant tortoise | U=115223.0<br>$p=2.066 \times 10^{-2}$<br>CLES:54.70% | U=266313.0<br>**$p=3.175 \times 10^{-14}$**<br>CLES:62.19% | U=88690.5<br>$p=2.988 \times 10^{-4}$<br>CLES:57.69% |
| Japanese quail | U=151830.0<br>$p=1.517 \times 10^{-2}$<br>CLES:54.60% | U=328801.0<br>**$p=1.386 \times 10^{-9}$**<br>CLES:59.12% | U=109215.0<br>$p=1.852 \times 10^{-2}$<br>CLES:54.69% |
| Budgerigar | U=121887.0<br>$p=1.703 \times 10^{-2}$<br>CLES:54.78% | U=271768.5<br>**$p=2.836 \times 10^{-14}$**<br>CLES:62.17% | U=90445.5<br>$p=1.873 \times 10^{-4}$<br>CLES:57.89% |
| Turkey | U=111327.0<br>p=0.982<br>CLES:50.04% | U=251292.0<br>**$p=3.719 \times 10^{-6}$**<br>CLES:57.42% | U=86201.0<br>$p=4.303 \times 10^{-4}$<br>CLES:57.52% |
| Tuatara | U=112627.5<br>$p=7.85 \times 10^{-3}$<br>CLES:55.48% | U=256934.0<br>**$p=1.696 \times 10^{-12}$**<br>CLES:61.42% | U=80893.5<br>$p=3.504 \times 10^{-3}$<br>CLES:56.33% |
| Bengalese finch | U=127017.0<br>p=0.104<br>CLES:53.19% | U=295827.0<br>**$p=8.939 \times 10^{-14}$**<br>CLES:61.63% | U=104717.0<br>**$p=1.001 \times 10^{-5}$**<br>CLES:59.06% |
| Agassiz's desert tortoise | U=123584.0<br>$p=2.866 \times 10^{-2}$<br>CLES:54.37% | U=277613.0<br>**$p=4.584 \times 10^{-12}$**<br>CLES:60.96% | U=90686.0<br>$p=1.022 \times 10^{-3}$<br>CLES:56.92% |
| Common canary | U=121275.5<br>$p=1.41 \times 10^{-2}$<br>CLES:54.92% | U=279469.0<br>**$p=7.204 \times 10^{-14}$**<br>CLES:61.86% | U=93105.0<br>$p=5.814 \times 10^{-4}$<br>CLES:57.22% |
| Flycatcher | U=134168.0<br>$p=2.715 \times 10^{-2}$<br>CLES:54.31% | U=302673.5<br>**$p=1.329 \times 10^{-14}$**<br>CLES:61.99% | U=102325.5<br>**$p=5.32 \times 10^{-5}$**<br>CLES:58.30% |
| Painted turtle | U=132781.0<br>$p=5.35 \times 10^{-2}$<br>CLES:53.78% | U=306200.0<br>**$p=3.092 \times 10^{-13}$**<br>CLES:61.28% | U=101586.0<br>$p=1.864 \times 10^{-4}$<br>CLES:57.68% |
| Great Tit | U=132480.0<br>$p=5.729 \times 10^{-2}$<br>CLES:53.72% | U=325067.5<br>**$p=2.187 \times 10^{-18}$**<br>CLES:63.43% | U=108357.5<br>**$p=1.253 \times 10^{-6}$**<br>CLES:59.92% |
| Duck | U=127700.0<br>$p=1.976 \times 10^{-3}$<br>CLES:56.15% | U=278745.0<br>**$p=2.137 \times 10^{-14}$**<br>CLES:62.14% | U=92665.0<br>$p=2.721 \times 10^{-3}$<br>CLES:56.25% |
| Blue tit | U=115303.5<br>$p=5.472 \times 10^{-2}$<br>CLES:53.89% | U=264378.0<br>**$p=2.384 \times 10^{-13}$**<br>CLES:61.78% | U=88119.0<br>$p=1.079 \times 10^{-4}$<br>CLES:58.26% |

| | | | |
|---|---|---|---|
| Pink-footed goose | U=148649.0<br>p=$1.245 \times 10^{-2}$<br>CLES:54.76% | U=335672.0<br>**p=$3.486 \times 10^{-15}$**<br>CLES:61.93% | U=111506.0<br>p=$2.359 \times 10^{-4}$<br>CLES:57.37% |
| Okarito brown kiwi | U=135375.0<br>p=$1.486 \times 10^{-2}$<br>CLES:54.75% | U=300635.5<br>**p=$2.554 \times 10^{-15}$**<br>CLES:62.35% | U=101349.0<br>**p=$7.43 \times 10^{-5}$**<br>CLES:58.14% |
| Anole lizard | U=101025.0<br>p=0.218<br>CLES:52.57% | U=237769.5<br>**p=$3.256 \times 10^{-11}$**<br>CLES:60.92% | U=80168.0<br>**p=$5.482 \times 10^{-5}$**<br>CLES:58.85% |
| Chilean tinamou | U=119654.5<br>p=$9.372 \times 10^{-3}$<br>CLES:55.23% | U=271580.0<br>**p=$1.662 \times 10^{-16}$**<br>CLES:63.24% | U=91139.5<br>**p=$9.872 \times 10^{-5}$**<br>CLES:58.23% |
| Emu | U=134285.0<br>p=$7.743 \times 10^{-2}$<br>CLES:53.43% | U=314315.5<br>**p=$6.329 \times 10^{-18}$**<br>CLES:63.37% | U=108723.5<br>**p=$1.227 \times 10^{-7}$**<br>CLES:60.81% |
| Australian saltwater crocodile | U=113393.5<br>p=$1.435 \times 10^{-2}$<br>CLES:55.00% | U=262285.0<br>**p=$2.762 \times 10^{-16}$**<br>CLES:63.26% | U=86987.0<br>**p=$7.818 \times 10^{-5}$**<br>CLES:58.47% |
| Ruff | U=141495.0<br>p=$1.369 \times 10^{-2}$<br>CLES:54.76% | U=327331.0<br>**p=$3.441 \times 10^{-18}$**<br>CLES:63.34% | U=108912.0<br>**p=$1.089 \times 10^{-5}$**<br>CLES:58.92% |
| Blue-crowned manakin | U=131863.0<br>p=$2.955 \times 10^{-2}$<br>CLES:54.26% | U=306844.0<br>**p=$4.409 \times 10^{-14}$**<br>CLES:61.67% | U=103397.0<br>p=$1.905 \times 10^{-4}$<br>CLES:57.64% |
| Argentine black and white tegu | U=127917.5<br>p=$8.327 \times 10^{-2}$<br>CLES:53.41% | U=310155.0<br>**p=$1.487 \times 10^{-13}$**<br>CLES:61.38% | U=104509.0<br>**p=$3.616 \times 10^{-5}$**<br>CLES:58.49% |
| Dark-eyed junco | U=119849.5<br>p=$2.664 \times 10^{-2}$<br>CLES:54.42% | U=268724.0<br>**p=$9.578 \times 10^{-15}$**<br>CLES:62.42% | U=95650.5<br>**p=$4.013 \times 10^{-5}$**<br>CLES:58.58% |
| Golden-collared manakin | U=110250.5<br>p=0.109<br>CLES:53.26% | U=254492.0<br>**p=$3.357 \times 10^{-13}$**<br>CLES:61.81% | U=89256.5<br>**p=$2.937 \times 10^{-5}$**<br>CLES:58.91% |
| Great spotted kiwi | U=123382.0<br>p=$4.034 \times 10^{-2}$<br>CLES:54.09% | U=286575.0<br>**p=$4.81 \times 10^{-15}$**<br>CLES:62.38% | U=94167.5<br>**p=$6.664 \times 10^{-5}$**<br>CLES:58.38% |
| Zebra Finch | U=80609.0<br>p=$2.776 \times 10^{-2}$<br>CLES:54.90% | U=200302.5<br>**p=$2.96 \times 10^{-13}$**<br>CLES:62.58% | U=65693.0<br>p=$5.595 \times 10^{-4}$<br>CLES:57.97% |
| Chinese softshell turtle | U=112801.5<br>p=$2.869 \times 10^{-2}$<br>CLES:54.46% | U=260717.5<br>**p=$2.788 \times 10^{-16}$**<br>CLES:63.28% | U=88598.5<br>**p=$1.291 \times 10^{-5}$**<br>CLES:59.33% |
| Little spotted kiwi | U=131102.0<br>p=$5.825 \times 10^{-2}$<br>CLES:53.72% | U=294017.0<br>**p=$2.305 \times 10^{-15}$**<br>CLES:62.47% | U=98146.0<br>**p=$7.978 \times 10^{-6}$**<br>CLES:59.29% |
| Helmeted guineafowl | U=149326.5<br>p=$3.732 \times 10^{-2}$<br>CLES:53.96% | U=333221.5<br>**p=$7.167 \times 10^{-10}$**<br>CLES:59.25% | U=109171.0<br>p=$9.359 \times 10^{-3}$<br>CLES:55.19% |

### Chicken Reference Genome, Zeisel et al., 2018

| MWU Test | CNS_Glia vs CNS_Endothelia | CNS_Glia vs CNS_Neuron | CNS_Endothelia vs CNS_Neuron |
|---|---|---|---|
| Spoon-billed sandpiper | U=32392.0<br>p=0.191<br>CLES:54.04% | U=949935.0<br>**$p=1.974 \times 10^{-23}$**<br>CLES:63.21% | U=159015.5<br>$p=7.372 \times 10^{-4}$<br>CLES:59.79% |
| White-throated sparrow | U=24012.0<br>p=0.24<br>CLES:53.93% | U=756955.0<br>**$p=3.473 \times 10^{-24}$**<br>CLES:64.37% | U=124526.0<br>$p=5.153 \times 10^{-4}$<br>CLES:60.89% |
| Abingdon island giant tortoise | U=26433.0<br>p=0.483<br>CLES:52.25% | U=765022.0<br>**$p=1.087 \times 10^{-14}$**<br>CLES:60.74% | U=137484.0<br>$p=9.631 \times 10^{-4}$<br>CLES:59.92% |
| Japanese quail | U=36209.5<br>$p=9.907 \times 10^{-2}$<br>CLES:55.01% | U=1028797.0<br>**$p=8.123 \times 10^{-22}$**<br>CLES:62.35% | U=162336.0<br>$p=7.555 \times 10^{-3}$<br>CLES:57.64% |
| Budgerigar | U=30646.0<br>$p=6.328 \times 10^{-3}$<br>CLES:58.67% | U=837977.0<br>**$p=2.089 \times 10^{-24}$**<br>CLES:64.07% | U=137053.0<br>$p=1.879 \times 10^{-2}$<br>CLES:56.99% |
| Turkey | U=27583.5<br>p=0.124<br>CLES:54.94% | U=761685.0<br>**$p=6.887 \times 10^{-19}$**<br>CLES:62.40% | U=130167.5<br>$p=7.104 \times 10^{-3}$<br>CLES:58.09% |
| Tuatara | U=23768.0<br>p=0.686<br>CLES:51.35% | U=756475.5<br>**$p=1.229 \times 10^{-17}$**<br>CLES:61.88% | U=123691.5<br>$p=1.676 \times 10^{-4}$<br>CLES:61.87% |
| Bengalese finch | U=30327.0<br>$p=6.608 \times 10^{-2}$<br>CLES:55.80% | U=898737.5<br>**$p=2.552 \times 10^{-25}$**<br>CLES:64.12% | U=149428.0<br>$p=2.042 \times 10^{-3}$<br>CLES:59.12% |
| Agassiz's desert tortoise | U=29187.0<br>p=0.434<br>CLES:52.45% | U=823482.5<br>**$p=3.886 \times 10^{-18}$**<br>CLES:61.83% | U=146473.0<br>$p=6.485 \times 10^{-4}$<br>CLES:60.00% |
| Common canary | U=27191.0<br>$p=4.882 \times 10^{-2}$<br>CLES:56.39% | U=809434.5<br>**$p=2.419 \times 10^{-25}$**<br>CLES:64.59% | U=136907.0<br>$p=1.529 \times 10^{-3}$<br>CLES:59.62% |
| Flycatcher | U=31502.5<br>$p=2.485 \times 10^{-2}$<br>CLES:57.02% | U=925128.0<br>**$p=1.819 \times 10^{-34}$**<br>CLES:66.66% | U=156171.0<br>$p=1.863 \times 10^{-4}$<br>CLES:60.95% |
| Painted turtle | U=32926.0<br>p=0.136<br>CLES:54.59% | U=914615.0<br>**$p=5.77 \times 10^{-19}$**<br>CLES:61.82% | U=154103.0<br>$p=5.122 \times 10^{-3}$<br>CLES:58.09% |
| Great Tit | U=30764.0<br>p=0.119<br>CLES:54.90% | U=921721.5<br>**$p=1.819 \times 10^{-26}$**<br>CLES:64.29% | U=150347.0<br>$p=7.645 \times 10^{-4}$<br>CLES:59.95% |
| Duck | U=28144.0<br>$p=6.256 \times 10^{-2}$<br>CLES:56.04% | U=818755.5<br>**$p=4.62 \times 10^{-22}$**<br>CLES:63.28% | U=127205.0<br>$p=2.022 \times 10^{-2}$<br>CLES:57.09% |
| Blue tit | U=24443.0<br>p=0.196 | U=797587.0<br>**$p=1.029 \times 10^{-23}$** | U=124179.0<br>$p=8.3 \times 10^{-4}$ |

| | | | |
|---|---|---|---|
| | CLES:54.34% | CLES:64.00% | CLES:60.59% |
| Pink-footed goose | U=34750.0<br>p=0.106<br>CLES:54.96% | U=997434.0<br>**p=4.302 × $10^{-24}$**<br>CLES:63.17% | U=157931.5<br>p=3.337 × $10^{-3}$<br>CLES:58.47% |
| Okarito brown kiwi | U=30432.0<br>p=0.307<br>CLES:53.20% | U=899467.0<br>**p=3.204 × $10^{-21}$**<br>CLES:62.65% | U=151797.5<br>p=3.731 × $10^{-4}$<br>CLES:60.48% |
| Anole lizard | U=22752.0<br>p=0.493<br>CLES:52.30% | U=685608.5<br>**p=1.145 × $10^{-11}$**<br>CLES:59.67% | U=115602.0<br>p=1.312 × $10^{-2}$<br>CLES:57.82% |
| Chilean tinamou | U=25186.0<br>p=0.26<br>CLES:53.71% | U=750085.0<br>**p=1.162 × $10^{-17}$**<br>CLES:61.98% | U=124010.0<br>p=2.674 × $10^{-3}$<br>CLES:59.32% |
| Emu | U=32687.0<br>p=0.197<br>CLES:53.95% | U=933472.0<br>**p=5.527 × $10^{-24}$**<br>CLES:63.44% | U=160184.0<br>p=6.32 × $10^{-4}$<br>CLES:59.83% |
| Australian saltwater crocodile | U=25668.0<br>p=0.495<br>CLES:52.21% | U=783977.5<br>**p=5.783 × $10^{-25}$**<br>CLES:64.42% | U=138929.0<br>**p=8.637 × $10^{-6}$**<br>CLES:63.51% |
| Ruff | U=31127.0<br>p=0.18<br>CLES:54.21% | U=947639.5<br>**p=1.018 × $10^{-25}$**<br>CLES:63.92% | U=154399.0<br>p=1.954 × $10^{-4}$<br>CLES:61.02% |
| Blue-crowned manakin | U=30729.0<br>p=0.118<br>CLES:54.88% | U=896318.0<br>**p=1.188 × $10^{-25}$**<br>CLES:64.23% | U=155443.0<br>p=6.428 × $10^{-4}$<br>CLES:59.95% |
| Argentine black and white tegu | U=29915.0<br>p=0.523<br>CLES:52.01% | U=931345.0<br>**p=4.897 × $10^{-18}$**<br>CLES:61.46% | U=154009.0<br>p=1.1 × $10^{-3}$<br>CLES:59.65% |
| Dark-eyed junco | U=27283.0<br>p=5.977 × $10^{-2}$<br>CLES:56.05% | U=773873.0<br>**p=3.549 × $10^{-25}$**<br>CLES:64.65% | U=134810.0<br>p=1.618 × $10^{-3}$<br>CLES:59.48% |
| Golden-collared manakin | U=24736.0<br>p=9.41 × $10^{-2}$<br>CLES:55.63% | U=784321.5<br>**p=4.082 × $10^{-27}$**<br>CLES:65.14% | U=121168.0<br>p=9.324 × $10^{-4}$<br>CLES:60.50% |
| Great spotted kiwi | U=28759.0<br>p=0.206<br>CLES:54.06% | U=867127.0<br>**p=1.362 × $10^{-22}$**<br>CLES:63.23% | U=139209.5<br>p=1.124 × $10^{-3}$<br>CLES:59.84% |
| Zebra Finch | U=18217.0<br>p=0.218<br>CLES:54.32% | U=568399.5<br>**p=4.834 × $10^{-23}$**<br>CLES:65.35% | U=106515.0<br>p=1.179 × $10^{-4}$<br>CLES:62.59% |
| Chinese softshell turtle | U=24464.0<br>p=0.343<br>CLES:53.15% | U=781790.0<br>**p=9.835 × $10^{-22}$**<br>CLES:63.37% | U=126417.0<br>p=5.089 × $10^{-4}$<br>CLES:60.90% |
| Little spotted kiwi | U=31146.0<br>p=8.393 × $10^{-2}$<br>CLES:55.42% | U=875456.0<br>**p=5.112 × $10^{-21}$**<br>CLES:62.69% | U=143671.5<br>p=7.956 × $10^{-3}$<br>CLES:57.82% |
| Helmeted guineafowl | U=37643.0<br>p=1.522 × $10^{-2}$ | U=1031562.0<br>**p=1.886 × $10^{-22}$** | U=156120.0<br>p=6.448 × $10^{-2}$ |

| | | |
|---|---|---|
| CLES:57.37% | CLES:62.56% | CLES:55.29% |

### Chicken Reference Genome, Saunders et al., 2018

| MWU Test | Vasculature vs Glia | Vasculature vs Neuron | Glia vs Neuron |
|---|---|---|---|
| Spoon-billed sandpiper | U=31760.0<br>p=0.304<br>CLES:52.69% | U=84829.0<br>**$p=1.36\times10^{-7}$**<br>CLES:61.43% | U=66770.0<br>**$p=1.201\times10^{-5}$**<br>CLES:60.22% |
| White-throated sparrow | U=25146.0<br>p=0.199<br>CLES:53.59% | U=69762.5<br>**$p=1.252\times10^{-5}$**<br>CLES:59.86% | U=47761.0<br>$p=7.035\times10^{-3}$<br>CLES:56.80% |
| Abingdon island giant tortoise | U=27686.0<br>$p=9.72\times10^{-2}$<br>CLES:54.54% | U=74887.0<br>**$p=1.359\times10^{-7}$**<br>CLES:61.77% | U=51295.0<br>$p=2.006\times10^{-3}$<br>CLES:57.65% |
| Japanese quail | U=36061.0<br>p=0.235<br>CLES:53.02% | U=88724.0<br>**$p=2.995\times10^{-5}$**<br>CLES:58.81% | U=65261.0<br>$p=7.988\times10^{-3}$<br>CLES:56.10% |
| Budgerigar | U=28646.0<br>$p=8.132\times10^{-2}$<br>CLES:54.74% | U=79226.0<br>**$p=8.505\times10^{-9}$**<br>CLES:62.73% | U=54908.0<br>$p=1.702\times10^{-4}$<br>CLES:59.23% |
| Turkey | U=24585.0<br>p=0.976<br>CLES:50.08% | U=63877.0<br>$p=2.955\times10^{-3}$<br>CLES:56.75% | U=47294.0<br>$p=9.105\times10^{-3}$<br>CLES:56.53% |
| Tuatara | U=26425.0<br>p=0.507<br>CLES:51.82% | U=69574.0<br>$p=1.639\times10^{-4}$<br>CLES:58.43% | U=49281.0<br>$p=1.676\times10^{-3}$<br>CLES:57.85% |
| Bengalese finch | U=34541.0<br>$p=1.796\times10^{-2}$<br>CLES:56.17% | U=90788.5<br>**$p=6.222\times10^{-10}$**<br>CLES:63.24% | U=63009.0<br>$p=3.727\times10^{-4}$<br>CLES:58.38% |
| Agassiz's desert tortoise | U=28331.0<br>p=0.122<br>CLES:54.20% | U=77232.0<br>**$p=1.845\times10^{-8}$**<br>CLES:62.51% | U=55009.0<br>$p=1.318\times10^{-4}$<br>CLES:59.36% |
| Common canary | U=27191.5<br>p=0.388<br>CLES:52.35% | U=79478.5<br>**$p=5.931\times10^{-8}$**<br>CLES:61.97% | U=57167.5<br>**$p=2.72\times10^{-5}$**<br>CLES:60.26% |
| Flycatcher | U=31592.0<br>$p=3.018\times10^{-2}$<br>CLES:55.78% | U=87838.0<br>**$p=6.569\times10^{-10}$**<br>CLES:63.35% | U=58566.0<br>$p=8.235\times10^{-4}$<br>CLES:58.05% |
| Painted turtle | U=30772.0<br>p=0.136<br>CLES:53.96% | U=84377.0<br>**$p=7.719\times10^{-9}$**<br>CLES:62.61% | U=63616.0<br>**$p=2.908\times10^{-5}$**<br>CLES:59.90% |
| Great Tit | U=32291.0<br>p=0.205<br>CLES:53.32% | U=88059.0<br>**$p=3.096\times10^{-8}$**<br>CLES:61.86% | U=62816.0<br>**$p=2.438\times10^{-5}$**<br>CLES:60.02% |
| Duck | U=28174.0<br>p=0.116<br>CLES:54.27% | U=72333.0<br>**$p=8.897\times10^{-7}$**<br>CLES:61.05% | U=53562.0<br>$p=2.042\times10^{-3}$<br>CLES:57.52% |
| Blue tit | U=26631.5<br>$p=5.133\times10^{-2}$<br>CLES:55.41% | U=73200.0<br>**$p=2.972\times10^{-7}$**<br>CLES:61.52% | U=49265.0<br>$p=5.843\times10^{-3}$<br>CLES:56.90% |

| | | | |
|---|---|---|---|
| Pink-footed goose | U=34098.0<br>p=0.369<br>CLES:52.31% | U=86125.0<br>**p=2.025 × 10$^{-5}$**<br>CLES:59.07% | U=64684.0<br>p=1.003 × 10$^{-3}$<br>CLES:57.64% |
| Okarito brown kiwi | U=31919.0<br>p=6.707 × 10$^{-2}$<br>CLES:54.84% | U=84014.0<br>**p=3.999 × 10$^{-10}$**<br>CLES:63.69% | U=62906.0<br>**p=7.649 × 10$^{-6}$**<br>CLES:60.61% |
| Anole lizard | U=26728.0<br>p=1.584 × 10$^{-2}$<br>CLES:56.72% | U=68516.0<br>**p=1.791 × 10$^{-7}$**<br>CLES:61.94% | U=46719.0<br>p=4.034 × 10$^{-2}$<br>CLES:55.13% |
| Chilean tinamou | U=27339.0<br>p=0.146<br>CLES:53.97% | U=70426.0<br>**p=1.536 × 10$^{-6}$**<br>CLES:60.89% | U=53284.0<br>p=1.506 × 10$^{-3}$<br>CLES:57.75% |
| Emu | U=32148.5<br>p=3.442 × 10$^{-2}$<br>CLES:55.61% | U=89189.0<br>**p=1.144 × 10$^{-10}$**<br>CLES:63.89% | U=61009.0<br>**p=2.459 × 10$^{-5}$**<br>CLES:60.12% |
| Australian saltwater crocodile | U=26416.0<br>p=1.61 × 10$^{-2}$<br>CLES:56.73% | U=74441.0<br>**p=1.382 × 10$^{-10}$**<br>CLES:64.57% | U=50810.0<br>p=5.217 × 10$^{-4}$<br>CLES:58.68% |
| Ruff | U=30818.5<br>p=0.222<br>CLES:53.23% | U=86463.0<br>**p=2.398 × 10$^{-9}$**<br>CLES:62.93% | U=62689.0<br>**p=1.102 × 10$^{-5}$**<br>CLES:60.47% |
| Blue-crowned manakin | U=32698.0<br>p=4.597 × 10$^{-2}$<br>CLES:55.25% | U=87102.0<br>**p=2.79 × 10$^{-10}$**<br>CLES:63.71% | U=64041.0<br>**p=6.285 × 10$^{-5}$**<br>CLES:59.42% |
| Argentine black and white tegu | U=30192.0<br>p=0.244<br>CLES:53.10% | U=81900.0<br>**p=4.066 × 10$^{-5}$**<br>CLES:58.88% | U=58798.0<br>p=5.501 × 10$^{-3}$<br>CLES:56.63% |
| Dark-eyed junco | U=27538.0<br>p=0.453<br>CLES:52.04% | U=76442.0<br>**p=2.526 × 10$^{-7}$**<br>CLES:61.41% | U=54373.0<br>**p=3.616 × 10$^{-5}$**<br>CLES:60.18% |
| Golden-collared manakin | U=26406.0<br>p=0.133<br>CLES:54.15% | U=76787.0<br>**p=1.165 × 10$^{-11}$**<br>CLES:65.27% | U=54102.0<br>**p=3.2 × 10$^{-7}$**<br>CLES:62.77% |
| Great spotted kiwi | U=29438.5<br>p=0.296<br>CLES:52.80% | U=79813.0<br>**p=8.296 × 10$^{-8}$**<br>CLES:61.77% | U=58548.5<br>**p=2.733 × 10$^{-5}$**<br>CLES:60.13% |
| Zebra Finch | U=20967.0<br>p=0.146<br>CLES:54.26% | U=59758.0<br>**p=7.891 × 10$^{-8}$**<br>CLES:62.76% | U=41398.0<br>p=3.156 × 10$^{-4}$<br>CLES:59.52% |
| Chinese softshell turtle | U=27160.0<br>p=2.314 × 10$^{-2}$<br>CLES:56.29% | U=77656.0<br>**p=1.117 × 10$^{-12}$**<br>CLES:66.09% | U=55439.0<br>**p=6.77 × 10$^{-6}$**<br>CLES:61.12% |
| Little spotted kiwi | U=30975.0<br>p=0.118<br>CLES:54.15% | U=82767.0<br>**p=6.094 × 10$^{-9}$**<br>CLES:62.70% | U=59641.0<br>**p=4.909 × 10$^{-5}$**<br>CLES:59.74% |
| Helmeted guineafowl | U=35019.0<br>p=0.144<br>CLES:53.75% | U=87909.0<br>**p=1.073 × 10$^{-5}$**<br>CLES:59.36% | U=65860.0<br>p=6.787 × 10$^{-3}$<br>CLES:56.23% |

### Mouse Reference Genome, Zhang et al., 2014

| MWU Test | Endothelia vs Glia | Endothelia vs Neuron | Glia vs Neuron |
|---|---|---|---|
| Microbat | U=268227.0<br>p=0.583<br>CLES:49.17% | U=386130.0<br>**p=8.646 × 10$^{-20}$**<br>CLES:63.40% | U=487527.0<br>**p=5.016 × 10$^{-25}$**<br>CLES:64.31% |
| Hedgehog | U=20862.0<br>p=0.98<br>CLES:50.07% | U=24745.0<br>p=1.326 × 10$^{-4}$<br>CLES:61.03% | U=29768.5<br>**p=6.716 × 10$^{-5}$**<br>CLES:60.96% |
| Naked mole-rat male | U=317383.5<br>p=0.902<br>CLES:50.18% | U=462666.5<br>**p=8.944 × 10$^{-32}$**<br>CLES:66.75% | U=619676.0<br>**p=1.703 × 10$^{-38}$**<br>CLES:67.10% |
| Sooty mangabey | U=405106.0<br>p=0.811<br>CLES:50.33% | U=598994.5<br>**p=2.942 × 10$^{-35}$**<br>CLES:66.58% | U=783150.5<br>**p=1.665 × 10$^{-39}$**<br>CLES:66.30% |
| Olive baboon | U=405323.0<br>p=0.745<br>CLES:49.56% | U=595418.0<br>**p=9.967 × 10$^{-33}$**<br>CLES:65.91% | U=787019.0<br>**p=3.892 × 10$^{-41}$**<br>CLES:66.65% |
| American bison | U=259433.0<br>p=0.767<br>CLES:49.55% | U=360671.5<br>**p=2.358 × 10$^{-24}$**<br>CLES:65.33% | U=455738.0<br>**p=3.123 × 10$^{-30}$**<br>CLES:66.20% |
| Leopard | U=366639.5<br>p=0.936<br>CLES:49.89% | U=550137.0<br>**p=6.339 × 10$^{-31}$**<br>CLES:65.75% | U=704001.0<br>**p=8.625 × 10$^{-37}$**<br>CLES:66.12% |
| Bolivian squirrel monkey | U=369438.0<br>p=0.588<br>CLES:50.76% | U=560933.0<br>**p=2.697 × 10$^{-38}$**<br>CLES:67.66% | U=710342.0<br>**p=4.282 × 10$^{-41}$**<br>CLES:67.11% |
| Crab-eating macaque | U=404026.0<br>p=0.953<br>CLES:50.08% | U=589181.0<br>**p=1.429 × 10$^{-32}$**<br>CLES:65.92% | U=777741.0<br>**p=2.263 × 10$^{-39}$**<br>CLES:66.30% |
| Human | U=425381.0<br>p=0.94<br>CLES:49.90% | U=634896.0<br>**p=1.743 × 10$^{-33}$**<br>CLES:65.85% | U=820011.5<br>**p=3.24 × 10$^{-40}$**<br>CLES:66.27% |
| Red fox | U=336016.0<br>p=0.771<br>CLES:49.58% | U=488131.0<br>**p=5.665 × 10$^{-25}$**<br>CLES:64.38% | U=609752.0<br>**p=1.543 × 10$^{-29}$**<br>CLES:64.81% |
| Shrew mouse | U=496797.0<br>p=0.801<br>CLES:49.67% | U=701001.0<br>**p=2.856 × 10$^{-29}$**<br>CLES:64.30% | U=919672.0<br>**p=6.23 × 10$^{-35}$**<br>CLES:64.58% |
| Arctic ground squirrel | U=402787.5<br>p=0.833<br>CLES:50.29% | U=596271.5<br>**p=2.751 × 10$^{-39}$**<br>CLES:67.61% | U=773267.0<br>**p=3.18 × 10$^{-45}$**<br>CLES:67.63% |
| Chinese hamster CriGri | U=389859.0<br>p=0.314<br>CLES:48.62% | U=558032.5<br>**p=9.332 × 10$^{-29}$**<br>CLES:65.04% | U=746400.5<br>**p=2.153 × 10$^{-37}$**<br>CLES:66.01% |
| Opossum | U=315436.0<br>p=0.228<br>CLES:51.77% | U=424670.0<br>**p=1.153 × 10$^{-14}$**<br>CLES:61.01% | U=521585.5<br>**p=4.673 × 10$^{-12}$**<br>CLES:59.23% |

| | | | |
|---|---|---|---|
| Dolphin | U=114401.0<br>p=7.044 × $10^{-2}$<br>CLES:46.67% | U=155841.0<br>**p=2.712 × $10^{-11}$**<br>CLES:62.18% | U=191654.0<br>**p=3.534 × $10^{-19}$**<br>CLES:65.73% |
| Capuchin | U=390021.5<br>p=0.668<br>CLES:50.59% | U=583318.0<br>**p=2.891 × $10^{-40}$**<br>CLES:67.99% | U=765894.0<br>**p=8.144 × $10^{-45}$**<br>CLES:67.59% |
| Mouse Lemur | U=376768.0<br>p=0.603<br>CLES:50.73% | U=555318.5<br>**p=1.131 × $10^{-35}$**<br>CLES:67.01% | U=707826.0<br>**p=7.268 × $10^{-39}$**<br>CLES:66.60% |
| Damara mole rat | U=317871.0<br>p=0.949<br>CLES:49.91% | U=480094.5<br>**p=9.43 × $10^{-34}$**<br>CLES:67.15% | U=623253.0<br>**p=8.603 × $10^{-41}$**<br>CLES:67.64% |
| Marmoset | U=392146.0<br>p=0.439<br>CLES:51.07% | U=576031.0<br>**p=8.551 × $10^{-36}$**<br>CLES:66.90% | U=746167.0<br>**p=8.104 × $10^{-37}$**<br>CLES:65.88% |
| Goat | U=359086.5<br>p=0.97<br>CLES:50.05% | U=546964.5<br>**p=2.368 × $10^{-32}$**<br>CLES:66.18% | U=693777.0<br>**p=1.026 × $10^{-36}$**<br>CLES:66.17% |
| Common wombat | U=291160.5<br>p=0.602<br>CLES:50.78% | U=409402.0<br>**p=1.403 × $10^{-19}$**<br>CLES:63.12% | U=505679.0<br>**p=1.483 × $10^{-19}$**<br>CLES:62.32% |
| American black bear | U=305271.0<br>p=0.47<br>CLES:51.06% | U=396971.0<br>**p=1.025 × $10^{-24}$**<br>CLES:65.06% | U=498026.0<br>**p=5.489 × $10^{-24}$**<br>CLES:63.88% |
| Sloth | U=7044.0<br>p=0.538<br>CLES:47.70% | U=8634.0<br>**p=1.055 × $10^{-5}$**<br>CLES:66.95% | U=12009.0<br>**p=3.061 × $10^{-7}$**<br>CLES:68.20% |
| Polar bear | U=289347.0<br>p=0.784<br>CLES:49.59% | U=391873.0<br>**p=3.209 × $10^{-21}$**<br>CLES:63.85% | U=486917.0<br>**p=3.153 × $10^{-25}$**<br>CLES:64.37% |
| Alpine marmot | U=358073.5<br>p=0.55<br>CLES:49.16% | U=515433.0<br>**p=2.861 × $10^{-30}$**<br>CLES:65.79% | U=672056.5<br>**p=1.91 × $10^{-39}$**<br>CLES:66.96% |
| Prairie vole | U=430333.5<br>p=0.313<br>CLES:48.65% | U=641588.0<br>**p=2.025 × $10^{-33}$**<br>CLES:65.80% | U=866799.0<br>**p=2.492 × $10^{-44}$**<br>CLES:66.91% |
| Donkey | U=373077.0<br>p=0.785<br>CLES:49.62% | U=548747.5<br>**p=5.024 × $10^{-27}$**<br>CLES:64.61% | U=717655.5<br>**p=5.664 × $10^{-33}$**<br>CLES:65.09% |
| Golden Hamster | U=390004.5<br>p=0.344<br>CLES:48.70% | U=584198.0<br>**p=5.466 × $10^{-33}$**<br>CLES:66.07% | U=784853.5<br>**p=7.398 × $10^{-44}$**<br>CLES:67.27% |
| Armadillo | U=314130.5<br>p=0.997<br>CLES:49.99% | U=440700.5<br>**p=2.305 × $10^{-25}$**<br>CLES:64.92% | U=580474.0<br>**p=3.899 × $10^{-30}$**<br>CLES:65.17% |
| Ugandan red Colobus | U=388782.0<br>p=0.957<br>CLES:50.08% | U=577166.0<br>**p=7.469 × $10^{-34}$**<br>CLES:66.36% | U=756027.5<br>**p=6.128 × $10^{-40}$**<br>CLES:66.56% |

| | | | |
|---|---|---|---|
| Pig-tailed macaque | U=408995.0<br>p=0.777<br>CLES:50.39% | U=606237.0<br>**p=2.921 × $10^{-34}$**<br>CLES:66.25% | U=776705.0<br>**p=1.952 × $10^{-38}$**<br>CLES:66.09% |
| Panda | U=370549.0<br>p=0.802<br>CLES:49.65% | U=542427.5<br>**p=2.804 × $10^{-31}$**<br>CLES:65.87% | U=690671.0<br>**p=2.649 × $10^{-39}$**<br>CLES:66.81% |
| Ferret | U=344909.0<br>p=0.234<br>CLES:48.32% | U=519833.0<br>**p=8.232 × $10^{-28}$**<br>CLES:65.05% | U=678948.5<br>**p=8.208 × $10^{-39}$**<br>CLES:66.77% |
| Koala | U=332243.0<br>p=0.772<br>CLES:50.42% | U=473061.0<br>**p=1.157 × $10^{-18}$**<br>CLES:62.31% | U=597267.0<br>**p=2.541 × $10^{-20}$**<br>CLES:62.06% |
| Wallaby | U=29802.0<br>p=0.675<br>CLES:48.90% | U=35456.0<br>p=2.932 × $10^{-3}$<br>CLES:57.78% | U=51594.0<br>**p=4.772 × $10^{-5}$**<br>CLES:59.69% |
| Tarsier | U=257686.0<br>p=0.755<br>CLES:49.52% | U=364955.5<br>**p=3.398 × $10^{-24}$**<br>CLES:65.25% | U=470245.0<br>**p=1.333 × $10^{-30}$**<br>CLES:66.18% |
| Degu | U=294274.0<br>p=0.496<br>CLES:51.01% | U=470486.5<br>**p=1.197 × $10^{-36}$**<br>CLES:68.15% | U=614722.0<br>**p=8.354 × $10^{-39}$**<br>CLES:67.24% |
| Chinese hamster CHOK1GS | U=483795.0<br>p=0.825<br>CLES:49.71% | U=703306.5<br>**p=1.284 × $10^{-35}$**<br>CLES:65.96% | U=922571.0<br>**p=1.305 × $10^{-41}$**<br>CLES:66.06% |
| Drill | U=368099.0<br>p=0.682<br>CLES:49.43% | U=531031.5<br>**p=4.013 × $10^{-29}$**<br>CLES:65.36% | U=707188.0<br>**p=1.642 × $10^{-37}$**<br>CLES:66.27% |
| Gibbon | U=365055.0<br>p=0.552<br>CLES:49.17% | U=537148.0<br>**p=2.434 × $10^{-29}$**<br>CLES:65.38% | U=712475.5<br>**p=1.781 × $10^{-37}$**<br>CLES:66.23% |
| Rat | U=455325.0<br>p=0.366<br>CLES:48.81% | U=647343.0<br>**p=2.264 × $10^{-27}$**<br>CLES:64.07% | U=868212.5<br>**p=4.692 × $10^{-37}$**<br>CLES:65.30% |
| Macaque | U=428423.0<br>p=0.923<br>CLES:49.87% | U=621027.0<br>**p=9.088 × $10^{-31}$**<br>CLES:65.20% | U=818091.0<br>**p=1.159 × $10^{-38}$**<br>CLES:65.92% |
| American mink | U=338310.0<br>p=0.4<br>CLES:48.81% | U=497245.0<br>**p=9.129 × $10^{-26}$**<br>CLES:64.57% | U=641647.5<br>**p=2.165 × $10^{-33}$**<br>CLES:65.65% |
| Gelada | U=410593.5<br>p=0.993<br>CLES:50.01% | U=611952.0<br>**p=9.805 × $10^{-33}$**<br>CLES:65.82% | U=800519.5<br>**p=4.412 × $10^{-39}$**<br>CLES:66.11% |
| Sheep | U=337567.5<br>p=0.965<br>CLES:49.94% | U=491409.0<br>**p=4.146 × $10^{-25}$**<br>CLES:64.42% | U=627612.5<br>**p=4.108 × $10^{-30}$**<br>CLES:64.86% |
| Tasmanian devil | U=280667.0<br>p=0.89<br>CLES:50.21% | U=374758.5<br>**p=4.582 × $10^{-17}$**<br>CLES:62.40% | U=479688.0<br>**p=7.027 × $10^{-19}$**<br>CLES:62.23% |

| | | | |
|---|---|---|---|
| Elephant | U=352705.5<br>p=0.557<br>CLES:49.17% | U=524276.0<br>**p=1.499 × 10$^{-26}$**<br>CLES:64.62% | U=661570.0<br>**p=3.016 × 10$^{-33}$**<br>CLES:65.49% |
| Algerian mouse | U=500618.5<br>p=0.758<br>CLES:50.40% | U=674722.0<br>**p=4.93 × 10$^{-24}$**<br>CLES:62.80% | U=894698.5<br>**p=2.176 × 10$^{-26}$**<br>CLES:62.44% |
| Dingo | U=364238.0<br>p=0.913<br>CLES:49.85% | U=531435.0<br>**p=4.225 × 10$^{-29}$**<br>CLES:65.34% | U=675751.0<br>**p=5.521 × 10$^{-33}$**<br>CLES:65.33% |
| Golden snub-nosed monkey | U=382334.5<br>p=0.643<br>CLES:49.36% | U=554269.5<br>**p=1.447 × 10$^{-32}$**<br>CLES:66.13% | U=708280.0<br>**p=7.035 × 10$^{-41}$**<br>CLES:67.06% |
| Horse | U=386845.5<br>p=0.774<br>CLES:50.40% | U=582662.0<br>**p=8.238 × 10$^{-32}$**<br>CLES:65.77% | U=744422.0<br>**p=7.541 × 10$^{-35}$**<br>CLES:65.42% |
| Black snub-nosed monkey | U=376040.5<br>p=0.945<br>CLES:49.90% | U=535428.0<br>**p=1.98 × 10$^{-27}$**<br>CLES:64.81% | U=694421.0<br>**p=3.19 × 10$^{-33}$**<br>CLES:65.28% |
| Brazilian guinea pig | U=193676.0<br>p=0.963<br>CLES:50.08% | U=269969.0<br>**p=4.472 × 10$^{-20}$**<br>CLES:64.89% | U=368619.0<br>**p=1.978 × 10$^{-23}$**<br>CLES:64.84% |
| Orangutan | U=378067.5<br>p=0.961<br>CLES:49.93% | U=557746.5<br>**p=9.307 × 10$^{-32}$**<br>CLES:65.90% | U=708655.0<br>**p=3.679 × 10$^{-38}$**<br>CLES:66.42% |
| Northern American deer mouse | U=454214.0<br>p=0.427<br>CLES:48.95% | U=636392.0<br>**p=8.841 × 10$^{-33}$**<br>CLES:65.62% | U=851634.5<br>**p=1.397 × 10$^{-42}$**<br>CLES:66.61% |
| Naked mole-rat female | U=367729.0<br>p=0.507<br>CLES:50.93% | U=543998.5<br>**p=6.853 × 10$^{-37}$**<br>CLES:67.45% | U=706838.5<br>**p=7.907 × 10$^{-40}$**<br>CLES:66.83% |
| Daurian ground squirrel | U=281856.0<br>p=0.855<br>CLES:49.73% | U=393420.5<br>**p=1.177 × 10$^{-27}$**<br>CLES:66.09% | U=490253.0<br>**p=2.37 × 10$^{-33}$**<br>CLES:66.79% |
| Tiger | U=244430.0<br>p=0.277<br>CLES:48.33% | U=342879.0<br>**p=1.572 × 10$^{-21}$**<br>CLES:64.48% | U=446058.5<br>**p=8.751 × 10$^{-31}$**<br>CLES:66.46% |
| Lesser hedgehog tenrec | U=26729.0<br>p=0.177<br>CLES:53.70% | U=28531.0<br>**p=5.675 × 10$^{-8}$**<br>CLES:65.35% | U=32448.0<br>**p=7.265 × 10$^{-6}$**<br>CLES:62.14% |
| Mongolian gerbil | U=379537.0<br>p=0.344<br>CLES:48.69% | U=512806.0<br>**p=4.634 × 10$^{-26}$**<br>CLES:64.51% | U=680383.0<br>**p=6.237 × 10$^{-35}$**<br>CLES:65.79% |
| Ma's night monkey | U=389689.0<br>p=0.245<br>CLES:51.62% | U=575314.0<br>**p=7.776 × 10$^{-40}$**<br>CLES:67.95% | U=737593.0<br>**p=2.283 × 10$^{-38}$**<br>CLES:66.30% |
| Upper Galilee mountains blind mole rat | U=401260.5<br>p=0.986<br>CLES:49.98% | U=600029.0<br>**p=1.043 × 10$^{-31}$**<br>CLES:65.64% | U=796606.5<br>**p=3.522 × 10$^{-37}$**<br>CLES:65.69% |

| | | | |
|---|---|---|---|
| Kangaroo rat | U=298457.0<br>p=0.964<br>CLES:49.93% | U=453020.0<br>**p=3.519 × 10$^{-27}$**<br>CLES:65.44% | U=586578.5<br>**p=1.305 × 10$^{-31}$**<br>CLES:65.55% |
| Chimpanzee | U=379475.5<br>p=0.848<br>CLES:50.27% | U=571169.5<br>**p=8.891 × 10$^{-37}$**<br>CLES:67.19% | U=742427.5<br>**p=1.629 × 10$^{-41}$**<br>CLES:67.00% |
| Guinea Pig | U=363048.0<br>p=0.575<br>CLES:49.22% | U=537024.5<br>**p=5.794 × 10$^{-31}$**<br>CLES:65.83% | U=686091.0<br>**p=1.499 × 10$^{-39}$**<br>CLES:66.90% |
| American beaver | U=351629.5<br>p=0.717<br>CLES:49.49% | U=486525.5<br>**p=1.01 × 10$^{-24}$**<br>CLES:64.30% | U=630312.0<br>**p=2.344 × 10$^{-30}$**<br>CLES:64.90% |
| Wild yak | U=286302.0<br>p=0.495<br>CLES:48.99% | U=400034.0<br>**p=8.793 × 10$^{-24}$**<br>CLES:64.71% | U=502174.0<br>**p=1.521 × 10$^{-31}$**<br>CLES:66.17% |
| Coquerel's sifaka | U=359338.0<br>p=0.611<br>CLES:50.72% | U=537858.0<br>**p=1.03 × 10$^{-32}$**<br>CLES:66.36% | U=682816.0<br>**p=1.002 × 10$^{-35}$**<br>CLES:65.99% |
| Alpaca | U=13900.5<br>p=0.814<br>CLES:50.75% | U=20067.5<br>**p=1.139 × 10$^{-7}$**<br>CLES:66.46% | U=19937.5<br>**p=4.555 × 10$^{-7}$**<br>CLES:65.63% |
| Lesser Egyptian jerboa | U=318611.5<br>p=0.908<br>CLES:49.83% | U=491618.0<br>**p=3.235 × 10$^{-34}$**<br>CLES:67.21% | U=648522.5<br>**p=3.408 × 10$^{-41}$**<br>CLES:67.56% |
| Long-tailed chinchilla | U=356742.0<br>p=0.629<br>CLES:49.32% | U=536062.0<br>**p=1.533 × 10$^{-30}$**<br>CLES:65.75% | U=702659.5<br>**p=5.289 × 10$^{-39}$**<br>CLES:66.66% |
| Angola colobus | U=382542.5<br>p=0.965<br>CLES:50.06% | U=559513.0<br>**p=9.987 × 10$^{-35}$**<br>CLES:66.72% | U=728045.5<br>**p=3.604 × 10$^{-41}$**<br>CLES:67.01% |
| Ryukyu mouse | U=500925.5<br>p=0.788<br>CLES:50.35% | U=706369.5<br>**p=2.068 × 10$^{-29}$**<br>CLES:64.28% | U=901596.5<br>**p=6.851 × 10$^{-33}$**<br>CLES:64.15% |
| Chinese hamster PICR | U=476909.5<br>p=0.814<br>CLES:50.31% | U=700391.0<br>**p=6.132 × 10$^{-43}$**<br>CLES:67.72% | U=902176.5<br>**p=3.582 × 10$^{-46}$**<br>CLES:67.12% |
| Vervet-AGM | U=412745.0<br>p=0.919<br>CLES:50.14% | U=603721.0<br>**p=2.74 × 10$^{-36}$**<br>CLES:66.78% | U=776561.5<br>**p=1.767 × 10$^{-41}$**<br>CLES:66.79% |
| Cow | U=368644.5<br>p=0.861<br>CLES:49.75% | U=554058.5<br>**p=4.491 × 10$^{-32}$**<br>CLES:66.03% | U=702747.0<br>**p=2.646 × 10$^{-38}$**<br>CLES:66.50% |
| Bushbaby | U=384647.5<br>p=0.627<br>CLES:49.33% | U=583573.5<br>**p=1.587 × 10$^{-34}$**<br>CLES:66.51% | U=778063.0<br>**p=1.413 × 10$^{-43}$**<br>CLES:67.25% |
| Bonobo | U=393610.0<br>p=0.331<br>CLES:51.35% | U=555375.0<br>**p=3.131 × 10$^{-34}$**<br>CLES:66.62% | U=721756.5<br>**p=4.699 × 10$^{-35}$**<br>CLES:65.58% |

| | | | |
|---|---|---|---|
| Hyrax | U=30066.0<br>p=$7.558 \times 10^{-2}$<br>CLES:45.45% | U=36459.5<br>p=$1.205 \times 10^{-2}$<br>CLES:56.46% | U=51465.0<br>**p=$4.537 \times 10^{-6}$**<br>CLES:60.99% |
| Rabbit | U=309481.0<br>p=0.787<br>CLES:50.40% | U=419604.0<br>**p=$1.507 \times 10^{-17}$**<br>CLES:62.24% | U=538212.0<br>**p=$6.028 \times 10^{-19}$**<br>CLES:61.90% |
| Megabat | U=114412.5<br>p=0.74<br>CLES:49.38% | U=144357.0<br>**p=$1.888 \times 10^{-12}$**<br>CLES:63.22% | U=193440.0<br>**p=$4.069 \times 10^{-16}$**<br>CLES:64.18% |
| Dog | U=371255.0<br>p=0.696<br>CLES:50.55% | U=528274.0<br>**p=$2.751 \times 10^{-28}$**<br>CLES:65.13% | U=679626.0<br>**p=$2.211 \times 10^{-29}$**<br>CLES:64.34% |
| Steppe mouse | U=490179.5<br>p=0.429<br>CLES:51.03% | U=595238.0<br>**p=$2.346 \times 10^{-11}$**<br>CLES:58.60% | U=780511.0<br>**p=$5.763 \times 10^{-11}$**<br>CLES:57.80% |
| Greater bamboo lemur | U=397699.5<br>p=0.826<br>CLES:50.30% | U=580411.5<br>**p=$4.654 \times 10^{-34}$**<br>CLES:66.36% | U=729486.0<br>**p=$2.465 \times 10^{-38}$**<br>CLES:66.34% |
| Pig | U=378406.0<br>p=0.591<br>CLES:49.26% | U=574256.0<br>**p=$7.648 \times 10^{-34}$**<br>CLES:66.37% | U=739596.0<br>**p=$1.956 \times 10^{-41}$**<br>CLES:67.00% |
| Tree Shrew | U=18231.5<br>p=0.467<br>CLES:47.87% | U=21164.5<br>**p=$1.257 \times 10^{-5}$**<br>CLES:63.20% | U=24308.5<br>**p=$7.642 \times 10^{-7}$**<br>CLES:64.52% |
| Pika | U=46346.0<br>p=$7.366 \times 10^{-2}$<br>CLES:45.88% | U=56035.0<br>**p=$1.012 \times 10^{-6}$**<br>CLES:61.52% | U=78893.0<br>**p=$2.835 \times 10^{-12}$**<br>CLES:65.32% |
| Shrew | U=13252.5<br>p=0.157<br>CLES:54.64% | U=16093.5<br>**p=$1.967 \times 10^{-6}$**<br>CLES:65.56% | U=16857.5<br>p=$4.288 \times 10^{-4}$<br>CLES:61.18% |
| Squirrel | U=381474.5<br>p=0.688<br>CLES:50.56% | U=579407.0<br>**p=$6.502 \times 10^{-35}$**<br>CLES:66.64% | U=745119.0<br>**p=$5.08 \times 10^{-40}$**<br>CLES:66.65% |
| Gorilla | U=385900.0<br>p=0.778<br>CLES:50.39% | U=566831.5<br>**p=$9.648 \times 10^{-34}$**<br>CLES:66.42% | U=745790.5<br>**p=$2.111 \times 10^{-38}$**<br>CLES:66.26% |
| Cat | U=394647.5<br>p=0.843<br>CLES:50.27% | U=580090.0<br>**p=$8.233 \times 10^{-33}$**<br>CLES:66.04% | U=741008.0<br>**p=$3.989 \times 10^{-38}$**<br>CLES:66.22% |

### Mouse Reference Genome, Zeisel et al., 2015

| MWU Test | Glia vs Vasculature | Glia vs Neuron | Vasculature vs Neuron |
|---|---|---|---|
| Microbat | U=189035.0<br>p=2.689 × $10^{-2}$<br>CLES:53.98% | U=419580.5<br>**p=1.083 × $10^{-17}$**<br>CLES:62.35% | U=134403.5<br>**p=4.911 × $10^{-6}$**<br>CLES:58.76% |
| Hedgehog | U=10644.5<br>p=0.145<br>CLES:55.47% | U=18503.0<br>p=0.33<br>CLES:52.97% | U=5217.0<br>p=0.478<br>CLES:47.10% |
| Naked mole-rat male | U=190430.0<br>p=3.949 × $10^{-3}$<br>CLES:55.20% | U=443910.0<br>**p=4.707 × $10^{-27}$**<br>CLES:65.47% | U=140386.0<br>**p=1.568 × $10^{-7}$**<br>CLES:60.02% |
| Sooty mangabey | U=258624.0<br>p=1.698 × $10^{-3}$<br>CLES:55.26% | U=611170.0<br>**p=1.684 × $10^{-30}$**<br>CLES:65.21% | U=191626.0<br>**p=4.278 × $10^{-9}$**<br>CLES:60.41% |
| Olive baboon | U=255806.5<br>p=2.518 × $10^{-3}$<br>CLES:55.07% | U=594897.5<br>**p=1.009 × $10^{-27}$**<br>CLES:64.52% | U=185875.5<br>**p=4.751 × $10^{-8}$**<br>CLES:59.72% |
| American bison | U=175708.0<br>p=7.386 × $10^{-3}$<br>CLES:54.94% | U=383311.0<br>**p=3.081 × $10^{-22}$**<br>CLES:64.46% | U=118467.0<br>**p=1.3 × $10^{-6}$**<br>CLES:59.59% |
| Leopard | U=229932.0<br>p=4.56 × $10^{-3}$<br>CLES:54.89% | U=566196.0<br>**p=1.066 × $10^{-28}$**<br>CLES:64.98% | U=175179.0<br>**p=1.958 × $10^{-8}$**<br>CLES:60.20% |
| Bolivian squirrel monkey | U=231471.0<br>p=1.05 × $10^{-2}$<br>CLES:54.39% | U=561607.5<br>**p=1.68 × $10^{-30}$**<br>CLES:65.55% | U=178724.5<br>**p=2.387 × $10^{-10}$**<br>CLES:61.48% |
| Crab-eating macaque | U=258912.5<br>p=2.637 × $10^{-3}$<br>CLES:55.04% | U=602704.5<br>**p=1.658 × $10^{-30}$**<br>CLES:65.29% | U=187496.5<br>**p=2.415 × $10^{-9}$**<br>CLES:60.63% |
| Human | U=276284.0<br>p=5.472 × $10^{-4}$<br>CLES:55.70% | U=645569.5<br>**p=4.078 × $10^{-31}$**<br>CLES:65.15% | U=202382.0<br>**p=2.736 × $10^{-8}$**<br>CLES:59.69% |
| Red fox | U=221618.0<br>p=4.876 × $10^{-3}$<br>CLES:54.90% | U=508519.0<br>**p=1.89 × $10^{-24}$**<br>CLES:64.12% | U=157860.0<br>**p=4.519 × $10^{-7}$**<br>CLES:59.33% |
| Shrew mouse | U=301177.0<br>p=1.734 × $10^{-3}$<br>CLES:55.07% | U=726162.0<br>**p=1.297 × $10^{-34}$**<br>CLES:65.61% | U=220338.0<br>**p=2.53 × $10^{-10}$**<br>CLES:60.85% |
| Arctic ground squirrel | U=244954.5<br>p=1.464 × $10^{-2}$<br>CLES:54.11% | U=612263.0<br>**p=2.706 × $10^{-34}$**<br>CLES:66.20% | U=198823.0<br>**p=3.018 × $10^{-12}$**<br>CLES:62.37% |
| Chinese hamster CriGri | U=249077.5<br>p=1.361 × $10^{-3}$<br>CLES:55.45% | U=578158.0<br>**p=1.468 × $10^{-24}$**<br>CLES:63.69% | U=171170.5<br>**p=1.783 × $10^{-6}$**<br>CLES:58.65% |
| Opossum | U=199228.0<br>p=1.005 × $10^{-2}$ | U=448997.0<br>**p=8.611 × $10^{-16}$** | U=142163.0<br>p=2.548 × $10^{-4}$ |

| | | | |
|---|---|---|---|
| | CLES:54.58% | CLES:61.33% | CLES:56.88% |
| Dolphin | U=72582.5 | U=166110.5 | U=50476.5 |
| | p=$6.396 \times 10^{-2}$ | **p=$3.11 \times 10^{-16}$** | **p=$1.971 \times 10^{-5}$** |
| | CLES:54.26% | CLES:65.05% | CLES:60.53% |
| Capuchin | U=244631.0 | U=576614.0 | U=184251.0 |
| | p=$4.677 \times 10^{-3}$ | **p=$4.66 \times 10^{-30}$** | **p=$9.603 \times 10^{-10}$** |
| | CLES:54.79% | CLES:65.32% | CLES:60.96% |
| Mouse Lemur | U=234564.0 | U=566581.5 | U=178412.5 |
| | p=$9.598 \times 10^{-3}$ | **p=$2.32 \times 10^{-30}$** | **p=$2.006 \times 10^{-10}$** |
| | CLES:54.43% | CLES:65.48% | CLES:61.53% |
| Damara mole rat | U=207364.0 | U=491414.0 | U=156209.0 |
| | p=$5.373 \times 10^{-3}$ | **p=$3.744 \times 10^{-25}$** | **p=$1.954 \times 10^{-7}$** |
| | CLES:54.91% | CLES:64.44% | CLES:59.69% |
| Marmoset | U=248908.5 | U=594429.0 | U=190380.0 |
| | p=$1.467 \times 10^{-3}$ | **p=$3.577 \times 10^{-30}$** | **p=$3.813 \times 10^{-9}$** |
| | CLES:55.37% | CLES:65.20% | CLES:60.48% |
| Goat | U=229010.0 | U=547362.0 | U=169410.0 |
| | p=$5.208 \times 10^{-2}$ | **p=$4.404 \times 10^{-23}$** | **p=$7.942 \times 10^{-9}$** |
| | CLES:53.34% | CLES:63.41% | CLES:60.56% |
| Common wombat | U=178720.0 | U=421492.0 | U=135952.0 |
| | p=$1.249 \times 10^{-2}$ | **p=$8.265 \times 10^{-21}$** | **p=$2.416 \times 10^{-6}$** |
| | CLES:54.56% | CLES:63.49% | CLES:59.07% |
| American black bear | U=185625.0 | U=413772.0 | U=131326.0 |
| | p=$4.178 \times 10^{-2}$ | **p=$5.695 \times 10^{-22}$** | **p=$2.429 \times 10^{-8}$** |
| | CLES:53.68% | CLES:64.07% | CLES:60.83% |
| Sloth | U=4862.5 | U=11967.0 | U=3056.0 |
| | p=0.434 | **p=$1.774 \times 10^{-6}$** | p=$3.275 \times 10^{-3}$ |
| | CLES:53.64% | CLES:67.38% | CLES:64.97% |
| Polar bear | U=188587.5 | U=423807.5 | U=131059.5 |
| | p=$1.437 \times 10^{-2}$ | **p=$1.344 \times 10^{-22}$** | **p=$1.419 \times 10^{-7}$** |
| | CLES:54.43% | CLES:64.20% | CLES:60.21% |
| Alpine marmot | U=225379.0 | U=523903.0 | U=169248.0 |
| | p=$1.689 \times 10^{-2}$ | **p=$6.651 \times 10^{-27}$** | **p=$5.035 \times 10^{-9}$** |
| | CLES:54.11% | CLES:64.77% | CLES:60.67% |
| Prairie vole | U=265248.0 | U=648864.5 | U=194787.0 |
| | p=$4.167 \times 10^{-4}$ | **p=$3.952 \times 10^{-32}$** | **p=$1.585 \times 10^{-8}$** |
| | CLES:55.92% | CLES:65.40% | CLES:60.00% |
| Donkey | U=255742.0 | U=579398.5 | U=176255.5 |
| | p=$1.119 \times 10^{-4}$ | **p=$5.367 \times 10^{-25}$** | **p=$1.616 \times 10^{-5}$** |
| | CLES:56.53% | CLES:63.79% | CLES:57.72% |
| Golden Hamster | U=241336.5 | U=595783.0 | U=183994.0 |
| | p=$5.363 \times 10^{-3}$ | **p=$1.099 \times 10^{-28}$** | **p=$6.406 \times 10^{-9}$** |
| | CLES:54.75% | CLES:64.78% | CLES:60.43% |
| Armadillo | U=210092.5 | U=478041.5 | U=147136.0 |
| | p=$1.792 \times 10^{-3}$ | **p=$7.098 \times 10^{-24}$** | **p=$4.307 \times 10^{-6}$** |
| | CLES:55.52% | CLES:64.15% | CLES:58.63% |
| Ugandan red Colobus | U=234314.5 | U=570003.5 | U=175137.5 |
| | p=$4.306 \times 10^{-3}$ | **p=$7.251 \times 10^{-28}$** | **p=$3.311 \times 10^{-8}$** |

| | | | |
|---|---|---|---|
| | CLES:54.91% | CLES:64.72% | CLES:60.02% |
| Pig-tailed macaque | U=259989.0<br>p=$2.718 \times 10^{-3}$<br>CLES:55.00% | U=610692.5<br>**p=$1.7 \times 10^{-31}$**<br>CLES:65.49% | U=195096.0<br>**p=$6.876 \times 10^{-10}$**<br>CLES:60.89% |
| Panda | U=231900.0<br>p=$7.107 \times 10^{-3}$<br>CLES:54.63% | U=537866.0<br>**p=$2.579 \times 10^{-24}$**<br>CLES:63.86% | U=165823.0<br>**p=$5.986 \times 10^{-7}$**<br>CLES:59.12% |
| Ferret | U=225518.0<br>p=$5.551 \times 10^{-3}$<br>CLES:54.82% | U=537708.5<br>**p=$3.671 \times 10^{-25}$**<br>CLES:64.13% | U=163701.5<br>**p=$1.664 \times 10^{-7}$**<br>CLES:59.63% |
| Koala | U=218551.0<br>**p=$6.396 \times 10^{-5}$**<br>CLES:57.02% | U=499879.0<br>**p=$1.025 \times 10^{-22}$**<br>CLES:63.55% | U=153505.0<br>p=$4.716 \times 10^{-4}$<br>CLES:56.45% |
| Wallaby | U=17390.0<br>p=0.297<br>CLES:53.36% | U=37219.0<br>**p=$4.847 \times 10^{-5}$**<br>CLES:60.62% | U=13508.0<br>p=$3.421 \times 10^{-2}$<br>CLES:57.16% |
| Tarsier | U=159281.0<br>p=$3.018 \times 10^{-2}$<br>CLES:54.11% | U=369011.5<br>**p=$1.202 \times 10^{-18}$**<br>CLES:63.19% | U=109070.0<br>**p=$1.658 \times 10^{-5}$**<br>CLES:58.72% |
| Degu | U=176222.0<br>p=$7.731 \times 10^{-2}$<br>CLES:53.21% | U=432075.0<br>**p=$1.174 \times 10^{-24}$**<br>CLES:64.76% | U=144032.5<br>**p=$6.37 \times 10^{-10}$**<br>CLES:61.84% |
| Chinese hamster CHOK1GS | U=291684.5<br>p=$2.392 \times 10^{-3}$<br>CLES:54.95% | U=707504.5<br>**p=$3.602 \times 10^{-32}$**<br>CLES:65.08% | U=214746.5<br>**p=$2.354 \times 10^{-9}$**<br>CLES:60.30% |
| Drill | U=244704.0<br>p=$4.04 \times 10^{-4}$<br>CLES:56.05% | U=554665.5<br>**p=$1.622 \times 10^{-26}$**<br>CLES:64.45% | U=166650.0<br>**p=$2.583 \times 10^{-6}$**<br>CLES:58.55% |
| Gibbon | U=244270.5<br>p=$1.118 \times 10^{-4}$<br>CLES:56.59% | U=558091.0<br>**p=$8.012 \times 10^{-31}$**<br>CLES:65.68% | U=171931.0<br>**p=$3.452 \times 10^{-7}$**<br>CLES:59.22% |
| Rat | U=277151.0<br>p=$1.048 \times 10^{-2}$<br>CLES:54.21% | U=672987.5<br>**p=$1.155 \times 10^{-28}$**<br>CLES:64.32% | U=206738.0<br>**p=$2.46 \times 10^{-9}$**<br>CLES:60.39% |
| Macaque | U=268287.0<br>p=$7.806 \times 10^{-4}$<br>CLES:55.58% | U=619399.0<br>**p=$8.927 \times 10^{-27}$**<br>CLES:64.08% | U=193554.0<br>**p=$8.553 \times 10^{-7}$**<br>CLES:58.64% |
| American mink | U=211177.5<br>p=$3.328 \times 10^{-3}$<br>CLES:55.18% | U=497888.0<br>**p=$1.225 \times 10^{-24}$**<br>CLES:64.24% | U=152378.0<br>**p=$1.321 \times 10^{-6}$**<br>CLES:59.04% |
| Gelada | U=257705.0<br>p=$4.671 \times 10^{-3}$<br>CLES:54.73% | U=610137.5<br>**p=$1.935 \times 10^{-28}$**<br>CLES:64.63% | U=191099.0<br>**p=$9.583 \times 10^{-9}$**<br>CLES:60.17% |
| Sheep | U=215705.0<br>p=$3.489 \times 10^{-2}$<br>CLES:53.70% | U=508839.0<br>**p=$1.474 \times 10^{-21}$**<br>CLES:63.16% | U=154370.0<br>**p=$1.887 \times 10^{-7}$**<br>CLES:59.72% |
| Tasmanian devil | U=180753.0<br>p=$3.122 \times 10^{-3}$ | U=383837.0<br>**p=$7.359 \times 10^{-16}$** | U=120684.5<br>p=$1.421 \times 10^{-3}$ |

| | | | |
|---|---|---|---|
| | CLES:55.40% | CLES:61.86% | CLES:56.21% |
| Elephant | U=229492.5<br>p=$4.189 \times 10^{-2}$<br>CLES:53.49% | U=531273.0<br>**p=$2.523 \times 10^{-21}$**<br>CLES:62.92% | U=166878.0<br>**p=$1.249 \times 10^{-7}$**<br>CLES:59.66% |
| Algerian mouse | U=298427.0<br>p=$1.437 \times 10^{-2}$<br>CLES:53.94% | U=720343.5<br>**p=$8.339 \times 10^{-28}$**<br>CLES:63.74% | U=217067.5<br>**p=$1.19 \times 10^{-8}$**<br>CLES:59.67% |
| Dingo | U=217307.0<br>p=$1.442 \times 10^{-2}$<br>CLES:54.27% | U=530336.0<br>**p=$1.253 \times 10^{-26}$**<br>CLES:64.63% | U=165110.5<br>**p=$4.881 \times 10^{-9}$**<br>CLES:60.80% |
| Golden snub-nosed monkey | U=234408.5<br>p=$5.235 \times 10^{-3}$<br>CLES:54.78% | U=556090.5<br>**p=$1.953 \times 10^{-30}$**<br>CLES:65.59% | U=175602.5<br>**p=$3.946 \times 10^{-10}$**<br>CLES:61.37% |
| Horse | U=252932.0<br>p=$2.846 \times 10^{-3}$<br>CLES:55.01% | U=584524.0<br>**p=$1.712 \times 10^{-26}$**<br>CLES:64.22% | U=186116.0<br>**p=$2.309 \times 10^{-8}$**<br>CLES:59.95% |
| Black snub-nosed monkey | U=235775.5<br>p=$4.912 \times 10^{-3}$<br>CLES:54.81% | U=555556.0<br>**p=$2.643 \times 10^{-29}$**<br>CLES:65.26% | U=174616.0<br>**p=$2.844 \times 10^{-9}$**<br>CLES:60.78% |
| Brazilian guinea pig | U=122682.0<br>p=0.276<br>CLES:52.17% | U=282001.0<br>**p=$6.174 \times 10^{-14}$**<br>CLES:61.93% | U=92216.0<br>**p=$7.646 \times 10^{-6}$**<br>CLES:59.46% |
| Orangutan | U=242403.0<br>p=$1.945 \times 10^{-3}$<br>CLES:55.25% | U=555581.5<br>**p=$1.093 \times 10^{-28}$**<br>CLES:65.07% | U=177941.0<br>**p=$4.91 \times 10^{-8}$**<br>CLES:59.80% |
| Northern American deer mouse | U=276643.0<br>p=$7.501 \times 10^{-4}$<br>CLES:55.58% | U=655088.5<br>**p=$1.365 \times 10^{-34}$**<br>CLES:66.06% | U=196149.5<br>**p=$1.745 \times 10^{-9}$**<br>CLES:60.61% |
| Naked mole-rat female | U=209447.0<br>p=$3.187 \times 10^{-3}$<br>CLES:55.20% | U=509991.5<br>**p=$1.884 \times 10^{-30}$**<br>CLES:65.92% | U=160751.0<br>**p=$5.523 \times 10^{-9}$**<br>CLES:60.84% |
| Daurian ground squirrel | U=185868.5<br>p=$7.211 \times 10^{-3}$<br>CLES:54.88% | U=399148.5<br>**p=$3.381 \times 10^{-23}$**<br>CLES:64.67% | U=124390.5<br>**p=$6.243 \times 10^{-7}$**<br>CLES:59.74% |
| Tiger | U=158433.5<br>p=$1.768 \times 10^{-2}$<br>CLES:54.51% | U=369974.0<br>**p=$1.06 \times 10^{-21}$**<br>CLES:64.38% | U=109976.0<br>**p=$1.384 \times 10^{-6}$**<br>CLES:59.80% |
| Lesser hedgehog tenrec | U=13137.0<br>p=0.656<br>CLES:51.55% | U=26093.0<br>p=$1.211 \times 10^{-4}$<br>CLES:61.17% | U=8422.0<br>p=$1.326 \times 10^{-2}$<br>CLES:59.44% |
| Mongolian gerbil | U=243991.0<br>**p=$3.06 \times 10^{-5}$**<br>CLES:57.18% | U=536632.5<br>**p=$3.872 \times 10^{-27}$**<br>CLES:64.80% | U=155880.5<br>**p=$1.296 \times 10^{-5}$**<br>CLES:58.04% |
| Ma's night monkey | U=230826.0<br>p=$9.802 \times 10^{-3}$<br>CLES:54.43% | U=563340.0<br>**p=$6.693 \times 10^{-31}$**<br>CLES:65.64% | U=180081.0<br>**p=$1.369 \times 10^{-10}$**<br>CLES:61.63% |

| | | | |
|---|---|---|---|
| Upper Galilee mountains blind mole rat | U=235349.5<br>p=2.575 × $10^{-2}$<br>CLES:53.82% | U=591914.0<br>**p=6.665 × $10^{-28}$**<br>CLES:64.59% | U=180180.0<br>**p=2.327 × $10^{-9}$**<br>CLES:60.81% |
| Kangaroo rat | U=186163.0<br>p=3.087 × $10^{-2}$<br>CLES:53.90% | U=447527.5<br>**p=6.64 × $10^{-23}$**<br>CLES:64.04% | U=143930.5<br>**p=4.034 × $10^{-8}$**<br>CLES:60.46% |
| Chimpanzee | U=236228.5<br>p=5.294 × $10^{-4}$<br>CLES:55.95% | U=564797.5<br>**p=8.105 × $10^{-34}$**<br>CLES:66.45% | U=176729.0<br>**p=9.128 × $10^{-10}$**<br>CLES:61.11% |
| Guinea Pig | U=226285.5<br>p=6.057 × $10^{-3}$<br>CLES:54.73% | U=533420.5<br>**p=5.049 × $10^{-25}$**<br>CLES:64.08% | U=172121.0<br>**p=1.555 × $10^{-7}$**<br>CLES:59.52% |
| American beaver | U=206608.0<br>p=5.908 × $10^{-2}$<br>CLES:53.31% | U=463666.0<br>**p=1.541 × $10^{-22}$**<br>CLES:63.82% | U=152228.0<br>**p=1.3 × $10^{-8}$**<br>CLES:60.63% |
| Wild yak | U=195594.5<br>p=2.196 × $10^{-3}$<br>CLES:55.54% | U=416346.0<br>**p=5.896 × $10^{-20}$**<br>CLES:63.32% | U=123737.0<br>**p=3.825 × $10^{-5}$**<br>CLES:58.02% |
| Coquerel's sifaka | U=224170.5<br>p=1.66 × $10^{-2}$<br>CLES:54.14% | U=542852.0<br>**p=1.751 × $10^{-25}$**<br>CLES:64.18% | U=170662.0<br>**p=1.417 × $10^{-8}$**<br>CLES:60.36% |
| Alpaca | U=11031.5<br>p=2.292 × $10^{-2}$<br>CLES:58.69% | U=26911.5<br>**p=5.109 × $10^{-13}$**<br>CLES:71.59% | U=7123.5<br>p=2.614 × $10^{-4}$<br>CLES:65.04% |
| Lesser Egyptian jerboa | U=202374.5<br>p=1.762 × $10^{-3}$<br>CLES:55.61% | U=504463.0<br>**p=8.977 × $10^{-27}$**<br>CLES:64.86% | U=148513.0<br>**p=6.679 × $10^{-7}$**<br>CLES:59.40% |
| Long-tailed chinchilla | U=229152.0<br>p=2.104 × $10^{-3}$<br>CLES:55.30% | U=551525.5<br>**p=1.462 × $10^{-29}$**<br>CLES:65.34% | U=174081.5<br>**p=9.376 × $10^{-9}$**<br>CLES:60.44% |
| Angola colobus | U=244230.0<br>p=4.486 × $10^{-3}$<br>CLES:54.83% | U=573735.0<br>**p=8.732 × $10^{-29}$**<br>CLES:64.99% | U=176563.5<br>**p=2.176 × $10^{-8}$**<br>CLES:60.11% |
| Ryukyu mouse | U=290770.0<br>p=3.485 × $10^{-2}$<br>CLES:53.41% | U=702832.0<br>**p=5.739 × $10^{-28}$**<br>CLES:63.95% | U=221211.0<br>**p=4.437 × $10^{-10}$**<br>CLES:60.66% |
| Chinese hamster PICR | U=278008.0<br>p=5.23 × $10^{-3}$<br>CLES:54.60% | U=695351.0<br>**p=4.758 × $10^{-34}$**<br>CLES:65.63% | U=213614.0<br>**p=1.984 × $10^{-11}$**<br>CLES:61.67% |
| Vervet-AGM | U=257922.5<br>p=2.802 × $10^{-3}$<br>CLES:55.01% | U=608602.5<br>**p=9.135 × $10^{-31}$**<br>CLES:65.31% | U=189837.5<br>**p=8.603 × $10^{-10}$**<br>CLES:60.91% |
| Cow | U=240601.0<br>p=8.518 × $10^{-3}$<br>CLES:54.49% | U=562684.0<br>**p=3.574 × $10^{-28}$**<br>CLES:64.91% | U=172817.0<br>**p=4.818 × $10^{-9}$**<br>CLES:60.64% |
| Bushbaby | U=246553.5<br>p=6.537 × $10^{-3}$<br>CLES:54.59% | U=589692.0<br>**p=9.491 × $10^{-32}$**<br>CLES:65.71% | U=186716.0<br>**p=1.564 × $10^{-10}$**<br>CLES:61.46% |

| | | | |
|---|---|---|---|
| Bonobo | U=243753.5<br>p=$4.943 \times 10^{-3}$<br>CLES:54.77% | U=559710.5<br>**p=$8.248 \times 10^{-28}$**<br>CLES:64.81% | U=175029.5<br>**p=$1.543 \times 10^{-8}$**<br>CLES:60.22% |
| Hyrax | U=20857.5<br>p=$7.547 \times 10^{-2}$<br>CLES:55.60% | U=39996.0<br>**p=$2.927 \times 10^{-5}$**<br>CLES:60.84% | U=12460.0<br>p=$9.524 \times 10^{-2}$<br>CLES:55.69% |
| Rabbit | U=198229.5<br>p=$3.365 \times 10^{-3}$<br>CLES:55.25% | U=449244.0<br>**p=$3.63 \times 10^{-20}$**<br>CLES:63.06% | U=140119.0<br>**p=$1.326 \times 10^{-5}$**<br>CLES:58.27% |
| Megabat | U=77378.0<br>p=0.287<br>CLES:52.36% | U=147606.0<br>**p=$5.998 \times 10^{-10}$**<br>CLES:61.63% | U=51677.0<br>p=$1.277 \times 10^{-4}$<br>CLES:59.24% |
| Dog | U=220454.0<br>p=$7.724 \times 10^{-2}$<br>CLES:53.05% | U=532801.0<br>**p=$1.075 \times 10^{-23}$**<br>CLES:63.68% | U=170206.5<br>**p=$1.427 \times 10^{-9}$**<br>CLES:61.09% |
| Steppe mouse | U=284735.0<br>p=$4.919 \times 10^{-3}$<br>CLES:54.58% | U=638344.0<br>**p=$4.263 \times 10^{-15}$**<br>CLES:60.03% | U=195516.0<br>p=$9.343 \times 10^{-4}$<br>CLES:55.69% |
| Greater bamboo lemur | U=247892.0<br>p=$1.3 \times 10^{-3}$<br>CLES:55.46% | U=580000.5<br>**p=$1.838 \times 10^{-27}$**<br>CLES:64.54% | U=177653.5<br>**p=$2.274 \times 10^{-7}$**<br>CLES:59.31% |
| Pig | U=251443.0<br>p=$7.784 \times 10^{-4}$<br>CLES:55.68% | U=591833.5<br>**p=$4.622 \times 10^{-32}$**<br>CLES:65.78% | U=184678.5<br>**p=$2.852 \times 10^{-9}$**<br>CLES:60.63% |
| Tree Shrew | U=9949.5<br>p=$7.571 \times 10^{-2}$<br>CLES:56.73% | U=19724.0<br>p=$2.602 \times 10^{-4}$<br>CLES:61.27% | U=6282.0<br>p=0.232<br>CLES:54.84% |
| Pika | U=34341.0<br>p=$2.162 \times 10^{-2}$<br>CLES:56.35% | U=54563.0<br>**p=$1.298 \times 10^{-6}$**<br>CLES:61.76% | U=17696.0<br>p=0.106<br>CLES:54.98% |
| Shrew | U=6263.0<br>p=0.283<br>CLES:54.67% | U=15029.5<br>p=$4.459 \times 10^{-3}$<br>CLES:59.34% | U=3907.0<br>p=0.342<br>CLES:54.40% |
| Squirrel | U=244689.0<br>p=$1.846 \times 10^{-3}$<br>CLES:55.28% | U=587353.0<br>**p=$2.38 \times 10^{-31}$**<br>CLES:65.59% | U=186472.0<br>**p=$3.039 \times 10^{-9}$**<br>CLES:60.60% |
| Gorilla | U=243221.0<br>p=$3.045 \times 10^{-3}$<br>CLES:55.03% | U=569837.0<br>**p=$1.472 \times 10^{-27}$**<br>CLES:64.63% | U=178955.0<br>**p=$3.738 \times 10^{-8}$**<br>CLES:59.90% |
| Cat | U=243988.0<br>p=$2.18 \times 10^{-2}$<br>CLES:53.89% | U=600247.0<br>**p=$1.034 \times 10^{-28}$**<br>CLES:64.77% | U=186332.0<br>**p=$5.956 \times 10^{-10}$**<br>CLES:61.10% |

### Mouse Reference Genome, Zeisel et al., 2018

| MWU Test | CNS_Glia vs CNS_Endothelia | CNS_Glia vs CNS_Neuron | CNS_Endothelia vs CNS_Neuron |
|---|---|---|---|
| Microbat | U=38709.0<br>p=0.436<br>CLES:47.77% | U=1238984.5<br>**p=$2.081 \times 10^{-35}$**<br>CLES:65.33% | U=221816.0<br>**p=$3.751 \times 10^{-10}$**<br>CLES:66.93% |
| Hedgehog | U=2682.0<br>p=0.919<br>CLES:49.44% | U=57805.0<br>**p=$4.889 \times 10^{-5}$**<br>CLES:60.54% | U=13108.0<br>p=$3.152 \times 10^{-2}$<br>CLES:60.80% |
| Naked mole-rat male | U=49834.5<br>p=0.841<br>CLES:50.54% | U=1341272.5<br>**p=$1.012 \times 10^{-45}$**<br>CLES:67.10% | U=248998.0<br>**p=$9.823 \times 10^{-11}$**<br>CLES:66.41% |
| Sooty mangabey | U=67275.0<br>p=0.364<br>CLES:52.29% | U=1777227.0<br>**p=$4.46 \times 10^{-52}$**<br>CLES:67.08% | U=322237.0<br>**p=$1.874 \times 10^{-10}$**<br>CLES:65.13% |
| Olive baboon | U=66302.0<br>p=0.515<br>CLES:51.64% | U=1765491.5<br>**p=$3.399 \times 10^{-53}$**<br>CLES:67.33% | U=326646.5<br>**p=$3.126 \times 10^{-11}$**<br>CLES:65.72% |
| American bison | U=40454.0<br>p=0.473<br>CLES:47.98% | U=1164808.5<br>**p=$2.824 \times 10^{-39}$**<br>CLES:66.32% | U=217967.0<br>**p=$1.039 \times 10^{-11}$**<br>CLES:68.05% |
| Leopard | U=59229.0<br>p=0.81<br>CLES:49.38% | U=1689520.5<br>**p=$2.477 \times 10^{-49}$**<br>CLES:66.84% | U=314109.0<br>**p=$1.448 \times 10^{-12}$**<br>CLES:67.14% |
| Bolivian squirrel monkey | U=54355.0<br>p=0.541<br>CLES:48.39% | U=1637645.5<br>**p=$1.432 \times 10^{-52}$**<br>CLES:67.57% | U=299367.0<br>**p=$3.249 \times 10^{-14}$**<br>CLES:68.81% |
| Crab-eating macaque | U=65932.0<br>p=0.582<br>CLES:51.40% | U=1768115.0<br>**p=$1.665 \times 10^{-53}$**<br>CLES:67.32% | U=320120.0<br>**p=$2.314 \times 10^{-11}$**<br>CLES:65.93% |
| Human | U=67402.0<br>p=0.876<br>CLES:50.39% | U=1857569.5<br>**p=$8.454 \times 10^{-56}$**<br>CLES:67.60% | U=353421.0<br>**p=$1.856 \times 10^{-13}$**<br>CLES:67.20% |
| Red fox | U=52950.0<br>p=0.75<br>CLES:49.15% | U=1527964.0<br>**p=$1.581 \times 10^{-40}$**<br>CLES:65.50% | U=277735.5<br>**p=$6.145 \times 10^{-11}$**<br>CLES:66.38% |
| Shrew mouse | U=78025.0<br>p=0.212<br>CLES:53.07% | U=2013558.0<br>**p=$5.518 \times 10^{-48}$**<br>CLES:65.73% | U=346518.0<br>**p=$5.182 \times 10^{-8}$**<br>CLES:62.60% |
| Arctic ground squirrel | U=64678.5<br>p=0.711<br>CLES:50.93% | U=1736663.5<br>**p=$1.709 \times 10^{-50}$**<br>CLES:66.94% | U=327197.0<br>**p=$2.412 \times 10^{-11}$**<br>CLES:65.81% |
| Chinese hamster CriGri | U=59918.5<br>p=0.855<br>CLES:49.53% | U=1689834.0<br>**p=$4.617 \times 10^{-46}$**<br>CLES:66.13% | U=304311.0<br>**p=$7.874 \times 10^{-12}$**<br>CLES:66.67% |
| Opossum | U=54495.0<br>p=0.156 | U=1293630.5<br>**p=$4.062 \times 10^{-21}$** | U=225935.0<br>p=$6.049 \times 10^{-3}$ |

| | | | |
|---|---|---|---|
| | CLES:53.80% | CLES:61.26% | CLES:56.91% |
| Dolphin | U=20558.0<br>$p=0.976$<br>CLES:50.10% | U=494064.0<br>**$p=7.614 \times 10^{-21}$**<br>CLES:64.37% | U=102000.0<br>**$p=3.053 \times 10^{-6}$**<br>CLES:64.42% |
| Capuchin | U=60744.0<br>$p=0.903$<br>CLES:49.69% | U=1718303.0<br>**$p=1.618 \times 10^{-54}$**<br>CLES:67.66% | U=317242.0<br>**$p=1.303 \times 10^{-13}$**<br>CLES:67.87% |
| Mouse Lemur | U=60546.0<br>$p=0.478$<br>CLES:51.83% | U=1616901.0<br>**$p=6.117 \times 10^{-49}$**<br>CLES:66.98% | U=300366.0<br>**$p=1.676 \times 10^{-10}$**<br>CLES:65.47% |
| Damara mole rat | U=48460.0<br>$p=0.814$<br>CLES:50.65% | U=1453794.5<br>**$p=5.977 \times 10^{-44}$**<br>CLES:66.53% | U=255018.0<br>**$p=5.221 \times 10^{-10}$**<br>CLES:66.07% |
| Marmoset | U=62793.0<br>$p=0.601$<br>CLES:51.34% | U=1706342.5<br>**$p=9.461 \times 10^{-51}$**<br>CLES:67.05% | U=310809.0<br>**$p=1.132 \times 10^{-10}$**<br>CLES:65.51% |
| Goat | U=53212.0<br>$p=0.791$<br>CLES:49.29% | U=1606565.0<br>**$p=1.511 \times 10^{-42}$**<br>CLES:65.75% | U=282774.0<br>**$p=5.03 \times 10^{-11}$**<br>CLES:66.56% |
| Common wombat | U=46669.0<br>$p=0.623$<br>CLES:51.36% | U=1225531.5<br>**$p=4.053 \times 10^{-25}$**<br>CLES:62.64% | U=222599.0<br>**$p=1.603 \times 10^{-5}$**<br>CLES:61.18% |
| American black bear | U=45504.0<br>$p=0.425$<br>CLES:47.83% | U=1252478.5<br>**$p=2.573 \times 10^{-31}$**<br>CLES:64.15% | U=243923.0<br>**$p=2.301 \times 10^{-10}$**<br>CLES:66.19% |
| Sloth | U=1096.0<br>$p=0.833$<br>CLES:48.50% | U=32129.5<br>**$p=4.763 \times 10^{-7}$**<br>CLES:65.36% | U=5822.0<br>$p=1.049 \times 10^{-2}$<br>CLES:66.92% |
| Polar bear | U=42211.0<br>$p=0.276$<br>CLES:46.98% | U=1290889.0<br>**$p=1.445 \times 10^{-40}$**<br>CLES:66.27% | U=242230.0<br>**$p=3.15 \times 10^{-12}$**<br>CLES:68.20% |
| Alpine marmot | U=53697.0<br>$p=0.461$<br>CLES:48.08% | U=1561975.0<br>**$p=7.56 \times 10^{-48}$**<br>CLES:66.97% | U=303546.0<br>**$p=1.037 \times 10^{-13}$**<br>CLES:68.19% |
| Prairie vole | U=69077.0<br>$p=0.732$<br>CLES:50.85% | U=1844328.5<br>**$p=3.206 \times 10^{-50}$**<br>CLES:66.58% | U=341058.0<br>**$p=5.523 \times 10^{-11}$**<br>CLES:65.33% |
| Donkey | U=58068.0<br>$p=0.689$<br>CLES:51.05% | U=1672845.0<br>**$p=5.688 \times 10^{-42}$**<br>CLES:65.49% | U=291569.5<br>**$p=6.245 \times 10^{-9}$**<br>CLES:64.39% |
| Golden Hamster | U=62691.5<br>$p=0.908$<br>CLES:49.71% | U=1667224.0<br>**$p=7.787 \times 10^{-44}$**<br>CLES:65.75% | U=312182.5<br>**$p=8.431 \times 10^{-11}$**<br>CLES:65.48% |
| Armadillo | U=48486.0<br>$p=0.572$<br>CLES:51.56% | U=1357968.5<br>**$p=2.826 \times 10^{-36}$**<br>CLES:65.06% | U=234839.0<br>**$p=1.715 \times 10^{-7}$**<br>CLES:63.59% |
| Ugandan red Colobus | U=61805.0<br>$p=0.699$ | U=1676911.0<br>**$p=4.454 \times 10^{-50}$** | U=307615.0<br>**$p=2.068 \times 10^{-11}$** |

| | | | |
|---|---|---|---|
| | CLES:50.99% | CLES:66.97% | CLES:66.18% |
| Pig-tailed macaque | U=67576.0<br>p=0.51<br>CLES:51.65% | U=1776384.5<br>**$p=9.999 \times 10^{-54}$**<br>CLES:67.38% | U=329816.5<br>**$p=3.782 \times 10^{-11}$**<br>CLES:65.56% |
| Panda | U=54738.0<br>p=0.612<br>CLES:48.67% | U=1614315.0<br>**$p=1.01 \times 10^{-44}$**<br>CLES:66.21% | U=303215.0<br>**$p=1.348 \times 10^{-12}$**<br>CLES:67.45% |
| Ferret | U=54622.0<br>p=0.761<br>CLES:50.81% | U=1605324.5<br>**$p=8.522 \times 10^{-47}$**<br>CLES:66.63% | U=282142.0<br>**$p=1.458 \times 10^{-10}$**<br>CLES:66.10% |
| Koala | U=59370.0<br>p=0.188<br>CLES:53.44% | U=1450983.5<br>**$p=6.581 \times 10^{-26}$**<br>CLES:62.31% | U=260944.0<br>$p=5.866 \times 10^{-4}$<br>CLES:58.41% |
| Wallaby | U=5829.0<br>p=0.569<br>CLES:47.53% | U=104194.0<br>$p=3.186 \times 10^{-2}$<br>CLES:54.69% | U=27573.0<br>$p=9.774 \times 10^{-2}$<br>CLES:56.59% |
| Tarsier | U=42751.0<br>p=0.834<br>CLES:50.59% | U=1200998.0<br>**$p=1.186 \times 10^{-44}$**<br>CLES:67.37% | U=214113.0<br>**$p=1.446 \times 10^{-10}$**<br>CLES:67.08% |
| Degu | U=50091.0<br>p=0.112<br>CLES:54.35% | U=1285540.0<br>**$p=7.447 \times 10^{-43}$**<br>CLES:66.82% | U=234222.0<br>**$p=1.573 \times 10^{-7}$**<br>CLES:63.44% |
| Chinese hamster CHOK1GS | U=72866.0<br>p=0.87<br>CLES:49.60% | U=1994063.5<br>**$p=3.288 \times 10^{-53}$**<br>CLES:66.74% | U=377151.0<br>**$p=4.827 \times 10^{-14}$**<br>CLES:67.30% |
| Drill | U=58711.0<br>p=0.89<br>CLES:50.36% | U=1657687.0<br>**$p=8.673 \times 10^{-50}$**<br>CLES:66.96% | U=298663.5<br>**$p=5.709 \times 10^{-12}$**<br>CLES:66.90% |
| Gibbon | U=60156.0<br>p=0.498<br>CLES:51.76% | U=1644300.0<br>**$p=4.559 \times 10^{-53}$**<br>CLES:67.62% | U=294448.0<br>**$p=5.84 \times 10^{-11}$**<br>CLES:66.02% |
| Rat | U=68871.0<br>p=0.704<br>CLES:50.96% | U=1923739.5<br>**$p=3.159 \times 10^{-50}$**<br>CLES:66.37% | U=336558.0<br>**$p=1.053 \times 10^{-10}$**<br>CLES:65.32% |
| Macaque | U=67907.0<br>p=0.767<br>CLES:50.74% | U=1814559.0<br>**$p=2.446 \times 10^{-47}$**<br>CLES:66.10% | U=331864.0<br>**$p=3.075 \times 10^{-10}$**<br>CLES:64.81% |
| American mink | U=54069.0<br>p=0.894<br>CLES:50.35% | U=1546235.5<br>**$p=3.596 \times 10^{-47}$**<br>CLES:66.84% | U=279861.0<br>**$p=6.933 \times 10^{-11}$**<br>CLES:66.28% |
| Gelada | U=65757.5<br>p=0.557<br>CLES:51.49% | U=1780196.5<br>**$p=4.038 \times 10^{-55}$**<br>CLES:67.64% | U=328339.0<br>**$p=6.273 \times 10^{-12}$**<br>CLES:66.32% |
| Sheep | U=49831.0<br>p=0.456<br>CLES:47.98% | U=1539288.5<br>**$p=5.474 \times 10^{-40}$**<br>CLES:65.38% | U=274145.0<br>**$p=1.013 \times 10^{-11}$**<br>CLES:67.34% |
| Tasmanian devil | U=50145.0<br>$p=7.845 \times 10^{-2}$ | U=1190644.5<br>**$p=1.129 \times 10^{-23}$** | U=204749.0<br>$p=6.231 \times 10^{-3}$ |

| | | | |
|---|---|---|---|
| | CLES:54.84% | CLES:62.28% | CLES:57.06% |
| Elephant | U=52795.0<br>$p=0.895$<br>CLES:49.65% | U=1527962.0<br>**$p=1.597 \times 10^{-37}$**<br>CLES:64.93% | U=280261.0<br>**$p=5.288 \times 10^{-10}$**<br>CLES:65.55% |
| Algerian mouse | U=79745.0<br>$p=0.127$<br>CLES:53.73% | U=1963361.0<br>**$p=5.979 \times 10^{-36}$**<br>CLES:63.38% | U=329412.0<br>$p=1.394 \times 10^{-4}$<br>CLES:58.68% |
| Dingo | U=54738.0<br>$p=0.628$<br>CLES:48.74% | U=1540989.0<br>**$p=2.804 \times 10^{-39}$**<br>CLES:65.30% | U=301714.0<br>**$p=1.509 \times 10^{-11}$**<br>CLES:66.44% |
| Golden snub-nosed monkey | U=66323.0<br>$p=0.301$<br>CLES:52.61% | U=1653873.5<br>**$p=1.703 \times 10^{-56}$**<br>CLES:68.15% | U=312731.0<br>**$p=1.655 \times 10^{-11}$**<br>CLES:65.91% |
| Horse | U=59910.0<br>$p=0.611$<br>CLES:51.33% | U=1713585.5<br>**$p=2.833 \times 10^{-47}$**<br>CLES:66.44% | U=301290.0<br>**$p=1.372 \times 10^{-9}$**<br>CLES:64.85% |
| Black snub-nosed monkey | U=58809.0<br>$p=0.872$<br>CLES:49.58% | U=1626178.5<br>**$p=9.257 \times 10^{-48}$**<br>CLES:66.68% | U=305060.0<br>**$p=3.797 \times 10^{-12}$**<br>CLES:66.82% |
| Brazilian guinea pig | U=29918.0<br>$p=0.752$<br>CLES:50.98% | U=824027.0<br>**$p=3.325 \times 10^{-19}$**<br>CLES:62.02% | U=142047.0<br>**$p=9.017 \times 10^{-5}$**<br>CLES:61.50% |
| Orangutan | U=55688.0<br>$p=0.533$<br>CLES:48.38% | U=1620198.0<br>**$p=1.236 \times 10^{-45}$**<br>CLES:66.36% | U=311058.0<br>**$p=2.277 \times 10^{-13}$**<br>CLES:67.87% |
| Northern American deer mouse | U=69634.0<br>$p=0.998$<br>CLES:49.99% | U=1900278.5<br>**$p=3.903 \times 10^{-55}$**<br>CLES:67.20% | U=345519.0<br>**$p=1.528 \times 10^{-13}$**<br>CLES:67.32% |
| Naked mole-rat female | U=60465.0<br>$p=0.5$<br>CLES:51.73% | U=1519753.5<br>**$p=5.838 \times 10^{-49}$**<br>CLES:67.20% | U=293951.0<br>**$p=5.357 \times 10^{-11}$**<br>CLES:65.76% |
| Daurian ground squirrel | U=42557.0<br>$p=0.765$<br>CLES:49.16% | U=1235115.0<br>**$p=2.019 \times 10^{-43}$**<br>CLES:67.06% | U=227220.0<br>**$p=5.41 \times 10^{-11}$**<br>CLES:67.26% |
| Tiger | U=37437.0<br>$p=0.177$<br>CLES:46.11% | U=1176063.5<br>**$p=7.724 \times 10^{-40}$**<br>CLES:66.33% | U=209281.0<br>**$p=2.303 \times 10^{-13}$**<br>CLES:70.01% |
| Lesser hedgehog tenrec | U=3269.0<br>$p=0.503$<br>CLES:46.57% | U=77796.0<br>**$p=1.15 \times 10^{-7}$**<br>CLES:62.82% | U=17775.0<br>$p=6.352 \times 10^{-4}$<br>CLES:66.25% |
| Mongolian gerbil | U=63905.0<br>$p=0.511$<br>CLES:51.67% | U=1570130.5<br>**$p=3.508 \times 10^{-43}$**<br>CLES:65.85% | U=298022.5<br>**$p=1.613 \times 10^{-9}$**<br>CLES:64.35% |
| Ma's night monkey | U=58357.0<br>$p=0.877$<br>CLES:50.41% | U=1666418.0<br>**$p=6.286 \times 10^{-52}$**<br>CLES:67.34% | U=297208.0<br>**$p=4.263 \times 10^{-12}$**<br>CLES:67.05% |

| | | | |
|---|---|---|---|
| Upper Galilee mountains blind mole rat | U=61997.0<br>p=0.779<br>CLES:50.71% | U=1679514.5<br>**p=7.106 × $10^{-47}$**<br>CLES:66.44% | U=318416.0<br>**p=4.084 × $10^{-11}$**<br>CLES:65.78% |
| Kangaroo rat | U=45960.0<br>p=0.773<br>CLES:50.80% | U=1304297.0<br>**p=1.093 × $10^{-34}$**<br>CLES:64.98% | U=240464.0<br>**p=4.731 × $10^{-8}$**<br>CLES:64.14% |
| Chimpanzee | U=62798.0<br>p=0.198<br>CLES:53.33% | U=1651682.5<br>**p=2.161 × $10^{-54}$**<br>CLES:67.81% | U=290639.5<br>**p=7.337 × $10^{-10}$**<br>CLES:65.02% |
| Guinea Pig | U=56403.0<br>p=0.77<br>CLES:50.77% | U=1561860.0<br>**p=3.93 × $10^{-41}$**<br>CLES:65.68% | U=297039.0<br>**p=6.469 × $10^{-10}$**<br>CLES:65.10% |
| American beaver | U=57694.0<br>p=0.985<br>CLES:49.95% | U=1412631.0<br>**p=1.287 × $10^{-36}$**<br>CLES:64.95% | U=289639.0<br>**p=4.926 × $10^{-10}$**<br>CLES:64.85% |
| Wild yak | U=45244.0<br>p=0.525<br>CLES:48.24% | U=1313943.0<br>**p=3.772 × $10^{-42}$**<br>CLES:66.40% | U=237608.0<br>**p=1.113 × $10^{-11}$**<br>CLES:67.67% |
| Coquerel's sifaka | U=56405.0<br>p=0.983<br>CLES:50.06% | U=1600354.0<br>**p=1.332 × $10^{-43}$**<br>CLES:65.96% | U=289737.0<br>**p=1.562 × $10^{-10}$**<br>CLES:65.80% |
| Alpaca | U=2507.0<br>p=0.767<br>CLES:51.75% | U=81558.0<br>**p=2.566 × $10^{-14}$**<br>CLES:68.72% | U=13090.0<br>p=1.122 × $10^{-3}$<br>CLES:68.15% |
| Lesser Egyptian jerboa | U=54391.0<br>p=0.407<br>CLES:52.20% | U=1421932.5<br>**p=5.706 × $10^{-42}$**<br>CLES:66.14% | U=263341.0<br>**p=1.166 × $10^{-8}$**<br>CLES:64.20% |
| Long-tailed chinchilla | U=60037.5<br>p=0.212<br>CLES:53.28% | U=1651872.0<br>**p=3.032 × $10^{-49}$**<br>CLES:66.97% | U=288687.5<br>**p=4.3 × $10^{-9}$**<br>CLES:64.50% |
| Angola colobus | U=59032.0<br>p=0.834<br>CLES:49.46% | U=1711429.0<br>**p=5.155 × $10^{-56}$**<br>CLES:67.95% | U=310861.0<br>**p=8.887 × $10^{-14}$**<br>CLES:68.17% |
| Ryukyu mouse | U=78478.0<br>p=0.306<br>CLES:52.49% | U=1977309.0<br>**p=1.663 × $10^{-45}$**<br>CLES:65.36% | U=358631.5<br>**p=1.857 × $10^{-8}$**<br>CLES:62.80% |
| Chinese hamster PICR | U=71330.5<br>p=0.881<br>CLES:49.63% | U=1934751.0<br>**p=3.741 × $10^{-51}$**<br>CLES:66.53% | U=368974.0<br>**p=1.69 × $10^{-13}$**<br>CLES:66.97% |
| Vervet-AGM | U=66265.0<br>p=0.817<br>CLES:50.58% | U=1787784.5<br>**p=3.922 × $10^{-54}$**<br>CLES:67.46% | U=342227.0<br>**p=5.337 × $10^{-13}$**<br>CLES:66.93% |
| Cow | U=55490.0<br>p=0.585<br>CLES:48.57% | U=1631965.5<br>**p=8.409 × $10^{-42}$**<br>CLES:65.51% | U=298941.0<br>**p=5.069 × $10^{-12}$**<br>CLES:67.05% |
| Bushbaby | U=59169.0<br>p=0.922<br>CLES:50.25% | U=1719956.0<br>**p=3.582 × $10^{-51}$**<br>CLES:67.09% | U=306801.5<br>**p=3.878 × $10^{-12}$**<br>CLES:67.02% |

| | | | |
|---|---|---|---|
| Bonobo | U=61864.0<br>p=0.739<br>CLES:50.85% | U=1670084.5<br>**p=$5.468 \times 10^{-51}$**<br>CLES:67.12% | U=303877.5<br>**p=$3.134 \times 10^{-11}$**<br>CLES:66.03% |
| Hyrax | U=6532.5<br>p=0.588<br>CLES:52.33% | U=116382.5<br>**p=$8.211 \times 10^{-11}$**<br>CLES:64.26% | U=28994.5<br>p=$3.622 \times 10^{-3}$<br>CLES:61.51% |
| Rabbit | U=44345.0<br>p=0.838<br>CLES:49.43% | U=1274002.0<br>**p=$4.699 \times 10^{-28}$**<br>CLES:63.29% | U=227253.0<br>**p=$1.53 \times 10^{-7}$**<br>CLES:63.80% |
| Megabat | U=17950.0<br>p=0.516<br>CLES:47.78% | U=464522.0<br>**p=$5.186 \times 10^{-27}$**<br>CLES:66.73% | U=90526.0<br>**p=$4.263 \times 10^{-8}$**<br>CLES:67.62% |
| Dog | U=58150.0<br>p=0.671<br>CLES:48.91% | U=1611873.0<br>**p=$6.578 \times 10^{-43}$**<br>CLES:65.75% | U=306629.0<br>**p=$2.305 \times 10^{-12}$**<br>CLES:66.99% |
| Steppe mouse | U=79176.5<br>p=$3.153 \times 10^{-3}$<br>CLES:57.37% | U=1762520.0<br>**p=$2.781 \times 10^{-21}$**<br>CLES:60.31% | U=272899.0<br>p=0.33<br>CLES:52.28% |
| Greater bamboo lemur | U=61686.0<br>p=0.85<br>CLES:50.49% | U=1710429.0<br>**p=$1.109 \times 10^{-46}$**<br>CLES:66.24% | U=308537.0<br>**p=$7.519 \times 10^{-11}$**<br>CLES:65.76% |
| Pig | U=59990.0<br>p=0.822<br>CLES:50.58% | U=1654057.0<br>**p=$3.798 \times 10^{-42}$**<br>CLES:65.57% | U=306846.0<br>**p=$5.963 \times 10^{-10}$**<br>CLES:64.98% |
| Tree Shrew | U=2625.0<br>p=0.568<br>CLES:53.26% | U=58323.0<br>**p=$6.743 \times 10^{-9}$**<br>CLES:65.04% | U=10808.0<br>p=$2.666 \times 10^{-2}$<br>CLES:61.82% |
| Pika | U=8670.0<br>p=0.421<br>CLES:46.80% | U=174320.0<br>**p=$1.284 \times 10^{-9}$**<br>CLES:61.78% | U=41973.0<br>**p=$3.835 \times 10^{-5}$**<br>CLES:65.23% |
| Shrew | U=1352.0<br>p=0.316<br>CLES:43.61% | U=44902.0<br>**p=$2.292 \times 10^{-6}$**<br>CLES:63.53% | U=9783.0<br>p=$1.584 \times 10^{-3}$<br>CLES:68.65% |
| Squirrel | U=60531.0<br>p=0.697<br>CLES:51.00% | U=1681079.0<br>**p=$1.007 \times 10^{-51}$**<br>CLES:67.41% | U=317998.0<br>**p=$9.931 \times 10^{-12}$**<br>CLES:66.37% |
| Gorilla | U=60399.0<br>p=0.688<br>CLES:51.04% | U=1689021.0<br>**p=$1.434 \times 10^{-53}$**<br>CLES:67.60% | U=302488.5<br>**p=$2.299 \times 10^{-11}$**<br>CLES:66.29% |
| Cat | U=63516.0<br>p=0.822<br>CLES:49.43% | U=1780861.0<br>**p=$3.26 \times 10^{-50}$**<br>CLES:66.76% | U=334016.0<br>**p=$8.964 \times 10^{-13}$**<br>CLES:66.97% |

### Mouse Reference Genome, Saunders et al., 2018

| MWU Test | Vasculature vs Glia | Vasculature vs Neuron | Glia vs Neuron |
|---|---|---|---|
| Microbat | U=36956.5<br>$p=1.894 \times 10^{-2}$<br>CLES:56.05% | U=106515.0<br>**$p=4.684 \times 10^{-17}$**<br>CLES:67.44% | U=66531.0<br>**$p=4.476 \times 10^{-8}$**<br>CLES:62.98% |
| Hedgehog | U=2766.0<br>p=0.153<br>CLES:57.09% | U=6720.0<br>**$p=2.051 \times 10^{-5}$**<br>CLES:67.57% | U=4128.0<br>$p=1.1 \times 10^{-2}$<br>CLES:61.90% |
| Naked mole-rat male | U=41204.0<br>$p=2.788 \times 10^{-2}$<br>CLES:55.49% | U=128426.0<br>**$p=1.588 \times 10^{-25}$**<br>CLES:71.02% | U=82851.0<br>**$p=5.573 \times 10^{-14}$**<br>CLES:67.19% |
| Sooty mangabey | U=55933.0<br>$p=1.058 \times 10^{-3}$<br>CLES:57.63% | U=157172.0<br>**$p=7.4 \times 10^{-25}$**<br>CLES:69.60% | U=103486.0<br>**$p=1.918 \times 10^{-11}$**<br>CLES:64.29% |
| Olive baboon | U=54752.0<br>$p=2.514 \times 10^{-3}$<br>CLES:57.07% | U=157448.0<br>**$p=3.903 \times 10^{-24}$**<br>CLES:69.27% | U=103083.5<br>**$p=1.773 \times 10^{-11}$**<br>CLES:64.36% |
| American bison | U=37501.0<br>$p=1.988 \times 10^{-2}$<br>CLES:55.98% | U=106559.0<br>**$p=2.072 \times 10^{-20}$**<br>CLES:69.30% | U=66715.0<br>**$p=2.72 \times 10^{-11}$**<br>CLES:65.91% |
| Leopard | U=53505.5<br>$p=3.534 \times 10^{-3}$<br>CLES:56.86% | U=149508.0<br>**$p=3.538 \times 10^{-22}$**<br>CLES:68.56% | U=95264.0<br>**$p=1.028 \times 10^{-9}$**<br>CLES:63.21% |
| Bolivian squirrel monkey | U=52804.0<br>$p=1.956 \times 10^{-4}$<br>CLES:58.86% | U=150750.0<br>**$p=3.959 \times 10^{-28}$**<br>CLES:71.29% | U=97090.0<br>**$p=7.709 \times 10^{-12}$**<br>CLES:64.86% |
| Crab-eating macaque | U=53331.0<br>$p=5.072 \times 10^{-3}$<br>CLES:56.59% | U=155060.5<br>**$p=6.854 \times 10^{-23}$**<br>CLES:68.74% | U=100189.0<br>**$p=5.45 \times 10^{-12}$**<br>CLES:64.88% |
| Human | U=61985.0<br>$p=2.316 \times 10^{-4}$<br>CLES:58.39% | U=169687.0<br>**$p=3.351 \times 10^{-25}$**<br>CLES:69.34% | U=110640.0<br>**$p=1.095 \times 10^{-10}$**<br>CLES:63.46% |
| Red fox | U=45140.0<br>$p=2.043 \times 10^{-2}$<br>CLES:55.67% | U=135666.0<br>**$p=1.943 \times 10^{-19}$**<br>CLES:67.68% | U=85128.0<br>**$p=2.121 \times 10^{-9}$**<br>CLES:63.43% |
| Shrew mouse | U=68981.0<br>$p=2.851 \times 10^{-3}$<br>CLES:56.56% | U=179618.5<br>**$p=9.27 \times 10^{-21}$**<br>CLES:66.97% | U=118352.0<br>**$p=1.61 \times 10^{-8}$**<br>CLES:61.42% |
| Arctic ground squirrel | U=57148.0<br>$p=5.987 \times 10^{-4}$<br>CLES:57.97% | U=159711.0<br>**$p=2.593 \times 10^{-26}$**<br>CLES:70.15% | U=103654.0<br>**$p=5.745 \times 10^{-11}$**<br>CLES:63.91% |
| Chinese hamster CriGri | U=55945.5<br>$p=5.899 \times 10^{-4}$<br>CLES:58.03% | U=144770.0<br>**$p=1.169 \times 10^{-20}$**<br>CLES:67.94% | U=93433.0<br>**$p=7.054 \times 10^{-8}$**<br>CLES:61.59% |
| Opossum | U=43036.0<br>p=0.122 | U=110527.0<br>**$p=3.553 \times 10^{-8}$** | U=72955.0<br>$p=4.757 \times 10^{-4}$ |

| | | | |
|---|---|---|---|
| | CLES:53.79% | CLES:61.06% | CLES:57.89% |
| Dolphin | U=15326.0 | U=45574.0 | U=25304.0 |
| | $p=9.216\times10^{-2}$ | **$p=8.47\times10^{-14}$** | **$p=1.132\times10^{-6}$** |
| | CLES:55.43% | CLES:69.18% | CLES:64.82% |
| Capuchin | U=54956.0 | U=152985.0 | U=101834.5 |
| | **$p=5.778\times10^{-5}$** | **$p=2.005\times10^{-29}$** | **$p=4.535\times10^{-12}$** |
| | CLES:59.48% | CLES:71.79% | CLES:64.81% |
| Mouse Lemur | U=54529.0 | U=148187.0 | U=96571.5 |
| | $p=3.143\times10^{-4}$ | **$p=2.621\times10^{-23}$** | **$p=3.851\times10^{-9}$** |
| | CLES:58.48% | CLES:69.18% | CLES:62.67% |
| Damara mole rat | U=44990.5 | U=138264.0 | U=89033.0 |
| | $p=6.244\times10^{-3}$ | **$p=1.493\times10^{-23}$** | **$p=1.681\times10^{-11}$** |
| | CLES:56.72% | CLES:69.72% | CLES:65.04% |
| Marmoset | U=53136.0 | U=152082.0 | U=97825.0 |
| | $p=1.53\times10^{-4}$ | **$p=4.144\times10^{-26}$** | **$p=2.325\times10^{-10}$** |
| | CLES:59.00% | CLES:70.38% | CLES:63.70% |
| Goat | U=53225.5 | U=150939.5 | U=94563.5 |
| | $p=4.958\times10^{-4}$ | **$p=1.09\times10^{-25}$** | **$p=2.665\times10^{-10}$** |
| | CLES:58.26% | CLES:70.16% | CLES:63.75% |
| Common wombat | U=38541.0 | U=109625.0 | U=70278.0 |
| | $p=6.159\times10^{-2}$ | **$p=8.57\times10^{-14}$** | **$p=2.656\times10^{-7}$** |
| | CLES:54.73% | CLES:65.27% | CLES:62.00% |
| American black bear | U=43641.0 | U=106133.5 | U=67516.0 |
| | $p=3.026\times10^{-3}$ | **$p=4.897\times10^{-14}$** | $p=1.065\times10^{-4}$ |
| | CLES:57.35% | CLES:65.50% | CLES:58.91% |
| Sloth | U=816.0 | U=2442.0 | U=1686.0 |
| | $p=1$ | **$p=3.023\times10^{-5}$** | $p=5.956\times10^{-4}$ |
| | CLES:50.00% | CLES:72.68% | CLES:70.84% |
| Polar bear | U=41763.5 | U=117901.5 | U=74840.5 |
| | $p=1.621\times10^{-2}$ | **$p=2.889\times10^{-18}$** | **$p=3.048\times10^{-8}$** |
| | CLES:56.00% | CLES:67.67% | CLES:62.72% |
| Alpine marmot | U=47813.0 | U=136497.0 | U=87484.0 |
| | $p=2.637\times10^{-2}$ | **$p=8.413\times10^{-20}$** | **$p=2.147\times10^{-10}$** |
| | CLES:55.34% | CLES:67.79% | CLES:64.10% |
| Prairie vole | U=59214.0 | U=164247.0 | U=119529.5 |
| | $p=4.738\times10^{-2}$ | **$p=1.018\times10^{-22}$** | **$p=4.638\times10^{-15}$** |
| | CLES:54.48% | CLES:68.39% | CLES:66.13% |
| Donkey | U=56428.0 | U=148927.5 | U=96554.5 |
| | $p=1.823\times10^{-3}$ | **$p=1.689\times10^{-20}$** | **$p=4.872\times10^{-8}$** |
| | CLES:57.24% | CLES:67.74% | CLES:61.65% |
| Golden Hamster | U=54474.0 | U=149015.0 | U=103449.0 |
| | $p=4.098\times10^{-2}$ | **$p=7.68\times10^{-21}$** | **$p=2.653\times10^{-12}$** |
| | CLES:54.73% | CLES:67.92% | CLES:64.85% |
| Armadillo | U=43886.0 | U=119527.0 | U=73541.0 |
| | $p=6.157\times10^{-3}$ | **$p=2.161\times10^{-15}$** | **$p=6.681\times10^{-6}$** |
| | CLES:56.78% | CLES:65.90% | CLES:60.29% |
| Ugandan red Colobus | U=54250.0 | U=150011.5 | U=98167.0 |
| | $p=4.631\times10^{-4}$ | **$p=7.337\times10^{-25}$** | **$p=9.291\times10^{-11}$** |

| | | | |
|---|---|---|---|
| | CLES:58.25% | CLES:69.84% | CLES:63.95% |
| Pig-tailed macaque | U=56512.0<br>p=1.751 × $10^{-3}$<br>CLES:57.26% | U=159699.0<br>**p=1.145 × $10^{-25}$**<br>CLES:69.88% | U=106957.5<br>**p=5.67 × $10^{-13}$**<br>CLES:65.28% |
| Panda | U=52691.0<br>p=5.392 × $10^{-4}$<br>CLES:58.22% | U=144735.0<br>**p=8.886 × $10^{-26}$**<br>CLES:70.39% | U=91884.0<br>**p=1.97 × $10^{-11}$**<br>CLES:64.71% |
| Ferret | U=48518.5<br>p=3.263 × $10^{-3}$<br>CLES:57.11% | U=142731.5<br>**p=1.264 × $10^{-23}$**<br>CLES:69.52% | U=89198.5<br>**p=2.145 × $10^{-10}$**<br>CLES:64.08% |
| Koala | U=43959.0<br>p=0.471<br>CLES:51.74% | U=124621.0<br>**p=2.887 × $10^{-12}$**<br>CLES:63.78% | U=89966.0<br>**p=1.398 × $10^{-10}$**<br>CLES:64.13% |
| Wallaby | U=4215.0<br>p=0.388<br>CLES:46.37% | U=7913.0<br>p=9.261 × $10^{-2}$<br>CLES:56.36% | U=7572.5<br>p=7.154 × $10^{-3}$<br>CLES:60.53% |
| Tarsier | U=39439.5<br>p=3.009 × $10^{-3}$<br>CLES:57.54% | U=104832.5<br>**p=6.768 × $10^{-23}$**<br>CLES:70.78% | U=71543.5<br>**p=7.417 × $10^{-12}$**<br>CLES:65.99% |
| Degu | U=38761.0<br>p=5.903 × $10^{-3}$<br>CLES:57.02% | U=125682.0<br>**p=6.621 × $10^{-26}$**<br>CLES:71.48% | U=82178.0<br>**p=8.895 × $10^{-12}$**<br>CLES:65.65% |
| Chinese hamster CHOK1GS | U=69741.5<br>**p=1.746 × $10^{-5}$**<br>CLES:59.56% | U=183574.0<br>**p=1.778 × $10^{-26}$**<br>CLES:69.43% | U=115020.5<br>**p=7.247 × $10^{-9}$**<br>CLES:61.85% |
| Drill | U=49594.0<br>p=3.045 × $10^{-3}$<br>CLES:57.11% | U=140420.0<br>**p=5.096 × $10^{-23}$**<br>CLES:69.30% | U=91225.5<br>**p=3.242 × $10^{-11}$**<br>CLES:64.61% |
| Gibbon | U=50876.0<br>p=1.322 × $10^{-4}$<br>CLES:59.19% | U=144626.0<br>**p=4.259 × $10^{-24}$**<br>CLES:69.73% | U=91250.0<br>**p=4.176 × $10^{-8}$**<br>CLES:61.99% |
| Rat | U=63872.5<br>p=1.144 × $10^{-3}$<br>CLES:57.32% | U=167315.0<br>**p=1.541 × $10^{-21}$**<br>CLES:67.71% | U=111685.5<br>**p=5.599 × $10^{-9}$**<br>CLES:62.01% |
| Macaque | U=58107.0<br>p=1.078 × $10^{-3}$<br>CLES:57.54% | U=161257.0<br>**p=1.567 × $10^{-24}$**<br>CLES:69.32% | U=108258.0<br>**p=6.409 × $10^{-12}$**<br>CLES:64.46% |
| American mink | U=50557.5<br>p=9.658 × $10^{-3}$<br>CLES:56.16% | U=140341.5<br>**p=4.27 × $10^{-21}$**<br>CLES:68.31% | U=89471.5<br>**p=4.194 × $10^{-10}$**<br>CLES:63.74% |
| Gelada | U=57584.0<br>p=1.62 × $10^{-4}$<br>CLES:58.77% | U=160149.5<br>**p=2.893 × $10^{-26}$**<br>CLES:70.16% | U=106050.0<br>**p=5.486 × $10^{-11}$**<br>CLES:63.86% |
| Sheep | U=49845.5<br>p=1.148 × $10^{-4}$<br>CLES:59.34% | U=138294.0<br>**p=2.18 × $10^{-22}$**<br>CLES:69.08% | U=84971.0<br>**p=8.566 × $10^{-8}$**<br>CLES:61.90% |
| Tasmanian devil | U=39177.0<br>p=6.639 × $10^{-2}$ | U=104679.0<br>**p=2.615 × $10^{-12}$** | U=70786.0<br>**p=3.589 × $10^{-6}$** |

| | | | |
|---|---|---|---|
| | CLES:54.62% | CLES:64.45% | CLES:60.67% |
| Elephant | U=54463.0<br>$p=3.527 \times 10^{-4}$<br>CLES:58.41% | U=145687.0<br>**$p=1.044 \times 10^{-24}$**<br>CLES:69.89% | U=94615.0<br>**$p=1.331 \times 10^{-9}$**<br>CLES:63.09% |
| Algerian mouse | U=66485.5<br>$p=1.016 \times 10^{-2}$<br>CLES:55.62% | U=165569.5<br>**$p=5.234 \times 10^{-13}$**<br>CLES:62.92% | U=109010.0<br>$p=1.061 \times 10^{-4}$<br>CLES:57.68% |
| Dingo | U=51418.0<br>$p=3.583 \times 10^{-4}$<br>CLES:58.53% | U=140297.5<br>**$p=4.008 \times 10^{-21}$**<br>CLES:68.40% | U=89496.0<br>**$p=6.44 \times 10^{-7}$**<br>CLES:60.85% |
| Golden snub-nosed monkey | U=47784.0<br>$p=2.116 \times 10^{-2}$<br>CLES:55.55% | U=139533.0<br>**$p=3.542 \times 10^{-23}$**<br>CLES:69.39% | U=91133.5<br>**$p=4.539 \times 10^{-13}$**<br>CLES:66.03% |
| Horse | U=52887.0<br>$p=4.805 \times 10^{-3}$<br>CLES:56.65% | U=158205.0<br>**$p=1.967 \times 10^{-25}$**<br>CLES:69.83% | U=99982.0<br>**$p=2.818 \times 10^{-11}$**<br>CLES:64.36% |
| Black snub-nosed monkey | U=52387.5<br>$p=9.549 \times 10^{-4}$<br>CLES:57.84% | U=142410.5<br>**$p=1.887 \times 10^{-22}$**<br>CLES:68.94% | U=91942.5<br>**$p=2.514 \times 10^{-9}$**<br>CLES:62.98% |
| Brazilian guinea pig | U=28077.0<br>$p=1.232 \times 10^{-2}$<br>CLES:56.90% | U=74569.0<br>**$p=4.299 \times 10^{-11}$**<br>CLES:64.91% | U=50279.0<br>$p=5.749 \times 10^{-4}$<br>CLES:58.58% |
| Orangutan | U=52628.0<br>$p=4.523 \times 10^{-3}$<br>CLES:56.70% | U=150489.5<br>**$p=6.734 \times 10^{-24}$**<br>CLES:69.37% | U=98466.0<br>**$p=7.801 \times 10^{-12}$**<br>CLES:64.79% |
| Northern American deer mouse | U=65224.0<br>$p=1.387 \times 10^{-3}$<br>CLES:57.16% | U=173700.0<br>**$p=1.453 \times 10^{-25}$**<br>CLES:69.32% | U=115975.0<br>**$p=1.52 \times 10^{-12}$**<br>CLES:64.57% |
| Naked mole-rat female | U=50276.0<br>$p=7.352 \times 10^{-4}$<br>CLES:58.12% | U=150001.0<br>**$p=1.156 \times 10^{-28}$**<br>CLES:71.54% | U=93804.0<br>**$p=1.716 \times 10^{-12}$**<br>CLES:65.54% |
| Daurian ground squirrel | U=35187.0<br>p=0.11<br>CLES:54.14% | U=107662.0<br>**$p=4.228 \times 10^{-20}$**<br>CLES:69.13% | U=68372.5<br>**$p=3.881 \times 10^{-12}$**<br>CLES:66.59% |
| Tiger | U=34237.0<br>$p=1.116 \times 10^{-2}$<br>CLES:56.69% | U=101556.0<br>**$p=1.593 \times 10^{-20}$**<br>CLES:69.66% | U=61127.5<br>**$p=2.311 \times 10^{-9}$**<br>CLES:64.59% |
| Lesser hedgehog tenrec | U=3685.0<br>$p=6.515 \times 10^{-2}$<br>CLES:58.49% | U=5998.0<br>**$p=1.118 \times 10^{-5}$**<br>CLES:68.66% | U=4714.0<br>$p=1.739 \times 10^{-2}$<br>CLES:60.44% |
| Mongolian gerbil | U=55510.0<br>**$p=6.038 \times 10^{-5}$**<br>CLES:59.46% | U=133033.5<br>**$p=1.421 \times 10^{-17}$**<br>CLES:66.64% | U=79932.0<br>$p=1.348 \times 10^{-4}$<br>CLES:58.40% |
| Ma's night monkey | U=53363.0<br>**$p=3.168 \times 10^{-5}$**<br>CLES:59.90% | U=150819.0<br>**$p=1.643 \times 10^{-28}$**<br>CLES:71.52% | U=99243.0<br>**$p=8.638 \times 10^{-11}$**<br>CLES:63.97% |

| | | | |
|---|---|---|---|
| Upper Galilee mountains blind mole rat | U=56913.0<br>p=1.054 × $10^{-3}$<br>CLES:57.59% | U=157147.0<br>**p=1.502 × $10^{-23}$**<br>CLES:69.03% | U=107096.5<br>**p=6.073 × $10^{-11}$**<br>CLES:63.77% |
| Kangaroo rat | U=41611.0<br>p=0.135<br>CLES:53.68% | U=120751.0<br>**p=2.585 × $10^{-16}$**<br>CLES:66.52% | U=86698.0<br>**p=1.454 × $10^{-10}$**<br>CLES:64.28% |
| Chimpanzee | U=52189.0<br>p=6.671 × $10^{-4}$<br>CLES:58.08% | U=146754.5<br>**p=1.862 × $10^{-25}$**<br>CLES:70.27% | U=97788.0<br>**p=4.813 × $10^{-11}$**<br>CLES:64.20% |
| Guinea Pig | U=50689.0<br>p=1.433 × $10^{-2}$<br>CLES:55.81% | U=150092.0<br>**p=5.347 × $10^{-25}$**<br>CLES:69.91% | U=99391.0<br>**p=4.34 × $10^{-13}$**<br>CLES:65.69% |
| American beaver | U=50584.0<br>p=1.564 × $10^{-2}$<br>CLES:55.74% | U=122257.0<br>**p=8.834 × $10^{-18}$**<br>CLES:67.09% | U=78848.0<br>**p=9.673 × $10^{-9}$**<br>CLES:62.83% |
| Wild yak | U=39823.0<br>p=6.553 × $10^{-3}$<br>CLES:56.91% | U=117760.0<br>**p=1.734 × $10^{-22}$**<br>CLES:69.93% | U=72802.0<br>**p=8.229 × $10^{-12}$**<br>CLES:66.06% |
| Coquerel's sifaka | U=52810.0<br>p=2.034 × $10^{-4}$<br>CLES:58.84% | U=147248.0<br>**p=1.024 × $10^{-22}$**<br>CLES:68.93% | U=92854.5<br>**p=1.154 × $10^{-8}$**<br>CLES:62.42% |
| Alpaca | U=1678.0<br>p=9.197 × $10^{-2}$<br>CLES:59.59% | U=5364.0<br>**p=5.993 × $10^{-10}$**<br>CLES:78.33% | U=3203.5<br>p=5.078 × $10^{-4}$<br>CLES:68.04% |
| Lesser Egyptian jerboa | U=43705.5<br>p=0.182<br>CLES:53.24% | U=130776.5<br>**p=1.029 × $10^{-20}$**<br>CLES:68.59% | U=95596.5<br>**p=1.225 × $10^{-14}$**<br>CLES:66.92% |
| Long-tailed chinchilla | U=49789.5<br>p=2.036 × $10^{-3}$<br>CLES:57.41% | U=148400.0<br>**p=9.261 × $10^{-26}$**<br>CLES:70.34% | U=94065.5<br>**p=4.77 × $10^{-11}$**<br>CLES:64.42% |
| Angola colobus | U=53230.5<br>p=8.828 × $10^{-4}$<br>CLES:57.86% | U=148776.5<br>**p=4.484 × $10^{-26}$**<br>CLES:70.44% | U=99700.5<br>**p=2.948 × $10^{-13}$**<br>CLES:65.75% |
| Ryukyu mouse | U=69089.5<br>p=1.262 × $10^{-3}$<br>CLES:57.09% | U=180185.0<br>**p=5.246 × $10^{-22}$**<br>CLES:67.43% | U=115373.0<br>**p=7.051 × $10^{-8}$**<br>CLES:60.86% |
| Chinese hamster PICR | U=69310.0<br>**p=3.037 × $10^{-7}$**<br>CLES:61.51% | U=185378.5<br>**p=1.231 × $10^{-33}$**<br>CLES:72.23% | U=112978.0<br>**p=5.529 × $10^{-10}$**<br>CLES:62.82% |
| Vervet-AGM | U=60032.5<br>p=2.017 × $10^{-4}$<br>CLES:58.54% | U=162664.0<br>**p=2.594 × $10^{-26}$**<br>CLES:70.04% | U=106933.0<br>**p=4.942 × $10^{-11}$**<br>CLES:63.82% |
| Cow | U=54156.0<br>**p=7.368 × $10^{-5}$**<br>CLES:59.39% | U=147401.0<br>**p=7.065 × $10^{-24}$**<br>CLES:69.47% | U=93262.0<br>**p=2.193 × $10^{-8}$**<br>CLES:62.13% |
| Bushbaby | U=55896.0<br>p=1.833 × $10^{-3}$<br>CLES:57.24% | U=154676.0<br>**p=1.106 × $10^{-24}$**<br>CLES:69.62% | U=105253.0<br>**p=4.271 × $10^{-11}$**<br>CLES:63.93% |

| | | | |
|---|---|---|---|
| Bonobo | U=51907.5<br>$p=2.069 \times 10^{-3}$<br>CLES:57.31% | U=147326.0<br>$\mathbf{p=3.026 \times 10^{-25}}$<br>CLES:70.11% | U=96236.5<br>$\mathbf{p=9.541 \times 10^{-12}}$<br>CLES:64.80% |
| Hyrax | U=4870.0<br>$p=3.534 \times 10^{-2}$<br>CLES:59.11% | U=9986.0<br>$\mathbf{p=6.323 \times 10^{-7}}$<br>CLES:68.62% | U=6473.0<br>$p=4.219 \times 10^{-3}$<br>CLES:61.81% |
| Rabbit | U=40664.0<br>$p=8.446 \times 10^{-3}$<br>CLES:56.63% | U=115854.0<br>$\mathbf{p=4.782 \times 10^{-15}}$<br>CLES:65.90% | U=72785.0<br>$\mathbf{p=2.05 \times 10^{-5}}$<br>CLES:59.78% |
| Megabat | U=15400.5<br>$p=4.203 \times 10^{-3}$<br>CLES:59.29% | U=36942.0<br>$\mathbf{p=3.701 \times 10^{-12}}$<br>CLES:68.83% | U=23587.0<br>$p=1.038 \times 10^{-4}$<br>CLES:61.72% |
| Dog | U=51949.0<br>$p=1.022 \times 10^{-3}$<br>CLES:57.81% | U=146553.5<br>$\mathbf{p=2.274 \times 10^{-23}}$<br>CLES:69.27% | U=92511.0<br>$\mathbf{p=1.337 \times 10^{-8}}$<br>CLES:62.36% |
| Steppe mouse | U=58606.5<br>p=0.167<br>CLES:53.09% | U=145368.5<br>$\mathbf{p=8.243 \times 10^{-8}}$<br>CLES:59.87% | U=102400.0<br>$p=6.305 \times 10^{-4}$<br>CLES:56.92% |
| Greater bamboo lemur | U=54219.5<br>$p=2.349 \times 10^{-2}$<br>CLES:55.27% | U=151120.0<br>$\mathbf{p=2.43 \times 10^{-21}}$<br>CLES:68.09% | U=101819.0<br>$\mathbf{p=1.866 \times 10^{-12}}$<br>CLES:65.07% |
| Pig | U=56307.5<br>$p=1.525 \times 10^{-3}$<br>CLES:57.37% | U=153433.0<br>$\mathbf{p=1.681 \times 10^{-24}}$<br>CLES:69.53% | U=102119.0<br>$\mathbf{p=2.564 \times 10^{-11}}$<br>CLES:64.20% |
| Tree Shrew | U=1920.0<br>p=0.16<br>CLES:57.81% | U=4237.0<br>$p=1.363 \times 10^{-3}$<br>CLES:64.58% | U=1918.0<br>p=0.164<br>CLES:57.75% |
| Pika | U=7291.0<br>$p=4.293 \times 10^{-2}$<br>CLES:57.88% | U=13962.0<br>$\mathbf{p=1.225 \times 10^{-7}}$<br>CLES:68.10% | U=8819.0<br>$p=2.834 \times 10^{-3}$<br>CLES:61.32% |
| Shrew | U=1301.0<br>$p=2.95 \times 10^{-2}$<br>CLES:63.40% | U=3308.0<br>$p=1.656 \times 10^{-4}$<br>CLES:68.83% | U=1947.0<br>p=0.179<br>CLES:57.57% |
| Squirrel | U=52713.0<br>$p=5.506 \times 10^{-3}$<br>CLES:56.53% | U=150741.0<br>$\mathbf{p=2.671 \times 10^{-22}}$<br>CLES:68.64% | U=100182.0<br>$\mathbf{p=3.847 \times 10^{-10}}$<br>CLES:63.41% |
| Gorilla | U=53645.0<br>$p=6.742 \times 10^{-4}$<br>CLES:58.02% | U=150216.0<br>$\mathbf{p=3.425 \times 10^{-25}}$<br>CLES:70.01% | U=99550.0<br>$\mathbf{p=8.071 \times 10^{-11}}$<br>CLES:63.95% |
| Cat | U=54372.0<br>$p=4.603 \times 10^{-3}$<br>CLES:56.64% | U=156832.0<br>$\mathbf{p=2.225 \times 10^{-23}}$<br>CLES:68.90% | U=99129.0<br>$\mathbf{p=1.479 \times 10^{-10}}$<br>CLES:63.79% |

### Rat Reference Genome, Zhang et al., 2014

| MWU Test | Endothelia vs Glia | Endothelia vs Neuron | Glia vs Neuron |
|---|---|---|---|
| Microbat | U=242979.0<br>p=0.9<br>CLES:50.20% | U=349709.0<br>**$p=3.091 \times 10^{-22}$**<br>CLES:64.71% | U=440321.0<br>**$p=2.038 \times 10^{-24}$**<br>CLES:64.50% |
| Hedgehog | U=19278.0<br>p=0.314<br>CLES:52.99% | U=22646.0<br>**$p=1.042 \times 10^{-5}$**<br>CLES:63.13% | U=27364.0<br>$p=1.565 \times 10^{-4}$<br>CLES:60.60% |
| Naked mole-rat male | U=286780.0<br>p=0.434<br>CLES:51.17% | U=405301.0<br>**$p=3.243 \times 10^{-31}$**<br>CLES:67.19% | U=550378.0<br>**$p=4.551 \times 10^{-34}$**<br>CLES:66.48% |
| Sooty mangabey | U=347370.0<br>p=0.857<br>CLES:50.26% | U=518380.5<br>**$p=3.782 \times 10^{-33}$**<br>CLES:66.66% | U=686865.0<br>**$p=7.056 \times 10^{-37}$**<br>CLES:66.25% |
| Olive baboon | U=347513.5<br>p=0.586<br>CLES:49.23% | U=512309.0<br>**$p=3.073 \times 10^{-30}$**<br>CLES:65.86% | U=694250.5<br>**$p=1.851 \times 10^{-39}$**<br>CLES:66.82% |
| American bison | U=227912.0<br>p=0.999<br>CLES:50.00% | U=316289.0<br>**$p=1.419 \times 10^{-23}$**<br>CLES:65.59% | U=403202.0<br>**$p=2.378 \times 10^{-27}$**<br>CLES:65.82% |
| Leopard | U=329887.0<br>p=0.47<br>CLES:51.04% | U=479909.0<br>**$p=3.963 \times 10^{-29}$**<br>CLES:65.80% | U=617974.0<br>**$p=2.52 \times 10^{-30}$**<br>CLES:64.98% |
| Bolivian squirrel monkey | U=318871.0<br>p=0.759<br>CLES:50.45% | U=484133.0<br>**$p=4.678 \times 10^{-34}$**<br>CLES:67.21% | U=620637.0<br>**$p=2.938 \times 10^{-37}$**<br>CLES:66.79% |
| Crab-eating macaque | U=348094.0<br>p=0.988<br>CLES:49.98% | U=509126.0<br>**$p=3.654 \times 10^{-30}$**<br>CLES:65.87% | U=684732.5<br>**$p=2.511 \times 10^{-36}$**<br>CLES:66.12% |
| Human | U=363973.5<br>p=0.935<br>CLES:49.88% | U=537755.0<br>**$p=1.043 \times 10^{-30}$**<br>CLES:65.80% | U=714111.0<br>**$p=5.353 \times 10^{-37}$**<br>CLES:66.11% |
| Red fox | U=306901.0<br>p=0.675<br>CLES:50.62% | U=434913.5<br>**$p=4.242 \times 10^{-26}$**<br>CLES:65.21% | U=550742.0<br>**$p=1.216 \times 10^{-27}$**<br>CLES:64.65% |
| Shrew mouse | U=415608.5<br>p=0.404<br>CLES:48.87% | U=589993.5<br>**$p=2.472 \times 10^{-25}$**<br>CLES:63.82% | U=800271.0<br>**$p=5.264 \times 10^{-33}$**<br>CLES:64.66% |
| Arctic ground squirrel | U=345543.0<br>p=0.986<br>CLES:49.97% | U=512228.5<br>**$p=4.545 \times 10^{-34}$**<br>CLES:66.96% | U=684549.0<br>**$p=4.066 \times 10^{-41}$**<br>CLES:67.28% |
| Chinese hamster CriGri | U=342072.0<br>p=0.451<br>CLES:48.93% | U=488128.0<br>**$p=4.443 \times 10^{-29}$**<br>CLES:65.69% | U=665183.5<br>**$p=6.807 \times 10^{-37}$**<br>CLES:66.38% |
| Opossum | U=277462.0<br>p=0.111<br>CLES:52.42% | U=380032.0<br>**$p=3.589 \times 10^{-17}$**<br>CLES:62.44% | U=471947.0<br>**$p=3.119 \times 10^{-13}$**<br>CLES:60.01% |

| | | | |
|---|---|---|---|
| Dolphin | U=102795.0<br>p=0.286<br>CLES:47.97% | U=138897.0<br>**p=4.739 × $10^{-12}$**<br>CLES:63.07% | U=169585.5<br>**p=4.098 × $10^{-17}$**<br>CLES:65.21% |
| Capuchin | U=340002.0<br>p=0.804<br>CLES:50.35% | U=505541.0<br>**p=2.007 × $10^{-35}$**<br>CLES:67.40% | U=668037.0<br>**p=1.052 × $10^{-39}$**<br>CLES:67.06% |
| Mouse Lemur | U=334398.5<br>p=0.37<br>CLES:51.30% | U=482979.0<br>**p=1.313 × $10^{-32}$**<br>CLES:66.80% | U=623702.5<br>**p=1.058 × $10^{-32}$**<br>CLES:65.58% |
| Damara mole rat | U=279323.0<br>p=0.712<br>CLES:50.56% | U=420899.0<br>**p=7.028 × $10^{-33}$**<br>CLES:67.52% | U=547692.0<br>**p=1.331 × $10^{-36}$**<br>CLES:67.19% |
| Marmoset | U=343875.0<br>p=0.5<br>CLES:50.96% | U=501461.5<br>**p=8.455 × $10^{-33}$**<br>CLES:66.70% | U=653351.0<br>**p=1.035 × $10^{-33}$**<br>CLES:65.66% |
| Goat | U=323940.5<br>p=0.637<br>CLES:50.68% | U=487457.0<br>**p=8.895 × $10^{-33}$**<br>CLES:66.81% | U=616695.5<br>**p=1.848 × $10^{-34}$**<br>CLES:66.10% |
| Common wombat | U=252574.5<br>p=0.941<br>CLES:50.11% | U=361410.5<br>**p=1.346 × $10^{-18}$**<br>CLES:63.18% | U=458583.5<br>**p=6.734 × $10^{-21}$**<br>CLES:63.13% |
| American black bear | U=273593.0<br>p=0.183<br>CLES:52.03% | U=344554.0<br>**p=1.696 × $10^{-25}$**<br>CLES:65.92% | U=438643.0<br>**p=3.088 × $10^{-22}$**<br>CLES:63.75% |
| Sloth | U=5663.0<br>p=0.302<br>CLES:45.97% | U=7317.5<br>**p=9.259 × $10^{-5}$**<br>CLES:65.60% | U=9987.0<br>**p=8 × $10^{-7}$**<br>CLES:68.38% |
| Polar bear | U=259728.0<br>p=0.634<br>CLES:50.73% | U=346693.0<br>**p=6.126 × $10^{-23}$**<br>CLES:64.97% | U=432385.0<br>**p=7.53 × $10^{-23}$**<br>CLES:64.02% |
| Alpine marmot | U=317759.0<br>p=0.865<br>CLES:49.75% | U=454894.0<br>**p=6.749 × $10^{-30}$**<br>CLES:66.23% | U=599190.0<br>**p=9.58 × $10^{-36}$**<br>CLES:66.55% |
| Prairie vole | U=376485.0<br>p=0.563<br>CLES:49.20% | U=553667.0<br>**p=2.184 × $10^{-30}$**<br>CLES:65.59% | U=760873.5<br>**p=1.574 × $10^{-39}$**<br>CLES:66.43% |
| Donkey | U=333690.0<br>p=0.768<br>CLES:50.42% | U=486780.0<br>**p=1.846 × $10^{-29}$**<br>CLES:65.85% | U=644822.0<br>**p=2.903 × $10^{-32}$**<br>CLES:65.33% |
| Golden Hamster | U=342841.0<br>p=0.629<br>CLES:49.31% | U=500286.5<br>**p=7.976 × $10^{-28}$**<br>CLES:65.24% | U=686026.5<br>**p=9.618 × $10^{-36}$**<br>CLES:65.97% |
| Armadillo | U=284840.0<br>p=0.42<br>CLES:51.21% | U=391478.0<br>**p=2.548 × $10^{-26}$**<br>CLES:65.73% | U=510993.0<br>**p=8.607 × $10^{-26}$**<br>CLES:64.37% |
| Ugandan red Colobus | U=334662.0<br>p=0.991<br>CLES:49.98% | U=498698.0<br>**p=3.509 × $10^{-31}$**<br>CLES:66.26% | U=665170.0<br>**p=1.636 × $10^{-37}$**<br>CLES:66.54% |

| | | | |
|---|---|---|---|
| Pig-tailed macaque | U=352615.0<br>p=0.871<br>CLES:50.23% | U=524079.0<br>**p=3.382 × $10^{-31}$**<br>CLES:66.05% | U=685034.0<br>**p=2.064 × $10^{-35}$**<br>CLES:65.90% |
| Panda | U=330525.0<br>p=0.641<br>CLES:50.67% | U=479078.0<br>**p=9.423 × $10^{-32}$**<br>CLES:66.57% | U=618678.0<br>**p=4.007 × $10^{-35}$**<br>CLES:66.25% |
| Ferret | U=311875.0<br>p=0.702<br>CLES:49.44% | U=459938.5<br>**p=2.299 × $10^{-28}$**<br>CLES:65.73% | U=601888.0<br>**p=1.286 × $10^{-34}$**<br>CLES:66.24% |
| Koala | U=298303.0<br>p=0.512<br>CLES:50.97% | U=422195.0<br>**p=4.173 × $10^{-19}$**<br>CLES:62.86% | U=534942.0<br>**p=3.465 × $10^{-19}$**<br>CLES:62.01% |
| Wallaby | U=25725.0<br>p=0.707<br>CLES:48.98% | U=29393.5<br>p=1.946 × $10^{-2}$<br>CLES:56.37% | U=43748.0<br>p=8.058 × $10^{-4}$<br>CLES:58.27% |
| Tarsier | U=228114.0<br>p=0.83<br>CLES:49.66% | U=321792.5<br>**p=2.363 × $10^{-22}$**<br>CLES:65.08% | U=421153.0<br>**p=2.209 × $10^{-27}$**<br>CLES:65.65% |
| Degu | U=259934.0<br>p=0.371<br>CLES:51.38% | U=404885.0<br>**p=3.546 × $10^{-32}$**<br>CLES:67.56% | U=534315.5<br>**p=2.373 × $10^{-33}$**<br>CLES:66.44% |
| Chinese hamster CHOK1GS | U=421896.0<br>p=0.992<br>CLES:49.99% | U=610744.0<br>**p=6.372 × $10^{-34}$**<br>CLES:66.14% | U=816832.5<br>**p=2.596 × $10^{-39}$**<br>CLES:66.07% |
| Drill | U=318815.0<br>p=0.65<br>CLES:49.34% | U=465892.0<br>**p=1.666 × $10^{-28}$**<br>CLES:65.73% | U=626334.5<br>**p=4.785 × $10^{-36}$**<br>CLES:66.44% |
| Gibbon | U=309460.0<br>p=0.358<br>CLES:48.66% | U=453129.0<br>**p=4.144 × $10^{-25}$**<br>CLES:64.76% | U=623704.0<br>**p=3.845 × $10^{-34}$**<br>CLES:65.96% |
| Macaque | U=376712.5<br>p=0.969<br>CLES:49.95% | U=546165.0<br>**p=7.996 × $10^{-30}$**<br>CLES:65.47% | U=729032.0<br>**p=4.259 × $10^{-37}$**<br>CLES:66.04% |
| American mink | U=298836.0<br>p=0.681<br>CLES:49.40% | U=427168.5<br>**p=1.947 × $10^{-23}$**<br>CLES:64.39% | U=564972.0<br>**p=7.977 × $10^{-29}$**<br>CLES:64.90% |
| Gelada | U=353789.0<br>p=0.953<br>CLES:50.08% | U=526257.0<br>**p=8.48 × $10^{-31}$**<br>CLES:65.93% | U=700765.0<br>**p=2.154 × $10^{-36}$**<br>CLES:66.04% |
| Sheep | U=296602.0<br>p=0.753<br>CLES:50.47% | U=434792.0<br>**p=1.185 × $10^{-25}$**<br>CLES:65.10% | U=562955.5<br>**p=1.366 × $10^{-29}$**<br>CLES:65.15% |
| Tasmanian devil | U=253639.0<br>p=0.334<br>CLES:51.50% | U=337957.0<br>**p=1.045 × $10^{-18}$**<br>CLES:63.44% | U=426650.0<br>**p=1.482 × $10^{-17}$**<br>CLES:62.10% |
| Elephant | U=310630.5<br>p=0.885<br>CLES:49.79% | U=454417.0<br>**p=2.122 × $10^{-26}$**<br>CLES:65.15% | U=588502.0<br>**p=3.369 × $10^{-30}$**<br>CLES:65.14% |

| | | | |
|---|---|---|---|
| Algerian mouse | U=419547.0<br>p=0.515<br>CLES:49.12% | U=595688.0<br>**p=1.149 × $10^{-25}$**<br>CLES:63.89% | U=806711.0<br>**p=3.673 × $10^{-33}$**<br>CLES:64.67% |
| Dingo | U=320829.0<br>p=0.781<br>CLES:50.40% | U=466788.0<br>**p=7.149 × $10^{-30}$**<br>CLES:66.13% | U=599437.0<br>**p=5.331 × $10^{-32}$**<br>CLES:65.56% |
| Golden snub-nosed monkey | U=328160.0<br>p=0.609<br>CLES:49.27% | U=477123.0<br>**p=3.206 × $10^{-29}$**<br>CLES:65.82% | U=626784.5<br>**p=3.04 × $10^{-37}$**<br>CLES:66.73% |
| Horse | U=341261.0<br>p=0.639<br>CLES:50.67% | U=512657.0<br>**p=2.92 × $10^{-31}$**<br>CLES:66.17% | U=661418.5<br>**p=1.062 × $10^{-32}$**<br>CLES:65.35% |
| Black snub-nosed monkey | U=330494.5<br>p=0.913<br>CLES:50.16% | U=467772.5<br>**p=6.6 × $10^{-25}$**<br>CLES:64.55% | U=618557.0<br>**p=1.536 × $10^{-29}$**<br>CLES:64.75% |
| Brazilian guinea pig | U=172579.0<br>p=0.834<br>CLES:50.36% | U=231252.0<br>**p=2.596 × $10^{-17}$**<br>CLES:64.25% | U=325711.0<br>**p=3.722 × $10^{-19}$**<br>CLES:63.66% |
| Orangutan | U=327603.0<br>p=0.795<br>CLES:49.63% | U=483083.0<br>**p=1.393 × $10^{-29}$**<br>CLES:65.89% | U=628388.5<br>**p=1.435 × $10^{-36}$**<br>CLES:66.56% |
| Northern American deer mouse | U=389500.0<br>p=0.535<br>CLES:49.15% | U=538602.5<br>**p=1.397 × $10^{-28}$**<br>CLES:65.13% | U=739591.5<br>**p=2.207 × $10^{-37}$**<br>CLES:66.04% |
| Naked mole-rat female | U=328399.0<br>p=0.293<br>CLES:51.53% | U=472226.0<br>**p=4.915 × $10^{-34}$**<br>CLES:67.33% | U=620936.5<br>**p=5.478 × $10^{-34}$**<br>CLES:65.94% |
| Mouse | U=452305.0<br>p=0.359<br>CLES:48.79% | U=644968.5<br>**p=2.713 × $10^{-27}$**<br>CLES:64.06% | U=865846.0<br>**p=5.331 × $10^{-37}$**<br>CLES:65.30% |
| Daurian ground squirrel | U=248643.0<br>p=0.891<br>CLES:50.21% | U=346769.0<br>**p=1.87 × $10^{-26}$**<br>CLES:66.24% | U=433090.5<br>**p=7.478 × $10^{-30}$**<br>CLES:66.31% |
| Tiger | U=219347.0<br>p=0.529<br>CLES:49.00% | U=302669.0<br>**p=4.418 × $10^{-21}$**<br>CLES:64.80% | U=397734.0<br>**p=1.02 × $10^{-27}$**<br>CLES:65.99% |
| Lesser hedgehog tenrec | U=21939.0<br>p=0.198<br>CLES:53.71% | U=23930.0<br>**p=1.048 × $10^{-6}$**<br>CLES:64.38% | U=27582.0<br>**p=4.239 × $10^{-5}$**<br>CLES:61.51% |
| Mongolian gerbil | U=328749.5<br>p=0.372<br>CLES:48.72% | U=440248.5<br>**p=7.813 × $10^{-23}$**<br>CLES:64.03% | U=599421.5<br>**p=3.638 × $10^{-31}$**<br>CLES:65.32% |
| Ma's night monkey | U=337613.0<br>p=0.338<br>CLES:51.38% | U=492916.5<br>**p=1.211 × $10^{-35}$**<br>CLES:67.58% | U=642576.0<br>**p=4.687 × $10^{-35}$**<br>CLES:66.07% |
| Upper Galilee mountains blind mole rat | U=355077.0<br>p=0.848<br>CLES:50.27% | U=523080.5<br>**p=7.839 × $10^{-30}$**<br>CLES:65.69% | U=709765.5<br>**p=4.27 × $10^{-34}$**<br>CLES:65.42% |

| | | | |
|---|---|---|---|
| Kangaroo rat | U=256034.5<br>p=0.925<br>CLES:49.86% | U=392406.0<br>**p=6.27 × $10^{-25}$**<br>CLES:65.30% | U=518420.0<br>**p=2.501 × $10^{-29}$**<br>CLES:65.41% |
| Chimpanzee | U=328656.0<br>p=0.951<br>CLES:49.91% | U=496879.0<br>**p=9.197 × $10^{-33}$**<br>CLES:66.74% | U=658058.5<br>**p=5.059 × $10^{-38}$**<br>CLES:66.72% |
| Guinea Pig | U=322099.0<br>p=0.945<br>CLES:49.90% | U=468397.0<br>**p=1.033 × $10^{-29}$**<br>CLES:66.05% | U=606822.0<br>**p=7.506 × $10^{-35}$**<br>CLES:66.27% |
| American beaver | U=308734.0<br>p=0.968<br>CLES:50.06% | U=419114.5<br>**p=7.215 × $10^{-24}$**<br>CLES:64.59% | U=553245.0<br>**p=3.493 × $10^{-27}$**<br>CLES:64.49% |
| Wild yak | U=259727.0<br>p=0.75<br>CLES:49.51% | U=352218.5<br>**p=3.767 × $10^{-23}$**<br>CLES:64.97% | U=443829.5<br>**p=4.988 × $10^{-29}$**<br>CLES:65.94% |
| Coquerel's sifaka | U=306848.5<br>p=0.414<br>CLES:51.20% | U=459173.0<br>**p=1.204 × $10^{-31}$**<br>CLES:66.76% | U=584019.0<br>**p=4.025 × $10^{-32}$**<br>CLES:65.71% |
| Alpaca | U=12212.0<br>p=0.649<br>CLES:51.50% | U=17760.0<br>**p=9.167 × $10^{-8}$**<br>CLES:67.15% | U=17648.5<br>**p=2.47 × $10^{-6}$**<br>CLES:65.02% |
| Lesser Egyptian jerboa | U=283369.5<br>p=0.927<br>CLES:49.86% | U=428562.0<br>**p=3.907 × $10^{-31}$**<br>CLES:66.94% | U=581684.5<br>**p=1.897 × $10^{-37}$**<br>CLES:67.13% |
| Long-tailed chinchilla | U=311507.5<br>p=0.972<br>CLES:50.05% | U=469975.5<br>**p=5.886 × $10^{-31}$**<br>CLES:66.45% | U=615397.5<br>**p=3.353 × $10^{-36}$**<br>CLES:66.56% |
| Angola colobus | U=329684.0<br>p=0.949<br>CLES:49.91% | U=480121.0<br>**p=1.606 × $10^{-31}$**<br>CLES:66.51% | U=638941.5<br>**p=5.182 × $10^{-37}$**<br>CLES:66.59% |
| Ryukyu mouse | U=415076.0<br>p=0.733<br>CLES:49.54% | U=593329.0<br>**p=3.857 × $10^{-26}$**<br>CLES:64.04% | U=785800.5<br>**p=6.287 × $10^{-32}$**<br>CLES:64.46% |
| Chinese hamster PICR | U=416745.0<br>p=0.869<br>CLES:50.22% | U=600981.0<br>**p=8.063 × $10^{-37}$**<br>CLES:66.96% | U=798153.0<br>**p=5.597 × $10^{-41}$**<br>CLES:66.55% |
| Vervet-AGM | U=357068.5<br>p=0.989<br>CLES:49.98% | U=525307.5<br>**p=2.658 × $10^{-35}$**<br>CLES:67.17% | U=695976.0<br>**p=1.565 × $10^{-41}$**<br>CLES:67.30% |
| Cow | U=325126.0<br>p=0.833<br>CLES:49.70% | U=482725.0<br>**p=1.035 × $10^{-29}$**<br>CLES:65.93% | U=621752.0<br>**p=1.339 × $10^{-35}$**<br>CLES:66.36% |
| Bushbaby | U=342863.0<br>p=0.861<br>CLES:50.25% | U=510622.0<br>**p=2.141 × $10^{-34}$**<br>CLES:67.09% | U=688076.5<br>**p=3.737 × $10^{-39}$**<br>CLES:66.79% |
| Bonobo | U=344154.0<br>p=0.437<br>CLES:51.11% | U=485145.5<br>**p=2.461 × $10^{-33}$**<br>CLES:66.98% | U=640786.5<br>**p=3.19 × $10^{-34}$**<br>CLES:65.87% |

| | | | |
|---|---|---|---|
| Hyrax | U=25947.5<br>p=0.108<br>CLES:45.71% | U=31792.5<br>p=$1.633 \times 10^{-2}$<br>CLES:56.41% | U=47837.5<br>**p=$6.43 \times 10^{-6}$**<br>CLES:61.02% |
| Rabbit | U=274095.0<br>p=0.729<br>CLES:50.52% | U=365523.0<br>**p=$2.414 \times 10^{-16}$**<br>CLES:62.20% | U=485586.0<br>**p=$2.516 \times 10^{-17}$**<br>CLES:61.61% |
| Megabat | U=99525.5<br>p=0.588<br>CLES:48.95% | U=128892.0<br>**p=$6.64 \times 10^{-12}$**<br>CLES:63.28% | U=177501.0<br>**p=$3.863 \times 10^{-16}$**<br>CLES:64.52% |
| Dog | U=327507.0<br>p=0.678<br>CLES:50.60% | U=463657.5<br>**p=$1.304 \times 10^{-25}$**<br>CLES:64.82% | U=600697.5<br>**p=$2.434 \times 10^{-26}$**<br>CLES:63.94% |
| Steppe mouse | U=411149.0<br>p=0.777<br>CLES:49.61% | U=561649.0<br>**p=$6.753 \times 10^{-25}$**<br>CLES:63.85% | U=760959.0<br>**p=$2.502 \times 10^{-30}$**<br>CLES:64.16% |
| Greater bamboo lemur | U=343751.0<br>p=0.696<br>CLES:50.56% | U=494310.0<br>**p=$1.592 \times 10^{-31}$**<br>CLES:66.37% | U=642176.5<br>**p=$1.799 \times 10^{-34}$**<br>CLES:65.92% |
| Pig | U=332643.0<br>p=0.693<br>CLES:49.43% | U=498670.0<br>**p=$1.087 \times 10^{-30}$**<br>CLES:66.09% | U=647013.5<br>**p=$1.419 \times 10^{-36}$**<br>CLES:66.43% |
| Tree Shrew | U=16060.0<br>p=0.661<br>CLES:48.67% | U=17891.0<br>**p=$5.602 \times 10^{-5}$**<br>CLES:62.68% | U=21974.0<br>**p=$6.779 \times 10^{-6}$**<br>CLES:63.51% |
| Pika | U=40112.5<br>p=$3.151 \times 10^{-2}$<br>CLES:44.89% | U=49266.5<br>**p=$2.556 \times 10^{-5}$**<br>CLES:60.20% | U=70846.5<br>**p=$2.715 \times 10^{-11}$**<br>CLES:64.98% |
| Shrew | U=11447.0<br>p=$9.844 \times 10^{-2}$<br>CLES:55.66% | U=13868.0<br>**p=$3.535 \times 10^{-7}$**<br>CLES:67.43% | U=15199.5<br>p=$3.522 \times 10^{-4}$<br>CLES:61.66% |
| Squirrel | U=333113.5<br>p=0.525<br>CLES:50.92% | U=501909.0<br>**p=$8.07 \times 10^{-33}$**<br>CLES:66.72% | U=655732.5<br>**p=$5.985 \times 10^{-36}$**<br>CLES:66.22% |
| Gorilla | U=335645.0<br>p=0.829<br>CLES:50.31% | U=495856.5<br>**p=$6.654 \times 10^{-33}$**<br>CLES:66.79% | U=666335.0<br>**p=$2.463 \times 10^{-37}$**<br>CLES:66.49% |
| Cat | U=355636.0<br>p=0.43<br>CLES:51.12% | U=508771.0<br>**p=$1.908 \times 10^{-30}$**<br>CLES:65.94% | U=656692.0<br>**p=$5.054 \times 10^{-32}$**<br>CLES:65.19% |

### Rat Reference Genome, Zeisel et al., 2015

| MWU Test | Glia vs Vasculature | Glia vs Neuron | Vasculature vs Neuron |
|---|---|---|---|
| Microbat | U=167902.5<br>$p=5.133 \times 10^{-2}$<br>CLES:53.60% | U=372203.0<br>**$p=7.211 \times 10^{-16}$**<br>CLES:61.96% | U=121445.0<br>**$p=6.596 \times 10^{-6}$**<br>CLES:58.86% |
| Hedgehog | U=8804.0<br>p=0.319<br>CLES:53.91% | U=17881.0<br>p=0.124<br>CLES:54.75% | U=5121.0<br>p=0.851<br>CLES:50.79% |
| Naked mole-rat male | U=164586.0<br>$p=4.344 \times 10^{-2}$<br>CLES:53.75% | U=392584.0<br>**$p=2.294 \times 10^{-23}$**<br>CLES:64.71% | U=126986.5<br>**$p=1.984 \times 10^{-8}$**<br>CLES:61.05% |
| Sooty mangabey | U=224556.0<br>$p=4.559 \times 10^{-3}$<br>CLES:54.92% | U=533001.5<br>**$p=2.127 \times 10^{-25}$**<br>CLES:64.23% | U=165400.5<br>**$p=8.335 \times 10^{-8}$**<br>CLES:59.85% |
| Olive baboon | U=222854.5<br>$p=6.224 \times 10^{-3}$<br>CLES:54.75% | U=524186.0<br>**$p=7.753 \times 10^{-24}$**<br>CLES:63.79% | U=162562.0<br>**$p=2.816 \times 10^{-7}$**<br>CLES:59.45% |
| American bison | U=156052.0<br>$p=6.417 \times 10^{-3}$<br>CLES:55.19% | U=351827.0<br>**$p=6.635 \times 10^{-22}$**<br>CLES:64.65% | U=107861.5<br>**$p=1.319 \times 10^{-6}$**<br>CLES:59.85% |
| Leopard | U=201279.0<br>$p=3.76 \times 10^{-2}$<br>CLES:53.69% | U=502080.5<br>**$p=3.428 \times 10^{-23}$**<br>CLES:63.71% | U=156587.0<br>**$p=6.338 \times 10^{-8}$**<br>CLES:60.11% |
| Bolivian squirrel monkey | U=207103.0<br>$p=2.25 \times 10^{-2}$<br>CLES:54.02% | U=500046.0<br>**$p=1.292 \times 10^{-26}$**<br>CLES:64.85% | U=159643.0<br>**$p=1.539 \times 10^{-9}$**<br>CLES:61.25% |
| Crab-eating macaque | U=230166.0<br>$p=1.505 \times 10^{-3}$<br>CLES:55.49% | U=532009.0<br>**$p=5.827 \times 10^{-27}$**<br>CLES:64.74% | U=162728.0<br>**$p=1.435 \times 10^{-7}$**<br>CLES:59.68% |
| Human | U=234551.0<br>$p=5.941 \times 10^{-3}$<br>CLES:54.72% | U=566135.0<br>**$p=1.639 \times 10^{-26}$**<br>CLES:64.33% | U=176169.0<br>**$p=2.811 \times 10^{-8}$**<br>CLES:60.06% |
| Red fox | U=191464.5<br>$p=3.107 \times 10^{-2}$<br>CLES:53.88% | U=462750.0<br>**$p=2.523 \times 10^{-23}$**<br>CLES:64.08% | U=143455.0<br>**$p=2.1 \times 10^{-8}$**<br>CLES:60.71% |
| Shrew mouse | U=255916.5<br>$p=6.897 \times 10^{-3}$<br>CLES:54.54% | U=621747.0<br>**$p=5.995 \times 10^{-27}$**<br>CLES:64.12% | U=189693.0<br>**$p=1.651 \times 10^{-8}$**<br>CLES:60.04% |
| Arctic ground squirrel | U=216645.5<br>$p=3.953 \times 10^{-2}$<br>CLES:53.57% | U=539167.0<br>**$p=2.706 \times 10^{-27}$**<br>CLES:64.74% | U=175034.5<br>**$p=1.269 \times 10^{-10}$**<br>CLES:61.75% |
| Chinese hamster CriGri | U=218623.0<br>$p=1.075 \times 10^{-2}$<br>CLES:54.47% | U=509579.5<br>**$p=5.22 \times 10^{-21}$**<br>CLES:62.95% | U=154654.5<br>**$p=8.343 \times 10^{-7}$**<br>CLES:59.17% |
| Opossum | U=171366.0<br>$p=5.178 \times 10^{-2}$ | U=400820.0<br>**$p=7.346 \times 10^{-15}$** | U=127410.5<br>**$p=5.013 \times 10^{-5}$** |

| | | | |
|---|---|---|---|
| | CLES:53.59% | CLES:61.27% | CLES:57.89% |
| Dolphin | U=62715.0 | U=146833.5 | U=44026.0 |
| | $p=6.379 \times 10^{-2}$ | **$p=5.403 \times 10^{-14}$** | **$p=6.821 \times 10^{-5}$** |
| | CLES:54.44% | CLES:64.23% | CLES:60.18% |
| Capuchin | U=215910.0 | U=510914.0 | U=164185.5 |
| | $p=1.418 \times 10^{-2}$ | **$p=2 \times 10^{-25}$** | **$p=5.036 \times 10^{-9}$** |
| | CLES:54.27% | CLES:64.39% | CLES:60.78% |
| Mouse Lemur | U=207029.0 | U=498087.0 | U=156498.0 |
| | $p=1.496 \times 10^{-2}$ | **$p=1.452 \times 10^{-26}$** | **$p=1.536 \times 10^{-9}$** |
| | CLES:54.30% | CLES:64.86% | CLES:61.31% |
| Damara mole rat | U=179035.0 | U=434335.0 | U=141459.5 |
| | $p=4.093 \times 10^{-2}$ | **$p=2.632 \times 10^{-21}$** | **$p=5.843 \times 10^{-8}$** |
| | CLES:53.72% | CLES:63.56% | CLES:60.39% |
| Marmoset | U=215638.0 | U=525199.0 | U=167452.0 |
| | $p=1.485 \times 10^{-2}$ | **$p=2.819 \times 10^{-25}$** | **$p=5.016 \times 10^{-9}$** |
| | CLES:54.25% | CLES:64.22% | CLES:60.76% |
| Goat | U=203604.0 | U=492700.0 | U=148958.0 |
| | $p=3.869 \times 10^{-2}$ | **$p=5.846 \times 10^{-22}$** | **$p=1.08 \times 10^{-7}$** |
| | CLES:53.68% | CLES:63.40% | CLES:60.03% |
| Common wombat | U=157263.5 | U=378368.5 | U=122997.5 |
| | $p=7.442 \times 10^{-2}$ | **$p=9.088 \times 10^{-19}$** | **$p=7.621 \times 10^{-7}$** |
| | CLES:53.35% | CLES:63.09% | CLES:59.78% |
| American black bear | U=163590.5 | U=374764.5 | U=121307.5 |
| | p=0.181 | **$p=6.948 \times 10^{-22}$** | **$p=2.034 \times 10^{-10}$** |
| | CLES:52.48% | CLES:64.42% | CLES:62.68% |
| Sloth | U=4997.0 | U=10519.0 | U=2580.0 |
| | $p=7.703 \times 10^{-2}$ | **$p=3.844 \times 10^{-5}$** | p=0.159 |
| | CLES:58.27% | CLES:65.34% | CLES:57.23% |
| Polar bear | U=162675.0 | U=378897.0 | U=117345.0 |
| | p=0.1 | **$p=5.545 \times 10^{-20}$** | **$p=1.968 \times 10^{-8}$** |
| | CLES:53.08% | CLES:63.64% | CLES:61.26% |
| Alpine marmot | U=198945.0 | U=468221.5 | U=152444.0 |
| | $p=4.937 \times 10^{-2}$ | **$p=1.574 \times 10^{-23}$** | **$p=6.16 \times 10^{-9}$** |
| | CLES:53.48% | CLES:64.10% | CLES:60.91% |
| Prairie vole | U=230185.5 | U=566677.5 | U=170331.5 |
| | $p=1.661 \times 10^{-3}$ | **$p=9.376 \times 10^{-27}$** | **$p=1.296 \times 10^{-7}$** |
| | CLES:55.46% | CLES:64.40% | CLES:59.66% |
| Donkey | U=219419.0 | U=512468.0 | U=154824.5 |
| | $p=1.681 \times 10^{-3}$ | **$p=5.373 \times 10^{-21}$** | **$p=1.776 \times 10^{-5}$** |
| | CLES:55.51% | CLES:62.90% | CLES:57.96% |
| Golden Hamster | U=211303.0 | U=519948.5 | U=157951.0 |
| | $p=9.025 \times 10^{-3}$ | **$p=5.733 \times 10^{-22}$** | **$p=6.605 \times 10^{-7}$** |
| | CLES:54.61% | CLES:63.17% | CLES:59.25% |
| Armadillo | U=182495.0 | U=422688.5 | U=131243.0 |
| | $p=2.108 \times 10^{-2}$ | **$p=8.001 \times 10^{-20}$** | **$p=2.168 \times 10^{-6}$** |
| | CLES:54.21% | CLES:63.15% | CLES:59.18% |
| Ugandan red Colobus | U=206322.0 | U=507525.0 | U=155730.0 |
| | $p=1.073 \times 10^{-2}$ | **$p=8.01 \times 10^{-24}$** | **$p=1.326 \times 10^{-7}$** |

| | | | |
|---|---|---|---|
| | CLES:54.52% | CLES:63.89% | CLES:59.87% |
| Pig-tailed macaque | U=224409.5<br>$p=7.329 \times 10^{-3}$<br>CLES:54.64% | U=533558.0<br>**$p=6.307 \times 10^{-27}$**<br>CLES:64.70% | U=168419.0<br>**$p=7.124 \times 10^{-9}$**<br>CLES:60.61% |
| Panda | U=200163.0<br>$p=4.308 \times 10^{-2}$<br>CLES:53.60% | U=481288.0<br>**$p=3.963 \times 10^{-21}$**<br>CLES:63.17% | U=148686.0<br>**$p=1.868 \times 10^{-7}$**<br>CLES:59.84% |
| Ferret | U=201693.5<br>$p=1.94 \times 10^{-2}$<br>CLES:54.16% | U=479903.0<br>**$p=7.572 \times 10^{-21}$**<br>CLES:63.08% | U=147341.5<br>**$p=1.048 \times 10^{-6}$**<br>CLES:59.21% |
| Koala | U=191771.0<br>$p=1.215 \times 10^{-3}$<br>CLES:55.85% | U=453138.0<br>**$p=1.682 \times 10^{-21}$**<br>CLES:63.48% | U=140816.0<br>**$p=3.155 \times 10^{-5}$**<br>CLES:57.91% |
| Wallaby | U=14821.0<br>p=0.523<br>CLES:52.14% | U=32988.0<br>**$p=4.752 \times 10^{-5}$**<br>CLES:60.98% | U=11891.0<br>$p=1.145 \times 10^{-2}$<br>CLES:58.90% |
| Tarsier | U=141366.0<br>p=0.111<br>CLES:53.10% | U=335023.0<br>**$p=1.75 \times 10^{-16}$**<br>CLES:62.60% | U=100557.0<br>**$p=4.979 \times 10^{-6}$**<br>CLES:59.48% |
| Degu | U=157011.5<br>p=0.117<br>CLES:52.94% | U=374331.5<br>**$p=9.191 \times 10^{-18}$**<br>CLES:62.71% | U=124087.5<br>**$p=6.074 \times 10^{-7}$**<br>CLES:59.84% |
| Chinese hamster CHOK1GS | U=253566.5<br>$p=1.508 \times 10^{-2}$<br>CLES:54.09% | U=623284.0<br>**$p=1.955 \times 10^{-27}$**<br>CLES:64.26% | U=192000.0<br>**$p=1.806 \times 10^{-9}$**<br>CLES:60.70% |
| Drill | U=215488.0<br>$p=8.399 \times 10^{-4}$<br>CLES:55.89% | U=491990.5<br>**$p=3.324 \times 10^{-24}$**<br>CLES:64.16% | U=148095.0<br>**$p=3.065 \times 10^{-6}$**<br>CLES:58.76% |
| Gibbon | U=210912.0<br>$p=1.058 \times 10^{-3}$<br>CLES:55.79% | U=492633.0<br>**$p=8.636 \times 10^{-27}$**<br>CLES:64.97% | U=152514.5<br>**$p=2.394 \times 10^{-7}$**<br>CLES:59.67% |
| Macaque | U=236099.0<br>$p=9.646 \times 10^{-4}$<br>CLES:55.67% | U=553847.0<br>**$p=3.207 \times 10^{-24}$**<br>CLES:63.69% | U=170399.0<br>**$p=6.566 \times 10^{-6}$**<br>CLES:58.17% |
| American mink | U=184442.5<br>$p=1.973 \times 10^{-2}$<br>CLES:54.25% | U=451237.0<br>**$p=2.793 \times 10^{-23}$**<br>CLES:64.15% | U=139063.0<br>**$p=9.959 \times 10^{-8}$**<br>CLES:60.25% |
| Gelada | U=226913.0<br>$p=7.091 \times 10^{-3}$<br>CLES:54.66% | U=540779.0<br>**$p=2.051 \times 10^{-25}$**<br>CLES:64.18% | U=167418.0<br>**$p=5.97 \times 10^{-8}$**<br>CLES:59.93% |
| Sheep | U=189817.0<br>$p=3.742 \times 10^{-2}$<br>CLES:53.78% | U=461622.0<br>**$p=2.85 \times 10^{-20}$**<br>CLES:63.02% | U=137013.0<br>**$p=7.286 \times 10^{-7}$**<br>CLES:59.54% |
| Tasmanian devil | U=153937.0<br>$p=1.646 \times 10^{-2}$<br>CLES:54.56% | U=347605.0<br>**$p=2.771 \times 10^{-15}$**<br>CLES:61.91% | U=107923.0<br>$p=3.854 \times 10^{-4}$<br>CLES:57.16% |
| Elephant | U=193811.5<br>p=0.17 | U=467158.0<br>**$p=4.97 \times 10^{-18}$** | U=145375.0<br>**$p=1.297 \times 10^{-7}$** |

| | | | |
|---|---|---|---|
| | CLES:52.45% | CLES:62.14% | CLES:60.03% |
| Algerian mouse | U=260177.0<br>$p=8.676 \times 10^{-3}$<br>CLES:54.39% | U=643501.0<br>**$p=4.797 \times 10^{-30}$**<br>CLES:64.88% | U=195470.0<br>**$p=1.41 \times 10^{-9}$**<br>CLES:60.72% |
| Dingo | U=192498.0<br>$p=3.414 \times 10^{-2}$<br>CLES:53.81% | U=479125.5<br>**$p=4.776 \times 10^{-25}$**<br>CLES:64.51% | U=148545.0<br>**$p=4.143 \times 10^{-9}$**<br>CLES:61.17% |
| Golden snub-nosed monkey | U=207431.5<br>$p=5.389 \times 10^{-3}$<br>CLES:54.92% | U=488581.5<br>**$p=6.794 \times 10^{-27}$**<br>CLES:65.04% | U=153799.5<br>**$p=1.211 \times 10^{-8}$**<br>CLES:60.68% |
| Horse | U=221232.0<br>$p=4.112 \times 10^{-3}$<br>CLES:55.00% | U=519512.0<br>**$p=1.665 \times 10^{-23}$**<br>CLES:63.71% | U=161744.0<br>**$p=2.203 \times 10^{-7}$**<br>CLES:59.56% |
| Black snub-nosed monkey | U=204725.0<br>$p=1.522 \times 10^{-2}$<br>CLES:54.30% | U=491510.0<br>**$p=7.362 \times 10^{-25}$**<br>CLES:64.37% | U=152675.5<br>**$p=2.352 \times 10^{-8}$**<br>CLES:60.49% |
| Brazilian guinea pig | U=110876.0<br>$p=0.3$<br>CLES:52.11% | U=245932.0<br>**$p=5.885 \times 10^{-11}$**<br>CLES:60.72% | U=81109.0<br>$p=1.006 \times 10^{-4}$<br>CLES:58.44% |
| Orangutan | U=210187.0<br>$p=3.579 \times 10^{-2}$<br>CLES:53.67% | U=494480.0<br>**$p=1.171 \times 10^{-24}$**<br>CLES:64.27% | U=160258.0<br>**$p=4.177 \times 10^{-9}$**<br>CLES:60.90% |
| Northern American deer mouse | U=236956.5<br>$p=3.967 \times 10^{-3}$<br>CLES:54.94% | U=563962.5<br>**$p=1.392 \times 10^{-29}$**<br>CLES:65.30% | U=172950.0<br>**$p=1.213 \times 10^{-9}$**<br>CLES:61.09% |
| Naked mole-rat female | U=182373.0<br>$p=3.19 \times 10^{-2}$<br>CLES:53.90% | U=455684.5<br>**$p=8.943 \times 10^{-26}$**<br>CLES:64.93% | U=144289.0<br>**$p=4.228 \times 10^{-9}$**<br>CLES:61.25% |
| Mouse | U=277021.0<br>$p=8.729 \times 10^{-3}$<br>CLES:54.32% | U=672987.5<br>**$p=1.155 \times 10^{-28}$**<br>CLES:64.32% | U=205932.0<br>**$p=3.448 \times 10^{-9}$**<br>CLES:60.30% |
| Daurian ground squirrel | U=165525.0<br>$p=2.705 \times 10^{-2}$<br>CLES:54.11% | U=356967.0<br>**$p=2.759 \times 10^{-22}$**<br>CLES:64.77% | U=115127.0<br>**$p=2.027 \times 10^{-8}$**<br>CLES:61.24% |
| Tiger | U=140945.0<br>$p=9.73 \times 10^{-2}$<br>CLES:53.22% | U=332276.0<br>**$p=5.036 \times 10^{-19}$**<br>CLES:63.72% | U=101301.0<br>**$p=2.322 \times 10^{-7}$**<br>CLES:60.75% |
| Lesser hedgehog tenrec | U=11344.0<br>$p=0.711$<br>CLES:51.33% | U=22581.0<br>$p=7.566 \times 10^{-4}$<br>CLES:60.07% | U=7656.0<br>$p=1.554 \times 10^{-2}$<br>CLES:59.46% |
| Mongolian gerbil | U=210681.0<br>$p=2.764 \times 10^{-4}$<br>CLES:56.50% | U=472143.0<br>**$p=8.884 \times 10^{-23}$**<br>CLES:63.86% | U=136169.5<br>**$p=2.553 \times 10^{-5}$**<br>CLES:58.04% |
| Ma's night monkey | U=204837.0<br>$p=3.218 \times 10^{-2}$<br>CLES:53.78% | U=501578.5<br>**$p=6.022 \times 10^{-27}$**<br>CLES:64.94% | U=160964.5<br>**$p=4.907 \times 10^{-10}$**<br>CLES:61.59% |

| | | | |
|---|---|---|---|
| Upper Galilee mountains blind mole rat | U=209748.0<br>p=2.861 × $10^{-2}$<br>CLES:53.86% | U=522086.0<br>**p=1.262 × $10^{-24}$**<br>CLES:64.05% | U=159275.5<br>**p=2.499 × $10^{-8}$**<br>CLES:60.39% |
| Kangaroo rat | U=161453.5<br>p=7.04 × $10^{-2}$<br>CLES:53.39% | U=401008.0<br>**p=4.317 × $10^{-21}$**<br>CLES:63.79% | U=127089.0<br>**p=4.043 × $10^{-8}$**<br>CLES:60.84% |
| Chimpanzee | U=208378.0<br>p=7.192 × $10^{-4}$<br>CLES:56.01% | U=494987.5<br>**p=2.833 × $10^{-28}$**<br>CLES:65.41% | U=152431.0<br>**p=6.599 × $10^{-8}$**<br>CLES:60.14% |
| Guinea Pig | U=194996.0<br>p=3.933 × $10^{-2}$<br>CLES:53.68% | U=475014.0<br>**p=1.878 × $10^{-22}$**<br>CLES:63.65% | U=152514.0<br>**p=4.734 × $10^{-8}$**<br>CLES:60.27% |
| American beaver | U=178175.0<br>p=0.149<br>CLES:52.61% | U=404013.0<br>**p=5.879 × $10^{-19}$**<br>CLES:62.99% | U=132430.0<br>**p=6.187 × $10^{-8}$**<br>CLES:60.48% |
| Wild yak | U=172674.0<br>p=5.897 × $10^{-3}$<br>CLES:55.13% | U=375332.0<br>**p=1.675 × $10^{-18}$**<br>CLES:63.10% | U=111208.0<br>**p=3.656 × $10^{-5}$**<br>CLES:58.27% |
| Coquerel's sifaka | U=191124.0<br>p=7.692 × $10^{-2}$<br>CLES:53.17% | U=461310.0<br>**p=1.683 × $10^{-20}$**<br>CLES:63.08% | U=148664.0<br>**p=4.56 × $10^{-8}$**<br>CLES:60.34% |
| Alpaca | U=9833.0<br>p=1.488 × $10^{-2}$<br>CLES:59.63% | U=24925.0<br>**p=4.761 × $10^{-12}$**<br>CLES:70.94% | U=6540.0<br>p=6.087 × $10^{-4}$<br>CLES:64.48% |
| Lesser Egyptian jerboa | U=179565.0<br>p=6.914 × $10^{-3}$<br>CLES:54.98% | U=451778.5<br>**p=2.529 × $10^{-24}$**<br>CLES:64.49% | U=135288.0<br>**p=2.125 × $10^{-7}$**<br>CLES:60.08% |
| Long-tailed chinchilla | U=200400.0<br>p=7.098 × $10^{-3}$<br>CLES:54.79% | U=483026.5<br>**p=5.126 × $10^{-24}$**<br>CLES:64.12% | U=152727.0<br>**p=1.54 × $10^{-7}$**<br>CLES:59.85% |
| Angola colobus | U=212235.0<br>p=7.064 × $10^{-3}$<br>CLES:54.74% | U=500542.5<br>**p=1.265 × $10^{-25}$**<br>CLES:64.55% | U=154727.0<br>**p=7.829 × $10^{-8}$**<br>CLES:60.03% |
| Ryukyu mouse | U=250795.0<br>p=2.357 × $10^{-2}$<br>CLES:53.81% | U=615559.5<br>**p=1.304 × $10^{-25}$**<br>CLES:63.75% | U=191849.0<br>**p=2.735 × $10^{-9}$**<br>CLES:60.58% |
| Chinese hamster PICR | U=245305.0<br>p=7.339 × $10^{-3}$<br>CLES:54.56% | U=611937.0<br>**p=8.19 × $10^{-28}$**<br>CLES:64.43% | U=187351.0<br>**p=1.888 × $10^{-9}$**<br>CLES:60.77% |
| Vervet-AGM | U=221726.0<br>p=1.18 × $10^{-2}$<br>CLES:54.38% | U=542811.0<br>**p=5.263 × $10^{-28}$**<br>CLES:64.97% | U=166771.5<br>**p=1.846 × $10^{-9}$**<br>CLES:61.08% |
| Cow | U=209821.0<br>p=1.319 × $10^{-2}$<br>CLES:54.38% | U=501664.0<br>**p=1.632 × $10^{-24}$**<br>CLES:64.19% | U=151474.0<br>**p=9.793 × $10^{-8}$**<br>CLES:60.01% |
| Bushbaby | U=211854.0<br>p=3.277 × $10^{-2}$<br>CLES:53.74% | U=517804.0<br>**p=8.376 × $10^{-27}$**<br>CLES:64.78% | U=163646.0<br>**p=4.167 × $10^{-10}$**<br>CLES:61.59% |

| | | | |
|---|---|---|---|
| Bonobo | U=212264.0<br>$p=1.973\times10^{-2}$<br>CLES:54.10% | U=499272.0<br>**$p=2.452\times10^{-24}$**<br>CLES:64.16% | U=154667.5<br>**$p=1.892\times10^{-8}$**<br>CLES:60.51% |
| Hyrax | U=18169.0<br>p=0.157<br>CLES:54.64% | U=38703.0<br>**$p=3.184\times10^{-5}$**<br>CLES:60.88% | U=11210.0<br>$p=7.198\times10^{-2}$<br>CLES:56.36% |
| Rabbit | U=177036.0<br>$p=1.091\times10^{-2}$<br>CLES:54.69% | U=407538.0<br>**$p=5.092\times10^{-18}$**<br>CLES:62.56% | U=126403.0<br>**$p=1.709\times10^{-5}$**<br>CLES:58.39% |
| Megabat | U=68394.5<br>p=0.421<br>CLES:51.84% | U=134727.5<br>**$p=4.977\times10^{-9}$**<br>CLES:61.21% | U=46916.5<br>**$p=8.841\times10^{-5}$**<br>CLES:59.73% |
| Dog | U=199712.0<br>$p=7.325\times10^{-2}$<br>CLES:53.17% | U=477552.0<br>**$p=6.002\times10^{-22}$**<br>CLES:63.48% | U=153671.0<br>**$p=6.707\times10^{-9}$**<br>CLES:60.88% |
| Steppe mouse | U=256029.0<br>$p=6.183\times10^{-4}$<br>CLES:55.78% | U=603165.0<br>**$p=5.568\times10^{-28}$**<br>CLES:64.54% | U=183371.0<br>**$p=2.452\times10^{-7}$**<br>CLES:59.22% |
| Greater bamboo lemur | U=214467.0<br>$p=8.034\times10^{-3}$<br>CLES:54.66% | U=513331.5<br>**$p=1.151\times10^{-23}$**<br>CLES:63.81% | U=156543.5<br>**$p=2.715\times10^{-7}$**<br>CLES:59.57% |
| Pig | U=221628.0<br>$p=1.218\times10^{-3}$<br>CLES:55.66% | U=525902.0<br>**$p=5.163\times10^{-27}$**<br>CLES:64.78% | U=160595.0<br>**$p=1.241\times10^{-7}$**<br>CLES:59.79% |
| Tree Shrew | U=8341.0<br>$p=6.92\times10^{-2}$<br>CLES:57.29% | U=17572.0<br>$p=1.698\times10^{-3}$<br>CLES:59.91% | U=5067.0<br>p=0.507<br>CLES:52.83% |
| Pika | U=29976.0<br>$p=4.515\times10^{-2}$<br>CLES:55.72% | U=50006.0<br>**$p=1.863\times10^{-6}$**<br>CLES:61.83% | U=16295.5<br>$p=5.422\times10^{-2}$<br>CLES:56.09% |
| Shrew | U=5378.0<br>p=0.217<br>CLES:55.60% | U=13861.5<br>$p=2\times10^{-4}$<br>CLES:62.63% | U=3620.0<br>$p=7.765\times10^{-2}$<br>CLES:58.50% |
| Squirrel | U=215628.0<br>$p=5.404\times10^{-3}$<br>CLES:54.86% | U=521696.5<br>**$p=1.559\times10^{-27}$**<br>CLES:64.95% | U=166915.0<br>**$p=7.3\times10^{-9}$**<br>CLES:60.64% |
| Gorilla | U=210480.0<br>$p=1.001\times10^{-2}$<br>CLES:54.54% | U=503958.0<br>**$p=6.737\times10^{-24}$**<br>CLES:63.95% | U=156300.5<br>**$p=7.021\times10^{-8}$**<br>CLES:60.06% |
| Cat | U=218572.0<br>$p=4.95\times10^{-2}$<br>CLES:53.42% | U=540937.0<br>**$p=7.266\times10^{-25}$**<br>CLES:64.00% | U=168047.0<br>**$p=5.339\times10^{-9}$**<br>CLES:60.73% |

### Rat Reference Genome, Zeisel et al., 2018

| MWU Test | CNS_Glia vs CNS_Endothelia | CNS_Glia vs CNS_Neuron | CNS_Endothelia vs CNS_Neuron |
|---|---|---|---|
| Microbat | U=33295.0<br>p=0.339<br>CLES:47.14% | U=1109348.5<br>**p=2.914 × $10^{-32}$**<br>CLES:64.97% | U=193613.0<br>**p=6.863 × $10^{-10}$**<br>CLES:67.41% |
| Hedgehog | U=2447.0<br>p=0.874<br>CLES:49.11% | U=52972.0<br>**p=4.327 × $10^{-5}$**<br>CLES:60.80% | U=11939.0<br>p=1.406 × $10^{-2}$<br>CLES:62.70% |
| Naked mole-rat male | U=43815.0<br>p=0.908<br>CLES:50.32% | U=1202883.5<br>**p=6.312 × $10^{-41}$**<br>CLES:66.54% | U=219086.0<br>**p=4.024 × $10^{-10}$**<br>CLES:66.47% |
| Sooty mangabey | U=56314.0<br>p=0.45<br>CLES:52.01% | U=1566168.0<br>**p=4.669 × $10^{-46}$**<br>CLES:66.53% | U=273490.0<br>**p=1.62 × $10^{-9}$**<br>CLES:65.11% |
| Olive baboon | U=55949.0<br>p=0.614<br>CLES:51.33% | U=1557025.0<br>**p=6.448 × $10^{-47}$**<br>CLES:66.74% | U=279279.0<br>**p=3.323 × $10^{-10}$**<br>CLES:65.63% |
| American bison | U=34716.0<br>p=0.511<br>CLES:48.06% | U=1041457.0<br>**p=1.836 × $10^{-34}$**<br>CLES:65.64% | U=186479.0<br>**p=2.32 × $10^{-10}$**<br>CLES:67.68% |
| Leopard | U=49977.0<br>p=0.676<br>CLES:48.86% | U=1511849.5<br>**p=8.226 × $10^{-43}$**<br>CLES:66.02% | U=264590.0<br>**p=5.144 × $10^{-11}$**<br>CLES:66.80% |
| Bolivian squirrel monkey | U=47265.0<br>p=0.273<br>CLES:47.03% | U=1465388.5<br>**p=5.325 × $10^{-48}$**<br>CLES:67.21% | U=272926.0<br>**p=3.69 × $10^{-15}$**<br>CLES:70.06% |
| Crab-eating macaque | U=56323.0<br>p=0.535<br>CLES:51.64% | U=1557321.5<br>**p=2.373 × $10^{-48}$**<br>CLES:66.97% | U=274281.0<br>**p=3.814 × $10^{-10}$**<br>CLES:65.63% |
| Human | U=58190.0<br>p=0.835<br>CLES:50.54% | U=1650334.5<br>**p=2.872 × $10^{-49}$**<br>CLES:66.94% | U=299305.0<br>**p=1.845 × $10^{-11}$**<br>CLES:66.48% |
| Red fox | U=45915.0<br>p=0.822<br>CLES:49.37% | U=1392205.5<br>**p=5.087 × $10^{-38}$**<br>CLES:65.31% | U=236934.0<br>**p=9.052 × $10^{-10}$**<br>CLES:66.16% |
| Shrew mouse | U=65032.0<br>p=0.374<br>CLES:52.29% | U=1781383.5<br>**p=3.565 × $10^{-46}$**<br>CLES:65.93% | U=298820.0<br>**p=2.114 × $10^{-8}$**<br>CLES:63.64% |
| Arctic ground squirrel | U=54450.0<br>p=0.819<br>CLES:50.61% | U=1549044.0<br>**p=8.531 × $10^{-45}$**<br>CLES:66.34% | U=275338.0<br>**p=4.084 × $10^{-10}$**<br>CLES:65.66% |
| Chinese hamster CriGri | U=52630.5<br>p=0.767<br>CLES:49.21% | U=1488271.5<br>**p=5.196 × $10^{-39}$**<br>CLES:65.23% | U=266085.5<br>**p=7.205 × $10^{-10}$**<br>CLES:65.49% |
| Opossum | U=46004.0<br>p=0.394 | U=1169830.5<br>**p=8.717 × $10^{-21}$** | U=201104.0<br>p=1.332 × $10^{-3}$ |

| | | | |
|---|---|---|---|
| | CLES:52.38% | CLES:61.46% | CLES:58.44% |
| Dolphin | U=17994.0<br>p=0.824<br>CLES:50.77% | U=429080.0<br>**$p=4.309 \times 10^{-18}$**<br>CLES:63.75% | U=85996.0<br>**$p=7.914 \times 10^{-5}$**<br>CLES:62.68% |
| Capuchin | U=51010.0<br>p=0.514<br>CLES:48.26% | U=1516146.0<br>**$p=9.732 \times 10^{-47}$**<br>CLES:66.80% | U=280956.0<br>**$p=9.937 \times 10^{-14}$**<br>CLES:68.71% |
| Mouse Lemur | U=51560.0<br>p=0.611<br>CLES:51.37% | U=1434371.5<br>**$p=2.836 \times 10^{-43}$**<br>CLES:66.40% | U=261057.0<br>**$p=7.662 \times 10^{-10}$**<br>CLES:65.58% |
| Damara mole rat | U=43226.0<br>p=0.516<br>CLES:51.85% | U=1288405.0<br>**$p=1.187 \times 10^{-40}$**<br>CLES:66.38% | U=221461.0<br>**$p=2.333 \times 10^{-8}$**<br>CLES:64.97% |
| Marmoset | U=53161.0<br>p=0.96<br>CLES:50.13% | U=1518969.0<br>**$p=1.219 \times 10^{-42}$**<br>CLES:65.94% | U=268743.0<br>**$p=2.822 \times 10^{-10}$**<br>CLES:65.91% |
| Goat | U=47503.0<br>p=0.762<br>CLES:49.16% | U=1431157.0<br>**$p=2.175 \times 10^{-37}$**<br>CLES:65.10% | U=249452.0<br>**$p=9.594 \times 10^{-10}$**<br>CLES:65.89% |
| Common wombat | U=39456.0<br>p=0.654<br>CLES:51.30% | U=1106926.0<br>**$p=1.068 \times 10^{-24}$**<br>CLES:62.86% | U=188951.0<br>**$p=3.527 \times 10^{-5}$**<br>CLES:61.33% |
| American black bear | U=39346.0<br>p=0.644<br>CLES:48.68% | U=1111901.0<br>**$p=5.382 \times 10^{-29}$**<br>CLES:64.00% | U=205736.0<br>**$p=1.038 \times 10^{-8}$**<br>CLES:65.37% |
| Sloth | U=925.0<br>p=0.94<br>CLES:49.41% | U=27923.0<br>**$p=2.225 \times 10^{-7}$**<br>CLES:66.46% | U=4764.0<br>$p=2.595 \times 10^{-2}$<br>CLES:65.51% |
| Polar bear | U=35582.0<br>p=0.16<br>CLES:45.92% | U=1146462.0<br>**$p=4.114 \times 10^{-34}$**<br>CLES:65.23% | U=208126.0<br>**$p=3.094 \times 10^{-11}$**<br>CLES:68.19% |
| Alpine marmot | U=46501.0<br>p=0.411<br>CLES:47.77% | U=1379670.0<br>**$p=1.829 \times 10^{-42}$**<br>CLES:66.42% | U=265575.0<br>**$p=6.441 \times 10^{-13}$**<br>CLES:68.28% |
| Prairie vole | U=58952.0<br>p=0.716<br>CLES:50.95% | U=1625452.5<br>**$p=1.275 \times 10^{-43}$**<br>CLES:65.88% | U=288228.0<br>**$p=5.745 \times 10^{-9}$**<br>CLES:64.29% |
| Donkey | U=48165.0<br>p=0.802<br>CLES:50.70% | U=1496619.0<br>**$p=2.588 \times 10^{-38}$**<br>CLES:65.16% | U=244950.0<br>**$p=1.771 \times 10^{-8}$**<br>CLES:64.84% |
| Golden Hamster | U=53871.5<br>p=0.953<br>CLES:49.84% | U=1469565.0<br>**$p=1.7 \times 10^{-37}$**<br>CLES:64.95% | U=265825.0<br>**$p=3.258 \times 10^{-9}$**<br>CLES:64.78% |
| Armadillo | U=40057.0<br>p=0.867<br>CLES:50.49% | U=1188362.0<br>**$p=1.47 \times 10^{-28}$**<br>CLES:63.69% | U=202436.0<br>**$p=3.321 \times 10^{-7}$**<br>CLES:63.97% |
| Ugandan red Colobus | U=51934.0<br>p=0.927 | U=1475314.0<br>**$p=6.254 \times 10^{-43}$** | U=265007.0<br>**$p=9.65 \times 10^{-11}$** |

| | | | |
|---|---|---|---|
| | CLES:50.25% | CLES:66.14% | CLES:66.39% |
| Pig-tailed macaque | U=56355.0<br>p=0.746<br>CLES:50.85% | U=1563236.5<br>**p=$1.24 \times 10^{-47}$**<br>CLES:66.86% | U=285786.0<br>**p=$4.935 \times 10^{-11}$**<br>CLES:66.24% |
| Panda | U=47771.0<br>p=0.591<br>CLES:48.54% | U=1436838.5<br>**p=$6.976 \times 10^{-40}$**<br>CLES:65.69% | U=264004.0<br>**p=$2.06 \times 10^{-11}$**<br>CLES:67.15% |
| Ferret | U=47351.0<br>p=0.944<br>CLES:50.19% | U=1422419.0<br>**p=$2.561 \times 10^{-39}$**<br>CLES:65.60% | U=246816.5<br>**p=$2.492 \times 10^{-9}$**<br>CLES:65.54% |
| Koala | U=51568.0<br>p=0.133<br>CLES:54.11% | U=1323537.0<br>**p=$4.517 \times 10^{-28}$**<br>CLES:63.19% | U=224534.0<br>p=$7.776 \times 10^{-4}$<br>CLES:58.64% |
| Wallaby | U=5142.0<br>p=0.516<br>CLES:47.09% | U=93981.0<br>p=$1.89 \times 10^{-2}$<br>CLES:55.25% | U=24195.0<br>p=$7.171 \times 10^{-2}$<br>CLES:57.44% |
| Tarsier | U=36939.0<br>p=0.897<br>CLES:49.62% | U=1068432.0<br>**p=$3.382 \times 10^{-37}$**<br>CLES:66.19% | U=189220.0<br>**p=$5.692 \times 10^{-10}$**<br>CLES:67.15% |
| Degu | U=43576.0<br>p=0.278<br>CLES:53.06% | U=1138234.0<br>**p=$9.399 \times 10^{-37}$**<br>CLES:65.95% | U=209075.0<br>**p=$2.321 \times 10^{-7}$**<br>CLES:63.67% |
| Chinese hamster CHOK1GS | U=62422.0<br>p=0.866<br>CLES:49.57% | U=1752000.5<br>**p=$2.895 \times 10^{-45}$**<br>CLES:65.85% | U=318584.0<br>**p=$1.921 \times 10^{-11}$**<br>CLES:66.14% |
| Drill | U=50673.0<br>p=0.794<br>CLES:50.71% | U=1468556.5<br>**p=$1.253 \times 10^{-44}$**<br>CLES:66.51% | U=256099.0<br>**p=$1.305 \times 10^{-10}$**<br>CLES:66.51% |
| Gibbon | U=52029.0<br>p=0.514<br>CLES:51.77% | U=1465185.5<br>**p=$8.409 \times 10^{-49}$**<br>CLES:67.36% | U=256782.0<br>**p=$2.929 \times 10^{-10}$**<br>CLES:66.08% |
| Macaque | U=58616.0<br>p=0.7<br>CLES:51.01% | U=1609097.0<br>**p=$4.204 \times 10^{-42}$**<br>CLES:65.60% | U=284183.0<br>**p=$8.017 \times 10^{-9}$**<br>CLES:64.20% |
| American mink | U=46694.5<br>p=0.949<br>CLES:50.18% | U=1372540.5<br>**p=$1.934 \times 10^{-41}$**<br>CLES:66.17% | U=240318.5<br>**p=$1.364 \times 10^{-9}$**<br>CLES:65.81% |
| Gelada | U=55056.0<br>p=0.597<br>CLES:51.41% | U=1568286.0<br>**p=$9.572 \times 10^{-49}$**<br>CLES:67.05% | U=274710.0<br>**p=$1.72 \times 10^{-10}$**<br>CLES:66.05% |
| Sheep | U=44537.0<br>p=0.493<br>CLES:48.08% | U=1377326.0<br>**p=$6.995 \times 10^{-34}$**<br>CLES:64.43% | U=238608.5<br>**p=$8.88 \times 10^{-10}$**<br>CLES:66.17% |
| Tasmanian devil | U=42315.0<br>p=0.116<br>CLES:54.55% | U=1057478.0<br>**p=$4.588 \times 10^{-19}$**<br>CLES:61.20% | U=170965.0<br>p=$2.606 \times 10^{-2}$<br>CLES:56.07% |
| Elephant | U=44845.0<br>p=0.646 | U=1355932.0<br>**p=$3.634 \times 10^{-34}$** | U=247721.0<br>**p=$3.629 \times 10^{-10}$** |

| | | | |
|---|---|---|---|
| | CLES:48.73% | CLES:64.65% | CLES:66.35% |
| Algerian mouse | U=64339.0<br>p=0.731<br>CLES:50.88% | U=1810896.5<br>**p=5.444 × $10^{-49}$**<br>CLES:66.42% | U=316336.5<br>**p=2.825 × $10^{-10}$**<br>CLES:65.21% |
| Dingo | U=47715.0<br>p=0.792<br>CLES:49.28% | U=1394022.5<br>**p=6.195 × $10^{-37}$**<br>CLES:65.18% | U=256315.0<br>**p=1.11 × $10^{-9}$**<br>CLES:65.59% |
| Golden snub-nosed monkey | U=57012.0<br>p=0.44<br>CLES:52.03% | U=1463226.5<br>**p=3.159 × $10^{-48}$**<br>CLES:67.18% | U=271046.0<br>**p=1.612 × $10^{-10}$**<br>CLES:65.77% |
| Horse | U=52028.0<br>p=0.694<br>CLES:51.07% | U=1532811.5<br>**p=2.163 × $10^{-43}$**<br>CLES:66.13% | U=263181.0<br>**p=1.768 × $10^{-9}$**<br>CLES:65.39% |
| Black snub-nosed monkey | U=50752.0<br>p=0.739<br>CLES:49.10% | U=1453105.5<br>**p=9.066 × $10^{-42}$**<br>CLES:65.93% | U=263219.0<br>**p=3.897 × $10^{-11}$**<br>CLES:66.74% |
| Brazilian guinea pig | U=25976.0<br>p=0.581<br>CLES:51.81% | U=743679.0<br>**p=8.308 × $10^{-19}$**<br>CLES:62.20% | U=119913.0<br>p=5.183 × $10^{-4}$<br>CLES:60.72% |
| Orangutan | U=49474.0<br>p=0.62<br>CLES:48.67% | U=1442730.0<br>**p=3.657 × $10^{-41}$**<br>CLES:65.93% | U=270647.5<br>**p=1.663 × $10^{-11}$**<br>CLES:66.99% |
| Northern American deer mouse | U=57766.0<br>p=0.989<br>CLES:50.04% | U=1632935.5<br>**p=8.05 × $10^{-46}$**<br>CLES:66.20% | U=284843.5<br>**p=7.981 × $10^{-11}$**<br>CLES:66.12% |
| Naked mole-rat female | U=51723.0<br>p=0.711<br>CLES:50.99% | U=1358274.5<br>**p=7.834 × $10^{-43}$**<br>CLES:66.43% | U=253806.0<br>**p=1.577 × $10^{-10}$**<br>CLES:66.11% |
| Mouse | U=68870.0<br>p=0.688<br>CLES:51.01% | U=1923569.5<br>**p=7.856 × $10^{-51}$**<br>CLES:66.48% | U=336538.0<br>**p=9.545 × $10^{-11}$**<br>CLES:65.36% |
| Daurian ground squirrel | U=37456.0<br>p=0.639<br>CLES:48.65% | U=1093726.0<br>**p=4.305 × $10^{-39}$**<br>CLES:66.63% | U=204522.0<br>**p=5.672 × $10^{-11}$**<br>CLES:67.74% |
| Tiger | U=32910.0<br>p=0.442<br>CLES:47.66% | U=1062220.0<br>**p=1.145 × $10^{-36}$**<br>CLES:66.00% | U=172949.0<br>**p=1.239 × $10^{-10}$**<br>CLES:68.60% |
| Lesser hedgehog tenrec | U=2968.0<br>p=0.751<br>CLES:48.32% | U=68142.0<br>**p=1.085 × $10^{-7}$**<br>CLES:63.35% | U=15669.0<br>p=1.668 × $10^{-3}$<br>CLES:65.35% |
| Mongolian gerbil | U=54690.0<br>p=0.48<br>CLES:51.88% | U=1379647.0<br>**p=3.25 × $10^{-36}$**<br>CLES:64.88% | U=248616.0<br>**p=2.556 × $10^{-7}$**<br>CLES:62.88% |
| Ma's night monkey | U=49057.0<br>p=0.635<br>CLES:48.71% | U=1471910.0<br>**p=3.361 × $10^{-43}$**<br>CLES:66.18% | U=261051.0<br>**p=3.78 × $10^{-12}$**<br>CLES:67.84% |

| | | | |
|---|---|---|---|
| Upper Galilee mountains blind mole rat | U=53897.0<br>p=0.748<br>CLES:50.86% | U=1501665.5<br>**p=$8.6 \times 10^{-43}$**<br>CLES:66.10% | U=274777.0<br>**p=$6.857 \times 10^{-10}$**<br>CLES:65.40% |
| Kangaroo rat | U=40201.0<br>p=0.661<br>CLES:51.26% | U=1176900.5<br>**p=$8.585 \times 10^{-34}$**<br>CLES:65.18% | U=209924.0<br>**p=$1.04 \times 10^{-7}$**<br>CLES:64.38% |
| Chimpanzee | U=54731.0<br>p=0.159<br>CLES:53.81% | U=1473999.5<br>**p=$3.239 \times 10^{-47}$**<br>CLES:66.99% | U=248186.0<br>**p=$5.21 \times 10^{-8}$**<br>CLES:63.89% |
| Guinea Pig | U=48747.0<br>p=0.611<br>CLES:51.40% | U=1398412.5<br>**p=$5.328 \times 10^{-38}$**<br>CLES:65.43% | U=248625.0<br>**p=$2.958 \times 10^{-8}$**<br>CLES:64.29% |
| American beaver | U=49675.0<br>p=0.909<br>CLES:49.69% | U=1254391.0<br>**p=$3.289 \times 10^{-31}$**<br>CLES:64.10% | U=246266.5<br>**p=$1.46 \times 10^{-8}$**<br>CLES:64.18% |
| Wild yak | U=39873.0<br>p=0.46<br>CLES:47.90% | U=1162518.0<br>**p=$2.53 \times 10^{-36}$**<br>CLES:65.59% | U=208912.0<br>**p=$2.317 \times 10^{-10}$**<br>CLES:67.02% |
| Coquerel's sifaka | U=45666.0<br>p=0.744<br>CLES:49.10% | U=1346971.0<br>**p=$2.362 \times 10^{-37}$**<br>CLES:65.37% | U=244994.0<br>**p=$2.447 \times 10^{-10}$**<br>CLES:66.45% |
| Alpaca | U=1880.0<br>p=0.868<br>CLES:51.09% | U=73449.0<br>**p=$5.4 \times 10^{-14}$**<br>CLES:69.13% | U=10498.0<br>p=$2.228 \times 10^{-3}$<br>CLES:68.74% |
| Lesser Egyptian jerboa | U=47299.0<br>p=0.39<br>CLES:52.38% | U=1271320.5<br>**p=$1.263 \times 10^{-37}$**<br>CLES:65.66% | U=226838.0<br>**p=$1.799 \times 10^{-7}$**<br>CLES:63.57% |
| Long-tailed chinchilla | U=50298.0<br>p=0.55<br>CLES:51.63% | U=1447200.5<br>**p=$8.169 \times 10^{-43}$**<br>CLES:66.29% | U=255538.0<br>**p=$1.813 \times 10^{-9}$**<br>CLES:65.45% |
| Angola colobus | U=49750.0<br>p=0.556<br>CLES:48.41% | U=1499100.0<br>**p=$2.278 \times 10^{-47}$**<br>CLES:66.93% | U=268544.5<br>**p=$2.782 \times 10^{-13}$**<br>CLES:68.63% |
| Ryukyu mouse | U=63356.0<br>p=0.667<br>CLES:51.11% | U=1775055.5<br>**p=$4.776 \times 10^{-50}$**<br>CLES:66.71% | U=311817.0<br>**p=$1.762 \times 10^{-10}$**<br>CLES:65.44% |
| Chinese hamster PICR | U=61436.0<br>p=0.946<br>CLES:49.83% | U=1705327.0<br>**p=$4.905 \times 10^{-44}$**<br>CLES:65.74% | U=311748.0<br>**p=$5.014 \times 10^{-11}$**<br>CLES:65.85% |
| Vervet-AGM | U=56011.0<br>p=0.935<br>CLES:50.22% | U=1589997.5<br>**p=$1.49 \times 10^{-48}$**<br>CLES:66.97% | U=290693.0<br>**p=$1.101 \times 10^{-11}$**<br>CLES:66.78% |
| Cow | U=47642.0<br>p=0.547<br>CLES:48.35% | U=1444776.0<br>**p=$3.703 \times 10^{-36}$**<br>CLES:64.77% | U=254551.0<br>**p=$1.664 \times 10^{-10}$**<br>CLES:66.53% |
| Bushbaby | U=51476.0<br>p=0.808<br>CLES:50.66% | U=1511888.5<br>**p=$2.185 \times 10^{-44}$**<br>CLES:66.36% | U=261376.5<br>**p=$5.126 \times 10^{-10}$**<br>CLES:65.90% |

| | | | |
|---|---|---|---|
| Bonobo | U=53683.0<br>p=0.74<br>CLES:50.89% | U=1490996.0<br>**p=$1.296 \times 10^{-45}$**<br>CLES:66.59% | U=262150.5<br>**p=$3.848 \times 10^{-10}$**<br>CLES:65.80% |
| Hyrax | U=5652.0<br>p=0.37<br>CLES:54.09% | U=108277.0<br>**p=$9.626 \times 10^{-11}$**<br>CLES:64.52% | U=24249.5<br>p=$1.354 \times 10^{-2}$<br>CLES:60.40% |
| Rabbit | U=38811.0<br>p=1<br>CLES:50.00% | U=1129299.5<br>**p=$1.613 \times 10^{-23}$**<br>CLES:62.43% | U=193280.0<br>**p=$3.774 \times 10^{-6}$**<br>CLES:62.71% |
| Megabat | U=15039.0<br>p=0.406<br>CLES:47.00% | U=412131.0<br>**p=$1.156 \times 10^{-22}$**<br>CLES:65.61% | U=76796.0<br>**p=$1.789 \times 10^{-7}$**<br>CLES:67.73% |
| Dog | U=48996.0<br>p=0.471<br>CLES:48.05% | U=1446153.5<br>**p=$1.07 \times 10^{-37}$**<br>CLES:65.08% | U=263232.0<br>**p=$1.268 \times 10^{-11}$**<br>CLES:67.27% |
| Steppe mouse | U=62788.5<br>p=0.264<br>CLES:52.92% | U=1701533.0<br>**p=$6.322 \times 10^{-47}$**<br>CLES:66.27% | U=283367.5<br>**p=$9.061 \times 10^{-8}$**<br>CLES:63.15% |
| Greater bamboo lemur | U=52040.0<br>p=0.866<br>CLES:50.46% | U=1501533.5<br>**p=$7.822 \times 10^{-42}$**<br>CLES:65.86% | U=264236.5<br>**p=$8.364 \times 10^{-10}$**<br>CLES:65.59% |
| Pig | U=54282.0<br>p=0.403<br>CLES:52.24% | U=1479948.5<br>**p=$1.779 \times 10^{-39}$**<br>CLES:65.47% | U=263679.0<br>**p=$5.647 \times 10^{-8}$**<br>CLES:63.65% |
| Tree Shrew | U=2546.0<br>p=0.258<br>CLES:56.64% | U=54602.0<br>**p=$1.824 \times 10^{-9}$**<br>CLES:65.85% | U=9214.0<br>p=$8.847 \times 10^{-2}$<br>CLES:59.39% |
| Pika | U=7564.0<br>p=0.594<br>CLES:47.77% | U=159055.5<br>**p=$3.35 \times 10^{-8}$**<br>CLES:60.94% | U=34925.0<br>p=$8.845 \times 10^{-4}$<br>CLES:62.99% |
| Shrew | U=1140.0<br>p=0.502<br>CLES:45.45% | U=39450.0<br>**p=$3.972 \times 10^{-6}$**<br>CLES:63.73% | U=7932.0<br>p=$9.077 \times 10^{-3}$<br>CLES:66.40% |
| Squirrel | U=53357.0<br>p=0.74<br>CLES:50.88% | U=1506349.5<br>**p=$2.175 \times 10^{-46}$**<br>CLES:66.86% | U=276888.0<br>**p=$2.799 \times 10^{-10}$**<br>CLES:65.75% |
| Gorilla | U=52043.0<br>p=0.713<br>CLES:50.99% | U=1493370.0<br>**p=$4.889 \times 10^{-48}$**<br>CLES:67.14% | U=262478.0<br>**p=$2.448 \times 10^{-10}$**<br>CLES:66.09% |
| Cat | U=54049.0<br>p=0.704<br>CLES:48.99% | U=1580816.5<br>**p=$2.452 \times 10^{-42}$**<br>CLES:65.73% | U=284498.0<br>**p=$3.024 \times 10^{-11}$**<br>CLES:66.58% |

### Rat Reference Genome, Saunders et al., 2018

| MWU Test | Vasculature vs Glia | Vasculature vs Neuron | Glia vs Neuron |
|---|---|---|---|
| Microbat | U=32236.0<br>p=$2.118 \times 10^{-2}$<br>CLES:56.14% | U=94906.0<br>**p=$4.5 \times 10^{-17}$**<br>CLES:68.04% | U=60486.0<br>**p=$3.051 \times 10^{-8}$**<br>CLES:63.51% |
| Hedgehog | U=2496.0<br>p=$7.143 \times 10^{-2}$<br>CLES:59.26% | U=5861.0<br>**p=$1.033 \times 10^{-5}$**<br>CLES:68.94% | U=3640.0<br>p=$1.406 \times 10^{-2}$<br>CLES:61.84% |
| Naked mole-rat male | U=36677.0<br>p=$2.034 \times 10^{-2}$<br>CLES:55.97% | U=109816.0<br>**p=$1.352 \times 10^{-22}$**<br>CLES:70.47% | U=74020.0<br>**p=$5.548 \times 10^{-12}$**<br>CLES:66.11% |
| Sooty mangabey | U=49876.0<br>p=$2.17 \times 10^{-4}$<br>CLES:58.91% | U=136294.0<br>**p=$1.911 \times 10^{-23}$**<br>CLES:69.72% | U=89770.0<br>**p=$6.82 \times 10^{-9}$**<br>CLES:62.70% |
| Olive baboon | U=48806.0<br>p=$9.974 \times 10^{-4}$<br>CLES:57.95% | U=137613.0<br>**p=$5.299 \times 10^{-23}$**<br>CLES:69.46% | U=91184.0<br>**p=$6.788 \times 10^{-10}$**<br>CLES:63.54% |
| American bison | U=32100.0<br>p=$2.063 \times 10^{-2}$<br>CLES:56.18% | U=93422.0<br>**p=$1.211 \times 10^{-18}$**<br>CLES:69.01% | U=58738.0<br>**p=$7.247 \times 10^{-10}$**<br>CLES:65.20% |
| Leopard | U=47781.0<br>p=$1.584 \times 10^{-3}$<br>CLES:57.66% | U=129525.0<br>**p=$5.153 \times 10^{-20}$**<br>CLES:68.19% | U=84278.0<br>**p=$3.325 \times 10^{-8}$**<br>CLES:62.27% |
| Bolivian squirrel monkey | U=47649.0<br>p=$1.149 \times 10^{-4}$<br>CLES:59.42% | U=132146.0<br>**p=$7.379 \times 10^{-27}$**<br>CLES:71.51% | U=87649.0<br>**p=$8.36 \times 10^{-11}$**<br>CLES:64.41% |
| Crab-eating macaque | U=47853.0<br>p=$8.65 \times 10^{-4}$<br>CLES:58.09% | U=132896.0<br>**p=$4.506 \times 10^{-21}$**<br>CLES:68.66% | U=87087.0<br>**p=$7.967 \times 10^{-9}$**<br>CLES:62.76% |
| Human | U=54034.0<br>p=$1.714 \times 10^{-4}$<br>CLES:58.87% | U=145214.0<br>**p=$3.421 \times 10^{-23}$**<br>CLES:69.25% | U=96648.5<br>**p=$4.719 \times 10^{-9}$**<br>CLES:62.58% |
| Red fox | U=41702.0<br>p=$7.088 \times 10^{-3}$<br>CLES:56.73% | U=119993.0<br>**p=$3.386 \times 10^{-20}$**<br>CLES:68.72% | U=79244.0<br>**p=$2.21 \times 10^{-9}$**<br>CLES:63.61% |
| Shrew mouse | U=60282.5<br>p=$1.689 \times 10^{-3}$<br>CLES:57.16% | U=154946.0<br>**p=$4.151 \times 10^{-19}$**<br>CLES:66.87% | U=105185.0<br>**p=$3.478 \times 10^{-8}$**<br>CLES:61.50% |
| Arctic ground squirrel | U=49677.0<br>p=$4.941 \times 10^{-4}$<br>CLES:58.38% | U=137568.0<br>**p=$1.495 \times 10^{-24}$**<br>CLES:70.21% | U=91919.0<br>**p=$9.488 \times 10^{-10}$**<br>CLES:63.36% |
| Chinese hamster CriGri | U=49545.0<br>p=$1.121 \times 10^{-3}$<br>CLES:57.83% | U=126025.5<br>**p=$2.573 \times 10^{-18}$**<br>CLES:67.39% | U=84413.0<br>**p=$6.961 \times 10^{-7}$**<br>CLES:60.90% |
| Opossum | U=37454.0<br>p=0.185 | U=95696.0<br>**p=$5.893 \times 10^{-7}$** | U=65892.0<br>p=$9.463 \times 10^{-4}$ |

| | | | |
|---|---|---|---|
| | CLES:53.35% | CLES:60.38% | CLES:57.65% |
| Dolphin | U=12978.0<br>p=0.173<br>CLES:54.55% | U=40372.0<br>**$p=2.542 \times 10^{-13}$**<br>CLES:69.48% | U=24080.0<br>**$p=1.737 \times 10^{-7}$**<br>CLES:66.23% |
| Capuchin | U=49656.0<br>**$p=1.704 \times 10^{-5}$**<br>CLES:60.43% | U=134991.0<br>**$p=3.294 \times 10^{-28}$**<br>CLES:72.02% | U=90546.0<br>**$p=1.699 \times 10^{-10}$**<br>CLES:64.02% |
| Mouse Lemur | U=49681.0<br>$p=1.129 \times 10^{-4}$<br>CLES:59.33% | U=132472.5<br>**$p=1.417 \times 10^{-21}$**<br>CLES:68.92% | U=86179.0<br>**$p=1.625 \times 10^{-7}$**<br>CLES:61.53% |
| Damara mole rat | U=38599.0<br>$p=6.331 \times 10^{-3}$<br>CLES:56.96% | U=117657.0<br>**$p=5.161 \times 10^{-21}$**<br>CLES:69.33% | U=77670.0<br>**$p=1.003 \times 10^{-9}$**<br>CLES:64.07% |
| Marmoset | U=48203.0<br>**$p=1.606 \times 10^{-5}$**<br>CLES:60.57% | U=136630.0<br>**$p=3.584 \times 10^{-27}$**<br>CLES:71.48% | U=86463.0<br>**$p=4.689 \times 10^{-9}$**<br>CLES:63.03% |
| Goat | U=47258.0<br>$p=7.828 \times 10^{-4}$<br>CLES:58.19% | U=131154.0<br>**$p=6.95 \times 10^{-22}$**<br>CLES:69.13% | U=85260.0<br>**$p=7.77 \times 10^{-9}$**<br>CLES:62.84% |
| Common wombat | U=35001.0<br>$p=6.273 \times 10^{-2}$<br>CLES:54.83% | U=96144.5<br>**$p=2.762 \times 10^{-11}$**<br>CLES:64.02% | U=62968.0<br>**$p=6.951 \times 10^{-6}$**<br>CLES:60.70% |
| American black bear | U=38994.0<br>$p=4.312 \times 10^{-3}$<br>CLES:57.26% | U=93981.5<br>**$p=1.869 \times 10^{-13}$**<br>CLES:65.64% | U=62232.0<br>$p=1.27 \times 10^{-4}$<br>CLES:58.99% |
| Sloth | U=810.0<br>p=0.92<br>CLES:50.69% | U=2194.0<br>**$p=8.614 \times 10^{-5}$**<br>CLES:71.82% | U=1594.0<br>$p=3.187 \times 10^{-4}$<br>CLES:72.13% |
| Polar bear | U=37835.5<br>$p=6.179 \times 10^{-3}$<br>CLES:57.02% | U=103292.5<br>**$p=9.221 \times 10^{-18}$**<br>CLES:68.04% | U=67848.5<br>**$p=4.221 \times 10^{-7}$**<br>CLES:61.84% |
| Alpine marmot | U=43410.5<br>$p=7.373 \times 10^{-3}$<br>CLES:56.63% | U=119659.5<br>**$p=4.065 \times 10^{-20}$**<br>CLES:68.62% | U=78138.5<br>**$p=5.024 \times 10^{-9}$**<br>CLES:63.28% |
| Prairie vole | U=52085.0<br>$p=3.397 \times 10^{-2}$<br>CLES:54.96% | U=142467.0<br>**$p=2.05 \times 10^{-20}$**<br>CLES:67.99% | U=104821.0<br>**$p=1.293 \times 10^{-12}$**<br>CLES:65.03% |
| Donkey | U=49827.0<br>$p=2.619 \times 10^{-3}$<br>CLES:57.19% | U=129413.0<br>**$p=8.561 \times 10^{-19}$**<br>CLES:67.57% | U=89522.0<br>**$p=8.259 \times 10^{-8}$**<br>CLES:61.65% |
| Golden Hamster | U=48263.0<br>$p=1.898 \times 10^{-2}$<br>CLES:55.61% | U=126827.0<br>**$p=4.504 \times 10^{-17}$**<br>CLES:66.69% | U=89702.5<br>**$p=1.07 \times 10^{-8}$**<br>CLES:62.46% |
| Armadillo | U=40059.0<br>$p=8.684 \times 10^{-4}$<br>CLES:58.47% | U=107425.0<br>**$p=1.387 \times 10^{-16}$**<br>CLES:67.16% | U=67745.0<br>**$p=5.113 \times 10^{-5}$**<br>CLES:59.41% |
| Ugandan red Colobus | U=49075.0<br>$p=1.789 \times 10^{-4}$ | U=130579.0<br>**$p=4.788 \times 10^{-23}$** | U=87195.0<br>**$p=4.864 \times 10^{-9}$** |

| | | | |
|---|---|---|---|
| | CLES:59.07% | CLES:69.73% | CLES:62.90% |
| Pig-tailed macaque | U=49100.0<br>$p=1.156 \times 10^{-3}$<br>CLES:57.83% | U=137312.0<br>**$p=1.14 \times 10^{-22}$**<br>CLES:69.31% | U=92167.5<br>**$p=3.707 \times 10^{-10}$**<br>CLES:63.71% |
| Panda | U=47532.0<br>$p=7.439 \times 10^{-4}$<br>CLES:58.20% | U=126205.0<br>**$p=1.483 \times 10^{-23}$**<br>CLES:70.14% | U=85533.0<br>**$p=1.866 \times 10^{-10}$**<br>CLES:64.15% |
| Ferret | U=43858.0<br>$p=8.616 \times 10^{-4}$<br>CLES:58.28% | U=123070.0<br>**$p=1.608 \times 10^{-20}$**<br>CLES:68.78% | U=80213.0<br>**$p=7.334 \times 10^{-8}$**<br>CLES:62.14% |
| Koala | U=39197.5<br>p=0.538<br>CLES:51.52% | U=107751.0<br>**$p=8.05 \times 10^{-11}$**<br>CLES:63.30% | U=82100.0<br>**$p=1.001 \times 10^{-9}$**<br>CLES:63.70% |
| Wallaby | U=4013.0<br>p=0.819<br>CLES:49.01% | U=6611.0<br>p=0.218<br>CLES:54.85% | U=6569.5<br>p=0.111<br>CLES:56.35% |
| Tarsier | U=36462.0<br>$p=6.461 \times 10^{-4}$<br>CLES:58.89% | U=94609.0<br>**$p=1.552 \times 10^{-21}$**<br>CLES:70.65% | U=64261.0<br>**$p=2.07 \times 10^{-9}$**<br>CLES:64.28% |
| Degu | U=32996.0<br>$p=5.616 \times 10^{-3}$<br>CLES:57.36% | U=106117.0<br>**$p=1.097 \times 10^{-21}$**<br>CLES:70.34% | U=70122.5<br>**$p=2.038 \times 10^{-9}$**<br>CLES:64.25% |
| Chinese hamster CHOK1GS | U=60558.5<br>$p=1.695 \times 10^{-4}$<br>CLES:58.63% | U=158844.5<br>**$p=2.133 \times 10^{-23}$**<br>CLES:68.85% | U=103804.5<br>**$p=1.432 \times 10^{-8}$**<br>CLES:61.90% |
| Drill | U=43386.5<br>$p=3.003 \times 10^{-3}$<br>CLES:57.37% | U=123026.0<br>**$p=9.849 \times 10^{-21}$**<br>CLES:68.87% | U=80862.0<br>**$p=1.833 \times 10^{-9}$**<br>CLES:63.60% |
| Gibbon | U=44841.5<br>$p=1.555 \times 10^{-4}$<br>CLES:59.38% | U=124712.0<br>**$p=3.66 \times 10^{-22}$**<br>CLES:69.59% | U=80770.5<br>**$p=7.11 \times 10^{-7}$**<br>CLES:61.12% |
| Macaque | U=51660.0<br>$p=6.675 \times 10^{-4}$<br>CLES:58.10% | U=140751.5<br>**$p=6.247 \times 10^{-23}$**<br>CLES:69.30% | U=96287.0<br>**$p=1.561 \times 10^{-10}$**<br>CLES:63.83% |
| American mink | U=45612.5<br>$p=4.763 \times 10^{-3}$<br>CLES:56.90% | U=121684.5<br>**$p=7.633 \times 10^{-19}$**<br>CLES:67.84% | U=80580.0<br>**$p=2.359 \times 10^{-8}$**<br>CLES:62.52% |
| Gelada | U=50336.0<br>$p=2.245 \times 10^{-4}$<br>CLES:58.86% | U=136340.0<br>**$p=4.615 \times 10^{-23}$**<br>CLES:69.55% | U=91935.0<br>**$p=6.127 \times 10^{-9}$**<br>CLES:62.65% |
| Sheep | U=42183.5<br>$p=5.424 \times 10^{-4}$<br>CLES:58.70% | U=118781.5<br>**$p=1.625 \times 10^{-19}$**<br>CLES:68.41% | U=76581.5<br>**$p=1.3 \times 10^{-7}$**<br>CLES:62.06% |
| Tasmanian devil | U=34577.0<br>$p=6.652 \times 10^{-2}$<br>CLES:54.76% | U=90318.5<br>**$p=9.083 \times 10^{-10}$**<br>CLES:63.08% | U=63210.0<br>**$p=9.224 \times 10^{-5}$**<br>CLES:59.21% |
| Elephant | U=48619.0<br>$p=1.735 \times 10^{-4}$ | U=127153.5<br>**$p=3.459 \times 10^{-22}$** | U=83976.0<br>**$p=9.29 \times 10^{-8}$** |

| | | | |
|---|---|---|---|
| | CLES:59.11% | CLES:69.43% | CLES:61.82% |
| Algerian mouse | U=60637.5<br>$p=4.188 \times 10^{-4}$<br>CLES:58.07% | U=157562.0<br>**$p=6.67 \times 10^{-22}$**<br>CLES:68.18% | U=105145.0<br>**$p=1.175 \times 10^{-8}$**<br>CLES:61.92% |
| Dingo | U=45213.5<br>$p=9.337 \times 10^{-4}$<br>CLES:58.14% | U=123616.0<br>**$p=8.404 \times 10^{-20}$**<br>CLES:68.37% | U=82071.0<br>**$p=4.186 \times 10^{-7}$**<br>CLES:61.28% |
| Golden snub-nosed monkey | U=43161.0<br>$p=5.456 \times 10^{-3}$<br>CLES:56.89% | U=123416.0<br>**$p=2.835 \times 10^{-21}$**<br>CLES:69.12% | U=81160.0<br>**$p=5.116 \times 10^{-10}$**<br>CLES:64.07% |
| Horse | U=47832.0<br>$p=1.998 \times 10^{-3}$<br>CLES:57.49% | U=138300.0<br>**$p=2.886 \times 10^{-23}$**<br>CLES:69.57% | U=90096.0<br>**$p=1.406 \times 10^{-9}$**<br>CLES:63.34% |
| Black snub-nosed monkey | U=47090.0<br>$p=3.234 \times 10^{-4}$<br>CLES:58.79% | U=124429.5<br>**$p=2.252 \times 10^{-20}$**<br>CLES:68.60% | U=81343.5<br>**$p=3.672 \times 10^{-7}$**<br>CLES:61.34% |
| Brazilian guinea pig | U=25277.0<br>$p=6.166 \times 10^{-3}$<br>CLES:57.77% | U=62588.0<br>**$p=6.397 \times 10^{-9}$**<br>CLES:63.68% | U=44298.5<br>$p=1.172 \times 10^{-2}$<br>CLES:56.40% |
| Orangutan | U=47984.5<br>$p=1.995 \times 10^{-3}$<br>CLES:57.48% | U=131062.0<br>**$p=6.888 \times 10^{-23}$**<br>CLES:69.62% | U=87980.0<br>**$p=2.671 \times 10^{-10}$**<br>CLES:63.95% |
| Northern American deer mouse | U=54810.0<br>$p=4.6 \times 10^{-3}$<br>CLES:56.60% | U=144267.0<br>**$p=1.717 \times 10^{-20}$**<br>CLES:67.92% | U=99834.0<br>**$p=1.261 \times 10^{-9}$**<br>CLES:62.90% |
| Naked mole-rat female | U=43883.0<br>$p=1.157 \times 10^{-3}$<br>CLES:58.06% | U=126541.5<br>**$p=3.585 \times 10^{-25}$**<br>CLES:70.97% | U=83405.0<br>**$p=1.004 \times 10^{-10}$**<br>CLES:64.57% |
| Mouse | U=63872.5<br>$p=1.144 \times 10^{-3}$<br>CLES:57.32% | U=166932.5<br>**$p=2.03 \times 10^{-21}$**<br>CLES:67.66% | U=111410.0<br>**$p=6.626 \times 10^{-9}$**<br>CLES:61.95% |
| Daurian ground squirrel | U=31941.0<br>$p=7.772 \times 10^{-2}$<br>CLES:54.68% | U=96671.0<br>**$p=3.861 \times 10^{-20}$**<br>CLES:69.72% | U=62045.5<br>**$p=9.652 \times 10^{-12}$**<br>CLES:66.67% |
| Tiger | U=30023.0<br>$p=1.786 \times 10^{-2}$<br>CLES:56.43% | U=87687.0<br>**$p=1.664 \times 10^{-18}$**<br>CLES:69.28% | U=54987.5<br>**$p=1.956 \times 10^{-8}$**<br>CLES:64.03% |
| Lesser hedgehog tenrec | U=2986.0<br>p=0.145<br>CLES:57.03% | U=5099.0<br>$p=1.054 \times 10^{-3}$<br>CLES:64.29% | U=4121.0<br>$p=5.093 \times 10^{-2}$<br>CLES:58.84% |
| Mongolian gerbil | U=49032.5<br>$p=2.782 \times 10^{-4}$<br>CLES:58.80% | U=114903.5<br>**$p=1.872 \times 10^{-15}$**<br>CLES:66.07% | U=73908.5<br>$p=1.316 \times 10^{-4}$<br>CLES:58.57% |
| Ma's night monkey | U=48231.0<br>**$p=6.963 \times 10^{-6}$**<br>CLES:61.01% | U=132035.0<br>**$p=1.068 \times 10^{-26}$**<br>CLES:71.49% | U=86932.0<br>**$p=1.525 \times 10^{-8}$**<br>CLES:62.51% |

| | | | |
|---|---|---|---|
| Upper Galilee mountains blind mole rat | U=50950.0<br>$p=8.177 \times 10^{-4}$<br>CLES:57.98% | U=137826.0<br>**$p=1.717 \times 10^{-22}$**<br>CLES:69.23% | U=96693.0<br>**$p=9.976 \times 10^{-10}$**<br>CLES:63.14% |
| Kangaroo rat | U=36699.5<br>$p=7.117 \times 10^{-2}$<br>CLES:54.60% | U=103709.5<br>**$p=2.86 \times 10^{-15}$**<br>CLES:66.61% | U=77875.5<br>**$p=1.051 \times 10^{-8}$**<br>CLES:63.01% |
| Chimpanzee | U=44547.0<br>$p=1.276 \times 10^{-3}$<br>CLES:57.94% | U=125806.0<br>**$p=3.964 \times 10^{-23}$**<br>CLES:70.05% | U=86576.5<br>**$p=5.307 \times 10^{-10}$**<br>CLES:63.81% |
| Guinea Pig | U=44040.5<br>$p=9.118 \times 10^{-3}$<br>CLES:56.42% | U=126160.5<br>**$p=4.507 \times 10^{-21}$**<br>CLES:68.93% | U=84972.5<br>**$p=4.033 \times 10^{-10}$**<br>CLES:63.98% |
| American beaver | U=44865.0<br>$p=5.029 \times 10^{-3}$<br>CLES:56.88% | U=104338.0<br>**$p=4.281 \times 10^{-16}$**<br>CLES:66.84% | U=69071.0<br>**$p=1.29 \times 10^{-6}$**<br>CLES:61.10% |
| Wild yak | U=35596.0<br>$p=6.422 \times 10^{-3}$<br>CLES:57.12% | U=103311.5<br>**$p=7.921 \times 10^{-20}$**<br>CLES:69.21% | U=65327.0<br>**$p=5.531 \times 10^{-10}$**<br>CLES:64.90% |
| Coquerel's sifaka | U=47669.0<br>**$p=2.761 \times 10^{-5}$**<br>CLES:60.29% | U=131273.0<br>**$p=4.415 \times 10^{-23}$**<br>CLES:69.73% | U=82350.0<br>**$p=6.837 \times 10^{-8}$**<br>CLES:62.09% |
| Alpaca | U=1623.0<br>p=0.133<br>CLES:58.53% | U=4908.0<br>**$p=3.474 \times 10^{-9}$**<br>CLES:77.74% | U=3552.0<br>**$p=4.726 \times 10^{-5}$**<br>CLES:70.63% |
| Lesser Egyptian jerboa | U=38399.0<br>p=0.201<br>CLES:53.20% | U=110705.0<br>**$p=2.088 \times 10^{-16}$**<br>CLES:67.02% | U=85251.0<br>**$p=1.116 \times 10^{-11}$**<br>CLES:65.21% |
| Long-tailed chinchilla | U=42539.0<br>$p=5.266 \times 10^{-3}$<br>CLES:56.94% | U=126875.0<br>**$p=7.201 \times 10^{-23}$**<br>CLES:69.86% | U=84432.0<br>**$p=1.135 \times 10^{-10}$**<br>CLES:64.53% |
| Angola colobus | U=47403.0<br>$p=3.31 \times 10^{-4}$<br>CLES:58.75% | U=127426.0<br>**$p=4.488 \times 10^{-24}$**<br>CLES:70.38% | U=87519.5<br>**$p=6.755 \times 10^{-11}$**<br>CLES:64.45% |
| Ryukyu mouse | U=60398.0<br>$p=9.064 \times 10^{-4}$<br>CLES:57.57% | U=155138.5<br>**$p=1.191 \times 10^{-20}$**<br>CLES:67.64% | U=104364.0<br>**$p=5.776 \times 10^{-8}$**<br>CLES:61.32% |
| Chinese hamster PICR | U=59020.0<br>**$p=1.982 \times 10^{-5}$**<br>CLES:59.90% | U=157358.5<br>**$p=5.27 \times 10^{-28}$**<br>CLES:70.96% | U=100486.0<br>**$p=2.831 \times 10^{-9}$**<br>CLES:62.62% |
| Vervet-AGM | U=53038.0<br>$p=1.196 \times 10^{-4}$<br>CLES:59.13% | U=140999.5<br>**$p=6.743 \times 10^{-25}$**<br>CLES:70.22% | U=95811.0<br>**$p=6.394 \times 10^{-10}$**<br>CLES:63.32% |
| Cow | U=46742.0<br>$p=3.395 \times 10^{-4}$<br>CLES:58.77% | U=127061.0<br>**$p=1.418 \times 10^{-20}$**<br>CLES:68.63% | U=83883.5<br>**$p=6.186 \times 10^{-8}$**<br>CLES:62.04% |
| Bushbaby | U=48945.0<br>$p=2.784 \times 10^{-3}$<br>CLES:57.18% | U=133592.0<br>**$p=2.062 \times 10^{-21}$**<br>CLES:68.82% | U=92252.0<br>**$p=1.967 \times 10^{-9}$**<br>CLES:63.06% |

| | | | |
|---|---|---|---|
| Bonobo | U=45777.0<br>p=$4.452 \times 10^{-4}$<br>CLES:58.64% | U=128859.0<br>**p=$1.419 \times 10^{-25}$**<br>CLES:71.04% | U=84359.5<br>**p=$3.081 \times 10^{-10}$**<br>CLES:64.10% |
| Hyrax | U=4424.0<br>p=$4.645 \times 10^{-2}$<br>CLES:58.80% | U=9144.0<br>**p=$2.781 \times 10^{-6}$**<br>CLES:67.91% | U=6354.0<br>p=$5.645 \times 10^{-3}$<br>CLES:61.47% |
| Rabbit | U=36268.0<br>p=$9.062 \times 10^{-3}$<br>CLES:56.76% | U=100729.5<br>**p=$1.191 \times 10^{-13}$**<br>CLES:65.58% | U=64585.0<br>**p=$9.036 \times 10^{-5}$**<br>CLES:59.22% |
| Megabat | U=14090.0<br>p=$1.199 \times 10^{-3}$<br>CLES:60.78% | U=32243.0<br>**p=$2.21 \times 10^{-12}$**<br>CLES:69.80% | U=21932.0<br>p=$2.221 \times 10^{-4}$<br>CLES:61.30% |
| Dog | U=46904.0<br>p=$4.639 \times 10^{-4}$<br>CLES:58.56% | U=127945.5<br>**p=$2.09 \times 10^{-21}$**<br>CLES:69.03% | U=83326.0<br>**p=$2.355 \times 10^{-7}$**<br>CLES:61.49% |
| Steppe mouse | U=53498.5<br>p=$2.848 \times 10^{-2}$<br>CLES:55.09% | U=140029.5<br>**p=$7.052 \times 10^{-17}$**<br>CLES:66.14% | U=100956.5<br>**p=$2.618 \times 10^{-9}$**<br>CLES:62.59% |
| Greater bamboo lemur | U=48060.0<br>p=$1.626 \times 10^{-2}$<br>CLES:55.76% | U=131388.0<br>**p=$5.605 \times 10^{-20}$**<br>CLES:68.12% | U=91635.5<br>**p=$3.767 \times 10^{-11}$**<br>CLES:64.47% |
| Pig | U=50126.5<br>p=$2.772 \times 10^{-3}$<br>CLES:57.14% | U=133732.0<br>**p=$3.108 \times 10^{-22}$**<br>CLES:69.17% | U=90417.0<br>**p=$1.941 \times 10^{-9}$**<br>CLES:63.09% |
| Tree Shrew | U=1764.0<br>p=0.115<br>CLES:58.94% | U=3568.0<br>p=$1.165 \times 10^{-2}$<br>CLES:61.87% | U=1744.0<br>p=0.493<br>CLES:53.84% |
| Pika | U=6478.0<br>p=$4.474 \times 10^{-2}$<br>CLES:58.05% | U=12600.0<br>**p=$2.889 \times 10^{-8}$**<br>CLES:69.60% | U=8285.0<br>p=$7.639 \times 10^{-4}$<br>CLES:63.05% |
| Shrew | U=1087.0<br>p=$2.302 \times 10^{-2}$<br>CLES:64.70% | U=2959.0<br>**p=$6.297 \times 10^{-5}$**<br>CLES:70.86% | U=1784.0<br>p=0.14<br>CLES:58.59% |
| Squirrel | U=46614.0<br>p=$2.473 \times 10^{-3}$<br>CLES:57.37% | U=130430.0<br>**p=$1.696 \times 10^{-20}$**<br>CLES:68.48% | U=86849.0<br>**p=$3.097 \times 10^{-8}$**<br>CLES:62.23% |
| Gorilla | U=46974.0<br>p=$1.095 \times 10^{-4}$<br>CLES:59.48% | U=130746.0<br>**p=$1.015 \times 10^{-25}$**<br>CLES:71.08% | U=86926.0<br>**p=$2.538 \times 10^{-9}$**<br>CLES:63.20% |
| Cat | U=48686.0<br>p=$4.11 \times 10^{-3}$<br>CLES:56.90% | U=135969.5<br>**p=$4.242 \times 10^{-21}$**<br>CLES:68.54% | U=90236.0<br>**p=$1.437 \times 10^{-9}$**<br>CLES:63.28% |

### Human Reference Genome, Zhang et al., 2014

| MWU Test | Endothelia vs Glia | Endothelia vs Neuron | Glia vs Neuron |
|---|---|---|---|
| Microbat | U=273750.0<br>p=0.798<br>CLES:50.39% | U=390748.0<br>**$p=1.111 \times 10^{-19}$**<br>CLES:63.32% | U=481179.5<br>**$p=2.475 \times 10^{-22}$**<br>CLES:63.47% |
| Hedgehog | U=20192.0<br>p=0.327<br>CLES:52.88% | U=24449.5<br>**$p=4.378 \times 10^{-6}$**<br>CLES:63.46% | U=30544.0<br>**$p=3.006 \times 10^{-5}$**<br>CLES:61.42% |
| Naked mole-rat male | U=311851.0<br>p=0.511<br>CLES:50.97% | U=443349.0<br>**$p=2.888 \times 10^{-30}$**<br>CLES:66.50% | U=607595.5<br>**$p=1.214 \times 10^{-35}$**<br>CLES:66.46% |
| Sooty mangabey | U=419622.5<br>p=0.229<br>CLES:51.65% | U=574596.5<br>**$p=6.774 \times 10^{-25}$**<br>CLES:63.78% | U=746171.5<br>**$p=1.674 \times 10^{-23}$**<br>CLES:62.34% |
| Olive baboon | U=416375.0<br>p=0.523<br>CLES:50.87% | U=572835.0<br>**$p=2.443 \times 10^{-23}$**<br>CLES:63.29% | U=752125.5<br>**$p=1.786 \times 10^{-25}$**<br>CLES:62.89% |
| American bison | U=264531.0<br>p=0.652<br>CLES:50.69% | U=362311.0<br>**$p=1.05 \times 10^{-25}$**<br>CLES:65.80% | U=465672.5<br>**$p=9.295 \times 10^{-30}$**<br>CLES:65.97% |
| Leopard | U=377751.0<br>p=0.296<br>CLES:51.46% | U=555870.5<br>**$p=2.955 \times 10^{-33}$**<br>CLES:66.38% | U=718287.5<br>**$p=7.63 \times 10^{-35}$**<br>CLES:65.56% |
| Bolivian squirrel monkey | U=390929.0<br>$p=2.956 \times 10^{-2}$<br>CLES:53.05% | U=551568.5<br>**$p=1.678 \times 10^{-34}$**<br>CLES:66.74% | U=693993.5<br>**$p=5.26 \times 10^{-30}$**<br>CLES:64.44% |
| Crab-eating macaque | U=424970.5<br>p=0.126<br>CLES:52.09% | U=568664.0<br>**$p=1.977 \times 10^{-22}$**<br>CLES:63.02% | U=733529.5<br>**$p=9.083 \times 10^{-21}$**<br>CLES:61.56% |
| Red fox | U=352994.0<br>p=0.319<br>CLES:51.42% | U=485067.0<br>**$p=1.121 \times 10^{-21}$**<br>CLES:63.30% | U=601441.5<br>**$p=1.576 \times 10^{-21}$**<br>CLES:62.43% |
| Shrew mouse | U=385604.0<br>p=0.628<br>CLES:49.33% | U=578146.0<br>**$p=2.388 \times 10^{-30}$**<br>CLES:65.38% | U=758645.0<br>**$p=9.614 \times 10^{-39}$**<br>CLES:66.27% |
| Arctic ground squirrel | U=399111.0<br>p=0.159<br>CLES:51.95% | U=576738.5<br>**$p=2.946 \times 10^{-37}$**<br>CLES:67.26% | U=736170.5<br>**$p=1.829 \times 10^{-37}$**<br>CLES:66.09% |
| Chinese hamster CriGri | U=326602.0<br>p=0.846<br>CLES:49.72% | U=470270.0<br>**$p=1.165 \times 10^{-26}$**<br>CLES:65.08% | U=610106.5<br>**$p=3.618 \times 10^{-32}$**<br>CLES:65.53% |
| Opossum | U=295782.0<br>$p=8.135 \times 10^{-2}$<br>CLES:52.61% | U=394432.0<br>**$p=2.518 \times 10^{-14}$**<br>CLES:61.08% | U=485663.0<br>**$p=8.644 \times 10^{-11}$**<br>CLES:58.80% |
| Dolphin | U=111409.0<br>p=0.954<br>CLES:49.89% | U=145801.0<br>**$p=1.061 \times 10^{-12}$**<br>CLES:63.31% | U=171168.0<br>**$p=3.943 \times 10^{-14}$**<br>CLES:63.56% |

| | | | |
|---|---|---|---|
| Capuchin | U=409663.0<br>$p=7.02 \times 10^{-2}$<br>CLES:52.50% | U=582344.0<br>**$p=9.055 \times 10^{-38}$**<br>CLES:67.37% | U=756461.5<br>**$p=4.822 \times 10^{-36}$**<br>CLES:65.63% |
| Mouse Lemur | U=385941.0<br>p=0.163<br>CLES:51.95% | U=556378.5<br>**$p=2.305 \times 10^{-35}$**<br>CLES:66.93% | U=711476.0<br>**$p=3.704 \times 10^{-35}$**<br>CLES:65.67% |
| Damara mole rat | U=319381.0<br>p=0.444<br>CLES:51.12% | U=472539.5<br>**$p=1.908 \times 10^{-32}$**<br>CLES:66.85% | U=609592.0<br>**$p=1.759 \times 10^{-36}$**<br>CLES:66.67% |
| Marmoset | U=398370.5<br>$p=2.62 \times 10^{-2}$<br>CLES:53.10% | U=559102.0<br>**$p=9.877 \times 10^{-33}$**<br>CLES:66.21% | U=711754.0<br>**$p=2.874 \times 10^{-28}$**<br>CLES:63.87% |
| Goat | U=382130.0<br>p=0.257<br>CLES:51.58% | U=560099.0<br>**$p=2.826 \times 10^{-32}$**<br>CLES:66.06% | U=706348.5<br>**$p=1.415 \times 10^{-31}$**<br>CLES:64.79% |
| Common wombat | U=281213.0<br>p=0.145<br>CLES:52.21% | U=381031.0<br>**$p=1.851 \times 10^{-19}$**<br>CLES:63.33% | U=472354.0<br>**$p=1.512 \times 10^{-16}$**<br>CLES:61.39% |
| American black bear | U=317911.0<br>$p=6.817 \times 10^{-2}$<br>CLES:52.68% | U=388892.0<br>**$p=2.057 \times 10^{-20}$**<br>CLES:63.59% | U=490242.0<br>**$p=4.948 \times 10^{-16}$**<br>CLES:61.07% |
| Sloth | U=6157.0<br>p=0.774<br>CLES:48.88% | U=6784.0<br>$p=9.68 \times 10^{-4}$<br>CLES:63.31% | U=9811.0<br>$p=1.14 \times 10^{-4}$<br>CLES:64.22% |
| Polar bear | U=291638.0<br>p=0.515<br>CLES:50.97% | U=375235.0<br>**$p=4.034 \times 10^{-19}$**<br>CLES:63.21% | U=473442.5<br>**$p=1.1 \times 10^{-19}$**<br>CLES:62.57% |
| Alpine marmot | U=352752.0<br>p=0.461<br>CLES:51.05% | U=486467.0<br>**$p=5.655 \times 10^{-28}$**<br>CLES:65.35% | U=623755.0<br>**$p=1.821 \times 10^{-30}$**<br>CLES:64.97% |
| Prairie vole | U=355279.0<br>p=0.424<br>CLES:48.88% | U=540134.0<br>**$p=5.111 \times 10^{-30}$**<br>CLES:65.58% | U=723771.5<br>**$p=1.189 \times 10^{-40}$**<br>CLES:66.92% |
| Donkey | U=390565.0<br>p=0.242<br>CLES:51.63% | U=545518.0<br>**$p=2.094 \times 10^{-26}$**<br>CLES:64.45% | U=710400.5<br>**$p=4.992 \times 10^{-26}$**<br>CLES:63.25% |
| Golden Hamster | U=324886.0<br>p=0.826<br>CLES:49.68% | U=496203.0<br>**$p=4.678 \times 10^{-31}$**<br>CLES:66.24% | U=649200.5<br>**$p=3.826 \times 10^{-38}$**<br>CLES:66.81% |
| Armadillo | U=339549.0<br>p=0.229<br>CLES:51.73% | U=454927.0<br>**$p=2.311 \times 10^{-26}$**<br>CLES:65.11% | U=587981.0<br>**$p=2.063 \times 10^{-25}$**<br>CLES:63.73% |
| Ugandan red Colobus | U=402325.0<br>p=0.238<br>CLES:51.63% | U=563640.0<br>**$p=5.723 \times 10^{-27}$**<br>CLES:64.49% | U=734391.0<br>**$p=6.012 \times 10^{-29}$**<br>CLES:63.94% |
| Pig-tailed macaque | U=431122.5<br>$p=6.002 \times 10^{-2}$<br>CLES:52.56% | U=584672.5<br>**$p=4.917 \times 10^{-25}$**<br>CLES:63.75% | U=736330.0<br>**$p=2.657 \times 10^{-21}$**<br>CLES:61.71% |

| | | | |
|---|---|---|---|
| Panda | U=388909.0<br>p=0.475<br>CLES:50.99% | U=538844.0<br>**p=1.101 × $10^{-28}$**<br>CLES:65.16% | U=700208.0<br>**p=3.619 × $10^{-32}$**<br>CLES:64.97% |
| Ferret | U=371377.0<br>p=0.839<br>CLES:50.29% | U=519435.5<br>**p=6.458 × $10^{-26}$**<br>CLES:64.47% | U=689967.0<br>**p=2.717 × $10^{-30}$**<br>CLES:64.53% |
| Koala | U=317615.0<br>p=0.134<br>CLES:52.20% | U=441730.0<br>**p=1.519 × $10^{-20}$**<br>CLES:63.25% | U=551462.0<br>**p=6.923 × $10^{-17}$**<br>CLES:61.07% |
| Wallaby | U=29389.0<br>p=0.728<br>CLES:50.93% | U=33067.0<br>p=4.876 × $10^{-4}$<br>CLES:59.34% | U=47751.0<br>p=2.75 × $10^{-4}$<br>CLES:58.81% |
| Tarsier | U=263469.0<br>p=0.79<br>CLES:50.41% | U=351438.5<br>**p=7.925 × $10^{-18}$**<br>CLES:62.93% | U=451978.0<br>**p=4.935 × $10^{-21}$**<br>CLES:63.22% |
| Degu | U=302916.0<br>p=0.139<br>CLES:52.20% | U=456793.0<br>**p=5.554 × $10^{-35}$**<br>CLES:67.82% | U=608299.5<br>**p=1.886 × $10^{-34}$**<br>CLES:66.16% |
| Chinese hamster CHOK1GS | U=404779.0<br>p=0.53<br>CLES:50.86% | U=593943.0<br>**p=1.639 × $10^{-34}$**<br>CLES:66.42% | U=767904.0<br>**p=4.678 × $10^{-37}$**<br>CLES:65.82% |
| Drill | U=382336.0<br>p=0.53<br>CLES:50.88% | U=504269.0<br>**p=3.638 × $10^{-18}$**<br>CLES:61.91% | U=669844.0<br>**p=6.366 × $10^{-20}$**<br>CLES:61.56% |
| Gibbon | U=382947.0<br>p=0.523<br>CLES:50.89% | U=499052.5<br>**p=9.238 × $10^{-14}$**<br>CLES:60.17% | U=653177.0<br>**p=5.218 × $10^{-14}$**<br>CLES:59.50% |
| Rat | U=360483.5<br>p=0.864<br>CLES:49.76% | U=533799.0<br>**p=1.846 × $10^{-30}$**<br>CLES:65.76% | U=710304.0<br>**p=5.038 × $10^{-37}$**<br>CLES:66.14% |
| Macaque | U=406454.5<br>p=0.95<br>CLES:49.91% | U=566878.5<br>**p=2.614 × $10^{-21}$**<br>CLES:62.66% | U=753787.5<br>**p=1.169 × $10^{-26}$**<br>CLES:63.21% |
| American mink | U=367417.5<br>p=0.609<br>CLES:50.72% | U=505317.5<br>**p=2.196 × $10^{-26}$**<br>CLES:64.71% | U=658784.5<br>**p=9.04 × $10^{-29}$**<br>CLES:64.29% |
| Gelada | U=413106.5<br>p=0.81<br>CLES:50.33% | U=582662.5<br>**p=6.161 × $10^{-23}$**<br>CLES:63.12% | U=775774.5<br>**p=9.913 × $10^{-28}$**<br>CLES:63.41% |
| Sheep | U=346261.0<br>p=0.783<br>CLES:50.39% | U=497071.5<br>**p=3.786 × $10^{-25}$**<br>CLES:64.39% | U=638972.0<br>**p=1.269 × $10^{-28}$**<br>CLES:64.37% |
| Tasmanian devil | U=272312.0<br>p=0.399<br>CLES:51.28% | U=358190.5<br>**p=1.395 × $10^{-17}$**<br>CLES:62.77% | U=457708.0<br>**p=4.816 × $10^{-17}$**<br>CLES:61.68% |
| Elephant | U=361367.5<br>p=0.654<br>CLES:50.63% | U=514008.5<br>**p=4.998 × $10^{-26}$**<br>CLES:64.54% | U=657710.0<br>**p=1.718 × $10^{-28}$**<br>CLES:64.22% |

| | | | |
|---|---|---|---|
| Algerian mouse | U=393037.5<br>p=0.821<br>CLES:49.69% | U=585835.0<br>**p=6.165 × $10^{-31}$**<br>CLES:65.51% | U=775785.5<br>**p=6.548 × $10^{-39}$**<br>CLES:66.21% |
| Dingo | U=370907.0<br>p=0.352<br>CLES:51.31% | U=524810.5<br>**p=3.085 × $10^{-27}$**<br>CLES:64.83% | U=658814.5<br>**p=3.722 × $10^{-27}$**<br>CLES:63.84% |
| Golden snub-nosed monkey | U=400511.0<br>p=0.298<br>CLES:51.44% | U=533145.5<br>**p=1.717 × $10^{-23}$**<br>CLES:63.56% | U=669818.0<br>**p=4.717 × $10^{-23}$**<br>CLES:62.55% |
| Horse | U=392360.5<br>p=0.235<br>CLES:51.65% | U=575096.5<br>**p=8.652 × $10^{-32}$**<br>CLES:65.83% | U=731979.0<br>**p=3.586 × $10^{-31}$**<br>CLES:64.55% |
| Black snub-nosed monkey | U=390388.0<br>p=0.282<br>CLES:51.49% | U=509179.5<br>**p=1.631 × $10^{-15}$**<br>CLES:60.84% | U=648287.0<br>**p=1.017 × $10^{-14}$**<br>CLES:59.80% |
| Brazilian guinea pig | U=197667.0<br>p=0.472<br>CLES:51.19% | U=266711.5<br>**p=1.991 × $10^{-18}$**<br>CLES:64.23% | U=361497.0<br>**p=1.237 × $10^{-18}$**<br>CLES:63.08% |
| Orangutan | U=382891.0<br>p=0.81<br>CLES:50.33% | U=490680.5<br>**p=4.815 × $10^{-10}$**<br>CLES:58.46% | U=640679.0<br>**p=2.548 × $10^{-11}$**<br>CLES:58.42% |
| Northern American deer mouse | U=377066.5<br>p=0.904<br>CLES:49.83% | U=534666.5<br>**p=1.979 × $10^{-30}$**<br>CLES:65.70% | U=700691.0<br>**p=1.539 × $10^{-36}$**<br>CLES:66.07% |
| Naked mole-rat female | U=361766.5<br>p=0.392<br>CLES:51.21% | U=530163.0<br>**p=1.069 × $10^{-34}$**<br>CLES:67.00% | U=705676.0<br>**p=1.47 × $10^{-38}$**<br>CLES:66.54% |
| Mouse | U=422037.0<br>p=0.854<br>CLES:49.75% | U=629188.0<br>**p=9.74 × $10^{-33}$**<br>CLES:65.69% | U=818061.5<br>**p=5.709 × $10^{-40}$**<br>CLES:66.22% |
| Daurian ground squirrel | U=275277.5<br>p=0.36<br>CLES:51.39% | U=374655.0<br>**p=6.612 × $10^{-26}$**<br>CLES:65.71% | U=451445.5<br>**p=3.712 × $10^{-26}$**<br>CLES:64.97% |
| Tiger | U=248672.0<br>p=0.788<br>CLES:50.42% | U=338295.5<br>**p=4.339 × $10^{-23}$**<br>CLES:65.14% | U=433265.5<br>**p=4.361 × $10^{-27}$**<br>CLES:65.43% |
| Lesser hedgehog tenrec | U=24035.5<br>p=0.146<br>CLES:54.10% | U=28174.0<br>**p=2.327 × $10^{-10}$**<br>CLES:68.18% | U=31840.0<br>**p=2.815 × $10^{-8}$**<br>CLES:65.28% |
| Mongolian gerbil | U=310338.5<br>p=0.842<br>CLES:49.71% | U=427301.5<br>**p=5.08 × $10^{-25}$**<br>CLES:64.90% | U=556343.5<br>**p=6.198 × $10^{-31}$**<br>CLES:65.56% |
| Ma's night monkey | U=408078.0<br>p=8.182 × $10^{-3}$<br>CLES:53.68% | U=567103.0<br>**p=6.331 × $10^{-35}$**<br>CLES:66.74% | U=712133.0<br>**p=2.226 × $10^{-26}$**<br>CLES:63.34% |
| Upper Galilee mountains blind mole rat | U=359114.0<br>p=0.506<br>CLES:50.94% | U=540364.5<br>**p=1.628 × $10^{-33}$**<br>CLES:66.60% | U=711669.0<br>**p=5.754 × $10^{-37}$**<br>CLES:66.12% |

| | | | |
|---|---|---|---|
| Kangaroo rat | U=292269.5<br>p=0.918<br>CLES:50.15% | U=436828.0<br>**p=8.643 × $10^{-25}$**<br>CLES:64.81% | U=573608.5<br>**p=1.458 × $10^{-28}$**<br>CLES:64.78% |
| Chimpanzee | U=382454.0<br>p=0.519<br>CLES:50.90% | U=493630.0<br>**p=1.449 × $10^{-9}$**<br>CLES:58.16% | U=627255.5<br>**p=9.521 × $10^{-10}$**<br>CLES:57.71% |
| Guinea Pig | U=363336.5<br>p=0.982<br>CLES:50.03% | U=522035.0<br>**p=1.288 × $10^{-28}$**<br>CLES:65.27% | U=679005.5<br>**p=1.991 × $10^{-35}$**<br>CLES:65.93% |
| American beaver | U=344136.5<br>p=0.406<br>CLES:51.19% | U=465527.5<br>**p=4.332 × $10^{-27}$**<br>CLES:65.23% | U=586724.0<br>**p=8.033 × $10^{-28}$**<br>CLES:64.46% |
| Wild yak | U=300065.0<br>p=0.768<br>CLES:50.44% | U=402398.0<br>**p=1.047 × $10^{-24}$**<br>CLES:65.01% | U=510129.5<br>**p=4.478 × $10^{-29}$**<br>CLES:65.38% |
| Coquerel's sifaka | U=369415.5<br>p=0.153<br>CLES:52.02% | U=532779.0<br>**p=8.777 × $10^{-32}$**<br>CLES:66.14% | U=678859.5<br>**p=4.634 × $10^{-31}$**<br>CLES:64.81% |
| Alpaca | U=11348.0<br>p=0.867<br>CLES:49.44% | U=15865.0<br>**p=2.899 × $10^{-7}$**<br>CLES:66.91% | U=15719.0<br>**p=4.025 × $10^{-7}$**<br>CLES:66.73% |
| Lesser Egyptian jerboa | U=292564.0<br>p=0.875<br>CLES:49.77% | U=448199.0<br>**p=1.103 × $10^{-30}$**<br>CLES:66.60% | U=598276.0<br>**p=6.295 × $10^{-37}$**<br>CLES:66.88% |
| Long-tailed chinchilla | U=353052.5<br>p=0.637<br>CLES:49.33% | U=519945.0<br>**p=6.081 × $10^{-27}$**<br>CLES:64.81% | U=703586.0<br>**p=2.204 × $10^{-36}$**<br>CLES:66.02% |
| Angola colobus | U=405857.0<br>p=7.187 × $10^{-2}$<br>CLES:52.49% | U=547432.0<br>**p=1.758 × $10^{-25}$**<br>CLES:64.12% | U=687630.5<br>**p=7.812 × $10^{-23}$**<br>CLES:62.40% |
| Ryukyu mouse | U=385678.0<br>p=0.934<br>CLES:49.89% | U=585095.5<br>**p=3.069 × $10^{-31}$**<br>CLES:65.59% | U=755211.0<br>**p=5.426 × $10^{-38}$**<br>CLES:66.12% |
| Chinese hamster PICR | U=401842.0<br>p=0.423<br>CLES:51.10% | U=590995.0<br>**p=3.382 × $10^{-37}$**<br>CLES:67.13% | U=747057.0<br>**p=1.03 × $10^{-38}$**<br>CLES:66.32% |
| Vervet-AGM | U=422427.0<br>p=0.441<br>CLES:51.05% | U=572871.5<br>**p=2.135 × $10^{-22}$**<br>CLES:62.97% | U=736413.0<br>**p=3.335 × $10^{-23}$**<br>CLES:62.29% |
| Cow | U=377830.5<br>p=0.646<br>CLES:50.64% | U=548856.5<br>**p=3.268 × $10^{-29}$**<br>CLES:65.26% | U=710264.5<br>**p=2.667 × $10^{-33}$**<br>CLES:65.21% |
| Bushbaby | U=389995.0<br>p=0.67<br>CLES:50.59% | U=576229.0<br>**p=3.117 × $10^{-33}$**<br>CLES:66.22% | U=756947.5<br>**p=5.283 × $10^{-37}$**<br>CLES:65.86% |
| Bonobo | U=381671.5<br>p=0.787<br>CLES:50.38% | U=453046.5<br>p=3.843 × $10^{-4}$<br>CLES:54.83% | U=589029.0<br>p=1.621 × $10^{-4}$<br>CLES:54.77% |

| | | | |
|---|---|---|---|
| Hyrax | U=30392.0<br>p=0.569<br>CLES:48.52% | U=34965.0<br>$p=1.296 \times 10^{-3}$<br>CLES:58.44% | U=48326.5<br>**$p=2.849 \times 10^{-5}$**<br>CLES:60.16% |
| Rabbit | U=299963.5<br>p=0.272<br>CLES:51.63% | U=399104.0<br>**$p=8.063 \times 10^{-18}$**<br>CLES:62.52% | U=509522.5<br>**$p=1.894 \times 10^{-16}$**<br>CLES:61.13% |
| Megabat | U=105013.0<br>p=0.985<br>CLES:50.04% | U=125586.5<br>**$p=2.403 \times 10^{-11}$**<br>CLES:62.95% | U=166663.0<br>**$p=1.249 \times 10^{-13}$**<br>CLES:63.37% |
| Dog | U=364818.0<br>p=0.173<br>CLES:51.93% | U=500602.0<br>**$p=4.022 \times 10^{-25}$**<br>CLES:64.36% | U=639514.5<br>**$p=1.639 \times 10^{-22}$**<br>CLES:62.54% |
| Steppe mouse | U=377383.5<br>p=0.596<br>CLES:50.74% | U=545821.5<br>**$p=6.81 \times 10^{-32}$**<br>CLES:66.07% | U=716518.0<br>**$p=2.859 \times 10^{-34}$**<br>CLES:65.42% |
| Greater bamboo lemur | U=407329.0<br>p=0.141<br>CLES:52.03% | U=573282.0<br>**$p=1.057 \times 10^{-32}$**<br>CLES:66.06% | U=723512.0<br>**$p=2.527 \times 10^{-31}$**<br>CLES:64.62% |
| Pig | U=377095.0<br>p=0.908<br>CLES:50.16% | U=562100.0<br>**$p=6.218 \times 10^{-33}$**<br>CLES:66.22% | U=725447.5<br>**$p=2.615 \times 10^{-38}$**<br>CLES:66.36% |
| Tree Shrew | U=16626.0<br>p=0.902<br>CLES:50.38% | U=17703.0<br>**$p=5.376 \times 10^{-5}$**<br>CLES:62.74% | U=19594.0<br>$p=1.692 \times 10^{-4}$<br>CLES:61.53% |
| Pika | U=43754.5<br>$p=5.964 \times 10^{-2}$<br>CLES:45.60% | U=53851.0<br>**$p=2.208 \times 10^{-6}$**<br>CLES:61.26% | U=76851.0<br>**$p=2.638 \times 10^{-12}$**<br>CLES:65.45% |
| Shrew | U=12372.5<br>p=0.292<br>CLES:53.51% | U=14574.0<br>**$p=2.071 \times 10^{-5}$**<br>CLES:64.20% | U=15844.0<br>$p=1.815 \times 10^{-3}$<br>CLES:60.01% |
| Squirrel | U=381565.5<br>p=0.124<br>CLES:52.16% | U=565383.5<br>**$p=7.252 \times 10^{-35}$**<br>CLES:66.74% | U=710381.0<br>**$p=8.942 \times 10^{-34}$**<br>CLES:65.34% |
| Gorilla | U=394924.0<br>p=0.54<br>CLES:50.85% | U=504305.5<br>**$p=1.772 \times 10^{-10}$**<br>CLES:58.61% | U=657216.5<br>**$p=7.075 \times 10^{-11}$**<br>CLES:58.15% |
| Cat | U=405143.5<br>$p=8.307 \times 10^{-2}$<br>CLES:52.40% | U=571603.0<br>**$p=6.309 \times 10^{-34}$**<br>CLES:66.41% | U=718706.5<br>**$p=1.691 \times 10^{-31}$**<br>CLES:64.70% |

### Human Reference Genome, Zeisel et al., 2015

| MWU Test | Glia vs Vasculature | Glia vs Neuron | Vasculature vs Neuron |
|---|---|---|---|
| Microbat | U=199887.0<br>$p=6.764 \times 10^{-3}$<br>CLES:54.80% | U=421742.5<br>**$p=5.302 \times 10^{-17}$**<br>CLES:62.05% | U=137872.0<br>**$p=8.883 \times 10^{-5}$**<br>CLES:57.41% |
| Hedgehog | U=10315.0<br>p=0.196<br>CLES:54.86% | U=20075.0<br>$p=1.054 \times 10^{-2}$<br>CLES:57.77% | U=6068.0<br>p=0.519<br>CLES:52.61% |
| Naked mole-rat male | U=188884.5<br>$p=1.28 \times 10^{-3}$<br>CLES:55.84% | U=434967.0<br>**$p=3.366 \times 10^{-28}$**<br>CLES:65.94% | U=136226.5<br>**$p=1.635 \times 10^{-7}$**<br>CLES:60.07% |
| Sooty mangabey | U=264281.5<br>$p=9.285 \times 10^{-4}$<br>CLES:55.52% | U=600857.5<br>**$p=1.78 \times 10^{-20}$**<br>CLES:62.19% | U=184956.5<br>$p=1.555 \times 10^{-4}$<br>CLES:56.66% |
| Olive baboon | U=256189.0<br>$p=1.67 \times 10^{-2}$<br>CLES:54.00% | U=585109.5<br>**$p=1.24 \times 10^{-16}$**<br>CLES:60.90% | U=182581.0<br>**$p=9.248 \times 10^{-5}$**<br>CLES:56.92% |
| American bison | U=183070.0<br>$p=7.917 \times 10^{-4}$<br>CLES:56.15% | U=401901.5<br>**$p=2.67 \times 10^{-26}$**<br>CLES:65.69% | U=123379.0<br>**$p=8.777 \times 10^{-7}$**<br>CLES:59.65% |
| Leopard | U=232394.0<br>$p=1.815 \times 10^{-3}$<br>CLES:55.38% | U=577696.0<br>**$p=2.339 \times 10^{-29}$**<br>CLES:65.08% | U=175766.5<br>**$p=8.391 \times 10^{-8}$**<br>CLES:59.72% |
| Bolivian squirrel monkey | U=233584.5<br>$p=2.001 \times 10^{-2}$<br>CLES:53.97% | U=561865.0<br>**$p=2.521 \times 10^{-25}$**<br>CLES:63.99% | U=179292.5<br>**$p=1.318 \times 10^{-8}$**<br>CLES:60.25% |
| Crab-eating macaque | U=260459.0<br>$p=4.342 \times 10^{-3}$<br>CLES:54.76% | U=591261.5<br>**$p=9.369 \times 10^{-20}$**<br>CLES:62.00% | U=182619.0<br>**$p=4.601 \times 10^{-5}$**<br>CLES:57.21% |
| Red fox | U=224063.5<br>$p=3.216 \times 10^{-3}$<br>CLES:55.11% | U=513535.5<br>**$p=5.054 \times 10^{-24}$**<br>CLES:63.92% | U=161109.5<br>**$p=1.198 \times 10^{-6}$**<br>CLES:58.92% |
| Shrew mouse | U=249899.5<br>$p=3.574 \times 10^{-3}$<br>CLES:54.91% | U=599139.0<br>**$p=4.345 \times 10^{-30}$**<br>CLES:65.16% | U=188955.5<br>**$p=3.115 \times 10^{-9}$**<br>CLES:60.55% |
| Arctic ground squirrel | U=244318.0<br>$p=5.641 \times 10^{-3}$<br>CLES:54.66% | U=594320.0<br>**$p=1.856 \times 10^{-32}$**<br>CLES:65.82% | U=195977.0<br>**$p=2.116 \times 10^{-10}$**<br>CLES:61.24% |
| Chinese hamster CriGri | U=214135.0<br>$p=4.042 \times 10^{-3}$<br>CLES:55.03% | U=496396.5<br>**$p=2.104 \times 10^{-25}$**<br>CLES:64.50% | U=156822.0<br>**$p=2.959 \times 10^{-7}$**<br>CLES:59.51% |
| Opossum | U=189815.0<br>$p=2.28 \times 10^{-3}$<br>CLES:55.51% | U=429348.5<br>**$p=6.265 \times 10^{-18}$**<br>CLES:62.33% | U=135699.0<br>$p=2.79 \times 10^{-4}$<br>CLES:56.92% |
| Dolphin | U=63792.5<br>$p=6.262 \times 10^{-2}$ | U=155687.0<br>**$p=3.226 \times 10^{-16}$** | U=47273.0<br>**$p=1.192 \times 10^{-5}$** |

| | | | |
|---|---|---|---|
| | CLES:54.43% | CLES:65.25% | CLES:61.06% |
| Capuchin | U=242510.0<br>p=$1.982 \times 10^{-2}$<br>CLES:53.94% | U=585506.5<br>**p=$3.592 \times 10^{-28}$**<br>CLES:64.70% | U=188503.5<br>**p=$8.535 \times 10^{-10}$**<br>CLES:60.95% |
| Mouse Lemur | U=236835.0<br>p=$6.281 \times 10^{-3}$<br>CLES:54.67% | U=576836.0<br>**p=$3.554 \times 10^{-31}$**<br>CLES:65.63% | U=180656.5<br>**p=$3.535 \times 10^{-10}$**<br>CLES:61.34% |
| Damara mole rat | U=208782.0<br>p=$3.874 \times 10^{-3}$<br>CLES:55.09% | U=490102.5<br>**p=$8.651 \times 10^{-26}$**<br>CLES:64.65% | U=156861.5<br>**p=$1.167 \times 10^{-7}$**<br>CLES:59.85% |
| Marmoset | U=249840.0<br>p=$7.615 \times 10^{-4}$<br>CLES:55.68% | U=593686.0<br>**p=$3.396 \times 10^{-29}$**<br>CLES:64.92% | U=190130.0<br>**p=$3.379 \times 10^{-8}$**<br>CLES:59.79% |
| Goat | U=237840.0<br>p=$1.412 \times 10^{-2}$<br>CLES:54.19% | U=571859.5<br>**p=$1.774 \times 10^{-26}$**<br>CLES:64.30% | U=178098.0<br>**p=$5.388 \times 10^{-9}$**<br>CLES:60.55% |
| Common wombat | U=171213.0<br>p=$1.964 \times 10^{-2}$<br>CLES:54.29% | U=405749.0<br>**p=$1.083 \times 10^{-21}$**<br>CLES:63.95% | U=132989.0<br>**p=$7.018 \times 10^{-7}$**<br>CLES:59.61% |
| American black bear | U=192098.0<br>p=$4.207 \times 10^{-2}$<br>CLES:53.64% | U=425044.5<br>**p=$2.415 \times 10^{-20}$**<br>CLES:63.36% | U=135568.0<br>**p=$1.421 \times 10^{-7}$**<br>CLES:60.11% |
| Sloth | U=4335.5<br>p=0.759<br>CLES:51.44% | U=9619.0<br>p=$1.753 \times 10^{-4}$<br>CLES:64.28% | U=2615.0<br>p=$2.857 \times 10^{-2}$<br>CLES:61.34% |
| Polar bear | U=187975.0<br>p=$9.058 \times 10^{-3}$<br>CLES:54.74% | U=418673.5<br>**p=$3.824 \times 10^{-21}$**<br>CLES:63.72% | U=126782.0<br>**p=$3.432 \times 10^{-6}$**<br>CLES:59.04% |
| Alpine marmot | U=219754.0<br>p=$8.281 \times 10^{-3}$<br>CLES:54.57% | U=507541.5<br>**p=$5.117 \times 10^{-27}$**<br>CLES:64.92% | U=165686.5<br>**p=$1.575 \times 10^{-8}$**<br>CLES:60.36% |
| Prairie vole | U=231675.0<br>p=$5.618 \times 10^{-4}$<br>CLES:55.96% | U=566116.5<br>**p=$4.751 \times 10^{-31}$**<br>CLES:65.65% | U=174496.5<br>**p=$5.171 \times 10^{-8}$**<br>CLES:59.89% |
| Donkey | U=254715.0<br>**p=$9.635 \times 10^{-5}$**<br>CLES:56.60% | U=576203.0<br>**p=$1.242 \times 10^{-22}$**<br>CLES:63.04% | U=175056.0<br>p=$1.87 \times 10^{-4}$<br>CLES:56.67% |
| Golden Hamster | U=205250.5<br>p=$3.133 \times 10^{-2}$<br>CLES:53.79% | U=516372.5<br>**p=$9.135 \times 10^{-28}$**<br>CLES:65.05% | U=167072.5<br>**p=$3.278 \times 10^{-10}$**<br>CLES:61.63% |
| Armadillo | U=209102.5<br>p=$5.99 \times 10^{-3}$<br>CLES:54.85% | U=489186.0<br>**p=$1.011 \times 10^{-24}$**<br>CLES:64.33% | U=152920.5<br>**p=$3.734 \times 10^{-7}$**<br>CLES:59.49% |
| Ugandan red Colobus | U=238809.0<br>p=$2.303 \times 10^{-3}$<br>CLES:55.23% | U=578297.5<br>**p=$4.31 \times 10^{-23}$**<br>CLES:63.17% | U=173336.0<br>**p=$1.186 \times 10^{-5}$**<br>CLES:57.91% |
| Pig-tailed macaque | U=256758.0<br>p=$1.437 \times 10^{-2}$ | U=592343.0<br>**p=$3.854 \times 10^{-19}$** | U=188006.0<br>**p=$1.608 \times 10^{-5}$** |

| | | | |
|---|---|---|---|
| | CLES:54.08% | CLES:61.77% | CLES:57.60% |
| Panda | U=236027.5<br>$p=5.297 \times 10^{-3}$<br>CLES:54.77% | U=550095.5<br>**$p=3.068 \times 10^{-24}$**<br>CLES:63.74% | U=170258.0<br>**$p=9.941 \times 10^{-7}$**<br>CLES:58.88% |
| Ferret | U=239026.0<br>$p=1.385 \times 10^{-3}$<br>CLES:55.47% | U=555739.0<br>**$p=1.181 \times 10^{-25}$**<br>CLES:64.15% | U=172308.5<br>**$p=5.931 \times 10^{-7}$**<br>CLES:59.04% |
| Koala | U=207271.0<br>$p=1.35 \times 10^{-4}$<br>CLES:56.79% | U=483805.0<br>**$p=1.683 \times 10^{-23}$**<br>CLES:63.93% | U=149473.0<br>**$p=5.651 \times 10^{-5}$**<br>CLES:57.53% |
| Wallaby | U=16955.0<br>p=0.242<br>CLES:53.78% | U=35837.0<br>**$p=7.816 \times 10^{-6}$**<br>CLES:61.86% | U=13286.0<br>$p=1.1 \times 10^{-2}$<br>CLES:58.67% |
| Tarsier | U=161480.0<br>$p=9.552 \times 10^{-3}$<br>CLES:54.91% | U=377256.5<br>**$p=8.995 \times 10^{-17}$**<br>CLES:62.31% | U=109748.0<br>$p=5.755 \times 10^{-4}$<br>CLES:56.93% |
| Degu | U=182301.0<br>$p=2.469 \times 10^{-2}$<br>CLES:54.07% | U=433837.0<br>**$p=5.47 \times 10^{-24}$**<br>CLES:64.52% | U=142243.0<br>**$p=2.662 \times 10^{-8}$**<br>CLES:60.63% |
| Chinese hamster CHOK1GS | U=251544.5<br>$p=4.847 \times 10^{-3}$<br>CLES:54.73% | U=605314.0<br>**$p=3.909 \times 10^{-32}$**<br>CLES:65.68% | U=194789.0<br>**$p=4.715 \times 10^{-10}$**<br>CLES:61.02% |
| Drill | U=244228.0<br>$p=7.745 \times 10^{-4}$<br>CLES:55.74% | U=540012.5<br>**$p=1.981 \times 10^{-16}$**<br>CLES:61.06% | U=161039.0<br>$p=3.739 \times 10^{-3}$<br>CLES:55.25% |
| Gibbon | U=249431.0<br>$p=3.291 \times 10^{-4}$<br>CLES:56.09% | U=532866.0<br>**$p=2.252 \times 10^{-13}$**<br>CLES:59.84% | U=162658.5<br>$p=1.851 \times 10^{-2}$<br>CLES:54.23% |
| Rat | U=233967.0<br>$p=5.867 \times 10^{-3}$<br>CLES:54.73% | U=563070.0<br>**$p=2.107 \times 10^{-26}$**<br>CLES:64.32% | U=175708.0<br>**$p=2.871 \times 10^{-8}$**<br>CLES:60.06% |
| Macaque | U=257190.5<br>$p=6.783 \times 10^{-4}$<br>CLES:55.70% | U=574340.5<br>**$p=6.345 \times 10^{-18}$**<br>CLES:61.42% | U=177993.5<br>$p=1.136 \times 10^{-3}$<br>CLES:55.76% |
| American mink | U=227610.0<br>$p=2.663 \times 10^{-4}$<br>CLES:56.33% | U=524410.0<br>**$p=3.06 \times 10^{-26}$**<br>CLES:64.55% | U=160537.5<br>**$p=5.972 \times 10^{-6}$**<br>CLES:58.32% |
| Gelada | U=266482.5<br>$p=1.789 \times 10^{-4}$<br>CLES:56.27% | U=606396.0<br>**$p=4.857 \times 10^{-22}$**<br>CLES:62.68% | U=181982.0<br>$p=4.102 \times 10^{-4}$<br>CLES:56.24% |
| Sheep | U=222905.0<br>$p=3.649 \times 10^{-3}$<br>CLES:55.07% | U=529462.5<br>**$p=3.436 \times 10^{-26}$**<br>CLES:64.52% | U=159721.5<br>**$p=1.281 \times 10^{-7}$**<br>CLES:59.78% |
| Tasmanian devil | U=178591.5<br>$p=4.496 \times 10^{-4}$<br>CLES:56.45% | U=373413.0<br>**$p=3.943 \times 10^{-17}$**<br>CLES:62.49% | U=117008.5<br>$p=2.744 \times 10^{-3}$<br>CLES:55.86% |
| Elephant | U=232955.5<br>$p=5.426 \times 10^{-3}$ | U=537456.0<br>**$p=1.55 \times 10^{-22}$** | U=165647.5<br>**$p=2.06 \times 10^{-6}$** |

| | | | |
|---|---|---|---|
| | CLES:54.78% | CLES:63.28% | CLES:58.66% |
| Algerian mouse | U=255236.0<br>$p=6.375 \times 10^{-4}$<br>CLES:55.75% | U=618460.5<br>**$p=1.788 \times 10^{-33}$**<br>CLES:65.95% | U=192015.0<br>**$p=4.806 \times 10^{-9}$**<br>CLES:60.39% |
| Dingo | U=219609.5<br>$p=1.144 \times 10^{-2}$<br>CLES:54.40% | U=530027.0<br>**$p=5.592 \times 10^{-26}$**<br>CLES:64.42% | U=166148.5<br>**$p=2.882 \times 10^{-8}$**<br>CLES:60.20% |
| Golden snub-nosed monkey | U=235308.0<br>$p=1.346 \times 10^{-2}$<br>CLES:54.22% | U=548297.0<br>**$p=4.338 \times 10^{-20}$**<br>CLES:62.36% | U=171105.0<br>**$p=6.17 \times 10^{-6}$**<br>CLES:58.17% |
| Horse | U=254840.0<br>$p=1.507 \times 10^{-4}$<br>CLES:56.39% | U=581084.0<br>**$p=5.701 \times 10^{-27}$**<br>CLES:64.37% | U=181989.5<br>**$p=1.08 \times 10^{-6}$**<br>CLES:58.69% |
| Black snub-nosed monkey | U=228564.0<br>$p=7.608 \times 10^{-2}$<br>CLES:53.05% | U=533003.0<br>**$p=9.106 \times 10^{-14}$**<br>CLES:60.01% | U=165181.0<br>$p=1.217 \times 10^{-4}$<br>CLES:56.98% |
| Brazilian guinea pig | U=129037.5<br>$p=9.253 \times 10^{-2}$<br>CLES:53.31% | U=281562.0<br>**$p=1.559 \times 10^{-12}$**<br>CLES:61.20% | U=93859.5<br>$p=1.501 \times 10^{-4}$<br>CLES:57.92% |
| Orangutan | U=249168.0<br>$p=5.524 \times 10^{-4}$<br>CLES:55.85% | U=540582.5<br>**$p=4.716 \times 10^{-17}$**<br>CLES:61.29% | U=166902.0<br>$p=2.137 \times 10^{-3}$<br>CLES:55.51% |
| Northern American deer mouse | U=238405.5<br>$p=1.592 \times 10^{-3}$<br>CLES:55.39% | U=553383.5<br>**$p=4.03 \times 10^{-32}$**<br>CLES:66.08% | U=175886.5<br>**$p=1.275 \times 10^{-9}$**<br>CLES:60.99% |
| Naked mole-rat female | U=209784.5<br>$p=1.25 \times 10^{-3}$<br>CLES:55.70% | U=501342.5<br>**$p=3.784 \times 10^{-30}$**<br>CLES:65.91% | U=156008.5<br>**$p=3.706 \times 10^{-8}$**<br>CLES:60.27% |
| Mouse | U=275350.0<br>$p=4.649 \times 10^{-4}$<br>CLES:55.78% | U=645569.5<br>**$p=4.078 \times 10^{-31}$**<br>CLES:65.15% | U=201097.0<br>**$p=3.859 \times 10^{-8}$**<br>CLES:59.60% |
| Daurian ground squirrel | U=178810.5<br>$p=3.05 \times 10^{-3}$<br>CLES:55.43% | U=380925.5<br>**$p=2.926 \times 10^{-23}$**<br>CLES:64.85% | U=121197.0<br>**$p=2.146 \times 10^{-6}$**<br>CLES:59.31% |
| Tiger | U=161133.5<br>$p=1.245 \times 10^{-3}$<br>CLES:56.14% | U=374809.5<br>**$p=1.643 \times 10^{-23}$**<br>CLES:64.98% | U=110253.0<br>**$p=9.342 \times 10^{-6}$**<br>CLES:58.97% |
| Lesser hedgehog tenrec | U=12142.0<br>p=0.84<br>CLES:50.72% | U=24430.0<br>$p=1.388 \times 10^{-3}$<br>CLES:59.36% | U=8080.0<br>$p=1.55 \times 10^{-2}$<br>CLES:59.34% |
| Mongolian gerbil | U=204877.0<br>$p=5.217 \times 10^{-4}$<br>CLES:56.18% | U=451481.0<br>**$p=2.05 \times 10^{-26}$**<br>CLES:65.24% | U=139289.5<br>**$p=1.196 \times 10^{-6}$**<br>CLES:59.24% |
| Ma's night monkey | U=231119.0<br>$p=2.522 \times 10^{-2}$<br>CLES:53.83% | U=561377.0<br>**$p=1.981 \times 10^{-24}$**<br>CLES:63.70% | U=180801.0<br>**$p=8.675 \times 10^{-9}$**<br>CLES:60.37% |

| | | | |
|---|---|---|---|
| Upper Galilee mountains blind mole rat | U=216659.0<br>$p=9.917 \times 10^{-3}$<br>CLES:54.51% | U=546405.0<br>**$p=3.614 \times 10^{-31}$**<br>CLES:65.84% | U=169673.0<br>**$p=7.439 \times 10^{-10}$**<br>CLES:61.33% |
| Kangaroo rat | U=184685.5<br>$p=2.721 \times 10^{-2}$<br>CLES:54.00% | U=445063.5<br>**$p=6.782 \times 10^{-24}$**<br>CLES:64.40% | U=143408.5<br>**$p=7.827 \times 10^{-9}$**<br>CLES:61.04% |
| Chimpanzee | U=233282.5<br>$p=5.82 \times 10^{-3}$<br>CLES:54.71% | U=512521.0<br>**$p=4.127 \times 10^{-11}$**<br>CLES:58.86% | U=160884.5<br>$p=1.131 \times 10^{-2}$<br>CLES:54.56% |
| Guinea Pig | U=231710.5<br>$p=1.539 \times 10^{-3}$<br>CLES:55.44% | U=539527.0<br>**$p=8.065 \times 10^{-26}$**<br>CLES:64.29% | U=172711.0<br>**$p=3.2 \times 10^{-7}$**<br>CLES:59.26% |
| American beaver | U=199419.0<br>$p=3.06 \times 10^{-2}$<br>CLES:53.83% | U=440847.0<br>**$p=2.564 \times 10^{-21}$**<br>CLES:63.57% | U=144076.0<br>**$p=2.624 \times 10^{-7}$**<br>CLES:59.72% |
| Wild yak | U=199322.5<br>$p=4.618 \times 10^{-4}$<br>CLES:56.31% | U=426158.0<br>**$p=9.784 \times 10^{-23}$**<br>CLES:64.25% | U=127075.0<br>**$p=1.751 \times 10^{-5}$**<br>CLES:58.32% |
| Coquerel's sifaka | U=227919.5<br>$p=4.981 \times 10^{-3}$<br>CLES:54.85% | U=552665.0<br>**$p=1.22 \times 10^{-27}$**<br>CLES:64.77% | U=172487.5<br>**$p=1.294 \times 10^{-8}$**<br>CLES:60.37% |
| Alpaca | U=8210.0<br>$p=6.955 \times 10^{-2}$<br>CLES:57.41% | U=21032.0<br>**$p=4.537 \times 10^{-11}$**<br>CLES:70.81% | U=5797.0<br>$p=2.377 \times 10^{-4}$<br>CLES:66.06% |
| Lesser Egyptian jerboa | U=192943.0<br>$p=3.381 \times 10^{-4}$<br>CLES:56.52% | U=484656.5<br>**$p=8.829 \times 10^{-31}$**<br>CLES:66.23% | U=143629.5<br>**$p=1.468 \times 10^{-7}$**<br>CLES:60.06% |
| Long-tailed chinchilla | U=231918.5<br>$p=5.01 \times 10^{-4}$<br>CLES:56.01% | U=552341.0<br>**$p=3.807 \times 10^{-29}$**<br>CLES:65.22% | U=169438.0<br>**$p=1.732 \times 10^{-7}$**<br>CLES:59.54% |
| Angola colobus | U=245527.5<br>$p=8.533 \times 10^{-3}$<br>CLES:54.46% | U=564435.5<br>**$p=2.522 \times 10^{-18}$**<br>CLES:61.64% | U=172232.0<br>$p=1.181 \times 10^{-4}$<br>CLES:56.91% |
| Ryukyu mouse | U=247417.0<br>$p=1.257 \times 10^{-3}$<br>CLES:55.46% | U=591086.5<br>**$p=2.259 \times 10^{-29}$**<br>CLES:65.00% | U=186225.5<br>**$p=4.426 \times 10^{-8}$**<br>CLES:59.76% |
| Chinese hamster PICR | U=243978.0<br>$p=4.187 \times 10^{-3}$<br>CLES:54.84% | U=596187.0<br>**$p=8.34 \times 10^{-33}$**<br>CLES:65.92% | U=193232.5<br>**$p=8.924 \times 10^{-11}$**<br>CLES:61.54% |
| Vervet-AGM | U=256268.0<br>$p=6.482 \times 10^{-3}$<br>CLES:54.57% | U=597984.0<br>**$p=9.887 \times 10^{-22}$**<br>CLES:62.63% | U=183792.0<br>**$p=7.574 \times 10^{-6}$**<br>CLES:57.94% |
| Cow | U=247401.0<br>$p=1.933 \times 10^{-3}$<br>CLES:55.25% | U=581059.0<br>**$p=2.827 \times 10^{-31}$**<br>CLES:65.65% | U=180140.0<br>**$p=4.812 \times 10^{-9}$**<br>CLES:60.53% |
| Bushbaby | U=251346.5<br>$p=4.848 \times 10^{-4}$<br>CLES:55.90% | U=596587.0<br>**$p=9.78 \times 10^{-33}$**<br>CLES:65.92% | U=186142.5<br>**$p=4.321 \times 10^{-9}$**<br>CLES:60.49% |

| | | | |
|---|---|---|---|
| Bonobo | U=224819.5<br>p=0.382<br>CLES:51.49% | U=492834.5<br>**p=$1.302 \times 10^{-5}$**<br>CLES:55.85% | U=160283.5<br>p=$1.019 \times 10^{-2}$<br>CLES:54.64% |
| Hyrax | U=20997.5<br>p=$4.709 \times 10^{-2}$<br>CLES:56.22% | U=40617.5<br>**p=$2.778 \times 10^{-7}$**<br>CLES:63.39% | U=13242.5<br>p=$2.923 \times 10^{-2}$<br>CLES:57.37% |
| Rabbit | U=193804.5<br>p=$5.799 \times 10^{-4}$<br>CLES:56.22% | U=438823.0<br>**p=$2.541 \times 10^{-20}$**<br>CLES:63.18% | U=135577.0<br>**p=$7.991 \times 10^{-5}$**<br>CLES:57.54% |
| Megabat | U=70534.0<br>p=0.145<br>CLES:53.31% | U=135012.0<br>**p=$3.609 \times 10^{-11}$**<br>CLES:62.75% | U=47801.0<br>**p=$9.083 \times 10^{-5}$**<br>CLES:59.65% |
| Dog | U=213264.0<br>p=$2.939 \times 10^{-2}$<br>CLES:53.80% | U=521691.5<br>**p=$9.679 \times 10^{-26}$**<br>CLES:64.40% | U=166272.0<br>**p=$5.652 \times 10^{-9}$**<br>CLES:60.73% |
| Steppe mouse | U=245000.5<br>p=$1.844 \times 10^{-4}$<br>CLES:56.37% | U=576835.0<br>**p=$1.089 \times 10^{-31}$**<br>CLES:65.76% | U=179091.0<br>**p=$1.051 \times 10^{-7}$**<br>CLES:59.56% |
| Greater bamboo lemur | U=251592.5<br>p=$2.392 \times 10^{-4}$<br>CLES:56.24% | U=590404.5<br>**p=$4.736 \times 10^{-28}$**<br>CLES:64.64% | U=177918.0<br>**p=$9.817 \times 10^{-7}$**<br>CLES:58.79% |
| Pig | U=252787.0<br>p=$2.535 \times 10^{-4}$<br>CLES:56.18% | U=580712.0<br>**p=$1.371 \times 10^{-30}$**<br>CLES:65.44% | U=181885.0<br>**p=$5.523 \times 10^{-8}$**<br>CLES:59.72% |
| Tree Shrew | U=6748.0<br>p=0.73<br>CLES:51.43% | U=15525.5<br>p=$1.254 \times 10^{-3}$<br>CLES:60.59% | U=4763.0<br>p=$3.216 \times 10^{-2}$<br>CLES:59.54% |
| Pika | U=34442.5<br>p=$1.281 \times 10^{-2}$<br>CLES:56.90% | U=52275.0<br>**p=$9.477 \times 10^{-6}$**<br>CLES:60.86% | U=16777.0<br>p=0.2<br>CLES:53.98% |
| Shrew | U=6344.0<br>p=0.258<br>CLES:54.86% | U=13957.0<br>p=$5.326 \times 10^{-3}$<br>CLES:59.34% | U=3909.0<br>p=0.258<br>CLES:55.21% |
| Squirrel | U=240704.5<br>p=$1.024 \times 10^{-3}$<br>CLES:55.59% | U=579553.0<br>**p=$9.994 \times 10^{-32}$**<br>CLES:65.74% | U=185668.0<br>**p=$3.516 \times 10^{-9}$**<br>CLES:60.57% |
| Gorilla | U=242273.5<br>p=$3.992 \times 10^{-2}$<br>CLES:53.46% | U=542238.0<br>**p=$8.36 \times 10^{-12}$**<br>CLES:59.08% | U=174424.0<br>p=$6.825 \times 10^{-4}$<br>CLES:56.04% |
| Cat | U=239651.5<br>p=$1.223 \times 10^{-2}$<br>CLES:54.27% | U=593826.0<br>**p=$1.811 \times 10^{-29}$**<br>CLES:65.02% | U=183677.5<br>**p=$1.573 \times 10^{-9}$**<br>CLES:60.86% |

### Human Reference Genome, Zeisel et al., 2018

| MWU Test | CNS_Glia vs CNS_Endothelia | CNS_Glia vs CNS_Neuron | CNS_Endothelia vs CNS_Neuron |
|---|---|---|---|
| Microbat | U=40034.0<br>p=0.461<br>CLES:47.90% | U=1238917.5<br>**$p=2.2 \times 10^{-35}$**<br>CLES:65.29% | U=225031.0<br>**$p=5.268 \times 10^{-10}$**<br>CLES:66.59% |
| Hedgehog | U=2429.0<br>p=0.539<br>CLES:46.59% | U=59760.0<br>**$p=7.892 \times 10^{-8}$**<br>CLES:63.89% | U=12783.0<br>$p=2.826 \times 10^{-3}$<br>CLES:65.43% |
| Naked mole-rat male | U=51142.5<br>p=0.8<br>CLES:50.68% | U=1332154.0<br>**$p=7.206 \times 10^{-47}$**<br>CLES:67.34% | U=251698.5<br>**$p=5.662 \times 10^{-11}$**<br>CLES:66.45% |
| Sooty mangabey | U=71888.5<br>$p=5.688 \times 10^{-2}$<br>CLES:54.79% | U=1723432.5<br>**$p=4.489 \times 10^{-33}$**<br>CLES:63.39% | U=296639.5<br>$p=1.942 \times 10^{-4}$<br>CLES:58.82% |
| Olive baboon | U=69918.0<br>p=0.111<br>CLES:54.02% | U=1742983.5<br>**$p=1.236 \times 10^{-38}$**<br>CLES:64.57% | U=303206.0<br>**$p=9.556 \times 10^{-6}$**<br>CLES:60.51% |
| American bison | U=41515.5<br>p=0.709<br>CLES:48.95% | U=1210821.5<br>**$p=1.028 \times 10^{-49}$**<br>CLES:68.40% | U=222640.0<br>**$p=2.868 \times 10^{-13}$**<br>CLES:69.38% |
| Leopard | U=60215.0<br>p=0.807<br>CLES:50.63% | U=1728978.5<br>**$p=1.897 \times 10^{-56}$**<br>CLES:68.02% | U=311180.0<br>**$p=9.973 \times 10^{-13}$**<br>CLES:67.37% |
| Bolivian squirrel monkey | U=59766.0<br>p=0.834<br>CLES:49.46% | U=1668410.0<br>**$p=4.831 \times 10^{-49}$**<br>CLES:66.77% | U=305482.0<br>**$p=8.395 \times 10^{-12}$**<br>CLES:66.54% |
| Crab-eating macaque | U=69306.0<br>p=0.181<br>CLES:53.39% | U=1724474.0<br>**$p=1.838 \times 10^{-35}$**<br>CLES:63.89% | U=297903.0<br>**$p=1.222 \times 10^{-5}$**<br>CLES:60.41% |
| Red fox | U=54227.0<br>p=0.624<br>CLES:48.71% | U=1530370.5<br>**$p=6.595 \times 10^{-39}$**<br>CLES:65.11% | U=281629.5<br>**$p=7.277 \times 10^{-11}$**<br>CLES:66.15% |
| Shrew mouse | U=63429.0<br>p=0.603<br>CLES:51.33% | U=1724314.5<br>**$p=6.88 \times 10^{-51}$**<br>CLES:67.03% | U=315532.5<br>**$p=7.476 \times 10^{-11}$**<br>CLES:65.61% |
| Arctic ground squirrel | U=63652.0<br>p=0.668<br>CLES:51.09% | U=1709207.0<br>**$p=8.319 \times 10^{-53}$**<br>CLES:67.41% | U=321058.5<br>**$p=4.116 \times 10^{-12}$**<br>CLES:66.52% |
| Chinese hamster CriGri | U=51757.0<br>p=0.716<br>CLES:49.03% | U=1452832.5<br>**$p=7.272 \times 10^{-43}$**<br>CLES:66.19% | U=271796.5<br>**$p=2.361 \times 10^{-11}$**<br>CLES:66.69% |
| Opossum | U=50461.0<br>p=0.137<br>CLES:54.07% | U=1208412.0<br>**$p=1.664 \times 10^{-20}$**<br>CLES:61.28% | U=208579.0<br>$p=9.65 \times 10^{-3}$<br>CLES:56.66% |
| Dolphin | U=17576.0<br>p=0.646 | U=445052.0<br>**$p=1.082 \times 10^{-21}$** | U=93419.0<br>**$p=9.457 \times 10^{-8}$** |

| | | | |
|---|---|---|---|
| | CLES:48.43% | CLES:65.10% | CLES:67.05% |
| Capuchin | U=64241.0 | U=1724802.0 | U=313045.5 |
| | p=0.817 | **p=$1.439 \times 10^{-47}$** | **p=$1.518 \times 10^{-10}$** |
| | CLES:50.59% | CLES:66.33% | CLES:65.31% |
| Mouse Lemur | U=63053.0 | U=1619940.5 | U=307140.5 |
| | p=0.279 | **p=$8.538 \times 10^{-52}$** | **p=$1.389 \times 10^{-10}$** |
| | CLES:52.77% | CLES:67.52% | CLES:65.35% |
| Damara mole rat | U=47618.0 | U=1453006.5 | U=260209.0 |
| | p=0.73 | **p=$7.699 \times 10^{-45}$** | **p=$1.025 \times 10^{-11}$** |
| | CLES:49.05% | CLES:66.67% | CLES:67.54% |
| Marmoset | U=64818.0 | U=1696478.0 | U=307081.0 |
| | p=0.337 | **p=$2.089 \times 10^{-49}$** | **p=$9.516 \times 10^{-10}$** |
| | CLES:52.45% | CLES:66.80% | CLES:64.67% |
| Goat | U=60215.0 | U=1636727.0 | U=294068.0 |
| | p=0.486 | **p=$3.366 \times 10^{-46}$** | **p=$8.211 \times 10^{-10}$** |
| | CLES:51.81% | CLES:66.37% | CLES:65.01% |
| Common wombat | U=43827.0 | U=1148956.5 | U=203376.0 |
| | p=0.388 | **p=$3.582 \times 10^{-25}$** | **p=$7.506 \times 10^{-5}$** |
| | CLES:52.44% | CLES:62.88% | CLES:60.50% |
| American black bear | U=47125.0 | U=1231206.5 | U=238858.0 |
| | p=0.838 | **p=$3.345 \times 10^{-29}$** | **p=$9.774 \times 10^{-9}$** |
| | CLES:49.44% | CLES:63.67% | CLES:64.60% |
| Sloth | U=802.0 | U=26411.0 | U=5065.0 |
| | p=0.308 | **p=$2.528 \times 10^{-5}$** | p=$2.73 \times 10^{-3}$ |
| | CLES:42.43% | CLES:63.36% | CLES:70.88% |
| Polar bear | U=41465.0 | U=1268206.5 | U=233517.0 |
| | p=0.447 | **p=$2.83 \times 10^{-43}$** | **p=$3.895 \times 10^{-12}$** |
| | CLES:47.87% | CLES:66.94% | CLES:68.33% |
| Alpine marmot | U=50848.0 | U=1519005.5 | U=298977.0 |
| | p=0.26 | **p=$6.141 \times 10^{-50}$** | **p=$2.956 \times 10^{-16}$** |
| | CLES:47.03% | CLES:67.47% | CLES:70.21% |
| Prairie vole | U=60435.0 | U=1589233.5 | U=301506.0 |
| | p=0.473 | **p=$9.749 \times 10^{-49}$** | **p=$4.123 \times 10^{-10}$** |
| | CLES:51.84% | CLES:67.06% | CLES:65.04% |
| Donkey | U=62582.0 | U=1665447.0 | U=293631.0 |
| | p=0.258 | **p=$7.154 \times 10^{-44}$** | **p=$1.672 \times 10^{-7}$** |
| | CLES:52.92% | CLES:65.88% | CLES:62.71% |
| Golden Hamster | U=55333.0 | U=1440895.5 | U=280130.0 |
| | p=0.922 | **p=$3.157 \times 10^{-42}$** | **p=$2.227 \times 10^{-10}$** |
| | CLES:50.26% | CLES:66.11% | CLES:65.49% |
| Armadillo | U=47423.0 | U=1383373.0 | U=252309.0 |
| | p=0.641 | **p=$1.894 \times 10^{-36}$** | **p=$2.428 \times 10^{-11}$** |
| | CLES:48.72% | CLES:64.99% | CLES:67.23% |
| Ugandan red Colobus | U=64948.0 | U=1689950.5 | U=291418.0 |
| | p=0.259 | **p=$4.949 \times 10^{-44}$** | **p=$1.589 \times 10^{-7}$** |
| | CLES:52.90% | CLES:65.73% | CLES:62.69% |
| Pig-tailed macaque | U=68841.5 | U=1719332.5 | U=308853.0 |
| | p=0.33 | **p=$6.64 \times 10^{-34}$** | **p=$4.409 \times 10^{-6}$** |

| | | | |
|---|---|---|---|
| | CLES:52.45% | CLES:63.59% | CLES:60.83% |
| Panda | U=57703.0<br>p=0.803<br>CLES:49.35% | U=1625121.5<br>**p=3.308 × $10^{-46}$**<br>CLES:66.42% | U=307618.0<br>**p=1.861 × $10^{-12}$**<br>CLES:67.11% |
| Ferret | U=58943.0<br>p=0.672<br>CLES:51.10% | U=1627839.0<br>**p=6.642 × $10^{-46}$**<br>CLES:66.36% | U=298062.5<br>**p=1.183 × $10^{-10}$**<br>CLES:65.75% |
| Koala | U=53284.0<br>p=0.401<br>CLES:52.25% | U=1366535.5<br>**p=5.094 × $10^{-25}$**<br>CLES:62.26% | U=245463.0<br>**p=8.376 × $10^{-5}$**<br>CLES:59.89% |
| Wallaby | U=5249.0<br>p=0.63<br>CLES:47.84% | U=94747.0<br>p=0.127<br>CLES:53.39% | U=24205.0<br>p=0.195<br>CLES:55.35% |
| Tarsier | U=42786.0<br>p=0.829<br>CLES:49.39% | U=1197316.5<br>**p=2.072 × $10^{-40}$**<br>CLES:66.43% | U=217543.5<br>**p=1.591 × $10^{-10}$**<br>CLES:66.91% |
| Degu | U=50313.5<br>p=0.305<br>CLES:52.77% | U=1279653.5<br>**p=1.329 × $10^{-42}$**<br>CLES:66.72% | U=240606.5<br>**p=7.454 × $10^{-9}$**<br>CLES:64.66% |
| Chinese hamster CHOK1GS | U=62082.0<br>p=0.767<br>CLES:49.25% | U=1704252.0<br>**p=1.24 × $10^{-47}$**<br>CLES:66.50% | U=332730.0<br>**p=3.3 × $10^{-13}$**<br>CLES:67.24% |
| Drill | U=61472.5<br>p=0.424<br>CLES:52.08% | U=1608746.0<br>**p=2.612 × $10^{-32}$**<br>CLES:63.45% | U=277359.0<br>**p=3.291 × $10^{-6}$**<br>CLES:61.41% |
| Gibbon | U=64101.5<br>p=0.139<br>CLES:53.84% | U=1612824.0<br>**p=2.866 × $10^{-32}$**<br>CLES:63.45% | U=278385.5<br>**p=2.142 × $10^{-5}$**<br>CLES:60.35% |
| Rat | U=57796.0<br>p=0.758<br>CLES:50.81% | U=1643856.5<br>**p=1.698 × $10^{-49}$**<br>CLES:67.02% | U=296516.0<br>**p=3.363 × $10^{-11}$**<br>CLES:66.32% |
| Macaque | U=68011.0<br>p=0.206<br>CLES:53.21% | U=1672235.5<br>**p=1.363 × $10^{-32}$**<br>CLES:63.38% | U=294210.0<br>**p=2.246 × $10^{-5}$**<br>CLES:60.10% |
| American mink | U=56741.0<br>p=0.968<br>CLES:50.10% | U=1584131.0<br>**p=1.285 × $10^{-49}$**<br>CLES:67.16% | U=294788.0<br>**p=1.718 × $10^{-12}$**<br>CLES:67.32% |
| Gelada | U=70971.5<br>p=9.852 × $10^{-2}$<br>CLES:54.16% | U=1799118.5<br>**p=1.116 × $10^{-48}$**<br>CLES:66.39% | U=314650.5<br>**p=1.199 × $10^{-7}$**<br>CLES:62.52% |
| Sheep | U=53216.0<br>p=0.702<br>CLES:48.98% | U=1563440.0<br>**p=5.784 × $10^{-45}$**<br>CLES:66.34% | U=288168.0<br>**p=1.061 × $10^{-12}$**<br>CLES:67.78% |
| Tasmanian devil | U=43680.0<br>p=0.679<br>CLES:51.16% | U=1129312.0<br>**p=2.332 × $10^{-22}$**<br>CLES:62.04% | U=202646.0<br>**p=4.144 × $10^{-5}$**<br>CLES:60.83% |
| Elephant | U=52693.0<br>p=0.935 | U=1539510.0<br>**p=5.704 × $10^{-42}$** | U=278983.0<br>**p=8.667 × $10^{-11}$** |

| | | | |
|---|---|---|---|
| | CLES:49.78% | CLES:65.83% | CLES:66.30% |
| Algerian mouse | U=63092.0<br>p=0.74<br>CLES:50.85% | U=1750252.0<br>**p=1.177 × $10^{-53}$**<br>CLES:67.51% | U=325602.0<br>**p=4.112 × $10^{-12}$**<br>CLES:66.57% |
| Dingo | U=56870.0<br>p=0.95<br>CLES:50.16% | U=1527599.5<br>**p=6.577 × $10^{-40}$**<br>CLES:65.43% | U=294582.0<br>**p=3.038 × $10^{-10}$**<br>CLES:65.30% |
| Golden snub-nosed monkey | U=67284.0<br>p=0.329<br>CLES:52.46% | U=1634780.5<br>**p=4.047 × $10^{-39}$**<br>CLES:64.86% | U=301110.0<br>**p=2.567 × $10^{-7}$**<br>CLES:62.17% |
| Horse | U=62046.0<br>p=0.366<br>CLES:52.34% | U=1713847.5<br>**p=2.27 × $10^{-53}$**<br>CLES:67.56% | U=305711.0<br>**p=3.62 × $10^{-10}$**<br>CLES:65.22% |
| Black snub-nosed monkey | U=61694.0<br>p=0.818<br>CLES:50.59% | U=1577306.0<br>**p=2.786 × $10^{-28}$**<br>CLES:62.54% | U=285358.5<br>**p=2.68 × $10^{-6}$**<br>CLES:61.33% |
| Brazilian guinea pig | U=31649.0<br>p=0.519<br>CLES:51.99% | U=826367.5<br>**p=6.808 × $10^{-18}$**<br>CLES:61.49% | U=139613.0<br>p=9.572 × $10^{-4}$<br>CLES:59.61% |
| Orangutan | U=61772.0<br>p=0.464<br>CLES:51.89% | U=1537073.0<br>**p=2.415 × $10^{-23}$**<br>CLES:61.38% | U=281777.0<br>**p=1.909 × $10^{-5}$**<br>CLES:60.34% |
| Northern American deer mouse | U=61500.0<br>p=0.631<br>CLES:51.23% | U=1627184.0<br>**p=3.63 × $10^{-52}$**<br>CLES:67.48% | U=303859.0<br>**p=1.117 × $10^{-11}$**<br>CLES:66.35% |
| Naked mole-rat female | U=60787.5<br>p=0.389<br>CLES:52.21% | U=1509509.5<br>**p=3.039 × $10^{-52}$**<br>CLES:67.84% | U=290709.0<br>**p=4.333 × $10^{-11}$**<br>CLES:65.84% |
| Mouse | U=67224.0<br>p=0.819<br>CLES:50.57% | U=1857569.5<br>**p=8.454 × $10^{-56}$**<br>CLES:67.60% | U=350417.0<br>**p=3.544 × $10^{-13}$**<br>CLES:67.05% |
| Daurian ground squirrel | U=41262.5<br>p=0.775<br>CLES:49.20% | U=1181884.0<br>**p=1.769 × $10^{-45}$**<br>CLES:67.69% | U=222802.0<br>**p=2.592 × $10^{-12}$**<br>CLES:68.50% |
| Tiger | U=36370.0<br>p=0.44<br>CLES:47.71% | U=1197344.5<br>**p=7.286 × $10^{-49}$**<br>CLES:68.18% | U=195459.0<br>**p=8.322 × $10^{-13}$**<br>CLES:70.12% |
| Lesser hedgehog tenrec | U=2965.0<br>p=0.422<br>CLES:45.79% | U=75839.0<br>**p=4.299 × $10^{-9}$**<br>CLES:64.39% | U=17317.0<br>**p=6.173 × $10^{-5}$**<br>CLES:69.54% |
| Mongolian gerbil | U=53890.0<br>p=0.422<br>CLES:52.13% | U=1343869.5<br>**p=6.204 × $10^{-42}$**<br>CLES:66.31% | U=254340.0<br>**p=7.852 × $10^{-9}$**<br>CLES:64.34% |
| Ma's night monkey | U=62047.5<br>p=0.45<br>CLES:51.95% | U=1654214.5<br>**p=1.432 × $10^{-45}$**<br>CLES:66.17% | U=292836.5<br>**p=1.137 × $10^{-8}$**<br>CLES:63.87% |

| | | | |
|---|---|---|---|
| Upper Galilee mountains blind mole rat | U=56855.5<br>p=0.495<br>CLES:51.78% | U=1531501.5<br>**$p=6.899 \times 10^{-51}$**<br>CLES:67.66% | U=292066.0<br>**$p=1.727 \times 10^{-11}$**<br>CLES:66.46% |
| Kangaroo rat | U=46178.0<br>p=0.498<br>CLES:51.88% | U=1290549.5<br>**$p=4.268 \times 10^{-38}$**<br>CLES:65.81% | U=237155.0<br>**$p=1.597 \times 10^{-8}$**<br>CLES:64.69% |
| Chimpanzee | U=64564.5<br>$p=1.002 \times 10^{-2}$<br>CLES:56.76% | U=1497740.5<br>**$p=6.329 \times 10^{-20}$**<br>CLES:60.40% | U=238341.0<br>$p=8.647 \times 10^{-2}$<br>CLES:54.21% |
| Guinea Pig | U=57989.0<br>p=0.561<br>CLES:51.52% | U=1548786.5<br>**$p=8.192 \times 10^{-41}$**<br>CLES:65.56% | U=287022.0<br>**$p=7.03 \times 10^{-9}$**<br>CLES:64.16% |
| American beaver | U=56080.0<br>p=0.928<br>CLES:49.77% | U=1333138.0<br>**$p=5.848 \times 10^{-37}$**<br>CLES:65.26% | U=284115.0<br>**$p=1.618 \times 10^{-10}$**<br>CLES:65.23% |
| Wild yak | U=47235.0<br>p=0.806<br>CLES:49.33% | U=1342675.0<br>**$p=1.735 \times 10^{-48}$**<br>CLES:67.60% | U=242861.0<br>**$p=1.668 \times 10^{-12}$**<br>CLES:68.23% |
| Coquerel's sifaka | U=58489.0<br>p=0.685<br>CLES:51.06% | U=1592808.5<br>**$p=1.984 \times 10^{-41}$**<br>CLES:65.53% | U=287282.0<br>**$p=4.499 \times 10^{-9}$**<br>CLES:64.39% |
| Alpaca | U=1998.0<br>p=0.672<br>CLES:47.44% | U=64296.5<br>**$p=1.47 \times 10^{-10}$**<br>CLES:66.58% | U=11635.0<br>$p=5.54 \times 10^{-4}$<br>CLES:69.62% |
| Lesser Egyptian jerboa | U=53674.0<br>$p=7.514 \times 10^{-2}$<br>CLES:54.79% | U=1345670.0<br>**$p=3.491 \times 10^{-44}$**<br>CLES:66.86% | U=242675.5<br>**$p=3.107 \times 10^{-7}$**<br>CLES:62.93% |
| Long-tailed chinchilla | U=60438.0<br>p=0.187<br>CLES:53.47% | U=1651481.0<br>**$p=6.081 \times 10^{-51}$**<br>CLES:67.23% | U=282431.0<br>**$p=5.49 \times 10^{-9}$**<br>CLES:64.45% |
| Angola colobus | U=61569.0<br>p=0.695<br>CLES:51.02% | U=1677860.5<br>**$p=1.502 \times 10^{-40}$**<br>CLES:65.09% | U=292773.0<br>**$p=3.528 \times 10^{-8}$**<br>CLES:63.43% |
| Ryukyu mouse | U=60992.0<br>p=0.986<br>CLES:49.95% | U=1714653.5<br>**$p=2.112 \times 10^{-55}$**<br>CLES:67.93% | U=326887.0<br>**$p=2.103 \times 10^{-13}$**<br>CLES:67.55% |
| Chinese hamster PICR | U=60697.5<br>p=0.849<br>CLES:49.52% | U=1672035.5<br>**$p=1.581 \times 10^{-48}$**<br>CLES:66.77% | U=327095.5<br>**$p=1.98 \times 10^{-13}$**<br>CLES:67.51% |
| Vervet-AGM | U=71371.0<br>p=0.266<br>CLES:52.75% | U=1734822.5<br>**$p=3.393 \times 10^{-40}$**<br>CLES:64.91% | U=326549.0<br>**$p=1.347 \times 10^{-7}$**<br>CLES:62.22% |
| Cow | U=61096.0<br>p=0.586<br>CLES:51.41% | U=1663885.5<br>**$p=2.387 \times 10^{-46}$**<br>CLES:66.33% | U=301915.5<br>**$p=2.99 \times 10^{-10}$**<br>CLES:65.30% |
| Bushbaby | U=60071.0<br>p=0.641<br>CLES:51.21% | U=1676321.0<br>**$p=2.131 \times 10^{-50}$**<br>CLES:67.09% | U=305801.5<br>**$p=1.501 \times 10^{-11}$**<br>CLES:66.44% |

| | | | |
|---|---|---|---|
| Bonobo | U=65611.0<br>p=$8.151 \times 10^{-2}$<br>CLES:54.47% | U=1453234.0<br>**p=$3.842 \times 10^{-13}$**<br>CLES:58.27% | U=251970.0<br>p=$8.91 \times 10^{-2}$<br>CLES:54.08% |
| Hyrax | U=6263.0<br>p=0.767<br>CLES:51.29% | U=112516.0<br>**p=$1.4 \times 10^{-10}$**<br>CLES:64.07% | U=26908.0<br>p=$3.263 \times 10^{-3}$<br>CLES:61.85% |
| Rabbit | U=42342.0<br>p=0.653<br>CLES:48.74% | U=1239310.0<br>**p=$1.295 \times 10^{-31}$**<br>CLES:64.33% | U=228171.0<br>**p=$1.928 \times 10^{-9}$**<br>CLES:65.85% |
| Megabat | U=15763.0<br>p=0.311<br>CLES:46.45% | U=405764.0<br>**p=$7.006 \times 10^{-23}$**<br>CLES:65.74% | U=81412.0<br>**p=$3.153 \times 10^{-8}$**<br>CLES:68.23% |
| Dog | U=55130.0<br>p=0.753<br>CLES:49.17% | U=1582714.0<br>**p=$5.229 \times 10^{-48}$**<br>CLES:66.85% | U=294254.0<br>**p=$2.136 \times 10^{-13}$**<br>CLES:68.14% |
| Steppe mouse | U=60743.0<br>p=0.597<br>CLES:51.37% | U=1659002.5<br>**p=$9.563 \times 10^{-54}$**<br>CLES:67.70% | U=300636.5<br>**p=$2.388 \times 10^{-11}$**<br>CLES:66.23% |
| Greater bamboo lemur | U=65517.0<br>p=0.524<br>CLES:51.61% | U=1716667.5<br>**p=$1.305 \times 10^{-49}$**<br>CLES:66.77% | U=316765.5<br>**p=$2.038 \times 10^{-10}$**<br>CLES:65.11% |
| Pig | U=62081.0<br>p=0.686<br>CLES:51.03% | U=1637774.0<br>**p=$9.93 \times 10^{-46}$**<br>CLES:66.32% | U=313909.0<br>**p=$1.76 \times 10^{-10}$**<br>CLES:65.21% |
| Tree Shrew | U=2257.0<br>p=0.98<br>CLES:50.16% | U=49296.5<br>**p=$8.221 \times 10^{-7}$**<br>CLES:63.20% | U=9884.0<br>p=$1.383 \times 10^{-2}$<br>CLES:63.36% |
| Pika | U=7377.0<br>p=0.131<br>CLES:43.82% | U=162668.0<br>**p=$3.02 \times 10^{-7}$**<br>CLES:60.08% | U=39563.0<br>**p=$2.36 \times 10^{-5}$**<br>CLES:66.11% |
| Shrew | U=1479.0<br>p=0.264<br>CLES:43.13% | U=43322.5<br>**p=$3.945 \times 10^{-5}$**<br>CLES:61.69% | U=10114.0<br>p=$1.843 \times 10^{-3}$<br>CLES:67.74% |
| Squirrel | U=60151.0<br>p=0.674<br>CLES:51.08% | U=1657325.5<br>**p=$4.818 \times 10^{-52}$**<br>CLES:67.49% | U=312040.5<br>**p=$7.959 \times 10^{-12}$**<br>CLES:66.50% |
| Gorilla | U=67014.5<br>p=0.122<br>CLES:53.96% | U=1599199.0<br>**p=$8.655 \times 10^{-22}$**<br>CLES:60.77% | U=272091.0<br>p=$3.228 \times 10^{-3}$<br>CLES:57.08% |
| Cat | U=62052.0<br>p=0.991<br>CLES:50.03% | U=1722684.5<br>**p=$3.834 \times 10^{-49}$**<br>CLES:66.70% | U=319314.0<br>**p=$8.485 \times 10^{-12}$**<br>CLES:66.37% |

### Human Reference Genome, Saunders et al., 2018

| MWU Test | Vasculature vs Glia | Vasculature vs Neuron | Glia vs Neuron |
|---|---|---|---|
| Microbat | U=39117.0<br>$p=1.506 \times 10^{-2}$<br>CLES:56.17% | U=105606.5<br>**$p=6.034 \times 10^{-15}$**<br>CLES:66.15% | U=67813.0<br>**$p=5.919 \times 10^{-7}$**<br>CLES:61.68% |
| Hedgehog | U=2803.0<br>$p=8.392 \times 10^{-2}$<br>CLES:58.54% | U=5583.0<br>$p=1.02 \times 10^{-4}$<br>CLES:66.78% | U=4134.0<br>$p=3.495 \times 10^{-2}$<br>CLES:59.65% |
| Naked mole-rat male | U=41118.0<br>$p=1.514 \times 10^{-2}$<br>CLES:56.08% | U=124378.0<br>**$p=6.684 \times 10^{-25}$**<br>CLES:70.92% | U=82431.0<br>**$p=1.455 \times 10^{-13}$**<br>CLES:66.88% |
| Sooty mangabey | U=56865.0<br>$p=3.273 \times 10^{-2}$<br>CLES:54.89% | U=150154.5<br>**$p=1.601 \times 10^{-15}$**<br>CLES:65.07% | U=104578.5<br>**$p=8.231 \times 10^{-8}$**<br>CLES:61.19% |
| Olive baboon | U=56933.5<br>$p=1.282 \times 10^{-2}$<br>CLES:55.73% | U=154789.5<br>**$p=7.468 \times 10^{-18}$**<br>CLES:66.22% | U=103972.0<br>**$p=1.569 \times 10^{-8}$**<br>CLES:61.88% |
| American bison | U=37599.5<br>$p=2.402 \times 10^{-2}$<br>CLES:55.78% | U=108223.0<br>**$p=3.403 \times 10^{-21}$**<br>CLES:69.69% | U=70973.0<br>**$p=2.803 \times 10^{-13}$**<br>CLES:67.27% |
| Leopard | U=56353.0<br>$p=1.877 \times 10^{-4}$<br>CLES:58.74% | U=150797.5<br>**$p=1.498 \times 10^{-21}$**<br>CLES:68.22% | U=96352.5<br>**$p=3.159 \times 10^{-8}$**<br>CLES:61.87% |
| Bolivian squirrel monkey | U=55637.5<br>$p=1.237 \times 10^{-4}$<br>CLES:59.02% | U=152270.0<br>**$p=1.208 \times 10^{-26}$**<br>CLES:70.54% | U=99427.0<br>**$p=7.315 \times 10^{-12}$**<br>CLES:64.74% |
| Crab-eating macaque | U=57007.5<br>$p=1.399 \times 10^{-2}$<br>CLES:55.65% | U=152426.0<br>**$p=1.534 \times 10^{-16}$**<br>CLES:65.57% | U=101544.5<br>**$p=2.063 \times 10^{-7}$**<br>CLES:60.93% |
| Red fox | U=47694.5<br>$p=8.28 \times 10^{-3}$<br>CLES:56.38% | U=135109.5<br>**$p=5.317 \times 10^{-17}$**<br>CLES:66.37% | U=86236.5<br>**$p=1.273 \times 10^{-7}$**<br>CLES:61.69% |
| Shrew mouse | U=58554.0<br>$p=1.04 \times 10^{-4}$<br>CLES:58.99% | U=157267.0<br>**$p=5.984 \times 10^{-24}$**<br>CLES:69.19% | U=103792.0<br>**$p=2.143 \times 10^{-9}$**<br>CLES:62.64% |
| Arctic ground squirrel | U=58207.0<br>**$p=1.581 \times 10^{-5}$**<br>CLES:60.06% | U=156788.5<br>**$p=2.435 \times 10^{-28}$**<br>CLES:71.16% | U=102941.5<br>**$p=1.858 \times 10^{-10}$**<br>CLES:63.52% |
| Chinese hamster CriGri | U=47869.0<br>$p=1.241 \times 10^{-3}$<br>CLES:57.84% | U=126576.0<br>**$p=7.33 \times 10^{-19}$**<br>CLES:67.66% | U=81347.0<br>**$p=2.817 \times 10^{-7}$**<br>CLES:61.45% |
| Opossum | U=42418.0<br>$p=1.175 \times 10^{-2}$<br>CLES:56.26% | U=106016.0<br>**$p=3.727 \times 10^{-9}$**<br>CLES:62.02% | U=68701.0<br>$p=5.675 \times 10^{-3}$<br>CLES:56.30% |
| Dolphin | U=14550.5<br>$p=1.286 \times 10^{-2}$ | U=39903.5<br>**$p=7.945 \times 10^{-13}$** | U=22591.5<br>**$p=5.089 \times 10^{-5}$** |

| | | | |
|---|---|---|---|
| | CLES:58.20% | CLES:69.04% | CLES:62.54% |
| Capuchin | U=57934.0 | U=154031.5 | U=103153.5 |
| | **p=$1.314 \times 10^{-5}$** | **p=$1.52 \times 10^{-28}$** | **p=$1.359 \times 10^{-10}$** |
| | CLES:60.16% | CLES:71.35% | CLES:63.59% |
| Mouse Lemur | U=59057.5 | U=151583.0 | U=98563.0 |
| | **p=$3.004 \times 10^{-6}$** | **p=$4.053 \times 10^{-25}$** | **p=$7.895 \times 10^{-8}$** |
| | CLES:60.87% | CLES:69.91% | CLES:61.40% |
| Damara mole rat | U=47418.5 | U=135453.5 | U=92158.0 |
| | p=$3.7 \times 10^{-3}$ | **p=$4.415 \times 10^{-22}$** | **p=$3.162 \times 10^{-11}$** |
| | CLES:57.03% | CLES:69.09% | CLES:64.59% |
| Marmoset | U=56439.0 | U=154740.0 | U=93885.5 |
| | **p=$7.891 \times 10^{-7}$** | **p=$2.112 \times 10^{-27}$** | **p=$4.099 \times 10^{-8}$** |
| | CLES:61.70% | CLES:70.81% | CLES:61.89% |
| Goat | U=54750.0 | U=149937.0 | U=97427.5 |
| | p=$5.867 \times 10^{-4}$ | **p=$2.464 \times 10^{-22}$** | **p=$3.393 \times 10^{-9}$** |
| | CLES:58.07% | CLES:68.65% | CLES:62.69% |
| Common wombat | U=36501.0 | U=101726.5 | U=64814.0 |
| | p=$3.765 \times 10^{-2}$ | **p=$3.086 \times 10^{-11}$** | **p=$5.674 \times 10^{-6}$** |
| | CLES:55.35% | CLES:63.77% | CLES:60.76% |
| American black bear | U=46945.0 | U=105822.5 | U=66074.0 |
| | p=$4.173 \times 10^{-4}$ | **p=$2.078 \times 10^{-11}$** | p=$1.165 \times 10^{-2}$ |
| | CLES:58.63% | CLES:63.69% | CLES:55.74% |
| Sloth | U=1005.0 | U=2248.0 | U=1409.0 |
| | p=0.129 | **p=$6.516 \times 10^{-5}$** | p=$5.018 \times 10^{-2}$ |
| | CLES:59.82% | CLES:72.05% | CLES:61.93% |
| Polar bear | U=43756.0 | U=113549.5 | U=73008.5 |
| | p=$1.703 \times 10^{-3}$ | **p=$7.735 \times 10^{-16}$** | **p=$2.554 \times 10^{-5}$** |
| | CLES:57.78% | CLES:66.42% | CLES:59.57% |
| Alpine marmot | U=49873.0 | U=136088.0 | U=85101.0 |
| | p=$3.067 \times 10^{-4}$ | **p=$4.102 \times 10^{-23}$** | **p=$4.726 \times 10^{-9}$** |
| | CLES:58.71% | CLES:69.48% | CLES:63.02% |
| Prairie vole | U=50726.0 | U=143126.0 | U=104389.5 |
| | p=$2.705 \times 10^{-2}$ | **p=$3.073 \times 10^{-22}$** | **p=$2.872 \times 10^{-14}$** |
| | CLES:55.21% | CLES:68.89% | CLES:66.23% |
| Donkey | U=59405.5 | U=147966.0 | U=97657.0 |
| | **p=$7.6 \times 10^{-5}$** | **p=$1.257 \times 10^{-19}$** | **p=$2.597 \times 10^{-6}$** |
| | CLES:59.13% | CLES:67.35% | CLES:59.92% |
| Golden Hamster | U=45822.0 | U=130293.5 | U=87777.5 |
| | p=$2.056 \times 10^{-2}$ | **p=$3.583 \times 10^{-20}$** | **p=$1.693 \times 10^{-10}$** |
| | CLES:55.62% | CLES:68.27% | CLES:64.16% |
| Armadillo | U=48272.0 | U=120867.0 | U=75104.0 |
| | p=$2.931 \times 10^{-4}$ | **p=$1.584 \times 10^{-14}$** | p=$4.636 \times 10^{-4}$ |
| | CLES:58.81% | CLES:65.33% | CLES:57.84% |
| Ugandan red Colobus | U=57786.0 | U=152708.5 | U=99581.0 |
| | p=$2.941 \times 10^{-4}$ | **p=$6.929 \times 10^{-22}$** | **p=$7.302 \times 10^{-9}$** |
| | CLES:58.40% | CLES:68.30% | CLES:62.29% |
| Pig-tailed macaque | U=57369.5 | U=152251.0 | U=104477.0 |
| | p=$1.178 \times 10^{-2}$ | **p=$7.378 \times 10^{-17}$** | **p=$5.352 \times 10^{-8}$** |

| | | | |
|---|---|---|---|
| | CLES:55.78% | CLES:65.77% | CLES:61.38% |
| Panda | U=56078.0<br>$p=2.116\times10^{-4}$<br>CLES:58.68% | U=146509.0<br>**$p=2.665\times10^{-22}$**<br>CLES:68.69% | U=93983.0<br>**$p=3.276\times10^{-9}$**<br>CLES:62.78% |
| Ferret | U=52603.0<br>$p=1.847\times10^{-3}$<br>CLES:57.37% | U=144283.0<br>**$p=2.258\times10^{-20}$**<br>CLES:67.86% | U=93995.0<br>**$p=3.29\times10^{-9}$**<br>CLES:62.83% |
| Koala | U=44909.0<br>$p=4.656\times10^{-2}$<br>CLES:54.84% | U=117243.5<br>**$p=2.025\times10^{-10}$**<br>CLES:62.69% | U=80617.0<br>**$p=1.084\times10^{-5}$**<br>CLES:59.78% |
| Wallaby | U=4198.0<br>p=0.653<br>CLES:48.09% | U=7588.0<br>$p=3.916\times10^{-2}$<br>CLES:57.95% | U=7360.5<br>$p=7.225\times10^{-3}$<br>CLES:60.58% |
| Tarsier | U=40238.0<br>$p=4.805\times10^{-3}$<br>CLES:57.11% | U=99412.0<br>**$p=2.605\times10^{-17}$**<br>CLES:67.87% | U=68007.0<br>**$p=2.817\times10^{-8}$**<br>CLES:62.93% |
| Degu | U=40843.0<br>$p=7.942\times10^{-4}$<br>CLES:58.48% | U=121489.0<br>**$p=2.912\times10^{-24}$**<br>CLES:70.88% | U=82920.0<br>**$p=1.865\times10^{-10}$**<br>CLES:64.43% |
| Chinese hamster CHOK1GS | U=59039.0<br>**$p=2.061\times10^{-5}$**<br>CLES:59.89% | U=159550.5<br>**$p=3.853\times10^{-26}$**<br>CLES:70.08% | U=100939.0<br>**$p=8.309\times10^{-9}$**<br>CLES:62.24% |
| Drill | U=51019.0<br>$p=4.655\times10^{-2}$<br>CLES:54.69% | U=138283.5<br>**$p=2.924\times10^{-15}$**<br>CLES:65.23% | U=92140.5<br>**$p=1.35\times10^{-7}$**<br>CLES:61.39% |
| Gibbon | U=52068.5<br>$p=1.012\times10^{-2}$<br>CLES:56.06% | U=134831.0<br>**$p=3.762\times10^{-11}$**<br>CLES:62.74% | U=87941.0<br>$p=9.658\times10^{-4}$<br>CLES:57.09% |
| Rat | U=52908.0<br>$p=2.222\times10^{-4}$<br>CLES:58.76% | U=143359.0<br>**$p=4.646\times10^{-23}$**<br>CLES:69.27% | U=95927.0<br>**$p=3.742\times10^{-9}$**<br>CLES:62.70% |
| Macaque | U=55226.0<br>$p=1.095\times10^{-2}$<br>CLES:55.90% | U=150758.0<br>**$p=7.176\times10^{-18}$**<br>CLES:66.36% | U=100960.0<br>**$p=4.382\times10^{-8}$**<br>CLES:61.58% |
| American mink | U=54226.0<br>$p=1.714\times10^{-3}$<br>CLES:57.36% | U=141065.5<br>**$p=1.099\times10^{-19}$**<br>CLES:67.57% | U=92221.0<br>**$p=1.495\times10^{-8}$**<br>CLES:62.25% |
| Gelada | U=56869.0<br>$p=1.779\times10^{-2}$<br>CLES:55.44% | U=156122.5<br>**$p=1.519\times10^{-19}$**<br>CLES:67.08% | U=108233.0<br>**$p=3.81\times10^{-10}$**<br>CLES:63.07% |
| Sheep | U=49967.5<br>$p=5.315\times10^{-4}$<br>CLES:58.34% | U=138210.5<br>**$p=3.521\times10^{-21}$**<br>CLES:68.48% | U=87325.5<br>**$p=2.907\times10^{-8}$**<br>CLES:62.24% |
| Tasmanian devil | U=39031.0<br>$p=5.05\times10^{-3}$<br>CLES:57.13% | U=98063.0<br>**$p=2.107\times10^{-11}$**<br>CLES:64.05% | U=64933.0<br>$p=8.484\times10^{-4}$<br>CLES:57.74% |
| Elephant | U=54258.5<br>$p=1.023\times10^{-4}$ | U=138860.0<br>**$p=2.39\times10^{-19}$** | U=90764.5<br>**$p=1.748\times10^{-6}$** |

| | | | |
|---|---|---|---|
| | CLES:59.18% | CLES:67.51% | CLES:60.32% |
| Algerian mouse | U=58408.0<br>$p=2.443 \times 10^{-4}$<br>CLES:58.49% | U=160636.0<br>**$p=2.042 \times 10^{-25}$**<br>CLES:69.73% | U=104725.0<br>**$p=1.042 \times 10^{-10}$**<br>CLES:63.67% |
| Dingo | U=52944.5<br>**$p=7.232 \times 10^{-5}$**<br>CLES:59.45% | U=136573.5<br>**$p=7.861 \times 10^{-18}$**<br>CLES:66.78% | U=86619.5<br>**$p=4.073 \times 10^{-5}$**<br>CLES:58.93% |
| Golden snub-nosed monkey | U=50650.0<br>$p=4.022 \times 10^{-2}$<br>CLES:54.85% | U=140183.0<br>**$p=6.653 \times 10^{-17}$**<br>CLES:66.10% | U=92153.5<br>**$p=5.285 \times 10^{-9}$**<br>CLES:62.69% |
| Horse | U=56022.5<br>$p=1.798 \times 10^{-4}$<br>CLES:58.78% | U=157469.0<br>**$p=7.618 \times 10^{-26}$**<br>CLES:70.03% | U=99140.0<br>**$p=3.511 \times 10^{-9}$**<br>CLES:62.65% |
| Black snub-nosed monkey | U=55755.0<br>$p=3.298 \times 10^{-3}$<br>CLES:56.84% | U=139670.5<br>**$p=5.434 \times 10^{-13}$**<br>CLES:63.79% | U=89235.0<br>$p=2.707 \times 10^{-4}$<br>CLES:57.80% |
| Brazilian guinea pig | U=29916.0<br>$p=9.067 \times 10^{-3}$<br>CLES:57.09% | U=73799.0<br>**$p=8.56 \times 10^{-9}$**<br>CLES:62.91% | U=49217.0<br>$p=7.223 \times 10^{-3}$<br>CLES:56.65% |
| Orangutan | U=52956.0<br>p=0.253<br>CLES:52.64% | U=134373.0<br>**$p=6.515 \times 10^{-8}$**<br>CLES:60.28% | U=92850.0<br>$p=1.868 \times 10^{-4}$<br>CLES:57.91% |
| Northern American deer mouse | U=54986.0<br>$p=3.649 \times 10^{-3}$<br>CLES:56.79% | U=151014.0<br>**$p=3.294 \times 10^{-23}$**<br>CLES:68.99% | U=99570.0<br>**$p=8.653 \times 10^{-12}$**<br>CLES:64.66% |
| Naked mole-rat female | U=49148.0<br>$p=8.566 \times 10^{-4}$<br>CLES:58.05% | U=143428.0<br>**$p=4.834 \times 10^{-26}$**<br>CLES:70.66% | U=92874.0<br>**$p=7.711 \times 10^{-12}$**<br>CLES:65.07% |
| Mouse | U=61552.0<br>$p=2.819 \times 10^{-4}$<br>CLES:58.29% | U=167603.0<br>**$p=1.117 \times 10^{-24}$**<br>CLES:69.17% | U=109935.0<br>**$p=1.569 \times 10^{-10}$**<br>CLES:63.35% |
| Daurian ground squirrel | U=35629.0<br>$p=2.35 \times 10^{-2}$<br>CLES:55.88% | U=105439.0<br>**$p=9.162 \times 10^{-22}$**<br>CLES:70.15% | U=66828.0<br>**$p=1.56 \times 10^{-11}$**<br>CLES:66.16% |
| Tiger | U=35590.0<br>$p=4.131 \times 10^{-4}$<br>CLES:59.32% | U=98329.0<br>**$p=1.431 \times 10^{-19}$**<br>CLES:69.29% | U=58742.0<br>**$p=6.917 \times 10^{-7}$**<br>CLES:62.10% |
| Lesser hedgehog tenrec | U=3456.5<br>$p=4.36 \times 10^{-2}$<br>CLES:59.49% | U=5974.5<br>**$p=3.22 \times 10^{-6}$**<br>CLES:69.89% | U=4362.5<br>$p=1.922 \times 10^{-2}$<br>CLES:60.51% |
| Mongolian gerbil | U=45585.0<br>$p=5.263 \times 10^{-4}$<br>CLES:58.56% | U=114322.0<br>**$p=2.277 \times 10^{-16}$**<br>CLES:66.65% | U=70361.0<br>**$p=1.753 \times 10^{-5}$**<br>CLES:59.84% |
| Ma's night monkey | U=57483.0<br>**$p=2.459 \times 10^{-6}$**<br>CLES:61.05% | U=150855.0<br>**$p=3.872 \times 10^{-26}$**<br>CLES:70.44% | U=99188.0<br>**$p=4.358 \times 10^{-8}$**<br>CLES:61.63% |

| | | | |
|---|---|---|---|
| Upper Galilee mountains blind mole rat | U=51950.5<br>$p=5.636 \times 10^{-4}$<br>CLES:58.20% | U=143697.0<br>**$p=2.071 \times 10^{-23}$**<br>CLES:69.45% | U=98298.0<br>**$p=1.849 \times 10^{-10}$**<br>CLES:63.71% |
| Kangaroo rat | U=42834.5<br>$p=2.005 \times 10^{-2}$<br>CLES:55.73% | U=121314.0<br>**$p=2.264 \times 10^{-19}$**<br>CLES:68.28% | U=88284.0<br>**$p=1.615 \times 10^{-10}$**<br>CLES:64.15% |
| Chimpanzee | U=50153.5<br>p=0.102<br>CLES:53.82% | U=123848.5<br>**$p=7.253 \times 10^{-6}$**<br>CLES:58.57% | U=86601.0<br>$p=2.309 \times 10^{-2}$<br>CLES:54.76% |
| Guinea Pig | U=51029.0<br>$p=1.746 \times 10^{-2}$<br>CLES:55.62% | U=144208.5<br>**$p=8.677 \times 10^{-22}$**<br>CLES:68.58% | U=98018.0<br>**$p=8.382 \times 10^{-12}$**<br>CLES:64.75% |
| American beaver | U=49411.0<br>$p=4.152 \times 10^{-3}$<br>CLES:56.88% | U=117925.0<br>**$p=3.685 \times 10^{-19}$**<br>CLES:68.02% | U=74828.0<br>**$p=1.033 \times 10^{-8}$**<br>CLES:62.97% |
| Wild yak | U=40324.0<br>$p=2.984 \times 10^{-2}$<br>CLES:55.46% | U=118066.0<br>**$p=2.871 \times 10^{-21}$**<br>CLES:69.28% | U=76436.5<br>**$p=5.003 \times 10^{-13}$**<br>CLES:66.77% |
| Coquerel's sifaka | U=56714.0<br>**$p=1.412 \times 10^{-5}$**<br>CLES:60.19% | U=146094.0<br>**$p=1.737 \times 10^{-21}$**<br>CLES:68.36% | U=93687.5<br>**$p=7.222 \times 10^{-7}$**<br>CLES:60.63% |
| Alpaca | U=1362.0<br>$p=8.773 \times 10^{-2}$<br>CLES:60.35% | U=4659.0<br>**$p=8.51 \times 10^{-9}$**<br>CLES:77.15% | U=2350.0<br>$p=1.131 \times 10^{-2}$<br>CLES:64.16% |
| Lesser Egyptian jerboa | U=39972.0<br>p=0.243<br>CLES:52.89% | U=118930.0<br>**$p=1.588 \times 10^{-18}$**<br>CLES:67.92% | U=91393.0<br>**$p=3.485 \times 10^{-14}$**<br>CLES:66.80% |
| Long-tailed chinchilla | U=50854.0<br>$p=2.489 \times 10^{-3}$<br>CLES:57.20% | U=141527.0<br>**$p=1.868 \times 10^{-21}$**<br>CLES:68.54% | U=94473.0<br>**$p=1.788 \times 10^{-9}$**<br>CLES:63.04% |
| Angola colobus | U=54874.0<br>$p=3.2 \times 10^{-3}$<br>CLES:56.89% | U=149059.0<br>**$p=5.695 \times 10^{-19}$**<br>CLES:66.96% | U=95970.5<br>**$p=7.627 \times 10^{-8}$**<br>CLES:61.52% |
| Ryukyu mouse | U=57363.0<br>$p=2.805 \times 10^{-4}$<br>CLES:58.44% | U=156659.0<br>**$p=8.019 \times 10^{-24}$**<br>CLES:69.14% | U=102823.5<br>**$p=9.895 \times 10^{-10}$**<br>CLES:62.96% |
| Chinese hamster PICR | U=59426.0<br>**$p=9.635 \times 10^{-7}$**<br>CLES:61.44% | U=161613.0<br>**$p=1.324 \times 10^{-31}$**<br>CLES:72.29% | U=98608.5<br>**$p=9.821 \times 10^{-10}$**<br>CLES:63.11% |
| Vervet-AGM | U=62024.0<br>$p=1.316 \times 10^{-3}$<br>CLES:57.28% | U=154353.5<br>**$p=2.777 \times 10^{-16}$**<br>CLES:65.36% | U=102280.0<br>**$p=1.525 \times 10^{-5}$**<br>CLES:58.97% |
| Cow | U=54472.0<br>$p=4.909 \times 10^{-4}$<br>CLES:58.20% | U=147717.0<br>**$p=4.363 \times 10^{-22}$**<br>CLES:68.62% | U=98279.0<br>**$p=7.438 \times 10^{-10}$**<br>CLES:63.21% |
| Bushbaby | U=56929.0<br>$p=7.468 \times 10^{-4}$<br>CLES:57.82% | U=150022.5<br>**$p=1.601 \times 10^{-21}$**<br>CLES:68.27% | U=103656.5<br>**$p=1.318 \times 10^{-9}$**<br>CLES:62.78% |

| | | | |
|---|---|---|---|
| Bonobo | U=49766.0<br>p=0.168<br>CLES:53.23% | U=123739.0<br>**$p=2.89 \times 10^{-5}$**<br>CLES:58.00% | U=84271.0<br>$p=1.771 \times 10^{-2}$<br>CLES:55.05% |
| Hyrax | U=4396.0<br>$p=4.671 \times 10^{-2}$<br>CLES:58.82% | U=9548.0<br>**$p=1.041 \times 10^{-6}$**<br>CLES:68.50% | U=6481.0<br>$p=1.245 \times 10^{-3}$<br>CLES:63.46% |
| Rabbit | U=41027.0<br>$p=1.61 \times 10^{-3}$<br>CLES:57.96% | U=111088.0<br>**$p=1.138 \times 10^{-14}$**<br>CLES:65.83% | U=70387.0<br>$p=1.201 \times 10^{-4}$<br>CLES:58.84% |
| Megabat | U=15070.0<br>$p=6.734 \times 10^{-4}$<br>CLES:61.19% | U=32070.0<br>**$p=2.125 \times 10^{-10}$**<br>CLES:67.69% | U=19344.0<br>$p=5.767 \times 10^{-3}$<br>CLES:58.60% |
| Dog | U=50979.0<br>$p=1.079 \times 10^{-4}$<br>CLES:59.31% | U=139676.0<br>**$p=1.023 \times 10^{-21}$**<br>CLES:68.73% | U=88252.0<br>**$p=3.402 \times 10^{-7}$**<br>CLES:61.18% |
| Steppe mouse | U=51817.0<br>$p=3.341 \times 10^{-3}$<br>CLES:56.94% | U=144555.0<br>**$p=3.357 \times 10^{-23}$**<br>CLES:69.30% | U=100545.0<br>**$p=4.112 \times 10^{-12}$**<br>CLES:64.88% |
| Greater bamboo lemur | U=59124.5<br>$p=3.274 \times 10^{-4}$<br>CLES:58.28% | U=153648.5<br>**$p=3.773 \times 10^{-23}$**<br>CLES:68.88% | U=102403.5<br>**$p=8.578 \times 10^{-10}$**<br>CLES:62.96% |
| Pig | U=54501.5<br>$p=1.813 \times 10^{-3}$<br>CLES:57.31% | U=148517.0<br>**$p=5.396 \times 10^{-22}$**<br>CLES:68.53% | U=99661.0<br>**$p=1.477 \times 10^{-10}$**<br>CLES:63.72% |
| Tree Shrew | U=2203.0<br>$p=5.518 \times 10^{-3}$<br>CLES:65.10% | U=3836.0<br>$p=2.209 \times 10^{-4}$<br>CLES:67.44% | U=1951.0<br>p=0.635<br>CLES:52.55% |
| Pika | U=7576.0<br>$p=6.576 \times 10^{-3}$<br>CLES:60.60% | U=13897.0<br>**$p=9.541 \times 10^{-8}$**<br>CLES:68.29% | U=8519.5<br>$p=1.486 \times 10^{-2}$<br>CLES:59.24% |
| Shrew | U=1402.0<br>$p=1.158 \times 10^{-2}$<br>CLES:65.36% | U=3082.5<br>$p=2.819 \times 10^{-4}$<br>CLES:68.35% | U=1723.0<br>p=0.493<br>CLES:53.88% |
| Squirrel | U=53554.0<br>$p=3.71 \times 10^{-4}$<br>CLES:58.41% | U=149783.5<br>**$p=5.989 \times 10^{-25}$**<br>CLES:69.94% | U=99125.0<br>**$p=6.316 \times 10^{-10}$**<br>CLES:63.26% |
| Gorilla | U=48783.0<br>p=0.862<br>CLES:50.40% | U=129326.0<br>**$p=1.6 \times 10^{-5}$**<br>CLES:58.18% | U=91595.0<br>$p=2.947 \times 10^{-4}$<br>CLES:57.66% |
| Cat | U=55691.5<br>$p=2.753 \times 10^{-4}$<br>CLES:58.52% | U=151186.0<br>**$p=4.104 \times 10^{-22}$**<br>CLES:68.50% | U=96432.0<br>**$p=7.878 \times 10^{-8}$**<br>CLES:61.51% |
