## Supplementary Table S8 for "Neuron-specific coding and regulatory sequences are the most highly conserved in amniote brains despite neuron-specific cell size diversity"

| *Species* | *Neuron-expressed Genes median dN/dS ($CI_{95\%}$)* | *Glia-expressed Genes median dN/dS ($CI_{95\%}$)* | *Mann-Whitney U Test p-value* |
|---|---|---|---|
| Human | 0.079 (0.077-0.082) | 0.085 (0.083-0.087) | $1.94 \times 10^{-5}$ |
| Rat | 0.093 (0.090-0.095) | 0.099 (0.096-0.102) | $2.11 \times 10^{-5}$ |
| Opossum | 0.071 (0.069-0.073) | 0.074 (0.072-0.076) | 0.011 |
| Megabat | 0.093 (0.090-0.096) | 0.098 (0.095-0.101) | 0.007 |
| Tasmanian devil | 0.069 (0.067-0.070) | 0.072 (0.070-0.074) | 0.006 |
| Cat | 0.081 (0.080-0.084) | 0.087 (0.085-0.089) | $3.74\times10^{-5}$ |
| Pig | 0.082 (0.080-0.084) | 0.087 (0.085-0.089) | $1.68\times10^{-5}$ |
| Average of 92 species | 0.088 (0.086-0.090) | 0.093 (0.091-0.095) | $1.29\times10^{-6}$ |
