## Supplementary Table S10 for "Neuron-specific coding and regulatory sequences are the most highly conserved in amniote brains despite neuron-specific cell size diversity"

| Dataset | ANCOVA p-value | F-statistics |
|---|---|---|
| Zhang et al., 2014 | $7.68\times10^{-31}$ | $F(2, 3442)=70.76$ |
| Zeisel et al., 2018 | $2.65\times10^{-155}$ | $F(2, 4682)=384.41$ |
| Saunders et al., 2018 | $3.97\times10^{-36}$ | $F(2, 1403)=86.44$ |
